# *Escherichia coli σ*^38^ promoters use two UP elements instead of a −35 element: resolution of a paradox and discovery that *σ*^38^ transcribes ribosomal promoters

**DOI:** 10.1101/2020.02.05.936344

**Authors:** Kevin S. Franco, Zhe Sun, Yixiong Chen, Cedric Cagliero, Yuhong Zuo, Yan Ning Zhou, Mikhail Kashlev, Ding Jun Jin, Thomas D. Schneider

## Abstract

In *E. coli*, one RNA polymerase (RNAP) transcribes all RNA species, and different regulons are transcribed by employing different sigma (*σ*) factors. RNAP containing *σ*^38^ (*σ*^*S*^) activates genes responding to stress conditions such as stationary phase. The structure of *σ*^38^ promoters has been controversial for more than two decades. To construct a model of *σ*^38^ promoters using information theory, we aligned proven transcriptional start sites to maximize the sequence information, in bits, and identified a −10 element similar to *σ*^70^ promoters. We could not align any −35 sequence logo; instead we found two patterns upstream of the −35 region. These patterns have dyad symmetry sequences and correspond to the location of UP elements in ribosomal RNA (rRNA) promoters. Additionally the UP element dyad symmetry suggests that the two polymerase *α* subunits, which bind to the UPs, should have two-fold dyad axis of symmetry on the polymerase and this is indeed observed in an X-ray crystal structure. Curiously the *α*CTDs should compete for overlapping UP elements. *In vitro* experiments confirm that *σ*^38^ recognizes the *rrnB* P1 promoter, requires a −10, UP elements and no −35. This clarifies the long-standing paradox of how *σ*^38^ promoters differ from those of *σ*^70^.

## 2 Introduction

### Current Models of *σ*^70^ and *σ*^38^

In *Escherichia coli* a *σ* factor is required for the initiation of transcription to recognize specific promoters [1]. The *σ* factor binds to the core enzyme of the RNA polymerase that consists of five subunits (*α*_2_*ββ*′*ω*) to form the holoenzyme. *E. coli* has seven different *σ* factors [2, 3]. *σ*^70^ RNAP transcribes most of the genome and the house-keeping genes responsible for basic cellular functions and it is capable of recognizing −10 and −35 elements that are approximately 10 and 35 bp upstream of the transcriptional start site, respectively [4]. It also transcribes from promoters which have an extended −10 element but no −35 element [5, 4] In some cases there are also UP sequence elements upstream of the −35 [6]. In contrast *σ*^38^ responds to stress and it can regulate the expression of nearly 15% of the genome [7, 8, 9]. Specifically, *σ*^38^ RNAP transcribes genes at the beginning of stationary phase and responds to osmotic and oxidative stress.

Current understanding suggests that *σ*^38^ is capable of recognizing −10 and −35 elements, and the differences between *σ*^38^ and *σ*^70^ are thought to be in those elements [10, 7, 11, 12, 13]. However, the promoter sequences recognized by *σ*^38^ remain to be defined precisely and the difference between *σ*^70^ and *σ*^38^ has been paradoxical for more than 20 years [14, 15, 16, 17, 12, 1]. Constructing a model of how *σ*^38^ binds to DNA may reveal the motifs it uses and resolve the paradox. It has been suggested that global transcription factors such as CRP, Fis, LRP or HNS affect the selectivity of *σ*^38^ [11]. A *σ*^38^ model could also clarify the relationship between the different global transcription factors and *σ*^38^ sites.

Both of the −10 and −35 elements of *σ*^38^ are said to be in a similar location as those of the *σ*^70^ elements, but they are not identical to those of *σ*^70^. The differences said to favor the transcription of *σ*^38^ are the presence of a C at position −13, a T at −14, an A/T rich region upstream of the −10 region or downstream of the −35, and a distal UP element [12]. The differences said to favor the transcription of *σ*^70^ are the presence of a strongly conserved −35 and a proximal UP element upstream of the −35 region [11]. There is still a mystery as to exactly what is needed to favor *σ*^38^ in terms of the −35 region. A wide variety of possible explanations have been proposed such as *σ*^38^ is favored by the presence of a −35 [17], or a lack of the −35 or even by the presence of a degenerate −35 element [11, 18]. Some experiments have been interpreted as implying that *σ*^38^ can use a variety of different sequences in the −35 region because there are different sequences for different *σ*^38^ controlled genes [19]. This has led to the conclusion that *σ*^38^ can use a degenerate −35 sequence. It has been hypothesized that the diverse sequences at the −35 makes *σ*^38^ able to respond to a variety of different stresses [19]. In previous attempts to identify a conserved sequence, 31 promoters controlled by *σ*^38^ located by microarray experiments were aligned to make a sequence logo, but only a conserved −10 and no −35 or UP elements could be found [18]. The interactions of *σ*^38^ and its promoter sequences remain to be determined. Here we apply information theory to define *σ*^38^ promoter sequences.

### Information Theory of Binding Sites

The programs we used to build *σ*^38^ models and to perform sequence analyses are based on information theory. In this 1948 theory Claude Shannon showed how to measure information transferred during communication in bits, the choice between two equally likely possibilities [20, 21]. Information theory is also a method to quantify the base sequence patterns in DNA or RNA that sequence recognizers bind to [22]. In this case, bits are related by an inequality to binding energy [23, 24]. When we make a sequence logo to graphically depict a set of binding sites [25] we can also calculate an average information content for the aligned sequences by adding together the information of all the base positions. The average information content has units of bits per site and is called *R*_*sequence*_ [22]. This average information content does not help us to interpret the information content for specific sequences, so we also make sequence walkers, another graphical display whose individual information contents are guaranteed to average to *R*_*sequence*_ [26, 27]. Sequence walkers allow us to locate specific binding sites on a sequence by **scan**ning models built from an information-theory generated weight matrix [27]. The information content for these binding sites is calculated and in general the higher the information content of a located binding site the higher the chance of the site being actually present there and being strongly bound [28], but too strong binding can be deleterious [29]. This method allows us to quantitatively predict the way proteins bind to and interact with the DNA sequence.

When a binding site has several parts that can reside at variable distances, such as the −10 and −35, each distance *d* has a different cost and this is reflected by its probability of use, *p*_*d*_. The cost can be expressed in bits by taking the negative log of the probability of a given distance. This is called the ‘gap surprisal’, *u*_*d*_ = −log_2_ *p*_*d*_ [30, 31]. With this definition, the average cost for all sites is the uncertainty of the gap, *H* = −∑_*d*_ *p*_*d*_ log_2_ *p*_*d*_. We originally used the gap surprisal to characterize the distance between the *E. coli* ribosomal binding site initiation codon and the Shine-Dalgarno [31]. Later we used this method to describe *σ*^70^ binding sites [4]. In both cases individual information weight matrices for the rigid binding sites were combined with a histogram of the distances between the sites and the results are displayed using sequence walkers.

## 3 Materials and Methods

The source code and documentation of all programs are available from: https://alum.mit.edu/www/toms/delila/delilaprograms.html

We began building a *σ*^38^ model by aligning the sequences of 78 transcription start sites determined by transcription initiation mapping, as recorded in the RegulonDB database (Release: 8.2 Date: April-22-2013) [32] for *E. coli* K-12 MG1655 in GenBank Accession NC000913 version 2 (Fig. 1; Note: the PDFs of all figures are in the supplement so that readers can examine their details). An unaligned set of sequences of a binding site will, in general, have a low information content in a specified range. However, by sliding any one sequence left and right an alignment with a higher information content may be found. The **malign** program repeats this shuffling process for all sequences until a global maximum is found. The program is fast enough that the sequences can be randomly misaligned repeatedly and then realigned to determine many different alignments. Generally we select the alignment with the highest information content, often the one found most frequently by **malign** [33].

**Figure 1:**
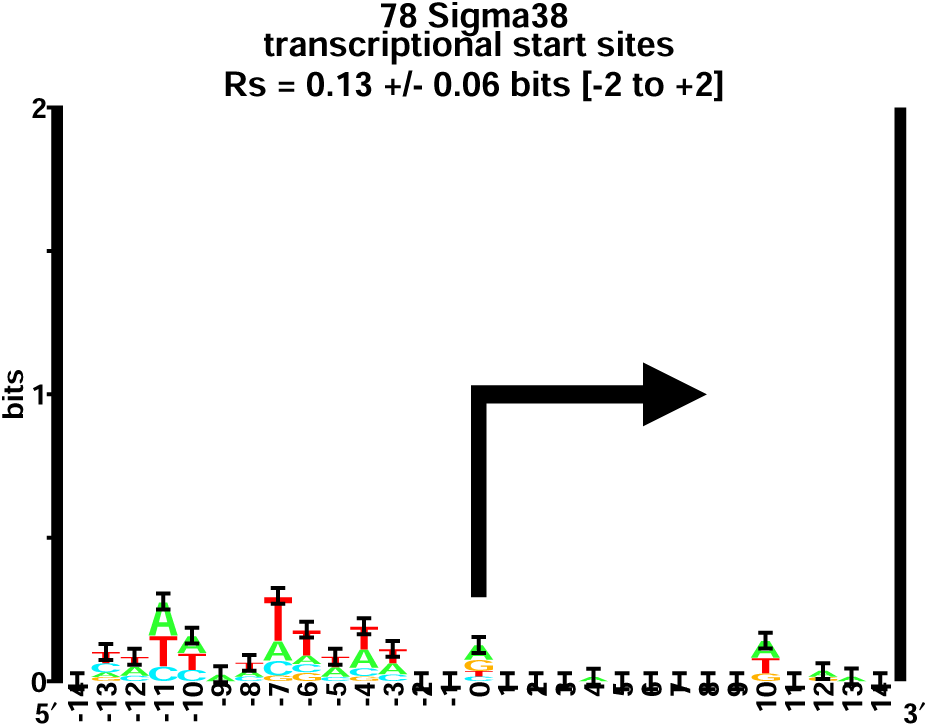
Sequence logo of *σ*^38^ promoter initiation regions. Sequence logos are a graphical representation of an aligned set of sequences. A logo shows a collection of sequences by stacks of letters in which the height of each letter is proportional to its frequency at that position and the total height of the stack is the information measured in bits [25, 72]. On top of each stack are uncertainty bars based on the number of sequences [22]. Conservation on the left, in positions −13 to −3, is from unaligned −10 patterns. As indicated by the arrow, transcription starts from base zero.

**Figure 2:**
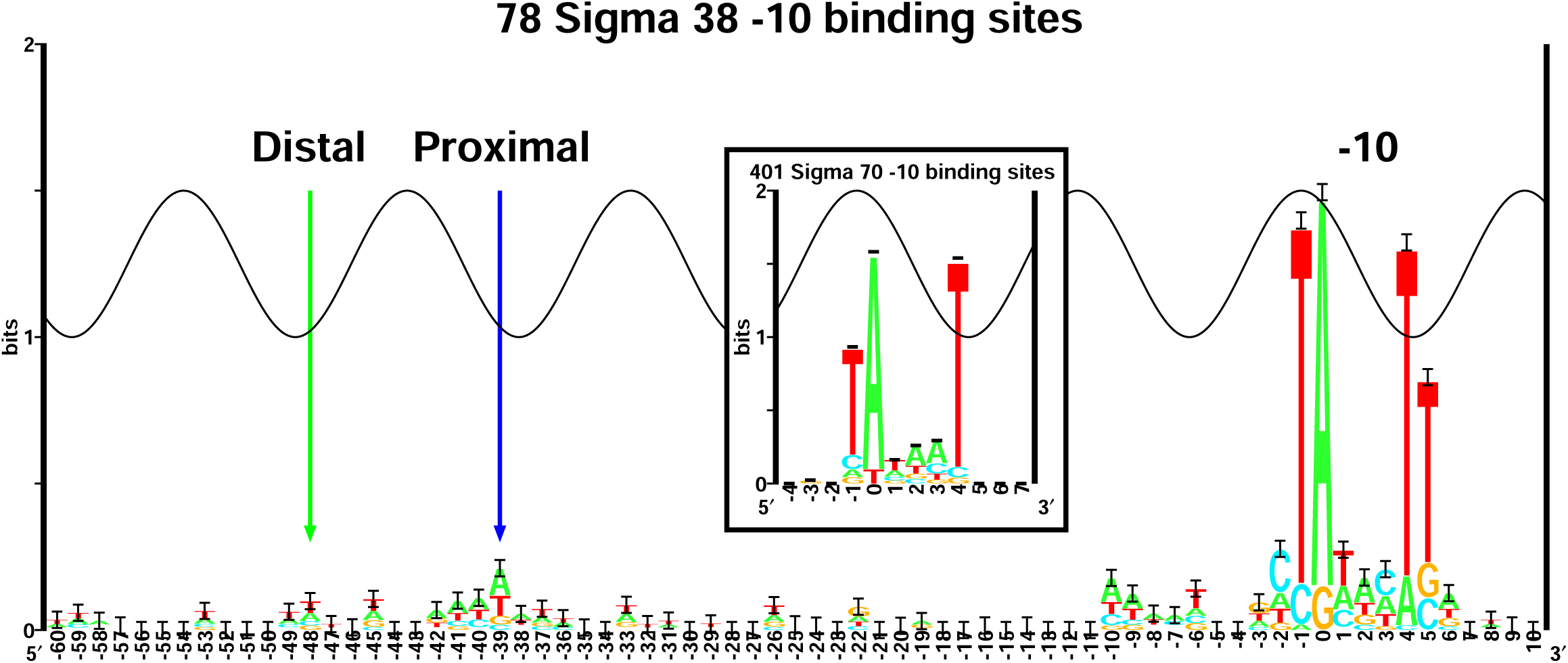
Alignment of *σ*^38^ −10 element shows lumps in the sequence logo upstream of the −10. The blue arrow labels a conserved lump at −39 and the green arrow labels the conserved lump at −48. The peak of the sine wave represents the major groove facing the protein and the phase of the wave for *σ*^38^ was chosen to match that of *σ*^70^ [4]. Inset: *σ*^70^ −10 from reference [4].

We took the best alignments made by the program **malign** and made sequence logos, which are visual representations of the average sequence conservation of a set of sequences [25, 34]. This helps us to understand the overall interaction between DNA and protein, but it does not allow us to understand the way a protein would interact with individual sequences. In contrast, a sequence walker graphically displays a single binding site. Sequence walkers present a model based on the conservation of the aligned sequences used to make the sequence logo [26]. This model can then be used to **scan** specific DNA sequences to visualize how the protein interacts with that sequence [27, 26].

The production of sequence walkers is an extension of the steps followed to make a sequence logo. The **ri** program uses the same set of aligned sequences to make a weight matrix of the binding sites, based on information theory [27]. **Scan** then uses the weight matrix produced by **ri** to evaluate the sequences at every possible position. Identified sites are then displayed as sequence walkers by **lister** [26]. RNA polymerase uses multiple binding sites that are separated by variable spacing, which means we have to **scan** multiple weight matrices that are separated by variable distances. For example, *σ*^70^ binds to two sites, the −10 and −35 elements [4]. To correctly locate binding sites on DNA sequences we have to use the information content for all of the parts that are used by the RNA polymerase. The total information content is then calculated by adding together the information content of each part and by subtracting the gap surprisal of the model. The gap is the distance between the parts, and it costs a certain amount of information [26]. After the multipart sites are located by **multiscan** they can then be visually represented as sequence walkers using the program **lister**.

On purpose, our numbering system is not the same as the conventional numbering system which records the number of bases between the −35 and −10 hexamers. For example ttgacaNNNNtataat would have a spacing of four. Instead, as described by Shultzaberger *et al.* [4], we assign the second base in both hexamers to be zero and the spacing is the difference between those two coordinates. By numbering this way, our spacing results in 6 bases greater than the conventional numbering used. In our numbering scheme, the previous example has a spacing of ten instead of four: tTgacannnntAtaat. Our zero-based numbering system allows us to analyze the sequences using sequence walkers [26, 4] because the location of binding sites is defined clearly and precisely by having the zero base inside the conserved region.

In this paper we introduce a new feature for multi-part sequence walkers. For example, in Fig. 4 there are four sequence walkers for the *σ*^70^ model: the distalUP, proximalUP, −35 (noted in this paper as ‘p35’) and −10 (noted in this paper as ‘p10’). Previously these were connected by horizontal bars that linearly interpolate the colors between the two colored rectangles (‘petals’) so that a viewer could tell which walkers were related by a flexible distance (see figures 5 and 6 of [4]). However, the viewer had to know that the end of the colored bar was associated with the zero coordinate of the corresponding sequence walker at the same location. The new feature consists of vertical extensions on these colored bars that complete the path. In Fig. 4, the rectangular petal for the distalUP is a deep blue and this walker is connected from the zero coordinate (light green rectangle) down to the colored Gap line and then up to the zero coordinate of the red p10 sequence walker petal.

**Figure 3:**
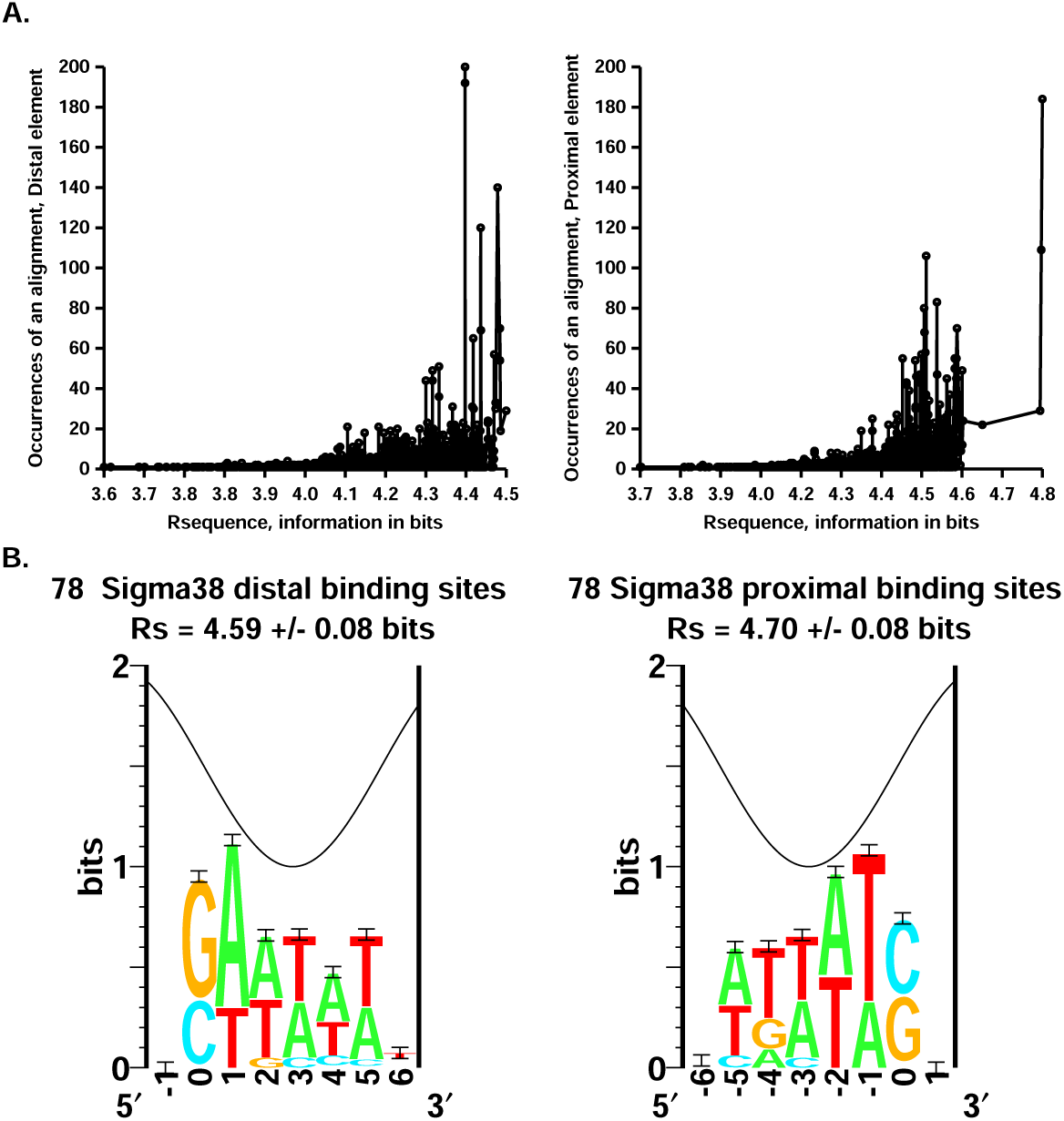
*σ*^38^ distal and proximal UP element sequence logos with a graphical representation of their multiple alignment. **A:** The graphs of occurrences of an alignment *versus* information of that alignment represent the distribution of 10, 000 random realignments of 78 *σ*^38^ binding sites to maximize their information content [33]. **B:** Sequence logos for the single best alignment of both the proximal and distal elements. The logos have dyad symmetry with respect to each other since both have C/G bases at one end followed by A/T rich regions. The bottom of the sine wave indicates the center of the minor groove assigned from NMR [60] and x-ray crystal structures [67, 73].

**Figure 4:**
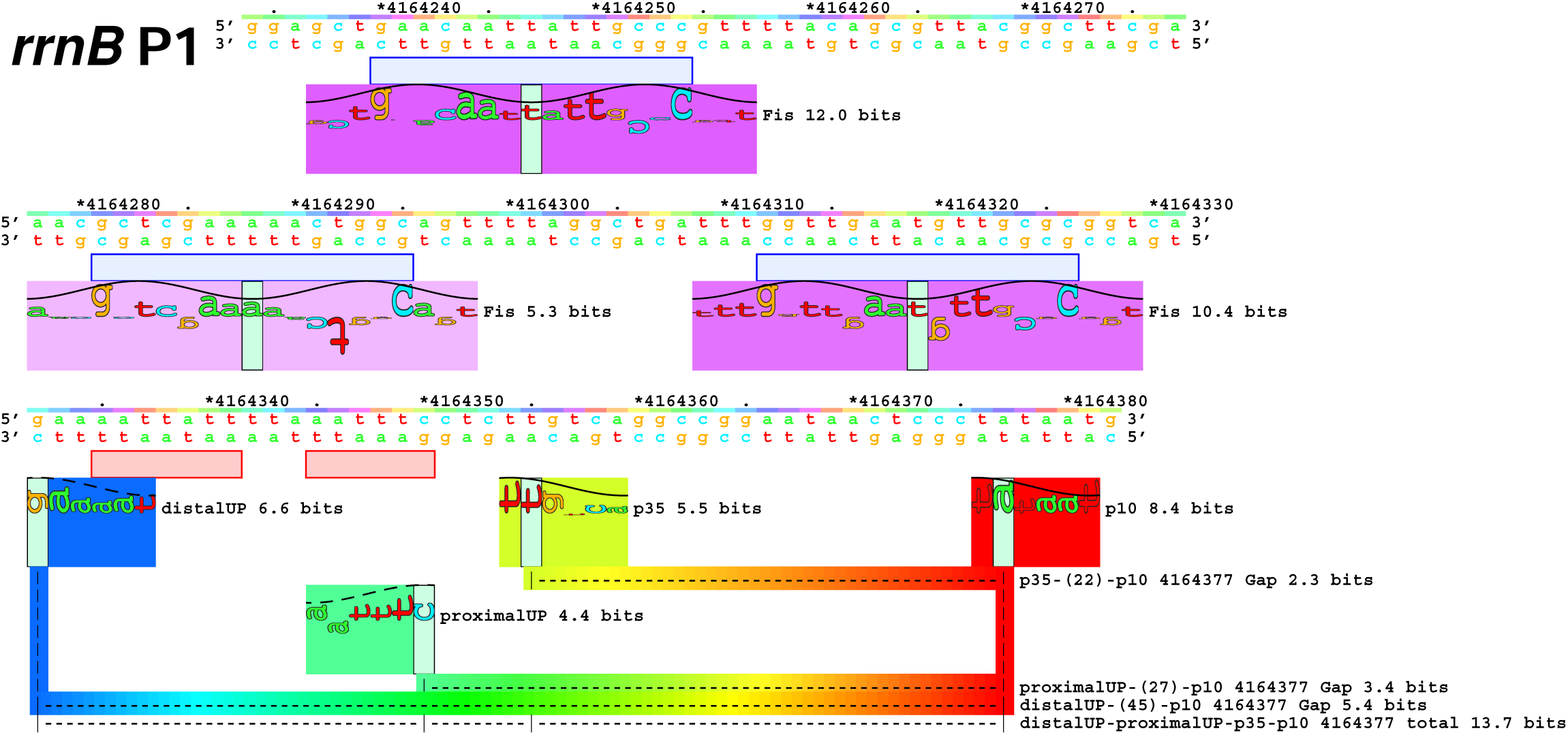
Sequence walkers of a flexible *σ*^70^ model with UP element patterns and Fis **scan**ned on the ribosomal promoter *rrnB* P1. A sequence walker graphically represents a single binding site in which the height of each letter indicates the information content at that position [26]. The colored rectangles, behind the sequence walker letters, called ‘petals’, indicate the kind of site by hue. The strength of the binding site is indicated by the petal’s saturation. The label to the right of each rectangle gives the kind of site and the bits of information of the individual site. The light blue rectangles are areas where the transcription factor Fis was located on the *rrnB* P1 promoter by DNase I footprinting [47, 48]. They match our Fis model [49] exactly (purple walkers). The pink rectangles are areas where UP elements were located on the *rrnB* P1 promoter by DNase I and hydroxyl radical footprinting [46]. The multi-part flexible sequence walker visualizes the locations of the UP elements, −35 and −10 that *σ*^70^ binds to. To indicate the relationships there are connecting bars that shade from the color of one site to the color of the other. The dashed line within the colored bridges has a label to the right that names the two kinds of sites, the distance between the two and the gap surprisal for the distance between the two sites. The dashed line labeled ‘total’ gives the kind of sites, the coordinate of the farthest downstream site, and the total information for the entire site found by the flexible model. The red rectangle represents the −10 element and the yellow one represents the −35 element found on the ribosomal *rrnB* P1 by *σ*^70^. The green rectangle represents what we hypothesize to be the proximal UP element and the blue rectangle represents the distal UP element. The UP elements models came from the *σ*^38^ dataset (Fig. 3).

**Figure 5:**
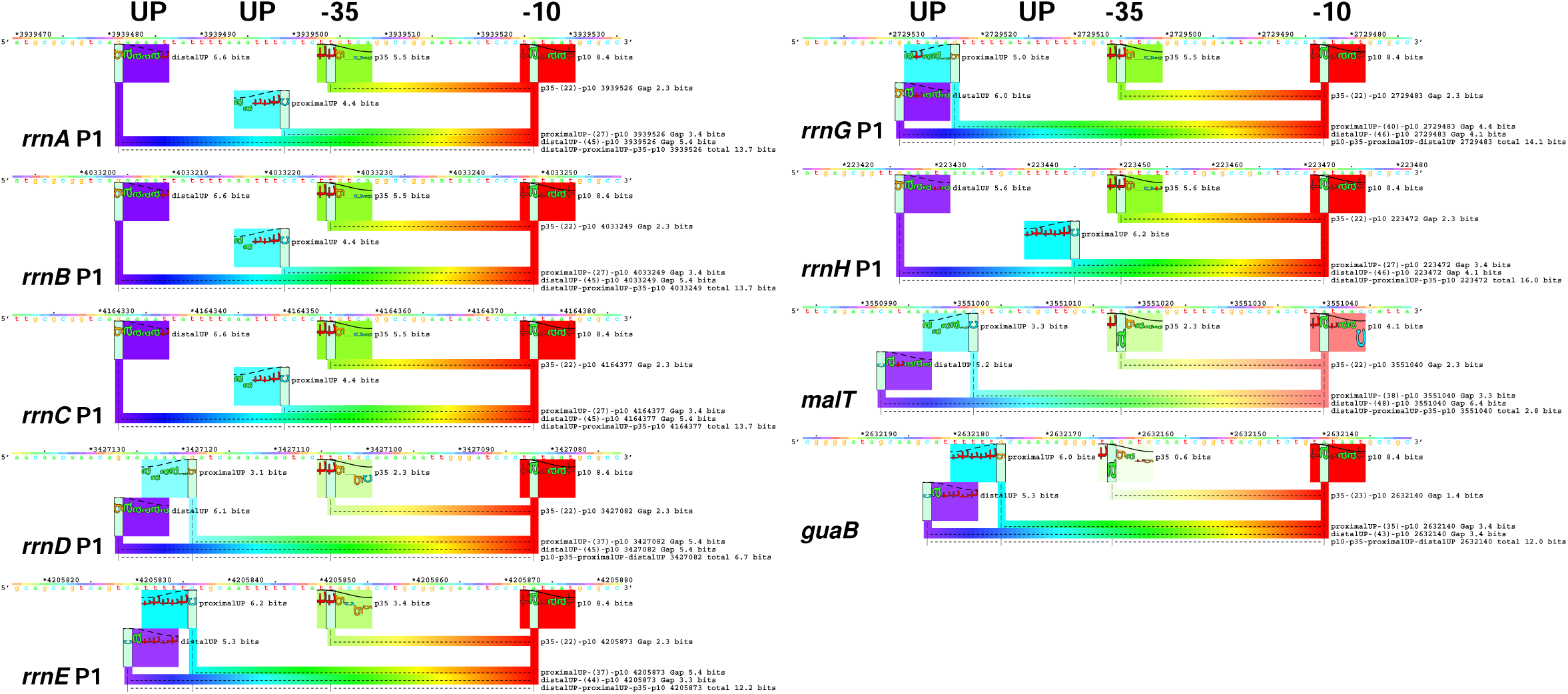
Sequence walkers of *σ*^70^ (green: −35 and red: −10) and validation that the information theory models correctly identify UP elements (distal: magenta and proximal: cyan) for *rrnA* P1, *rrnB* P1, *rrnC* P1, *rrnD* P1, *rrnE* P1, *rrnG* P1 *rrnH* P1, *guaB*, and *malT* [48].

**Figure 6:**
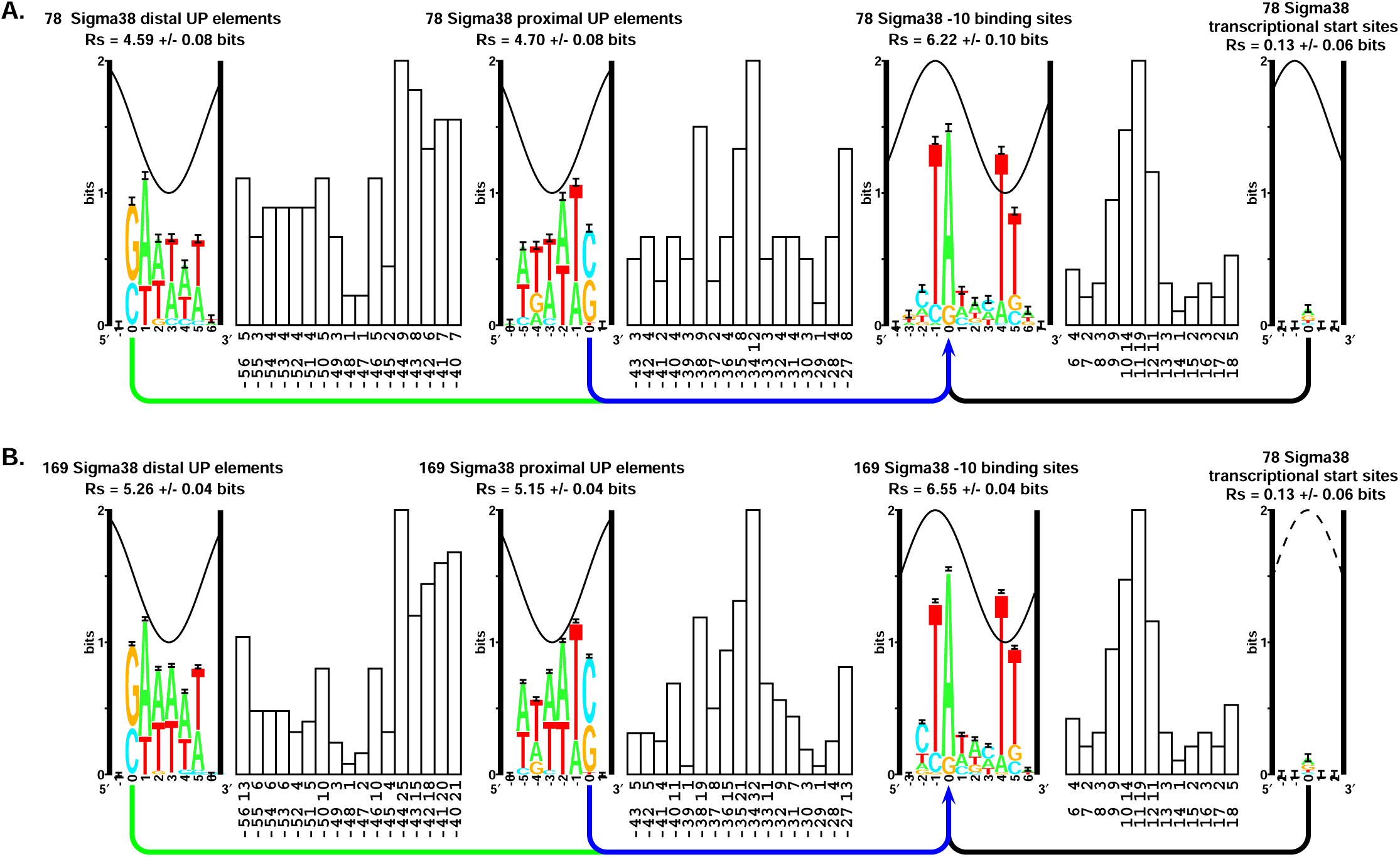
Sequence logos for *σ*^38^ promoter models. **A:** Sequence logos for 78 *σ*^38^ promoters that have experimentally proven transcriptional start sites. From right to left: sequence logo of the transcription start sites, spacing distribution from transcription start sites to the −10 binding sites, sequence logo of the −10 binding sites, spacing distribution from −10 binding sites to the proximal UP element, sequence logo of the proximal UP element binding sites, spacing distribution from −10 binding sites to the distal UP element binding sites, and the sequence logo of the distal UP element binding sites. **B:** Sequence logos for 169 experimentally proven *σ*^38^ promoters. From right to left: sequence logo of the transcription start sites (only 78 sites), sequence logo of the −10 binding sites, spacing distribution from −10 binding sites to the proximal UP element, sequence logo of the proximal UP element binding sites, spacing distribution from −10 binding sites to the distal UP element binding sites, and the sequence logo of the distal UP element binding sites. Because the information in the start logo (0.13 bits) is less than the uncertainty of the start histogram (3.2 bits) [4] we did not use a transcription start sites component for the 169 based model. The 10.6 base helical twist of B-form DNA [74] is represented by sine waves on each logo. The peak represents the major groove facing the binding protein [4]. The sine wave for the binding sites has the minor groove positioned at −2.5 for the proximal and +2.5 for the distal UP element because it was identified through DNase I and hydroxyl radical protection experiments [46, 47, 56] and an X-ray crystal structure [73] to be bound to the minor groove. The eight logos have variable spacing to the −10 indicated by the bar graphs. Below each bar graph are the distance between the zero coordinate of the sequence logos and the number of cases observed.

In our previous work on ribosome binding sites [31] and *σ*^70^ promoters [4] we showed that the total information of a multi-part binding site can be computed from the sum of the individual information of each binding site part followed by subtraction of the cost of the ‘gaps’ between the parts. For example, for the *rrnB* P1 promoter shown in Fig. 4, the total information of 13.7 bits is found by adding the distalUP (6.6 bits), proximalUP (4.4 bits), p35 (5.5 bits) and p10 (8.4 bits) and then subtracting the three gap costs (2.3, 3.4 and 5.4 bits). In this case the numbers sum to 13.8 which is within rounding error.

### *In vitro* transcription assay

The *rrnB* P1, *bolA* P1 promoters and the derived mutants were obtained by PCR and overlap PCR from *E. coli* MG1655 genomic DNA to introduce mutations (see supplementary materials). Then these promoter sequences were used to replace the EcoRI to HindIII fragment of plasmid pRLG1617 [35, 36], a gift by Wilma Ross and Richard Gourse, to make pDJ plasmids. Supercoiled templates were used for *in vitro* transcription assays. *E. coli* RNA polymerase core enzyme, *α*CTD truncated core enzyme, *σ*^38^, *σ*^70^ and Fis were purified as previously reported [37, 38, 39, 40, 41]. Then the corresponding core enzymes were mixed with either *σ*^38^ or *σ*^70^ with a core to *σ* molar ratio of 1 : 3 at 37°C for 20 min to reconstitute the holoenzymes E*σ*^38^, E*σ*^38^Δ*α*CTD, E*σ*^70^ and E*σ*^70^Δ*α*CTD [41]. For *in vitro* transcription, 20 nM reconstituted holoenzymes were mixed with 3 nM supercoiling plasmids in 5× transcription buffer (200 mM Tris-HCl at pH 8.0, 5 mM dithiothreitol, 0.5 mg/mL purified calf BSA, 50 mM MgCl_2_ and 50 mM KCl). After pre-incubating at 37°C for 15 min, the reactions were started by adding 5× NTP mixture (1 mM for ATP, CTP and GTP, 0.1 mM for UTP) containing 2 µCi of [*α*−^32^P] UTP (PerkinElmer). Transcription reactions were performed at 37°C for 20 min. Then the reactions were terminated by adding equal volume of stop buffer (250 mM EDTA, pH 8.0, 10 M Urea, 0.05% xylene cyanol and bromphenol blue). The samples were run on 8% sequencing gel (National Diagnostics) and visualized by phosphorimager analysis.

## 4 Results

### The *σ*^38^ model

Although RegulonDB contained 134 transcription start sites for *σ*^38^ [32], for our initial analysis we chose only the 78 start sites that were precisely defined by transcription initiation mapping. We aligned the start sites and made a sequence logo to calculate the information content. The transcription start sites only contained 0.13 ± 0.06 bits of information within the range −2 to +2 [22]. This is shown as a sequence logo [25] in Fig. 1. We saw a small preference for adenine and guanine at the transcription start.

Next, using the program **malign** [33], we shuffled the sequences (slid them back and forth) to maximize the −10 region information in the range −13 to −7 upstream of the transcription start. We shuffled up to 6 bases in either direction. Unlike *σ*^70^ −10 sites, which contain 4.78 ± 0.11 bits [4], in the range −4 to −7 the *σ*^38^ −10 information was 6.22 ± 0.10 bits (Fig. 2).

A recent X-ray crystal structure of *σ*^38^ bound to DNA in an initiation complex [13] showed that the structure is nearly identical to that of *σ*^70^. Indeed, the aligned *σ*^38^ −10 logo (Fig. 2) partially resembles the *σ*^70^ −10 logo [4] (Fig. 2 inset). The *σ*^38^ logo has a strongly conserved T at position +4, similar to the strongly conserved T from the *σ*^70^ logo. RNA polymerase is facing the minor groove there, implying that this position is being flipped out of the helix [42], an information theory prediction that has been confirmed experimentally [43, 44]. In logos for DNA binding proteins [42], positions conserved in the minor groove generally do not exceed an information content of 1 bit because a protein cannot distinguish all 4 bases from each other in the minor groove of B-form DNA; it can only differentiate A and T from C and G [45]. The T that is strongly conserved in our logo at position +4 has an information content that exceeds 1 bit of information, consistent with base flipping [4]. Other similarities are the strongly conserved T at position −1 and the strongly conserved A at position 0. On *σ*^38^, there is a strongly conserved T at position +5 that is not found on the logo for *σ*^70^. This extra T is said to favor *σ*^38^ because of the possibility of it making promoter melting easier [12] so it may also involve base flipping. In addition there is frequently a C at position −2 in our logo; this conserved C improves transcription by *σ*^38^ over *σ*^70^ [11].

Once we had obtained the −10 alignment we then attempted to repeat the same steps to locate a pattern in the −35 region. The optimal spacing for *σ*^70^ is 23 bases relative to the −10, and *σ*^38^ has been said to utilize spacer lengths that differ by 1 − 2 base pairs from that [11]. It has been proposed that *σ*^38^ can use a degenerate form of the −35 element, that it can function without a −35, or that it might recognize a different −35 element [12]. To locate the −35 we aligned the 78 RegulonDB sequences in the region where the −35 is said to be found [11]. Using **malign** we computed the information content over the range of −30 to −20 bases relative to the −10 element, set the shift window to shuffle up to 5 bases upstream and 7 bases downstream, and set **malign** to redo the alignment 1,000 times. We found a pattern having an information content of 3.71 ± 0.09 bits. In random sequence 5 + 7 + 1 bases wide we expect patterns of *R*_*f requency*_ = log_2_ 13 = 3.70 bits [22]. In comparison, the −35 element of *σ*^70^ has an information content of 4.02 ± 0.09 bits [4]. To test whether the information of the observed −35 pattern is significantly above noise, we used the markov program to generate 78 random sequences that had the composition of the *E. coli* K-12 MG1655 genome (GenBank Accession NC000913 version 2) and realigned them 1000 times by shuffling over 13 positions to maximize the information in a window 11 bases wide. This matches the parameters of the natural alignment attempt. We repeated this process 100 times and found that the best alignments were 3.74 ± 0.07 bits. Since the observed sequence pattern of 3.71 bits is 0.4 standard deviations below that produced by this Monte Carlo simulation, the pattern found by aligning the −35 *σ*^38^ region is not significant.

Next, we went back to the sequence logo of the 78 RegulonDB sequences aligned by the −10 element; when we increased the range to −100 to +10, we noticed possible patterns upstream that appeared as small lumps of conservation located −48 and −39 bases upstream from the −10 element (Fig. 2). We aligned the 78 RegulonDB sequences in the proximal −39 region around the first lump. Using **malign** we computed the information content over the range of −40 to −35, set the shift window to shuffle up to 9 bases in either direction, and set **malign** to redo the alignment 10,000 times. We found a pattern having an information content of 4.70 ± 0.08 bits (Fig. 3B, right side). In random sequence 2 × 9 + 1 wide we expect patterns of *R*_*f requency*_ = log_2_ 19 = 4.25 bits [22]. In comparison, the −35 element of *σ*^70^ has an information content of 4.02 ± 0.09 bits [4]. To test whether the information of the observed proximal pattern is significantly above noise, we used the **markov** program to generate 78 random sequences that had the composition of the *E. coli* K-12 MG1655 genome (GenBank Accession NC000913 version 2) and realigned them 1000 times by shuffling over 19 positions to maximize the information in a window 6 bases wide. We repeated this process 100 times and found that the best alignments were 3.93 ± 0.08 bits. Since the observed sequence pattern of 4.7 bits is 9.72 standard deviations higher than produced by this Monte Carlo simulation, the pattern found by aligning the proximal *σ*^38^ sequences is highly significant.

Likewise, for the distal −48 region around the second lump, over the range of −49 to −42 with a shift window up to 8 bases in either direction and 10, 000 alignments, we found a 4.59 ± 0.08 bit pattern (Fig. 3B, left side). To test the significance, 78 random sequences were realigned 1000 times by shuffling over 17 positions with a 6-base wide window. For 100 repeats the best alignments were 3.94 ± 0.08 bits which means that the observed pattern is 8.14 standard deviations above the noise. Similar to the proximal site, the pattern found by aligning the distal *σ*^38^ sequences is also highly significant.

Curiously the two patterns at −39 and −48 have inverted dyad axis symmetry sequences (Fig. 3). The patterns found through this procedure are too far upstream to be a conventional −35 element. Because these elements are upstream of the putative −35 region, we hypothesized that they represent UP elements defined by Newlands *et al.* [46]

### UP elements on Ribosomal RNA Promoters

To determine whether the logos we had found represent UP elements, we **scan**ned each putative UP element model attached to the current flexible two-part *σ*^70^ model for sites > 0 bits, using the programs **scan** and **multiscan**, on the promoter *rrnB* P1. *rrnB* P1 is the best characterized ribosomal RNA promoter known to have UP elements. On the *rrnB* P1 promoter the UP elements are located −60 to −40 bases upstream from the transcription start inbetween binding sites for the transcription factor Fis and the −35. UP elements were found by Rao *et al.* [47] to cause increased promoter activity by more than 30-fold [46, 6, 47, 48].

After the **scan** and **multiscan** programs have identified sites, the **lister** program displays their results in the form of sequence walkers that show the individual information content of each base in a binding site by the different heights of the letters. The heights of the letters correspond to bits of information [26, 27]. **Scan** identified the three Fis sites on *rrnB* P1 (Fig. 4) [49] and found that they match DNase I footprints exactly [48]. Using the *σ*^70^ UP model with attached putative UP elements, **multiscan** correctly identified the multiple locations of the *σ*^70^ −35 and −10 (Fig. 4). The *σ*^70^ UP model also uniquely identified locations of distal and proximal UPs in the region of DNase I and hydroxyl radical protection by *α*CTD [46, 47]. These sites are displayed as four linked sequence walkers, one for each of the elements. Two correspond to the *σ*^70^ −10 and −35 elements, and the other two to the proximal and distal UP elements. Thus our approach has correctly identified the known elements of *σ*^70^. These results indicate that the significant sequence elements we found upstream of the −35 region of *σ*^38^ promoters correspond to UP elements.

To further test whether the upstream logos represent UP elements we investigated a bigger set of sequences containing proven UP elements. The combined *σ*^70^ UP model again uniquely identified locations of distal and proximal UPs exactly in the region of DNase I and hydroxyl radical protection by *α*CTD on the seven *rrn* operons [46, 47, 48] and the genes *guaB* and *malT* [50, 51] (Fig. 5).

A DNase I protection experiment using the *aidB σ*^38^ promoter showed hypersensitivity −40 and −50 bases upstream of the transcriptional start (their Fig 5) [52]. These correspond to the −35 and both UP elements respectively (supplementary material).

### Building a complete *σ*^38^ model

We then constructed a three-part flexible model of *σ*^38^ consisting of the −10 element, the proximal UP element, and the distal UP element (Fig. 6A). The distances between the flexible parts of the model were calculated by scanning the *σ*^38^ −10, the proximal UP and the distal UP elements on the 78 proven *σ*^38^ genes and locating the coordinates of the individual scanned parts on each gene. We scanned our combined model on the original 78 site data set and found that it was able to locate 72 out of the 78 sequences (> 0 bits). We used a zero bit cutoff because, according to the second law of thermodynamics, sequences that have an information content less than zero should not be binding sites because they have a positive ΔG of binding [23, 27, 4].

To test whether the model represents *σ*^38^, we investigated a bigger dataset for sites of *σ*^38^ promoters. Weber *et al.* [8] performed a genome-wide expression profiling of *Escherichia coli* that located 140 positively *σ*^38^ controlled genes under the growth and stress conditions of osmotic upshift, stationary phase and acid stress. We multiscanned our flexible *σ*^38^ model on this set of 140 genes for sites > 0 bits, and located 139 sites out of the 140 genes (> 99%). This finding further confirmed that our model does identify *σ*^38^ binding sites that consist of a −10, proximal and distal UP elements.

To construct a model based on more sequences, we took the *σ*^38^ 78 gene model and scanned it on a new dataset made up of the RegulonDB 78 *σ*^38^ data set of proven sites combined with the 140 dataset from Weber *et al.* to form a new dataset of 169 positively *σ*^38^ controlled genes (49 sites are shared in both sets). We took the strongest site at each gene to make a new 169 site model. To confirm that the new *σ*^38^ model still located *σ*^38^ sites, we scanned the new model on the 169 genes; it was able to locate 166 sites (98%). We used the 169 site model from this point on (Fig. 6B). The 169 model and the scans are shown in the supplementary materials.

The individual information distribution of the 169 site *σ*^38^ model had a mean *R*_*sequence*_ = 8.40 bits/site with a standard deviation of 2.76 bits/site and standard error of the mean (sem) of 0.21 bits/site. There are γ = 169 known *σ*^38^ sites and two possible orientations of the polymerase on each base of the double stranded DNA of the *E. coli* K-12 MG1655 genome, giving *G* = 2 × 4639675 ways for the polymerase to sit on the DNA before binding. Therefore the information needed to locate these sites is *R*_*f requency*_ = log_2_ *G*/γ = 15.74 bits/site [22, 53]. This is significantly higher information than our model contains. An unlikely possibility is that there are *G*/2^8.40±0.21^ = 27470 ± 4000 sites yet to discover. Alternatively, the model may be missing part (15.74 − 8.4 = 7.4 bits) of the promoters or many promoters are being suppressed by factors such as H-NS [54].

### Comparing the parts of *σ*^38^ promoters with those of *σ*^70^ promoters

In the this section, we compare each of the components of *σ*^38^ promoters with those of *σ*^70^ promoters.

#### Distal and Proximal UP elements are Equivalent

The proximal and distal UP element sequence logos shown in Fig. 3 and Fig. 6 appear to be inverted sequences with respect to each other. We did several tests to determine if they are the same. First, for the 169 site model we compared the number of bases at each position in the distal sites (positions 0 to +5) to the numbers of the corresponding dyad symmetry proximal sites (positions 0 to −5) and found that they are linearly related with a correlation coefficient of 0.98 (Fig. 7). This explains why the logos appear similar.

**Figure 7:**
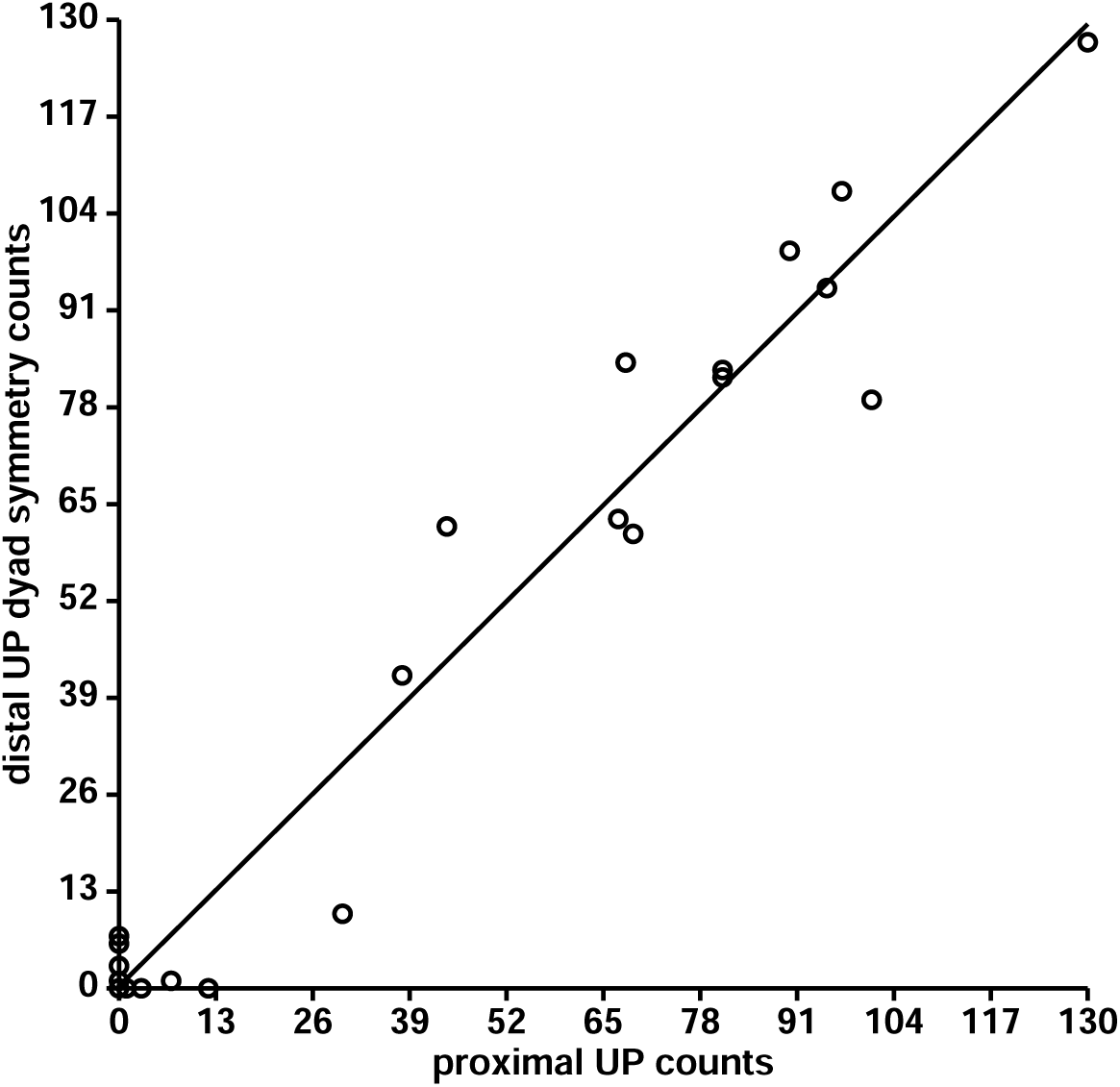
Comparison of distal and proximal UP element base counts. The 24 base counts for the 6 positions of the distal *vs.* proximal UP element sequence logos of Fig. 6B were plotted against each other. The correlation coefficient is *r* = 0.98 and *r*^2^ = 0.95 so 95% of the variation of the distal UPs is explained by the variation of the proximal UPs. The regression line is: distal UP dyad symmetry counts = 0.99 × proximal UP counts + 0.27.

Then we evaluated the individual information content of proximal sites using both the proximal and distal UP models; these had a correlation coefficient of 0.92. Likewise, distal sites evaluated with both the distal and proximal UP models had a correlation coefficient of 0.89.

To compare the total information content (area under the sequence logo, *R*_*sequence*_, [22]), of the two UP elements we performed a two-tailed Student’s *t*-test between the individual information distributions and found that they are the same (*p* = 0.76). Since the average of the individual information distributions is *R*_*sequence*_, the information contents of the two UP elements are identical.

All of these results indicate that the two models are indistinguishable, so we combined the proximal and distal UP models (Fig. 8) and then did Student’s *t*-tests between the individual information distributions of the proximal *vs.* combined UP model (*p* = 0.64) and the distal *vs.* combined UP model (*p* = 0.86). Therefore the 338 site combined UP model information content is the same as that of the distal and proximal UP models.

**Figure 8:**
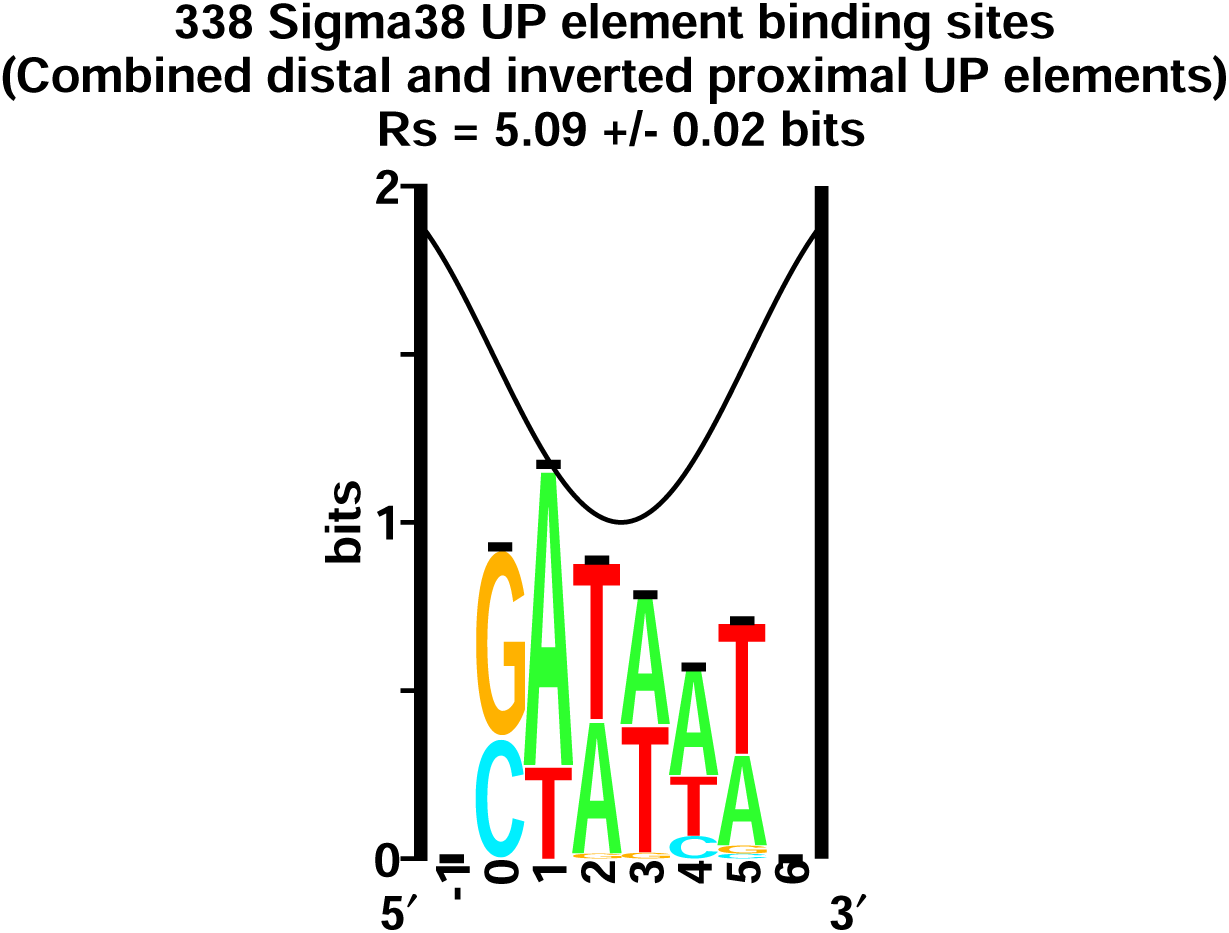
Sequence logo of the 169 sequences of both proximal and distal UP elements (Fig. 6B) were combined.

#### Absence of −35 on *σ*^38^ promoters

It has been proposed that *σ*^38^ uses a −35 to promote transcription [10], but multiple alignment to maximize information in that region did not locate a −35. In previous attempts others were also unable to locate a −35 [18, 11]. We decided to take a different approach to locate a potential −35.

We used two datasets, one dataset contained the proven RegulonDB *σ*^70^ sites. Our second dataset was of the 169 positively controlled *σ*^38^ genes. First, we scanned the −35 weight matrix from the *σ*^70^ flexible model [4] on the 1784 *σ*^70^ transcription start sites from RegulonDB within a range of −100 to −10 bases upstream. This first scan revealed that a majority of the −35 sites found were located 35 bases upstream with an information content between 5.5 and 6.5 bits of information (Fig. 9A). This result was expected as we already know that *σ*^70^ controlled genes use a −35 to promote transcription. We used the same −35 weight matrix to scan the 169 genes that are *σ*^38^ controlled. The scan did not locate a high density of −35 sites for the 169 genes (Fig. 9B). Most notably in the colored density plot there are no sites within the range of 5.5 to 6.5 bits of information compared to the approximately 130 sites that were found on the *σ*^70^ colored density plot. We should have seen about 130(169/1784) = 12 sites.

**Figure 9:**
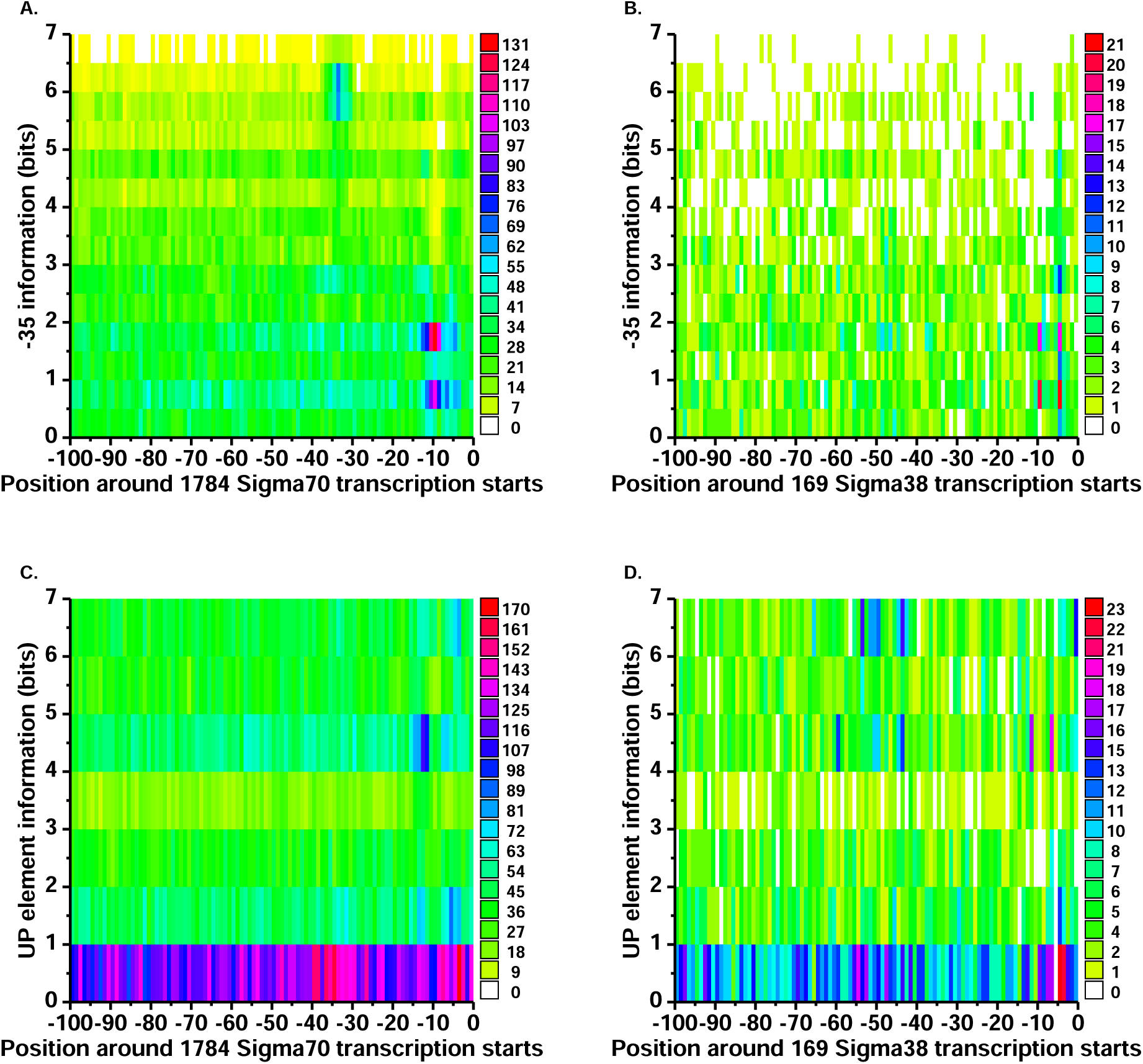
Scans of −35 (top) and UP (bottom) element models on *σ*^70^ (left) and *σ*^38^ (right) promoters. **A:** −35 model from *σ*^70^ sites scanned on *σ*^70^ promoters. We scanned the RegulonDB dataset of 1784 proven *σ*^70^ sites 100 bases upstream of the transcription start sites using the −35 weight matrix of our *σ*^70^ model [4]. The −35 model located a cluster of −35 sites approximately 35 bases upstream from the transcription start sites with an information content of 5.5 to 6.5 bits. The weak sites in the −10 region were caused by overlapping patterns between the −35 model and the sequences at that location. The colored legend on the side shows the number of sites found at every base for every half bit. The spectrum goes from white to red, white being zero sites to red being 131 sites. The vertical axis represents the information content of the located −35 sites between 0 and 7 bits. The horizontal axis represents the bases upstream from the transcription start sites at zero. **B:** −35 model from *σ*^70^ sites scanned on *σ*^38^ promoters. As in part A, the sites near the −10 region were caused by overlapping patterns between the −35 model and the sequences at those locations. At the location where the cluster was found on part A there is no −35 pattern on this graph. **C:** UP element model consisting of both proximal and distal UP elements from *σ*^38^ sites scanned on *σ*^70^ promoters. We scanned the RegulonDB dataset of 1784 proven *σ*^70^ sites 100 bases upstream of the transcription start sites using the combined UP element weight matrix. The combined UP element model located weak sites in the −10 region, caused by overlapping patterns of the UP elements and the patterns at that location. There is no cluster of UP element patterns on this graph at the UP element locations. **D:** UP element model consisting of both proximal and distal UP elements from *σ*^38^ sites scanned on *σ*^38^ promoters. As in part C, the weak sites near the −10 were caused by overlapping patterns of the UP elements and the patterns at that location. At the location in part C where we were unable to locate many UP element patterns, the combined UP element model located a cluster of UP element sites approximately between 40 and 50 bases upstream from the transcription start sites with an information content of 4 to 7 bits. Note that in (C) and (D) the red, purple and blue lines below 1 bit represent noise from sequences with low information content.

Since we were unable to locate −35 sites in the 169 set of *σ*^38^-controlled genes, we hypothesized that *σ*^38^ may have lost the ability to locate a −35 as the UP elements evolved. To test this hypothesis we repeated the scanning of the −35 weight matrix on the 169 positively *σ*^38^ controlled genes, but this time we lowered the cutoff to -20 bits. If our hypothesis were correct then we might be able to locate a high density of −35 sites 35 bases upstream but well below 0 bits of information. Our scan revealed nothing within the area of −35 that appeared to have a higher density then the rest of the range (Fig. 10).Furthermore, there was no overlap of start points between *σ*^70^ and *σ*^38^sites in RegulonDB promoters so the −35s of *σ*^70^ sites could not be contributing to putative *σ*^38^−35s. Our results indicate that *σ*^38^ does not use a −35 in natural *σ*^38^ controlled promoters.

**Figure 10:**
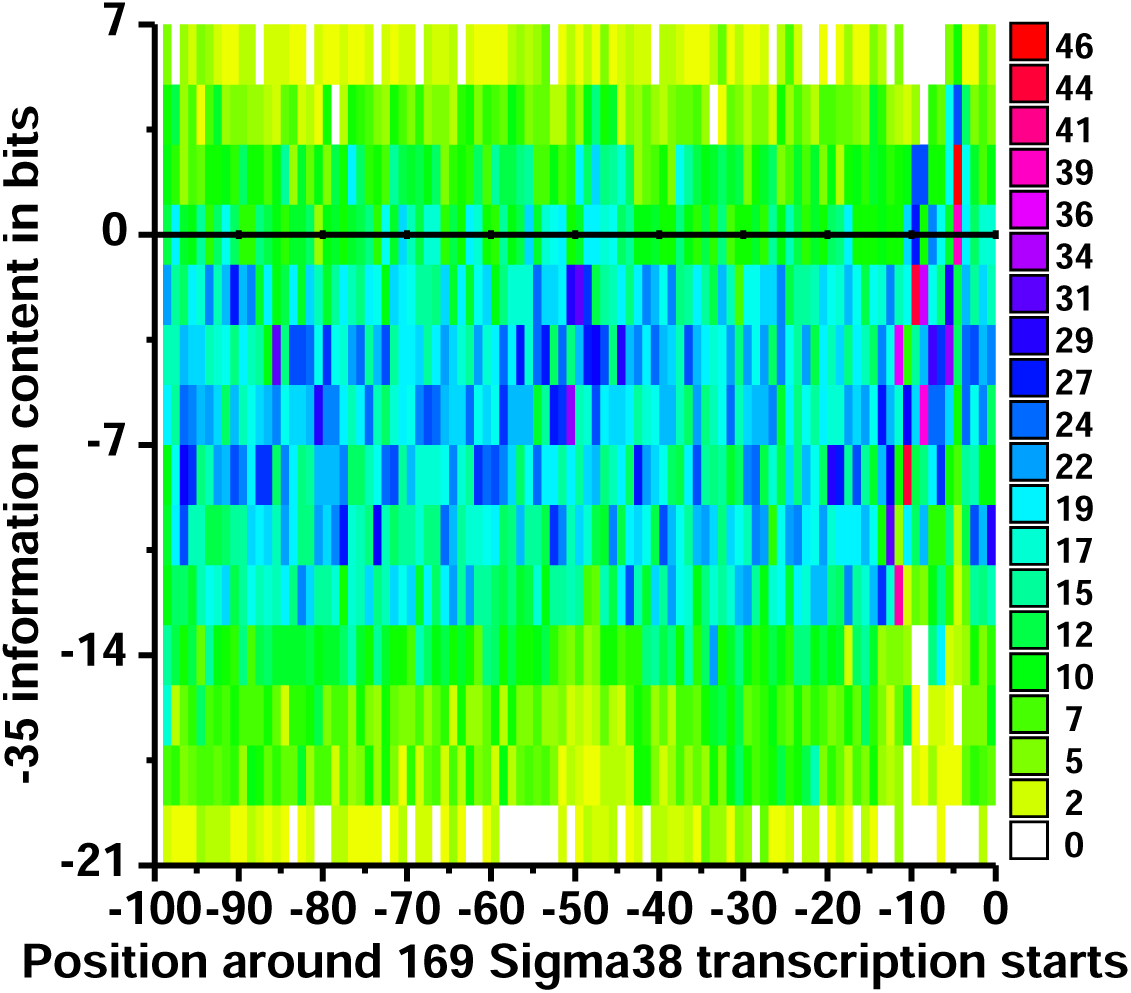
*σ*^70^ −35 model scanned on 169 *σ*^38^ controlled genes to show information below zero bits. No cluster of −35 sites is detectable. This figure is an extension of figure Fig. 9B.

#### Absence of UP elements on most *σ*^70^ promoters

Proven UP element binding locations on the seven *rrn* operons are shown in Fig. 5 [46, 47, 48]. They show that *σ*^70^ can use UP elements. To test whether or not *σ*^38^ uses UP elements more frequently than *σ*^70^, we used the same density method as above to test for the presence of UP elements. To do this, we scanned the combined UP element weight matrix (Fig. 8) on the 1784 *σ*^70^ transcription start sites from RegulonDB within a range of −100 to −10 bases upstream. This first scan did not locate a high density of UP element sites for the *σ*^70^ controlled genes in the place where the UP element sites are found on *rrn* (Fig. 9C). We used the combined UP element weight matrix to scan the 169 genes that are *σ*^38^ controlled. The second scan revealed that a majority of the UP element sites were located 40 to 50 bases upstream with an information content between 4 and 7 bits (Fig. 9D). Overall, Fig. 9 shows that *σ*^70^ promoters have −35s (A) but essentially no UPs (C), while *σ*^38^ promoters have UPs (D) but no −35s (B).

#### −10 elements of *σ*^38^ and *σ*^70^ are different

Because the −10 sequence logos are similar (Fig. 2), but *σ*^38^ has extra conservation on each end, we expect that the *σ*^70^ −10 model may be able to identify *σ*^38^ −10 elements. To determine the relationship between these two models we scanned both across *σ*^70^ and *σ*^38^ promoters (Fig. 11). Fig. 11A shows that on *σ*^70^ promoters the −10 from *σ*^70^ has a higher information content than the −10 from *σ*^38^. Fig. 11B shows that on *σ*^38^ promoters the −10 from *σ*^38^ has a higher information content then the −10 from *σ*^70^. In other words, each model will accept the other, but each *σ* prefers its own sequence element. These results confirm that the −10 elements of *σ*^38^ and *σ*^70^ differ in such a way such that they can start transcription for their corresponding promoters even though the elements appear similar and that our flexible *σ*^38^ model represents *σ*^38^ sites.

**Figure 11:**
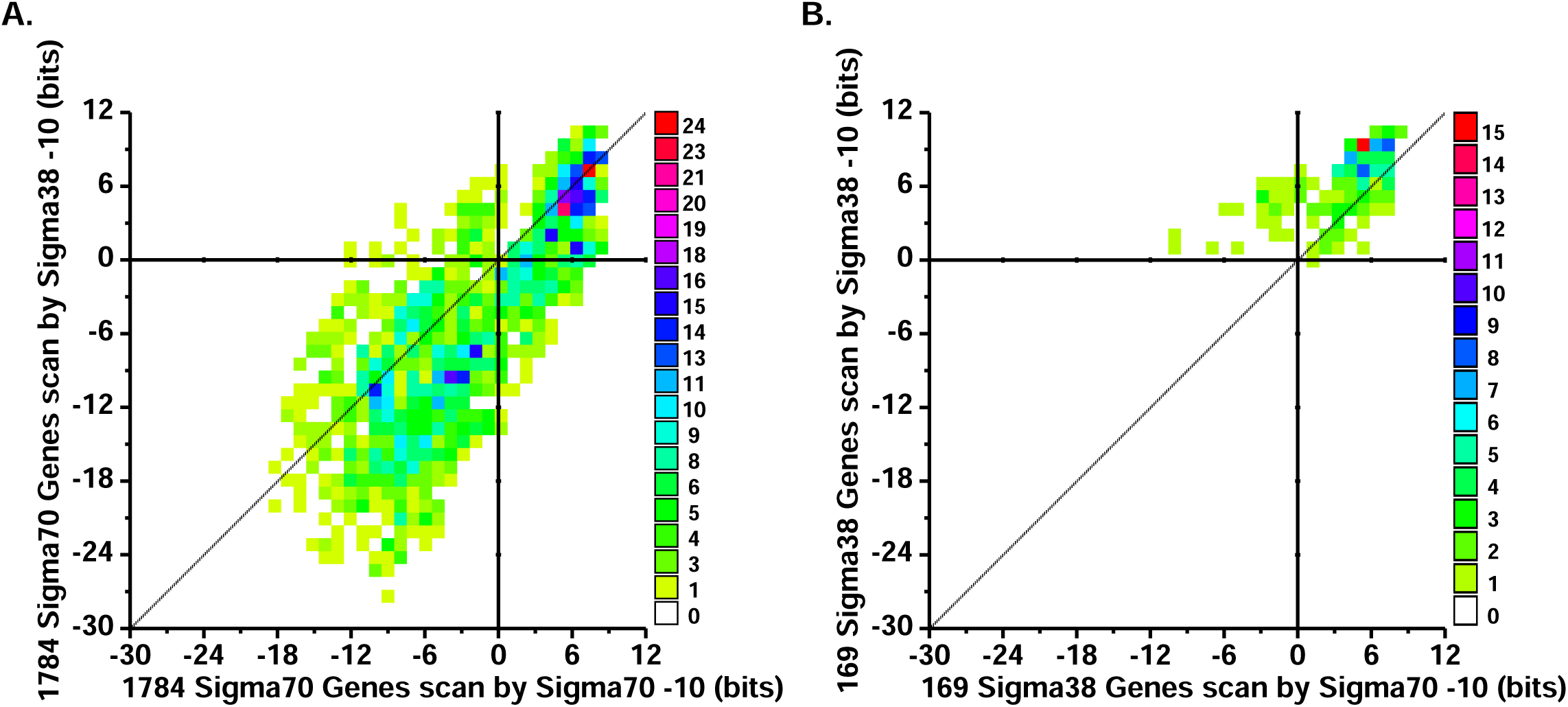
Scanning −10s of *σ*^70^ and *σ*^38^ models on *σ*^70^ and *σ*^38^ promoters. Each graph is a colored density plot in which the individual counts are placed in a small rectangle and the color designated to the rectangle depends on the total individual counts inside the rectangle. In these graphs the colors vary from white being the no counts to red being the highest number of counts. Inbetween these two is the rainbow spectrum representing the different total counts. Each graph has four quadrants and the Y and X axis that divide the quadrants are represented in thick black lines. The diagonal line represents what the trend would be if there were no difference in information content between the *σ*^70^ and *σ*^38^ −10 scans. The Y axis on both graphs represents the information content of the sites found with the −10 weight matrix from *σ*^38^ and the X axis represents the information content from *σ*^70^. **A:** The graph on the left represents scanning the −10 weight matrices from both *σ*^70^ and *σ*^38^ on 1784 *σ*^70^ promoters and plotting the information contents against each other. **B:** The graph on the right represents scanning the −10 weight matrix from both *σ*^70^ and *σ*^38^ on 169 *σ*^38^ promoters and plotting the information contents against each other.

### Analysis of the *bolA* P1 promoter

Our 169 site *σ*^38^ model (Fig. 6B) and several analyses (Figures 9A, 9B, 10 and 11) show that *σ*^38^ does not need a −35 element to function. In contrast, previous work by Gaal *et al.* [10] on *σ*^38^ using *in vitro* selection on the promoter *bolA* P1 had identified −10 and −35-like sequences similar to those of *σ*^70^. Nguyen *et al.* [55] ran footprinting experiments on the *σ*^38^ dependent *bolA* P1 promoter and identified protection sites for both *σ*^38^ and *σ*^70^ on the nontemplate and template strands. Our model was able to locate a 6.4 bit *σ*^38^ site on the natural *bolA* P1 sequence (Fig. 12).

**Figure 12:**
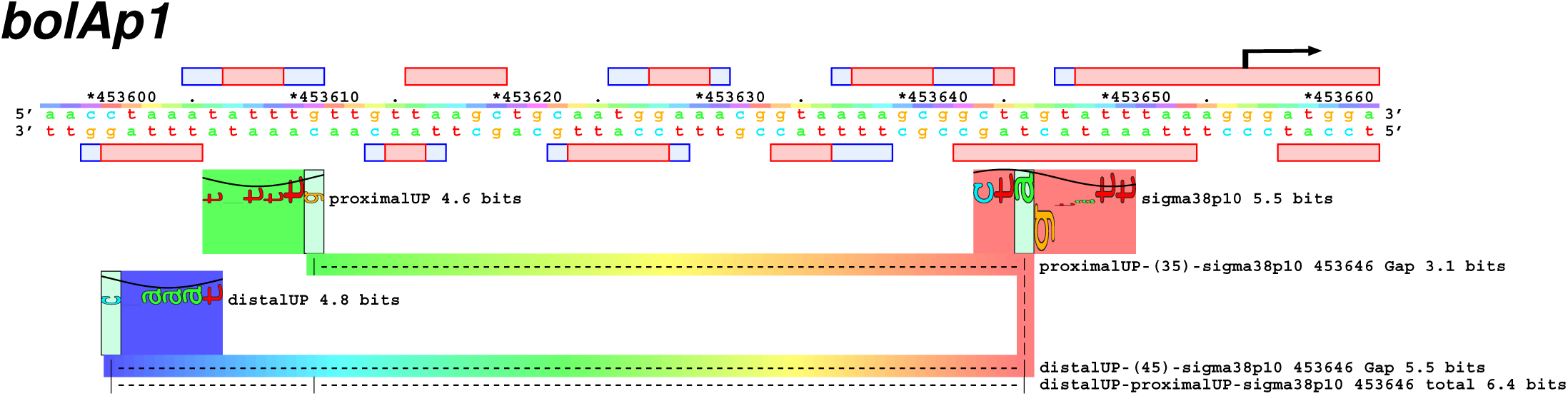
Sequence walker of our *σ*^38^ model scanned on the *bolA* P1 promoter along with hydroxy radical and DNase I footprinting data by Nguyen *et al.* [55]. The top strand represents the nontemplate strand and the bottom represents the template strand. The blue and red boxes above represent protection sites by *σ*^38^ on the nontemplate strand and those below represent protection sites by *σ*^38^ on the template strand. The pink boxes correspond to the locations with strong protection and the light blue boxes correspond to the locations with moderate protection. The black arrow labels the direction of transcription and the start location. The red sequence walker rectangle represents the −10 element, the green one represents the proximal UP element and the blue one represents the distal UP element found on the promoter *bolA* P1 by *σ*^38^. The colored line sweeps through the spectrum every 10.6 bases, which is one turn of B-form DNA [74].

The model picked up a 4.6 bit proximal UP element and a 4.8 distal UP element 11 bases apart. The distal and proximal UP elements were found where protection had been located. The proximal site matched the protection site found on the nontemplate strand and the distal UP element matched the protection site found on the template strand of *bolA* P1 This supports our claim that *σ*^38^ uses a −10, along with proximal and distal UP elements, since the locations of the UP elements were found at the locations of the protection sites. The sequence walker also gave us insight to the possible location of the UP elements corresponding to the binding in the minor groove. It has been shown that the *α*CTD binds the proximal UP element in the minor groove of the DNA [56]. The −1 coordinate of the −10 (the zero coordinate is at the light green vertical rectangle) corresponds to the protein facing the major groove [4] and it is on a yellow section of the spectrum line (program: **live**). That means that the minor groove is located at the magenta-blue sections of the spectrum. The proximal UP element is indeed located in this area of the spectrum, agreeing with the idea that the proximal UP element binds to the minor groove. Because the distal walker is centered on a yellow major groove face at 453603, the distal UP element should be bound by the *α*-CTD in the minor groove on the back side of the DNA from the proximal *α*-CTD.

### Analysis of *in vitro* experiments of the *bolA* P1 promoter

Gaal *et al.* [10] used *in vitro* selection to identify −10 and −35 sequences similar to those of *σ*^70^ on the *σ*^38^ *bolA* P1 promoter. Because we did not find a −35 on *σ*^38^ promoters, we decided to analyze the constructions of Gaal *et al.* in more detail. We first made a control by scanning two models on *bolA1* wildtype sequence (Fig. 13A). We scanned our 169 site *σ*^38^ model and our *σ*^38^ model with an attached −35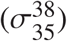 at *σ*^70^ spacing that belongs to our *σ*^70^ model. The *σ*^38^ model was able to locate a 9.4 bit site because of the artificial EcoRI site introduced during the cloning. This differs from the 6.4 bits of the wild type sequence in (Fig. 12). The 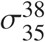 model was unable to locate any site on the ‘natural’ sequence because we made it require a −35. The Gaal *et al.* experiment began with the selection of the −10 promoter region. They constructed fragments that contained an EcoRI site and *bolA1* sequence (−54 to +15) which was randomized in the region −18 to +1, where zero is the first base transcribed (Fig. 13B). After 17 cycles of selection they sequenced 16 DNA fragments; 4 sequences were abnormal leaving 12 for analysis. The sequence logo for their selected −10 is shown in Fig. 14A. In addition to being strong, it resembles the −10 of both *σ*^70^and *σ*^38^. Positions −3 and −2 are TG, which apparently matches the extended −10 [4]. The TGTG pattern is also observed in bacteriophage P1 RepA binding sites [57] and related plasmid DNA replication sites [58].

**Figure 13:**
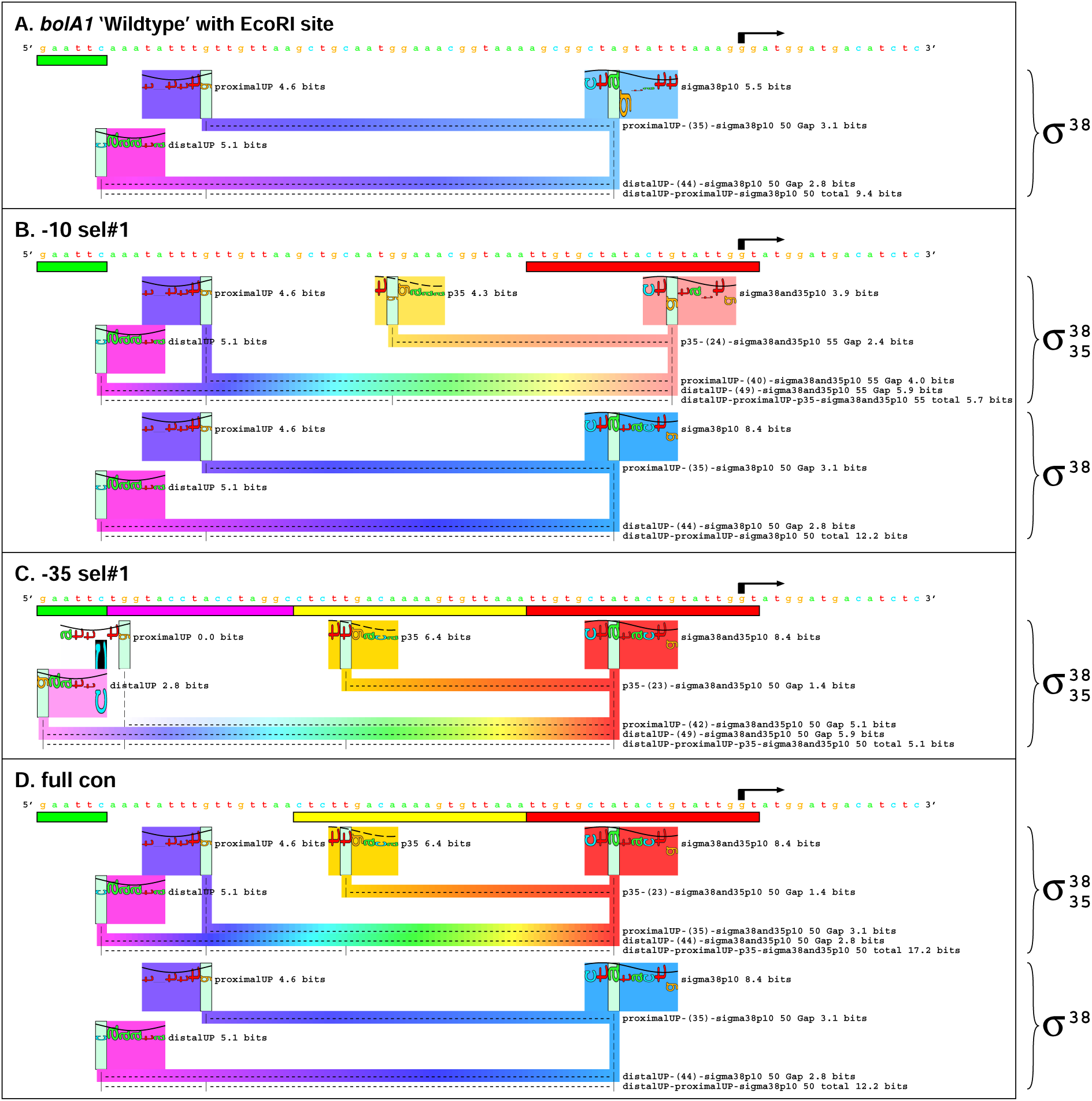
Sequence walkers **scan**ned on the constructed fragments made by Gaal *et al.* [10] for their experiments on σ^*38*^ recognition. The black arrow marks the direction and location of the transcription start sites on *bolA1* and presumably each of the other sequences. Each sequence walker has thin rectangular boxes directly under the sequence that represent different sections of the constructs: green marks the location of the EcoRI site added by Gaal *et al.*, purple represents the SUB sequence added to remove any interference from −35 like motifs. Yellow represents the −35 element selected by *σ*^38^ and red represents the −10 element selected by *σ*^38^. We scanned two different models on the sequences: our *σ*^38^ model and a *σ*^38^ model with a −35 part attached that was taken from our previous *σ*^70^ model 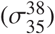. The rectangular colored boxes with letters inside of them (known as petals) represent the location where that part of the model binds. Red petal: −10 of 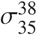 model. Yellow petal: −35 of 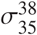 model. Light blue petal: −10 of *σ*^38^ model. Purple petal: Proximal UP element of both models. Pink petal: Distal UP element of both models. The colored bar under the petals represent the Gap and distance between the two sites. The color of the bar corresponds to the color of the sites and transitions from one color to the next. **A:** Sequence for the *bolA1* promoter with EcoRI site. The *σ*^38^ **scan** resulted in a site with a total of 9.4 bits of information. The 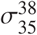 model found no site. **B:** The constructed sequence after selection of the −10 region. The 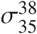 model found a 5.7 bit site. The *σ*^38^ model found a 12.2 bit site. **C:** The selection of the −35 region. The 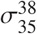 model found a 5.1 bit site. The *σ*^38^ model could not locate a site. **D:** The sequence represents the constructed fragment by Gaal *et al.* in which they kept the selected regions from the −10 and −35 selections and removed the SUB sequence. The 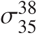 model found a site with a total of 17.2 bits of information. The *σ*^38^ model found a site with a total of 12.2 bits of information.

**Figure 14:**
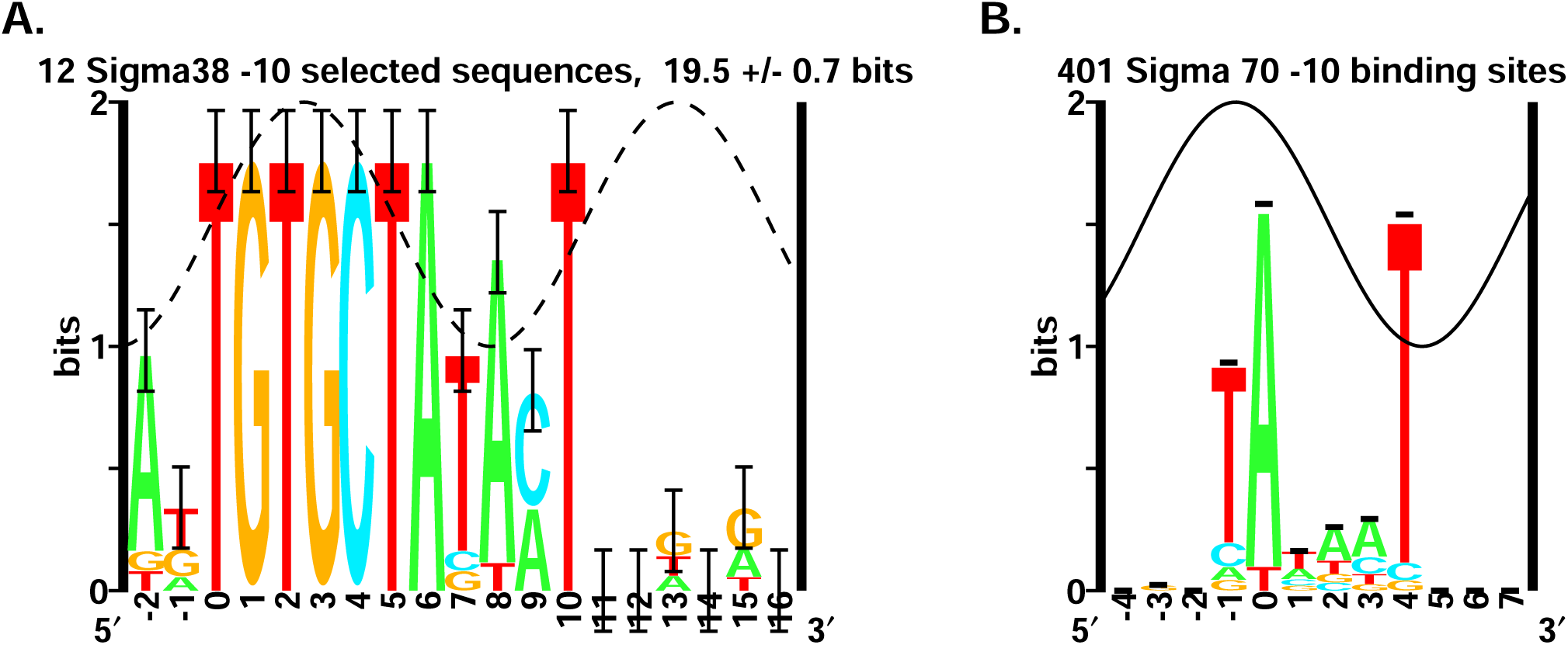
*In vitro* selection of *σ*^38^ −10 elements. After 17 cycles of selection in the −10 region by *σ*^38^ polymerase, Gaal *et al.* [10] reported 12 DNA fragments (Fig.2 of Gaal *et al.*) **A:** Sequence logo of 12 selected *σ*^38^ −10 elements, 19.5 ± 0.7 bits. **B:** Sequence logo of 401 natural *σ*^70^ −10 elements from [4], 4.78 ± 0.11 bits.

We reconstructed their resulting DNA (‘−10 sel#1’) and scanned the same two models on the sequence. The 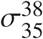 model was able to locate a 5.7 bit site and the *σ*^38^ model was able to locate a 12.2 bit site (Fig. 13B). Both models had different −10 sites from the wildtype. The *σ*^38^ model found a strong 8.4 bit −10. The 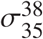 model was able to locate a −35 in the natural region, but the distance of the −35 causes the model to locate a weak −10 and the gap between the two lowers the total information content to 5.7 bits. The gap was 24 bases, which is one base off from the optimal range for a −35 of 23 bases when the −35 corresponds to a *σ*^70^ [4]. The −10 site found for the 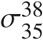 model was only 6 bases away from the transcription start which appears too close to be an actual −10 site, but transcription start sites were not reported in this experiment. The −10 site found for the *σ*^38^ model was 11 bases away from the transcription start which is the optimal distance for a −10 location when it corresponds to *σ*^70^ [4].

The next selections by Gaal *et al.* were to locate a −35. For these selections they constructed fragments that contained an EcoRI site, a ‘SUB’ sequence from −54 to −39, random sequence from −38 to −19, the previously selected −10 sequence from −18 to +1 and *bolA1* sequence from +2 to +15 (Fig. 13C, −35 sel#1). The SUB sequence from −54 to −39 was designed to remove −35-like sequences. After 17 cycles of selection they sequenced 22 DNA fragments; 3 sequences were abnormal leaving 19 for analysis. We built a model for the selected −35 sequences and found that they resemble but are 14 bits stronger than natural *σ*^70^ −35 sites (Fig. 15). We then scanned the *σ*^70^ and *σ*^38^ promoters using a model built from these selected sites (Fig. 16). On *σ*^70^ promoters we identified a high density of weak sites around the −35 region (Fig. 16A). In contrast when the model was scanned on *σ*^38^ promoters we did not find a high number of sites around the −35 region (Fig. 16B).

**Figure 15:**
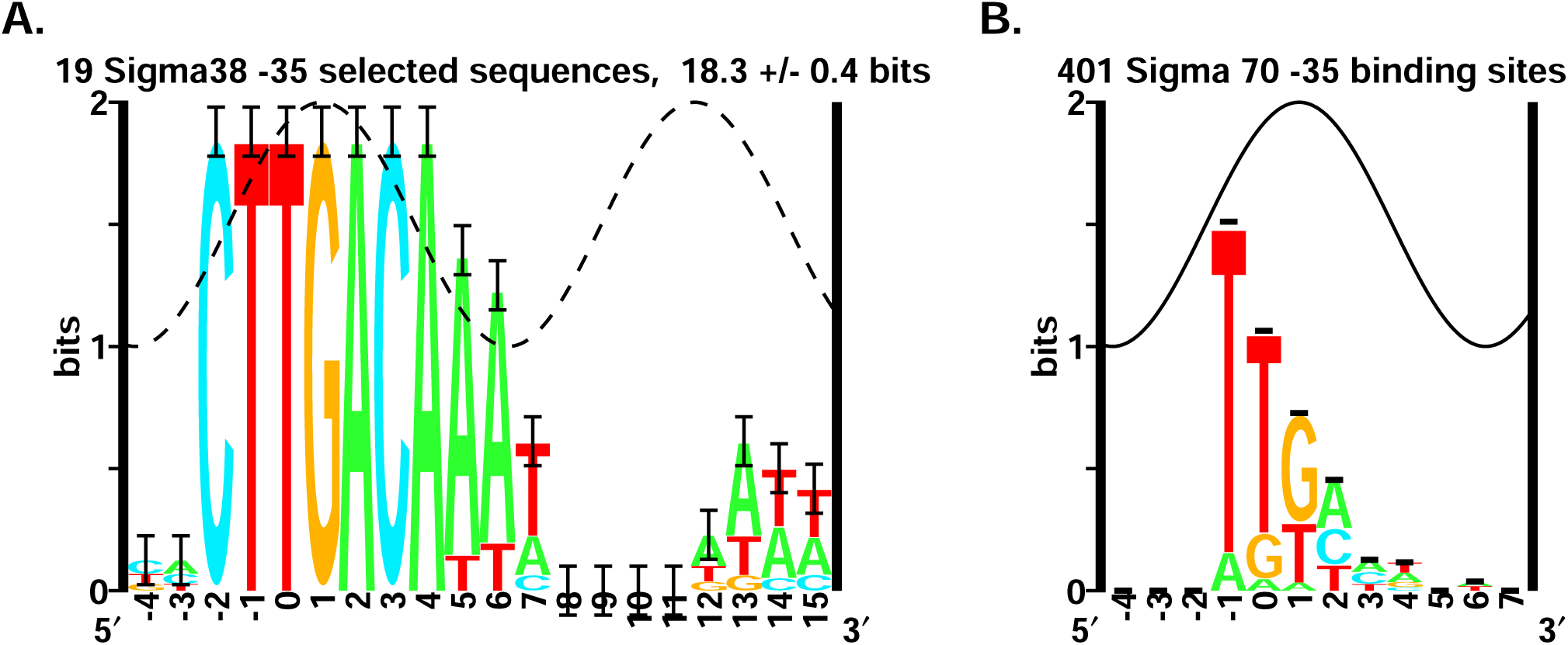
*In vitro* selection of *σ*^38^ −35 elements. After 17 cycles of selection in the −35 region by *σ*^38^ polymerase, Gaal *et al.* [10*] reported 19 DNA fragments (Fig.3 of Gaal et al.*) **A:** Sequence logo of 19 selected *σ*^38^ −35 elements, 18.3 ± 0.4 bits. **B:** Sequence logo of 401 natural *σ*^70^ −35 elements from [4], 4.14 ± 0.02 bits.

**Figure 16:**
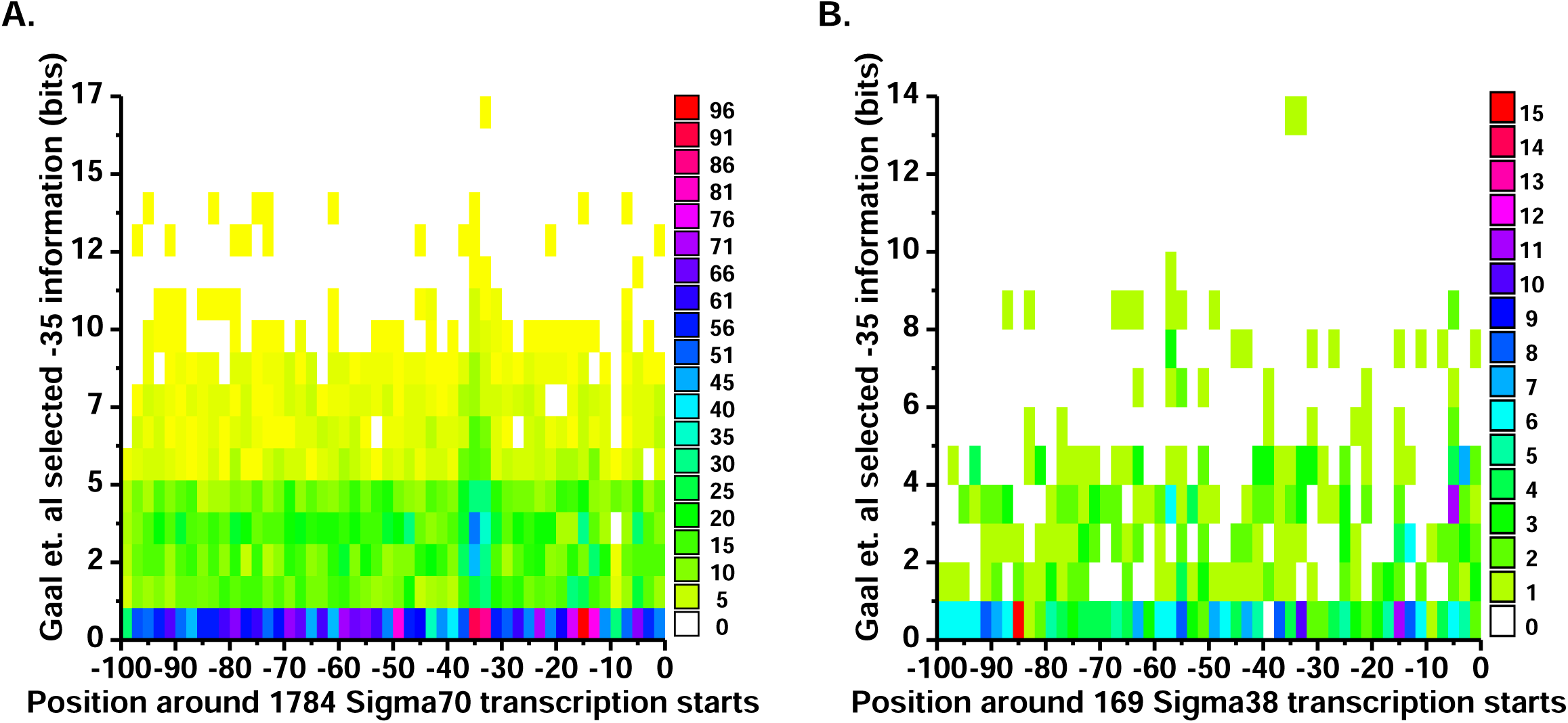
-35 weight matrix from Gaal *et al.’s* −35 *σ*^38^ selection experiment (Fig. 15A) scanned on *σ*^70^ and *σ*^38^promoters. **A:** We scanned the RegulonDB dataset of 1784 proven *σ*^70^ sites 100 bases upstream from their transcription start sites. The selected −35 model located a cluster of −35 sites approximately 35 bases upstream from the transcription start sites with an information content of 2 to 4 bits. **B:** −35 weight matrix scanned on *σ*^38^ promoters. We scanned the 169 *σ*^38^ promoters 100 bases upstream from their transcription start sites. The selected −35 model could only locate two −35 sites on the *σ*^38^ promoters, on genes ybgAp1 and ihfBp, out of 169 genes.

Gaal *et al.* chose one of the selected −35 sequences (‘−35 sel#1’), but the *σ*^38^ model was unable to locate a site (Fig. 13C). Instead, the 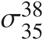 model located a 5.1 bit site with a strong 8.4 bit −10 which was the same as the −10 that the *σ*^38^ model had found in the selection for the −10. In the −35 selection (Fig. 13C), since the −35 element was improved, the 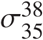 model was able to identify a better −10 than it found in the −10 selection (Fig. 13B). The 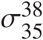 model was also able to locate a strong 6.4 bit −35 site in the region selected for a −35. Notably, this sequence lacked UP elements. By adding the SUB sequence upstream from the −35, the area where the UP elements were was ruined. The 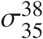 model was only able to locate a proximal UP element just above 0 bits and the distal UP element that was located was the artificial EcoRI site upstream from the SUB fragment. Apparently, in the absence of strong UP elements *σ*^38^ was forced to use a −35.

Once all selections were completed Gaal *et al.* constructed a promoter that contained the selected −35 and −10 regions (‘full con’). This construct had selected sequence from −38 to +1. They replaced the SUB sequence with wildtype *bolA1* sequence from −54 to −39. This brought the modified UP elements back into the construct (Fig. 13D). Our *σ*^38^ model was able to locate a 12.2 bit site and the 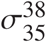 model found a 17.2 bit site. By removing the SUB sequence and returning to almost natural sequence, the UP elements were able to improve the information content from a 5.1 bit site to a 17.2 bit site. Compared to the average *σ*^38^ promoter in our set of 169 sites, a 17.2 bit site is unusually strong (Z=2.33, p=0.01). We predict that due to strong binding, the *σ*^38^ will be unable to escape the promoter. Indeed, supporting our result of strong binding, Gaal *et al.* reported that the full con is relatively inactive.

These observations explain why we were unable to locate a −35 element in our *σ*^38^ dataset sequences. Natural *σ*^38^ promoters do not have a −35 element, but when *σ*^38^ is forced into choosing a −35 by removing the UP elements, it can use one. Even though it can use a −35, using one in the presence of UP elements makes the promoter too strong and therefore inactive [29]. This effect should select −35 element mutations for the *σ*^38^ promoters to stay functional.

### Analysis of *in vitro* experiments on the *rrnB* P1 promoter

Estrem *et al.* [59] randomized the region upstream of the natural *E. coli rrnB* P1 promoter and selected for strong transcription. The resulting sequence logo shows 23.7 ± 0.3 bits (Supplementary figure Estrem.Gourse1998-logo.pdf), which is more than the information required to locate a single site in the *E. coli* genome (log_2_(2 × 4639675) = 23.1 bits), suggesting that the selection obtained unnaturally strong sequences [22, 53]. We analyzed the individual mutant sequences with our 169 site model (Supplementary figure Estrem.Gourse1998-rrnB-P1-selection.pdf constructed from their Figure 2) and found that most of the conservation observed in the logo (−7 to +3) is explained by UP elements at various positions. We do not know why +7 to +9 is conserved as our model does not provide for proximal UP elements in this region, but it may prevent binding of the *α*CTD in ways that reduce transcription. Some of the distal UP elements use a C 5′ to the randomized region. This analysis using the two-UP element model suggests that the interpretation of the original randomization experiment is complicated and that its representation as a sequence logo does not directly represent UP element binding since they are not rigidly aligned relative to the rest of the promoter.

Estrem *et al.* [59] performed DNase I footprinting on the 4192 selected variant and the protected and enhanced DNA cleavage they observed corresponds to the zero base of both the distal and proximal UP elements predicted by our model (Supplementary figure Estrem.Gourse1998-rrnB-P1-selection.pdf).

Our model predicts that the −46 DNA sequence 5′CGCGAAATTTCGCG3′ used by Yasuno *et al.* [60] contains two oppositely oriented 6.1 bit UP elements that would compete for *α*CTD in the minor groove [61]. This is consistent with their analysis of NMR NOEs which show that two *α*CTDs bind across the central A/T region, including the ending G and C. In addition, their −49 sequence 5′CGCGTTTAAACGCG3′ contains two weaker (5.0 bit) sites, explaining why this sequence bound more weakly in their experiments. (See Supplementary materials Yasuno.Kyogoku2001-figure1b.pdf)

### Test of the 169 site model on new data

Since completing the results presented above, a new version of the RegulonDB database has been released (Release: 9.2 Date: 09-08-2016). This version now provides strength of evidence for the *σ*^38^ promoters: in our 78 sequence set, 3 are now labeled ‘confirmed’ and 75 are ‘strong’. The data set also provides 25 new sequences that are not in our original 169 site set. As a test, we **scan**ned these with the 169 site model and 22 (88%) are identified above zero bits (see supplementary figure new22map.pdf).

We also analyzed the sequences and experimental data from Typas *et al.* [17] (See Supplementary materials). *β*-galactosidase measurements of UP element sequence constructs matched our model in 8 of 9 cases. The one exceptional construct could be explained by other factors influencing the *in vivo* experiment.

Finally, in the Supplementary materials we provide sequence walker analysis of the ChIP-seq data from Peano *et al.* [62]. 100 of the 104 ChIP-seq peak sequences had predicted *σ*^38^ sites.

In summary analyses of many experiments already performed on *σ*^38^ promoters are consistent with our information-theory based model.

### Confirmation of the *σ*^38^ model by *in vitro* transcription experiments

Based on our model, *σ*^38^ should recognize the *rrnB* P1 promoter and indeed, our *in vitro* transcription assays confirmed this prediction (Fig. 17). This first experiment also shows that *α*CTD is essential for *σ*^38^ and important for *σ*^70^. In both cases Fis improves transcription. To test our model we designed several mutations of the *rrnB* P1 promoter to determine the importance of the −10, −35 and UP elements (see the supplementary materials). In our second experiment (Fig. 18) the −10 and UP elements were required for transcription, but the −35 was not needed. In addition, the *α*CTD was essential for *σ*^38^ transcription. These results are fully consistent with our *σ*^38^ model (Fig. 6). Finally, in our third experiment we examined the *bolA* P1 promoter (Fig. 12, Fig. 19). Transcription by *σ*^38^ required the −10 and the *α*CTD but not the −35. However, in this case removal of the UPs did not affect *σ*^38^ transcription.

**Figure 17:**
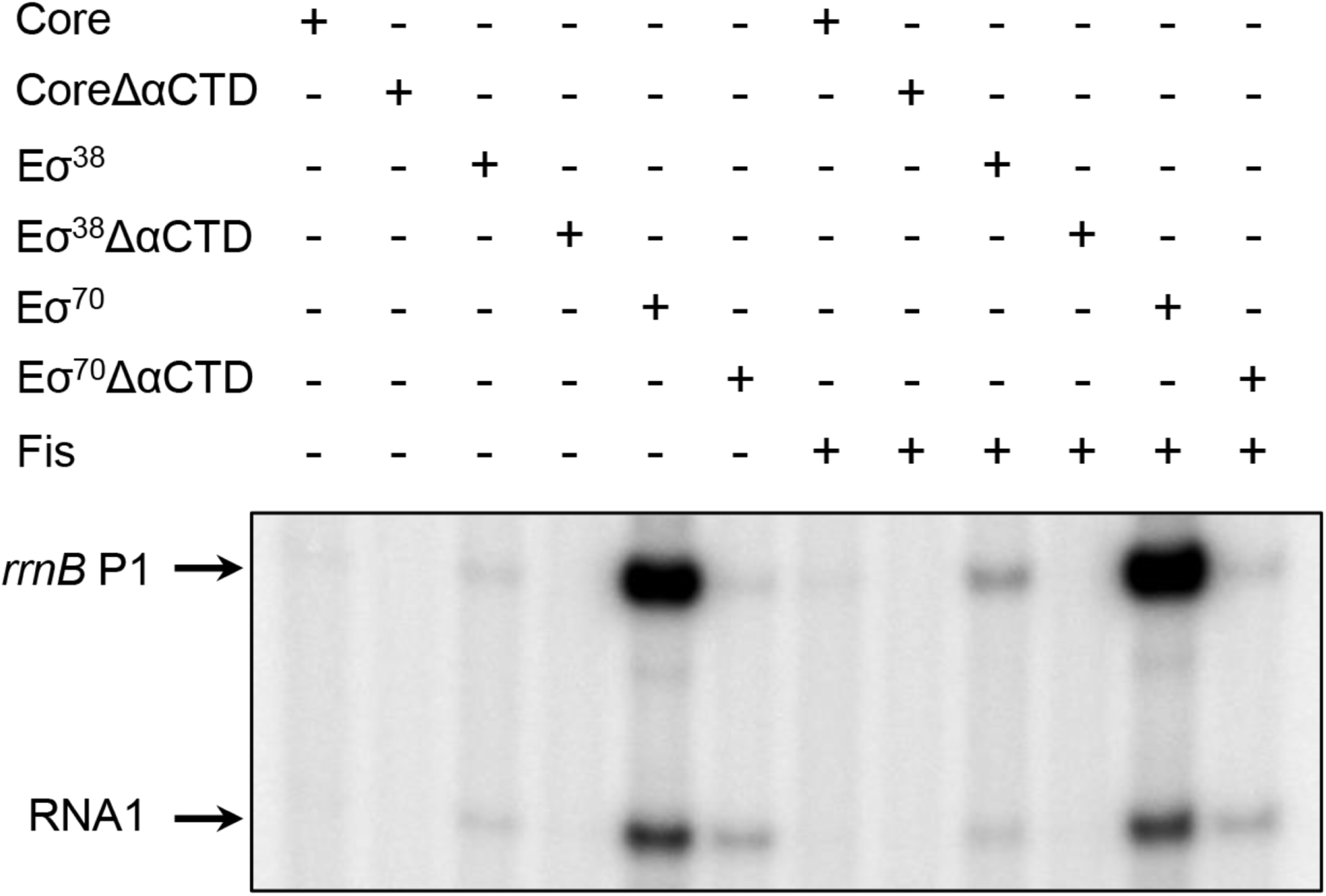
*In vitro* transcription of the *rrnB* P1 promoter by RNA polymerases E*σ*^38^, E*σ*^70^, E*σ*^38^Δ*α*CTD and E*σ*^70^Δ*α*CTD. The supercoiled template used in the assays is plasmid pDJ-*rrnB* P1. Arrows indicate the transcripts from the target promoter and internal control RNA1 from the ColE1 plasmid.

**Figure 18:**
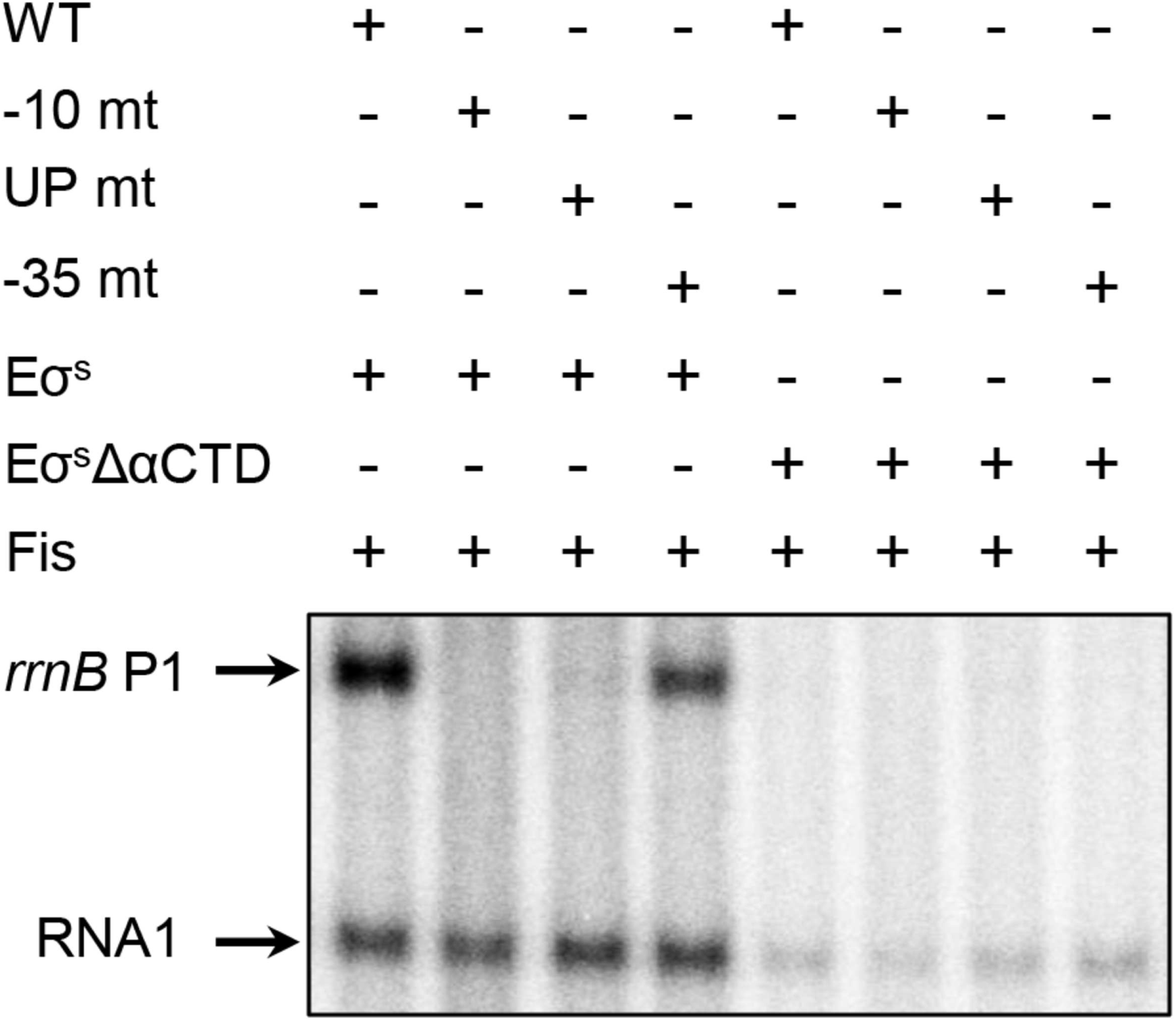
*In vitro* transcription of the *rrnB* P1 promoter and mutants by RNA polymerases E*σ*^38^ and E*σ*^38^Δ*α*CTD. WT, wild-type *rrnB* P1 promoter; −10 mt, mutated −10 region; UP mt, mutated UP elements; −35 mt, mutated −35 region. See supplementary materials for lister maps of the mutations.

**Figure 19:**
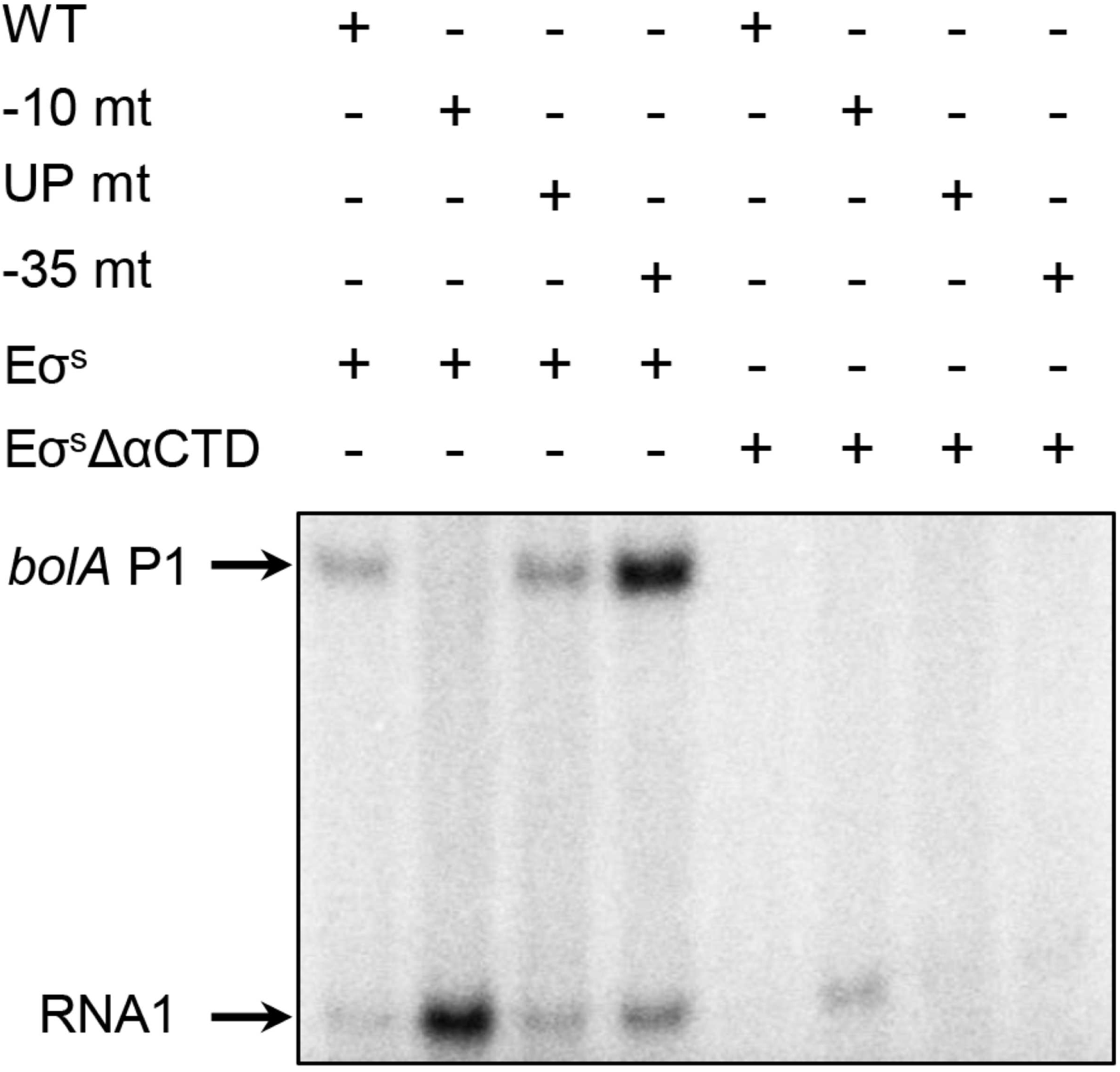
*In vitro* transcription of the *bolA* P1 promoter and mutants by RNA polymerases E*σ*^38^ and E*σ*^38^Δ*α*CTD. WT, wild-type *bolA* P1 promoter; −10 mt, mutated −10 region; UP mt, mutated UP elements; −35 mt, mutated −35 region. See supplementary materials for lister maps of the mutations.

## 5 Discussion

Although natural *σ*^38^ promoters have a −10 element that is easily found by multiple alignment to maximize the information content (Fig. 2), they do not have a detectable −35 (Fig. 9, Fig. 10). However, instead of a −35 we found that two UP elements are always present upstream of *σ*^38^ promoters. In addition to the subtle variation between the −10 elements (Fig. 2, Fig. 11) our analysis resolves the paradox of why *σ*^38^ promoters function without the −35 element which is naturally required by *σ*^70^ [14, 15, 16, 17, 12, 1].

Our quantitative results were obtained using information theory [20, 21]. We have successfully used this mathematics to align simple single-protein OxyR and Fis binding sites [33, 49]. We then applied this method to the more complicated flexible *E. coli* ribosome binding sites by aligning the Shine and Dalgarno region relative to the known initiation codon [31]. This revealed a sequence logo that matches the 3′ end of the 16S ribosomal RNA, as expected for the Shine and Dalgarno element. Next, we applied the same method to *σ*^70^ promoters and generated a two-part flexible model [4] that was able to identify small RNAs and proteins [63]. In other words, alignment of sequences by maximizing information works and this explains why we were able to discover that essentially all *σ*^38^ promoters contain distal and proximal UP elements, but no −35.

Previous authors have instead proposed that *σ*^38^ RNAP can transcribe with the presence of a degenerate −35 element. It was called degenerate because it does not match a consensus sequence for the −35 of *σ*^70^ [11, 18]. It had also been noticed that genes that respond to *σ*^38^ have completely different sequences in the −35 region [19]. These experiments led to the belief that *σ*^38^ RNAP can transcribe stress genes simply through the differences in the −10 element from *σ*^70^ [64, 65, 8]. Our results show that *σ*^38^ RNAP does not use a −35 in natural promoters and instead recognizes the differences in the −10 element along with two UP elements.

### UP elements and the *α*CTD of RNAP

The 329-amino-acid long *α* subunit of RNA polymerase contains an N-terminal domain (*α*NTD) and a C-terminal domain (*α*CTD) [50, 66]. These two domains are separated by a 13 amino acid flexible linker. The *α*CTD can increase the rate of transcription initiation by interacting with a DNA sequence upstream from the −35 known as the UP element [6]. The two *α*CTD domains can bind to a ‘proximal’ UP element −46 to −38 positions upstream or a ‘distal’ UP element −59 to −47 positions upstream as found on the *rrnB* P1 promoter (Fig. 4) [47, 46, 48]. The *rrnB* P1 promoter has the best characterized UP elements; they increase the promoter activity > 30-fold [6, 47].

Footprinting, cross-linking, drug binding and X-ray crystallography have shown that the *α*CTD binds to the minor groove of UP-elements [56, 67, 60]. The majority of the sequence logo positions of both the distal and proximal UP elements have about equal numbers of A and T in a combined UP element model (Fig. 8). In B-form DNA this could represent minor groove contacts in which the N2 moiety of G in the minor groove is blocked [45, 34] and it could be provided by protein moiety or a spine of water in the minor groove [67]. By blocking the guanine N2, only A and T bases are allowed but their orientation is not determined and so they should appear at approximately equal frequencies, as observed. In contrast, the 5′ end of the distal UP and the 3′ end of the proximal UP (both are position 0 in Figs 3, 6 and 8) contain nearly equal numbers of G and C. We have not seen this configuration before, but it could represent a protein or water spine contact in the minor groove to the guanine N2 hydrogen donor [67]. Because such a hydrogen bond is on the dyad axis of the DNA, this ambiguity leads to approximately equal numbers of G and C. Since there is ambiguity in minor groove contacts leading to the equal numbers of G and C or A and T [45, 34], all positions of the logos being near or less than 1 bit is consistent with minor groove contacts [42]. Thus all of the UP element sequence logo bases can be accounted for by contacts in the minor groove, as observed by footprinting data [56]. The string of As and Ts would allow multiple binding positions but the G or C bases would anchor the *α*CTD to specific positions on the DNA, allowing for precise positioning relative to the RNA polymerase and activators.

However, a closer inspection of the logos reveals that position +1 in the distal UPs, position −1 in the proximal UPs (Fig. 3) and position +1 in the combined UP model (Fig. 8) all have more than 1 bit of information. This is not a nearest-neighbor effect since a logo of G or C (*i.e.*, ‘S’) followed by A or T (*i.e.*, ‘W’) in the first 100,000 bases of the *E. coli* genome show no such bias (data not shown). Likewise, logos for the 91428 cases of SWWWWW or WWWWWS are flat at 1 bit except that the base next to the S is 1.025 bits, so the observed 1.176 bits are not caused by general sequence biases although they are in the same direction (data not shown). Since there is only one contact that can distinguish bases in the minor groove and it is close to the dyad axis [45] no more than 1 bit should be observed in contacts made into the minor groove of B-form DNA [57, 42]. By a binomial test between the 257 As and the 80 Ts (ignoring the one G), this position is highly unusual (*p* < 2.2 × 10^−16^), suggesting that non-B-form DNA exists in at least some of the UP binding sites, as found in IHF, TATA and RepA which also bind the minor groove [42, 43].

The UP region is said to be A/T rich [6]. Such a region must eventually end in a G or a C, so when we aligned the sequences to maximize the information in an 8 bp wide window of the distal region we would get G or C followed by a string of A and T, as observed (Fig 3 and 6). Likewise a pattern related by dyad symmetry would appear on the proximal side. How can we distinguish this A/T tract model from the specific binding motif models shown in figures 3, 6 and 8? First, the information curves (height of sequence logo stacks) of the proximal and distal logos match so that position ±1 (*i.e.* distal position +1 and proximal −1) is anomalously high, as discussed above. This suggests that a specific alignment exists beyond the terminal G or C. Furthermore, position ±4 is lower than ±2, ±3 and ±5 for both regions. Thus the distal and proximal information curves are similar even though they come from different sequences. This would not be likely according to a tract model. Furthermore, position 6 has no information for either the distal or proximal regions and this extends to ±20 (0.01 ± 0.01 bits for the combined UP, not shown), as expected from a finite length motif. On the other hand, a tract with indefinite length should show an A/T pattern from position ±6 onward. Finally, a test of the motif hypothesis is to ask whether the region between the two UPs is A/T rich or consists of equiprobable bases. The base composition between UP motifs that do not overlap is close to equiprobable (A 114, C 104, G 105, T 109) so the A/T tract model is not supported. In an A/T tract the position of the *α*CTD would not be well determined. However, a specific motif suggests that the *α*CTD is anchored by the G/C at position 0 of the UP (Fig. 8) allowing it to be precisely positioned.

### Dyad symmetry of UP elements in *σ*^38^ promoters correspond to the dyad axis of symmetry of *α*NTDs bound to core polymerase

When we aligned two sequence regions upstream of the *σ*^38^ −35 region, we found that they were inverted sequences with respect to each other (Fig. 3, Fig. 6). Since these regions correspond exactly to UP elements (Figures 4, 12), they are the DNA binding sites of the *α*CTD domains of the RNA polymerase. The identified UP elements have dyad symmetry, which implies that the *α*CTD subunits bind DNA with a two fold axis of symmetry. In no case did we observe UP elements in the same orientation. Does this dyad axis of symmetry reflect the symmetry of the other end of the *α* subunits?

Indeed, the first step of assembly of RNA polymerase is the dimerization of the *α* subunits, followed by addition of the *β* and *β*′ subunits [68]. Furthermore, the *α*NTD subunits do bind to the polymerase with dyad symmetry, as shown by the holoenzyme structure (Fig. 20). Since *α* subunits dimerize, they may require that the UP elements to be of opposite orientation on the DNA, as we discovered. Notably, it is somewhat mysterious that the 13 amino acid long linker between the *α*NTD and the *α*CTD subunits apparently does not allow significant 180° rotation of the *α*CTD for the UP elements to bind in the other orientation.

**Figure 20:**
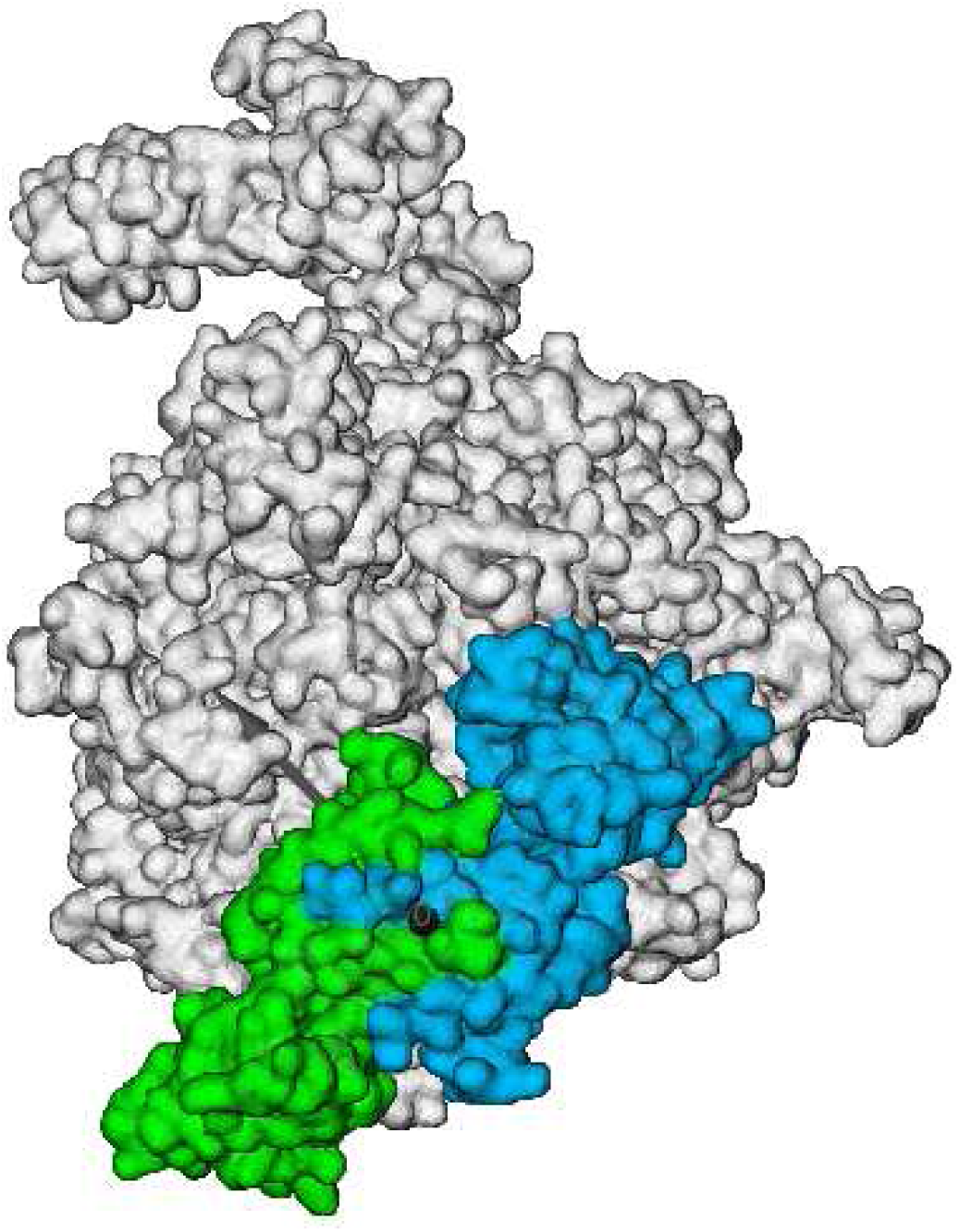
Dyad symmetry of *α*NTD subunits bound to RNAP. The two *α*-N-terminal domains (*α*NTD) of the *σ*^70^ RNA polymerase are shown in green and cyan. The *σ*^70^, *β, β*′ and *ω* subunits are represented as white. The structure is one of two polymerases in PDB entry 4IGC [75]. The black dot shows the dyad axis of symmetry for the two *α*-NTD subunits pointing directly at the viewer. The figure was generated using VMD 1.9.2 [76]. (4IGC has been superseded by 4YG2 but the new structure still shows *α* dyad symmetry. The *α*CTD in 4IGC was removed for this figure.)

A DNA binding protein may halve its required genetic coding region requirement by dimerizing. This explains the frequent dimerization of transcription factors and their palindromic dyad axis of symmetry binding sites (see Fig. 6 in [57]). In contrast, a directionally oriented molecule such as an RNA polymerase must have an asymmetric binding site. A polymerase can be activated by a symmetrical transcriptional factor by recruitment [67]. Since the orientation is already determined by the *σ* factor, by having two symmetrical contacts of its own, the polymerase can get more information with a smaller protein coding gene. Perhaps this geometrical logic led to the evolution of two *α* subunits on RNAP.

Our results are consistent with the proposal by Gaal *et al.* [69] that *α*CTD binds DNA using twofold symmetry. However, our models are inconsistent with the proposed major groove binding since the sequence logos (Figures 3, 6 and 8) only show G/C and A/T, which are best explained as entirely minor groove binding.

### rpoS(*σ*^38^**)-***α***CTD contact**

Ross *et al.* [70] showed that when bound to the proximal UP element, the *α*CTD residues D259 and E261 contact R603 in the *σ*^70^ region 4.2 and that this contact increases transcription both *in vivo* and *in vitro*. We analyzed the relevant sequences (4547, 4549 and rrnBP1-Proximal in supplementary materials Estrem.Gourse1999-fig1-map.pdf and Figure 4) and found that our model fits this position with a gap of 27 bases. A referee of the present paper pointed out that the contact between the *α*CTD and rpoD (*σ*^70^) does not exist in rpoS (*σ*^38^) (gene *rpoD versus rpoS* in GenBank Accession NC_000913). This lack of a contact is consistent with the wide distribution of gap distances between the proximal UP element and the −10 shown in Figure 6B. This figure shows that the gap for the UP element with *σ*^70^ at −27 has an exceptionally high 13 examples, while the rest of the distribution is roughly Gaussian with a peak at −34. This suggests that some number of UP elements are positioned so as to make the supposedly non-existent contact in *σ*^38^ promoters. Ribosomal promoters could potentially use either *σ*^70^ or *σ*^38^ binding but Figure 5 shows that only *rrnA, rrnB, rrnC* and *rrnH* have the right spacing according to our model. Our 169 sample model of *σ*^38^ binding does not include the ribosomal promoters, yet of the 329 ways that our *σ*^38^ model binds in the known *σ*^38^ promoters (see Supplement 169sigma38map.pdf), the distribution is again roughly Gaussian but there is a spike of 18 cases that have a gap of 27 bases (data not shown). This appears to suggest that rpoS may also make a contact to the *α*CTD, but differently from rpoD.

### Flip-Flop model of *α*CTD-UP binding

An important feature of UP elements is revealed by the flexible sequence walker models. The UP walkers sometimes overlap in physical space, for example in Fig. 5 *rrnD* P1, *rrnE* P1, *rrnG* P1, possibly *malT* and *guaB*. Since the *α*CTD binds in the minor groove [60, 67], strongly overlapping sequence walkers imply that the two *α*CTD molecules cannot bind at the same time. We previously reported this self-regulating effect at Fis binding sites [61]. In other words, the *α*CTDs bind as a molecular ‘flip-flop’. If only one molecule can bind at one time but both are able to bind, the number of ways of binding is doubled and so the association constant is doubled. Experimental work could distinguish the flip-flop binding mode from both binding simultaneously since instead of the sum of the two UP elements the information should be only about one bit more. Despite this caveat, our model was able to identify the majority of sigma38 sites.

### Evolution of promoter elements

We found no −35 sequences in *σ*^38^ promoters, yet Gaal *et al.* [10] were able to force *σ*^38^ to use a −35 in their DNA randomization and selection experiment. A transcription initiation complex of *σ*^38^ lacking the *α*CTD did not reveal a −35 contact to DNA [13]. This was attributed to crystal packing. One hypothesis to explain this well-known paradox is that *rpoS*, the gene for *σ*^38^, and *rpoD*, the gene for *σ*^70^, were formed by gene duplication of a common ancestor that was able to use the −35 [1]. As time went on *σ*^38^ began to specialize by only functioning during stress. We hypothesize that the distinction between *σ*^38^ and *σ*^70^ occurred by *σ*^38^ DNA binding sites losing the −35 elements and gaining UP elements, a mix-and-match strategy [71]. Indeed, having both UP and −35 elements together makes the promoter non-functional, possibly due to very tight binding of polymerase to the promoter [10, 29]. However perhaps *σ*^38^ has not completely lost the ability to bind the −35 region, so if *σ*^38^ is forced to use the −35 region, it can do so in the absence of UP elements [29].

### Experimental confirmation of the *σ*^38^ −10, no −35 with UPs model

The *σ*^70^-driven P1 promoters of all *E. coli rrn* operons have two UP elements, as shown in Fig. 4 and Fig. 5. This raises a question as to whether RNA polymerase holoenzyme containing *σ*^38^ uses those UPs to recognize and initiate transcription from ribosomal promoters. Such promoters would use the same −10 and be activated during stress conditions. Therefore experiments were warranted to address this issue.

To test our *σ*^38^ binding site model (Fig. 6), we performed *in vitro* transcription on the *rrnB* P1 and the *bolA* P1 promoters (Figs. 17-19). First we showed that E*σ*^38^ does indeed transcribe the *rrnB* P1 promoter and that this depends on the *α*CTD (Fig. 17), as would be expected if E*σ*^38^ uses UP elements. Next, using the *rrnB* P1 promoter we demonstrated that E*σ*^38^ required a −10 and UP elements but does not does not need a −35 as predicted by our model (Fig. 18).

Finally, we found that E*σ*^38^ also required a −10 but does not need a −35 on the *bolA* promoter (Figs. 12, 19). However, unlike *rrnB* P1, deletion of the UP elements on this promoter did not prevent transcription (Fig. 19 lane 3). This result suggests that there could be additional binding modes for UP elements that our model does not accommodate. That may be related to the difficulties we had in designing mutations of UP elements in the first place. We frequently found that a single base mutation in an UP element would result in another UP being created nearby according to the model. We learned that when we created strings of Cs and Gs these alternative UP elements could be suppressed. However, our model was built from only 169 sequences, which limits its precision. In the model the UP elements are spread over 34 positions, so an average of only 5 UP elements are represented at each distance (Fig. 6). Because of this low count, it is possible that there are other positions to which the *α*CTD could bind that escaped our model and these might account for the insensitivity to this UP-region mutation. Alternatively, there may be additional binding modes not found in the 169 examples or such modes may be active only *in vitro*. More extensive mutagenesis may be needed to determine the precise range and modes that the *α*CTDs can bind.

Deletion of the UP elements in *rrnB* P1 is complicated since the natural *rrnB* P1 promoter has 3 Fis sites (Fig. 4). Our construction for testing *rrnB* P1 removed the most distal site and reduced the information content of the middle site from 5.3 bits to 3.2 bits. Since the efficiency of Fis binding is 69% (data not shown) this change should reduce binding by 2^(5.3−3.2)/0.69^ = 8.2 fold [24]. Without Fis the *rrnB* P1 promoter fires weakly (Fig. 17) which implies that the one Fis site and the reduced site are needed for transcription. Perhaps this was lucky since if all three Fis sites had been in our construction maybe *σ*^38^ could have avoided using the UPs. But *σ*^38^ needs the UPs according to the experimental result. This is also consistent with our binding site model.

Our information theory-based computer modeling and experimental results imply that *σ*^38^ requires UP elements or an activator and a −10 but that in natural promoters it does not use a −35. The new finding that *σ*^38^ recognizes ribosomal RNA promoters demonstrates an additional dimension in the regulation of ribosomal RNA which warrants further study.

## Supporting information

Supplement_for_sigma38_paper-directory

## 7 Acknowledgments

We thank Vi Black for managing the NIH CCR Cancer Research Interns Summer Internship Program https://ccr.cancer.gov/training/ccr-interns. This project would not have been possible without the well-curated RegulonDB database created by Julio Collado-Vides [32]. We thank Jeff Strathern for useful discussions; Amar Klar, Jerome Izard and Maria Kireeva for comments on the manuscript; John Eberwein and Gary Stormo for writing the **markov** program; an anonymous referee for pointing out the interesting paper Ross *et al.* [70]; Stephan Lacour for updated data from Peano *et al.* [62]; and the Advanced Biomedical Computing Center (ABCC) for support. This material was presented by KSF at the 2017 Mechanisms and Regulation of Prokaryotic Transcription meeting hosted by the Federation of American Societies for Experimental Biology. This research was supported in part by the Intramural Research Program of the NIH, National Cancer Institute, Center for Cancer Research. Yuhong Zuo was supported by NIH grant GM22778 to Thomas A. Steitz.

## References

[1] K. B. Decker and D. M. Hinton. Transcription regulation at the core: similarities among bacterial, archaeal, and eukaryotic RNA polymerases. Annu Rev Microbiol, 67:113–139, 2013. https://doi.org/10.1146/annurev-micro-092412-155756.

[2] T. M. Gruber and C. A. Gross. Multiple sigma subunits and the partitioning of bacterial transcription space. Annu Rev Microbiol, 57:441–466, 2003. https://doi.org/10.1146/annurev.micro.57.030502.090913.

[3] M. S. Paget and J. D. Helmann. The *σ*^70^ family of sigma factors. Genome Biol, 4:203, 2003. https://doi.org/10.1186/gb-2003-4-1-203.

[4] R. K. Shultzaberger, Zehua Chen, Karen A. Lewis, and T. D. Schneider. Anatomy of *Escherichia coli σ*^70^ promoters. Nucleic Acids Res., 35:771–788, 2007. https://doi.org/10.1093/nar/gkl956, https://alum.mit.edu/www/toms/papers/flexprom/.

[5] S. Keilty and M. Rosenberg. Constitutive function of a positively regulated promoter reveals new sequences essential for activity. J. Biol. Chem., 262:6389–6395. http://www.jbc.org/content/262/13/6389.abstract.

[6] W. Ross, K. K. Gosink, J. Salomon, K. Igarashi, C. Zou, A. Ishihama, K. Severinov, and R. L. Gourse. A third recognition element in bacterial promoters: DNA binding by the α subunit of RNA polymerase. Science, 262:1407–1413, 1993. https://doi.org/10.1126/science.8248780.

[7] S. Lacour and P. Landini. *σ^S^* -Dependent Gene Expression at the Onset of Stationary Phase in *Escherichia coli*: Function of *σ^S^* -Dependent Genes and Identification of Their Promoter Sequences. J. Bacteriol., 186:7186–7195, 2004. https://doi.org/10.1128/JB.186.21.7186-7195.2004.

[8] H. Weber, T. Polen, J. Heuveling, V. F. Wendisch, and R. Hengge. Genome-wide analysis of the general stress response network in *Escherichia coli: σ^S^* -dependent genes, promoters, and sigma factor selectivity. J. Bacteriol., 187:1591–1603, 2005. https://doi.org/10.1128/JB.187.5.1591-1603.2005.

[9] R. Hengge. Proteolysis of *σ^S^* (RpoS) and the general stress response in *Escherichia coli*. Res Microbiol, 160:667–676, 2009. https://doi.org/10.1016/j.resmic.2009.08.014.

[10] T. Gaal, W. Ross, S. T. Estrem, L. H. Nguyen, R. R. Burgess, and R. L. Gourse. Promoter recognition and discrimination by E*σ*^S^ RNA polymerase. Mol. Microbiol., 42:939–954, 2001. https://doi.org/10.1046/j.1365-2958.2001.02703.x.

[11] A. Typas and R. Hengge. Role of the spacer between the −35 and −10 regions in *σ^s^* promoter selectivity in *Escherichia coli*. Mol. Microbiol., 59:1037–1051, 2006. https://doi.org/10.1111/j.1365-2958.2005.04998.x.

[12] A. Typas, G. Becker, and R. Hengge. The molecular basis of selective promoter activation by the *σ^S^* subunit of RNA polymerase. Mol. Microbiol., 63:1296–1306, 2007. https://doi.org/10.1111/j.1365-2958.2007.05601.x.

[13] B. Liu, Y. Zuo, and T. A. Steitz. Structures of *E. coli σ^S^* -transcription initiation complexes provide new insights into polymerase mechanism. Proc. Natl. Acad. Sci. USA, 113:4051–4056, 2016. https://doi.org/10.1073/pnas.1520555113.

[14] M. Espinosa-Urgel and A. Tormo. *σ^s^*-dependent promoters in *Escherichia coli* are located in DNA regions with intrinsic curvature. Nucleic Acids Res., 21:3667–3670, 1993. https://doi.org/10.1093/nar/21.16.3667.

[15] M. Espinosa-Urgel, C. Chamizo, and A. Tormo. A consensus structure for *σ^S^* -dependent promoters. Mol. Microbiol., 21:657–659, 1996. https://doi.org/10.1111/j.1365-2958.1996.tb02573.x.

[16] R. Hengge-Aronis. Stationary phase gene regulation: what makes an *Escherichia coli* promoter *σ*^S^-selective? Curr Opin Microbiol, 5:591–595, 2002. https://doi.org/10.1016/S1369-5274(02)00372-7.

[17] A. Typas and R. Hengge. Differential ability of *σ^s^* and *σ*^70^ of *Escherichia coli* to utilize promoters containing half or full UP-element sites. Mol. Microbiol., 55:250–260, 2005. https://doi.org/10.1111/j.1365-2958.2004.04382.x.

[18] A. Maciag, C. Peano, A. Pietrelli, T. Egli, G. De Bellis, and P. Landini. *In vitro* transcription profiling of the *σ^S^* subunit of bacterial RNA polymerase: re-definition of the *σ^S^* regulon and identification of *σ^S^* specific promoter sequence elements. Nucleic Acids Res., 39:5338–5355, 2011. https://doi.org/10.1093/nar/gkr129.

[19] A. Z. Rosenthal, M. Hu, and J. D. Gralla. Osmolyte-induced transcription: −35 region elements and recognition by sigma38 (rpoS). Mol. Microbiol., 59:1052–1061, 2006. https://doi.org/10.1111/j.1365-2958.2005.04999.x.

[20] C. E. Shannon. A Mathematical Theory of Communication. Bell System Tech. J., 27:379–423, 623–656, 1948. https://doi.org/10.1002/j.1538-7305.1948.tb01338.x https://doi.org/10.1002/j.1538-7305.1948.tb00917.x.

[21] J. R. Pierce. An Introduction to Information Theory: Symbols, Signals and Noise. Dover Publications, Inc., NY, 2nd edition, 1980. http://store.doverpublications.com/0486240614.html, https://www.amazon.com/gp/product/0486240614/, https://archive.org/details/symbolssignalsan002575mbp.

[22] T. D. Schneider, G. D. Stormo, L. Gold, and A. Ehrenfeucht. Information content of binding sites on nucleotide sequences. J. Mol. Biol., 188:415–431, 1986. https://doi.org/10.1016/0022-2836(86)90165-8, https://alum.mit.edu/www/toms/papers/schneider1986/.

[23] T. D. Schneider. Theory of molecular machines. II. Energy dissipation from molecular machines. J. Theor. Biol., 148:125–137, 1991. https://doi.org/10.1016/S0022-5193(05)80467-9, https://alum.mit.edu/www/toms/papers/edmm/.

[24] T. D. Schneider. 70% efficiency of bistate molecular machines explained by information theory, high dimensional geometry and evolutionary convergence. Nucleic Acids Res., 38:5995–6006, 2010. https://doi.org/doi:10.1093/nar/gkq389, https://alum.mit.edu/www/toms/papers/emmgeo/.

[25] T. D. Schneider and R. M. Stephens. Sequence logos: A new way to display consensus sequences. Nucleic Acids Res., 18:6097–6100, 1990. https://doi.org/10.1093/nar/18.20.6097, https://alum.mit.edu/www/toms/papers/logopaper/.

[26] T. D. Schneider. Sequence walkers: a graphical method to display how binding proteins interact with DNA or RNA sequences. Nucleic Acids Res., 25:4408–4415, 1997. https://doi.org/10.1093/nar/25.21.4408, https://alum.mit.edu/www/toms/papers/walker/, erratum: NAR 26(4): 1135, 1998.

[27] T. D. Schneider and J. Spouge. Information content of individual genetic sequences. J. Theor. Biol., 189:427–441, 1997. https://doi.org/10.1006/jtbi.1997.0540, https://alum.mit.edu/www/toms/papers/ri/.

[28] R. K. Shultzaberger, L. R. Roberts, I. G. Lyakhov, I. A. Sidorov, A. G. Stephen, R. J. Fisher, and T. D. Schneider. Correlation between binding rate constants and individual information of *E. coli* Fis binding sites. Nucleic Acids Res., 35:5275–5283, 2007. https://doi.org/10.1093/nar/gkm471, https://alum.mit.edu/www/toms/papers/fisbc/.

[29] N. S. Miroslavova and S. J. Busby. Investigations of the modular structure of bacterial promoters. Biochem Soc Symp, 73:1–10, 2006. https://doi.org/10.1042/bss0730001.

[30] M. Tribus. Thermostatics and Thermodynamics. D. van Nostrand Company, Inc., Princeton, N. J., 1961.

[31] R. K. Shultzaberger, R. E. Bucheimer, K. E. Rudd, and T. D. Schneider. Anatomy of *Escherichia coli* Ribosome Binding Sites. J. Mol. Biol., 313:215–228, 2001. https://doi.org/10.1006/jmbi.2001.5040, https://alum.mit.edu/www/toms/papers/flexrbs/.

[32] H. Salgado, M. Peralta-Gil, S. Gama-Castro, A. Santos-Zavaleta, L. Muniz-Rascado, J. S. Garcia-Sotelo, V. Weiss, H. Solano-Lira, I. Martinez-Flores, A. Medina-Rivera, G. Salgado-Osorio, S. Alquicira-Hernandez, K. Alquicira-Hernandez, A. Lopez-Fuentes, L. Porron-Sotelo, A. M. Huerta, C. Bonavides-Martinez, Y. I. Balderas-Martinez, L. Pannier, M. Olvera, A. Labastida, V. Jimenez-Jacinto, L. Vega-Alvarado, V. Del Moral-Chavez, A. Hernandez-Alvarez, E. Morett, and J. Collado-Vides. RegulonDB v8.0: omics data sets, evolutionary conservation, regulatory phrases, cross-validated gold standards and more. Nucleic Acids Res., 41:D203–13, 2013. https://doi.org/10.1093/nar/gks1201.

[33] T. D. Schneider and D. N. Mastronarde. Fast multiple alignment of ungapped DNA sequences using information theory and a relaxation method. Discrete Applied Mathematics, 71:259–268, 1996. https://alum.mit.edu/www/toms/papers/malign, http://www.ncbi.nlm.nih.gov/pmc/articles/PMC2785095/, https://doi.org/10.1016/S0166-218X(96)00068-6.

[34] T. D. Schneider. Reading of DNA sequence logos: Prediction of major groove binding by information theory. Meth. Enzym., 274:445–455, 1996. https://alum.mit.edu/www/toms/papers/oxyr/, https://doi.org/10.1016/S0076-6879(96)74036-3.

[35] W. Ross, J. F. Thompson, J. T. Newlands, and R. L. Gourse. *E.coli* Fis protein activates ribosomal RNA transcription *in vitro* and *in vivo*. EMBO J, 9:3733–3742, 1990. https://doi.org/10.1002/j.1460-2075.1990.tb07586.x.

[36] Y. N. Zhou and D. J. Jin. The *rpoB* mutants destabilizing initiation complexes at stringently controlled promoters behave like “stringent” RNA polymerases in *Escherichia coli*. Proc. Natl. Acad. Sci. USA, 95:2908–2913, 1998. https://doi.org/10.1073/pnas.95.6.2908.

[37] H. Zhi, W. Yang, and D. J. Jin. *Escherichia coli* proteins eluted from mono Q chromatography, a final step during RNA polymerase purification procedure. Methods Enzymol, 370:291–300, 2003. https://doi.org/10.1016/S0076-6879(03)70026-3.

[38] K. Tanaka, Y. Takayanagi, N. Fujita, A. Ishihama, and H. Takahashi. Heterogeneity of the principal *σ* factor in *Escherichia coli*: the *rpoS* gene product, *σ*^38^, is a second principal *σ* factor of RNA polymerase in stationary-phase *Escherichia coli*. Proc. Natl. Acad. Sci. USA, 90:3511–3515, 1993. https://doi.org/10.1073/pnas.90.8.3511.

[39] H. Zhi and D. J. Jin. Purification of highly-active and soluble *Escherichia coli σ*^70^ polypeptide overproduced at low temperature. Methods Enzymol, 370:174–180, 2003. https://doi.org/10.1016/S0076-6879(03)70015-9.

[40] C. Q. Pan, S. E. Finkel, S. E. Cramton, J. A. Feng, D. S. Sigman, and R. C. Johnson. Variable structures of Fis-DNA complexes determined by flanking DNA-protein contacts. J. Mol. Biol., 264:675–695, 1996. https://doi.org/10.1006/jmbi.1996.0669.

[41] Y. Zuo and T. A. Steitz. Crystal Structures of the *E. coli* Transcription Initiation Complexes with a Complete Bubble. Mol Cell, 58:534–540, 2015. https://doi.org/10.1016/j.molcel.2015.03.010.

[42] T. D. Schneider. Strong minor groove base conservation in sequence logos implies DNA distortion or base flipping during replication and transcription initiation. Nucleic Acids Res., 29:4881–4891, 2001. https://doi.org/10.1093/nar/29.23.4881, https://alum.mit.edu/www/toms/papers/baseflip/.

[43] I. G. Lyakhov, P. N. Hengen, D. Rubens, and T. D. Schneider. The P1 Phage Replication Protein RepA Contacts an Otherwise Inaccessible Thymine N3 Proton by DNA Distortion or Base Flipping. Nucleic Acids Res., 29:4892–4900, 2001. https://doi.org/10.1093/nar/29.23.4892, https://alum.mit.edu/www/toms/papers/repan3/.

[44] S. A. Darst, A. Feklistov, and C. A. Gross. Promoter melting by an alternative *σ*, one base at a time. Nat Struct Mol Biol, 21:350–351, 2014. https://doi.org/10.1038/nsmb.2798.

[45] N. C. Seeman, J. M. Rosenberg, and A. Rich. Sequence-specific recognition of double helical nucleic acids by proteins. Proc. Natl. Acad. Sci. USA, 73:804–808, 1976. https://doi.org/10.1073/pnas.73.3.804.

[46] J. T. Newlands, W. Ross, K. K. Gosink, and R. L. Gourse. Factor-independent activation of *Escherichia coli* rRNA transcription. II. characterization of complexes of *rrnB* P1 promoters containing or lacking the upstream activator region with *Escherichia coli* RNA polymerase. J. Mol. Biol., 220:569–583, 1991. https://doi.org/10.1016/0022-2836(91)90101-B.

[47] L. Rao, W. Ross, J. A. Appleman, T. Gaal, S. Leirmo, P. J. Schlax, M. T. Record Jr, and R. L. Gourse. Factor independent activation of *rrnB* P1. An “extended” promoter with an upstream element that dramatically increases promoter strength. J. Mol. Biol., 235:1421–1435, 1994. https://doi.org/10.1006/jmbi.1994.1098.

[48] C. A. Hirvonen, W. Ross, C. E. Wozniak, E. Marasco, J. R. Anthony, S. E. Aiyar, V. H. Newburn, and R. L. Gourse. Contributions of UP elements and the transcription factor FIS to expression from the seven *rrn* P1 promoters in *Escherichia coli*. J. Bacteriol., 183:6305–6314, 2001. https://doi.org/10.1128/JB.183.21.6305-6314.2001.

[49] P. N. Hengen, S. L. Bartram, L. E. Stewart, and T. D. Schneider. Information analysis of Fis binding sites. Nucleic Acids Res., 25:4994–5002, 1997. https://doi.org/10.1093/nar/25.24.4994, https://alum.mit.edu/www/toms/papers/fisinfo/.

[50] S. I. Husnain and M. S. Thomas. The UP Element Is Necessary but Not Sufficient for Growth Rate-Dependent Control of the *Escherichia coli guaB* Promoter. J. Bacteriol., 190:2450–2457, 2008. https://doi.org/10.1128/JB.01732-07.

[51] H. Tagami and H. Aiba. An inactive open complex mediated by an UP element at *Escherichia coli* promoters. Proc. Natl. Acad. Sci. USA, 96:7202–7207, 1999. https://doi.org/10.1073/pnas.96.13.7202.

[52] S. Lacour, A. Kolb, A. J. Boris Zehnder, and P. Landini. Mechanism of specific recognition of the *aidB* promoter by *σ^S^* -RNA polymerase. Biochem. Biophys. Res. Commun., 292:922–930, 2002. https://doi.org/10.1006/bbrc.2002.6744.

[53] T. D. Schneider. Evolution of biological information. Nucleic Acids Res., 28:2794–2799, 2000. https://doi.org/10.1093/nar/28.14.2794, https://alum.mit.edu/www/toms/papers/ev/.

[54] B. Lang, N. Blot, E. Bouffartigues, M. Buckle, M. Geertz, C. O. Gualerzi, R. Mavathur, G. Muskhelishvili, C. L. Pon, S. Rimsky, S. Stella, M. M. Babu, and A. Travers. High-affinity DNA binding sites for H-NS provide a molecular basis for selective silencing within proteobacterial genomes. Nucleic Acids Res., 35:6330–6337, 2007. https://doi.org/10.1093/nar/gkm712.

[55] L. H. Nguyen and R. R. Burgess. Comparative analysis of the interactions of *Escherichia coli σ^S^* and *σ*^70^ RNA polymerase holoenzyme with the stationary-phase-specific *bolAp1* promoter. Biochemistry, 36:1748–1754, 1997. https://doi.org/10.1021/bi961175h.

[56] W. Ross, A. Ernst, and R. L. Gourse. Fine structure of *E. coli* RNA polymerase-promoter interactions: α subunit binding to the UP element minor groove. Genes Dev, 15:491–506, 2001. https://doi.org/10.1101/gad.870001.

[57] P. P. Papp, D. K. Chattoraj, and T. D. Schneider. Information analysis of sequences that bind the replication initiator RepA. J. Mol. Biol., 233:219–230, 1993. https://doi.org/10.1006/jmbi.1993.1501 https://alum.mit.edu/www/toms/papers/helixrepa/.

[58] D. K. Chattoraj and T. D. Schneider. Replication control of plasmid P1 and its host chromosome: the common ground. Prog. Nucl. Acid Res. Mol. Biol., 57:145–186, 1997. http://www.sciencedirect.com/science/article/pii/S0079660308602809, https://doi.org/10.1016/S0079-6603(08)60280-9.

[59] S. T. Estrem, T. Gaal, W. Ross, and R. L. Gourse. Identification of an UP element consensus sequence for bacterial promoters. Proc. Natl. Acad. Sci. USA, 95:9761–9766, 1998. https://doi.org/10.1073/pnas.95.17.9761.

[60] K. Yasuno, T. Yamazaki, Y. Tanaka, T. S. Kodama, A. Matsugami, M. Katahira, A. Ishihama, and Y. Kyogoku. Interaction of the C-terminal domain of the *E. coli* RNA polymerase α subunit with the UP element: recognizing the backbone structure in the minor groove surface. J. Mol. Biol., 306:213–225, 2001. https://doi.org/10.1006/jmbi.2000.4369.

[61] P. N. Hengen, I. G. Lyakhov, L. E. Stewart, and T. D. Schneider. Molecular flip-flops formed by overlapping Fis sites. Nucleic Acids Res., 31(22):6663–6673, 2003. https://doi.org/10.1093/nar/gkg877.

[62] C. Peano, J. Wolf, J. Demol, E. Rossi, L. Petiti, G. De Bellis, J. Geiselmann, T. Egli, S. Lacour, and P. Landini. Characterization of the *Escherichia coli σ^S^* core regulon by Chromatin Immunoprecipitation-sequencing (ChIP-seq) analysis. Sci Rep, 5:10469, 2015. https://doi.org/10.1038/srep10469.

[63] M. R. Hemm, B. J. Paul, T. D. Schneider, G. Storz, and K. E. Rudd. Small membrane proteins found by comparative genomics and ribosome binding site models. Mol. Microbiol., 70:1487–1501, 2008. https://doi.org/10.1111/j.1365-2958.2008.06495.x, https://alum.mit.edu/www/toms/papers/smallproteins/.

[64] S. J. Lee and J. D. Gralla. Sigma38 (*rpoS*) RNA Polymerase Promoter Engagement via −10 Region Nucleotides. J. Biol. Chem., 276:30064–30071, 2001. https://doi.org/10.1074/jbc.M102886200.

[65] S. J. Lee and J. D. Gralla. Osmo-regulation of bacterial transcription via poised RNA polymerase. Mol Cell, 14:153–162, 2004. https://doi.org/10.1016/S1097-2765(04)00202-3.

[66] E. E. Blatter, W. Ross, H. Tang, R. L. Gourse, and R. H. Ebright. Domain organization of RNA polymerase α subunit: C-terminal 85 amino acids constitute a domain capable of dimerization and DNA binding. Cell, 78:889–896, 1994. https://doi.org/10.1016/S0092-8674(94)90682-3.

[67] B. Benoff, H. Yang, C. L. Lawson, G. Parkinson, J. Liu, E. Blatter, Y. W. Ebright, H. M. Berman, and R. H. Ebright. Structural basis of transcription activation: the CAP-αCTD-DNA complex. Science, 297:1562–1566, 2002. https://doi.org/10.1126/science.1076376.

[68] A. Ishihama. Subunit of assembly of *Escherichia coli* RNA polymerase. Adv Biophys, 14:1–35, 1981. https://www.sciencedirect.com/journal/advances-in-biophysics/issues.

[69] T. Gaal, W. Ross, E. E. Blatter, H. Tang, X. Jia, V. V. Krishnan, N. Assa-Munt, R. H. Ebright, and R. L. Gourse. DNA-binding determinants of the alpha subunit of RNA polymerase: novel DNA-binding domain architecture. Genes Dev, 10:16–26, 1996. https://doi.org/10.1101/gad.10.1.16.

[70] W. Ross, D. A. Schneider, B. J. Paul, A. Mertens, and R. L. Gourse. An intersubunit contact stimulating transcription initiation by E coli RNA polymerase: interaction of the alpha C-terminal domain and sigma region 4. Genes Dev, 17:1293–1307, 2003. https://doi.org/10.1101/gad.1079403.

[71] I. G. Hook-Barnard and D. M. Hinton. Transcription initiation by mix and match elements: flexibility for polymerase binding to bacterial promoters. Gene Regul Syst Bio, 1:275–293, 2007. https://doi.org/10.1177/117762500700100020.

[72] G. E. Crooks, G. Hon, J. M. Chandonia, and S. E. Brenner. WebLogo: a sequence logo generator. Genome Res., 14:1188–1190, 2004. https://doi.org/10.1101/gr.849004.

[73] B. Liu, C. Hong, R. K. Huang, Z. Yu, and T. A. Steitz. Structural basis of bacterial transcription activation. Science, 358:947–951, 2017. https://doi.org/10.1126/science.aao1923.

[74] L. J. Peck and J. C. Wang. Sequence dependence of the helical repeat of DNA in solution. Nature, 292:375–378, 1981. https://doi.org/10.1038/292375a0.

[75] K. S. Murakami. X-ray crystal structure of *Escherichia coli* RNA polymerase *σ*^70^ holoenzyme. J. Biol. Chem., 288:9126–9134, 2013. https://doi.org/10.1074/jbc.M112.430900.

[76] W. Humphrey, A. Dalke, and K. Schulten. VMD: visual molecular dynamics. J Mol Graph, 14:33–38, 27–28, 1996. http://www.ks.uiuc.edu/Research/vmd/, https://doi.org/10.1016/0263-7855(96)00018-5.

