## Supplement_for_sigma38_paper-directory for "*Escherichia coli σ*^38^ promoters use two UP elements instead of a −35 element: resolution of a paradox and discovery that *σ*^38^ transcribes ribosomal promoters": 169sigma38map.pdf

piece 1, NC_000913, adhP, config: linear, direction: -, begin: 1552172, end: 1552087

```
      *1552170  .        *1552160  .        *1552150  .        *1552140  .        *1552130  .        *1552120  .        *1552110  .        *1552100  .        *1552090
5' g t a c t g c c g g a a c t g a a a g a t a t t c a g a a a g t c t c c c g c g c a a t t g c g t t t g c g g t t g g c a a a a t g g c g c a g c a g c a a g g c g t g g c 3'
3' c a t g a c g g c c t t g a c t t t c t a t a a g t c t t t c a g a g g g c g c g t t a a c g c a a a c g c c a a c c g t t t t a c c g c g t c g t c g t t c c g c a c c g 5'
```

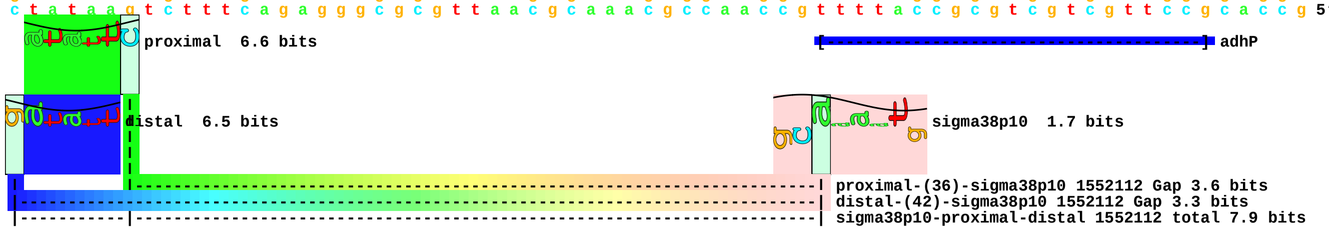

piece 2, NC_000913, amyA, config: linear, direction: +, begin: 2004085, end: 2004170

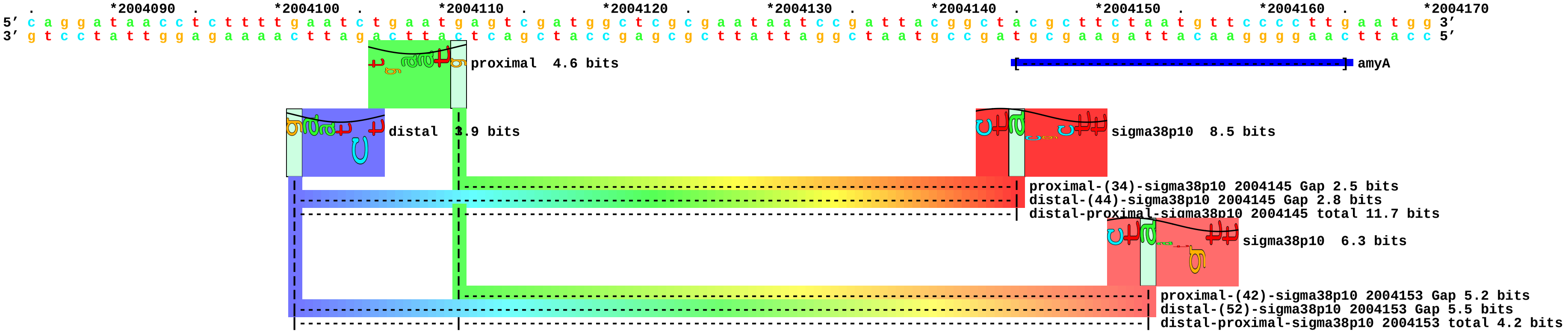

piece 3, NC_000913, dkgA, config: linear, direction: +, begin: 3154479, end: 3154564

```
     *3154480 .         *3154490 .         *3154500 .         *3154510 .         *3154520 .         *3154530 .         *3154540 .         *3154550 .         *3154560
5' t g g g c g a a a a t c a t g a c a t t a c g t t g g a t g t c a g c c g c c g t a t a t a c g a a g c c g c c c g c t a a g c t t t t t a c g c c t c a a a c t t t c g t 3'
3' a c c c g c t t t t a g t a c t g t a a t g c a a c c t a c a g t c g g c g g c a t a t a t g c t t c g g c g g g c g a t t c g a a a a a t g c g g a g t t t g a a a g c a 5'
```

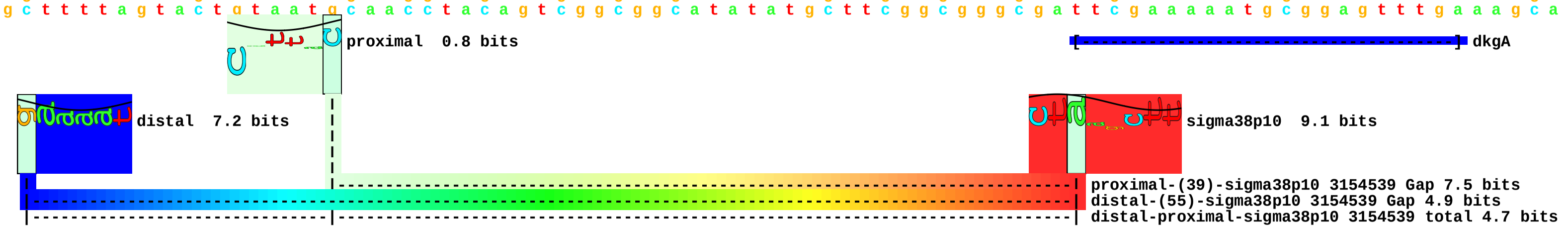

piece 4, NC_000913, gabD, config: linear, direction: +, begin: 2788861, end: 2788946

```
         .         *2788870  .         *2788880  .         *2788890  .         *2788900  .         *2788910  .         *2788920  .         *2788930  .         *2788940  .
5' t a g c g a c a c g c t g g a g a t t t t g g g c t c g t c g g g g a t t c g c c g g g t g c t g c a a a a c c a t c t a c g c t c a g g a c t g g g c g a g a t g a a a a 3'
3' a t c g c t g t g c g a c c t c t a a a a c c c g a g c a g c c c c t a a g c g g c c c a c g a c g t t t t g g t a g a t g c g a g t c c t g a c c c g c t c t a c t t t t 5'
```

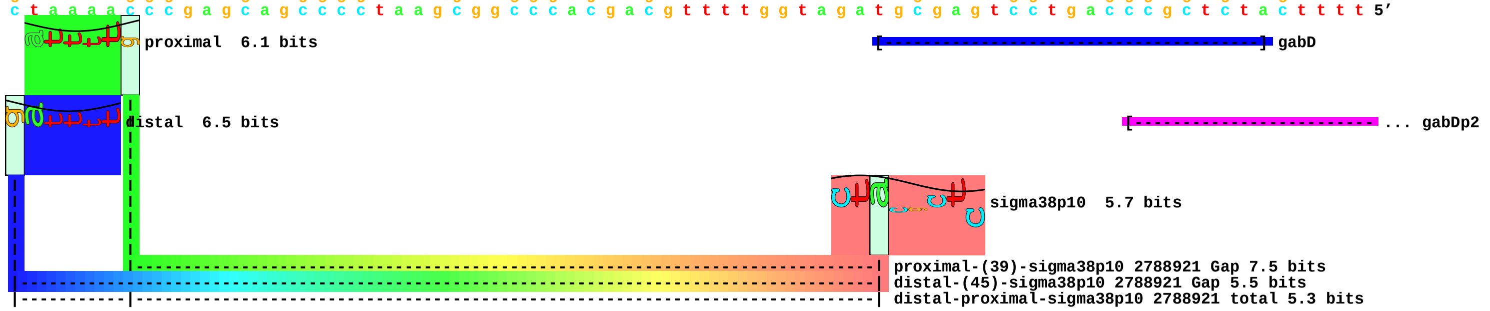

piece 5, NC_000913, hdhA, config: linear, direction: -, begin: 1696144, end: 1696059

```
             *1696140  .         *1696130  .         *1696120  .         *1696110  .         *1696100  .         *1696090  .         *1696080  .         *1696070  .         *1696060
5' c c t t t c c t c a a g g t t a a t g a t a a a a a a t g t g g c a c a c a g g c g g t g c c t t t t a t t t t t a c t g a a g t t c a c a g g a g g t t t a t g t g t t t 3'
3' g g a a a g g a g t t c c a a t t a c t a t t t t t t a c a c c g t g t g t c c g c c a c g g a a a a t a a a a a t g a c t t c a a g t g t c c t c c a a a t a c a c a a a a 5'
```

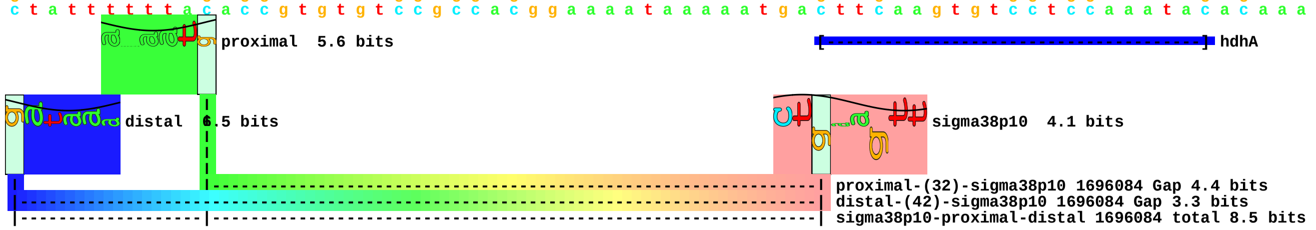

piece 6, NC_000913, narU, config: linear, direction: -, begin: 1542258, end: 1542173

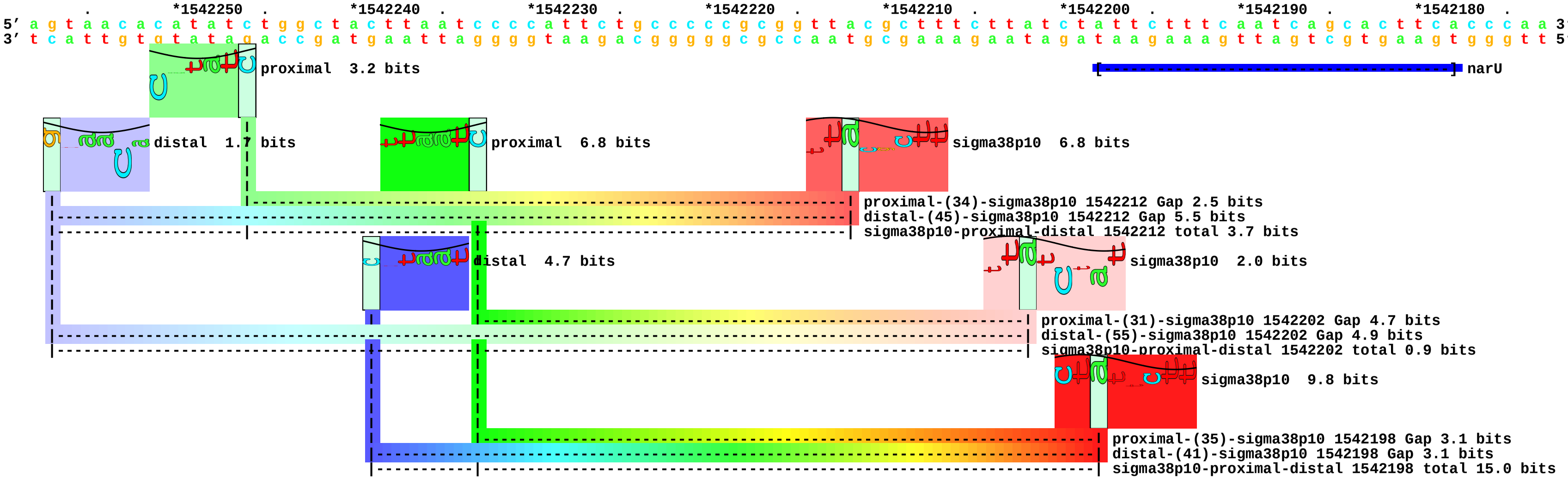

piece 7, NC_000913, narY, config: linear, direction: -, begin: 1537086, end: 1537001

```
     .         *1537080   .         *1537070   .         *1537060   .         *1537050   .         *1537040   .         *1537030   .         *1537020   .         *1537010   .
5' a c t g g c a t g c g c g g c g g c a t t c a t a a c t c g g t t a c c c g c g t t t g c c c g a a a c c a a c g c a t a t g a t t g g c g g t t a c g c g c a g c t g g c 3'
3' t g a c c g t a c g c g c c g c c g t a a g t a t t g a g c c a a t g g g c g c a a a c g g g c t t t g g t t g c g t a t a c t a a c c g c c a a t g c g c g t c g a c c g 5'
```

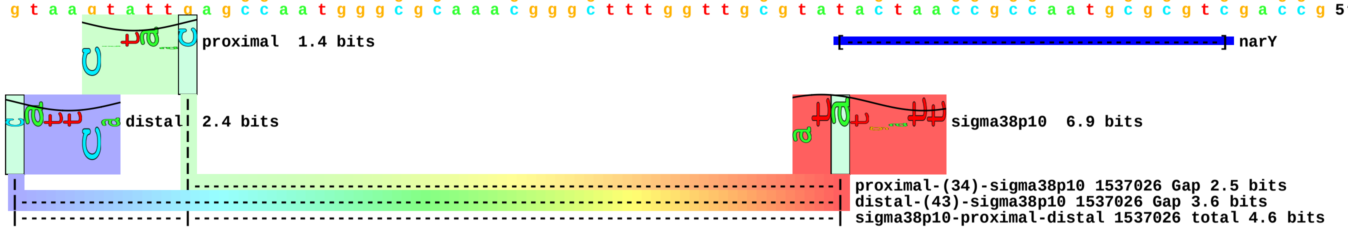

piece 8, NC_000913, poxB, config: linear, direction: -, begin: 910717, end: 910632

```
         .         *910710    .         *910700    .         *910690    .         *910680    .         *910670    .         *910660    .         *910650    .         *910640    .
5’ g a c g c g t t t c t t t a a a g a g a a a t t c t t c a c c c c a g t a g c a g a a g c g g c g a c c a g c g g t c t g a a a t t c a c c a a a c t g c a a c c g g c a c 3’
3’ c t g c g c a a a g a a a t t t c t c t t t a a g a a g t g g g g t c a t c g t c t t c g c c g c t g g t c g c c a g a c t t t a a g t g g t t t g a c g t t g g c c g t g 5’
```

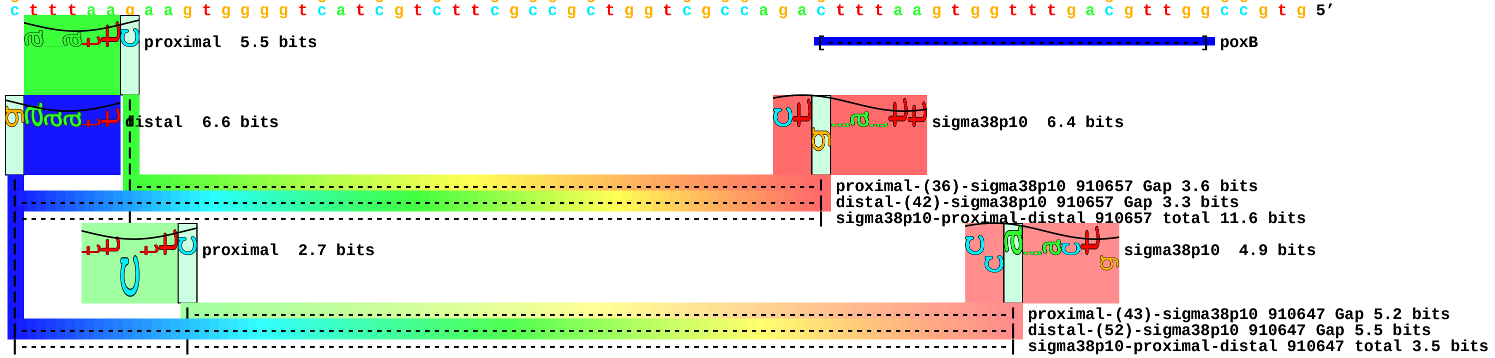

piece 9, NC_000913, qorA, config: linear, direction: -, begin: 4262362, end: 4262277

```
     *4262360  .         *4262350  .         *4262340  .         *4262330  .         *4262320  .         *4262310  .         *4262300  .         *4262290  .         *4262280
5’ t t g t t g a a g g g t t t a t t t c c t g c c a t a g t g a a t g g a g t t a c c g a g a t a a a g a a t g g g t c g a a a c t t t a t c a t a t g a a g c a g a c c c t 3’
3’ a a c a a c t t c c c a a a t a a a g g a c g g t a t c a c t t a c c t c a a t g g c t c t a t t t c t t a c c c a g c t t t g a a a t a g t a t a c t t c g t c t g g g a 5’
```

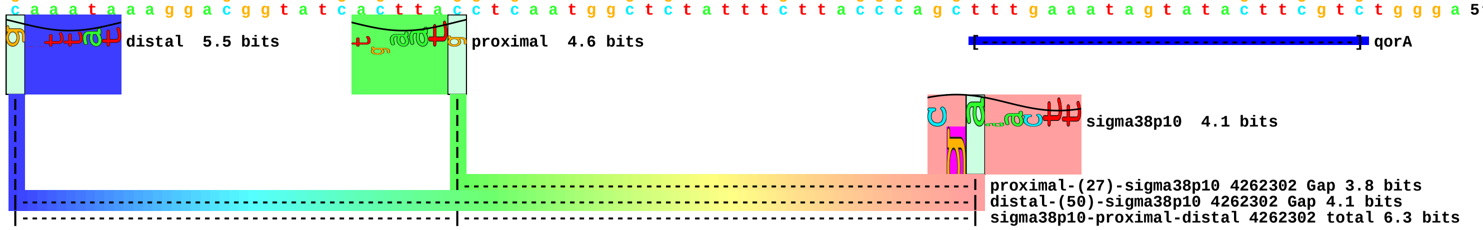

piece 10, NC_000913, talAp1, config: linear, direction: +, begin: 2576541, end: 2576626

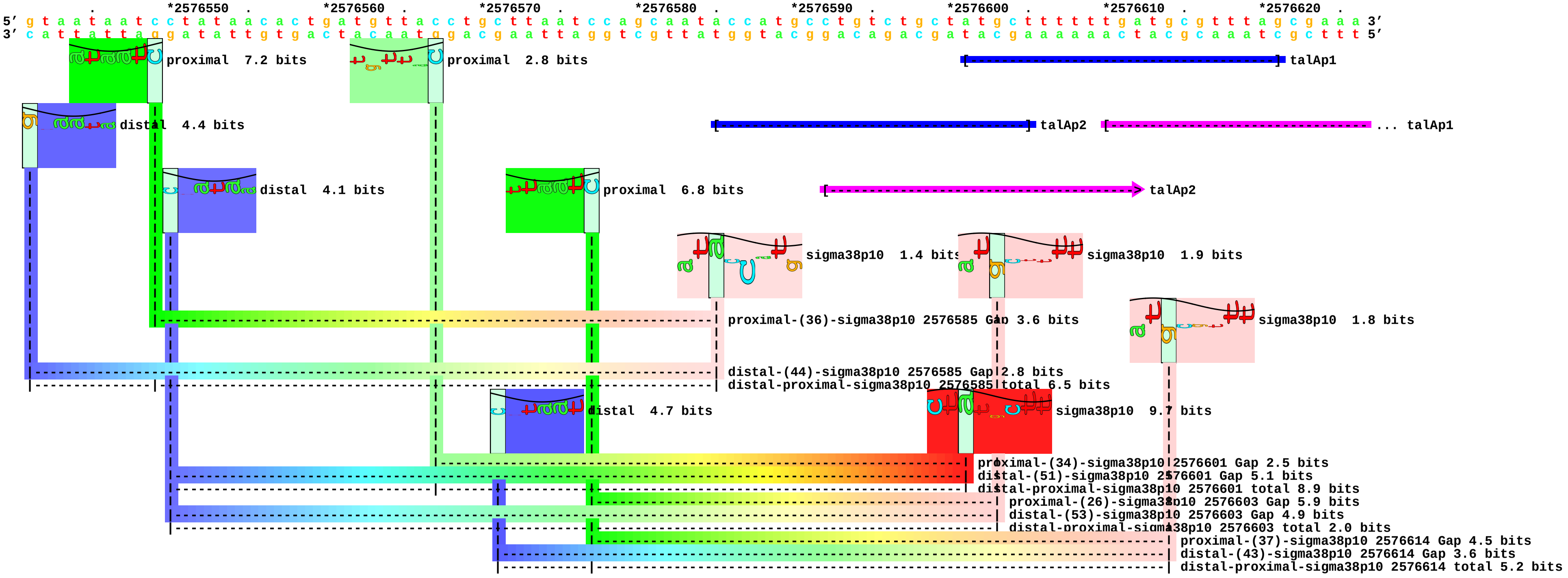

piece 11, NC_000913, talAp2, config: linear, direction: +, begin: 2576525, end: 2576610

```
   .         *2576530    .         *2576540    .         *2576550    .         *2576560    .         *2576570    .         *2576580    .         *2576590    .         *2576600    .         *2576610
5' t t a a t a t a a t g t t t g g t a a t a a t c c t a t a a c a c t g a t g t t a c c t g c t t a a t c c a g c a a t a c c a t g c c t g t c t g c t a t g c t t t t t t 3'
3' a a t t a t a t t a c a a a c c a t t a t t a g g a t a t t g t g a c t a c a a t g g a c g a a t t a g g t c g t t a t g g t a c g g a c a g a c g a t a c g a a a a a a 5'
```

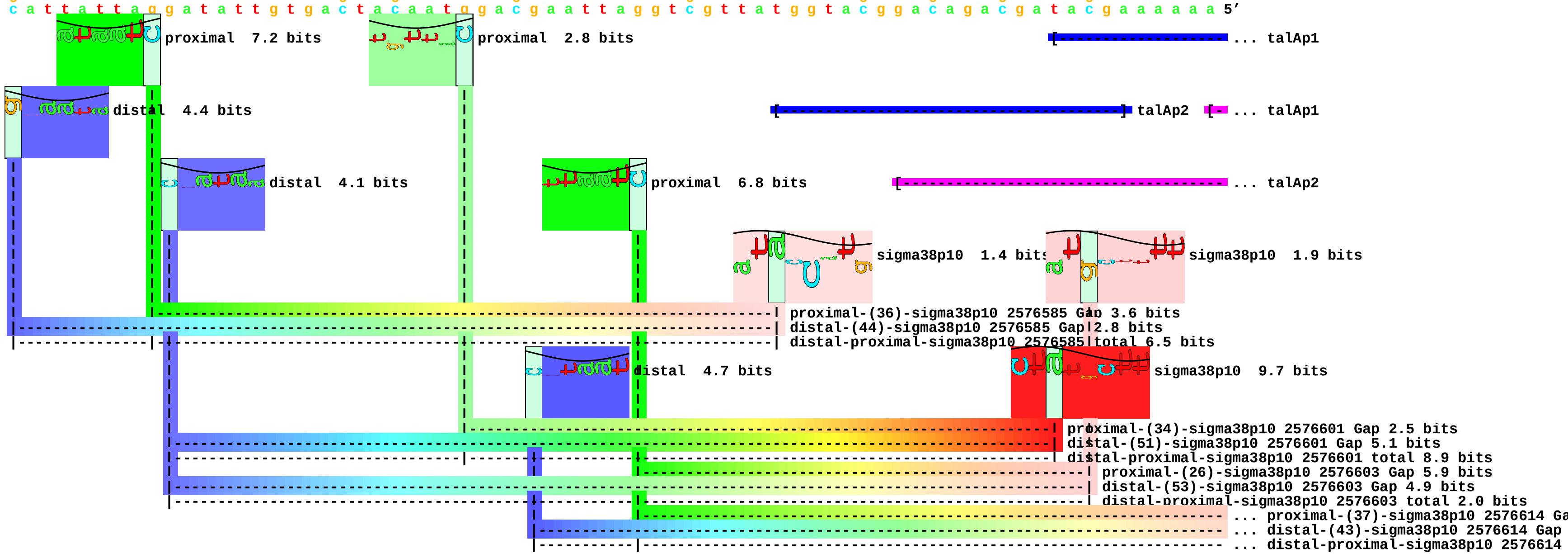

piece 12, NC_000913, tam, config: linear, direction: +, begin: 1604972, end: 1605057

```
         .         *1604980  .         *1604990  .         *1605000  .         *1605010  .         *1605020  .         *1605030  .         *1605040  .         *1605050  .
5' t g a a c t c t a t c t g a g t g a a a a a c a g t a a c t g c g g a t c t a a g g a g a a g a a t t a t g c a c g t c a c a c t g g t t g a a a t t a a c g t t c a t g a 3'
3' a c t t g a g a t a g a c t c a c t t t t t g t c a t t g a c g c c t a g a t t c c t c t t c t t a a t a c g t g c a g t g t g a c c a a c t t t a a t t g c a a g t a c t 5'
```

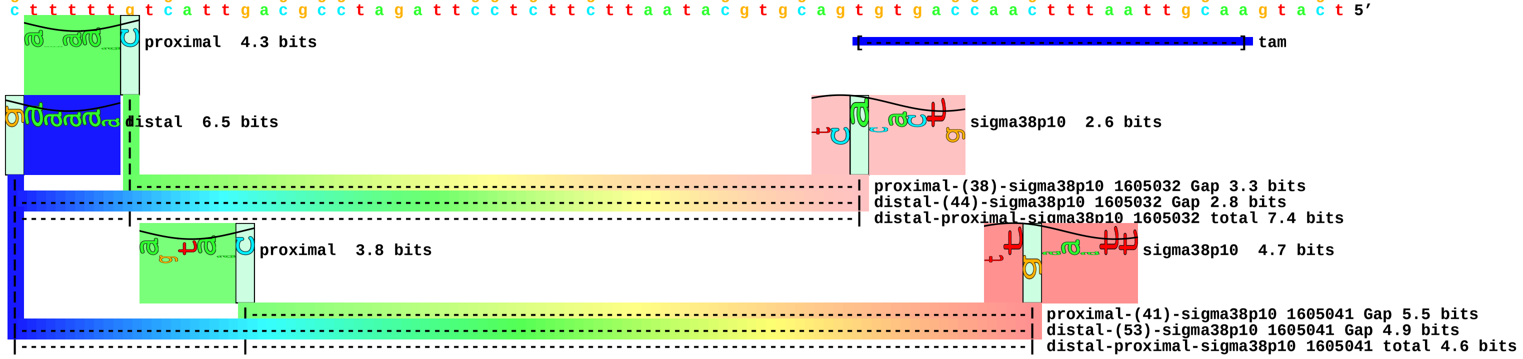

piece 13, NC_000913, tktB, config: linear, direction: +, begin: 2577285, end: 2577370

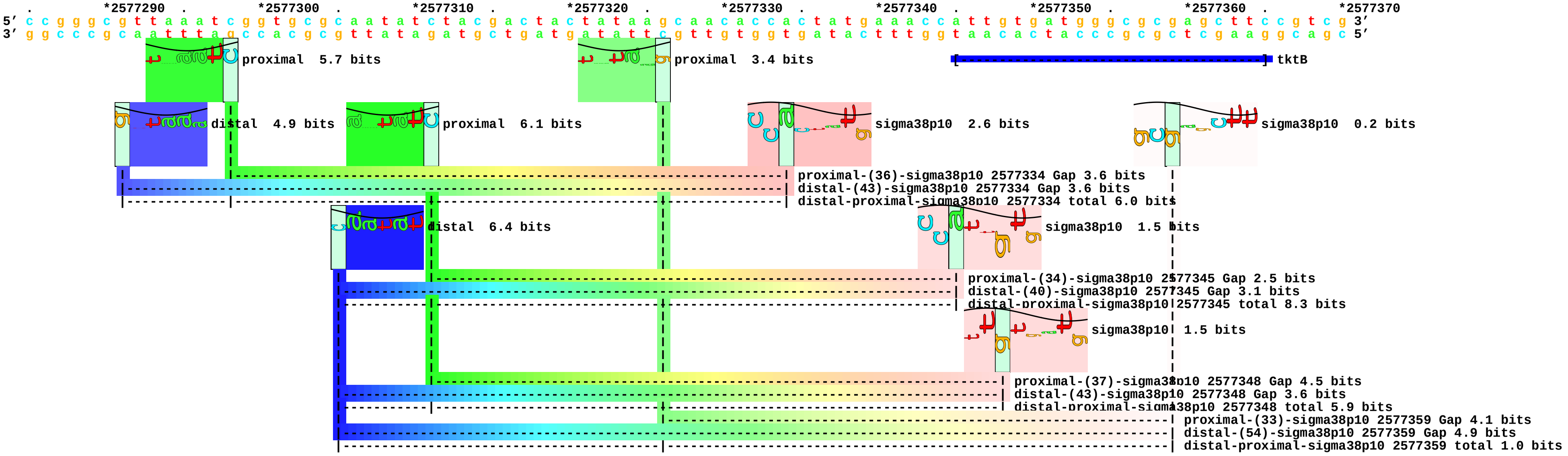

piece 14, NC_000913, ugpQ, config: linear, direction: -, begin: 3586403, end: 3586318

```
         *3586400  .         *3586390  .         *3586380  .         *3586370  .         *3586360  .         *3586350  .         *3586340  .         *3586330  .         *3586320
5' c t a c c g t c a g t a t g c c g g g c g t a a a a t g a c t c t c g g c a t t c g c c c g g a a c a t a t t g c g c t a a g c t c g c a g g c a g a a g g c g g c g t a c 3'
3' g a t g g c a g t c a t a c g g c c c g c a t t t t a c t g a g a g c c g t a a g c g g g c c t t g t a t a a c g c g a t t c g a g c g t c c g t c t t c c g c c g c a t g 5'
```

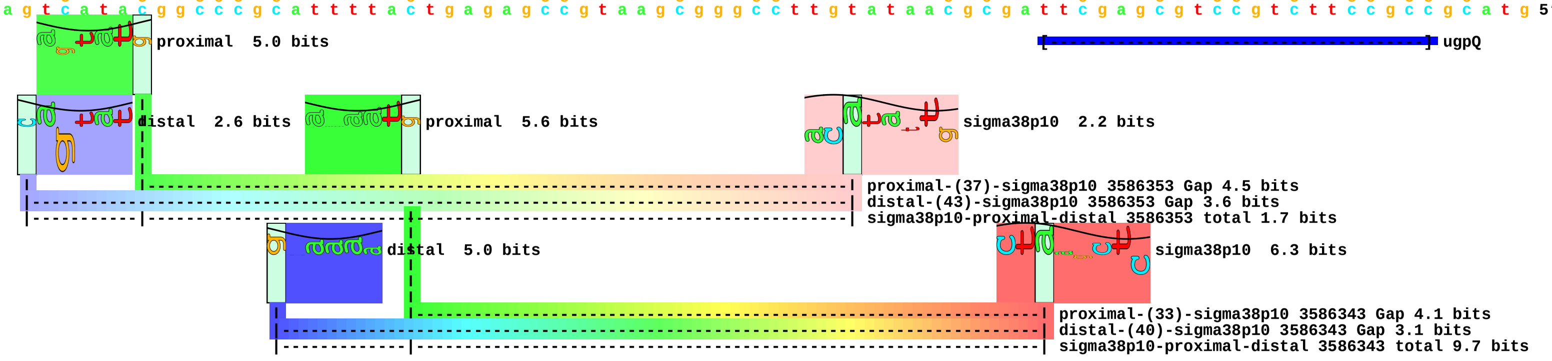

piece 15, NC_000913, glsA, config: linear, direction: +, begin: 510786, end: 510871

```
     *510790   .       *510800   .       *510810   .       *510820   .       *510830   .       *510840   .       *510850   .       *510860   .       *510870
5' a a g t t a a a t c a g g a t g c c t g a a a a t c g g c a c c g g g g t g a g g a a t t a c c t c c c g c a t c t a t a a a a a g g a g t t a a c a a a a g a t g t t a g 3'
3' t t c a a t t t a g t c c t a c g g a c t t t t a g c c g t g g c c c c a c t c c t t a a t g g a g g g c g t a g a t a t t t t t c c t c a a t t g t t t t c t a c a a t c 5'
```

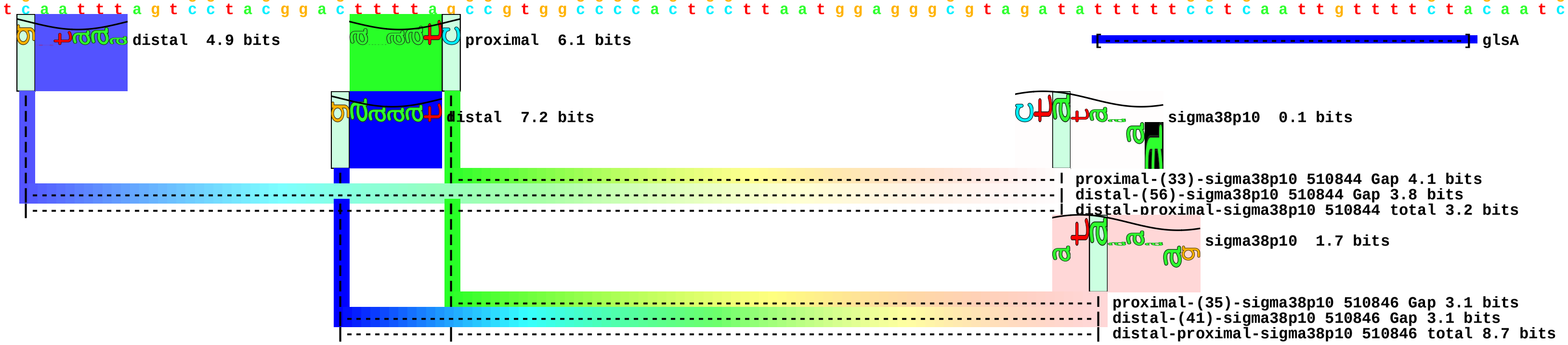

piece 16, NC_000913, ybaY, config: linear, direction: +, begin: 474458, end: 474543

```
      *474460   .       *474470   .       *474480   .       *474490   .       *474500   .       *474510   .       *474520   .       *474530   .       *474540
5' c g a t g g g t g c a a t g g t c t g a a t t a c g g g c t a a t t a c a g g c a g a a a t g c g t g a t g t g t g c c a c a c t t g t t g a c g t t a c t a t t t t g t t 3'
3' g c t a c c c a c g t t a c c a g a c t t a a t g c c c g a t t a a t g t c c g t c t t t a c g c a c t a c a c a c g g t g t g a a c a a c t g c a a t g a t a a a a c a a 5'
```

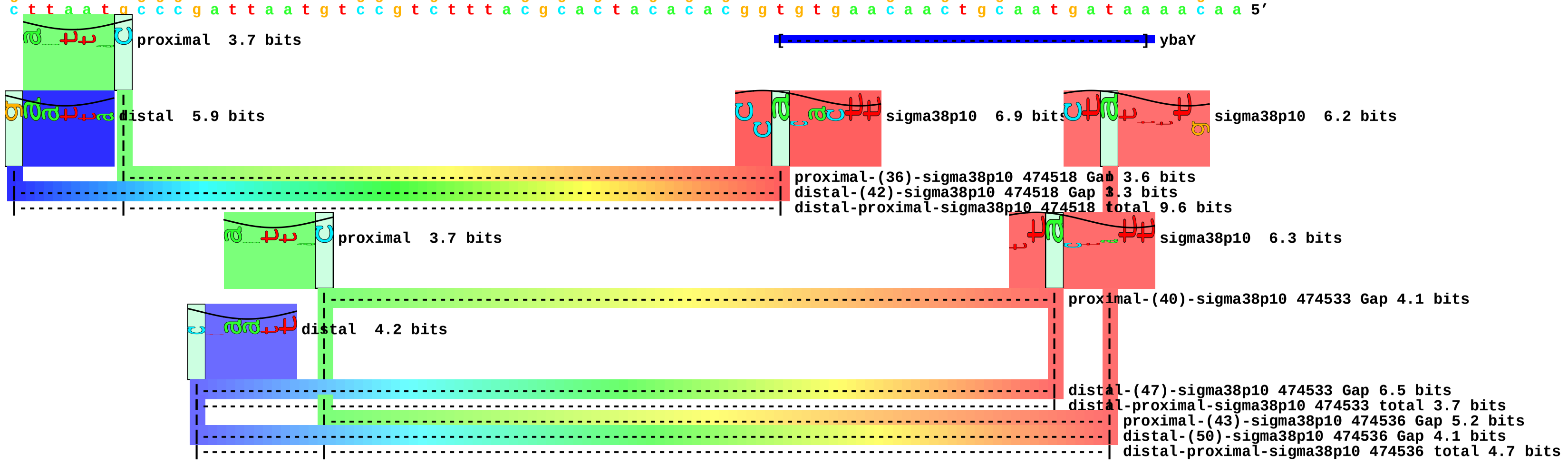

piece 17, NC_000913, ycaC, config: linear, direction: -, begin: 944910, end: 944825

```
   *944910   .       *944900   .       *944890   .       *944880   .       *944870   .       *944860   .       *944850   .       *944840   .       *944830   .
5' t t c a t a c g c t t c g t t c a g t c g a a a a a t t t c a t g g t c a c a g c a a a c t c a t a t t t c c c g t c t a t g c t t t c a g a g c a a t g c a t c g c g t c 3'
3' a a g t a t g c g a a g c a a g t c a g c t t t t t a a a g t a c c a g t g t c g t t t g a g t a t a a a g g g c a g a t a c g a a a g t c t c g t t a c g t a g c g c a g 5'
```

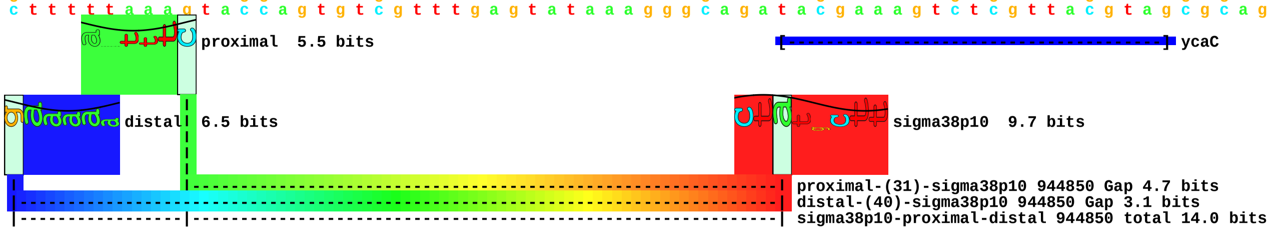

piece 18, NC_000913, yfcF, config: linear, direction: -, begin: 2418747, end: 2418662

```
        .         *2418740  .         *2418730  .         *2418720  .         *2418710  .         *2418700  .         *2418690  .         *2418680  .         *2418670  .
5' t t t c c c c a g g t c t a c c t t a a t c a a g c g a t a a t c c a g c t c t g c t t c t t c g a g a a a c a g c g t a a t t t t g t g a c c a t t g g g t g t c g g g g 3'
3' a a a g g g g t c c a g a t g g a a t t a g t t c g c t a t t a g g t c g a g a c g a a g a a g c t c t t t g t c g c a t t a a a a c a c t g g t a a c c c a c a g c c c c 5'
```

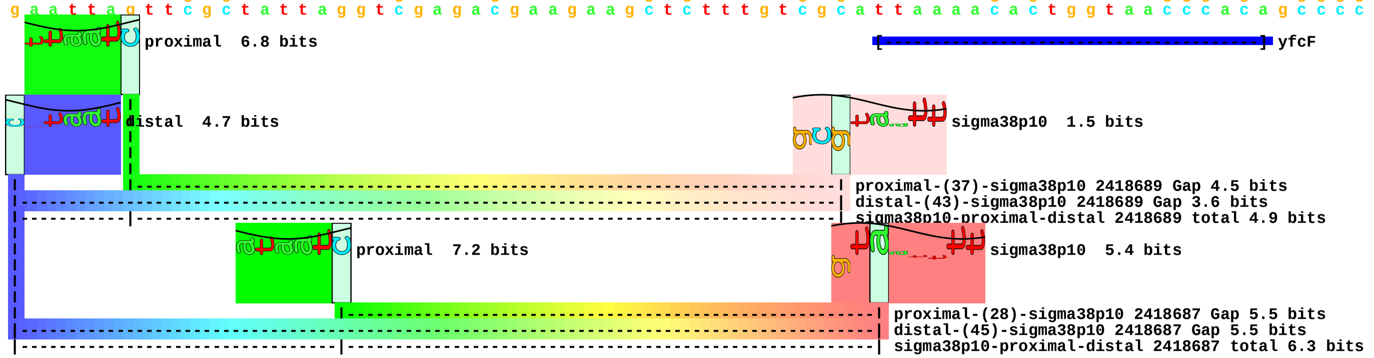

piece 19, NC_000913, patA, config: linear, direction: +, begin: 3217410, end: 3217495

```
   *3217410  .         *3217420  .         *3217430  .         *3217440  .         *3217450  .         *3217460  .         *3217470  .         *3217480  .         *3217490  .
5' a a c c g a a t t t g t g t t t a t c c c g a t t t t c g c g a t c g c a g c c g g a g t g g c g c a a t c c c t g c a a t a c t t a a a t c g g t a t c a t g t g a t a c 3'
3' t t g g c t t a a a c a c a a a t a g g g c t a a a a g c g c t a g c g t c g g c c t c a c c g c g t t a g g g a c g t t a t g a a t t t a g c c a t a g t a c a c t a t g 5'
```

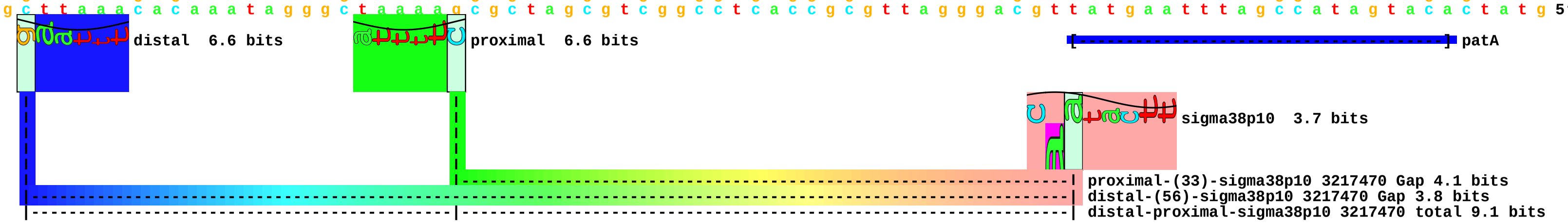

piece 20, NC_000913, yhfW, config: linear, direction: -, begin: 3507960, end: 3507875

```
   *3507960  .         *3507950  .         *3507940  .         *3507930  .         *3507920  .         *3507910  .         *3507900  .         *3507890  .         *3507880  .
5' g t g c t c t g t t t t c g c a c g c a g a t t t t t g t c a c c c g t a g c g a t g t g g t g c t g g t g t c c g g t a t t c a t c g t g g c g a a c c g g a a a t c g t 3'
3' c a c g a g a c a a a a g c g t g c g t c t a a a a a c a g t g g g c a t c g c t a c a c c a c g a c c a c a g g c c a t a a g t a g c a c c g c t t g g c c t t t a g c a 5'
```

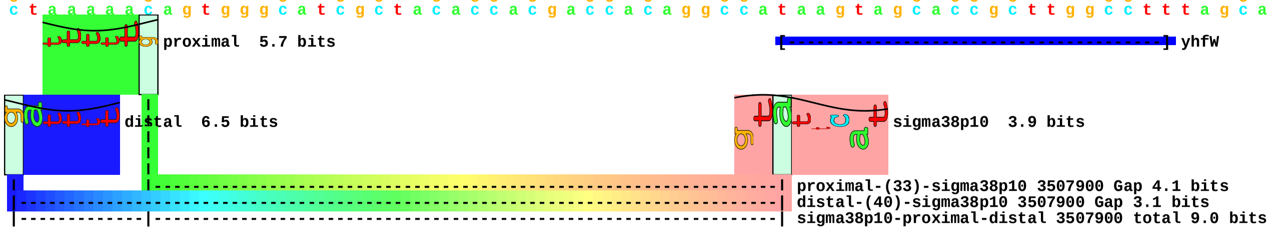

piece 21, NC_000913, yjcS, config: linear, direction: -, begin: 4304810, end: 4304725

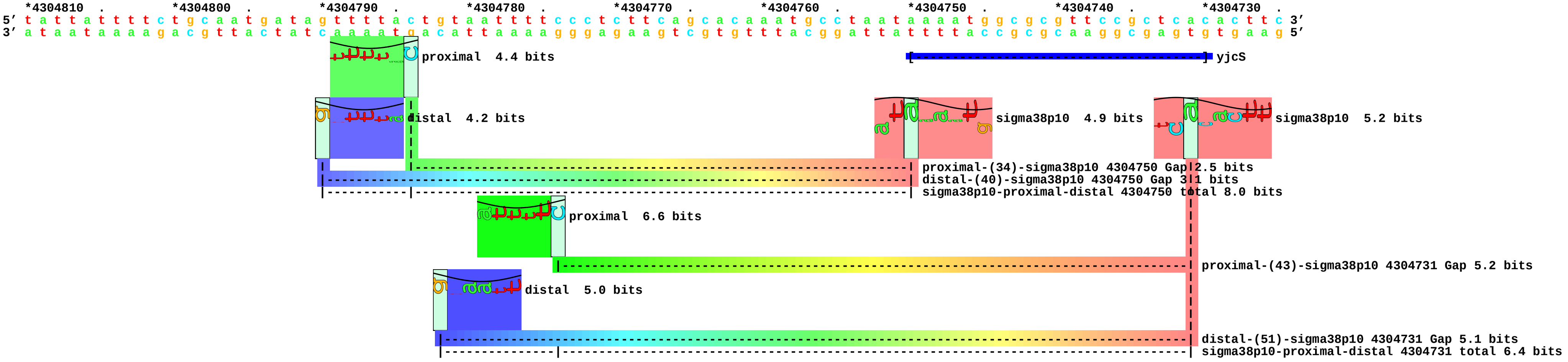

piece 22, NC_000913, yjgB, config: linear, direction: -, begin: 4494327, end: 4494242

```
        .         *4494320  .         *4494310  .         *4494300  .         *4494290  .         *4494280  .         *4494270  .         *4494260  .         *4494250  .
5' a t t g c a t a c a g c g a t g t g t a a c c t t t g t c a c a c t c c a g g c a c c c c g c c c t g c c a t g c t c t a c a c t t c c c a a a c a a c a c c a g a g a a g 3'
3' t a a c g t a t g t c g c t a c a c a t t g g a a a c a g t g t g a g g t c c g t g g g g c g g g a c g g t a c g a g a t g t g a a g g g t t t g t t g t g g t c t c t t c 5'
```

proximal 3.3 bits

yjgB

distal 2.4 bits

sigma38p10 9.7 bits

proximal-(39)-sigma38p10 4494267 Gap 7.5 bits
distal-(56)-sigma38p10 4494267 Gap 3.8 bits
sigma38p10-proximal-distal 4494267 total 4.2 bits

piece 23, NC_000913, yncG, config: linear, direction: +, begin: 1524179, end: 1524264

```
    *1524180  .       *1524190  .       *1524200  .       *1524210  .       *1524220  .       *1524230  .       *1524240  .       *1524250  .       *1524260
5' a t t a a t a c a a c t g a c g a a a a a a t g a c a a a t c c t t t t g c t g g t t a a c c t g t g t a c t g t c c t a c a c t t a a t c t t t a a a a g a t t g t g a g 3'
3' t a a t t a t g t t g a c t g c t t t t t t a c t g t t t a g g a a a a c g a c c a a t t g g a c a c a t g a c a g g a t g t g a a t t a g a a a t t t t c t a a c a c t c 5'
```

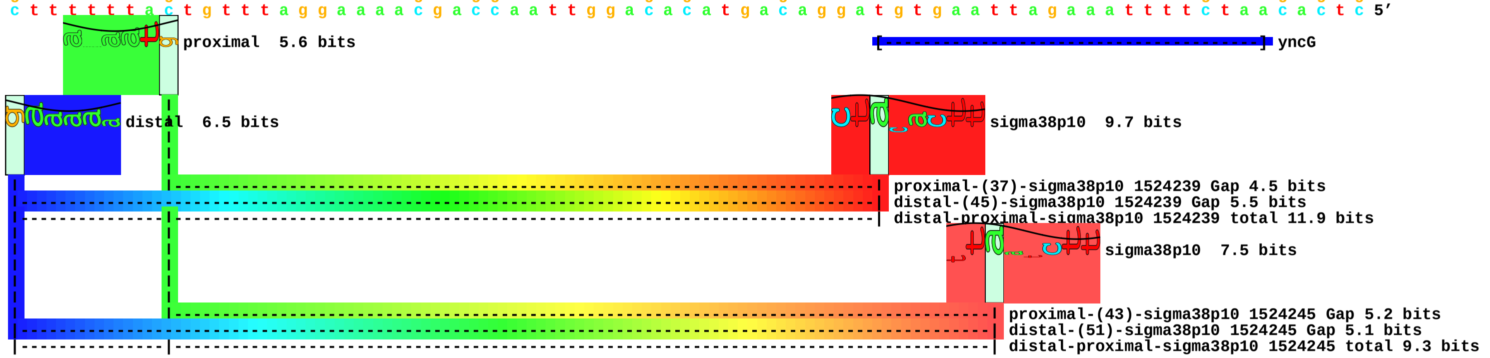

piece 24, NC_000913, ysgA, config: linear, direction: -, begin: 4014404, end: 4014319

```
          *4014400 .         *4014390 .         *4014380 .         *4014370 .         *4014360 .         *4014350 .         *4014340 .         *4014330 .         *4014320
5' t t a c c g a a g g g t t t a a c a t t t t t t c g t g a t a c t c a t c a c c a t g a c g c a a a t g c g t t g c a t a a a t t t a c c c a c c a a a a g t g a t g c a a 3'
3' a a t g g c t t c c c a a a t t g t a a a a a a g c a c t a t g a g t a g t g g t a c t g c g t t t a c g c a a c g t a t t t a a a t g g g t g g t t t t c a c t a c g t t 5'
```

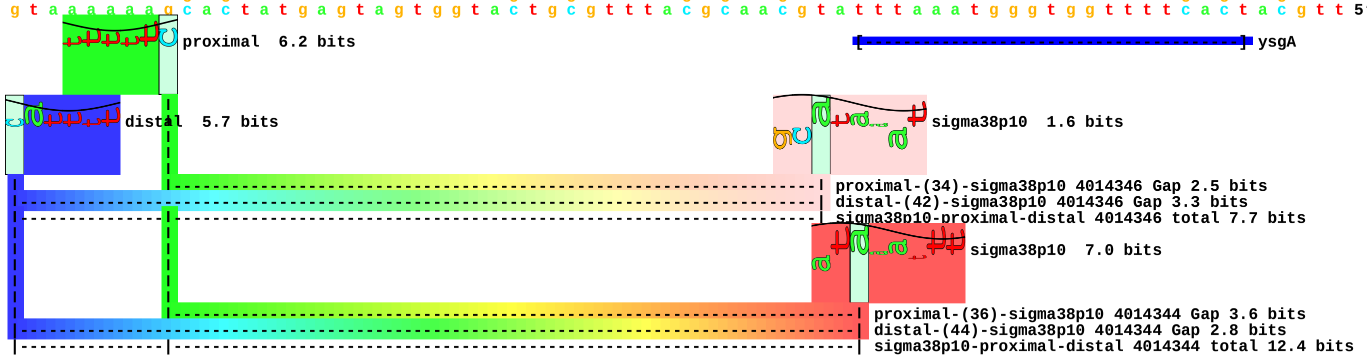

piece 25, NC_000913, bolA, config: linear, direction: +, begin: 453482, end: 453567

```
         .         *453490    .         *453500    .         *453510    .         *453520    .         *453530    .         *453540    .         *453550    .         *453560    .
5’ t a a a t t c a t c a t g t t g a c g a a a t a a t c g c c c c t g g t a a a a g a a a c a c t g a t g c g a g g c c t g t g t t t c a a t c t t t a a a t c a g t a a a c 3’
3’ a t t t a a g t a g t a c a a c t g c t t t a t t a g c g g g g a c c a t t t t c t t t g t g a c t a c g c t c c g g a c a c a a a g t t a g a a a t t t a g t c a t t t g 5’
```

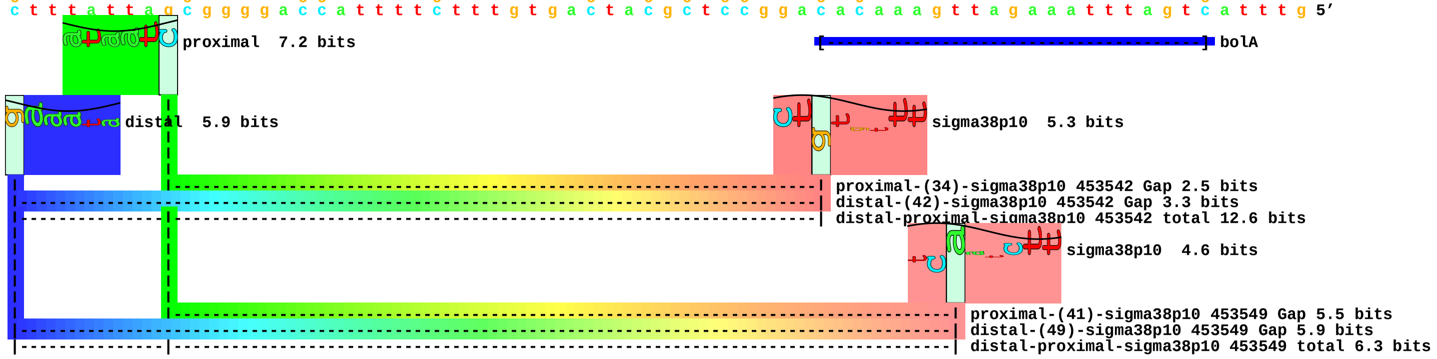

piece 26, NC_000913, chaB, config: linear, direction: +, begin: 1271217, end: 1271302

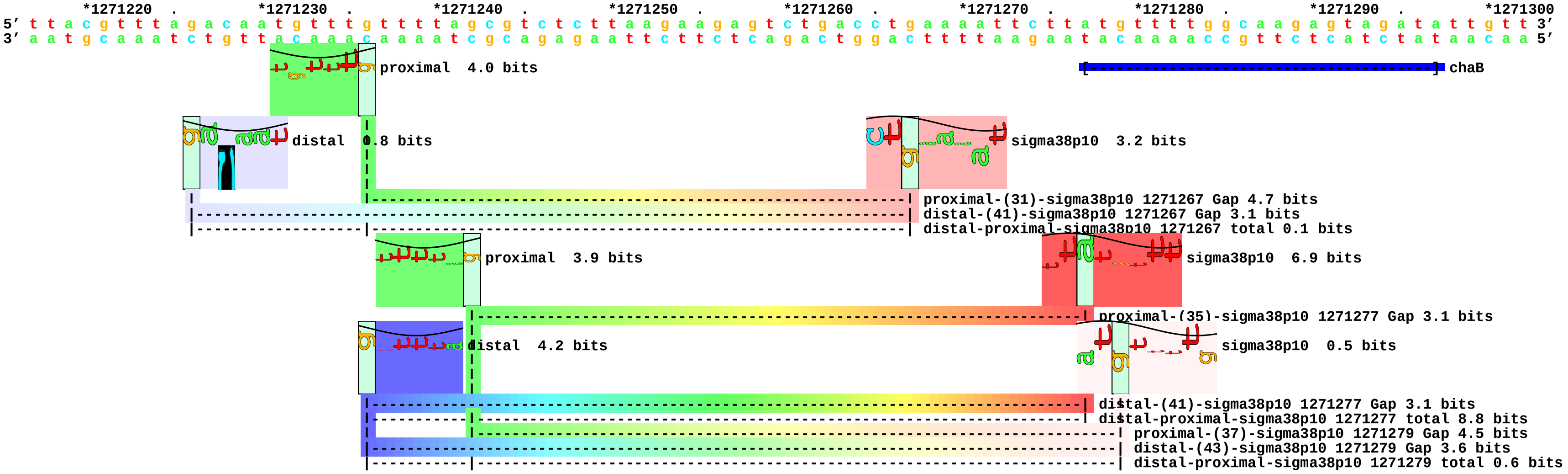

piece 27, NC_000913, csiR, config: linear, direction: +, begin: 2793576, end: 2793661

```
      *2793580  .         *2793590  .         *2793600  .         *2793610  .         *2793620  .         *2793630  .         *2793640  .         *2793650  .         *2793660
5' c g g c t t a t t t a g c g a t a g g g g a t t a t c t g t a c c g t g c c a a t t a t g g c t c g c t g g a a a a a g c t g g t a t t g t g g c a a a a a a c a c c c g t t c 3'
3' g c c g a a t a a a t c g c t a t c c c c t a a t a g a c a t g g c a c g g t t a a t a c c g a g c g a c c t t t t t c g a c c a t a a c a c c g t t t t t t g t g g g c a a g 5'
```

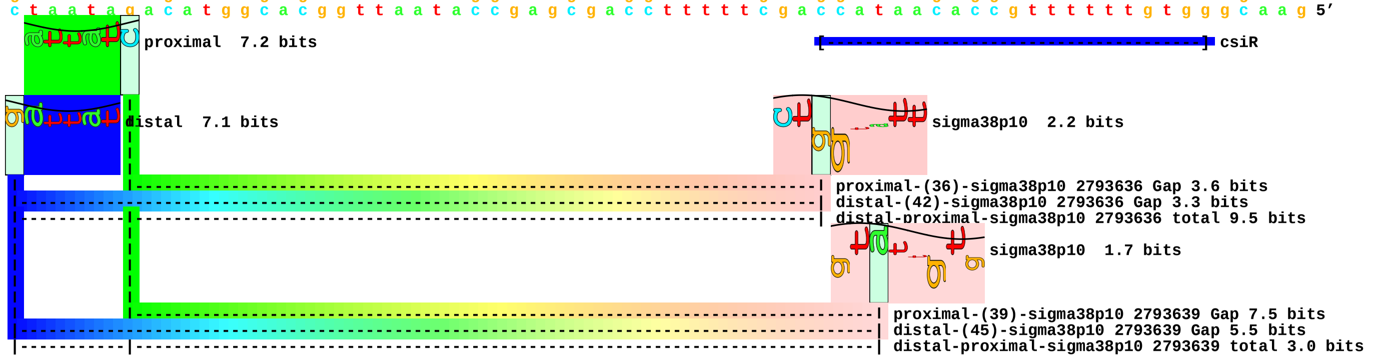

piece 28, NC_000913, gadW, config: linear, direction: -, begin: 3662746, end: 3662661

```
    .         *3662740  .         *3662730  .         *3662720  .         *3662710  .         *3662700  .         *3662690  .         *3662680  .         *3662670  .
5' t g g t c c t a c c a a a t c t g g c a g t t t t t g c g c t a a g a a a c a g t c t g g c a t c a t t t c a t t a g t a t a c t g a a a t t g a a a t a a t c g c a g t a 3'
3' a c c a g g a t g g t t t a g a c c g t c a a a a a c g c g a t t c t t t g t c a g a c c g t a g t a a a g t a a t c a t a t g a c t t t a a c t t t a t t a g c g t c a t 5'
```

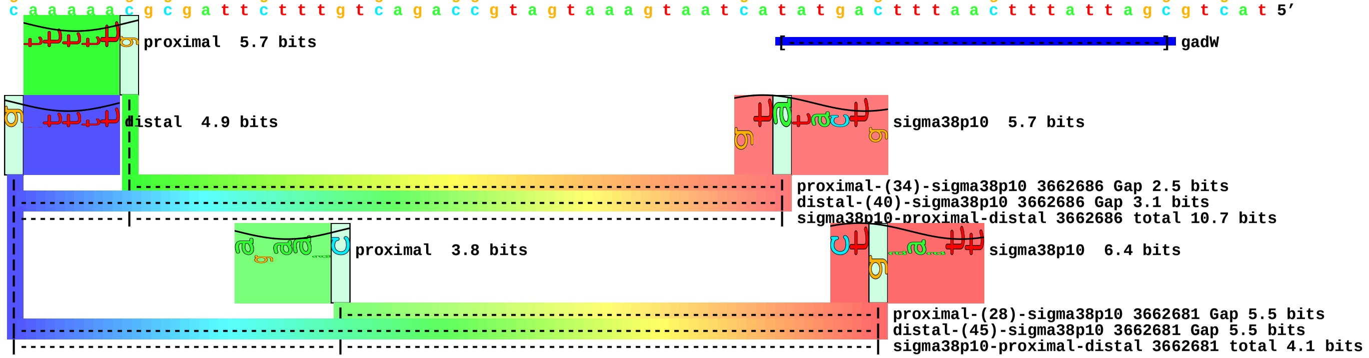

piece 29, NC_000913, gadX, config: linear, direction: -, begin: 3664079, end: 3663994

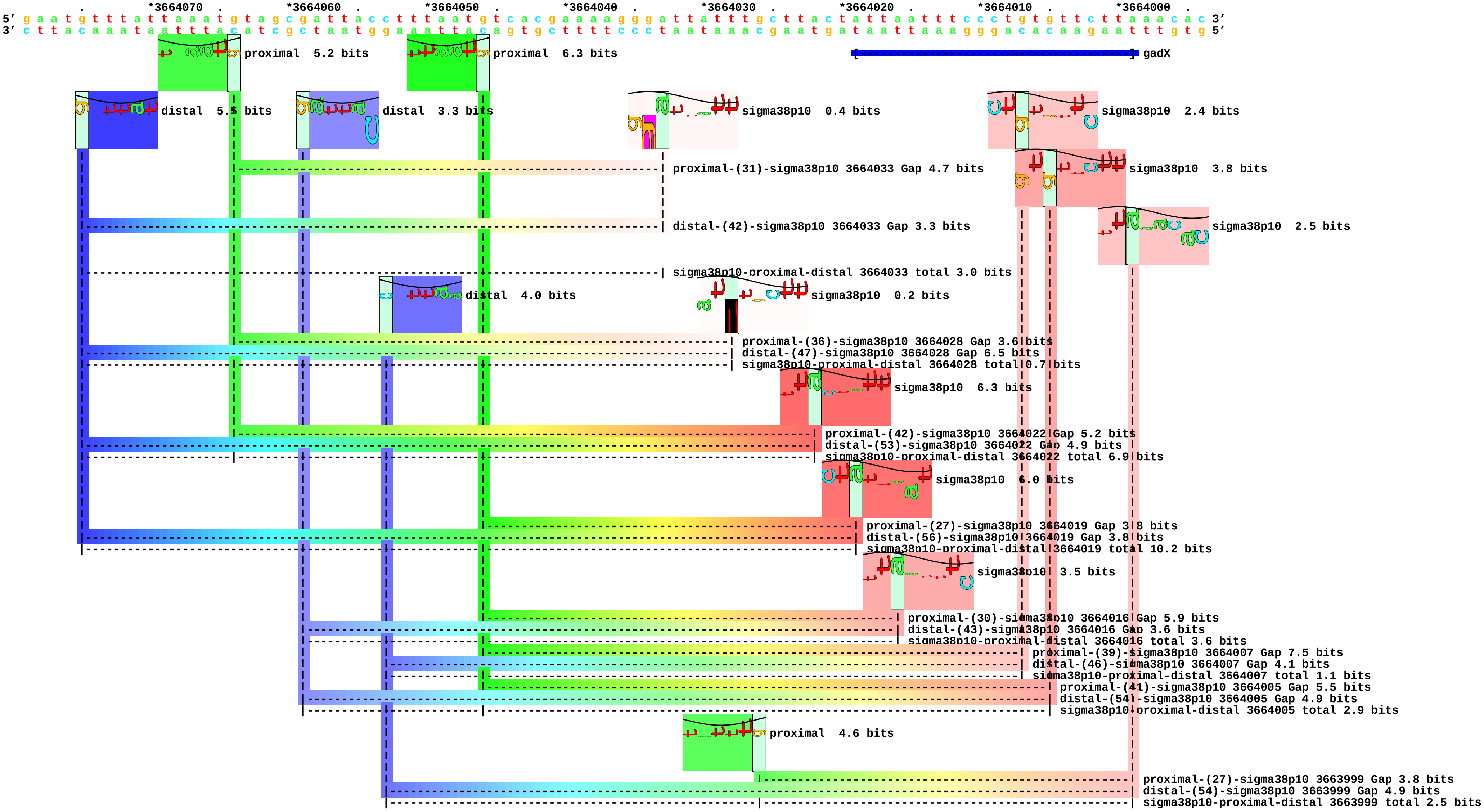

piece 30, NC_000913, gmr, config: linear, direction: -, begin: 1344938, end: 1344853

```
          .        *1344930  .        *1344920  .        *1344910  .        *1344900  .        *1344890  .        *1344880  .        *1344870  .        *1344860  .
5' t c c a g g c a g a t t t t t t a a c t g a c t t t c g t t t g a a a a c t g g c g t t t t t c c a a a a c t g g a t t a c a a a t a a a t a t a t a a a a a a a a c a a t 3'
3' a g g t c c g t c t a a a a a a t t g a c t g a a a g c a a a c t t t g a c c g c a a a a a g g t t t t g a c c t a a t g t t t a t t t a t a t a t t t t t t t t g t t a 5'
```

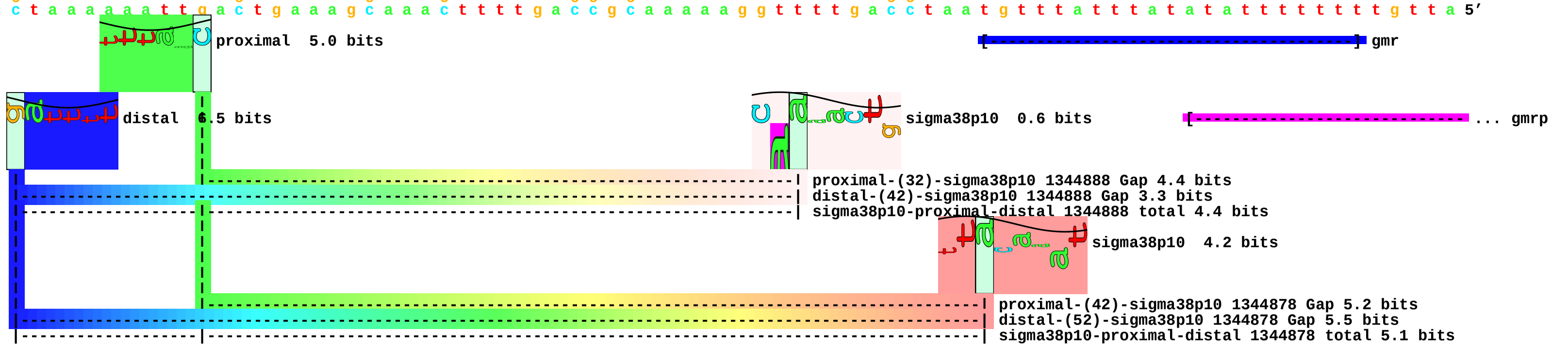

piece 31, NC_000913, pdhR, config: linear, direction: +, begin: 121851, end: 121936

```
               .         *121860    .         *121870    .         *121880    .         *121890    .         *121900    .         *121910    .         *121920    .         *121930    .
5' t t c a a t t c g g a t t t t t a t a g t t t a a t a a t c g t t a a a a a a c t c c t t t c c t a c g t a a a g t c t a c a t t t g t g c a t a g t t a c a a c t t t g a 3'
3' a a g t t a a g c c t a a a a a t a t c a a a t t a t t a g c a a t t t t t t g a g g a a a g g a t g c a t t t c a g a t g t a a a c a c g t a t c a a t g t t g a a a c t 5'
```

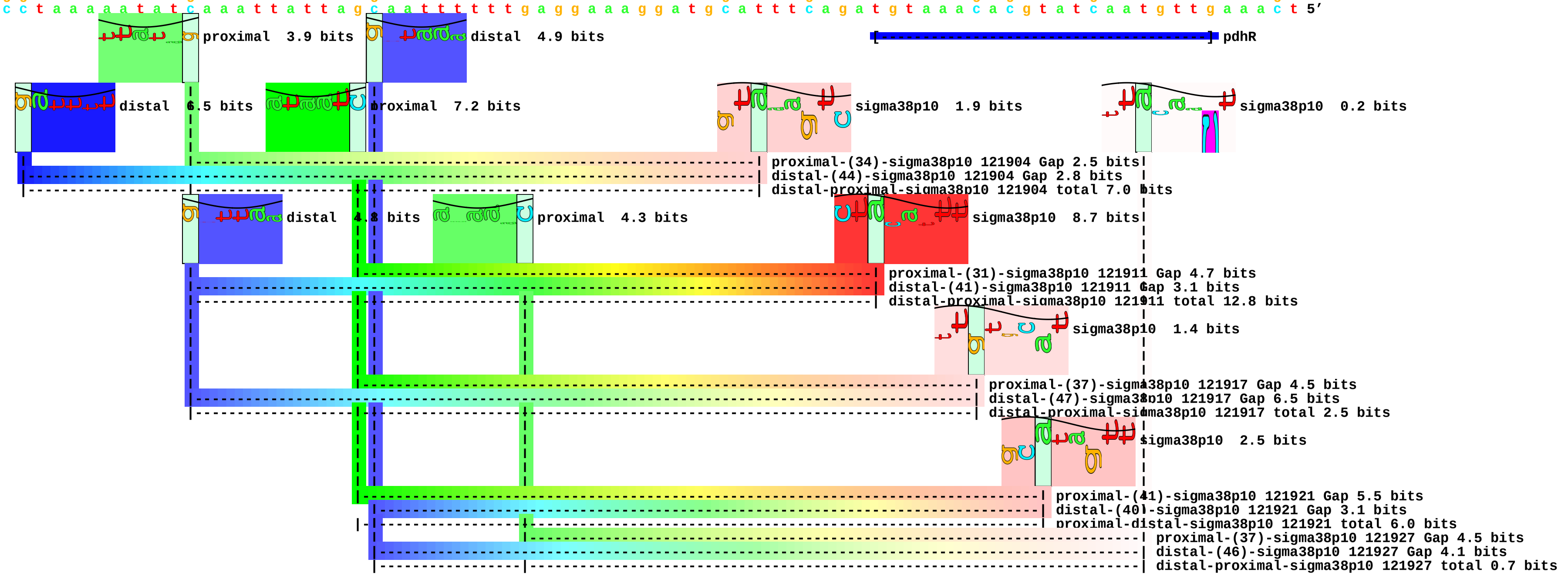

piece 32, NC_000913, rssB, config: linear, direction: +, begin: 1289373, end: 1289458

```
         .          *1289380  .          *1289390  .          *1289400  .          *1289410  .          *1289420  .          *1289430  .          *1289440  .          *1289450  .
5' a c c a g a a t t t t t a t c t a c a c t t a a g t t a a t t c t g a c a g g c g c a g g t g g c a a t a g c a t g c c a c t a t t g a g t a a a g c c a g t c a g g g g a 3'
3' t g g t c t t a a a a a t a g a t g t g a a t t c a a t t a a g a c t g t c c g c g t c c a c c g t t a t c g t a c g g t g a t a a c t c a t t t c g g t c a g t c c c c t 5'
```

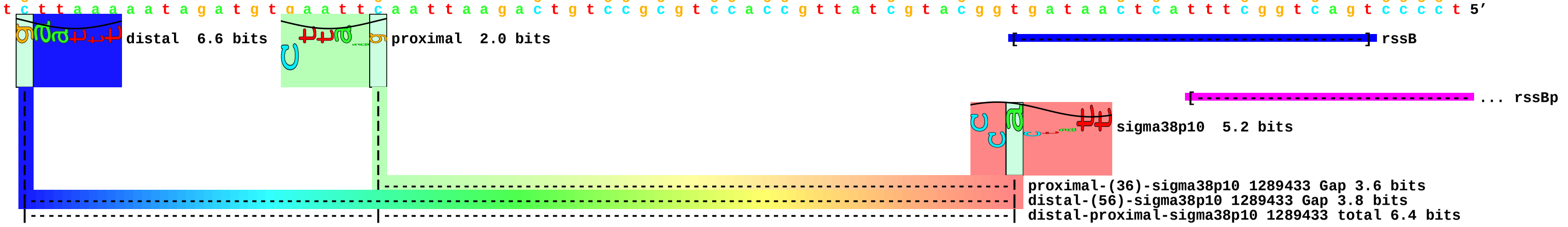

piece 33, NC_000913, wrbA, config: linear, direction: -, begin: 1067101, end: 1067016

```
   *1067100  .       *1067090  .       *1067080  .       *1067070  .       *1067060  .       *1067050  .       *1067040  .       *1067030  .       *1067020
5' a a c g c g a t g a g a t a a a a a t g t g t c a t t t t g c g a c a a a a t t a c g t g c t t g t g a a a g t t g t t a t a a a t c a a a t a a g t g g t t g t g a a a t 3'
3' t t g c g c t a c t c t a t t t t t a c a c a g t a a a a c g c t g t t t t a a t g c a c g a a c a c t t t c a a c a a t a t t t a g t t t a t t c a c c a a c a c t t t a 5'
```

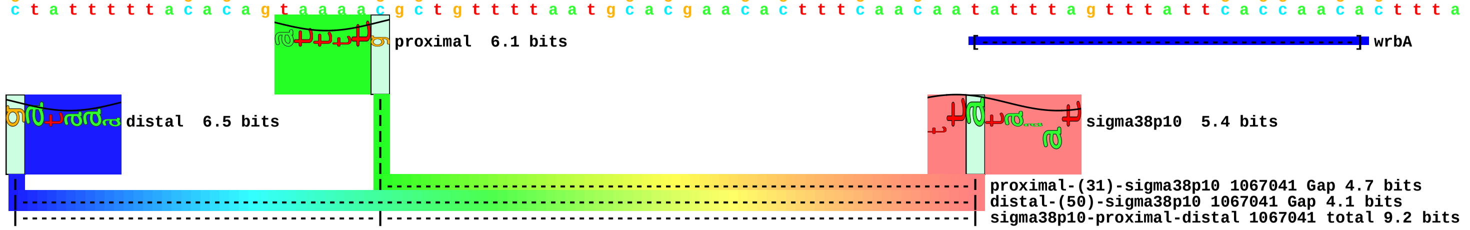

piece 34, NC_000913, yiaG, config: linear, direction: +, begin: 3717383, end: 3717468

```
          .         *3717390  .         *3717400  .         *3717410  .         *3717420  .         *3717430  .         *3717440  .         *3717450  .         *3717460  .
5' a a t c g a a t c a t a g c c a g a g c a t g c c c t g a c t t c a c c c c g c t g t g t c t g c t t t t c c c g a c t a t t c t t a a t g a g c t t c g a t g c a a t t c 3'
3' t t a g c t t a g t a t c g g t c t c g t a c g g g a c t g a a g t g g g g c g a c a c a g a c g a a a a g g g c t g a t a a g a a t t a c t c g a a g c t a c g t t a a g 5'
```

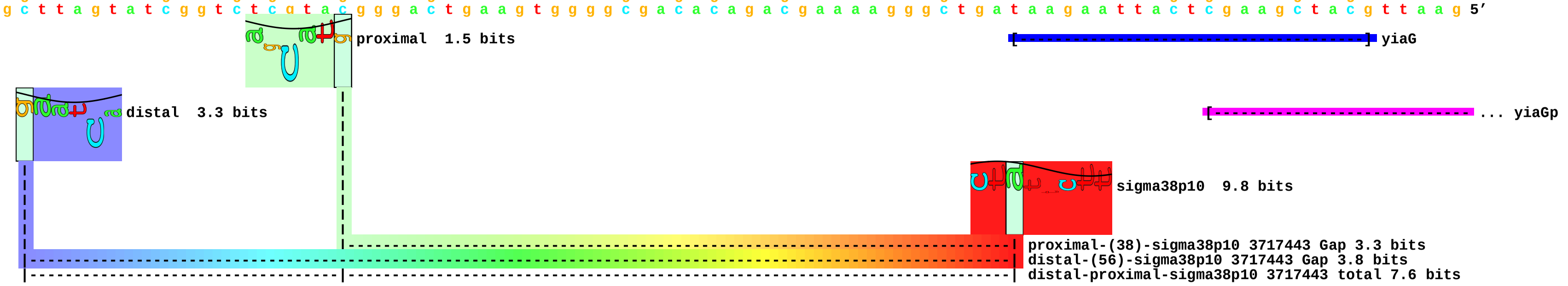

piece 35, NC_000913, artPp1, config: linear, direction: -, begin: 903108, end: 903023

```
          .         *903100   .         *903090   .         *903080   .         *903070   .         *903060   .         *903050   .         *903040   .         *903030   .
5' t c g c c g a c c a c c g c c c c c g t t a t t t t g t g c t a t g t t t a t t g a a t a a t g c g c t t t g c t t t t a a c t t t t a a a g c a g a a a t a t t g c a t a 3'
3' a g c g g c t g g t g g c g g g g g c a a t a a a a c a c g a t a c a a a t a a c t t a t t a c g c g a a a c g a a a a t t g a a a a t t t c g t c t t t a t a a c g t a t 5'
```

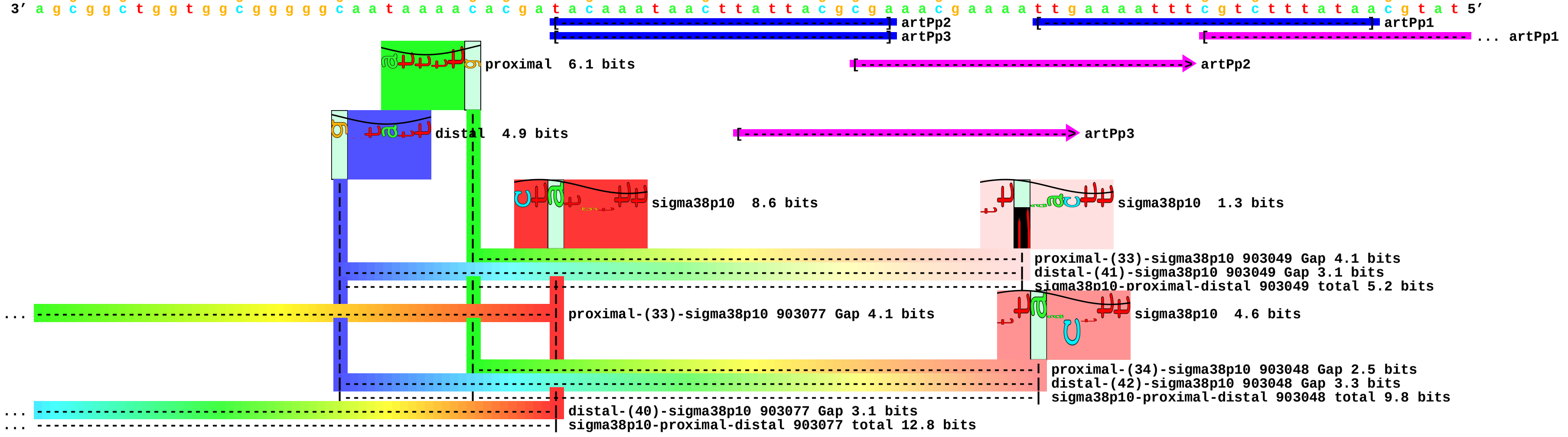

piece 36, NC_000913, artPp2, config: linear, direction: -, begin: 903137, end: 903052

```
        .         *903130    .         *903120    .         *903110    .         *903100    .         *903090    .         *903080    .         *903070    .         *903060    .
5' c a t t t c g t g a g g a a t g c c g a c a t t t a t g c t c g c c g a c c a c c g c c c c c g t t a t t t t g t g c t a t g t t t a t t g a a t a a t g c g c t t t g c t 3'
3' g t a a a g c a c t c c t t a c g g c t g t a a a t a c g a g c g g c t g g t g g c g g g g g c a a t a a a a c a c g a t a c a a a t a a c t t a t t a c g c g a a a c g a 5'
```

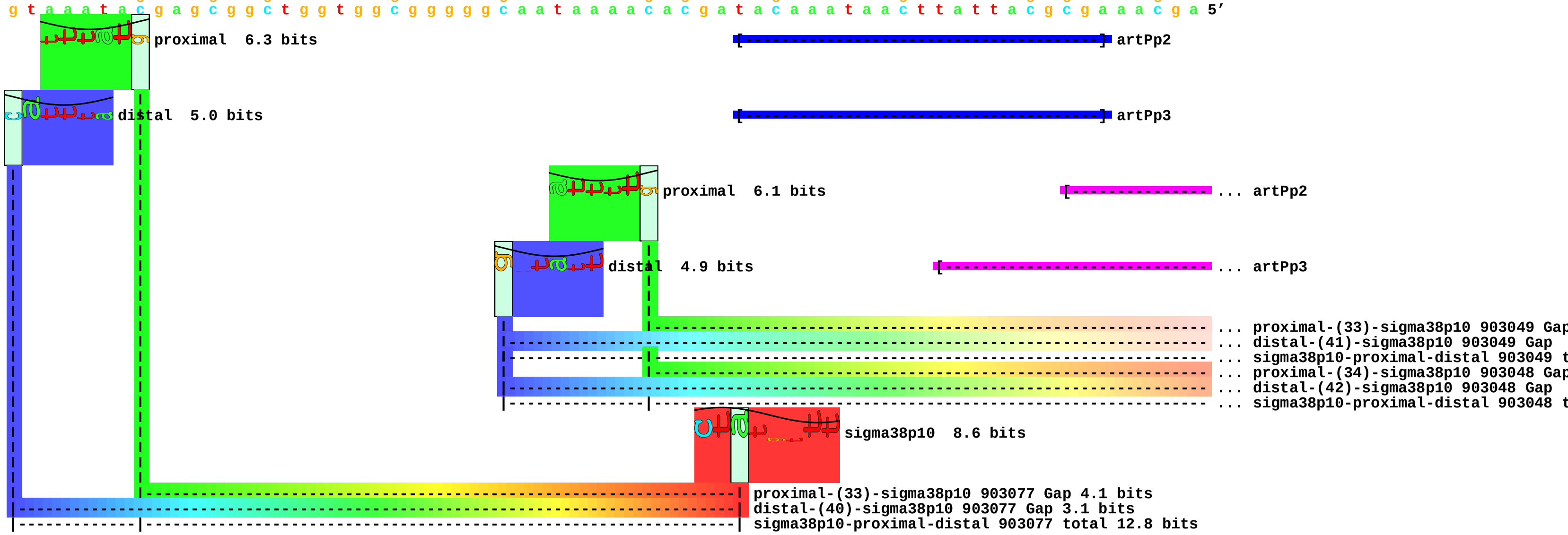

piece 37, NC_000913, artPp3, config: linear, direction: -, begin: 903137, end: 903052

```
        .         *903130    .         *903120    .         *903110    .         *903100    .         *903090    .         *903080    .         *903070    .         *903060    .
5' c a t t t c g t g a g g a a t g c c g a c a t t t a t g c t c g c c g a c c a c c g c c c c g t t a t t t t g t g c t a t g t t t a t t g a a t a a t g c g c t t t g c t 3'
3' g t a a a g c a c t c c t t a c g g c t g t a a a t a c g a g c g g c t g g t g g c g g g g c a a t a a a a c a c g a t a c a a a t a a c t t a t t a c g c g a a a c g a 5'
```

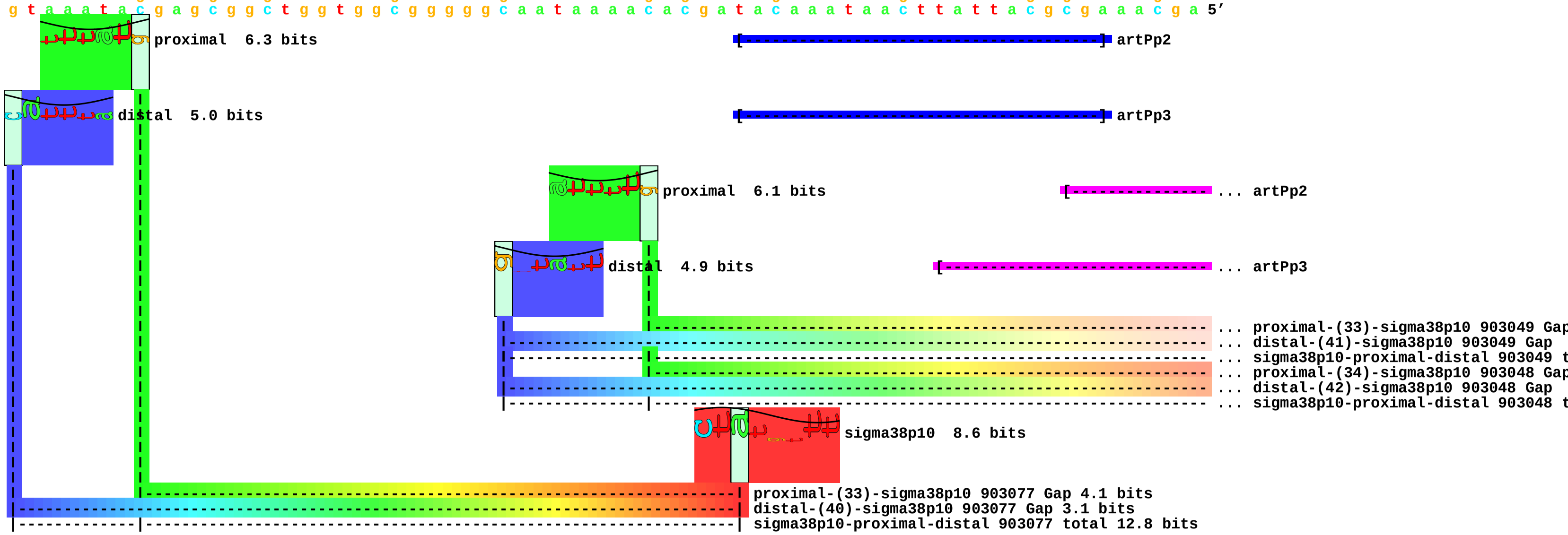

piece 38, NC_000913, blc, config: linear, direction: -, begin: 4375837, end: 4375752

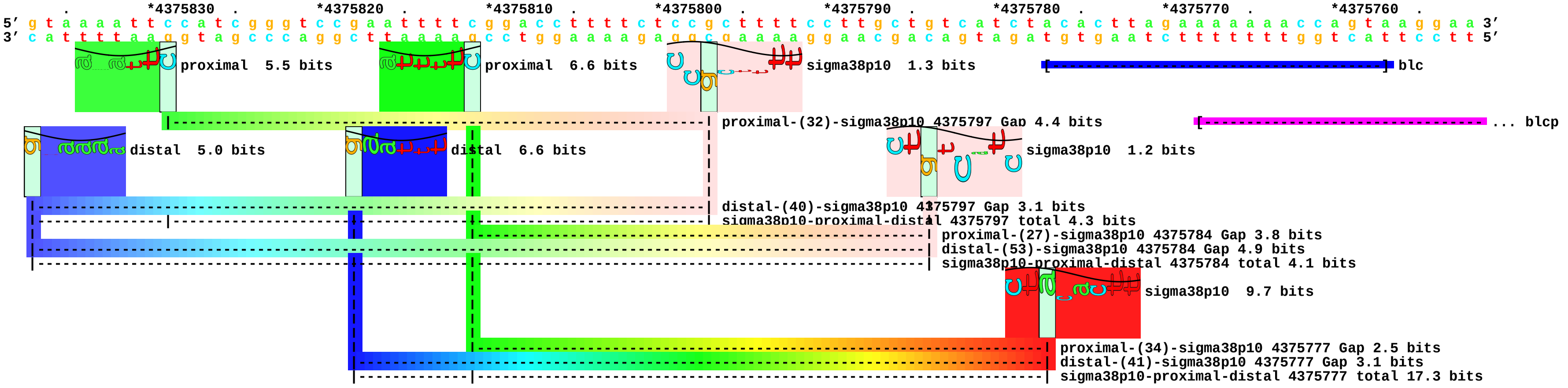

piece 39, NC_000913, mscL, config: linear, direction: +, begin: 3435941, end: 3436026

```
               .         *3435950  .         *3435960  .         *3435970  .         *3435980  .         *3435990  .         *3436000  .         *3436010  .         *3436020  .
5' t a t t a t t t c c c t t t g a t t a t c a a g g a t t a a t t a a a t t c a t t c c t g g c a g g a a a a t g g c t t a a c a t t t g t t a g a c t t a t g g t t g t c g 3'
3' a t a a t a a a g g g a a a c t a a t a g t t c c t a a t t a a t t t a a g t a a g g a c c g t c c t t t t a c c g a a t t g t a a a c a a t c t g a a t a c c a a c a g c 5'
```

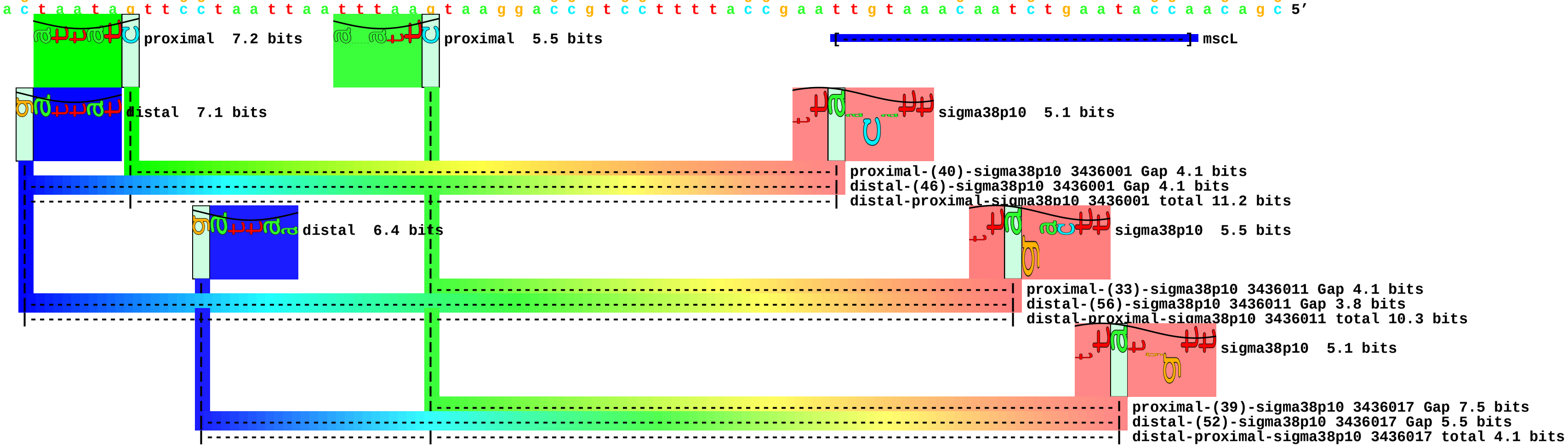

piece 40, NC_000913, potF, config: linear, direction: +, begin: 892793, end: 892878

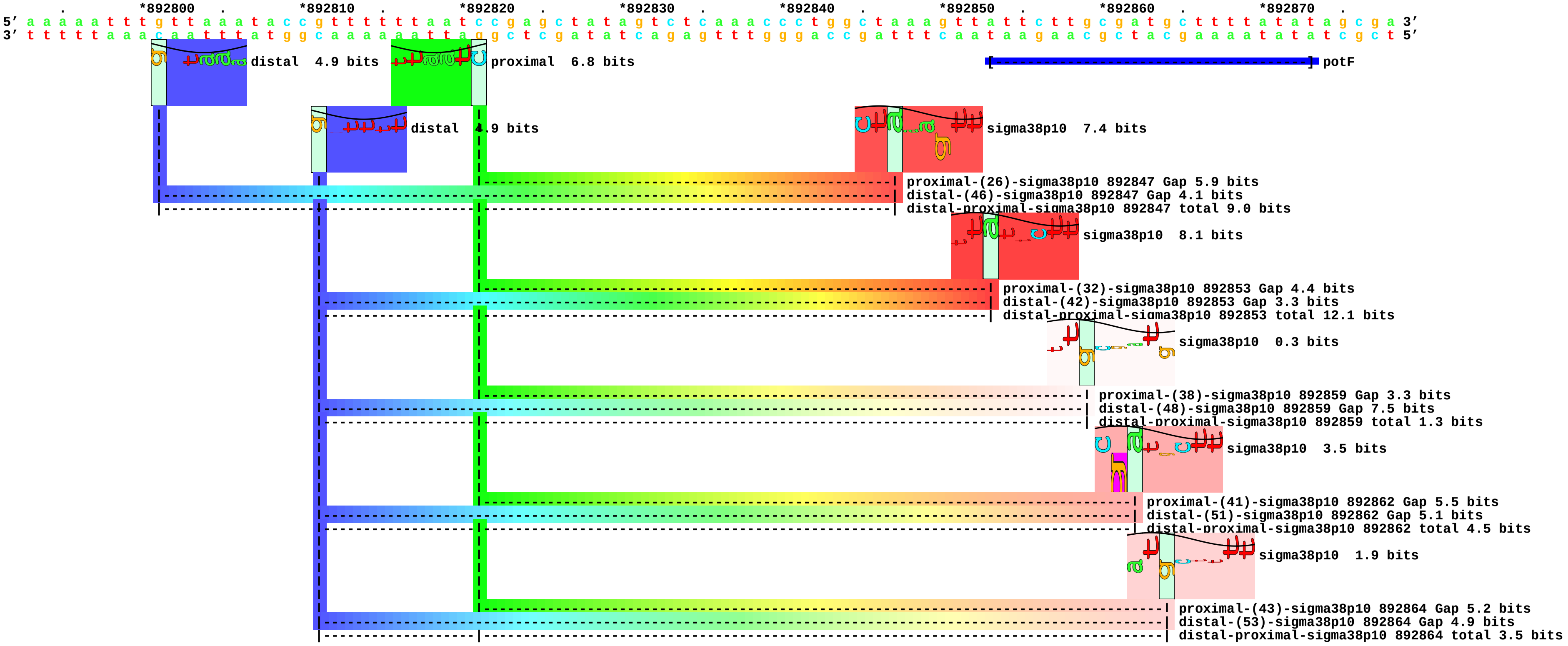

piece 41, NC_000913, rssA, config: linear, direction: +, begin: 1288069, end: 1288154

```
    *1288070  .        *1288080  .        *1288090  .        *1288100  .        *1288110  .        *1288120  .        *1288130  .        *1288140  .        *1288150
5' a a a c t g a c a a a t c c g a t a t t g t c g g c g t c c t g c c a a c a a t g t t c a a a t a c g g t c a a t c c c a a c c a t t c a g t a t g c g c a a a t c c g g c 3'
3' t t t g a c t g t t t a g g c t a t a a c a g c c g c a g g a c g g t t g t t a c a a g t t t a t g c c a g t t a g g g t t g g t a a g t c a t a c g c g t t t a g g c c g 5'
```

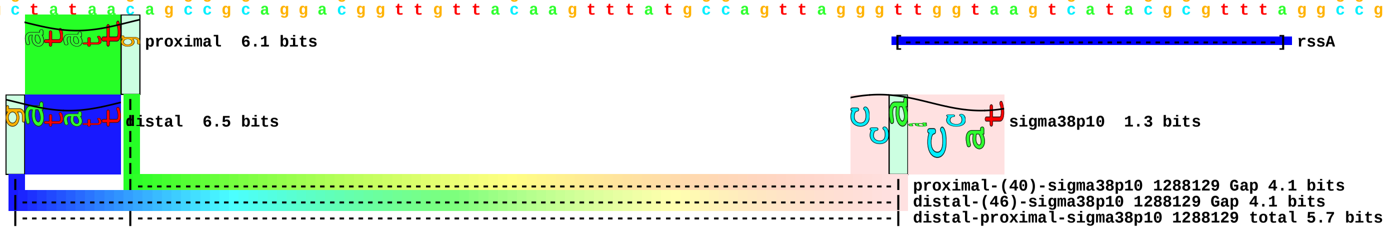

piece 42, NC_000913, ugpB, config: linear, direction: -, begin: 3590604, end: 3590519

```
          *3590600  .         *3590590  .         *3590580  .         *3590570  .         *3590560  .         *3590550  .         *3590540  .         *3590530  .         *3590520
5' g t t t a c g c c g c a t c c g g c a t g a a a c g a t g a g c a a t c t g t a g a g t t t g a t t c a g a c c t t c t a t a t t t t c c c g c t t a t c c g t g c c c c a 3'
3' c a a a t g c g g c g t a g g c c g t a c t t t g c t a c t c g t t a g a c a t c t c a a a c t a a g t c t g g a a g a t a t a a a a g g g c g a a t a g g c a c g g g g t 5'
                                                                                                                                [-----------------------------------------] ugpB
```

piece 43, NC_000913, ugpC, config: linear, direction: -, begin: 3587450, end: 3587365

```
   *3587450  .         *3587440  .         *3587430  .         *3587420  .         *3587410  .         *3587400  .         *3587390  .         *3587380  .         *3587370  .
5' g c g g c g c t g t t t g t g a t c a c c t t t a t c t a c g g c t g g a a t c a g t a t t t g t g g c c g t t g t t g a t t a t t a c c g a t g t g g a t c t c g g c a c 3'
3' c g c c g c g a c a a a c a c t a g t g g a a a t a g a t g c c g a c c t t a g t c a t a a a c a c c g g c a a c a a c t a a t a a t g g c t a c a c c t a g a g c c g t g 5'
```

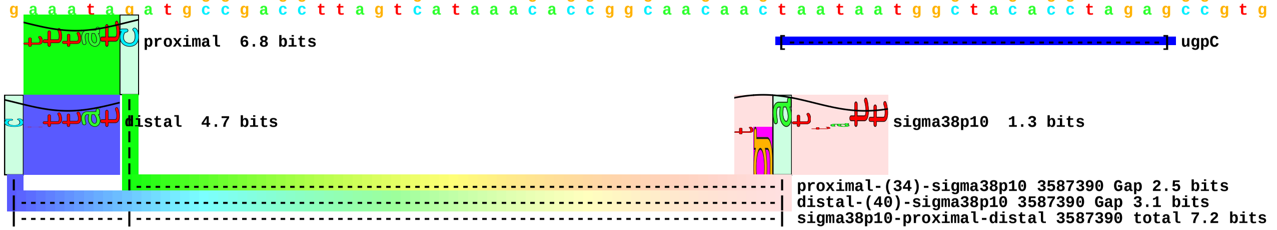

piece 44, NC_000913, ybiO, config: linear, direction: -, begin: 844802, end: 844717

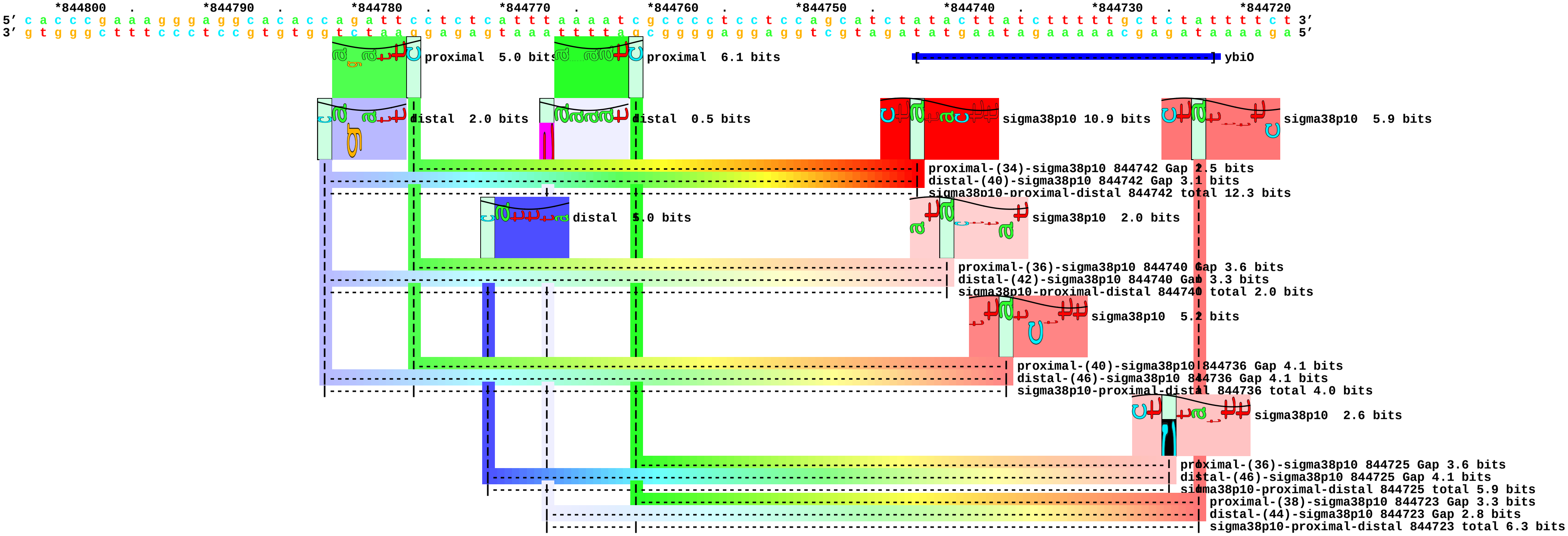

piece 45, NC_000913, ybiP, config: linear, direction: -, begin: 851955, end: 851870

```
   .         *851950   .         *851940   .         *851930   .         *851920   .         *851910   .         *851900   .         *851890   .         *851880   .         *851870
5' c t c g c t c t a t c t t c t t g c a g c c c t g c g t a t a t t g c g g c t c g c g g a t g c g g a c c c c t t t c c a c t c t t c a c g c a c t c t t g c a g g t a t t 3'
3' g a g c g a g a t a g a a g a a c g t c g g g a c g c a t a t a a c g c c g a g c g c c t a c g c c t g g g g a a a g g t g a g a a g t g c g t g a g a a c g t c c a t a a 5'
```

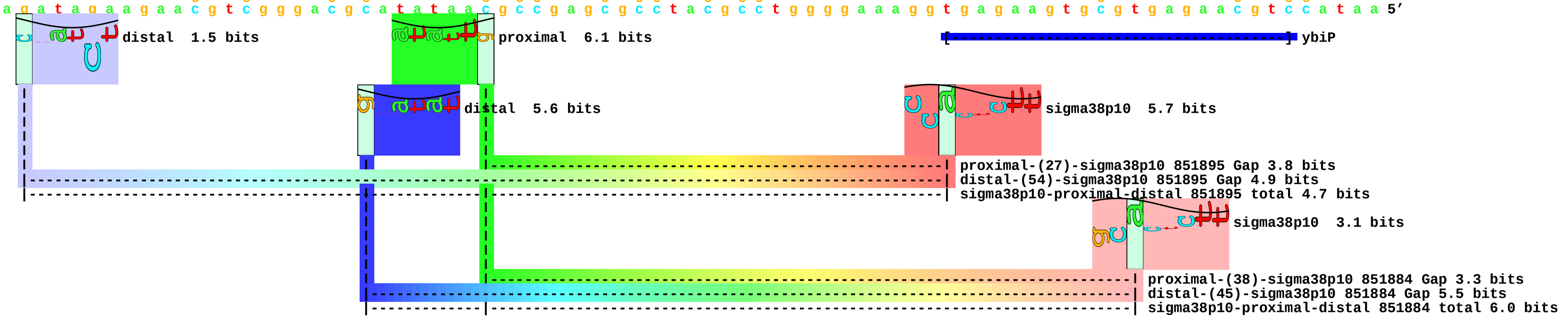

piece 46, NC_000913, ydcS, config: linear, direction: +, begin: 1509549, end: 1509634

```
   *1509550  .         *1509560  .         *1509570  .         *1509580  .         *1509590  .         *1509600  .         *1509610  .         *1509620  .         *1509630
5′ g t a g c t c a c a a c c t g a a t a a a t t t t c t c a g g g g c g a a g g t g t g c c t g c a a g c c g c c g t c t a t g g t t a a a c a a g g a g a t a t t t t t a c 3′
3′ c a t c g a g t g t t g g a c t t a t t t a a a a g a g t c c c c g c t t c c a c a c g g a c g t t c g g c g g c a g a t a c c a a t t t g t t c c t c t a t a a a a a t g 5′
```

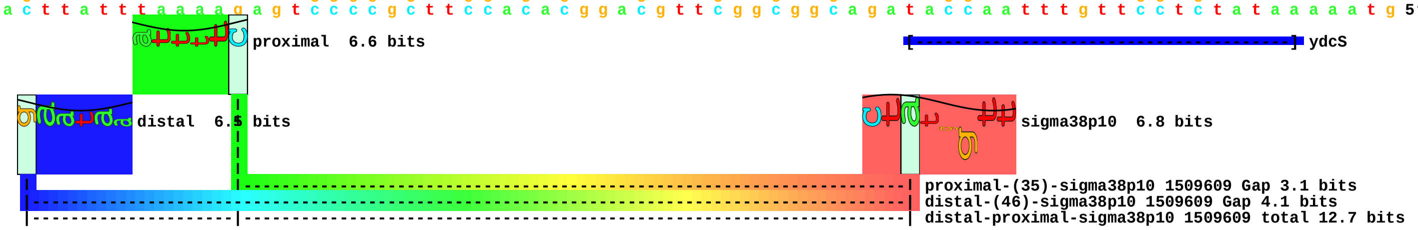

piece 47, NC_000913, ydhJ, config: linear, direction: +, begin: 1718962, end: 1719047

```
.         *1718970  .         *1718980  .         *1718990  .         *1719000  .         *1719010  .         *1719020  .         *1719030  .         *1719040  .
5' t t t t c a g a a a t t g c g c g t a a a t t t t t c g c a c t t a a a g a a t a t t t a t t a a t c t a a c g c a a t a t a t t c g g t c g t a a a g g a a t c t a c t t 3'
3' a a a a g t c t t t a a c g c g c a t t t a a a a a g c g t g a a t t t c t t a t a a a t a a t t a g a t t g c g t t a t a t a a g c c a g c a t t t c c t t a g a t g a a 5'
```

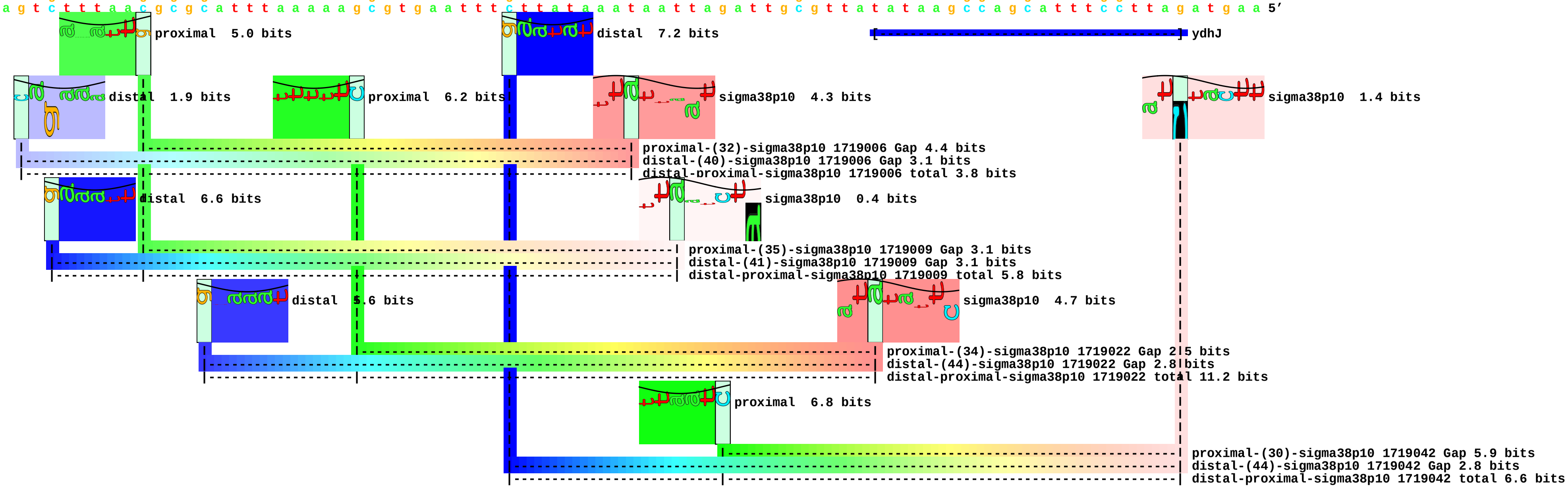

piece 48, NC_000913, yebS, config: linear, direction: +, begin: 1914186, end: 1914271

```
      *1914190  .       *1914200  .       *1914210  .       *1914220  .       *1914230  .       *1914240  .       *1914250  .       *1914260  .       *1914270
5' c c a g c c t a t t a a a a t a a g c a t t a a a t g c g t t a a t g c t c a a g a t c a t t c c c a t c a t g g g t t a a g a t t a a t g t t a a t t c t t a t t a c a t 3'
3' g g t c g g a t a a t t t t a t t c g t a a t t t a c g c a a t t a c g a g t t c t a g t a a g g g t a g t a c c c a a t t c t a a t t a c a a t t a a g a a t a a t g t a 5'
```

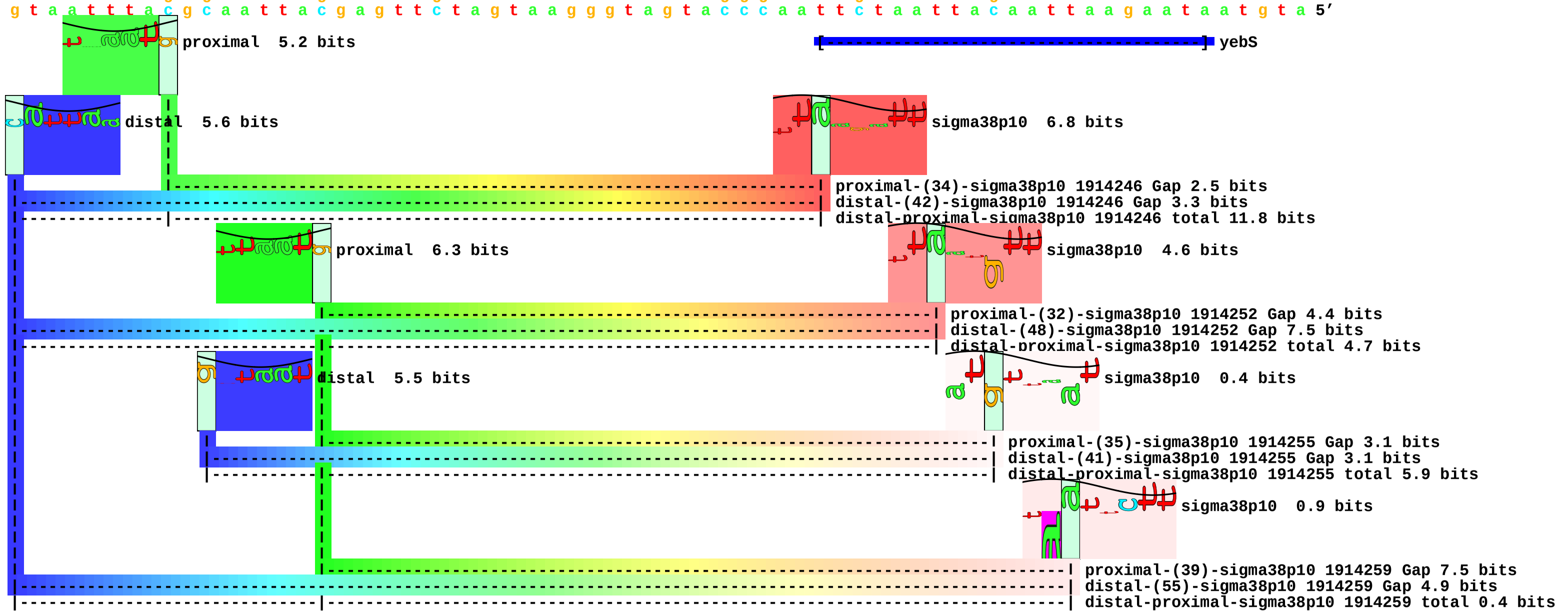

piece 49, NC_000913, yhhT, config: linear, direction: +, begin: 3609795, end: 3609880

```
   .         *3609800  .         *3609810  .         *3609820  .         *3609830  .         *3609840  .         *3609850  .         *3609860  .         *3609870  .         *3609880
5' c c a c g t g c t g a c g t g a a a a t c a g c a g t a a g a a t c a g a a t a t t g c t g g c g t g a c t c c c g c t a c a c t t a a t a c a a a a a g t c a c a a g g a 3'
3' g g t g c a c g a c t g c a c t t t t a g t c g t c a t t c t t a g t c t t a t a a c g a c c g c a c t g a g g g c g a t g t g a a t t a t g t t t t t c a g t g t t c c t 5'
```

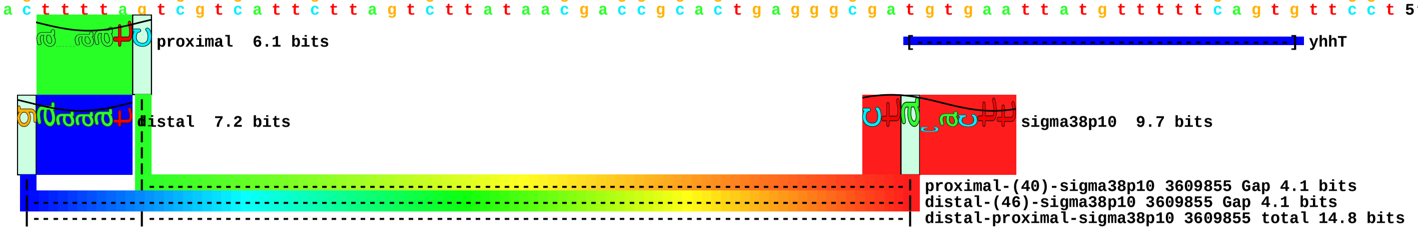

piece 50, NC_000913, yhjD, config: linear, direction: +, begin: 3671231, end: 3671316

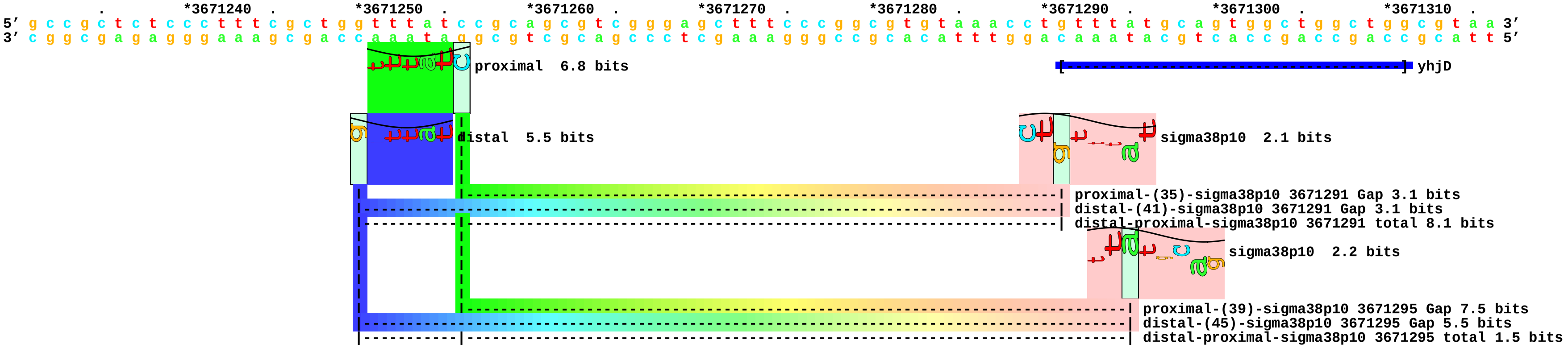

piece 51, NC_000913, yphA, config: linear, direction: +, begin: 2671272, end: 2671357

```
          .         *2671280  .         *2671290  .         *2671300  .         *2671310  .         *2671320  .         *2671330  .         *2671340  .         *2671350  .
5' c c a g g a t a a t t a g c g a a t a t c t c a a t g c c t g g g g c g t g g c g a g g t g c a a g a g t g t g t a t t a c g t t t a a a t c a c a t t a t c t t g c a a a 3'
3' g g t c c t a t t a a t c g c t t a t a g a g t t a c g g a c c c c g c a c c g c t c c a c g t t c t c a c a c a t a a t g c a a a t t t a g t g t a a t a g a a c g t t t 5'
```

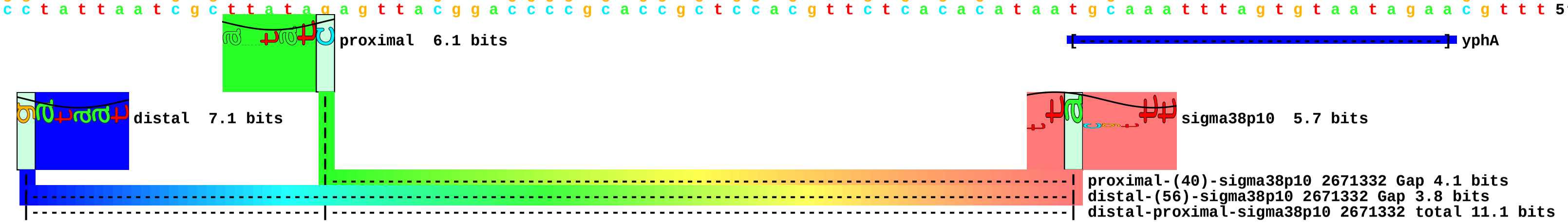

piece 52, NC_000913, yqaE, config: linear, direction: -, begin: 2795178, end: 2795093

```
       .        *2795170  .        *2795160  .        *2795150  .        *2795140  .        *2795130  .        *2795120  .        *2795110  .        *2795100  .
5' t a c c a g c c t c a g c a g c g t a a a t g a g a g t a a a a g c g t a a g c t g a a a c t g g c a g g c t c c g c t a a a a t t a c t a c g c t t a a g a g a t a a a a 3'
3' a t g g t c g g a g t c g t c g c a t t t a c t c t c a t t t t c g c a t t c g a c t t t g a c c g t c c g a g g c g a t t t t a a t g a t g c g a a t t c t c t a t t t t 5'
```

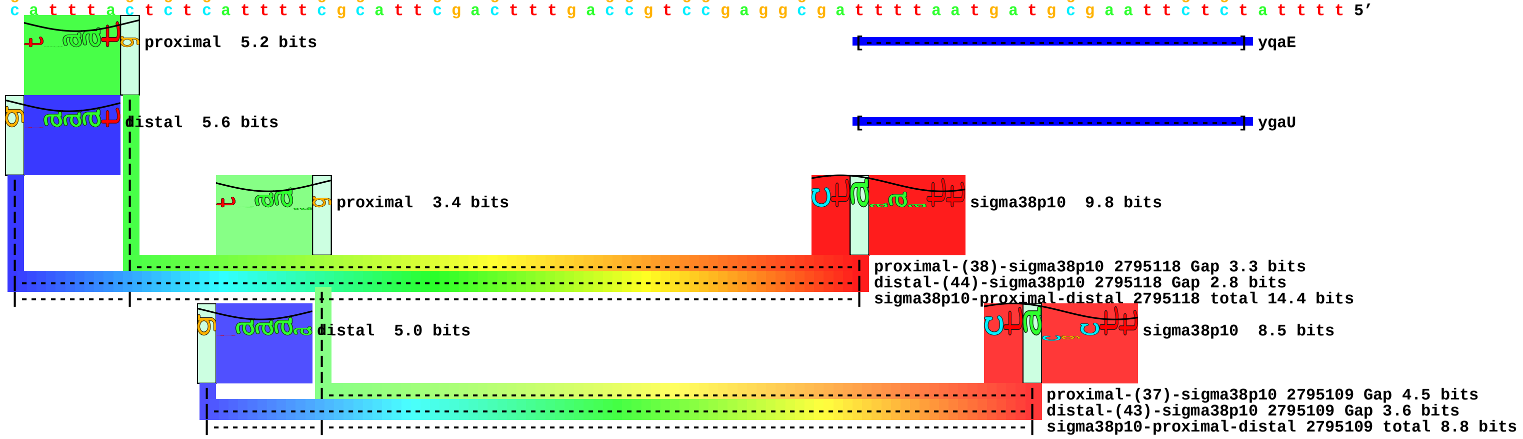

piece 53, NC_000913, aidB, config: linear, direction: +, begin: 4411828, end: 4411913

```
       *4411830  .         *4411840  .         *4411850  .         *4411860  .         *4411870  .         *4411880  .         *4411890  .         *4411900  .         *4411910
5' a g t a t c t t g a g t a a t a a a g g g c t a a t g c c g c t g c t t t g g c g c t t c t t c c c t g g t c a t c c t a a t c t t c t t g c g t c c t g g t t c g a t g g 3'
3' t c a t a g a a c t c a t t a t t t c c c g a t t a c g g c g a c g a a a c c g c g a a g a a g g g a c c a g t a g g a t t a g a a g a a c g c a g g a c c a a g c t a c c 5'
```

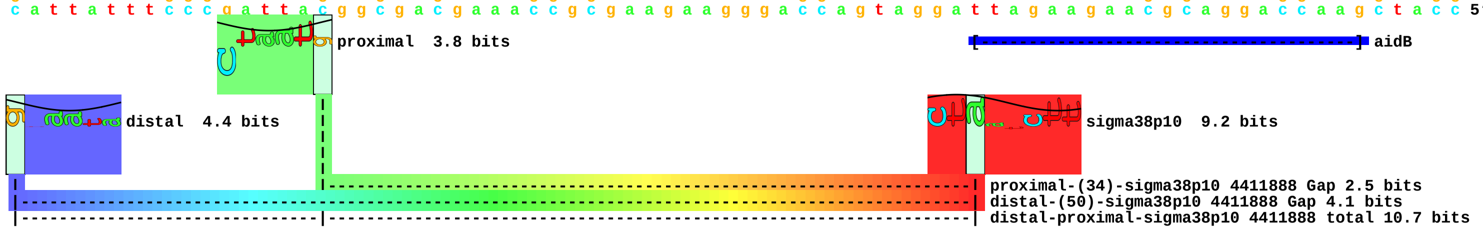

piece 54, NC_000913, bfr, config: linear, direction: -, begin: 3464841, end: 3464756

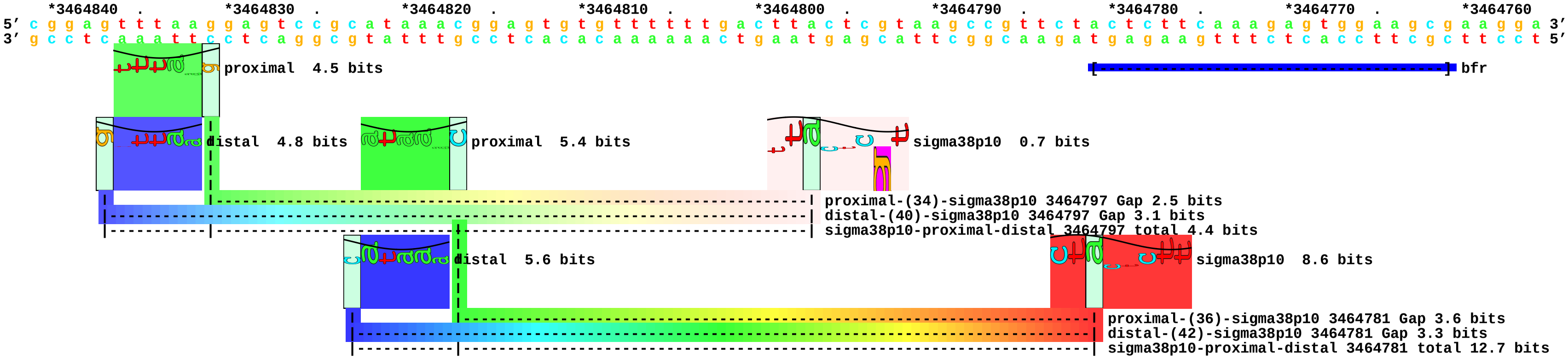

piece 55, NC_000913, dps, config: linear, direction: -, begin: 848358, end: 848273

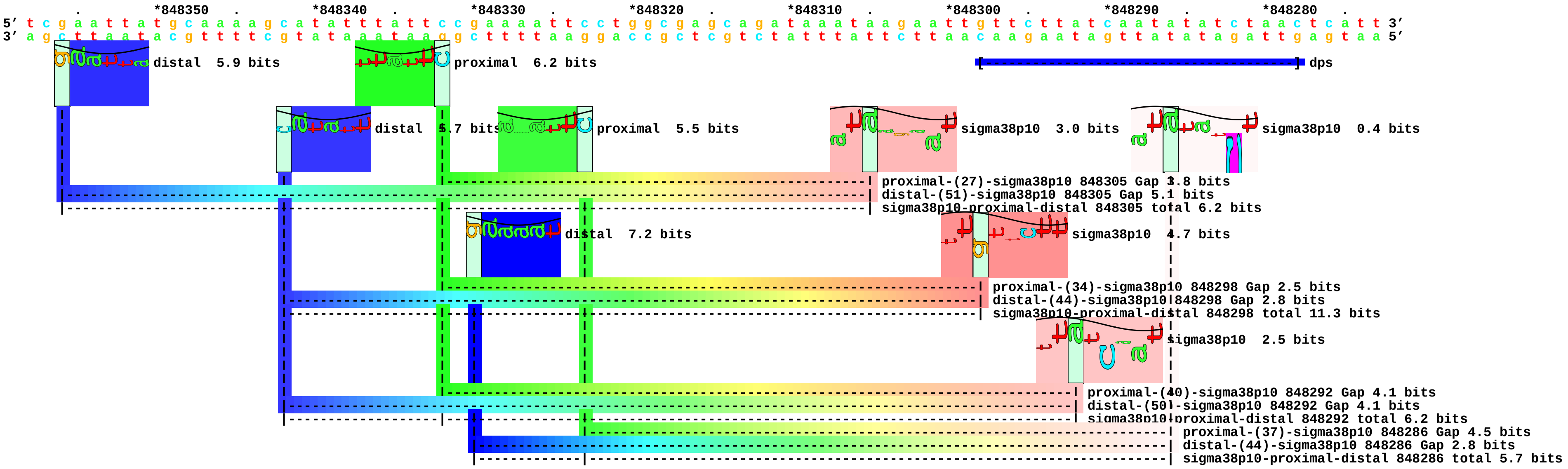

piece 56, NC_000913, gadA, config: linear, direction: -, begin: 3665702, end: 3665617

```
   *3665700  .       *3665690  .       *3665680  .       *3665670  .       *3665660  .       *3665650  .       *3665640  .       *3665630  .       *3665620
5' t t a g g a t t t t g t t a t t t a a a t t a a g c c t g t a a t g c c t t g c t t c c a t t g c g g a t a a a t c c t a c t t t t t t a t t g c c t t c a a a t a a a t t 3'
3' a a t c c t a a a a c a a t a a a t t t a a t t c g g a c a t t a c g g a a c g a a g g t a a c g c c t a t t t a g g a t g a a a a a a t a a c g g a a g t t t a t t t a a 5'
```

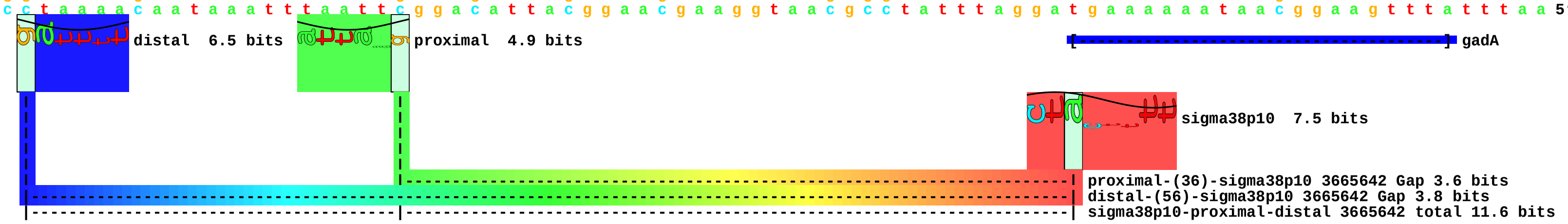

piece 57, NC_000913, gadB, config: linear, direction: -, begin: 1570168, end: 1570083

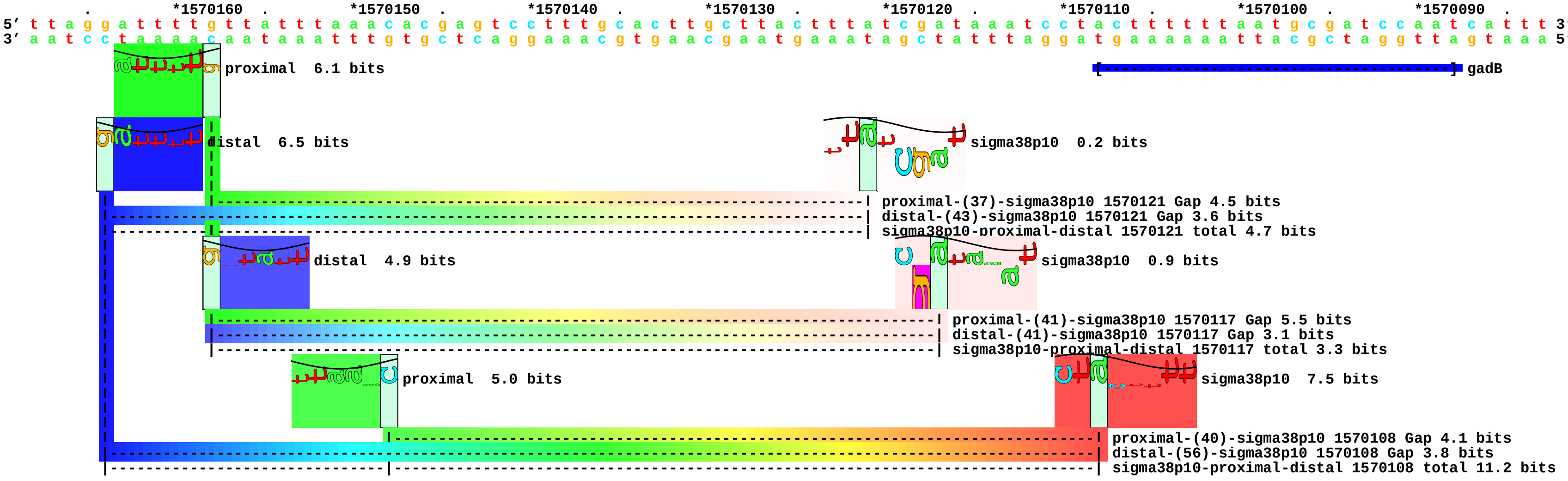

piece 58, NC_000913, katE, config: linear, direction: +, begin: 1811766, end: 1811851

```
          *1811770  .         *1811780  .         *1811790  .         *1811800  .         *1811810  .         *1811820  .         *1811830  .         *1811840  .         *1811850
5' c g t g t a t t t c a t a a c a c c g t t t c c a g a a t a g t c t c c g a a g c g g g a t c t g g c t g g t g g t c t a t a g t t a g a g a g t t t t t t g a c c a a a a 3'
3' g c a c a t a a a g t a t t g t g g c a a a g g t c t t a t c a g a g g c t t c g c c c t a g a c c g a c c a c c a g a t a t c a a t c t c t c a a a a a a c t g g t t t t 5'
```

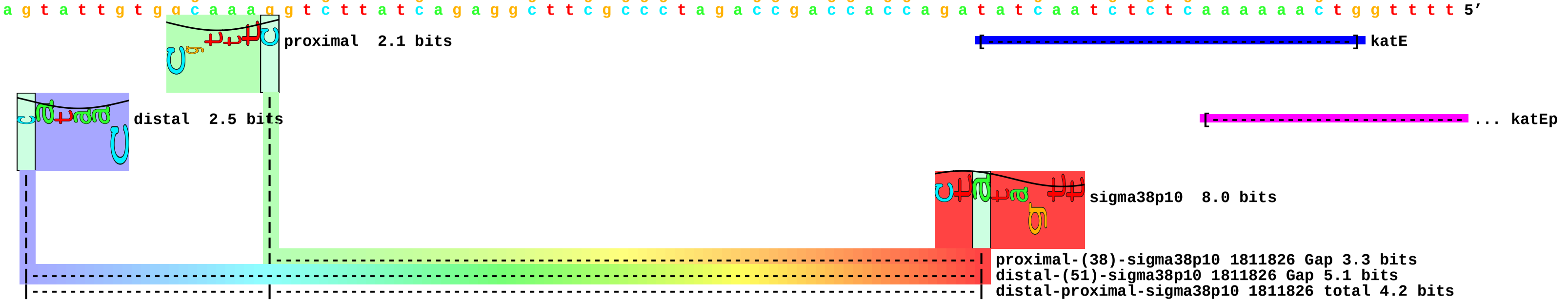

piece 59, NC_000913, osmB, config: linear, direction: -, begin: 1341571, end: 1341486

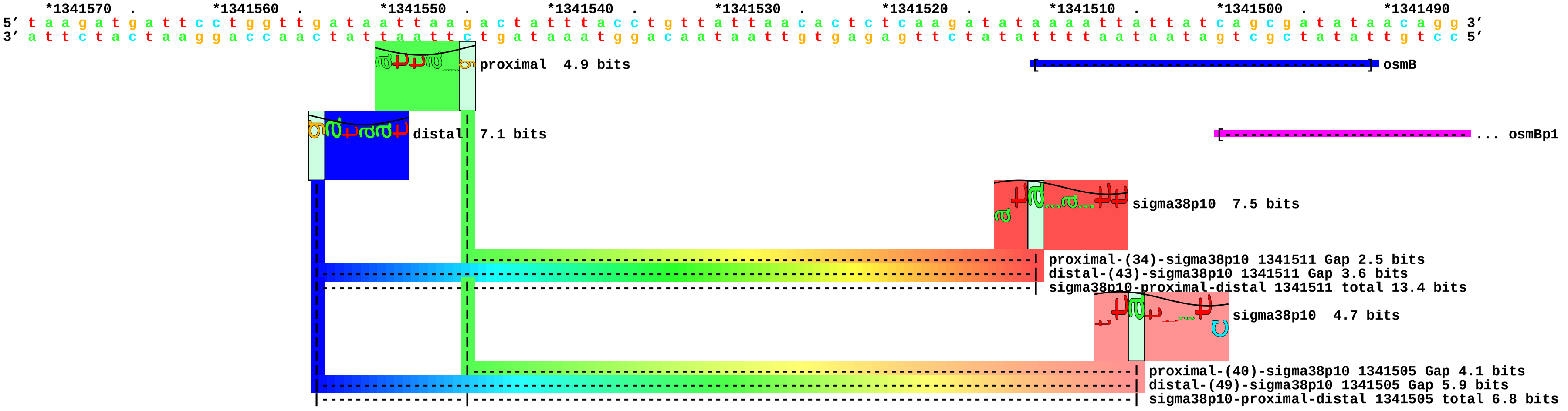

piece 60, NC_000913, osmC, config: linear, direction: +, begin: 1554555, end: 1554640

```
   .         *1554560  .         *1554570  .         *1554580  .         *1554590  .         *1554600  .         *1554610  .         *1554620  .         *1554630  .         *1554640
5' t t t t c g c c g g a t t t t a t t c g g a a t a t c c t g c t t a t c c t c g t g c t g t t t c t c a c g t a g t c t a t a a t t t c c t t t t t a a g c c c a c a g g a 3'
3' a a a a g c g g c c t a a a a t a a g c c t t a t a g g a c g a a t a g g a g c a c g a c a a a g a g t g c a t c a g a t a t t a a a g g a a a a a t t c g g g t g t c c t 5'
```

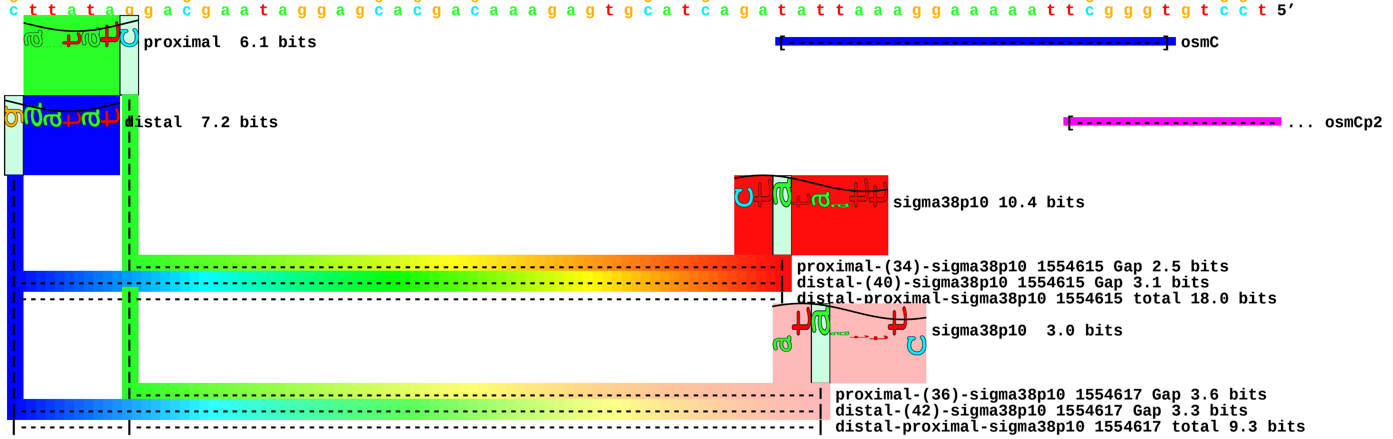

piece 61, NC_000913, osmY, config: linear, direction: +, begin: 4609104, end: 4609189

```
    .         *4609110  .         *4609120  .         *4609130  .         *4609140  .         *4609150  .         *4609160  .         *4609170  .         *4609180  .
5' a a a c a t t c a c t t t t g c t t a t g t t t t c g c t g a t a t c c c g a g c g g t t t c a a a a t t g t g a t c t a t a t t t a a c a a a g t g a t g a c a t t t c t 3'
3' t t t g t a a g t g a a a a c g a a t a c a a a a g c g a c t a t a g g g c t c g c c a a a g t t t t a a c a c t a g a t a t a a a t t g t t t c a c t a c t g t a a a g a 5'
```

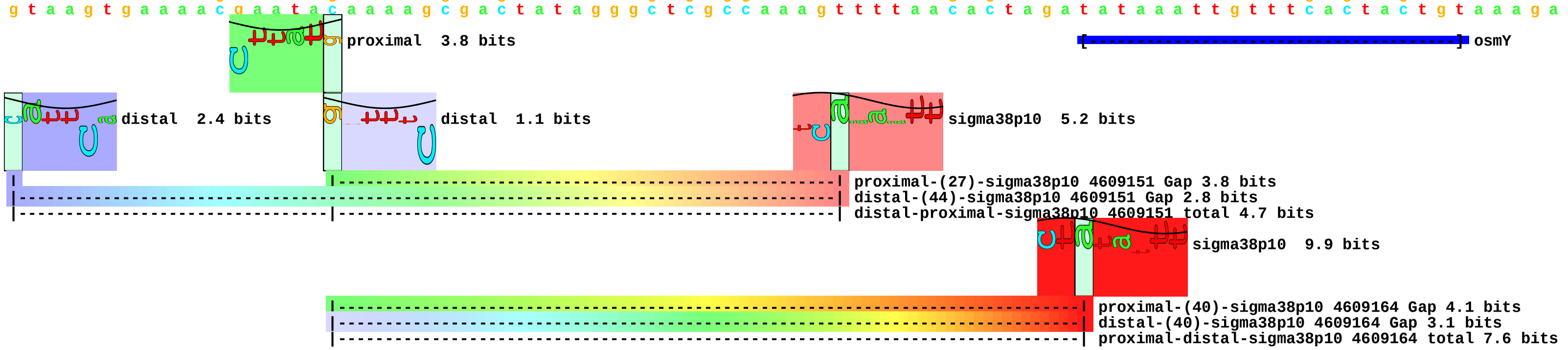

piece 62, NC_000913, otsA, config: linear, direction: -, begin: 1979821, end: 1979736

```
     *1979820  .         *1979810  .         *1979800  .         *1979790  .         *1979780  .         *1979770  .         *1979760  .         *1979750  .         *1979740
5' c g a t g a t t t a a c c g a t g a a t c t g g c t t c g c a g t c g t t a a c c g a c t g g g c g g a a t g t c a g t a a a a a t t g g c a c a g g t g c a a c t c a g g 3'
3' g c t a c t a a a t t g g c t a c t t a g a c c g a a g c g t c a g c a a t t g g c t g a c c c g c c t t a c a g t c a t t t t t a a c c g t g t c c a c g t t g a g t c c 5'
```

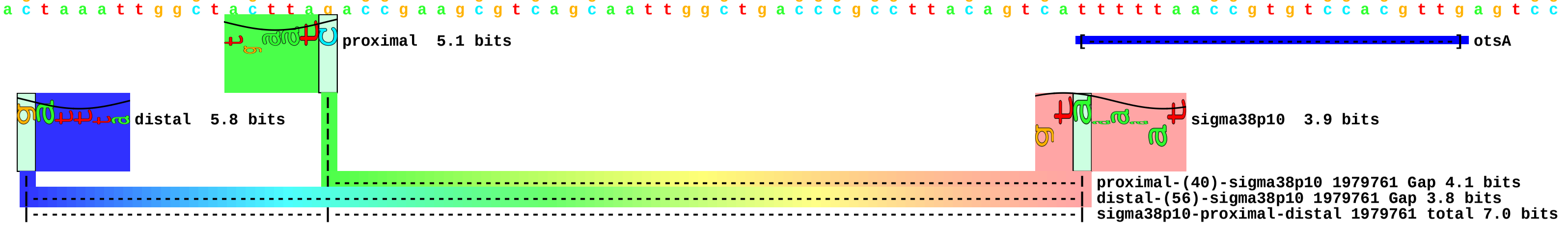

piece 63, NC_000913, otsBp, config: linear, direction: -, begin: 1980574, end: 1980489

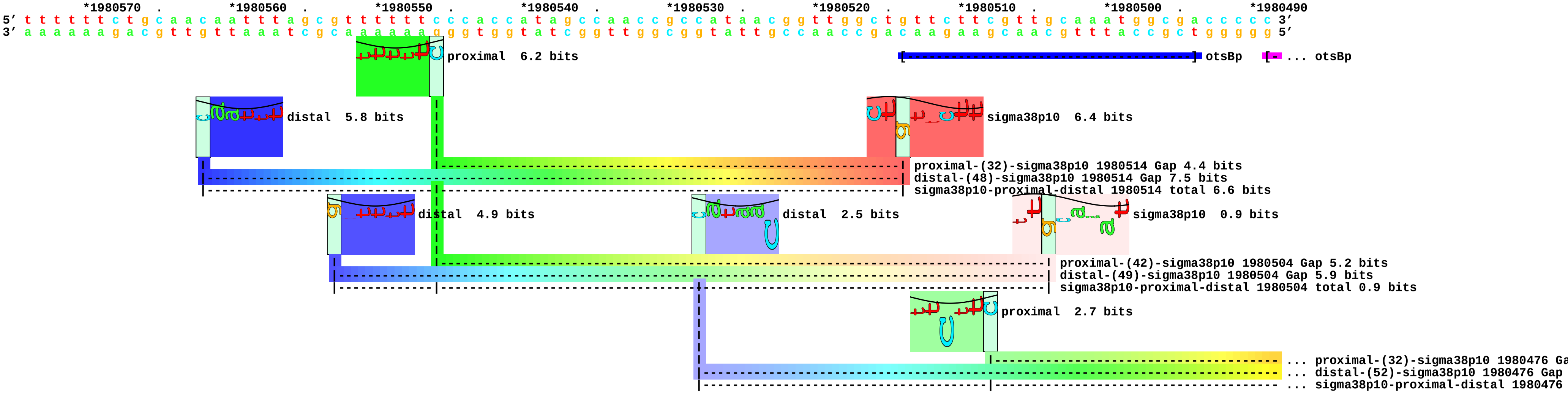

piece 64, NC_000913, otsBp2, config: linear, direction: -, begin: 1980536, end: 1980451

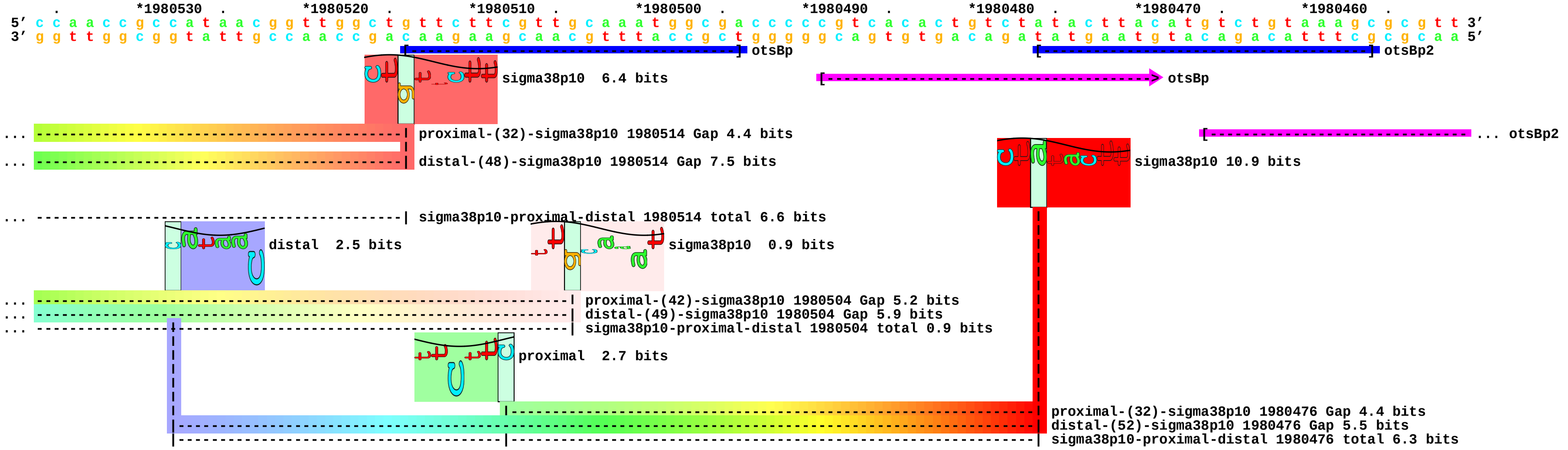

piece 65, NC_000913, psiF, config: linear, direction: +, begin: 402383, end: 402468

```
        .         *402390   .         *402400   .         *402410   .         *402420   .         *402430   .         *402440   .         *402450   .         *402460   .
5' a t a a a a c c g c g c c c g g c a g t g a a t t t t c g c t g c c g g g t g g t t t t t t g c t g t t a g c a a c c a g a c t t a a t g g c a g a t c a c g g g c g c a 3'
3' t a t t t t g g c g c g g g c c g t c a c t t a a a a g c g a c g g c c c a c c a a a a a a c g a c a a t c g t t g g t c t g a a t t a c c g t c t a g t g c c c g c g t 5'
```

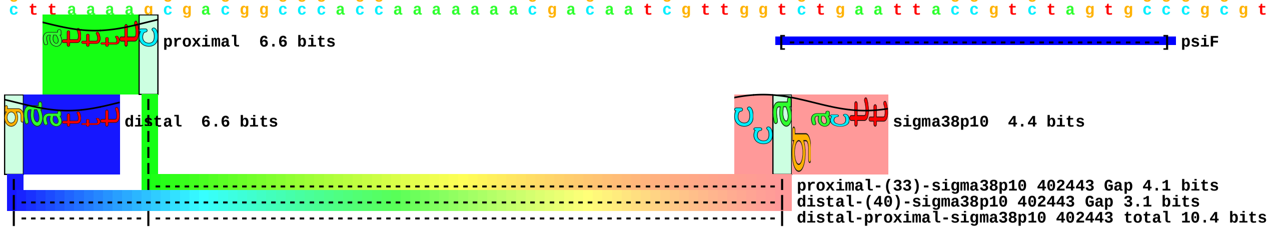

piece 66, NC_000913, treA, config: linear, direction: -, begin: 1246716, end: 1246631

```
     .         *1246710  .         *1246700  .         *1246690  .         *1246680  .         *1246670  .         *1246660  .         *1246650  .         *1246640  .
5' a a a a t g c c t g a c a g t t c g c a g a a t g a g a t t t c g a t c a t g c a g c t a g t g c g a t c c t g a a c t a a g g t t t t c t g a t a c t t g a a t a c c g t 3'
3' t t t t a c g g a c t g t c a a g c g t c t t a c t c t a a a g c t a g t a c g t c g a t c a c g c t a g g a c t t g a t t c c a a a a g a c t a t g a a c t t a t g g c a 5'
```

proximal 5.0 bits

treA

distal 2.6 bits

... treAp

sigma38p10 6.2 bits

proximal-(36)-sigma38p10 1246656 Gap 3.6 bits

distal-(42)-sigma38p10 1246656 Gap 3.3 bits

sigma38p10-proximal-distal 1246656 total 6.9 bits

piece 67, NC_000913, treF, config: linear, direction: +, begin: 3667433, end: 3667518

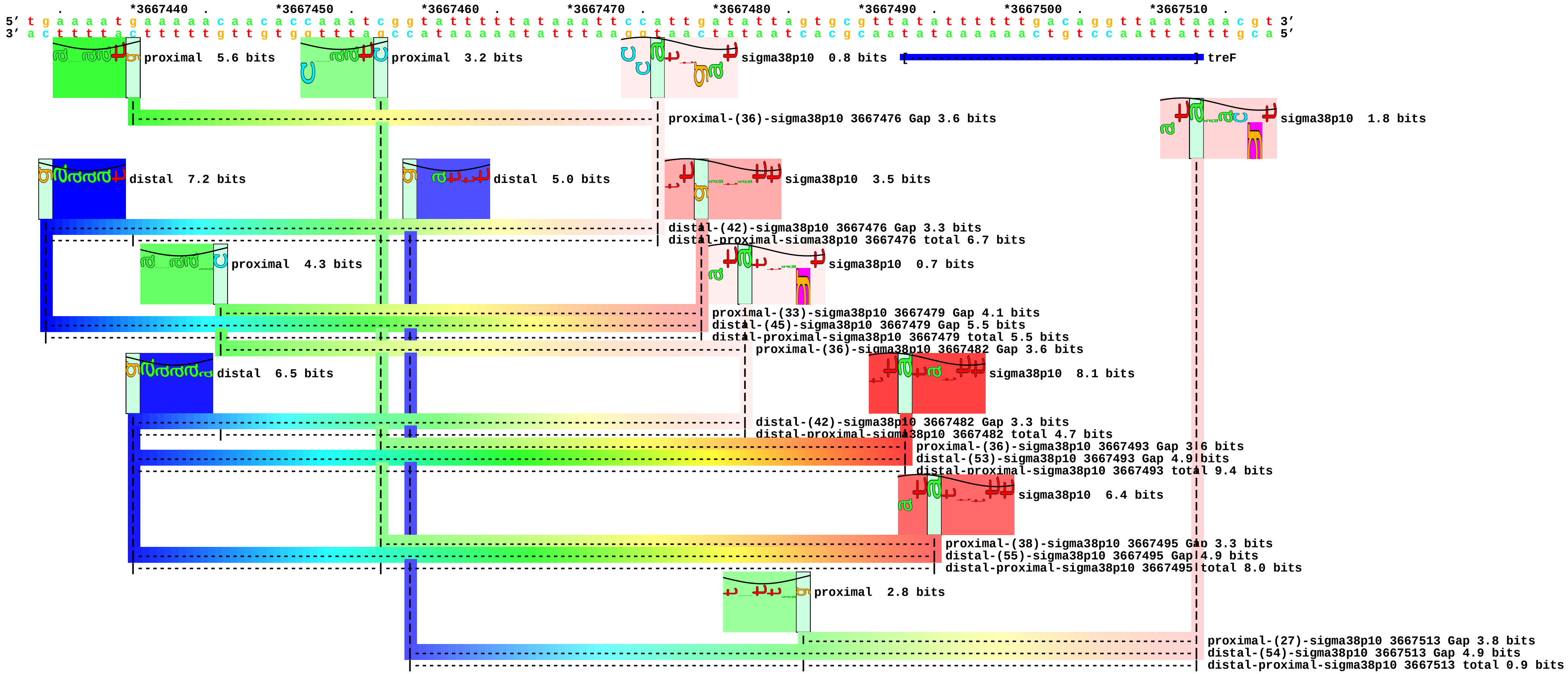

piece 68, NC_000913, uspB, config: linear, direction: -, begin: 3637948, end: 3637863

```
          .          *3637940  .          *3637930  .          *3637920  .          *3637910  .          *3637900  .          *3637890  .          *3637880  .          *3637870  .
5' a a a c a a a t t t t c g a c a t g a a t a t a t t g a t a g t g g t t a a c c t t c t g g a a a a a a a a c a a c c t g a t c t c c t a c a c t a t c t a t a g a g c c g 3'
3' t t t g t t t a a a a g c t g t a c t t a t a t a a c t a t c a c c a a t t g g a a g a c c t t t t t t t t g t t g g a c t a g a g g a t g t g a t a g a t a t c t c g g c 5'
```

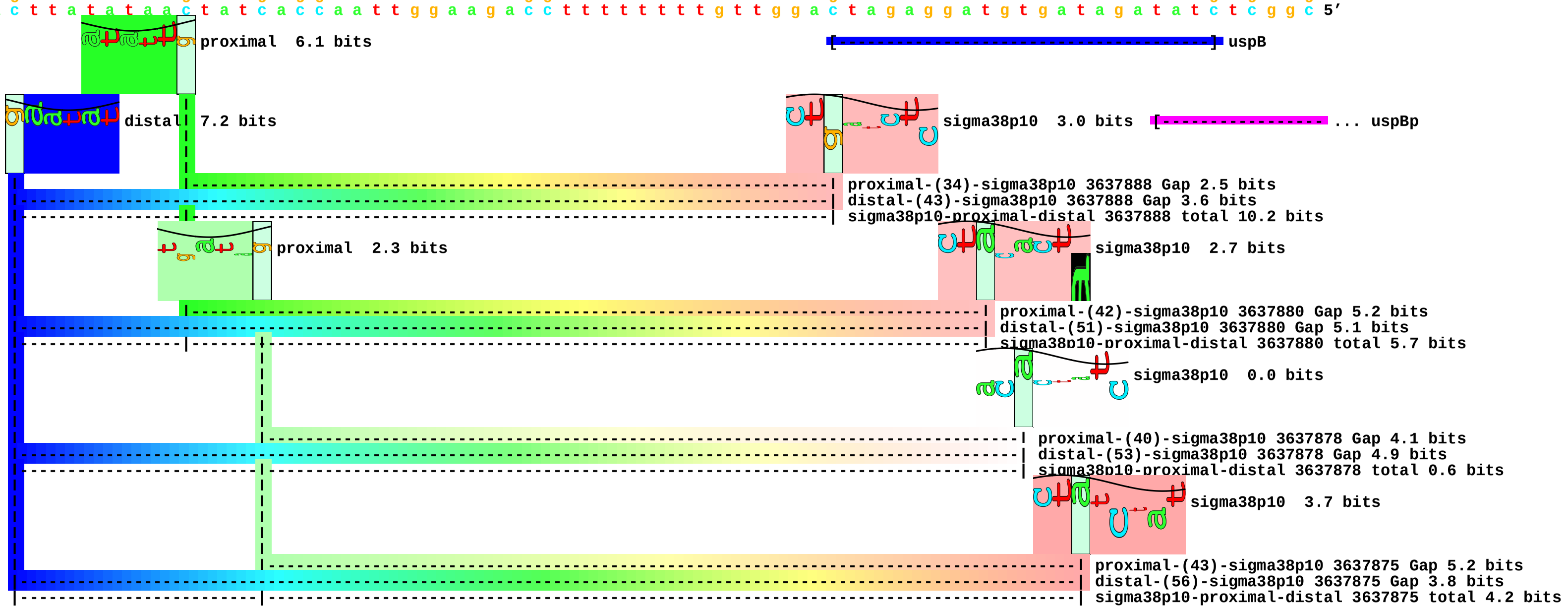

piece 69, NC_000913, gadC, config: linear, direction: -, begin: 1568642, end: 1568557

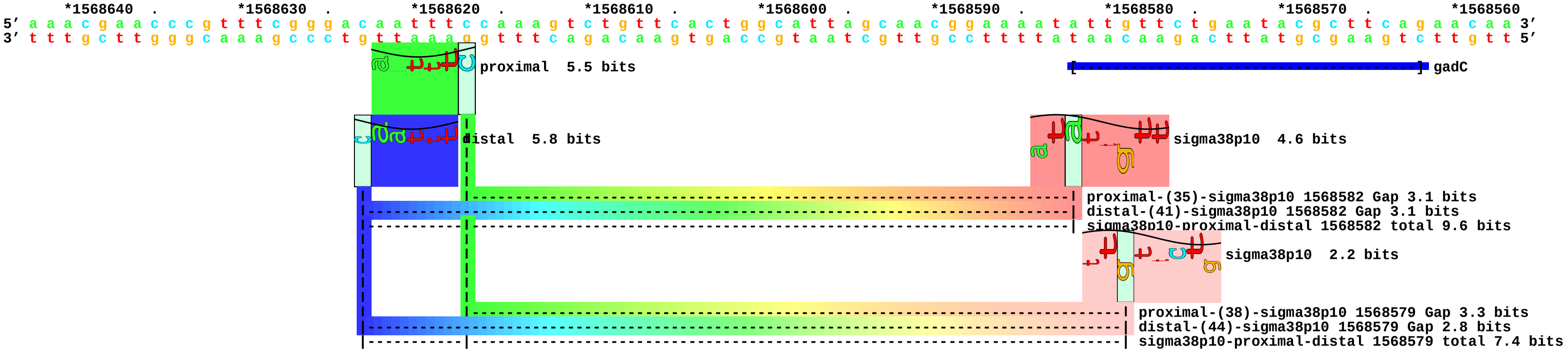

piece 70, NC_000913, cbpA, config: linear, direction: -, begin: 1063122, end: 1063037

```
      *1063120 .        *1063110 .        *1063100 .        *1063090 .        *1063080 .        *1063070 .        *1063060 .        *1063050 .        *1063040
5' c t c c g c g a t g a t t t c a a t a a c a t a t t c t g t g t t g g c a t a t g a a a t t t t g a g g a t t a c c c t a c a c t t a t a g g a g t t a c c t t a c a g g g 3'
3' g a g g c g c t a c t a a a g t t a t t g t a t a a g a c a c a a c c g t a t a c t t t a a a a c t c c t a a t g g g a t g t g a a t a t c c t c a a t g g a a t g t c c c 5'
```

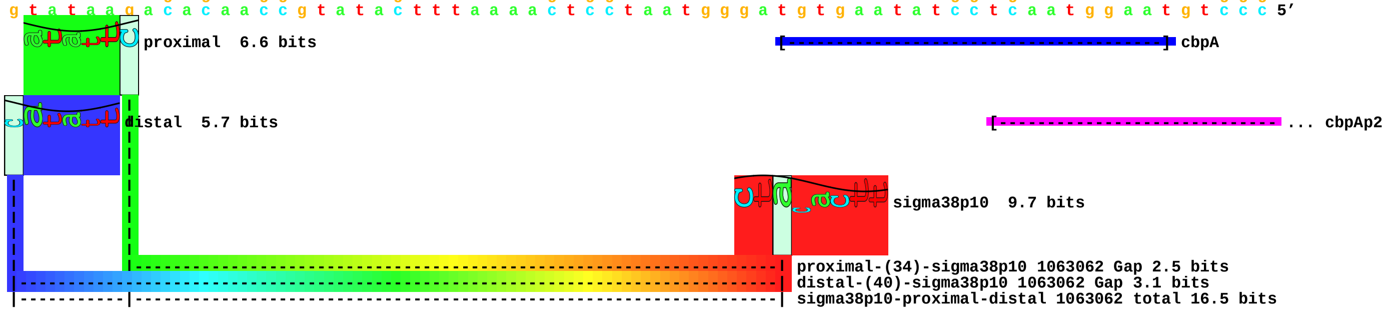

piece 71, NC_000913, hchA, config: linear, direction: +, begin: 2033584, end: 2033669

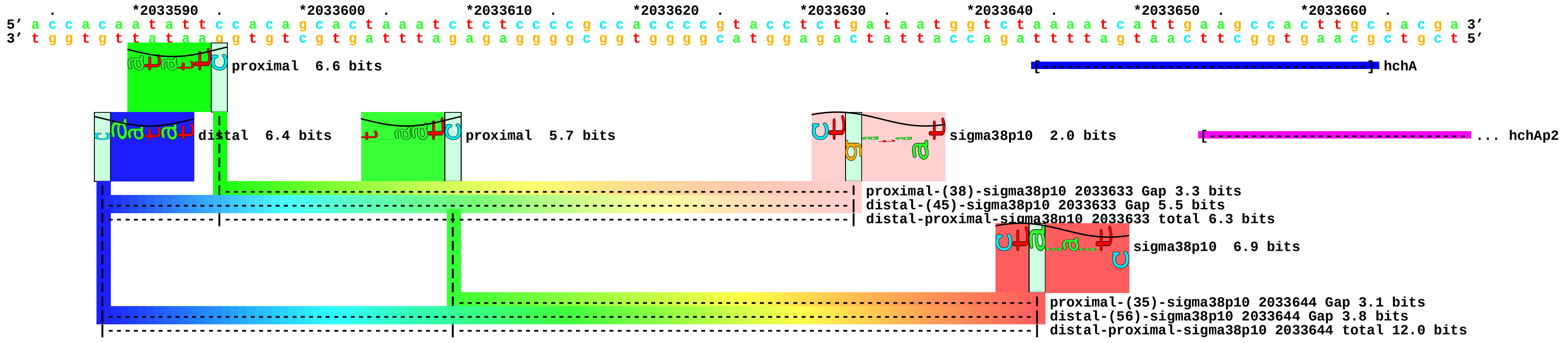

piece 72, NC_000913, sra, config: linear, direction: -, begin: 1554142, end: 1554057

```
       *1554140  .         *1554130  .         *1554120  .         *1554110  .         *1554100  .         *1554090  .         *1554080  .         *1554070  .         *1554060
5' a t c t a c c a g c a a a c g a t c a a t t a t g t g g t c a g t g g c c a g c a c c c t a c g c t t t a a g g t g c t a t g c t t g a t c g g c a a c c t a a t t t a g g 3'
3' t a g a t g g t c g t t t g c t a g t t a a t a c a c c a g t c a c c g g t c g t g g g a t g c g a a a t t c c a c g a t a c g a a c t a g c c g t t g g a t t a a a t c c 5'
```

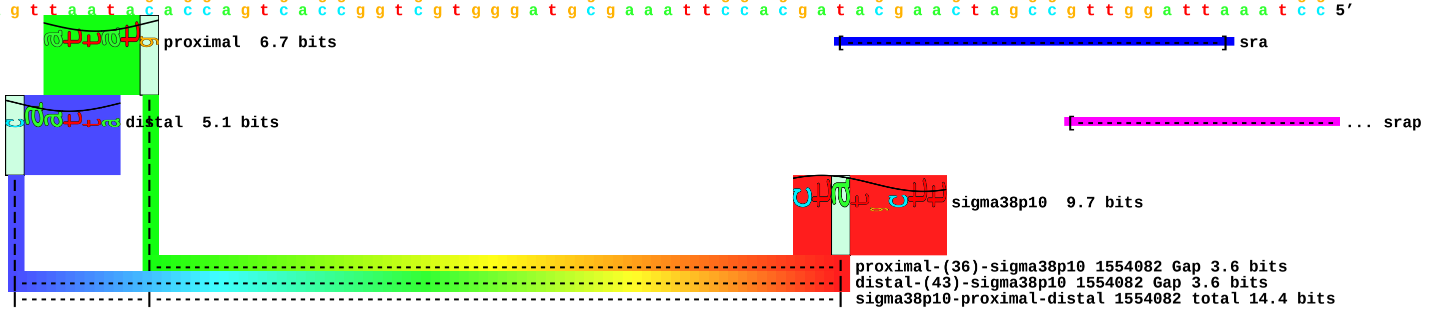

piece 73, NC_000913, hinT, config: linear, direction: +, begin: 1160831, end: 1160916

```
              .         *1160840   .         *1160850   .         *1160860   .         *1160870   .         *1160880   .         *1160890   .         *1160900   .         *1160910   .
5' g a t t a t a t t t a g t g t g c g a a t a a t t t t g t t t g t a a a t g c a a a t g a g a a a t a t a c g c a t t c a t a t t t g t c a t g t a a a c c a a a c a g a g 3'
3' c t a a t a t a a a t c a c a c g c t t a t t a a a a c a a a c a t t t a c g t t t a c t c t t t a t a t g c g t a a g t a t a a a c a g t a c a t t t g g t t t g t c t c 5'
```

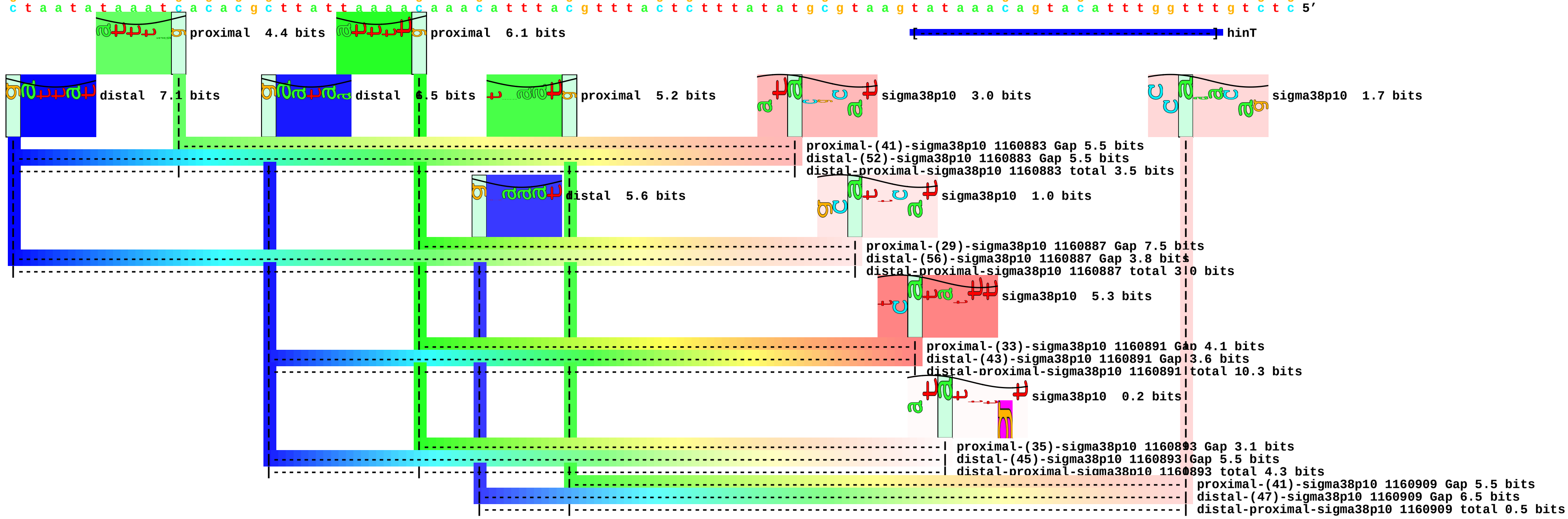

piece 74, NC_000913, yjgH, config: linear, direction: -, begin: 4471353, end: 4471268

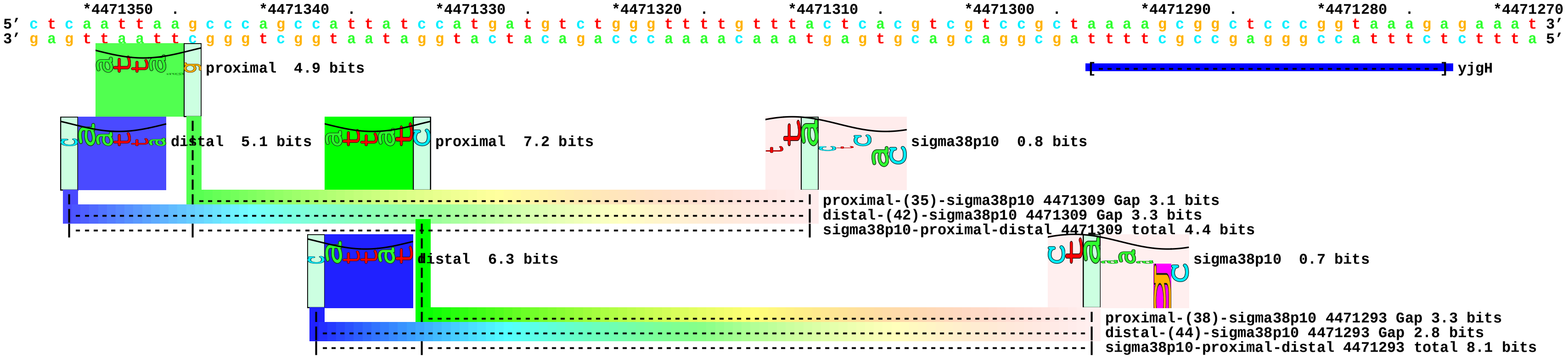

piece 75, NC_000913, yhbO, config: linear, direction: +, begin: 3296789, end: 3296874

```
     *3296790  .         *3296800  .         *3296810  .         *3296820  .         *3296830  .         *3296840  .         *3296850  .         *3296860  .         *3296870
5' a c t t t c g g t t c g t t a a t c a a c a t c c g c a g c a g g t g a c c g c c c a c c a g c c c c g t t g c g c c t g t a a t c a g t a c c t g a c t c a t c t t c g c 3'
3' t g a a a g c c a a g c a a t t a g t t g t a g g c g t c g t c c a c t g g c g g g t g g t c g g g g c a a c g c g g a c a t t a g t c a t g g a c t g a g t a g a a g c g 5'
```

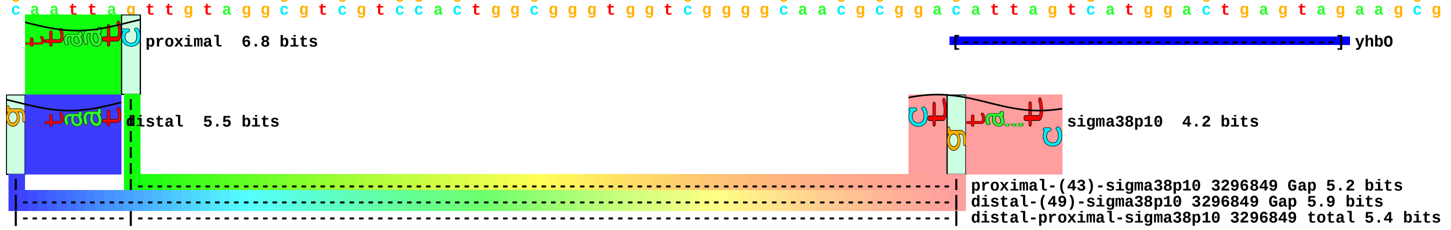

piece 76, NC_000913, csiD, config: linear, direction: +, begin: 2786878, end: 2786963

```
      *2786880  .      *2786890  .      *2786900  .      *2786910  .      *2786920  .      *2786930  .      *2786940  .      *2786950  .      *2786960
5' t t t t g a t c a c a a t a a g a a a a c a a t a t g t c g c t t t t g t g c g c a t t t t t c a g a a a t g t a g a t a t t t t t a g a t t a t g g c t a c g a a a t g a 3'
3' a a a a c t a g t g t t a t t c t t t t g t t a t a c a g c g a a a a c a c g c g t a a a a a g t c t t t a c a t c t a t a a a a a t c t a a t a c c g a t g c t t t a c t 5'
```

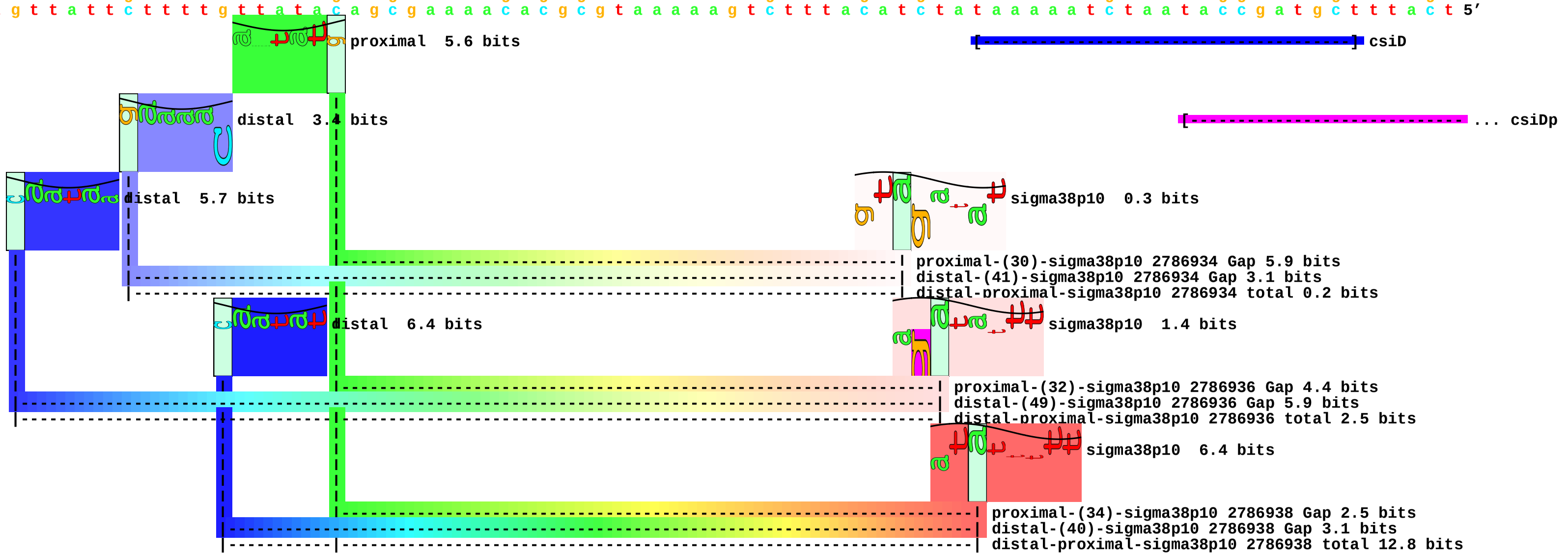

piece 77, NC_000913, elaB, config: linear, direction: -, begin: 2379200, end: 2379115

```
   *2379200  .         *2379190  .         *2379180  .         *2379170  .         *2379160  .         *2379150  .         *2379140  .         *2379130  .         *2379120  .
5' c t t c t a c c a g a g t t t c g g t t t t a t c c c t g t g a c g g a g g t c t a t g a a g a a g a t g g t a t c c c a c a c a t t g g g a t g g c a c g c g a g g t a a 3'
3' g a a g a t g g t c t c a a a g c c a a a a t a g g g a c a c t g c c t c c a g a t a c t t c t t c t a c c a t a g g g t g t g t a a c c c t a c c g t g c g c t c c a t t 5'
```

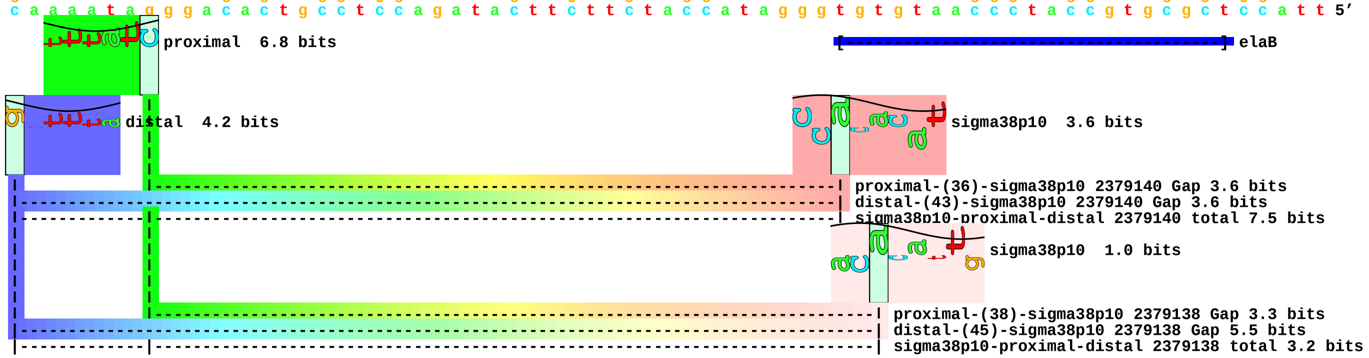

piece 78, NC_000913, erfK, config: linear, direction: -, begin: 2061448, end: 2061363

```
         .         *2061440  .         *2061430  .         *2061420  .         *2061410  .         *2061400  .         *2061390  .         *2061380  .         *2061370  .
5' t c t a c c g g g g a a t a c g a c t t c t g a t t t g a a c a g t t a a a a a c g a a a a g t c t g a t a a a a g c t a t a c t t a a c g g a t a g c t t t c g c g a c a 3'
3' a g a t g g c c c c t t a t g c t g a a g a c t a a a c t t g t c a a t t t t t g c t t t t c a g a c t a t t t t c g a t a t g a a t t g c c t a t c g a a a g c g c t g t 5'
```

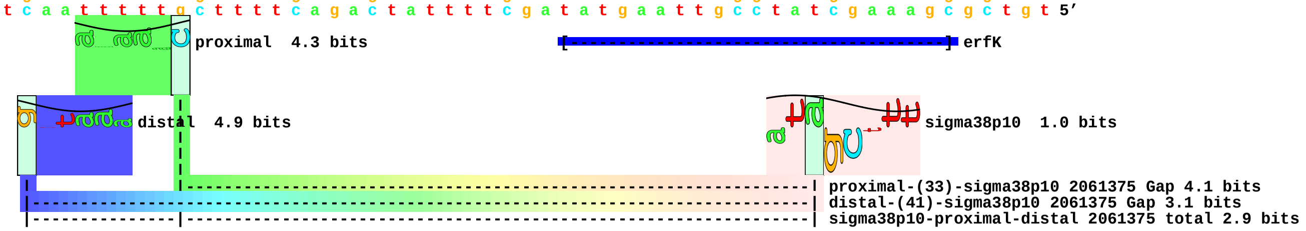

piece 79, NC_000913, fic, config: linear, direction: -, begin: 3489740, end: 3489655

```
   *3489740  .         *3489730  .         *3489720  .         *3489710  .         *3489700  .         *3489690  .         *3489680  .         *3489670  .         *3489660  .
5' c t c g c g c g g g c a a t c t t g c c c g c g c t t c t g c t c t c c c g g c g t a a c c c g g a t t t g c c g c t t a t a c t t g t g g c a a a t g g a c a c g t t c a 3'
3' g a g c g c g c c c g t t a g a a c g g g c g c g a a g a c g a g a g g g c c g c a t t g g g c c t a a a c g g c g a a t a t g a a c a c c g t t t a c c t g t g c a a g t 5'
```

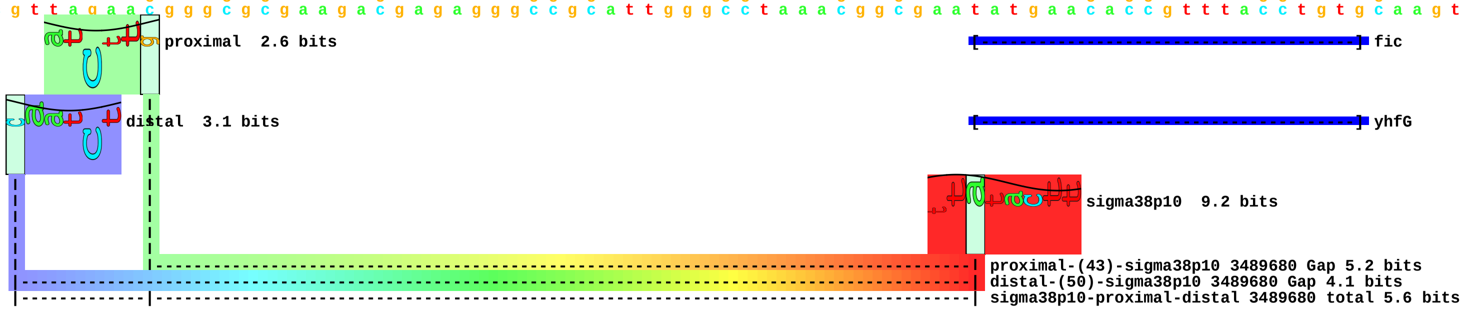

piece 80, NC_000913, msyB, config: linear, direction: -, begin: 1113507, end: 1113422

```
          .         *1113500  .         *1113490  .         *1113480  .         *1113470  .         *1113460  .         *1113450  .         *1113440  .         *1113430  .
5' a t a c c c c a g g t a t c g a a c t g a t t t t t c g c c t t t c a t a c t t g c a a a a g c g g a g a a t c a g c t a t c c t t t t c c c t g a a a c c t c a t c a a c 3'
3' t a t g g g g t c c a t a g c t t g a c t a a a a a g c g g a a a g t a t g a a c g t t t t c g c c t c t t a g t c g a t a g g a a a a g g g a c t t t g g a g t a g t t g 5'
```

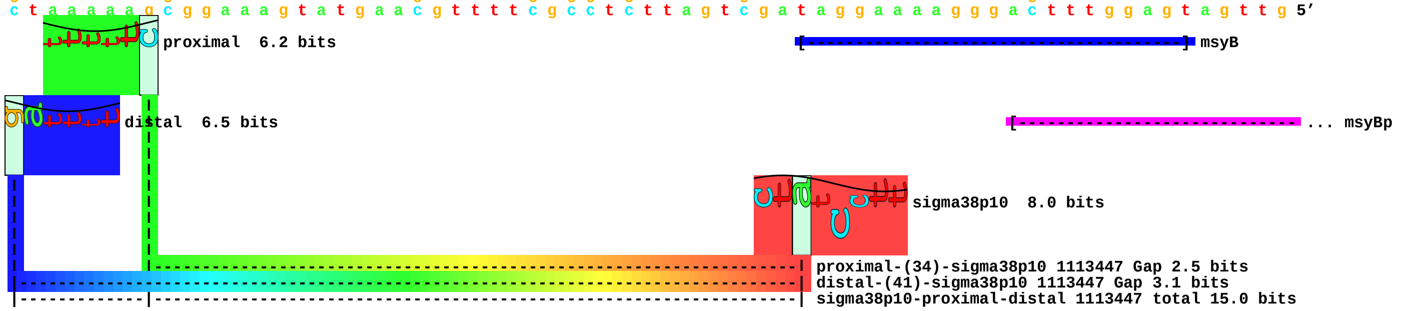

piece 81, NC_000913, yaiA, config: linear, direction: +, begin: 406055, end: 406140

```
   .         *406060   .         *406070   .         *406080   .         *406090   .         *406100   .         *406110   .         *406120   .         *406130   .         *406140
5' t a t c g c g a a g t t g c g c a t a t t a t c a t c g a c g c a a c a a a c g a a c c c a g c c a g g t g a t t t c t g a a a t t c g c a g c g c c c t g g c a c a g a c 3'
3' a t a g c g c t t c a a c g c g t a t a a t a g t a g c t g c g t t g t t t g c t t g g g t c g g t c c a c t a a a g a c t t t a a g c g t c g c g g g a c c g t g t c t g 5'
```

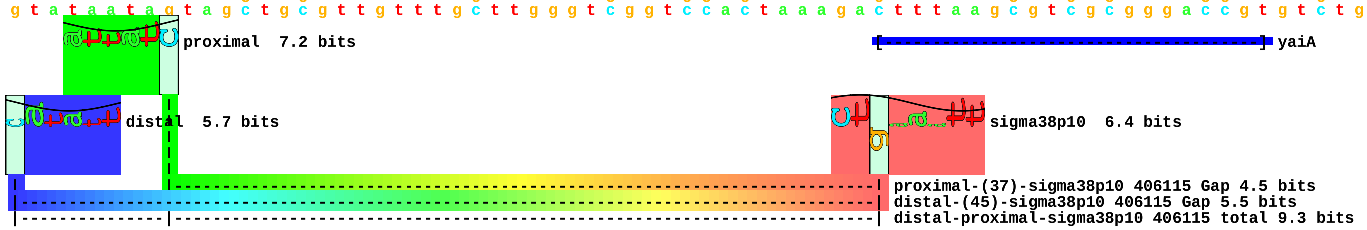

piece 82, NC_000913, yajO, config: linear, direction: -, begin: 437579, end: 437494

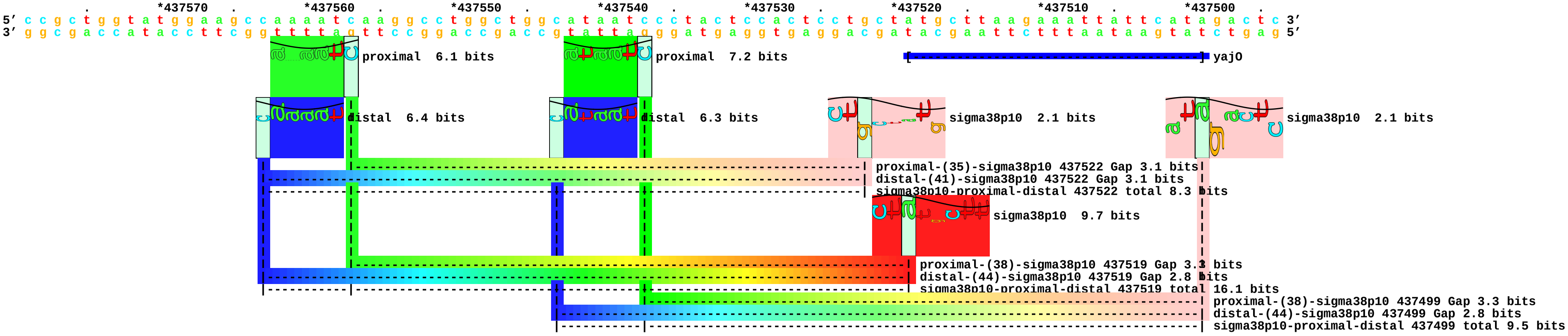

piece 83, NC_000913, ybdK, config: linear, direction: -, begin: 606701, end: 606616

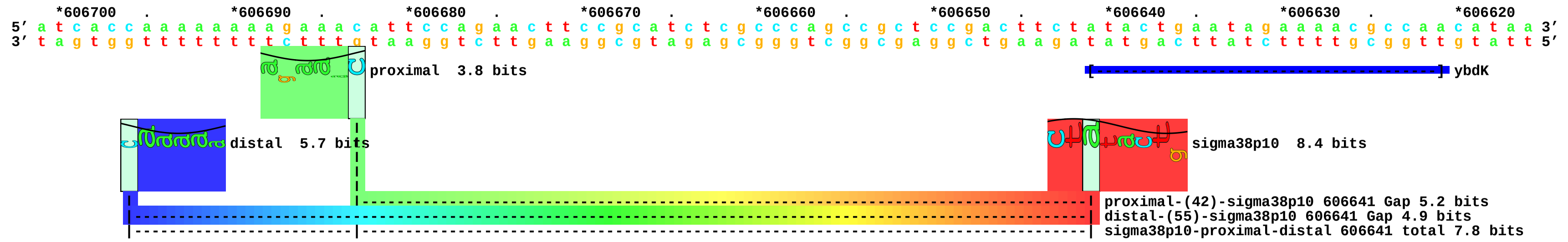

piece 84, NC_000913, ybeL, config: linear, direction: +, begin: 674144, end: 674229

```
    .         *674150   .         *674160   .         *674170   .         *674180   .         *674190   .         *674200   .         *674210   .         *674220   .
5' t c a t a c a t t a a t t t a c a g a g t t t g t g g g c g t a t t a g c a a a g c a a g g a a c a a a g a a c g t c t a t t a t t a t a g t c a g t t a a c g a c c c g g 3'
3' a g t a t g t a a t t a a a t g t c t c a a a c a c c c g c a t a a t c g t t t c g t t c c t t g t t t c t t g c a g a t a a t a a t a t c a g t c a a t t g c t g g g c c 5'
```

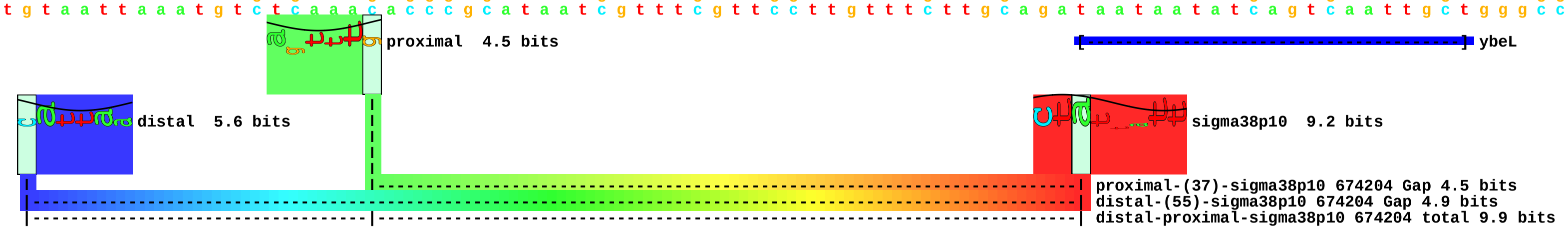

piece 85, NC_000913, ybgAp1, config: linear, direction: +, begin: 738131, end: 738216

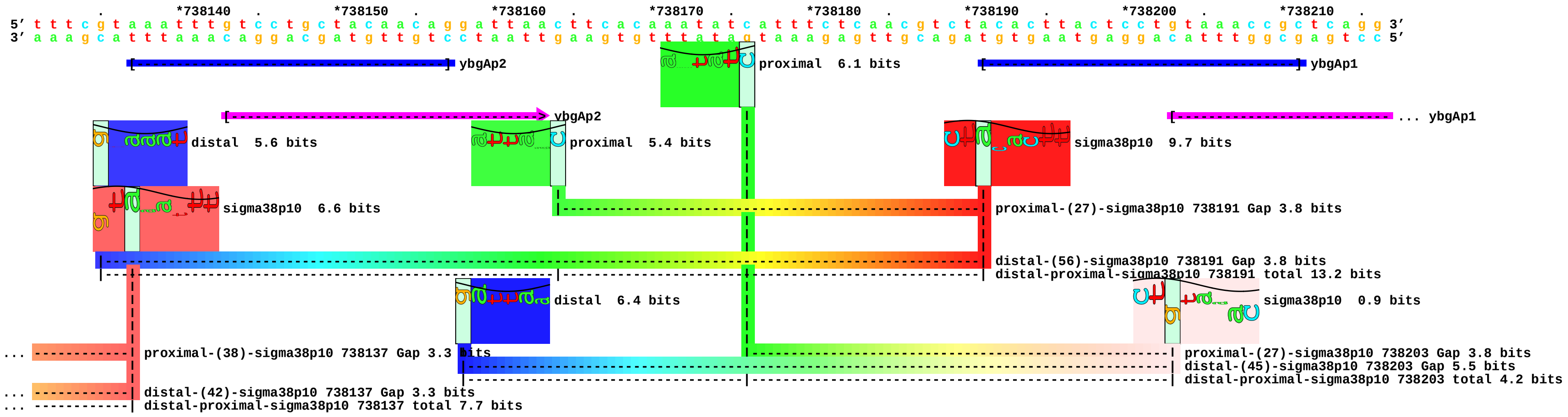

piece 86, NC_000913, ybgAp2, config: linear, direction: +, begin: 738077, end: 738162

```
          *738080   .         *738090   .         *738100   .         *738110   .         *738120   .         *738130   .         *738140   .         *738150   .         *738160
5' a a a g g g c a g c t a t g g t t a g a a a c t a c c t g a c g t c a g t c c t t g c g g g g a g c a g g c t t t c g t a a a t t t g t c c t g c t a c a a c a g g a t t a 3'
3' t t t c c c g t c g a t a c c a a t c t t t g a t g g a c t g c a g t c a g g a a c g c c c c t c g t c c g a a a g c a t t t a a a c a g g a c g a t g t t g t c c t a a t 5'
```

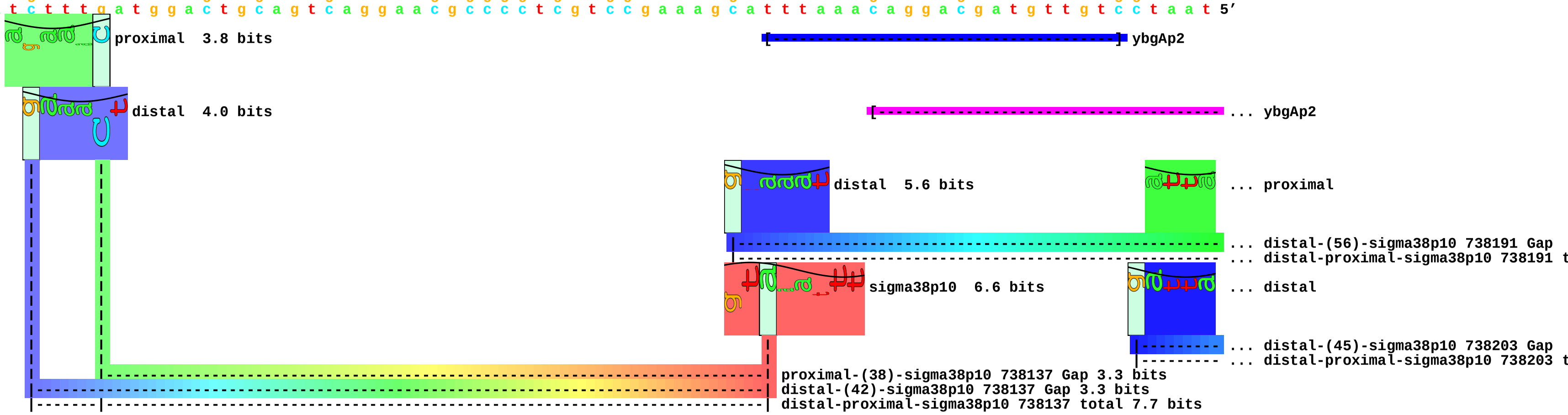

piece 87, NC_000913, ybgS, config: linear, direction: -, begin: 784758, end: 784673

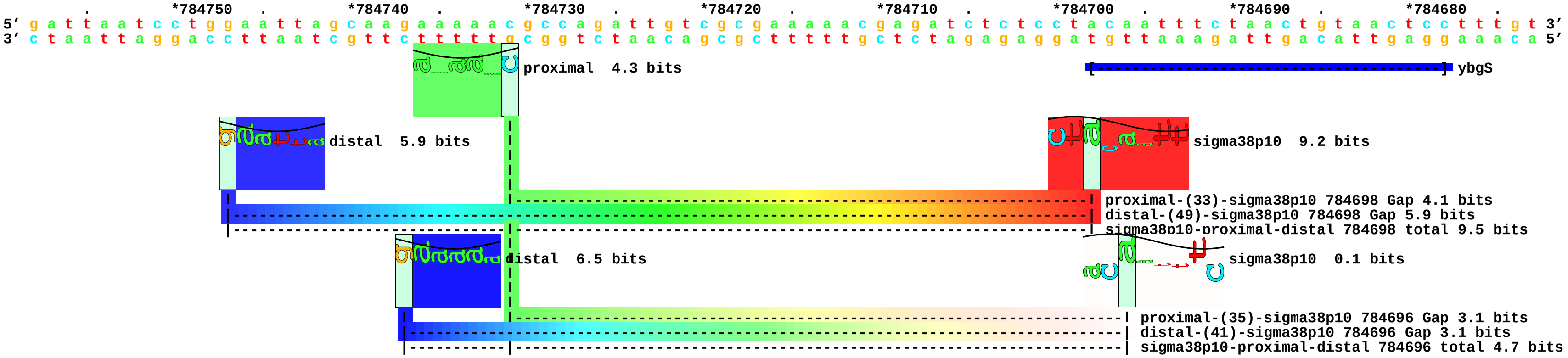

piece 88, NC_000913, pgl, config: linear, direction: +, begin: 797705, end: 797790

```
 .         *797710   .         *797720   .         *797730   .         *797740   .         *797750   .         *797760   .         *797770   .         *797780   .         *797790
5' c g a t a c c a c t a t c g g a c a a a a t t c t g c g t t t t a a t t c a g c a t t c a c c g c c a a a a g c g a c t a a t t t t a g c t g t t a c a g t c a g t t g c t 3'
3' g c t a t g g t g a t a g c c t g t t t t a a g a c g c a a a a t t a a g t c g t a a g t g g c g g t t t t c g c t g a t t a a a a t c g a c a a t g t c a g t c a a c g a 5'
```

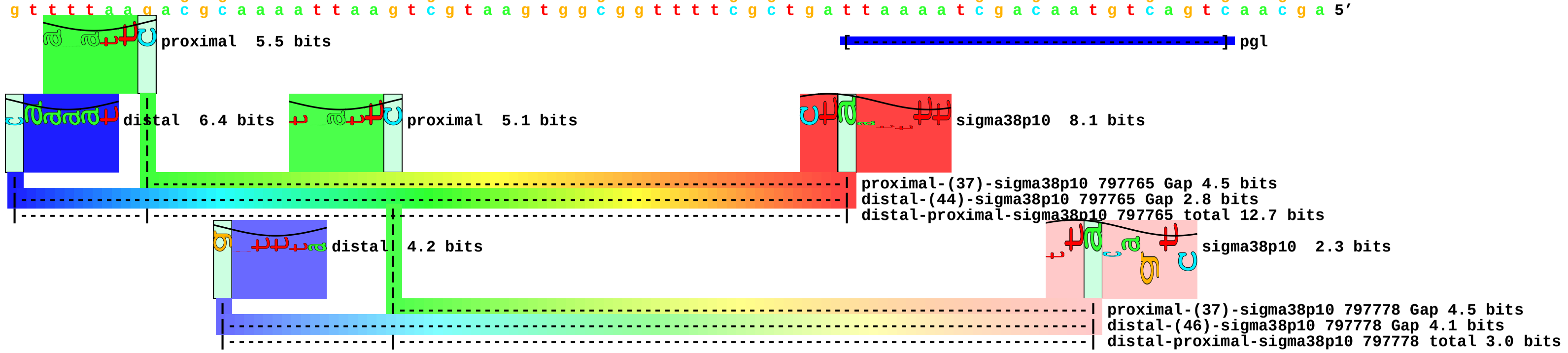

piece 89, NC_000913, ybjP, config: linear, direction: -, begin: 903796, end: 903711

```
    .         *903790   .         *903780   .         *903770   .         *903760   .         *903750   .         *903740   .         *903730   .         *903720   .
5’ t a a g c c t t c g a t c t c a a a a g c a t t a t c a g a c t g a t a c g c t a t t a t t g a a a g g a t a t c a t t a t t a t c g t a g a a c g c t t t c a g a g c g a 3’
3’ a t t c g g a a g c t a g a g t t t t c g t a a t a g t c t g a c t a t g c g a t a a t a a c t t t c c t a t a g t a a t a a t a g c a t c t t g c g a a a g t c t c g c t 5’
```

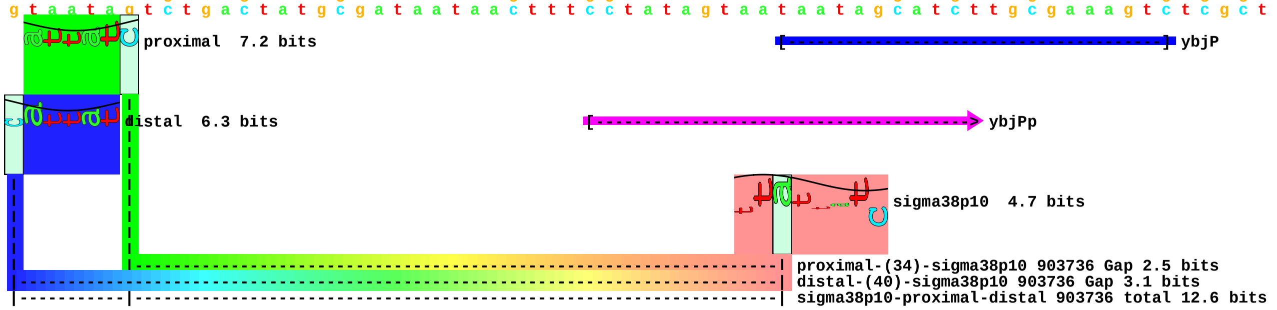

piece 90, NC_000913, yceK, config: linear, direction: +, begin: 1112718, end: 1112803

```
       *1112720  .         *1112730  .         *1112740  .         *1112750  .         *1112760  .         *1112770  .         *1112780  .         *1112790  .         *1112800
5' g c g t t c a c g c c g c a t c c g g c a a a a a c a a c g a g c c a a t a a a a a t a c c g g c g t t a t g c c g g t a t t t t t t t a c g a a a g a g g t a t c a a a t 3'
3' c g c a a g t g c g g c g t a g g c c g t t t t t g t t g c t c g g t t a t t t t t a t g g c c g c a a t a c g g c c a t a a a a a a a t g c t t t c t c c a t a g t t t a 5'
```

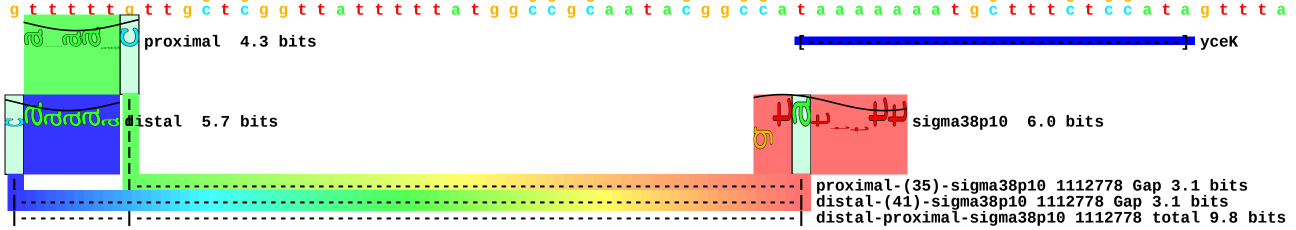

piece 91, NC_000913, ycgB, config: linear, direction: -, begin: 1236579, end: 1236494

```
              .         *1236570  .         *1236560  .         *1236550  .         *1236540  .         *1236530  .         *1236520  .         *1236510  .         *1236500  .
5' g a t a a t a a g a a a t c g c a a a g a a a a a a a t t c t c c g g c g a c g c g g t t t t t a a c a g g t g t t c t a t g c t t g a a a t g a g g t g c t c c a c a c a 3'
3' c t a t t a t t c t t t a g c g t t t c t t t t t t t a a g a g g c c g c t g c g c c a a a a a t t g t c c a c a a g a t a c g a a c t t t a c t c c a c g a g g t g t g t 5'
```

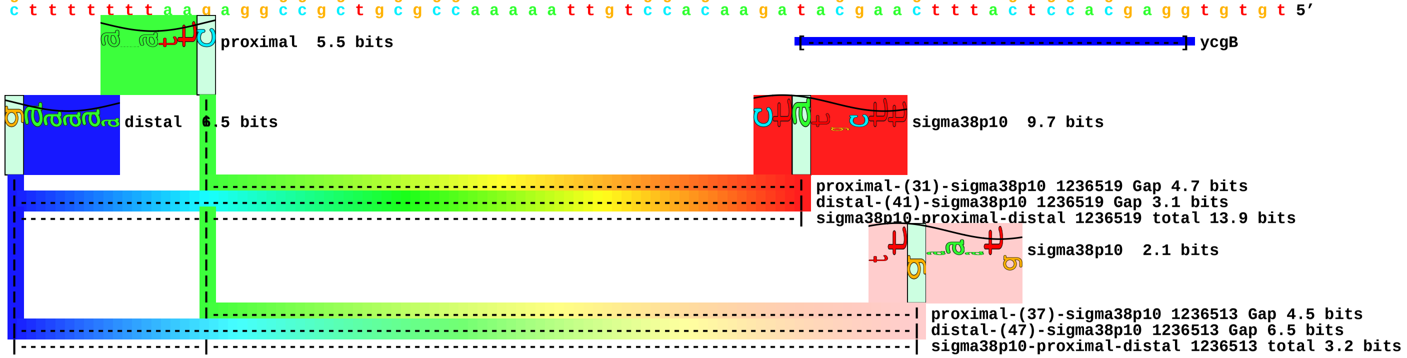

piece 92, NC_000913, ycgZ, config: linear, direction: +, begin: 1214904, end: 1214989

```
      .    *1214910   .    *1214920   .    *1214930   .    *1214940   .    *1214950   .    *1214960   .    *1214970   .    *1214980   .
5' t a g c a c t a a t t g c a a a a a a t t a a t t t a t c a t t c t g t a c a c a t a t t t c g t a c a a g t t t g c t a t t g t t a c t t c a c t t a a c a t t g a t t a 3'
3' a t c g t g a t t a a c g t t t t t t a a t t a a a t a g t a a g a c a t g t g t a t a a a g c a t g t t c a a a c g a t a a c a a t g a a g t g a a t t g t a a c t a a t 5'
```

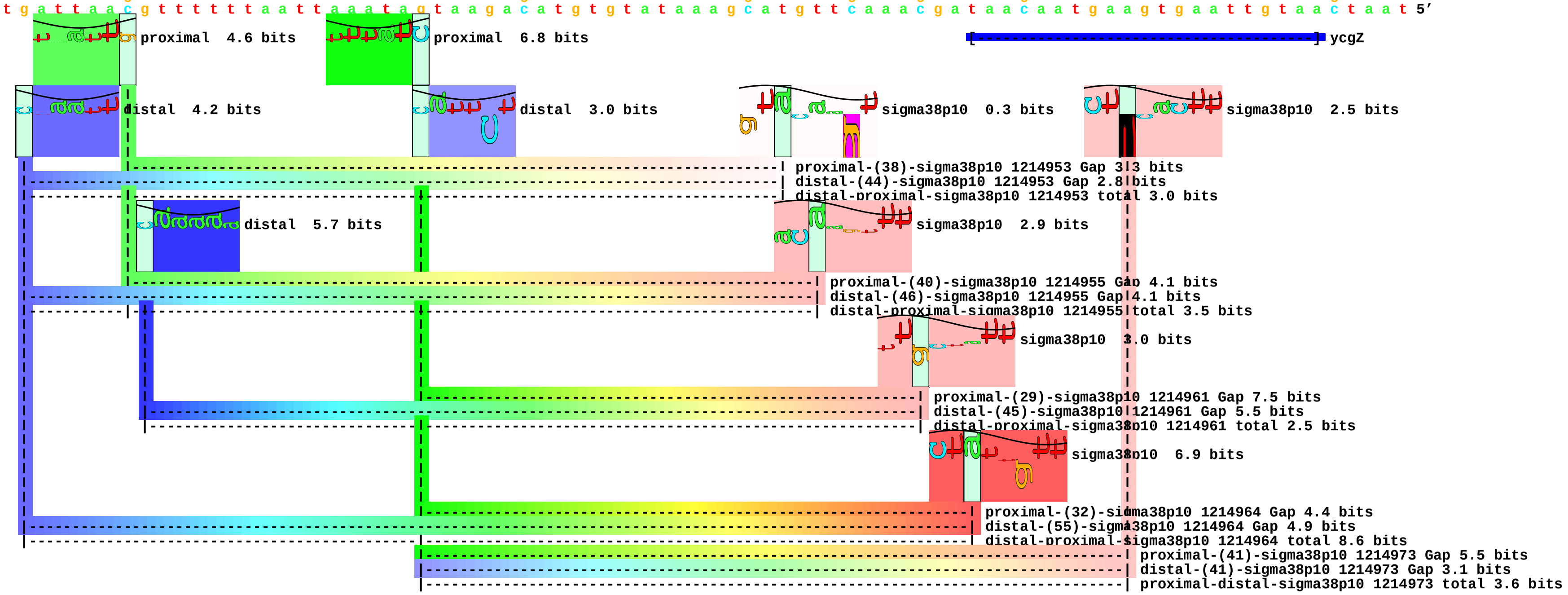

piece 93, NC_000913, yciF, config: linear, direction: -, begin: 1314243, end: 1314158

```
          *1314240  .         *1314230  .         *1314220  .         *1314210  .         *1314200  .         *1314190  .         *1314180  .         *1314170  .         *1314160
5' t t t c t c t a a g t c a c t t a t t c c t c a g g t a a t t g t t a a t a t a t c c a g a a t g t t c c t c a a a a t a t a t t t t c c c t c t a t c t t c t c g t t g c 3'
3' a a a g a g a t t c a g t g a a t a a g g a g t c c a t t a a c a a t t a t a t a g g t c t t a c a a g g a g t t t t a t a t a a a a g g g a g a t a g a a g a g c a a c g 5'
```

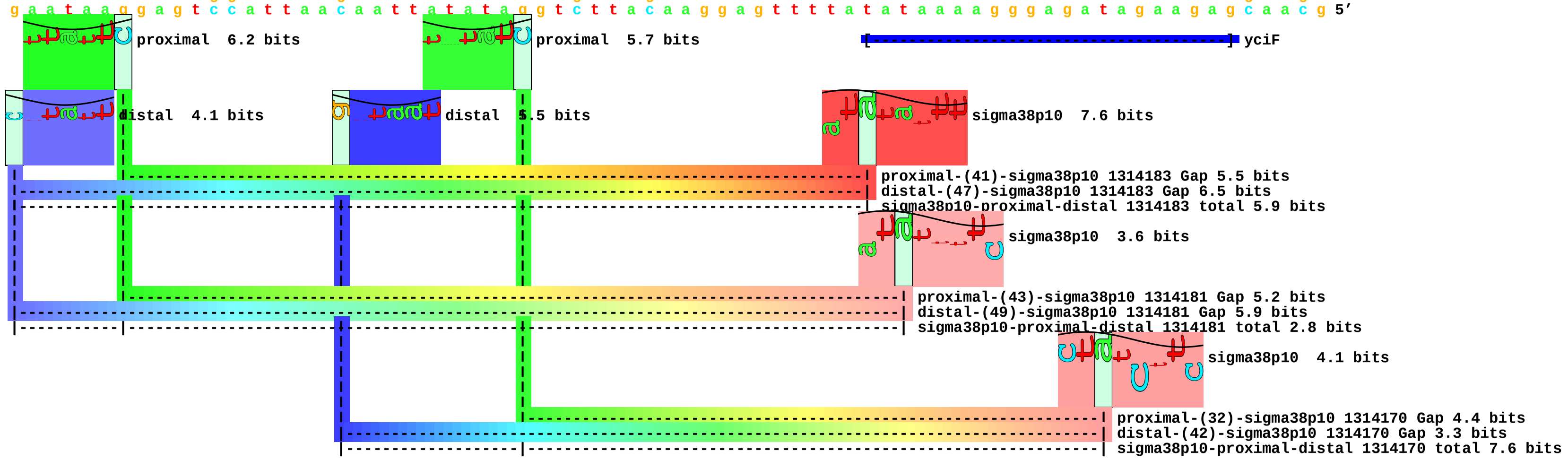

piece 94, NC_000913, yciG, config: linear, direction: -, begin: 1314426, end: 1314341

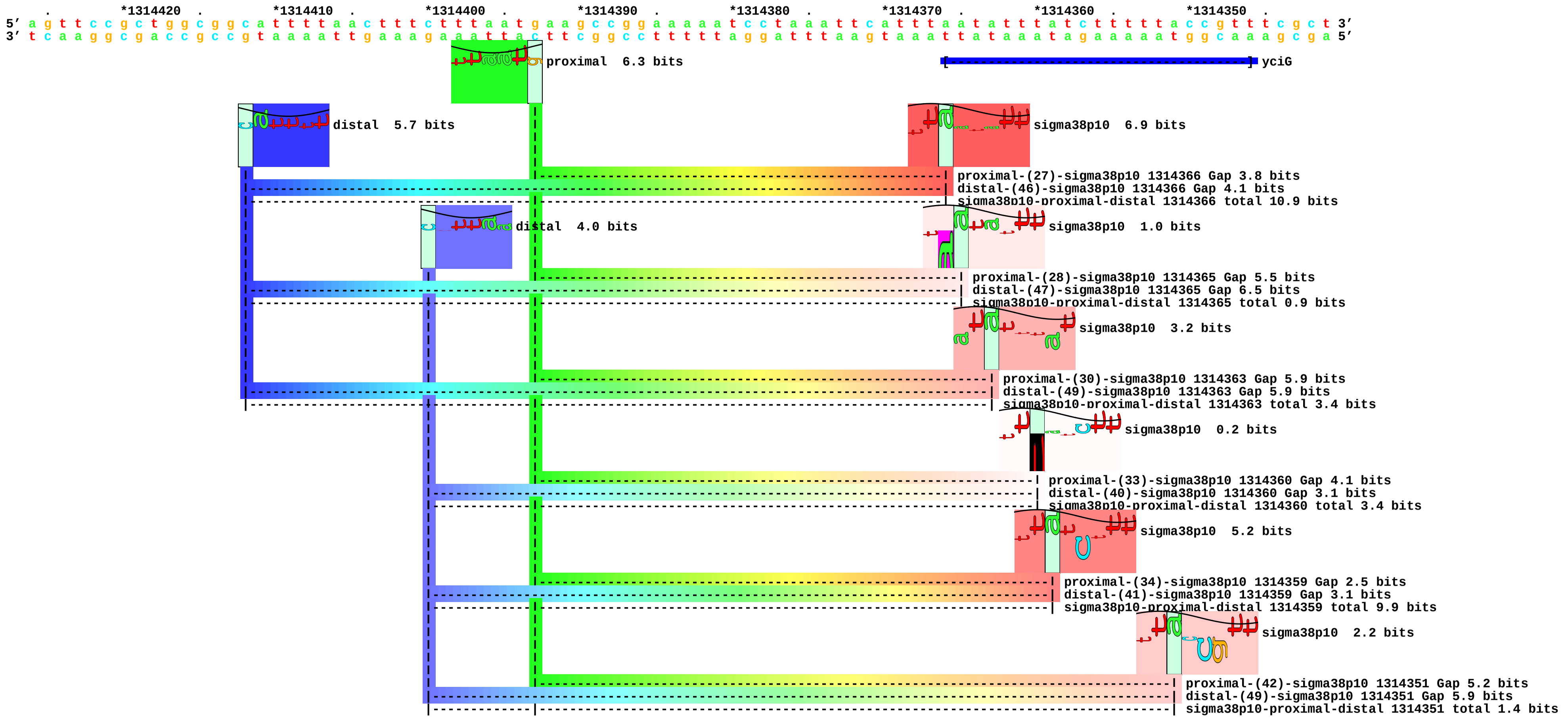

piece 95, NC_000913, ydaM, config: linear, direction: -, begin: 1405964, end: 1405879

```
  *1405960 .         *1405950 .         *1405940 .         *1405930 .         *1405920 .         *1405910 .         *1405900 .         *1405890 .         *1405880
5' g a a t a c a a a a a g c a a a a t g c t t t t c c g t a t a c a a a c c g t g t g a a g t g t t a a a t a g c g t c t a t c a t t a t c a g a a t t a t c t g a t c a t a 3'
3' c t t a t g t t t t t c g t t t t a c g a a a a g g c a t a t g t t t g g c a c a c t t c a c a a t t t a t c g c a g a t a g t a a t a g t c t t a a t a g a c t a g t a t 5'
```

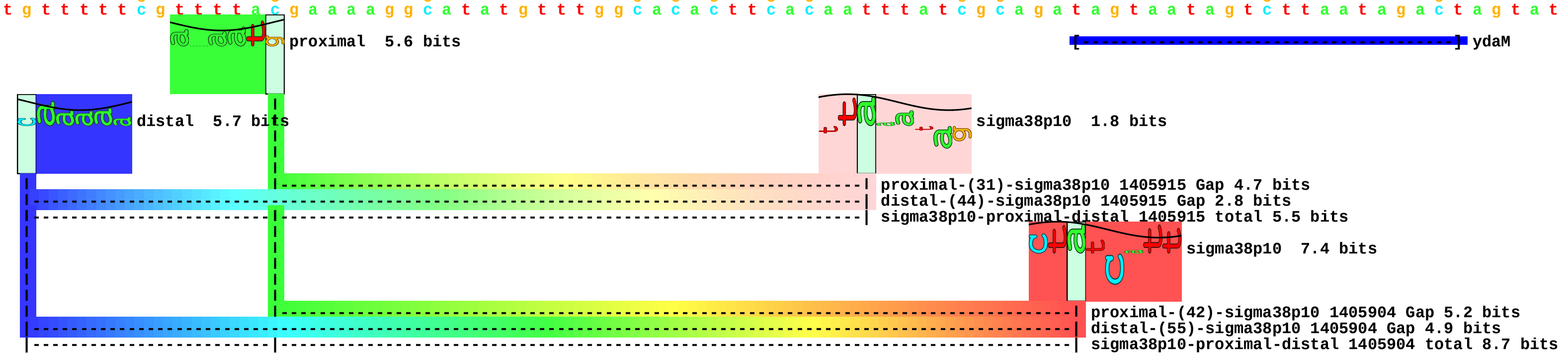

piece 96, NC_000913, stfQ, config: linear, direction: -, begin: 1634111, end: 1634026

```
     *1634110  .          *1634100  .          *1634090  .          *1634080  .          *1634070  .          *1634060  .          *1634050  .          *1634040  .          *1634030
5' c c g a c g c a c a g g a g c t g a a a a a t g c c a g a g a c t c c g c t g a a g t g g t g g a a a c c g c a t t c t g t a c t t t c g t g c t g t c g c g g a t c g c a 3'
3' g g c t g c g t g t c c t c g a c t t t t t a c g g t c t c t g a g g c g a c t t c a c c a c c t t t g g c g t a a g a c a t g a a a g c a c g a c a g c g c c t a g c g t 5'
```

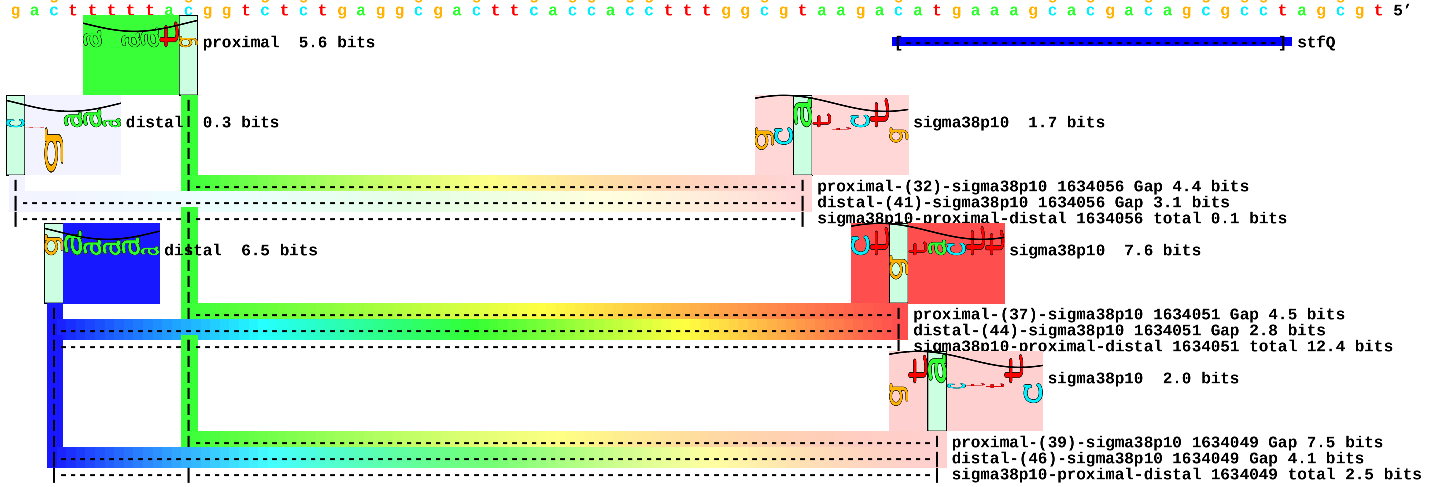

piece 97, NC_000913, ydgA, config: linear, direction: +, begin: 1687747, end: 1687832

```
         *1687750  .         *1687760  .         *1687770  .         *1687780  .         *1687790  .         *1687800  .         *1687810  .         *1687820  .         *1687830
5' g t t t a g c g c g t g t t t a c a a c a a g c t g t a a g a g c t t a c t g a a a a a a t t a a c a t c t c t t g c t a a g c t t a t t a a a g g c t t a t a a c a c c t 3'
3' c a a a t c g c g c a c a a a t g t t g t t c g a c a t t c t c g a a t g a c t t t t t t a a t t g t a g a g a a c g a t t c g a a t a a t t t c c g a a t a t t g t g g a 5'
```

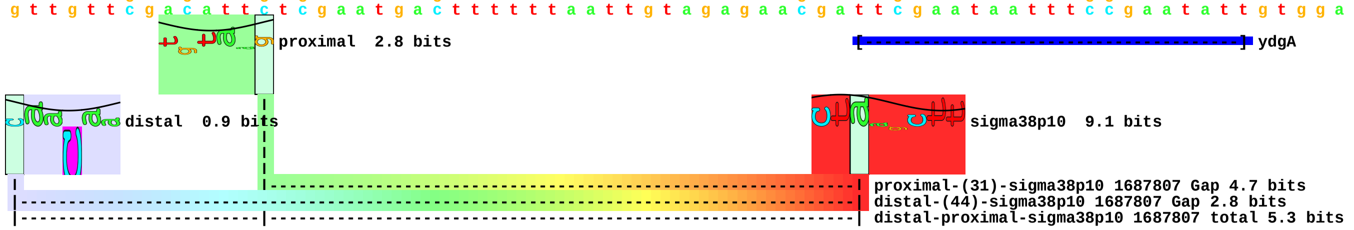

piece 98, NC_000913, yeaG, config: linear, direction: +, begin: 1864699, end: 1864784

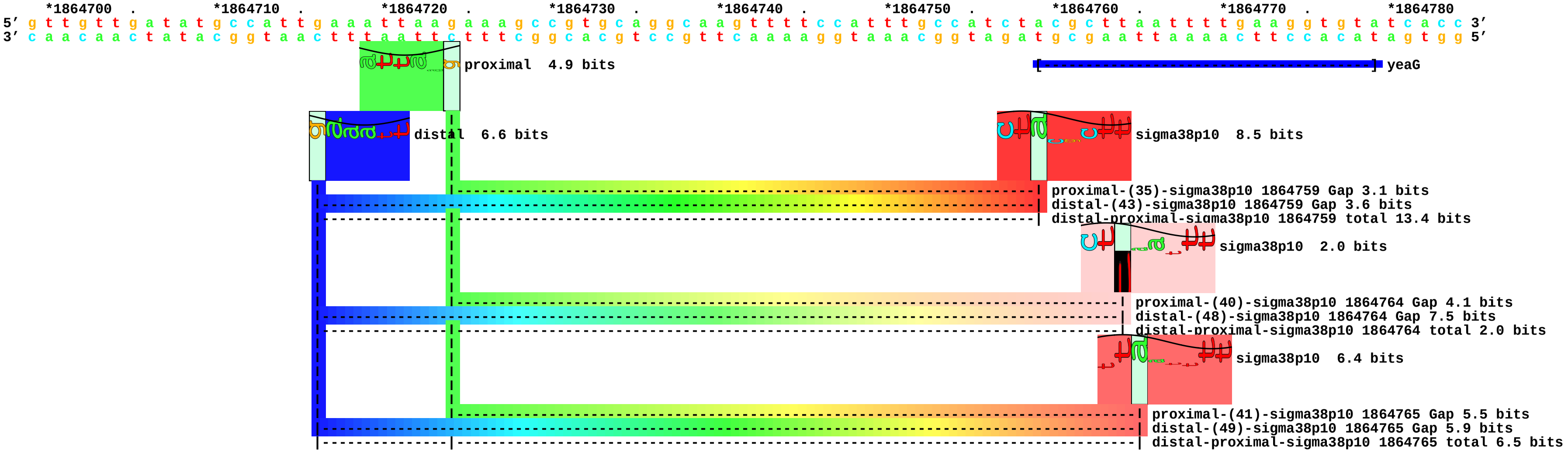

piece 99, NC_000913, yebF, config: linear, direction: -, begin: 1928869, end: 1928784

```
         .         *1928860  .         *1928850  .         *1928840  .         *1928830  .         *1928820  .         *1928810  .         *1928800  .         *1928790  .
5' a a a c g a a a a c g t t t g c g t g t g t c t t t a t t g c t g a t g t t g a t t t c t c a a c c g a a a a g a a a t a t a c t g t a t a a a a t c a c a g t t a t t a t 3'
3' t t t g c t t t t g c a a a c g c a c a c a g a a a t a a c g a c t a c a a c t a a a g a g t t g g c t t t t c t t t a t a t g a c a t a t t t t a g t g t c a a t a a t a 5'
```

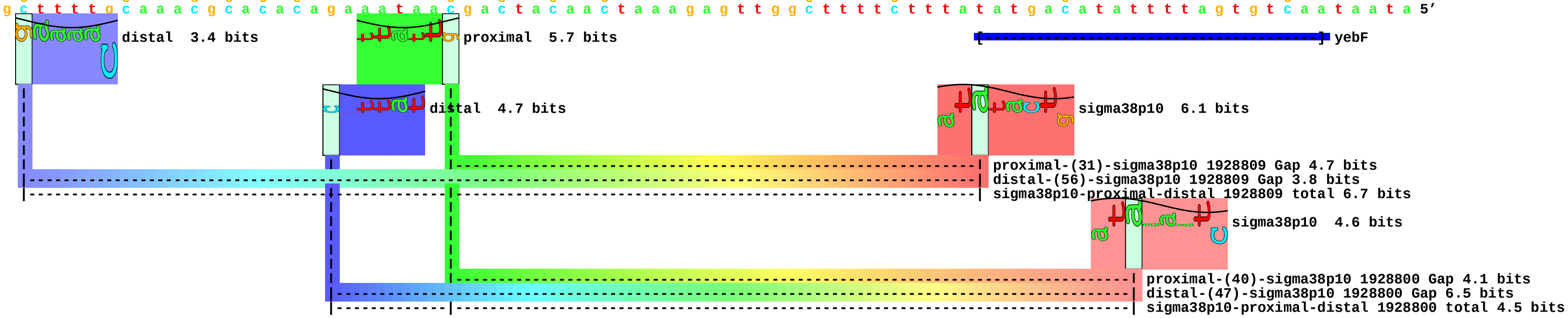

piece 100, NC_000913, yeeP, config: linear, direction: +, begin: 2068415, end: 2068500

```
 .         *2068420  .         *2068430  .         *2068440  .         *2068450  .         *2068460  .         *2068470  .         *2068480  .         *2068490  .         *2068500
5' g a g g t a g t g a a a g g t c c a t a t t c c g t a c t g t a c g g t t c a c a g g c a a t t g g c g g t a t t g t t a a c t t c a t c a c c a a a a a g g g t g a a t c 3'
3' c t c c a t c a c t t t c c a g g t a t a a g g c a t g a c a t g c c a a g t g t c c g t t a a c c g c c a t a a c a a t t g a a g t a g t g g t t t t t c c c a c t t a g 5'
```

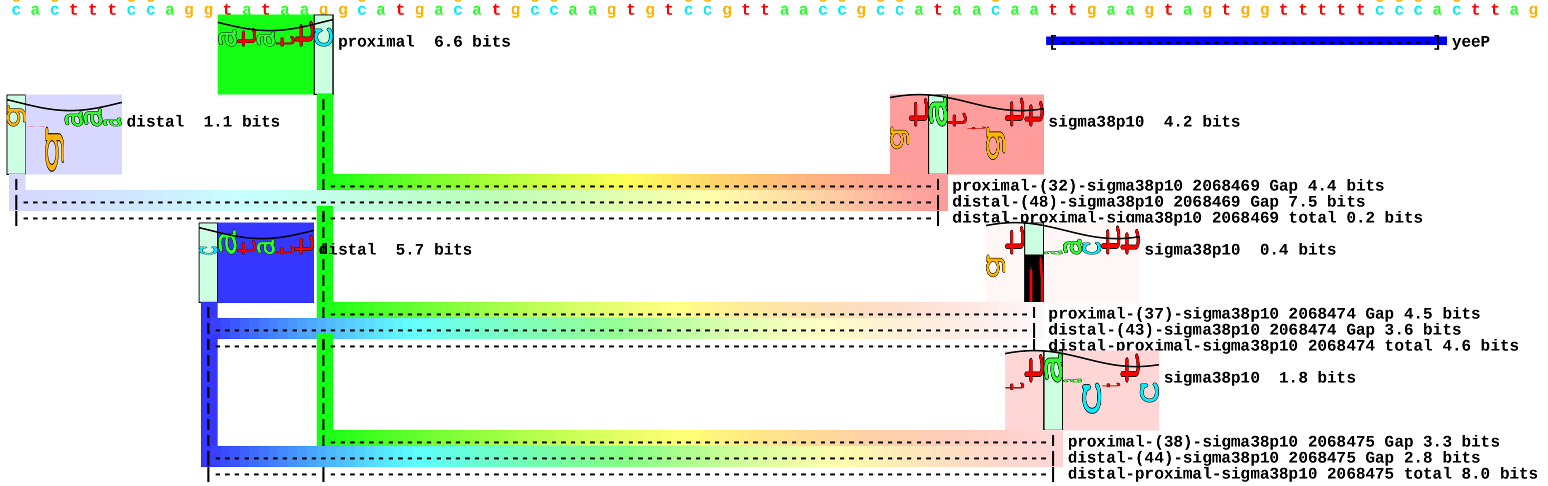

piece 101, NC_000913, yegP, config: linear, direction: +, begin: 2162708, end: 2162793

```
       *2162710  .         *2162720  .         *2162730  .         *2162740  .         *2162750  .         *2162760  .         *2162770  .         *2162780  .         *2162790
5' c a g g a t g c t g a a a g c c c g t t g a t t a t c g a c g a a g g t c g t t t t c a g g c t t c a t g g c g c g g t a a a a t g c t t g a c c t g a c g c c t g c g g a 3'
3' g t c c t a c g a c t t t c g g g c a a c t a a t a g c t g c t t c c a g c a a a a g t c c g a a g t a c c g c g c c a t t t t a c g a a c t g g a c t g c g g a c g c c t 5'
```

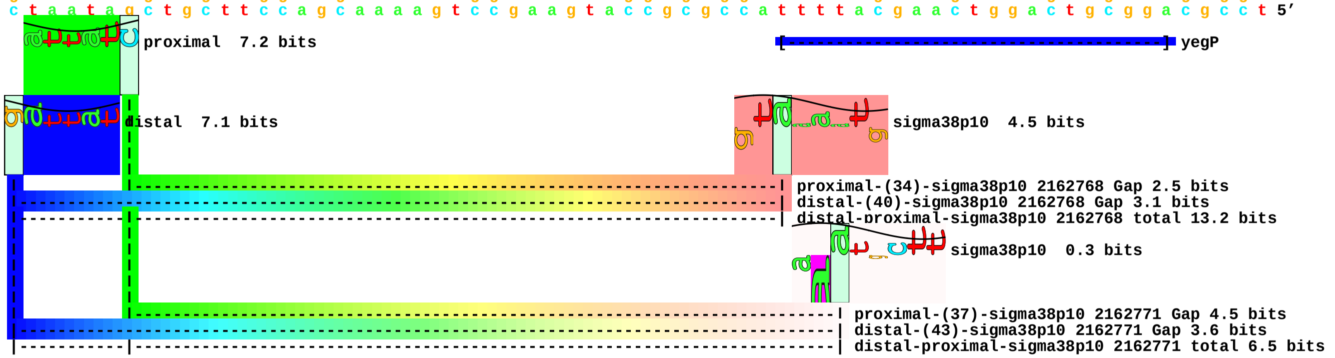

piece 102, NC_000913, yegS, config: linear, direction: +, begin: 2166484, end: 2166569

```
    .         *2166490  .         *2166500  .         *2166510  .         *2166520  .         *2166530  .         *2166540  .         *2166550  .         *2166560  .
5’ t c c a t g g g g g c t t t a a t a t a t t a t a t t g a a g a t g c c a c t g t t t a g t t g a a t a t t a g g t a t a t g c t c t t t t t t g a a a t t t a t c g g t g 3’
3’ a g g t a c c c c c g a a a t t a t a t a a t a t a a c t t c t a c g g t g a c a a a t c a a c t t a t a a t c c a t a t a c g a g a a a a a a c t t t a a a t a g c c a c 5’
```

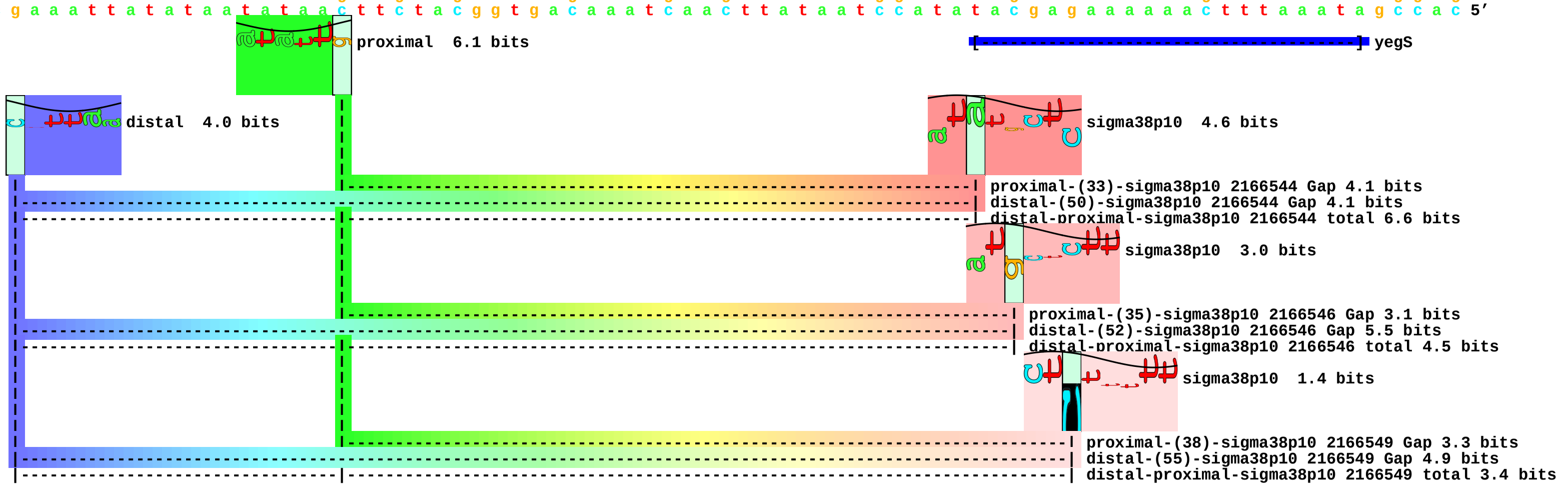

piece 103, NC_000913, yfiL, config: linear, direction: +, begin: 2739129, end: 2739214

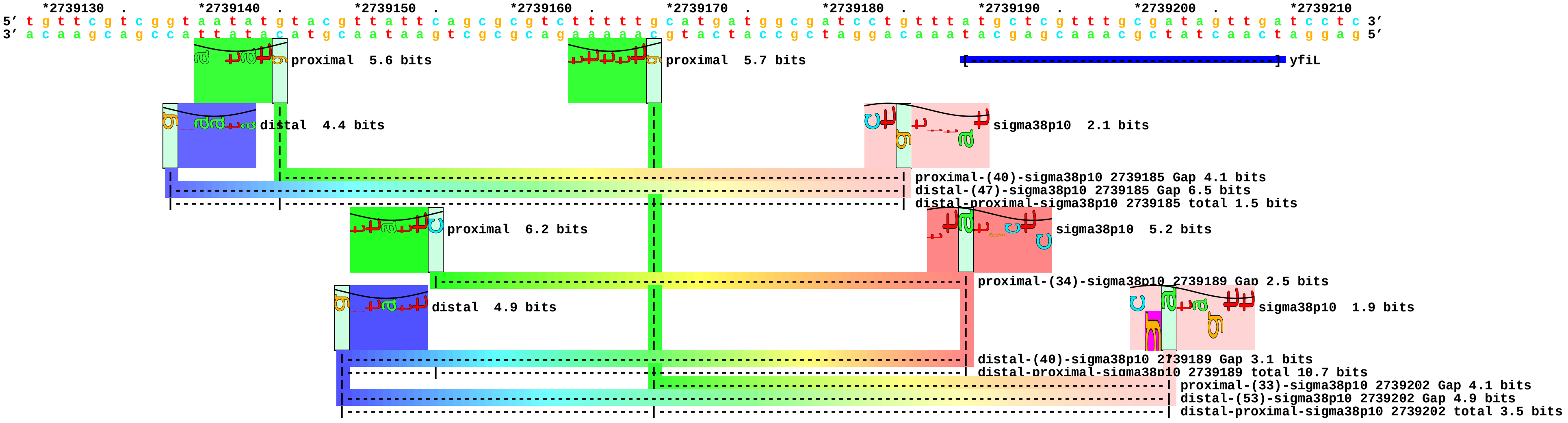

piece 104, NC_000913, ygaM, config: linear, direction: +, begin: 2798071, end: 2798156

```
         .         *2798080  .         *2798090  .         *2798100  .         *2798110  .         *2798120  .         *2798130  .         *2798140  .         *2798150  .
5’ c a a c g c t t t c a g a a a a g t c c a t a a a g c c g c g a t g a a c c t c a g t t a a g a a a a t t a t g g t c t a c a c t g a a a a t t a c a t c g a a t t c t a a 3’
3’ g t t g c g a a a g t c t t t t c a g g t a t t t c g g c g c t a c t t g g a g t c a a t t c t t t t a a t a c c a g a t g t g a c t t t t a a t g t a g c t t a a g a t t 5’
```

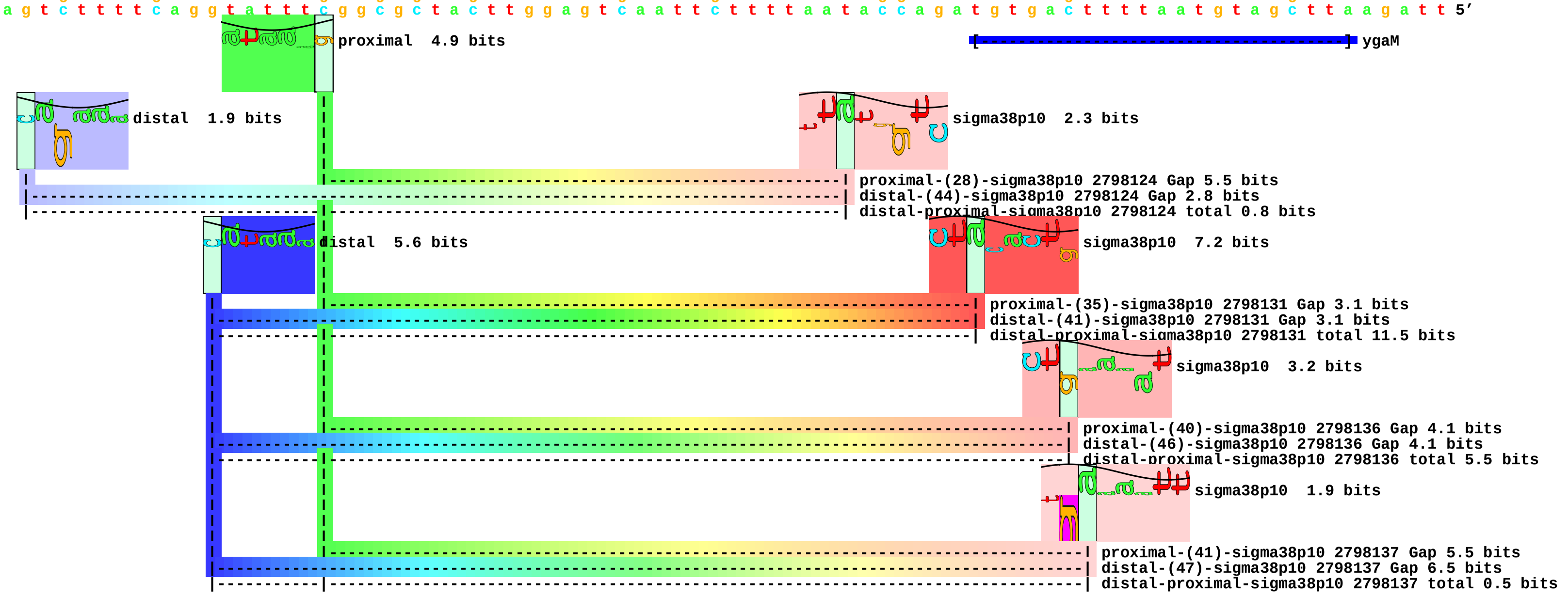

piece 105, NC_000913, ygaU, config: linear, direction: -, begin: 2795178, end: 2795093

```
         .         *2795170  .         *2795160  .         *2795150  .         *2795140  .         *2795130  .         *2795120  .         *2795110  .         *2795100  .
5' t a c c a g c c t c a g c a g c g t a a a t g a g a g t a a a a g c g t a a g c t g a a a c t g g c a g g c t c c g c t a a a a t t a c t a c g c t t a a g a g a t a a a a 3'
3' a t g g t c g g a g t c g t c g c a t t t a c t c t c a t t t t c g c a t t c g a c t t t g a c c g t c c g a g g c g a t t t t a a t g a t g c g a a t t c t c t a t t t t 5'
```

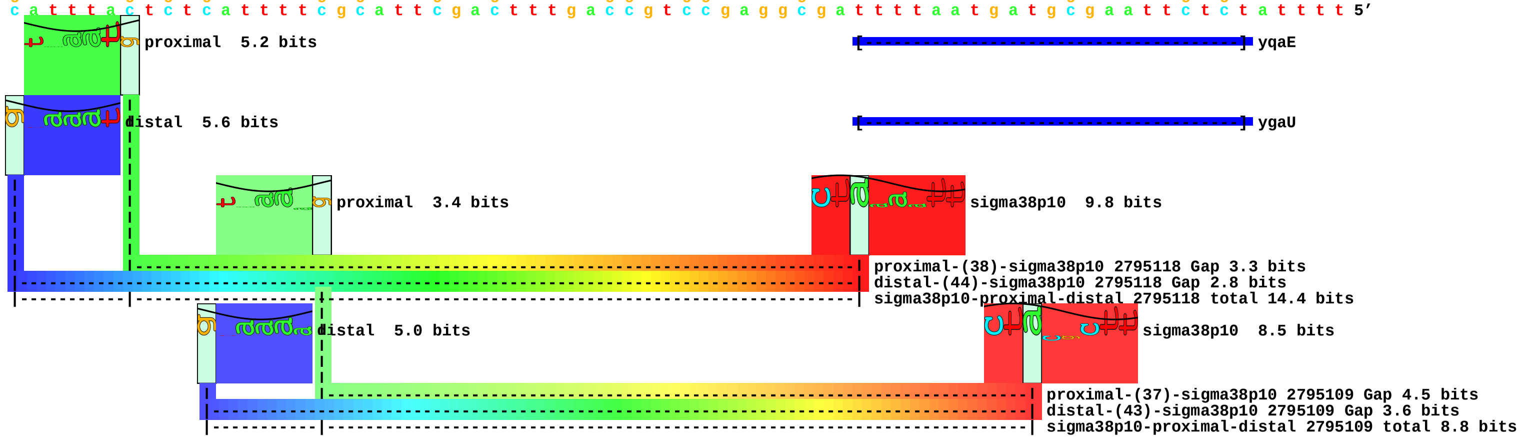

piece 106, NC_000913, ygbA, config: linear, direction: -, begin: 2854934, end: 2854849

```
           *2854930  .        *2854920  .        *2854910  .        *2854900  .        *2854890  .        *2854880  .        *2854870  .        *2854860  .        *2854850
5' c a a g t a c g c a a a a t t g t a t c g g t t g g c t a a c c c t t c t t c t t t g g a g t c a t a g t g c c c t t t a a g a t t a a g a t g t a a t a t a a a t a c a t 3'
3' g t t c a t g c g t t t t a a c a t a g c c a a c c g a t t g g g a a g a a g a a a c c t c a g t a t c a c g g g a a a t t c t a a t t c t a c a t t a t a t t t a t g t a 5'
```

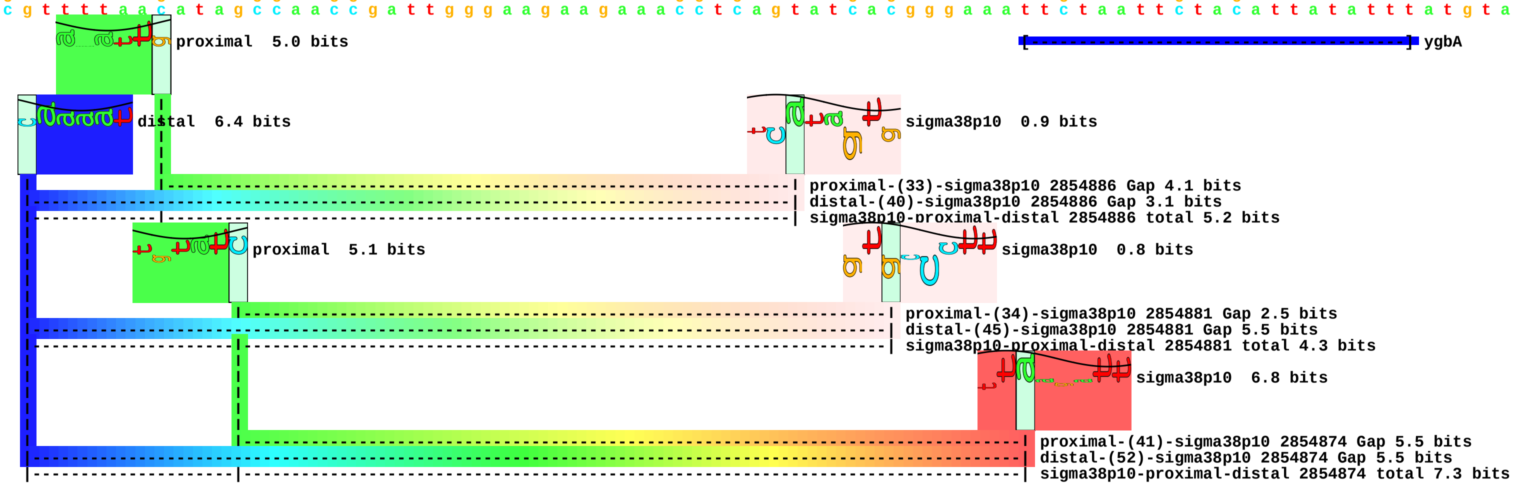

piece 107, NC_000913, ygfS, config: linear, direction: -, begin: 3027228, end: 3027143

```
         .         *3027220  .         *3027210  .         *3027200  .         *3027190  .         *3027180  .         *3027170  .         *3027160  .         *3027150  .
5' a g c g g a a t t a a a c t c g a t a a a t g g g g c c t g a t t c a a a c c g g t g a c g t c g g g t a t t t a c c t a c c c a g a c g c a t c t g a a a a a a g t c t t 3'
3' t c g c c t t a a t t t g a g c t a t t t a c c c c g g a c t a a g t t t g g c c a c t g c a g c c c a t a a a t g g a t g g g t c t g c g t a g a c t t t t t t c a g a a 5'
```

proximal 5.2 bits

ygfS

distal 5.9 bits

sigma38p10 1.0 bits

proximal-(38)-sigma38p10 3027168 Gap 3.3 bits
distal-(56)-sigma38p10 3027168 Gap 3.8 bits
sigma38p10-proximal-distal 3027168 total 5.1 bits

piece 108, NC_000913, yggEp1, config: linear, direction: -, begin: 3066220, end: 3066135

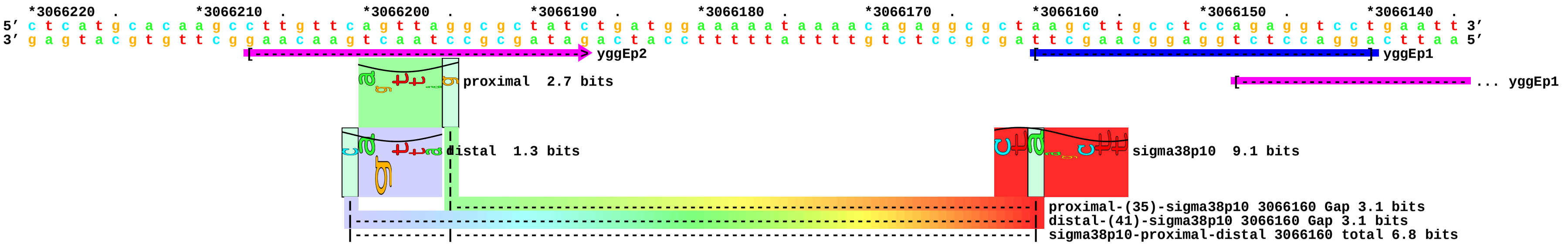

piece 109, NC_000913, yggEp2, config: linear, direction: -, begin: 3066520, end: 3066435

```
   *3066520  .         *3066510  .         *3066500  .         *3066490  .         *3066480  .         *3066470  .         *3066460  .         *3066450  .         *3066440  .
5' t g a a g c a a g g c a g a t g g a a a a t t a t c g c c a c c a t g t t g g c a g t g a c c t g g c t g a a t c c g c a t g t t t a c c t g g a t a c t t t t g t t g t a 3'
3' a c t t c g t t c c g t c t a c c t t t t a a t a g c g g t g g t a c a a c c g t c a c t g g a c c g a c t t a g g c g t a c a a a t g g a c c t a t g a a a a c a a c a t 5'
```

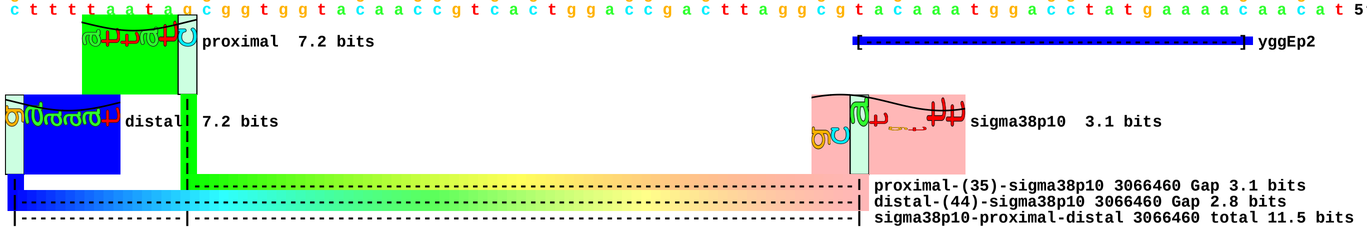

piece 110, NC_000913, yghA, config: linear, direction: +, begin: 3147510, end: 3147595

```
   *3147510  .         *3147520  .         *3147530  .         *3147540  .         *3147550  .         *3147560  .         *3147570  .         *3147580  .         *3147590  .
5' g t t a a a a c g g g c a a c a g t a t a t c g c a a t t g a g t a a a c a g c a c g c a a c a g c a a a t c a a c a t a t a c g t t a a a c c t t t g t c g c a t a a t t 3'
3' c a a t t t t g c c c g t t g t c a t a t a g c g t t a a c t c a t t t g t c g t g c g t t g t c g t t t a g t t g t a t a t g c a a t t t g g a a a c a g c g t a t t a a 5'
```

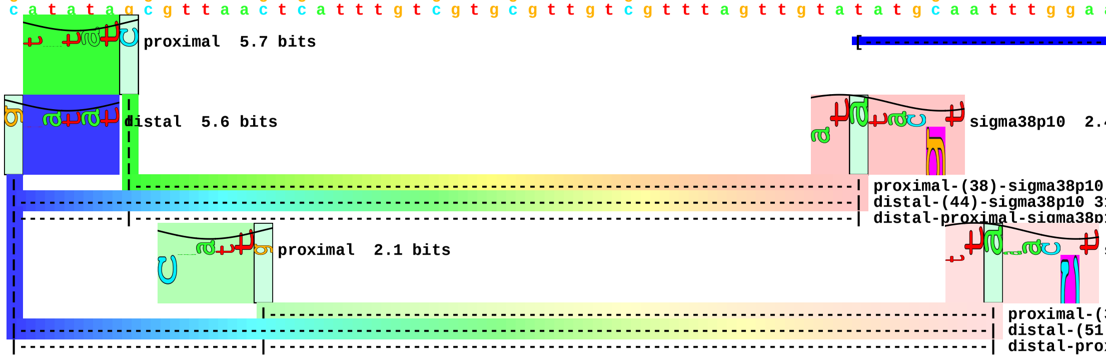

piece 111, NC_000913, ygiW, config: linear, direction: -, begin: 3167841, end: 3167756

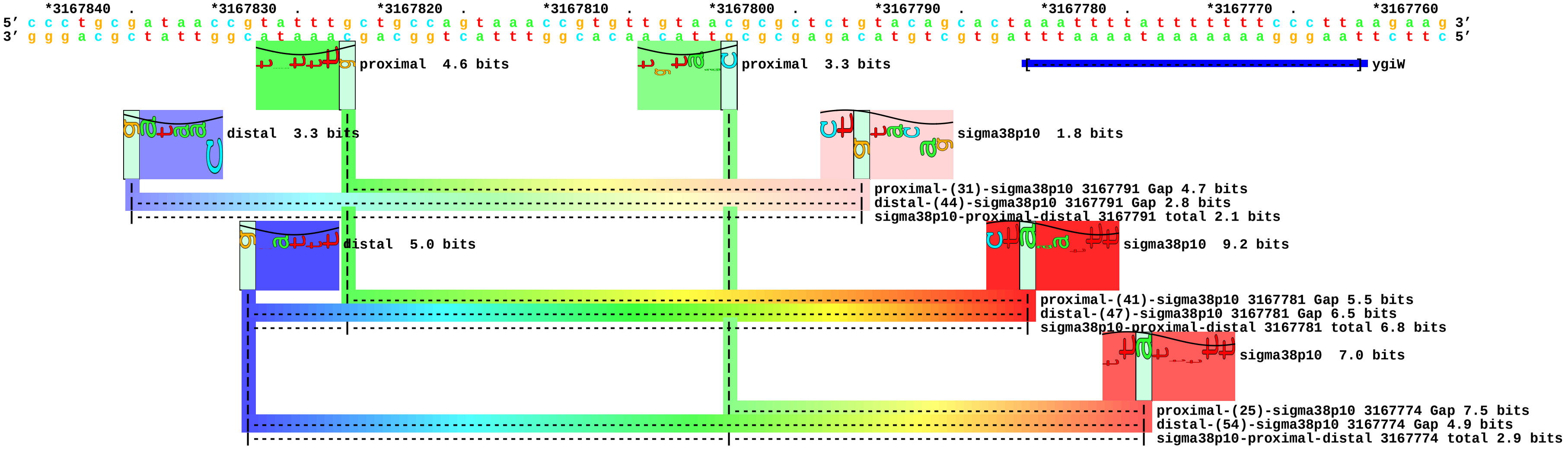

piece 112, NC_000913, yhfG, config: linear, direction: -, begin: 3489740, end: 3489655

```
   *3489740  .         *3489730  .         *3489720  .         *3489710  .         *3489700  .         *3489690  .         *3489680  .         *3489670  .         *3489660  .
5' c t c g c g c g g g c a a t c t t g c c c c g c g c t t c t g c t c t c c c g g c g t a a c c c g g a t t t g c c g c t t a t a c t t g t g g c a a a t g g a c a c g t t c a 3'
3' g a g c g c g c c c g t t a g a a c g g g g c g c g a a g a c g a g a g g g c c g c a t t g g g c c t a a a c g g c g a a t a t g a a c a c c g t t t a c c t g t g c a a g t 5'
```

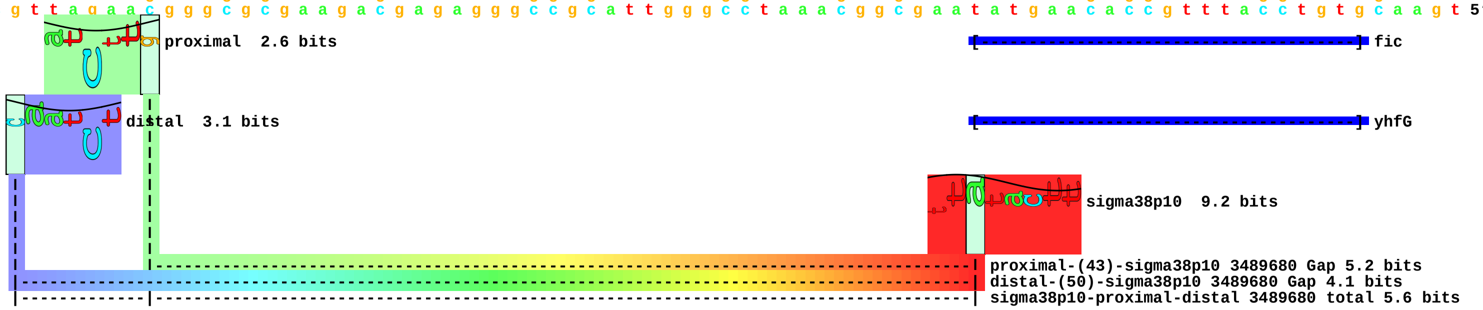

piece 113, NC_000913, yhhA, config: linear, direction: +, begin: 3584635, end: 3584720

```
   .         *3584640  .         *3584650  .         *3584660  .         *3584670  .         *3584680  .         *3584690  .         *3584700  .         *3584710  .         *3584720
5' t t c c c g c c c t c c t t g a g a a t a t c c a c c c c c a c c t g a g t g g c a g t g g c g t c c a c a g a c g c t a c c a t t c c c t g t t t c g c g c g t a c c g g 3'
3' a a g g g c g g g a g g a a c t c t t a t a g g t g g g g g t g g a c t c a c c g t c a c c g c a g g t g t c t g c g a t g g t a a g g g a c a a a g c g c g c a t g g c c 5'
```

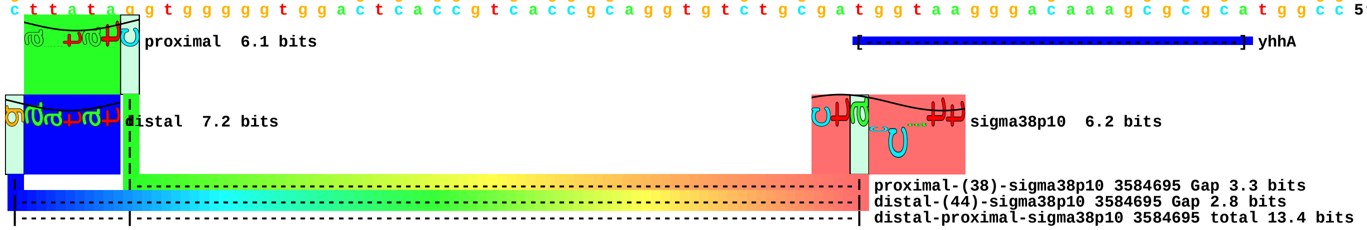

piece 114, NC_000913, yhjG, config: linear, direction: -, begin: 3676776, end: 3676691

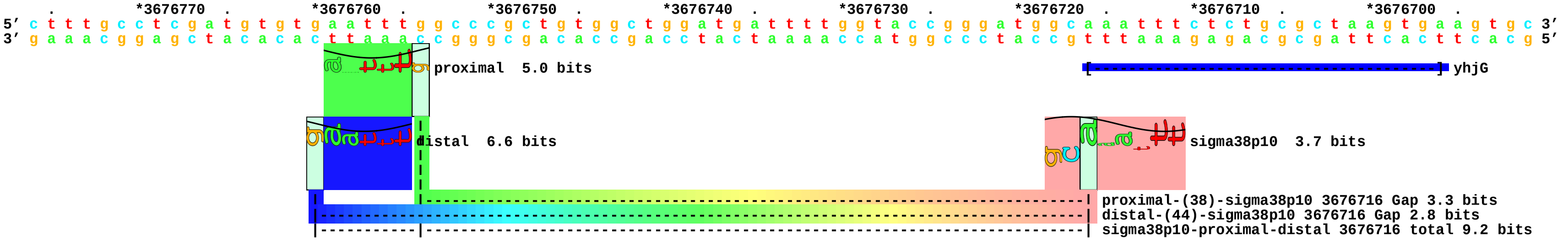

piece 115, NC_000913, yjbJ, config: linear, direction: +, begin: 4257130, end: 4257215

```
   *4257130  .         *4257140  .         *4257150  .         *4257160  .         *4257170  .         *4257180  .         *4257190  .         *4257200  .         *4257210  .
5' t t t g c c g c a a c g t g a c g g t t a a a a a t t c t g a a t a a a t a a t c c t a a g c c a a a t t g c t g a c t a c a c t t a a t c t c a c g t t c a g a a g a a a 3'
3' a a a c g g c g t t g c a c t g c c a a t t t t t a a g a c t t a t t t a t t a g g a t t c g g t t t a a c g a c t g a t g t g a a t t a g a g t g c a a g t c t t c t t t 5'
```

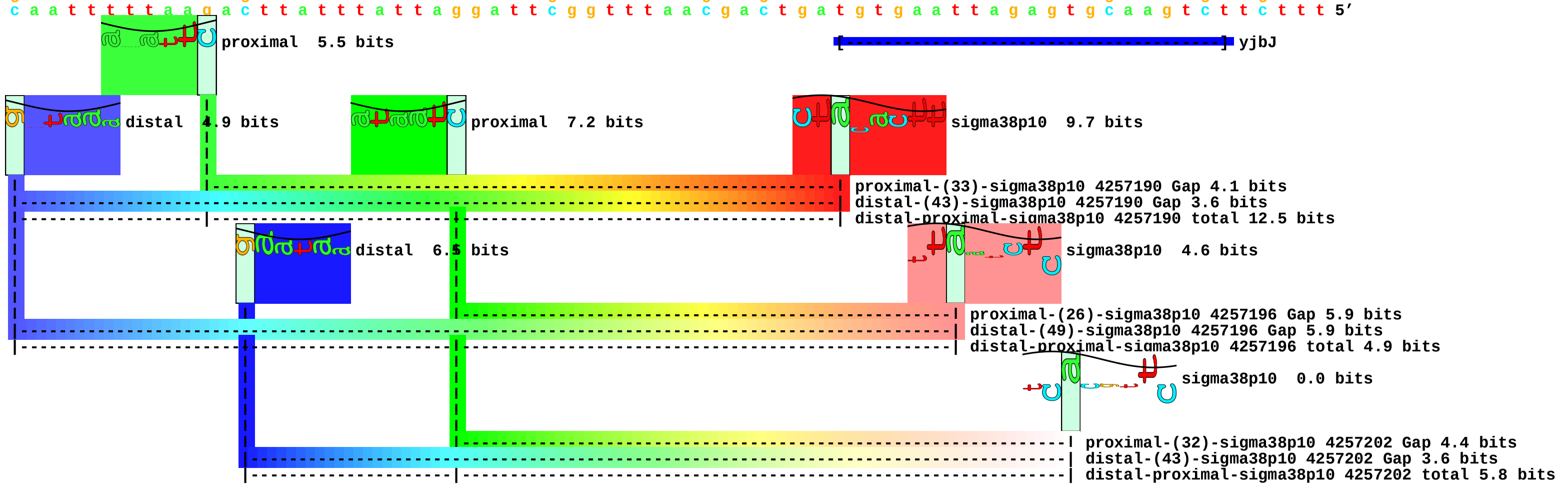

piece 116, NC_000913, nsrR, config: linear, direction: +, begin: 4404114, end: 4404199

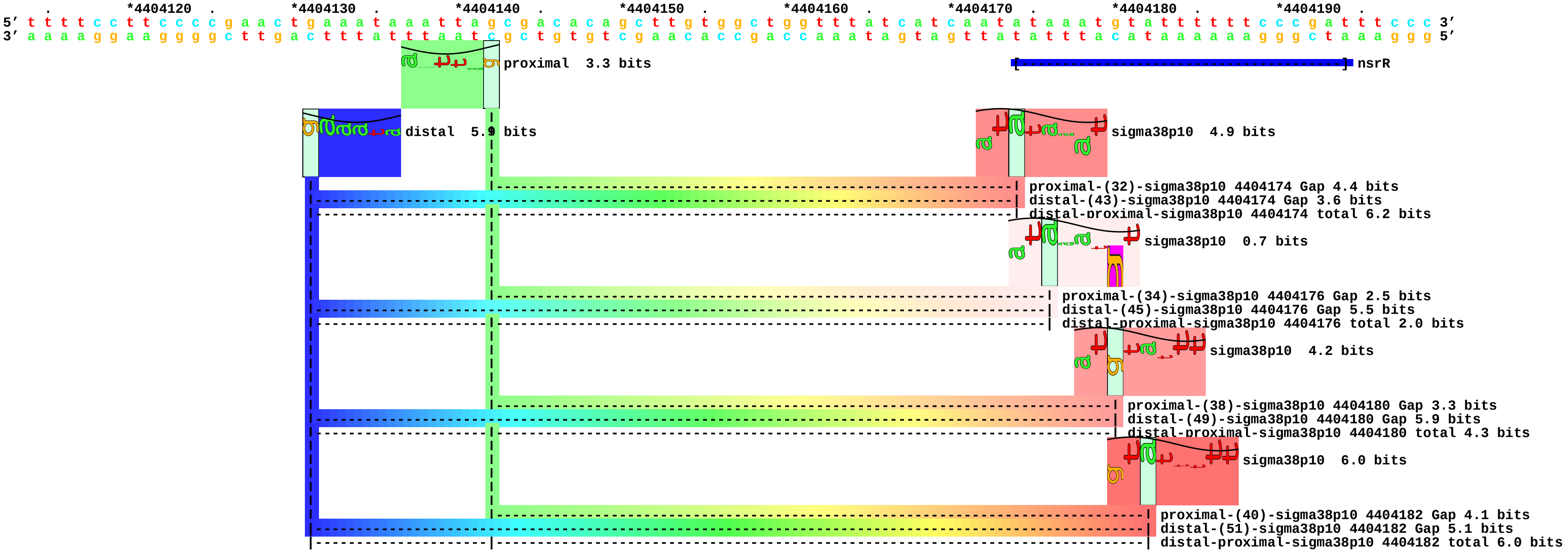

piece 117, NC_000913, yjgR, config: linear, direction: -, begin: 4488177, end: 4488092

```
          .         *4488170  .         *4488160  .         *4488150  .         *4488140  .         *4488130  .         *4488120  .         *4488110  .         *4488100  .
5' g c a a c a c g c t t t a g g t t a t t t t c t c t t c a t t a c a c g t a t t c t c t t t g c c g g a c g t t g c c t a t a c t c c a g g c a g g a a a c t g g a g g a t 3'
3' c g t t g t g c g a a a t c c a a t a a a a g a g a a g t a a t g t g c a t a a g a g a a a c g g c c t g c a a c g g a t a t g a g g t c c g t c c t t t g a c c t c c t a 5'
```

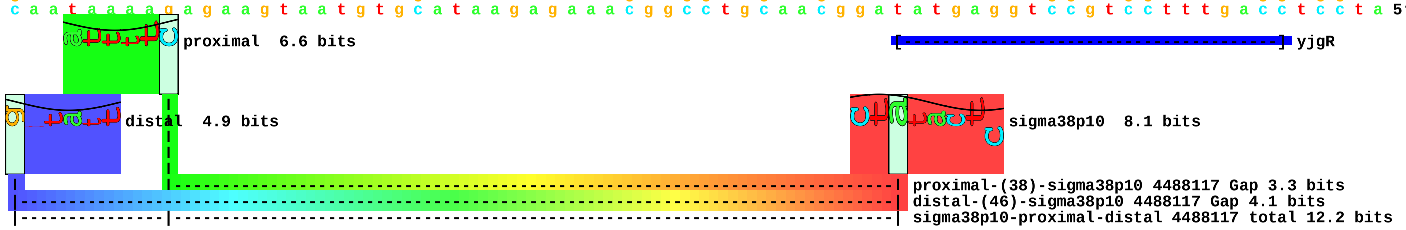

piece 118, NC_000913, nanM, config: linear, direction: -, begin: 4536876, end: 4536791

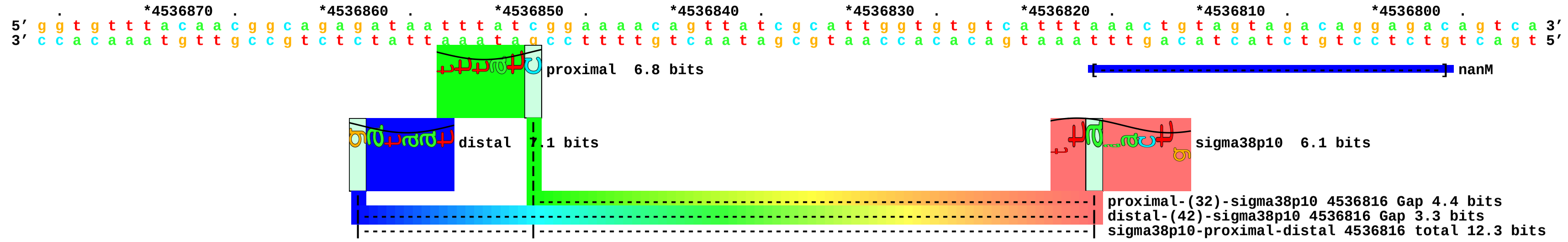

piece 119, NC_000913, ymgA, config: linear, direction: +, begin: 1215105, end: 1215190

```
   .         *1215110  .         *1215120  .         *1215130  .         *1215140  .         *1215150  .         *1215160  .         *1215170  .         *1215180  .         *1215190
5' g a a a t t g t g a c t g a g a t t t t g a a a g a t g g a c g t a a t c t g a g t c g a a a g t c g c t t t g t g c c a a a c t t c t t t g c c g a c t g g a a c a c g c 3'
3' c t t t a a c a c t g a c t c t a a a a c t t t c t a c c t g c a t t a g a c t c a g c t t t c a g c g a a a c a c g g t t t g a a g a a a c g g c t g a c c t t g t g c g 5'
```

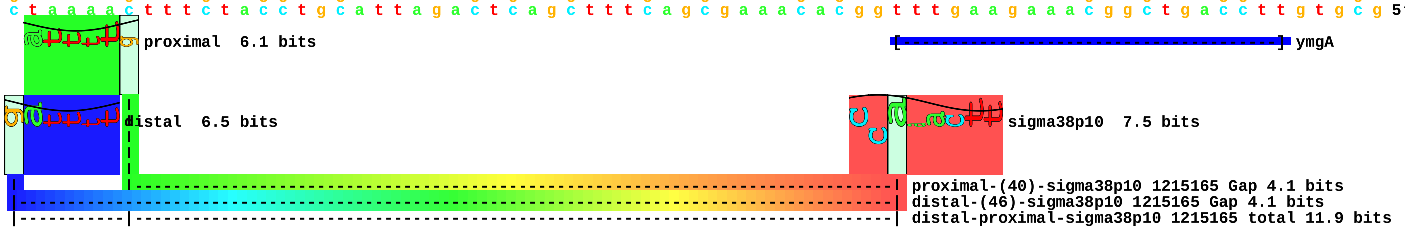

piece 120, NC_000913, ynhG, config: linear, direction: -, begin: 1756903, end: 1756818

```
         *1756900  .         *1756890  .         *1756880  .         *1756870  .         *1756860  .         *1756850  .         *1756840  .         *1756830  .         *1756820
5' a g c t a a a c t a a a a g g a c a g c t t t c a t c c g g t t g g c t c a c a t t g c c g g a t g g t c a a t g t t t a c a c t g a c c t g c a c a g g a g t g t t t t g 3'
3' t c g a t t t g a t t t t c c t g t c g a a a g t a g g c c a a c c g a g t g t a a c g g c c t a c c a g t t a c a a a t g t g a c t g g a c g t g t c c t c a c a a a a c 5'
```

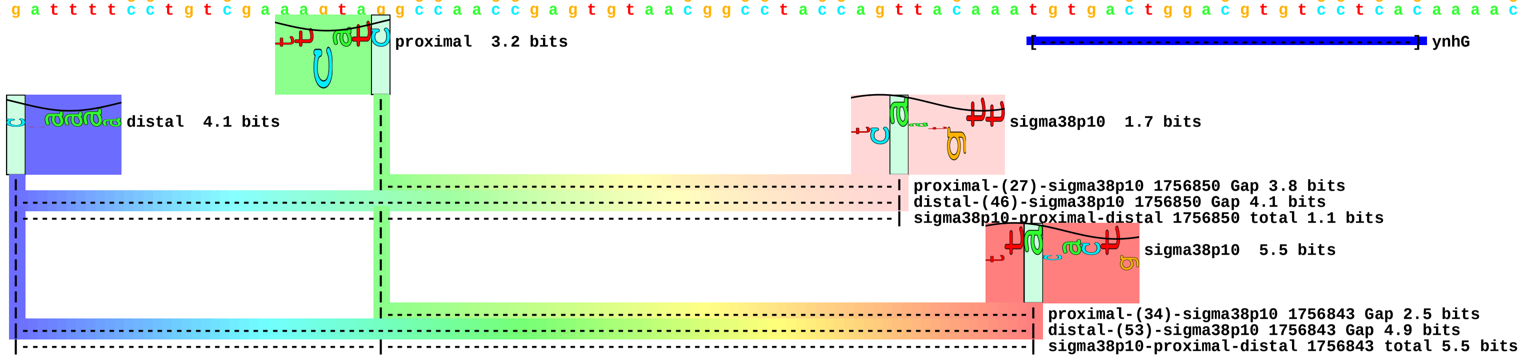

piece 121, NC_000913, ynjF, config: linear, direction: -, begin: 1839626, end: 1839541

```
   .         *1839620  .         *1839610  .         *1839600  .         *1839590  .         *1839580  .         *1839570  .         *1839560  .         *1839550  .
5' t t a c c a c c g g c a a a c t c c c a t a a t c c c g c c t g a t c g c t c t g g g c g g g g c g t t g c g c g a g t a a a a t t t t g c c a t c a c g t t c a a t g a t 3'
3' a a t g g t g g c c g t t t g a g g g t a t t a g g g c g g a c t a g c g a g a c c c g c c c c g c a a c g c g c t c a t t t t a a a a c g g t a g t g c a a g t t a c t a 5'
```

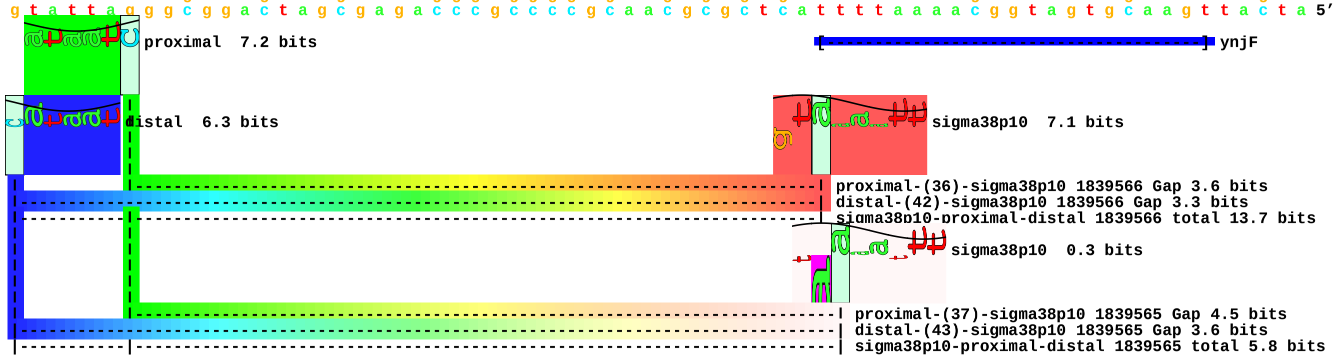

piece 122, NC_000913, yodC, config: linear, direction: -, begin: 2026518, end: 2026433

```
           .         *2026510  .         *2026500  .         *2026490  .         *2026480  .         *2026470  .         *2026460  .         *2026450  .         *2026440  .
5' g c t g g g a g t g a a a g c c g t a g a g a a a a t g c g c g g t c a g g c a c a t t a a t a c t c g c a c t g g c t a t t a t g c a a a a t t c g c c a t c t t c a a c 3'
3' c g a c c c t c a c t t t c g g c a t c t c t t t t a c g c g c c a g t c c g t g t a a t t a t g a g c g t g a c c g a t a a t a c g t t t t a a g c g g t a g a a g t t g 5'
```

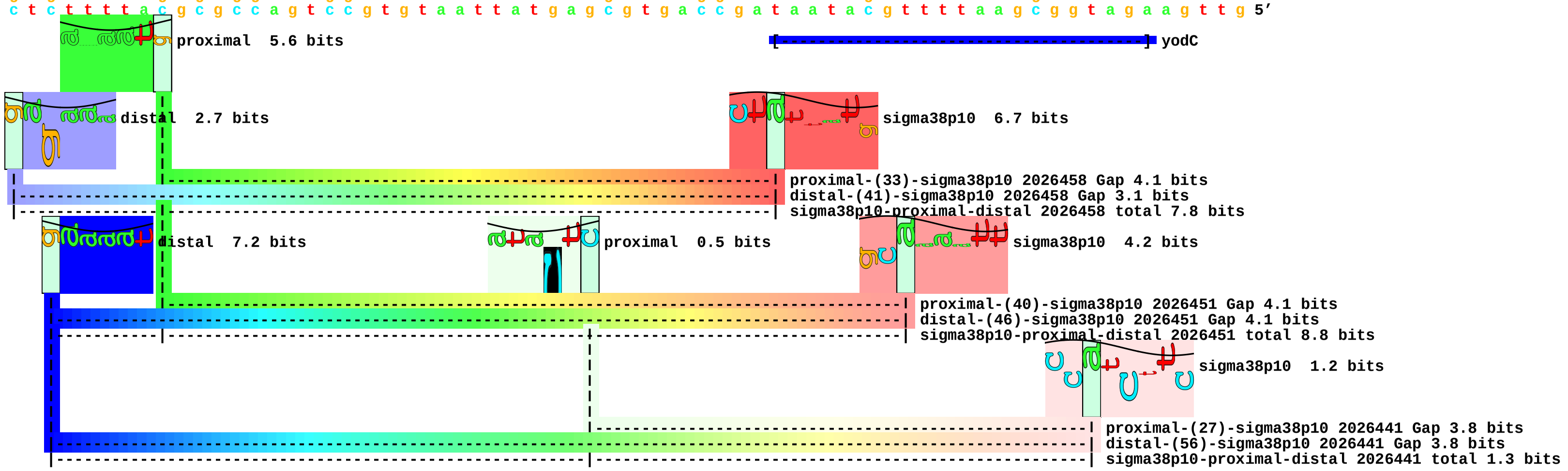

piece 123, NC_000913, yodD, config: linear, direction: +, begin: 2022863, end: 2022948

```
   .         *2022870  .         *2022880  .         *2022890  .         *2022900  .         *2022910  .         *2022920  .         *2022930  .         *2022940  .
5' g c t t t g t g c c a g t g t a g a a c a a t t t c g t t t t t c t g g c a g c g c c a g g c g c g c g c g a g t g c t g a t t t t c t c g a c g g t c t a t a c t t a a g 3'
3' c g a a a c a c g g t c a c a t c t t g t t a a a g c a a a a a g a c c g t c g c g g t c c g c g c g c g c t c a c g a c t a a a a g a g c t g c c a g a t a t g a a t t c 5'
```

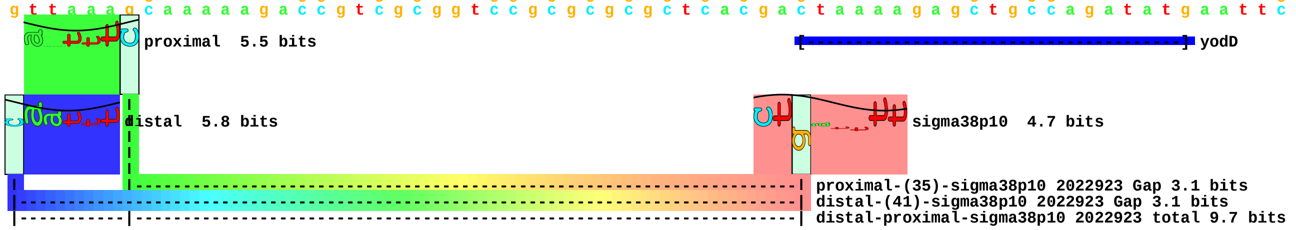

piece 124, NC_000913, yqjC, config: linear, direction: +, begin: 3246839, end: 3246924

```
    *3246840  .         *3246850  .         *3246860  .         *3246870  .         *3246880  .         *3246890  .         *3246900  .         *3246910  .         *3246920
5' g g a t a a c t t c c t g t t a t c c g a a a c a t a t c a c t c a c t t t g g t g a t t t c a c c g t a a c t g t c t a t g a t t a a t g a g g c g g t g g t t a a c a c 3'
3' c c t a t t g a a g g a c a a t a g g c t t t g t a t a g t g a g t g a a a c c a c t a a a g t g g c a t t g a c a g a t a c t a a t t a c t c c g c c a c c a a t t g t g 5'
```

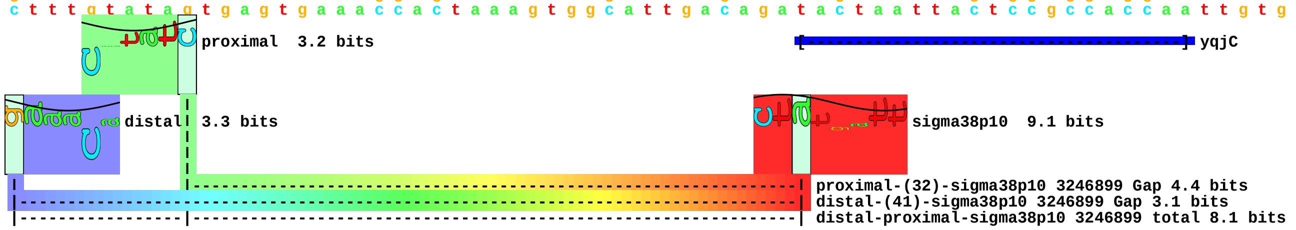

piece 125, NC_000913, yqjD, config: linear, direction: +, begin: 3247187, end: 3247272

```
         *3247190  .         *3247200  .         *3247210  .         *3247220  .         *3247230  .         *3247240  .         *3247250  .         *3247260  .         *3247270
5' t g c g t g c c g a t c a t c a g a a g a a a a t c g c a a a g c a g a a a g a t g a g g t g g c g g a a c g c c a g c a a g a t t t a g c c g a g g c g a a g c a a a a a 3'
3' a c g c a c g g c t a g t a g t c t t c t t t t a g c g t t t c g t c t t t c t a c t c c a c c g c c t t g c g g t c g t t c t a a a t c g g c t c c g c t t c g t t t t t 5'
```

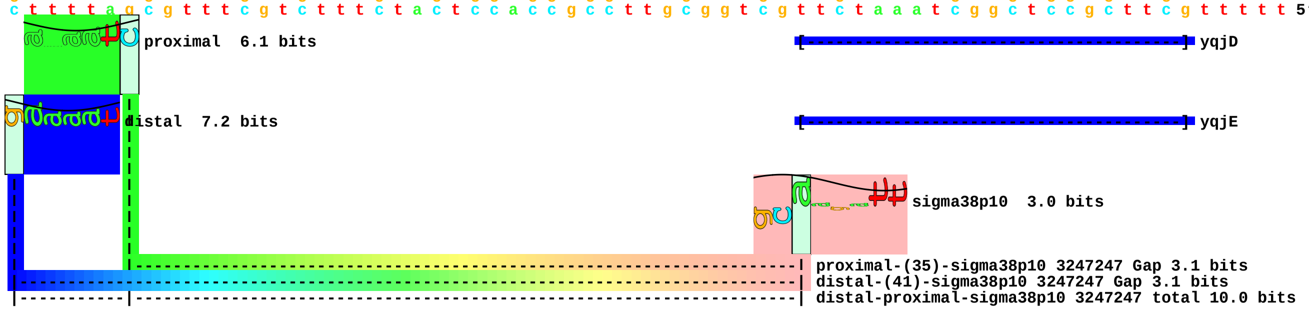

piece 126, NC_000913, yqjE, config: linear, direction: +, begin: 3247187, end: 3247272

```
        *3247190  .       *3247200  .       *3247210  .       *3247220  .       *3247230  .       *3247240  .       *3247250  .       *3247260  .       *3247270
5’ t g c g t g c c g a t c a t c a g a a g a a a a t c g c a a a g c a g a a a g a t g a g g t g g c g g a a c g c c a g c a a g a t t t a g c c g a g g c g a a g c a a a a a 3’
3’ a c g c a c g g c t a g t a g t c t t c t t t t a g c g t t t c g t c t t t c t a c t c c a c c g c c t t g c g g t c g t t c t a a a t c g g c t c c g c t t c g t t t t t 5’
```

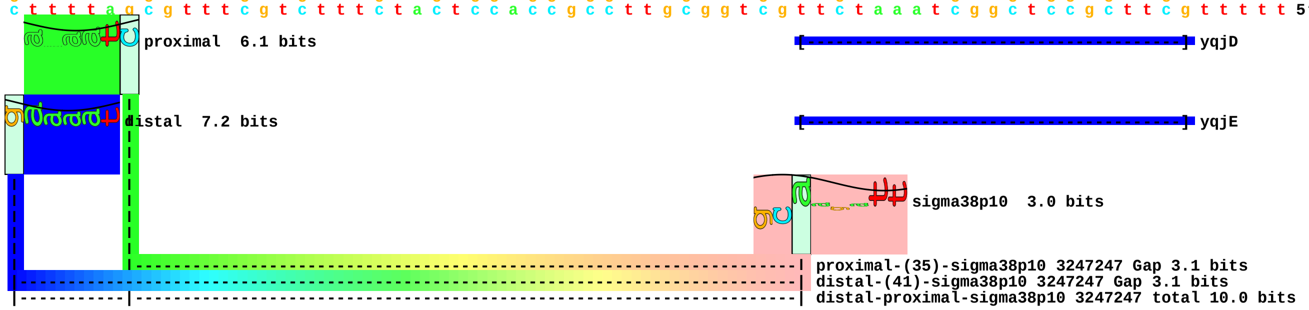

piece 127, NC_000913, ygjK, config: linear, direction: +, begin: 3226660, end: 3226745

```
   *3226660  .         *3226670  .         *3226680  .         *3226690  .         *3226700  .         *3226710  .         *3226720  .         *3226730  .         *3226740  .
5' g t g a g c g c c a t g t a t g a a g c g t t a a a g g g c g t t t a t c t c t c c a c c g c t t a c a c c t a c g c c a a a c a t c g c c c c t g a c c a c g c t g a c g a 3'
3' c a c t c g c g g t a c a t a c t t c g c a a t t t c c c g c a a a t a g a g a g g t g g c g a a t g t g g a t g c g g t t t g t a g c g g g g a c t g g t g c g a c t g c t 5'
```

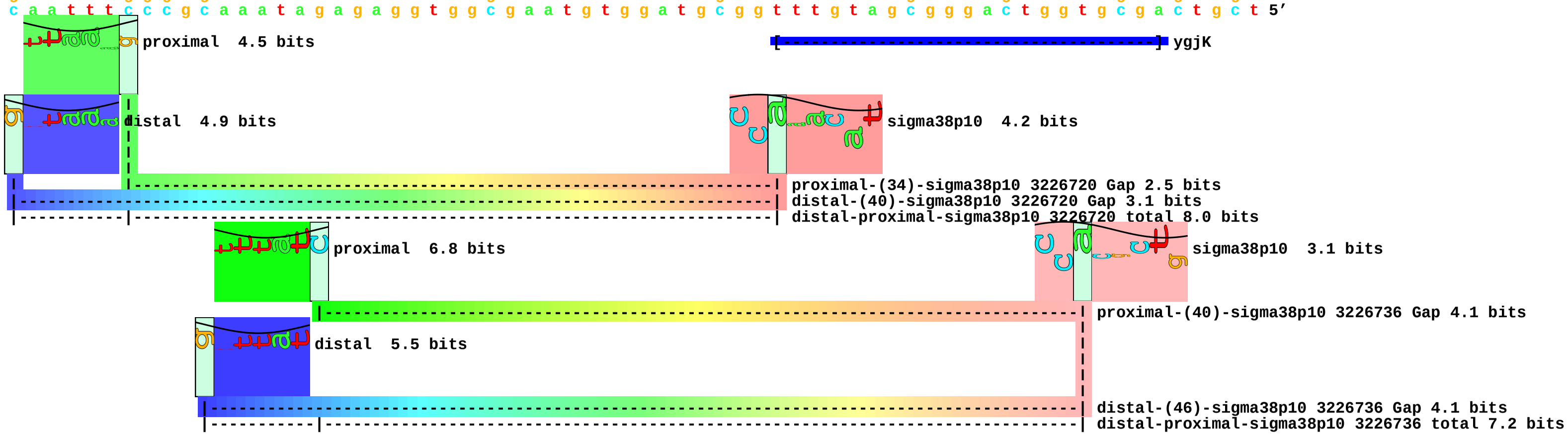

piece 128, NC_000913, adhEp2, config: linear, direction: -, begin: 1297604, end: 1297519

```
          *1297600  .         *1297590  .         *1297580  .         *1297570  .         *1297560  .         *1297550  .         *1297540  .         *1297530  .         *1297520
5' a a a g g t g t c a g c t t t g c a a a a a a t t t g a t t t g g a t c a c g t a a t c a g t a c c c a g a a g t g a g t a a t c t t g c t t a c g c c a c c t g g a a g t g 3'
3' t t t c c a c a g t c g a a a c g t t t t t t a a a c t a a a c c t a g t g c a t t a g t c a t g g g t c t t c a c t c a t t a g a a c g a a t g c g g t g g a c c t t c a c 5'
```

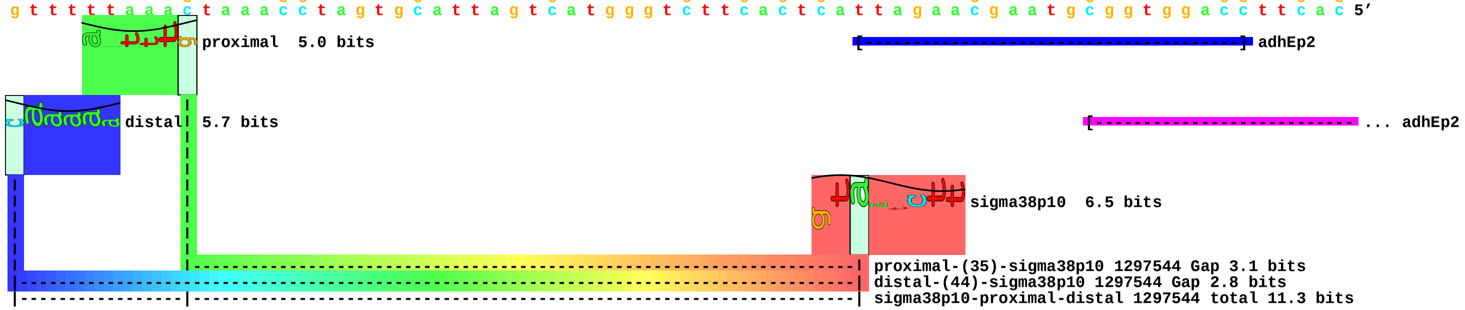

piece 129, NC_000913, adrAp, config: linear, direction: +, begin: 402843, end: 402928

```
         .         *402850    .         *402860    .         *402870    .         *402880    .         *402890    .         *402900    .         *402910    .         *402920    .
5' g g g t g a g c t a c g a a a a t g g c t c a c c c g a a a t a t c a t a c t t c t g c c t t t a g c t c c g t c t c t a t a a t t t g g g a a a a t t g t t t c t g a a t 3'
3' c c c a c t c g a t g c t t t t a c c g a g t g g g c t t t a t a g t a t g a a g a c g g a a a t c g a g g c a g a g a t a t t a a a c c c t t t t a a c a a a g a c t t a 5'
```

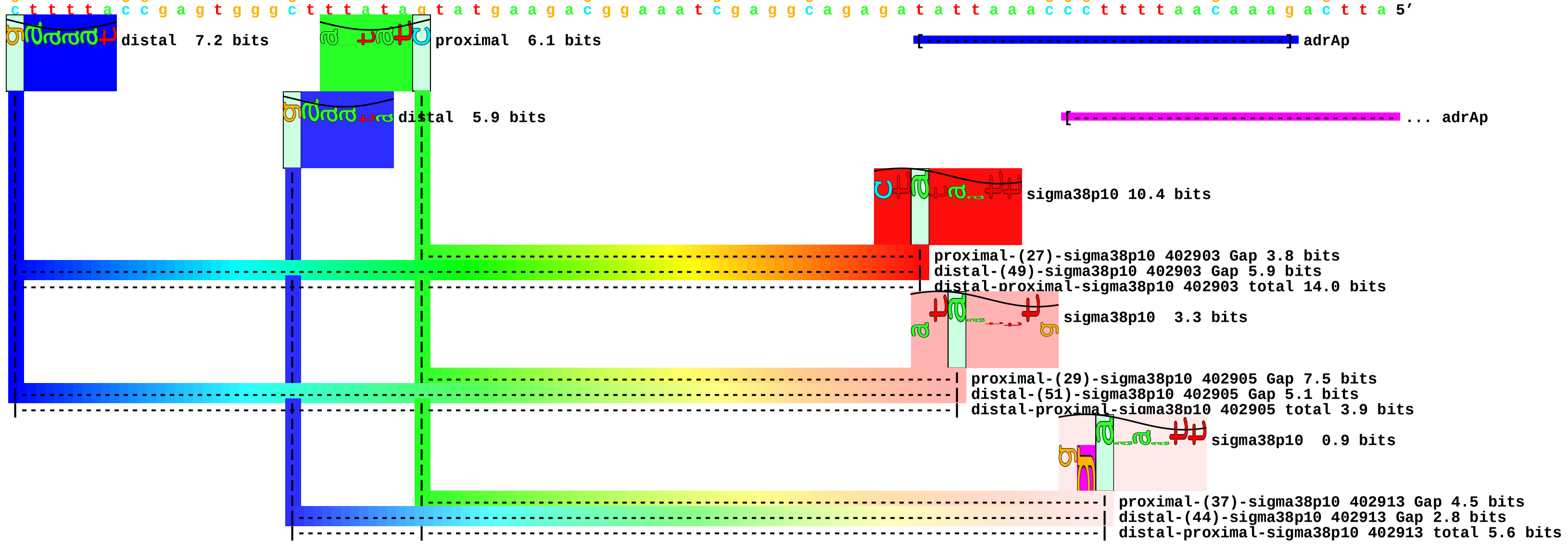

piece 130, NC_000913, aldBp, config: linear, direction: -, begin: 3754629, end: 3754544

```
          .         *3754620  .         *3754610  .         *3754600  .         *3754590  .         *3754580  .         *3754570  .         *3754560  .         *3754550  .
5' a a t t c g t g a t a g c t g t c g t a a a g c t g t t a c c g a c t g g c g a a a g a t t t c g c c a g t c a c g t c t a c c c t t g t t a t a c c t c a c a c c g c a a g 3'
3' t t a a g c a c t a t c g a c a g c a t t t c g a c a a t g g c t g a c c g c t t t c t a a a g c g g t c a g t g c a g a t g g g a a c a a t a t g g a g t g t g g c g t t c 5'
```

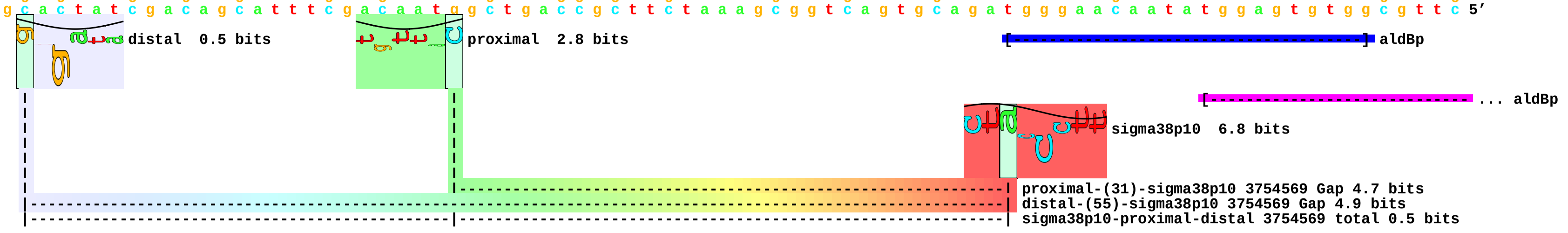

piece 131, NC_000913, ansPp1, config: linear, direction: -, begin: 1524113, end: 1524028

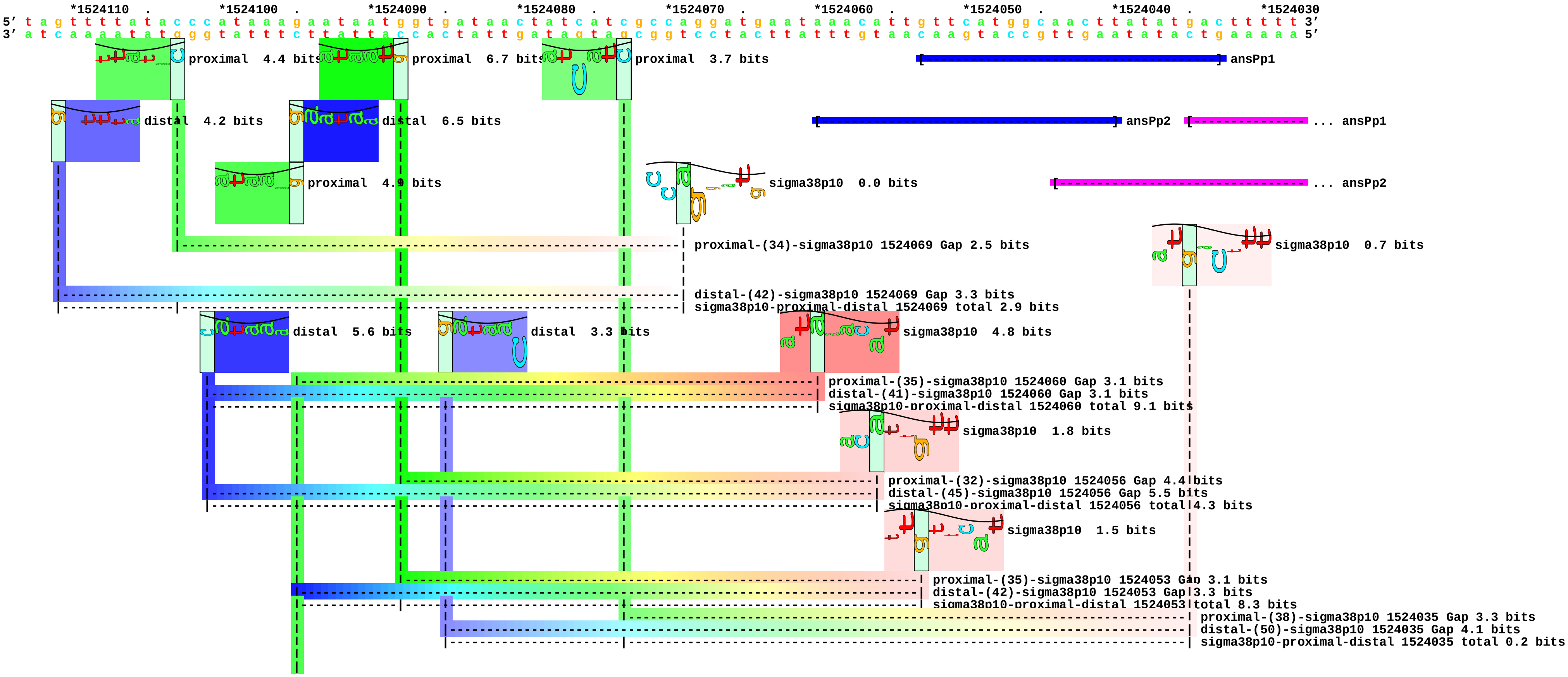

piece 132, NC_000913, ansPp2, config: linear, direction: -, begin: 1524120, end: 1524035

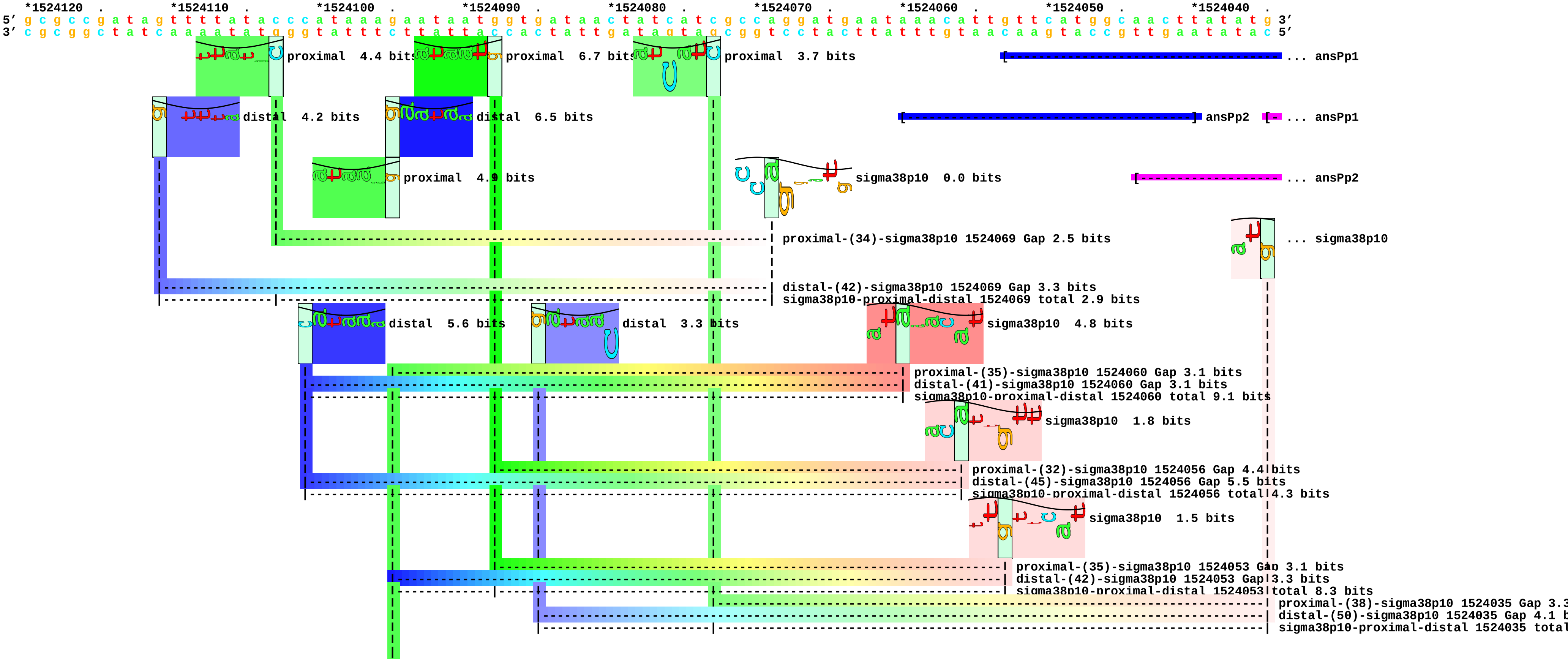

piece 133, NC_000913, astCp3, config: linear, direction: -, begin: 1830134, end: 1830049

```
      *1830130  .       *1830120  .       *1830110  .       *1830100  .       *1830090  .       *1830080  .       *1830070  .       *1830060  .       *1830050
5' t c c a g t t a g c c t c c g c c g t t t a t g c a c t t t t a t c a c t g g c t g g c a c g a a c c c t g c a a t c t a c a t t t a c a g c g c a a a c a t t a c t t a t 3'
3' a g g t c a a t c g g a g g c g g c a a a t a c g t g a a a a t a g t g a c c g a c c g t g c t t g g g a c g t t a g a t g t a a a t g t c g c g t t t g t a a t g a a t a 5'
```

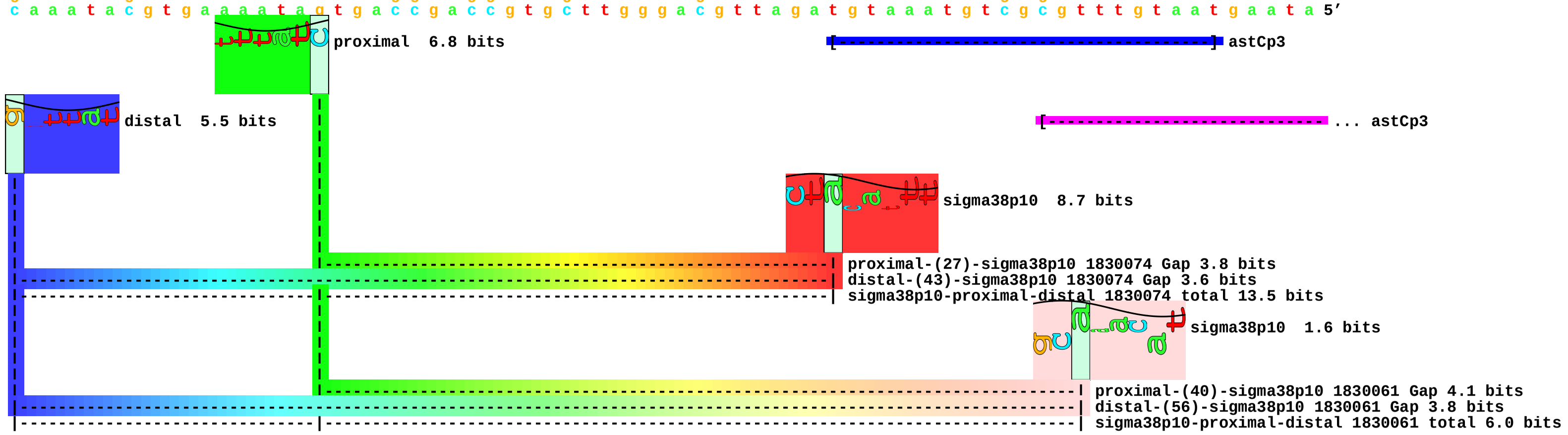

piece 134, NC_000913, cfap2, config: linear, direction: +, begin: 1739333, end: 1739418

```
         .         *1739340  .         *1739350  .         *1739360  .         *1739370  .         *1739380  .         *1739390  .         *1739400  .         *1739410  .
5' g t c g g a t t a t a g a c t c g c g g t t t t t t c t g c g a g a t t t c t c a c a a a g c c c a a a a a g c g t c t a c g c t g t t t t a a g g t t c t g a t c a c c g 3'
3' c a g c c t a a t a t c t g a g c g c c a a a a a a g a c g c t c t a a a g a g t g t t t c g g g t t t t t c g c a g a t g c g a c a a a a t t c c a a g a c t a g t g g c 5'
```

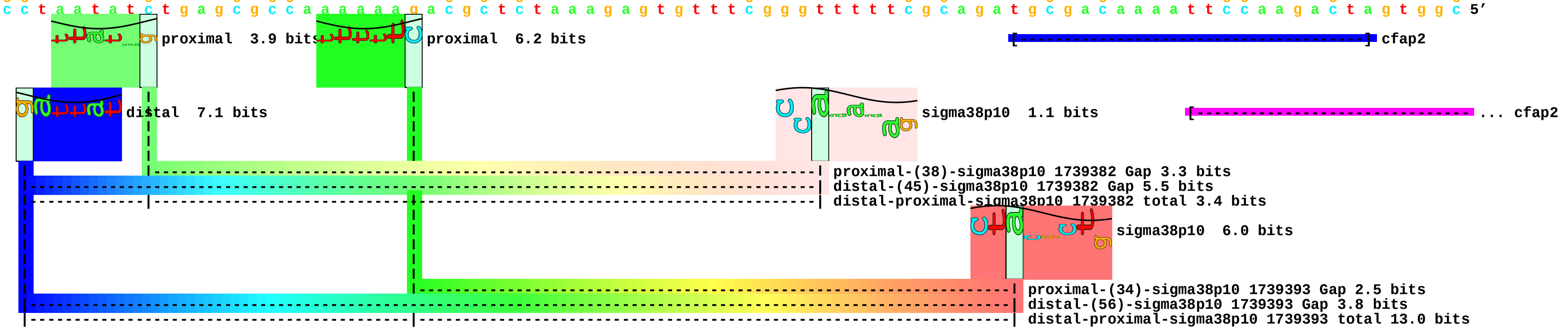

piece 135, NC_000913, csrAp3, config: linear, direction: -, begin: 2817367, end: 2817282

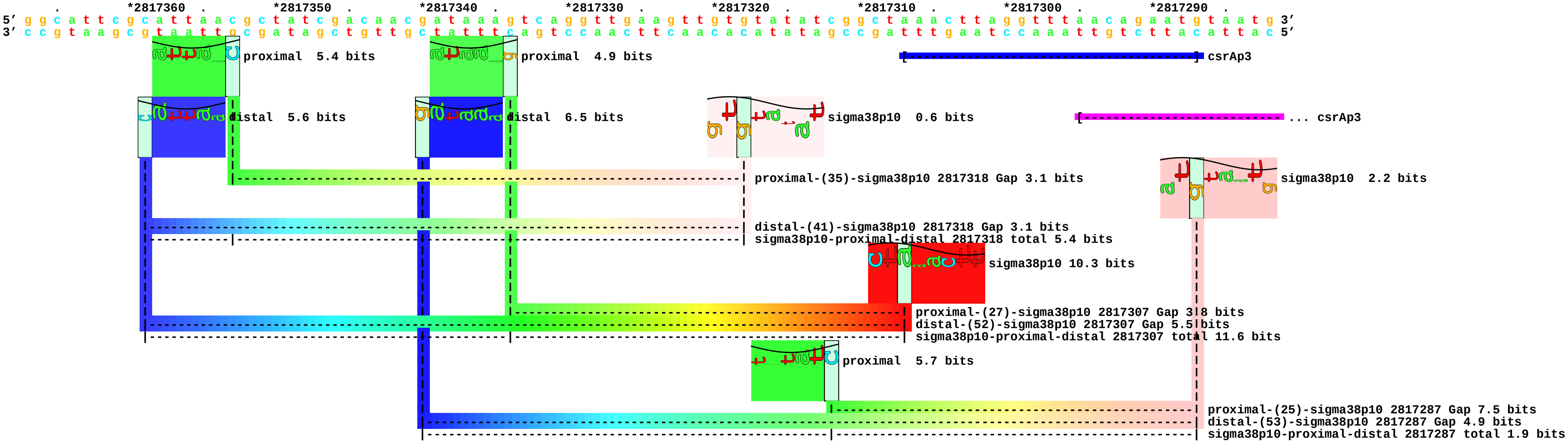

piece 136, NC_000913, ecnBp, config: linear, direction: +, begin: 4374462, end: 4374547

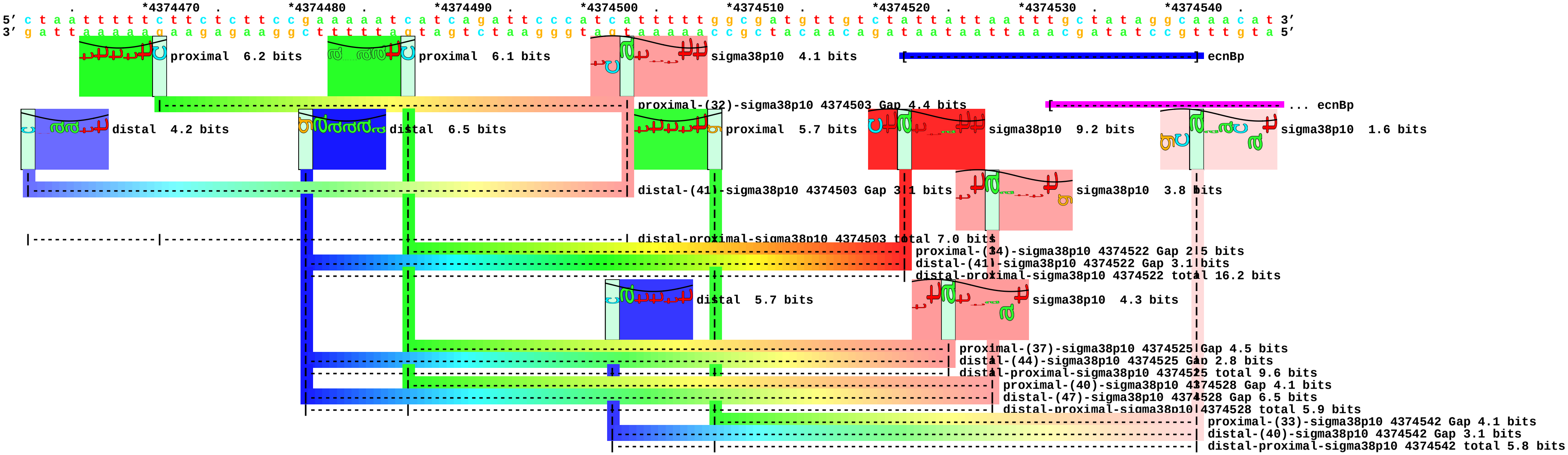

piece 137, NC_000913, evgAp1, config: linear, direction: +, begin: 2481592, end: 2481677

```
         .         *2481600  .         *2481610  .         *2481620  .         *2481630  .         *2481640  .         *2481650  .         *2481660  .         *2481670  .
5' c a t c g g t a c a a t c c c t g a t t t t a t t g t t g a c a t t t c a t t t a t g c c g a c t a t t t a t a t g g t a t a c t t g t c g a a t t a t c t t a a a g g a a 3'
3' g t a g c c a t g t t a g g g a c t a a a a t a a c a a c t g t a a a g t a a a t a c g g c t g a t a a a t a t a c c a t a t g a a c a g c t t a a t a g a a t t t c c t t 5'
```

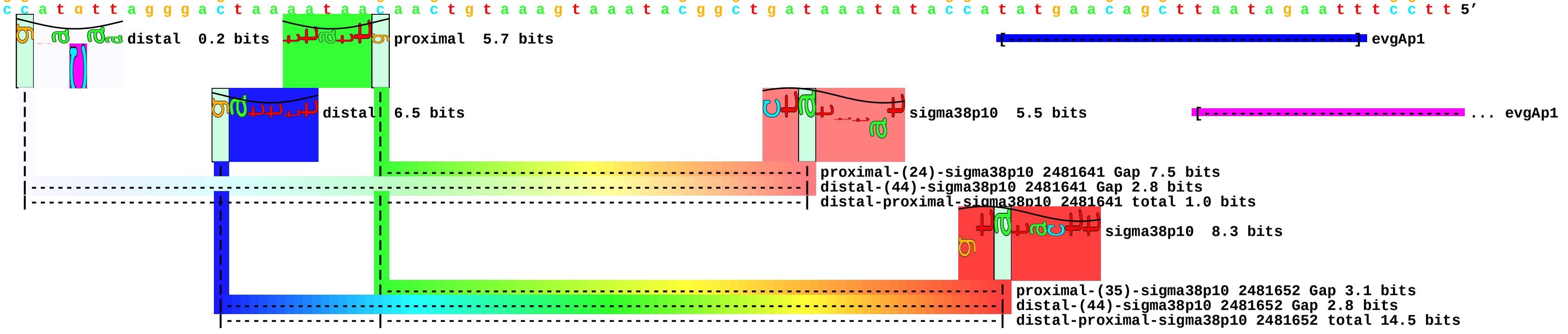

piece 138, NC_000913, fadLp, config: linear, direction: +, begin: 2459156, end: 2459241

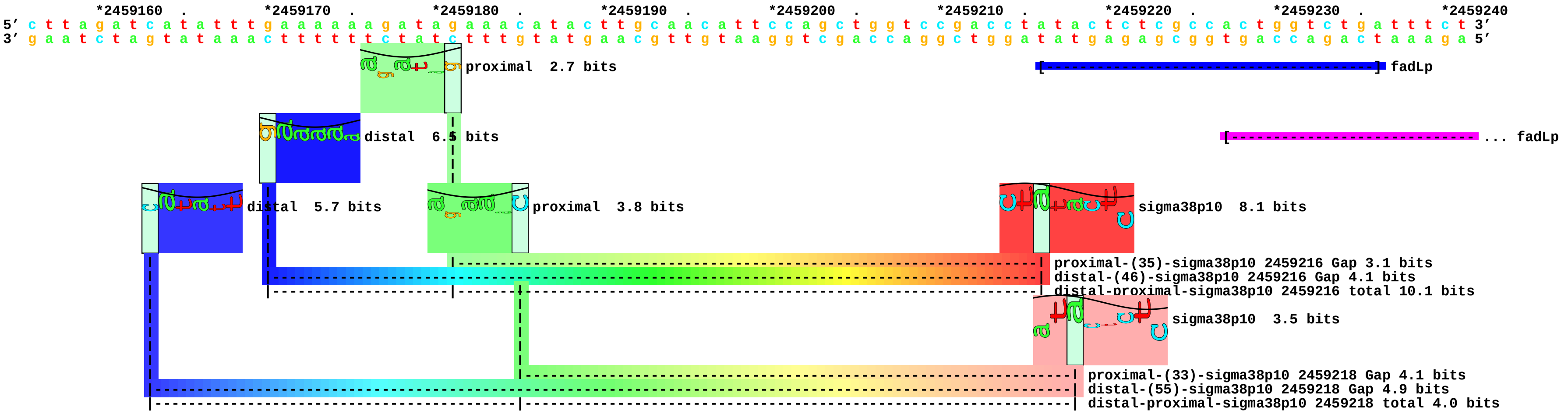

piece 139, NC_000913, ftsQp2, config: linear, direction: +, begin: 102673, end: 102758

```
        .         *102680   .         *102690   .         *102700   .         *102710   .         *102720   .         *102730   .         *102740   .         *102750   .
5' c g a a g g t t c c a g t g t g g g a a t g t c a a a a g t a g t a g c a g a a a a t g c t c t a c a a g a t g c a t t a a g a t t g g c a t t t c a g c a c g a t g a a g 3'
3' g c t t c c a a g g t c a c a c c c t t a c a g t t t t c a t c a t c g t c t t t t a c g a g a t g t t c t a c g t a a t t c t a a c c g t a a a g t c g t g c t a c t t c 5'
```

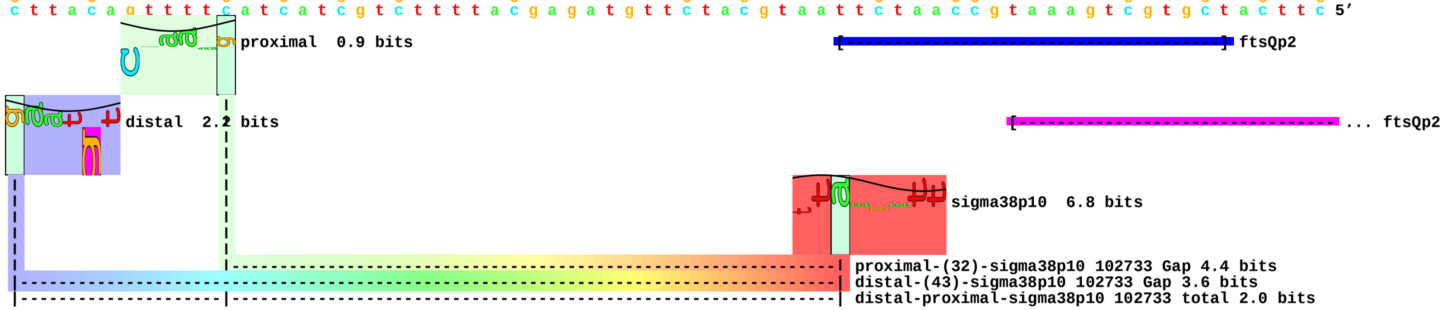

piece 140, NC_000913, gadEp, config: linear, direction: +, begin: 3656296, end: 3656381

```
          *3656300  .       *3656310  .       *3656320  .       *3656330  .       *3656340  .       *3656350  .       *3656360  .       *3656370  .       *3656380
5' t a g g c g t t t a c t a t a t t g a a c a a c g a t t c g g a c a a g g a t g t a a a t a a t g a a a a g g a t g a c a t a t t c g a a a c g a t a a c g g c t a a g g a 3'
3' a t c c g c a a a t g a t a t a a c t t g t t g c t a a g c c t g t t c c t a c a t t t a t t a c t t t t c c t a c t g t a t a a g c t t t g c t a t t g c c g a t t c c t 5'
```

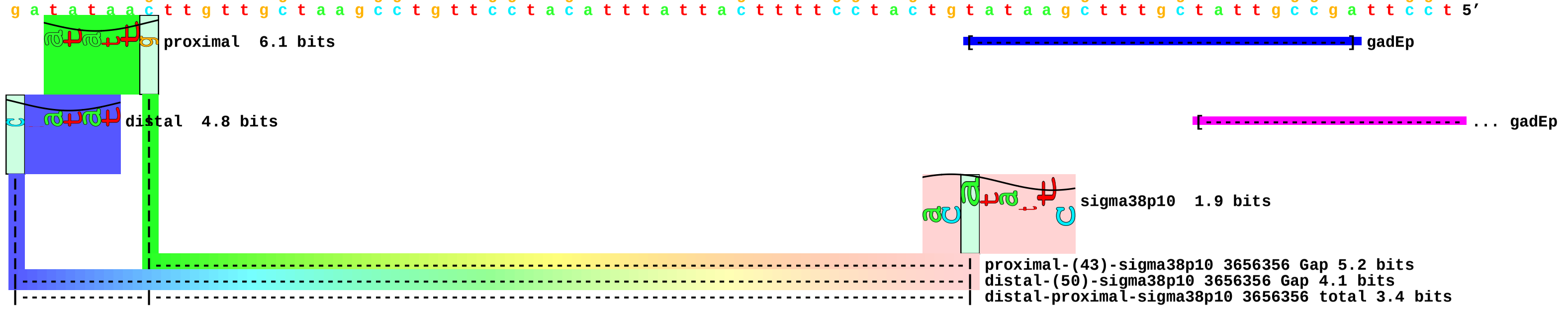

piece 141, NC_000913, gadEp1, config: linear, direction: +, begin: 3656194, end: 3656279

```
   .         *3656200  .         *3656210  .         *3656220  .         *3656230  .         *3656240  .         *3656250  .         *3656260  .         *3656270  .
5' a g g a a t c t t a c t t a g g a t c a a t a t a t g g a g t g c g t g a t g g a t a a a t c t g a a g t a t t g a t t a g t g t t a a t a g a c g t a t t a g t t c a c g 3'
3' t c c t t a g a a t g a a t c c t a g t t a t a t a c c t c a c g c a c t a c c t a t t t a g a c t t c a t a a c t a a t c a c a a t t a t c t g c a t a a t c a a g t g c 5'
```

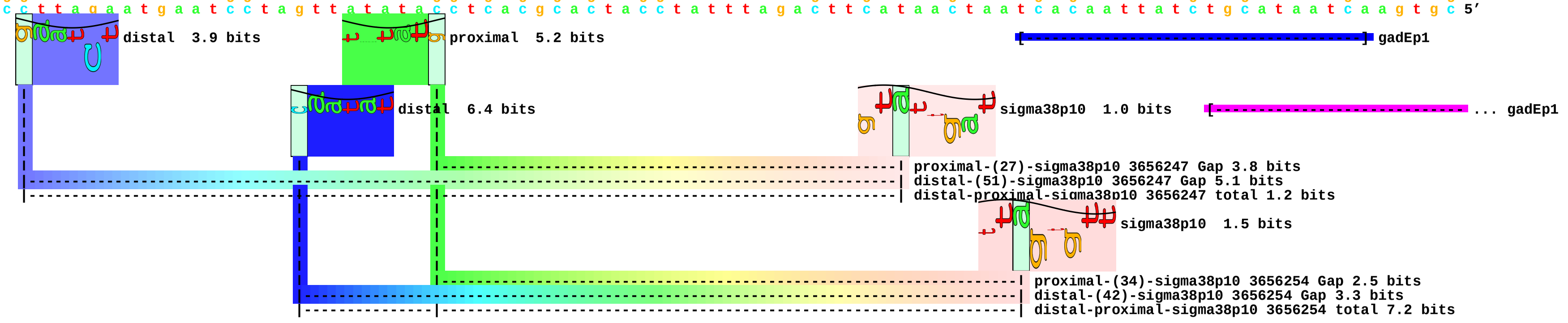

piece 142, NC_000913, gadYp, config: linear, direction: +, begin: 3662816, end: 3662901

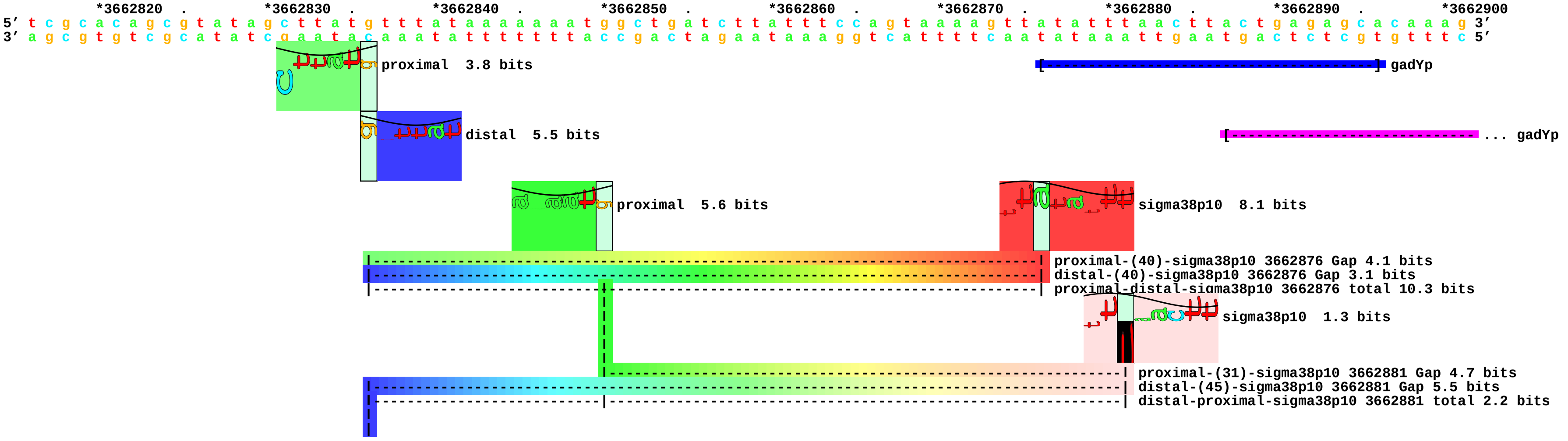

piece 143, NC_000913, glgCp3, config: linear, direction: -, begin: 3567663, end: 3567578

```
         *3567660  .         *3567650  .         *3567640  .         *3567630  .         *3567620  .         *3567610  .         *3567600  .         *3567590  .         *3567580
5’ c a t t c c c g c t t t g g t g g a g a a t c g c t g g t g g g a a g a a g g c g a c g g c a a t g t c c g t t g g c t a a a t c g a t a t g c t c a a c c t t t a a g c a 3’
3’ g t a a g g g c g a a a c c a c c t c t t a g c g a c c a c c c t t c t t c c g c t g c c g t t a c a g g c a a c c g a t t t a g c t a t a c g a g t t g g a a a t t c g t 5’
                                                                                                                                       [---------------------------------------] glgCp3
                                                                                                                                                         [---------------------------------- ... glgCp3
```

piece 144, NC_000913, glgSp1, config: linear, direction: -, begin: 3190111, end: 3190026

```
   *3190110  .       *3190100  .       *3190090  .       *3190080  .       *3190070  .       *3190060  .       *3190050  .       *3190040  .       *3190030
5' a a t a t g g t t t t t t a t t t t t c a c t a a a a a g t g t g a t c g g g g a c a a t a t a t t t a c g c a c g t t a t g t t t a a a g g c a c t a c a c t g a t t g g 3'
3' t t a t a c c a a a a a a t a a a a a g t g a t t t t t c a c a c t a g c c c c t g t t a t a t a a a t g c g t g c a a t a c a a a t t t c c g t g a t g t g a c t a a c c 5'
```

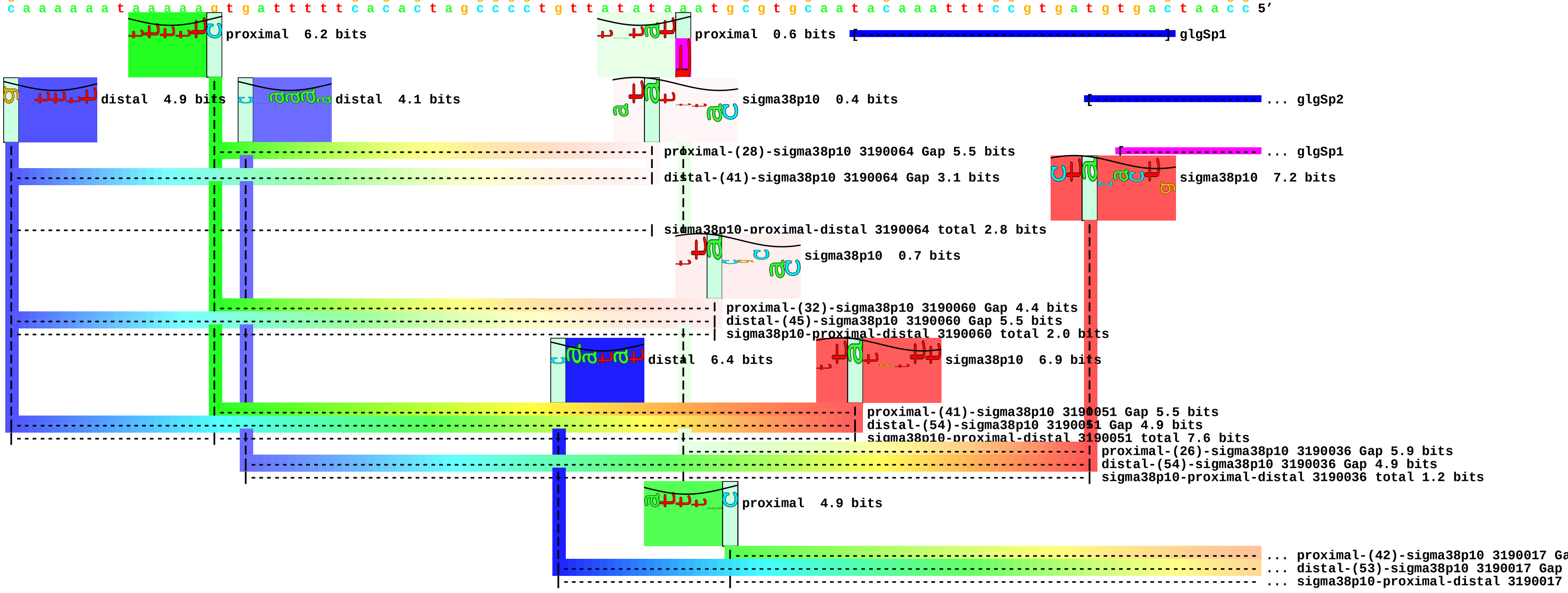

piece 145, NC_000913, glgSp2, config: linear, direction: -, begin: 3190096, end: 3190011

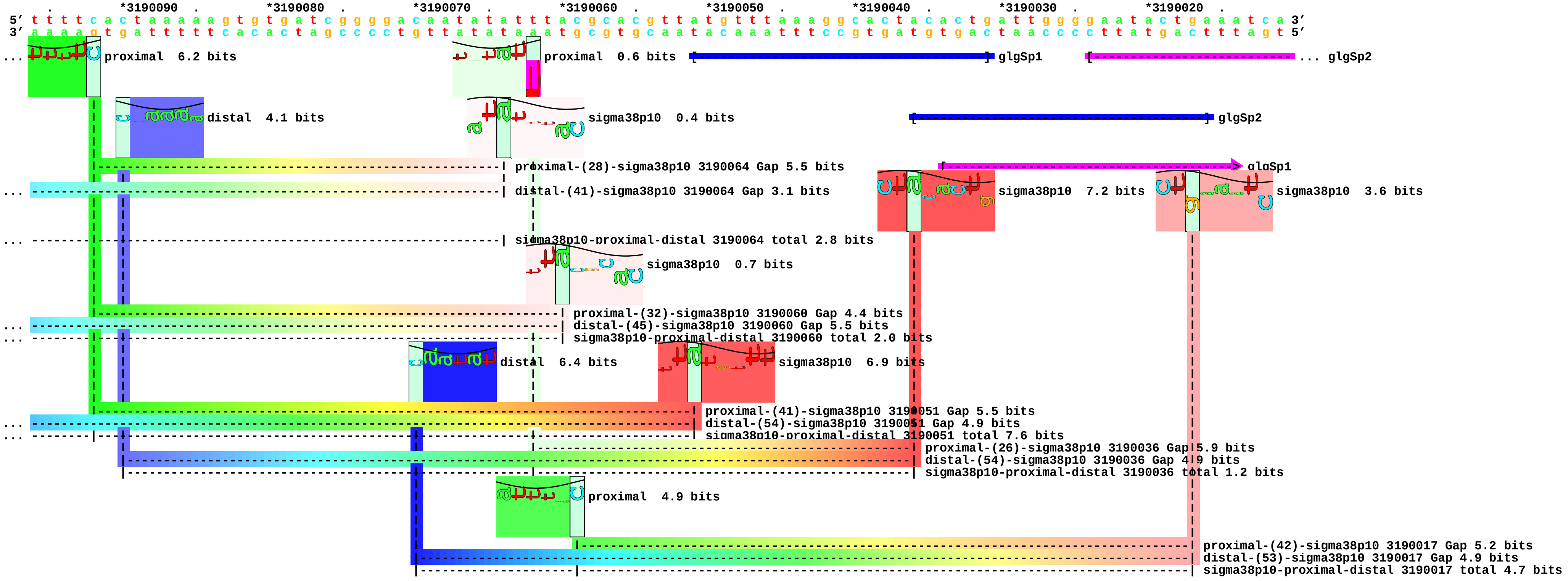

piece 146, NC_000913, hmpAp, config: linear, direction: +, begin: 2683748, end: 2683833

```
     *2683750  .        *2683760  .        *2683770  .        *2683780  .        *2683790  .        *2683800  .        *2683810  .        *2683820  .        *2683830
5' t c a t c a c a t t c t t t c t g a a a a a c a c c a a a g a a c c a t t t a c a t t g c a g g g c t a t t t t t t a t a a g a t g c a t t t g a g a t a c a t c a a t t a 3'
3' a g t a g t g t a a g a a a g a c t t t t t g t g g t t t c t t g g t a a a t g t a a c g t c c c g a t a a a a a a t a t t c t a c g t a a a c t c t a t g t a g t t a a t 5'
```

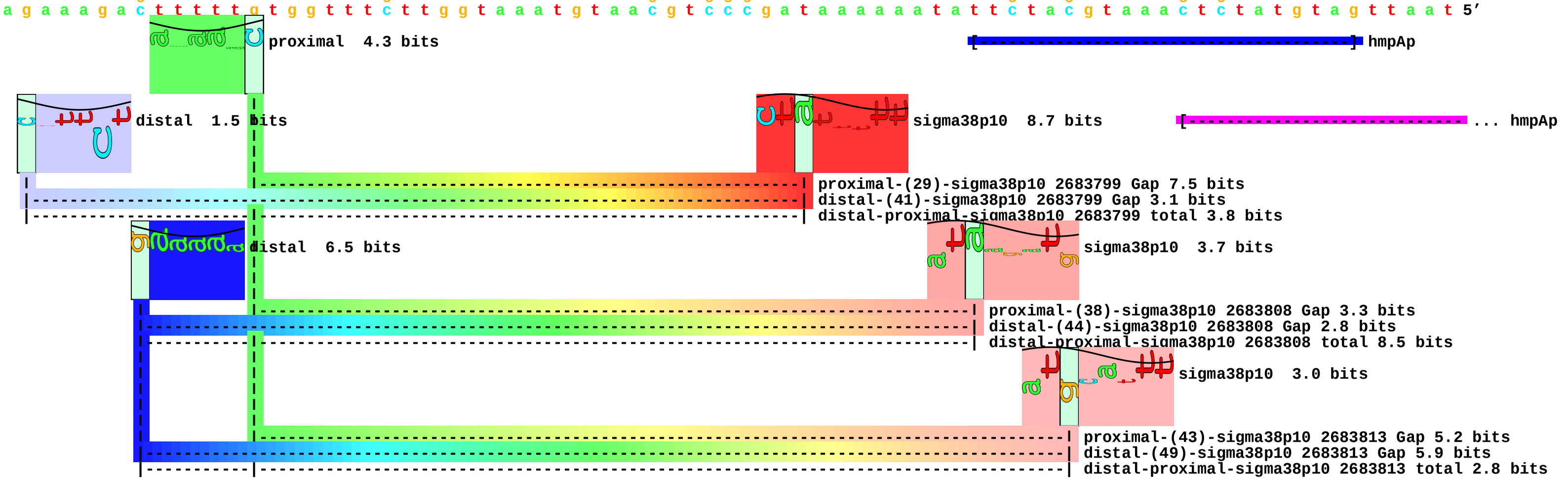

piece 147, NC_000913, ihfAp4, config: linear, direction: -, begin: 1793815, end: 1793730

```
   .         *1793810  .         *1793800  .         *1793790  .         *1793780  .         *1793770  .         *1793760  .         *1793750  .         *1793740  .         *1793730
5' t c c c g c a g c g g a t a t t t t a t c c g a a t g t a a g a a a g t t g g c g t a a a t c a g g t a g t t g g c g t a a a c t t a t t t g a c g t g t a c c g c g g t a 3'
3' a g g g c g t c g c c t a t a a a a t a g g c t t a c a t t c t t t c a a c c g c a t t t a g t c c a t c a a c c g c a t t t g a a t a a a c t g c a c a t g g c g c c a t 5'
```

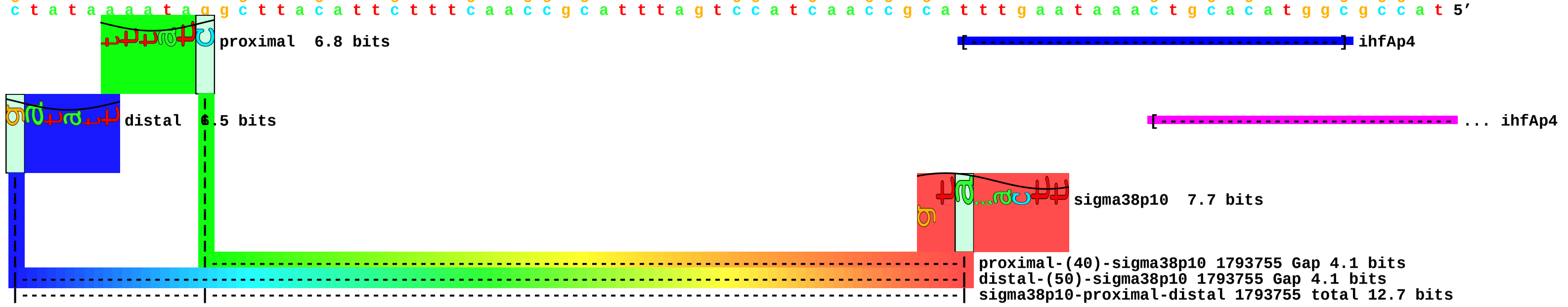

piece 148, NC_000913, ihfBp, config: linear, direction: +, begin: 962902, end: 962987

```
          .         *962910    .         *962920    .         *962930    .         *962940    .         *962950    .         *962960    .         *962970    .         *962980    .
5' c t t c g g g a t t t t t a t t c c g a a g t t t g t t g a g t t t a c t t g a c a g a t t g c a g g t t t c g t c c t g t a a t c a a g c a c t a a g g g c g g c t a c g 3'
3' g a a g c c c t a a a a a t a a g g c t t c a a a c a a c t c a a a t g a a c t g t c t a a c g t c c a a a g c a g g a c a t t a g t t c g t g a t t c c c g c c g a t g c 5'
```

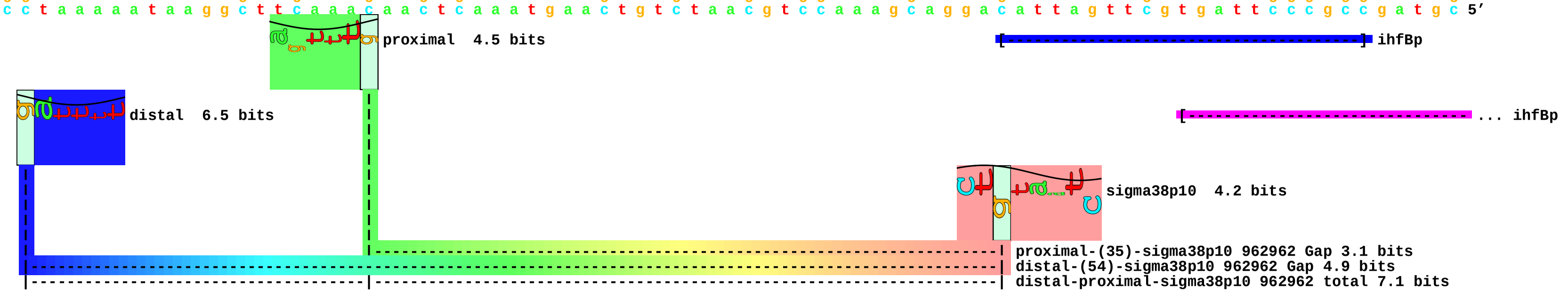

piece 149, NC_000913, luxSp1, config: linear, direction: -, begin: 2813159, end: 2813074

```
              .         *2813150   .         *2813140   .         *2813130   .         *2813120   .         *2813110   .         *2813100   .         *2813090   .         *2813080   .
5’ g a a c t g g t t g c c t g c t t c g a t a a t c c c g a t c t g a c t t t c t c t g c c c g t a t c t t a a g g t c t a t g a t t g a t a c t g g t a t t g g c g g a a c 3’
3’ c t t g a c c a a c g g a c g a a g c t a t t a g g g c t a g a c t g a a a g a g a c g g g c a t a g a a t t c c a g a t a c t a a c t a t g a c c a t a a c c g c c t t g 5’
```

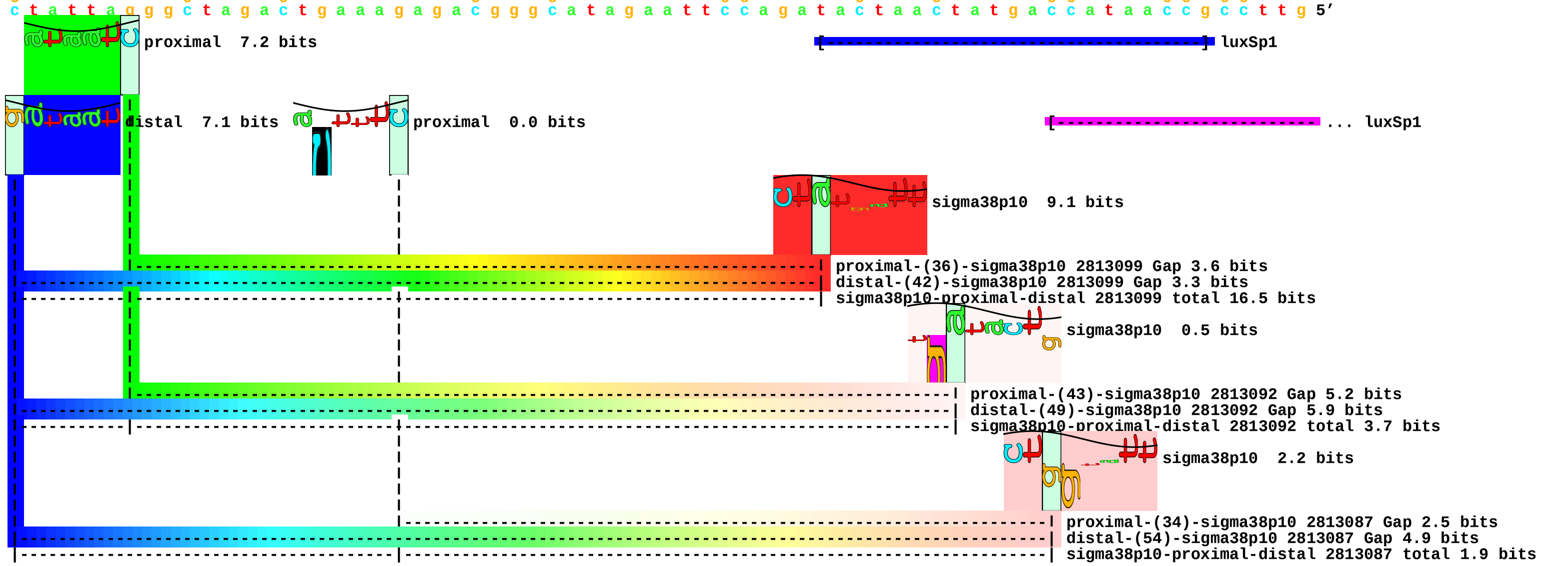

piece 150, NC_000913, nhaAp2, config: linear, direction: +, begin: 17239, end: 17324

```
    *17240    .    *17250    .    *17260    .    *17270    .    *17280    .    *17290    .    *17300    .    *17310    .    *17320
5' t g a c a a c a a a t t a a t t a t t a c t t t t c c t a a t t a a t c c c t c a g g a a t c c t c a c c t t a a g c t a t g a t t a t c t a g g c t t a g g g t c a c t c 3'
3' a c t g t t g t t t a a t t a a t a a t g a a a a g g a t t a a t t a g g g a g t c c t t a g g a g t g g a a t t c g a t a c t a a t a g a t c c g a a t c c c a g t g a g 5'
```

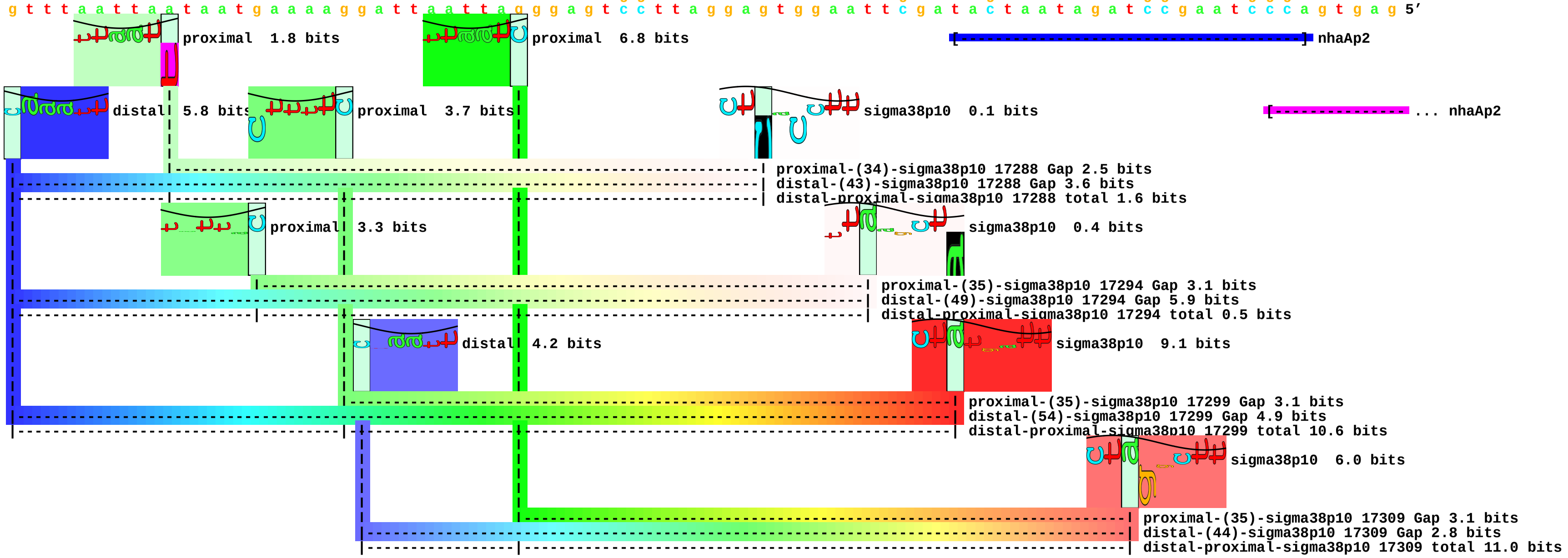

piece 151, NC_000913, osmFp, config: linear, direction: -, begin: 2217616, end: 2217531

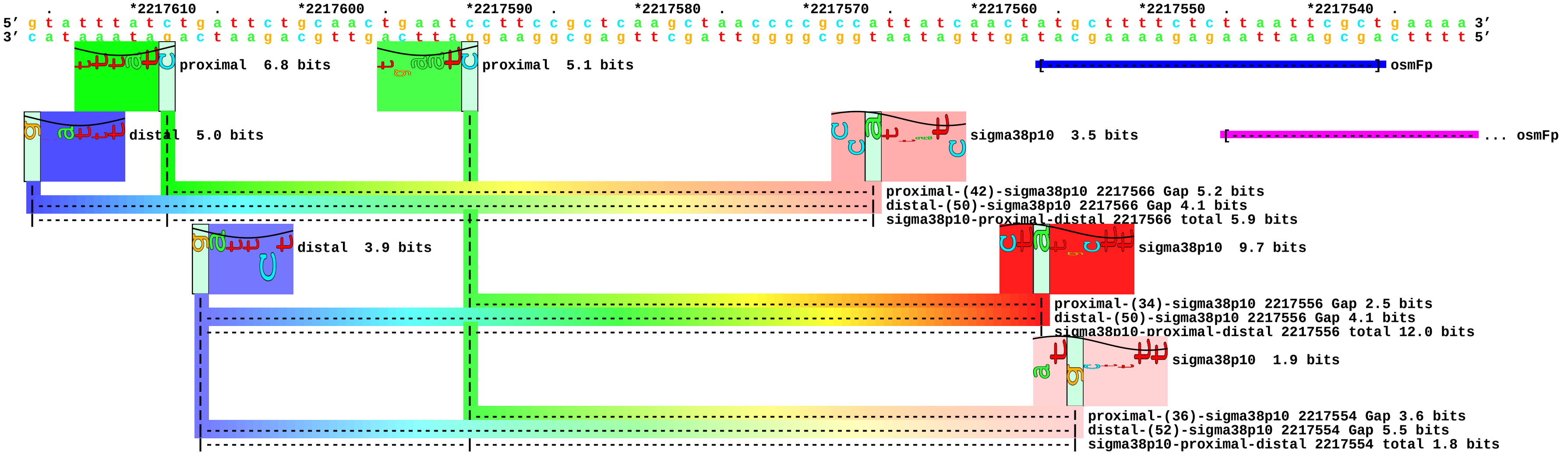

piece 152, NC_000913, pcnBp4, config: linear, direction: -, begin: 159461, end: 159376

```
   *159460   .      *159450   .      *159440   .      *159430   .      *159420   .      *159410   .      *159400   .      *159390   .      *159380
5' t t t g g c t g g a a a g t g c c a g a t t a c a t t c a t c t g c c g c t g g c g c t t a a t c c a c a a g g c g c t a a a c t t t c c a a g c a g a a t c a t g c g c c 3'
3' a a a c c g a c c t t t c a c g g t c t a a t g t a a g t a g a c g g c g a c c g c g a a t t a g g t g t t c c g c g a t t t g a a a g g t t c g t c t t a g t a c g c g g 5'
```

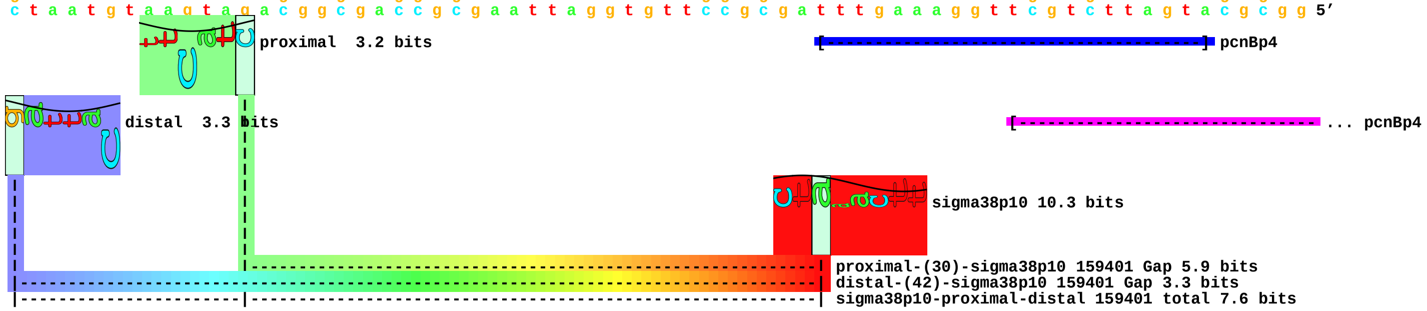

piece 153, NC_000913, pfkBp1, config: linear, direction: +, begin: 1804305, end: 1804390

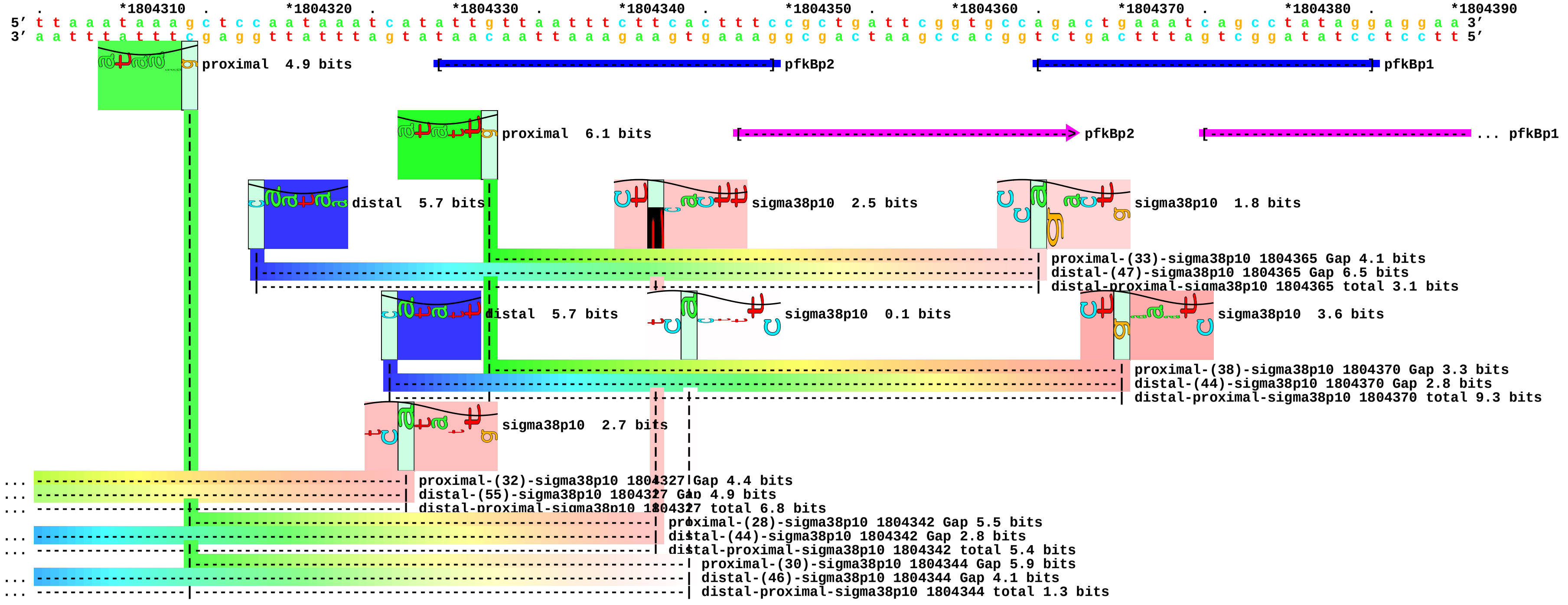

piece 154, NC_000913, pfkBp2, config: linear, direction: +, begin: 1804269, end: 1804354

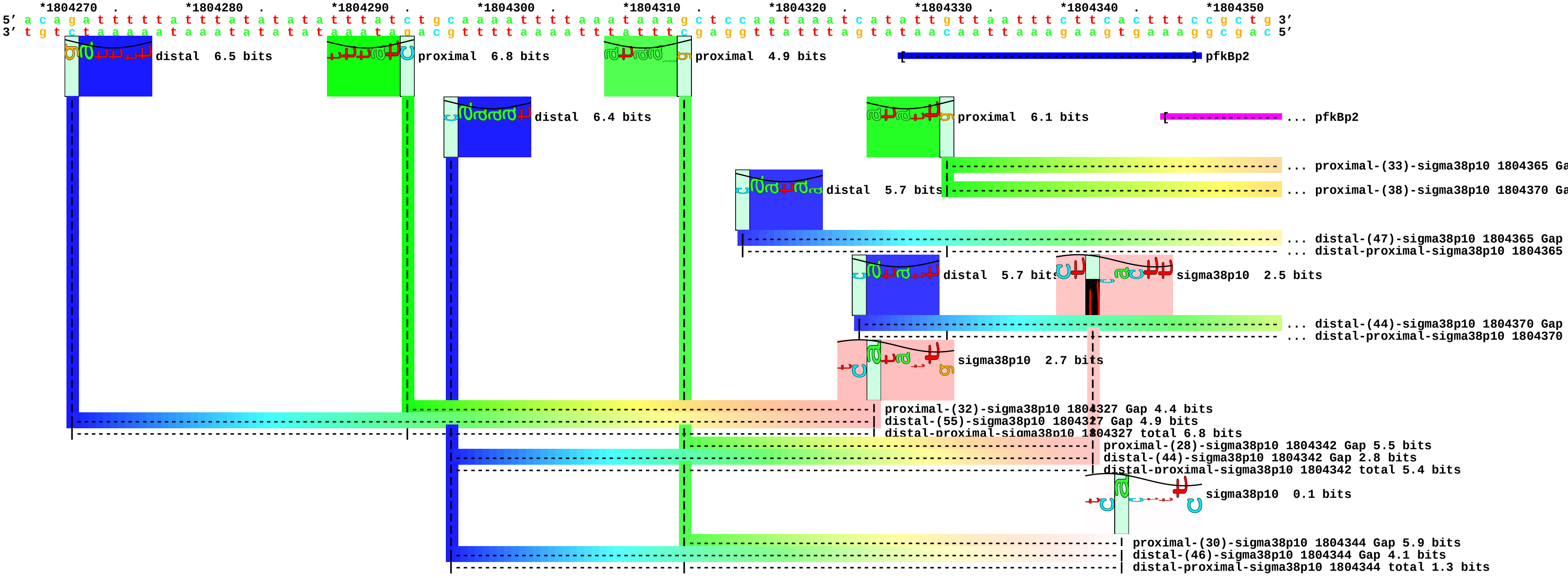

piece 155, NC_000913, proPp2, config: linear, direction: +, begin: 4328359, end: 4328444

```
    *4328360  .       *4328370  .       *4328380  .       *4328390  .       *4328400  .       *4328410  .       *4328420  .       *4328430  .       *4328440
5' t t c a g g c g t c a a c t g g t t t g a t t g t a c a t t c c t t a a c c g g a g g g t g t a a g c a a a c c c g c t a c g c t t g t t a c a g a g a t t g c a t c c t g 3'
3' a a g t c c g c a g t t g a c c a a a c t a a c a t g t a a g g a a t t g g c c t c c c a c a t t c g t t t g g g c g a t g c g a a c a a t g t c t c t a a c g t a g g a c 5'
```

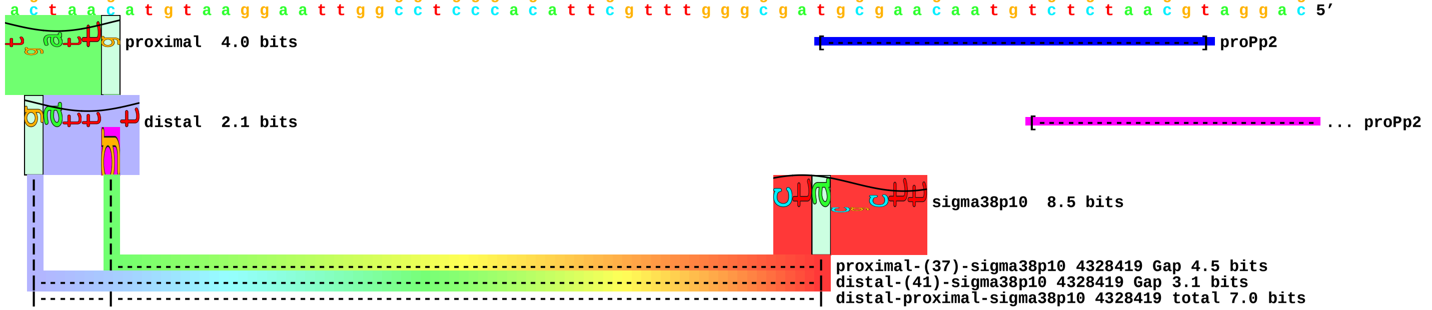

piece 156, NC_000913, puuDp, config: linear, direction: +, begin: 1359005, end: 1359090

```
   .       *1359010  .       *1359020  .       *1359030  .       *1359040  .       *1359050  .       *1359060  .       *1359070  .       *1359080  .       *1359090
5' a t g c a a a c t a a a t g t t t g t c a a a t g t t a a a t t g a g t t t g c a a a a a t g a a a a c c c a c t g c t a g a t t g a a a a a a a t a t t g a a c a t a a a a g 3'
3' t a c g t t t g a t t t a c a a a c a g t t t a c a a t t t a a c t c a a a c g t t t t t a c t t t t g g g t g a c g a t c t a a c t t t t t t t a t a a c t t g t a t t t t c 5'
```

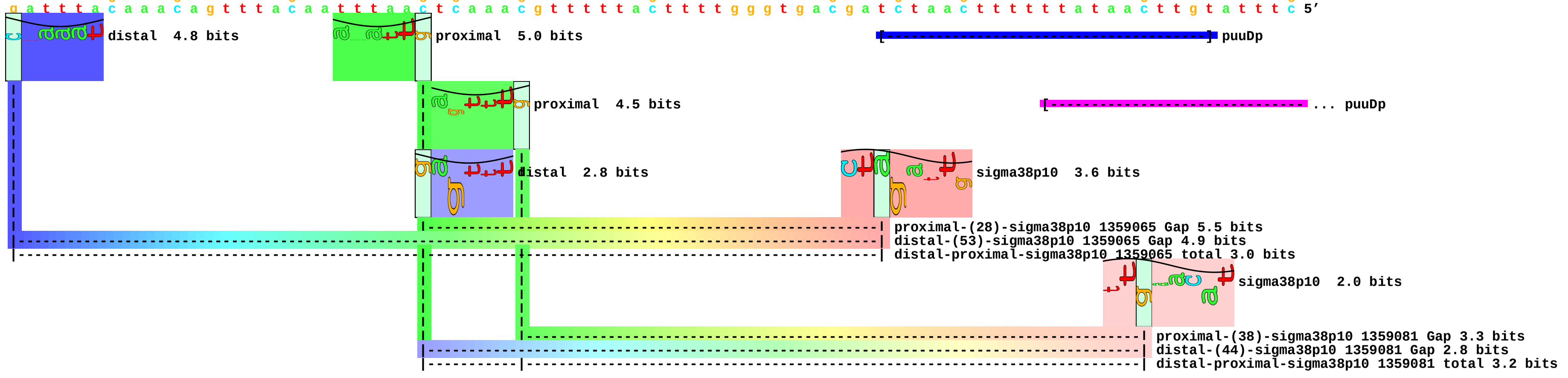

piece 157, NC_000913, rraAp, config: linear, direction: -, begin: 4117450, end: 4117365

```
   *4117450  .         *4117440  .         *4117430  .         *4117420  .         *4117410  .         *4117400  .         *4117390  .         *4117380  .         *4117370  .
5' c a t a a c t g a c a a a t a t c a a t t a a c a a t t g a t g a t t t t g c c a a c a g c c c a c a t a g c g c g a t a t a c t g a a a a t t c t c g c a g c a a c t g a 3'
3' g t a t t g a c t g t t t a t a g t t a a t t g t t a a c t a c t a a a a c g g t t g t c g g g t g t a t c g c g c t a t a t g a c t t t t a a g a g c g t c g t t g a c t 5'
```

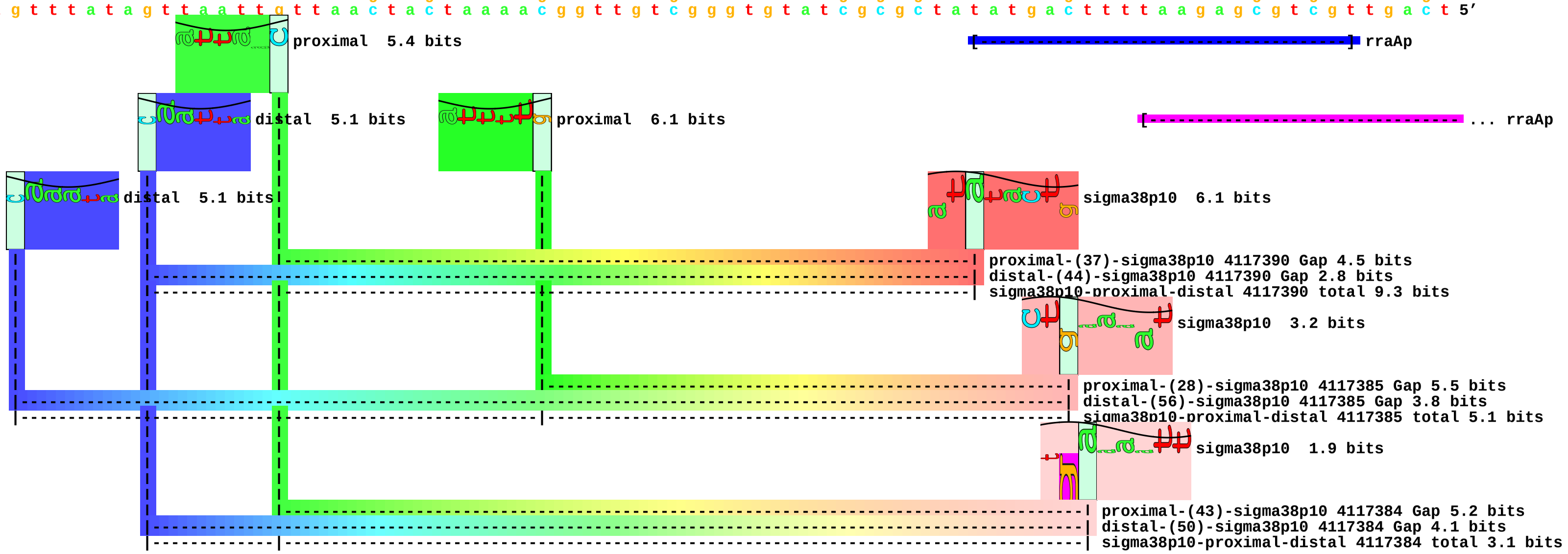

piece 158, NC_000913, rsdp1, config: linear, direction: -, begin: 4195049, end: 4194964

```
          .         *4195040  .         *4195030  .         *4195020  .         *4195010  .         *4195000  .         *4194990  .         *4194980  .         *4194970  .
5' g c t g a c c c a c a a g a t c g a a a t t t g c c g c t t c g c c a t a t g g c a a t t c t c c c t t c g g c a a c c a t a a t t t t t g t t c a t g g c t g a c g a c c 3'
3' c g a c t g g g t g t t c t a g c t t t a a a c g g c g a a g c g g t a t a c c g t t a a g a g g g a a g c c g t t g g t a t t a a a a a c a a g t a c c g a c t g c t g g 5'
```

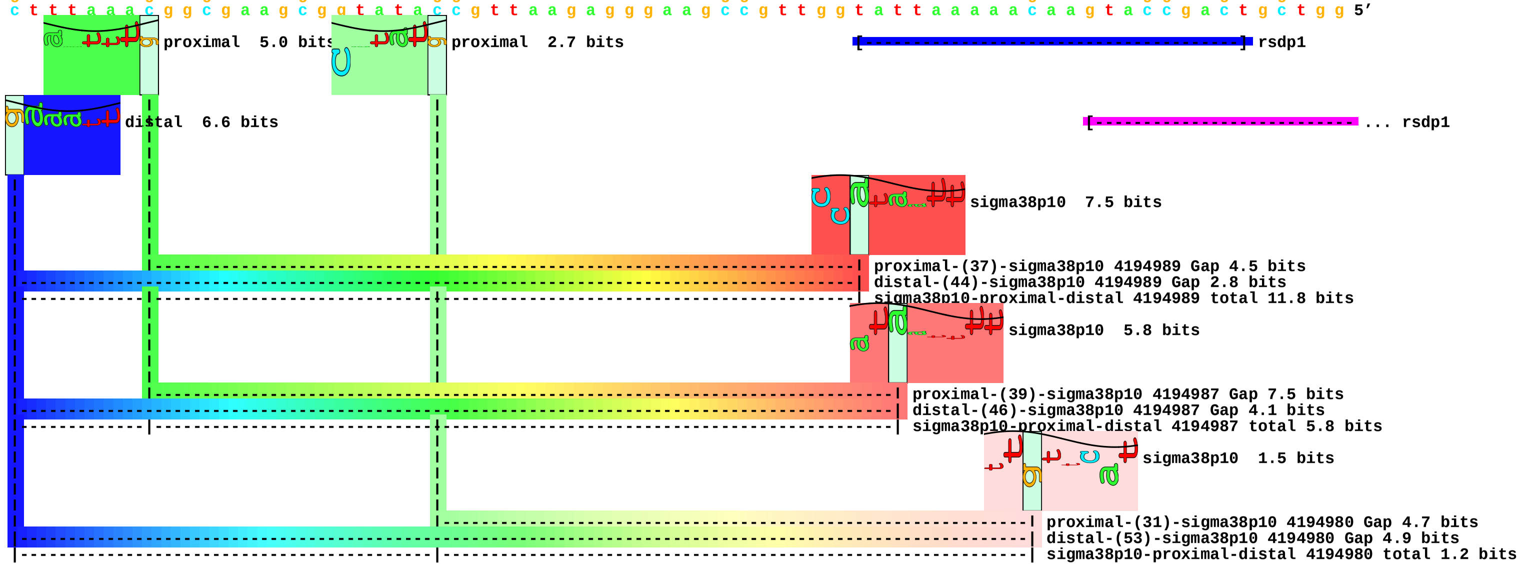

piece 159, NC_000913, sodCp, config: linear, direction: -, begin: 1722775, end: 1722690

```
   .       *1722770   .       *1722760   .       *1722750   .       *1722740   .       *1722730   .       *1722720   .       *1722710   .       *1722700   .       *1722690
5' g t a c g a c g t a c c g t a a g c a c t t t t a g g a a t a g c c g c c g t t c a a a a a t g t g t c a c t g g t t t a c a c t t a t t c a g g a a t g c a c a a t g a a 3'
3' c a t g c t g c a t g g c a t t c g t g a a a a t c c t t a t c g g c g g c a a g t t t t t a c a c a g t g a c c a a a t g t g a a t a a g t c c t t a c g t g t t a c t t 5'
```

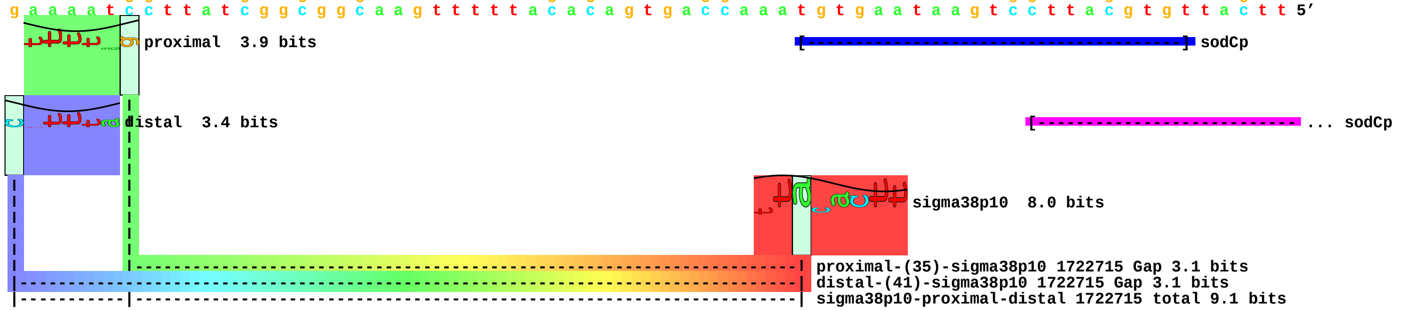

piece 160, NC_000913, speBp, config: linear, direction: -, begin: 3082039, end: 3081954

```
              .         *3082030  .         *3082020  .         *3082010  .         *3082000  .         *3081990  .         *3081980  .         *3081970  .         *3081960  .
5' a t c t t g a t g c t g a a c t g c a a c a a c a g t t c c t t g a a g a g t t c g a g g c a g g t t t g t a c g g t t a t a c t t a t c t t g a a g a t g a g t a a g t c 3'
3' t a g a a c t a c g a c t t g a c g t t g t t g t c a a g g a a c t t c t c a a g c t c c g t c c a a a c a t g c c a a t a t g a a t a g a a c t t c t a c t c a t t c a g 5'
```

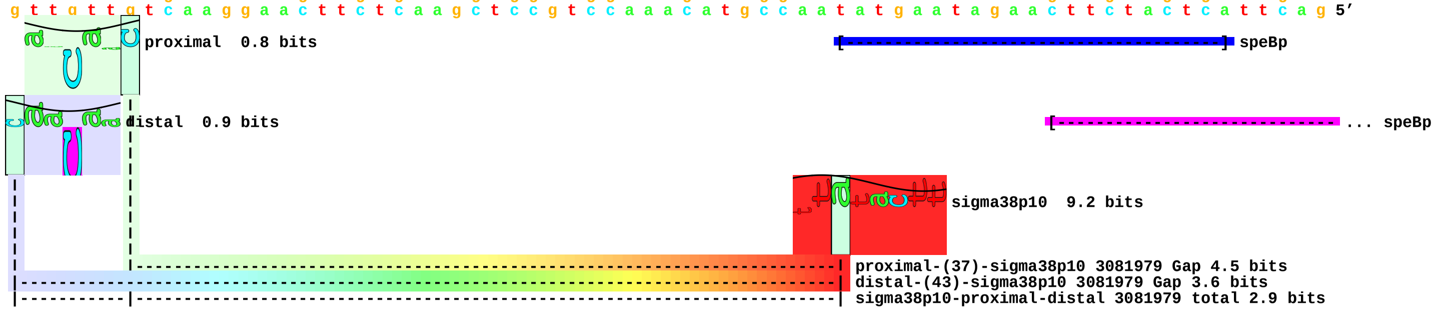

piece 161, NC_000913, tolCp4, config: linear, direction: +, begin: 3176015, end: 3176100

```
   .         *3176020  .         *3176030  .         *3176040  .         *3176050  .         *3176060  .         *3176070  .         *3176080  .         *3176090  .         *3176100
5' t g a g g c a c a t t a a c g c c c t a t g g c a c g t a a c g c c a a c c t t t t g c g g t a g c g g c t t c t g c t a g a a t c c g c a a t a a t t t t a c a g t t t g 3'
3' a c t c c g t g t a a t t g c g g g a t a c c g t g c a t t g c g g t t g g a a a a c g c c a t c g c c g a a g a c g a t c t t a g g c g t t a t t a a a a t g t c a a a c 5'
                                                                                                                                [---------------------------------------] tolCp4
                                                                                                                                                                         [---------- ... tolCp4
```

piece 162, NC_000913, xthAp, config: linear, direction: +, begin: 1830352, end: 1830437

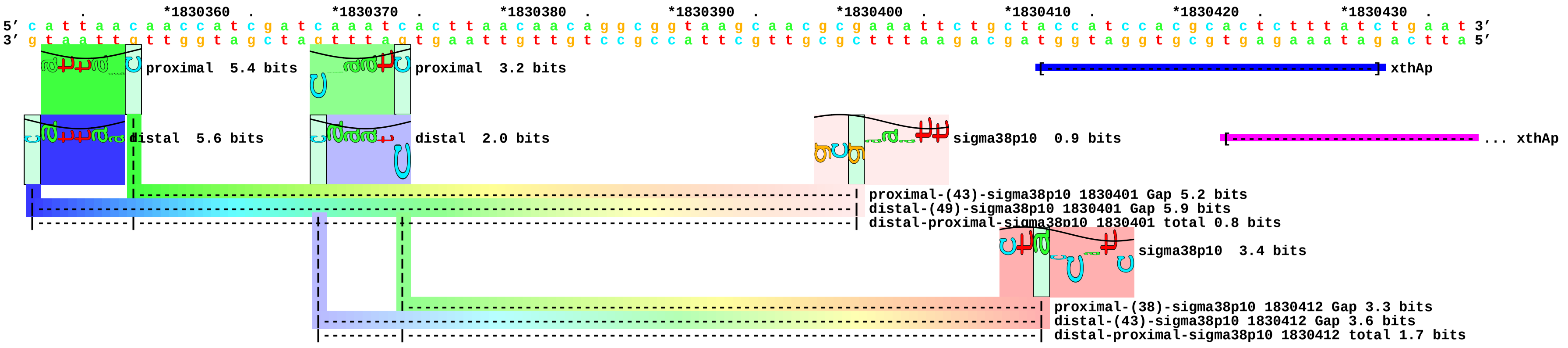

piece 163, NC_000913, yadVp2, config: linear, direction: -, begin: 156300, end: 156215

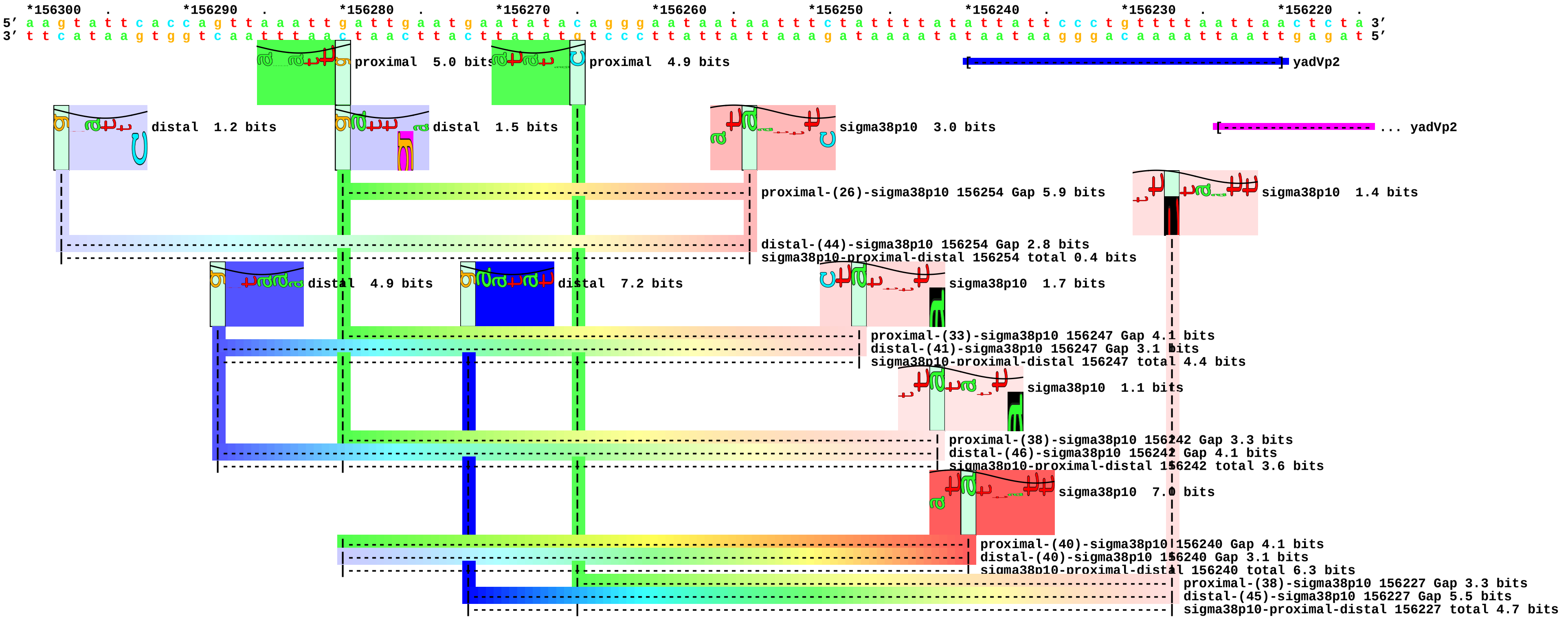

piece 164, NC_000913, yciZp, config: linear, direction: -, begin: 1342729, end: 1342644

```
            .        *1342720  .         *1342710  .         *1342700  .         *1342690  .         *1342680  .         *1342670  .         *1342660  .         *1342650  .
5' c c t g g a t g a t a a t t a g c c g t t g c t g t t a a a c a c t t c t c t t t t a g c g g c a a t t t t t g c a t t a t t c a t t t t c c c c c t t t t g c t a t a a g 3'
3' g g a c c t a c t a t t a a t c g g c a a c g a c a a t t t g t g a a g a g a a a a t c g c c g t t a a a a a c g t a a t a a g t a a a a g g g g g a a a a c g a t a t t c 5'
```

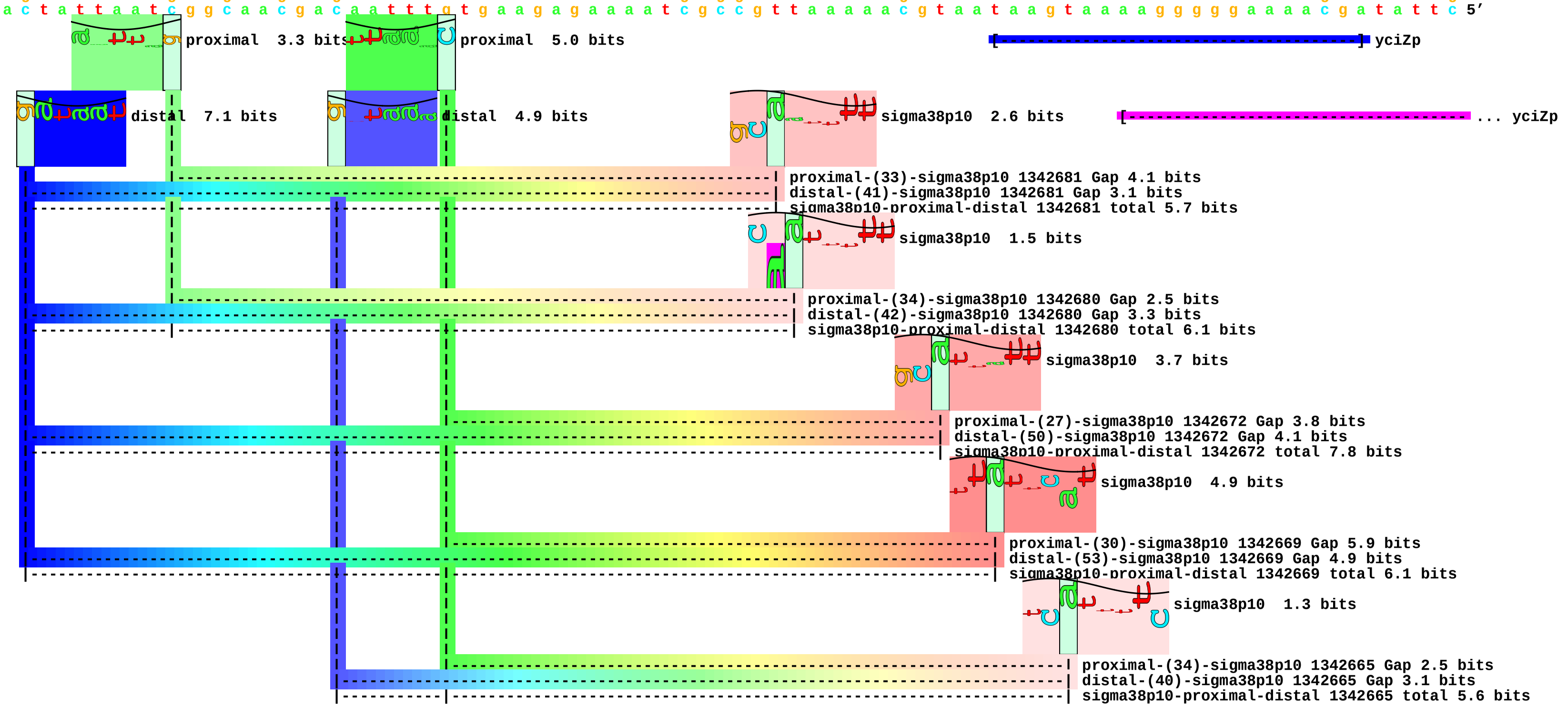

piece 165, NC_000913, yddGp, config: linear, direction: -, begin: 1545299, end: 1545214

```
              .         *1545290  .         *1545280  .         *1545270  .         *1545260  .         *1545250  .         *1545240  .         *1545230  .         *1545220  .
5' t a c c t c a a t a g c g g t a g a a a a a c g c a c c a c t g c c t g a c a g g c c a g t t a a a a a a a t g c t a t a a a a t t c a g c t t a a t t t t t a a c g g c a 3'
3' a t g g a g t t a t c g c c a t c t t t t t g c g t g g t g a c g g a c t g t c c g g t c a a t t t t t t t a c g a t a t t t t a a g t c g a a t t a a a a a t t g c c g t 5'
```

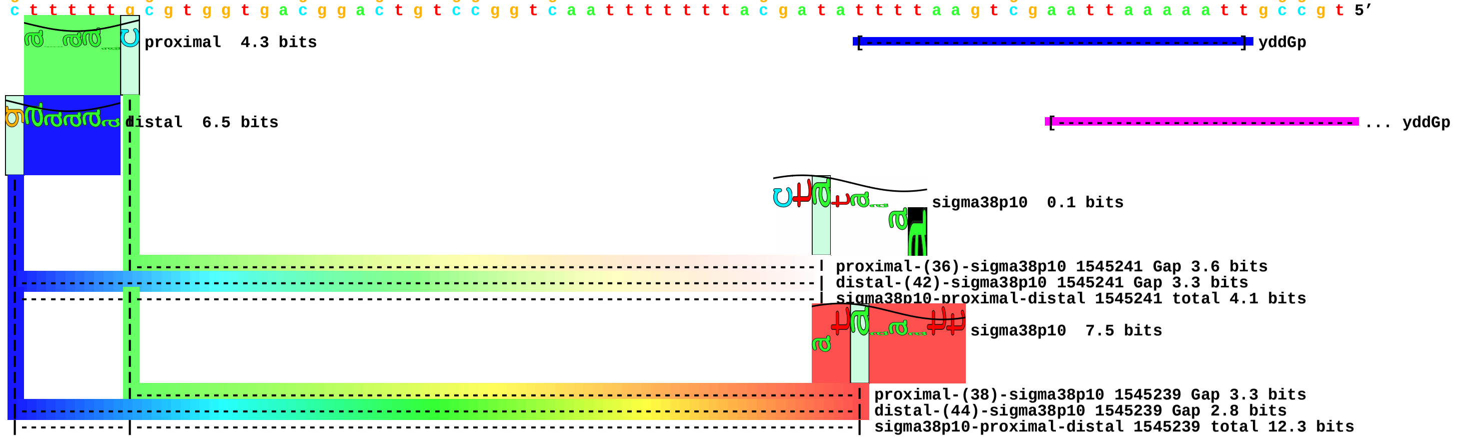

piece 166, NC_000913, ydhYp, config: linear, direction: -, begin: 1752735, end: 1752650

```
 .         *1752730 .         *1752720 .         *1752710 .         *1752700 .         *1752690 .         *1752680 .         *1752670 .         *1752660 .         *1752650
5' t t c t t g c t t t g t a a t t t t a a t t g a t a a c g a t c a a t g t t a a g g g c t c a g a a t a a t c a c a t t a t t t t t t c t c c t a a c c a t a a a g g a t t 3'
3' a a g a a c g a a a c a t t a a a a t t a a c t a t t g c t a g t t a c a a t t c c c g a g t c t t a t t a g t g t a a t a a a a a a g a g g a t t g g t a t t t c c t a a 5'
```

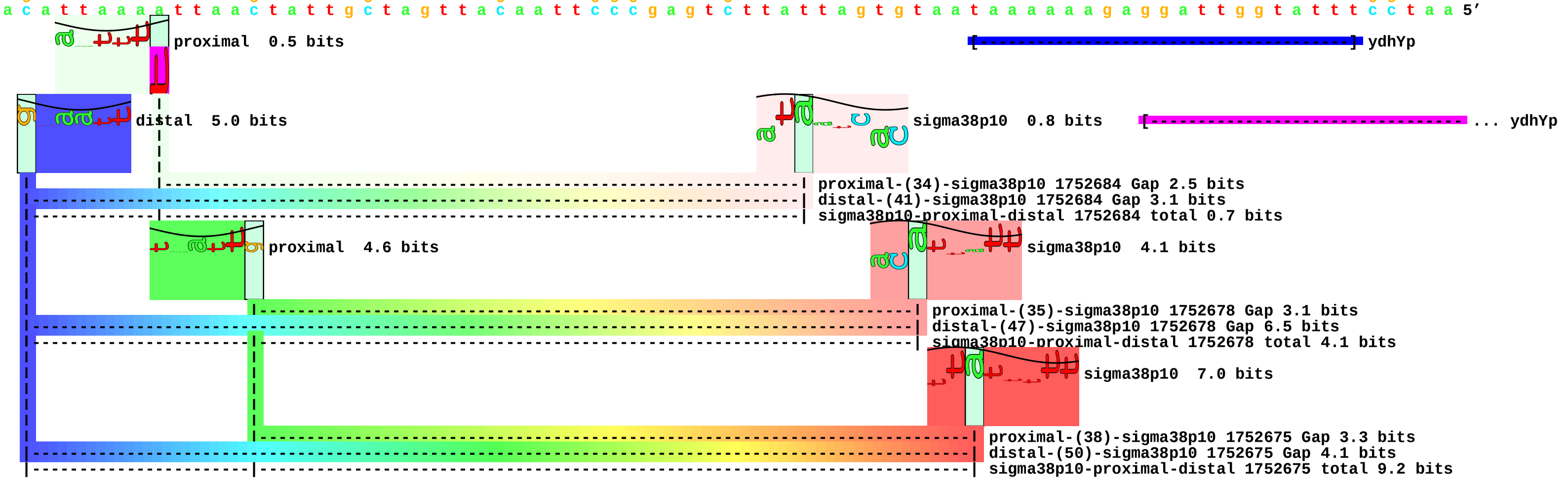

piece 167, NC_000913, yeiLp, config: linear, direction: +, begin: 2253246, end: 2253331

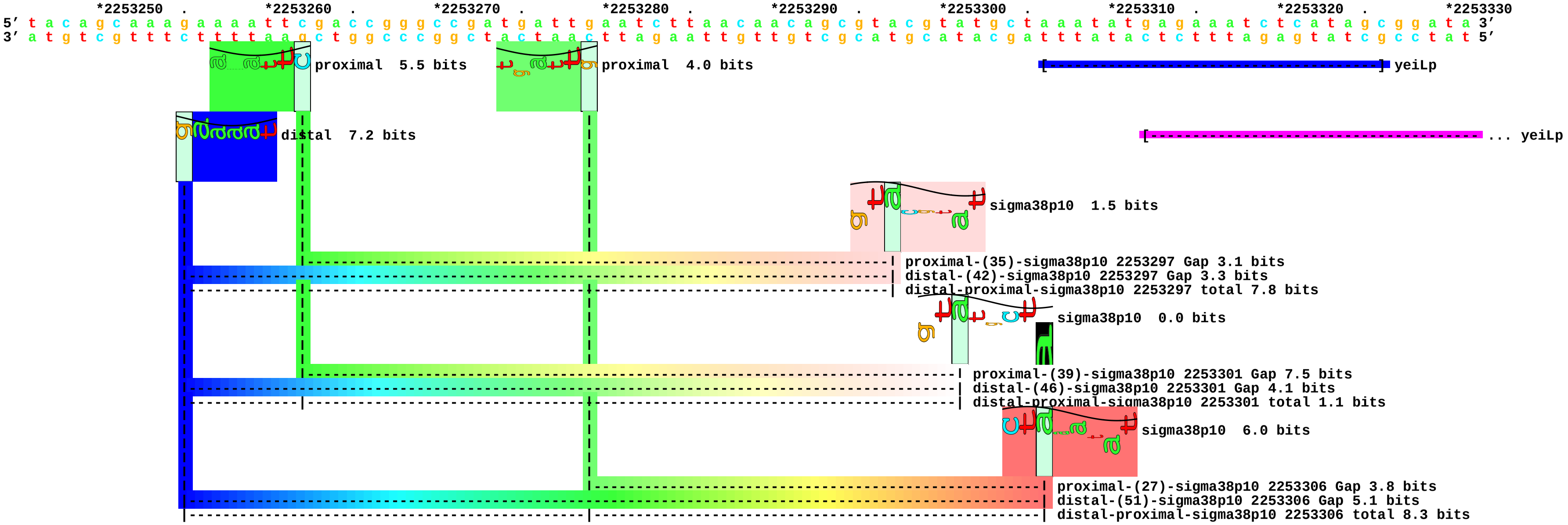

piece 168, NC_000913, yihGp, config: linear, direction: -, begin: 4044743, end: 4044658

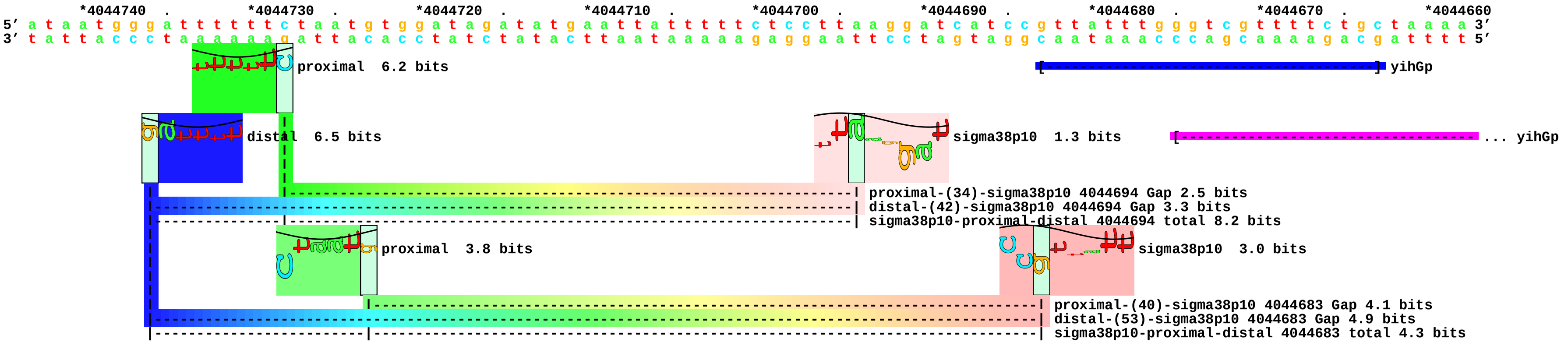

piece 169, NC_000913, ytfKp, config: linear, direction: +, begin: 4437463, end: 4437548

```
         .         *4437470  .         *4437480  .         *4437490  .         *4437500  .         *4437510  .         *4437520  .         *4437530  .         *4437540  .
5' a t a a t t g t a a c c t t t a g g t a a a a a a a g t t a t a c g c g g t g g a a a c a t t g c c c g g a t a g t c t a t a g t c a c t a a g c a t t a a a a t t t g c g 3'
3' t a t t a a c a t t g g a a a t c c a t t t t t t t c a a t a t g c g c c a c c t t t g t a a c g g g c c t a t c a g a t a t c a g t g a t t c g t a a t t t t a a a c g c 5'
```

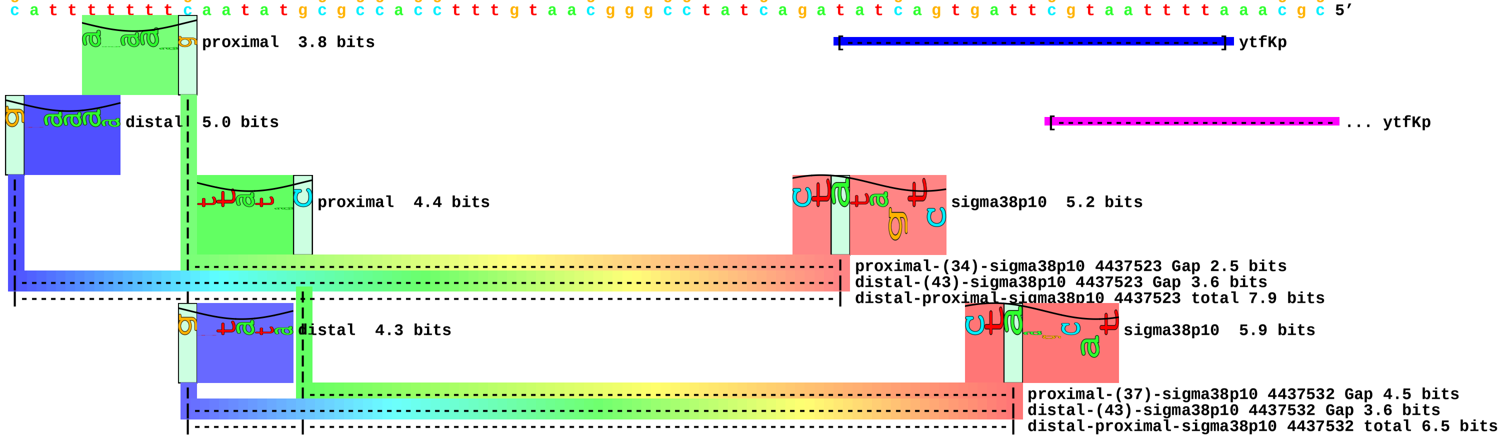
