## Supplement_for_sigma38_paper-directory for "*Escherichia coli σ*^38^ promoters use two UP elements instead of a −35 element: resolution of a paradox and discovery that *σ*^38^ transcribes ribosomal promoters": Estrem.Gourse1998-rrnB-P1-selection.pdf

piece 1, NC_000913.i4164330,4164353ggaaaatttttttcaaaagta, insert ggaaaatttttttcaaaagta between 4164330 and 4164353, config: linear, direction

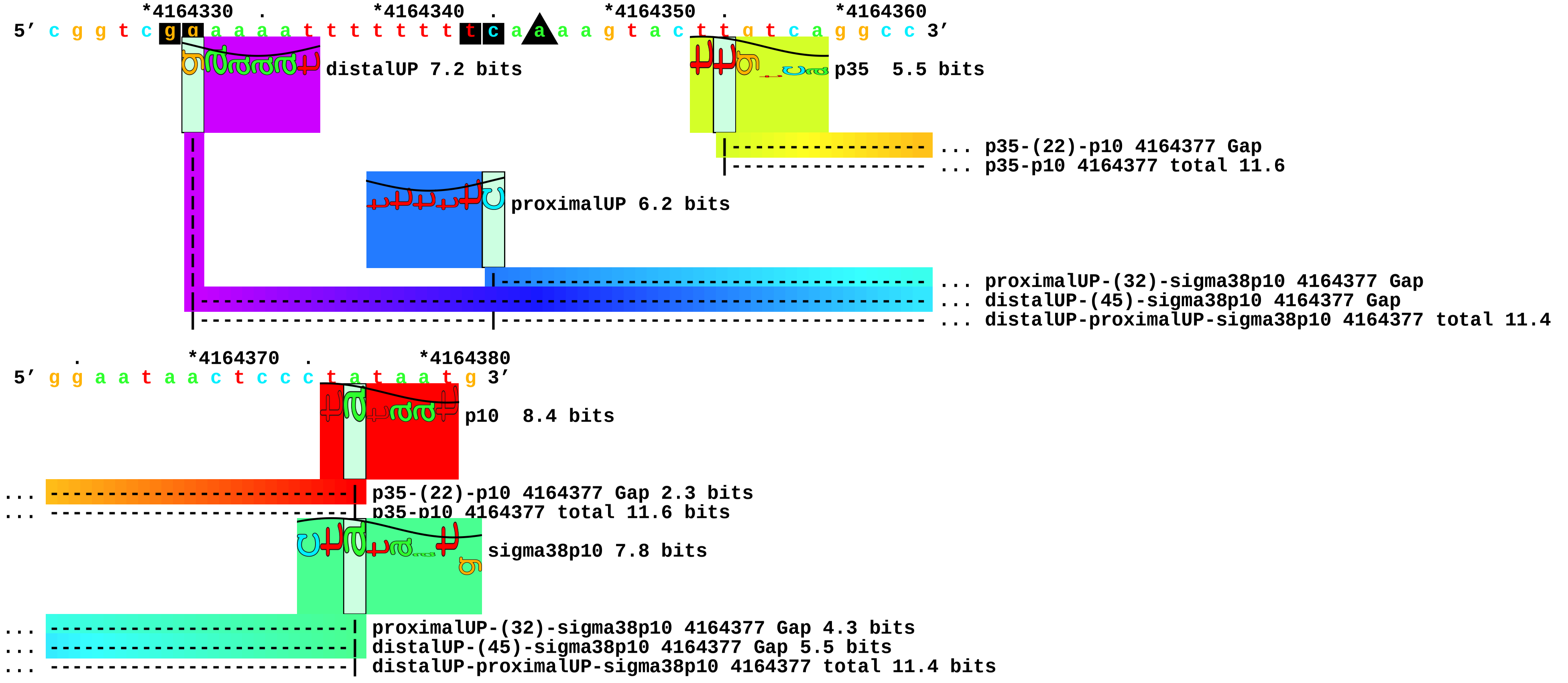

piece 2, NC_000913.i4164330,4164353agaaattttttttcgaaaaaca, insert agaaattttttttcgaaaaaca between 4164330 and 4164353, config: linear, direction

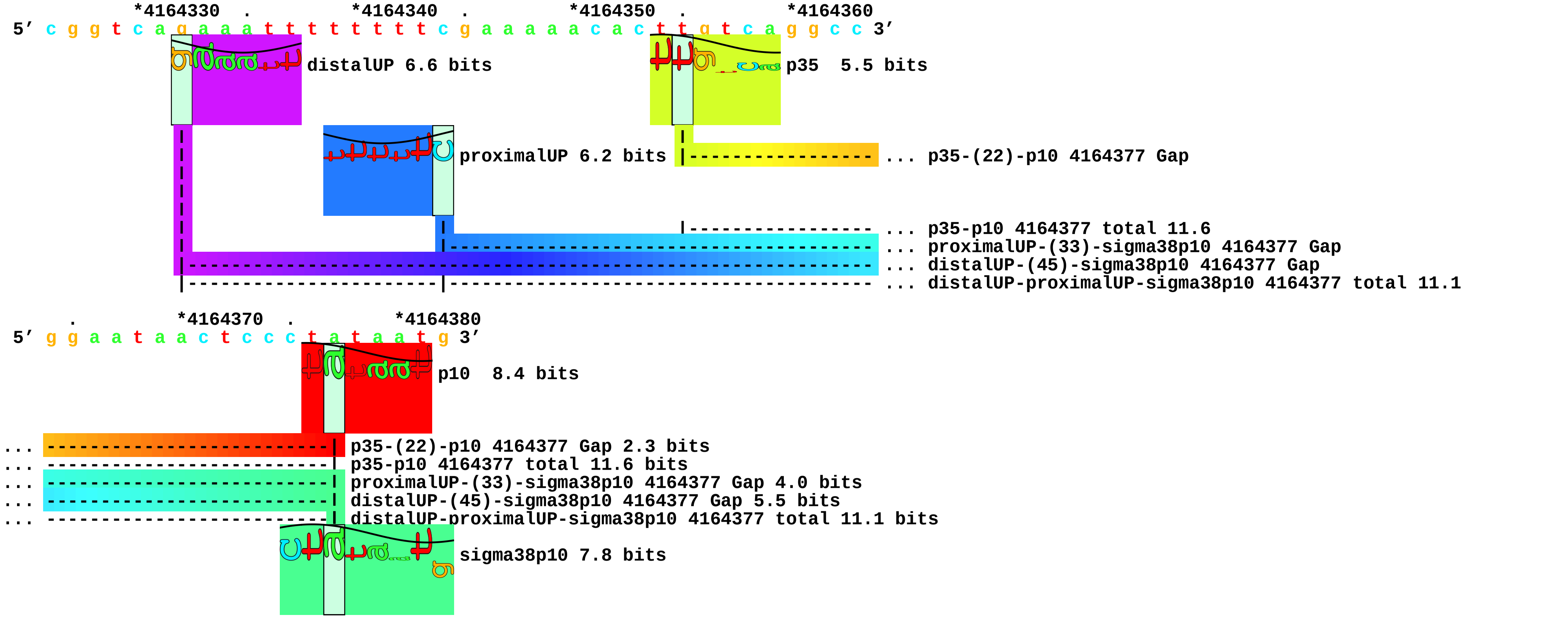

piece 3, NC_000913.i4164330,4164353taaaaattttttttgaaaaggg, insert taaaaattttttttgaaaaggg between 4164330 and 4164353, config: linear, direction

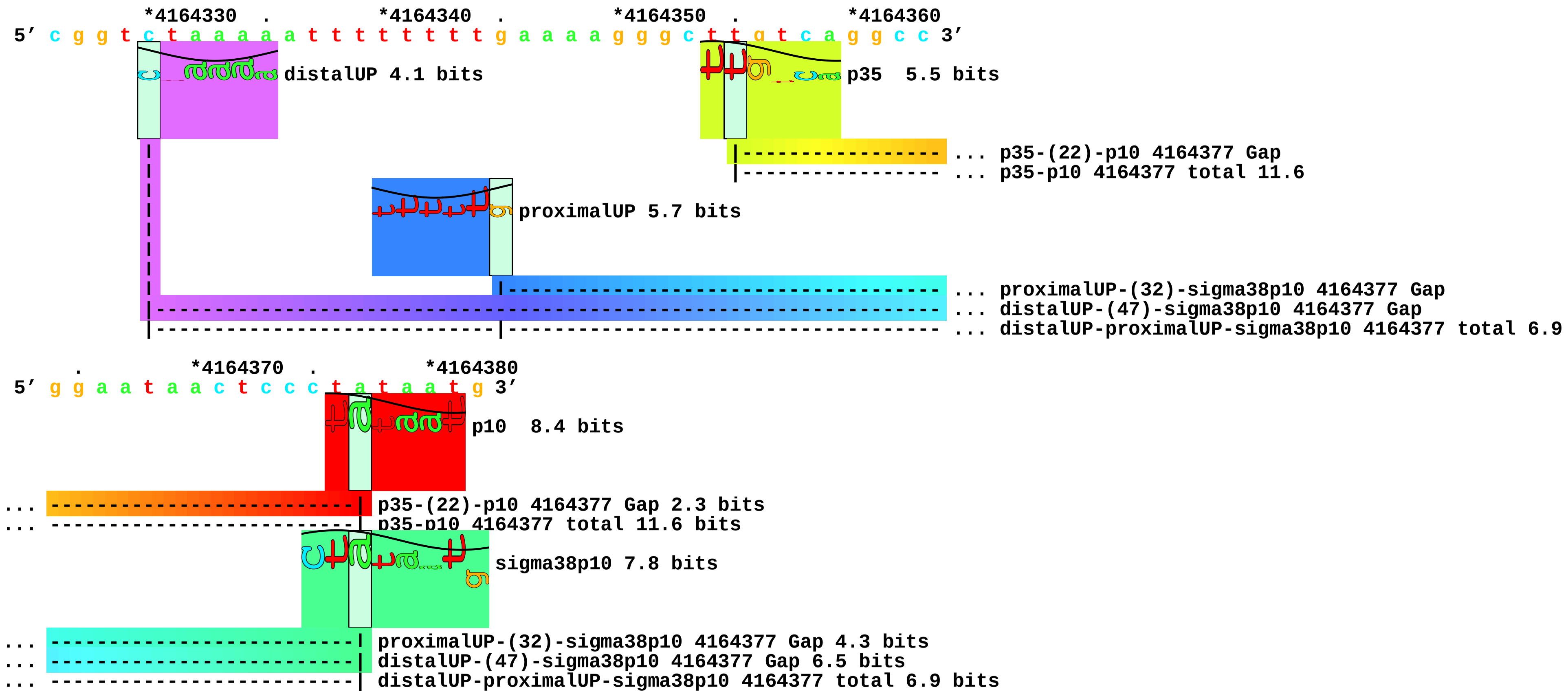

piece 4, NC_000913.i4164330,4164353caaaaatatttttgaaaaaaga, insert caaaaatatttttgaaaaaaga between 4164330 and 4164353, config: linear, direction

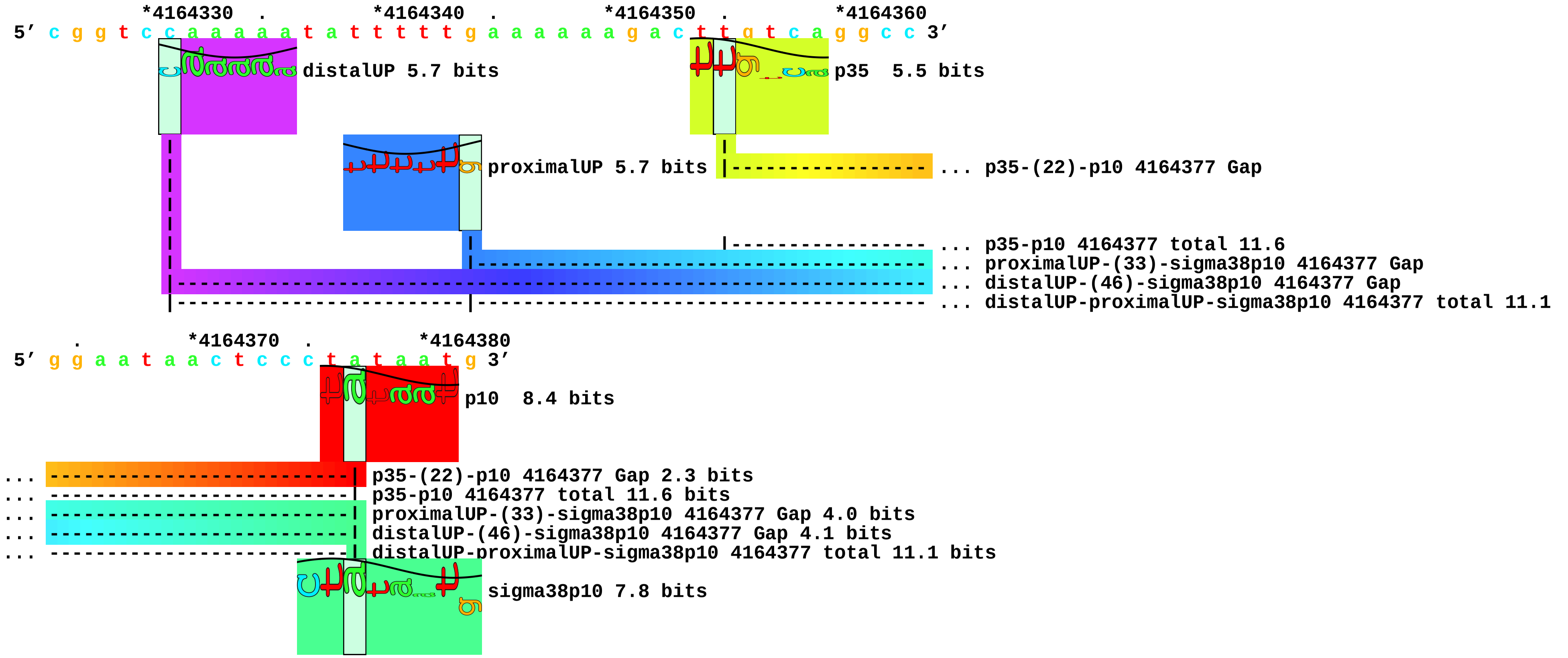

piece 5, NC_000913.i4164330,4164353ggaaatatttttcataaaccc, insert ggaaatatttttcataaaccc between 4164330 and 4164353, config: linear, direction

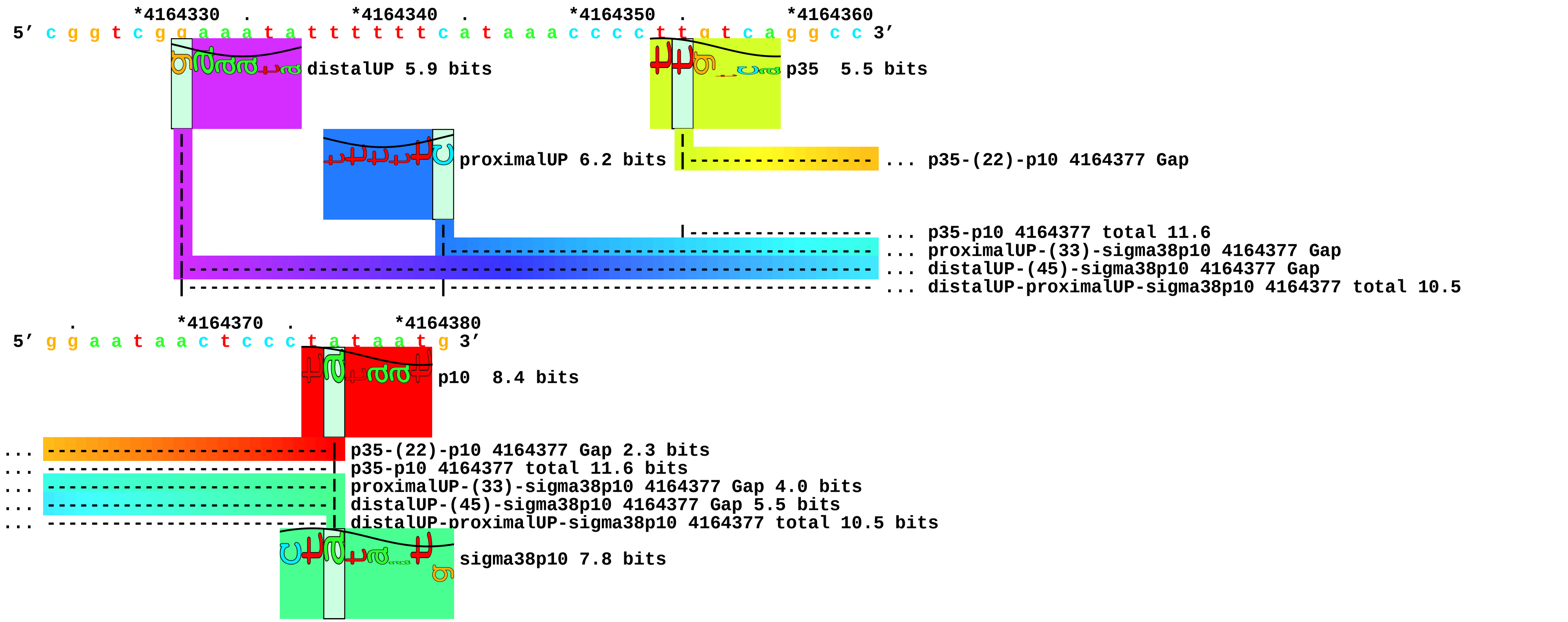

piece 6, NC_000913.i4164330,4164353agaaaaatattttcgaaaacta, insert agaaaaatattttcgaaaacta between 4164330 and 4164353, config: linear, direction

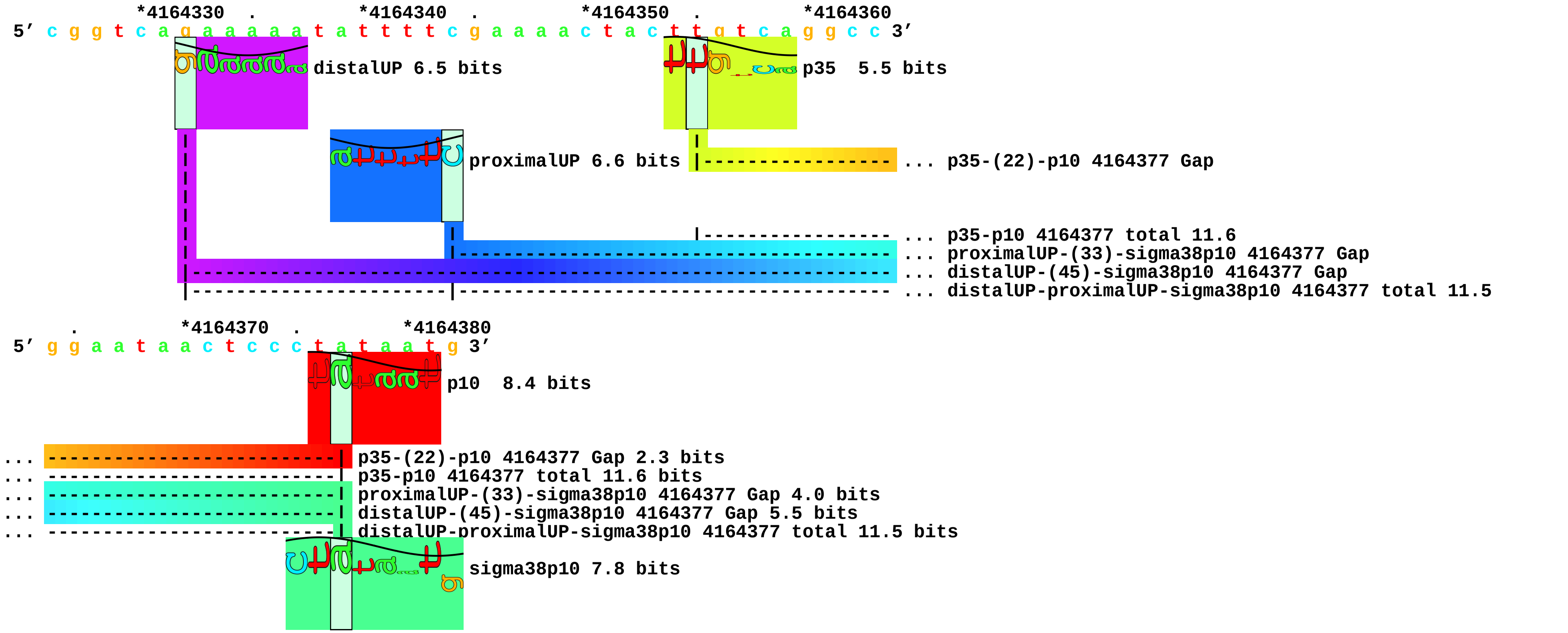

piece 7, NC_000913.i4164330,4164353aaaaatattttttcgaaaagta, insert aaaaatattttttcgaaaagta between 4164330 and 4164353, config: linear, direction

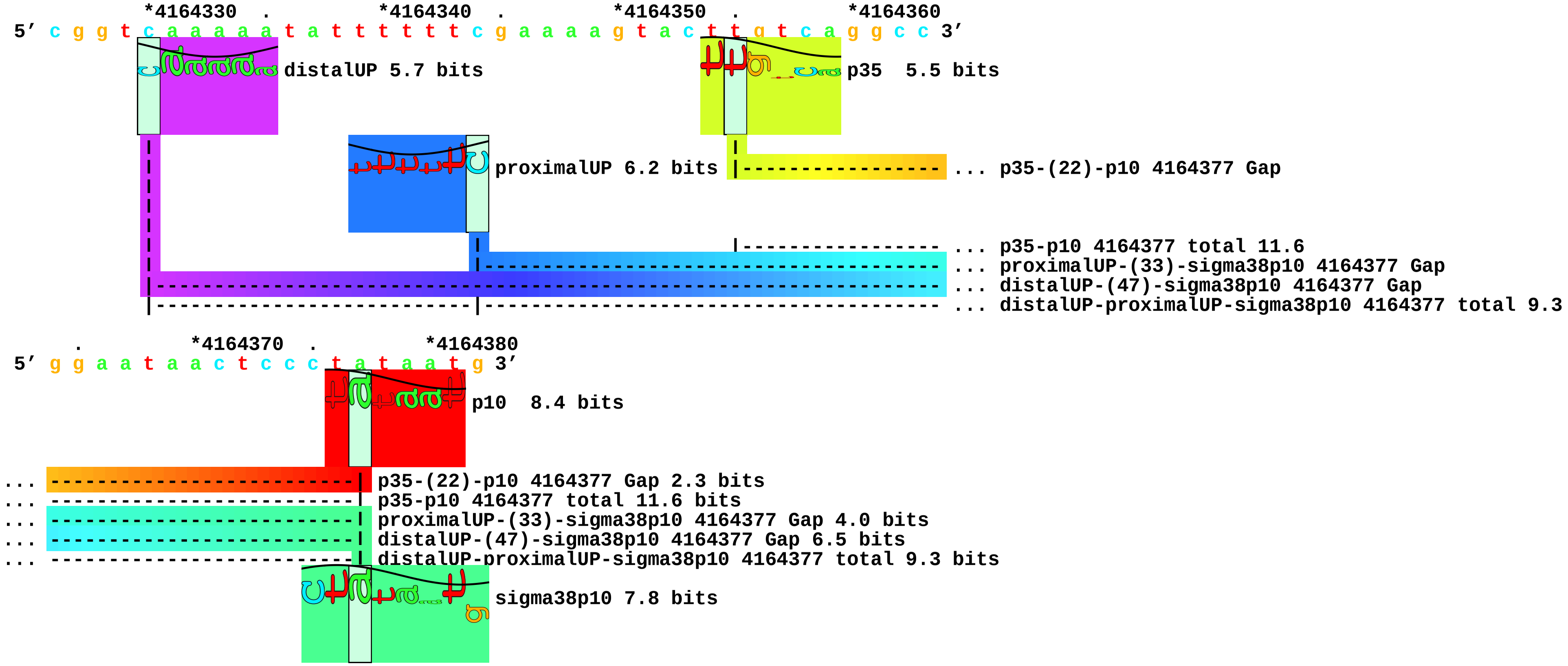

piece 8, NC_000913.i4164330,4164353taaatttttttttgcaaaagta, insert taaatttttttttgcaaaagta between 4164330 and 4164353, config: linear, direction

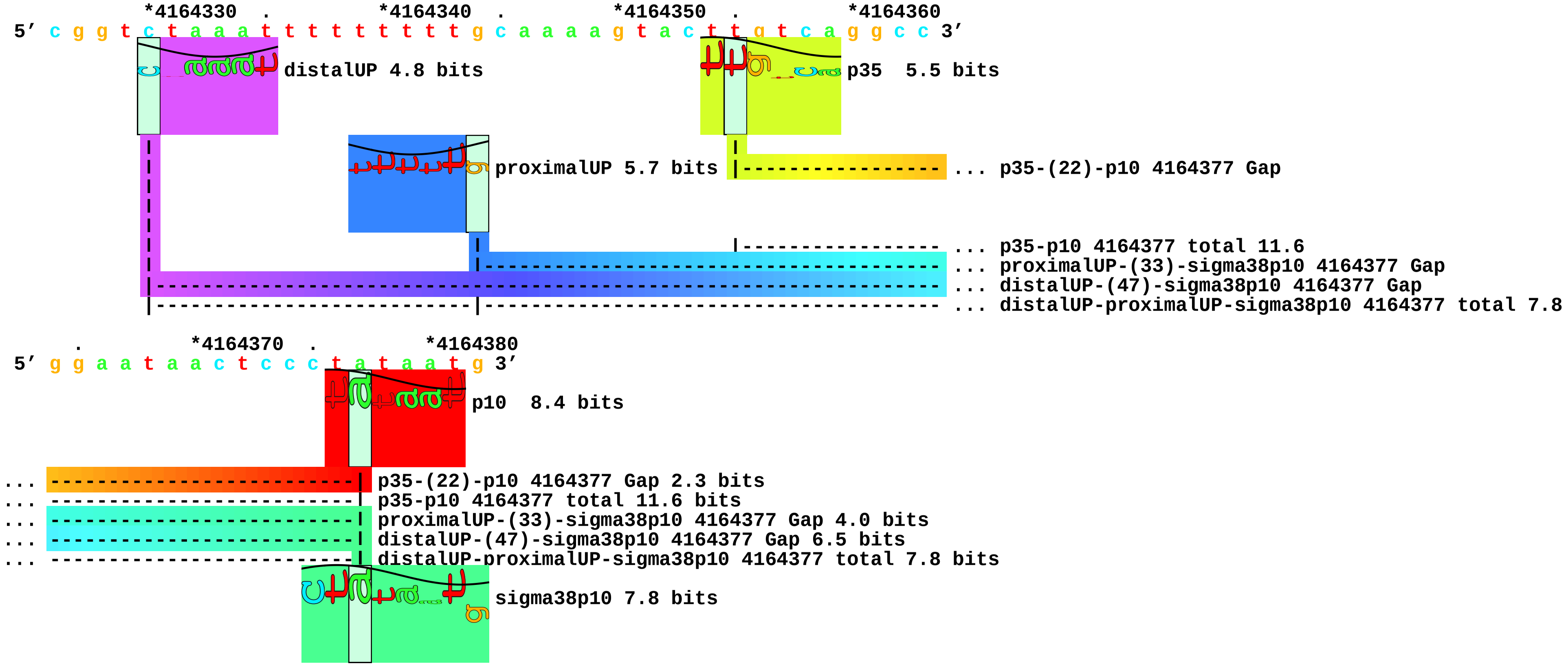

piece 9, NC_000913.i4164330,4164353acaaaaatatttttcaaaaccc, insert acaaaaatatttttcaaaaccc between 4164330 and 4164353, config: linear, direction

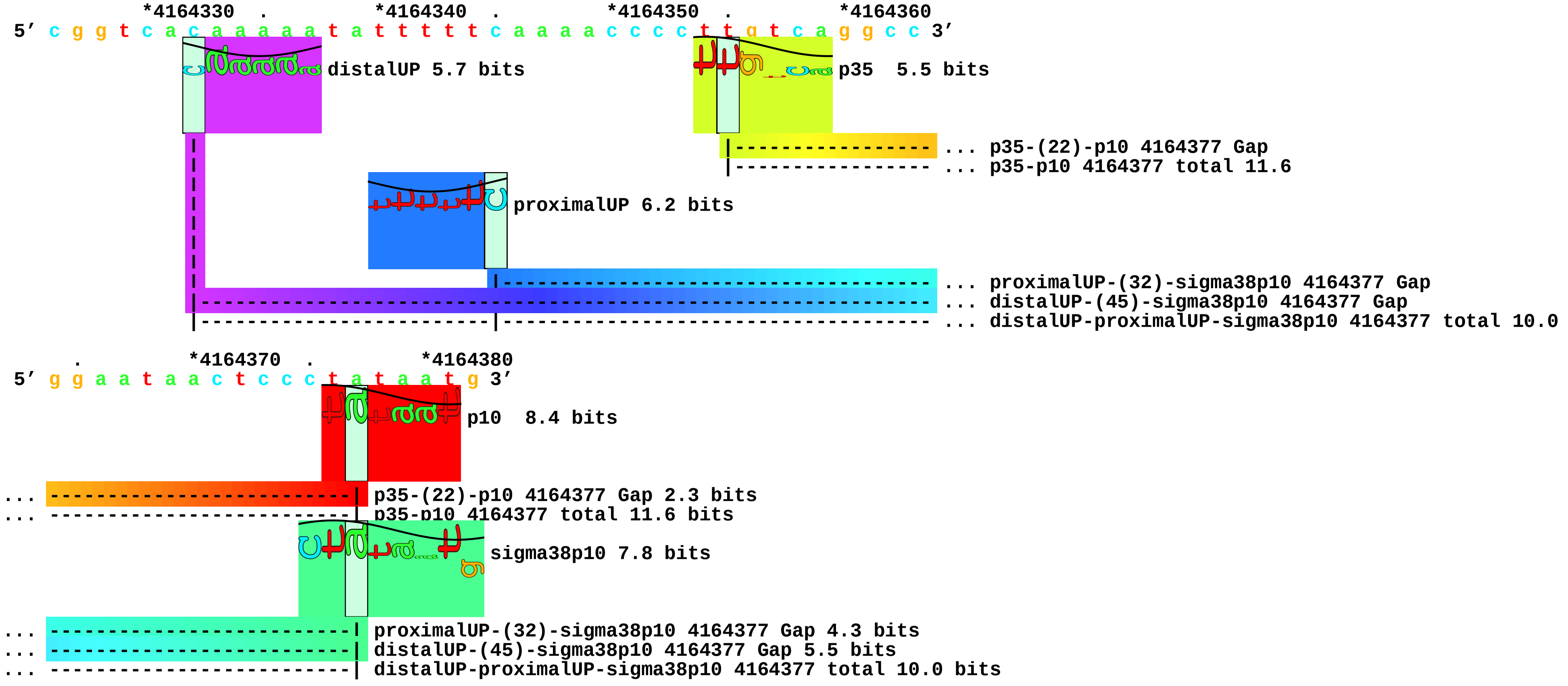

piece 10, NC_000913.i4164330,4164353ttaaatttttttttcgtaaaccc, insert ttaaatttttttttcgtaaaccc between 4164330 and 4164353, config: linear, directio

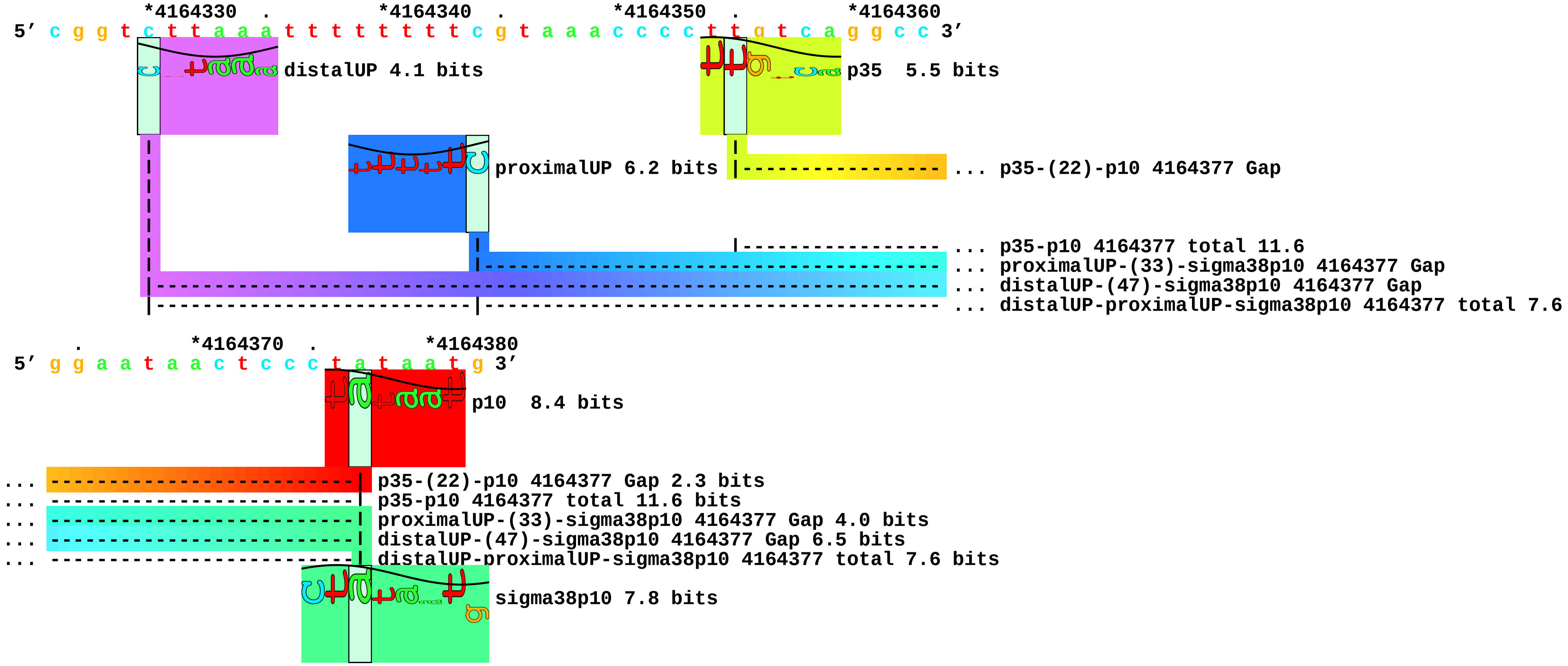

piece 11, NC_000913.i4164330,4164353ttaaatttttttcataaaccc, insert ttaaatttttttcataaaccc between 4164330 and 4164353, config: linear, directio

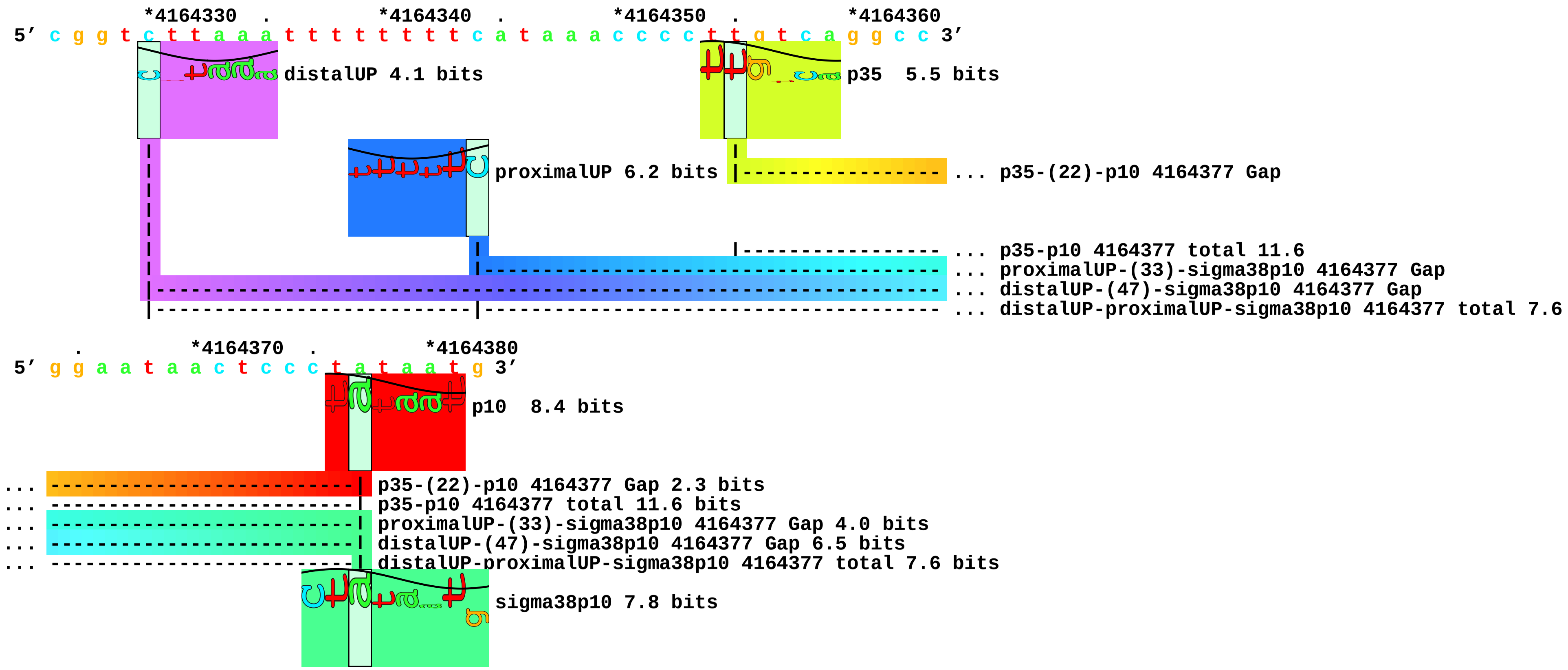

piece 12, NC_000913.i4164330,4164353tcaaatttttttttgcaaaccc, insert tcaaatttttttttgcaaaccc between 4164330 and 4164353, config: linear, directio

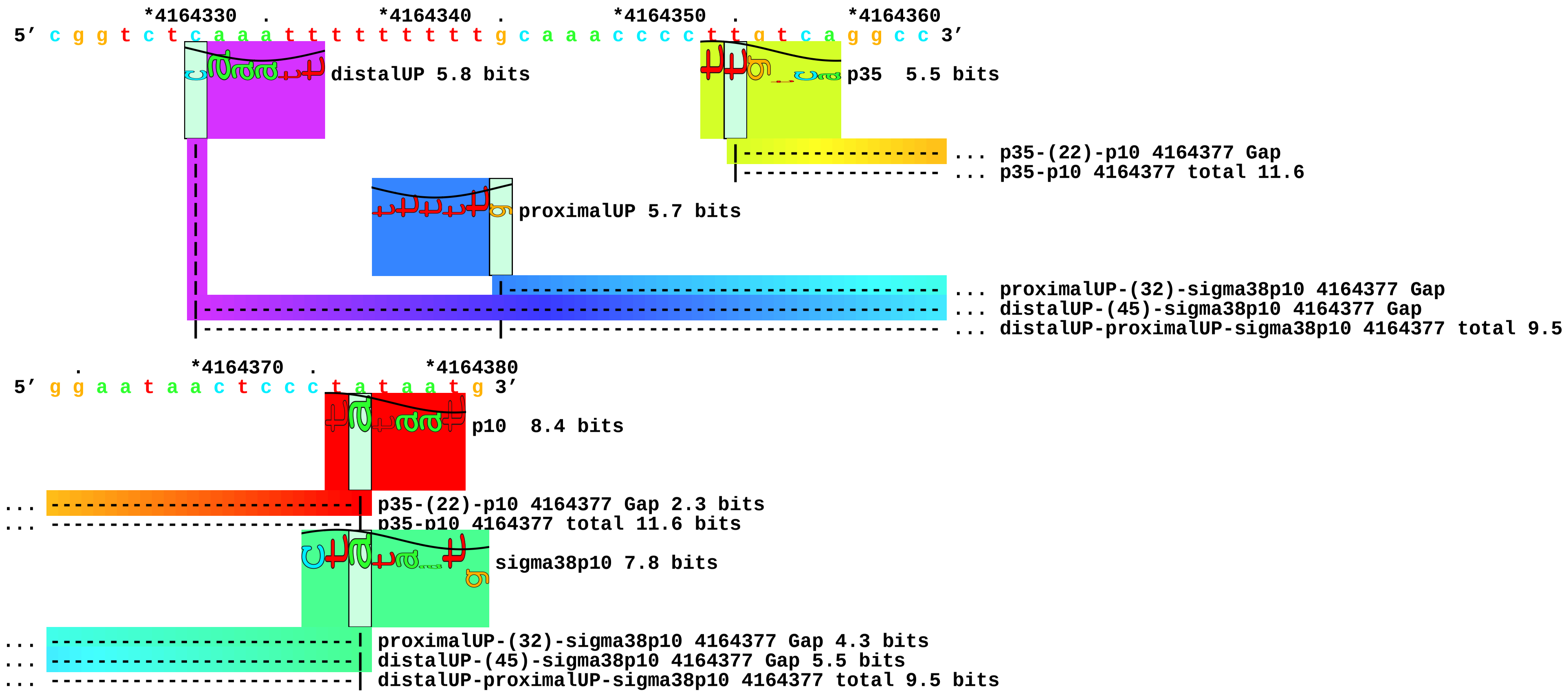

piece 13, NC_000913.i4164330,4164353caaatttttttttgctaaaccc, insert caaatttttttttgctaaaccc between 4164330 and 4164353, config: linear, directio

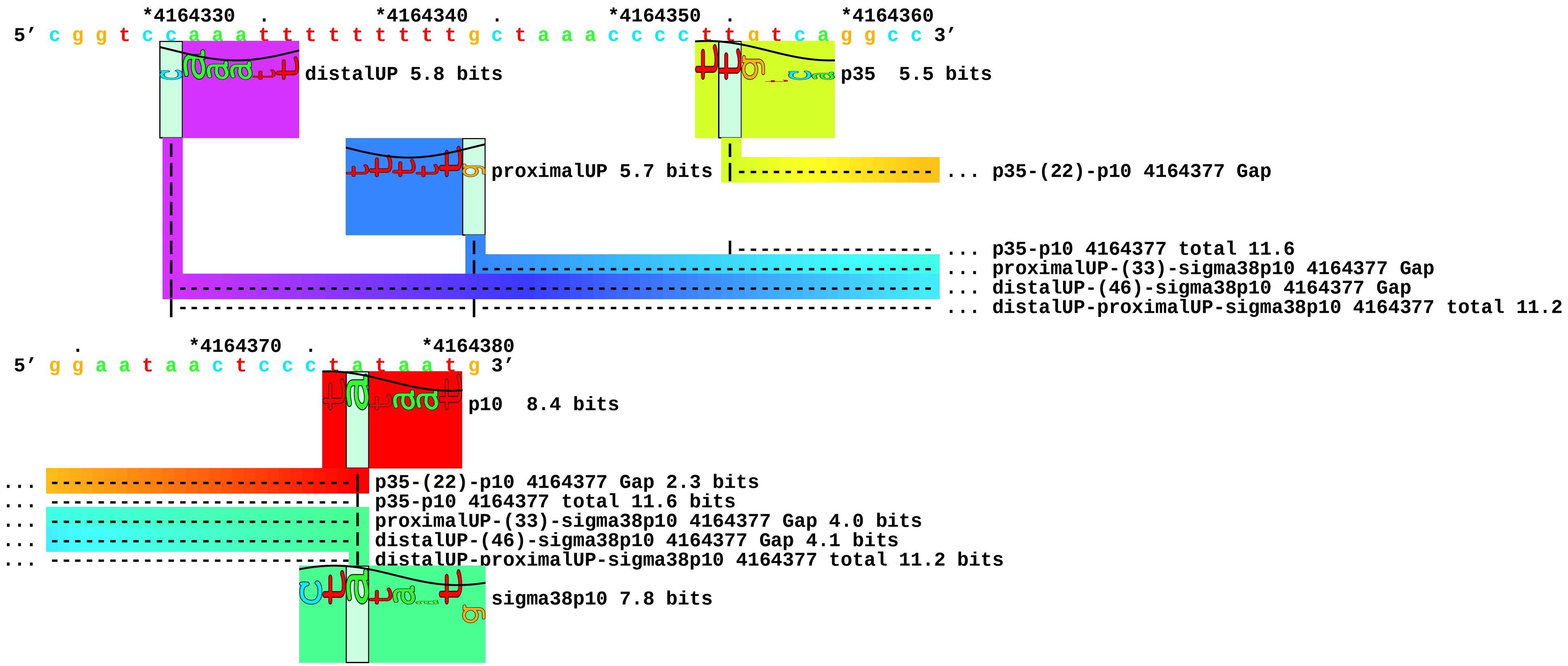

piece 14, NC_000913.i4164330,4164353aaaaatattttttgaaaagta, insert aaaaatattttttgaaaagta between 4164330 and 4164353, config: linear, directio

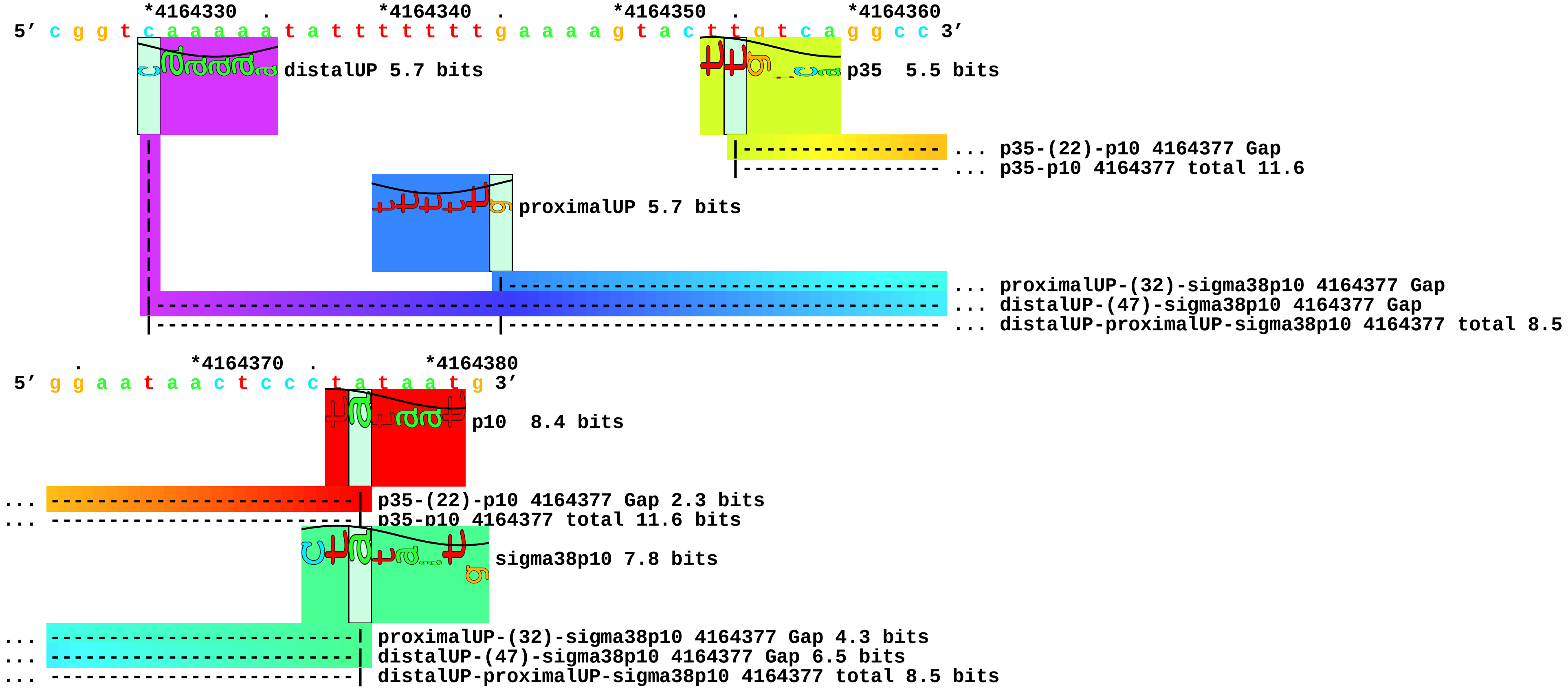

piece 15, NC_000913.i4164330,4164353taaaaatatttttcgtttaccc, insert taaaaatatttttcgtttaccc between 4164330 and 4164353, config: linear, directio

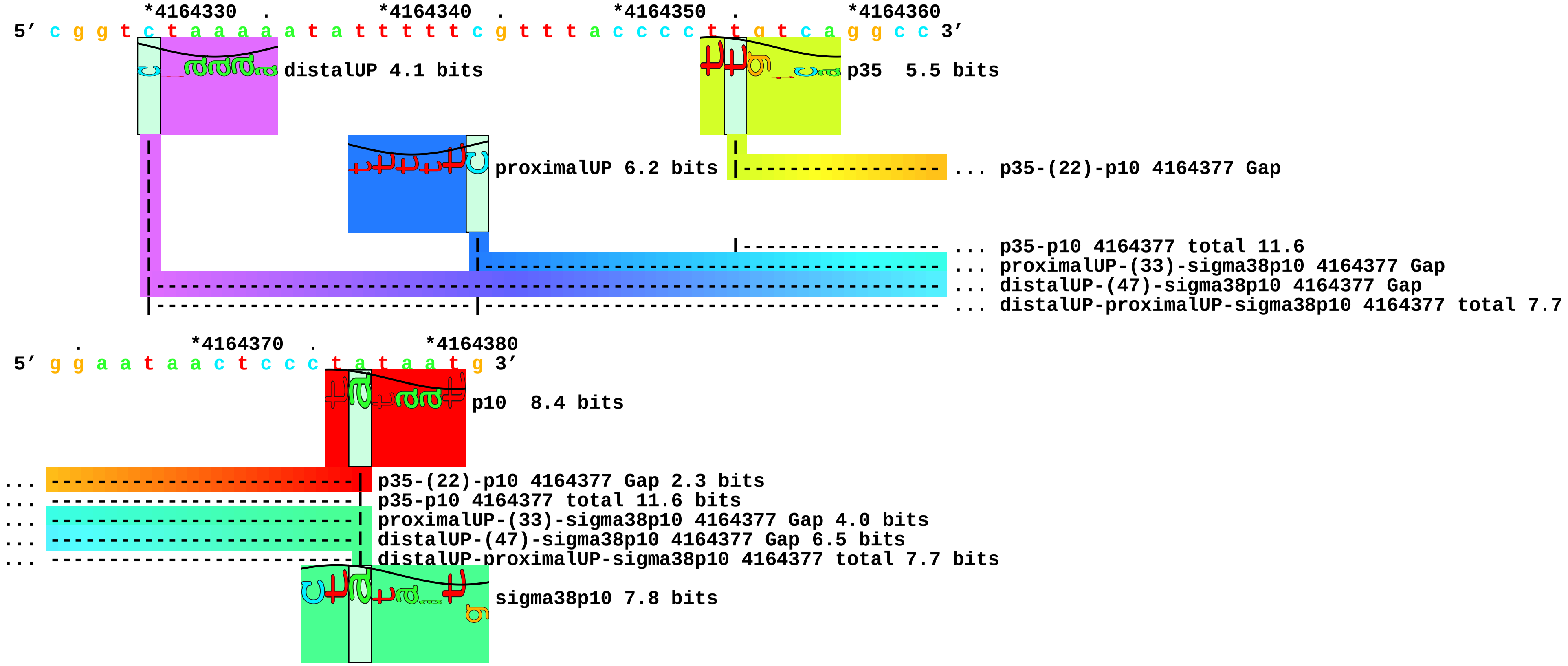

piece 16, NC_000913.i4164330,4164353acaaaaatatttttcgaaaccc, insert acaaaaatatttttcgaaaccc between 4164330 and 4164353, config: linear, directio

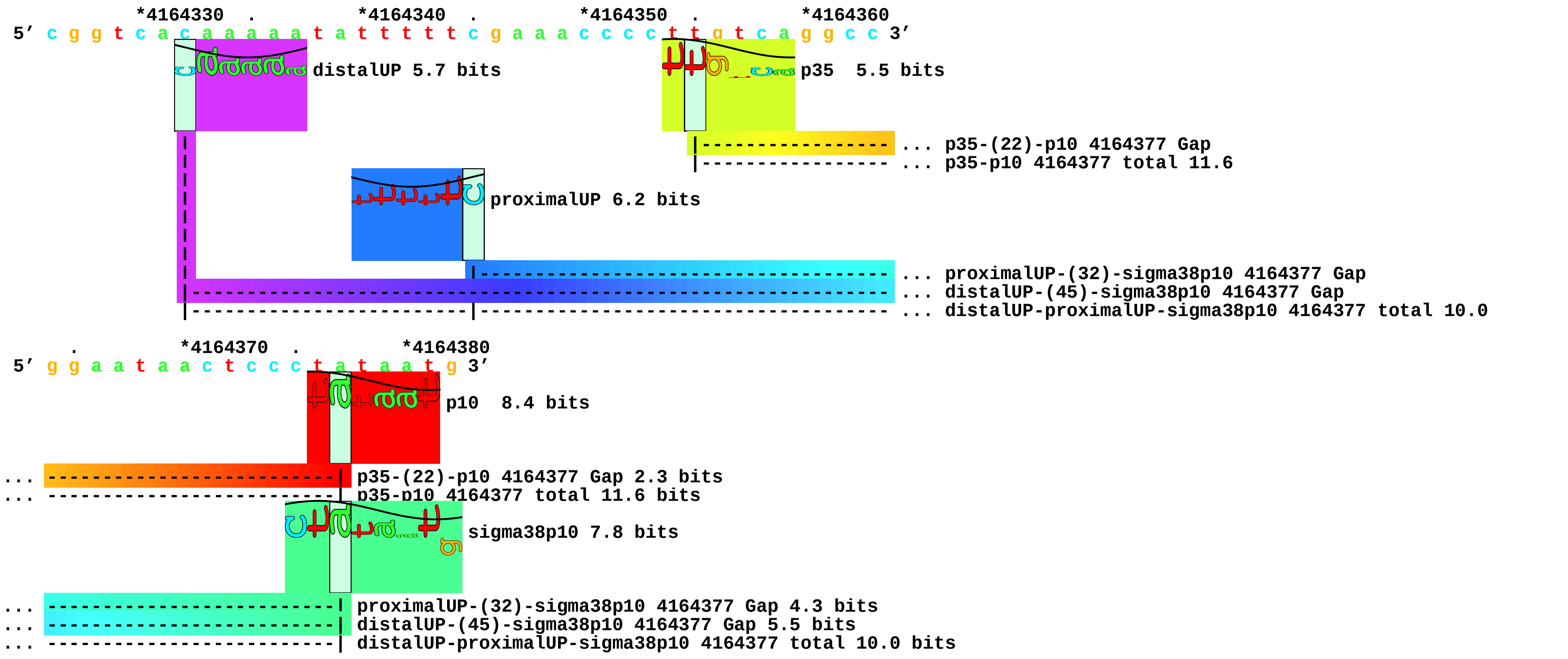

piece 17, NC_000913.i4164330,4164353tcaaaattttttttgcaaagta, insert tcaaaattttttttgcaaagta between 4164330 and 4164353, config: linear, directio

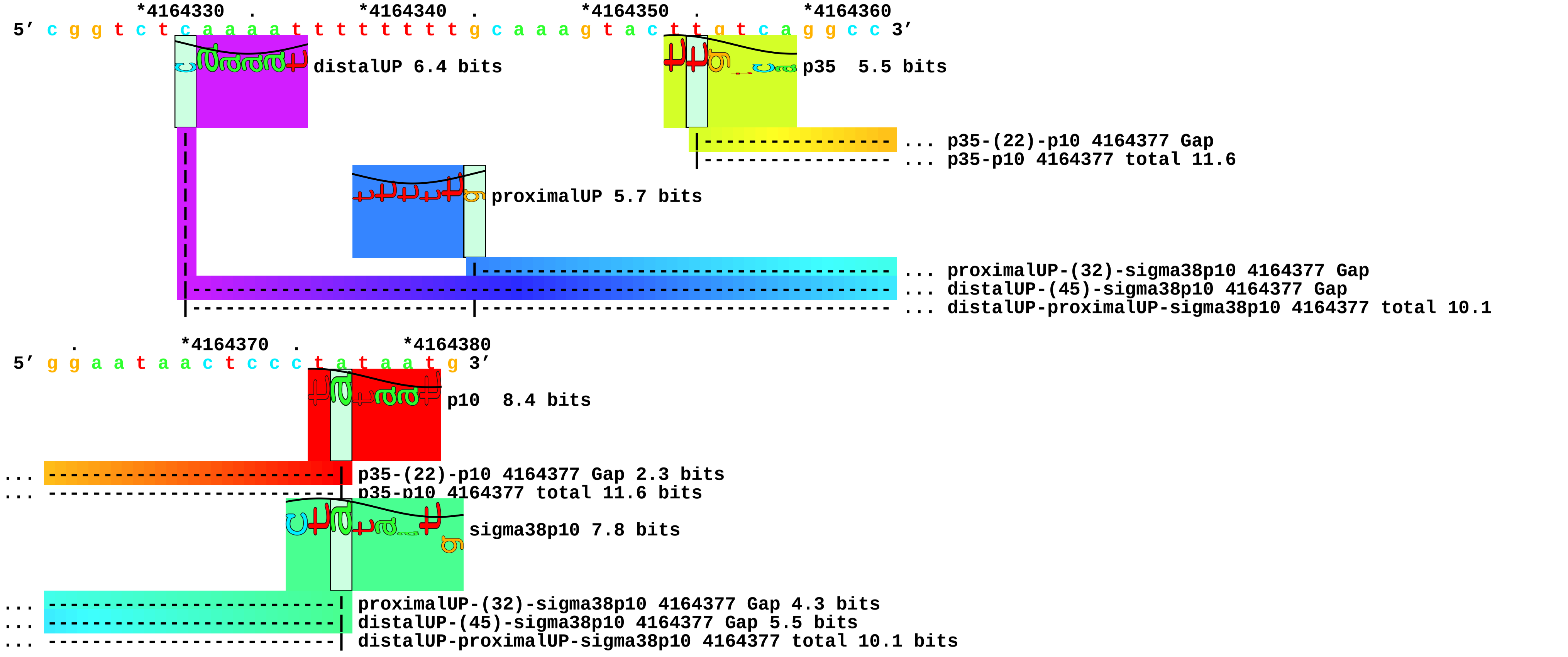

piece 18, NC_000913.i4164330,4164353tgaattttttttttcgtctaccc, insert tgaattttttttttcgtctaccc between 4164330 and 4164353, config: linear, directio

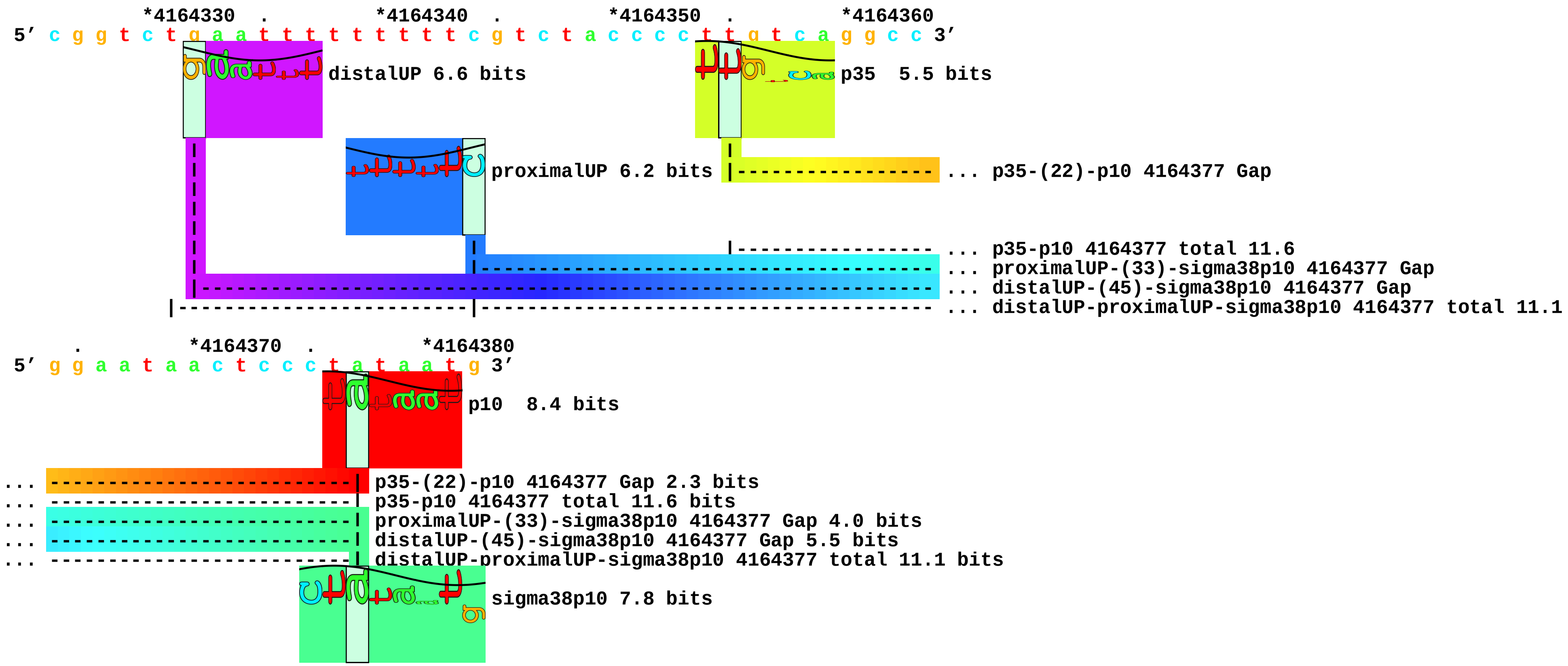

piece 19, NC_000913.i4164330,4164353agaaaaatatttttgaaaacta, insert agaaaaatatttttgaaaacta between 4164330 and 4164353, config: linear, directio

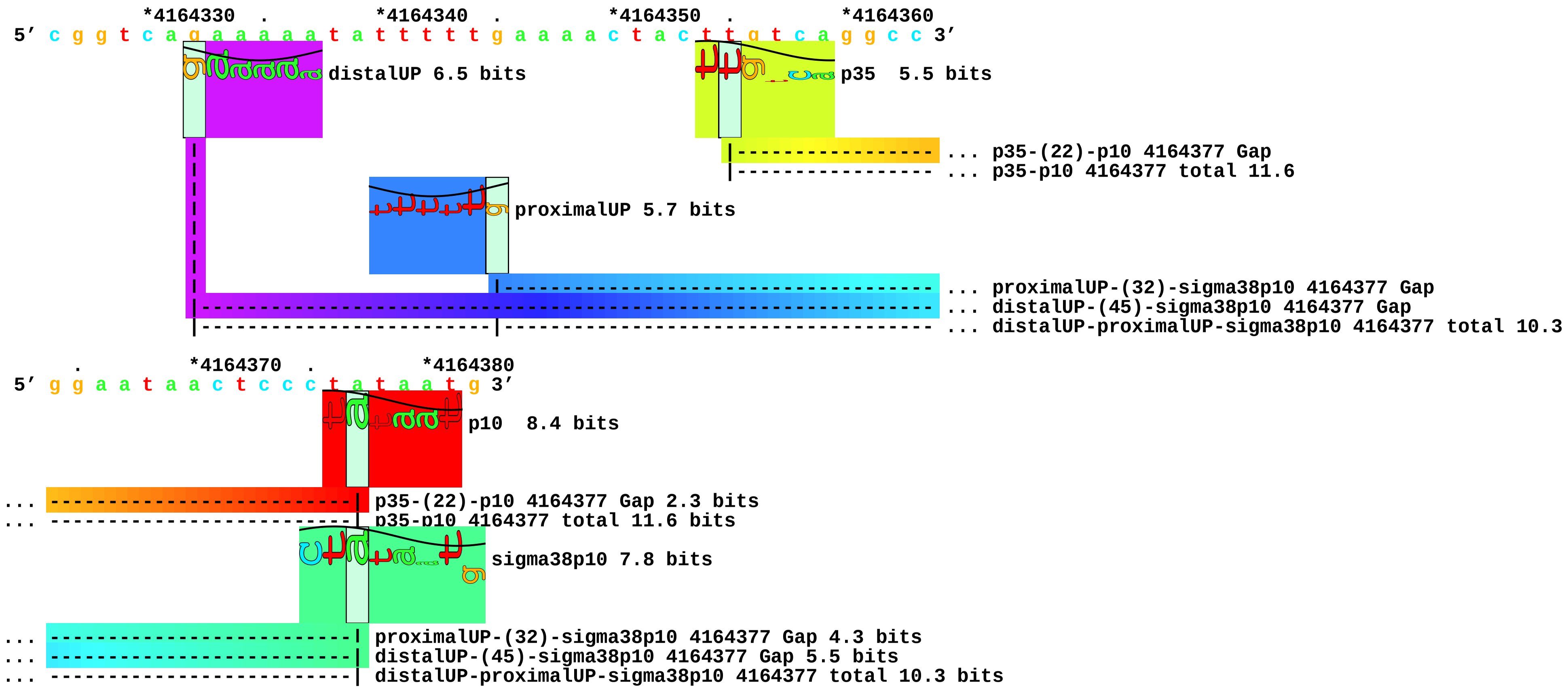

piece 20, NC_000913.i4164330,4164353gcaaaataattgtaaaaagta, insert gcaaaataattgtaaaaagta between 4164330 and 4164353, config: linear, directio

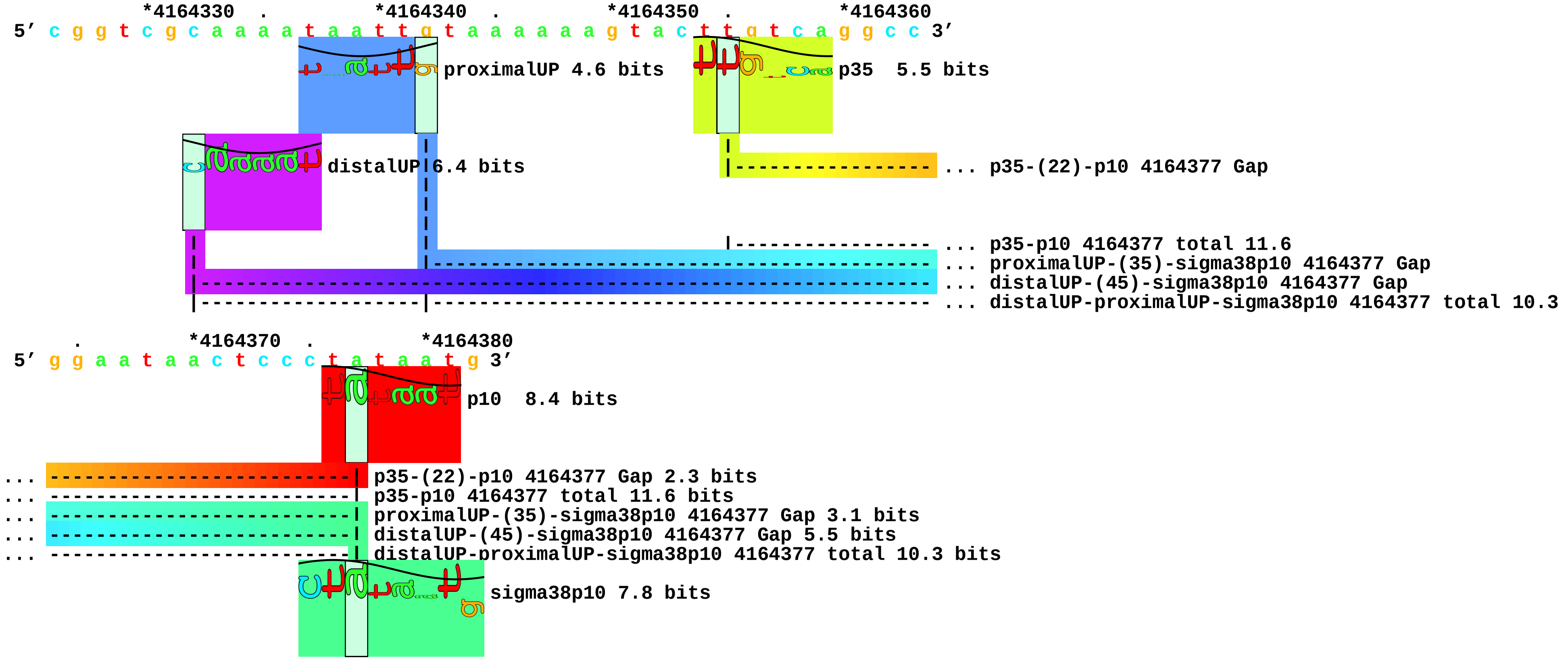

piece 21, NC_000913.i4164330,4164353agaaatttattttaaaaaaggg, insert agaaatttattttaaaaaaggg between 4164330 and 4164353, config: linear, directio

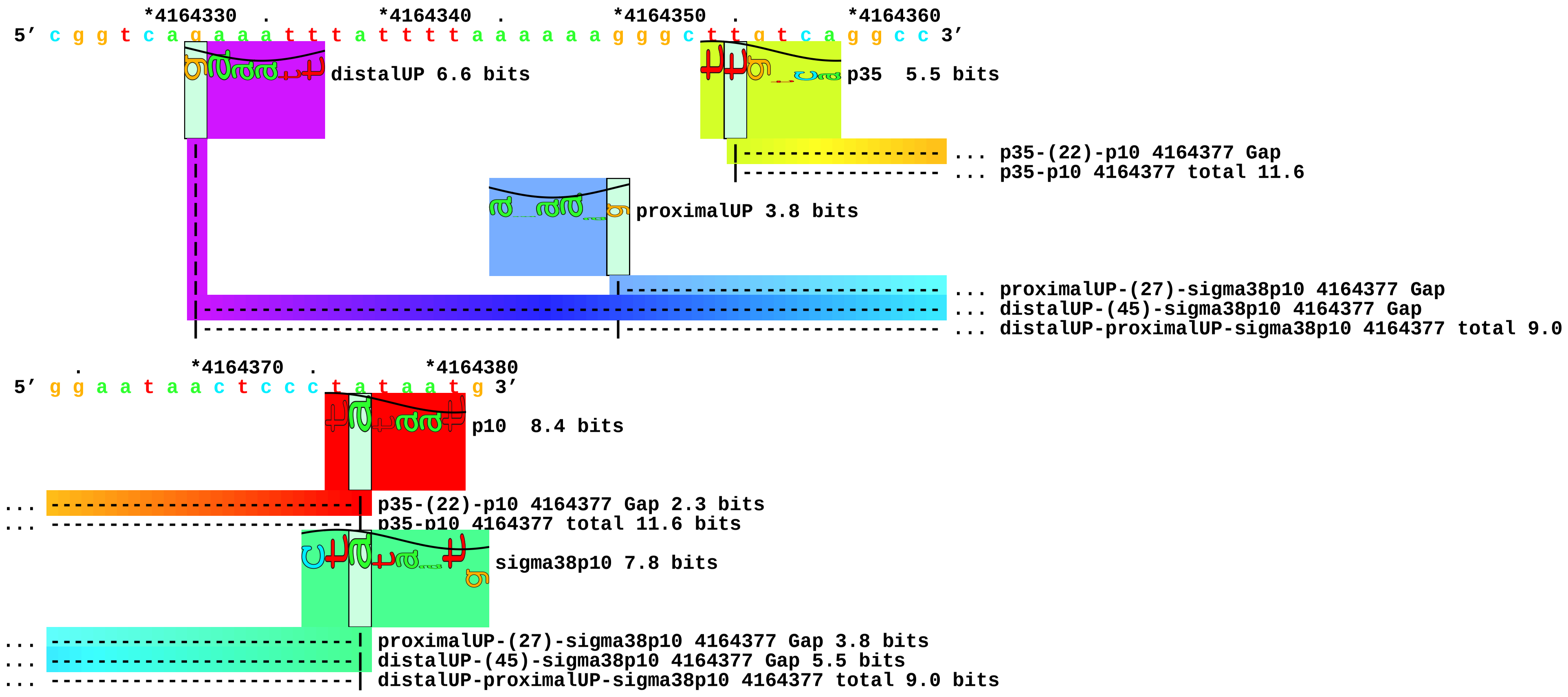

piece 22, NC_000913.i4164330,4164353tgaaaaatatttttgaaaacta, insert tgaaaaatatttttgaaaacta between 4164330 and 4164353, config: linear, directio

```
          *4164330  .         *4164340  .         *4164350  .         *4164360
5' c g g t c t g a a a a a t a t t t t t g a a a a c t a c t t g t c a g g c c 3'
```

distalUP 6.5 bits

p35 5.5 bits

... p35-(22)-p10 4164377 Gap
... p35-p10 4164377 total 11.6

proximalUP 5.7 bits

... proximalUP-(32)-sigma38p10 4164377 Gap
... distalUP-(45)-sigma38p10 4164377 Gap
... distalUP-proximalUP-sigma38p10 4164377 total 10.3

```
    .    *4164370  .         *4164380
5' g g a a t a a c t c c c t a t a a t g 3'
```

p10 8.4 bits

... p35-(22)-p10 4164377 Gap 2.3 bits
... p35-p10 4164377 total 11.6 bits

sigma38p10 7.8 bits

... proximalUP-(32)-sigma38p10 4164377 Gap 4.3 bits
... distalUP-(45)-sigma38p10 4164377 Gap 5.5 bits
... distalUP-proximalUP-sigma38p10 4164377 total 10.3 bits

piece 23, NC_000913.i4164330,4164353taaactattttttcaaaaagga, insert taaactattttttcaaaaagga between 4164330 and 4164353, config: linear, directio

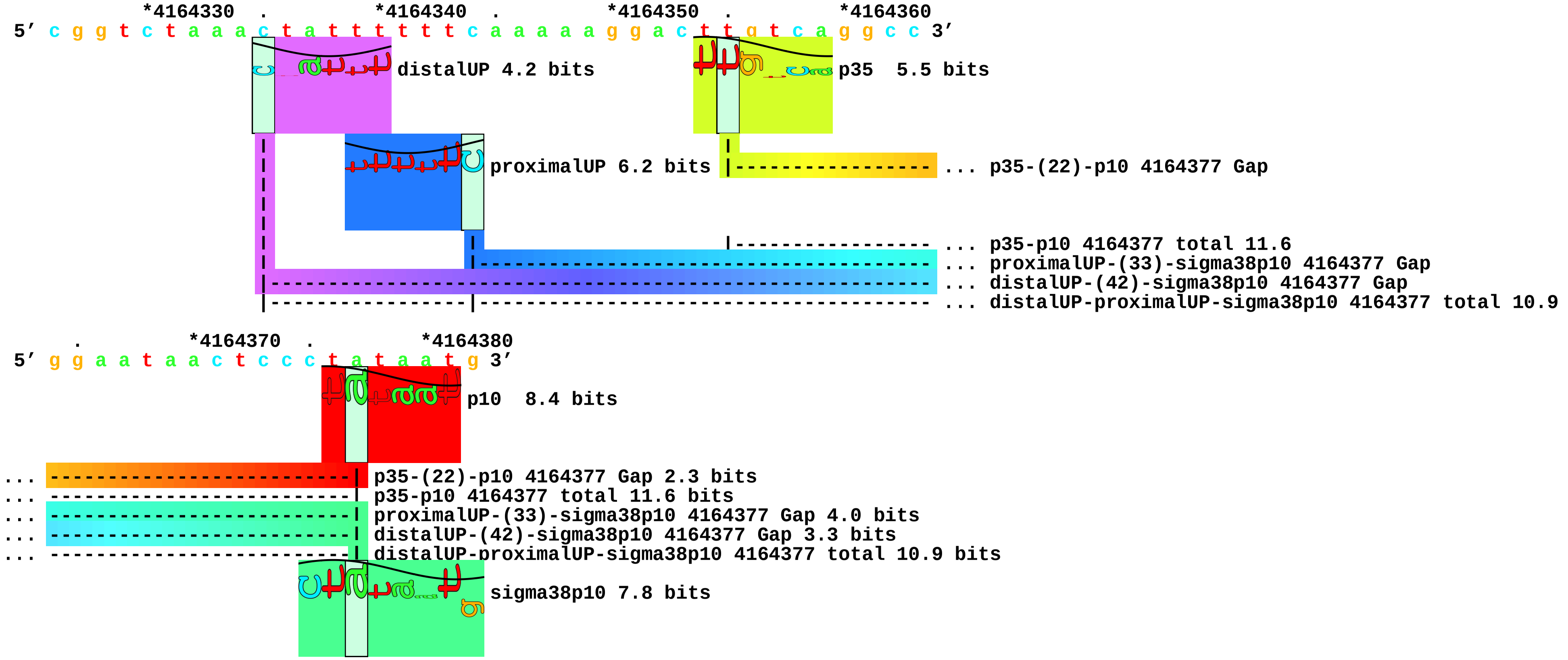

piece 24, NC_000913.i4164330,4164353tgaaatttattttgcgaaaggg, insert tgaaatttattttgcgaaaggg between 4164330 and 4164353, config: linear, directio

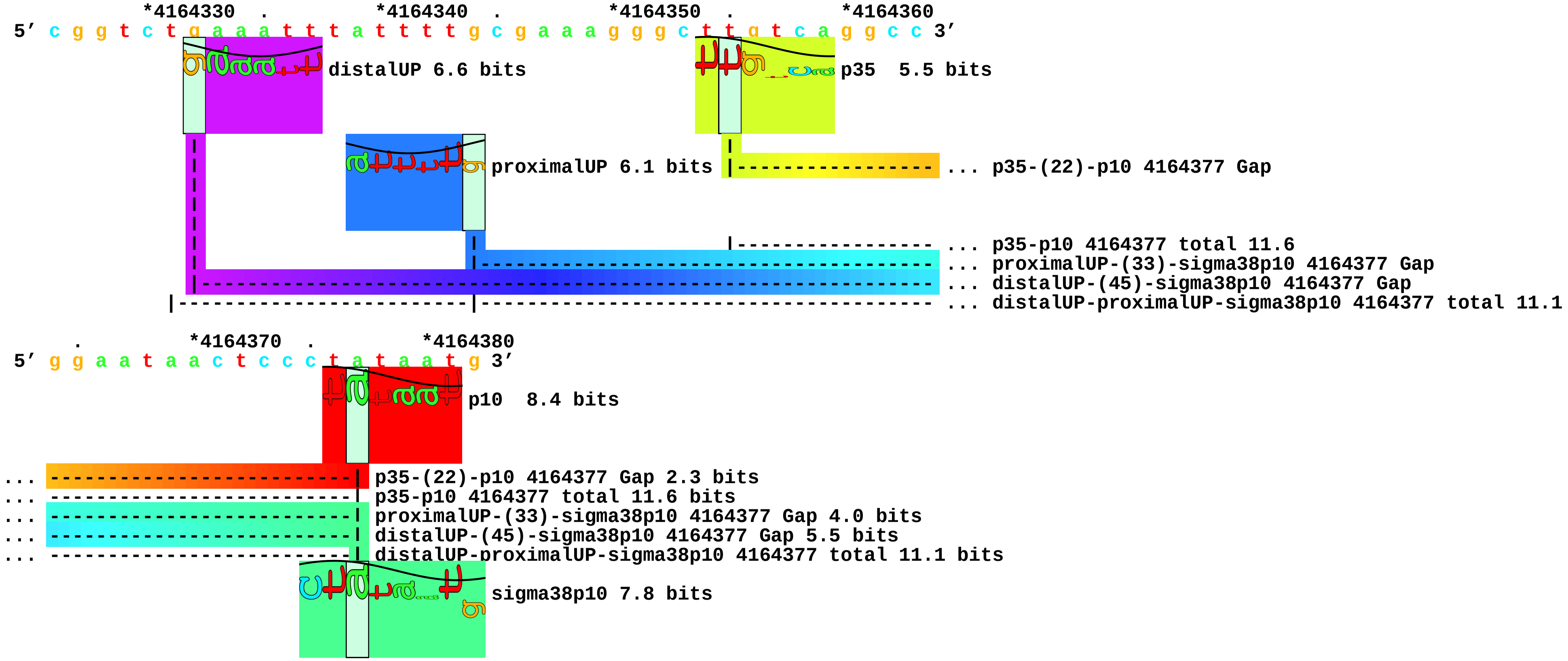

piece 25, NC_000913.i4164330,4164353taaacttttttttttcgaaagtg, insert taaacttttttttttcgaaagtg between 4164330 and 4164353, config: linear, directio

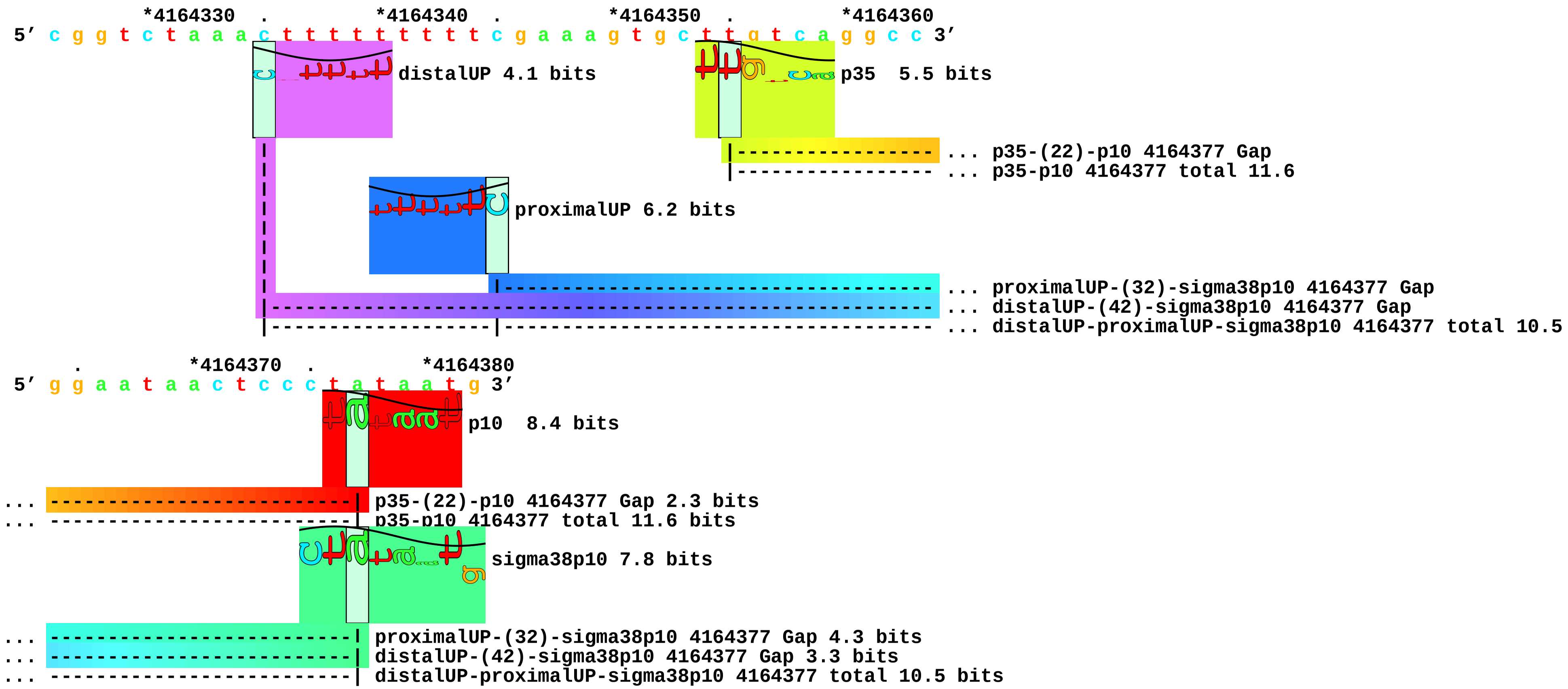

piece 26, NC_000913.i4164330,4164353tgaaatattttttgaaaaccc, insert tgaaatattttttgaaaaccc between 4164330 and 4164353, config: linear, directio

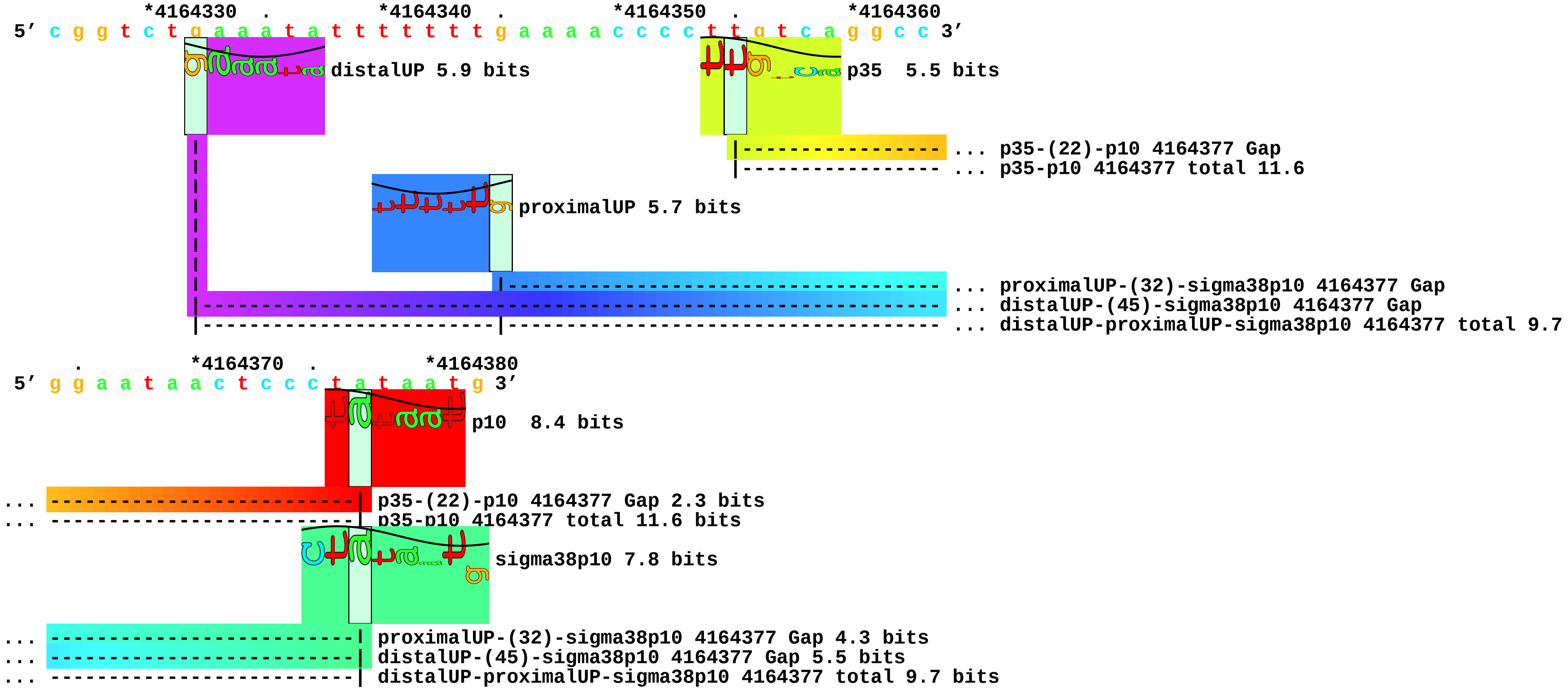

piece 27, NC_000913.i4164330,4164353gcaaaaatatttcgtcaaaccc, insert gcaaaaatatttcgtcaaaccc between 4164330 and 4164353, config: linear, directio

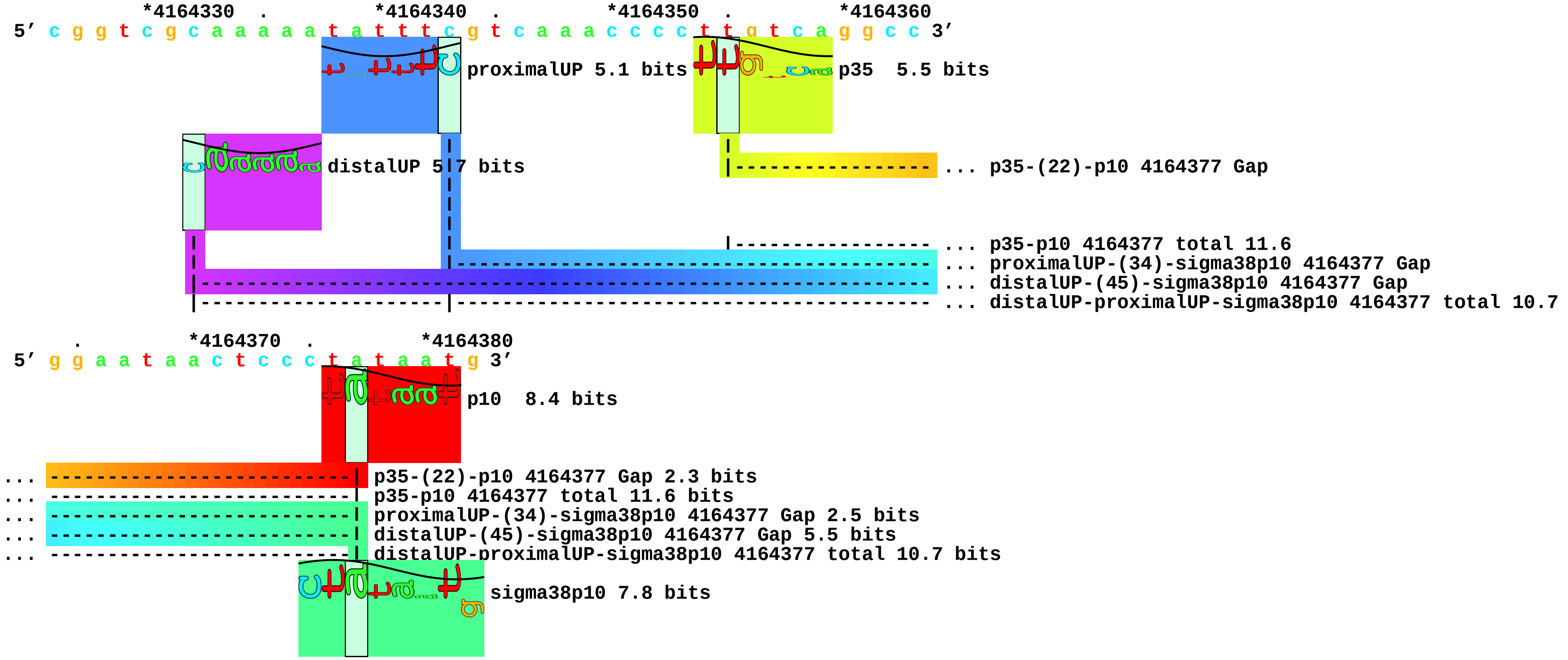

piece 28, NC_000913.i4164330,4164353gaaaaatatttttgataaagta, insert gaaaaatatttttgataaagta between 4164330 and 4164353, config: linear, directio

```
            *4164330  .         *4164340  .         *4164350  .         *4164360
5' c g g t c g a a a a a t a t t t t t g a t a a a g t a c t t g t c a g g c c 3'
```

distalUP 6.5 bits

p35 5.5 bits

proximalUP 5.7 bits

... p35-(22)-p10 4164377 Gap

... p35-p10 4164377 total 11.6
... proximalUP-(33)-sigma38p10 4164377 Gap
... distalUP-(46)-sigma38p10 4164377 Gap
... distalUP-proximalUP-sigma38p10 4164377 total 11.9

```
      .         *4164370  .         *4164380
5' g g a a t a a c t c c c t a t a a t g 3'
```

p10 8.4 bits

... p35-(22)-p10 4164377 Gap 2.3 bits
... p35-p10 4164377 total 11.6 bits
... proximalUP-(33)-sigma38p10 4164377 Gap 4.0 bits
... distalUP-(46)-sigma38p10 4164377 Gap 4.1 bits
... distalUP-proximalUP-sigma38p10 4164377 total 11.9 bits

sigma38p10 7.8 bits

piece 29, NC_000913.i4164330,4164353gcaaaattattttgctaaagta, insert gcaaaattattttgctaaagta between 4164330 and 4164353, config: linear, directio

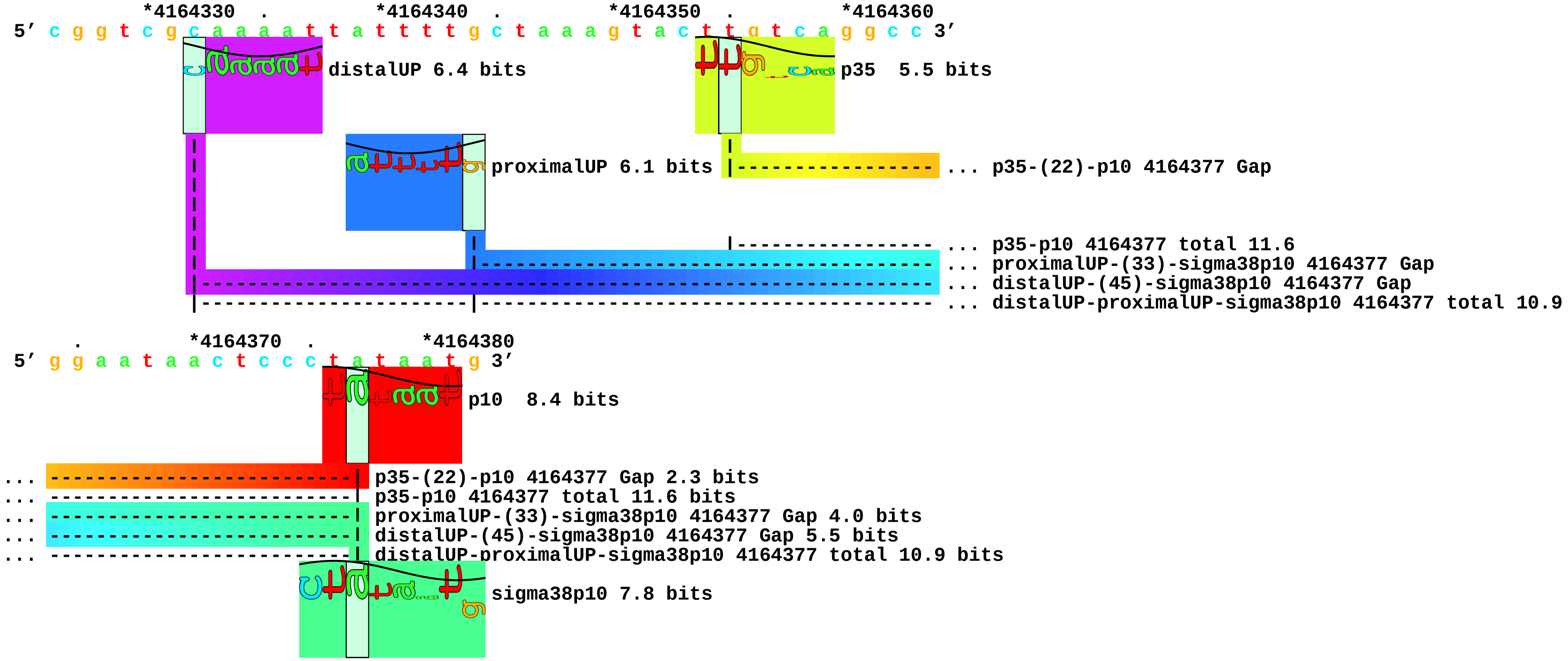

piece 30, NC_000913.i4164330,4164353gaaaatatatttttcaaaagta, insert gaaaatatatttttcaaaagta between 4164330 and 4164353, config: linear, directio

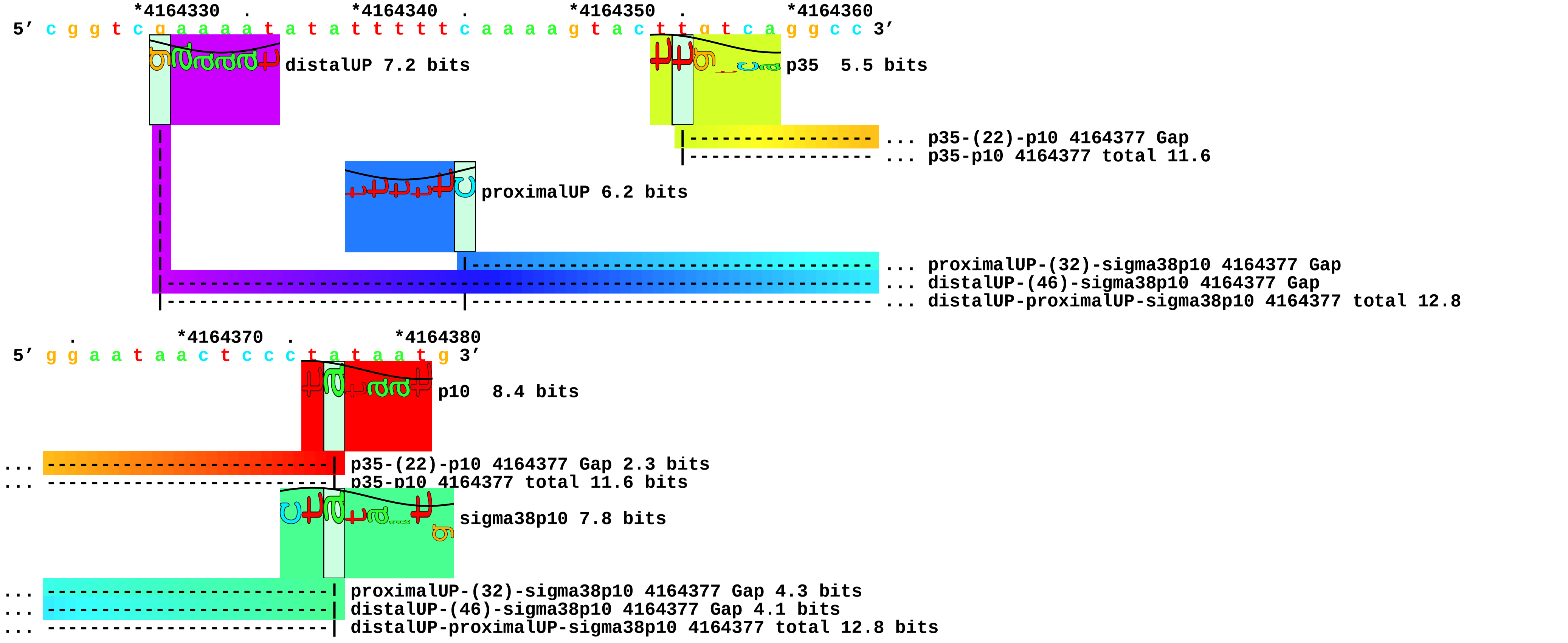
