## Supplement_for_sigma38_paper-directory for "*Escherichia coli σ*^38^ promoters use two UP elements instead of a −35 element: resolution of a paradox and discovery that *σ*^38^ transcribes ribosomal promoters": Kolb.Ishihama1995-fig2-map.pdf

piece 1, NC_000913.3, galEp1, config: linear, direction: -, begin: 792151, end: 792071

```
   *792150   .         *792140   .         *792130   .         *792120   .         *792110   .         *792100   .         *792090   .         *792080   .
5' t t g t g t a a a c g a t t c c a c t a a t t t a t t c c a t g t c a c a c t t t t c g c a t c t t t g t t a t g c t a t g g t t a t t t c a t a c c a t a a g c 3'
3' a a c a c a t t t g c t a a g g t g a t t a a a t a a g g t a c a g t g t g a a a a g c g t a g a a a c a a t a c g a t a c c a a t a a a g t a t g g t a t t c g 5'
```

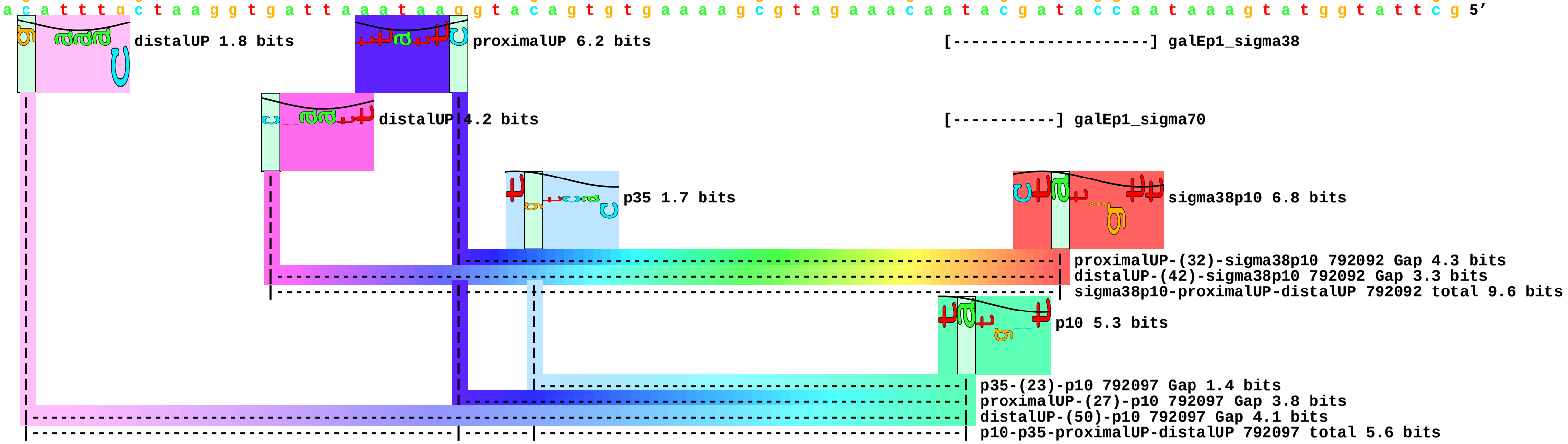
