## Supplement_for_sigma38_paper-directory for "*Escherichia coli σ*^38^ promoters use two UP elements instead of a −35 element: resolution of a paradox and discovery that *σ*^38^ transcribes ribosomal promoters": Lacour.Landini2002-map.pdf

piece 1, NC_000913, aidB, config: linear, direction: +, begin: 4412219, end: 4412268

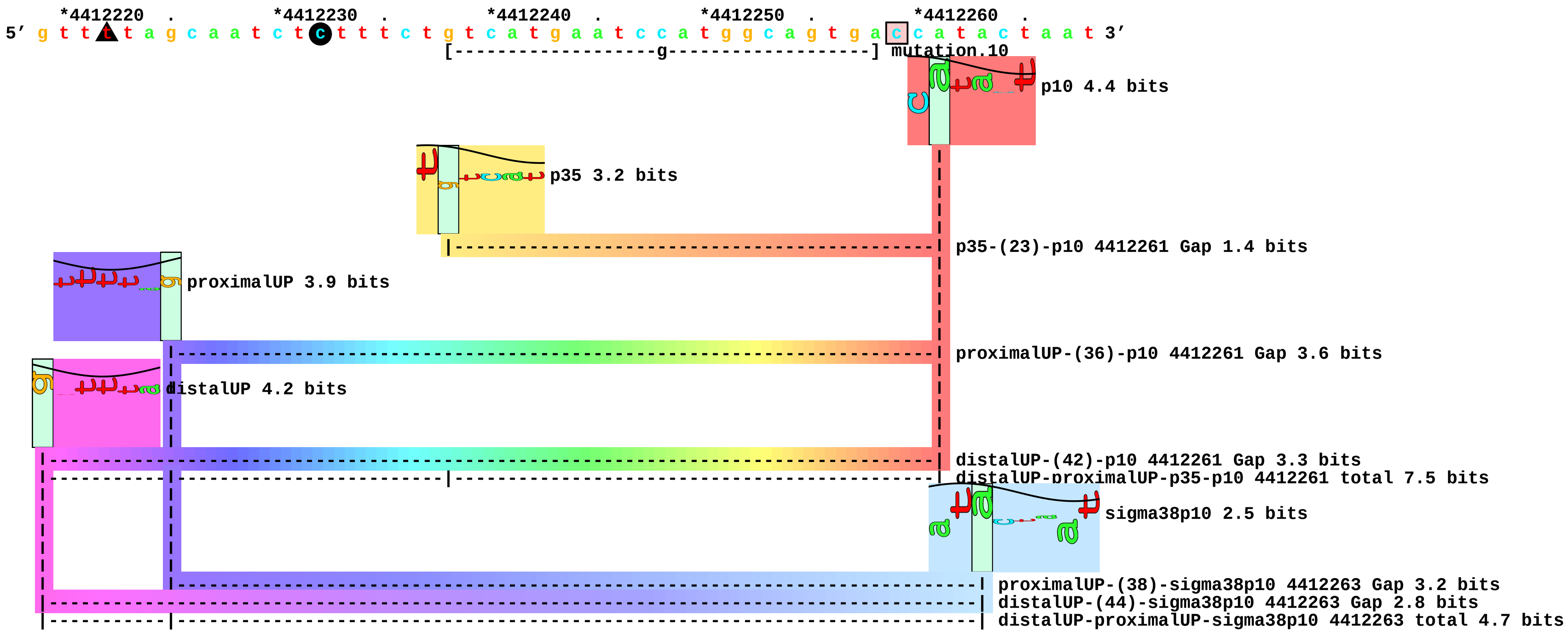

piece 2, NC_000913.d4412259,4412259, aidB, config: linear, direction: +, begin: 4412219, end: 4412267

```
     *4412220  .         *4412230  .         *4412240  .         *4412250  .         *4412260  .
5' g t t t t a g c a a t c t c t t t c t g t c a t g a a t c c a t g g c a g t g a c a t a c t a a t 3'
```

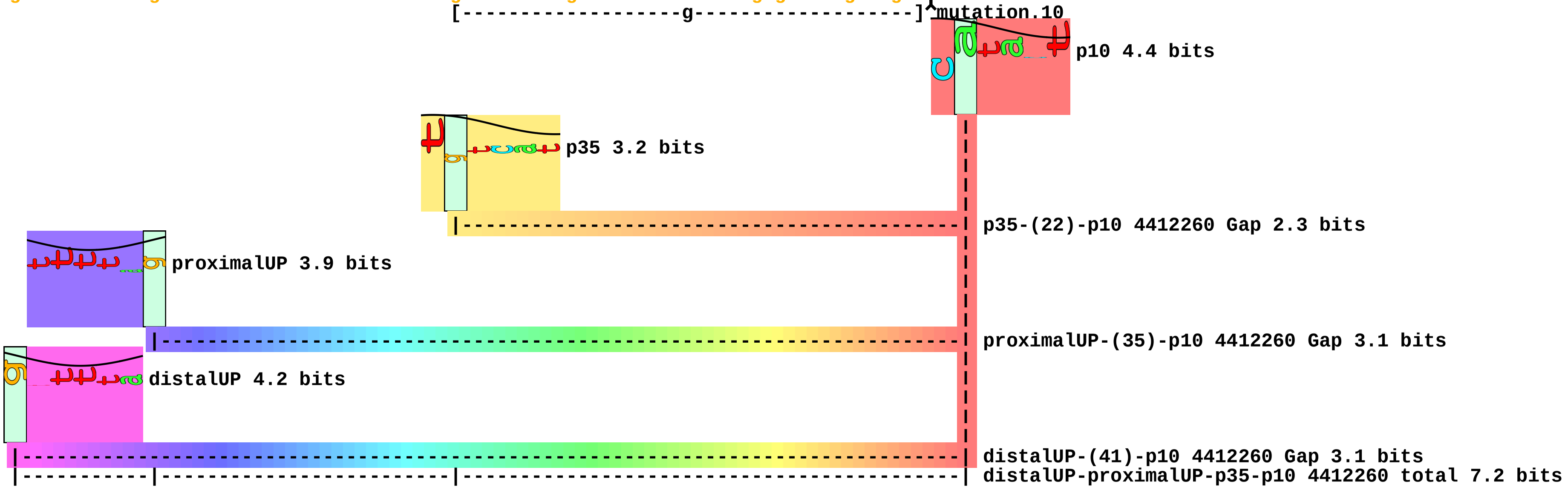
