## Supplement_for_sigma38_paper-directory for "*Escherichia coli σ*^38^ promoters use two UP elements instead of a −35 element: resolution of a paradox and discovery that *σ*^38^ transcribes ribosomal promoters": new22map.pdf

piece 1, NC_000913, adap, config: linear, direction: -, begin: 2310529, end: 2310459

```
          .         *2310520  .         *2310510  .         *2310500  .         *2310490  .         *2310480  .         *2310470  .         *2310460
5' c c a g g g c a a c a g c g c t g a a a a c g a a a a c a a c t c c t g g a c c c g t g t g g c a t t c g c a g g t c t g a a a t t c c a g g 3'
3' g g t c c c g t t g t c g c g a c t t t t g c t t t t g t t g a g g a c c t g g g c a c a c c g t a a g c g t c c a g a c t t t a a g g t c c 5'
```

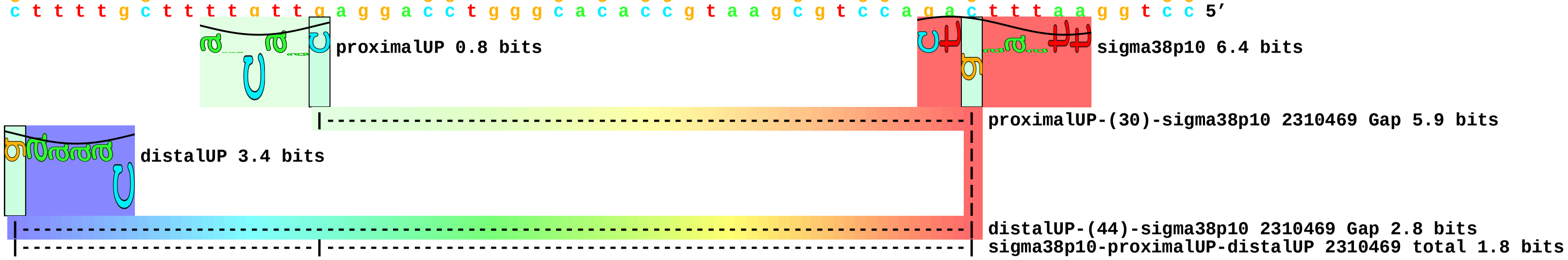

piece 2, NC_000913, alkAp, config: linear, direction: -, begin: 2147818, end: 2147748

```
              .         *2147810  .         *2147800  .         *2147790  .         *2147780  .         *2147770  .         *2147760  .         *2147750
5’ g a g a t c g c a c t g t c a g c a t t a t c g c t g c t g c t t t g c g g c c t t c a g g c a g a a c c g c g t t a t a t t a t t t t g g t 3’
3’ c t c t a g c g t g a c a g t c g t a a t a g c g a c g a c g a a a c g c c g g a a g t c c g t c t t g g c g c a a t a t a a t a a a a c c a 5’
```

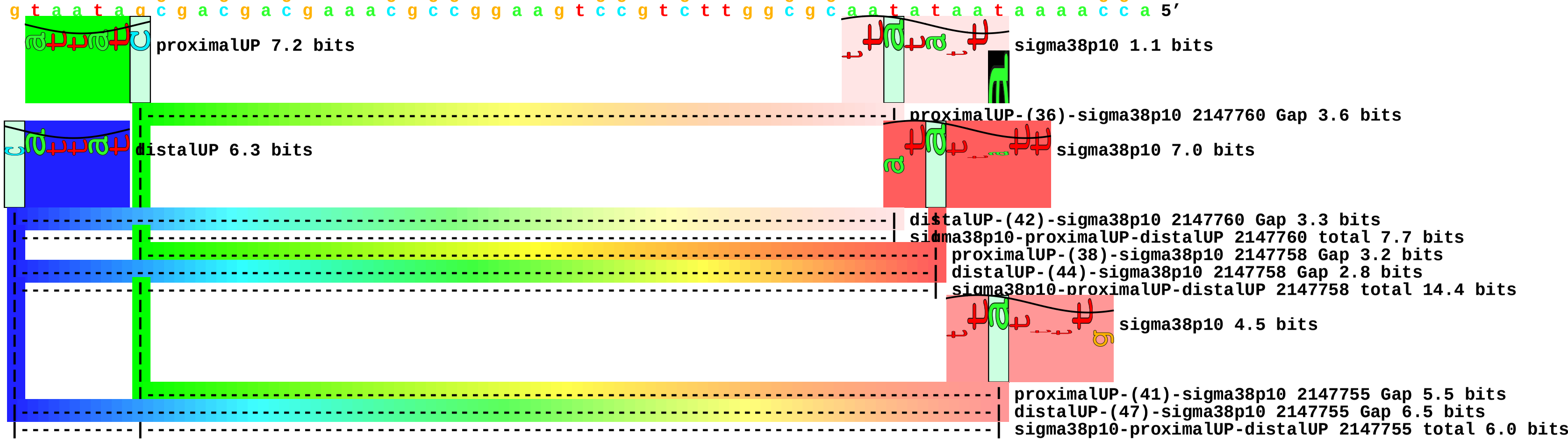

piece 3, NC_000913, appYp2, config: linear, direction: +, begin: 583592, end: 583662

```
           .         *583600   .         *583610   .         *583620   .         *583630   .         *583640   .         *583650   .         *583660
5' c a t c t c a t t a t t t t g a g a a a a t a a t c g g c g t c a c a g a t g g a a t a a a c a a a a c a a t t g a c t g a t a a t g t t t a 3'
3' g t a g a g t a a t a a a a c t c t t t t a t t a g c c g c a g t g t c t a c c t t a t t t g t t t t g t t a a c t g a c t a t t a c a a a t 5'
```

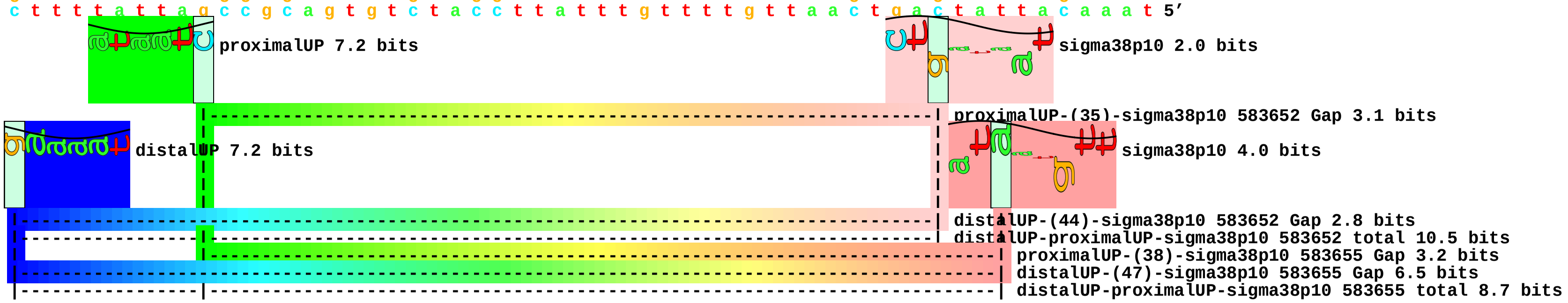

piece 4, NC_000913, araFp2, config: linear, direction: -, begin: 1986275, end: 1986205

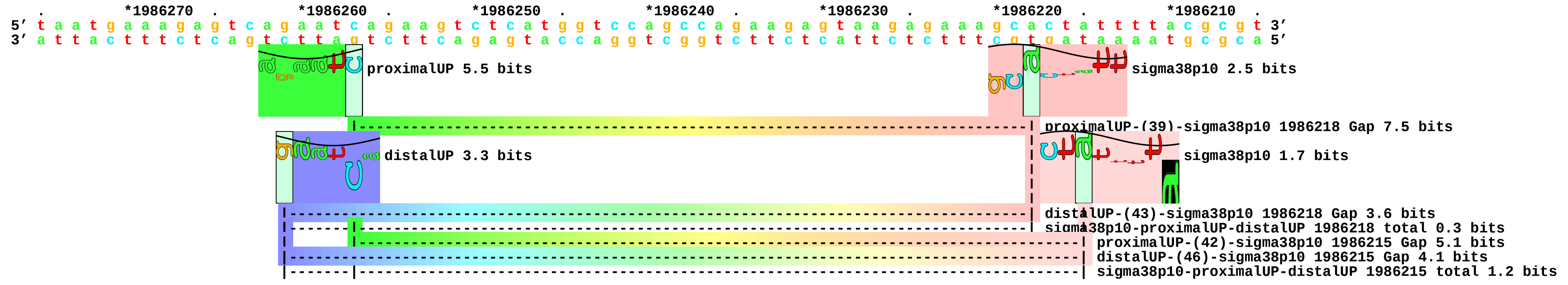

piece 5, NC_000913, bsmAp, config: linear, direction: -, begin: 4417014, end: 4416944

```
             *4417010  .         *4417000  .         *4416990  .         *4416980  .         *4416970  .         *4416960  .         *4416950  .
5' a c c t g t t c a a a a c g c c t g g c g g c t c c c t g t a t c a c a g c g g c g a c t c c c a c t a c t c t a a c t a t t a t g c g a a g 3'
3' t g g a c a a g t t t t g c g g a c c g c c g a g g g a c a t a g t g t c g c c g c t g a g g g t g a t g a g a t t g a t a a t a c g c t t c 5'
```

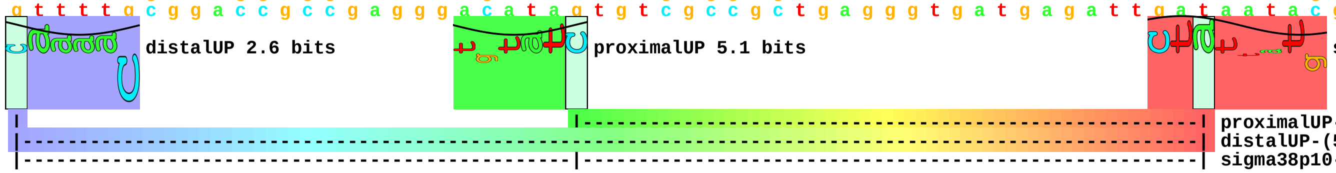

piece 6, NC_000913, crrpI, config: linear, direction: +, begin: 2535461, end: 2535531

```
          .         *2535470  .        *2535480  .        *2535490  .        *2535500  .        *2535510  .        *2535520  .        *2535530
5' t g g a g g g a t t a g c a c a g a a a a a g g a t t t g a t a t t t g t g c g c a a c g g t g a t g a a c a t a c c t g c t a t g a a g c g 3'
3' a c c t c c c t a a t c g t g t c t t t t t c c t a a a c t a t a a a c a c g c g t t g c c a c t a c t t g t a t g g a c g a t a c t t c g c 5'
```

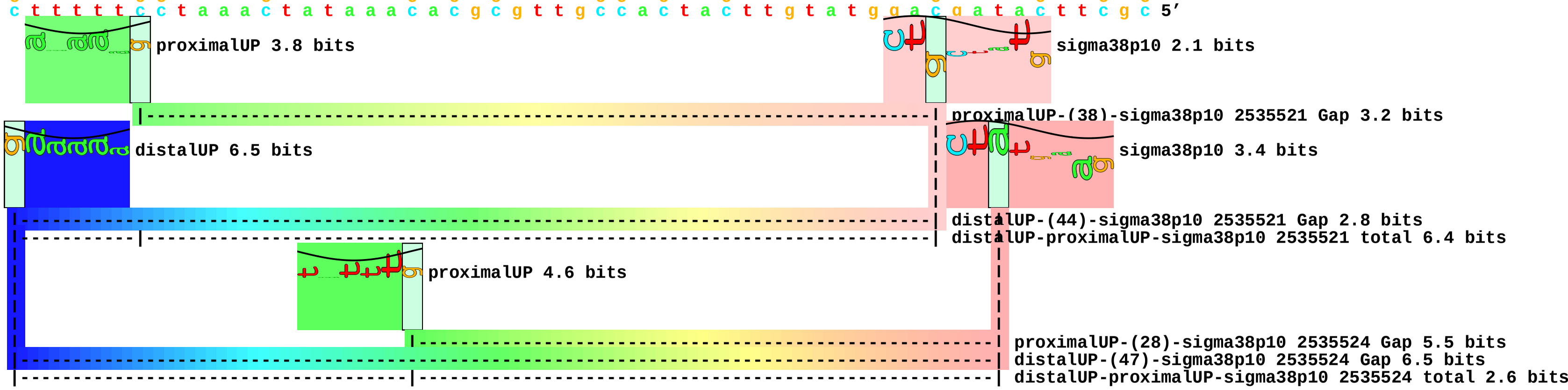

piece 7, NC_000913, csgBp, config: linear, direction: +, begin: 1103775, end: 1103845

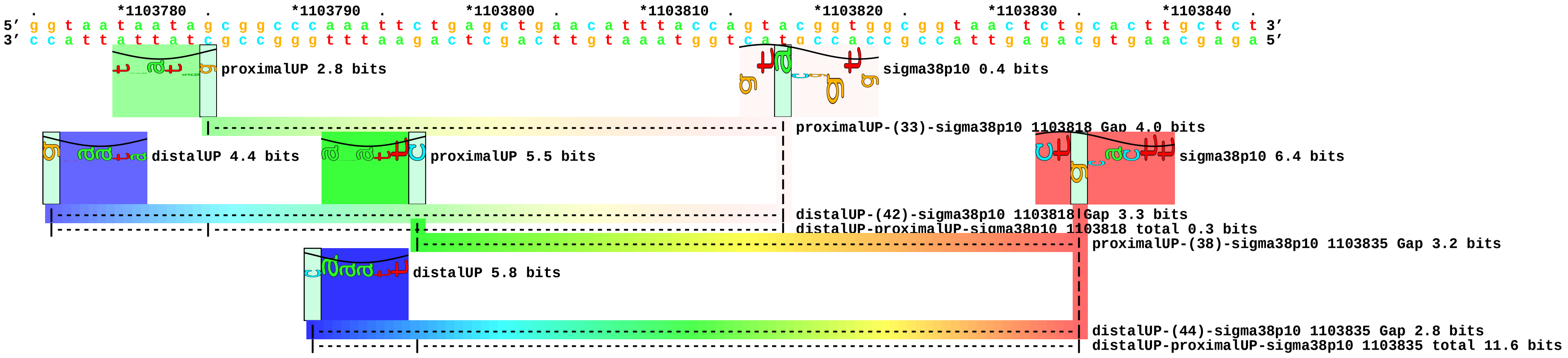

piece 8, NC_000913, csgDp1, config: linear, direction: -, begin: 1103458, end: 1103388

```
        .         *1103450  .         *1103440  .         *1103430  .         *1103420  .         *1103410  .         *1103400  .         *1103390
5' c t g a t c a a t a t a t g c a a g g t t a t a a t c t c c t g t c t g g t c a a t c t t t g c c c g g t t g c t a c t a c c t t c t t g c g 3'
3' g a c t a g t t a t a t a c g t t c c a a t a t t a g a g g a c a g a c c a g t t a g a a a c g g g c c a a c g a t g a t g g a a g a a c g c 5'
```

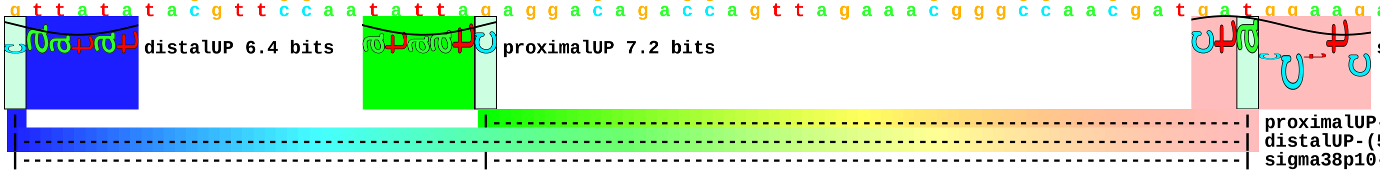


proximalUP-(34)-sigma38p10 1103398 Gap 2.5 bits
distalUP-(55)-sigma38p10 1103398 Gap 4.9 bits
sigma38p10-proximalUP-distalUP 1103398 total 9.1 bits

piece 9, NC_000913, csgDp2, config: linear, direction: -, begin: 1103458, end: 1103388

```
        .         *1103450  .         *1103440  .         *1103430  .         *1103420  .         *1103410  .         *1103400  .         *1103390
5' c t g a t c a a t a t a t g c a a g g t t a t a a t c t c c t g t c t g g t c a a t c t t t g c c c g g t t g c t a c t a c c t t c t t g c g 3'
3' g a c t a g t t a t a t a c g t t c c a a t a t t a g a g g a c a g a c c a g t t a g a a a c g g g c c a a c g a t g a t g g a a g a a c g c 5'
```

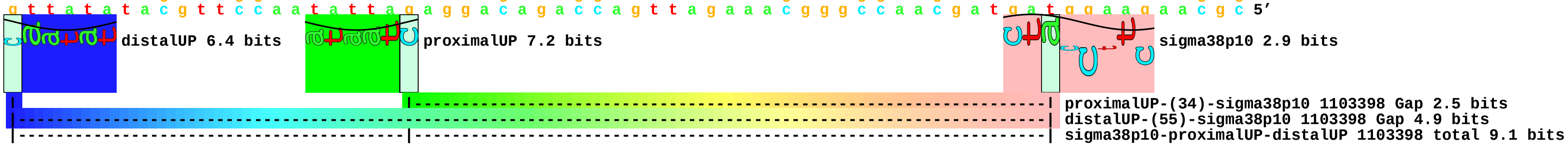

piece 10, NC_000913, fumCp, config: linear, direction: -, begin: 1686809, end: 1686739

```
              .          *1686800  .          *1686790  .          *1686780  .          *1686770  .          *1686760  .          *1686750  .          *1686740
5' g a g c a g a g t c g a t t t a t c a c t c t c a a t c a c a t c a c g c a g t g a a a c g a t a t c t c c g g c g g c a t t c t g c a c t c 3'
3' c t c g t c t c a g c t a a a t a g t g a g a g t t a g t g t a g t g c g t c a c t t t g c t a t a g a g g c c g c c g t a a g a c g t g a g 5'
```

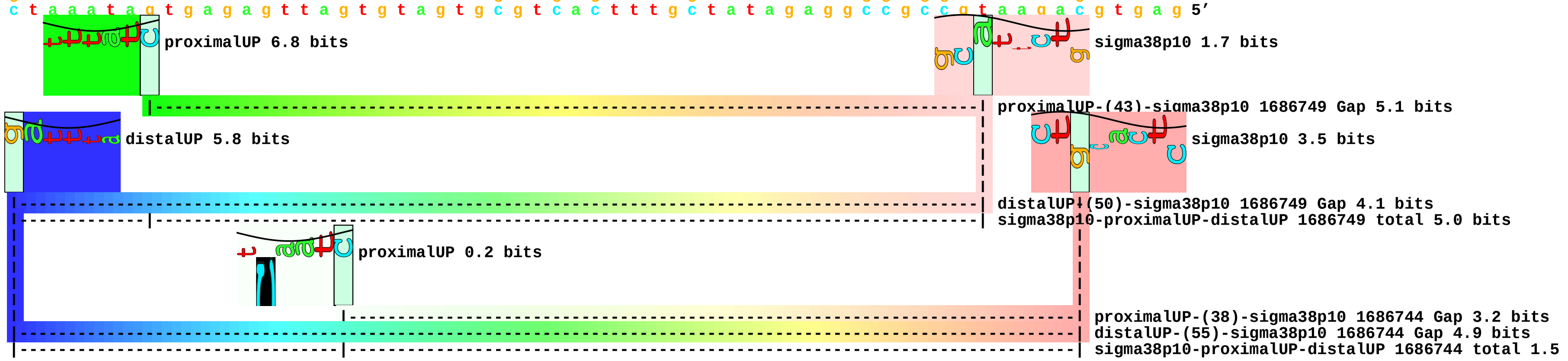

piece 11, NC_000913, gpmMp2, config: linear, direction: +, begin: 3785113, end: 3785183

```
          .         *3785120  .         *3785130  .         *3785140  .         *3785150  .         *3785160  .         *3785170  .         *3785180
5' a c g c a a a a c a c g c t c a a t c a a c t c a a t a a a c a g a t t g a t g a g a t g a a c g c g t c g a t t g c c a a a c t g g a g c a 3'
3' t g c g t t t t g t g c g a g t t a g t t g a g t t a t t t g t c t a a c t a c t c t a c t t g c g c a g c t a a c g g t t t g a c c t c g t 5'
```

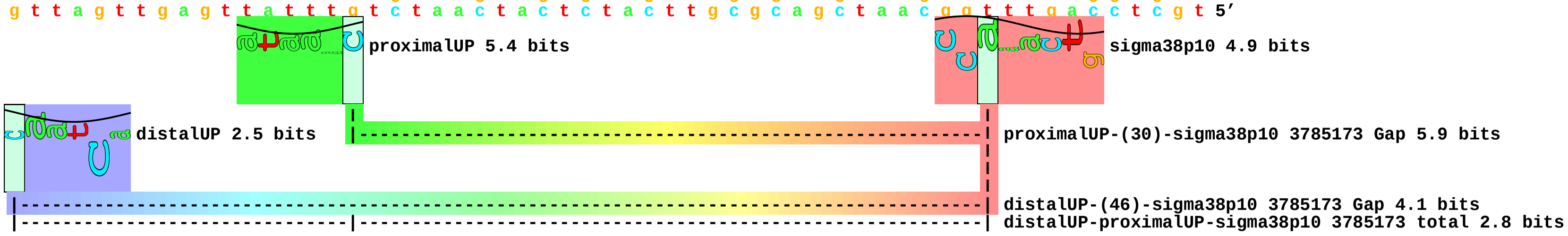

piece 12, NC_000913, insC-5p, config: linear, direction: +, begin: 3185923, end: 3185993

```
         .         *3185930  .         *3185940  .         *3185950  .         *3185960  .         *3185970  .         *3185980  .         *3185990
5' t a t g c a c c a c a t c a g g g c g a t a a t t c g c a a a a t a t a a g t t c c t a t g g g c a g g c t g g g g t t a a t c t t g g g g c 3'
3' a t a c g t g g t g t a g t c c c g c t a t t a a g c g t t t t a t a t t c a a g g a t a c c c g t c c g a c c c c a a t t a g a a c c c c g 5'
```

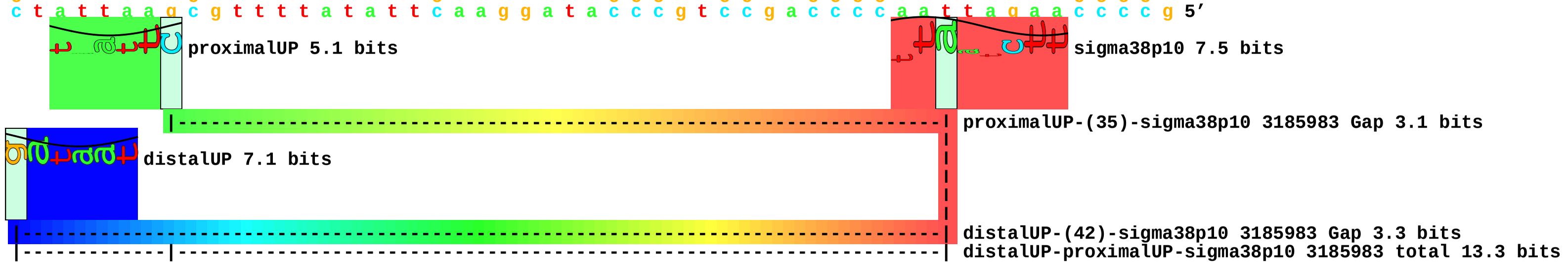

piece 13, NC_000913, insN-1p, config: linear, direction: +, begin: 270146, end: 270216

```
             *270150   .         *270160   .         *270170   .         *270180   .         *270190   .         *270200   .         *270210   .
5' t t g a t g c t a a t a a c c g a g c c a a c a g a a t g g c g a a a a g g c c a a a a c c g t g c t t a c t g g a t c a a a a t t t a c c a 3'
3' a a c t a c g a t t a t t g g c t c g g t t g t c t t a c c g c t t t t c c g g t t t t g g c a c g a a t g a c c t a g t t t t a a a t g g t 5'
```

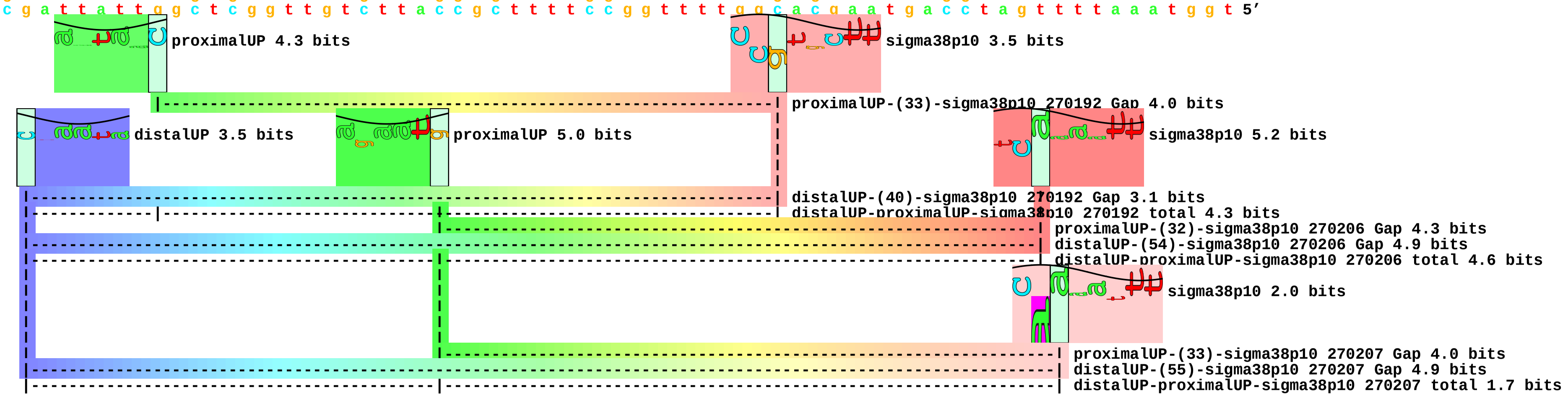

piece 14, NC_000913, mnmGp, config: linear, direction: -, begin: 3925852, end: 3925782

```
        *3925850  .         *3925840  .         *3925830  .         *3925820  .         *3925810  .         *3925800  .         *3925790  .
5' t g a c g g g g t g c t c c a g t c a t c a t a a t c c g g t g c g c g c a c g t c g t g g c g a t g a c c a t c g c t c a g c t t g c c g c 3'
3' a c t g c c c c a c g a g g t c a g t a g t a t t a g g c c a c g c g c g t g c a g c a c c g c t a c t g g t a g c g a g t c g a a c g g c g 5'
```

proximalUP 7.2 bits

sigma38p10 1.8 bits

proximalUP-(34)-sigma38p10 3925792 Gap 2.5 bits

distalUP 6.3 bits

distalUP-(40)-sigma38p10 3925792 Gap 3.1 bits

sigma38p10-proximalUP-distalUP 3925792 total 9.8 bits

piece 15, NC_000913, mutSp, config: linear, direction: +, begin: 2856879, end: 2856949

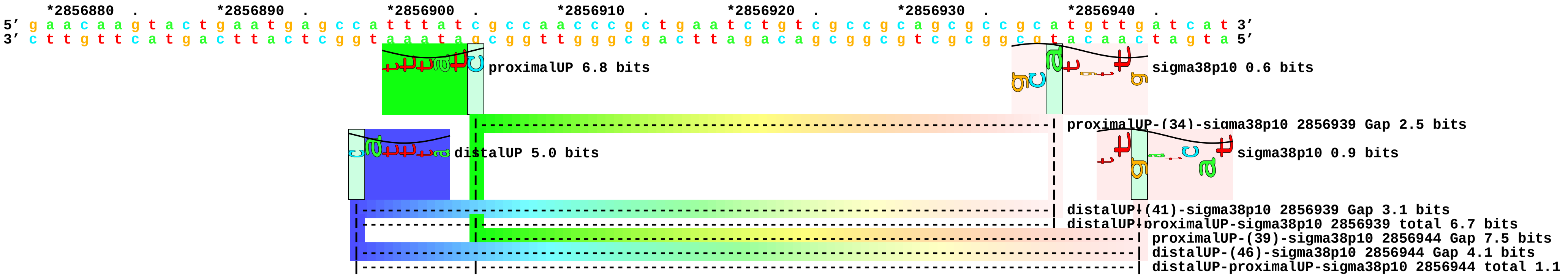

piece 16, NC_000913, ompFp, config: linear, direction: -, begin: 987350, end: 987280

```
   *987350   .         *987340   .         *987330   .         *987320   .         *987310   .         *987300   .         *987290   .         *987280
5' t t c t t c g c t g a a c g c g t a g a t a a a g a t g c c g t t t c a c g t c t g g a a c g c t t c a t t g a a g c c g a t t t t g c g c a 3'
3' a a g a a g c g a c t t g c g c a t c t a t t t c t a c g g c a a a g t g c a g a c c t t g c g a a g t a a c t t c g g c t a a a a c g c g t 5'
```

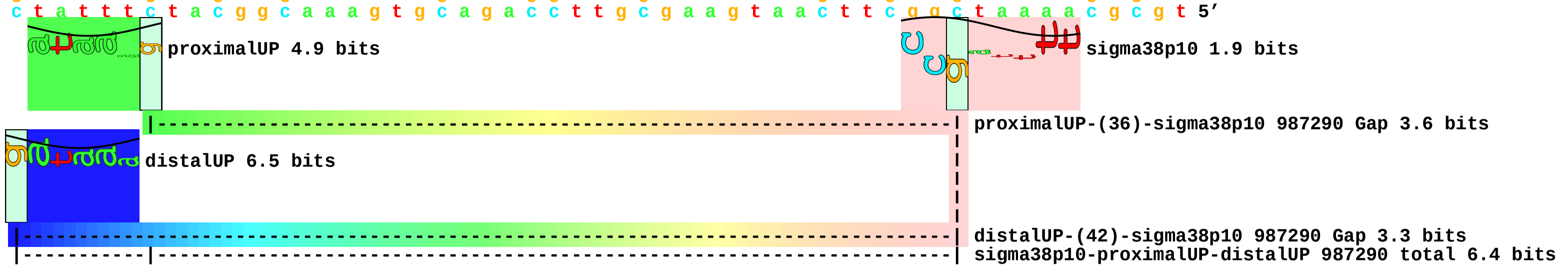

piece 17, NC_000913, ppkp, config: linear, direction: +, begin: 2622854, end: 2622924

```
     .         *2622860  .         *2622870  .         *2622880  .         *2622890  .         *2622900  .         *2622910  .         *2622920
5' a t t t t t g a a a a t g g c g g c g a t a a a a a g g t c t a c c t t t c t t c c g c c g a c t g g a t g a c g c g c a a t a t t g a t t a 3'
3' t a a a a a c t t t t a c c g c c g c t a t t t t t c c a g a t g g a a a g a a g g c g g c t g a c c t a c t g c g c g t t a t a a c t a a t 5'
```

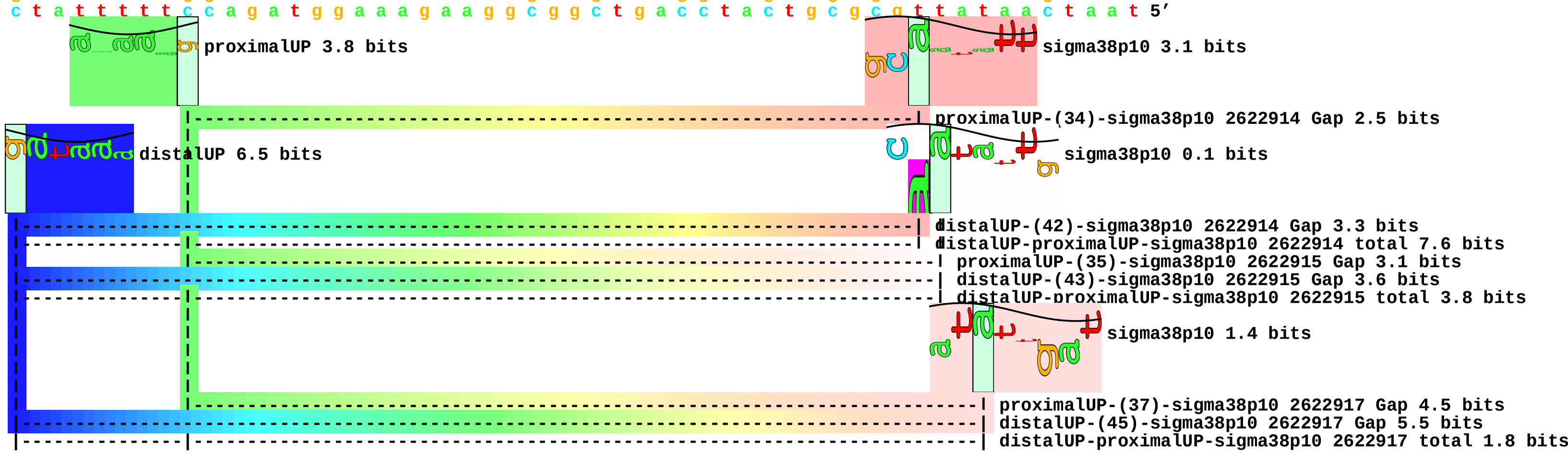

piece 18, NC_000913, puuCp, config: linear, direction: +, begin: 1362490, end: 1362560

```
   *1362490  .        *1362500  .        *1362510  .        *1362520  .        *1362530  .        *1362540  .        *1362550  .        *1362560
5' c g c g a c a t c g a c g t g a t c g a a a a a a g c t a c g g c a t g g a c a c c g c c c g t a t g c t c g g c a g c a t g a t g t t c g a 3'
3' g c g c t g t a g c t g c a c t a g c t t t t t t c g a t g c c g t a c c t g t g g c g g g c a t a c g a g c c g t c g t a c t a c a a g c t 5'
```

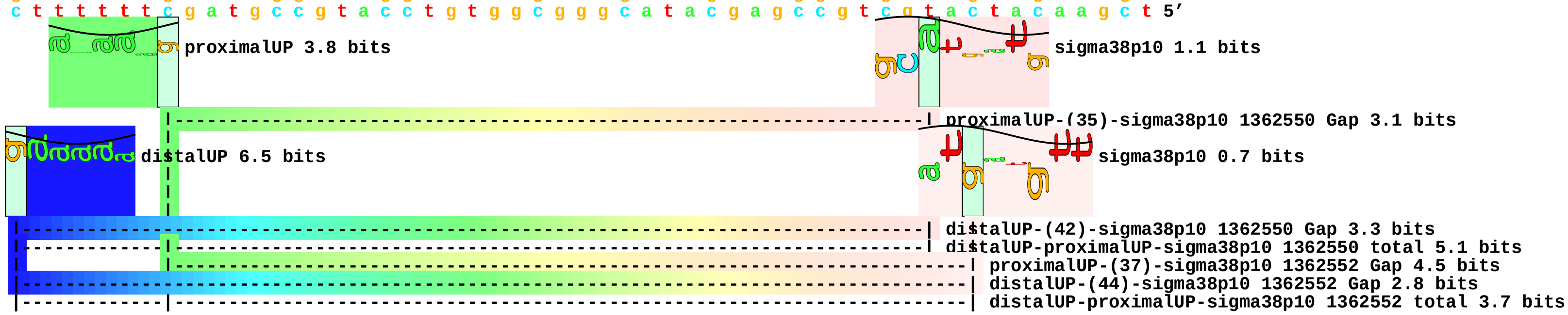

piece 19, NC_000913, pykFp2, config: linear, direction: +, begin: 1755397, end: 1755467

```
          *1755400  .         *1755410  .         *1755420  .         *1755430  .         *1755440  .         *1755450  .         *1755460  .
5' t a c t t g t a a c g c t a c a t g g a g a t t a a c t c a a t c t a g a g g g t a t t a a t a a t g a a a g c t a c t a a a c t g g t a c t 3'
3' a t g a a c a t t g c g a t g t a c c t c t a a t t g a g t t a g a t c t c c c a t a a t t a t t a c t t t c g a t g a t t t g a c c a t g a 5'
```

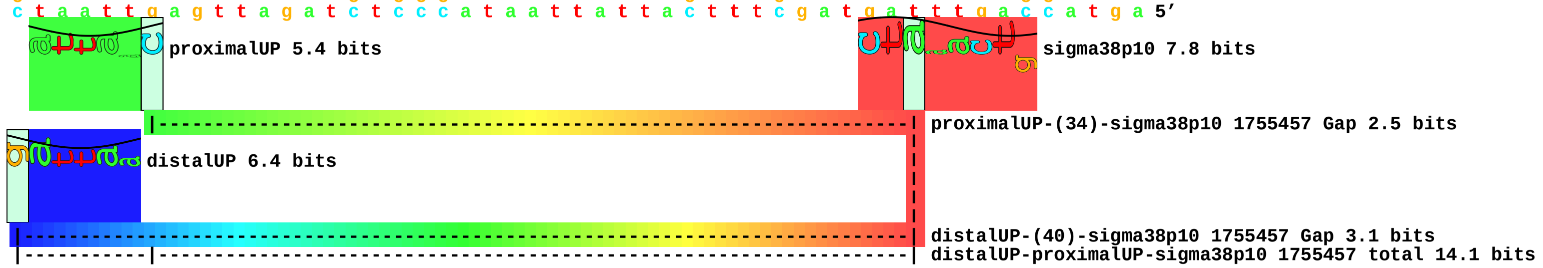

piece 20, NC_000913, yffOp, config: linear, direction: +, begin: 2561893, end: 2561963

```
         .         *2561900  .         *2561910  .         *2561920  .         *2561930  .         *2561940  .         *2561950  .         *2561960
5’ a a a g a g a a t a g c g a t g a a a a g a t t a a c c t a t t g a a c a a g a t g g g g g a t c t g g c t g g t c g t a t c g t t g c a g a 3’
3’ t t t c t c t t a t c g c t a c t t t t c t a a t t g g a t a a c t t g t t c t a c c c c c t a g a c c g a c c a g c a t a g c a a c g t c t 5’
```

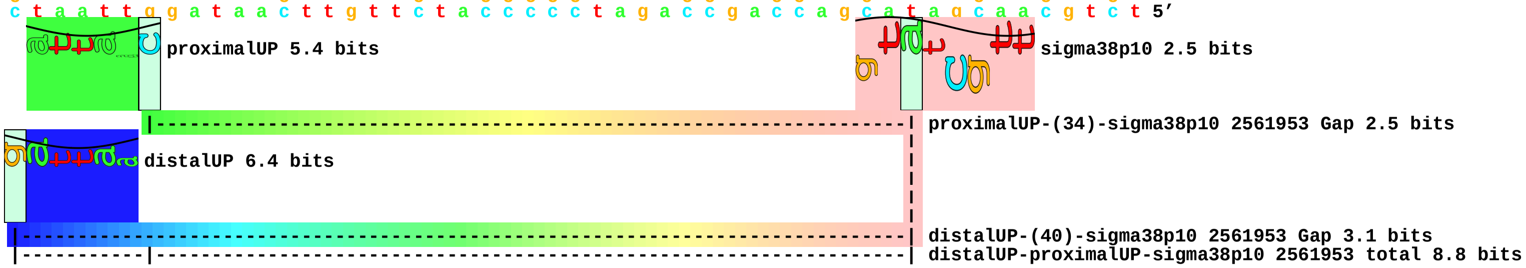

piece 21, NC_000913, yjcCp, config: linear, direction: +, begin: 4275296, end: 4275366

```
         *4275300  .       *4275310  .       *4275320  .       *4275330  .       *4275340  .       *4275350  .       *4275360  .
5' g t a c c a c t t t g a a t a g c c t g a t t t t t t t g c g a c t a c a t c a a t g t t a a g c g g c t g g t c a a t a t g c t c g t c a a 3'
3' c a t g g t g a a a c t t a t c g g a c t a a a a a a a c g c t g a t g t a g t t a c a a t t c g c c g a c c a g t t a t a c g a g c a g t t 5'
```

proximalUP 5.7 bits

sigma38p10 4.6 bits

proximalUP-(32)-sigma38p10 4275356 Gap 4.3 bits

distalUP 6.5 bits

distalUP-(41)-sigma38p10 4275356 Gap 3.1 bits

distalUP-proximalUP-sigma38p10 4275356 total 9.4 bits

piece 22, NC_000913, yjiYp, config: linear, direction: -, begin: 4591531, end: 4591461

```
     *4591530  .         *4591520  .         *4591510  .         *4591500  .         *4591490  .         *4591480  .         *4591470  .
5' t t t t g c g c c g a t c a t t a c t g g t t t t a t t g t t g a t a c c a c c c a c t c a t t c c g t c t g g c a c t a a t c a t c t g c g 3'
3' a a a a c g c g g c t a g t a a t g a c c a a a a t a a c a a c t a t g g t g g g t g a g t a a g g c a g a c c g t g a t t a g t a g a c g c 5'
```

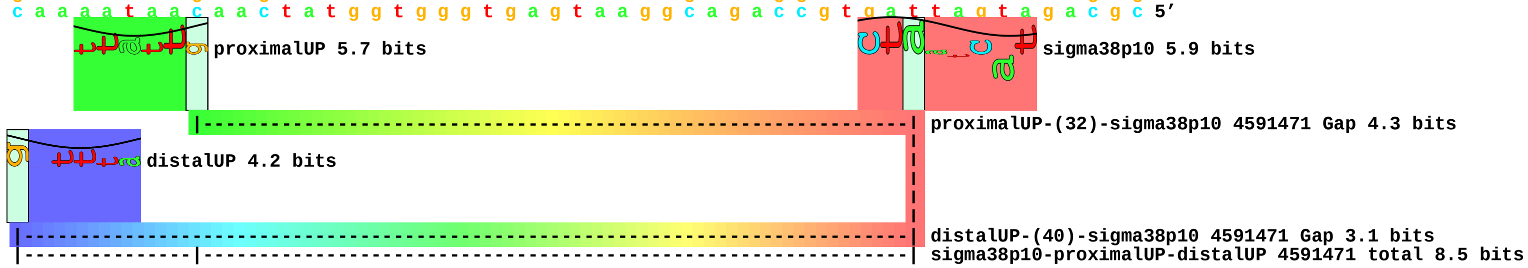
