## Supplement_for_sigma38_paper-directory for "*Escherichia coli σ*^38^ promoters use two UP elements instead of a −35 element: resolution of a paradox and discovery that *σ*^38^ transcribes ribosomal promoters": Peano.Landini2015-Table1-neg-map.pdf

piece 1, NC_000913, hepA, config: linear, direction: +, begin: 63350, end: 63588

*63350 . *63360 . *63370 . *63380 .
5′ a t g g a a a g g g c g c t a t g g t a c t g g a t g g c a a a g c a t t c 3′
T timestamp:_2018Sep20_11-35-24

*63390 . *63400 . *63410 . *63420 .
5′ g t c a c g c a t c a a a a a t g g t a t c t g g c g a a c t c t t t t t t t t 3′
proximalUP 5.6 bits ... proximalUP
... proximalUP-(38)
distalUP 6.4 bits distalUP 4.1 bits
... distalUP-(44)-s
... distalUP-proxin
... distalUP-(47)-s
... distalUP-proxin

*63430 . *63440 . *63450 . *63460
5′ t g c t c a a a a t a g c c c a a g t t g c c c g g t c a t a a g t g t a g 3′
... proximalUP 5.7 bits sigma38p10 2.6 bits ... sigma38p10
... proximalUP-(38)-sigma38p10 63441 Gap 3.2 bits
... proximalUP-(38)
... distalUP-(44)-sigma38p10 63441 Gap 2.8 bits
... distalUP-proximalUP-sigma38p10 63441 total 8.5 bits
... ... distalUP-(47)-s
... ... distalUP-proxin
proximalUP 3.3 bits
... proximalUP-(34)
distalUP 6.4 bits
... distalUP-(41)-s
... distalUP-proxin

. *63470 . *63480 . *63490 . *63500
5′ c a a a a t t a t c c t c a a t a a a a g g g a g t a t t c c c t c c g c c 3′
... sigma38p10 4.2 bits
sigma38p10 3.5 bits

```
... --| proximalUP-(38)-sigma38p10 63465 Gap 3.2 bits
      |           |
      |           |
... --| distalUP-(47)-sigma38p10 63465 Gap 6.5 bits
... --| distalUP-proximalUP-sigma38p10 63465 total 4.3 bits
                  |
... --------------| proximalUP-(34)-sigma38p10 63471 Gap 2.5 bits
                  |
... --------------| distalUP-(41)-sigma38p10 63471 Gap 3.1 bits
... --------------| distalUP-proximalUP-sigma38p10 63471 total 7.5 bits

             .         *63510    .         *63520    .         *63530    .
 5' a c g g g t t g t a g c t g g c g g g t c a g a t a g t g t t c g t a a t c 3'

    *63540    .         *63550    .         *63560    .         *63570    .
 5' c a g t g g t g a a c g t t g g t a g t c c a g c g g c t c c g g g c c g t 3'

        *63580    .
 5' t g g t g g t c c a t 3'
```

piece 2, NC_000913, lpxC, config: linear, direction: -, begin: 106666, end: 106386

```
     .         *106660   .         *106650   .         *106640   .         *106630
5' a c c c c g g t g t t g g c c g g c g c a g g g c g t a a c g t c a g g g t 3'
   T timestamp:_2018Sep20_11-35-24

       .         *106620   .         *106610   .         *106600   .
5' g a c t t t c t t g c c g g t a t g t a a a c c g a c a c c c g t c g c c t 3'
                                   proximalUP 2.2 bits
                           |------------------------------------------------------ ... proximalUP-(36)
                           distalUP 1.5 bits
       |---------------------------------------------------------------------- ... distalUP-(43)-s
       |-------------|---------------------------------------------------------- ... sigma38p10-prox

   *106590   .         *106580   .         *106570   .         *106560   .
5' g a a c g a t a c g t t t a a g t g t c c t t t g t t t g a t c a t c g t a 3'
                      sigma38p10 5.2 bits                                          ... proximalUP
... --------------| proximalUP-(36)-sigma38p10 106583 Gap 3.6 bits                 ... distalUP
                      proximalUP 4.5 bits                           |-----        ... distalUP-(40)-s
... --------------| distalUP-(43)-sigma38p10 106583 Gap 3.6 bits    |-----        ... sigma38p10-prox
... --------------| sigma38p10-proximalUP-distalUP 106583 total 1.7 bits |-----   ... distalUP-(42)-s
                           |---------------------------------------------------- ... proximalUP-(38)
                      distalUP 4.8 bits                             |-----        ... sigma38p10-prox
                           |---------------------------------------------------- ... distalUP-(44)-s
                           |-----------|---------------------------------------- ... sigma38p10-prox

       *106550   .         *106540   .         *106530   .         *106520   .
5' t t a t c t c g c c a a a t t a c c t a t c c a a c c g a a g t g t a c t a 3'
... proximalUP 7.2 bits                                                            ... sigma38p10
... distalUP 4.3 bits            sigma38p10 0.8 bits                               ... sigma38p10
```

```
... -----------------------------------------------------------------------| distalUP-(40)-sigma3
... --------|---------------------|-----------------------------------------| sigma38p10-proximalU
... ---------------------------------------------------------------------------- ... distalUP-(42)-s
... ------------------------------| proximalUP-(38)-sigma38p10 106537 Gap 3.2 bits
... --------|---------------------|-------------------------------------------- ... sigma38p10-prox
... ------------------------------| distalUP-(44)-sigma38p10 106537 Gap 2.8 bits
... --------|---------------------| sigma38p10-proximalUP-distalUP 106537 total 4.1 bits
            |-----------------------------------------------------------| proximalUP-(33)-sigm
            |-------------------------------------------------------------- ... proximalUP-(35)

          *106510    .         *106500    .         *106490    .         *106480
 5' t a c a t t c g g c g g g c c a g t t t a g c a c a a a g a g c c t c g a a 3'
```

... 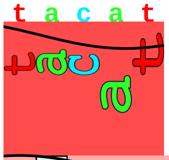 sigma38p10 7.7 bits

... 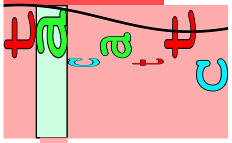 sigma38p10 3.5 bits

```
... --| distalUP-(42)-sigma38p10 106513 Gap 3.3 bits
... --| sigma38p10-proximalUP-distalUP 106513 total 8.7 bits
... --| proximalUP-(35)-sigma38p10 106513 Gap 3.1 bits

      .         *106470    .         *106460    .         *106450    .         *106440
 5' a c c c a a a t t c c a g t c a a t t c t t a a t c a g c t t g c t t a c g 3'

        .         *106430    .         *106420    .         *106410    .
 5' c a g g a a t g c t g g g a t a t c c a g a t a a t c c g g c t c t t t c g 3'

    *106400    .         *106390
 5' c a g t t t g c g g c g c a t 3'
```

piece 3, NC_000913, thrW, config: linear, direction: -, begin: 262252, end: 261990

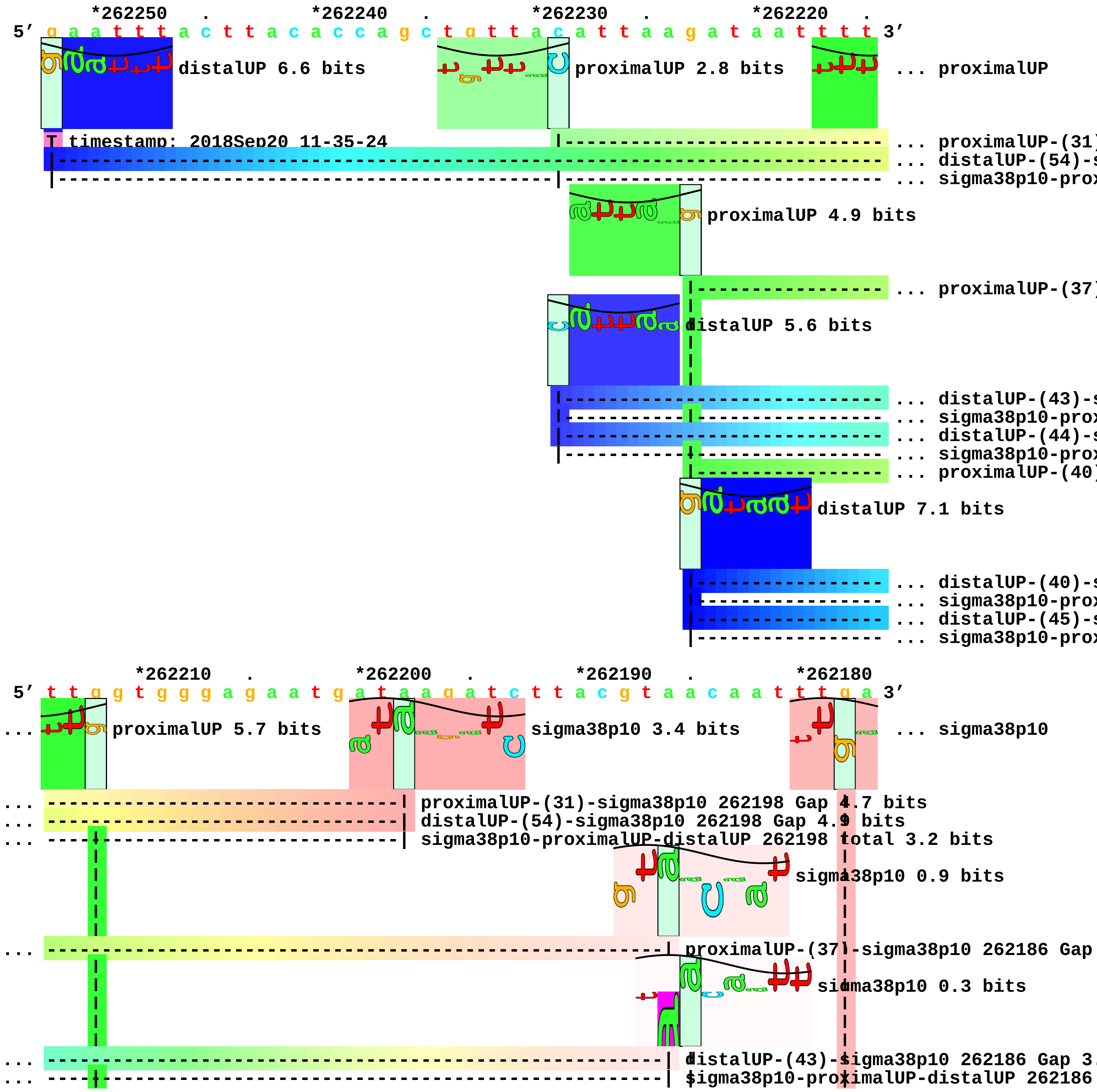

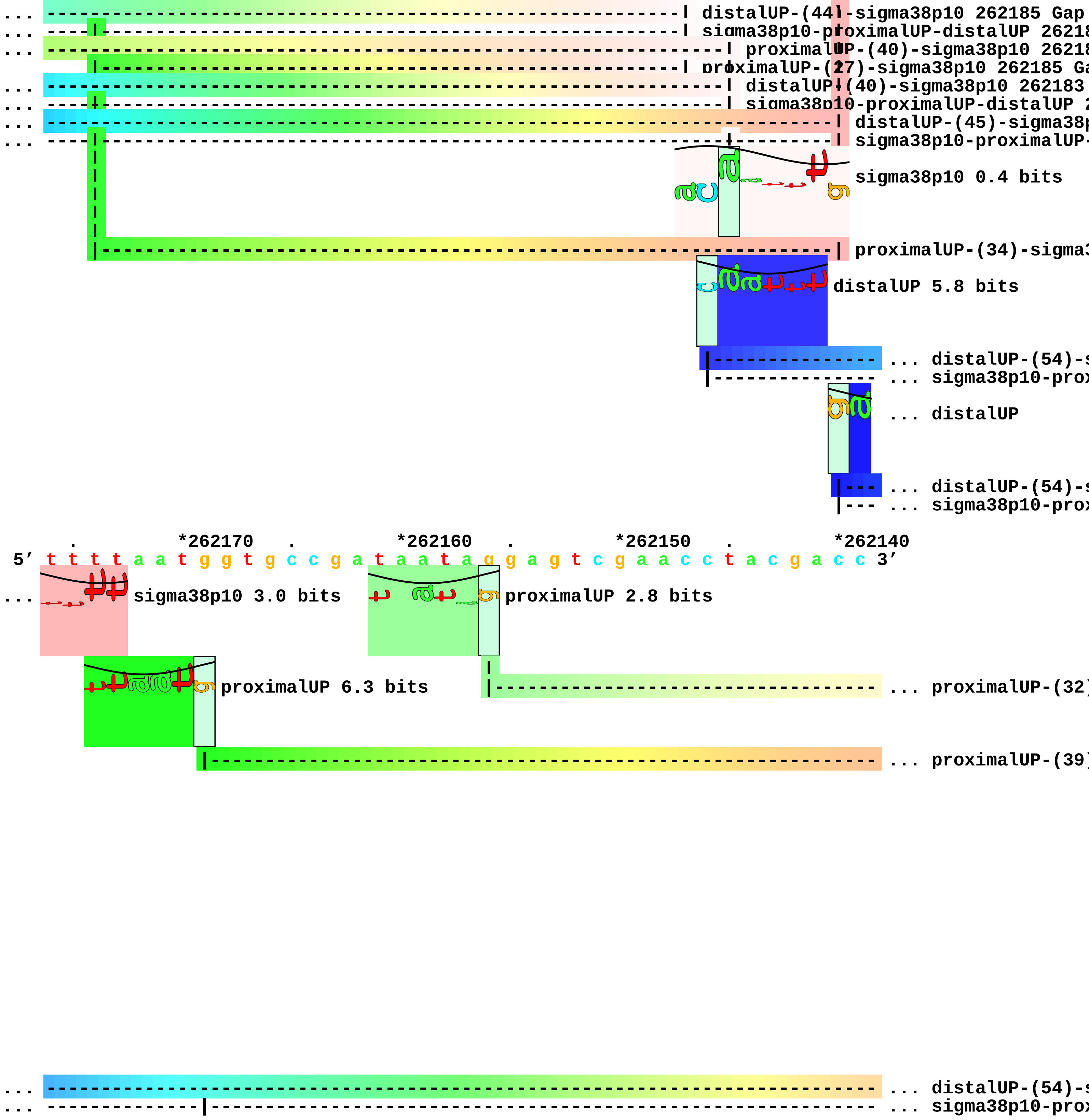
... distalUP-(44)-sigma38p10 262185 Gap
... sigma38p10-proximalUP-distalUP 26218
... proximalUP-(40)-sigma38p10 26218
proximalUP-(27)-sigma38p10 262185 Ga
... distalUP-(40)-sigma38p10 262183
... sigma38p10-proximalUP-distalUP 2
... distalUP-(45)-sigma38p
... sigma38p10-proximalUP-
sigma38p10 0.4 bits
proximalUP-(34)-sigma3
distalUP 5.8 bits
... distalUP-(54)-s
... sigma38p10-prox
... distalUP
... distalUP-(54)-s
... sigma38p10-prox
. *262170 . *262160 . *262150 . *262140
5′ t t t t a a t g g t g c c g a t a a t a g g a g t c g a a c c t a c g a c c 3′
... sigma38p10 3.0 bits
proximalUP 2.8 bits
proximalUP 6.3 bits
... proximalUP-(32)
... proximalUP-(39)
... distalUP-(54)-s
... sigma38p10-prox

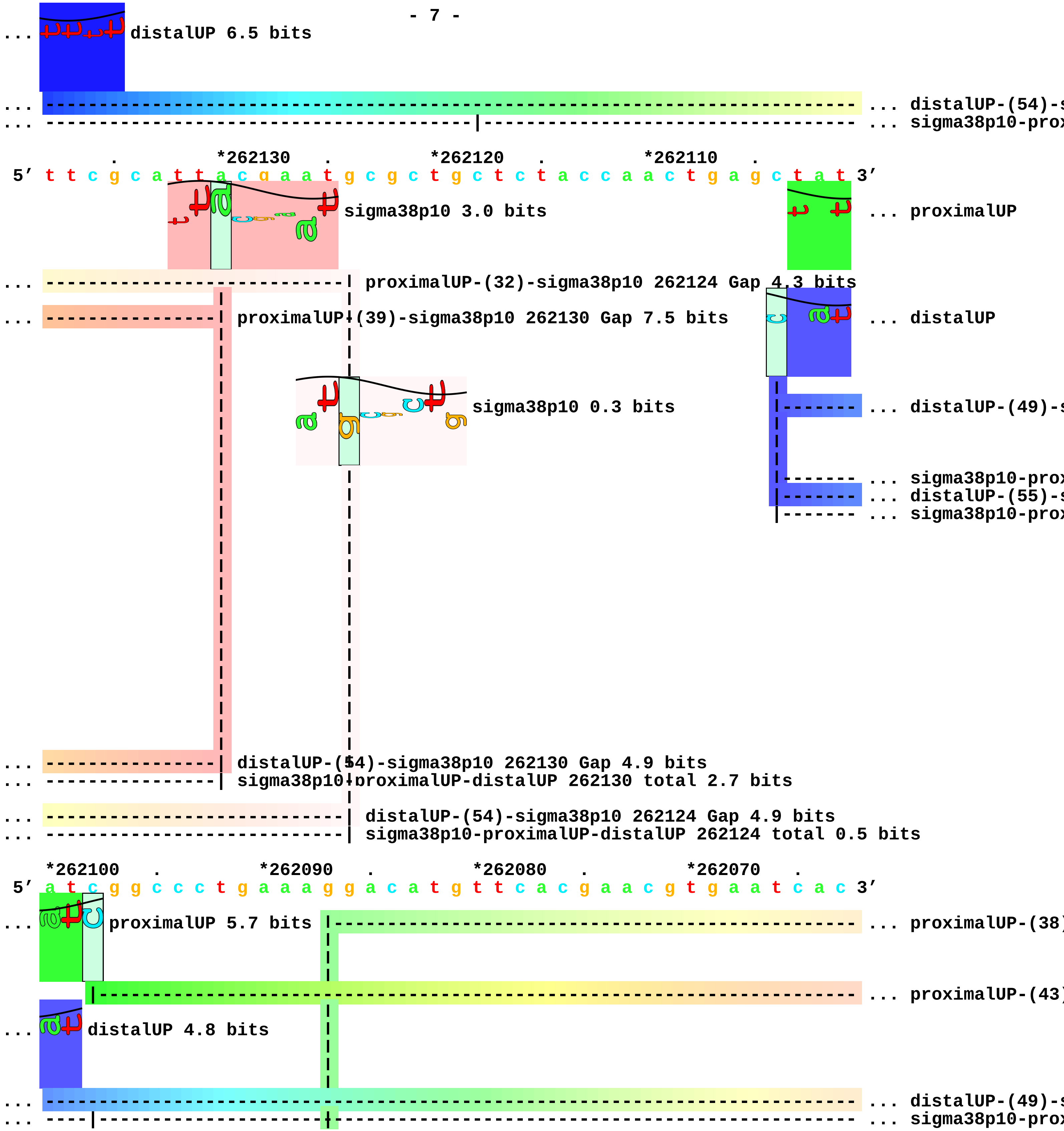
distalUP 6.5 bits
... distalUP-(54)-s
... sigma38p10-prox
*262130
*262120
*262110
5′ t t c g c a t t a c g a a t g c g c t g c t c t a c c a a c t g a g c t a t 3′
sigma38p10 3.0 bits
... proximalUP
proximalUP-(32)-sigma38p10 262124 Gap 4.3 bits
proximalUP-(39)-sigma38p10 262130 Gap 7.5 bits
... distalUP
sigma38p10 0.3 bits
... distalUP-(49)-s
... sigma38p10-prox
... distalUP-(55)-s
... sigma38p10-prox
distalUP-(54)-sigma38p10 262130 Gap 4.9 bits
sigma38p10-proximalUP-distalUP 262130 total 2.7 bits
distalUP-(54)-sigma38p10 262124 Gap 4.9 bits
sigma38p10-proximalUP-distalUP 262124 total 0.5 bits
*262100
*262090
*262080
*262070
5′ a t c g g c c c t g a a a g g a c a t g t t c a c g a a c g t g a a t c a c 3′
proximalUP 5.7 bits
... proximalUP-(38)
... proximalUP-(43)
distalUP 4.8 bits
... distalUP-(49)-s
... sigma38p10-prox

... distalUP-(55)-s
... sigma38p10-prox
proximalUP 2.8 bits

*262060 . *262050 . *262040 . *262030 .
5′ g g t g g a c a a g g t t a a a a c t a a c c g g g c g a t g c g t c a a t 3′
... proximalUP-(38)-sigma38p10 262049 Gap 3.2 bits
... proximalUP-(43)-sigma38p10 262055 Gap 5.1 bits
sigma38p10 1.1 bits
... distalUP-(49)-sigma38p10 262055 Gap 5.9 bits
... sigma38p10-proximalUP-distalUP 262055 total 0.5 bits
... distalUP-(55)-sigma38p10 262049 Gap 4.9 bits
... sigma38p10-proximalUP-distalUP 262049 total 0.3 bits
sigma38p10 0.8 bits
proximalUP 3.9 bits
... proximalUP-(42)
distalUP 4.9 bits
... distalUP-(49)-s
... sigma38p10-prox

*262020 . *262010 . *262000 . *261990
5′ g g c c t t g t g a a t c a a a t g g c t a c t t t t g c a t c a c c 3′
sigma38p10 7.5 bits
... proximalUP-(42)-sigma38p10 262003 Gap 5.1 bits
... distalUP-(49)-sigma38p10 262003 Gap 5.9 bits
... sigma38p10-proximalUP-distalUP 262003 total 5.2 bits

piece 4, NC_000913, insEF-2, config: linear, direction: +, begin: 392200, end: 392399

```
   *392200   .         *392210   .         *392220   .         *392230   .
5' c t g t a c g c c a a t a c t a t c a c c g t a a c t a a c g g c g g t g t 3'
   T timestamp:_2018Sep20_11-35-24

       *392240   .         *392250   .         *392260   .         *392270   .
5' a t t g g a t g t g a a c g t t g a t c a g t t c g a t a c t g a a g c t t 3'

           *392280   .         *392290   .         *392300   .         *392310
5' t c c g t a c t g a c a a a c t g g a a c t g a c c a g c g g c a a c a t c 3'

     .         *392320   .         *392330   .         *392340   .         *392350
5' g c t g a c c a t a a c g g t a a c g t a g t a t c t g g t g t g t t c g a 3'

         .         *392360   .         *392370   .         *392380   .
5' t a t c c a t a g c a g c g a t t a c g t t c t g a a c g c t g a t c t g g 3'

   *392390   .
5' t g a a c g a c c g 3'
```

piece 5, NC_000913, yaiA, config: linear, direction: -, begin: 406249, end: 406050

. *406240 . *406230 . *406220 .
5′ a t c g t c a c t a t a t a t g c t t c a c g a g g a t a a g g c g g t t t 3′
proximalUP 5.2 bits
... proximalUP-(33)
distalUP 4.8 bits
... distalUP-(41)-s
... sigma38p10-prox

T timestamp:_2018Sep20_11-35-24

*406210 . *406200 . *406190 . *406180 .
5′ c g t t g g c a t c g t t c t t c c t t a t t t c a c g g g a t g a a c g t 3′
sigma38p10 1.9 bits
... proximalUP-(33)-sigma38p10 406201 Gap 4.0 bits
proximalUP 5.1 bits
... distalUP-(41)-sigma38p10 406201 Gap 3.1 bits
... sigma38p10-proximalUP-distalUP 406201 total 4.6 bits
... proximalUP-(37)
distalUP 4.1 bits
... distalUP-(44)-s
... sigma38p10-prox

*406170 . *406160 . *406150 . *406140
5′ t a a g t a t a g g c g c t c g a a a a t c a a c a a t t g a t c g t c t g 3′
sigma38p10 2.0 bits
... proximalUP-(37)-sigma38p10 406150 Gap 4.5 bits
... distalUP-(44)-sigma38p10 406150 Gap 2.8 bits
... sigma38p10-proximalUP-distalUP 406150 total 3.9

. *406130 . *406120 . *406110 . *406100
5′ t g c c a g g g c g c t g c g a a t t t c a g a a a t c a c c t g g c t g g 3′

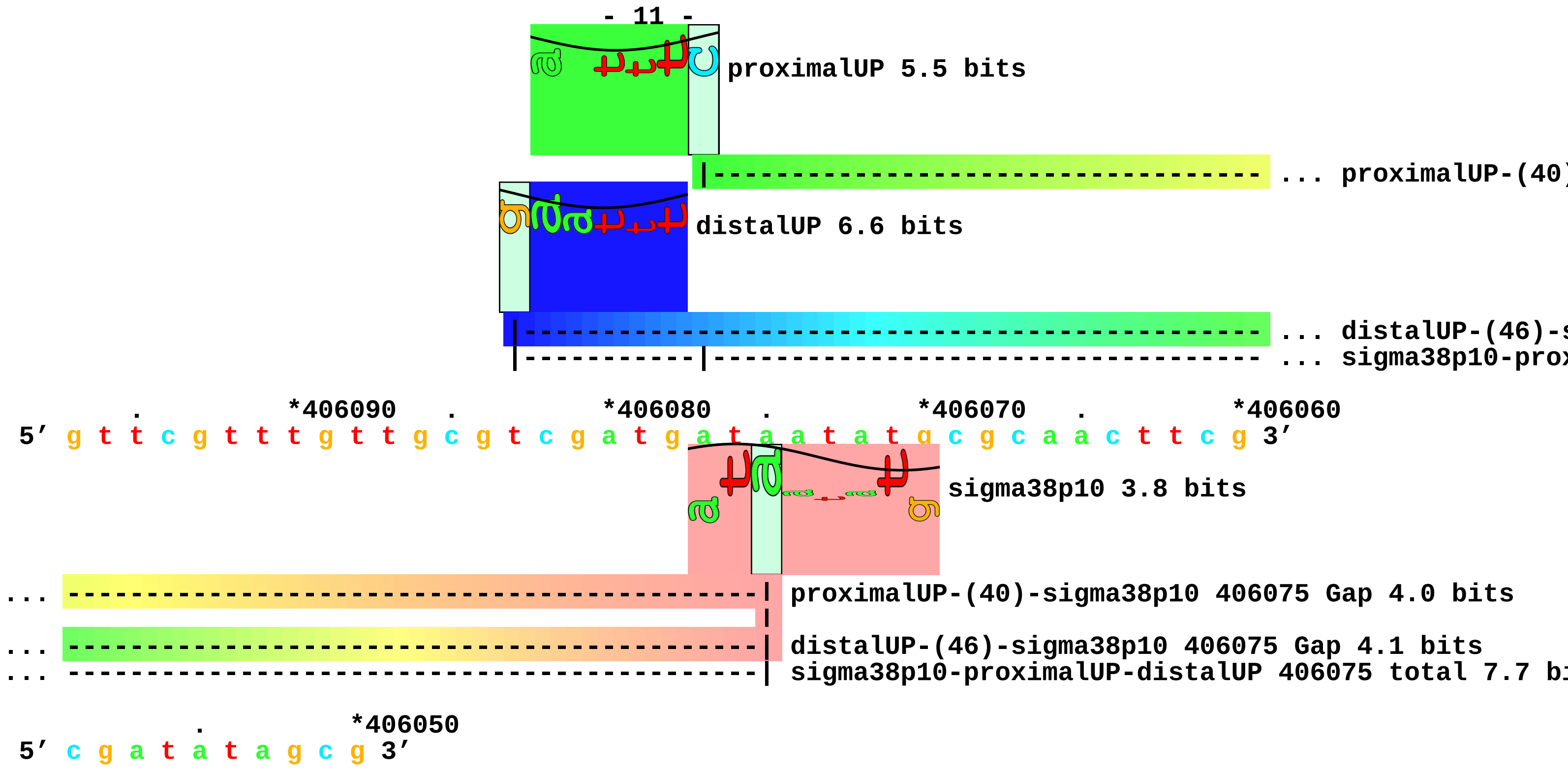
- 11 -
proximalUP 5.5 bits
... proximalUP-(40)
distalUP 6.6 bits
... distalUP-(46)-s
... sigma38p10-prox
. *406090 . *406080 . *406070 . *406060
5′ g t t c g t t t g t t g c g t c g a t g a t a a t a t g c g c a a c t t c g 3′
sigma38p10 3.8 bits
... proximalUP-(40)-sigma38p10 406075 Gap 4.0 bits
... distalUP-(46)-sigma38p10 406075 Gap 4.1 bits
... sigma38p10-proximalUP-distalUP 406075 total 7.7 bi
. *406050
5′ c g a t a t a g c g 3′

piece 6, NC_000913, yajO, config: linear, direction: +, begin: 437279, end: 437519

*437280 . *437290 . *437300 . *437310 .
5′ a t c t g g c t c g c c a a a g g t c a t a c a g c c g a g g c a a a g t c 3′
T timestamp:_2018Sep20_11-35-24

*437320 . *437330 . *437340 . *437350
5′ g g g a a a c g c g a a g g t c g g t t t t t c c t a a g g g g t t g t a t 3′

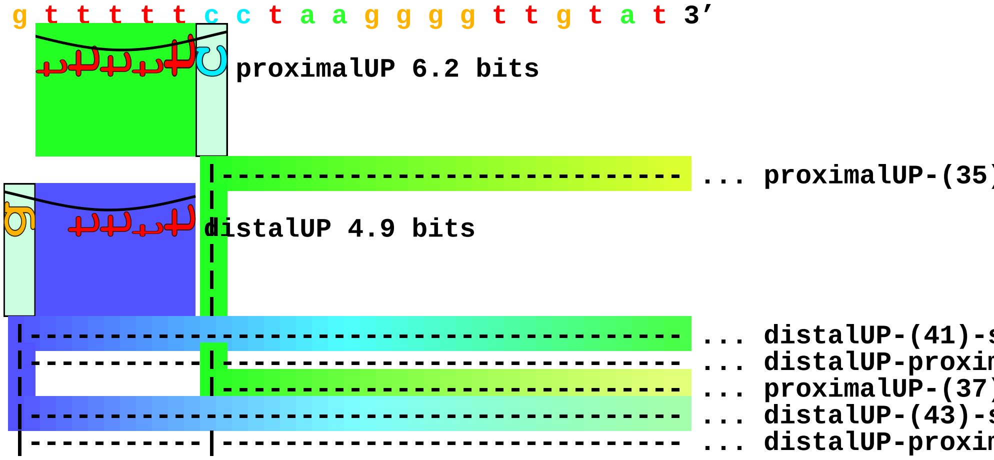


. *437360 . *437370 . *437380 . *437390
5′ t g c a t g c t g c c a c t c c t g c t a t a c t c g t c a t a c t t c a a 3′

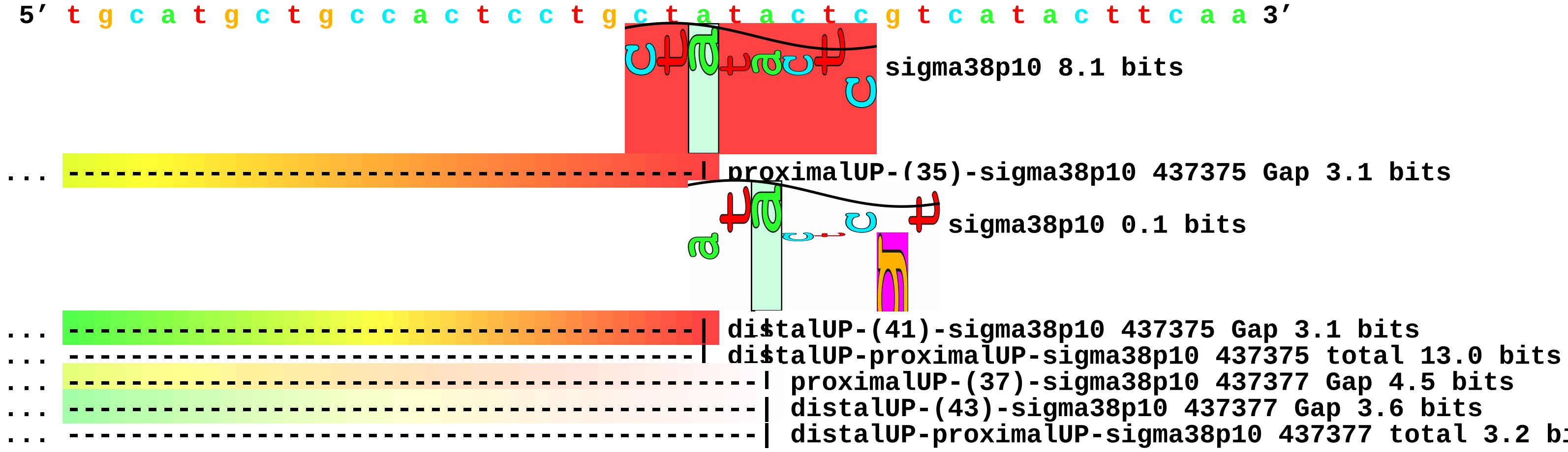


. *437400 . *437410 . *437420 . *437430
5′ g t t g c a t g t g c t g c g g c t g c a t t c g t t c a c c c c a g t c a 3′

. *437440 . *437450 . *437460 .
5′ c t t a c t t a t g t a a g c t c c t g g g g c t t c a c t c g t t t g c c 3′

*437470 . *437480 . *437490 . *437500 .
5′ g c c t t c c t g c a a c t c g a a t t a t t t a g a g t c t a t g a a t a 3′

*437510 .
5′ a t t t c t t a a g c a t 3′

piece 7, NC_000913, tomB, config: linear, direction: +, begin: 479870, end: 480165

```
   *479870   .       *479880   .       *479890   .       *479900   .
5′ c a g g g t t t c a c a g a g a a a c t t a a g c t g t g c g a t a t c a t 3′
```

proximalUP 3.8 bits

T timestamp:_2018Sep20_11-35-24

... proximalUP-(35)

distalUP 2.7 bits

... distalUP-(41)-s

... distalUP-proxin

proximalUP 2.0 bits

... proximalUP-(34)

distalUP 4.0 bits

... distalUP-(43)-s

... distalUP-proxin

```
     *479910   .       *479920   .       *479930   .       *479940   .
5′ g t c t t t t g g g t g a g t a t t c a t c c a t a a c g c g t c c c c t t 3′
```

sigma38p10 3.9 bits

... proximalUP-(35)-sigma38p10 479923 Gap 3.1 bits

sigma38p10 0.2 bits

... distalUP-(41)-sigma38p10 479923 Gap 3.1 bits

... distalUP-proximalUP-sigma38p10 479923 total 4.2 bits

... proximalUP-(34)-sigma38p10 479927 Gap 2.5 bits

... distalUP-(43)-sigma38p10 479927 Gap 3.6 bits

... distalUP-proximalUP-sigma38p10 479927 total 0.2 bits

```
       *479950   .       *479960   .       *479970   .       *479980
5′ c t t a g c g g t t g a a c t a a c g g a c a c c t t t c g g g a t g g a a 3′
```

... proximalUP

... distalUP

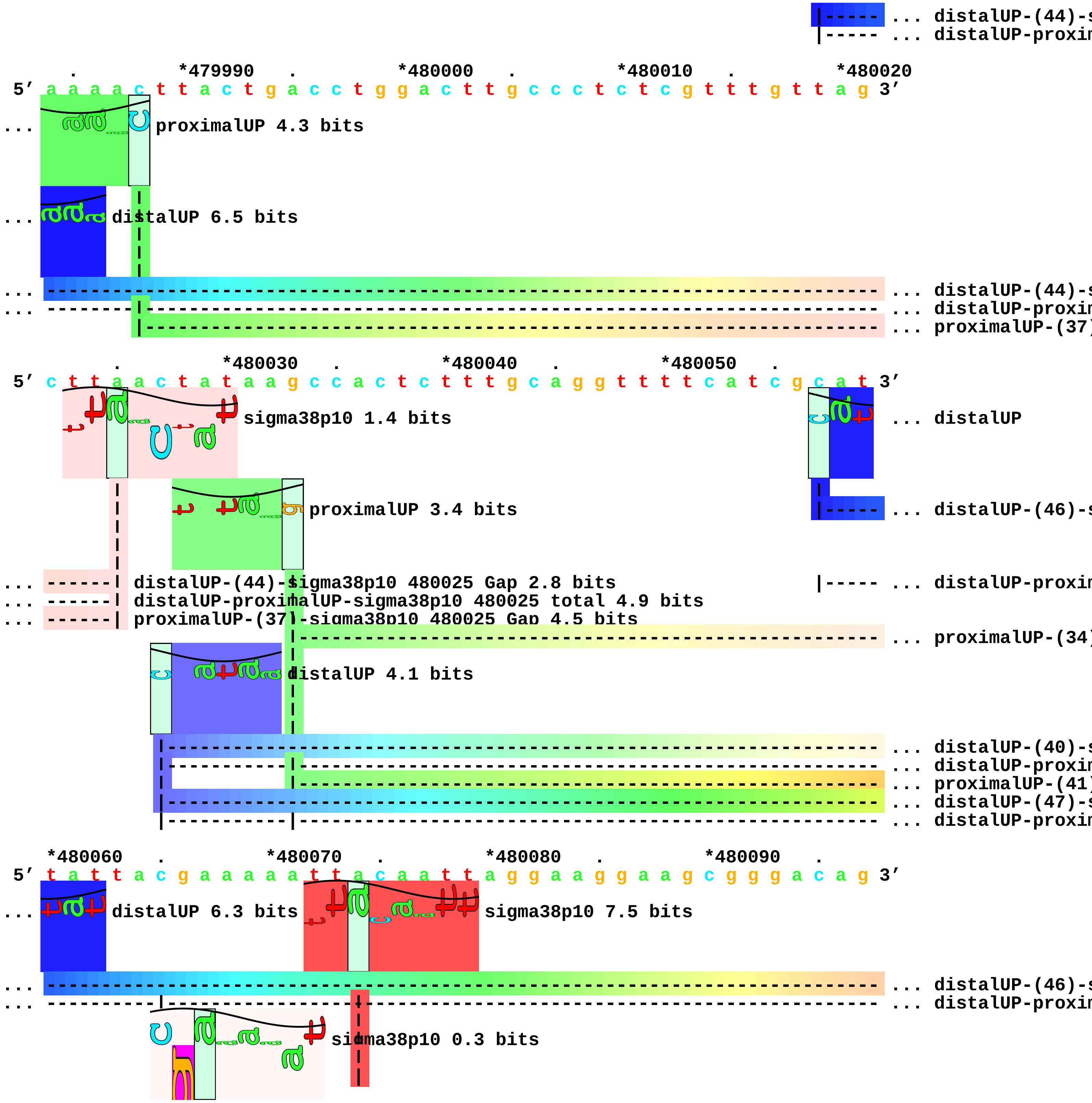
... distalUP-(44)-s
distalUP-proxim
*479990
*480000
*480010
*480020
5′ a a a a c t t a c t g a c c t g g a c t t g c c c t c t c g t t t g t t a g 3′
proximalUP 4.3 bits
distalUP 6.5 bits
... distalUP-(44)-s
... distalUP-proxim
... proximalUP-(37)
*480030
*480040
*480050
5′ c t t a a c t a t a a g c c a c t c t t t g c a g g t t t t c a t c g c a t 3′
sigma38p10 1.4 bits
... distalUP
proximalUP 3.4 bits
... distalUP-(46)-s
distalUP-(44)-sigma38p10 480025 Gap 2.8 bits
distalUP-proximalUP-sigma38p10 480025 total 4.9 bits
proximalUP-(37)-sigma38p10 480025 Gap 4.5 bits
... distalUP-proxim
... proximalUP-(34)
distalUP 4.1 bits
... distalUP-(40)-s
... distalUP-proxim
... proximalUP-(41)
... distalUP-(47)-s
... distalUP-proxim
*480060
*480070
*480080
*480090
5′ t a t t a c g a a a a a t t a c a a t t a g g a a g g a a g c g g g a c a g 3′
distalUP 6.3 bits
sigma38p10 7.5 bits
... distalUP-(46)-s
... distalUP-proxim
sigma38p10 0.3 bits

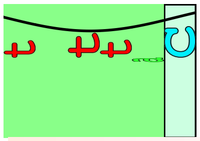

```
                      proximalUP 3.3 bits

... --------------| proximalUP-(34)-sigma38p10 480067 Gap 2.5 bits
          |------------------------------------------------------------ ... proximalUP-(38)
... --------------| distalUP-(40)-sigma38p10 480067 Gap 3.1 bits
... --------------| distalUP-proximalUP-sigma38p10 480067 total 2.3 bits
... ----------------------------| proximalUP-(41)-sigma38p10 480074 Gap 5.5 bits
... ----------------------------| distalUP-(47)-sigma38p10 480074 Gap 6.5 bits
... ----------------------------| distalUP-proximalUP-sigma38p10 480074 total 3.0 bits

        *480100   .        *480110   .         *480120   .         *480130   .
5' g a c t c a c t t t t t c a t a c t a g c a c t a a c c c g c a g c a g g t 3'
```

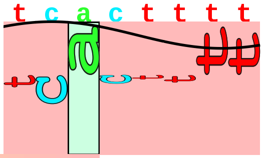

```
                             sigma38p10 2.9 bits

... ----------| distalUP-(46)-sigma38p10 480103 Gap 4.1 bits
... ----------| distalUP-proximalUP-sigma38p10 480103 total 5.2 bits

... ----------| proximalUP-(38)-sigma38p10 480103 Gap 3.2 bits

            *480140   .        *480150   .         *480160   .
5' a a a a g c a g t t t g c t c g a a c t a t t c a t a a c a 3'
```

piece 8, NC_000913, insH-2, config: linear, direction: +, begin: 574800, end: 575149

*574800 . *574810 . *574820 . *574830 .
5′ c t t g g g g t a a a a c g g c t c g a t g a c t t c c a c c a t g t t t t 3′
T timestamp:_2018Sep20_11-35-24

*574840 . *574850 . *574860 . *574870 .
5′ g c c a t g g c a g a a t c t g c t c c a t g c g g g a c a a g a a a a t c 3′

*574880 . *574890 . *574900 . *574910
5′ t c t t t t c t g g t c t g a c g g c g c t t a c t g c t g a a t t c a c t 3′

. *574920 . *574930 . *574940 . *574950
5′ g t c g g c g a a g g t a a g t t g a t g a c t c a t g a t g a a c c c t g 3′

. *574960 . *574970 . *574980 .
5′ t t c t a t g g c t c c a g a t g a c a a a c a t g a t c t c a t a t c a g 3′

proximalUP 3.2 bits
distalUP 2.5 bits
|----- ... proximalUP-(33)
|------------------------------------- ... distalUP-(50)-s
|----------------------------------|----- ... distalUP-proxin

*574990 . *575000 . *575010 . *575020 .
5′ g g a c t t g t t c g c a c c t t c c t t a g t g a a g t c a t t t t t g t 3′

sigma38p10 4.1 bits
... ------------------------------------------------------------| proximalUP-(33)-sigma38p10 575020
... ------------------------------------------------------------| distalUP-(50)-sigma38p10 575020 Ga
... ------------------------------------------------------------| distalUP-proximalUP-sigma38p10 575
proximalUP 5.7 bits
|--- ... proximalUP-(33)
distalUP 5.7 bits
|----------------- ... distalUP-(40)-s
|-------------|--- ... distalUP-proxin
|--- ... proximalUP-(37)
|----------------- ... distalUP-(44)-s
|-------------|--- ... distalUP-proxin
|--- ... proximalUP-(39)
|----------------- ... distalUP-(46)-s
|-------------|--- ... distalUP-proxin

*575030 . *575040 . *575050 . *575060 .

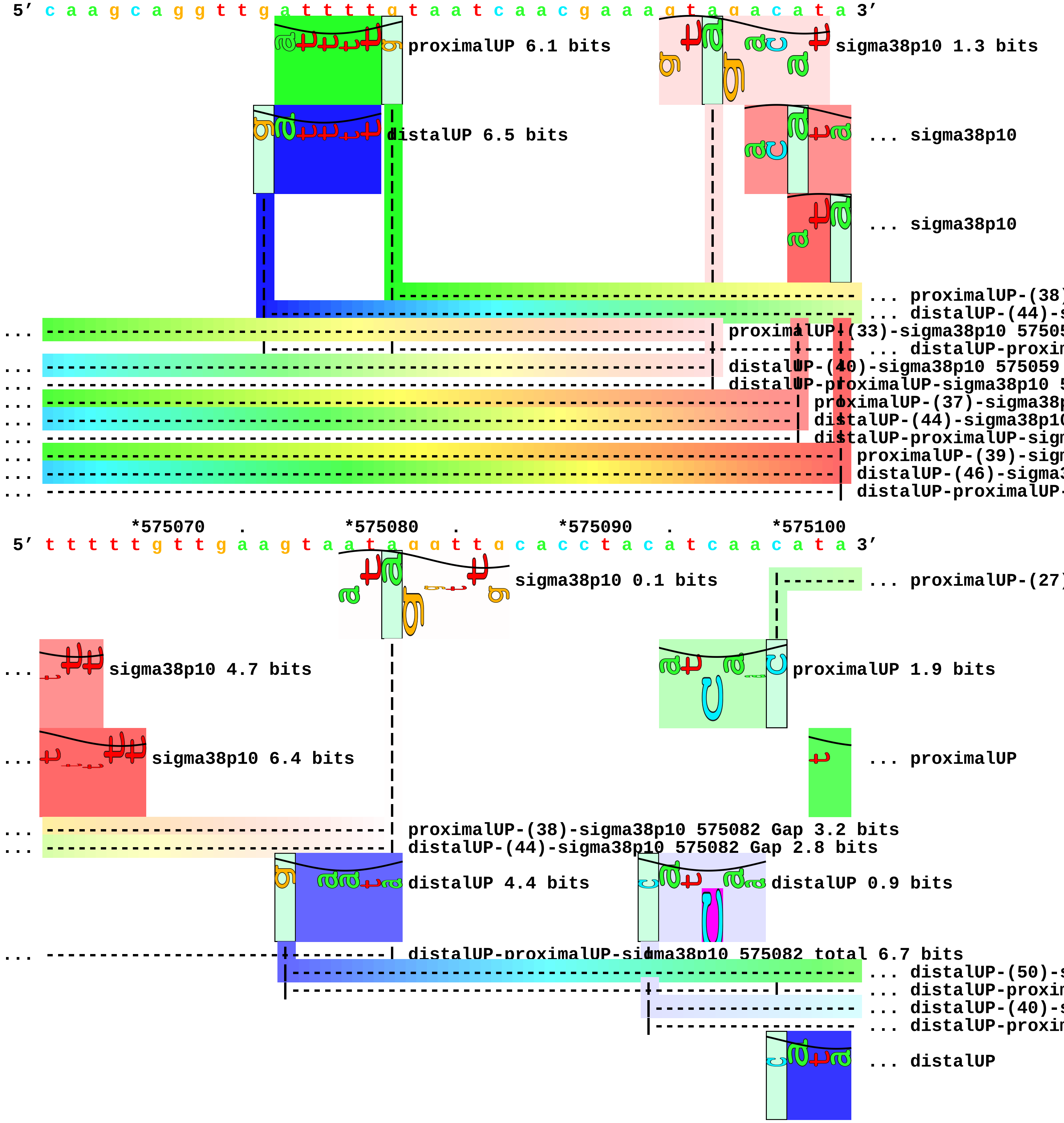
5′ c a a g c a g g t t g a t t t t g t a a t c a a c g a a a g t a g a c a t a 3′
proximalUP 6.1 bits
sigma38p10 1.3 bits
distalUP 6.5 bits
... sigma38p10
... sigma38p10
... proximalUP-(38)
... distalUP-(44)-s
... proximalUP-(33)-sigma38p10 57505
... distalUP-proxim
... distalUP-(40)-sigma38p10 575059
... distalUP-proximalUP-sigma38p10 5
... proximalUP-(37)-sigma38p
... distalUP-(44)-sigma38p10
... distalUP-proximalUP-sigm
... proximalUP-(39)-sigm
... distalUP-(46)-sigma3
... distalUP-proximalUP-
*575070 . *575080 . *575090 . *575100
5′ t t t t t g t t g a a g t a a t a g g t t g c a c c t a c a t c a a c a t a 3′
sigma38p10 0.1 bits
... proximalUP-(27)
... sigma38p10 4.7 bits
proximalUP 1.9 bits
... sigma38p10 6.4 bits
... proximalUP
... proximalUP-(38)-sigma38p10 575082 Gap 3.2 bits
... distalUP-(44)-sigma38p10 575082 Gap 2.8 bits
distalUP 4.4 bits
distalUP 0.9 bits
... distalUP-proximalUP-sigma38p10 575082 total 6.7 bits
... distalUP-(50)-s
... distalUP-proxim
... distalUP-(40)-s
... distalUP-proxim
... distalUP

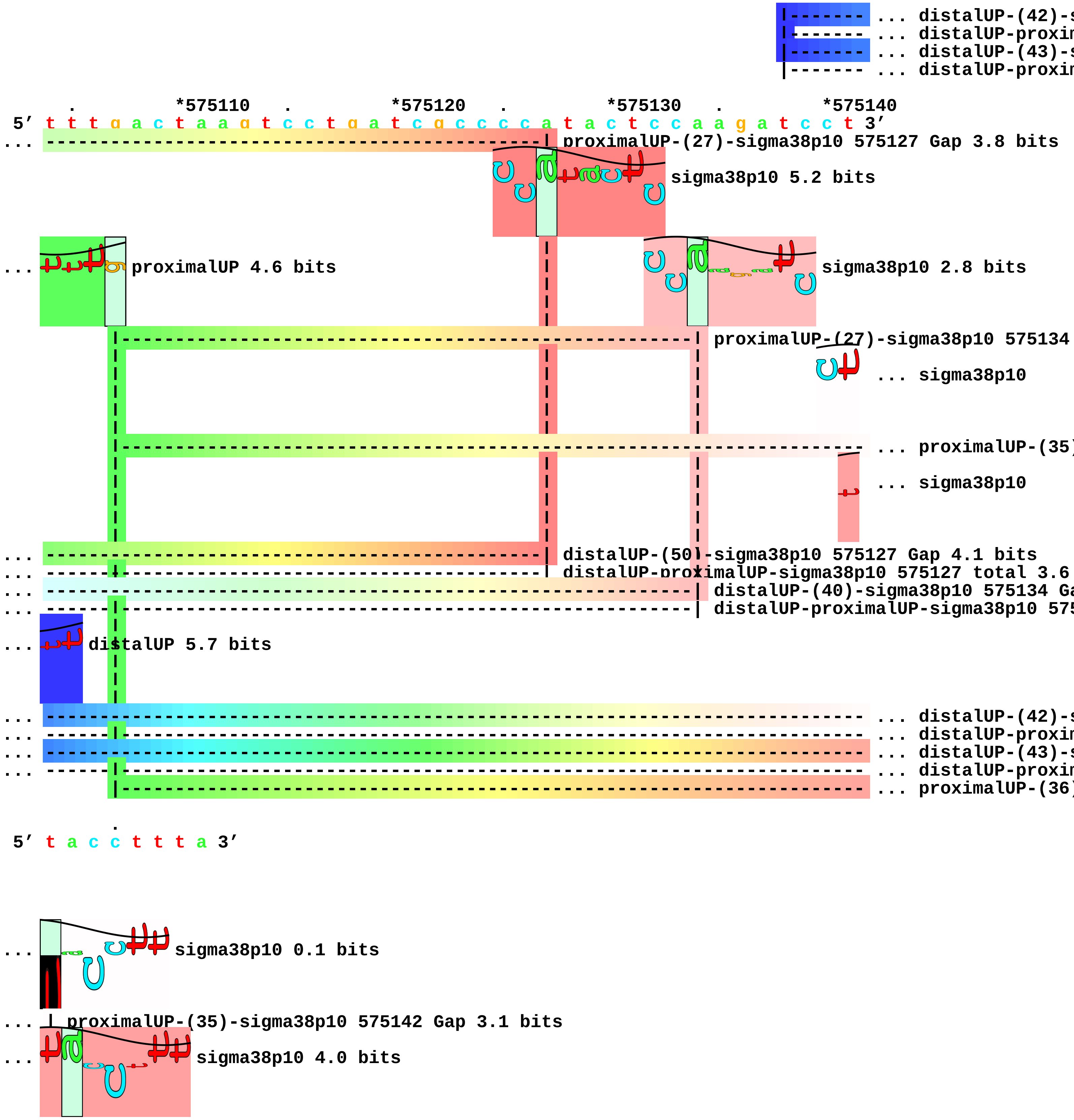
... distalUP-(42)-s
... distalUP-proxim
... distalUP-(43)-s
... distalUP-proxim
. *575110 . *575120 . *575130 . *575140
5' t t t g a c t a a g t c c t g a t c g c c c c a t a c t c c a a g a t c c t 3'
... proximalUP-(27)-sigma38p10 575127 Gap 3.8 bits
sigma38p10 5.2 bits
... proximalUP 4.6 bits
sigma38p10 2.8 bits
proximalUP-(27)-sigma38p10 575134
... sigma38p10
... proximalUP-(35)
... sigma38p10
... distalUP-(50)-sigma38p10 575127 Gap 4.1 bits
... distalUP-proximalUP-sigma38p10 575127 total 3.6
... distalUP-(40)-sigma38p10 575134 Ga
... distalUP-proximalUP-sigma38p10 575
... distalUP 5.7 bits
... distalUP-(42)-s
... distalUP-proxim
... distalUP-(43)-s
... distalUP-proxim
... proximalUP-(36)
5' t a c c t t t a 3'
... sigma38p10 0.1 bits
... proximalUP-(35)-sigma38p10 575142 Gap 3.1 bits
... sigma38p10 4.0 bits

```
    | |
    | |
    | |
    | |
    | |
... | distalUP-(42)-sigma38p10 575142 Gap 3.3 bits
... | distalUP-proximalUP-sigma38p10 575142 total 4.0 bits
... --| distalUP-(43)-sigma38p10 575143 Gap 3.6 bits
... --| distalUP-proximalUP-sigma38p10 575143 total 7.2 bits
... --| proximalUP-(36)-sigma38p10 575143 Gap 3.6 bits
```

piece 9, NC_000913, ybiI, config: linear, direction: +, begin: 837500, end: 837899

```
   *837500   .         *837510   .         *837520   .         *837530   .
5' a c a t a a g c g c a c g c c a g g a a t g g c t t c c c g a c g g g c c t 3'
   T timestamp:_2018Sep20_11-35-24

     *837540   .         *837550   .         *837560   .         *837570   .
5' g c g g g a t g g g g g c a c c g c a c t c t t c a c a t t c a t c c a g g 3'

       *837580   .         *837590   .         *837600   .         *837610
5' c t t t c g c c g c g c g g a a t t t c a c c c c g a g c g c g g g c a a t 3'
                                               proximalUP 5.5 bits
                                    |-------------------------------------- ... proximalUP-(34)
                                              distalUP 6.6 bits
                              |------------------------------------------------ ... distalUP-(40)-s
                              |---------- |------------------------------------ ... distalUP-proxin
                                          |------------------------------------ ... proximalUP-(37)
                              |------------------------------------------------ ... distalUP-(43)-s
                              |---------- |------------------------------------ ... distalUP-proxin
                                          |------------------------------------ ... proximalUP-(40)
                              |------------------------------------------------ ... distalUP-(46)-s
                              |---------- |------------------------------------ ... distalUP-proxin

   .         *837620   .         *837630   .         *837640   .         *837650
5' c g c a t c t t c a a t t g t a c t g t t g a t c t g t t c g t t g a c g g 3'
                                                       sigma38p10 3.0 bits
... -----------------------------| proximalUP-(34)-sigma38p10 837629 Gap 2.5 bits
                                                       sigma38p10 0.3 bits
... -----------------------------| distalUP-(40)-sigma38p10 837629 Gap 3.1 bits
... -----------------------------| distalUP-proximalUP-sigma38p10 837629 total 9.6 bits
... ----------------------------------| proximalUP-(37)-sigma38p10 837632 Gap 4.5 bits
... ----------------------------------| distalUP-(43)-sigma38p10 837632 Gap 3.6 bits
... ----------------------------------| distalUP-proximalUP-sigma38p10 837632 total 4.4 bits
... ---------------------------------------| proximalUP-(40)-sigma38p10 837635 Gap 4.0 bits
... ---------------------------------------| distalUP-(46)-sigma38p10 837635 Gap 4.1 bits
... ---------------------------------------| distalUP-proximalUP-sigma38p10 837635 total 5.5 bits
                                                       sigma38p10 1.6 bits

     .         *837660   .         *837670   .         *837680   .
5' c g t c a t c g t t a g c c c a a c c g g a t g c c a t a t c g a c c t c c 3'
```

```
   *837690   .         *837700   .         *837710   .         *837720   .
5' c c a t a t c a a t a c t t g t a c a g t t a a g t g t a g c t a a t c c a 3'
           <-------------------------------------#---------] ybiI

     *837730   .         *837740   .         *837750   .         *837760   .
5' g g g a c g a a c t c g g g c a g t t c a a g c a t c a g a t c t c c g a c 3'

       *837770   .         *837780   .         *837790   .         *837800
5' c a t t c c c g c a g c a g a t t a t g a t a a a g a t t a a g c a g c g a 3'

   .     *837810   .         *837820   .         *837830   .         *837840
5' c a g g a t c t c t t c a c t t t c g c c g t a g c g g c t t t t c a g c g 3'

     .     *837850   .         *837860   .         *837870   .
5' a c t g a a t a t t g t t g t c c a g t t c a a a c a g c a t g g c g c g c 3'
```

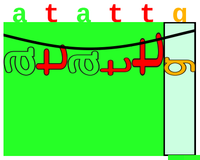

proximalUP 6.1 bits

```
            |------------------------------------------------------------ ... proximalUP-(33
```

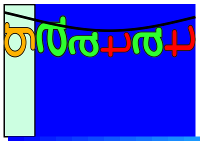

distalUP 7.2 bits

```
      |---------------------------------------------------------------------- ... distalUP-(40)-s
      |-------------|-------------------------------------------------------- ... distalUP-proxim
```

```
   *837880   .         *837890   .
5' t t t t t a t c a t c g c g g a t c a t 3'
```

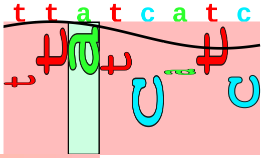

sigma38p10 2.9 bits

```
... ----------| proximalUP-(33)-sigma38p10 837885 Gap 4.0 bits
              |
... ----------| distalUP-(40)-sigma38p10 837885 Gap 3.1 bits
... ----------| distalUP-proximalUP-sigma38p10 837885 total 9.1 bits
```

piece 10, NC_000913, dps, config: linear, direction: +, begin: 848000, end: 848399

*848000 . *848010 . *848020 . *848030 .
5' c a a a g a a a g a t c a a t a a a c t g g a t a a c c t g g c g a t t c a 3'

proximalUP 3.3 bits
distalUP 3.2 bits
... proximalUP-(38)
distalUP 1.6 bits
... distalUP-(55)-s
... distalUP-(42)-s
... distalUP-proxim
proximalUP 5.4 bits
... distalUP-proxim
T timestamp:_2018Sep20 11-35-24
... proximalUP-(34)
distalUP 5.7 bits
... distalUP-(41)-s
... distalUP-proxim

*848040 . *848050 . *848060 . *848070 .
5' g c a a c t c t a c t g t t g c t t t t t c t c g c t g t c g g a g a c a 3'

sigma38p10 5.7 bits
... proximalUP-(38)-sigma38p10 848046 Gap 3.2 bits
... ... distalUP-(55)-s
... distalUP-(42)-sigma38p10 848046 Gap 3.3 bits
... distalUP-proximalUP-sigma38p10 848046 total 4.0 bits
... ... distalUP-proxim
... proximalUP-(34)-sigma38p10 848052 Gap 2.5 bits
sigma38p10 2.4 bits
... distalUP-(41)-sigma38p10 848052 Gap 3.1 bits
... distalUP-proximalUP-sigma38p10 848052 total 8.0 bits
proximalUP 6.2 bits
... proximalUP-(27)

*848080 . *848090 . *848100 . *848110
5' t c g t t g c g g g t a t a a a g c a g a t t g g t c g c t t t t g a t t t 3'

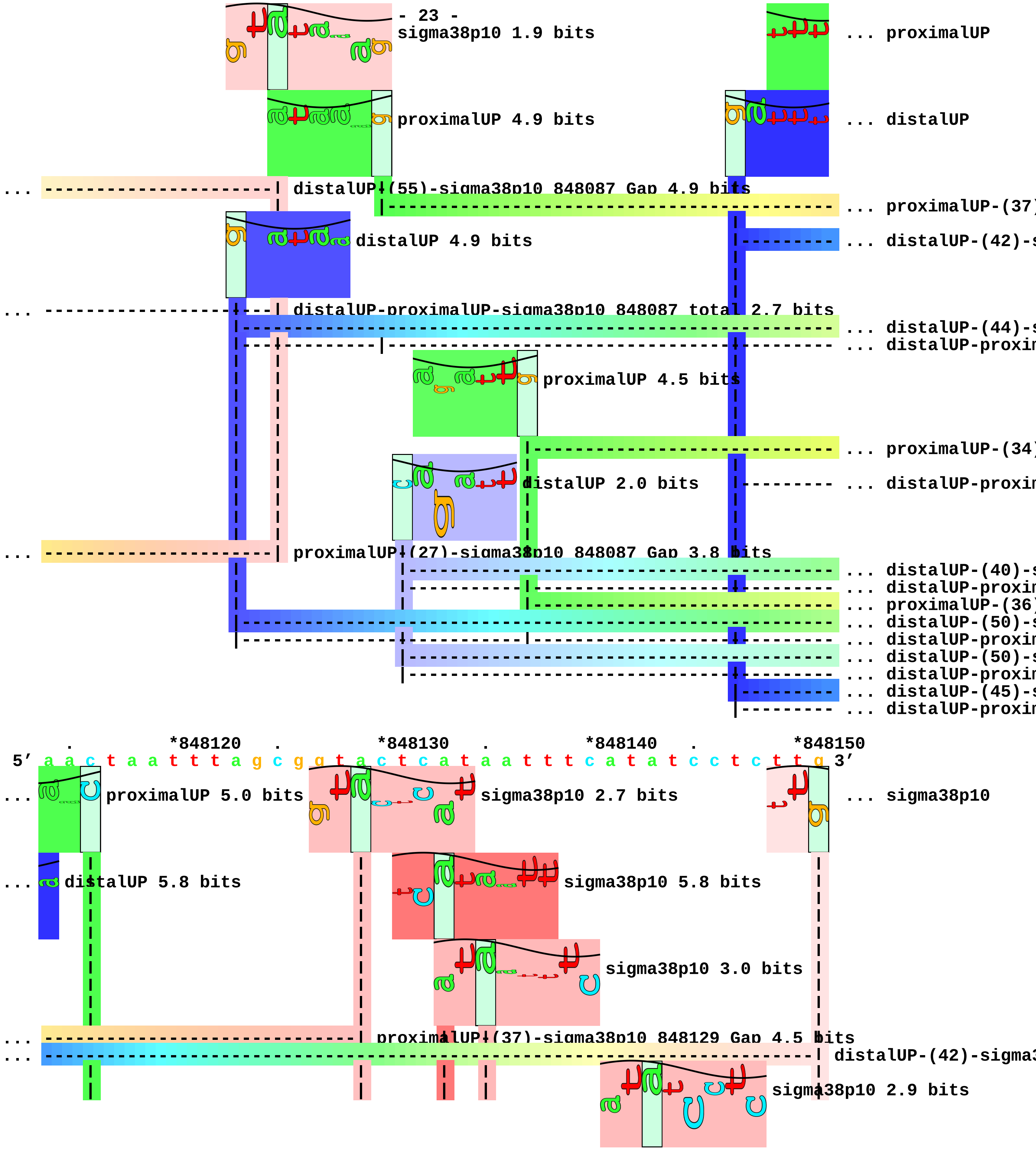

sigma38p10 1.9 bits
... proximalUP
proximalUP 4.9 bits
... distalUP
... distalUP-(55)-sigma38p10 848087 Gap 4.9 bits
... proximalUP-(37)
distalUP 4.9 bits
... distalUP-(42)-s
... distalUP-proximalUP-sigma38p10 848087 total 2.7 bits
... distalUP-(44)-s
... distalUP-proxim
proximalUP 4.5 bits
... proximalUP-(34)
distalUP 2.0 bits
... distalUP-proxim
... proximalUP-(27)-sigma38p10 848087 Gap 3.8 bits
... distalUP-(40)-s
... distalUP-proxim
... proximalUP-(36)
... distalUP-(50)-s
... distalUP-proxim
... distalUP-(50)-s
... distalUP-proxim
... distalUP-(45)-s
... distalUP-proxim
. *848120 . *848130 . *848140 . *848150
5' a a c t a a t t t a g c g g t a c t c a t a a t t t c a t a t c c t c t t g 3'
... proximalUP 5.0 bits
sigma38p10 2.7 bits
... sigma38p10
... distalUP 5.8 bits
sigma38p10 5.8 bits
sigma38p10 3.0 bits
... proximalUP-(37)-sigma38p10 848129 Gap 4.5 bits
... distalUP-(42)-sigma3
sigma38p10 2.9 bits

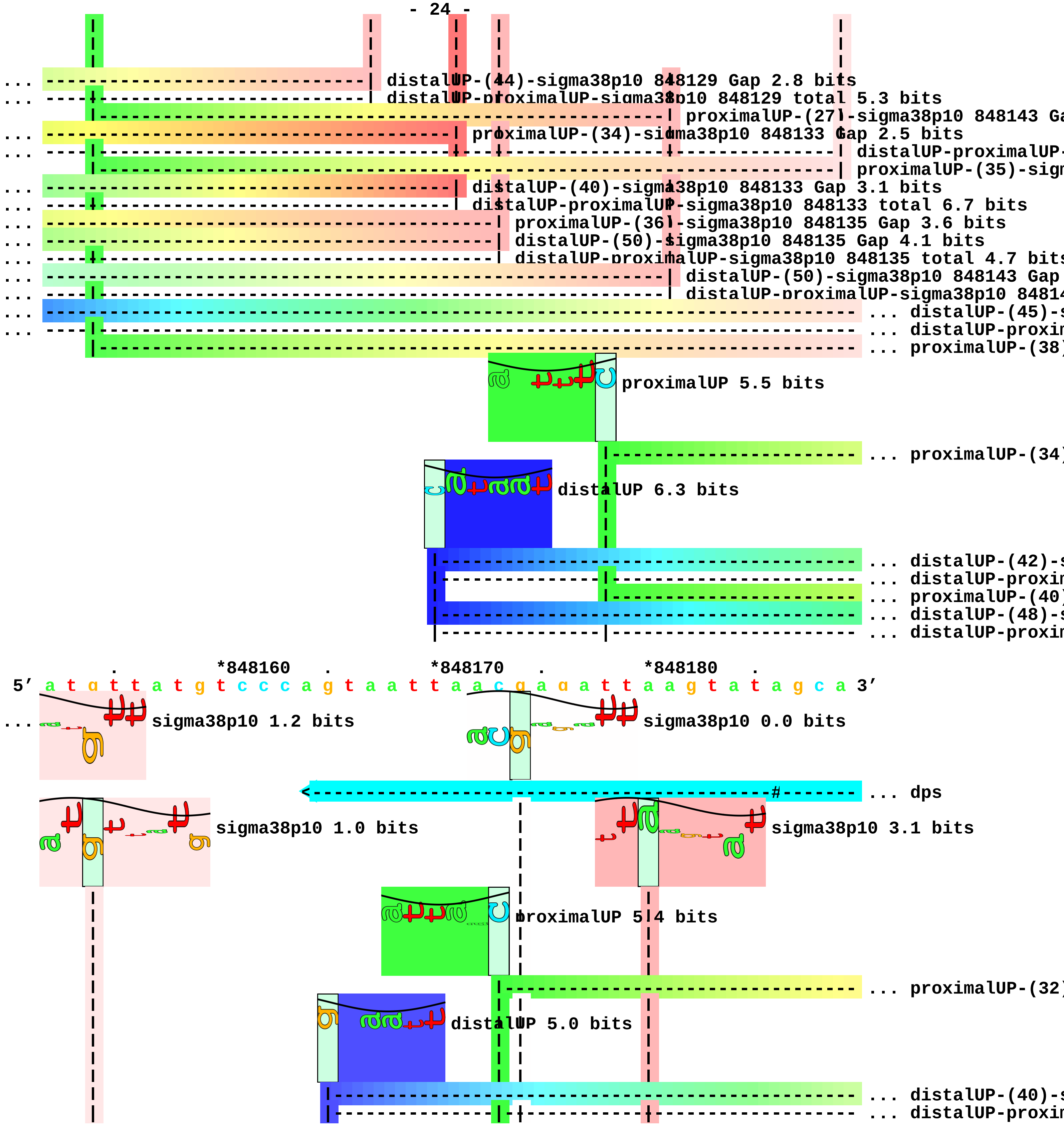
distalUP-(44)-sigma38p10 848129 Gap 2.8 bits
distalUP-proximalUP-sigma38p10 848129 total 5.3 bits
proximalUP-(27)-sigma38p10 848143 Ga
proximalUP-(34)-sigma38p10 848133 Gap 2.5 bits
distalUP-proximalUP-
proximalUP-(35)-sigm
distalUP-(40)-sigma38p10 848133 Gap 3.1 bits
distalUP-proximalUP-sigma38p10 848133 total 6.7 bits
proximalUP-(36)-sigma38p10 848135 Gap 3.6 bits
distalUP-(50)-sigma38p10 848135 Gap 4.1 bits
distalUP-proximalUP-sigma38p10 848135 total 4.7 bits
distalUP-(50)-sigma38p10 848143 Gap
distalUP-proximalUP-sigma38p10 8481
distalUP-(45)-s
distalUP-proxim
proximalUP-(38)
proximalUP 5.5 bits
proximalUP-(34)
distalUP 6.3 bits
distalUP-(42)-s
distalUP-proxim
proximalUP-(40)
distalUP-(48)-s
distalUP-proxim
*848160
*848170
*848180
5' a t g t t a t g t c c c a g t a a t t a a c g a g a t t a a g t a t a g c a 3'
sigma38p10 1.2 bits
sigma38p10 0.0 bits
dps
sigma38p10 1.0 bits
sigma38p10 3.1 bits
proximalUP 5.4 bits
proximalUP-(32)
distalUP 5.0 bits
distalUP-(40)-s
distalUP-proxim

... proximalUP-(35)
... distalUP-(43)-s
... distalUP-proxim

... distalUP-(45)-sigma38p10 848154 Gap 5.5 bits
... distalUP-proximalUP-sigma38p10 848154 total 3.2 bits
... proximalUP-(38)-sigma38p10 848154 Gap 3.2 bits
... proximalUP-(34)-sigma38p10 848174 Gap 2.5 bits
... distalUP-(42)-sigma38p10 848174 Gap 3.3 bits
... distalUP-proximalUP-sigma38p10 848174 total 6.1 bi
... proximalUP-(40)-sigma38p10 848180 Gap
... distalUP-(48)-sigma38p10 848180 Gap 7.
... distalUP-proximalUP-sigma38p10 848180

*848190 . *848200 . *848210 . *848220 .

5′ c c g g c t a t g t g t t c c g c t a t t c t g g c t g t t c c t a t c a c 3′

sigma38p10 1.8 bits

... --] dps

sigma38p10 7.2 bits

... proximalUP-(32)-sigma38p10 848205 Gap 4.3 bits
... distalUP-(40)-sigma38p10 848205 Gap 3.1 bits
... distalUP-proximalUP-sigma38p10 848205 total 4.8 bits
... proximalUP-(35)-sigma38p10 848208 Gap 3.1 bits
... distalUP-(43)-sigma38p10 848208 Gap 3.6 bits
... distalUP-proximalUP-sigma38p10 848208 total 11.0 bits

*848230 . *848240 . *848250 . *848260 .

5′ a c t a a t a g t g g t a a c a a g c g t g a a a a a c a a a a c t a a t a 3′

distalUP 3.5 bits

proximalUP 4.3 bits

... proximalUP-(30)
... distalUP-(56)-s
... distalUP-proxim
... proximalUP-(38)

distalUP 6.5 bits

... distalUP-(44)-s

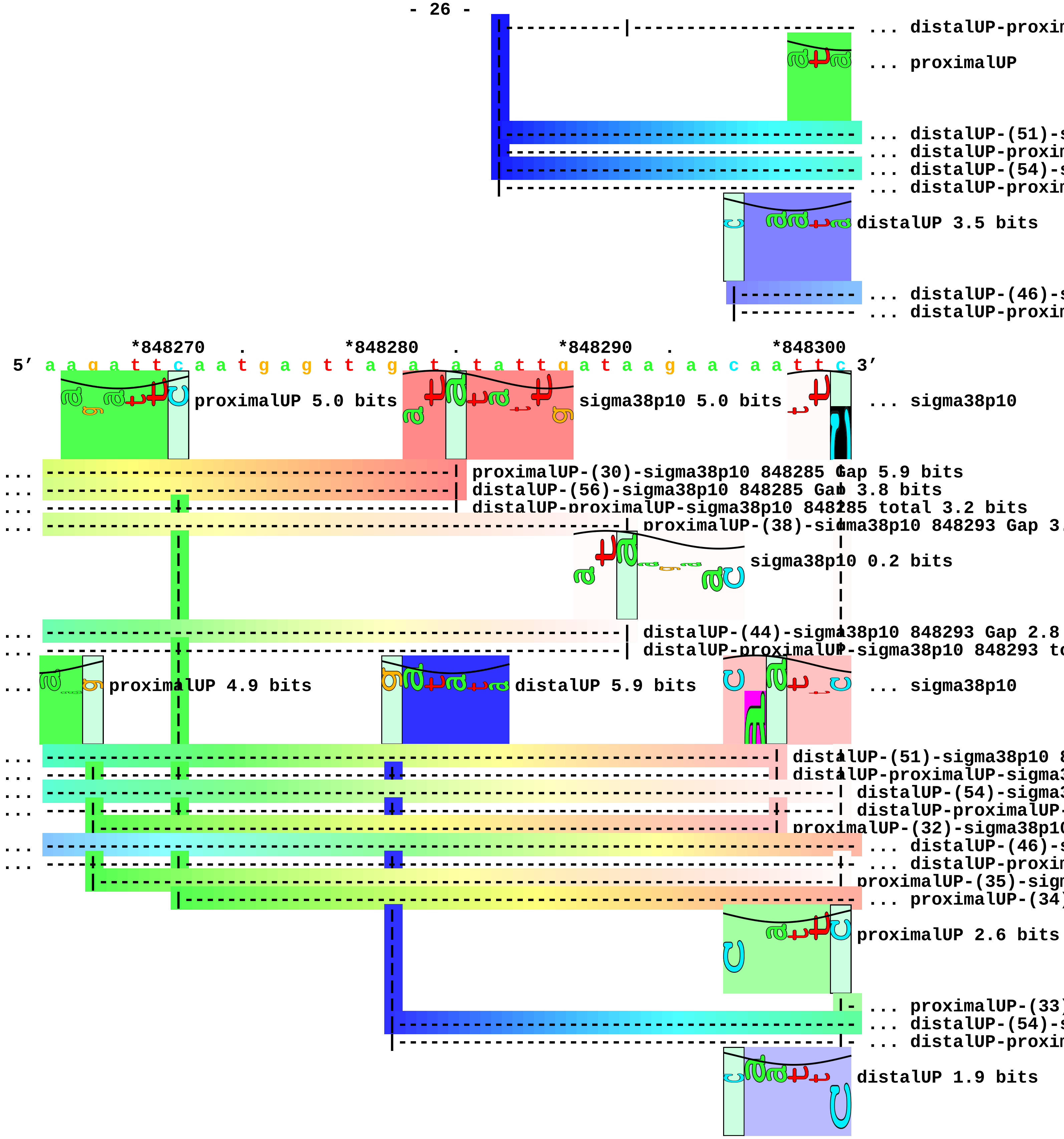
... distalUP-proxim
... proximalUP
... distalUP-(51)-s
... distalUP-proxim
... distalUP-(54)-s
... distalUP-proxim
distalUP 3.5 bits
... distalUP-(46)-s
... distalUP-proxim
*848270 . *848280 . *848290 . *848300
5′ a a g a t t c a a t g a g t t a g a t a t a t t g a t a a g a a c a a t t c 3′
proximalUP 5.0 bits
sigma38p10 5.0 bits
... sigma38p10
proximalUP-(30)-sigma38p10 848285 Gap 5.9 bits
distalUP-(56)-sigma38p10 848285 Gap 3.8 bits
distalUP-proximalUP-sigma38p10 848285 total 3.2 bits
proximalUP-(38)-sigma38p10 848293 Gap 3.
sigma38p10 0.2 bits
distalUP-(44)-sigma38p10 848293 Gap 2.8
distalUP-proximalUP-sigma38p10 848293 to
proximalUP 4.9 bits
distalUP 5.9 bits
... sigma38p10
distalUP-(51)-sigma38p10 8
distalUP-proximalUP-sigma3
distalUP-(54)-sigma3
distalUP-proximalUP-
proximalUP-(32)-sigma38p10
... distalUP-(46)-s
... distalUP-proxim
proximalUP-(35)-sigm
... proximalUP-(34)
proximalUP 2.6 bits
... proximalUP-(33)
... distalUP-(54)-s
... distalUP-proxim
distalUP 1.9 bits

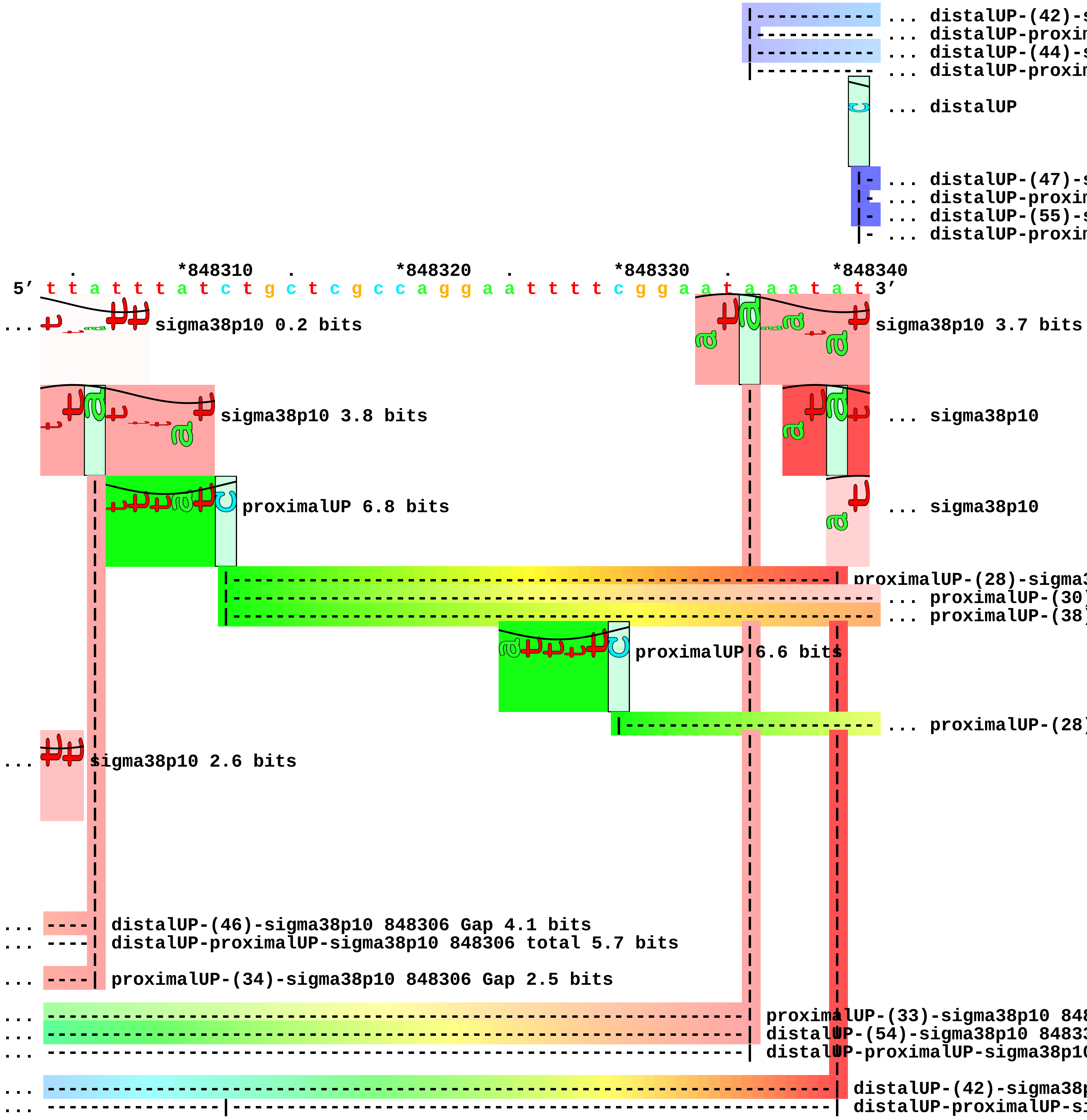
... distalUP-(42)-s
... distalUP-proxin
... distalUP-(44)-s
... distalUP-proxin
... distalUP
... distalUP-(47)-s
... distalUP-proxin
... distalUP-(55)-s
... distalUP-proxin
. *848310 . *848320 . *848330 . *848340
5' t t a t t t a t c t g c t c g c c a g g a a t t t t c g g a a t a a a t a t 3'
... sigma38p10 0.2 bits
sigma38p10 3.7 bits
sigma38p10 3.8 bits
... sigma38p10
proximalUP 6.8 bits
... sigma38p10
proximalUP-(28)-sigma3
... proximalUP-(30)
... proximalUP-(38)
proximalUP 6.6 bits
... proximalUP-(28)
... sigma38p10 2.6 bits
... distalUP-(46)-sigma38p10 848306 Gap 4.1 bits
... distalUP-proximalUP-sigma38p10 848306 total 5.7 bits
... proximalUP-(34)-sigma38p10 848306 Gap 2.5 bits
... proximalUP-(33)-sigma38p10 848
... distalUP-(54)-sigma38p10 84833
... distalUP-proximalUP-sigma38p10
... distalUP-(42)-sigma38p
... distalUP-proximalUP-si

```
... ---------------------------------------------------------------------------- ... distalUP-(44)-s
... ----------------|----------------------------------------------------------- ... distalUP-proxim
... distalUP 4.1 bits
... ---------------------------------------------------------------------------- ... distalUP-(47)-s
... ----------------|----------------------------------------------------------- ... distalUP-proxim
... ---------------------------------------------------------------------------- ... distalUP-(55)-s
... ---------------------------------------------------|------------------------ ... distalUP-proxim

           .         *848350    .         *848360    .         *848370    .
 5' g c t t t t g c a t a a t t c g a t a t a t a a a g a c g g t g t a g a g g 3'
                                    sigma38p10 4.8 bits
... sigma38p10 7.4 bits                   sigma38p10 0.4 bits
... sigma38p10 1.9 bits
... | proximalUP-(30)-sigma38p10 848342 Gap 5.9 bits
... ----------------| proximalUP-(38)-sigma38p10 848350 Gap 3.2 bits
... --------------------------------| proximalUP-(28)-sigma38p10 848358 Gap 5.5 bits
... | distalUP-(44)-sigma38p10 848342 Gap 2.8 bits
... | distalUP-proximalUP-sigma38p10 848342 total 1.9 bits
... ----------------| distalUP-(47)-sigma38p10 848350 Gap 6.5 bits
... ----------------| distalUP-proximalUP-sigma38p10 848350 total 6.0 bits
... --------------------------------| distalUP-(55)-sigma38p10 848358 Gap 4.9 bits
... --------------------------------| distalUP-proximalUP-sigma38p10 848358 total 0.8 bits

    *848380    .         *848390    .
 5' a a a a g t a g c g a g a a a t t c t g 3'
```

piece 11, NC_000913, ymgC, config: linear, direction: -, begin: 1216449, end: 1215850

```
          .         *1216440  .         *1216430  .         *1216420  .
5' c c g a a g a t a a a a t t t t c t c c a t c t t g a t c t a a c c c a t t 3'
```

proximalUP 6.6 bits

... proximalUP-(29

distalUP 6.5 bits

... distalUP-(40)-s
... sigma38p10-prox
... proximalUP-(33)
... distalUP-(44)-s
... sigma38p10-prox

T timestamp:_2018Sep20_11-35-24

```
  *1216410  .         *1216400  .         *1216390  .         *1216380  .
5' a a t t a t t g a g c a t a a t g t a g g c a g t a c a a a a t a a g t t a 3'
```

sigma38p10 0.8 bits

... proximalUP

... proximalUP-(29)-sigma38p10 1216404 Gap 7.5 bits

sigma38p10 2.3 bits

... distalUP-(40)-sigma38p10 1216404 Gap 3.1 bits
... sigma38p10-proximalUP-distalUP 1216404 total 3.3 bits
... proximalUP-(33)-sigma38p10 1216400 Gap 4.0 bits
... distalUP-(44)-sigma38p10 1216400 Gap 2.8 bits
... sigma38p10-proximalUP-distalUP 1216400 total 8.5 bits

proximalUP 6.7 bits

... proximalUP-(40

distalUP 6.3 bits

distalUP 6.4 bits

... distalUP-(46)-s
... sigma38p10-prox
... distalUP-(54)-s
... sigma38p10-prox

```
     *1216370  .         *1216360  .         *1216350  .         *1216340
5' g g c g g g a t a t c a g g c g t c a a g a a t g g a g a g c g a g a a c t 3'
```

... proximalUP 2.7 bits

sigma38p10 0.7 bits

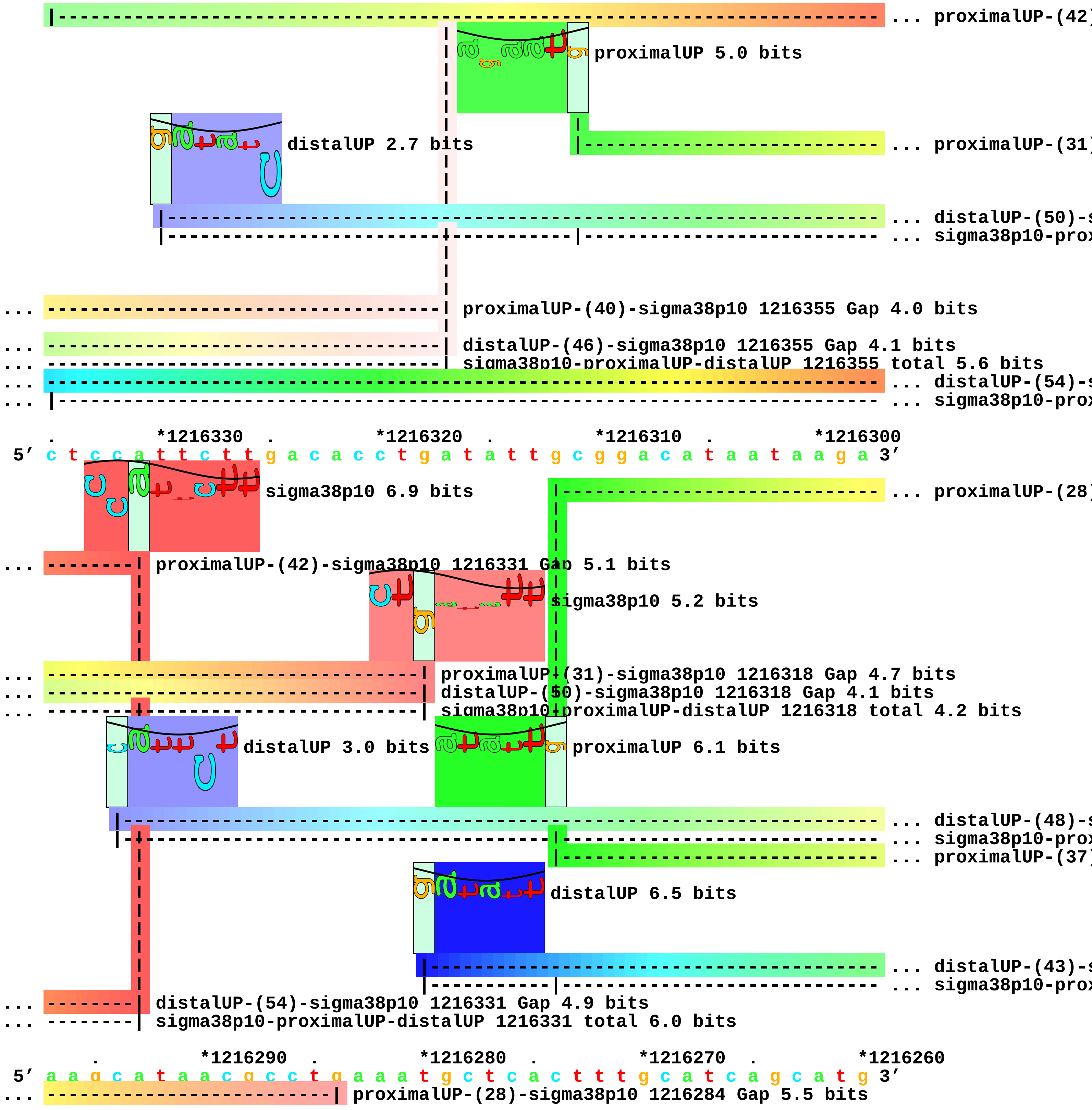
... proximalUP-(42)
proximalUP 5.0 bits
distalUP 2.7 bits
... proximalUP-(31)
... distalUP-(50)-s
... sigma38p10-prox
... proximalUP-(40)-sigma38p10 1216355 Gap 4.0 bits
... distalUP-(46)-sigma38p10 1216355 Gap 4.1 bits
... sigma38p10-proximalUP-distalUP 1216355 total 5.6 bits
... distalUP-(54)-s
... sigma38p10-prox
. *1216330 . *1216320 . *1216310 . *1216300
5' c t c c a t t c t t g a c a c c t g a t a t t g c g g a c a t a a t a a g a 3'
sigma38p10 6.9 bits
... proximalUP-(28)
... proximalUP-(42)-sigma38p10 1216331 Gap 5.1 bits
sigma38p10 5.2 bits
... proximalUP-(31)-sigma38p10 1216318 Gap 4.7 bits
... distalUP-(50)-sigma38p10 1216318 Gap 4.1 bits
... sigma38p10-proximalUP-distalUP 1216318 total 4.2 bits
distalUP 3.0 bits
proximalUP 6.1 bits
... distalUP-(48)-s
... sigma38p10-prox
... proximalUP-(37)
distalUP 6.5 bits
... distalUP-(43)-s
... sigma38p10-prox
... distalUP-(54)-sigma38p10 1216331 Gap 4.9 bits
... sigma38p10-proximalUP-distalUP 1216331 total 6.0 bits
. *1216290 . *1216280 . *1216270 . *1216260
5' a a g c a t a a c g c c t g a a a t g c t c a c t t t g c a t c a g c a t g 3'
... proximalUP-(28)-sigma38p10 1216284 Gap 5.5 bits

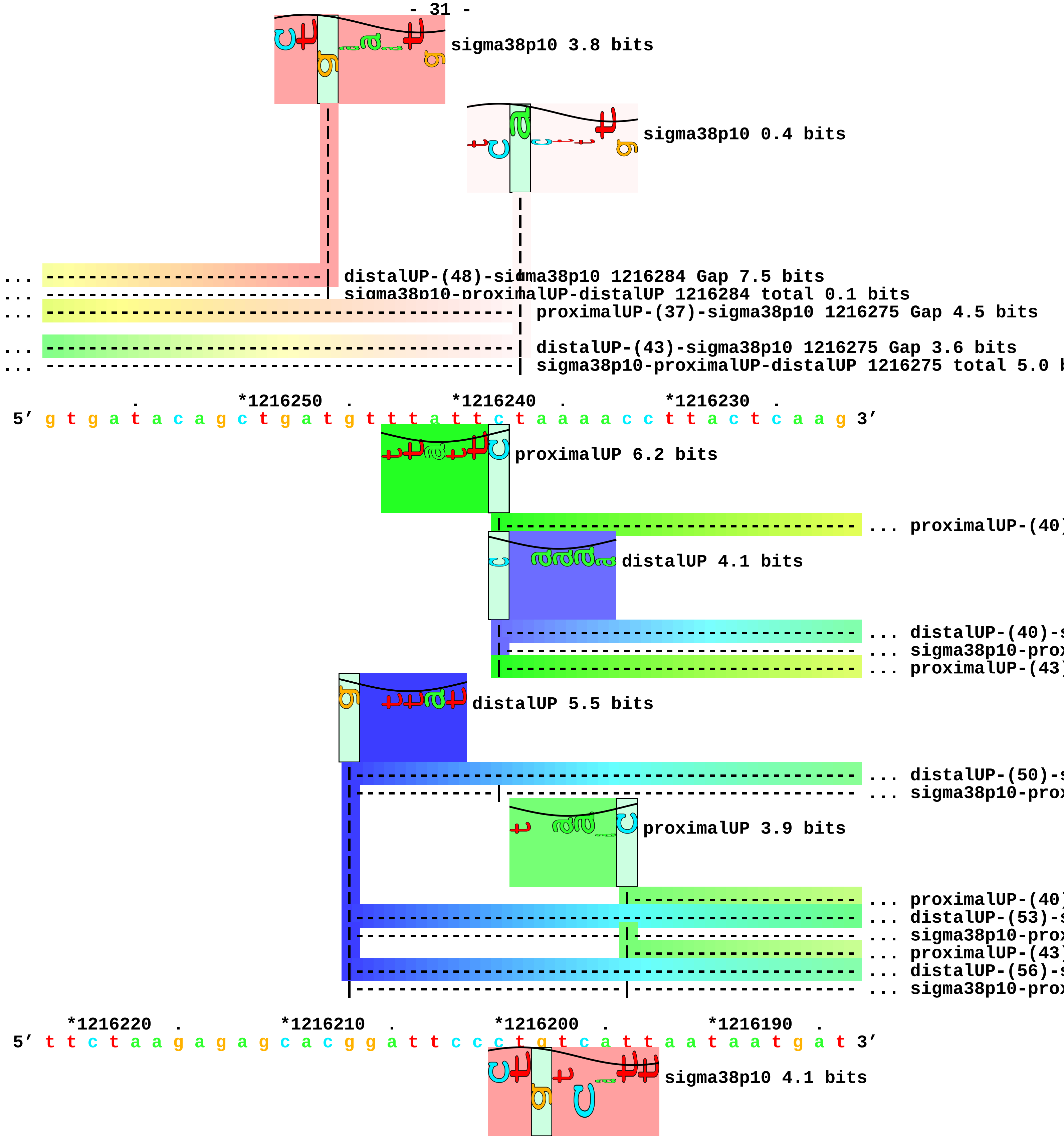
sigma38p10 3.8 bits
sigma38p10 0.4 bits
distalUP-(48)-sigma38p10 1216284 Gap 7.5 bits
sigma38p10-proximalUP-distalUP 1216284 total 0.1 bits
proximalUP-(37)-sigma38p10 1216275 Gap 4.5 bits
distalUP-(43)-sigma38p10 1216275 Gap 3.6 bits
sigma38p10-proximalUP-distalUP 1216275 total 5.0 b
*1216250
*1216240
*1216230
5′ g t g a t a c a g c t g a t g t t t a t t c t a a a a c c t t a c t c a a g 3′
proximalUP 6.2 bits
proximalUP-(40)
distalUP 4.1 bits
distalUP-(40)-s
sigma38p10-prox
proximalUP-(43)
distalUP 5.5 bits
distalUP-(50)-s
sigma38p10-prox
proximalUP 3.9 bits
proximalUP-(40)
distalUP-(53)-s
sigma38p10-prox
proximalUP-(43)
distalUP-(56)-s
sigma38p10-prox
*1216220
*1216210
*1216200
*1216190
5′ t t c t a a g a g a g c a c g g a t t c c c t g t c a t t a a t a a t g a t 3′
sigma38p10 4.1 bits

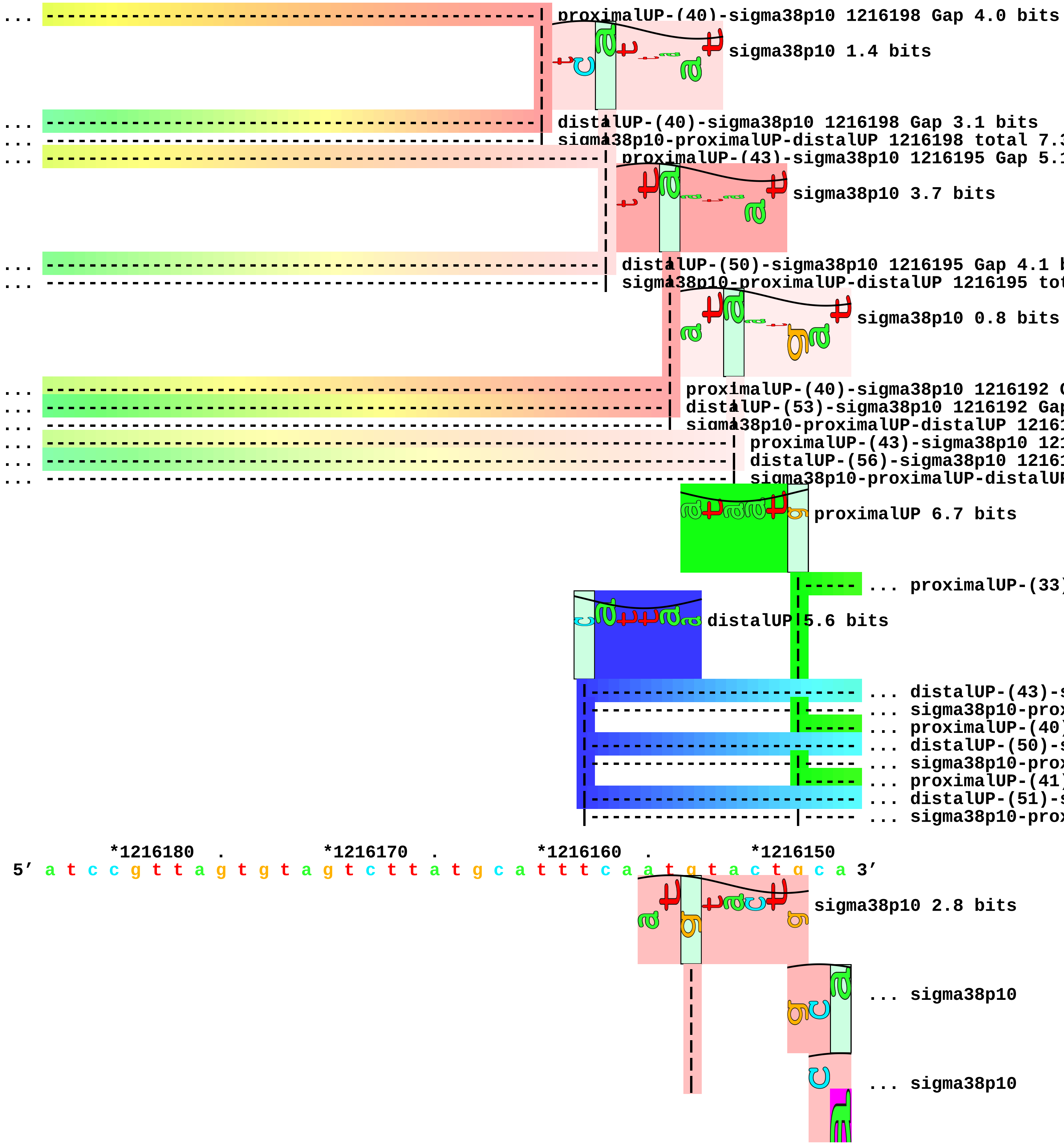
proximalUP-(40)-sigma38p10 1216198 Gap 4.0 bits
sigma38p10 1.4 bits
distalUP-(40)-sigma38p10 1216198 Gap 3.1 bits
sigma38p10-proximalUP-distalUP 1216198 total 7.3
proximalUP-(43)-sigma38p10 1216195 Gap 5.1
sigma38p10 3.7 bits
distalUP-(50)-sigma38p10 1216195 Gap 4.1 b
sigma38p10-proximalUP-distalUP 1216195 tot
sigma38p10 0.8 bits
proximalUP-(40)-sigma38p10 1216192 G
distalUP-(53)-sigma38p10 1216192 Gap
sigma38p10-proximalUP-distalUP 12161
proximalUP-(43)-sigma38p10 121
distalUP-(56)-sigma38p10 1216
sigma38p10-proximalUP-distalUP
proximalUP 6.7 bits
... proximalUP-(33)
distalUP 5.6 bits
... distalUP-(43)-s
... sigma38p10-prox
... proximalUP-(40)
... distalUP-(50)-s
... sigma38p10-prox
... proximalUP-(41)
... distalUP-(51)-s
... sigma38p10-prox
*1216180 . *1216170 . *1216160 . *1216150
5′ a t c c g t t a g t g t a g t c t t a t g c a t t t c a a t g t a c t g c a 3′
sigma38p10 2.8 bits
... sigma38p10
... sigma38p10

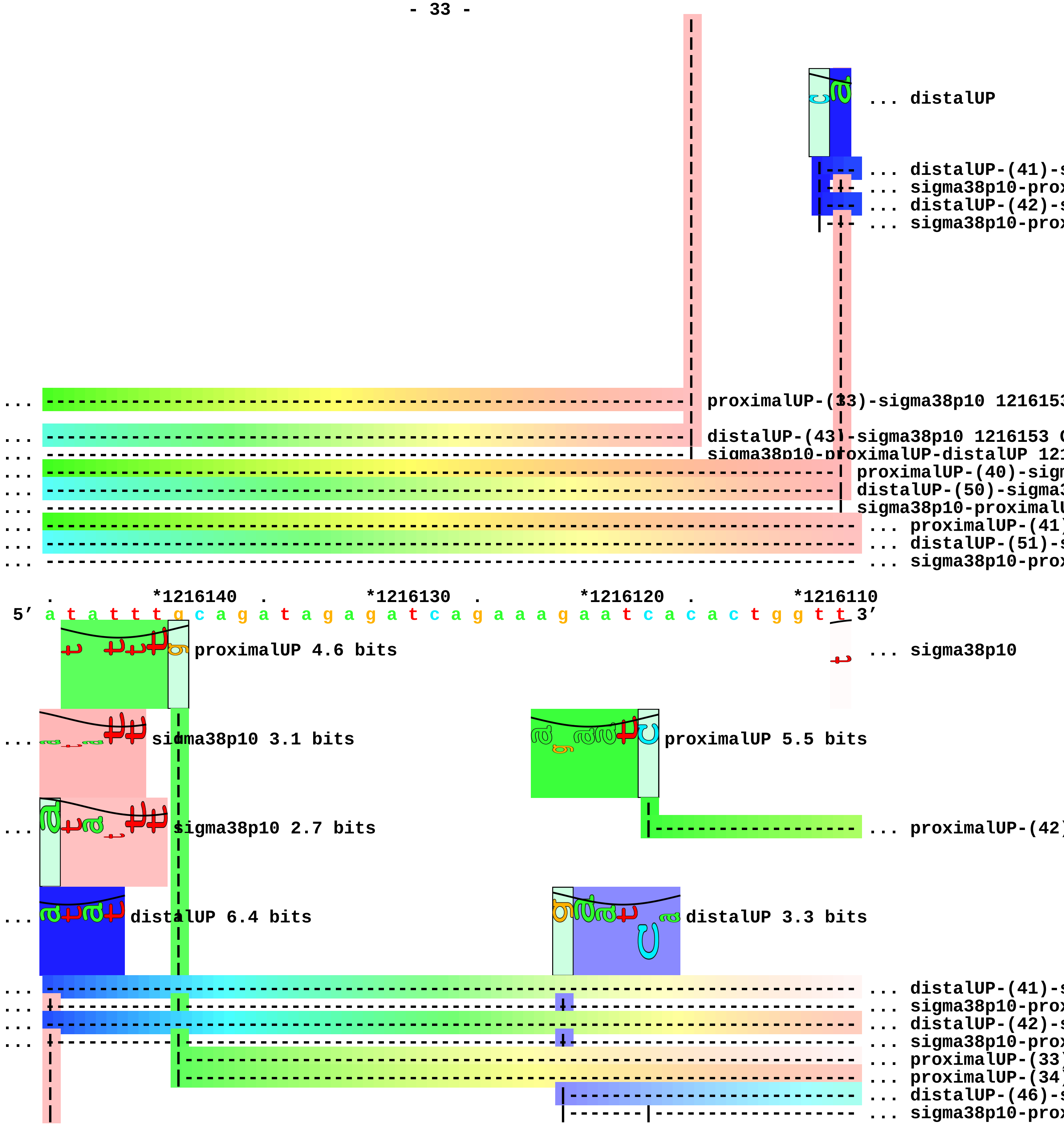
... distalUP
... distalUP-(41)-s
... sigma38p10-prox
... distalUP-(42)-s
... sigma38p10-prox
... proximalUP-(33)-sigma38p10 1216153
... distalUP-(43)-sigma38p10 1216153 (
... sigma38p10-proximalUP-distalUP 121
... proximalUP-(40)-sign
... distalUP-(50)-sigma3
... sigma38p10-proximalU
... proximalUP-(41)
... distalUP-(51)-s
... sigma38p10-prox
*1216140
*1216130
*1216120
*1216110
5′ a t a t t t g c a g a t a g a g a t c a g a a a g a a t c a c a c t g g t t 3′
proximalUP 4.6 bits
... sigma38p10
... sigma38p10 3.1 bits
proximalUP 5.5 bits
... sigma38p10 2.7 bits
... proximalUP-(42)
... distalUP 6.4 bits
distalUP 3.3 bits
... distalUP-(41)-s
... sigma38p10-prox
... distalUP-(42)-s
... sigma38p10-prox
... proximalUP-(33)
... proximalUP-(34)
... distalUP-(46)-s
... sigma38p10-prox

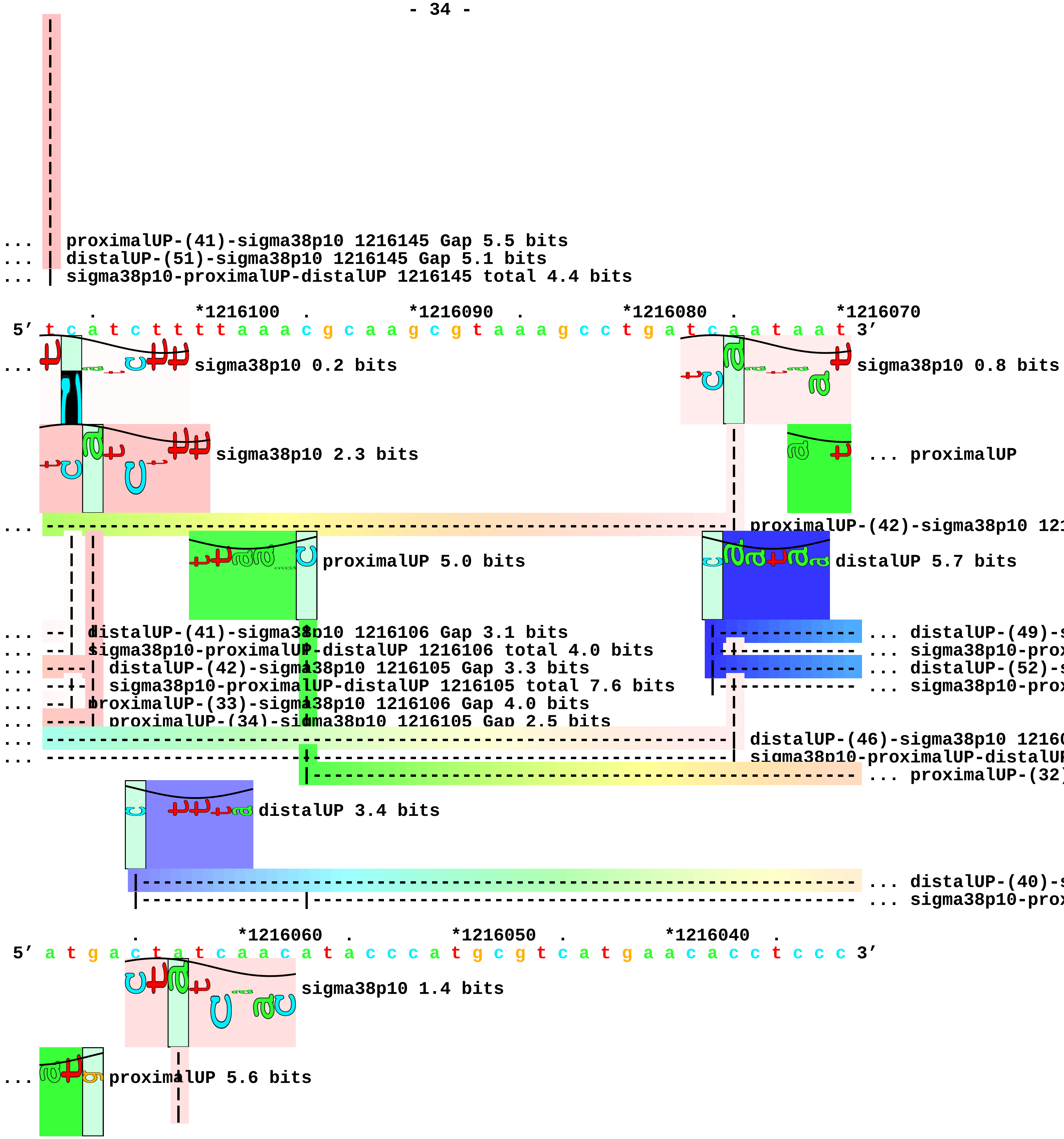
... proximalUP-(41)-sigma38p10 1216145 Gap 5.5 bits
... distalUP-(51)-sigma38p10 1216145 Gap 5.1 bits
... sigma38p10-proximalUP-distalUP 1216145 total 4.4 bits
*1216100
*1216090
*1216080
*1216070
5′ t c a t c t t t t a a a c g c a a g c g t a a a g c c t g a t c a a t a a t 3′
sigma38p10 0.2 bits
sigma38p10 0.8 bits
sigma38p10 2.3 bits
... proximalUP
proximalUP 5.0 bits
distalUP 5.7 bits
distalUP-(41)-sigma38p10 1216106 Gap 3.1 bits
sigma38p10-proximalUP-distalUP 1216106 total 4.0 bits
distalUP-(42)-sigma38p10 1216105 Gap 3.3 bits
sigma38p10-proximalUP-distalUP 1216105 total 7.6 bits
proximalUP-(33)-sigma38p10 1216106 Gap 4.0 bits
proximalUP-(34)-sigma38p10 1216105 Gap 2.5 bits
distalUP 3.4 bits
*1216060
*1216050
*1216040
5′ a t g a c t a t c a a c a t a c c c a t g c g t c a t g a a c a c c t c c c 3′
sigma38p10 1.4 bits
proximalUP 5.6 bits

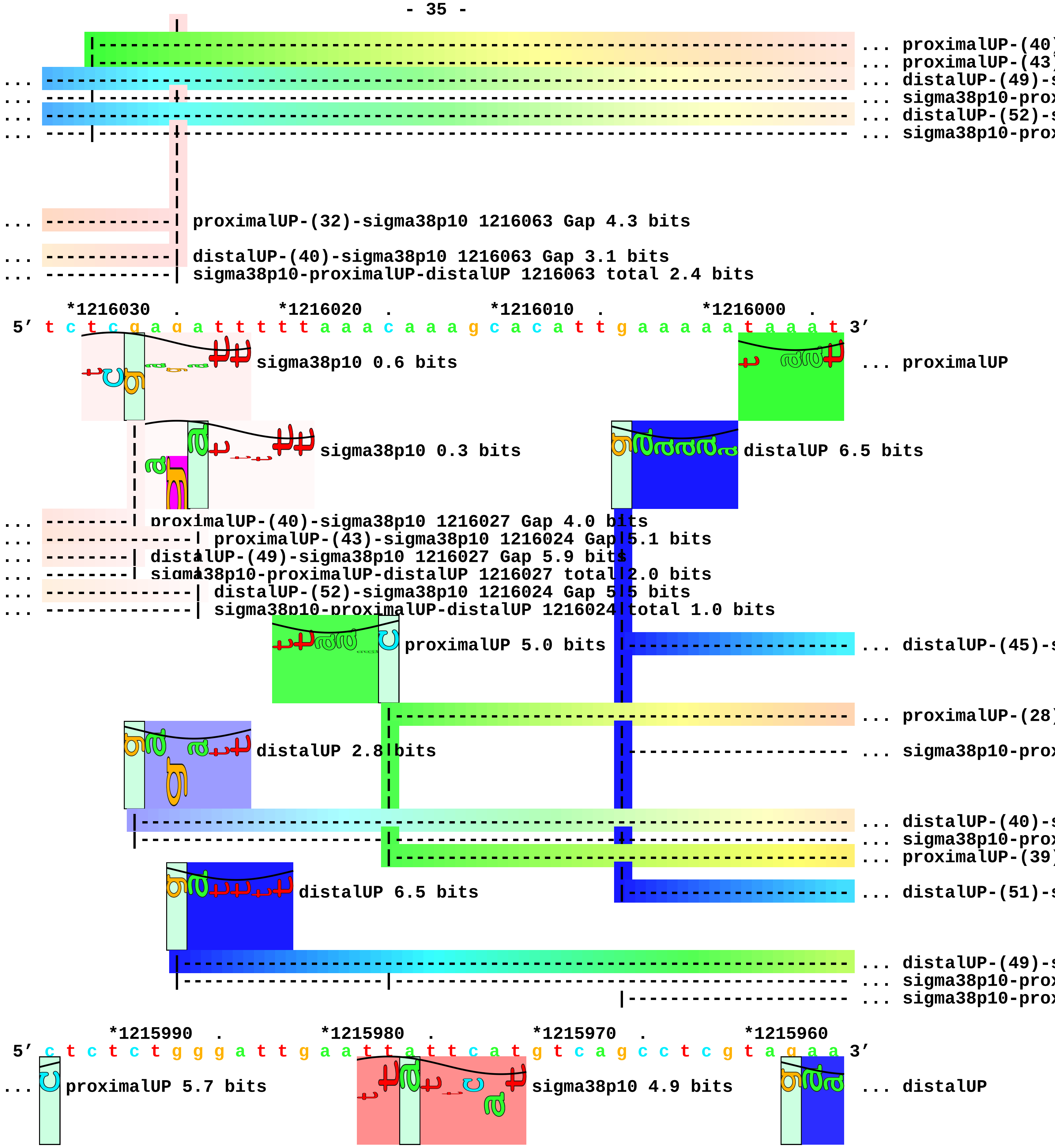

... proximalUP-(40)
... proximalUP-(43)
... distalUP-(49)-s
... sigma38p10-prox
... distalUP-(52)-s
... sigma38p10-prox
... proximalUP-(32)-sigma38p10 1216063 Gap 4.3 bits
... distalUP-(40)-sigma38p10 1216063 Gap 3.1 bits
... sigma38p10-proximalUP-distalUP 1216063 total 2.4 bits
*1216030 . *1216020 . *1216010 . *1216000 .
5' t c t c g a g a t t t t t a a a c a a a g c a c a t t g a a a a a t a a a t 3'
sigma38p10 0.6 bits
... proximalUP
sigma38p10 0.3 bits
distalUP 6.5 bits
... proximalUP-(40)-sigma38p10 1216027 Gap 4.0 bits
... proximalUP-(43)-sigma38p10 1216024 Gap 5.1 bits
... distalUP-(49)-sigma38p10 1216027 Gap 5.9 bits
... sigma38p10-proximalUP-distalUP 1216027 total 2.0 bits
... distalUP-(52)-sigma38p10 1216024 Gap 5.5 bits
... sigma38p10-proximalUP-distalUP 1216024 total 1.0 bits
proximalUP 5.0 bits
... distalUP-(45)-s
... proximalUP-(28)
distalUP 2.8 bits
... sigma38p10-prox
... distalUP-(40)-s
... sigma38p10-prox
... proximalUP-(39)
distalUP 6.5 bits
... distalUP-(51)-s
... distalUP-(49)-s
... sigma38p10-prox
... sigma38p10-prox
*1215990 . *1215980 . *1215970 . *1215960
5' c t c t c t g g g a t t g a a t t a t t c a t g t c a g c c t c g t a g a a 3'
... proximalUP 5.7 bits
sigma38p10 4.9 bits
... distalUP

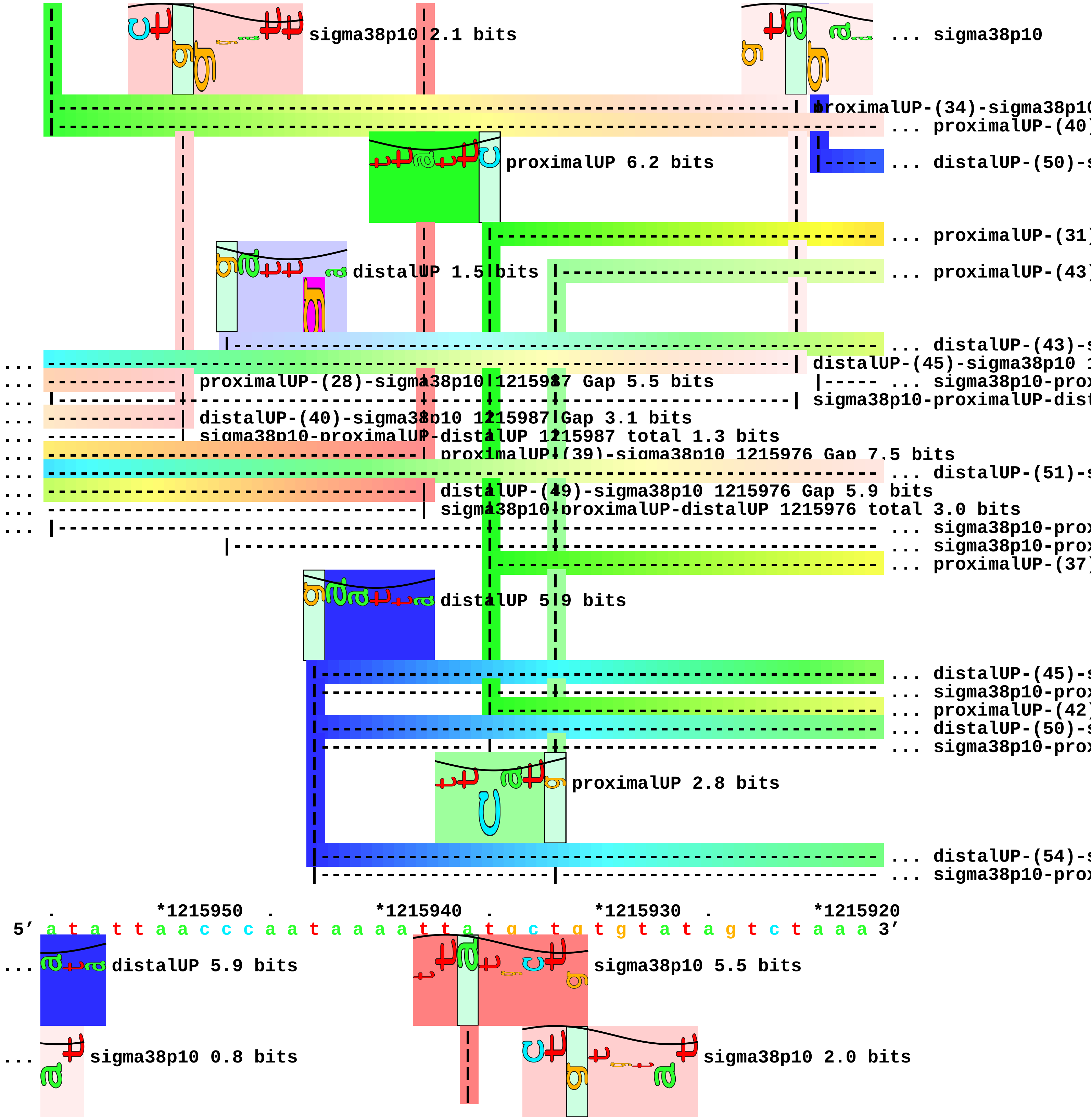
sigma38p10 2.1 bits
... sigma38p10
proximalUP-(34)-sigma38p10
... proximalUP-(40)
proximalUP 6.2 bits
... distalUP-(50)-s
... proximalUP-(31)
distalUP 1.5 bits
... proximalUP-(43)
... distalUP-(43)-s
distalUP-(45)-sigma38p10 1
proximalUP-(28)-sigma38p10 1215987 Gap 5.5 bits
... sigma38p10-prox
sigma38p10-proximalUP-dist
distalUP-(40)-sigma38p10 1215987 Gap 3.1 bits
sigma38p10-proximalUP-distalUP 1215987 total 1.3 bits
proximalUP-(39)-sigma38p10 1215976 Gap 7.5 bits
... distalUP-(51)-s
distalUP-(49)-sigma38p10 1215976 Gap 5.9 bits
sigma38p10-proximalUP-distalUP 1215976 total 3.0 bits
... sigma38p10-prox
... sigma38p10-prox
... proximalUP-(37)
distalUP 5.9 bits
... distalUP-(45)-s
... sigma38p10-prox
... proximalUP-(42)
... distalUP-(50)-s
... sigma38p10-prox
proximalUP 2.8 bits
... distalUP-(54)-s
... sigma38p10-prox
. *1215950 . *1215940 . *1215930 . *1215920
5' a t a t t a a c c c a a t a a a a t t a t g c t g t g t a t a g t c t a a a 3'
... distalUP 5.9 bits
sigma38p10 5.5 bits
... sigma38p10 0.8 bits
sigma38p10 2.0 bits

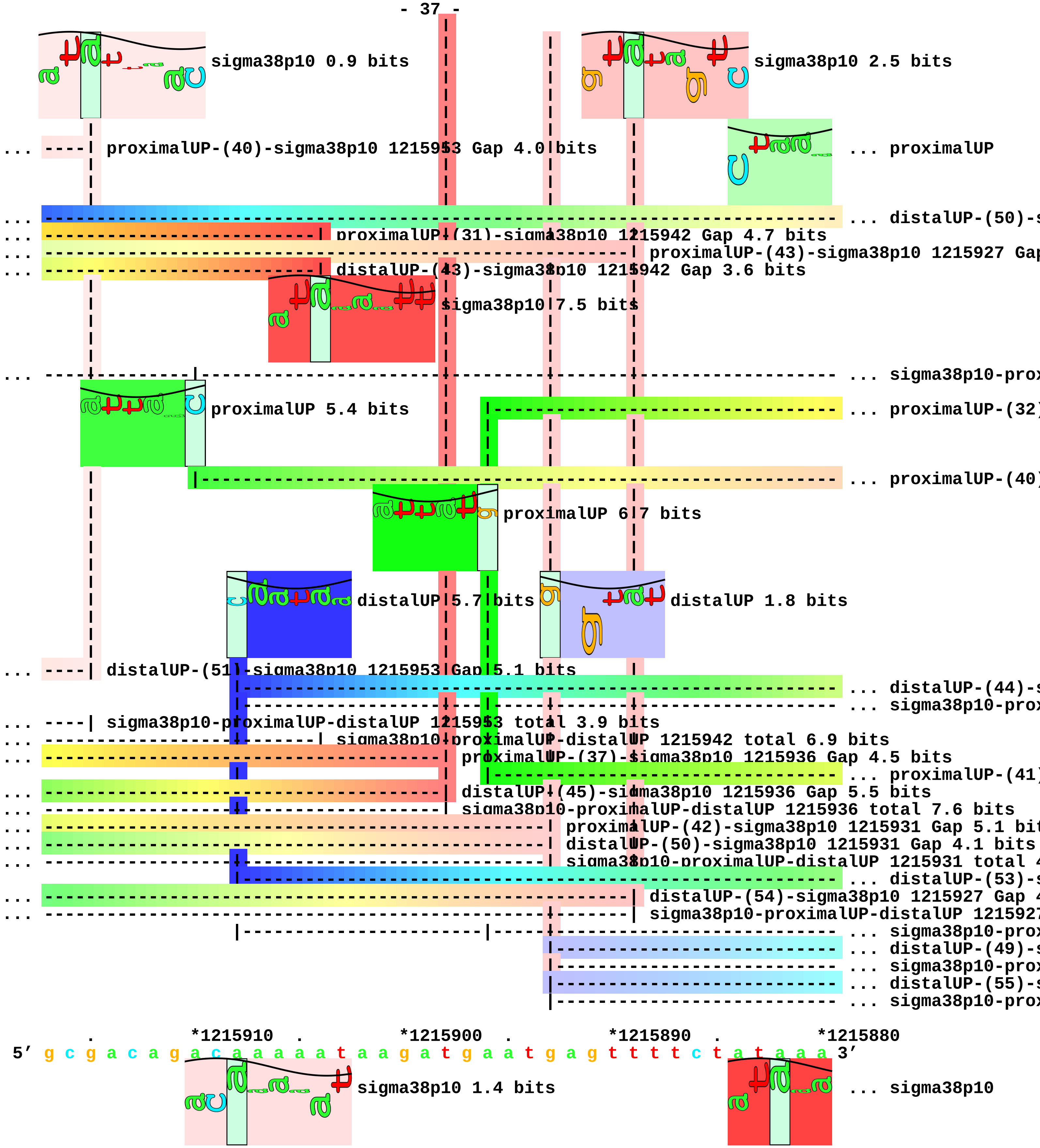
sigma38p10 0.9 bits
sigma38p10 2.5 bits
... ----| proximalUP-(40)-sigma38p10 1215953 Gap 4.0 bits
... proximalUP
... distalUP-(50)-s
proximalUP-(31)-sigma38p10 1215942 Gap 4.7 bits
proximalUP-(43)-sigma38p10 1215927 Gap
distalUP-(43)-sigma38p10 1215942 Gap 3.6 bits
sigma38p10 7.5 bits
... sigma38p10-prox
proximalUP 5.4 bits
... proximalUP-(32)
... proximalUP-(40)
proximalUP 6.7 bits
distalUP 5.7 bits
distalUP 1.8 bits
... ----| distalUP-(51)-sigma38p10 1215953 Gap 5.1 bits
... distalUP-(44)-s
... sigma38p10-prox
... ----| sigma38p10-proximalUP-distalUP 1215953 total 3.9 bits
sigma38p10-proximalUP-distalUP 1215942 total 6.9 bits
proximalUP-(37)-sigma38p10 1215936 Gap 4.5 bits
... proximalUP-(41)
distalUP-(45)-sigma38p10 1215936 Gap 5.5 bits
sigma38p10-proximalUP-distalUP 1215936 total 7.6 bits
proximalUP-(42)-sigma38p10 1215931 Gap 5.1 bit
distalUP-(50)-sigma38p10 1215931 Gap 4.1 bits
sigma38p10-proximalUP-distalUP 1215931 total 4
... distalUP-(53)-s
distalUP-(54)-sigma38p10 1215927 Gap 4
sigma38p10-proximalUP-distalUP 1215927
... sigma38p10-prox
... distalUP-(49)-s
... sigma38p10-prox
... distalUP-(55)-s
... sigma38p10-prox
. *1215910 . *1215900 . *1215890 . *1215880
5′ g c g a c a g a c a a a a a t a a g a t g a a t g a g t t t t c t a t a a a 3′
sigma38p10 1.4 bits
... sigma38p10

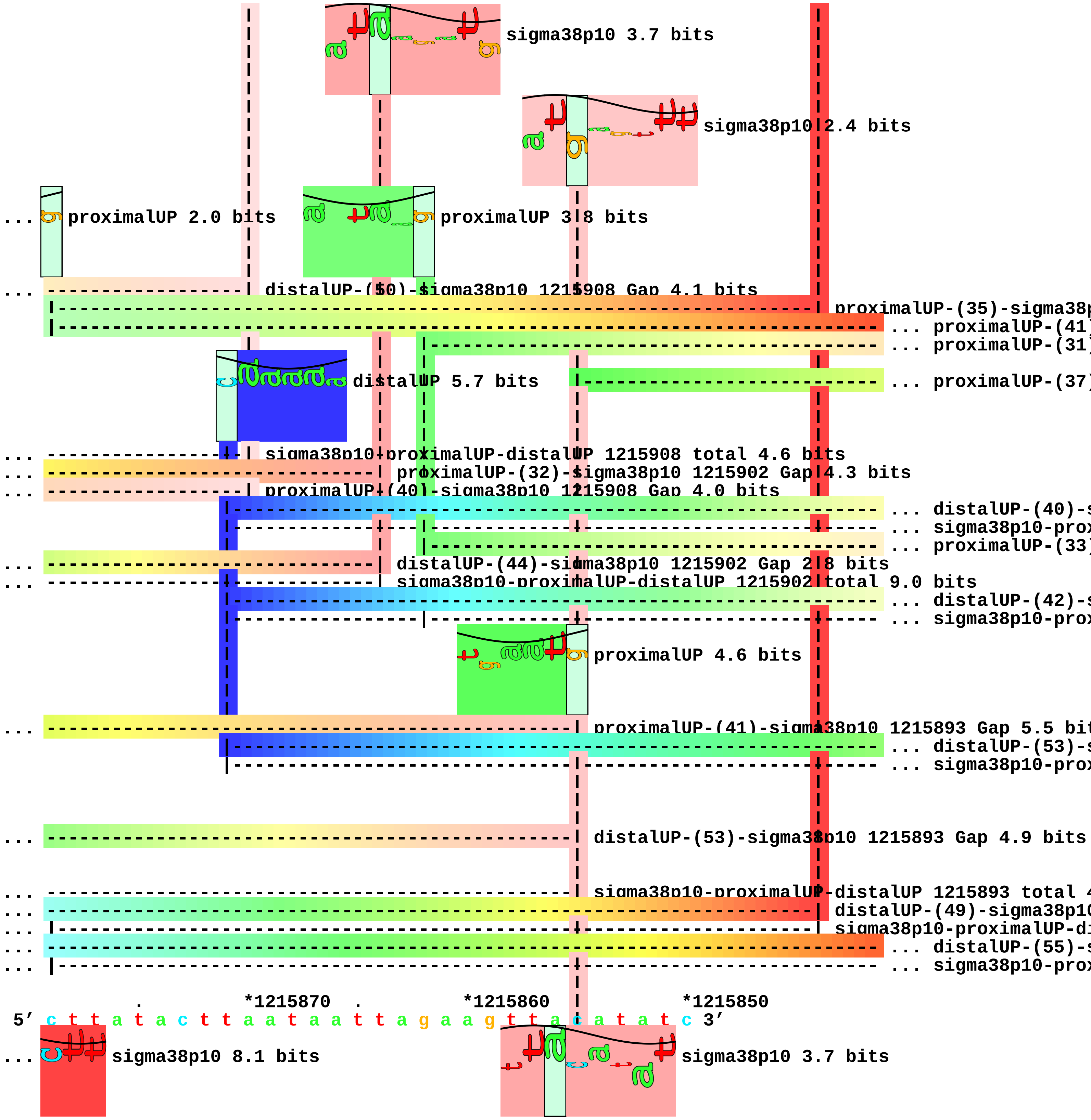
sigma38p10 3.7 bits
sigma38p10 2.4 bits
... proximalUP 2.0 bits
proximalUP 3.8 bits
... distalUP-(50)-sigma38p10 1215908 Gap 4.1 bits
proximalUP-(35)-sigma38p
... proximalUP-(41)
... proximalUP-(31)
distalUP 5.7 bits
... proximalUP-(37)
... sigma38p10-proximalUP-distalUP 1215908 total 4.6 bits
... proximalUP-(32)-sigma38p10 1215902 Gap 4.3 bits
... proximalUP-(40)-sigma38p10 1215908 Gap 4.0 bits
... distalUP-(40)-s
... sigma38p10-prox
... proximalUP-(33)
... distalUP-(44)-sigma38p10 1215902 Gap 2.8 bits
... sigma38p10-proximalUP-distalUP 1215902 total 9.0 bits
... distalUP-(42)-s
... sigma38p10-prox
proximalUP 4.6 bits
... proximalUP-(41)-sigma38p10 1215893 Gap 5.5 bit
... distalUP-(53)-s
... sigma38p10-prox
... distalUP-(53)-sigma38p10 1215893 Gap 4.9 bits
... sigma38p10-proximalUP-distalUP 1215893 total 4
... distalUP-(49)-sigma38p10
... sigma38p10-proximalUP-di
... distalUP-(55)-s
... sigma38p10-prox
. *1215870 . *1215860 . *1215850
5' c t t a t a c t t a a t a a t t a g a a g t t a c a t a t c 3'
... sigma38p10 8.1 bits
sigma38p10 3.7 bits

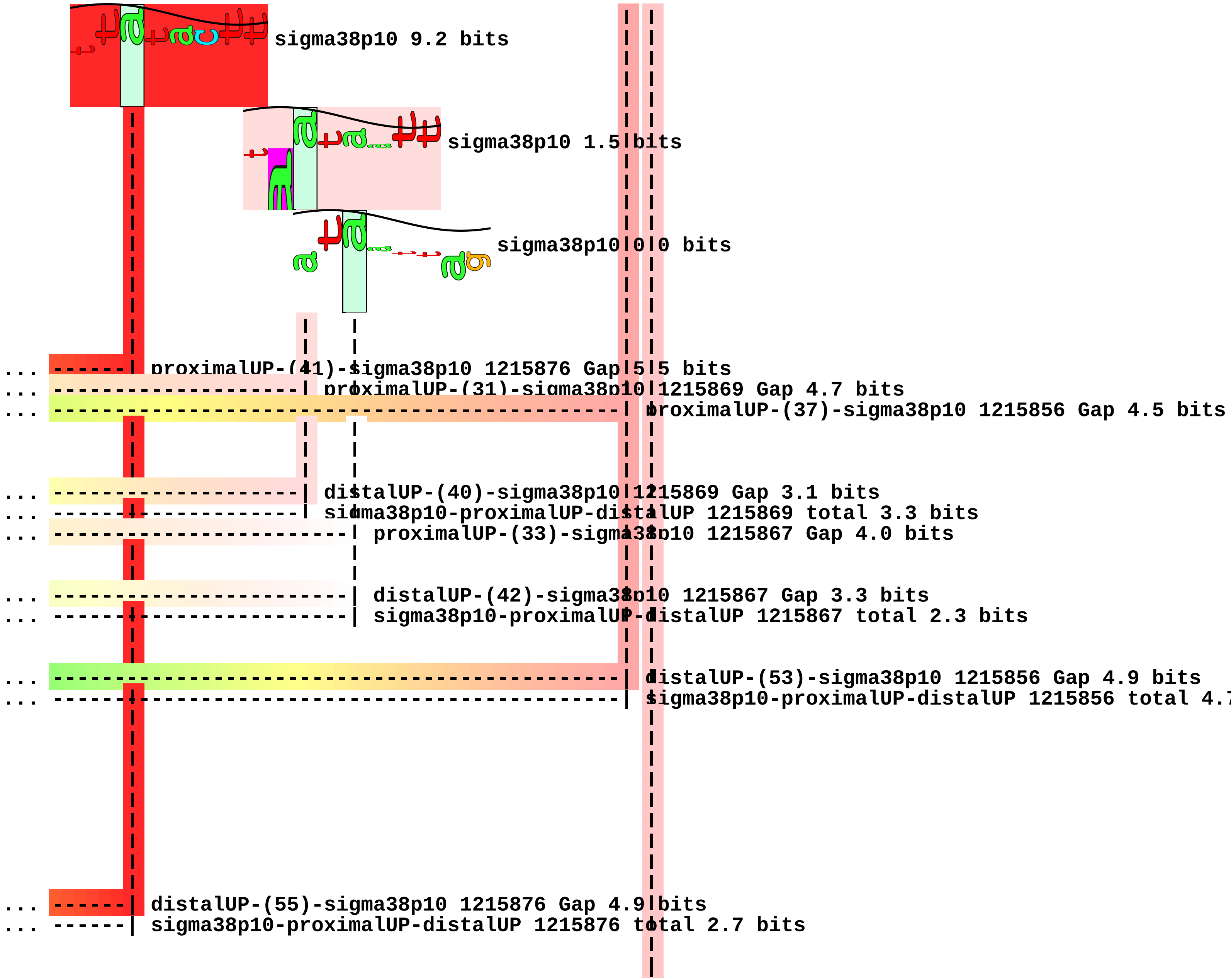
sigma38p10 9.2 bits
sigma38p10 1.5 bits
sigma38p10 0.0 bits
... proximalUP-(41)-sigma38p10 1215876 Gap 5.5 bits
... proximalUP-(31)-sigma38p10 1215869 Gap 4.7 bits
... proximalUP-(37)-sigma38p10 1215856 Gap 4.5 bits
... distalUP-(40)-sigma38p10 1215869 Gap 3.1 bits
... sigma38p10-proximalUP-distalUP 1215869 total 3.3 bits
... proximalUP-(33)-sigma38p10 1215867 Gap 4.0 bits
... distalUP-(42)-sigma38p10 1215867 Gap 3.3 bits
... sigma38p10-proximalUP-distalUP 1215867 total 2.3 bits
... distalUP-(53)-sigma38p10 1215856 Gap 4.9 bits
... sigma38p10-proximalUP-distalUP 1215856 total 4.
... distalUP-(55)-sigma38p10 1215876 Gap 4.9 bits
... sigma38p10-proximalUP-distalUP 1215876 total 2.7 bits

piece 12, NC_000913, ycgH, config: linear, direction: -, begin: 1219999, end: 1219350

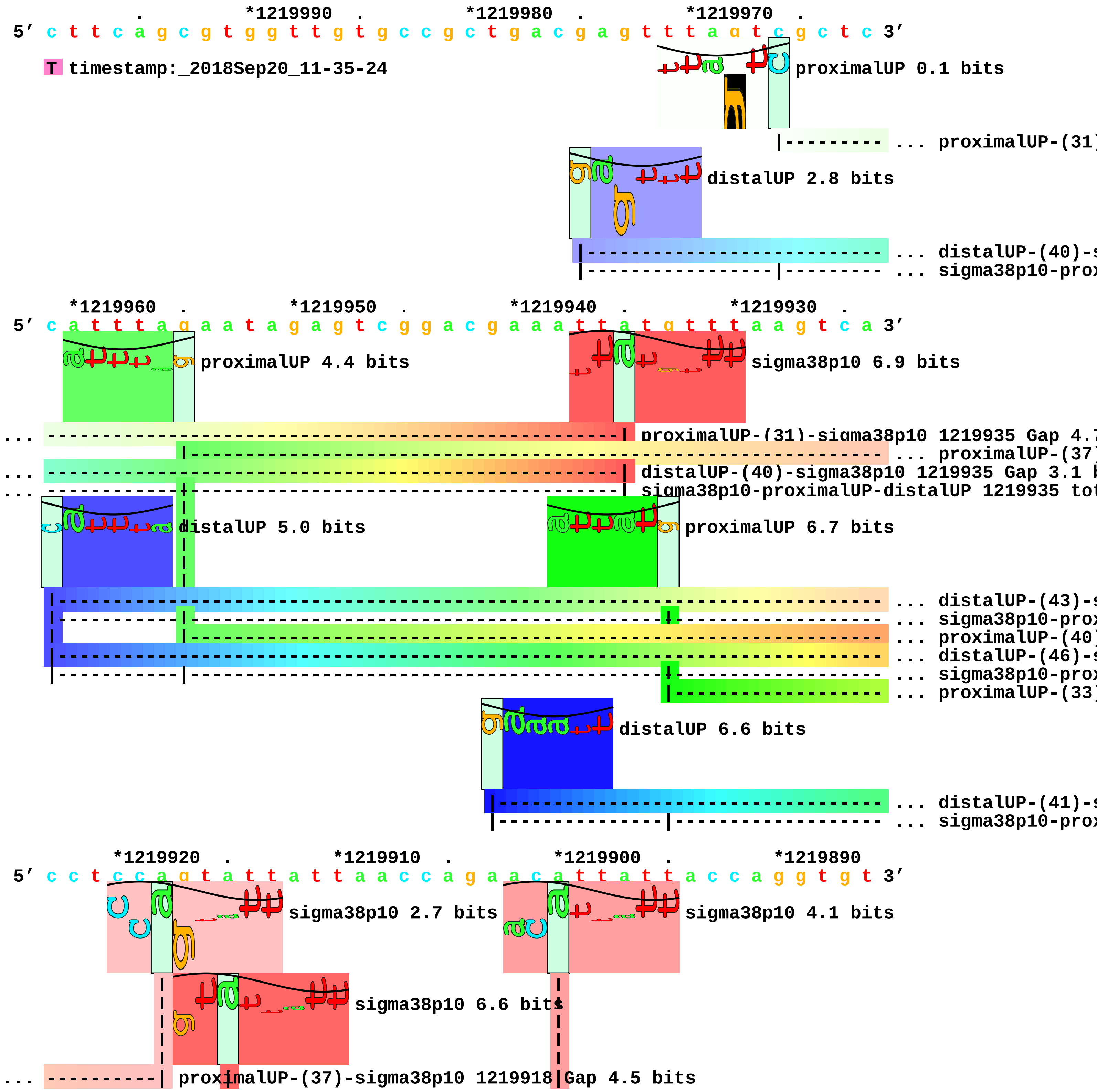

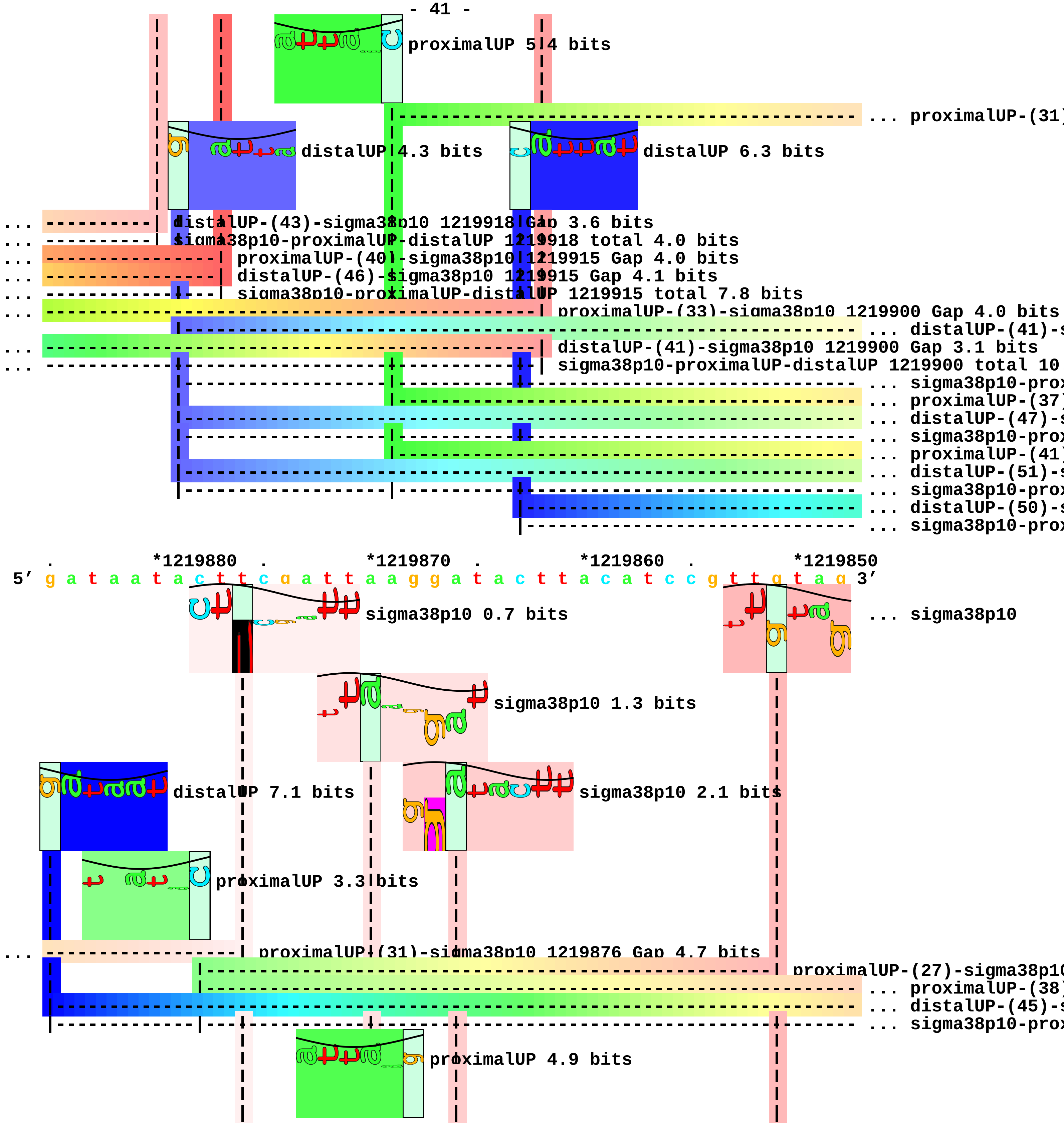
proximalUP 5.4 bits
... proximalUP-(31)
distalUP 4.3 bits
distalUP 6.3 bits
... distalUP-(43)-sigma38p10 1219918 Gap 3.6 bits
... sigma38p10-proximalUP-distalUP 1219918 total 4.0 bits
... proximalUP-(40)-sigma38p10 1219915 Gap 4.0 bits
... distalUP-(46)-sigma38p10 1219915 Gap 4.1 bits
... sigma38p10-proximalUP-distalUP 1219915 total 7.8 bits
... proximalUP-(33)-sigma38p10 1219900 Gap 4.0 bits
... distalUP-(41)-s
... distalUP-(41)-sigma38p10 1219900 Gap 3.1 bits
... sigma38p10-proximalUP-distalUP 1219900 total 10.
... sigma38p10-prox
... proximalUP-(37)
... distalUP-(47)-s
... sigma38p10-prox
... proximalUP-(41)
... distalUP-(51)-s
... sigma38p10-prox
... distalUP-(50)-s
... sigma38p10-prox
*1219880
*1219870
*1219860
*1219850
5' g a t a a t a c t t c g a t t a a g g a t a c t t a c a t c c g t t g t a g 3'
sigma38p10 0.7 bits
... sigma38p10
sigma38p10 1.3 bits
distalUP 7.1 bits
sigma38p10 2.1 bits
proximalUP 3.3 bits
... proximalUP-(31)-sigma38p10 1219876 Gap 4.7 bits
proximalUP-(27)-sigma38p10
... proximalUP-(38)
... distalUP-(45)-s
... sigma38p10-prox
proximalUP 4.9 bits

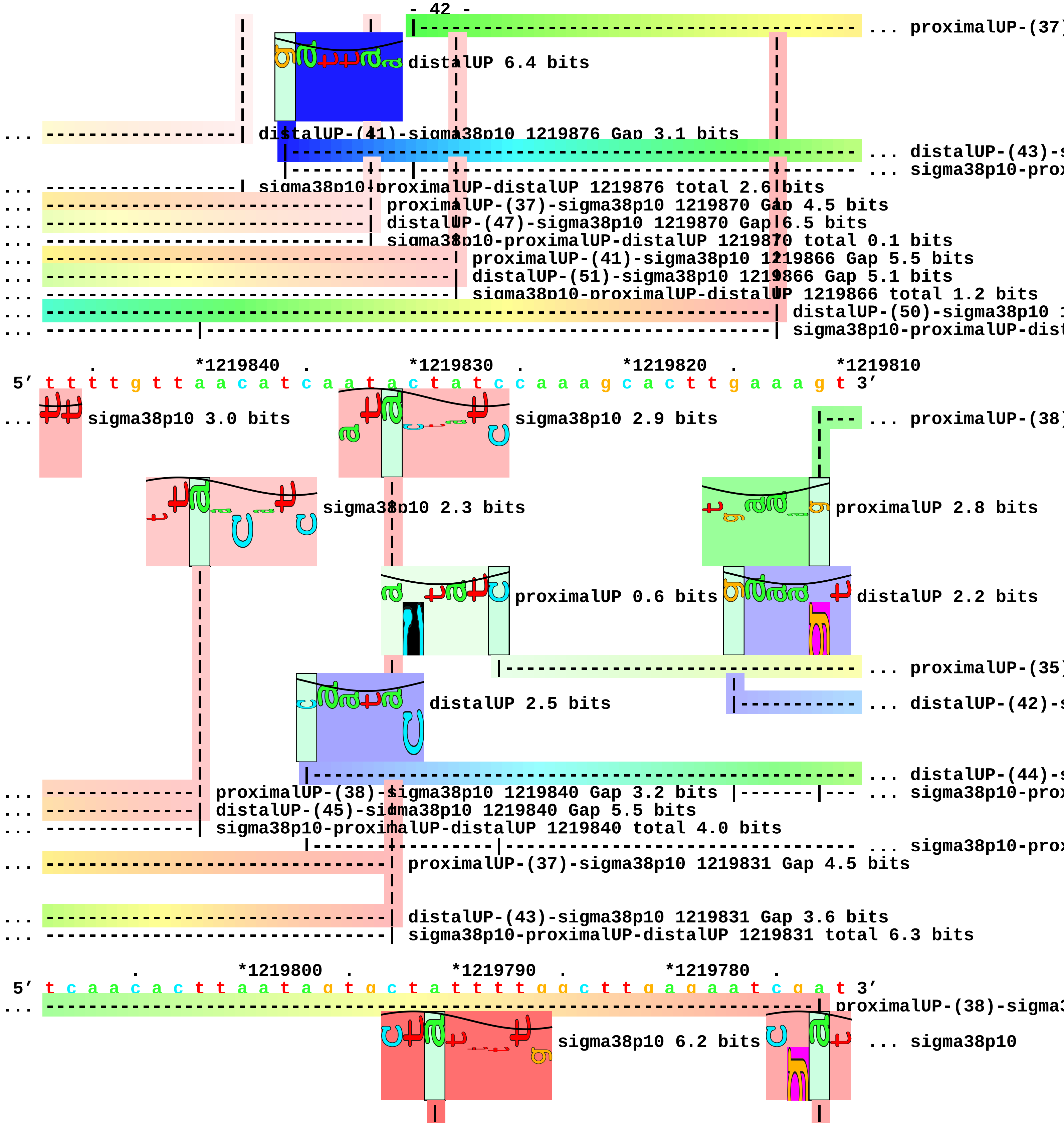

... proximalUP-(37)
distalUP 6.4 bits
... distalUP-(41)-sigma38p10 1219876 Gap 3.1 bits
... distalUP-(43)-s
... sigma38p10-prox
... sigma38p10-proximalUP-distalUP 1219876 total 2.6 bits
... proximalUP-(37)-sigma38p10 1219870 Gap 4.5 bits
... distalUP-(47)-sigma38p10 1219870 Gap 6.5 bits
... sigma38p10-proximalUP-distalUP 1219870 total 0.1 bits
... proximalUP-(41)-sigma38p10 1219866 Gap 5.5 bits
... distalUP-(51)-sigma38p10 1219866 Gap 5.1 bits
... sigma38p10-proximalUP-distalUP 1219866 total 1.2 bits
... distalUP-(50)-sigma38p10 1
... sigma38p10-proximalUP-dist
*1219840 *1219830 *1219820 *1219810
5' t t t t g t t a a c a t c a a t a c t a t c c a a a g c a c t t g a a a g t 3'
... sigma38p10 3.0 bits
sigma38p10 2.9 bits
... proximalUP-(38)
sigma38p10 2.3 bits
proximalUP 2.8 bits
proximalUP 0.6 bits
distalUP 2.2 bits
... proximalUP-(35)
distalUP 2.5 bits
... distalUP-(42)-s
... distalUP-(44)-s
... proximalUP-(38)-sigma38p10 1219840 Gap 3.2 bits
... sigma38p10-prox
... distalUP-(45)-sigma38p10 1219840 Gap 5.5 bits
... sigma38p10-proximalUP-distalUP 1219840 total 4.0 bits
... sigma38p10-prox
... proximalUP-(37)-sigma38p10 1219831 Gap 4.5 bits
... distalUP-(43)-sigma38p10 1219831 Gap 3.6 bits
... sigma38p10-proximalUP-distalUP 1219831 total 6.3 bits
*1219800 *1219790 *1219780
5' t c a a c a c t t a a t a g t g c t a t t t t g g c t t g a g a a t c g a t 3'
... proximalUP-(38)-sigma3
sigma38p10 6.2 bits
... sigma38p10

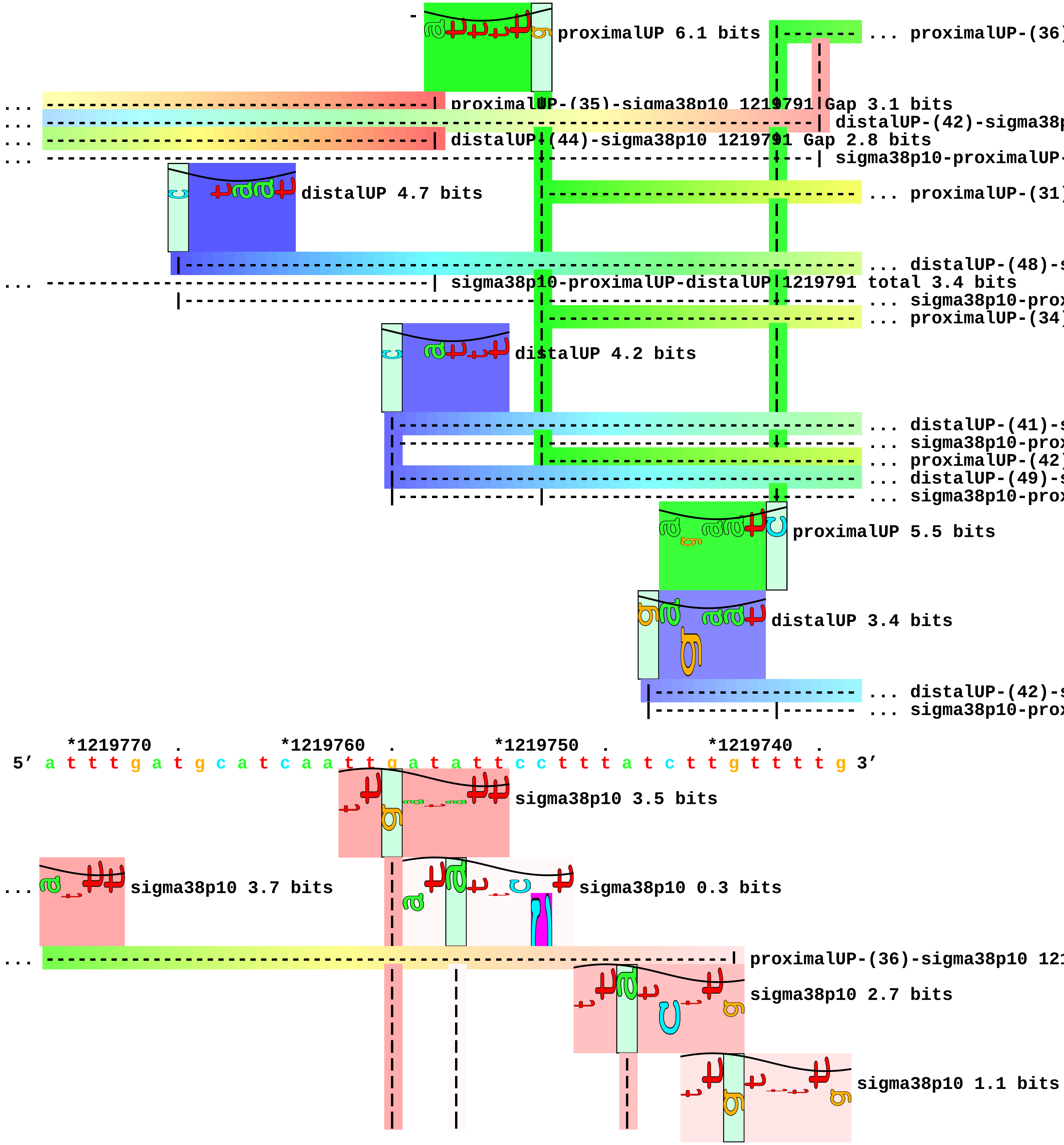
proximalUP 6.1 bits
... proximalUP-(36)
proximalUP-(35)-sigma38p10 1219791 Gap 3.1 bits
distalUP-(42)-sigma38p
distalUP-(44)-sigma38p10 1219791 Gap 2.8 bits
sigma38p10-proximalUP-
distalUP 4.7 bits
... proximalUP-(31)
... distalUP-(48)-s
sigma38p10-proximalUP-distalUP 1219791 total 3.4 bits
... sigma38p10-prox
... proximalUP-(34)
distalUP 4.2 bits
... distalUP-(41)-s
... sigma38p10-prox
... proximalUP-(42)
... distalUP-(49)-s
... sigma38p10-prox
proximalUP 5.5 bits
distalUP 3.4 bits
... distalUP-(42)-s
... sigma38p10-prox
*1219770 . *1219760 . *1219750 . *1219740 .
5′ a t t t g a t g c a t c a a t t g a t a t t c c t t t a t c t t g t t t t g 3′
sigma38p10 3.5 bits
sigma38p10 3.7 bits
sigma38p10 0.3 bits
proximalUP-(36)-sigma38p10 121
sigma38p10 2.7 bits
sigma38p10 1.1 bits

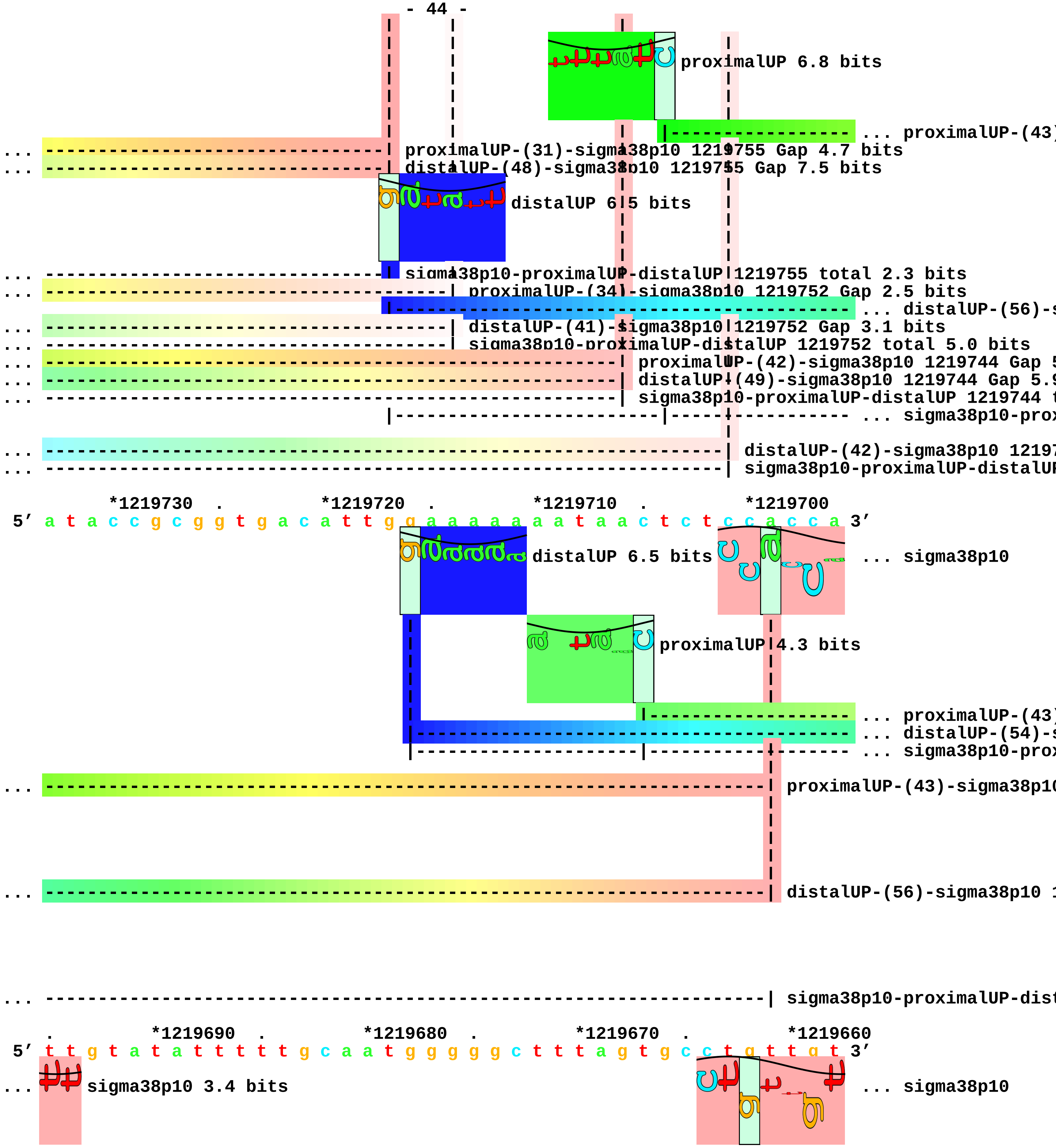
proximalUP 6.8 bits
... proximalUP-(43)
... proximalUP-(31)-sigma38p10 1219755 Gap 4.7 bits
... distalUP-(48)-sigma38p10 1219755 Gap 7.5 bits
distalUP 6.5 bits
... sigma38p10-proximalUP-distalUP 1219755 total 2.3 bits
... proximalUP-(34)-sigma38p10 1219752 Gap 2.5 bits
... distalUP-(56)-s
... distalUP-(41)-sigma38p10 1219752 Gap 3.1 bits
... sigma38p10-proximalUP-distalUP 1219752 total 5.0 bits
... proximalUP-(42)-sigma38p10 1219744 Gap 5
... distalUP-(49)-sigma38p10 1219744 Gap 5.9
... sigma38p10-proximalUP-distalUP 1219744 t
... sigma38p10-prox
... distalUP-(42)-sigma38p10 1219
... sigma38p10-proximalUP-distalUP
*1219730 . *1219720 . *1219710 . *1219700
5′ a t a c c g c g g t g a c a t t g g a a a a a a a t a a c t c t c c a c c a 3′
distalUP 6.5 bits
... sigma38p10
proximalUP 4.3 bits
... proximalUP-(43)
... distalUP-(54)-s
... sigma38p10-prox
... proximalUP-(43)-sigma38p10
... distalUP-(56)-sigma38p10 1
... sigma38p10-proximalUP-dist
. *1219690 . *1219680 . *1219670 . *1219660
5′ t t g t a t a t t t t t g c a a t g g g g g c t t t a g t g c c t g t t g t 3′
... sigma38p10 3.4 bits
... sigma38p10

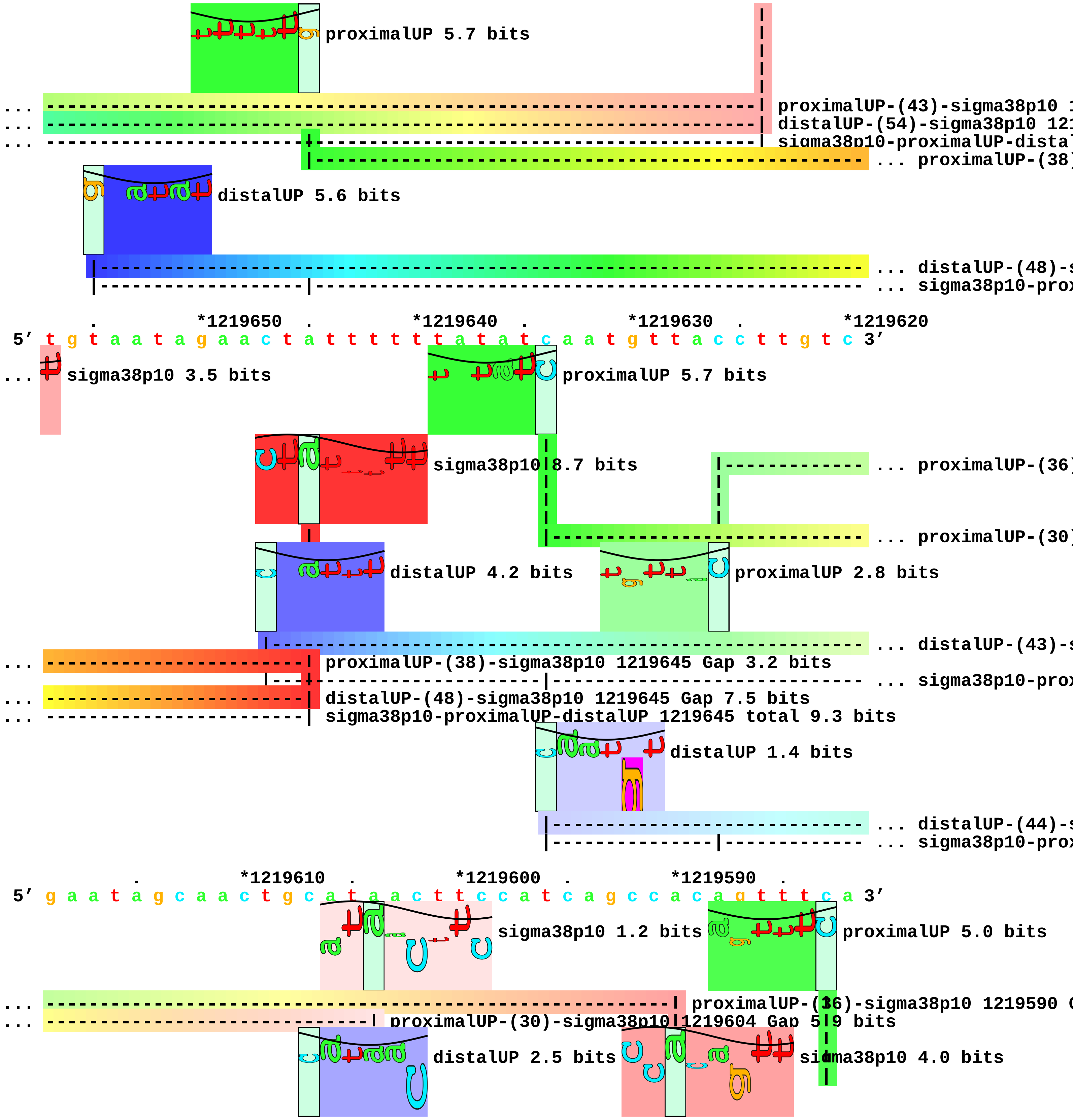
proximalUP 5.7 bits
proximalUP-(43)-sigma38p10 1
distalUP-(54)-sigma38p10 121
sigma38p10-proximalUP-dista
... proximalUP-(38)
distalUP 5.6 bits
... distalUP-(48)-s
... sigma38p10-prox
*1219650
*1219640
*1219630
*1219620
5' t g t a a t a g a a c t a t t t t t t a t a t c a a t g t t a c c t t g t c 3'
... sigma38p10 3.5 bits
proximalUP 5.7 bits
sigma38p10 8.7 bits
... proximalUP-(36)
... proximalUP-(30)
distalUP 4.2 bits
proximalUP 2.8 bits
... distalUP-(43)-s
proximalUP-(38)-sigma38p10 1219645 Gap 3.2 bits
... sigma38p10-prox
distalUP-(48)-sigma38p10 1219645 Gap 7.5 bits
sigma38p10-proximalUP-distalUP 1219645 total 9.3 bits
distalUP 1.4 bits
... distalUP-(44)-s
... sigma38p10-prox
*1219610
*1219600
*1219590
5' g a a t a g c a a c t g c a t a a c t t c c a t c a g c c a c a g t t t c a 3'
sigma38p10 1.2 bits
proximalUP 5.0 bits
proximalUP-(36)-sigma38p10 1219590 G
proximalUP-(30)-sigma38p10 1219604 Gap 5.9 bits
distalUP 2.5 bits
sigma38p10 4.0 bits

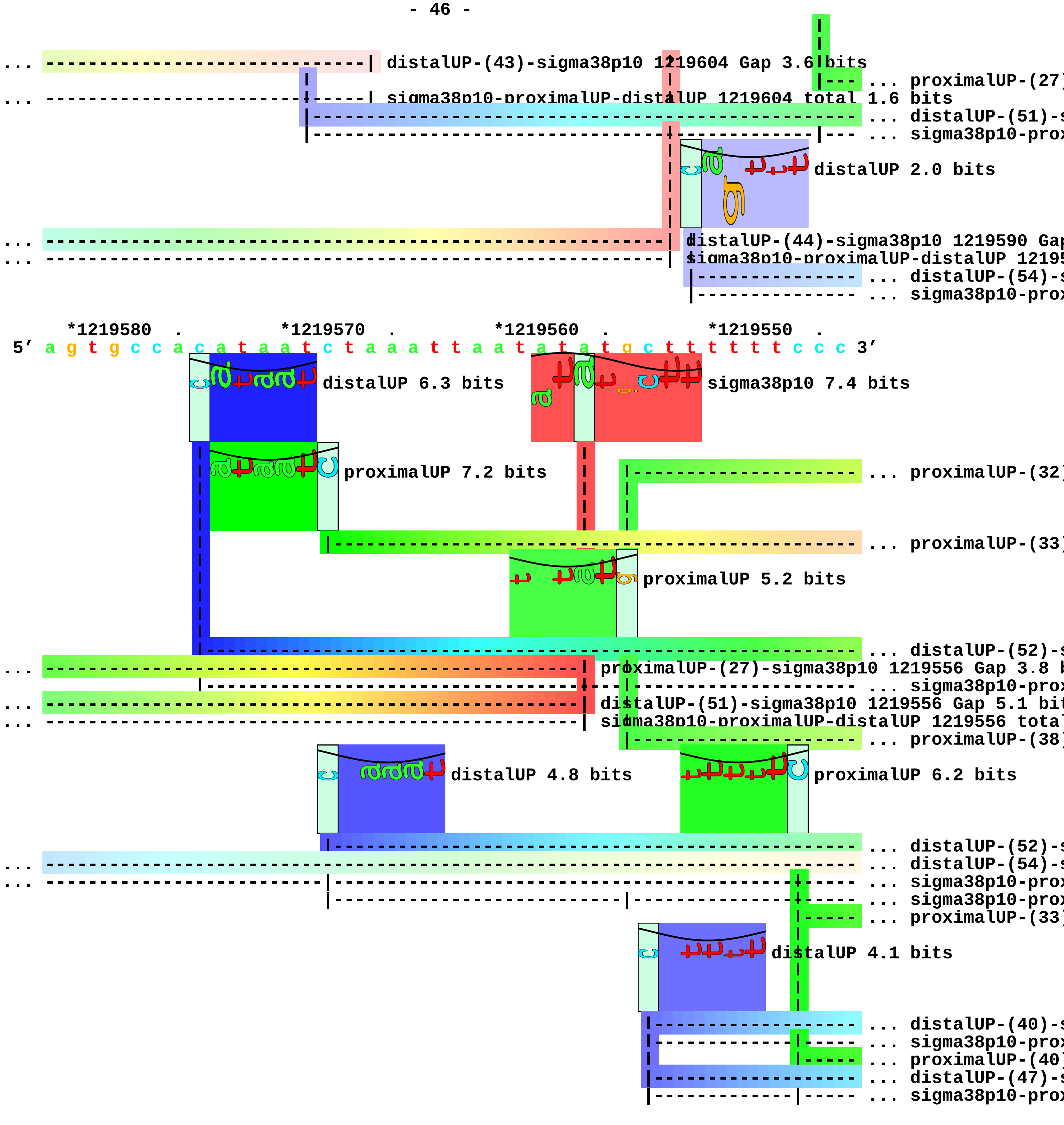
distalUP-(43)-sigma38p10 1219604 Gap 3.6 bits
sigma38p10-proximalUP-distalUP 1219604 total 1.6 bits
... proximalUP-(27)
... distalUP-(51)-s
... sigma38p10-prox
distalUP 2.0 bits
distalUP-(44)-sigma38p10 1219590 Gap
sigma38p10-proximalUP-distalUP 12195
... distalUP-(54)-s
... sigma38p10-prox
*1219580
*1219570
*1219560
*1219550
5' a g t g c c a c a t a a t c t a a a t t a a t a t a t g c t t t t t t c c c 3'
distalUP 6.3 bits
sigma38p10 7.4 bits
proximalUP 7.2 bits
... proximalUP-(32)
... proximalUP-(33)
proximalUP 5.2 bits
... distalUP-(52)-s
proximalUP-(27)-sigma38p10 1219556 Gap 3.8 b
... sigma38p10-prox
distalUP-(51)-sigma38p10 1219556 Gap 5.1 bit
sigma38p10-proximalUP-distalUP 1219556 tota
... proximalUP-(38)
distalUP 4.8 bits
proximalUP 6.2 bits
... distalUP-(52)-s
... distalUP-(54)-s
... sigma38p10-prox
... sigma38p10-prox
... proximalUP-(33)
distalUP 4.1 bits
... distalUP-(40)-s
... sigma38p10-prox
... proximalUP-(40)
... distalUP-(47)-s
... sigma38p10-prox

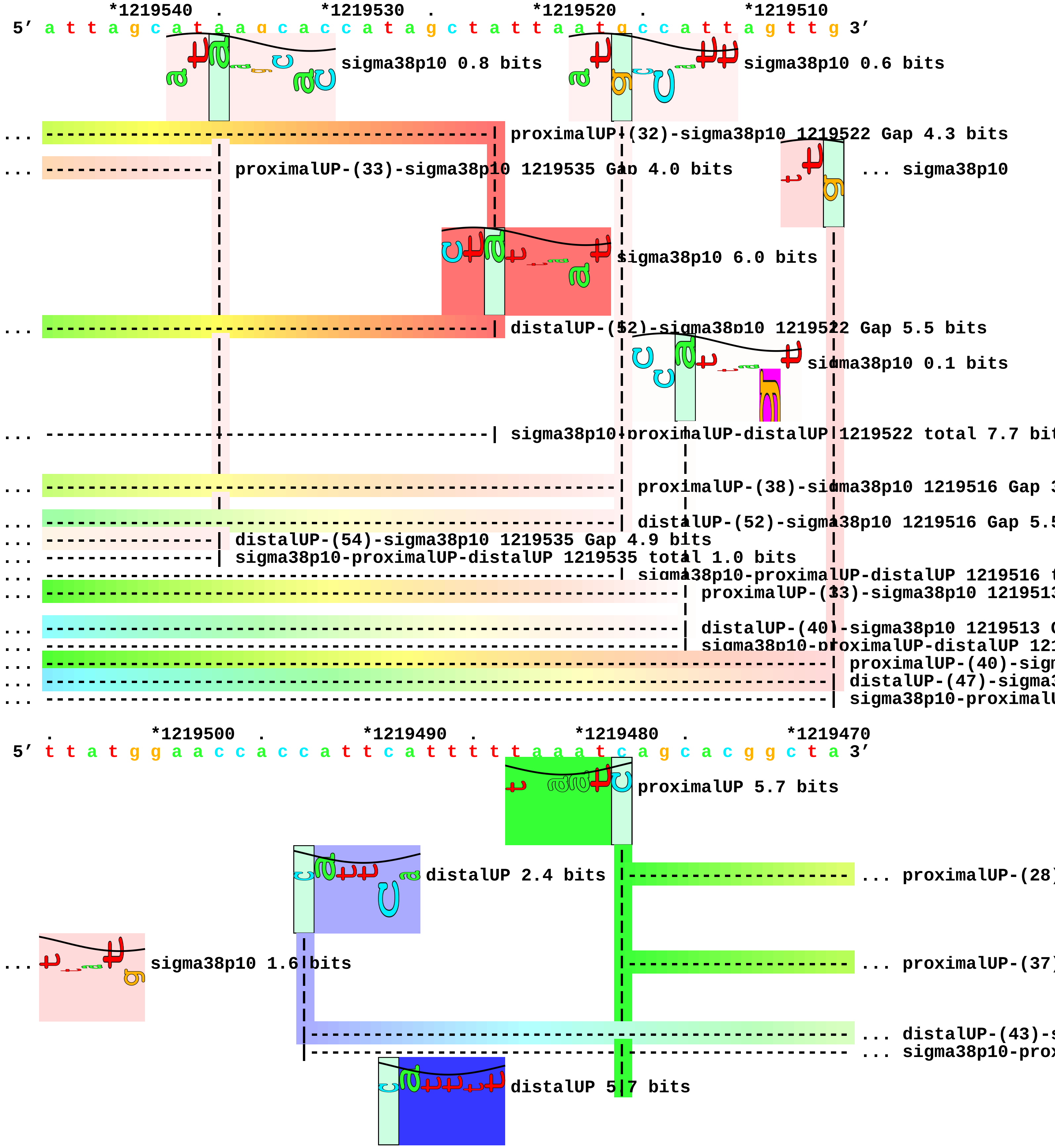
*1219540 . *1219530 . *1219520 . *1219510
5' a t t a g c a t a a g c a c c a t a g c t a t t a a t g c c a t t a g t t g 3'
sigma38p10 0.8 bits
sigma38p10 0.6 bits
... proximalUP-(32)-sigma38p10 1219522 Gap 4.3 bits
... proximalUP-(33)-sigma38p10 1219535 Gap 4.0 bits
... sigma38p10
sigma38p10 6.0 bits
... distalUP-(52)-sigma38p10 1219522 Gap 5.5 bits
sigma38p10 0.1 bits
... sigma38p10-proximalUP-distalUP 1219522 total 7.7 bit
... proximalUP-(38)-sigma38p10 1219516 Gap 3
... distalUP-(52)-sigma38p10 1219516 Gap 5.5
... distalUP-(54)-sigma38p10 1219535 Gap 4.9 bits
... sigma38p10-proximalUP-distalUP 1219535 total 1.0 bits
... sigma38p10-proximalUP-distalUP 1219516 t
... proximalUP-(33)-sigma38p10 121951
... distalUP-(40)-sigma38p10 1219513 G
... sigma38p10-proximalUP-distalUP 121
... proximalUP-(40)-sigm
... distalUP-(47)-sigma3
... sigma38p10-proximalU
. *1219500 . *1219490 . *1219480 . *1219470
5' t t a t g g a a c c a c c a t t c a t t t t t a a a t c a g c a c g g c t a 3'
proximalUP 5.7 bits
distalUP 2.4 bits
... proximalUP-(28)
... sigma38p10 1.6 bits
... proximalUP-(37)
... distalUP-(43)-s
... sigma38p10-prox
distalUP 5.7 bits

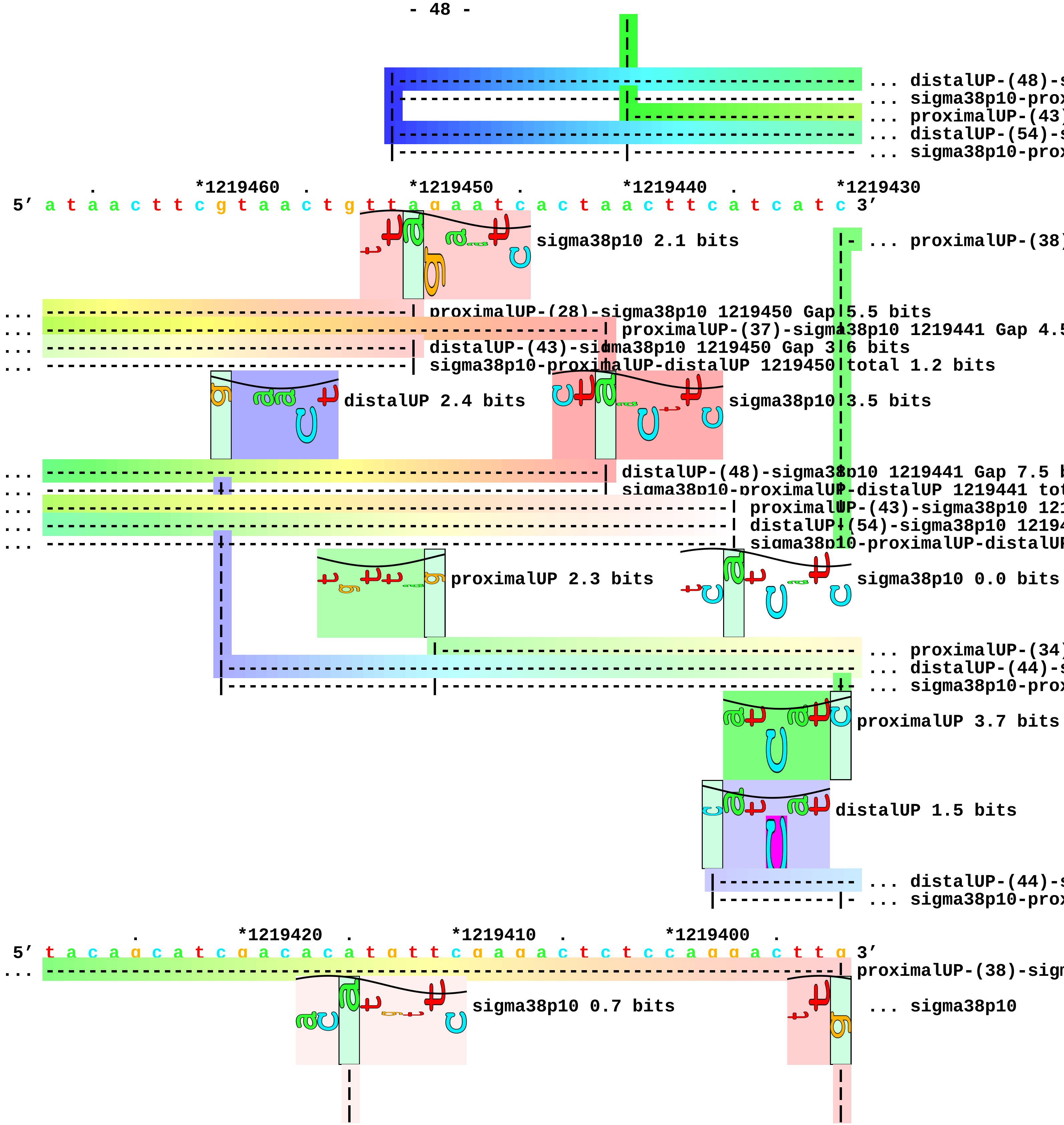

... distalUP-(48)-s
... sigma38p10-prox
... proximalUP-(43)
... distalUP-(54)-s
... sigma38p10-prox
. *1219460 . *1219450 . *1219440 . *1219430
5′ a t a a c t t c g t a a c t g t t a g a a t c a c t a a c t t c a t c a t c 3′
sigma38p10 2.1 bits
... proximalUP-(38)
... proximalUP-(28)-sigma38p10 1219450 Gap 5.5 bits
... proximalUP-(37)-sigma38p10 1219441 Gap 4.5
... distalUP-(43)-sigma38p10 1219450 Gap 3.6 bits
... sigma38p10-proximalUP-distalUP 1219450 total 1.2 bits
distalUP 2.4 bits
sigma38p10 3.5 bits
... distalUP-(48)-sigma38p10 1219441 Gap 7.5 b
... sigma38p10-proximalUP-distalUP 1219441 tot
... proximalUP-(43)-sigma38p10 121
... distalUP-(54)-sigma38p10 12194
... sigma38p10-proximalUP-distalUP
proximalUP 2.3 bits
sigma38p10 0.0 bits
... proximalUP-(34)
... distalUP-(44)-s
... sigma38p10-prox
proximalUP 3.7 bits
distalUP 1.5 bits
... distalUP-(44)-s
... sigma38p10-prox
. *1219420 . *1219410 . *1219400 .
5′ t a c a g c a t c g a c a c a t g t t c g a g a c t c t c c a g g a c t t g 3′
... proximalUP-(38)-sigm
sigma38p10 0.7 bits
... sigma38p10

```
... ----------------------------| proximalUP-(34)-sigma38p10 1219415 Gap 2.5 bits
... ----------------------------| distalUP-(44)-sigma38p10 1219415 Gap 2.8 bits
... ----------------------------| sigma38p10-proximalUP-distalUP 1219415 total 0.0 bits

... ---------------------------------------------------------------------------| distalUP-(44)-sigma3
... ---------------------------------------------------------------------------| sigma38p10-proximalU

     *1219390  .         *1219380  .         *1219370  .         *1219360  .
 5' a a c a t g c a a g a a c a c c t a c t g c a t t t t c c c c a g a g g t a 3'
```

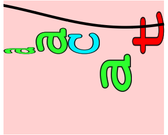

```
... sigma38p10 2.0 bits

         *1219350
 5' a c a a 3'
```

piece 13, NC_000913, ycgB, config: linear, direction: +, begin: 1236370, end: 1236576

```
   *1236370   .         *1236380   .         *1236390   .         *1236400   .
5' a t c t c t g c c a g a t a a a c a t c c a g c a g g t c g a a c g t c c a 3'
   T timestamp:_2018Sep20_11-35-24

     *1236410   .         *1236420   .         *1236430   .         *1236440   .
5' g t c g g g t c c a t c g c t c a a a c g t g t g g t g t c c t t a t t c a 3'

       *1236450   .         *1236460   .         *1236470   .         *1236480
5' t a g a a t c g a t c g t c g c c a t a c g c g c a c c t c a t t g t t g t 3'

    .         *1236490   .         *1236500   .         *1236510   .         *1236520
5' c g g c g c t c t g t g t g g a g c a c c t c a t t t c a a g c a t a g 3'
```

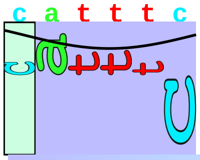

distalUP 1.9 bits

```
                                          |---------------------------- ... distalUP-(56)-s
                                          |---------------------------- ... distalUP-proxin

      .         *1236530   .         *1236540   .         *1236550   .
5' a a c a c c t g t t a a a a a c c g c g t c g c c g g a g a a t t t t t t t 3'
```

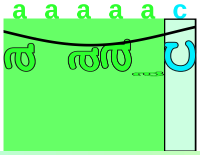

proximalUP 4.3 bits

```
... ------------------------------------------------------------------------ ... distalUP-(56)-s
... ------------------------------|----------------------------------------- ... distalUP-proxin
                                  |----------------------------------------- ... proximalUP-(27)
                                  |----------------------------------------- ... proximalUP-(34)
```

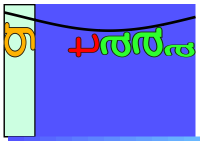

distalUP 4.9 bits

```
             |-------------------------------------------------------------- ... distalUP-(42)-s
             |---------------|---------------------------------------------- ... distalUP-proxin

   *1236560   .         *1236570   .
5' c t t t g c g a t t t c t t a t t 3'
```

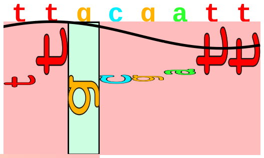

sigma38p10 2.9 bits

```
... --------| distalUP-(56)-sigma38p10 1236564 Gap 3.8 bits
... --------| distalUP-proximalUP-sigma38p10 1236564 total 1.5 bits
... --------| proximalUP-(27)-sigma38p10 1236564 Gap 3.8 bits
... ----------------------| proximalUP-(34)-sigma38p10 1236571 Gap 2.5 bits
```

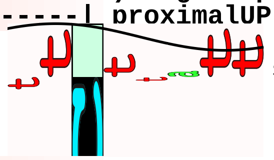

sigma38p10 0.2 bits

```
... ----------------------| distalUP-(42)-sigma38p10 1236571 Gap 3.3 bits
... ----------------------| distalUP-proximalUP-sigma38p10 1236571 total 3.6 bits
```

piece 14, NC_000913, osmB, config: linear, direction: +, begin: 1341254, end: 1341530

```
     .         *1341260  .         *1341270  .         *1341280  .         *1341290
5' g t t g c g g t c c c g t t t a g a c c a g t t a g a a c a g g c a c t c a 3'
   T timestamp:_2018Sep20_11-35-24

       .         *1341300  .         *1341310  .         *1341320  .
5' g a g a c a t t g c c a a a g t a a t t g c c a g a a c a g c c g c g g t c 3'
                                 proximalUP 4.6 bits                  ... distalUP
                                 |-----------------------------------  ... proximalUP-(35)
                               distalUP 5.0 bits                    |  
                                                                    |- ... distalUP-(43)-s
                               |-------------------------------------- ... distalUP-(41)-s
                               |-----------|-------------------------- ... distalUP-proxim
                                                                    |- ... distalUP-proxim

   *1341330  .         *1341340  .         *1341350  .         *1341360  .
5' a t t t t t t t g c t c g t t a c a a a c a t a a t a c t c t c c t g a 3'
... distalUP 5.7 bits                 sigma38p10 2.0 bits               ... distalUP
... ----------------------------------| proximalUP-(35)-sigma38p10 1341347 Gap 3.1 bits
... ------------------------------------------------------------------ ... distalUP-(43)-s
... ----------------------------------| distalUP-(41)-sigma38p10 1341347 Gap 3.1 bits
... ----------------------------------| distalUP-proximalUP-sigma38p10 1341347 total 5.3 bits
... ----------------|--------------------------------------------------- ... distalUP-proxim
      proximalUP 5.7 bits           proximalUP 3.3 bits
              |--------------------------------------------------------- ... proximalUP-(34)
                                        |------------------------------- ... proximalUP-(30)
                             distalUP 2.5 bits                    |
                                                                  |--- ... distalUP-(45)-s
                             |------------------------------------------ ... distalUP-(41)-s
                             |---------------------|-------------------- ... distalUP-proxim
                                     proximalUP 0.6 bits
                                              |------------------------- ... proximalUP-(40)
                                   distalUP 6.3 bits              |--- ... distalUP-proxim
                                   |------------------------------------ ... distalUP-(46)-s
```

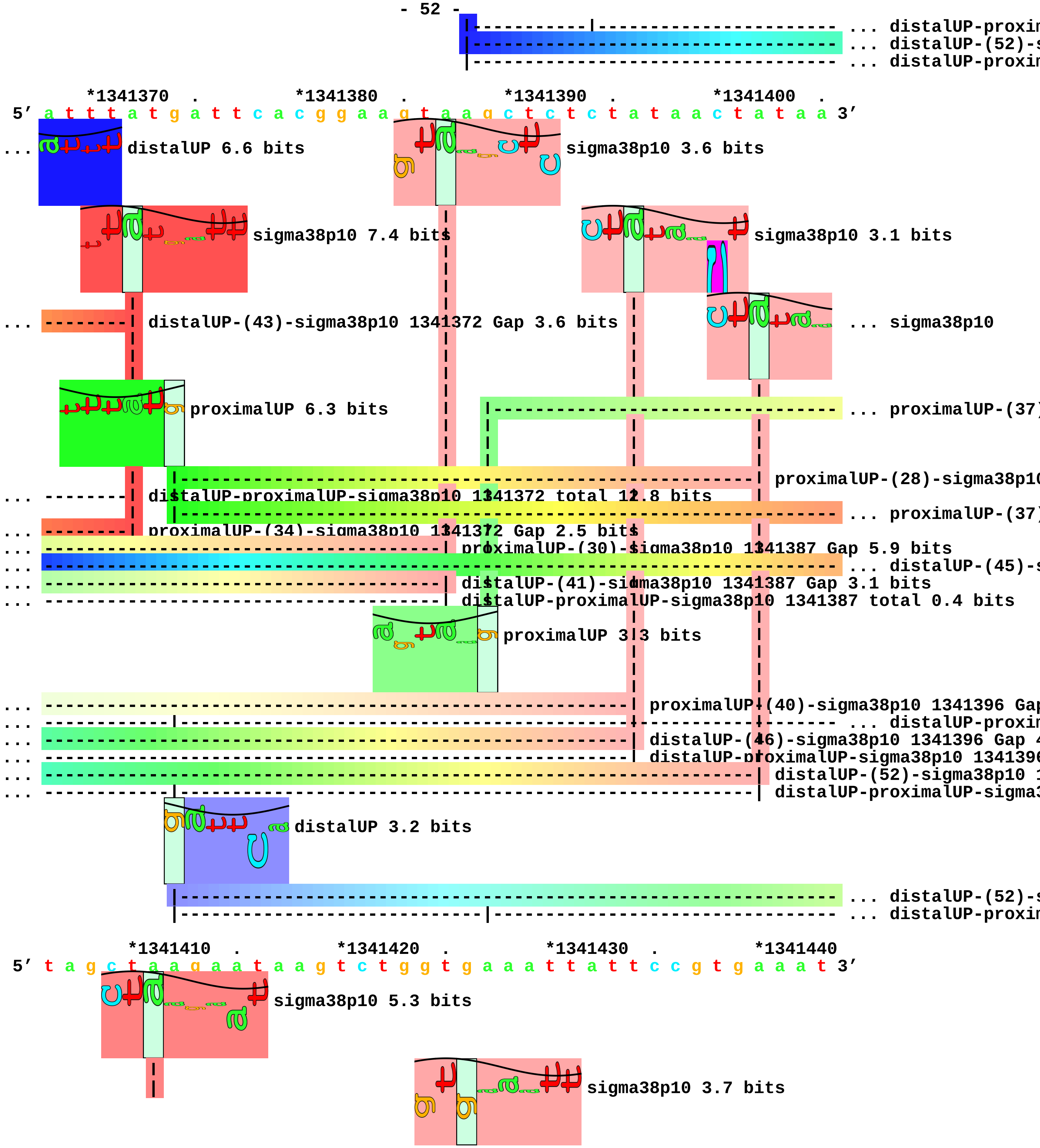
... distalUP-proxim
... distalUP-(52)-s
... distalUP-proxim
*1341370 . *1341380 . *1341390 . *1341400 .
5′ a t t t a t g a t t c a c g g a a g t a a g c t c t c t a t a a c t a t a a 3′
... distalUP 6.6 bits
sigma38p10 3.6 bits
sigma38p10 7.4 bits
sigma38p10 3.1 bits
... distalUP-(43)-sigma38p10 1341372 Gap 3.6 bits
... sigma38p10
proximalUP 6.3 bits
... proximalUP-(37)
proximalUP-(28)-sigma38p10
... distalUP-proximalUP-sigma38p10 1341372 total 12.8 bits
... proximalUP-(37)
... proximalUP-(34)-sigma38p10 1341372 Gap 2.5 bits
... proximalUP-(30)-sigma38p10 1341387 Gap 5.9 bits
... distalUP-(45)-s
... distalUP-(41)-sigma38p10 1341387 Gap 3.1 bits
... distalUP-proximalUP-sigma38p10 1341387 total 0.4 bits
proximalUP 3.3 bits
... proximalUP-(40)-sigma38p10 1341396 Gap
... distalUP-proxim
... distalUP-(46)-sigma38p10 1341396 Gap 4
... distalUP-proximalUP-sigma38p10 1341396
... distalUP-(52)-sigma38p10 1
... distalUP-proximalUP-sigma3
distalUP 3.2 bits
... distalUP-(52)-s
... distalUP-proxim
*1341410 . *1341420 . *1341430 . *1341440
5′ t a g c t a a g a a t a a g t c t g g t g a a a t t a t t c c g t g a a a t 3′
sigma38p10 5.3 bits
sigma38p10 3.7 bits

```
... sigma38p10 3.3 bits
... -----------------------------------------| proximalUP-(37)-sigma38p10 1341426 Gap 4.5 bits
... ----------| proximalUP-(37)-sigma38p10 1341411 Gap 4.5 bits
... ----------| distalUP-(45)-sigma38p10 1341411 Gap 5.5 bits
... ----------| distalUP-proximalUP-sigma38p10 1341411 total 8.2 bits
... -----------------------------------------| distalUP-(52)-sigma38p10 1341426 Gap 5.5 bits
... -----------------------------------------| distalUP-proximalUP-sigma38p10 1341426 total 0.3 bits

       .         *1341450  .         *1341460  .         *1341470  .         *1341480
 5' c t g c t c g c g c g c a g g g t t a t a t c a c g c a g g t g a t a a t g 3'
                                         proximalUP 5.7 bits              |- ... proximalUP-(31)
                                               |------------------------------- ... proximalUP-(34)
                                      distalUP 4.3 bits            proximalUP 6.7 bits
                                    |----------------------------------------- ... distalUP-(41)-s
                                    |-------------|-------------------------|- ... distalUP-proxim
                                                  |---------------------------- ... proximalUP-(37)
                                    |----------------------------------------- ... distalUP-(44)-s
                                    |-------------|-------------------------|- ... distalUP-proxim
                                                  |---------------------------- ... proximalUP-(38)
                                    |----------------------------------------- ... distalUP-(45)-s
                                    |-------------|-------------------------|- ... distalUP-proxim
                                                  |---------------------------- ... proximalUP-(40)
                                    |----------------------------------------- ... distalUP-(47)-s
                                    |-------------|-------------------------|- ... distalUP-proxim
                                    |----------------------------------------- ... distalUP-(53)-s
                                    |---------------------------------------|- ... distalUP-proxim
                                                                            |- ... proximalUP-(33)
                                    |----------------------------------------- ... distalUP-(55)-s
                                    |---------------------------------------|- ... distalUP-proxim

           .         *1341490  .         *1341500  .         *1341510  .
 5' a c t t c c t g t t a t a t c g c t g a t a a t a a t t t t a t a t c t t g 3'
... ------------------------------------------------------------| proximalUP-(31)-sigma38p10 1341512
... ------------------------------------| proximalUP-(34)-sigma38p10 1341500 Gap 2.5 bits
```

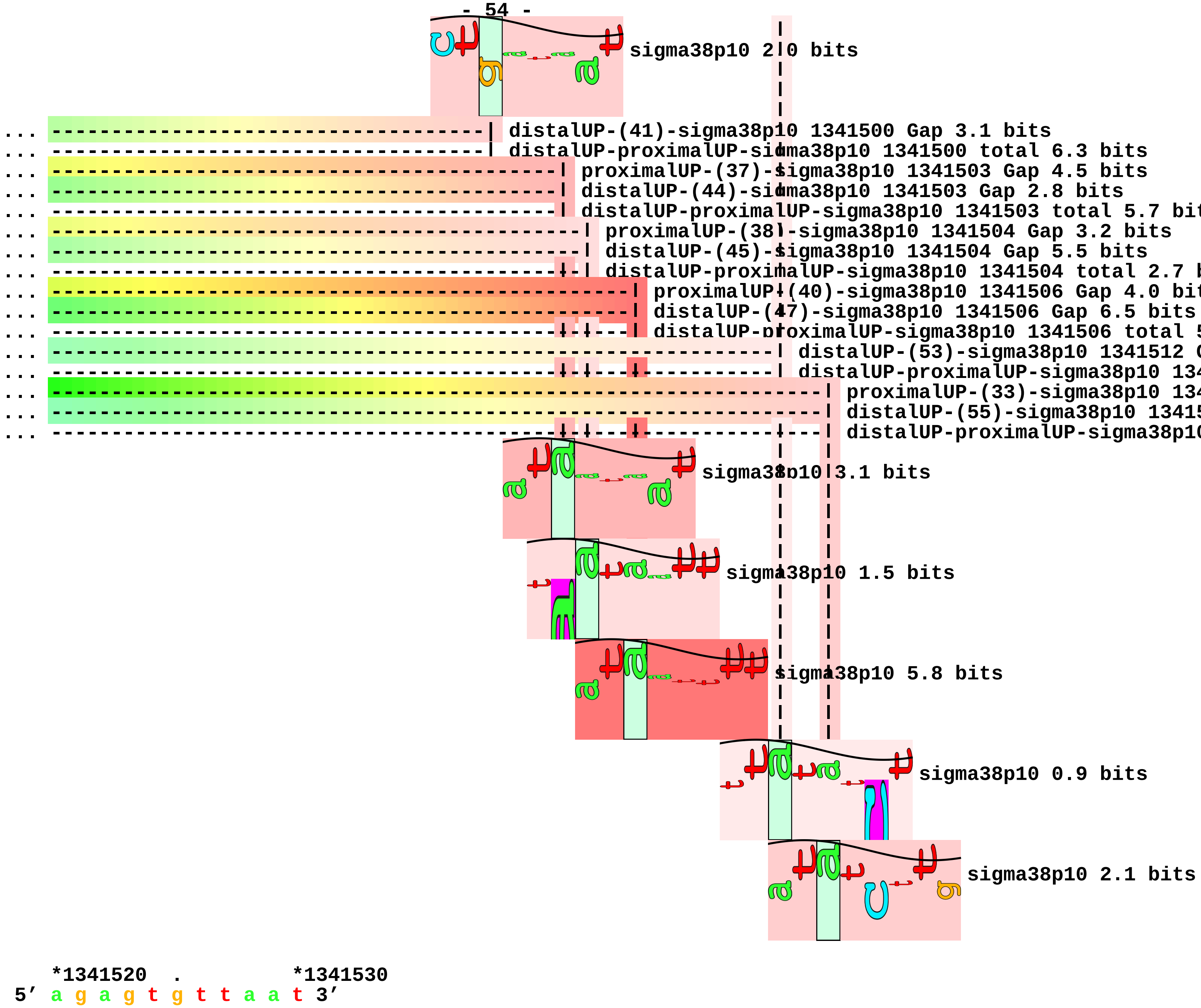
sigma38p10 2.0 bits
distalUP-(41)-sigma38p10 1341500 Gap 3.1 bits
distalUP-proximalUP-sigma38p10 1341500 total 6.3 bits
proximalUP-(37)-sigma38p10 1341503 Gap 4.5 bits
distalUP-(44)-sigma38p10 1341503 Gap 2.8 bits
distalUP-proximalUP-sigma38p10 1341503 total 5.7 bit
proximalUP-(38)-sigma38p10 1341504 Gap 3.2 bits
distalUP-(45)-sigma38p10 1341504 Gap 5.5 bits
distalUP-proximalUP-sigma38p10 1341504 total 2.7 b
proximalUP-(40)-sigma38p10 1341506 Gap 4.0 bit
distalUP-(47)-sigma38p10 1341506 Gap 6.5 bits
distalUP-proximalUP-sigma38p10 1341506 total 5
distalUP-(53)-sigma38p10 1341512 G
distalUP-proximalUP-sigma38p10 134
proximalUP-(33)-sigma38p10 134
distalUP-(55)-sigma38p10 13415
distalUP-proximalUP-sigma38p10
sigma38p10 3.1 bits
sigma38p10 1.5 bits
sigma38p10 5.8 bits
sigma38p10 0.9 bits
sigma38p10 2.1 bits
*1341520 . *1341530
5' a g a g t g t t a a t 3'

piece 15, NC_000913, tfaR, config: linear, direction: -, begin: 1430599, end: 1430200

. *1430590 . *1430580 . *1430570 .

5' c a c g a a g c c a g c c g g a a t a t c t g g c g g t g c a a t a t c g g 3'

T timestamp:_2018Sep20_11-35-24

proximalUP 6.1 bits

|----- ... proximalUP-(39)

distalUP 6.4 bits

|---------------- ... distalUP-(45)-s

|-----------|----- ... sigma38p10-prox

*1430560 . *1430550 . *1430540 . *1430530 .

5' t a c t g t t t g c t g g c a g a c c t g t a t g a g g c g g a a t a t a t 3'

... sigma38p10

... ------------------------------------------------------------------------| proximalUP-(39)-sigma3

... proximalUP

... ------------------------------------------------------------------------| distalUP-(45)-sigma38p

... ------------------------------------------------------------------------| sigma38p10-proximalUP-

distalUP 7.2 bits

|--------------- ... distalUP-(52)-s

|--------------- ... sigma38p10-prox

|--------------- ... distalUP-(54)-s

|--------------- ... sigma38p10-prox

*1430520 . *1430510 . *1430500 . *1430490

5' g c a t c a c c t t c a c c a a t a a a t t c a t t a g t t c c g g c c a g 3'

... sigma38p10 4.2 bits

proximalUP 5.5 bits

|--------------------------------------------------------------------- ... proximalUP-(40)

... proximalUP 1.6 bits

|---------------------------------- ... proximalUP-(32)

|--------------------------------------------------------------------- ... proximalUP-(42)

distalUP 5.7 bits

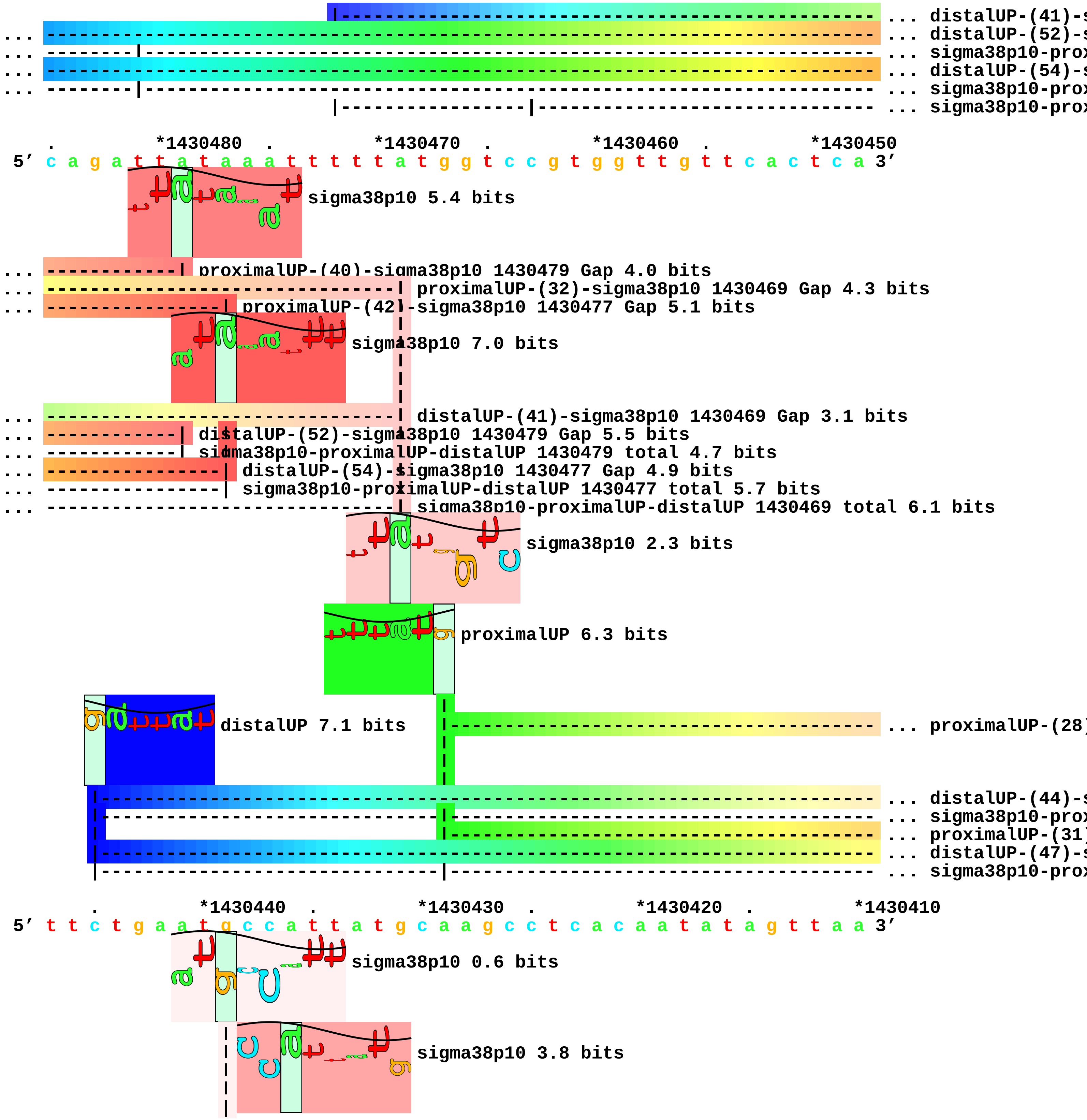
... distalUP-(41)-s
... distalUP-(52)-s
... sigma38p10-prox
... distalUP-(54)-s
... sigma38p10-prox
... sigma38p10-prox
*1430480
*1430470
*1430460
*1430450
5′ c a g a t t t a t a a a t t t t t a t g g t c c g t g g t t g t t c a c t c a 3′
sigma38p10 5.4 bits
proximalUP-(40)-sigma38p10 1430479 Gap 4.0 bits
proximalUP-(32)-sigma38p10 1430469 Gap 4.3 bits
proximalUP-(42)-sigma38p10 1430477 Gap 5.1 bits
sigma38p10 7.0 bits
distalUP-(41)-sigma38p10 1430469 Gap 3.1 bits
distalUP-(52)-sigma38p10 1430479 Gap 5.5 bits
sigma38p10-proximalUP-distalUP 1430479 total 4.7 bits
distalUP-(54)-sigma38p10 1430477 Gap 4.9 bits
sigma38p10-proximalUP-distalUP 1430477 total 5.7 bits
sigma38p10-proximalUP-distalUP 1430469 total 6.1 bits
sigma38p10 2.3 bits
proximalUP 6.3 bits
distalUP 7.1 bits
... proximalUP-(28)
... distalUP-(44)-s
... sigma38p10-prox
... proximalUP-(31)
... distalUP-(47)-s
... sigma38p10-prox
*1430440
*1430430
*1430420
*1430410
5′ t t c t g a a t g c c a t t t a t g c a a g c c t c a c a a t a t a g t t a a 3′
sigma38p10 0.6 bits
sigma38p10 3.8 bits

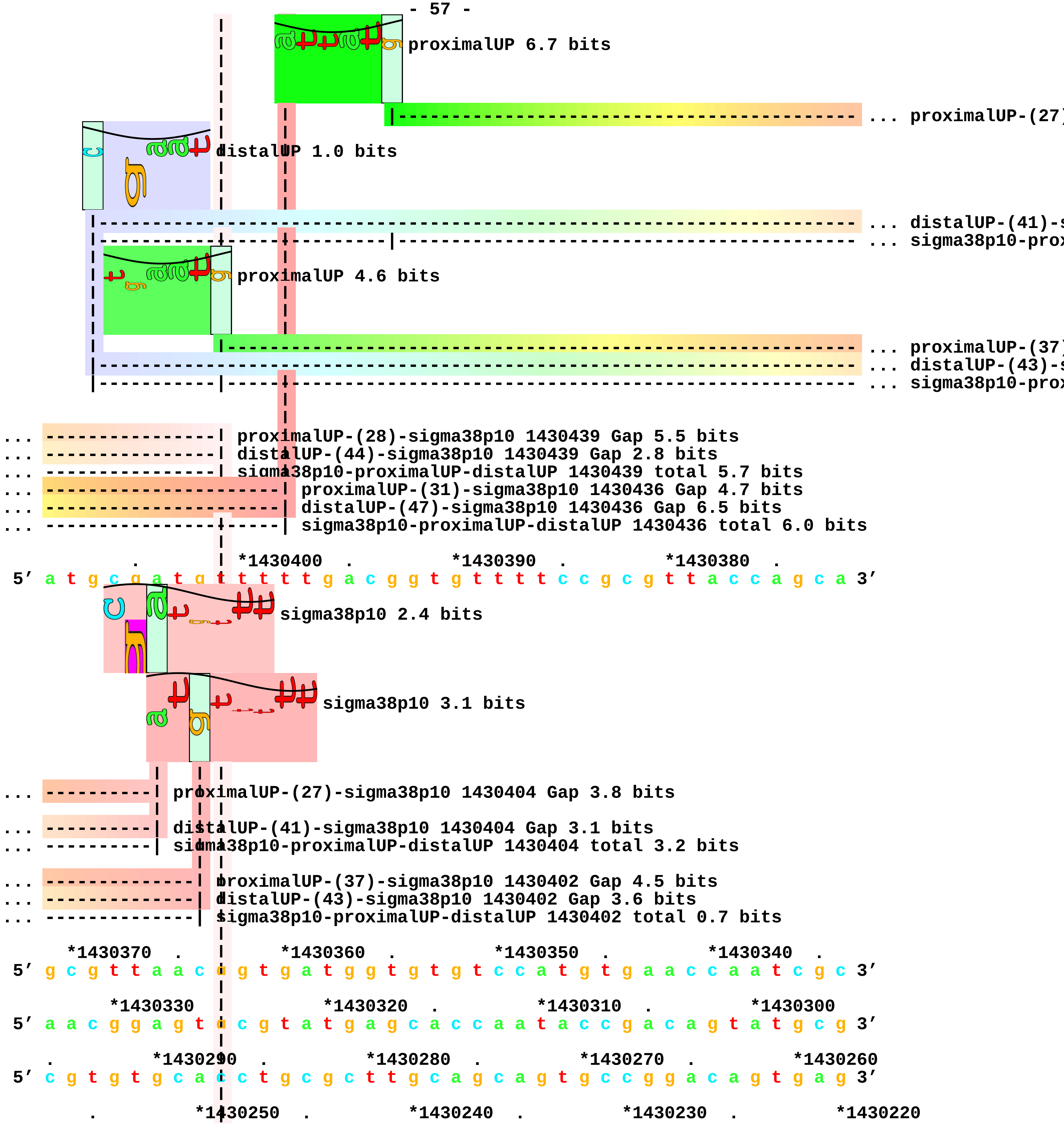
proximalUP 6.7 bits
... proximalUP-(27)
distalUP 1.0 bits
... distalUP-(41)-s
... sigma38p10-prox
proximalUP 4.6 bits
... proximalUP-(37)
... distalUP-(43)-s
... sigma38p10-prox
... proximalUP-(28)-sigma38p10 1430439 Gap 5.5 bits
... distalUP-(44)-sigma38p10 1430439 Gap 2.8 bits
... sigma38p10-proximalUP-distalUP 1430439 total 5.7 bits
... proximalUP-(31)-sigma38p10 1430436 Gap 4.7 bits
... distalUP-(47)-sigma38p10 1430436 Gap 6.5 bits
... sigma38p10-proximalUP-distalUP 1430436 total 6.0 bits
*1430400 . *1430390 . *1430380 .
5′ a t g c g a t g t t t t t g a c g g t g t t t t c c g c g t t a c c a g c a 3′
sigma38p10 2.4 bits
sigma38p10 3.1 bits
... proximalUP-(27)-sigma38p10 1430404 Gap 3.8 bits
... distalUP-(41)-sigma38p10 1430404 Gap 3.1 bits
... sigma38p10-proximalUP-distalUP 1430404 total 3.2 bits
... proximalUP-(37)-sigma38p10 1430402 Gap 4.5 bits
... distalUP-(43)-sigma38p10 1430402 Gap 3.6 bits
... sigma38p10-proximalUP-distalUP 1430402 total 0.7 bits
*1430370 . *1430360 . *1430350 . *1430340 .
5′ g c g t t a a c c g t g a t g g t g t g t c c a t g t g a a c c a a t c g c 3′
*1430330 *1430320 . *1430310 . *1430300
5′ a a c g g a g t c c g t a t g a g c a c c a a t a c c g a c a g t a t g c g 3′
. *1430290 . *1430280 . *1430270 . *1430260
5′ c g t g t g c a c c t g c g c t t g c a g c a g t g c c g g a c a g t g a g 3′
. *1430250 . *1430240 . *1430230 . *1430220

```
5’ t g g g t a t g t g c g c c a g c a g a t g a t g t t g c a t a g t t t t g 3’
                   |
       .           | *1430210  .         *1430200
5’ a t t a t g c a c a a c a g a c a a t c 3’
                   |
                   |
```

piece 16, NC_000913, ydcS, config: linear, direction: -, begin: 1509747, end: 1509476

```
       .         *1509740  .         *1509730  .         *1509720  .         *1509710
5' c g g c g g c g t g a g c g g t c a t t a t t g t c a t g c t g a g c g c a 3'
 T timestamp:_2018Sep20_11-35-24

           .         *1509700  .         *1509690  .         *1509680  .
5' c a c a g g c t g c t g c g g g c a a a t g t c t t g c t c a t a a g g t c 3'

     *1509670  .         *1509660  .         *1509650  .         *1509640  .
5' t t a c t c c t g t c g t a a t t a a t t g t t c a g c c g c c g t g c c g 3'
```

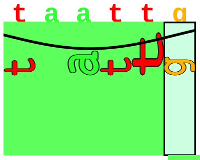

proximalUP 4.6 bits

|----------------------------------- ... proximalUP-(36)

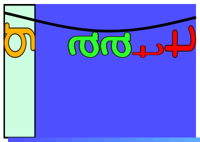

distalUP 5.0 bits

|------------------------------------------------------------ ... distalUP-(46)-s
|-------------------|------------------------------------ ... sigma38p10-prox

```
         *1509630  .         *1509620  .         *1509610  .         *1509600
5' t a a a a a t a t c t c c t t g t t t a a c c a t a g a c g g c g g c t t g 3'
```

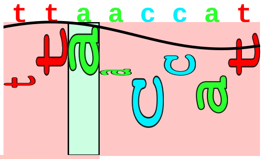

sigma38p10 2.5 bits

... --------------------------------------| proximalUP-(36)-sigma38p10 1509614 Gap 3.6 bits
... --------------------------------------| distalUP-(46)-sigma38p10 1509614 Gap 4.1 bits
... --------------------------------------| sigma38p10-proximalUP-distalUP 1509614 total 4.3 bits

```
   .         *1509590  .         *1509580  .         *1509570  .         *1509560
5' c a g g c a c a c c t t c g c c c c t g a g a a a a t t t a t t c a g g t t 3'
```

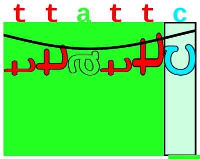

proximalUP 6.2 bits

|----------- ... proximalUP-(34)

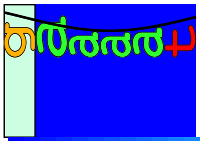

distalUP 7.2 bits

|-------------------------------------- ... distalUP-(45)-s
|---------------------|----------- ... sigma38p10-prox
|----------- ... proximalUP-(40)
|-------------------------------------- ... distalUP-(51)-s
|---------------------|----------- ... sigma38p10-prox

```
       .         *1509550  .         *1509540  .         *1509530  .         *1509520
5' g t g a g c t a c c g c g c a t t a t g c c g t c t g t a a g g t t a t c t 3'
```

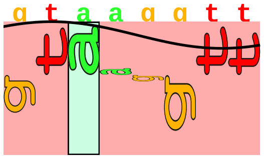

sigma38p10 3.6 bits

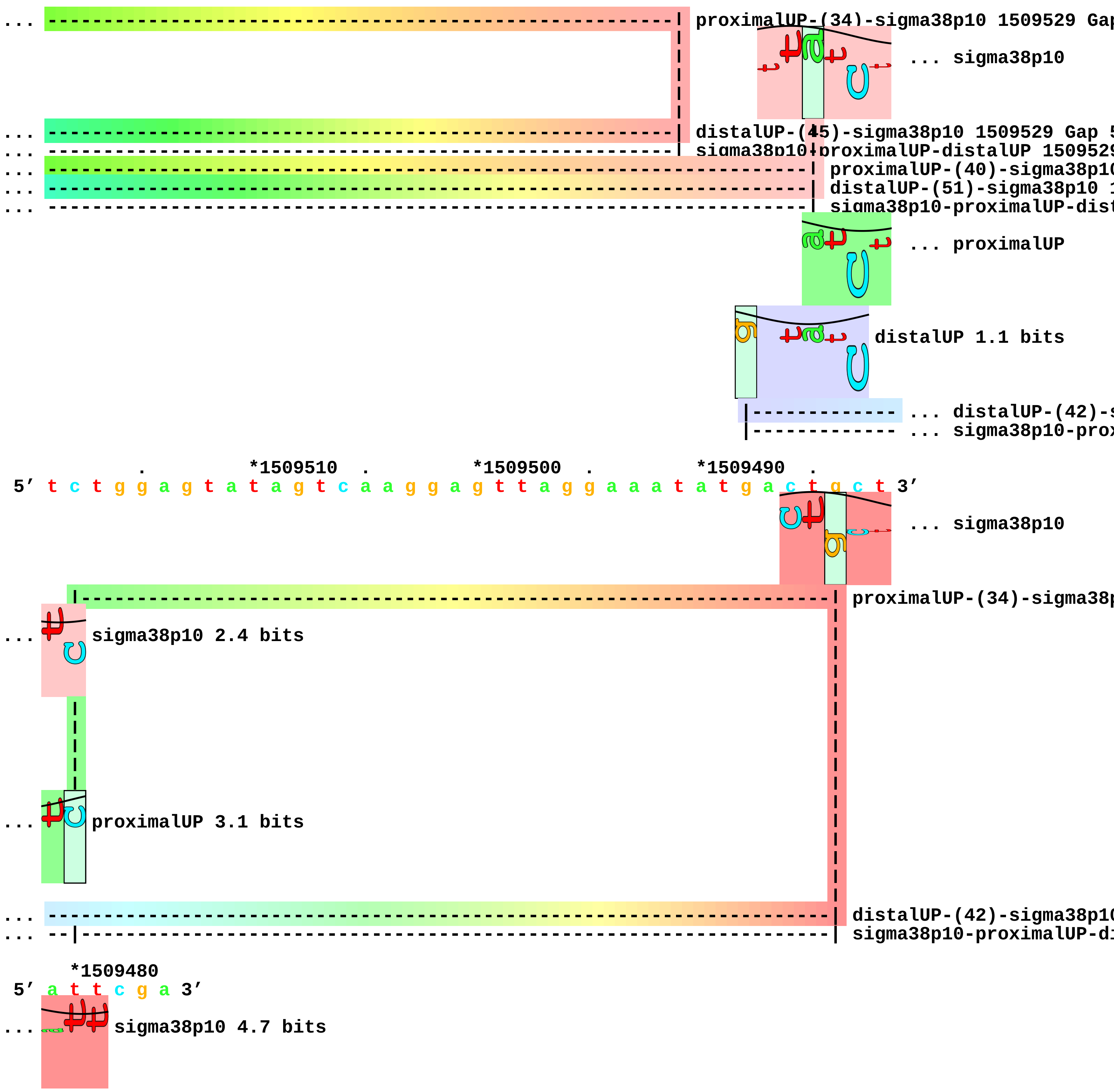

... proximalUP-(34)-sigma38p10 1509529 Gap
... sigma38p10
... distalUP-(45)-sigma38p10 1509529 Gap 5
... sigma38p10-proximalUP-distalUP 1509529
... proximalUP-(40)-sigma38p10
... distalUP-(51)-sigma38p10 1
... sigma38p10-proximalUP-dist
... proximalUP
distalUP 1.1 bits
... distalUP-(42)-s
... sigma38p10-prox
*1509510 *1509500 *1509490
5' t c t g g a g t a t a g t c a a g g a g t t a g g a a a t a t g a c t g c t 3'
... sigma38p10
proximalUP-(34)-sigma38p
... sigma38p10 2.4 bits
... proximalUP 3.1 bits
... distalUP-(42)-sigma38p10
... sigma38p10-proximalUP-di
*1509480
5' a t t c g a 3'
... sigma38p10 4.7 bits

piece 17, NC_000913, ansP, config: linear, direction: +, begin: 1523950, end: 1524249

```
   *1523950  .         *1523960  .         *1523970  .         *1523980  .
5' g c c a g c g g c g t t t c g c g g c g t g t t g a t c t g a a g t g t c g 3'
   T timestamp:_2018Sep20_11-35-24

     *1523990  .         *1524000  .         *1524010  .         *1524020  .
5' g t g t c g t g t t t a c t c a t t g c t c t c c c t g a t t g c t t t a a 3'
                                                                     ... proximalUP
                                                   distalUP 0.4 bits
                                                   |------------------------- ... distalUP-(41)-s
                                                   |------------------------- ... distalUP-proxim
                                                                  distalUP 4.0 bits
                                                                  |---------- ... distalUP-(41)-s
                                                                  |---------- ... distalUP-proxim
                                                                  |---------- ... distalUP-(45)-s
                                                                  |---------- ... distalUP-proxim

       *1524030  .         *1524040  .         *1524050  .         *1524060
5' t g a a a a a g t c a t a t a a g t t g c c a t g a a c a a t g t t t a t t 3'
... proximalUP 6.3 bits                                 sigma38p10 1.2 bits
  |-------------------------------------------------------| proximalUP-(27)-sigma38p10 1524054 Gap
... -------------------------------------------------------| distalUP-(41)-sigma38p10 1524054 Gap 3
... -------------------------------------------------------| distalUP-proximalUP-sigma38p10 1524054
                      proximalUP 3.4 bits                       ... sigma38p10
... ---------------------------------------------------------------------| distalUP-(41)-sigma38p10
... ---------------------------------------------------------------------| distalUP-proximalUP-sign
... ---------------------------------------------------------------------- ... distalUP-(45)-s
... ---------------------------------------------------------------------- ... distalUP-proxim
  |----------------------------------------------------------------------| proximalUP-(34)-sigma38p
  distalUP 6.5 bits                                               ... sigma38p10
  |----------------------------------------------------------------------- ... proximalUP-(38)
                                |----------------------------------------- ... proximalUP-(35)
  |----------------------------------------------------------------------- ... distalUP-(50)-s
  |-----------------------------|----------------------------------------- ... distalUP-proxin
                      distalUP 5.0 bits               proximalUP 0.1 bits
```

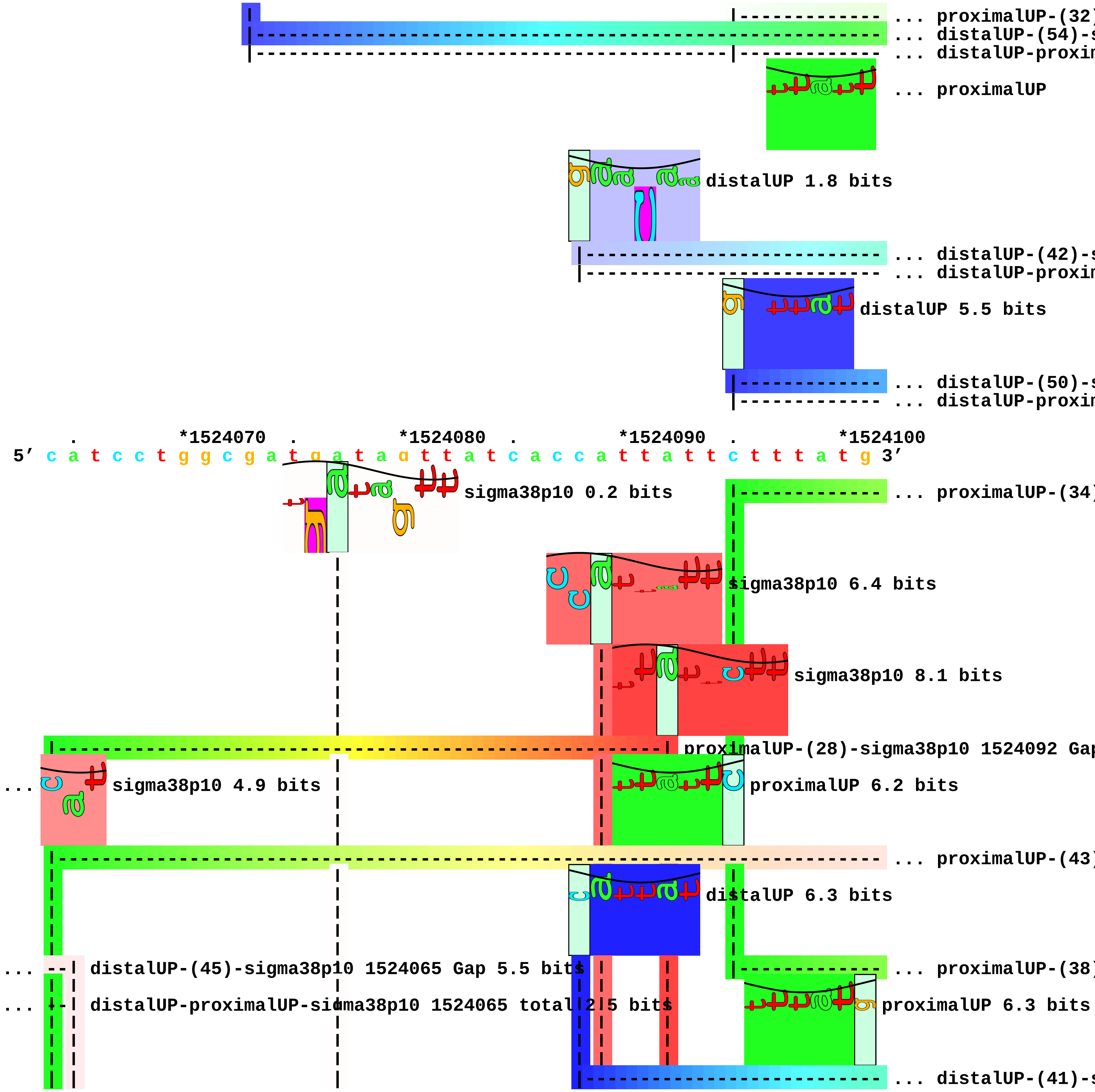
... proximalUP-(32)
... distalUP-(54)-s
... distalUP-proxim
... proximalUP
distalUP 1.8 bits
... distalUP-(42)-s
... distalUP-proxim
distalUP 5.5 bits
... distalUP-(50)-s
... distalUP-proxim
*1524070
*1524080
*1524090
*1524100
5′ c a t c c t g g c g a t g a t a g t t a t c a c c a t t a t t c t t t a t g 3′
sigma38p10 0.2 bits
... proximalUP-(34)
sigma38p10 6.4 bits
sigma38p10 8.1 bits
proximalUP-(28)-sigma38p10 1524092 Gap
... sigma38p10 4.9 bits
proximalUP 6.2 bits
... proximalUP-(43)
distalUP 6.3 bits
... distalUP-(45)-sigma38p10 1524065 Gap 5.5 bits
... proximalUP-(38)
... distalUP-proximalUP-sigma38p10 1524065 total 2 5 bits
proximalUP 6.3 bits
... distalUP-(41)-s

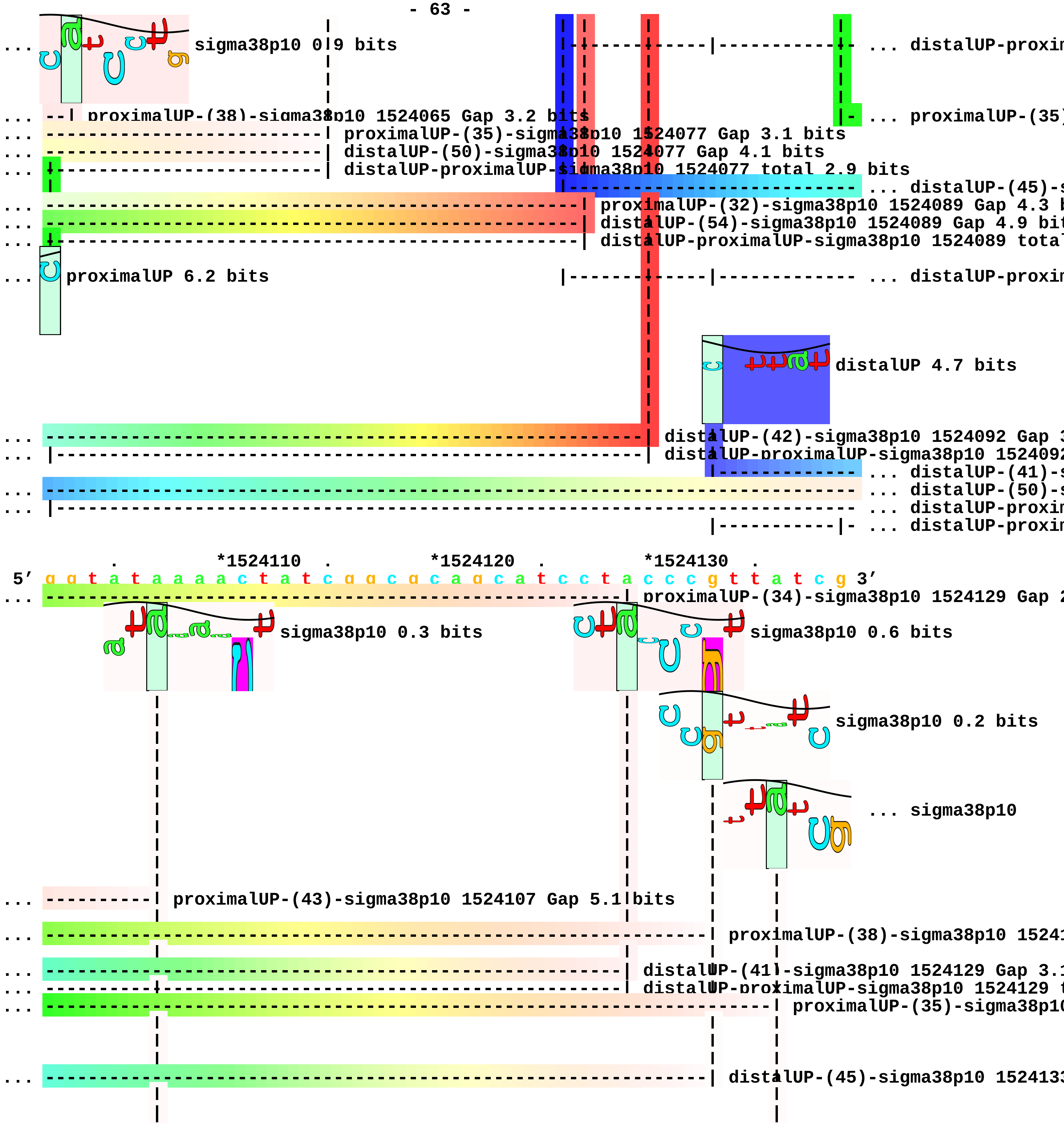
sigma38p10 0.9 bits
... distalUP-proxin
proximalUP-(38)-sigma38p10 1524065 Gap 3.2 bits
... proximalUP-(35)
proximalUP-(35)-sigma38p10 1524077 Gap 3.1 bits
distalUP-(50)-sigma38p10 1524077 Gap 4.1 bits
distalUP-proximalUP-sigma38p10 1524077 total 2.9 bits
... distalUP-(45)-s
proximalUP-(32)-sigma38p10 1524089 Gap 4.3 b
distalUP-(54)-sigma38p10 1524089 Gap 4.9 bit
distalUP-proximalUP-sigma38p10 1524089 tota
proximalUP 6.2 bits
... distalUP-proxin
distalUP 4.7 bits
distalUP-(42)-sigma38p10 1524092 Gap 3
distalUP-proximalUP-sigma38p10 1524092
... distalUP-(41)-s
... distalUP-(50)-s
... distalUP-proxin
... distalUP-proxin
*1524110 *1524120 *1524130
5′ g g t a t a a a a c t a t c g g c g c a g c a t c c t a c c c g t t a t c g 3′
proximalUP-(34)-sigma38p10 1524129 Gap 2
sigma38p10 0.3 bits
sigma38p10 0.6 bits
sigma38p10 0.2 bits
... sigma38p10
proximalUP-(43)-sigma38p10 1524107 Gap 5.1 bits
proximalUP-(38)-sigma38p10 15241
distalUP-(41)-sigma38p10 1524129 Gap 3.1
distalUP-proximalUP-sigma38p10 1524129 t
proximalUP-(35)-sigma38p10
distalUP-(45)-sigma38p10 152413

```
... ---------------------------------------------------------------| distalUP-proximalUP-sigma38p10 1
... ----------------------------------------------------------------------------| distalUP-(41)-sigma38p10 1
... ----------| distalUP-(50)-sigma38p10 1524107 Gap 4.1 bits
... ----------| distalUP-proximalUP-sigma38p10 1524107 total 2.7 bits
... ----------------------------------------------------------------------------| distalUP-proximalUP-sigma3

    *1524140  .         *1524150  .         *1524160  .         *1524170  .
 5' a t a a a c g a t g c a a a a c a t c c c c t t a c a a t c c t g a a g g g 3'

... sigma38p10 0.2 bits

        *1524180  .         *1524190  .         *1524200  .         *1524210  .
 5' g a t t a a t a c a a c t g a c g a a a a a a t g a c a a a t c c t t t t g 3'
      proximalUP 3.3 bits      proximalUP 5.6 bits
                 |--------------------------------------------------------------- ... proximalUP-(36)
    distalUP 6.4 bits                                  |-------------------------- ... proximalUP-(27)
 |--------------------------------------------------------------------------------- ... distalUP-(44)-s
 |---------------|-----------------------------------|----------------------------- ... distalUP-proxim
 |--------------------------------------------------------------------------------- ... distalUP-(51)-s
 |---------------------------------------------------|----------------------------- ... distalUP-proxim
                                                      |--------------------------- ... proximalUP-(37)
                                    distalUP 6.5 bits
                                   |---------------------------------------------- ... distalUP-(45)-s
                                   |---------------|------------------------------ ... distalUP-proxim

           *1524220  .         *1524230  .         *1524240  .
 5' c t g g t t a a c c t g t g t a c t g t c c t a c a c t t a a t c t 3'
                sigma38p10 3.1 bits
... ------------| proximalUP-(36)-sigma38p10 1524222 Gap 3.6 bits
... ---------------------------| proximalUP-(27)-sigma38p10 1524229 Gap 3.8 bits
... ------------| distalUP-(44)-sigma38p10 1524222 Gap 2.8 bits
... ------------| distalUP-proximalUP-sigma38p10 1524222 total 6.5 bits
... ---------------------------| distalUP-(51)-sigma38p10 1524229 Gap 5.1 bits
... ---------------------------| distalUP-proximalUP-sigma38p10 1524229 total 5.5 bits
... --------------------------------------------| proximalUP-(37)-sigma38p10 1524239 Gap 4.5 bits
```

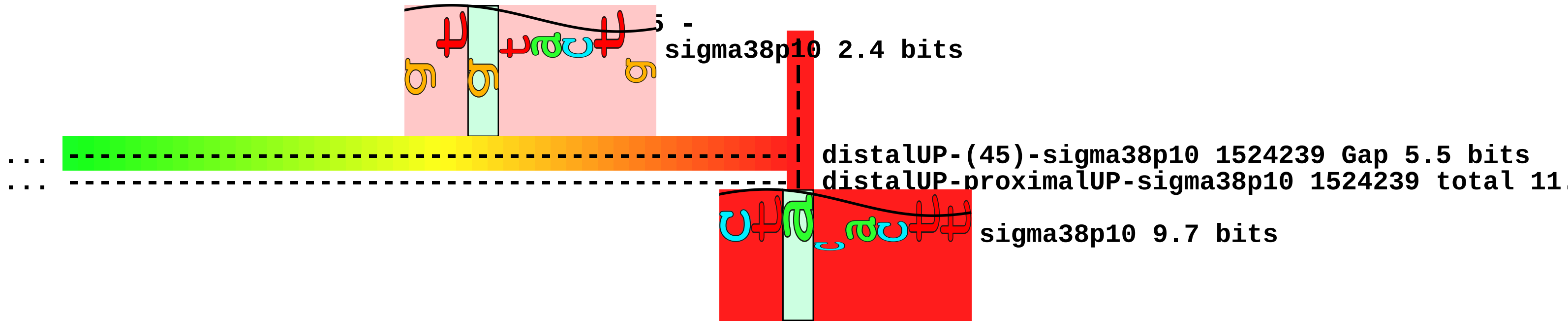

sigma38p10 2.4 bits
distalUP-(45)-sigma38p10 1524239 Gap 5.5 bits
distalUP-proximalUP-sigma38p10 1524239 total 11.
sigma38p10 9.7 bits

piece 18, NC_000913, uxaB, config: linear, direction: +, begin: 1608650, end: 1608999

```
   *1608650   .         *1608660   .         *1608670   .         *1608680   .
5' t g a t a c g t t c t g g a t a c t g t g c a c c g g g a a a a t c g c g a 3'
```

T timestamp:_2018Sep20_11-35-24

proximalUP 6.1 bits

... proximalUP-(35)

distalUP 7.2 bits

... distalUP-(41)-s
... distalUP-proxim
... proximalUP-(41)
... distalUP-(47)-s
... distalUP-proxim

```
     *1608690   .         *1608700   .         *1608710   .         *1608720   .
5' c g a t t t a g t g t t t t c a c a a t g g g t t c c c t t c t g a t t a g 3'
```

sigma38p10 0.2 bits

... proximalUP-(35)-sigma38p10 160871

... sigma38p10

... distalUP-(41)-sigma38p10 1608718 0
... distalUP-proximalUP-sigma38p10 160
... proximalUP-(41)-sigma3
... distalUP-(47)-sigma38p
... distalUP-proximalUP-si

```
       *1608730   .         *1608740   .         *1608750   .         *1608760
5' t c a t a c a a c c t g t t t g a a t t g g t a c g a c a g g t t a g c a a 3'
```

... sigma38p10 1.2 bits

```
   .         *1608770   .         *1608780   .         *1608790   .         *1608800
5' a c t t t a a t a c g c c g a a c c c c t g t t t t g a t c a a c t c c t g 3'
```

```
     .         *1608810   .         *1608820   .         *1608830   .
5' a t g a t t a a t g a g c a g t t t t a t g a g a a a a g t g t g g c g c g 3'
```

proximalUP 6.3 bits

... proximalUP-(33)

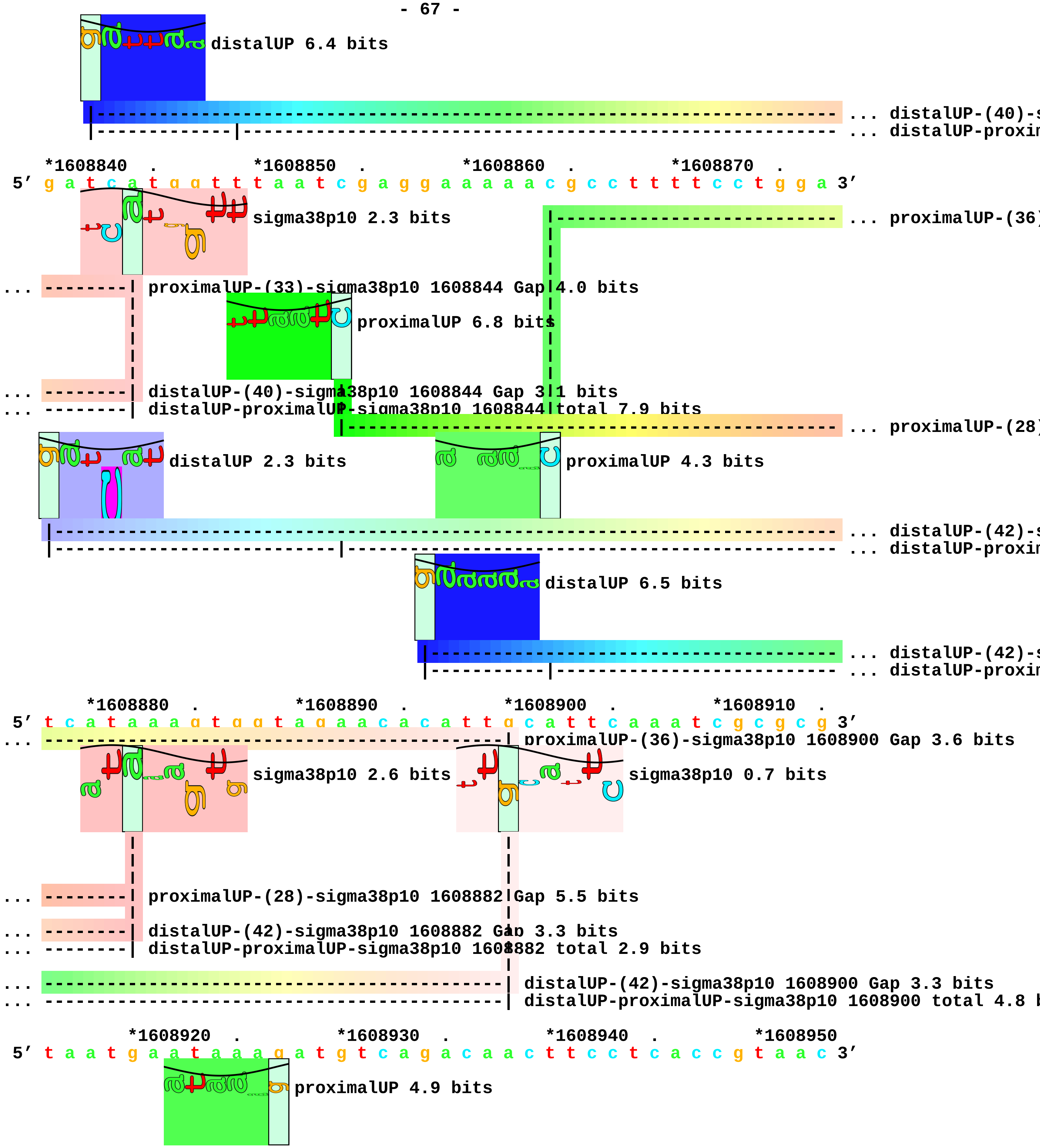
distalUP 6.4 bits
... distalUP-(40)-s
... distalUP-proxin
*1608840 . *1608850 . *1608860 . *1608870 .
5′ g a t c a t g g t t t t a a t c g a g g a a a a a c g c c t t t t c c t g g a 3′
sigma38p10 2.3 bits
... proximalUP-(36)
... proximalUP-(33)-sigma38p10 1608844 Gap 4.0 bits
proximalUP 6.8 bits
... distalUP-(40)-sigma38p10 1608844 Gap 3.1 bits
... distalUP-proximalUP-sigma38p10 1608844 total 7.9 bits
... proximalUP-(28)
distalUP 2.3 bits
proximalUP 4.3 bits
... distalUP-(42)-s
... distalUP-proxin
distalUP 6.5 bits
... distalUP-(42)-s
... distalUP-proxin
*1608880 . *1608890 . *1608900 . *1608910 .
5′ t c a t a a a g t g g t a g a a a c a c a t t t g c a t t c a a a t c g c g c g 3′
... proximalUP-(36)-sigma38p10 1608900 Gap 3.6 bits
sigma38p10 2.6 bits
sigma38p10 0.7 bits
... proximalUP-(28)-sigma38p10 1608882 Gap 5.5 bits
... distalUP-(42)-sigma38p10 1608882 Gap 3.3 bits
... distalUP-proximalUP-sigma38p10 1608882 total 2.9 bits
... distalUP-(42)-sigma38p10 1608900 Gap 3.3 bits
... distalUP-proximalUP-sigma38p10 1608900 total 4.8 b
*1608920 . *1608930 . *1608940 . *1608950
5′ t a a t g a a t a a a g a t g t c a g a c a a c t t c c t c a c c g t a a c 3′
proximalUP 4.9 bits

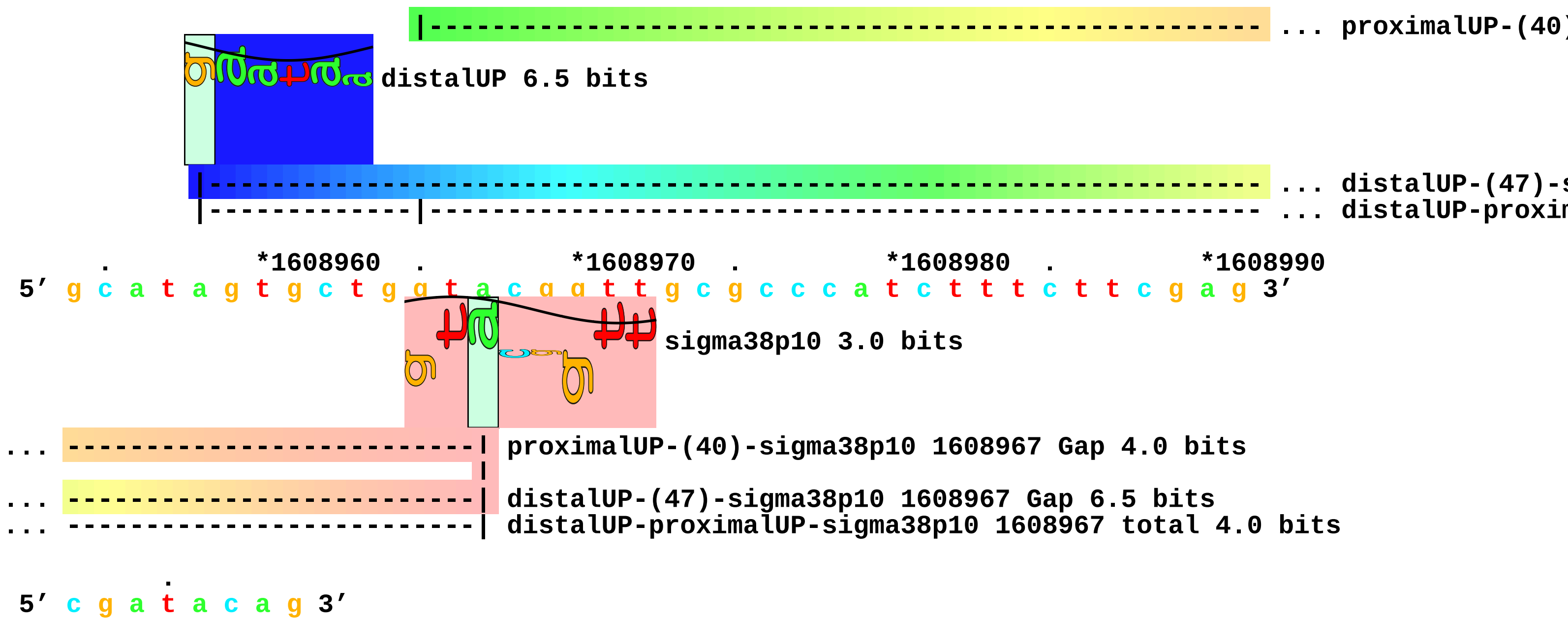
... proximalUP-(40)
distalUP 6.5 bits
... distalUP-(47)-s
... distalUP-proxim
*1608960
*1608970
*1608980
*1608990
5′ g c a t a g t g c t g g t a c g g t t g c g c c c a t c t t t c t t c g a g 3′
sigma38p10 3.0 bits
... proximalUP-(40)-sigma38p10 1608967 Gap 4.0 bits
... distalUP-(47)-sigma38p10 1608967 Gap 6.5 bits
... distalUP-proximalUP-sigma38p10 1608967 total 4.0 bits
5′ c g a t a c a g 3′

piece 19, NC_000913, ydgA, config: linear, direction: -, begin: 1687957, end: 1687694

```
      .         *1687950   .         *1687940   .         *1687930   .         *1687920
5' t c t t g c c t g t a t a c c a t g c g c c g c c t g t c c a g a c t a c g 3'
   T timestamp:_2018Sep20_11-35-24

        .         *1687910   .         *1687900   .         *1687890   .
5' c c t a g c g c a a c a a t g a c g c c t a c c g c t a c c g a t t t 3'

    *1687880  .         *1687870   .         *1687860   .         *1687850   .
5' a t t c a t a a t g a t t a t c c a t a a a a t g a a a t c a g g c g g a c 3'
                                     proximalUP 5.6 bits
                                     |---------------------------- ... proximalUP-(42)
                                distalUP 5.6 bits
                              |--------------------------------------------- ... distalUP-(50)-s
                              |---------------|---------------------------- ... sigma38p10-prox

      *1687840  .         *1687830   .         *1687820   .         *1687810
5' t g g c c g c c t g a a g g t g t t a t a a g c c t t t a a t a a g c t t a 3'
                                                    sigma38p10 1.1 bits
... ---------------------------------------------------| proximalUP-(42)-sigma38p10 1687815 Gap
                                  proximalUP 3.4 bits
... ---------------------------------------------------| distalUP-(50)-sigma38p10 1687815 Gap 4
... ---------------------------------------|---------| sigma38p10-proximalUP-distalUP 1687815
                                          |-------------------------------- ... proximalUP-(40)
                              distalUP 4.3 bits
                              |--------------------------------------------- ... distalUP-(47)-s
                              |-------------|------------------------------- ... sigma38p10-prox

   .         *1687800   .         *1687790   .         *1687780   .         *1687770
5' g c a a g a g a t g t t a a t t t t t t c a g t a a g c t c t t a c a g c t 3'
                                                sigma38p10 3.6 bits
                              proximalUP 6.2 bits
```

```
                                     |---------------------------------- ... proximalUP-(35)
```

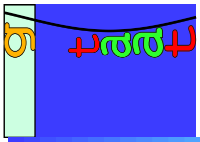

distalUP 5.5 bits

```
                  |----------------------------------------------------- ... distalUP-(46)-s
... ------------------------------------------------| proximalUP-(40)-sigma38p10 1687781 Gap 4.0 bit
                  |----------------------|-------------------------------- ... sigma38p10-prox
... ------------------------------------------------| distalUP-(47)-sigma38p10 1687781 Gap 6.5 bits
... ------------------------------------------------| sigma38p10-proximalUP-distalUP 1687781 total (

        .         *1687760  .         *1687750  .         *1687740  .         *1687730
5' t g t t g t a a a c a c g c g c t a a a c g g c c g t g g c c t t t g a c a 3'
```

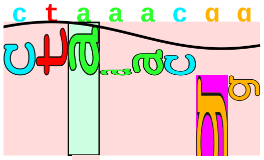

sigma38p10 1.5 bits

```
... ----------------------------------| proximalUP-(35)-sigma38p10 1687750 Gap 3.1 bits
... ----------------------------------| distalUP-(46)-sigma38p10 1687750 Gap 4.1 bits
... ----------------------------------| sigma38p10-proximalUP-distalUP 1687750 total 6.0 bits

          .         *1687720  .         *1687710  .         *1687700  .
5' g t c a c c g g t g a t t c g t t g g c g g c a a t a a a c g c t g a t 3'
```

piece 20, NC_000913, lpp, config: linear, direction: -, begin: 1755549, end: 1755300

```
           .         *1755540  .         *1755530  .         *1755520  .
5' c t g a a c g t c a g a a g a c a g c t g a t c g a t t t t a g c g t t g c 3'
   T timestamp:_2018Sep20_11-35-24                            proximalUP 3.9 bits
                                                                  |------------ ... proximalUP-(36)
                                                           distalUP 6.5 bits
                                                          |------------------------- ... distalUP-(43)-s
                                                          |------------|------------ ... sigma38p10-prox

    *1755510  .         *1755500  .         *1755490  .         *1755480  .
5' t g g a g c a a c c t g c c a g c a g a g t a g a a c c c a g g a t t a c c 3'
                                                                  sigma38p10 2.6 bits
... -----------------------------------------------------------| proximalUP-(36)-sigma38p10 1755482 G
                                                               |
... -----------------------------------------------------------| distalUP-(43)-sigma38p10 1755482 Gap
... -----------------------------------------------------------| sigma38p10-proximalUP-distalUP 17554

        *1755470  .         *1755460  .         *1755450  .         *1755440
5' g c g c c c a g t a c c a g t t t a g t a g c t t t c a t t a t t a a t a c 3'
                                                                  proximalUP 3.3 bits
                                                                      |- ... proximalUP-(33)
                                                           distalUP 6.3 bits
                                                          |---------------------- ... distalUP-(44)-s
                                                          |--------------------|- ... sigma38p10-prox

   .         *1755430  .         *1755420  .         *1755410  .         *1755400
5' c c t c t a g a t t g a g t t a a t c t c c a t g t a g c g t t a c a a g t 3'
                          proximalUP 6.8 bits                     sigma38p10 1.2 bits
... -----------------------------------------------------------------| proximalUP-(33)-sigma38p10 175
                                   |-------------------------------- ... proximalUP-(28)
... -----------------------------------------------------------------| distalUP-(44)-sigma38p10 17554
... -----------------------------------------------------------------| sigma38p10-proximalUP-distalUP
              distalUP 1.5 bits                                 proximalUP 0.3 bits
```

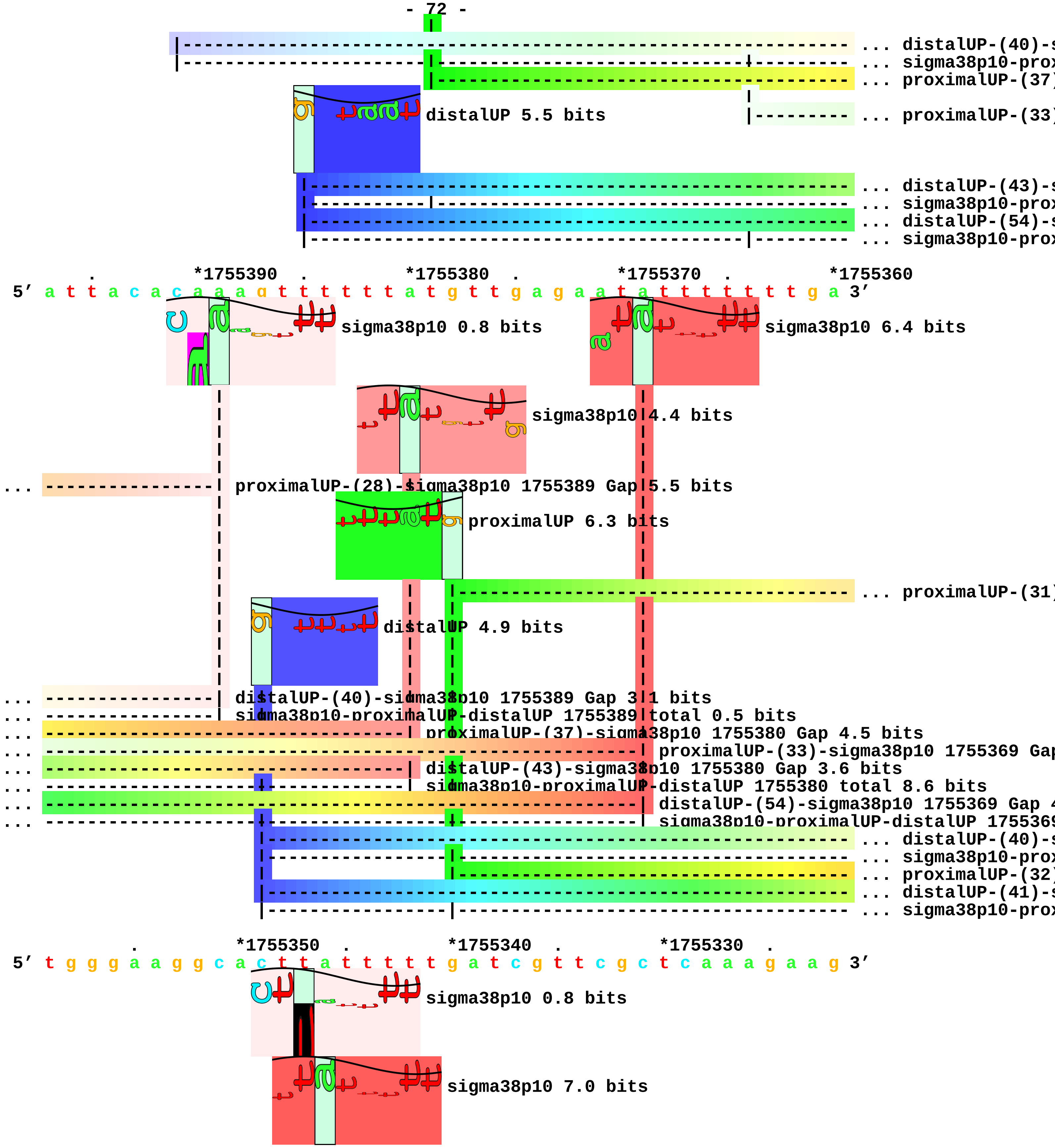
... distalUP-(40)-s
... sigma38p10-prox
... proximalUP-(37)
distalUP 5.5 bits
... proximalUP-(33)
... distalUP-(43)-s
... sigma38p10-prox
... distalUP-(54)-s
... sigma38p10-prox
. *1755390 . *1755380 . *1755370 . *1755360
5′ a t t a c a c a a a g t t t t t t a t g t t g a g a a t a t t t t t t t g a 3′
sigma38p10 0.8 bits
sigma38p10 6.4 bits
sigma38p10 4.4 bits
... proximalUP-(28)-sigma38p10 1755389 Gap 5.5 bits
proximalUP 6.3 bits
... proximalUP-(31)
distalUP 4.9 bits
... distalUP-(40)-sigma38p10 1755389 Gap 3.1 bits
... sigma38p10-proximalUP-distalUP 1755389 total 0.5 bits
... proximalUP-(37)-sigma38p10 1755380 Gap 4.5 bits
... proximalUP-(33)-sigma38p10 1755369 Gap
... distalUP-(43)-sigma38p10 1755380 Gap 3.6 bits
... sigma38p10-proximalUP-distalUP 1755380 total 8.6 bits
... distalUP-(54)-sigma38p10 1755369 Gap 4
... sigma38p10-proximalUP-distalUP 1755369
... distalUP-(40)-s
... sigma38p10-prox
... proximalUP-(32)
... distalUP-(41)-s
... sigma38p10-prox
. *1755350 . *1755340 . *1755330 .
5′ t g g g a a g g c a c t t a t t t t t t g a t c g t t c g c t c a a a g a a g 3′
sigma38p10 0.8 bits
sigma38p10 7.0 bits

```
... ------------------------| proximalUP-(31)-sigma38p10 1755347 Gap 4.7 bits
... ------------------------| distalUP-(40)-sigma38p10 1755347 Gap 3.1 bits
... ------------------------| sigma38p10-proximalUP-distalUP 1755347 total 4.3 bits
... --------------------------| proximalUP-(32)-sigma38p10 1755346 Gap 4.3 bits
... --------------------------| distalUP-(41)-sigma38p10 1755346 Gap 3.1 bits
... --------------------------| sigma38p10-proximalUP-distalUP 1755346 total 10.7 bits

      *1755320  .         *1755310  .         *1755300
 5' c a t c g a a c g c t g g g t t t t a g a t 3'
```

piece 21, NC_000913, ynhG, config: linear, direction: +, begin: 1756770, end: 1756935

```
   *1756770  .         *1756780  .         *1756790  .         *1756800  .
5’ c g g a g a c t c a c g c t g a a a c g t c g g a t g g c g c t t a t g t t 3’
   T timestamp:_2018Sep20_11-35-24

       *1756810  .         *1756820  .         *1756830  .         *1756840  .
5’ c a c c t g a a a c c a a a a c a c t c c t g t g c a g g t c a g t g t a a 3’

           *1756850  .         *1756860  .         *1756870  .         *1756880
5’ a c a t t g a c c a t c c g g c a a t g t g a g c c a a c c g g a t g a a a 3’

     .         *1756890  .         *1756900  .         *1756910  .         *1756920
5’ g c t g t c c t t t t a g t t t a g c t a a g t g c a g c g g c t t t g g c 3’

         .         *1756930  .
5’ g c g a a t t g c g c g a a 3’
```

piece 22, NC_000913, sdaA, config: linear, direction: -, begin: 1894946, end: 1894613

. *1894940 . *1894930 . *1894920 . *1894910

5′ c c t g a c a a g g t g a c t g g a c t t c c a g t a a c g a a c t a a t a 3′

T timestamp:_2018Sep20_11-35-24

... proximalUP

distalUP 3.5 bits

... distalUP-(44)-s

... sigma38p10-prox

. *1894900 . *1894890 . *1894880 .

5′ a c a g c g c a t a g t g t a a g a g g g a a c g c g g c g a c t g g c t t 3′

... proximalUP 4.3 bits

... sigma38p10

... proximalUP-(37)

... distalUP-(44)-s

... sigma38p10-prox

*1894870 . *1894860 . *1894850 . *1894840 .

5′ a a c t a t t c a c a t g a a t t a a a c t a a t c a a a a a c c g c g a t 3′

... sigma38p10 1.4 bits

... proximalUP-(33)

... proximalUP-(37)-sigma38p10 1894870 Gap 4.5 bits

proximalUP 4.3 bits

... distalUP-(44)-sigma38p10 1894870 Gap 2.8 bits

... proximalUP-(37)

... sigma38p10-proximalUP-distalUP 1894870 total 1.9 bits

... proximalUP-(40)

proximalUP 3.7 bits

distalUP 5.7 bits

... proximalUP-(37)

distalUP 0.4 bits

... distalUP-(43)-s

... distalUP-(42)-s

... sigma38p10-prox

proximalUP 5.0 bits

... distalUP

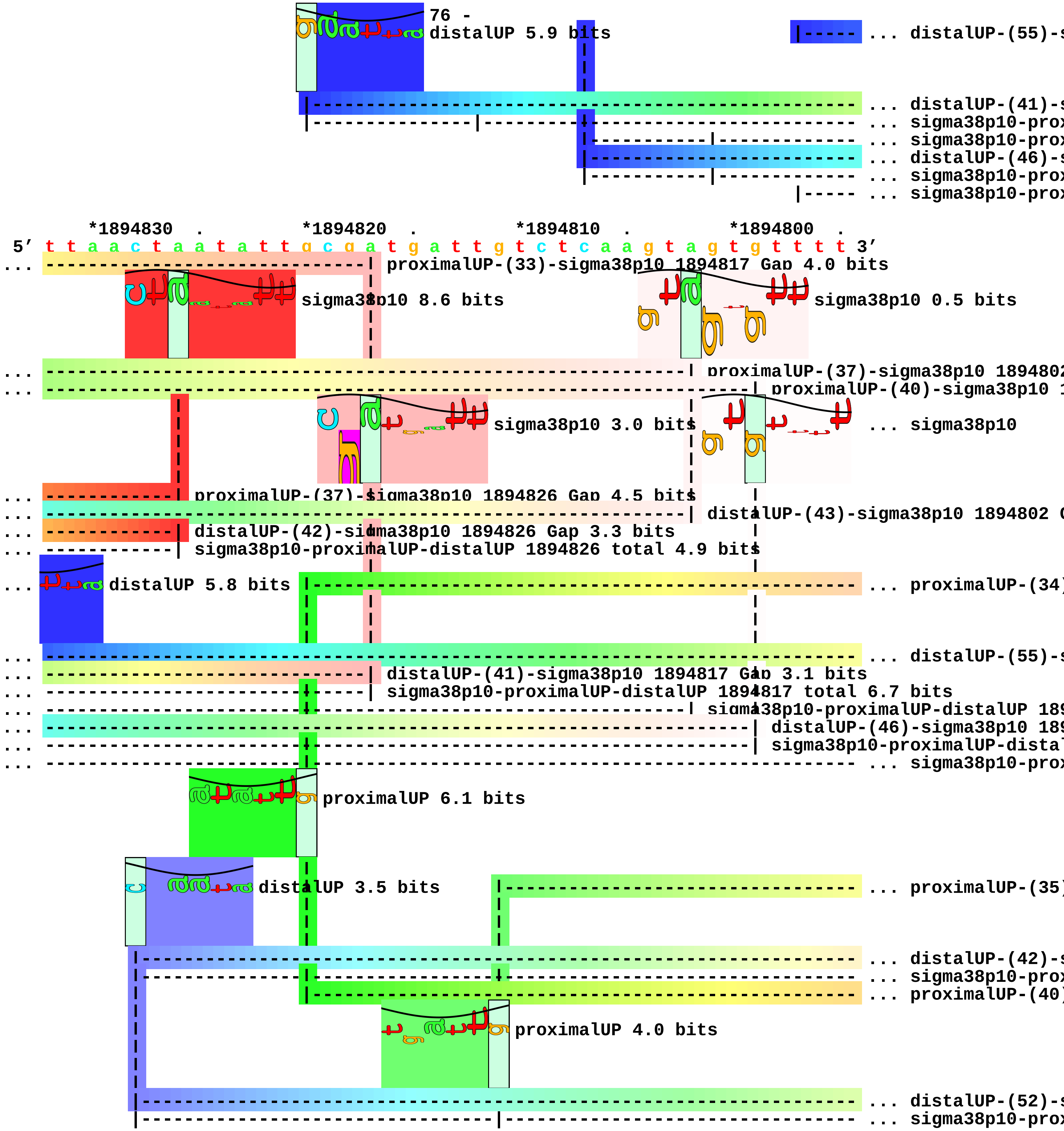
distalUP 5.9 bits
... distalUP-(55)-s
... distalUP-(41)-s
... sigma38p10-prox
... sigma38p10-prox
... distalUP-(46)-s
... sigma38p10-prox
... sigma38p10-prox
*1894830 .
*1894820 .
*1894810 .
*1894800 .
5′ t t a a c t a a t a t t g c g a t g a t t g t c t c a a g t a g t g t t t t 3′
... proximalUP-(33)-sigma38p10 1894817 Gap 4.0 bits
sigma38p10 8.6 bits
sigma38p10 0.5 bits
... proximalUP-(37)-sigma38p10 189480
... proximalUP-(40)-sigma38p10 1
sigma38p10 3.0 bits
... sigma38p10
... proximalUP-(37)-sigma38p10 1894826 Gap 4.5 bits
... distalUP-(43)-sigma38p10 1894802 G
... distalUP-(42)-sigma38p10 1894826 Gap 3.3 bits
... sigma38p10-proximalUP-distalUP 1894826 total 4.9 bits
... distalUP 5.8 bits
... proximalUP-(34)
... distalUP-(55)-s
... distalUP-(41)-sigma38p10 1894817 Gap 3.1 bits
... sigma38p10-proximalUP-distalUP 1894817 total 6.7 bits
... sigma38p10-proximalUP-distalUP 189
... distalUP-(46)-sigma38p10 189
... sigma38p10-proximalUP-distal
... sigma38p10-prox
proximalUP 6.1 bits
distalUP 3.5 bits
... proximalUP-(35)
... distalUP-(42)-s
... sigma38p10-prox
... proximalUP-(40)
proximalUP 4.0 bits
... distalUP-(52)-s
... sigma38p10-prox

*1894790 . *1894780 . *1894770 . *1894760
5′ g t a g a t g t a a a g a t a a g t a t a g t c a g g g t t t c a c a c c a 3′

sigma38p10 1.5 bits

sigma38p10 2.5 bits

sigma38p10 2.5 bits

... sigma38p10 0.1 bits

... ----------------| proximalUP-(34)-sigma38p10 1894786 Gap 2.5 bits
... ----------------------------| distalUP-(55)-sigma38p10 1894780 Gap 4.9 bits

... ----------------------------| sigma38p10-proximalUP-distalUP 1894780 total 5.6 bits

... ------------------------------------| proximalUP-(35)-sigma38p10 1894776 Gap 3.1 bits
... ----------------| distalUP-(42)-sigma38p10 1894786 Gap 3.3 bits
... ----------------| sigma38p10-proximalUP-distalUP 1894786 total 5.5 bits
... ----------------------------| proximalUP-(40)-sigma38p10 1894780 Gap 4.0 bits

... ------------------------------------| distalUP-(52)-sigma38p10 1894776 Gap 5.5 bits
... ------------------------------------| sigma38p10-proximalUP-distalUP 1894776 total 1.6 bits

. *1894750 . *1894740 . *1894730 . *1894720
5′ a t t t g c a g c g c c a g c t c a c g a a t t a t g c c t g c g g t c a t 3′

. *1894710 . *1894700 . *1894690 .
5′ t c c c c a t a c a a a a t a c t g t t c g t a c c a g g a c a g c c a t a 3′

*1894680 . *1894670 . *1894660 . *1894650 .
5′ c c c g a t g t g a a t c a c c a c g g c g g t a g a t a t c t a a a g g g 3′

proximalUP 5.1 bits

|----------------------------------------------- ... proximalUP-(42)

distalUP 3.3 bits

```
                      |------------------------------------------------------------ ... distalUP-(46)-s
                      |-------|---------------------------------------------------- ... sigma38p10-prox

          *1894640  .         *1894630  .         *1894620  .
5' t g a t a a c g a c c c a g a t g t a a t g c c t g g g c g 3'
```

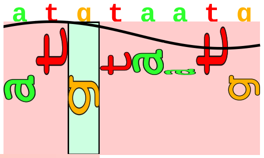

sigma38p10 2.2 bits

```
... --------------------------------| proximalUP-(42)-sigma38p10 1894626 Gap 5.1 bits
                                    |
... --------------------------------| distalUP-(46)-sigma38p10 1894626 Gap 4.1 bits
... --------------------------------| sigma38p10-proximalUP-distalUP 1894626 total 1.3 bits
```

piece 23, NC_000913, yobF, config: linear, direction: +, begin: 1905497, end: 1905834

```
         *1905500  .         *1905510  .         *1905520  .         *1905530
5' c a t a a c a t a g a t a a a a c t g a c a c a t t a c t g c a t g a g g c 3'
 T timestamp:_2018Sep20_11-35-24

 .         *1905540  .         *1905550  .         *1905560  .         *1905570
5' a c c a a t a t a a g g c t c g g c a g a g a a g c g g t a t t c a a c g t 3'
```

proximalUP 3.4 bits

... proximalUP-(32)

distalUP 6.4 bits

... distalUP-(40)-s
... distalUP-proxim
... proximalUP-(38)
... distalUP-(46)-s
... distalUP-proxim

```
   .         *1905580  .         *1905590  .         *1905600  .         *1905610
5' c a a c g t g t t t a c t c a g g a c t t c t t t a c t g a a a a t g c c a 3'
```

sigma38p10 0.1 bits

proximalUP 5.6 bits

... -------- proximalUP-(32)-sigma38p10 1905577 Gap 4.3 bits

... proximalUP-(35)

sigma38p10 1.1 bits

distalUP 7.2 bits

... -------- distalUP-(40)-sigma38p10 1905577 Gap 3.1 bits
... -------- distalUP-proximalUP-sigma38p10 1905577 total 2[illegible]5 bits
... -------------------- proximalUP-(38)-sigma38p10 1905583 Gap 3.2 bits
... -------------------- distalUP-(46)-sigma38p10 1905583 Ga[illegible] 4.1 bits
... -------------------- distalUP-proximalUP-sigma38p10 190[illegible]583 total 3.5 bits

... distalUP-(41)-s
... proximalUP-(42)
... distalUP-proxim
... distalUP-(48)-s
... distalUP-proxim

```
         .         *1905620  .         *1905630  .         *1905640  .
5' c a c a t a a a c a g a a c t g t a c c t c g t t t a a c c c g a a a t c t 3'
```

distalUP 5.6 bits

sigma38p10 0.7 bits

... proximalUP-(35)-sigma38p10 1905[illegible]

... sigma38p10

... distalUP-(41)-sigma38p10 190564[illegible]

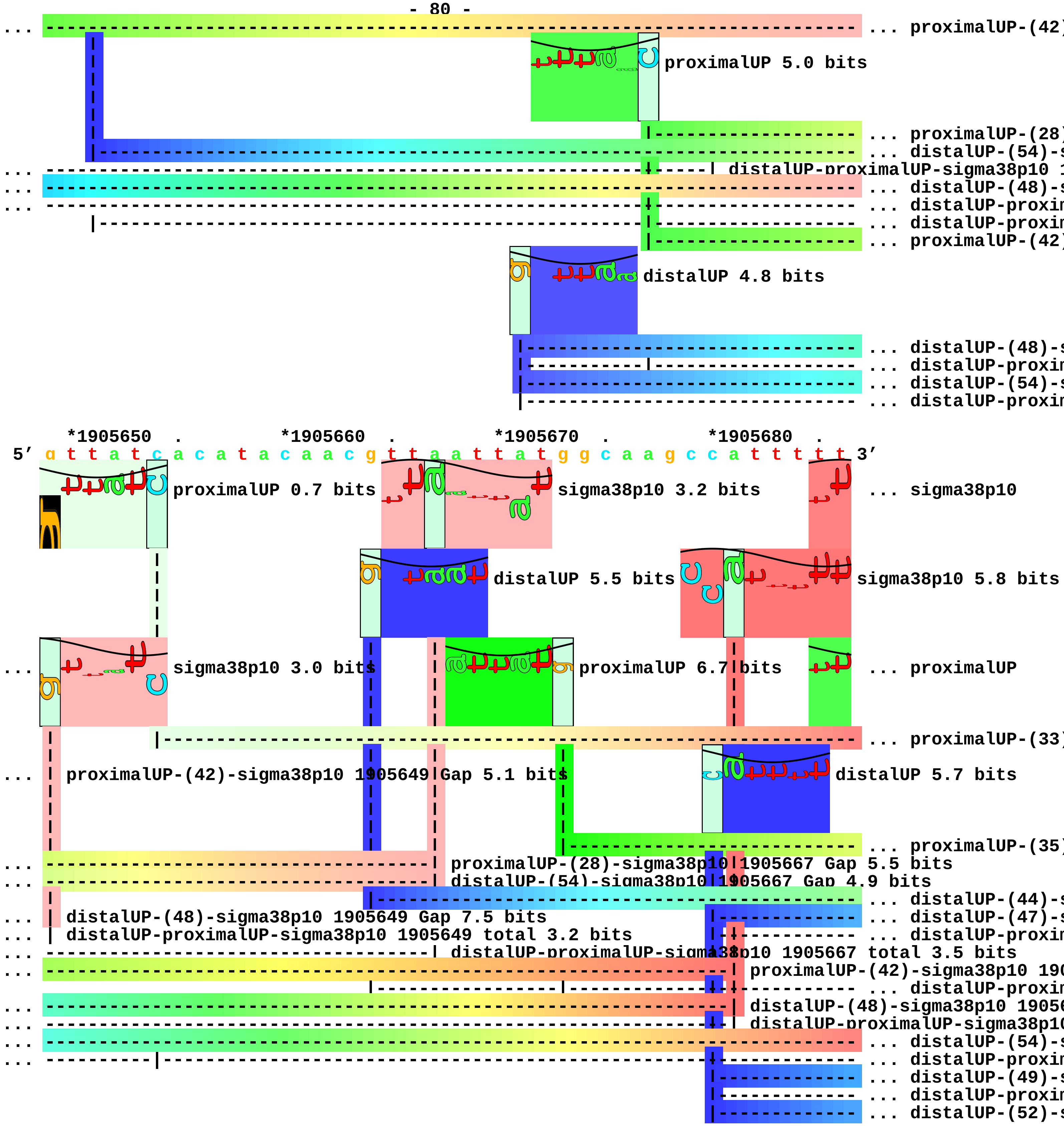
... proximalUP-(42)
proximalUP 5.0 bits
... proximalUP-(28)
... distalUP-(54)-s
distalUP-proximalUP-sigma38p10 1
... distalUP-(48)-s
... distalUP-proxim
... distalUP-proxim
... proximalUP-(42)
distalUP 4.8 bits
... distalUP-(48)-s
... distalUP-proxim
... distalUP-(54)-s
... distalUP-proxim
*1905650 . *1905660 . *1905670 . *1905680 .
5' g t t a t c a c a t a c a a c g t t a a t t a t g g c a a g c c a t t t t t 3'
proximalUP 0.7 bits
sigma38p10 3.2 bits
... sigma38p10
distalUP 5.5 bits
sigma38p10 5.8 bits
sigma38p10 3.0 bits
proximalUP 6.7 bits
... proximalUP
... proximalUP-(33)
proximalUP-(42)-sigma38p10 1905649 Gap 5.1 bits
distalUP 5.7 bits
... proximalUP-(35)
proximalUP-(28)-sigma38p10 1905667 Gap 5.5 bits
distalUP-(54)-sigma38p10 1905667 Gap 4.9 bits
... distalUP-(44)-s
distalUP-(48)-sigma38p10 1905649 Gap 7.5 bits
... distalUP-(47)-s
distalUP-proximalUP-sigma38p10 1905649 total 3.2 bits
... distalUP-proxim
distalUP-proximalUP-sigma38p10 1905667 total 3.5 bits
proximalUP-(42)-sigma38p10 190
... distalUP-proxim
distalUP-(48)-sigma38p10 1905
distalUP-proximalUP-sigma38p10
... distalUP-(54)-s
... distalUP-proxim
... distalUP-(49)-s
... distalUP-proxim
... distalUP-(52)-s

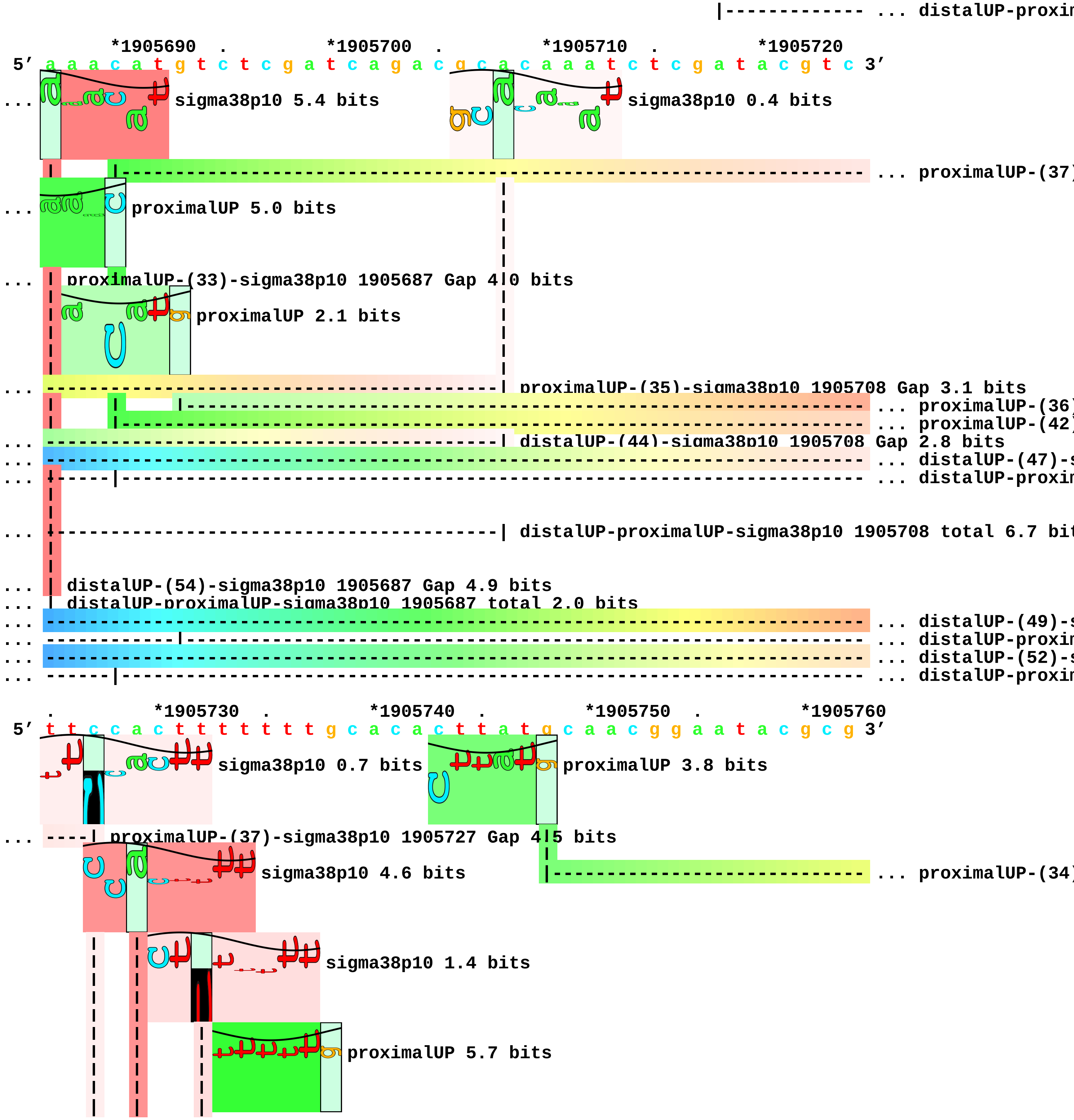
|------------- ... distalUP-proxin
*1905690 . *1905700 . *1905710 . *1905720
5′ a a a c a t g t c t c g a t c a g a c g c a c a a a t c t c g a t a c g t c 3′
... sigma38p10 5.4 bits
sigma38p10 0.4 bits
... proximalUP-(37)
... proximalUP 5.0 bits
... | proximalUP-(33)-sigma38p10 1905687 Gap 4.0 bits
proximalUP 2.1 bits
... proximalUP-(35)-sigma38p10 1905708 Gap 3.1 bits
... proximalUP-(36)
... proximalUP-(42)
... distalUP-(44)-sigma38p10 1905708 Gap 2.8 bits
... distalUP-(47)-s
... distalUP-proxin
... distalUP-proximalUP-sigma38p10 1905708 total 6.7 bit
... | distalUP-(54)-sigma38p10 1905687 Gap 4.9 bits
... | distalUP-proximalUP-sigma38p10 1905687 total 2.0 bits
... distalUP-(49)-s
... distalUP-proxin
... distalUP-(52)-s
... distalUP-proxin
. *1905730 . *1905740 . *1905750 . *1905760
5′ t t c c a c t t t t t t t t g c a c a c t t a t g c a a c g g a a t a c g c g 3′
sigma38p10 0.7 bits
proximalUP 3.8 bits
... ----| proximalUP-(37)-sigma38p10 1905727 Gap 4.5 bits
sigma38p10 4.6 bits
... proximalUP-(34)
sigma38p10 1.4 bits
proximalUP 5.7 bits

... proximalUP-(32)
... proximalUP-(36)-sigma38p10 1905729 Gap 3.6 bits
... proximalUP-(42)-sigma38p10 1905732 Gap 5.1 bits
distalUP 4.1 bits
... distalUP-(47)-sigma38p10 1905727 Gap 6.5 bits
... distalUP-proximalUP-sigma38p10 1905727 total 0.5 bits
... distalUP-(40)-s
... distalUP-proxim
... proximalUP-(38)
... distalUP-(46)-s
... distalUP-proxim
... distalUP-(52)-s
... distalUP-proxim
... distalUP-(49)-sigma38p10 1905729 Gap 5.9 bits
... distalUP-proximalUP-sigma38p10 1905729 total 2.9 bits
... distalUP-(52)-sigma38p10 1905732 Gap 5.5 bits
... distalUP-proximalUP-sigma38p10 1905732 total 1.5 bits

. *1905770 . *1905780 . *1905790 . *1905800
5′ c c g a t a t a t c a t t g t g c t t a a c c t t g c c a g t t c a g g c a 3′

sigma38p10 5.2 bits
sigma38p10 4.6 bits
... proximalUP-(34)-sigma38p10 1905782 Gap 2.5 bits
distalUP 5.9 bits
sigma38p10 5.7 bits
proximalUP 5.7 bits
... proximalUP-(34)
... proximalUP-(32)-sigma38p10 1905770 Gap 4.3 bits
... proximalUP-(36)
... distalUP-(43)-s
... distalUP-proxim
proximalUP 2.5 bits
... distalUP-(53)-s
... distalUP-(40)-sigma38p10 1905770 Gap 3.1 bits
... distalUP-proximalUP-sigma38p10 1905770 total 7.6 bits
... proximalUP-(38)-sigma38p10 1905776 Gap 3.2 bits
... distalUP-(46)-sigma38p10 1905776 Gap 4.1 bits
... distalUP-proximalUP-sigma38p10 1905776 total 7.1 bits
... distalUP-(52)-sigma38p10 1905782 Gap 5.5 bits
... distalUP-proximalUP-sigma38p10 1905782 total 5.6 bits

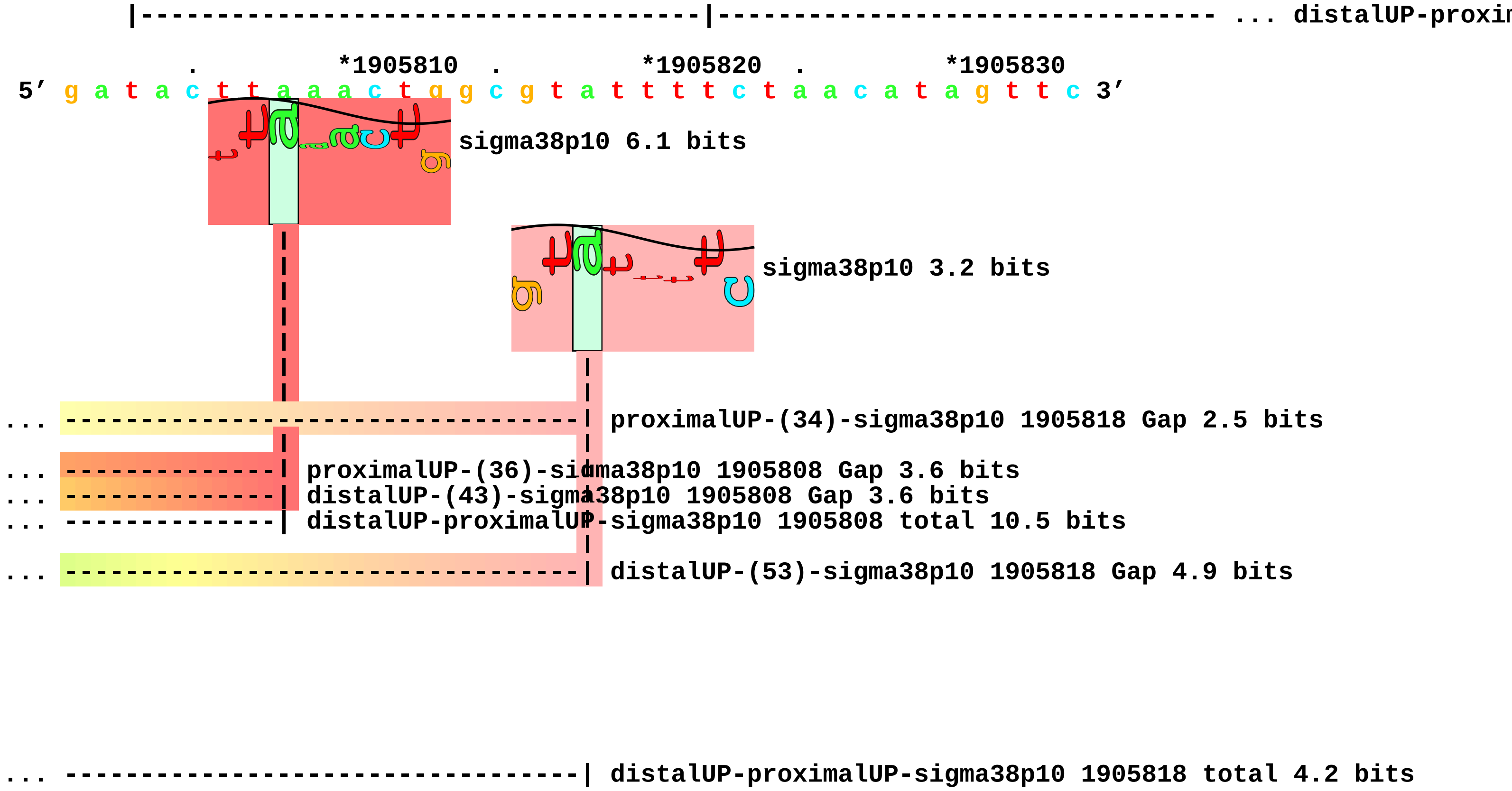
|-----------------------------------|-------------------------------- ... distalUP-proxin
. *1905810 . *1905820 . *1905830
5′ g a t a c t t a a a c t g g c g t a t t t t c t a a c a t a g t t c 3′
sigma38p10 6.1 bits
sigma38p10 3.2 bits
... ----------------------------------| proximalUP-(34)-sigma38p10 1905818 Gap 2.5 bits
... --------------| proximalUP-(36)-sigma38p10 1905808 Gap 3.6 bits
... --------------| distalUP-(43)-sigma38p10 1905808 Gap 3.6 bits
... --------------| distalUP-proximalUP-sigma38p10 1905808 total 10.5 bits
... ----------------------------------| distalUP-(53)-sigma38p10 1905818 Gap 4.9 bits
... ----------------------------------| distalUP-proximalUP-sigma38p10 1905818 total 4.2 bits

piece 24, NC_000913, yebW, config: linear, direction: -, begin: 1920253, end: 1919983

```
        *1920250   .        *1920240   .        *1920230   .        *1920220
5' g t t g a t c g c t c a t a g a a t a a a g a c a c t g c t g t t c c g t g 3'
```

proximalUP 4.9 bits

T timestamp:_2018Sep20_11-35-24

... proximalUP-(34)

distalUP 6.5 bits

... distalUP-(41)-s
... sigma38p10-prox
... proximalUP-(37)
... distalUP-(44)-s
... sigma38p10-prox

```
.        *1920210   .        *1920200   .        *1920190   .        *1920180
5' t t g t a g a c a t c c a c a a a c a a t a t c t t c a c a a c c g c c a t c 3'
```

sigma38p10 0.6 bits

... proximalUP-(34)-sigma38p10 1920198 Gap 2.5 bits

sigma38p10 1.8 bits

... distalUP-(41)-sigma38p10 1920198 Gap 3.1 bits
... sigma38p10-proximalUP-distalUP 1920198 total 6.5 bits
... proximalUP-(37)-sigma38p10 1920195 Gap 4.5 bits
... distalUP-(44)-sigma38p10 1920195 Gap 2.8 bits
... sigma38p10-proximalUP-distalUP 1920195 total 6.0 bits

proximalUP 3.1 bits

... proximalUP-(41)

distalUP 6.4 bits

... distalUP-(50)-s
... sigma38p10-prox

```
     .        *1920170   .        *1920160   .        *1920150   .        *1920140
5' c a g g t a g c a a a c a a a a a g t a c c a g c g c g a a c a t t t c a t 3'
```

distalUP 5.7 bits

sigma38p10 0.6 bits

85 -

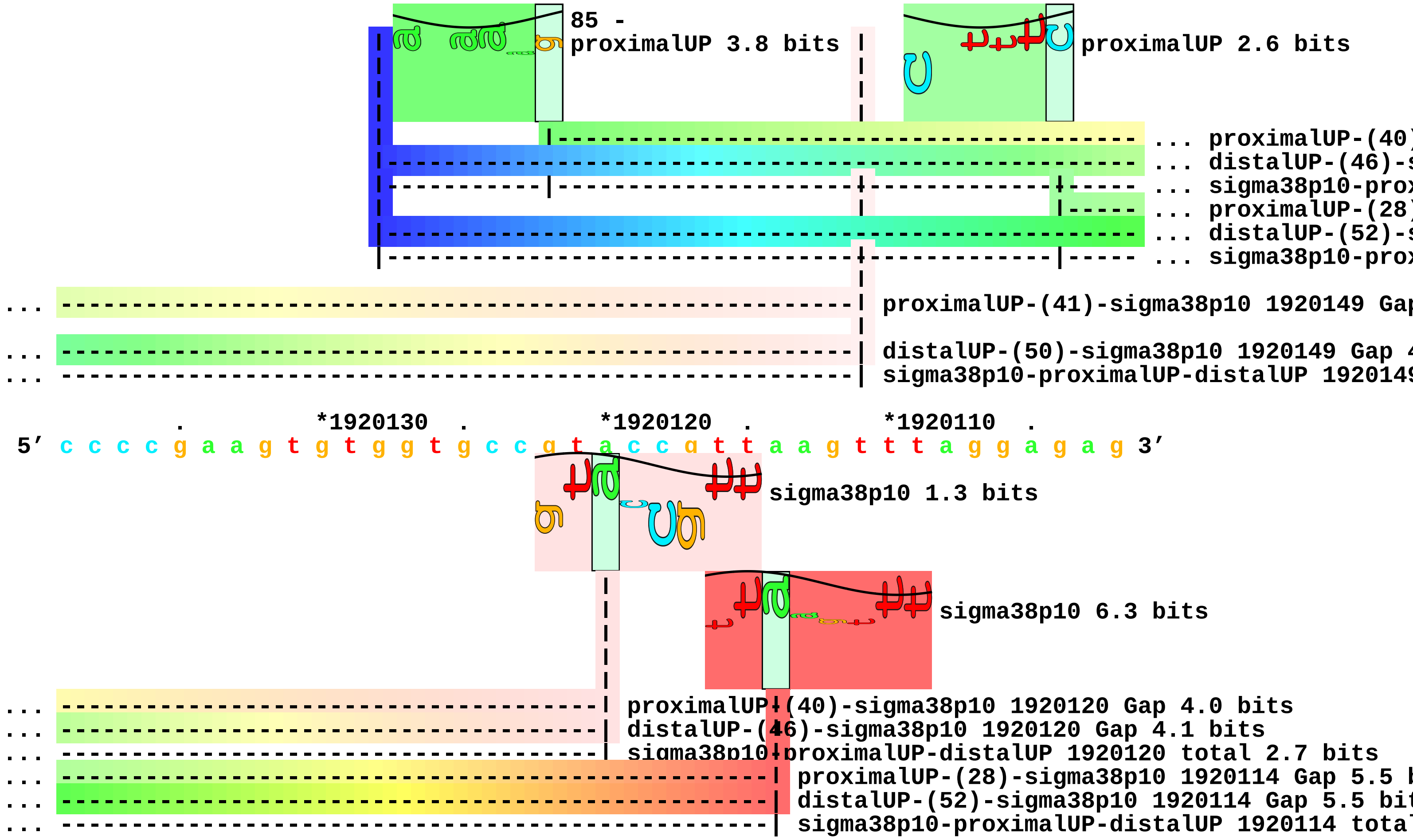


*1920100 . *1920090 . *1920080 . *1920070 .
5′ a t t t t a c a a c g g g g g a a t a a c c a g g a c a a a t a a c c c g c 3′

*1920060 . *1920050 . *1920040 . *1920030
5′ c a t g c c g a c g g g t t c t t t t t g g a t c a g g c a a g a c g c a t 3′

. *1920020 . *1920010 . *1920000 . *1919990
5′ a a t c c a g g c a t c a g a t t g t t g a t c g t a a c g g g t a c g g t 3′

.
5′ a a a g g 3′

piece 25, NC_000913, ryeB, config: linear, direction: +, begin: 1921100, end: 1921349

```
   *1921100   .          *1921110   .          *1921120   .          *1921130   .
5' a a g g a a a t g c t t c t g g c t t t t a a c a g a t a a a a a g a g a c 3'
   T timestamp:_2018Sep20_11-35-24

     *1921140   .          *1921150   .          *1921160   .          *1921170   .
5' c g a a c a c g a t t c c t g t a t t c g g t c c a g g g a a a t g g c t c 3'

       *1921180   .          *1921190   .          *1921200   .          *1921210
5' t t g g g a g a g a g c c g t g c g c t a a a a g t t g g c a t t a a t g c 3'
                                      proximalUP 3.4 bits          |--- ... proximalUP-(43)
                                      |--------------------------- ... proximalUP-(36)
                                     distalUP 4.1 bits
   |------------------------------------------------ ... distalUP-(42)-s
   |------------|------------------------------------ ... distalUP-proxim
                                      proximalUP 6.3 bits
                                     distalUP 5.6 bits
                                      |----------------- ... distalUP-(50)-s
                                      |-------------|--- ... distalUP-proxim

   .          *1921220   .          *1921230   .          *1921240   .          *1921250
5' a g g c t t a g t t g c c t t g c c c t t t a a g a a t a g a t g a c g a c 3'
... ---------------------------------------------------------------------------- ... proximalUP-(43)
... ------------------------------------------| proximalUP-(36)-sigma38p10 1921236 Gap 3.6 bits
                                              sigma38p10 3.6 bits
... ------------------------------------------| distalUP-(42)-sigma38p10 1921236 Gap 3.3 bits
... ------------------------------------------| distalUP-proximalUP-sigma38p10 1921236 total 4.3 b
                                      proximalUP 4.5 bits
                                      |--------------------------- ... proximalUP-(37)
... ---------------------------------------------------------------------------- ... distalUP-(50)-s
... ---------------------------------------------------------------------------- ... distalUP-proxim
                                     distalUP 4.0 bits
```

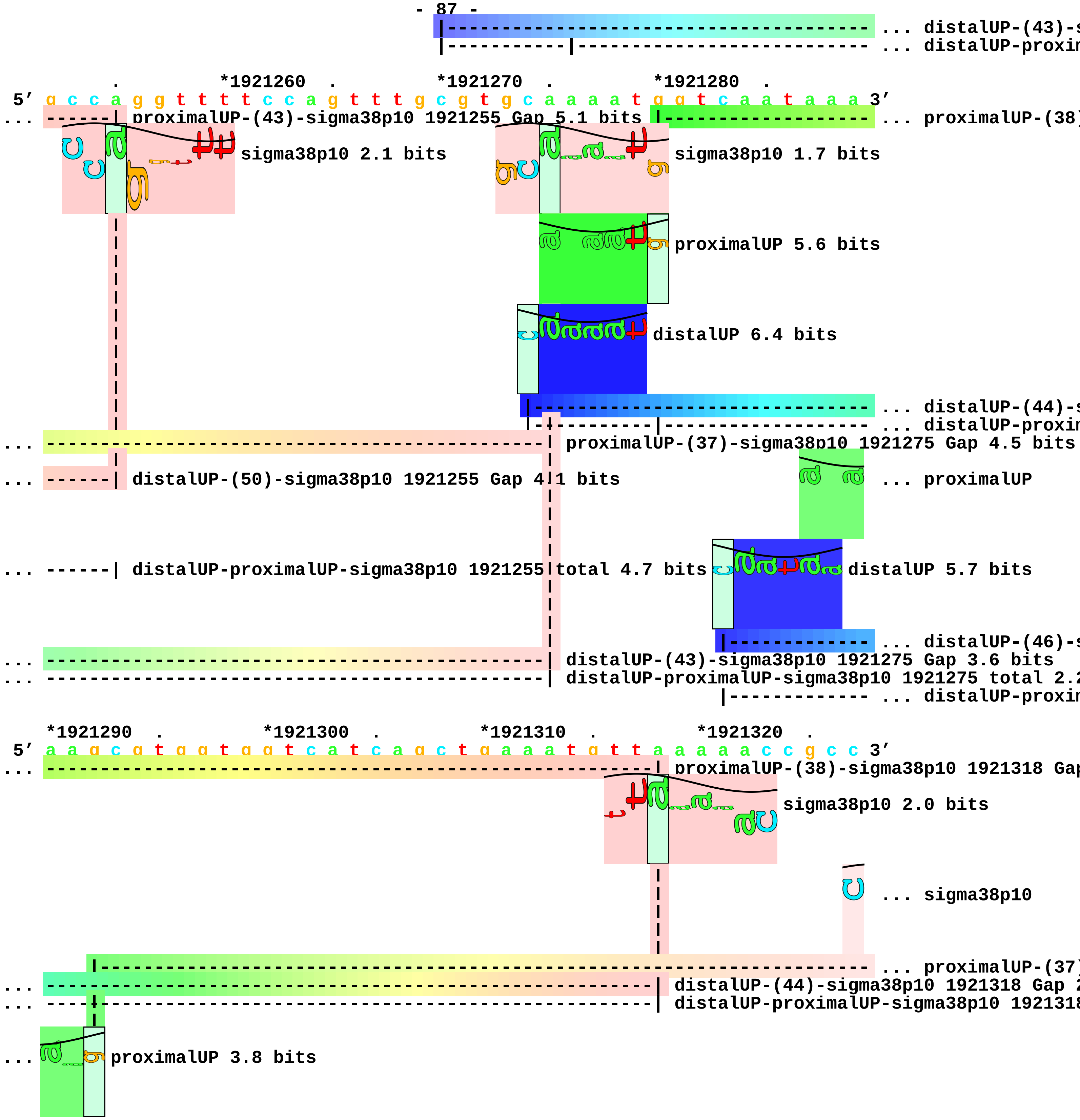
... distalUP-(43)-s
... distalUP-proxim
*1921260 . *1921270 . *1921280 .
5' g c c a g g t t t t c c a g t t t g c g t g c a a a a t g g t c a a t a a a 3'
... ------| proximalUP-(43)-sigma38p10 1921255 Gap 5.1 bits
... proximalUP-(38)
sigma38p10 2.1 bits
sigma38p10 1.7 bits
proximalUP 5.6 bits
distalUP 6.4 bits
... distalUP-(44)-s
... distalUP-proxim
... ---------------------------------------| proximalUP-(37)-sigma38p10 1921275 Gap 4.5 bits
... ------| distalUP-(50)-sigma38p10 1921255 Gap 4.1 bits
... proximalUP
... ------| distalUP-proximalUP-sigma38p10 1921255 total 4.7 bits
distalUP 5.7 bits
... distalUP-(46)-s
... ---------------------------------------| distalUP-(43)-sigma38p10 1921275 Gap 3.6 bits
... ---------------------------------------| distalUP-proximalUP-sigma38p10 1921275 total 2.2
... distalUP-proxim
*1921290 . *1921300 . *1921310 . *1921320 .
5' a a g c g t g g t g g t c a t c a g c t g a a a t g t t a a a a a c c g c c 3'
... proximalUP-(38)-sigma38p10 1921318 Gap
sigma38p10 2.0 bits
... sigma38p10
... proximalUP-(37)
... distalUP-(44)-sigma38p10 1921318 Gap 2
... distalUP-proximalUP-sigma38p10 1921318
... proximalUP 3.8 bits

... [dashed line] ... distalUP-(46)-s

... ----|[dashed line] ... distalUP-proxim

*1921330 . *1921340 .

5′ c g t t c t g g t g a a a g a a c t g a g g 3′

... 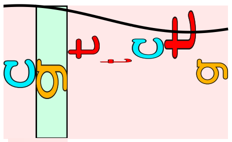 sigma38p10 1.0 bits

... --| proximalUP-(37)-sigma38p10 1921329 Gap 4.5 bits

... --| distalUP-(46)-sigma38p10 1921329 Gap 4.1 bits

... --| distalUP-proximalUP-sigma38p10 1921329 total 1.9 bits piece 26, NC_000913, yodC, config: linear, direction: +, begin: 2026334, end: 2026555

. *2026340 . *2026350 . *2026360 . *2026370

5′ t g a c a a t c a t c c g c g g g c c g c c c t c t t t a a c c g t a a c t 3′

proximalUP 3.7 bits

... proximalUP-(36)

distalUP 2.5 bits

... distalUP-(43)-s

... distalUP-proxim

T timestamp:_2018Sep20_11-35-24

. *2026380 . *2026390 . *2026400 .

5′ t c c t c a c t a a c c a t a a a g c t c a t a c t c g c c t c c t t t t t 3′

sigma38p10 4.2 bits

... proximalUP

... proximalUP-(36)-sigma38p10 2026380 Gap 3.6 bits

distalUP 4.1 bits

... distalUP-(43)-sigma38p10 2026380 Gap 3.6 bits ... distalUP-(41)-s

... distalUP-proximalUP-sigma38p10 2026380 total 3.2 bits

... distalUP-proxim

... distalUP-(42)-s

... distalUP-proxim

*2026410 . *2026420 . *2026430 . *2026440 .

5′ t c g a g t g a a a c t t g t t c a c c t t a g t t g a a g a t g g c g a a 3′

... proximalUP 6.2 bits

... sigma38p10

proximalUP-(34)-sigma38p

... sigma38p10

... distalUP-(41)-sigma38p10

proximalUP-(35)-sigma3

... distalUP-proximalUP-sigm

... distalUP-(42)-sigma38p

... distalUP-proximalUP-si

... proximalUP

... distalUP
... distalUP-(47)-s
... distalUP-proxim

*2026450 . *2026460 . *2026470 . *2026480 .
5' t t t t g c a t a a t a g c c a g t g c g a g t a t t a a t g t g c c t g a 3'
... sigma38p10 0.4 bits
... proximalUP-(40)
... sigma38p10 1.9 bits
... proximalUP 6.1 bits
... distalUP 6.6 bits
... distalUP-(47)-s
... distalUP-proxim

*2026490 . *2026500 . *2026510 . *2026520
5' c c g c g c a t t t t c t c t a c g g c t t t c a c t c c c a g c a c c a c 3'
sigma38p10 0.3 bits
... proximalUP-(40)-sigma38p10 2026492 Gap 4.0 bits
... distalUP-(47)-sigma38p10 2026492 Gap 6.5 bits
... distalUP-proximalUP-sigma38p10 2026492 total 2.6 bits

. *2026530 . *2026540 . *2026550 .
5' g a t g c c g c c g a t g a t a a a t c c a a g a a t c a g a t 3'

piece 27, NC_000913, erfK, config: linear, direction: +, begin: 2061211, end: 2061534

. *2061220 . *2061230 . *2061240 .
5′ t t g t g a t c a g g a a c a g t t a c a g t a a a c g a c t g c c c c a c 3′
T timestamp:_2018Sep20_11-35-24

*2061250 . *2061260 . *2061270 . *2061280 .
5′ t a a a c g g c t a c c c t c t g g a g g t a a t g g a t a a g t t a c c g 3′

*2061290 . *2061300 . *2061310 . *2061320
5′ c c a g g c t a g t a t g g c t g g c a a a a a g c a g a g c a a a t g a g 3′

. *2061330 . *2061340 . *2061350 . *2061360
5′ c a a a g a a t a t t t a c a c g a c g c a t c a t g t c c c t t t c c t a 3′

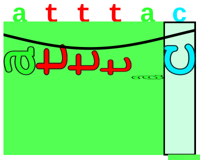
proximalUP 4.9 bits

... proximalUP-(32)
... proximalUP-(38)

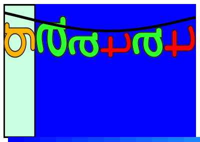
distalUP 7.2 bits

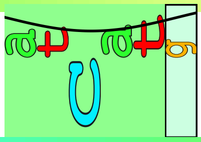
proximalUP 3.2 bits

... distalUP-(47)-s
... distalUP-proxim
... distalUP-(54)-s
... distalUP-proxim

. *2061370 . *2061380 . *2061390 . *2061400
5′ t g t c g c g a a a g c t a t c c g t t a a g t a t a g c t t t t a t c a g 3′
... proximalUP-(32)-sigma38p10 2061383 Gap 4.3 bits
... proximalUP-(38)-sigma38p10 2061376 Gap 3.2 bits

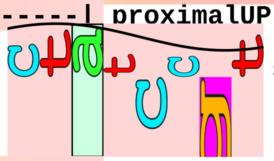
sigma38p10 1.8 bits

... distalUP-(47)-sigma38p10 2061376 Gap 6.5 bits
... distalUP-proximalUP-sigma38p10 2061376 total 4.1 bits
... distalUP-(54)-sigma38p10 2061383 Gap 4.9 bits
... distalUP-proximalUP-sigma38p10 2061383 total 4.3 bits

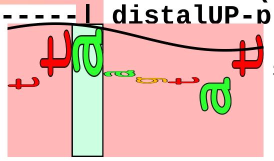
sigma38p10 3.1 bits

. *2061410 . *2061420 . *2061430 .
5′ a c t t t t c g t t t t t a a c t g t t c a a a t c a g a a g t c g t a t t 3′

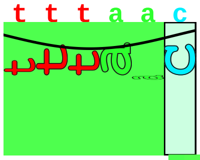
proximalUP 5.0 bits

... proximalUP-(35)

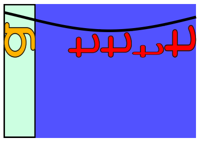
distalUP 4.9 bits

|------------------------------------------------------------ ... distalUP-(43)-s
|---------------|------------------------------------------ ... distalUP-proxim

*2061440 . *2061450 . *2061460 . *2061470 .
5′ c c c c g g t a g a a c a a t a t t a c t g g c a g c a a g t t c g c c c a 3′

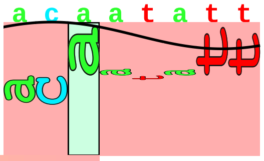

sigma38p10 3.5 bits

... -----------------------| proximalUP-(35)-sigma38p10 2061451 Gap 3.1 bits
... -----------------------| distalUP-(43)-sigma38p10 2061451 Gap 3.6 bits
... -----------------------| distalUP-proximalUP-sigma38p10 2061451 total 6.7 bits

*2061480 . *2061490 . *2061500 . *2061510
5′ t g t t g t t g t a t a t c g c a c a g g c a g c t t c g a t g a t g g g c 3′

. *2061520 . *2061530
5′ a t c g c c a g a g c t g c g c c a c t 3′

piece 28, NC_000913, wbbI, config: linear, direction: +, begin: 2103800, end: 2104249

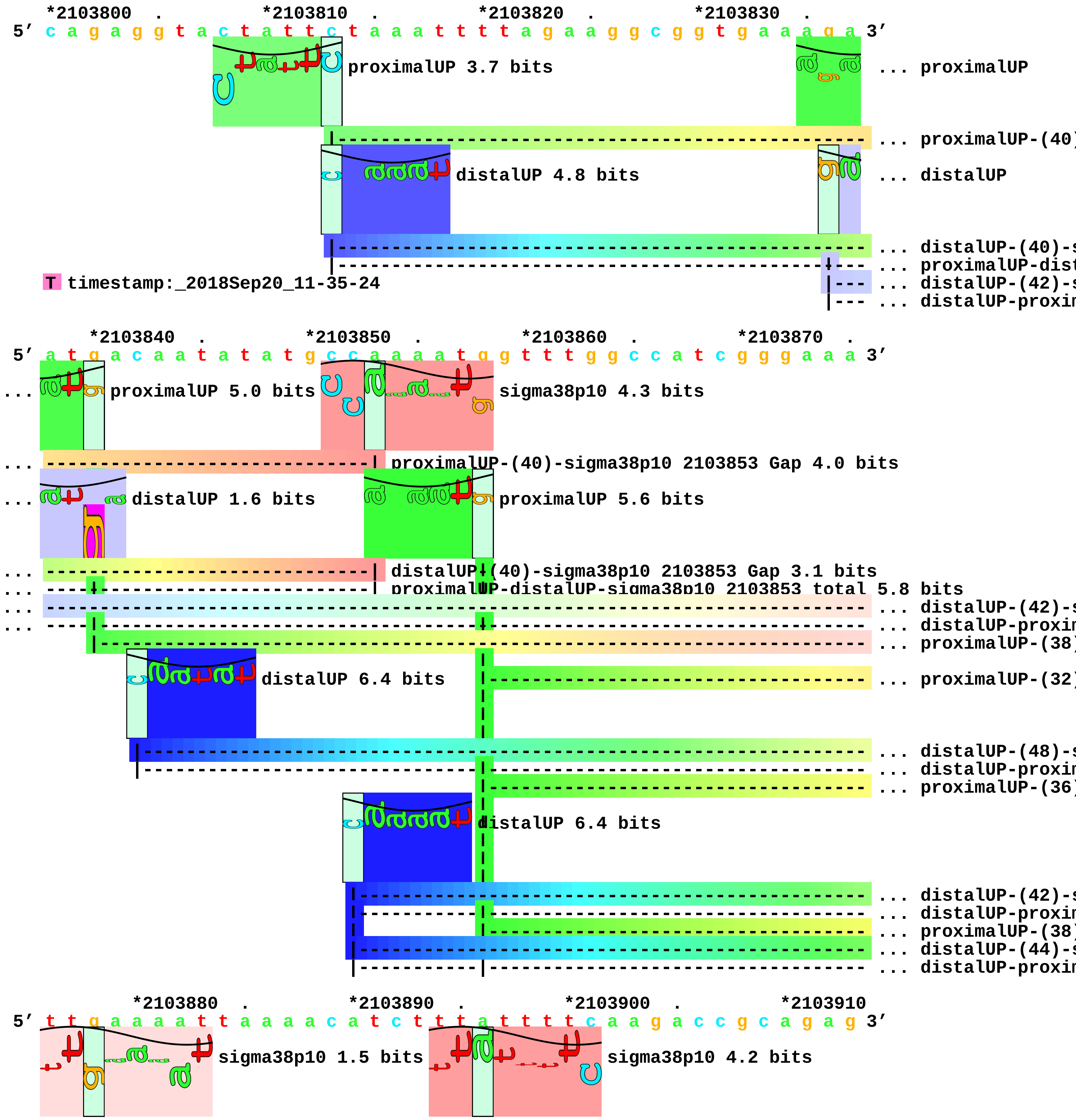

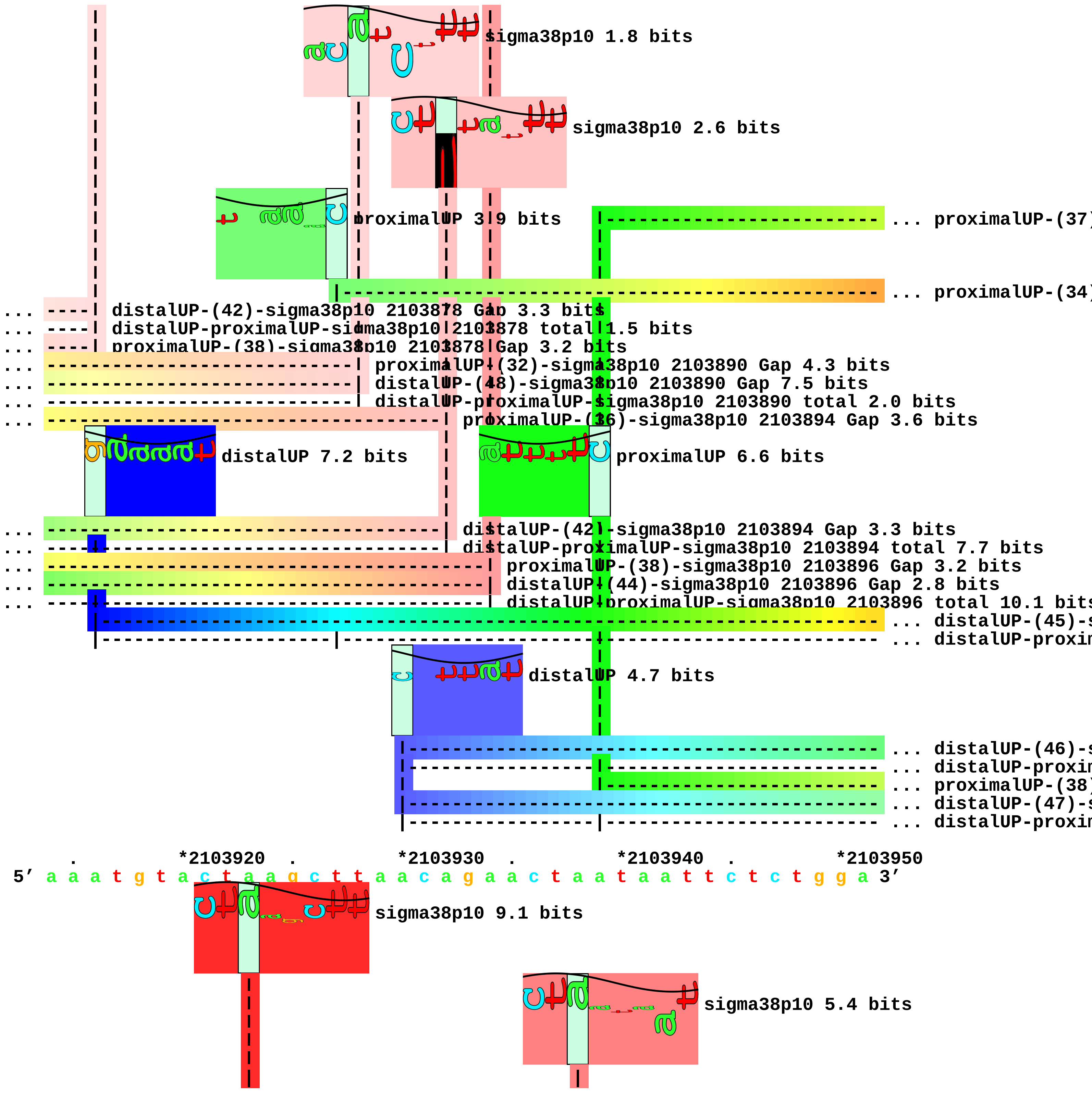
sigma38p10 1.8 bits
sigma38p10 2.6 bits
proximalUP 3.9 bits
... proximalUP-(37)
... proximalUP-(34)
... ---- distalUP-(42)-sigma38p10 2103878 Gap 3.3 bits
... ---- distalUP-proximalUP-sigma38p10 2103878 total 1.5 bits
... ---- proximalUP-(38)-sigma38p10 2103878 Gap 3.2 bits
... proximalUP-(32)-sigma38p10 2103890 Gap 4.3 bits
... distalUP-(48)-sigma38p10 2103890 Gap 7.5 bits
... distalUP-proximalUP-sigma38p10 2103890 total 2.0 bits
... proximalUP-(36)-sigma38p10 2103894 Gap 3.6 bits
distalUP 7.2 bits
proximalUP 6.6 bits
... distalUP-(42)-sigma38p10 2103894 Gap 3.3 bits
... distalUP-proximalUP-sigma38p10 2103894 total 7.7 bits
... proximalUP-(38)-sigma38p10 2103896 Gap 3.2 bits
... distalUP-(44)-sigma38p10 2103896 Gap 2.8 bits
... distalUP-proximalUP-sigma38p10 2103896 total 10.1 bits
... distalUP-(45)-s
... distalUP-proxin
distalUP 4.7 bits
... distalUP-(46)-s
... distalUP-proxin
... proximalUP-(38)
... distalUP-(47)-s
... distalUP-proxin
. *2103920 . *2103930 . *2103940 . *2103950
5' a a a t g t a c t a a g c t t a a c a g a a c t a a t a a t t c t c t g g a 3'
sigma38p10 9.1 bits
sigma38p10 5.4 bits

sigma38p10 1.5 bits

... ----------------------------------------| proximalUP-(37)-sigma38p10 2103938 Gap 4.5 bit
... ------------------| proximalUP-(34)-sigma38p10 2103923 Gap 2.5 bits

... ------------------| distalUP-(45)-sigma38p10 2103923 Gap 5.5 bits
... ------------------| distalUP-proximalUP-sigma38p10 2103923 total 12.2 bits

... ----------------------------------------| distalUP-(46)-sigma38p10 2103938 Gap 4.1 bits
... ----------------------------------------| distalUP-proximalUP-sigma38p10 2103938 total 8
... -----------------------------------------| proximalUP-(38)-sigma38p10 2103939 Gap 3.2 b
... -----------------------------------------| distalUP-(47)-sigma38p10 2103939 Gap 6.5 bit
... -----------------------------------------| distalUP-proximalUP-sigma38p10 2103939 tota

. *2103960 . *2103970 . *2103980 .
5' c t a c t c c a c c c c a t a g a g g a a t g t t a a c a a c a g a a a t g 3'

*2103990 . *2104000 . *2104010 . *2104020 .
5' t t t t c a t a a t c t g a a g c a a t g t c c a g t g c a t c t t t t c t 3'

*2104030 . *2104040 . *2104050 . *2104060 .
5' t g c t t t a a a t c c a g c a t c g c g t c t a g a g a a a t t t a a a t 3'

proximalUP 5.7 bits

|---------------------------------------------------- ... proximalUP-(40)

distalUP 4.0 bits

|----------------------------------------------------------------- ... distalUP-(48)-s
|---------------------------------------------------------------- ... distalUP-proxim
|---------------------------------------------------- ... proximalUP-(42)
|----------------------------------------------------------------- ... distalUP-(50)-s
|---------------|------------------------------------------------ ... distalUP-proxim

*2104070 . *2104080 . *2104090 . *2104100
5' c a t t c a a a a a a t a c a t t t t c a c t t t a t t t t c t g g g c c t 3'

sigma38p10 6.4 bits

|--------------- ... proximalUP-(28)

... -------------------------| proximalUP-(40)-sigma38p10 2104078 Gap 4.0 bits

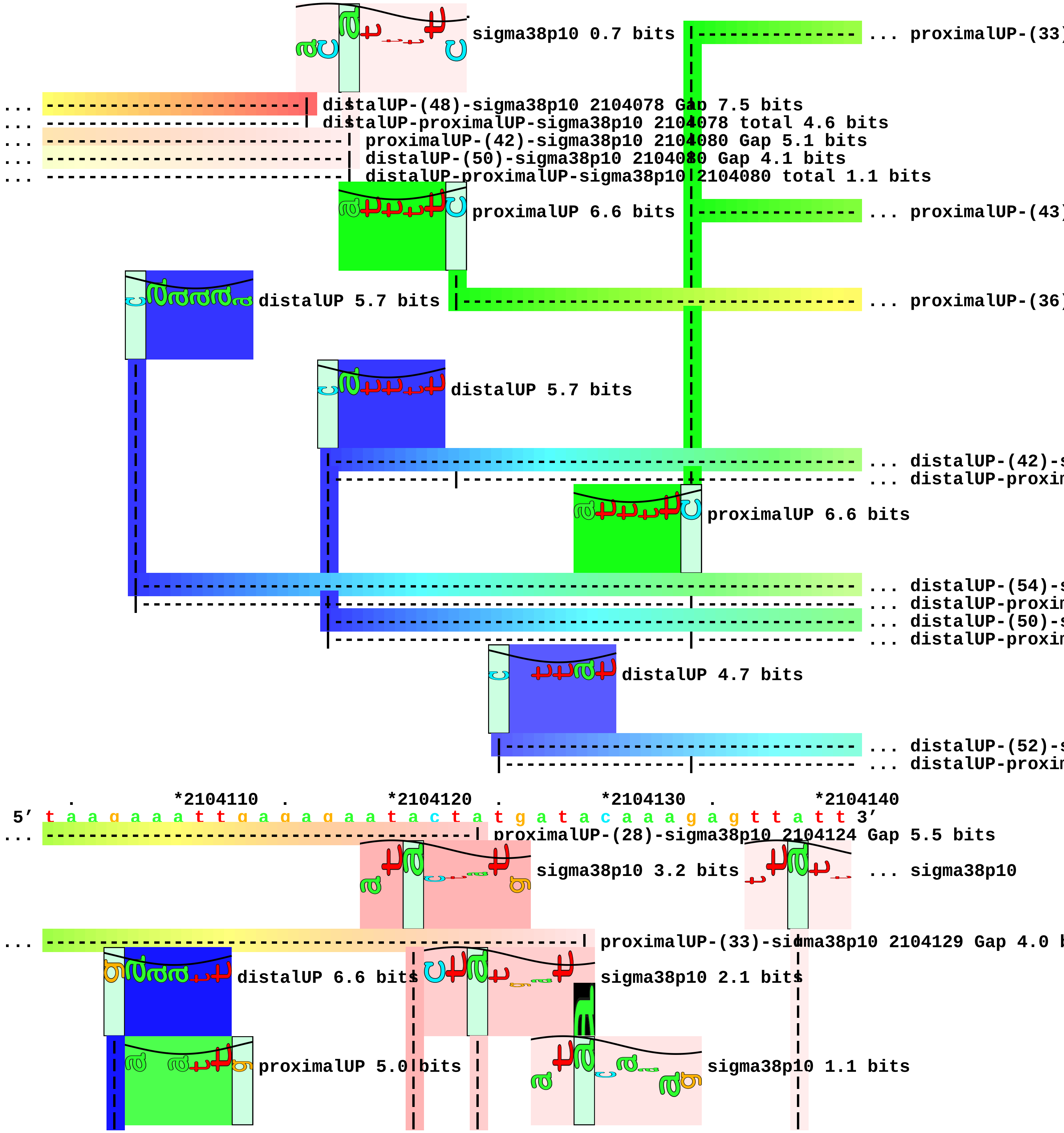

sigma38p10 0.7 bits
... proximalUP-(33)
... distalUP-(48)-sigma38p10 2104078 Gap 7.5 bits
... distalUP-proximalUP-sigma38p10 2104078 total 4.6 bits
... proximalUP-(42)-sigma38p10 2104080 Gap 5.1 bits
... distalUP-(50)-sigma38p10 2104080 Gap 4.1 bits
... distalUP-proximalUP-sigma38p10 2104080 total 1.1 bits
proximalUP 6.6 bits
... proximalUP-(43)
distalUP 5.7 bits
... proximalUP-(36)
distalUP 5.7 bits
... distalUP-(42)-s
... distalUP-proxin
proximalUP 6.6 bits
... distalUP-(54)-s
... distalUP-proxin
... distalUP-(50)-s
... distalUP-proxin
distalUP 4.7 bits
... distalUP-(52)-s
... distalUP-proxin
. *2104110 . *2104120 . *2104130 . *2104140
5' t a a g a a a t t g a g a g a a t a c t a t g a t a c a a a g a g t t a t t 3'
... proximalUP-(28)-sigma38p10 2104124 Gap 5.5 bits
sigma38p10 3.2 bits
... sigma38p10
... proximalUP-(33)-sigma38p10 2104129 Gap 4.0 b
distalUP 6.6 bits
sigma38p10 2.1 bits
proximalUP 5.0 bits
sigma38p10 1.1 bits

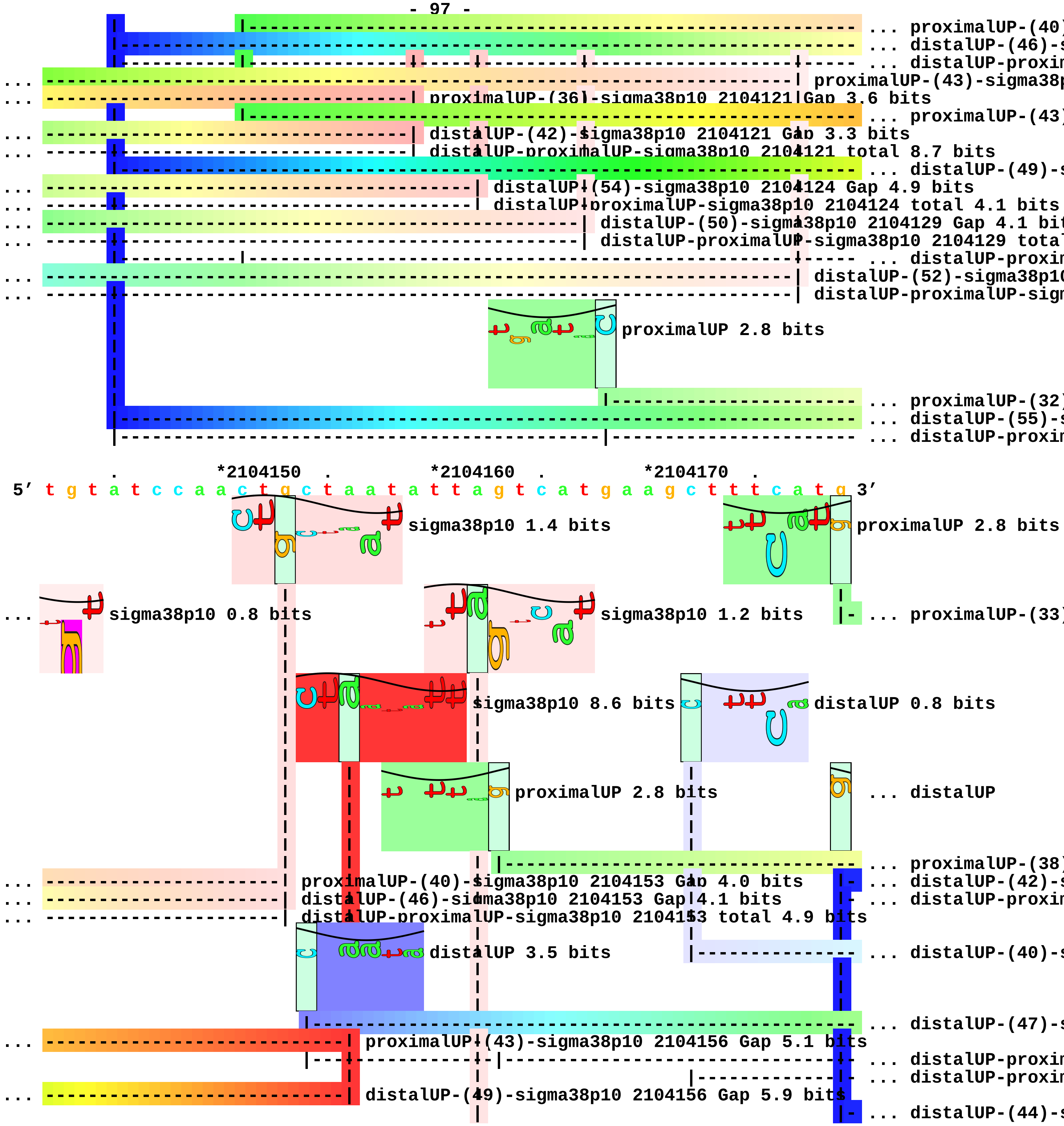
... proximalUP-(40)
... distalUP-(46)-s
... distalUP-proxim
... proximalUP-(43)-sigma38p
... proximalUP-(36)-sigma38p10 2104121 Gap 3.6 bits
... proximalUP-(43)
... distalUP-(42)-sigma38p10 2104121 Gap 3.3 bits
... distalUP-proximalUP-sigma38p10 2104121 total 8.7 bits
... distalUP-(49)-s
... distalUP-(54)-sigma38p10 2104124 Gap 4.9 bits
... distalUP-proximalUP-sigma38p10 2104124 total 4.1 bits
... distalUP-(50)-sigma38p10 2104129 Gap 4.1 bit
... distalUP-proximalUP-sigma38p10 2104129 total
... distalUP-proxim
... distalUP-(52)-sigma38p10
... distalUP-proximalUP-sigm
proximalUP 2.8 bits
... proximalUP-(32)
... distalUP-(55)-s
... distalUP-proxim
*2104150
*2104160
*2104170
5′ t g t a t c c a a c t g c t a a t a t t a g t c a t g a a g c t t t c a t g 3′
sigma38p10 1.4 bits
proximalUP 2.8 bits
sigma38p10 0.8 bits
sigma38p10 1.2 bits
... proximalUP-(33)
sigma38p10 8.6 bits
distalUP 0.8 bits
proximalUP 2.8 bits
... distalUP
... proximalUP-(38)
... proximalUP-(40)-sigma38p10 2104153 Gap 4.0 bits
... distalUP-(42)-s
... distalUP-(46)-sigma38p10 2104153 Gap 4.1 bits
... distalUP-proxim
... distalUP-proximalUP-sigma38p10 2104153 total 4.9 bits
distalUP 3.5 bits
... distalUP-(40)-s
... distalUP-(47)-s
... proximalUP-(43)-sigma38p10 2104156 Gap 5.1 bits
... distalUP-proxim
... distalUP-proxim
... distalUP-(49)-sigma38p10 2104156 Gap 5.9 bits
... distalUP-(44)-s

|- ... distalUP-proxin

... ----------------------------| distalUP-proximalUP-sigma38p10 2104156 total 9.2 bits

... ----------------------------------------| proximalUP-(32)-sigma38p10 2104162 Gap 4.3 bits
... ----------------------------------------| distalUP-(55)-sigma38p10 2104162 Gap 4.9 bits
... ----------------------------------------| distalUP-proximalUP-sigma38p10 2104162 total 1.3 bits

*2104180 . *2104190 . *2104200 . *2104210 .
5' a t a a a a a a t g a a a g a a a a g g t a t a a a t a a a a t a t g a a t 3'

sigma38p10 4.5 bits

... -------------------------------------------------------------| proximalUP-(33)-sigma38p10 210

proximalUP 5.6 bits

sigma38p10 3.6 bits

... distalUP 6.5 bits

proximalUP 0.4 bits

... ------------------------------------------| proximalUP-(38)-sigma38p10 2104201 Gap 3.2 bits
... ------------------------------------------------------------------------- ... distalUP-(42)-s
... ------------------------------------------------------------------------- ... distalUP-proxin
------------------------------------------------- ... proximalUP-(32)
... --------------------------------------------------------------| distalUP-(40)-sigma38p10 21042
... ------------------------------------------| distalUP-(47)-sigma38p10 2104201 Gap 6.5 bits
|---------------------------------------------------- ... proximalUP-(34)
... ------------------------------------------| distalUP-proximalUP-sigma38p10 2104201 total 1.2 bit
... --------------------------------------------------------------| distalUP-proximalUP-sigma38p10
|------------------------ ... proximalUP-(33)
... ------------------------------------------------------------------------- ... distalUP-(44)-s
... ----------------|-------------------------------------------------------- ... distalUP-proxin

distalUP 4.9 bits

|------------------------------------ ... distalUP-(40)-s
|-------------|------------------------ ... distalUP-proxin

*2104220 . *2104230 . *2104240 .
5' a a a a t a t t t t c a c a g a t a t g t a a t t t c g a g a c 3'

sigma38p10 0.4 bits

sigma38p10 2.6 bits

sigma38p10 3.6 bits

```
          |  |                                          |
          |  |                                          |
... ------| distalUP-(42)-sigma38p10 2104221 Gap 3.3 bits
... ------| distalUP-proximalUP-sigma38p10 2104221 total 4.9 bits
... ------| proximalUP-(32)-sigma38p10 2104221|Gap 4.3 bits
             |                                          |
             |                                          |
... ----------| proximalUP-(34)-sigma38p10 2104223 Gap 2.5 bits
             |                                          |
             |                                          |
... ------------------------------------------| proximalUP-(33)-sigma38p10 2104239 Gap 4.0 bits
... ----------| distalUP-(44)-sigma38p10 2104223 Gap 2.8 bits
... ----------| distalUP-proximalUP-sigma38p10|2104223 total 10.4 bits
                                                        |
... ------------------------------------------| distalUP-(40)-sigma38p10 2104239 Gap 3.1 bits
... ------------------------------------------| distalUP-proximalUP-sigma38p10 2104239 total 0.9 bit[illegible]
```

piece 29, NC_000913, wbbH, config: linear, direction: +, begin: 2104500, end: 2105649

*2104500 . *2104510 . *2104520 . *2104530 .
5' t a g g g c t g a c c a a a t a c a t g g a t a g a t a a t a t g c c a t c 3'

proximalUP 3.7 bits
... proximalUP-(40)
distalUP 5.1 bits
... distalUP-(46)-s
... distalUP-proxim

T timestamp:_2018Sep20_11-35-24

*2104540 . *2104550 . *2104560 . *2104570 .
5' c c c c c a g g c a g t c c a c g t a a a a a g a g c a t a t a c a a g g a 3'

sigma38p10 1.3 bits
... proximalUP-(40)-sigma38p10 2104556 Gap 4.0 bits
proximalUP 4.9 bits
... distalUP-(46)-sigma38p10 2104556 Gap 4.1 bits
... distalUP-proximalUP-sigma38p10 2104556 total 2.0 bits
distalUP 5.0 bits
... proximalUP-(27)
... distalUP-(43)-s
... distalUP-proxim
... proximalUP-(35)
distalUP 5.0 bits
... distalUP-(41)-s
... distalUP-proxim
... proximalUP-(39)
... distalUP-(55)-s
... distalUP-proxim
... proximalUP-(41)
... distalUP-(47)-s
... distalUP-proxim

*2104580 . *2104590 . *2104600 . *2104610
5' g a a t a g a a c a c c t a c a g c t g t a a t a a g a t a a a c a t a a t 3'

sigma38p10 0.2 bits

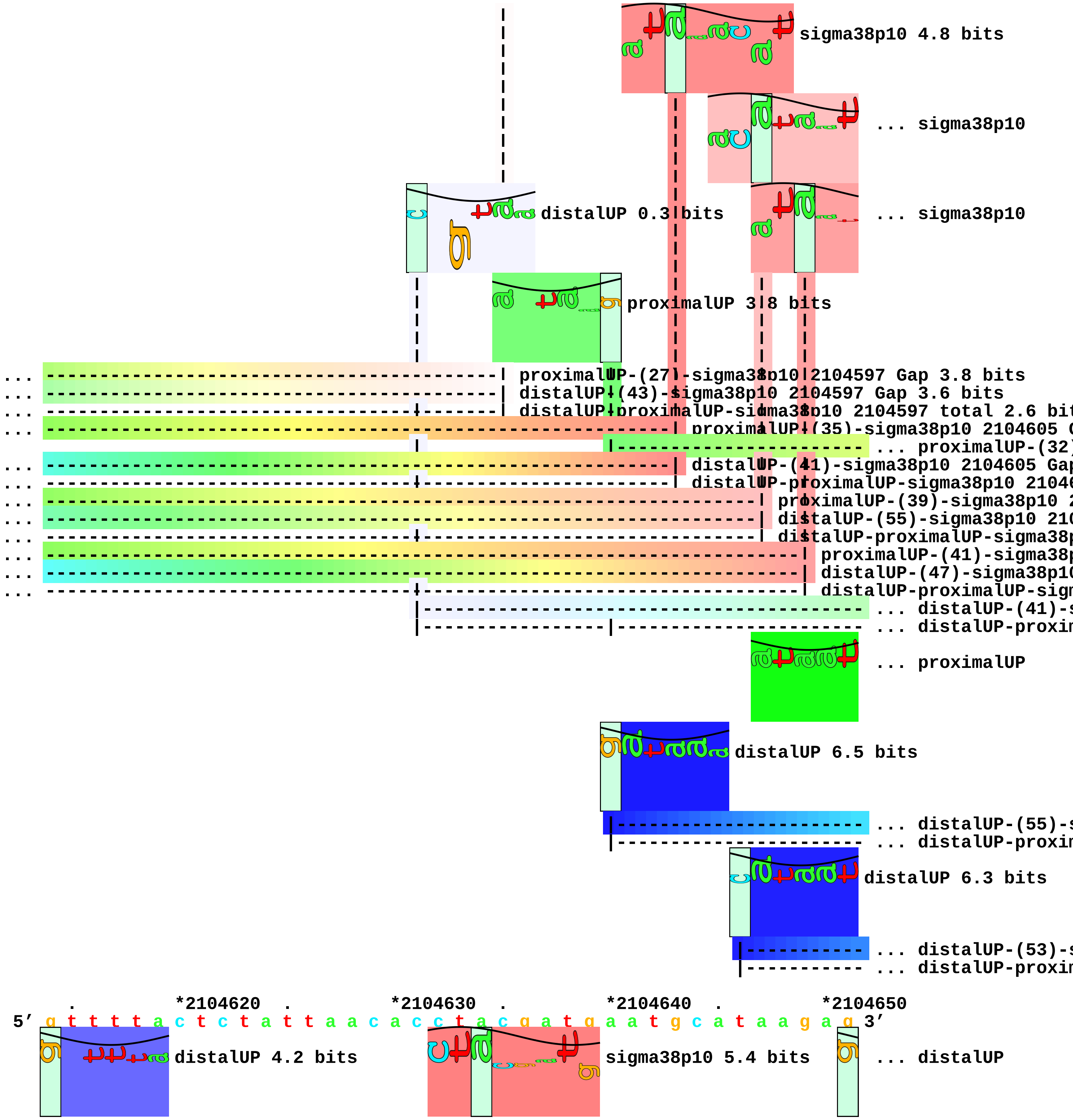
sigma38p10 4.8 bits
... sigma38p10
distalUP 0.3 bits
... sigma38p10
proximalUP 3.8 bits
... proximalUP-(27)-sigma38p10 2104597 Gap 3.8 bits
... distalUP-(43)-sigma38p10 2104597 Gap 3.6 bits
... distalUP-proximalUP-sigma38p10 2104597 total 2.6 bit
... proximalUP-(35)-sigma38p10 2104605 G
... proximalUP-(32)
... distalUP-(41)-sigma38p10 2104605 Gap
... distalUP-proximalUP-sigma38p10 21046
... proximalUP-(39)-sigma38p10 2
... distalUP-(55)-sigma38p10 210
... distalUP-proximalUP-sigma38p
... proximalUP-(41)-sigma38p
... distalUP-(47)-sigma38p10
... distalUP-proximalUP-sigm
... distalUP-(41)-s
... distalUP-proxim
... proximalUP
distalUP 6.5 bits
... distalUP-(55)-s
... distalUP-proxim
distalUP 6.3 bits
... distalUP-(53)-s
... distalUP-proxim
. *2104620 . *2104630 . *2104640 . *2104650
5′ g t t t t a c t c t a t t a a c a c c t a c g a t g a a t g c a t a a g a g 3′
distalUP 4.2 bits
sigma38p10 5.4 bits
... distalUP

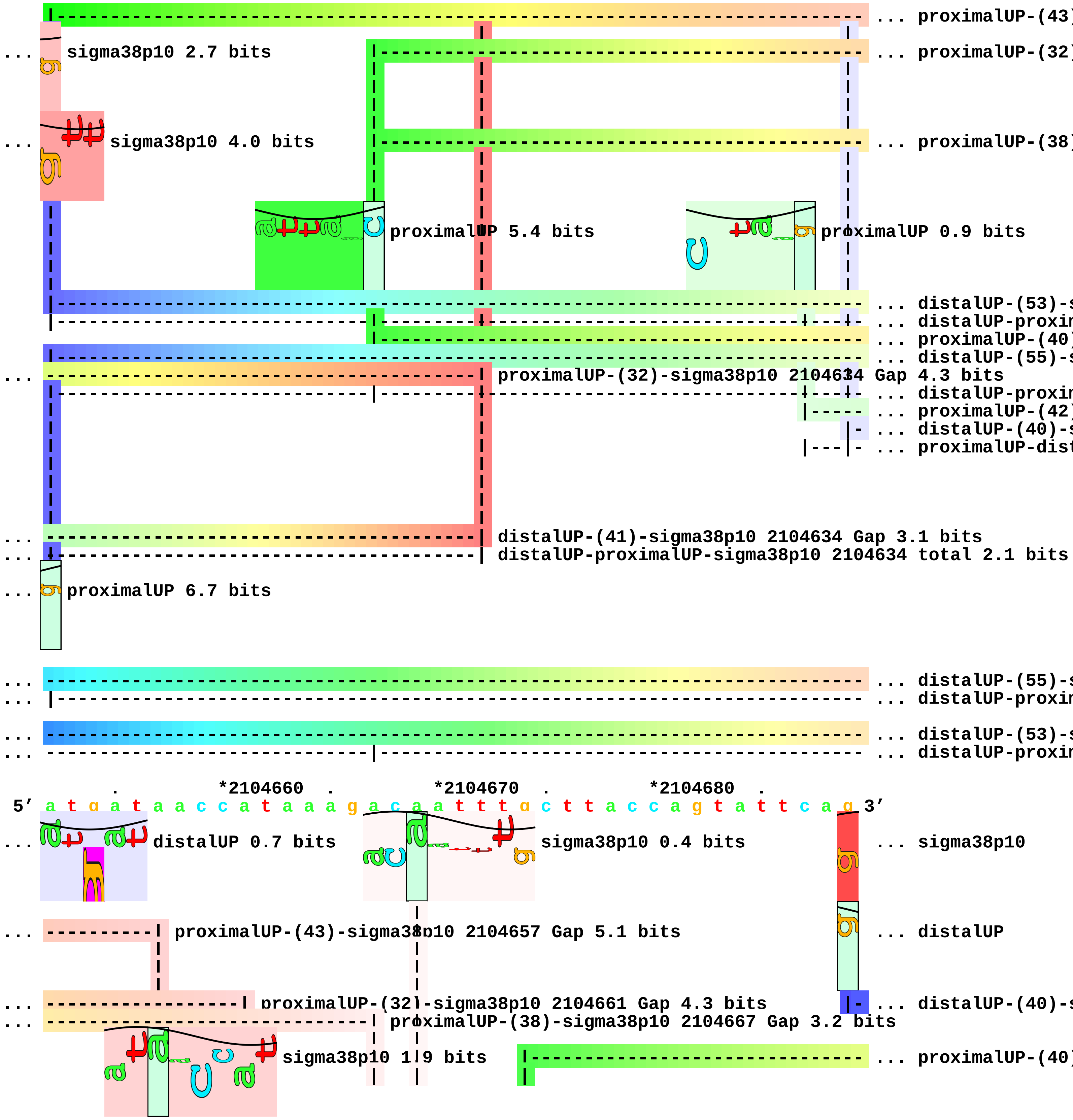
... proximalUP-(43)
... sigma38p10 2.7 bits
... proximalUP-(32)
... sigma38p10 4.0 bits
... proximalUP-(38)
proximalUP 5.4 bits
proximalUP 0.9 bits
... distalUP-(53)-s
... distalUP-proxin
... proximalUP-(40)
... distalUP-(55)-s
... proximalUP-(32)-sigma38p10 2104634 Gap 4.3 bits
... distalUP-proxin
... proximalUP-(42)
... distalUP-(40)-s
... proximalUP-dist
... distalUP-(41)-sigma38p10 2104634 Gap 3.1 bits
... distalUP-proximalUP-sigma38p10 2104634 total 2.1 bits
... proximalUP 6.7 bits
... distalUP-(55)-s
... distalUP-proxin
... distalUP-(53)-s
... distalUP-proxin
. *2104660 . *2104670 . *2104680 .
5' a t g a t a a c c a t a a a g a c a a t t t g c t t a c c a g t a t t c a g 3'
... distalUP 0.7 bits
sigma38p10 0.4 bits
... sigma38p10
... proximalUP-(43)-sigma38p10 2104657 Gap 5.1 bits
... distalUP
... proximalUP-(32)-sigma38p10 2104661 Gap 4.3 bits
... distalUP-(40)-s
... proximalUP-(38)-sigma38p10 2104667 Gap 3.2 bits
sigma38p10 1.9 bits
... proximalUP-(40)

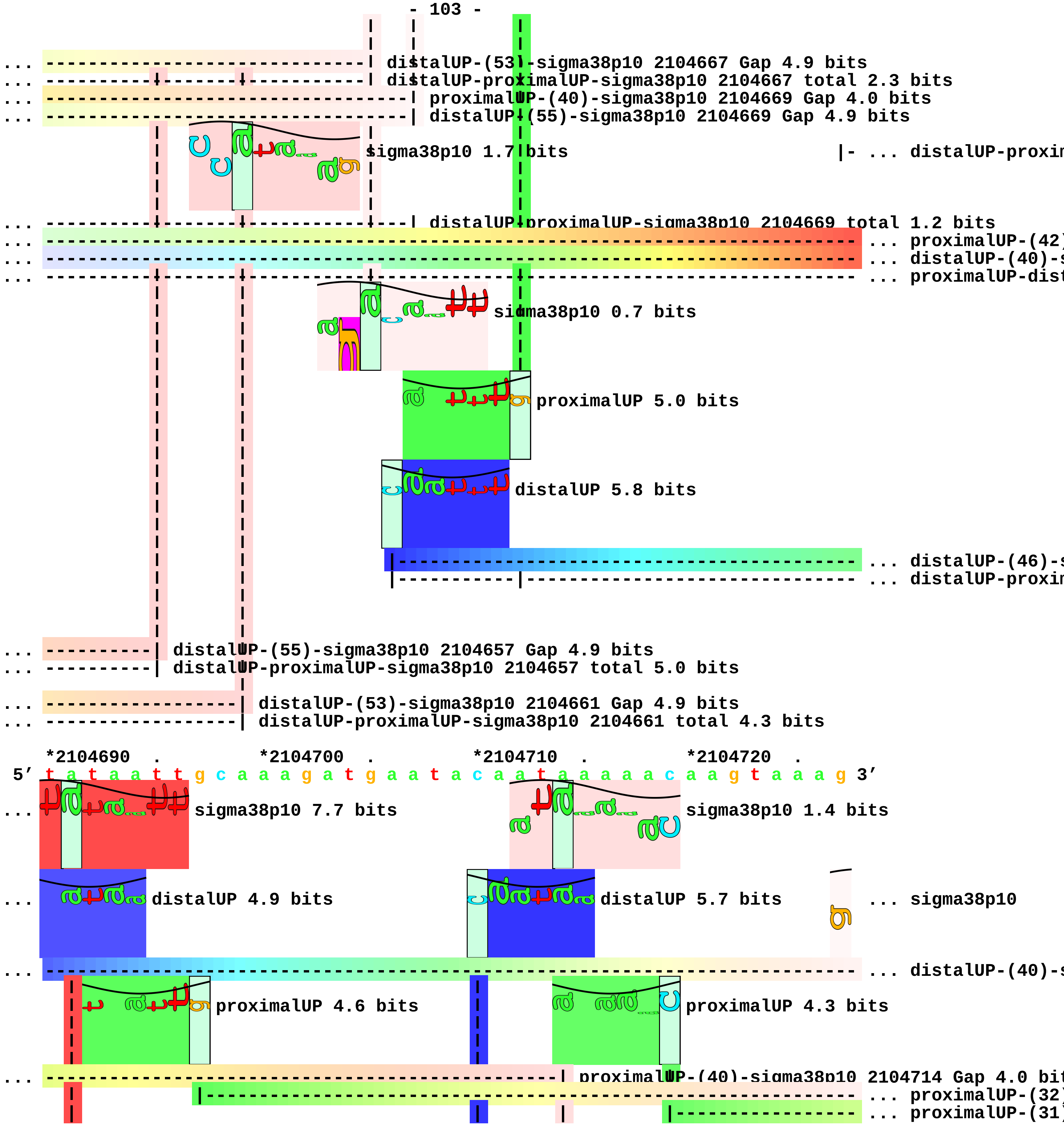
... distalUP-(53)-sigma38p10 2104667 Gap 4.9 bits
... distalUP-proximalUP-sigma38p10 2104667 total 2.3 bits
... proximalUP-(40)-sigma38p10 2104669 Gap 4.0 bits
... distalUP-(55)-sigma38p10 2104669 Gap 4.9 bits
sigma38p10 1.7 bits
|- ... distalUP-proxim
... distalUP-proximalUP-sigma38p10 2104669 total 1.2 bits
... proximalUP-(42)
... distalUP-(40)-s
... proximalUP-dist
sigma38p10 0.7 bits
proximalUP 5.0 bits
distalUP 5.8 bits
... distalUP-(46)-s
... distalUP-proxim
... distalUP-(55)-sigma38p10 2104657 Gap 4.9 bits
... distalUP-proximalUP-sigma38p10 2104657 total 5.0 bits
... distalUP-(53)-sigma38p10 2104661 Gap 4.9 bits
... distalUP-proximalUP-sigma38p10 2104661 total 4.3 bits
*2104690 . *2104700 . *2104710 . *2104720 .
5′ t a t a a t t g c a a a g a t g a a t a c a a t a a a a a c a a g t a a a g 3′
... sigma38p10 7.7 bits
sigma38p10 1.4 bits
... distalUP 4.9 bits
distalUP 5.7 bits
... sigma38p10
... distalUP-(40)-s
proximalUP 4.6 bits
proximalUP 4.3 bits
... proximalUP-(40)-sigma38p10 2104714 Gap 4.0 bit
... proximalUP-(32)
... proximalUP-(31)

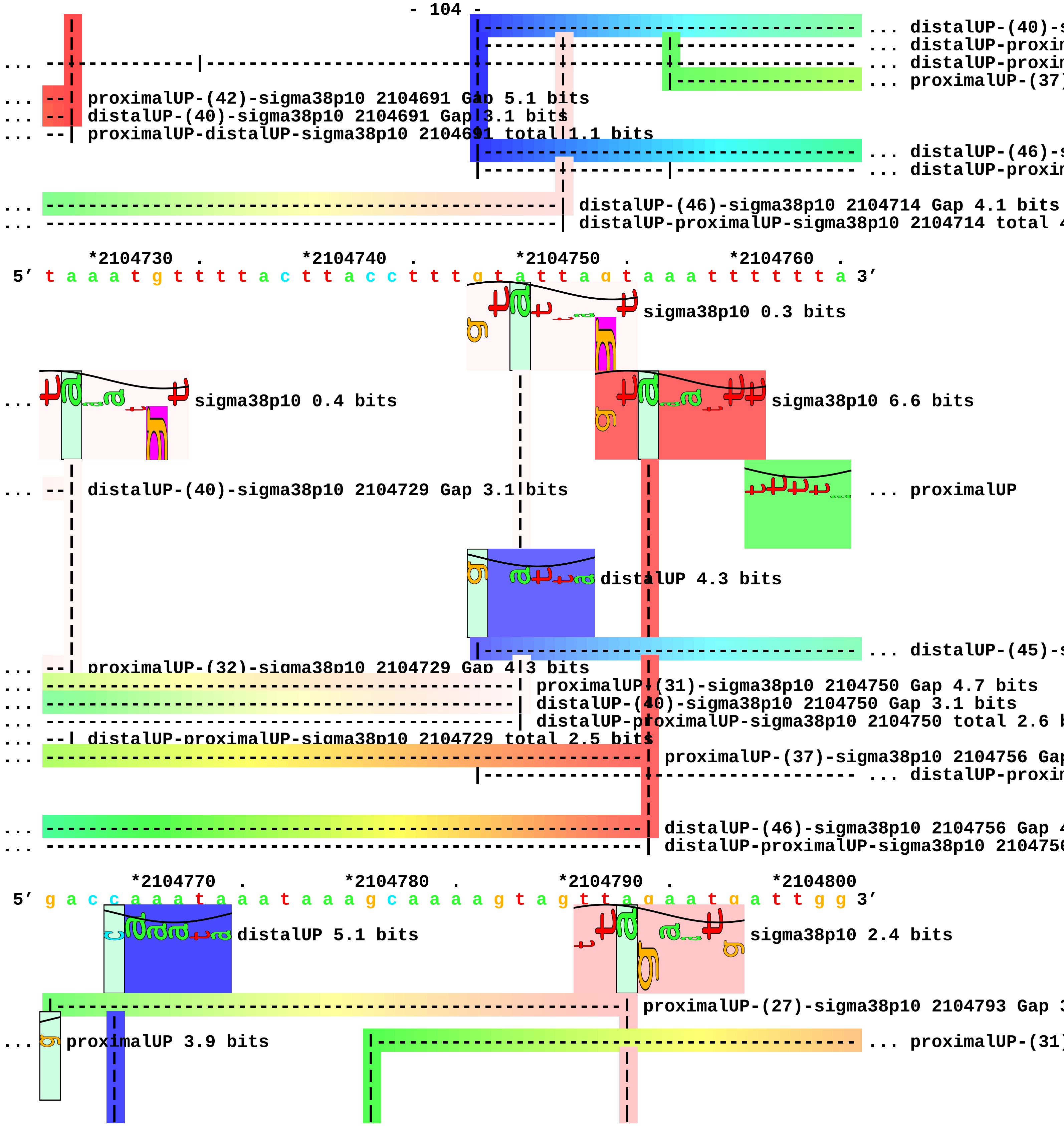
... distalUP-(40)-s
... distalUP-proxim
... distalUP-proxim
... proximalUP-(37)
... --| proximalUP-(42)-sigma38p10 2104691 Gap 5.1 bits
... --| distalUP-(40)-sigma38p10 2104691 Gap 3.1 bits
... --| proximalUP-distalUP-sigma38p10 2104691 total 1.1 bits
... distalUP-(46)-s
... distalUP-proxim
... distalUP-(46)-sigma38p10 2104714 Gap 4.1 bits
... distalUP-proximalUP-sigma38p10 2104714 total 4
*2104730 . *2104740 . *2104750 . *2104760 .
5' t a a a t g t t t a c t t a c c t t t g t a t t a g t a a a t t t t t t a 3'
sigma38p10 0.3 bits
... sigma38p10 0.4 bits
sigma38p10 6.6 bits
... --| distalUP-(40)-sigma38p10 2104729 Gap 3.1 bits
... proximalUP
distalUP 4.3 bits
... distalUP-(45)-s
... --| proximalUP-(32)-sigma38p10 2104729 Gap 4.3 bits
... proximalUP-(31)-sigma38p10 2104750 Gap 4.7 bits
... distalUP-(40)-sigma38p10 2104750 Gap 3.1 bits
... distalUP-proximalUP-sigma38p10 2104750 total 2.6 b
... --| distalUP-proximalUP-sigma38p10 2104729 total 2.5 bits
... proximalUP-(37)-sigma38p10 2104756 Gap
... distalUP-proxim
... distalUP-(46)-sigma38p10 2104756 Gap 4
... distalUP-proximalUP-sigma38p10 2104756
*2104770 . *2104780 . *2104790 . *2104800
5' g a c c a a a t a a a t a a a g c a a a a g t a g t t a g a a t g a t t g g 3'
distalUP 5.1 bits
sigma38p10 2.4 bits
proximalUP-(27)-sigma38p10 2104793 Gap 3
... proximalUP 3.9 bits
... proximalUP-(31)

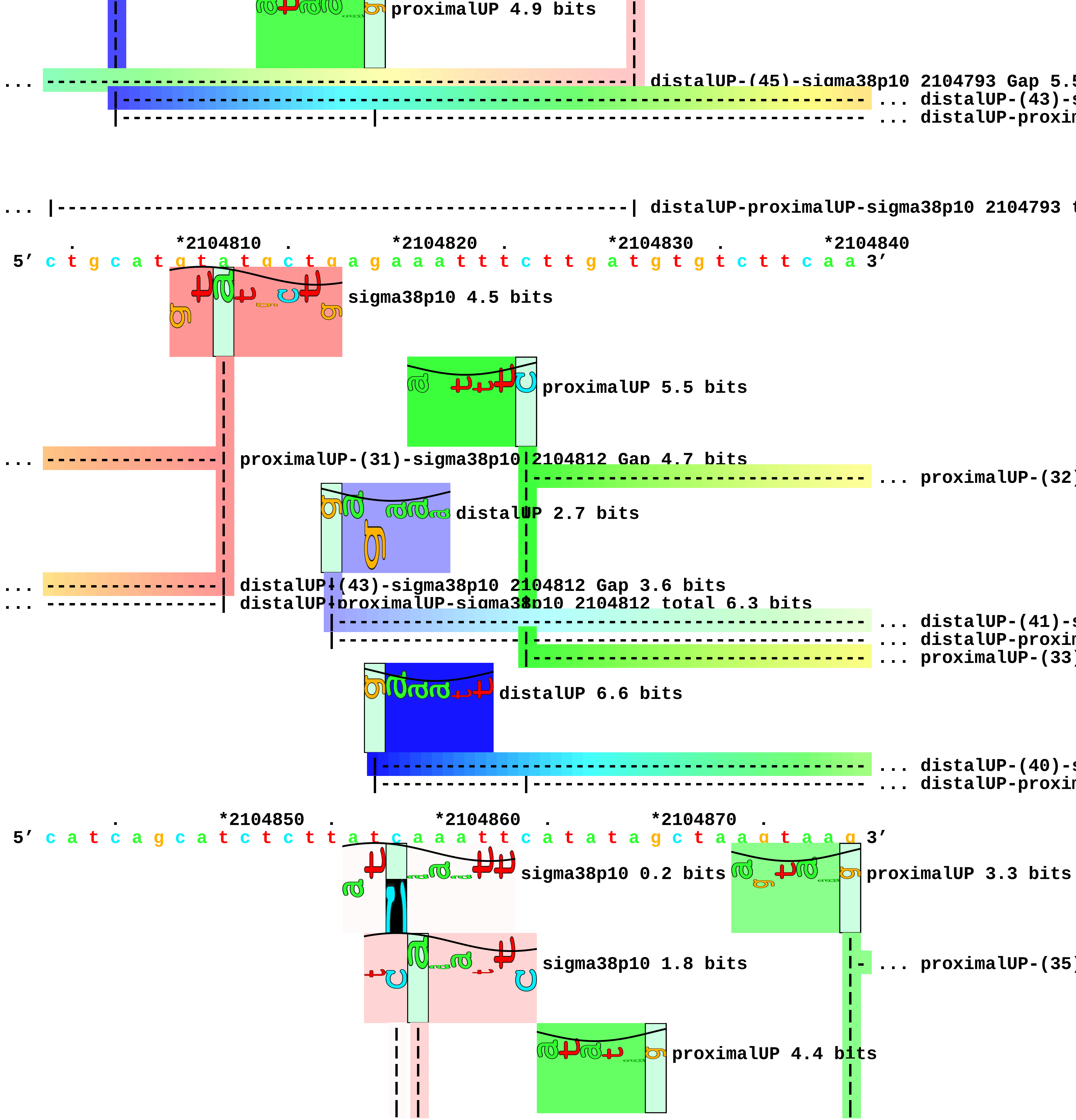
proximalUP 4.9 bits
... distalUP-(45)-sigma38p10 2104793 Gap 5.5
... distalUP-(43)-s
... distalUP-proxin
... distalUP-proximalUP-sigma38p10 2104793 t
*2104810
*2104820
*2104830
*2104840
5′ c t g c a t g t a t g c t g a g a a a t t t c t t g a t g t g t c t t c a a 3′
sigma38p10 4.5 bits
proximalUP 5.5 bits
... proximalUP-(31)-sigma38p10 2104812 Gap 4.7 bits
... proximalUP-(32)
distalUP 2.7 bits
... distalUP-(43)-sigma38p10 2104812 Gap 3.6 bits
... distalUP-proximalUP-sigma38p10 2104812 total 6.3 bits
... distalUP-(41)-s
... distalUP-proxin
... proximalUP-(33)
distalUP 6.6 bits
... distalUP-(40)-s
... distalUP-proxin
*2104850
*2104860
*2104870
5′ c a t c a g c a t c t c t t a t c a a a t t c a t a t a g c t a a g t a a g 3′
sigma38p10 0.2 bits
proximalUP 3.3 bits
sigma38p10 1.8 bits
... proximalUP-(35)
proximalUP 4.4 bits

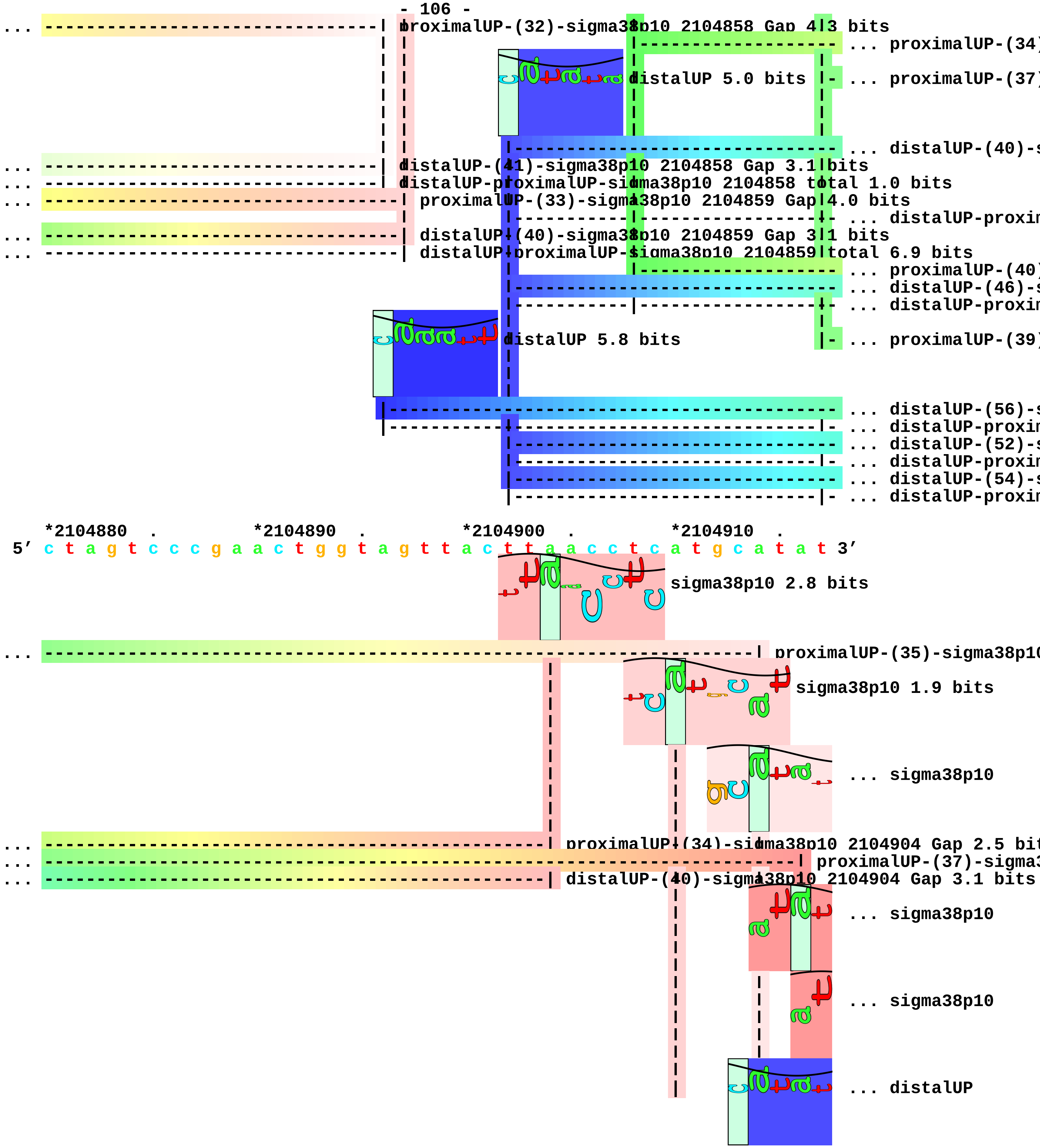
proximalUP-(32)-sigma38p10 2104858 Gap 4.3 bits
... proximalUP-(34)
distalUP 5.0 bits
... proximalUP-(37)
... distalUP-(40)-s
distalUP-(41)-sigma38p10 2104858 Gap 3.1 bits
distalUP-proximalUP-sigma38p10 2104858 total 1.0 bits
proximalUP-(33)-sigma38p10 2104859 Gap 4.0 bits
... distalUP-proxim
distalUP-(40)-sigma38p10 2104859 Gap 3.1 bits
distalUP-proximalUP-sigma38p10 2104859 total 6.9 bits
... proximalUP-(40)
... distalUP-(46)-s
... distalUP-proxim
distalUP 5.8 bits
... proximalUP-(39)
... distalUP-(56)-s
... distalUP-proxim
... distalUP-(52)-s
... distalUP-proxim
... distalUP-(54)-s
... distalUP-proxim
*2104880 . *2104890 . *2104900 . *2104910 .
5′ c t a g t c c c g a a c t g g t a g t t a c t t a a c c t c a t g c a t a t 3′
sigma38p10 2.8 bits
proximalUP-(35)-sigma38p10
sigma38p10 1.9 bits
... sigma38p10
proximalUP-(34)-sigma38p10 2104904 Gap 2.5 bit
proximalUP-(37)-sigma3
distalUP-(40)-sigma38p10 2104904 Gap 3.1 bits
... sigma38p10
... sigma38p10
... distalUP

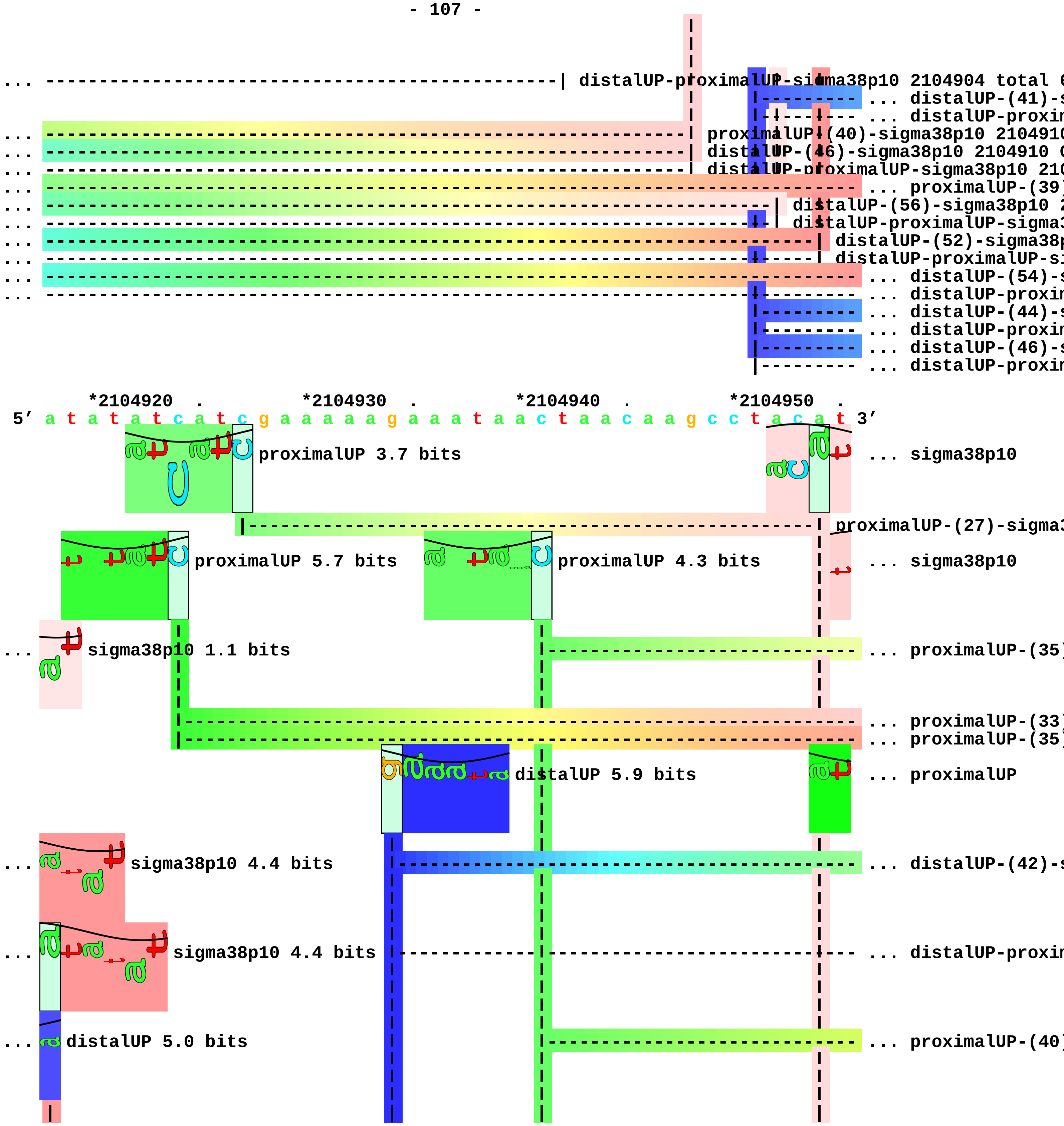
distalUP-proximalUP-sigma38p10 2104904 total
... distalUP-(41)-
... distalUP-proxi
proximalUP-(40)-sigma38p10 210491
distalUP-(46)-sigma38p10 2104910
distalUP-proximalUP-sigma38p10 210
... proximalUP-(39
distalUP-(56)-sigma38p10
distalUP-proximalUP-sigma
distalUP-(52)-sigma38
distalUP-proximalUP-s
... distalUP-(54)-
... distalUP-proxi
... distalUP-(44)-
... distalUP-proxi
... distalUP-(46)-
... distalUP-proxi
*2104920 . *2104930 . *2104940 . *2104950 .
5′ a t a t a t c a t c g a a a a a g a a a t a a c t a a c a a g c c t a c a t 3′
proximalUP 3.7 bits
... sigma38p10
proximalUP-(27)-sigma
proximalUP 5.7 bits
proximalUP 4.3 bits
... sigma38p10
sigma38p10 1.1 bits
... proximalUP-(35
... proximalUP-(33
... proximalUP-(35
distalUP 5.9 bits
... proximalUP
sigma38p10 4.4 bits
... distalUP-(42)-
sigma38p10 4.4 bits
... distalUP-proxi
distalUP 5.0 bits
... proximalUP-(40

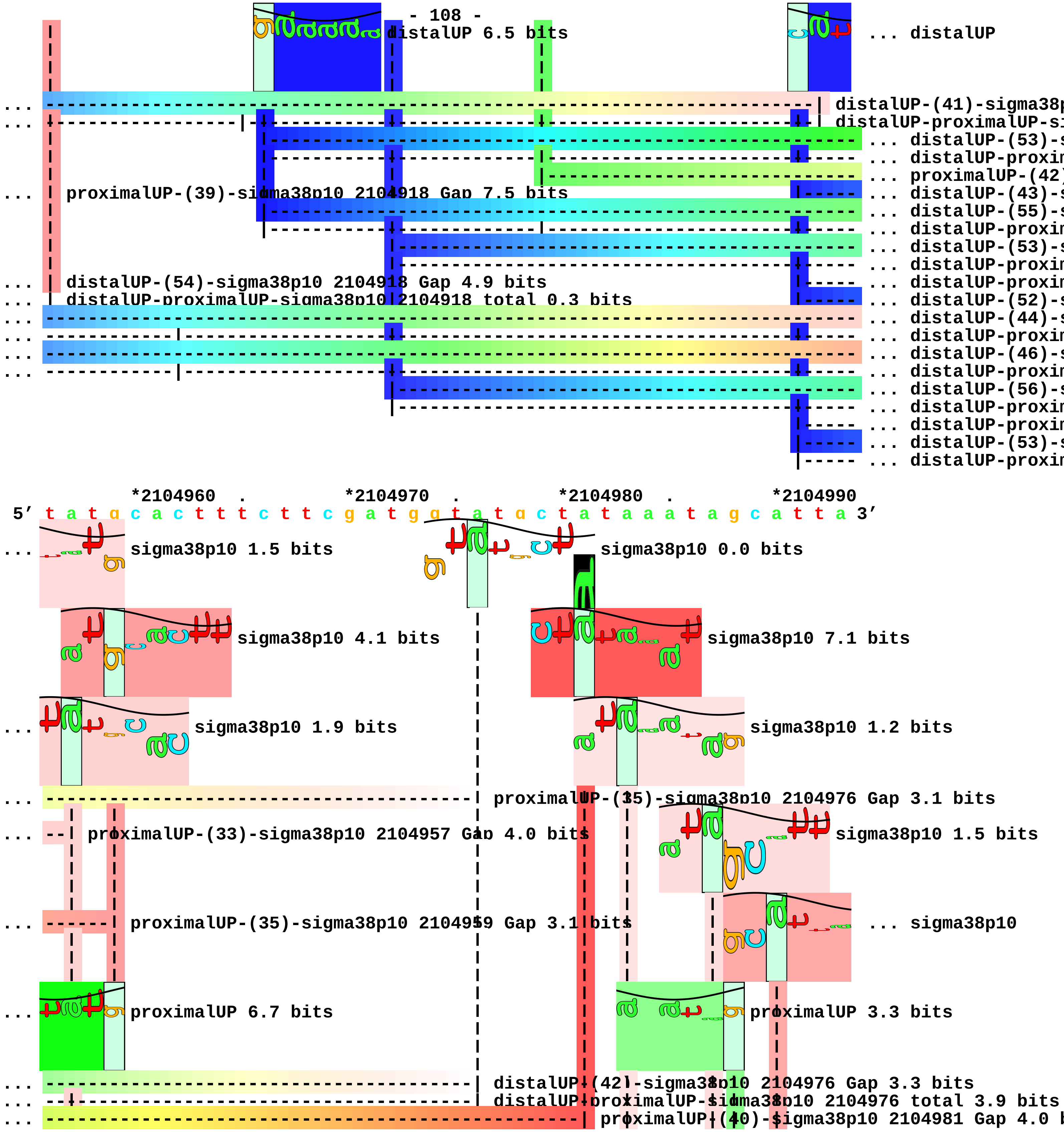
distalUP 6.5 bits
... distalUP
... distalUP-(41)-sigma38p
... distalUP-proximalUP-si
... distalUP-(53)-s
... distalUP-proxim
... proximalUP-(42)
... proximalUP-(39)-sigma38p10 2104918 Gap 7.5 bits
... distalUP-(43)-s
... distalUP-(55)-s
... distalUP-proxim
... distalUP-(53)-s
... distalUP-proxim
... distalUP-(54)-sigma38p10 2104918 Gap 4.9 bits
... distalUP-proxim
... distalUP-proximalUP-sigma38p10 2104918 total 0.3 bits
... distalUP-(52)-s
... distalUP-(44)-s
... distalUP-proxim
... distalUP-(46)-s
... distalUP-proxim
... distalUP-(56)-s
... distalUP-proxim
... distalUP-proxim
... distalUP-(53)-s
... distalUP-proxim
*2104960 . *2104970 . *2104980 . *2104990
5' t a t g c a c t t t c t t c g a t g g t a t g c t a t a a a t a g c a t t a 3'
... sigma38p10 1.5 bits
sigma38p10 0.0 bits
sigma38p10 4.1 bits
sigma38p10 7.1 bits
... sigma38p10 1.9 bits
sigma38p10 1.2 bits
... proximalUP-(35)-sigma38p10 2104976 Gap 3.1 bits
... proximalUP-(33)-sigma38p10 2104957 Gap 4.0 bits
sigma38p10 1.5 bits
... proximalUP-(35)-sigma38p10 2104959 Gap 3.1 bits
... sigma38p10
... proximalUP 6.7 bits
proximalUP 3.3 bits
... distalUP-(42)-sigma38p10 2104976 Gap 3.3 bits
... distalUP-proximalUP-sigma38p10 2104976 total 3.9 bits
... proximalUP-(40)-sigma38p10 2104981 Gap 4.0 b

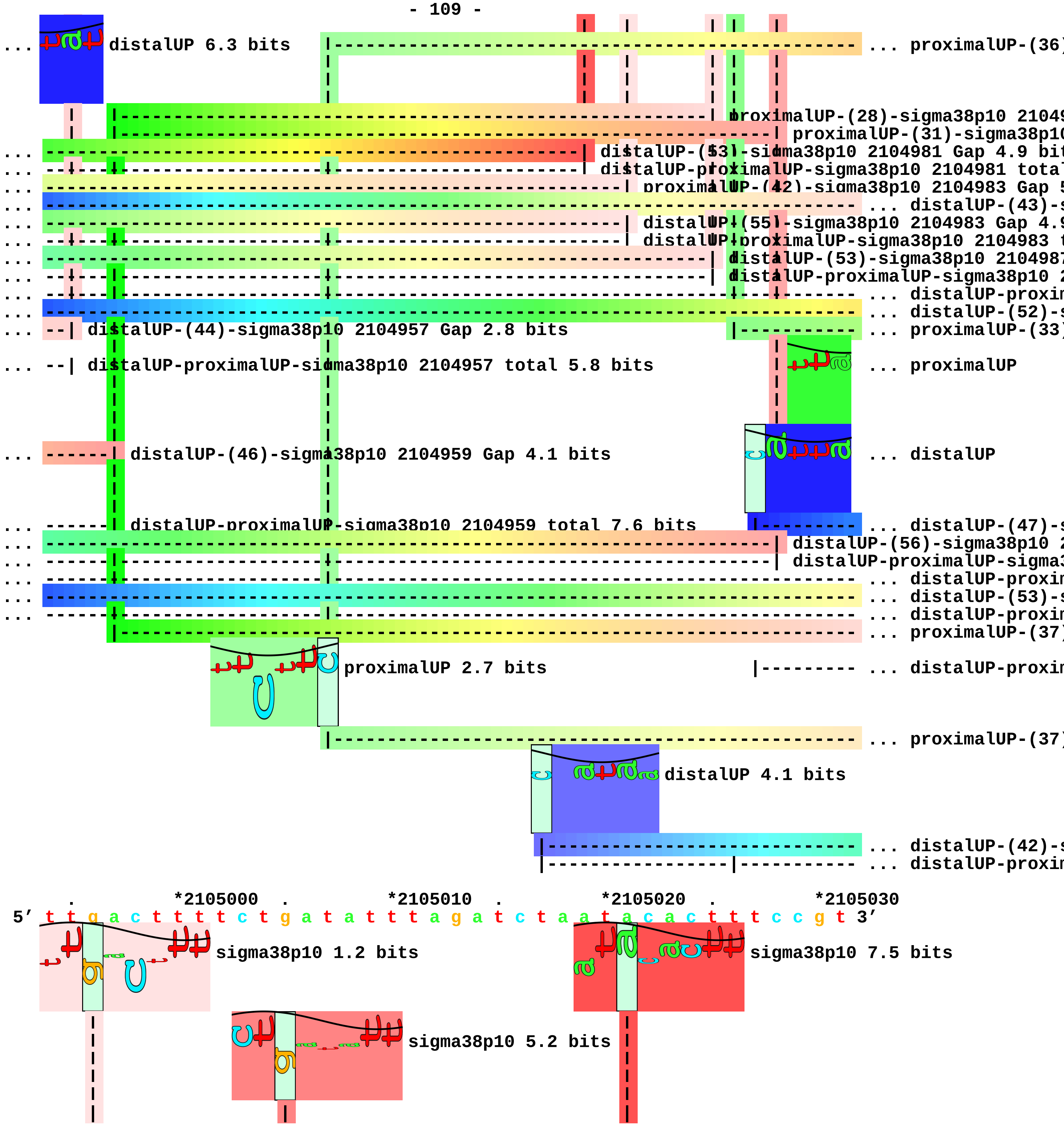
distalUP 6.3 bits
... proximalUP-(36)
proximalUP-(28)-sigma38p10 21049
proximalUP-(31)-sigma38p10
... distalUP-(53)-sigma38p10 2104981 Gap 4.9 bit
... distalUP-proximalUP-sigma38p10 2104981 tota
... proximalUP-(42)-sigma38p10 2104983 Gap 5
... distalUP-(43)-s
... distalUP-(55)-sigma38p10 2104983 Gap 4.9
... distalUP-proximalUP-sigma38p10 2104983 t
... distalUP-(53)-sigma38p10 2104987
... distalUP-proximalUP-sigma38p10 2
... distalUP-proxin
... distalUP-(52)-s
... distalUP-(44)-sigma38p10 2104957 Gap 2.8 bits
... proximalUP-(33)
... distalUP-proximalUP-sigma38p10 2104957 total 5.8 bits
... proximalUP
... distalUP-(46)-sigma38p10 2104959 Gap 4.1 bits
... distalUP
... distalUP-proximalUP-sigma38p10 2104959 total 7.6 bits
... distalUP-(47)-s
... distalUP-(56)-sigma38p10 2
... distalUP-proximalUP-sigma3
... distalUP-proxin
... distalUP-(53)-s
... distalUP-proxin
... proximalUP-(37)
proximalUP 2.7 bits
... distalUP-proxin
... proximalUP-(37)
distalUP 4.1 bits
... distalUP-(42)-s
... distalUP-proxin
*2105000 *2105010 *2105020 *2105030
5' t t g a c t t t t c t g a t a t t t a g a t c t a a t a c a c t t t c c g t 3'
sigma38p10 1.2 bits
sigma38p10 7.5 bits
sigma38p10 5.2 bits

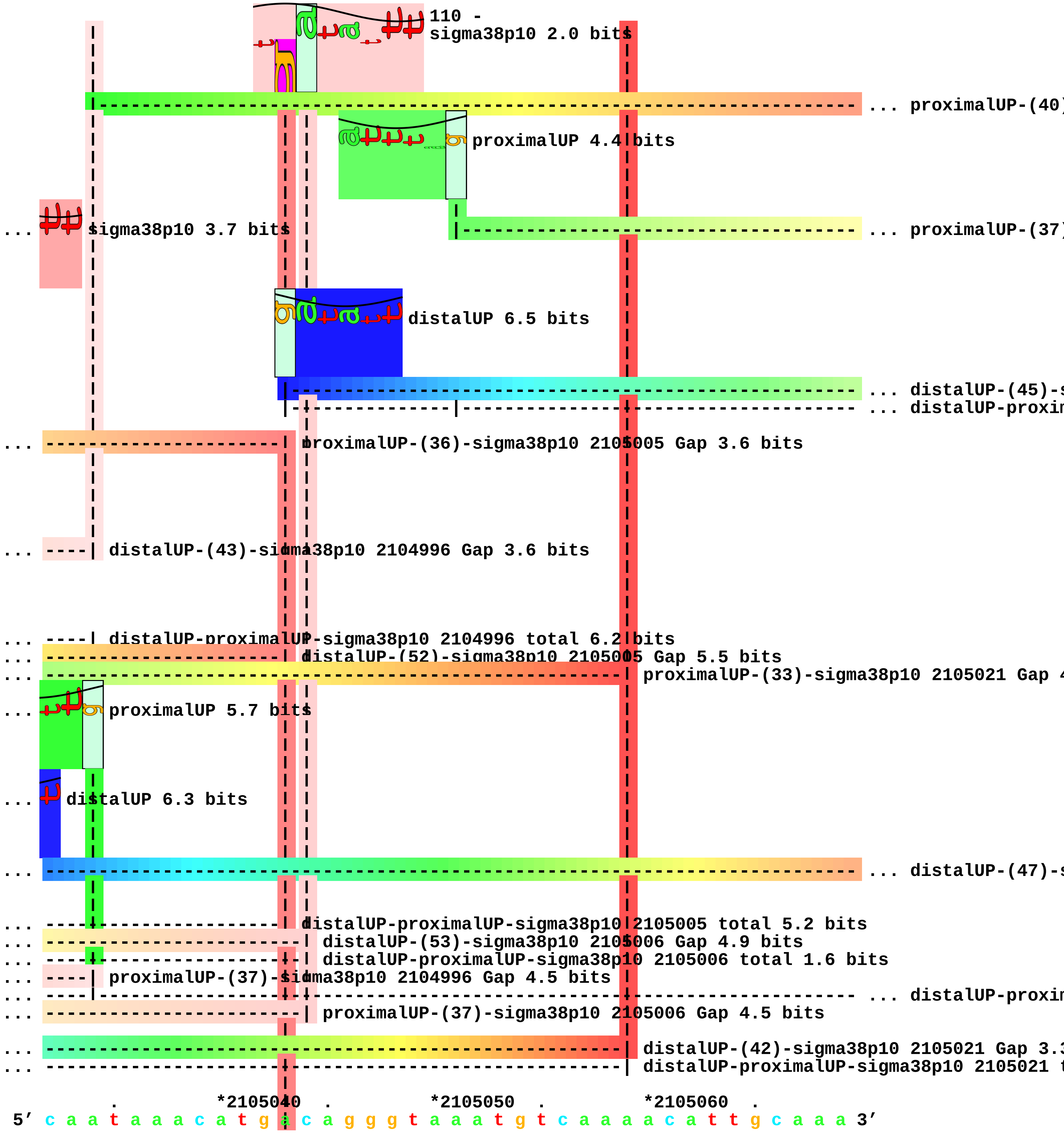

110 -
sigma38p10 2.0 bits
... proximalUP-(40)
proximalUP 4.4 bits
... sigma38p10 3.7 bits
... proximalUP-(37)
distalUP 6.5 bits
... distalUP-(45)-s
... distalUP-proxin
... proximalUP-(36)-sigma38p10 2105005 Gap 3.6 bits
... ----| distalUP-(43)-sigma38p10 2104996 Gap 3.6 bits
... ----| distalUP-proximalUP-sigma38p10 2104996 total 6.2 bits
... distalUP-(52)-sigma38p10 2105005 Gap 5.5 bits
... proximalUP-(33)-sigma38p10 2105021 Gap 4
... proximalUP 5.7 bits
... distalUP 6.3 bits
... distalUP-(47)-s
... distalUP-proximalUP-sigma38p10 2105005 total 5.2 bits
... distalUP-(53)-sigma38p10 2105006 Gap 4.9 bits
... distalUP-proximalUP-sigma38p10 2105006 total 1.6 bits
... ----| proximalUP-(37)-sigma38p10 2104996 Gap 4.5 bits
... distalUP-proxin
... proximalUP-(37)-sigma38p10 2105006 Gap 4.5 bits
... distalUP-(42)-sigma38p10 2105021 Gap 3.3
... distalUP-proximalUP-sigma38p10 2105021 t
. *2105040 . *2105050 . *2105060 .
5' c a a t a a a c a t g a c a g g g t a a a t g t c a a a a c a t t g c a a a 3'

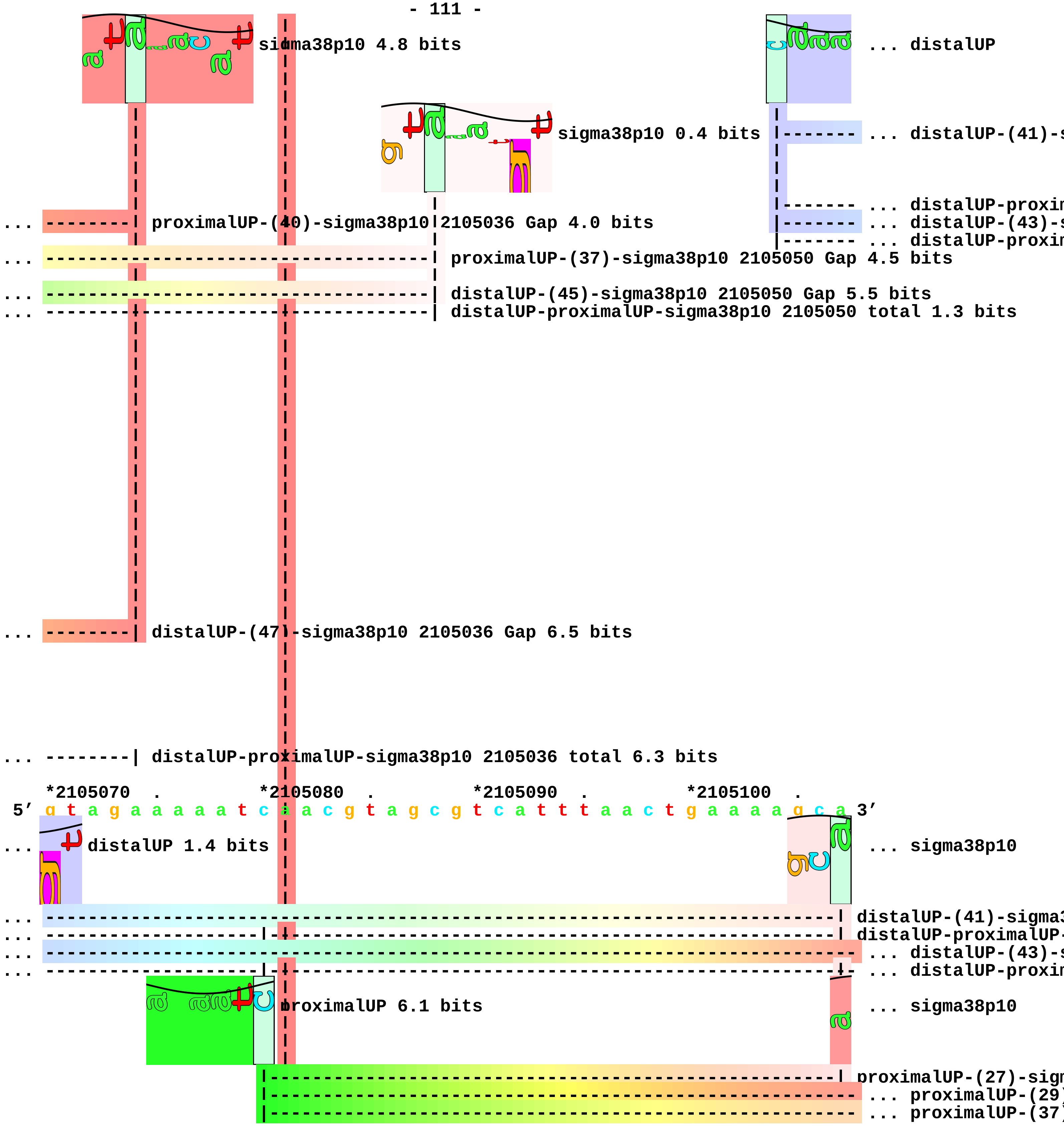
sigma38p10 4.8 bits
... distalUP
sigma38p10 0.4 bits
... distalUP-(41)-s
... distalUP-proxim
... proximalUP-(40)-sigma38p10 2105036 Gap 4.0 bits
... distalUP-(43)-s
... distalUP-proxim
... proximalUP-(37)-sigma38p10 2105050 Gap 4.5 bits
... distalUP-(45)-sigma38p10 2105050 Gap 5.5 bits
... distalUP-proximalUP-sigma38p10 2105050 total 1.3 bits
... distalUP-(47)-sigma38p10 2105036 Gap 6.5 bits
... distalUP-proximalUP-sigma38p10 2105036 total 6.3 bits
*2105070 . *2105080 . *2105090 . *2105100 .
5' g t a g a a a a a t c a a c g t a g c g t c a t t t a a c t g a a a a g c a 3'
... distalUP 1.4 bits
... sigma38p10
... distalUP-(41)-sigma3
... distalUP-proximalUP-
... distalUP-(43)-s
... distalUP-proxim
proximalUP 6.1 bits
... sigma38p10
proximalUP-(27)-sigm
... proximalUP-(29)
... proximalUP-(37)

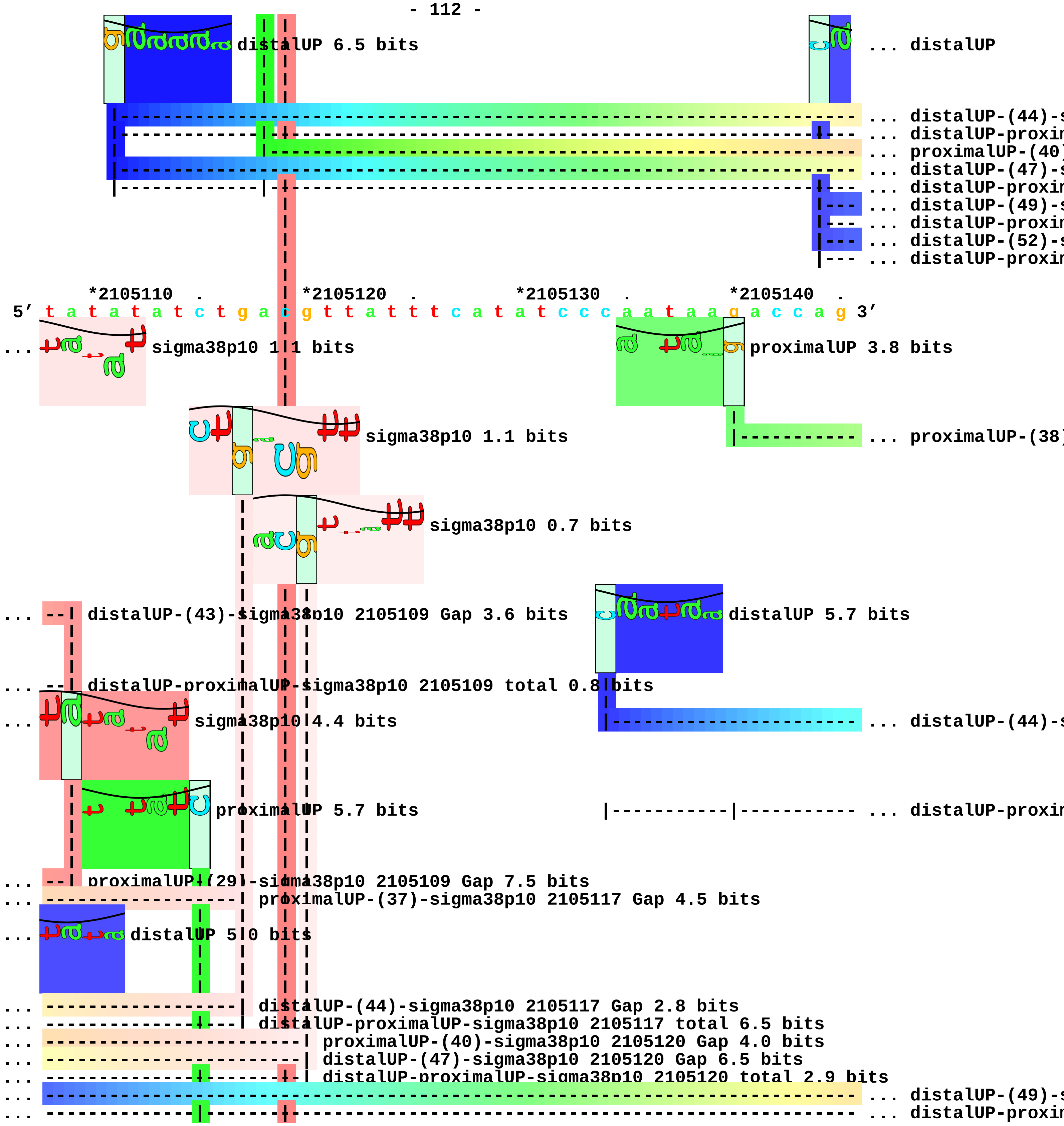
distalUP 6.5 bits
... distalUP
... distalUP-(44)-s
... distalUP-proxim
... proximalUP-(40)
... distalUP-(47)-s
... distalUP-proxim
... distalUP-(49)-s
... distalUP-proxim
... distalUP-(52)-s
... distalUP-proxim
*2105110 .
*2105120 .
*2105130 .
*2105140 .
5′ t a t a t a t c t g a c g t t a t t t c a t a t c c c a a t a a g a c c a g 3′
... sigma38p10 1.1 bits
proximalUP 3.8 bits
sigma38p10 1.1 bits
... proximalUP-(38)
sigma38p10 0.7 bits
... distalUP-(43)-sigma38p10 2105109 Gap 3.6 bits
distalUP 5.7 bits
... distalUP-proximalUP-sigma38p10 2105109 total 0.8 bits
... sigma38p10 4.4 bits
... distalUP-(44)-s
proximalUP 5.7 bits
... distalUP-proxim
... proximalUP-(29)-sigma38p10 2105109 Gap 7.5 bits
... proximalUP-(37)-sigma38p10 2105117 Gap 4.5 bits
... distalUP 5.0 bits
... distalUP-(44)-sigma38p10 2105117 Gap 2.8 bits
... distalUP-proximalUP-sigma38p10 2105117 total 6.5 bits
... proximalUP-(40)-sigma38p10 2105120 Gap 4.0 bits
... distalUP-(47)-sigma38p10 2105120 Gap 6.5 bits
... distalUP-proximalUP-sigma38p10 2105120 total 2.9 bits
... distalUP-(49)-s
... distalUP-proxim

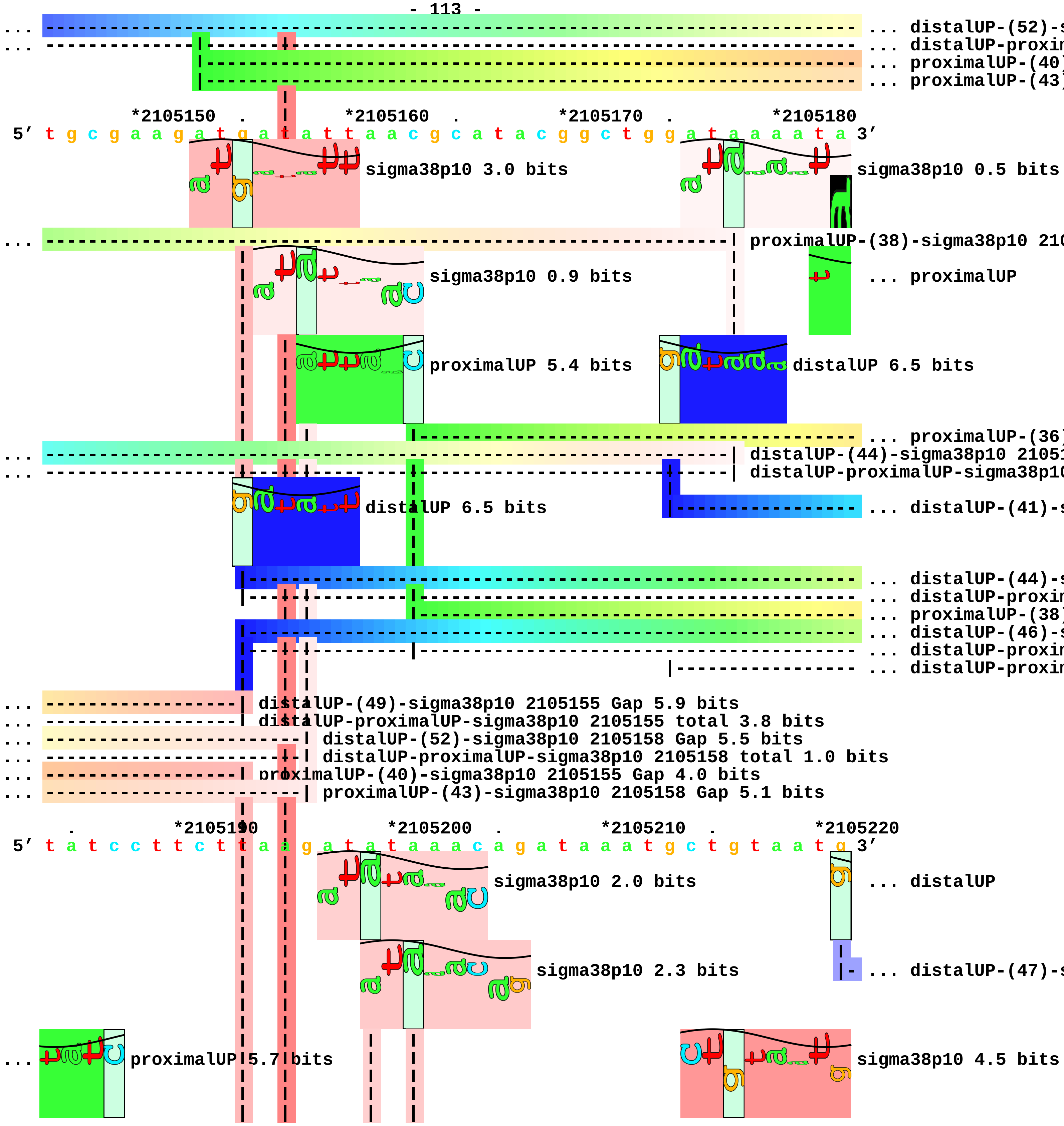
... distalUP-(52)-s
... distalUP-proxin
... proximalUP-(40)
... proximalUP-(43)
*2105150 . *2105160 . *2105170 . *2105180
5' t g c g a a g a t g a t a t t a a c g c a t a c g g c t g g a t a a a a t a 3'
sigma38p10 3.0 bits
sigma38p10 0.5 bits
... proximalUP-(38)-sigma38p10 210
sigma38p10 0.9 bits
... proximalUP
proximalUP 5.4 bits
distalUP 6.5 bits
... proximalUP-(36)
... distalUP-(44)-sigma38p10 21051
... distalUP-proximalUP-sigma38p10
distalUP 6.5 bits
... distalUP-(41)-s
... distalUP-(44)-s
... distalUP-proxin
... proximalUP-(38)
... distalUP-(46)-s
... distalUP-proxin
... distalUP-proxin
... distalUP-(49)-sigma38p10 2105155 Gap 5.9 bits
... distalUP-proximalUP-sigma38p10 2105155 total 3.8 bits
... distalUP-(52)-sigma38p10 2105158 Gap 5.5 bits
... distalUP-proximalUP-sigma38p10 2105158 total 1.0 bits
... proximalUP-(40)-sigma38p10 2105155 Gap 4.0 bits
... proximalUP-(43)-sigma38p10 2105158 Gap 5.1 bits
. *2105190 *2105200 . *2105210 . *2105220
5' t a t c c t t c t t a a g a t a t a a a c a g a t a a a t g c t g t a a t g 3'
sigma38p10 2.0 bits
... distalUP
sigma38p10 2.3 bits
... distalUP-(47)-s
... proximalUP 5.7 bits
sigma38p10 4.5 bits

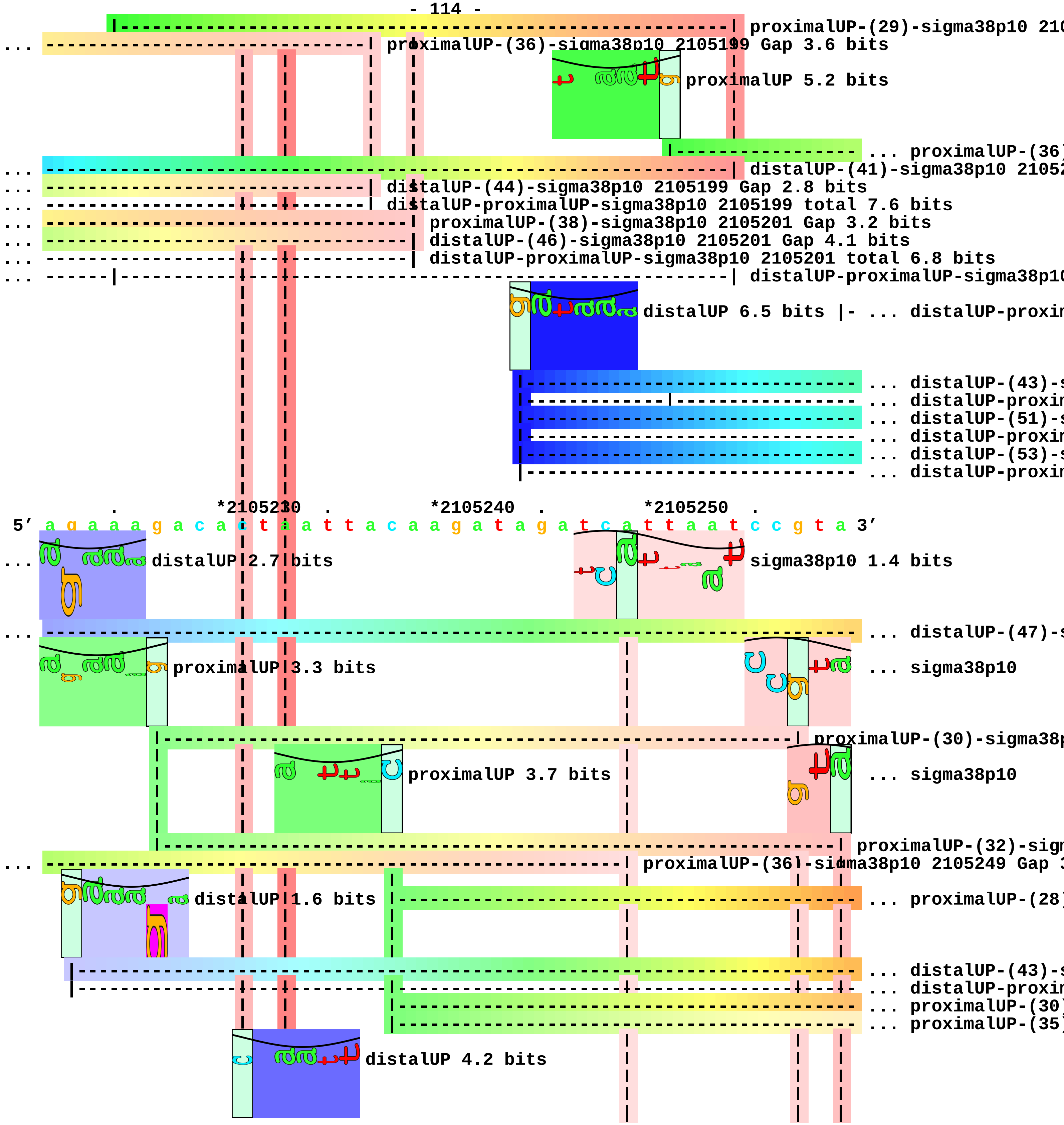
proximalUP-(29)-sigma38p10 210
proximalUP-(36)-sigma38p10 2105199 Gap 3.6 bits
proximalUP 5.2 bits
proximalUP-(36)
distalUP-(41)-sigma38p10 2105
distalUP-(44)-sigma38p10 2105199 Gap 2.8 bits
distalUP-proximalUP-sigma38p10 2105199 total 7.6 bits
proximalUP-(38)-sigma38p10 2105201 Gap 3.2 bits
distalUP-(46)-sigma38p10 2105201 Gap 4.1 bits
distalUP-proximalUP-sigma38p10 2105201 total 6.8 bits
distalUP-proximalUP-sigma38p10
distalUP 6.5 bits
distalUP-proxim
distalUP-(43)-s
distalUP-proxim
distalUP-(51)-s
distalUP-proxim
distalUP-(53)-s
distalUP-proxim
*2105230
*2105240
*2105250
5′ a g a a a g a c a c t a a t t a c a a g a t a g a t c a t t a a t c c g t a 3′
distalUP 2.7 bits
sigma38p10 1.4 bits
distalUP-(47)-s
proximalUP 3.3 bits
sigma38p10
proximalUP-(30)-sigma38p
proximalUP 3.7 bits
sigma38p10
proximalUP-(32)-sigm
proximalUP-(36)-sigma38p10 2105249 Gap 3
distalUP 1.6 bits
proximalUP-(28)
distalUP-(43)-s
distalUP-proxim
proximalUP-(30)
proximalUP-(35)
distalUP 4.2 bits

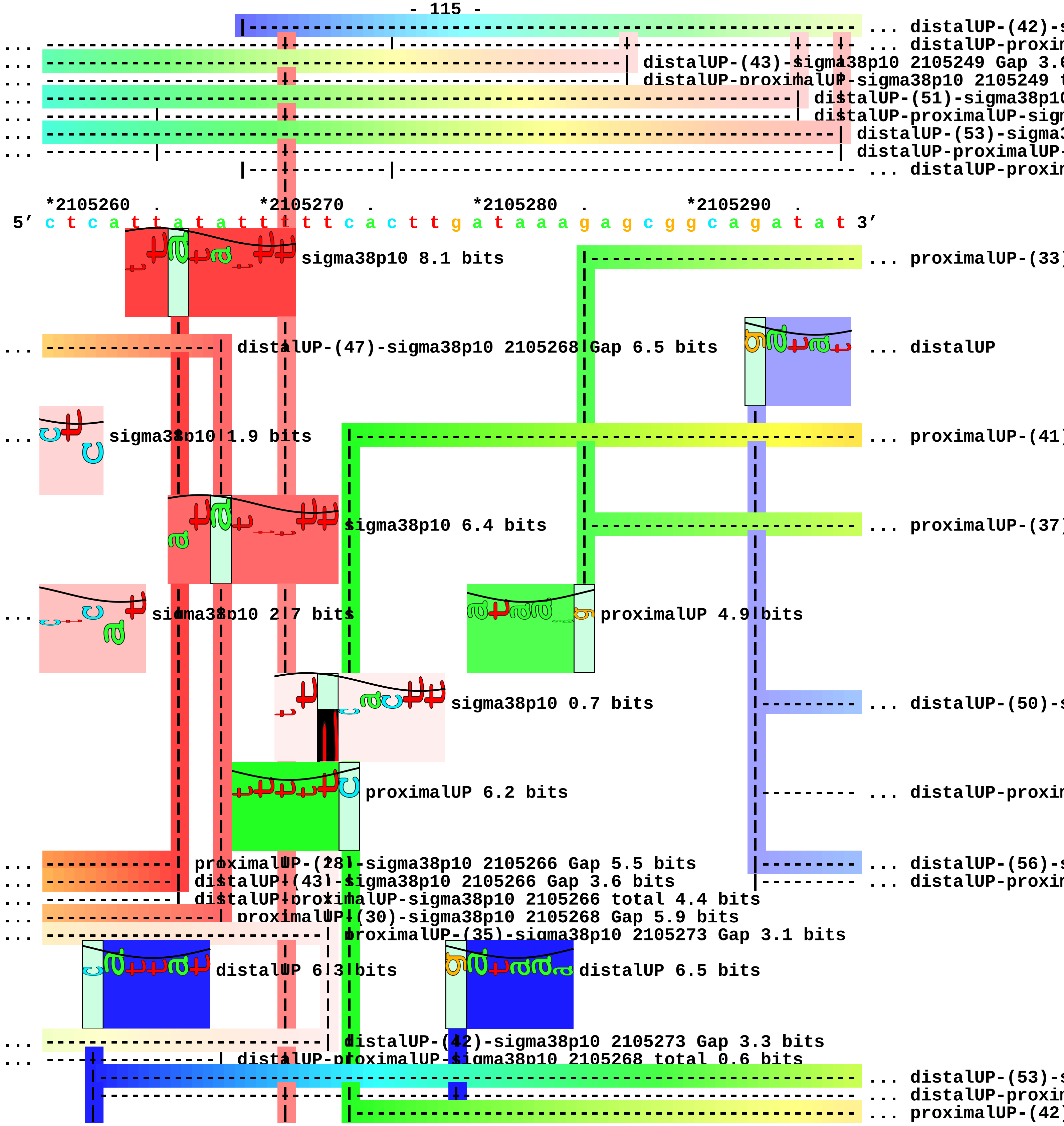
... distalUP-(42)-s
... distalUP-proxin
distalUP-(43)-sigma38p10 2105249 Gap 3.6
distalUP-proximalUP-sigma38p10 2105249 t
distalUP-(51)-sigma38p10
distalUP-proximalUP-sign
distalUP-(53)-sigma3
distalUP-proximalUP-
... distalUP-proxin
*2105260 . *2105270 . *2105280 . *2105290 .
5′ c t c a t t a t a t t t t t t c a c t t g a t a a a g a g c g g c a g a t a t 3′
sigma38p10 8.1 bits
... proximalUP-(33)
distalUP-(47)-sigma38p10 2105268 Gap 6.5 bits
... distalUP
sigma38p10 1.9 bits
... proximalUP-(41)
sigma38p10 6.4 bits
... proximalUP-(37)
sigma38p10 2.7 bits
proximalUP 4.9 bits
sigma38p10 0.7 bits
... distalUP-(50)-s
proximalUP 6.2 bits
... distalUP-proxin
proximalUP-(28)-sigma38p10 2105266 Gap 5.5 bits
... distalUP-(56)-s
distalUP-(43)-sigma38p10 2105266 Gap 3.6 bits
... distalUP-proxin
distalUP-proximalUP-sigma38p10 2105266 total 4.4 bits
proximalUP-(30)-sigma38p10 2105268 Gap 5.9 bits
proximalUP-(35)-sigma38p10 2105273 Gap 3.1 bits
distalUP 6.3 bits
distalUP 6.5 bits
distalUP-(42)-sigma38p10 2105273 Gap 3.3 bits
distalUP-proximalUP-sigma38p10 2105268 total 0.6 bits
... distalUP-(53)-s
... distalUP-proxin
... proximalUP-(42)

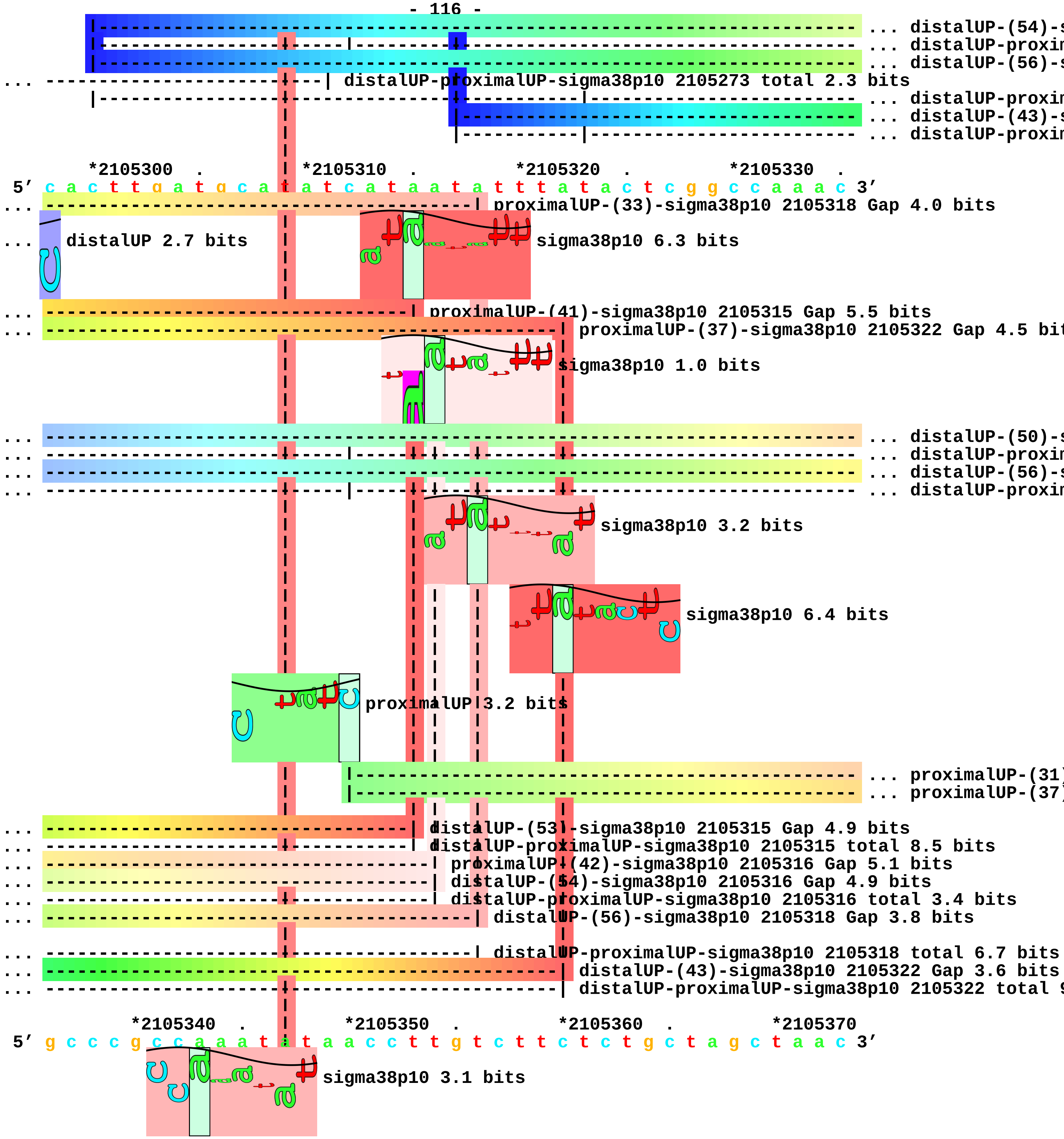
... distalUP-(54)-s
... distalUP-proxim
... distalUP-(56)-s
... distalUP-proximalUP-sigma38p10 2105273 total 2.3 bits
... distalUP-proxim
... distalUP-(43)-s
... distalUP-proxim
*2105300 . *2105310 . *2105320 . *2105330 .
5′ c a c t t g a t g c a t a t c a t a a t a t t t a t a c t c g g c c a a a c 3′
... proximalUP-(33)-sigma38p10 2105318 Gap 4.0 bits
... distalUP 2.7 bits
sigma38p10 6.3 bits
... proximalUP-(41)-sigma38p10 2105315 Gap 5.5 bits
... proximalUP-(37)-sigma38p10 2105322 Gap 4.5 bit
sigma38p10 1.0 bits
... distalUP-(50)-s
... distalUP-proxim
... distalUP-(56)-s
... distalUP-proxim
sigma38p10 3.2 bits
sigma38p10 6.4 bits
proximalUP 3.2 bits
... proximalUP-(31)
... proximalUP-(37)
... distalUP-(53)-sigma38p10 2105315 Gap 4.9 bits
... distalUP-proximalUP-sigma38p10 2105315 total 8.5 bits
... proximalUP-(42)-sigma38p10 2105316 Gap 5.1 bits
... distalUP-(54)-sigma38p10 2105316 Gap 4.9 bits
... distalUP-proximalUP-sigma38p10 2105316 total 3.4 bits
... distalUP-(56)-sigma38p10 2105318 Gap 3.8 bits
... distalUP-proximalUP-sigma38p10 2105318 total 6.7 bits
... distalUP-(43)-sigma38p10 2105322 Gap 3.6 bits
... distalUP-proximalUP-sigma38p10 2105322 total 9
*2105340 . *2105350 . *2105360 . *2105370
5′ g c c c g c c a a a t a t a a c c t t g t c t t c t c t g c t a g c t a a c 3′
sigma38p10 3.1 bits

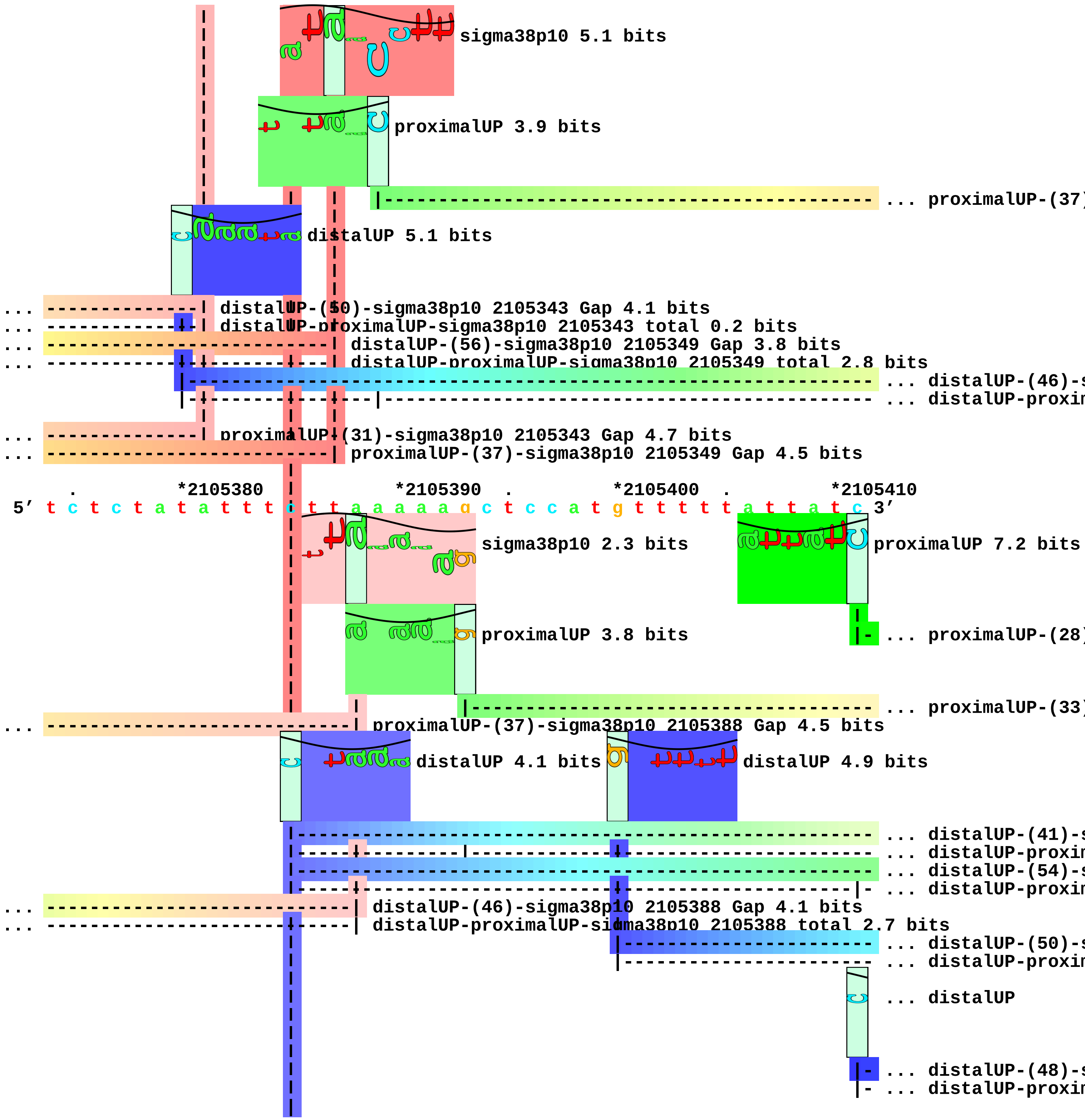
sigma38p10 5.1 bits
proximalUP 3.9 bits
... proximalUP-(37)
distalUP 5.1 bits
... distalUP-(50)-sigma38p10 2105343 Gap 4.1 bits
... distalUP-proximalUP-sigma38p10 2105343 total 0.2 bits
... distalUP-(56)-sigma38p10 2105349 Gap 3.8 bits
... distalUP-proximalUP-sigma38p10 2105349 total 2.8 bits
... distalUP-(46)-s
... distalUP-proxim
... proximalUP-(31)-sigma38p10 2105343 Gap 4.7 bits
... proximalUP-(37)-sigma38p10 2105349 Gap 4.5 bits
*2105380 *2105390 *2105400 *2105410
5′ t c t c t a t a t t t c t t a a a a a g c t c c a t g t t t t t a t t a t c 3′
sigma38p10 2.3 bits
proximalUP 7.2 bits
proximalUP 3.8 bits
... proximalUP-(28)
... proximalUP-(33)
... proximalUP-(37)-sigma38p10 2105388 Gap 4.5 bits
distalUP 4.1 bits
distalUP 4.9 bits
... distalUP-(41)-s
... distalUP-proxim
... distalUP-(54)-s
... distalUP-proxim
... distalUP-(46)-sigma38p10 2105388 Gap 4.1 bits
... distalUP-proximalUP-sigma38p10 2105388 total 2.7 bits
... distalUP-(50)-s
... distalUP-proxim
... distalUP
... distalUP-(48)-s
... distalUP-proxim

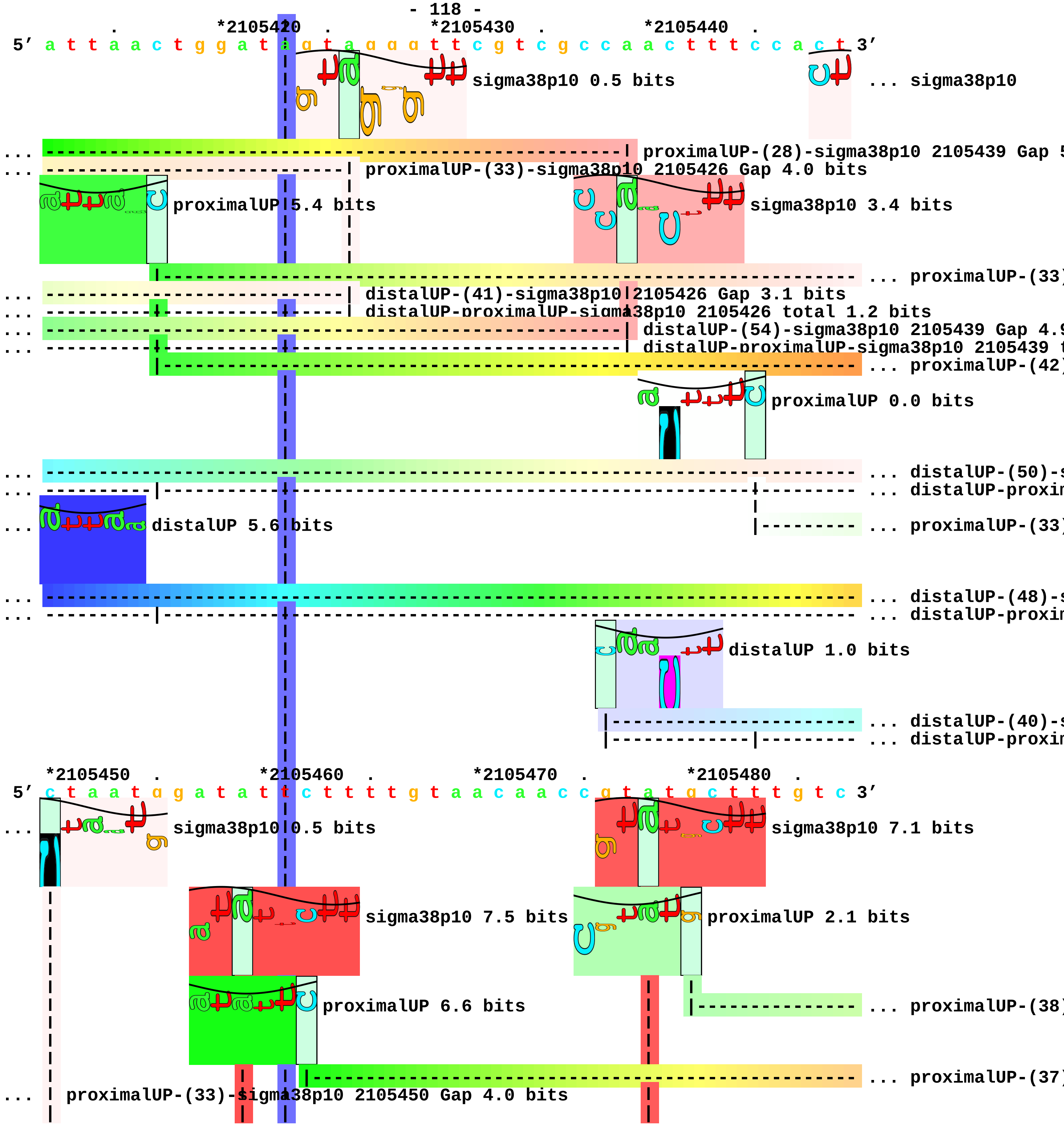

*2105420 *2105430 *2105440
5′ a t t a a c t g g a t a g t a g g g t t c g t c g c c a a c t t t c c a c t 3′
sigma38p10 0.5 bits
... sigma38p10
... proximalUP-(28)-sigma38p10 2105439 Gap 5
... proximalUP-(33)-sigma38p10 2105426 Gap 4.0 bits
proximalUP 5.4 bits
sigma38p10 3.4 bits
... proximalUP-(33)
... distalUP-(41)-sigma38p10 2105426 Gap 3.1 bits
... distalUP-proximalUP-sigma38p10 2105426 total 1.2 bits
... distalUP-(54)-sigma38p10 2105439 Gap 4.9
... distalUP-proximalUP-sigma38p10 2105439 t
... proximalUP-(42)
proximalUP 0.0 bits
... distalUP-(50)-s
... distalUP-proxin
distalUP 5.6 bits
... proximalUP-(33)
... distalUP-(48)-s
... distalUP-proxin
distalUP 1.0 bits
... distalUP-(40)-s
... distalUP-proxin
*2105450 *2105460 *2105470 *2105480
5′ c t a a t g g a t a t t c t t t t g t a a c a a c c g t a t g c t t t g t c 3′
... sigma38p10 0.5 bits
sigma38p10 7.1 bits
sigma38p10 7.5 bits
proximalUP 2.1 bits
proximalUP 6.6 bits
... proximalUP-(38)
... proximalUP-(37)
... proximalUP-(33)-sigma38p10 2105450 Gap 4.0 bits

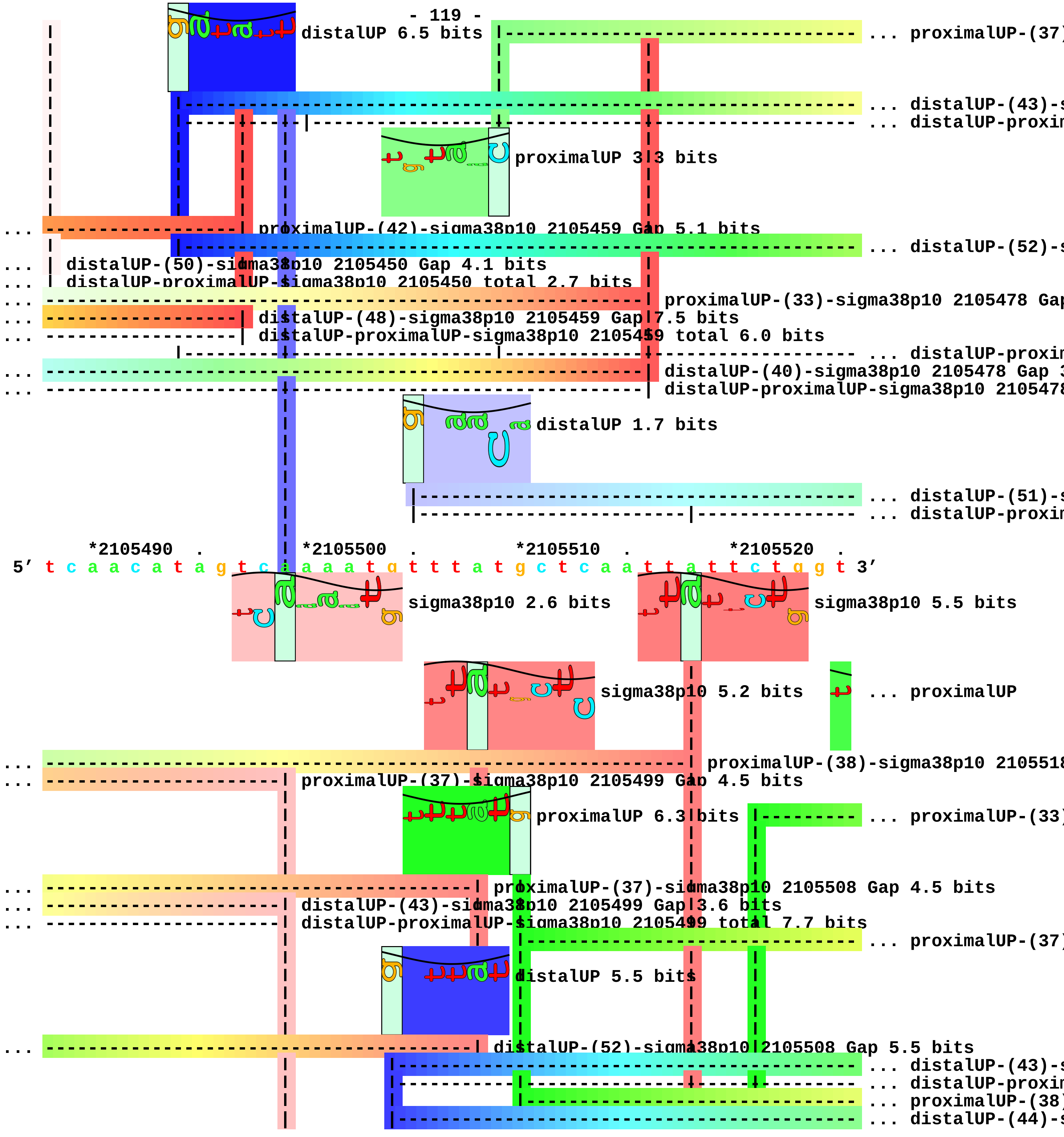
distalUP 6.5 bits
... proximalUP-(37)
... distalUP-(43)-s
... distalUP-proxim
proximalUP 3.3 bits
proximalUP-(42)-sigma38p10 2105459 Gap 5.1 bits
... distalUP-(52)-s
distalUP-(50)-sigma38p10 2105450 Gap 4.1 bits
distalUP-proximalUP-sigma38p10 2105450 total 2.7 bits
proximalUP-(33)-sigma38p10 2105478 Gap
distalUP-(48)-sigma38p10 2105459 Gap 7.5 bits
distalUP-proximalUP-sigma38p10 2105459 total 6.0 bits
... distalUP-proxim
distalUP-(40)-sigma38p10 2105478 Gap 3
distalUP-proximalUP-sigma38p10 2105478
distalUP 1.7 bits
... distalUP-(51)-s
... distalUP-proxim
*2105490 . *2105500 . *2105510 . *2105520 .
5′ t c a a c a t a g t c a a a a t g t t t a t g c t c a a t t a t t c t g g t 3′
sigma38p10 2.6 bits
sigma38p10 5.5 bits
sigma38p10 5.2 bits
... proximalUP
proximalUP-(38)-sigma38p10 2105518
proximalUP-(37)-sigma38p10 2105499 Gap 4.5 bits
proximalUP 6.3 bits
... proximalUP-(33)
proximalUP-(37)-sigma38p10 2105508 Gap 4.5 bits
distalUP-(43)-sigma38p10 2105499 Gap 3.6 bits
distalUP-proximalUP-sigma38p10 2105499 total 7.7 bits
... proximalUP-(37)
distalUP 5.5 bits
distalUP-(52)-sigma38p10 2105508 Gap 5.5 bits
... distalUP-(43)-s
... distalUP-proxim
... proximalUP-(38)
... distalUP-(44)-s

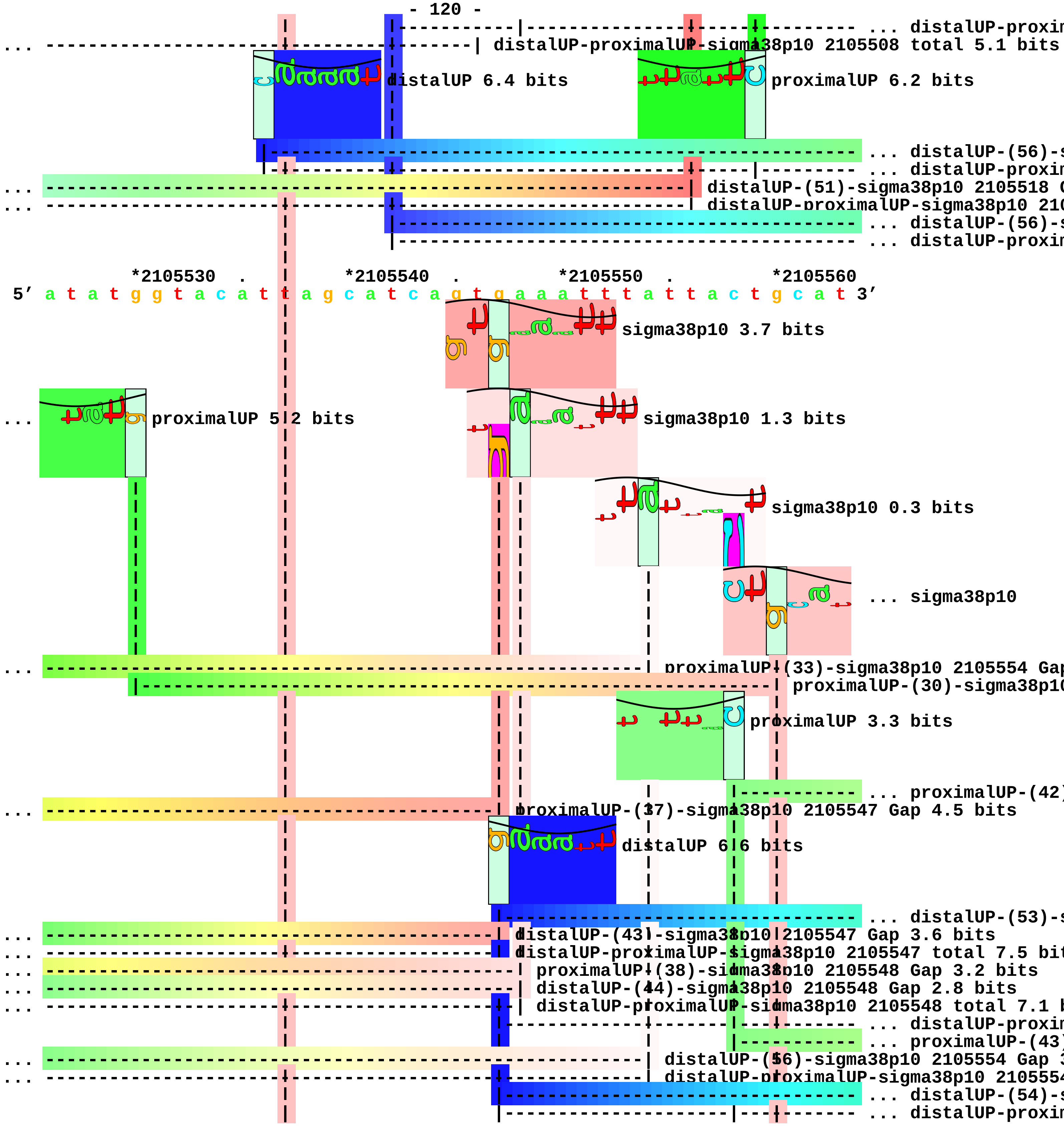
... distalUP-proxin
... distalUP-proximalUP-sigma38p10 2105508 total 5.1 bits
distalUP 6.4 bits
proximalUP 6.2 bits
... distalUP-(56)-s
... distalUP-proxin
... distalUP-(51)-sigma38p10 2105518 G
... distalUP-proximalUP-sigma38p10 210
... distalUP-(56)-s
... distalUP-proxin
*2105530 . *2105540 . *2105550 . *2105560
5' a t a t g g t a c a t t a g c a t c a g t g a a a t t t a t t a c t g c a t 3'
sigma38p10 3.7 bits
... proximalUP 5.2 bits
sigma38p10 1.3 bits
sigma38p10 0.3 bits
... sigma38p10
... proximalUP-(33)-sigma38p10 2105554 Gap
proximalUP-(30)-sigma38p10
proximalUP 3.3 bits
... proximalUP-(42)
... proximalUP-(37)-sigma38p10 2105547 Gap 4.5 bits
distalUP 6.6 bits
... distalUP-(53)-s
... distalUP-(43)-sigma38p10 2105547 Gap 3.6 bits
... distalUP-proximalUP-sigma38p10 2105547 total 7.5 bit
... proximalUP-(38)-sigma38p10 2105548 Gap 3.2 bits
... distalUP-(44)-sigma38p10 2105548 Gap 2.8 bits
... distalUP-proximalUP-sigma38p10 2105548 total 7.1 b
... distalUP-proxin
... proximalUP-(43)
... distalUP-(56)-sigma38p10 2105554 Gap 3
... distalUP-proximalUP-sigma38p10 210555
... distalUP-(54)-s
... distalUP-proxin

```
... -----------------------------------------------------------------------| distalUP-(56)-sigma38p10 2
... --------|--------------------------------------------------------------| distalUP-proximalUP-sigma3

       .            *2105570            *2105580   .        *2105590   .        *2105600
5' t c c c t t g g a a g t t t g g a a a t t c a t g g c g t t c c g t c t c a 3'
                                                                          ... sigma38p10
                                                                          ... sigma38p10
... sigma38p10 2.5 bits
... --------------------------------------------------------------------------| proximalUP-(42)-sigma3
... --------------------------------------------------------------------------| distalUP-(53)-sigma38p
... --------------------------------------------------------------------------| distalUP-proximalUP-si
... ----------------------------------------------------------------------------| proximalUP-(43)-sigm
... ----------------------------------------------------------------------------| distalUP-(54)-sigma3
... ----------------------------------------------------------------------------| distalUP-proximalUP-

       .            *2105610   .        *2105620   .        *2105630   .
5' a a t t t t a a a g a g c g a t a t t c t a a c g c t c c a a a c c t a t a 3'
... sigma38p10 2.5 bits
... sigma38p10 4.7 bits

   *2105640   .
5' g t c g a a g t a c 3'
```

piece 30, NC_000913, yehE, config: linear, direction: +, begin: 2190750, end: 2190999

```
   *2190750  .         *2190760  .         *2190770  .         *2190780  .
5' t g a a g a g a a t g c g g g a g a a g c c a a t c c a t a g g c a a g a a 3'
   T timestamp:_2018Sep20_11-35-24

     *2190790  .         *2190800  .         *2190810  .         *2190820  .
5' a a a t a a t t c c t g a c a g c c a a t a c t t a t t c a t t g a a c g t 3'

       *2190830  .         *2190840  .         *2190850  .         *2190860
5' t a t c c c t g t a g t a a a g g t t a t g c c t g g a t a g a a t g a g t 3'
                                                                proximalUP 5.0 bits
                                                                |------- ... proximalUP-(33)
                                                          distalUP 1.5 bits
                                                          |----------------------- ... distalUP-(41)-s
                                                          |---------------|------- ... distalUP-proxim

     .         *2190870  .         *2190880  .         *2190890  .         *2190900
5' g c a t a a c a a a c t a t a g c t g t a c a t c c a c t a c a c a g c c a 3'
                                                                sigma38p10 4.0 bits
... -------------------------------------------------------------| proximalUP-(33)-sigma38p10 2190893 G
                                                                 |
... -------------------------------------------------------------| distalUP-(41)-sigma38p10 2190893 Gap
... -------------------------------------------------------------| distalUP-proximalUP-sigma38p10 21908

       .         *2190910  .         *2190920  .         *2190930  .
5' c g a a g g a t g a t a a t g a a g c a t t g c c t g t a t g a t c a a t c 3'
                   proximalUP 6.7 bits
                            |------------------------------------------------ ... proximalUP-(32)
           distalUP 0.7 bits
           |------------------------------------------------------------------ ... distalUP-(41)-s
           |-----------------|------------------------------------------------ ... distalUP-proxim

   *2190940  .         *2190950  .         *2190960  .         *2190970  .
5' g a c t t t g t a g a g t t t c g g a c g a a g g t c c g c a g a a t a t t 3'
                  sigma38p10 1.7 bits
... ----------------| proximalUP-(32)-sigma38p10 2190948 Gap 4.3 bits
```

```
                   |
... ----------------| distalUP-(41)-sigma38p10 2190948 Gap 3.1 bits
... ----------------| distalUP-proximalUP-sigma38p10 2190948 total 1.7 bits

        *2190980  .         *2190990  .
 5' c g c a g t a t t a a a t a a g t g t t c a 3'
```

piece 31, NC_000913, yohF, config: linear, direction: +, begin: 2225229, end: 2225440

```
     *2225230   .         *2225240   .         *2225250   .         *2225260   .
5' g t a a t a a c g c g c a c t c t t t g c c g a t c c c c g a a t c g g a g 3'
```

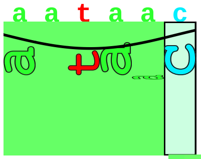

proximalUP 4.3 bits

|------------------------------------------------------------- ... proximalUP-(36)

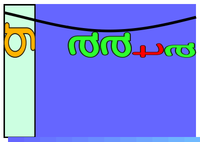

distalUP 4.4 bits

|------------------------------------------------------------- ... distalUP-(43)-s
|-------------|----------------------------------------------- ... distalUP-proxim

T timestamp:_2018Sep20_11-35-24

```
       *2225270   .         *2225280   .         *2225290   .         *2225300
5' g c g g t a a t a a t c g c a a c c t g t g c c a t c g a g t t c t c c a c 3'
```

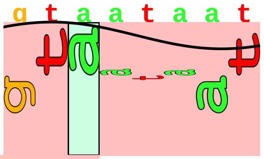

sigma38p10 2.7 bits

... ----------| proximalUP-(36)-sigma38p10 2225272 Gap 3.6 bits
... ----------| distalUP-(43)-sigma38p10 2225272 Gap 3.6 bits
... ----------| distalUP-proximalUP-sigma38p10 2225272 total 4.3 bits

```
   .         *2225310   .         *2225320   .         *2225330   .         *2225340
5' t t a a c g c t g a a t a a a c g t t a a g t a t a g a a g g c g c a t a t 3'

     .         *2225350   .         *2225360   .         *2225370   .         *2225380
5' c a t c a g c g t t t g t a c c c c c c g c c c a a c g c a c c a g t g a g 3'

       .         *2225390   .         *2225400   .         *2225410   .
5' t t g a a t g g a g g c a t c c a g c c a c t g c c c t t g c a a t a a c a 3'

     *2225420   .         *2225430   .         *2225440
5' g g c c a t t g g c c c g c t c a c g c a g 3'
```

piece 32, NC_000913, tfaS, config: linear, direction: -, begin: 2468932, end: 2468627

```
      *2468930   .         *2468920   .         *2468910   .         *2468900   .
5' t t c t g c t g c a t c t a c t g c g a c g c t a t g c t g t g c c t c g g 3'
   T timestamp:_2018Sep20_11-35-24

          *2468890   .         *2468880   .         *2468870   .         *2468860
5' t a t c c g t c a c c c a t t c t c a c c a t c c c a t t t a t c g t a t 3'
```

proximalUP 6.8 bits

... proximalUP-(37)

distalUP 5.0 bits

... distalUP-(44)-s

... sigma38p10-prox

... proximalUP

... distalUP-(46)-s

... sigma38p10-prox

```
    .         *2468850   .         *2468840   .         *2468830   .         *2468820
5' g g c g t t a a t g g g g c g a t a g t c g c g g g a c g c a t a c a a a c 3'
```

proximalUP 6.3 bits

sigma38p10 0.8 bits

... proximalUP-(37)-sigma38p10 246

distalUP 5.5 bits

... sigma38p10

... distalUP-(44)-sigma38p10 24688

... sigma38p10-proximalUP-distalUP

... proximalUP 2.1 bits

... distalUP-(46)-sigma38p10 2

... sigma38p10-proximalUP-dist

proximalUP-(34)-sigma38p10

... proximalUP-(42)

... distalUP-(48)-s

... sigma38p10-prox

```
      .         *2468810   .         *2468800   .         *2468790   .
5' a t a a c c c c t g g a t g t t a t g t g t c t a t c g a g a a t c a a a g 3'
```

sigma38p10 1.0 bits

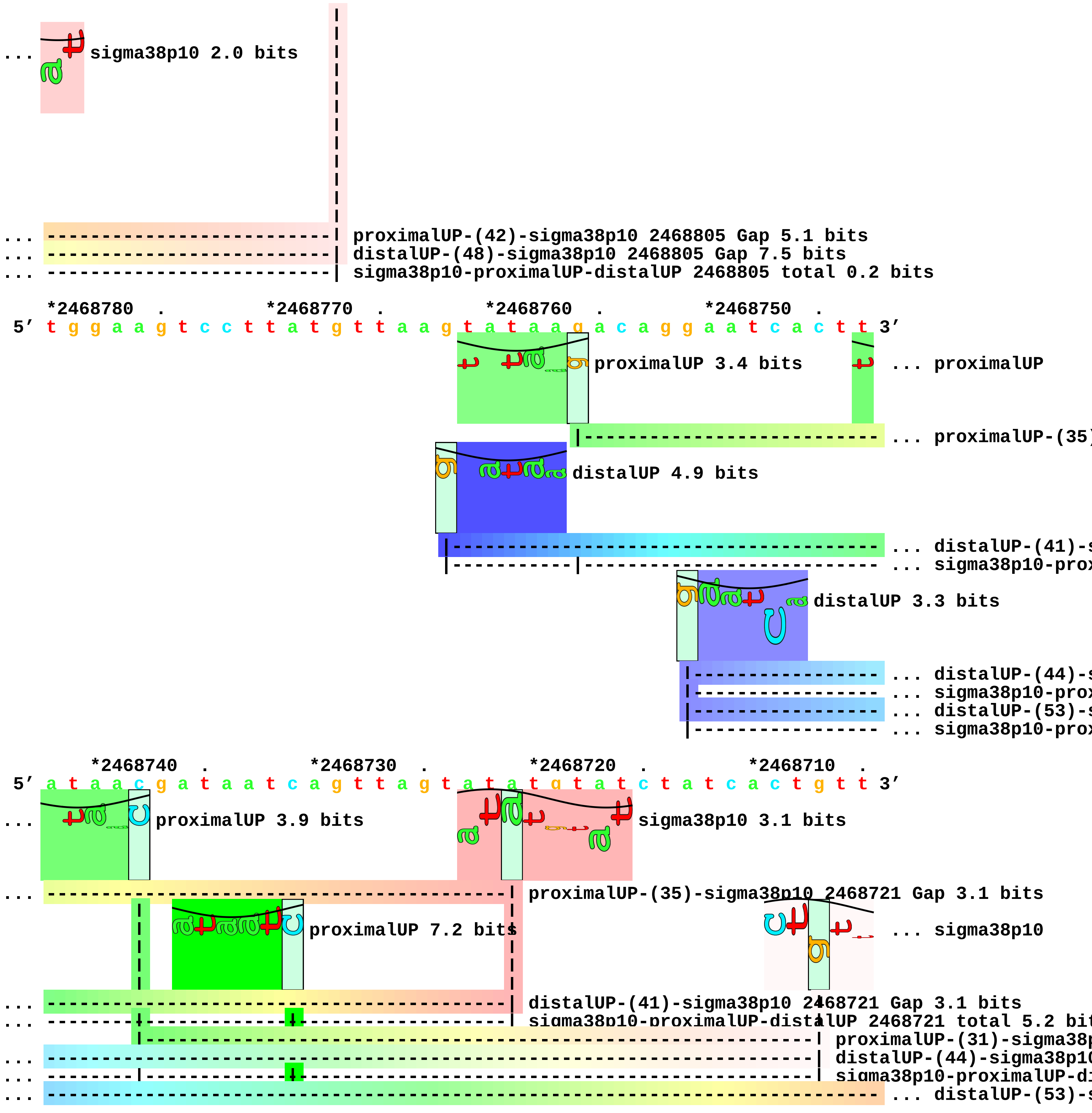
... sigma38p10 2.0 bits
... proximalUP-(42)-sigma38p10 2468805 Gap 5.1 bits
... distalUP-(48)-sigma38p10 2468805 Gap 7.5 bits
... sigma38p10-proximalUP-distalUP 2468805 total 0.2 bits
*2468780 . *2468770 . *2468760 . *2468750 .
5' t g g a a g t c c t t a t g t t a a g t a t a a g a c a g g a a t c a c t t 3'
proximalUP 3.4 bits
... proximalUP
... proximalUP-(35)
distalUP 4.9 bits
... distalUP-(41)-s
... sigma38p10-prox
distalUP 3.3 bits
... distalUP-(44)-s
... sigma38p10-prox
... distalUP-(53)-s
... sigma38p10-prox
*2468740 . *2468730 . *2468720 . *2468710 .
5' a t a a c g a t a a t c a g t t a g t a t a t g t a t c t a t c a c t g t t 3'
... proximalUP 3.9 bits
sigma38p10 3.1 bits
... proximalUP-(35)-sigma38p10 2468721 Gap 3.1 bits
proximalUP 7.2 bits
... sigma38p10
... distalUP-(41)-sigma38p10 2468721 Gap 3.1 bits
... sigma38p10-proximalUP-distalUP 2468721 total 5.2 bit
proximalUP-(31)-sigma38p
... distalUP-(44)-sigma38p10
... sigma38p10-proximalUP-di
... distalUP-(53)-s

... ------------------------|-------------------------------------------------------- ... sigma38p10-prox
... |-------------------------------------------------------- ... proximalUP-(33)

*2468700 . *2468690 . *2468680 . *2468670

5′ g a t g a t a a t a t c a g c a c t t g g t t c t g g a g g g g g t t t g t 3′

sigma38p10 3.5 bits

proximalUP 6.1 bits

... sigma38p10 0.3 bits

|-------------------------------------------------------- ... proximalUP-(40)

distalUP 7.1 bits

--------------------------------------------------------------- ... distalUP-(48)-s
|---------------|------------------------------------------------------ ... sigma38p10-prox

... ------------| distalUP-(53)-sigma38p10 2468698 Gap 4.9 bits
... ------------| sigma38p10-proximalUP-distalUP 2468698 total 5.1 bits
... ------------| proximalUP-(33)-sigma38p10 2468698 Gap 4.0 bits

. *2468660 . *2468650 . *2468640 . *2468630

5′ t g t g g g c a a t g a t g c a t t t a a g t t a t c g t c t g c a g a t a 3′

sigma38p10 3.0 bits

... --------------------------| proximalUP-(40)-sigma38p10 2468653 Gap 4.0 bits

... --------------------------| distalUP-(48)-sigma38p10 2468653 Gap 7.5 bits
... --------------------------| sigma38p10-proximalUP-distalUP 2468653 total 4.8 bits

5′ g a 3′

piece 33, NC_000913, csiE, config: linear, direction: -, begin: 2663551, end: 2663314

```
   *2663550   .         *2663540   .         *2663530   .         *2663520   .
5' g t g g c g g t c a a c c c c g g c t g a a a g a g c g t c a g c a a g a t 3'
 T timestamp:_2018Sep20_11-35-24

       *2663510   .         *2663500   .         *2663490   .         *2663480
5' c t g g c a g c g g c g c t g g g g a g c c g a a a g g a c a g a t g g t g 3'

   .         *2663470   .         *2663460   .         *2663450   .         *2663440
5' g a g c a a g c g t a g g c a t c a t c t g t a g a t c t c c c a t a a g c 3'
                                            proximalUP 3.7 bits                ... distalUP
                                            |-------------------------------------- ... proximalUP-(38)
                                            distalUP 1.5 bits                  |- ... distalUP-(54)-s
                               |-------------------------------------------------- ... distalUP-(44)-s
                               |-----------|-------------------------------------- ... sigma38p10-prox
                                                                               |- ... sigma38p10-prox

     .         *2663430   .         *2663420   .         *2663410   .         *2663400
5' a t a a t t g a t a a g a a t a g c t a a a g t t t g c t c a a g g a a a t 3'
... distalUP 6.3 bits                 sigma38p10 7.4 bits
... --------------------------------------| proximalUP-(38)-sigma38p10 2663418 Gap 3.2 bits
... ----------------------------------------------------------------------------- ... distalUP-(54)-s
... --------------------------------------| distalUP-(44)-sigma38p10 2663418 Gap 2.8 bits
... --------------------------------------| sigma38p10-proximalUP-distalUP 2663418 total 6.6 bits
... ----------------------------------------------------|------------------------ ... sigma38p10-prox
                                                        proximalUP 4.5 bits
                                                        |------------------------ ... proximalUP-(27)

         .         *2663390   .         *2663380   .         *2663370   .
5' g g c g a a g g g a a t g c t a a t c a t c a g a a a t g t t g g c a a g c 3'
                                sigma38p10 5.9 bits
... ------------------------------| distalUP-(54)-sigma38p10 2663384 Gap 4.9 bits
... ------------------------------| sigma38p10-proximalUP-distalUP 2663384 total 8.1 bits
... ------------------------------| proximalUP-(27)-sigma38p10 2663384 Gap 3.8 bits

   *2663360   .         *2663350   .         *2663340   .         *2663330   .
```

5′ g t c a c a g a a t t t t t c t c a t c t t a a a c t g t t c t g a c c a g 3′

*2663320 .
5′ t g g a t t a g c a 3′

piece 34, NC_000913, raiA, config: linear, direction: -, begin: 2735131, end: 2734860

```
     *2735130   .         *2735120   .         *2735110   .         *2735100   .
5' g t t c c t g a t g t c a a a a t g t g t g a t g a a a a t c t c a t a a t 3'

T timestamp:_2018Sep20_11-35-24
proximalUP 6.1 bits
distalUP 6.4 bits
... proximalUP-(28)
... distalUP-(47)-s
... sigma38p10-prox
distalUP 6.3 bits
... distalUP-(56)-s
... sigma38p10-prox

       *2735090   .         *2735080   .         *2735070   .         *2735060
5' t t t g t c a c t t t t t g t c a a c g a a t c t t t t t t g t g a g a a a 3'

sigma38p10 3.0 bits
... proximalUP-(28)-sigma38p10 2735073 Gap 5.5 bits
... distalUP-(47)-sigma38p10 2735073 Gap 6.5 bits
... sigma38p10-proximalUP-distalUP 2735073 total 3.5 bits
proximalUP 5.7 bits
proximalUP 5.7 bits
... distalUP-(56)-s
... sigma38p10-prox
... proximalUP-(37)
distalUP 4.1 bits
... proximalUP-(28)
... distalUP-(51)-s
... sigma38p10-prox
distalUP 2.7 bits
... distalUP-(54)-s
... sigma38p10-prox

   .         *2735050   .         *2735040   .         *2735030   .         *2735020
5' t c g c a g g a a g a g a t g t t t t t t t a c t t g a g g a a t a t c t c 3'

sigma38p10 0.2 bits
```

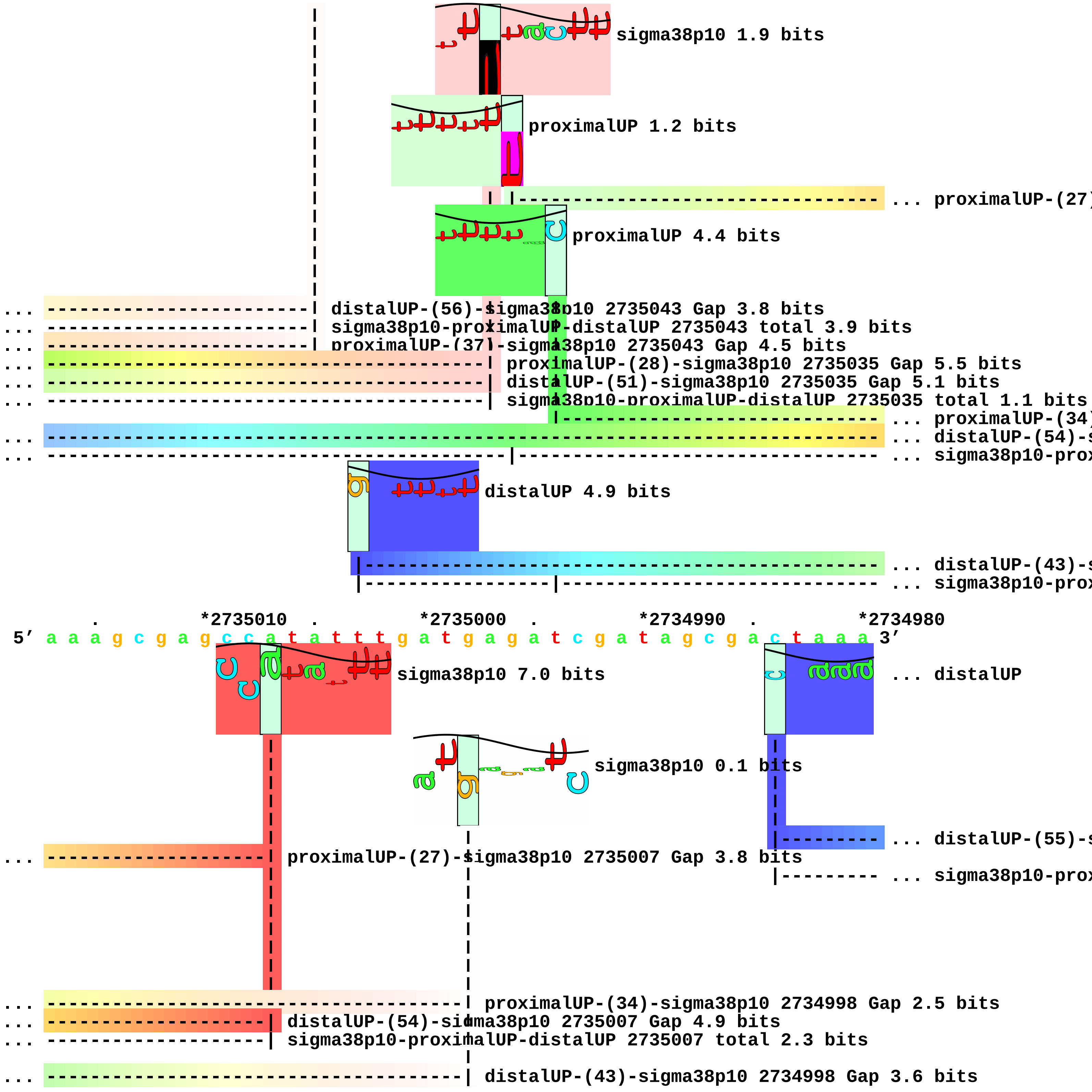
sigma38p10 1.9 bits
proximalUP 1.2 bits
... proximalUP-(27)
proximalUP 4.4 bits
... distalUP-(56)-sigma38p10 2735043 Gap 3.8 bits
... sigma38p10-proximalUP-distalUP 2735043 total 3.9 bits
... proximalUP-(37)-sigma38p10 2735043 Gap 4.5 bits
... proximalUP-(28)-sigma38p10 2735035 Gap 5.5 bits
... distalUP-(51)-sigma38p10 2735035 Gap 5.1 bits
... sigma38p10-proximalUP-distalUP 2735035 total 1.1 bits
... proximalUP-(34)
... distalUP-(54)-s
... sigma38p10-prox
distalUP 4.9 bits
... distalUP-(43)-s
... sigma38p10-prox
. *2735010 . *2735000 . *2734990 . *2734980
5′ a a a g c g a g c c a t a t t t g a t g a g a t c g a t a g c g a c t a a a 3′
sigma38p10 7.0 bits
... distalUP
sigma38p10 0.1 bits
... distalUP-(55)-s
... proximalUP-(27)-sigma38p10 2735007 Gap 3.8 bits
... sigma38p10-prox
... proximalUP-(34)-sigma38p10 2734998 Gap 2.5 bits
... distalUP-(54)-sigma38p10 2735007 Gap 4.9 bits
... sigma38p10-proximalUP-distalUP 2735007 total 2.3 bits
... distalUP-(43)-sigma38p10 2734998 Gap 3.6 bits

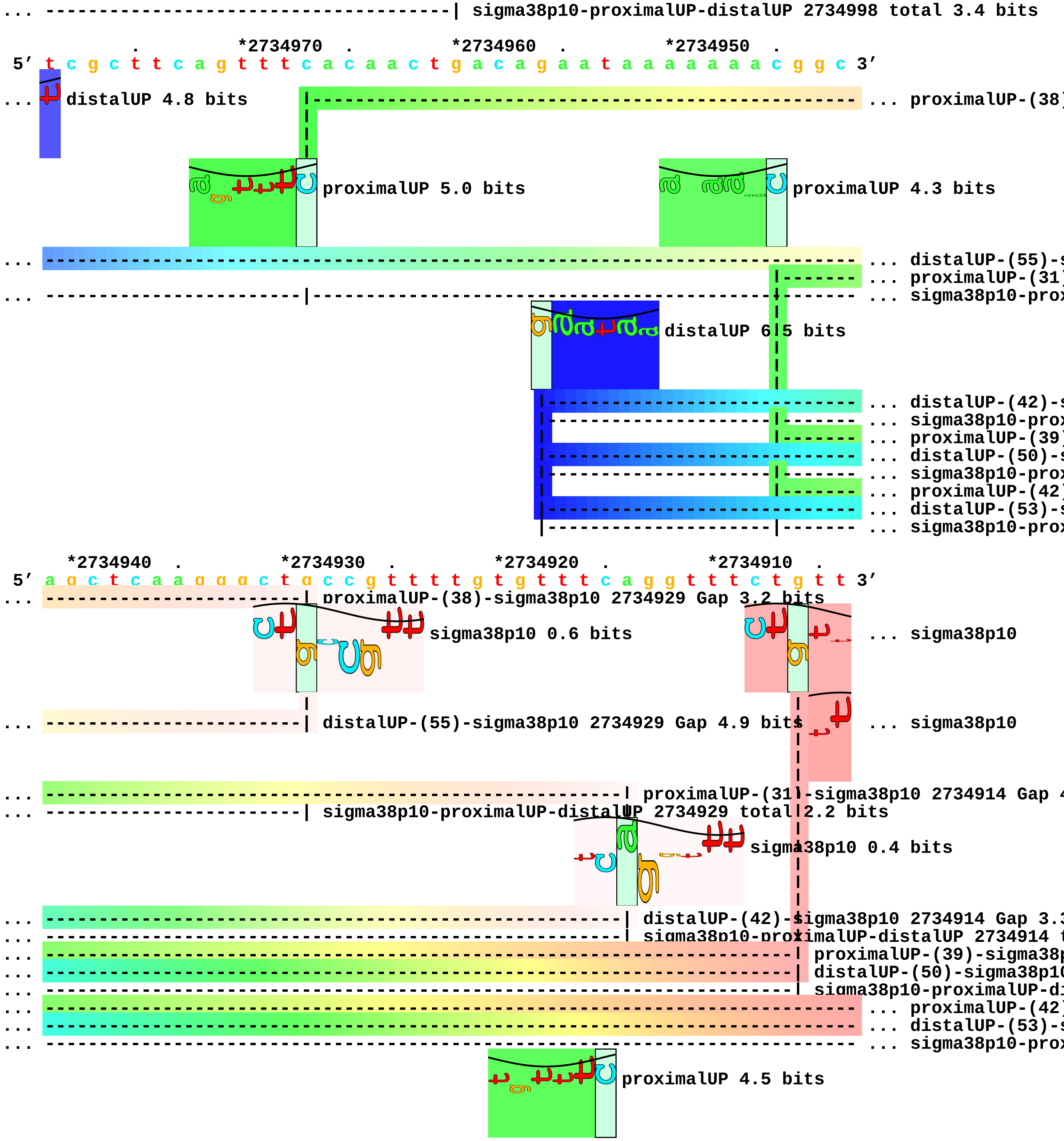
... -------------------------------------| sigma38p10-proximalUP-distalUP 2734998 total 3.4 bits
*2734970 . *2734960 . *2734950 .
5' t c g c t t c a g t t t c a c a a c t g a c a g a a t a a a a a a a c g g c 3'
... distalUP 4.8 bits
... proximalUP-(38)
proximalUP 5.0 bits
proximalUP 4.3 bits
... distalUP-(55)-s
... proximalUP-(31)
... sigma38p10-prox
distalUP 6.5 bits
... distalUP-(42)-s
... sigma38p10-prox
... proximalUP-(39)
... distalUP-(50)-s
... sigma38p10-prox
... proximalUP-(42)
... distalUP-(53)-s
... sigma38p10-prox
*2734940 . *2734930 . *2734920 . *2734910 .
5' a g c t c a a g g g c t g c c g t t t t g t g t t t c a g g t t t c t g t t 3'
proximalUP-(38)-sigma38p10 2734929 Gap 3.2 bits
sigma38p10 0.6 bits
... sigma38p10
distalUP-(55)-sigma38p10 2734929 Gap 4.9 bits
... sigma38p10
proximalUP-(31)-sigma38p10 2734914 Gap 4
sigma38p10-proximalUP-distalUP 2734929 total 2.2 bits
sigma38p10 0.4 bits
distalUP-(42)-sigma38p10 2734914 Gap 3.3
sigma38p10-proximalUP-distalUP 2734914 t
proximalUP-(39)-sigma38p
distalUP-(50)-sigma38p10
sigma38p10-proximalUP-di
... proximalUP-(42)
... distalUP-(53)-s
... sigma38p10-prox
proximalUP 4.5 bits

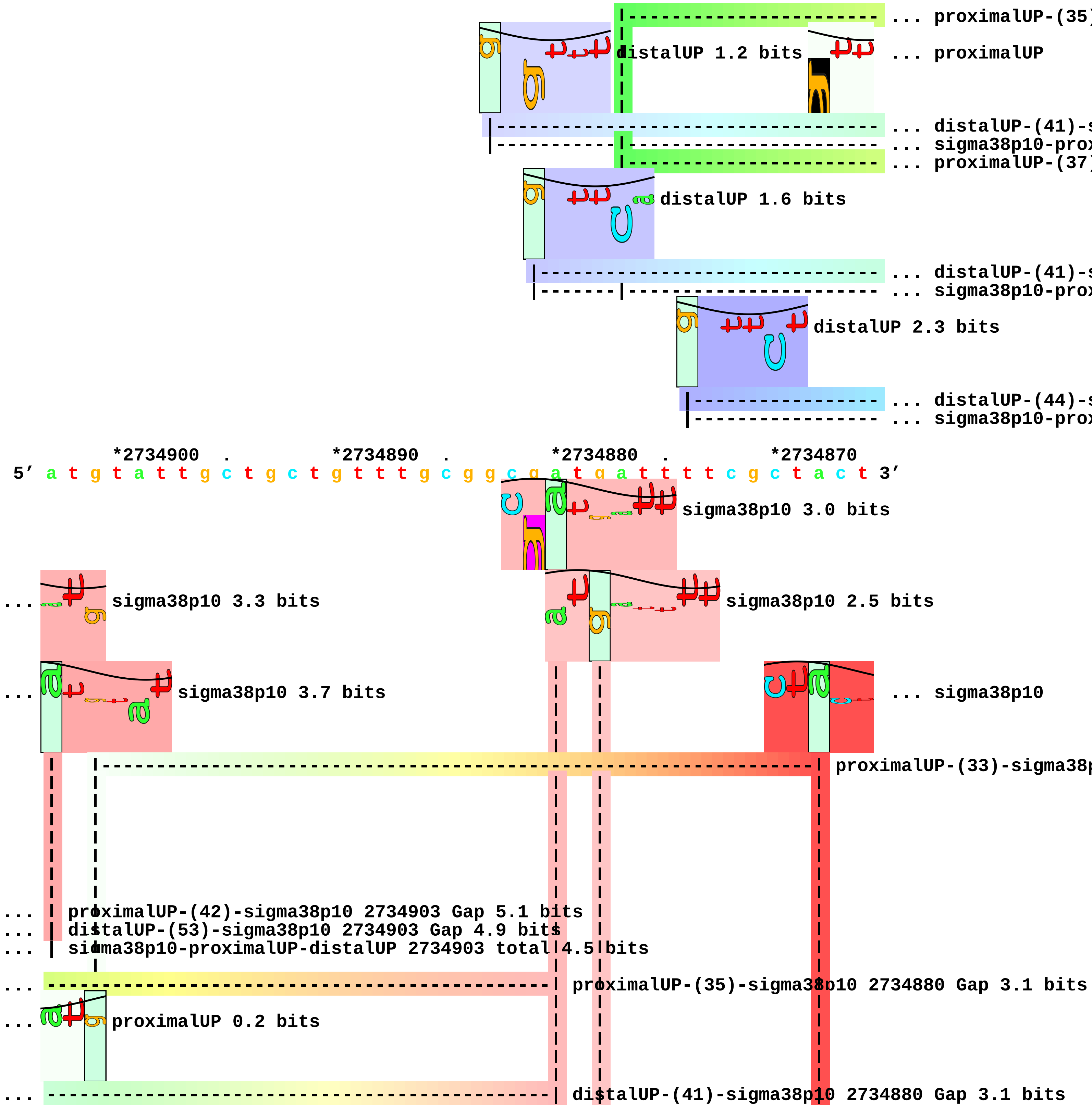
... proximalUP-(35)
distalUP 1.2 bits
... proximalUP
... distalUP-(41)-s
... sigma38p10-prox
... proximalUP-(37)
distalUP 1.6 bits
... distalUP-(41)-s
... sigma38p10-prox
distalUP 2.3 bits
... distalUP-(44)-s
... sigma38p10-prox
*2734900 . *2734890 . *2734880 . *2734870
5′ a t g t a t t g c t g c t g t t t g c g g c g a t g a t t t t c g c t a c t 3′
sigma38p10 3.0 bits
... sigma38p10 3.3 bits
sigma38p10 2.5 bits
... sigma38p10 3.7 bits
... sigma38p10
proximalUP-(33)-sigma38p
... proximalUP-(42)-sigma38p10 2734903 Gap 5.1 bits
... distalUP-(53)-sigma38p10 2734903 Gap 4.9 bits
... sigma38p10-proximalUP-distalUP 2734903 total 4.5 bits
... proximalUP-(35)-sigma38p10 2734880 Gap 3.1 bits
... proximalUP 0.2 bits
... distalUP-(41)-sigma38p10 2734880 Gap 3.1 bits

```
... ----------------------------------------------| sigma38p10-proximalUP-distalUP 2734880 total 2.4
... --------------------------------------------------| proximalUP-(37)-sigma38p10 2734878 Gap 4.5 b
                                                      |                   |
... --------------------------------------------------| distalUP-(41)-sigma38p10 2734878 Gap 3.1 bit
... --------------------------------------------------| sigma38p10-proximalUP-distalUP 2734878 total
                                                                          |
... ----------------------------------------------------------------------| distalUP-(44)-sigma38p10
... ----|-----------------------------------------------------------------| sigma38p10-proximalUP-di

     .         *2734860
 5' t t t t c a 3'
```

... 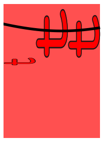 sigma38p10 7.5 bits piece 35, NC_000913, ssrA, config: linear, direction: -, begin: 2753757, end: 2753452

```
         .         *2753750   .         *2753740   .         *2753730   .         *2753720
5' g a g a g g g c t c t a a g c a g g t t a t t a a g c t g c t a a a g c g t 3'
T timestamp:_2018Sep20_11-35-24          proximalUP 4.9 bits
                                            |------------------------- ... proximalUP-(33)
                                 distalUP 4.9 bits
                               |---------------------------------------- ... distalUP-(41)-s
                               |---------------|------------------------- ... sigma38p10-prox

         .         *2753710   .         *2753700   .         *2753690   .
5' a g t t t t c g t c g t t t g c g a c t a t t t t t t g c g g c t t t t t a 3'
                                 sigma38p10 8.7 bits
... --------------------------------------| proximalUP-(33)-sigma38p10 2753699 Gap 4.0 bits
... --------------------------------------| distalUP-(41)-sigma38p10 2753699 Gap 3.1 bits
... --------------------------------------| sigma38p10-proximalUP-distalUP 2753699 total 11.4 bits

   *2753680   .         *2753670   .         *2753660   .         *2753650   .
5' c g a g g c c a a c c g c c c c t c g g c a t g c a c c t t g g g t t t c g 3'

      *2753640   .         *2753630   .         *2753620   .         *2753610
5' c a a a t c c c c g t c g a a t c c a g a a t c a g c c c c a a t g t g t a a 3'

   .         *2753600   .         *2753590   .         *2753580   .         *2753570
5' a g g t a a g t a t a c c a g a t t t a t g a g c g c c a t g a c c a g c c 3'
                                      proximalUP 6.3 bits
                       distalUP 1.8 bits |------------------------------- ... proximalUP-(28)
               |------------------------------------------------------- ... distalUP-(43)-s
               |------------------------------|------------------------- ... sigma38p10-prox
                                              |------------------------- ... proximalUP-(34)
                            distalUP 5.8 bits
                            |------------------------------------------- ... distalUP-(41)-s
                            |-------------|------------------------------ ... sigma38p10-prox

         .         *2753560   .         *2753550   .         *2753540   .         *2753530
5' t c a a t g g c g t t a t c g t t a a a g a t t t a g c a c c c a t g t a g 3'
```

sigma38p10 3.4 bits

... ----------------------| proximalUP-(28)-sigma38p10 2753556 Gap 5.5 bits
... ----------------------| distalUP-(43)-sigma38p10 2753556 Gap 3.6 bits
... ----------------------| sigma38p10-proximalUP-distalUP 2753556 total 2.4 bits
... ------------------------------------| proximalUP-(34)-sigma38p10 2753550 Gap 2.5 bits

sigma38p10 2.5 bits

... ------------------------------------| distalUP-(41)-sigma38p10 2753550 Gap 3.1 bits
... ------------------------------------| sigma38p10-proximalUP-distalUP 2753550 total 9.0 bits

. *2753520 . *2753510 . *2753500 .
5′ c c t g a t t t t t a t t c g a t t a a g c a a t g g g a t g g c a a c a t 3′

proximalUP 4.9 bits

|------------------------------------ ... proximalUP-(41)

distalUP 6.4 bits

|------------------------------------------------ ... distalUP-(47)-s
|-----------|------------------------------------ ... sigma38p10-prox

*2753490 . *2753480 . *2753470 . *2753460 .
5′ t t g t g t c g g a t g t g a t a g c c a a t a a g a t g t t c a t t c g c 3′

sigma38p10 3.7 bits

... ---------------------------------------------| proximalUP-(41)-sigma38p10 2753468 Gap 5.5 bits
... ---------------------------------------------| distalUP-(47)-sigma38p10 2753468 Gap 6.5 bits
... ---------------------------------------------| sigma38p10-proximalUP-distalUP 2753468 total 3.2

5′ g c 3′

piece 36, NC_000913, yfjJ, config: linear, direction: -, begin: 2759049, end: 2758250

```
             .          *2759040   .          *2759030   .          *2759020   .
5' g c g g g t a a t t a a c t a a c t t c c c c g a g c t a t c t a t a g g g 3'
```

proximalUP 5.4 bits

proximalUP 1.5 bits

... proximalUP-(35)

distalUP 5.0 bits

... proximalUP-(38)

... distalUP-(43)-s

... sigma38p10-prox

proximalUP 2.0 bits

... proximalUP-(33)

... distalUP-(53)-s

... sigma38p10-prox

T timestamp:_2018Sep20_11-35-24

distalUP 1.5 bits

... distalUP-(42)-s

... sigma38p10-prox

```
        *2759010   .          *2759000   .          *2758990   .          *2758980   .
5' a t g c c a a g t g c a c t g t a c c a a g c t g t c g t t a t c a t t g t 3'
```

sigma38p10 1.2 bits

sigma38p10 5.7 bits

... proximalUP-(35)-sigma38p10 2759002 Gap 3.1 bits

... proximalUP-(38)-sigma38p10 2758992 Gap 3.2 bits

... distalUP-(43)-sigma38p10 2759002 Gap 3.6 bits

... sigma38p10-proximalUP-distalUP 2759002 total 5.0 bits

... proximalUP-(33)-sigma38p10 2758981

... distalUP-(53)-sigma38p10 2758992 Gap 4.9 bits

... sigma38p10-proximalUP-distalUP 2758992 total 2.6 bits

sigma38p10 3.7 bits

... distalUP-(42)-sigma38p10 2758981 G

... sigma38p10-proximalUP-distalUP 275

```
           *2758970   .          *2758960   .          *2758950   .          *2758940
5' a c g t a a c g t g t t a c g g t t t a a g t c g t a a t c a c c t a a a t 3'

   .          *2758930   .          *2758920   .          *2758910   .          *2758900
5' g a t a a t a a g c a t c c t g a t t g a a a c a c a g t a t c a a a t g g 3'
```

distalUP 7.1 bits

proximalUP 3.3 bits

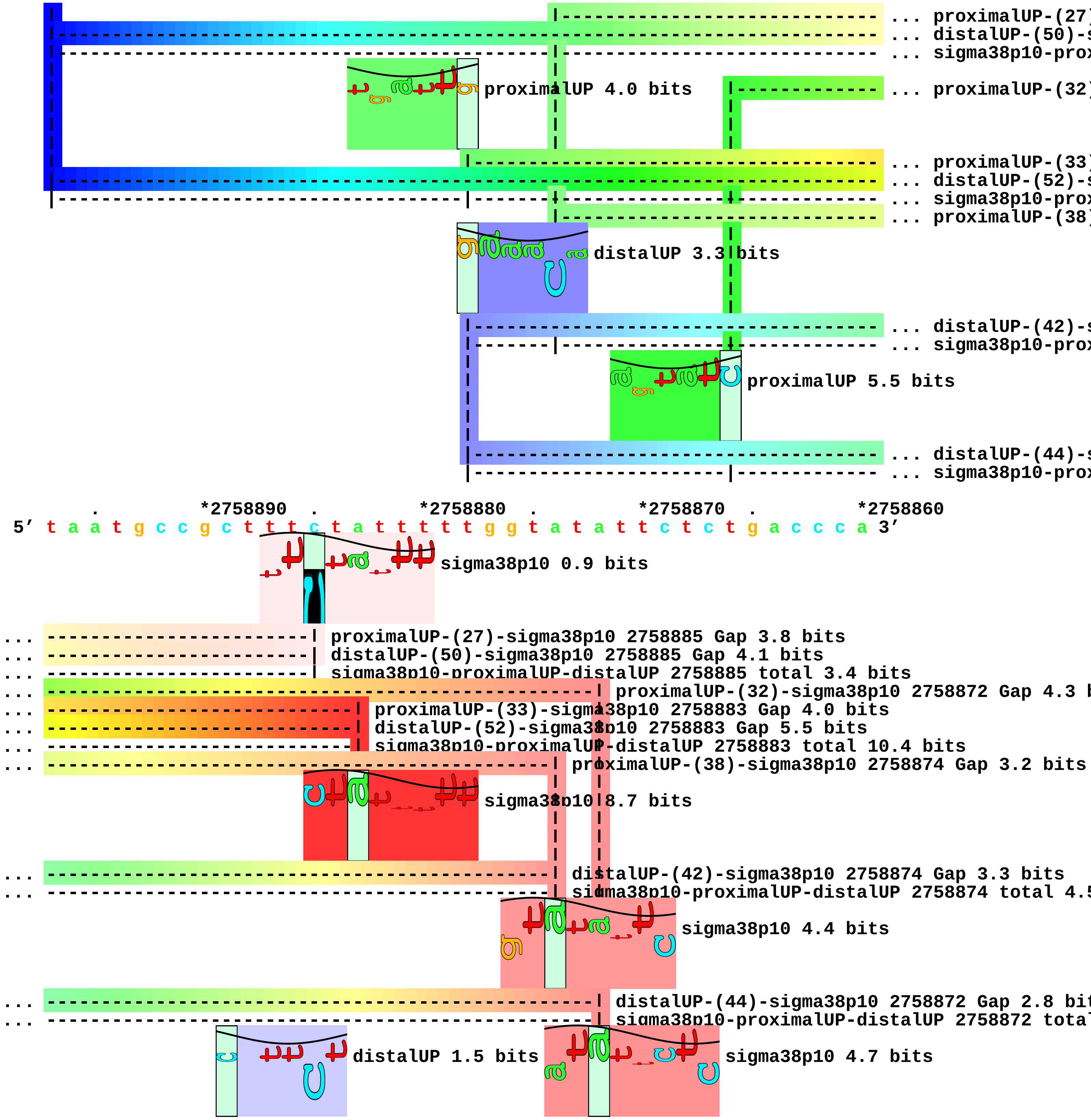
... proximalUP-(27)
... distalUP-(50)-s
... sigma38p10-prox
proximalUP 4.0 bits
... proximalUP-(32)
... proximalUP-(33)
... distalUP-(52)-s
... sigma38p10-prox
... proximalUP-(38)
distalUP 3.3 bits
... distalUP-(42)-s
... sigma38p10-prox
proximalUP 5.5 bits
... distalUP-(44)-s
... sigma38p10-prox
. *2758890 . *2758880 . *2758870 . *2758860
5′ t a a t g c c g c t t t c t a t t t t t g g t a t a t t c t c t g a c c c a 3′
sigma38p10 0.9 bits
... proximalUP-(27)-sigma38p10 2758885 Gap 3.8 bits
... distalUP-(50)-sigma38p10 2758885 Gap 4.1 bits
... sigma38p10-proximalUP-distalUP 2758885 total 3.4 bits
... proximalUP-(32)-sigma38p10 2758872 Gap 4.3 b
... proximalUP-(33)-sigma38p10 2758883 Gap 4.0 bits
... distalUP-(52)-sigma38p10 2758883 Gap 5.5 bits
... sigma38p10-proximalUP-distalUP 2758883 total 10.4 bits
... proximalUP-(38)-sigma38p10 2758874 Gap 3.2 bits
sigma38p10 8.7 bits
... distalUP-(42)-sigma38p10 2758874 Gap 3.3 bits
... sigma38p10-proximalUP-distalUP 2758874 total 4.5
sigma38p10 4.4 bits
... distalUP-(44)-sigma38p10 2758872 Gap 2.8 bit
... sigma38p10-proximalUP-distalUP 2758872 total
distalUP 1.5 bits
sigma38p10 4.7 bits

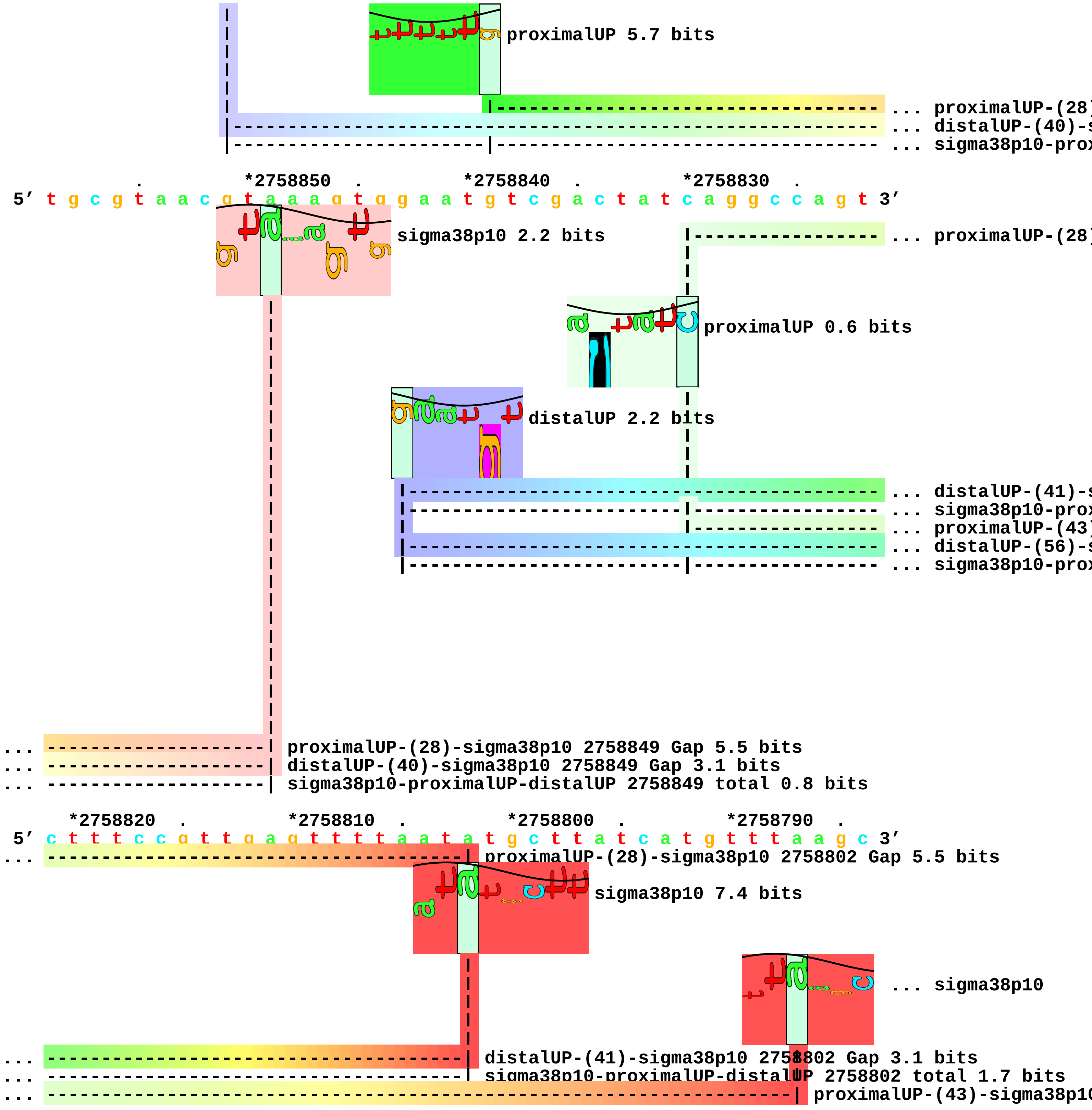
proximalUP 5.7 bits
... proximalUP-(28)
... distalUP-(40)-s
... sigma38p10-prox
. *2758850 . *2758840 . *2758830 .
5' t g c g t a a c g t a a a g t g g a a t g t c g a c t a t c a g g c c a g t 3'
sigma38p10 2.2 bits
... proximalUP-(28)
proximalUP 0.6 bits
distalUP 2.2 bits
... distalUP-(41)-s
... sigma38p10-prox
... proximalUP-(43)
... distalUP-(56)-s
... sigma38p10-prox
... proximalUP-(28)-sigma38p10 2758849 Gap 5.5 bits
... distalUP-(40)-sigma38p10 2758849 Gap 3.1 bits
... sigma38p10-proximalUP-distalUP 2758849 total 0.8 bits
*2758820 . *2758810 . *2758800 . *2758790 .
5' c t t t c c g t t g a g t t t t a a t a t g c t t a t c a t g t t t a a g c 3'
... proximalUP-(28)-sigma38p10 2758802 Gap 5.5 bits
sigma38p10 7.4 bits
... sigma38p10
... distalUP-(41)-sigma38p10 2758802 Gap 3.1 bits
... sigma38p10-proximalUP-distalUP 2758802 total 1.7 bits
... proximalUP-(43)-sigma38p10

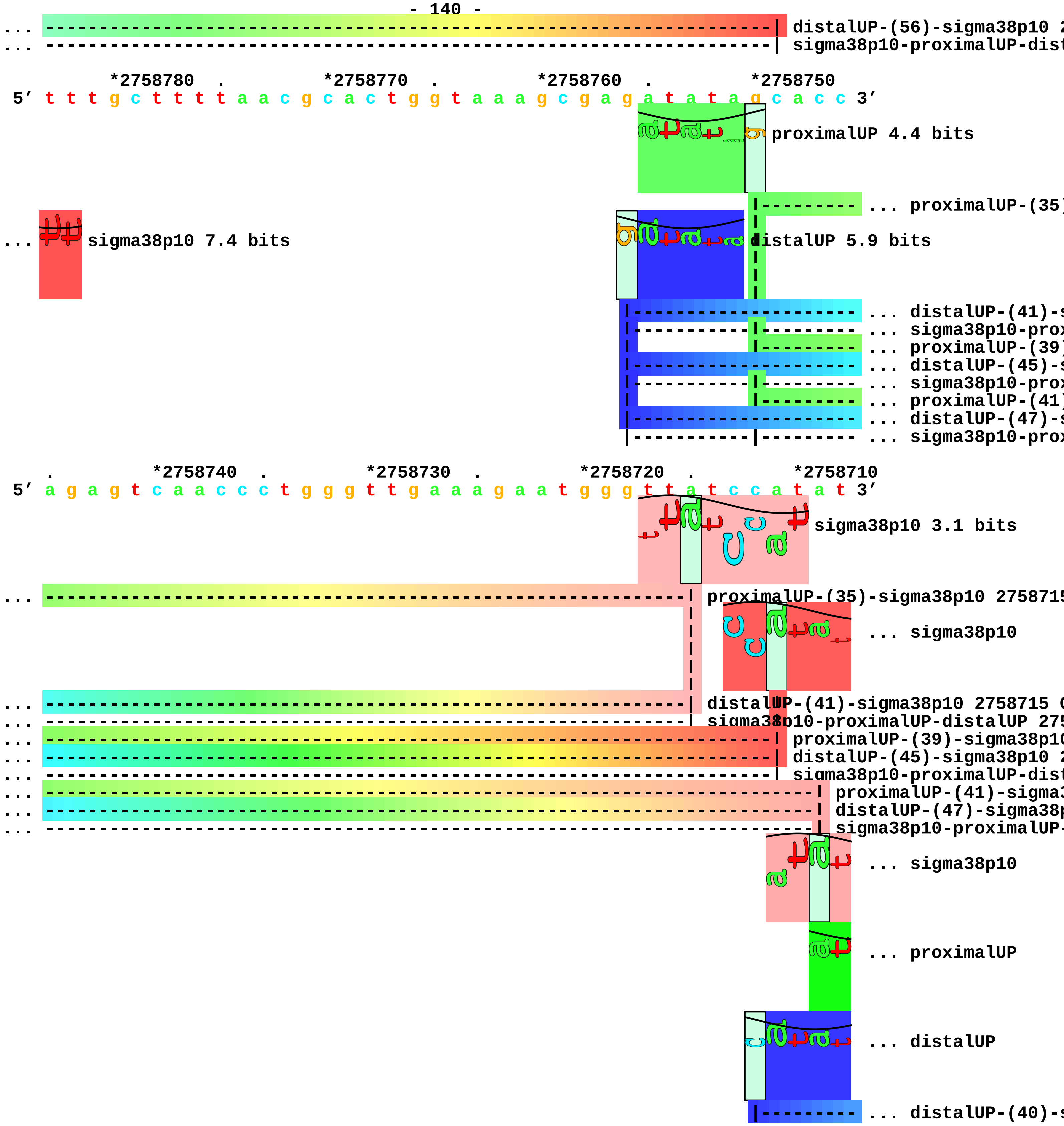
distalUP-(56)-sigma38p10
sigma38p10-proximalUP-dist
*2758780
*2758770
*2758760
*2758750
5′ t t t g c t t t t a a c g c a c t g g t a a a g c g a g a t a t a g c a c c 3′
proximalUP 4.4 bits
... proximalUP-(35)
... sigma38p10 7.4 bits
distalUP 5.9 bits
... distalUP-(41)-s
... sigma38p10-prox
... proximalUP-(39)
... distalUP-(45)-s
... sigma38p10-prox
... proximalUP-(41)
... distalUP-(47)-s
... sigma38p10-prox
*2758740
*2758730
*2758720
*2758710
5′ a g a g t c a a c c c t g g g t t g a a a g a a t g g g t t a t c c a t a t 3′
sigma38p10 3.1 bits
proximalUP-(35)-sigma38p10 275871
... sigma38p10
distalUP-(41)-sigma38p10 2758715
sigma38p10-proximalUP-distalUP 275
proximalUP-(39)-sigma38p10
distalUP-(45)-sigma38p10 2
sigma38p10-proximalUP-dist
proximalUP-(41)-sigma3
distalUP-(47)-sigma38p
sigma38p10-proximalUP-
... sigma38p10
... proximalUP
... distalUP
... distalUP-(40)-s

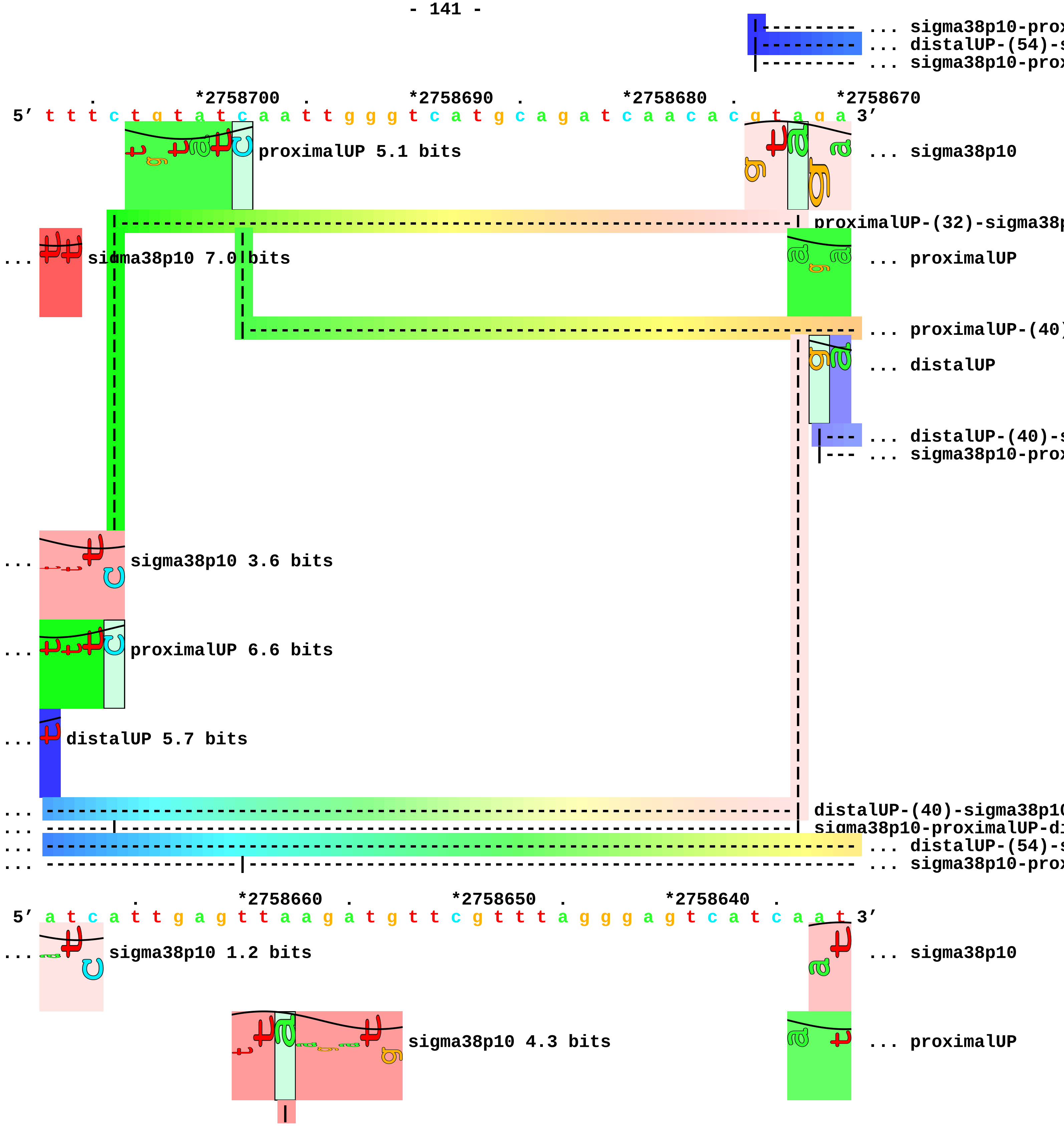
... sigma38p10-prox
... distalUP-(54)-s
... sigma38p10-prox
. *2758700 . *2758690 . *2758680 . *2758670
5′ t t t c t g t a t c a a t t g g g t c a t g c a g a t c a a c a c g t a g a 3′
proximalUP 5.1 bits
... sigma38p10
proximalUP-(32)-sigma38p
... sigma38p10 7.0 bits
... proximalUP
... proximalUP-(40)
... distalUP
... distalUP-(40)-s
... sigma38p10-prox
... sigma38p10 3.6 bits
... proximalUP 6.6 bits
... distalUP 5.7 bits
... distalUP-(40)-sigma38p10
... sigma38p10-proximalUP-di
... distalUP-(54)-s
... sigma38p10-prox
. *2758660 . *2758650 . *2758640 .
5′ a t c a t t g a g t t a a g a t g t t c g t t t a g g g a g t c a t c a a t 3′
... sigma38p10 1.2 bits
... sigma38p10
sigma38p10 4.3 bits
... proximalUP

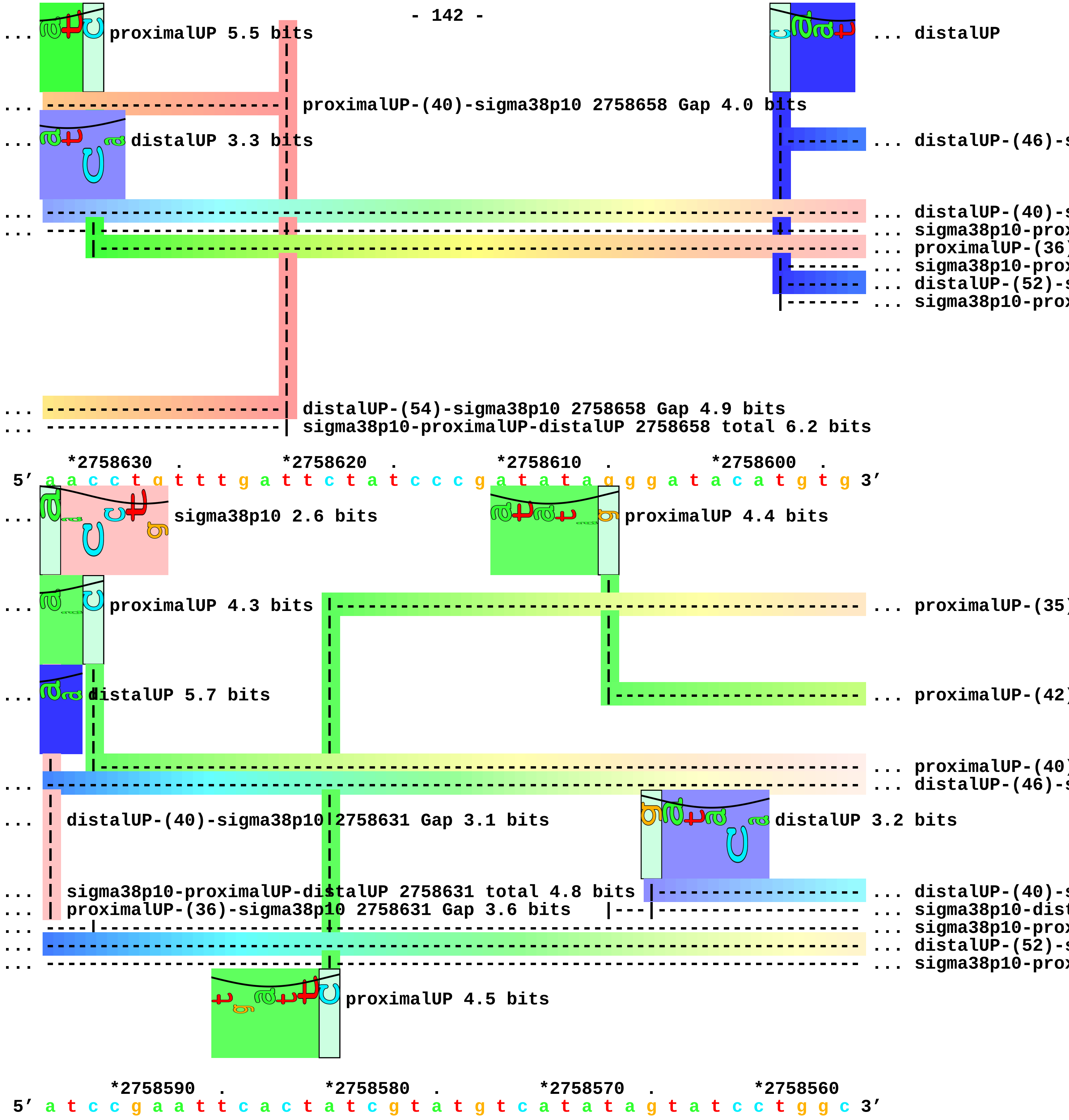
... proximalUP 5.5 bits
... distalUP
... proximalUP-(40)-sigma38p10 2758658 Gap 4.0 bits
... distalUP 3.3 bits
... distalUP-(46)-s
... distalUP-(40)-s
... sigma38p10-prox
... proximalUP-(36)
... sigma38p10-prox
... distalUP-(52)-s
... sigma38p10-prox
... distalUP-(54)-sigma38p10 2758658 Gap 4.9 bits
... sigma38p10-proximalUP-distalUP 2758658 total 6.2 bits
*2758630 . *2758620 . *2758610 . *2758600 .
5′ a a c c t g t t t g a t t c t a t c c c g a t a t a g g g a t a c a t g t g 3′
... sigma38p10 2.6 bits
proximalUP 4.4 bits
... proximalUP 4.3 bits
... proximalUP-(35)
... distalUP 5.7 bits
... proximalUP-(42)
... proximalUP-(40)
... distalUP-(46)-s
... distalUP-(40)-sigma38p10 2758631 Gap 3.1 bits
distalUP 3.2 bits
... sigma38p10-proximalUP-distalUP 2758631 total 4.8 bits
... distalUP-(40)-s
... proximalUP-(36)-sigma38p10 2758631 Gap 3.6 bits
... sigma38p10-dist
... sigma38p10-prox
... distalUP-(52)-s
... sigma38p10-prox
proximalUP 4.5 bits
*2758590 . *2758580 . *2758570 . *2758560
5′ a t c c g a a t t c a c t a t c g t a t g t c a t a t a g t a t c c t g g c 3′

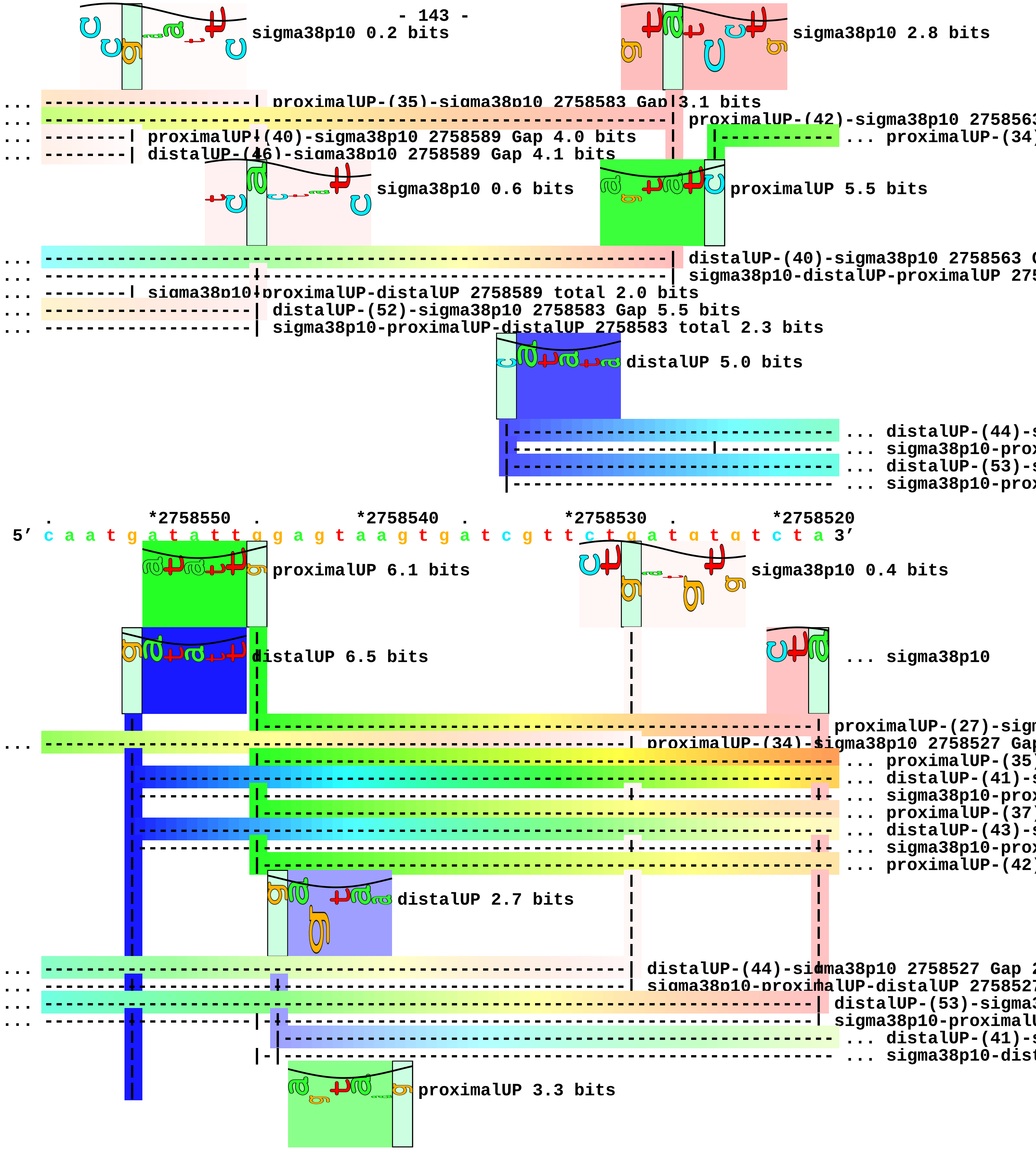
sigma38p10 0.2 bits
sigma38p10 2.8 bits
... proximalUP-(35)-sigma38p10 2758583 Gap 3.1 bits
... proximalUP-(42)-sigma38p10 275856
... proximalUP-(40)-sigma38p10 2758589 Gap 4.0 bits
... proximalUP-(34)
... distalUP-(46)-sigma38p10 2758589 Gap 4.1 bits
sigma38p10 0.6 bits
proximalUP 5.5 bits
... distalUP-(40)-sigma38p10 2758563
... sigma38p10-distalUP-proximalUP 275
... sigma38p10-proximalUP-distalUP 2758589 total 2.0 bits
... distalUP-(52)-sigma38p10 2758583 Gap 5.5 bits
... sigma38p10-proximalUP-distalUP 2758583 total 2.3 bits
distalUP 5.0 bits
... distalUP-(44)-s
... sigma38p10-prox
... distalUP-(53)-s
... sigma38p10-prox
. *2758550 . *2758540 . *2758530 . *2758520
5′ c a a t g a t a t t g g a g t a a g t g a t c g t t c t g a t g t g t c t a 3′
proximalUP 6.1 bits
sigma38p10 0.4 bits
distalUP 6.5 bits
... sigma38p10
proximalUP-(27)-sign
... proximalUP-(34)-sigma38p10 2758527 Gap
... proximalUP-(35)
... distalUP-(41)-s
... sigma38p10-prox
... proximalUP-(37)
... distalUP-(43)-s
... sigma38p10-prox
... proximalUP-(42)
distalUP 2.7 bits
... distalUP-(44)-sigma38p10 2758527 Gap 2
... sigma38p10-proximalUP-distalUP 275852
... distalUP-(53)-sigma3
... sigma38p10-proximalU
... distalUP-(41)-s
... sigma38p10-dist
proximalUP 3.3 bits

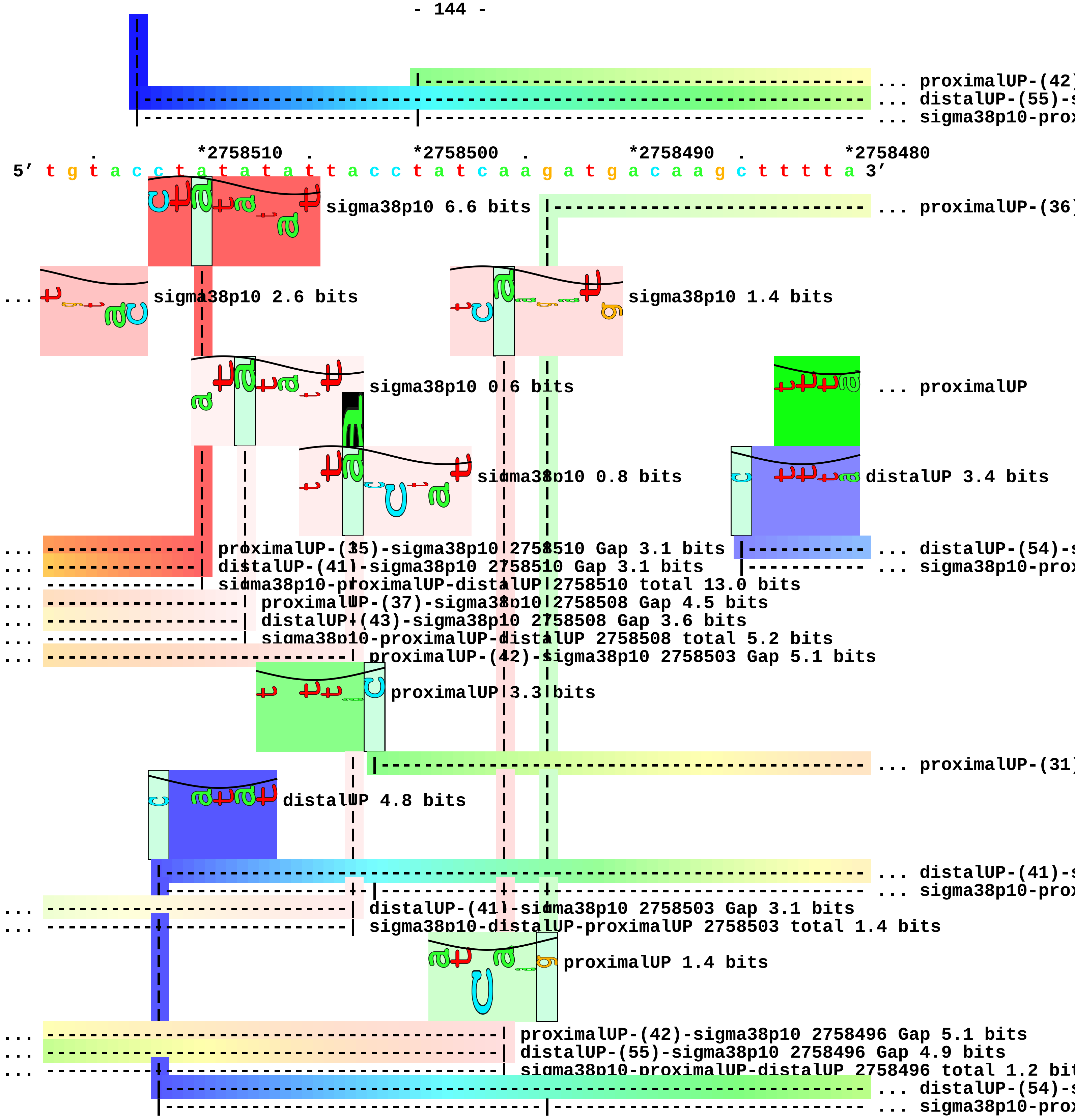
... proximalUP-(42)
... distalUP-(55)-s
... sigma38p10-prox
*2758510
*2758500
*2758490
*2758480
5′ t g t a c c t a t a t a t t a c c t a t c a a g a t g a c a a g c t t t t a 3′
sigma38p10 6.6 bits
... proximalUP-(36)
sigma38p10 2.6 bits
sigma38p10 1.4 bits
sigma38p10 0.6 bits
... proximalUP
sigma38p10 0.8 bits
distalUP 3.4 bits
... proximalUP-(35)-sigma38p10 2758510 Gap 3.1 bits
... distalUP-(41)-sigma38p10 2758510 Gap 3.1 bits
... sigma38p10-proximalUP-distalUP 2758510 total 13.0 bits
... distalUP-(54)-s
... sigma38p10-prox
... proximalUP-(37)-sigma38p10 2758508 Gap 4.5 bits
... distalUP-(43)-sigma38p10 2758508 Gap 3.6 bits
... sigma38p10-proximalUP-distalUP 2758508 total 5.2 bits
... proximalUP-(42)-sigma38p10 2758503 Gap 5.1 bits
proximalUP 3.3 bits
... proximalUP-(31)
distalUP 4.8 bits
... distalUP-(41)-s
... sigma38p10-prox
... distalUP-(41)-sigma38p10 2758503 Gap 3.1 bits
... sigma38p10-distalUP-proximalUP 2758503 total 1.4 bits
proximalUP 1.4 bits
... proximalUP-(42)-sigma38p10 2758496 Gap 5.1 bits
... distalUP-(55)-sigma38p10 2758496 Gap 4.9 bits
... sigma38p10-proximalUP-distalUP 2758496 total 1.2 bit
... distalUP-(54)-s
... sigma38p10-prox

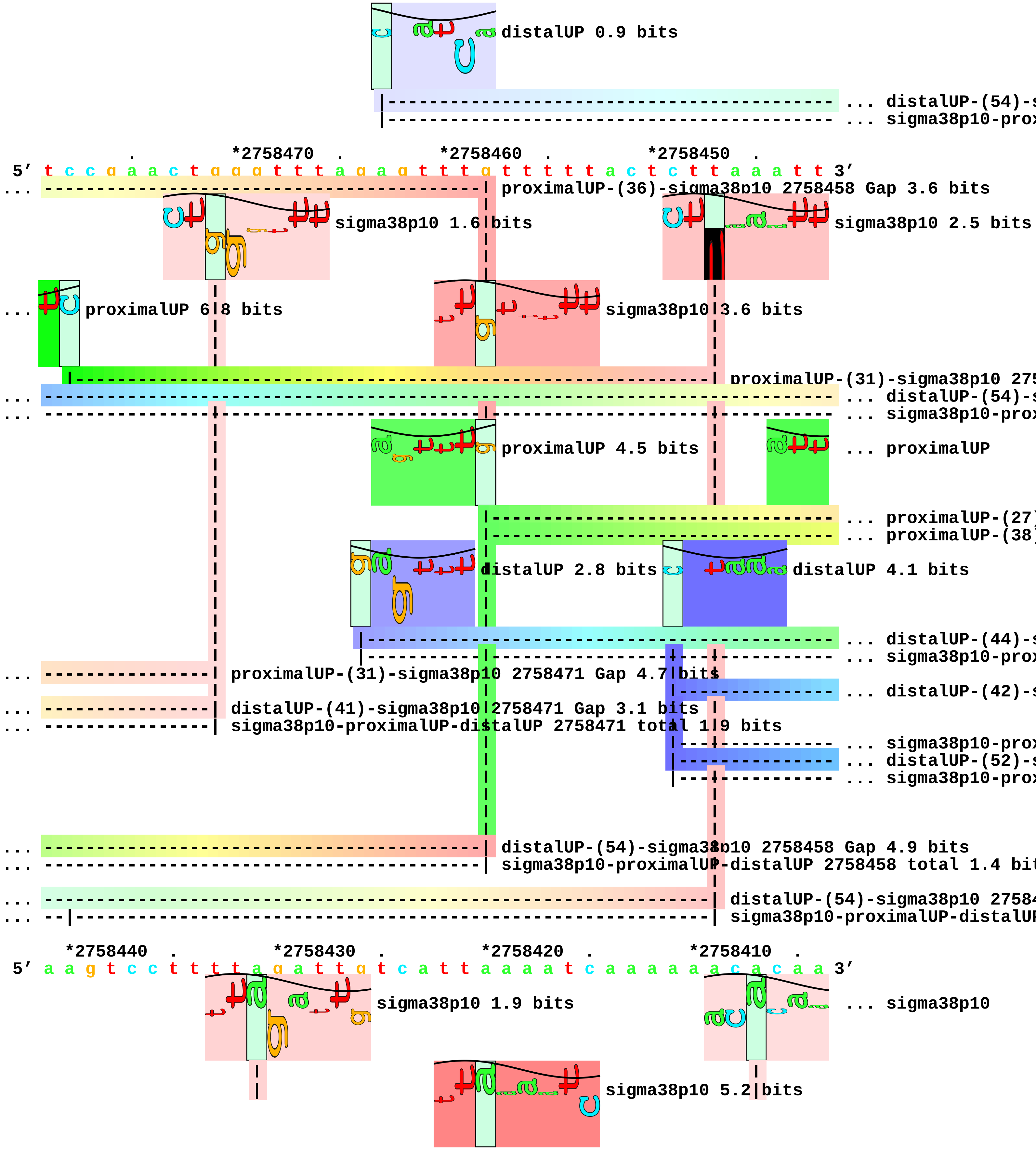
distalUP 0.9 bits
... distalUP-(54)-s
... sigma38p10-prox
*2758470
*2758460
*2758450
5′ t c c g a a c t g g g t t t a g a g t t t g t t t t t a c t c t t a a a t t 3′
proximalUP-(36)-sigma38p10 2758458 Gap 3.6 bits
sigma38p10 1.6 bits
sigma38p10 2.5 bits
proximalUP 6.8 bits
sigma38p10 3.6 bits
proximalUP-(31)-sigma38p10 275
... distalUP-(54)-s
... sigma38p10-prox
proximalUP 4.5 bits
... proximalUP
... proximalUP-(27)
... proximalUP-(38)
distalUP 2.8 bits
distalUP 4.1 bits
... distalUP-(44)-s
... sigma38p10-prox
proximalUP-(31)-sigma38p10 2758471 Gap 4.7 bits
... distalUP-(42)-s
distalUP-(41)-sigma38p10 2758471 Gap 3.1 bits
sigma38p10-proximalUP-distalUP 2758471 total 1.9 bits
... sigma38p10-prox
... distalUP-(52)-s
... sigma38p10-prox
distalUP-(54)-sigma38p10 2758458 Gap 4.9 bits
sigma38p10-proximalUP-distalUP 2758458 total 1.4 bit
distalUP-(54)-sigma38p10 27584
sigma38p10-proximalUP-distalUP
*2758440
*2758430
*2758420
*2758410
5′ a a g t c c t t t t a g a t t g t c a t t a a a a t c a a a a a a c a c a a 3′
sigma38p10 1.9 bits
... sigma38p10
sigma38p10 5.2 bits

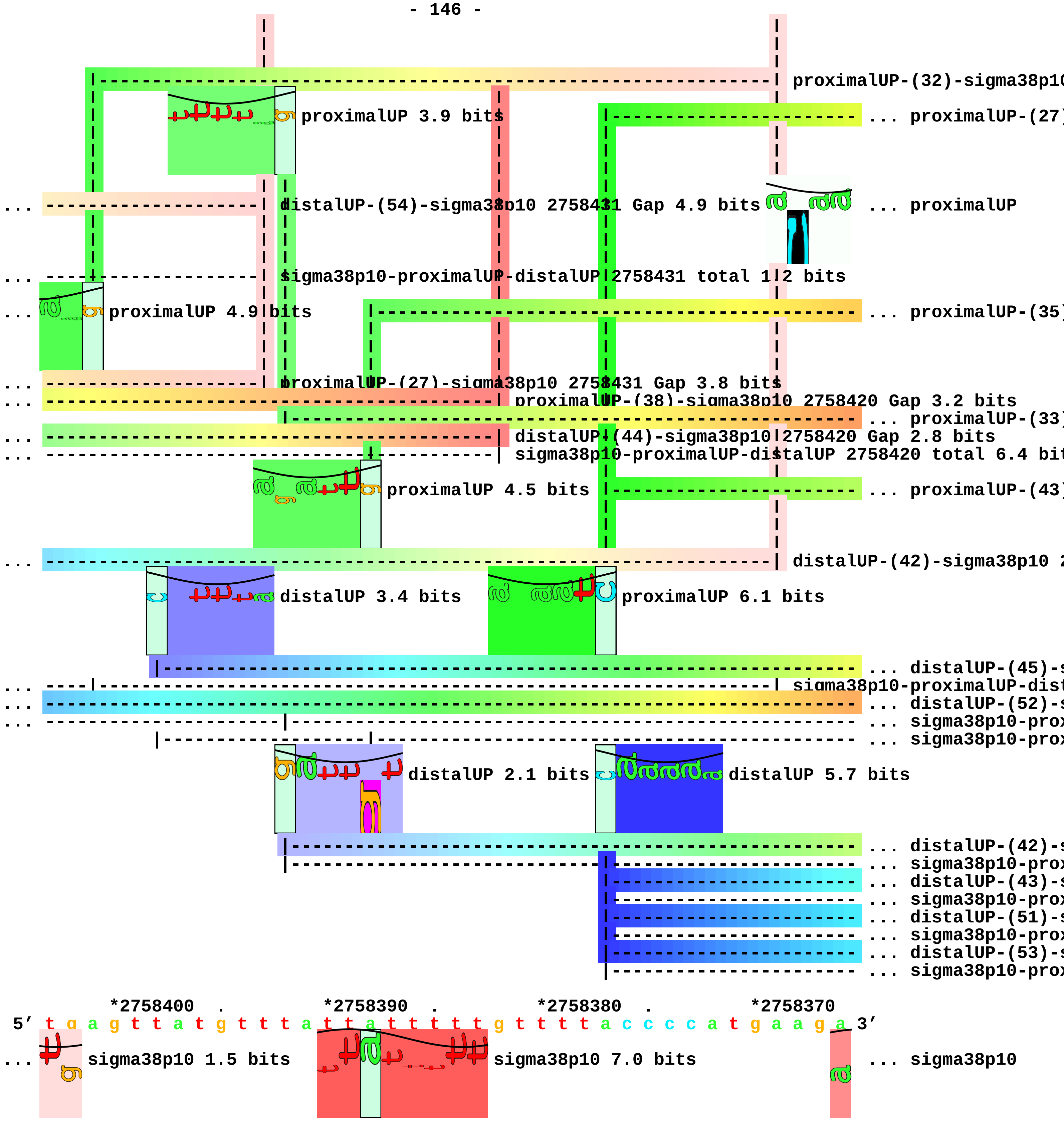

proximalUP-(32)-sigma38p10
proximalUP 3.9 bits
... proximalUP-(27)
... distalUP-(54)-sigma38p10 2758431 Gap 4.9 bits
... proximalUP
... sigma38p10-proximalUP-distalUP 2758431 total 1.2 bits
... proximalUP 4.9 bits
... proximalUP-(35)
... proximalUP-(27)-sigma38p10 2758431 Gap 3.8 bits
... proximalUP-(38)-sigma38p10 2758420 Gap 3.2 bits
... proximalUP-(33)
... distalUP-(44)-sigma38p10 2758420 Gap 2.8 bits
... sigma38p10-proximalUP-distalUP 2758420 total 6.4 bit
proximalUP 4.5 bits
... proximalUP-(43)
... distalUP-(42)-sigma38p10 2
distalUP 3.4 bits
proximalUP 6.1 bits
... distalUP-(45)-s
... sigma38p10-proximalUP-dist
... distalUP-(52)-s
... sigma38p10-prox
... sigma38p10-prox
distalUP 2.1 bits
distalUP 5.7 bits
... distalUP-(42)-s
... sigma38p10-prox
... distalUP-(43)-s
... sigma38p10-prox
... distalUP-(51)-s
... sigma38p10-prox
... distalUP-(53)-s
... sigma38p10-prox
*2758400 . *2758390 . *2758380 . *2758370
5' t g a g t t a t g t t t a t t a t t t t t g t t t t a c c c c a t g a a g a 3'
... sigma38p10 1.5 bits
sigma38p10 7.0 bits
... sigma38p10

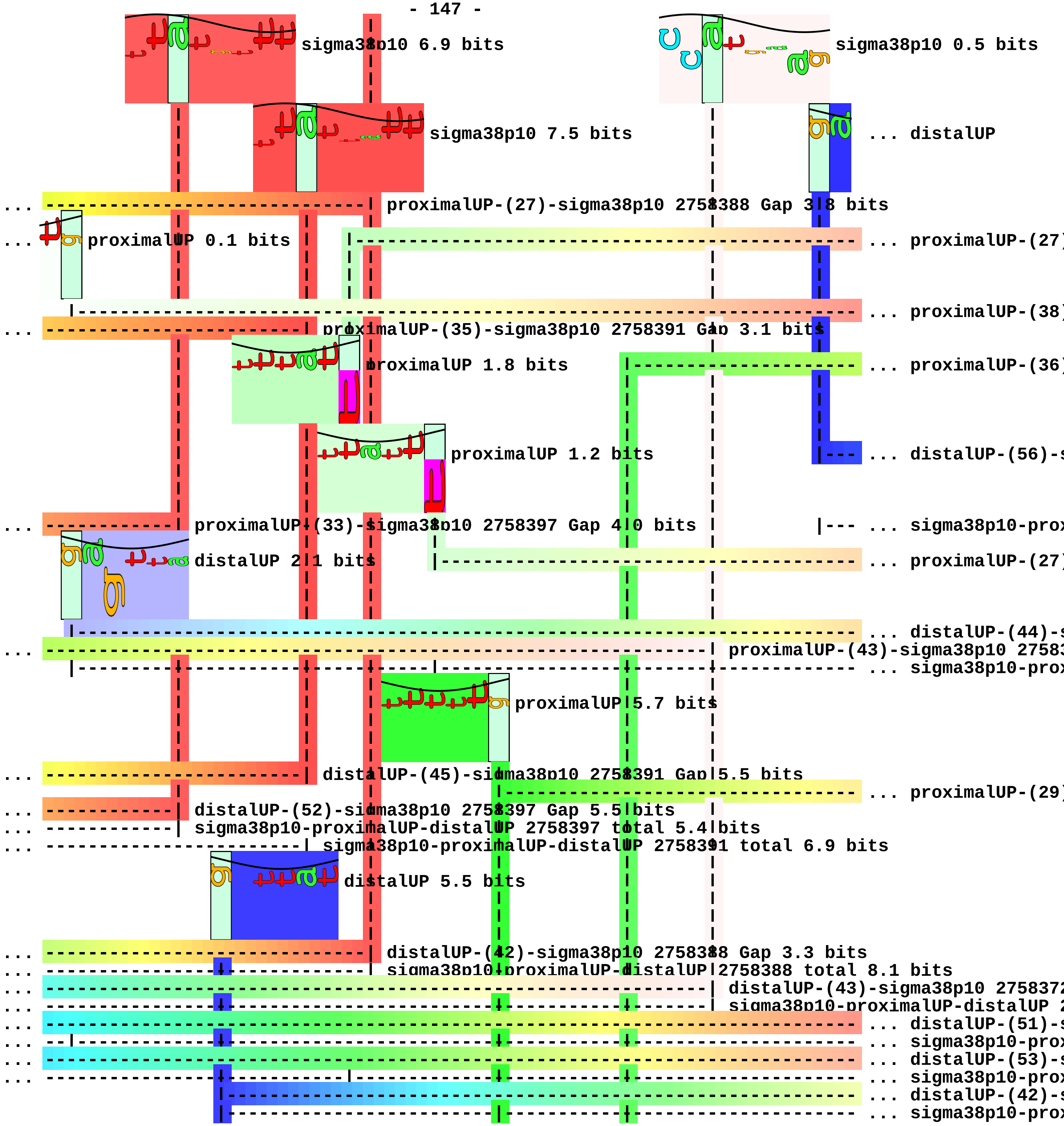
sigma38p10 6.9 bits
sigma38p10 0.5 bits
sigma38p10 7.5 bits
... distalUP
... proximalUP-(27)-sigma38p10 2758388 Gap 3.8 bits
... proximalUP 0.1 bits
... proximalUP-(27
... proximalUP-(38
... proximalUP-(35)-sigma38p10 2758391 Gap 3.1 bits
proximalUP 1.8 bits
... proximalUP-(36
proximalUP 1.2 bits
... distalUP-(56)-s
... proximalUP-(33)-sigma38p10 2758397 Gap 4.0 bits
... sigma38p10-prox
distalUP 2.1 bits
... proximalUP-(27
... distalUP-(44)-s
... proximalUP-(43)-sigma38p10 2758
... sigma38p10-prox
proximalUP 5.7 bits
... distalUP-(45)-sigma38p10 2758391 Gap 5.5 bits
... proximalUP-(29
... distalUP-(52)-sigma38p10 2758397 Gap 5.5 bits
... sigma38p10-proximalUP-distalUP 2758397 total 5.4 bits
... sigma38p10-proximalUP-distalUP 2758391 total 6.9 bits
distalUP 5.5 bits
... distalUP-(42)-sigma38p10 2758388 Gap 3.3 bits
... sigma38p10-proximalUP-distalUP 2758388 total 8.1 bits
... distalUP-(43)-sigma38p10 275837
... sigma38p10-proximalUP-distalUP 2
... distalUP-(51)-s
... sigma38p10-prox
... distalUP-(53)-s
... sigma38p10-prox
... distalUP-(42)-s
... sigma38p10-prox

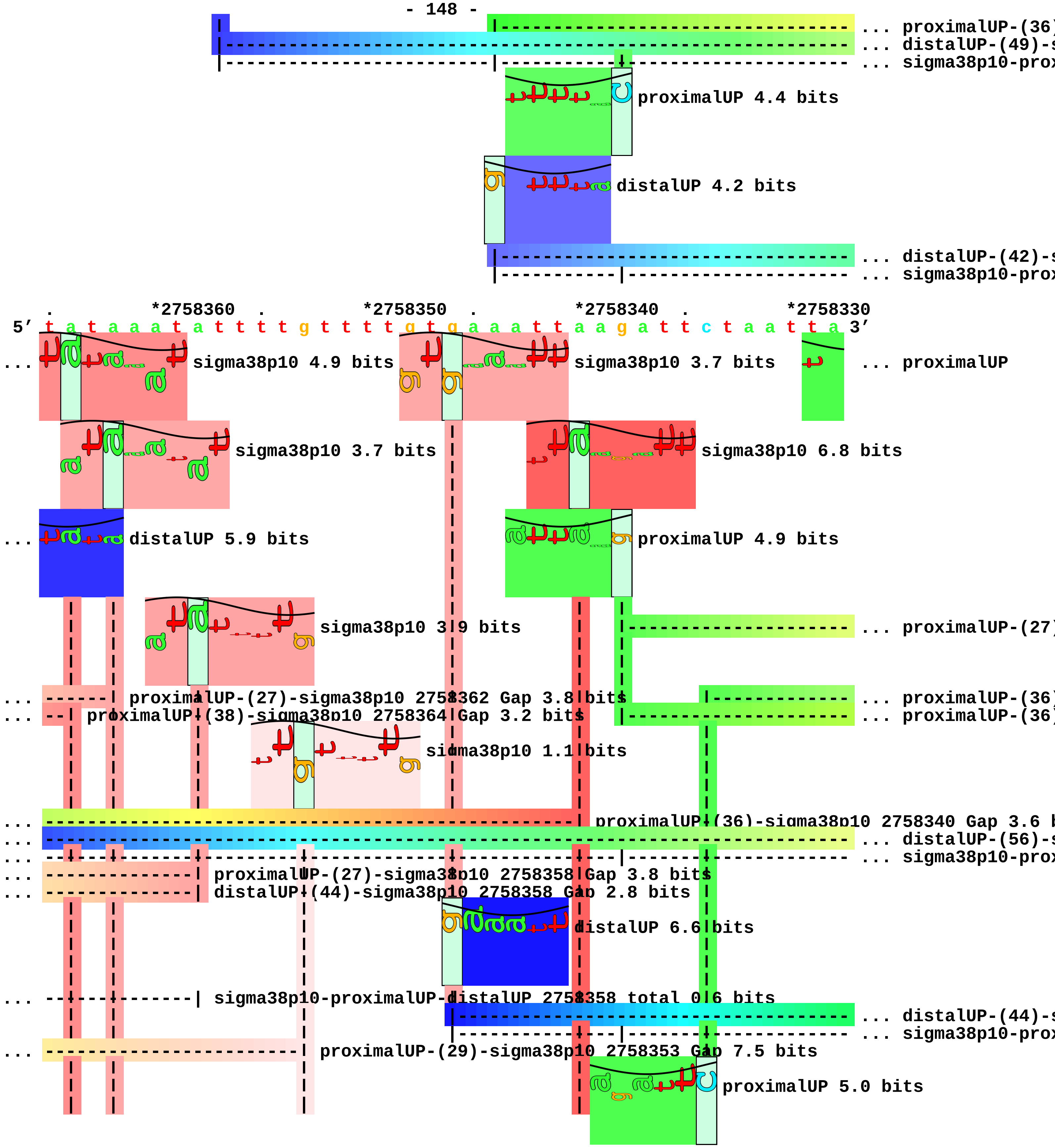
... proximalUP-(36)
... distalUP-(49)-s
... sigma38p10-prox
proximalUP 4.4 bits
distalUP 4.2 bits
... distalUP-(42)-s
... sigma38p10-prox
*2758360
*2758350
*2758340
*2758330
5' t a t a a a t a t t t t g t t t t g t g a a a t t a a g a t t c t a a t t a 3'
... sigma38p10 4.9 bits
sigma38p10 3.7 bits
... proximalUP
sigma38p10 3.7 bits
sigma38p10 6.8 bits
... distalUP 5.9 bits
proximalUP 4.9 bits
sigma38p10 3.9 bits
... proximalUP-(27)
... proximalUP-(27)-sigma38p10 2758362 Gap 3.8 bits
... proximalUP-(38)-sigma38p10 2758364 Gap 3.2 bits
... proximalUP-(36)
... proximalUP-(36)
sigma38p10 1.1 bits
... proximalUP-(36)-sigma38p10 2758340 Gap 3.6 b
... distalUP-(56)-s
... sigma38p10-prox
... proximalUP-(27)-sigma38p10 2758358 Gap 3.8 bits
... distalUP-(44)-sigma38p10 2758358 Gap 2.8 bits
distalUP 6.6 bits
... sigma38p10-proximalUP-distalUP 2758358 total 0.6 bits
... distalUP-(44)-s
... sigma38p10-prox
... proximalUP-(29)-sigma38p10 2758353 Gap 7.5 bits
proximalUP 5.0 bits

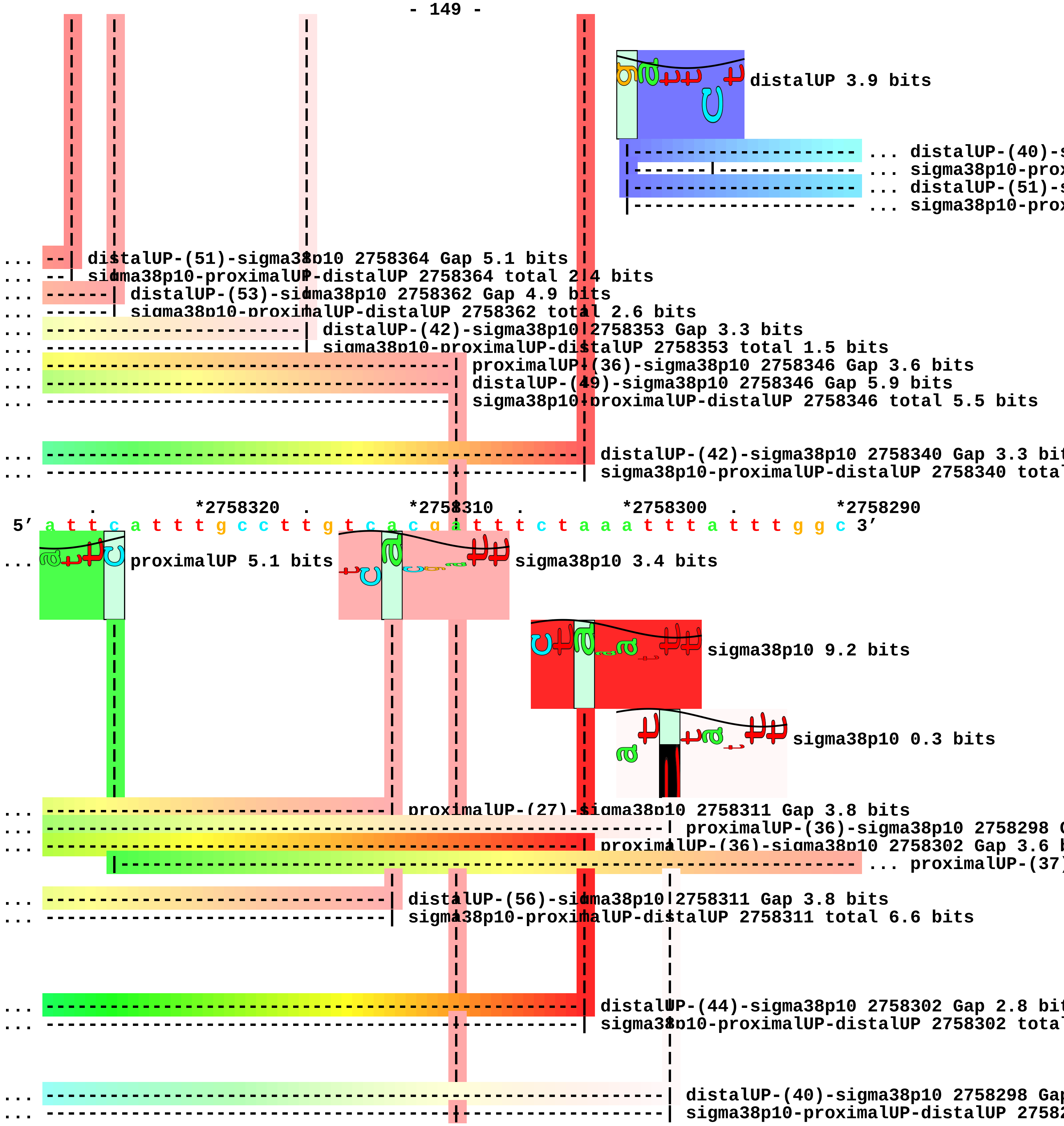
distalUP 3.9 bits
... distalUP-(40)-s
... sigma38p10-prox
... distalUP-(51)-s
... sigma38p10-prox
... distalUP-(51)-sigma38p10 2758364 Gap 5.1 bits
... sigma38p10-proximalUP-distalUP 2758364 total 2.4 bits
... distalUP-(53)-sigma38p10 2758362 Gap 4.9 bits
... sigma38p10-proximalUP-distalUP 2758362 total 2.6 bits
... distalUP-(42)-sigma38p10 2758353 Gap 3.3 bits
... sigma38p10-proximalUP-distalUP 2758353 total 1.5 bits
... proximalUP-(36)-sigma38p10 2758346 Gap 3.6 bits
... distalUP-(49)-sigma38p10 2758346 Gap 5.9 bits
... sigma38p10-proximalUP-distalUP 2758346 total 5.5 bits
... distalUP-(42)-sigma38p10 2758340 Gap 3.3 bit
... sigma38p10-proximalUP-distalUP 2758340 tota
. *2758320 . *2758310 . *2758300 . *2758290
5′ a t t c a t t t g c c t t g t c a c g a t t t c t a a a t t t a t t t g g c 3′
... proximalUP 5.1 bits
sigma38p10 3.4 bits
sigma38p10 9.2 bits
sigma38p10 0.3 bits
... proximalUP-(27)-sigma38p10 2758311 Gap 3.8 bits
... proximalUP-(36)-sigma38p10 2758298 G
... proximalUP-(36)-sigma38p10 2758302 Gap 3.6 b
... proximalUP-(37)
... distalUP-(56)-sigma38p10 2758311 Gap 3.8 bits
... sigma38p10-proximalUP-distalUP 2758311 total 6.6 bits
... distalUP-(44)-sigma38p10 2758302 Gap 2.8 bit
... sigma38p10-proximalUP-distalUP 2758302 tota
... distalUP-(40)-sigma38p10 2758298 Gap
... sigma38p10-proximalUP-distalUP 27582

- 150 -

... ---------------------------------------------------------------------------- ... distalUP-(51)-s

... ------|--------------------------------------------------------------------- ... sigma38p10-prox

. *2758280 . *2758270 . *2758260 .

5′ c c a a a t t g t a a g a t a a t a t t t t t c t t g a t g c a g g a a g a 3′

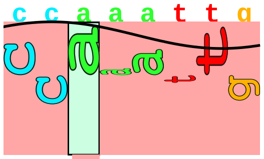

sigma38p10 3.8 bits

... ----| proximalUP-(37)-sigma38p10 2758287 Gap 4.5 bits

... ----| distalUP-(51)-sigma38p10 2758287 Gap 5.1 bits

... ----| sigma38p10-proximalUP-distalUP 2758287 total 3.2 bits

*2758250

5′ a a 3′

piece 37, NC_000913, alaE, config: linear, direction: -, begin: 2797299, end: 2797050

```
              .         *2797290  .         *2797280  .         *2797270  .
5' g a a g c t c a t t c c g g a g g a a a a c t t c a a t a c a c a t g t 3'
   T timestamp:_2018Sep20_11-35-24

     *2797260  .         *2797250  .         *2797240  .         *2797230  .
5' t c a c g a c a g a a c a g t a a a c a a c c a t c g c g a a c g t a t c t 3'

        *2797220  .         *2797210  .         *2797200  .         *2797190
5' g c a a c t g c a t g a c g c a a g c g t g a c t g c g g t g a g a a c a t 3'

   .         *2797180  .         *2797170  .         *2797160  .         *2797150
5' g c t a a t g c t c c t t a a t a a a a g c g t a a t g g g a t g t t a a t 3'
                                                                          ... proximalUP
                                                                 distalUP 0.1 bits
                                                                 |---------------- ... distalUP-(41)-s
                                                                 |---------------- ... sigma38p10-prox
                                                                       distalUP 5.5 bits
                                                                       |---------- ... distalUP-(40)-s
                                                                       |---------- ... sigma38p10-prox

      .         *2797140  .         *2797130  .         *2797120  .         *2797110
5' g g a g a t g a a a g c a g t g g a t c t c a t t g c c a t c t g c a t t t 3'
...  proximalUP 6.3 bits                                                sigma38p10 5.3 bits
   |-------------------------------------------------------------| proximalUP-(32)-sigma38p10 279
...  ------------------------------------------------------------| distalUP-(41)-sigma38p10 27971
...  |-----------------------------------------------------------| sigma38p10-proximalUP-distalUP
                                                                          ... sigma38p10
...  ----------------------------------------------------------------| distalUP-(40)-sigma38p10 2
...  |---------------------------------------------------------------| sigma38p10-proximalUP-dist
   |-----------------------------------------------------------------| proximalUP-(34)-sigma38p10
                                                                          ... distalUP
                                                                   |--------- ... distalUP-(53)-s
                                                                   |--------- ... sigma38p10-prox

          .         *2797100  .         *2797090  .         *2797080  .
```

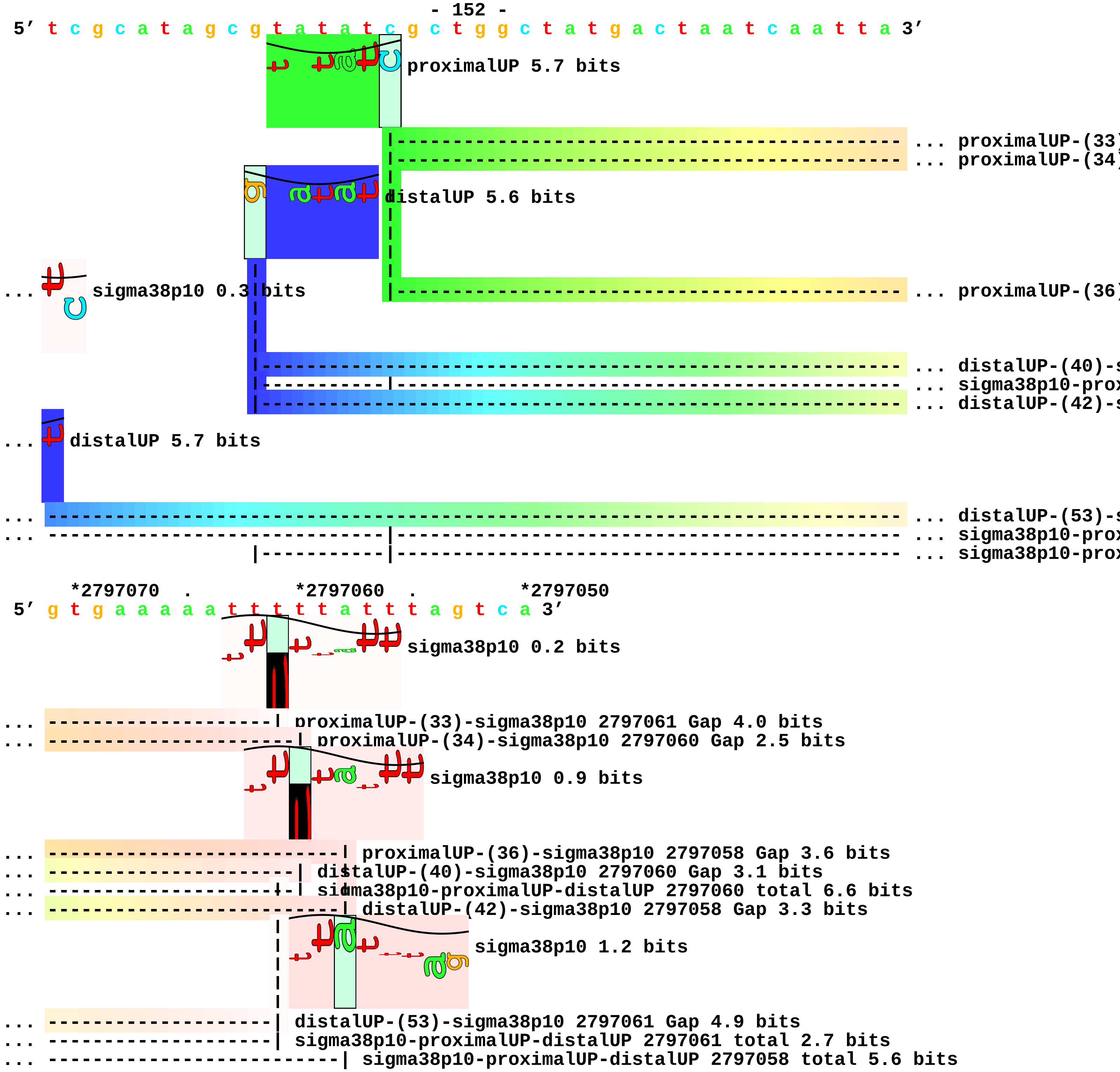
5′ t c g c a t a g c g t a t a t c g c t g g c t a t g a c t a a t c a a t t a 3′
proximalUP 5.7 bits
... proximalUP-(33
... proximalUP-(34
distalUP 5.6 bits
... sigma38p10 0.3 bits
... proximalUP-(36
... distalUP-(40)-s
... sigma38p10-prox
... distalUP-(42)-s
... distalUP 5.7 bits
... distalUP-(53)-s
... sigma38p10-prox
... sigma38p10-prox
*2797070 . *2797060 . *2797050
5′ g t g a a a a a t t t t t a t t t a g t c a 3′
sigma38p10 0.2 bits
... proximalUP-(33)-sigma38p10 2797061 Gap 4.0 bits
... proximalUP-(34)-sigma38p10 2797060 Gap 2.5 bits
sigma38p10 0.9 bits
... proximalUP-(36)-sigma38p10 2797058 Gap 3.6 bits
... distalUP-(40)-sigma38p10 2797060 Gap 3.1 bits
... sigma38p10-proximalUP-distalUP 2797060 total 6.6 bits
... distalUP-(42)-sigma38p10 2797058 Gap 3.3 bits
sigma38p10 1.2 bits
... distalUP-(53)-sigma38p10 2797061 Gap 4.9 bits
... sigma38p10-proximalUP-distalUP 2797061 total 2.7 bits
... sigma38p10-proximalUP-distalUP 2797058 total 5.6 bits

piece 38, NC_000913, csrA, config: linear, direction: +, begin: 2817177, end: 2817445

*2817180 . *2817190 . *2817200 . *2817210
5' c c t t g a a a g a t t a a a a g a g t c g g g t c t c t g t a t c c c 3'
proximalUP 2.8 bits
... proximalUP-(36)
distalUP 1.6 bits
... distalUP-(40)-s
... distalUP-proxin
T timestamp:_2018Sep20_11-35-24

. *2817220 . *2817230 . *2817240 . *2817250
5' g g c a t t a t c c a t c a t a t a a c g c c a a a a a g t a a g c a a t g 3'
sigma38p10 3.1 bits
proximalUP 3.8 bits
... proximalUP-(36)-sigma38p10 2817221 Gap 3.6 bits
... proximalUP-(41)
proximalUP 3.9 bits
... distalUP-(40)-sigma38p10 2817221 Gap 3.1 bits
... distalUP-proximalUP-sigma38p10 2817221 total 0.9 bits
... proximalUP-(36)
distalUP 5.0 bits
... proximalUP-(43)
... distalUP-(43)-s
... distalUP-proxin
distalUP 5.7 bits
... distalUP-(49)-s
... distalUP-proxin
proximalUP 3.3 bits
... distalUP-(51)-s
... distalUP-proxin

. *2817260 . *2817270 . *2817280 . *2817290
5' a c a a a c a c a t t a c a t c t a a g c a g t c a t g g c a t t a c a t t 3'
sigma38p10 3.3 bits
... sigma38p10

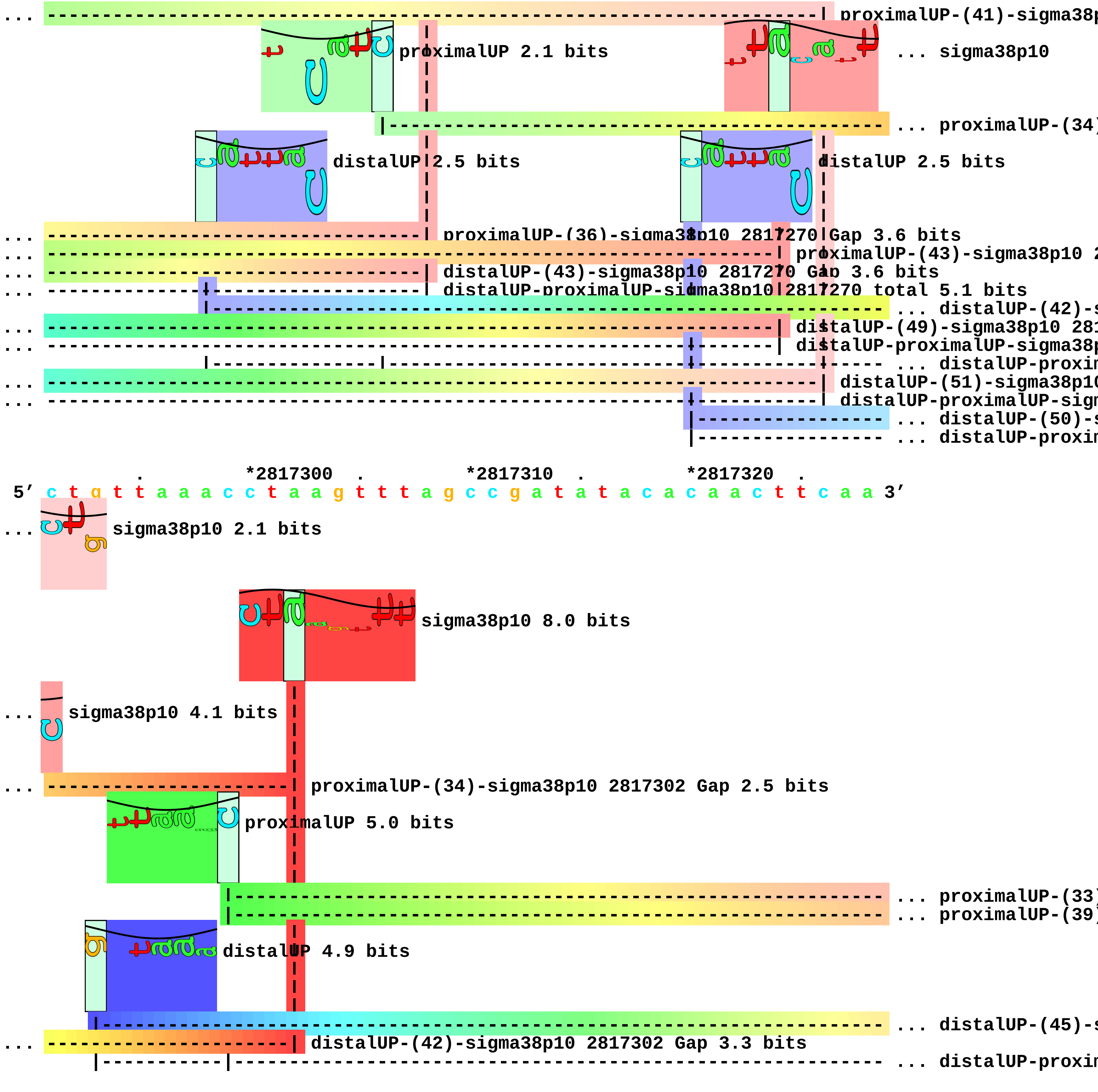

... proximalUP-(41)-sigma38p
proximalUP 2.1 bits
... sigma38p10
... proximalUP-(34)
distalUP 2.5 bits
distalUP 2.5 bits
... proximalUP-(36)-sigma38p10 2817270 Gap 3.6 bits
... proximalUP-(43)-sigma38p10 2
... distalUP-(43)-sigma38p10 2817270 Gap 3.6 bits
... distalUP-proximalUP-sigma38p10 2817270 total 5.1 bits
... distalUP-(42)-s
... distalUP-(49)-sigma38p10 281
... distalUP-proximalUP-sigma38p
... distalUP-proxin
... distalUP-(51)-sigma38p10
... distalUP-proximalUP-sigm
... distalUP-(50)-s
... distalUP-proxin
. *2817300 . *2817310 . *2817320 .
5' c t g t t a a a c c t a a g t t t a g c c g a t a t a c a c a a c t t c a a 3'
... sigma38p10 2.1 bits
sigma38p10 8.0 bits
... sigma38p10 4.1 bits
... proximalUP-(34)-sigma38p10 2817302 Gap 2.5 bits
proximalUP 5.0 bits
... proximalUP-(33)
... proximalUP-(39)
distalUP 4.9 bits
... distalUP-(45)-s
... distalUP-(42)-sigma38p10 2817302 Gap 3.3 bits
... distalUP-proxin

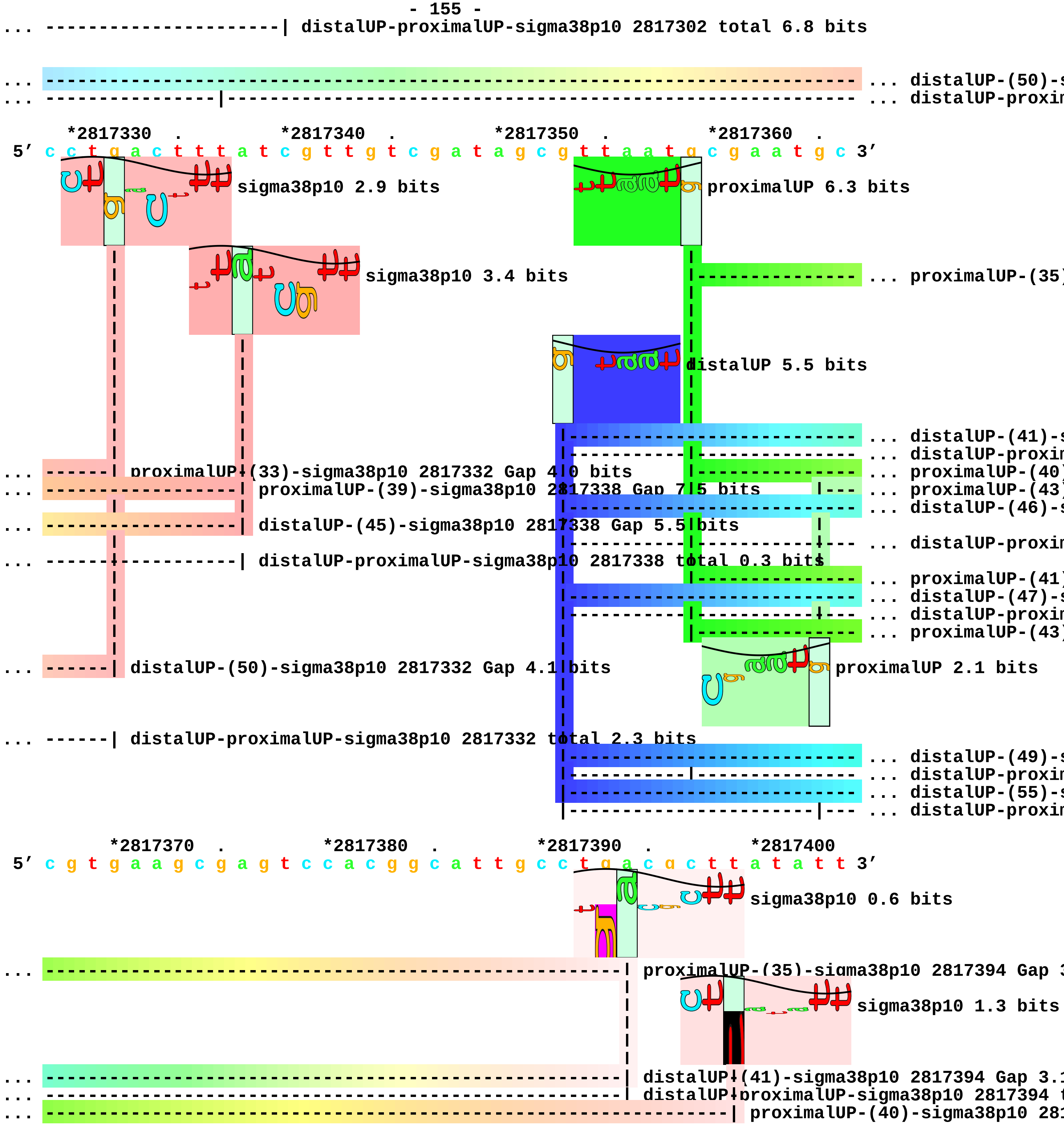
... ---------------------| distalUP-proximalUP-sigma38p10 2817302 total 6.8 bits
... distalUP-(50)-s
... distalUP-proxin
*2817330 . *2817340 . *2817350 . *2817360 .
5' c c t g a c t t t a t c g t t g t c g a t a g c g t t a a t g c g a a t g c 3'
sigma38p10 2.9 bits
proximalUP 6.3 bits
sigma38p10 3.4 bits
... proximalUP-(35)
distalUP 5.5 bits
... distalUP-(41)-s
... distalUP-proxin
... ------| proximalUP-(33)-sigma38p10 2817332 Gap 4.0 bits
... proximalUP-(40)
... ------------------| proximalUP-(39)-sigma38p10 2817338 Gap 7.5 bits
... proximalUP-(43)
... distalUP-(46)-s
... ------------------| distalUP-(45)-sigma38p10 2817338 Gap 5.5 bits
... distalUP-proxin
... ------------------| distalUP-proximalUP-sigma38p10 2817338 total 0.3 bits
... proximalUP-(41)
... distalUP-(47)-s
... distalUP-proxin
... proximalUP-(43)
... ------| distalUP-(50)-sigma38p10 2817332 Gap 4.1 bits
proximalUP 2.1 bits
... ------| distalUP-proximalUP-sigma38p10 2817332 total 2.3 bits
... distalUP-(49)-s
... distalUP-proxin
... distalUP-(55)-s
... distalUP-proxin
*2817370 . *2817380 . *2817390 . *2817400
5' c g t g a a g c g a g t c c a c g g c a t t g c c t g a c g c t t a t a t t 3'
sigma38p10 0.6 bits
... proximalUP-(35)-sigma38p10 2817394 Gap 3
sigma38p10 1.3 bits
... distalUP-(41)-sigma38p10 2817394 Gap 3.1
... distalUP-proximalUP-sigma38p10 2817394 t
... proximalUP-(40)-sigma38p10 281

```
... ---------------------------------------------------------------------------- ... proximalUP-(43)
... ----------------------------------------------------------------| distalUP-(46)-sigma38p10 28173
                                                                      ... sigma38p10
... ----------------------------------------------------------------| distalUP-proximalUP-sigma38p10
                                                                      ... sigma38p10
... ------------------------------------------------------------------| proximalUP-(41)-sigma38p10 2
... ------------------------------------------------------------------| distalUP-(47)-sigma38p10 281
... ------------------------------------------------------------------| distalUP-proximalUP-sigma38p
... ----------------------------------------------------------------------| proximalUP-(43)-sigma38p
... ----------------------------------------------------------------------| distalUP-(49)-sigma38p10
... ----------------------------------------------------------------------| distalUP-proximalUP-sigm
... ---------------------------------------------------------------------------- ... distalUP-(55)-s
... ---------------------------------------------------------------------------- ... distalUP-proxim

    .         *2817410  .         *2817420  .         *2817430  .         *2817440
5' a t t g c a a t t t c g c g c t g a c c c a g c c t t t c a c a c t g g c t 3'
                         sigma38p10 4.1 bits

... ------| proximalUP-(43)-sigma38p10 2817408 Gap 5.1 bits
... sigma38p10 1.1 bits
... sigma38p10 7.0 bits
... ------| distalUP-(55)-sigma38p10 2817408 Gap 4.9 bits
... ------| distalUP-proximalUP-sigma38p10 2817408 total 1.7 bits

       .
5' a a c 3'
```

piece 39, NC_000913, ygdH, config: linear, direction: -, begin: 2924420, end: 2924202

```
    *2924420   .          *2924410   .          *2924400   .          *2924390   .
5' a g a g g t c g c t g c t g g c g g t g c g t t t a a g c a t a t c c a c t 3'
```

T timestamp:_2018Sep20_11-35-24

proximalUP 3.2 bits

... proximalUP-(43)

distalUP 4.8 bits

... distalUP-(55)-s

... sigma38p10-prox

```
      *2924380   .          *2924370   .          *2924360   .          *2924350   .
5' t c c a g c t g c g a c a a c a t a t c c a t g g a g c c a a g c g g g c t 3'
```

... sigma38p10

... ... proximalUP-(43)

... distalUP

... ... distalUP-(55)-s

... ... sigma38p10-prox

... distalUP-(40)-s

... sigma38p10-prox

```
          *2924340   .          *2924330   .          *2924320   .          *2924310
5' a a t a t g t g t a a t c a a g a a a a a c t c c t t a t g g g a c g a a t 3'
```

... sigma38p10 6.1 bits proximalUP 4.3 bits ... sigma38p10

... proximalUP-(43)-sigma38p10 2924344 Gap 5.1 bits ... proximalUP

... distalUP 3.5 bits ... proximalUP-(36)

... distalUP-(55)-sigma38p10 2924344 Gap 4.9 bits ... distalUP

... sigma38p10-proximalUP-distalUP 2924344 total 4.1 bits ... distalUP-(41)-s

... ... distalUP-(40)-s

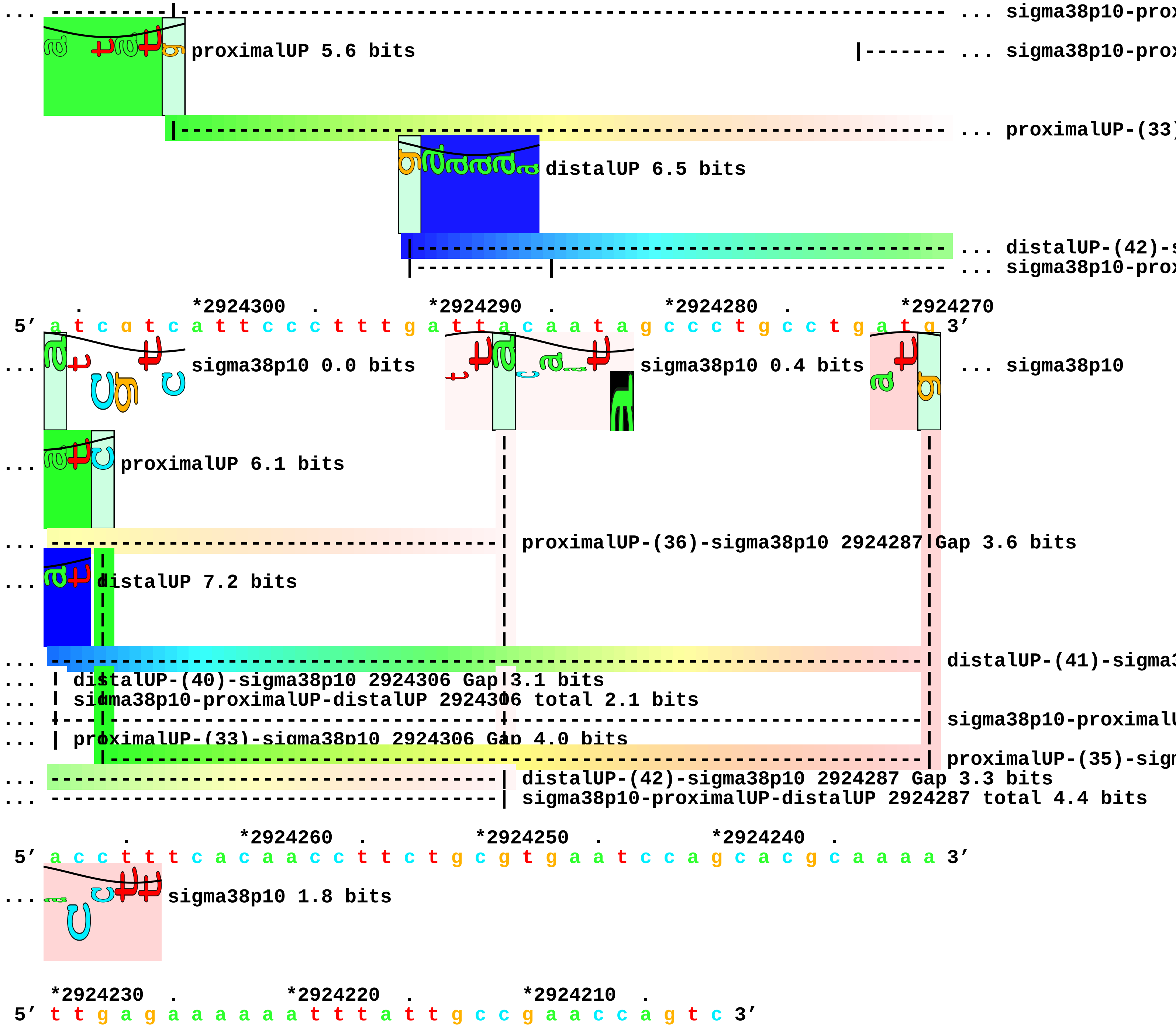
... sigma38p10-prox
proximalUP 5.6 bits
... sigma38p10-prox
... proximalUP-(33)
distalUP 6.5 bits
... distalUP-(42)-s
... sigma38p10-prox
. *2924300 . *2924290 . *2924280 . *2924270
5′ a t c a t c a t t c c c t t t g a t t a c a a t a g c c c t g c c t g a t g 3′
... sigma38p10 0.0 bits
sigma38p10 0.4 bits
... sigma38p10
... proximalUP 6.1 bits
... proximalUP-(36)-sigma38p10 2924287 Gap 3.6 bits
... distalUP 7.2 bits
... distalUP-(41)-sigma3
... distalUP-(40)-sigma38p10 2924306 Gap 3.1 bits
... sigma38p10-proximalUP-distalUP 2924306 total 2.1 bits
... sigma38p10-proximalU
... proximalUP-(33)-sigma38p10 2924306 Gap 4.0 bits
proximalUP-(35)-sigm
... distalUP-(42)-sigma38p10 2924287 Gap 3.3 bits
... sigma38p10-proximalUP-distalUP 2924287 total 4.4 bits
. *2924260 . *2924250 . *2924240 .
5′ a c c t t t c a c a a c c t t c t g c g t g a a t c c a g c a c g c a a a a 3′
... sigma38p10 1.8 bits
*2924230 . *2924220 . *2924210 .
5′ t t g a g a a a a a a t t t a t t g c c g a a c c a g t c 3′

piece 40, NC_000913, omrA, config: linear, direction: +, begin: 2974103, end: 2974328

```
       .         *2974110  .         *2974120  .         *2974130  .         *2974140
5' a g c g a c a g t a a a t t a g g t g c g a a a a a a a a c c t g c g c a t 3'
 T timestamp:_2018Sep20_11-35-24

           .         *2974150  .         *2974160  .         *2974170  .
5' c c g c g c a g g t t g g t g c a a g a g a c a g g g t a c g a a g a g c g 3'

     *2974180  .         *2974190  .         *2974200  .         *2974210  .
5' t a c c g a a t a a t c t c a c c a a t c a a t a c c t c t g g g a t c t t 3'
```

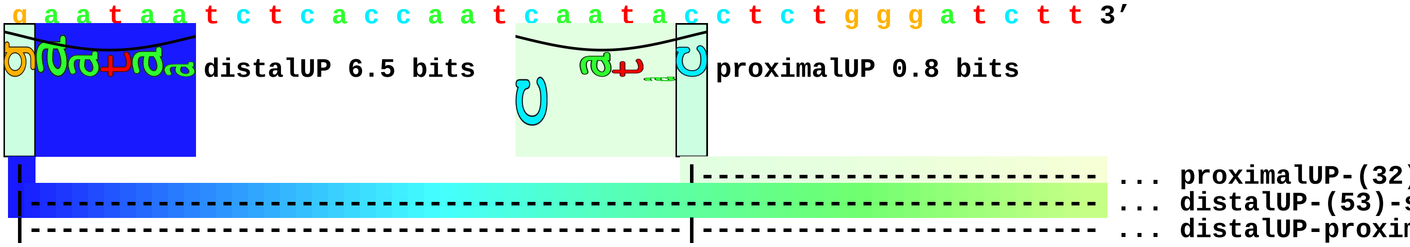


```
         *2974220  .         *2974230  .         *2974240  .         *2974250
5' g a t t g t g g t c t g c a c g a c g a c t c t t c g c c a g c g a g a a a 3'
```

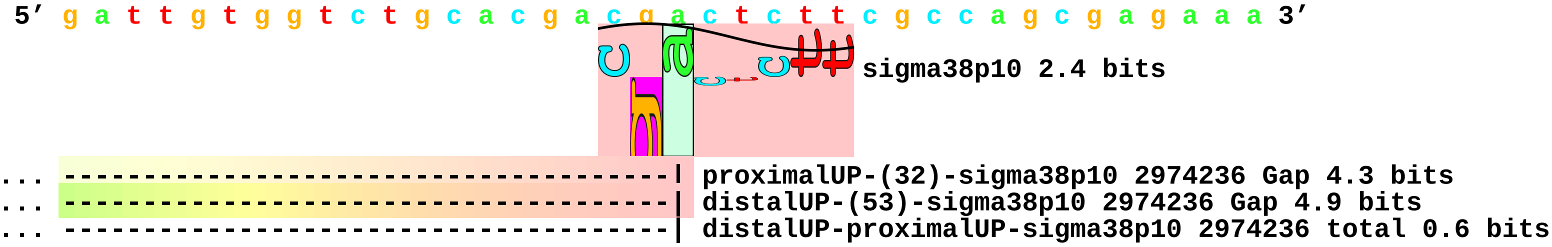


```
   .         *2974260  .         *2974270  .         *2974280  .         *2974290
5' a c g c a a a g g a a t g a a g g g a a a t g c a a c g a g g t g t g t a a 3'

       .         *2974300  .         *2974310  .         *2974320  .
5' a t t g t c g g t t a c t g t t a c a g a t t g a t g a c c g g c a a a 3'
```

piece 41, NC_000913, ygeI, config: linear, direction: -, begin: 2992349, end: 2991050

. *2992340 . *2992330 . *2992320 .
5′ t a a g t c c c c a g g c a t t g c c a c g t t a c t c c a c a a c g t g a 3′

T timestamp:_2018Sep20_11-35-24

distalUP 2.3 bits

... distalUP-(50)-s
... sigma38p10-prox

*2992310 . *2992300 . *2992290 . *2992280 .
5′ a a a a t g g t t a t a t a a a a t c t c g g t t c c t a t t t t a a t g t 3′

proximalUP 5.6 bits

... sigma38p10

... distalUP-(50)-sigma38p10 299
... sigma38p10-proximalUP-distal
proximalUP-(28)-sigma38p10 2

proximalUP 6.3 bits

... proximalUP-(32)

distalUP 4.2 bits

... distalUP-(42)-s
... sigma38p10-prox
... proximalUP-(33)
... distalUP-(43)-s
... sigma38p10-prox
... proximalUP-(37)
... distalUP-(47)-s
... sigma38p10-prox
... proximalUP-(40)
... distalUP-(50)-s
... sigma38p10-prox

*2992270 . *2992260 . *2992250 . *2992240
5′ t c a g a c a a g g g t t a t c t a a c a a a t c t c g t t c a t c t t t t 3′

... sigma38p10 4.6 bits

sigma38p10 0.2 bits

sigma38p10 2.3 bits

... sigma38p10

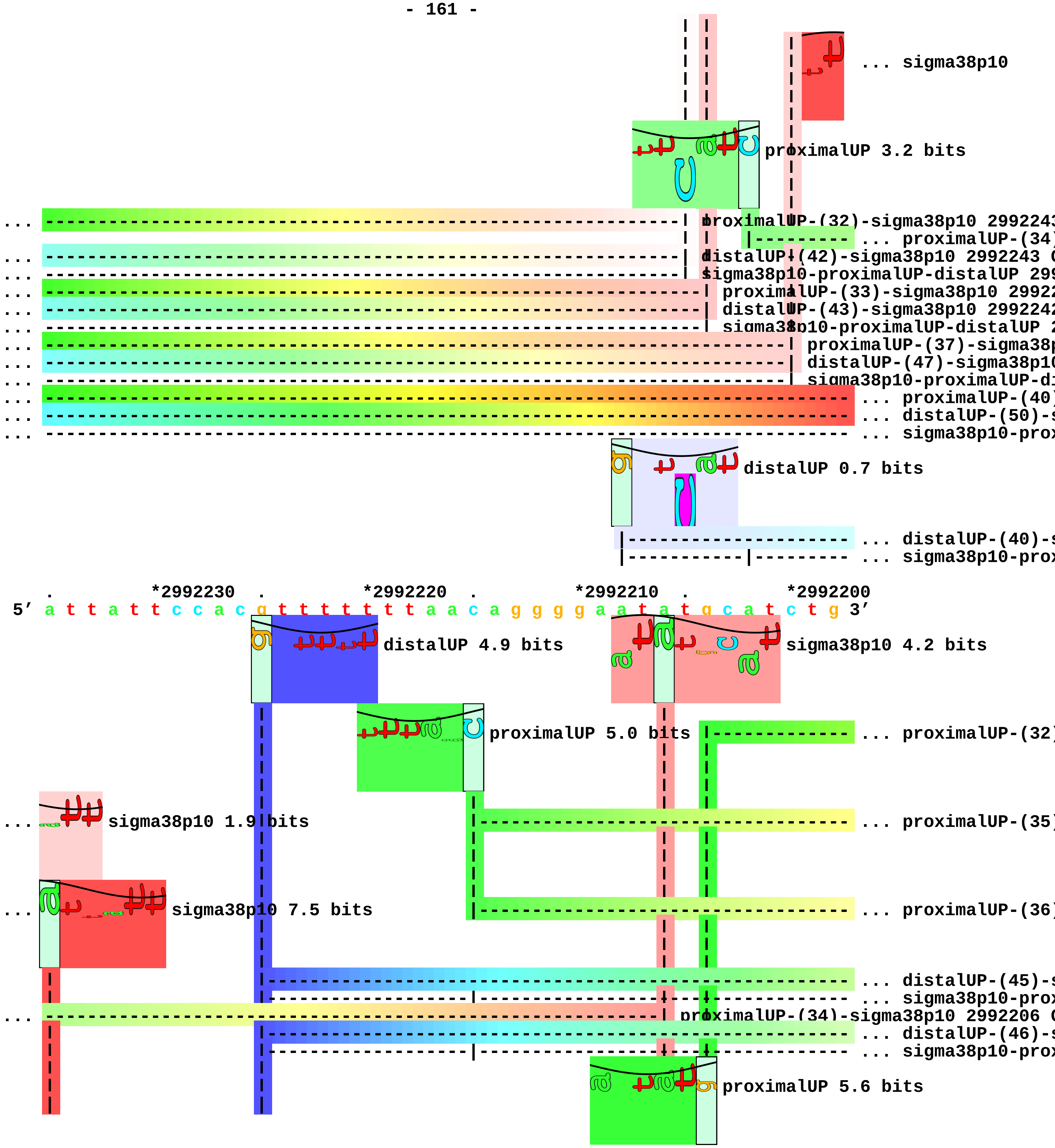

sigma38p10
proximalUP 3.2 bits
proximalUP-(32)-sigma38p10 299224
proximalUP-(34)
distalUP-(42)-sigma38p10 2992243
sigma38p10-proximalUP-distalUP 299
proximalUP-(33)-sigma38p10 2992
distalUP-(43)-sigma38p10 299224
sigma38p10-proximalUP-distalUP
proximalUP-(37)-sigma38
distalUP-(47)-sigma38p1
sigma38p10-proximalUP-d
proximalUP-(40)
distalUP-(50)-
sigma38p10-pro
distalUP 0.7 bits
distalUP-(40)-
sigma38p10-pro
*2992230
*2992220
*2992210
*2992200
5′ a t t a t t c c a c g t t t t t t t a a c a g g g g a a t a t g c a t c t g 3′
distalUP 4.9 bits
sigma38p10 4.2 bits
proximalUP 5.0 bits
proximalUP-(32)
sigma38p10 1.9 bits
proximalUP-(35)
sigma38p10 7.5 bits
proximalUP-(36)
distalUP-(45)-
sigma38p10-pro
proximalUP-(34)-sigma38p10 2992206
distalUP-(46)-
sigma38p10-pro
proximalUP 5.6 bits

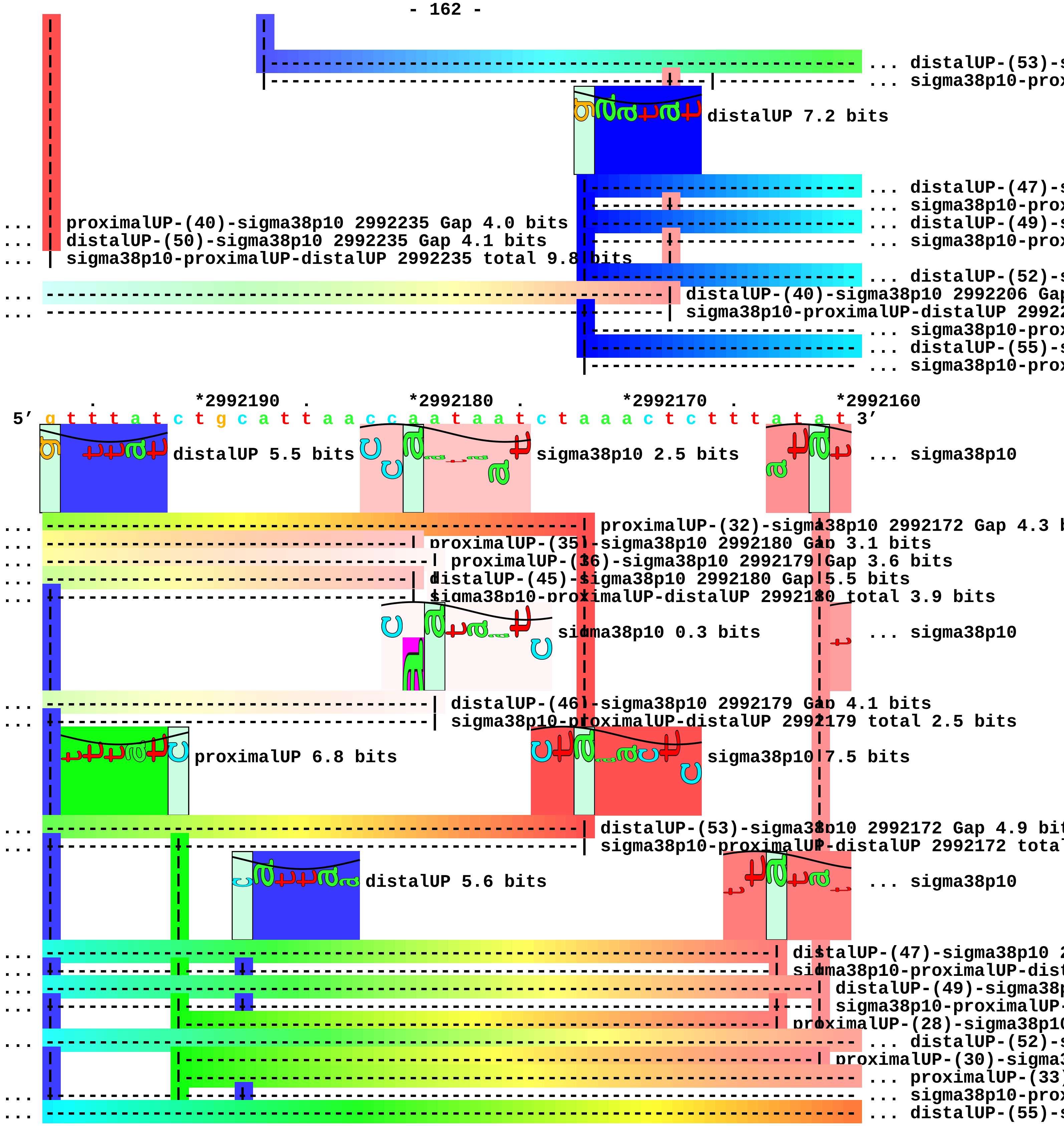

... distalUP-(53)-s
... sigma38p10-prox
distalUP 7.2 bits
... distalUP-(47)-s
... sigma38p10-prox
... distalUP-(49)-s
... sigma38p10-prox
... proximalUP-(40)-sigma38p10 2992235 Gap 4.0 bits
... distalUP-(50)-sigma38p10 2992235 Gap 4.1 bits
... sigma38p10-proximalUP-distalUP 2992235 total 9.8 bits
... distalUP-(52)-s
... distalUP-(40)-sigma38p10 2992206 Gap
... sigma38p10-proximalUP-distalUP 29922
... sigma38p10-prox
... distalUP-(55)-s
... sigma38p10-prox
*2992190 *2992180 *2992170 *2992160
5′ g t t t a t c t g c a t t a a c c a a t a a t c t a a a c t c t t t a t a t 3′
distalUP 5.5 bits
sigma38p10 2.5 bits
... sigma38p10
... proximalUP-(32)-sigma38p10 2992172 Gap 4.3 b
... proximalUP-(35)-sigma38p10 2992180 Gap 3.1 bits
... proximalUP-(36)-sigma38p10 2992179 Gap 3.6 bits
... distalUP-(45)-sigma38p10 2992180 Gap 5.5 bits
... sigma38p10-proximalUP-distalUP 2992180 total 3.9 bits
sigma38p10 0.3 bits
... sigma38p10
... distalUP-(46)-sigma38p10 2992179 Gap 4.1 bits
... sigma38p10-proximalUP-distalUP 2992179 total 2.5 bits
proximalUP 6.8 bits
sigma38p10 7.5 bits
... distalUP-(53)-sigma38p10 2992172 Gap 4.9 bit
... sigma38p10-proximalUP-distalUP 2992172 total
distalUP 5.6 bits
... sigma38p10
... distalUP-(47)-sigma38p10 2
... sigma38p10-proximalUP-dist
... distalUP-(49)-sigma38p
... sigma38p10-proximalUP-
proximalUP-(28)-sigma38p10
... distalUP-(52)-s
proximalUP-(30)-sigma3
... proximalUP-(33)
... sigma38p10-prox
... distalUP-(55)-s

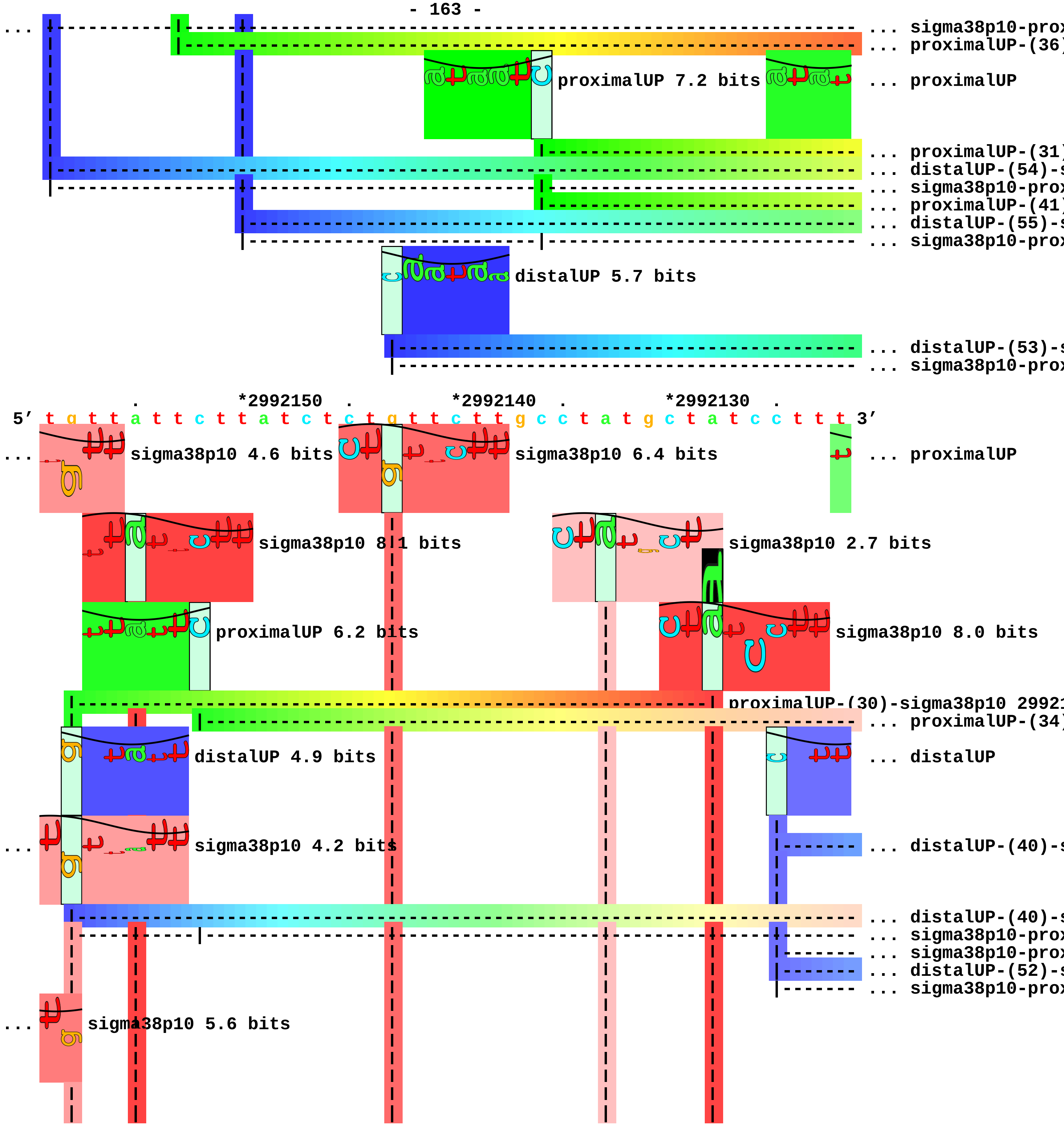
... sigma38p10-prox
... proximalUP-(36)
proximalUP 7.2 bits
... proximalUP
... proximalUP-(31)
... distalUP-(54)-s
... sigma38p10-prox
... proximalUP-(41)
... distalUP-(55)-s
... sigma38p10-prox
distalUP 5.7 bits
... distalUP-(53)-s
... sigma38p10-prox
*2992150
*2992140
*2992130
5′ t g t t a t t c t t a t c t c t g t t c t t g c c t a t g c t a t c c t t t 3′
sigma38p10 4.6 bits
sigma38p10 6.4 bits
... proximalUP
sigma38p10 8.1 bits
sigma38p10 2.7 bits
proximalUP 6.2 bits
sigma38p10 8.0 bits
proximalUP-(30)-sigma38p10 29921
... proximalUP-(34)
distalUP 4.9 bits
... distalUP
sigma38p10 4.2 bits
... distalUP-(40)-s
... distalUP-(40)-s
... sigma38p10-prox
... sigma38p10-prox
... distalUP-(52)-s
... sigma38p10-prox
sigma38p10 5.6 bits

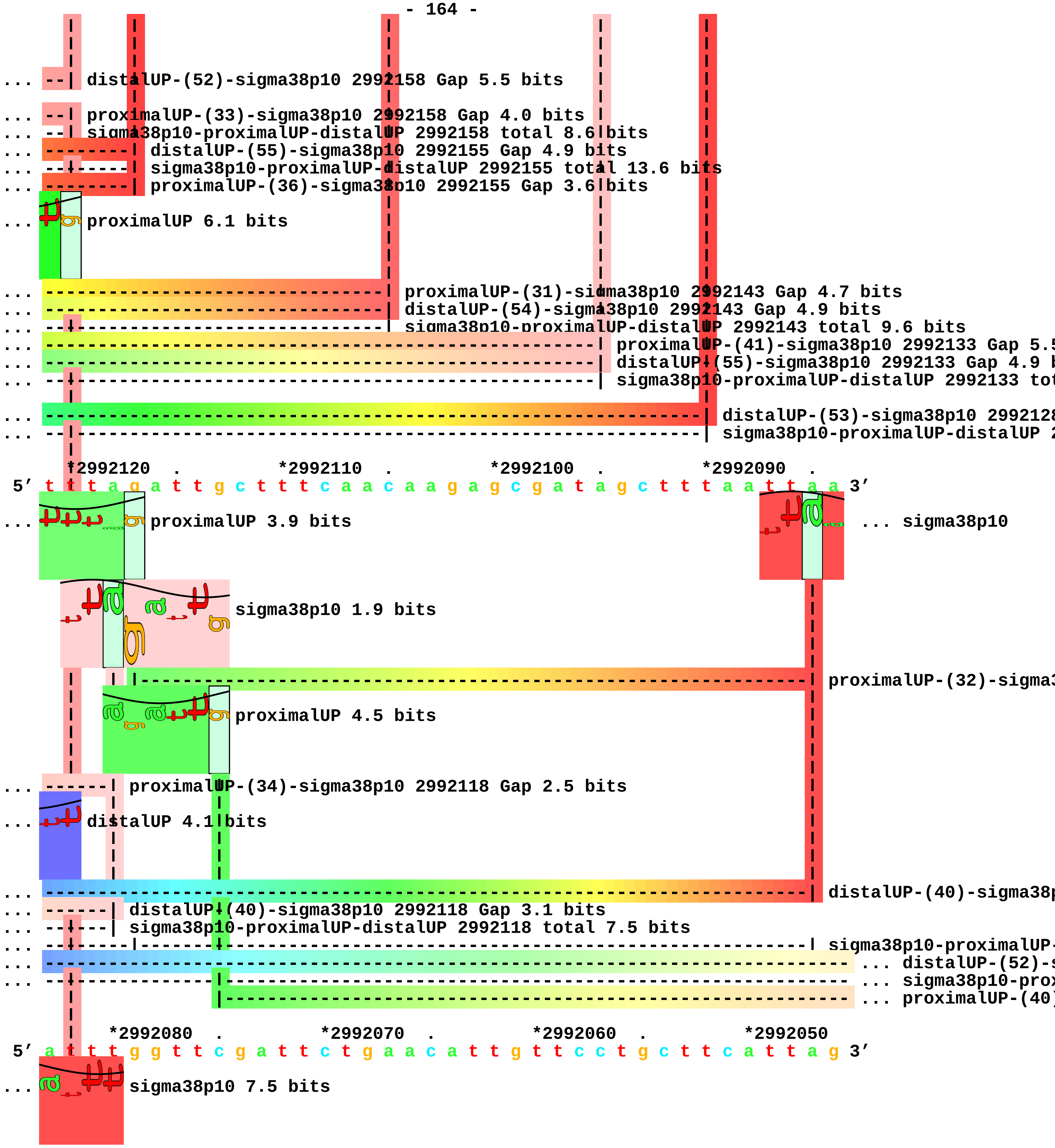
... distalUP-(52)-sigma38p10 2992158 Gap 5.5 bits
... proximalUP-(33)-sigma38p10 2992158 Gap 4.0 bits
... sigma38p10-proximalUP-distalUP 2992158 total 8.6 bits
... distalUP-(55)-sigma38p10 2992155 Gap 4.9 bits
... sigma38p10-proximalUP-distalUP 2992155 total 13.6 bits
... proximalUP-(36)-sigma38p10 2992155 Gap 3.6 bits
... proximalUP 6.1 bits
... proximalUP-(31)-sigma38p10 2992143 Gap 4.7 bits
... distalUP-(54)-sigma38p10 2992143 Gap 4.9 bits
... sigma38p10-proximalUP-distalUP 2992143 total 9.6 bits
... proximalUP-(41)-sigma38p10 2992133 Gap 5.5
... distalUP-(55)-sigma38p10 2992133 Gap 4.9 b
... sigma38p10-proximalUP-distalUP 2992133 tot
... distalUP-(53)-sigma38p10 299212
... sigma38p10-proximalUP-distalUP 2
*2992120 . *2992110 . *2992100 . *2992090 .
5′ t t t a g a t t g c t t t c a a c a a g a g c g a t a g c t t t a a t t a a 3′
... proximalUP 3.9 bits
... sigma38p10
sigma38p10 1.9 bits
proximalUP-(32)-sigma3
proximalUP 4.5 bits
... proximalUP-(34)-sigma38p10 2992118 Gap 2.5 bits
... distalUP 4.1 bits
... distalUP-(40)-sigma38p
... distalUP-(40)-sigma38p10 2992118 Gap 3.1 bits
... sigma38p10-proximalUP-distalUP 2992118 total 7.5 bits
... sigma38p10-proximalUP-
... distalUP-(52)-s
... sigma38p10-prox
... proximalUP-(40)
*2992080 . *2992070 . *2992060 . *2992050
5′ a t t t g g t t c g a t t c t g a a c a t t g t t c c t g c t t c a t t a g 3′
... sigma38p10 7.5 bits

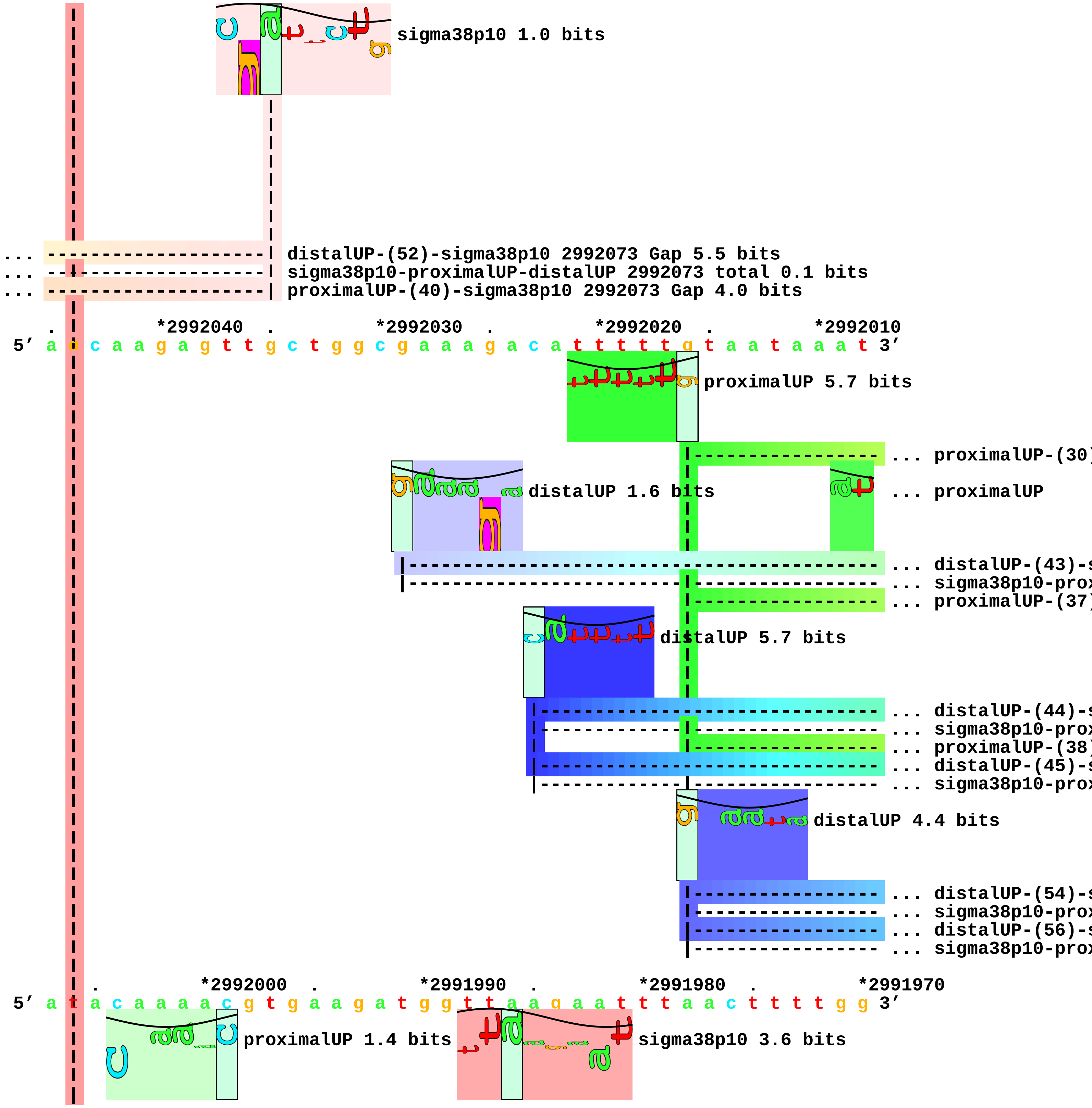
sigma38p10 1.0 bits
... distalUP-(52)-sigma38p10 2992073 Gap 5.5 bits
... sigma38p10-proximalUP-distalUP 2992073 total 0.1 bits
... proximalUP-(40)-sigma38p10 2992073 Gap 4.0 bits
. *2992040 . *2992030 . *2992020 . *2992010
5′ a a c a a g a g t t g c t g g c g a a a g a c a t t t t t g t a a t a a a t 3′
proximalUP 5.7 bits
... proximalUP-(30)
distalUP 1.6 bits
... proximalUP
... distalUP-(43)-s
... sigma38p10-prox
... proximalUP-(37)
distalUP 5.7 bits
... distalUP-(44)-s
... sigma38p10-prox
... proximalUP-(38)
... distalUP-(45)-s
... sigma38p10-prox
distalUP 4.4 bits
... distalUP-(54)-s
... sigma38p10-prox
... distalUP-(56)-s
... sigma38p10-prox
. *2992000 . *2991990 . *2991980 . *2991970
5′ a t a c a a a a c g t g a a g a t g g t t a a g a a t t t a a c t t t t g g 3′
proximalUP 1.4 bits
sigma38p10 3.6 bits

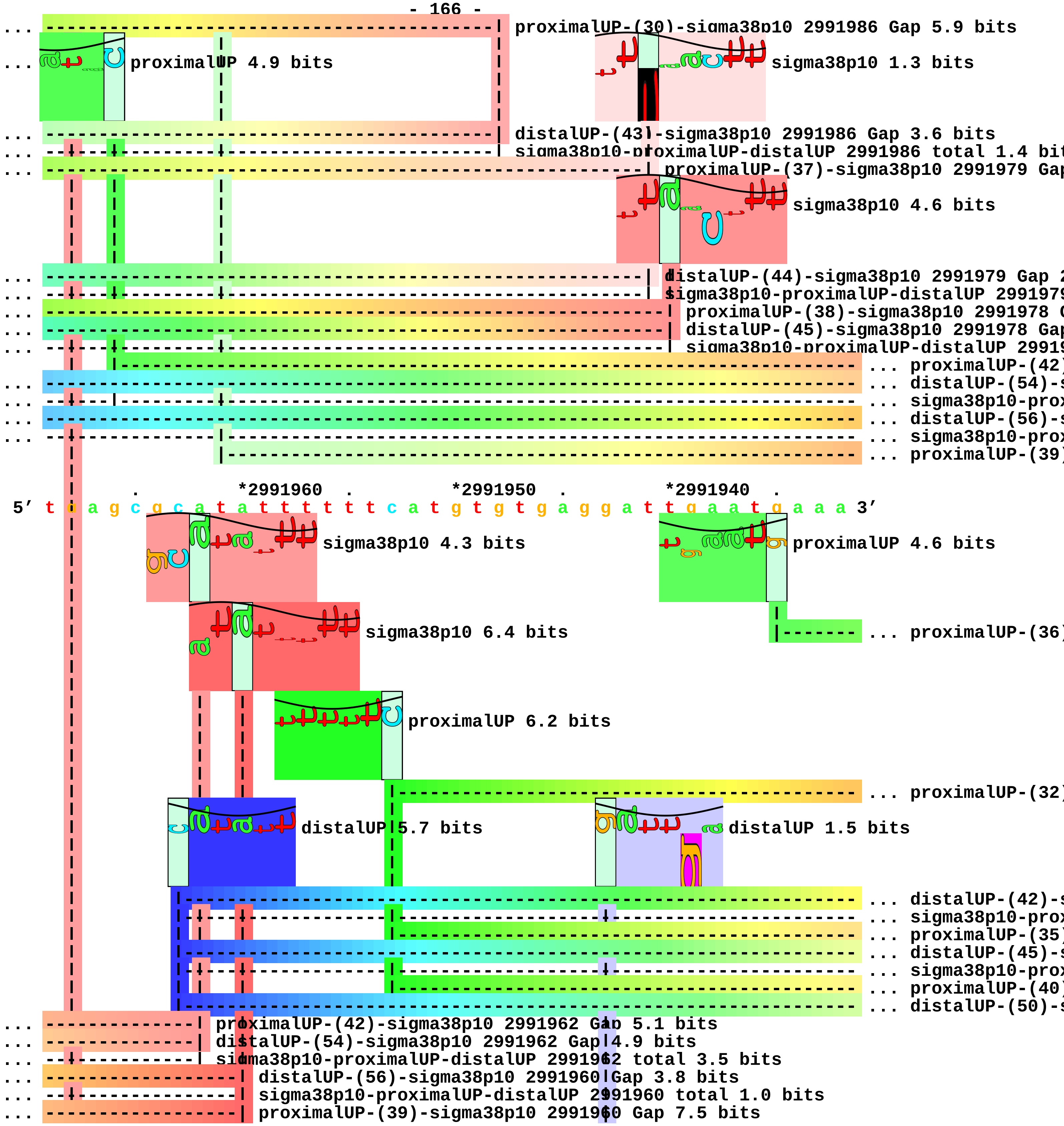
... proximalUP-(30)-sigma38p10 2991986 Gap 5.9 bits
... proximalUP 4.9 bits
sigma38p10 1.3 bits
... distalUP-(43)-sigma38p10 2991986 Gap 3.6 bits
... sigma38p10-proximalUP-distalUP 2991986 total 1.4 bit
... proximalUP-(37)-sigma38p10 2991979 Gap
sigma38p10 4.6 bits
... distalUP-(44)-sigma38p10 2991979 Gap 2
... sigma38p10-proximalUP-distalUP 2991979
... proximalUP-(38)-sigma38p10 2991978 G
... distalUP-(45)-sigma38p10 2991978 Gap
... sigma38p10-proximalUP-distalUP 29919
... proximalUP-(42)
... distalUP-(54)-s
... sigma38p10-prox
... distalUP-(56)-s
... sigma38p10-prox
... proximalUP-(39)
*2991960
*2991950
*2991940
5′ t a a g c g c a t a t t t t t t c a t g t g t g a g g a t t g a a t g a a a 3′
sigma38p10 4.3 bits
proximalUP 4.6 bits
sigma38p10 6.4 bits
... proximalUP-(36)
proximalUP 6.2 bits
... proximalUP-(32)
distalUP 5.7 bits
distalUP 1.5 bits
... distalUP-(42)-s
... sigma38p10-prox
... proximalUP-(35)
... distalUP-(45)-s
... sigma38p10-prox
... proximalUP-(40)
... distalUP-(50)-s
... proximalUP-(42)-sigma38p10 2991962 Gap 5.1 bits
... distalUP-(54)-sigma38p10 2991962 Gap 4.9 bits
... sigma38p10-proximalUP-distalUP 2991962 total 3.5 bits
... distalUP-(56)-sigma38p10 2991960 Gap 3.8 bits
... sigma38p10-proximalUP-distalUP 2991960 total 1.0 bits
... proximalUP-(39)-sigma38p10 2991960 Gap 7.5 bits

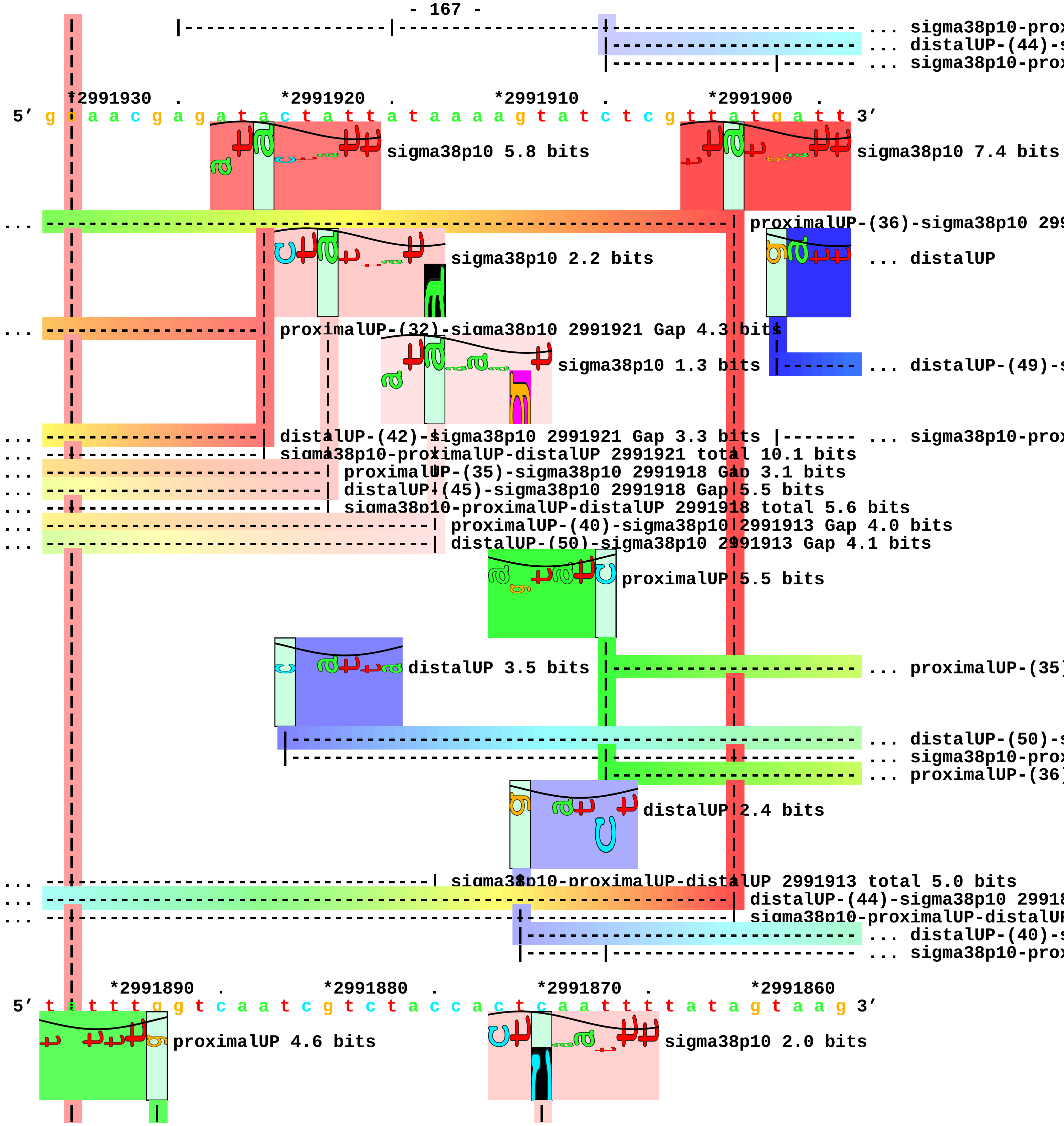

... sigma38p10-prox
... distalUP-(44)-s
... sigma38p10-prox
*2991930 . *2991920 . *2991910 . *2991900 .
5′ g a a a c g a g a t a c t a t t a t a a a a g t a t c t c g t t a t g a t t 3′
sigma38p10 5.8 bits
sigma38p10 7.4 bits
... proximalUP-(36)-sigma38p10 299
sigma38p10 2.2 bits
... distalUP
... proximalUP-(32)-sigma38p10 2991921 Gap 4.3 bits
sigma38p10 1.3 bits
... distalUP-(49)-s
... distalUP-(42)-sigma38p10 2991921 Gap 3.3 bits
... sigma38p10-prox
... sigma38p10-proximalUP-distalUP 2991921 total 10.1 bits
... proximalUP-(35)-sigma38p10 2991918 Gap 3.1 bits
... distalUP-(45)-sigma38p10 2991918 Gap 5.5 bits
... sigma38p10-proximalUP-distalUP 2991918 total 5.6 bits
... proximalUP-(40)-sigma38p10 2991913 Gap 4.0 bits
... distalUP-(50)-sigma38p10 2991913 Gap 4.1 bits
proximalUP 5.5 bits
distalUP 3.5 bits
... proximalUP-(35)
... distalUP-(50)-s
... sigma38p10-prox
... proximalUP-(36)
distalUP 2.4 bits
... sigma38p10-proximalUP-distalUP 2991913 total 5.0 bits
... distalUP-(44)-sigma38p10 29918
... sigma38p10-proximalUP-distalUP
... distalUP-(40)-s
... sigma38p10-prox
*2991890 . *2991880 . *2991870 . *2991860
5′ t a t t t g g t c a a t c g t c t a c c a c t c a a t t t t a t a g t a a g 3′
proximalUP 4.6 bits
sigma38p10 2.0 bits

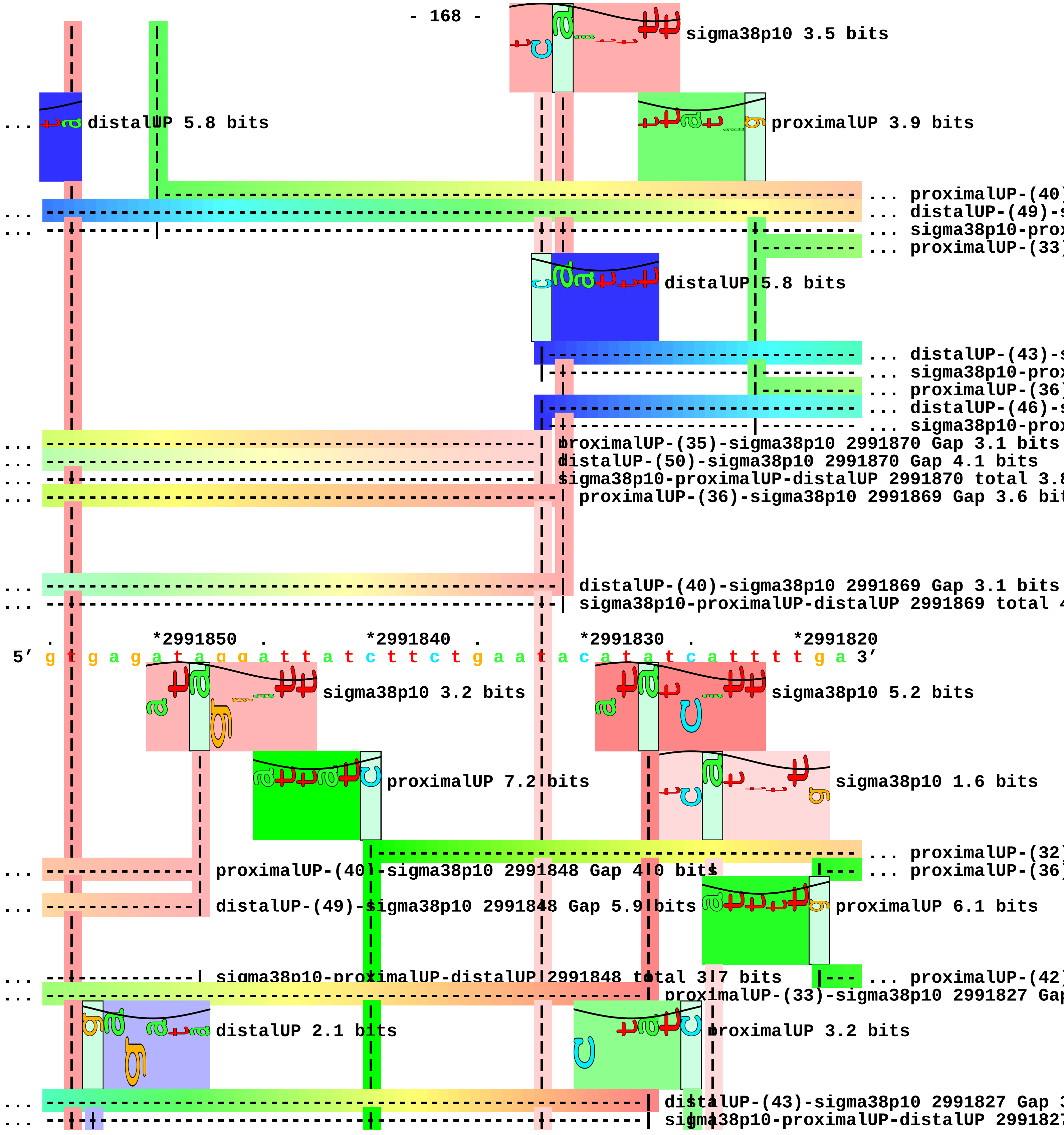

sigma38p10 3.5 bits
... distalUP 5.8 bits
proximalUP 3.9 bits
... proximalUP-(40)
... distalUP-(49)-s
... sigma38p10-prox
... proximalUP-(33)
distalUP 5.8 bits
... distalUP-(43)-s
... sigma38p10-prox
... proximalUP-(36)
... distalUP-(46)-s
... sigma38p10-prox
... proximalUP-(35)-sigma38p10 2991870 Gap 3.1 bits
... distalUP-(50)-sigma38p10 2991870 Gap 4.1 bits
... sigma38p10-proximalUP-distalUP 2991870 total 3.8
... proximalUP-(36)-sigma38p10 2991869 Gap 3.6 bit
... distalUP-(40)-sigma38p10 2991869 Gap 3.1 bits
... sigma38p10-proximalUP-distalUP 2991869 total 4
. *2991850 . *2991840 . *2991830 . *2991820
5′ g t g a g a t a g g a t t a t c t t c t g a a t a c a t a t c a t t t t g a 3′
sigma38p10 3.2 bits
sigma38p10 5.2 bits
proximalUP 7.2 bits
sigma38p10 1.6 bits
... proximalUP-(32)
... proximalUP-(40)-sigma38p10 2991848 Gap 4.0 bits
... proximalUP-(36)
... distalUP-(49)-sigma38p10 2991848 Gap 5.9 bits
proximalUP 6.1 bits
... sigma38p10-proximalUP-distalUP 2991848 total 3.7 bits
... proximalUP-(42)
... proximalUP-(33)-sigma38p10 2991827 Gap
distalUP 2.1 bits
proximalUP 3.2 bits
... distalUP-(43)-sigma38p10 2991827 Gap 3
... sigma38p10-proximalUP-distalUP 2991827

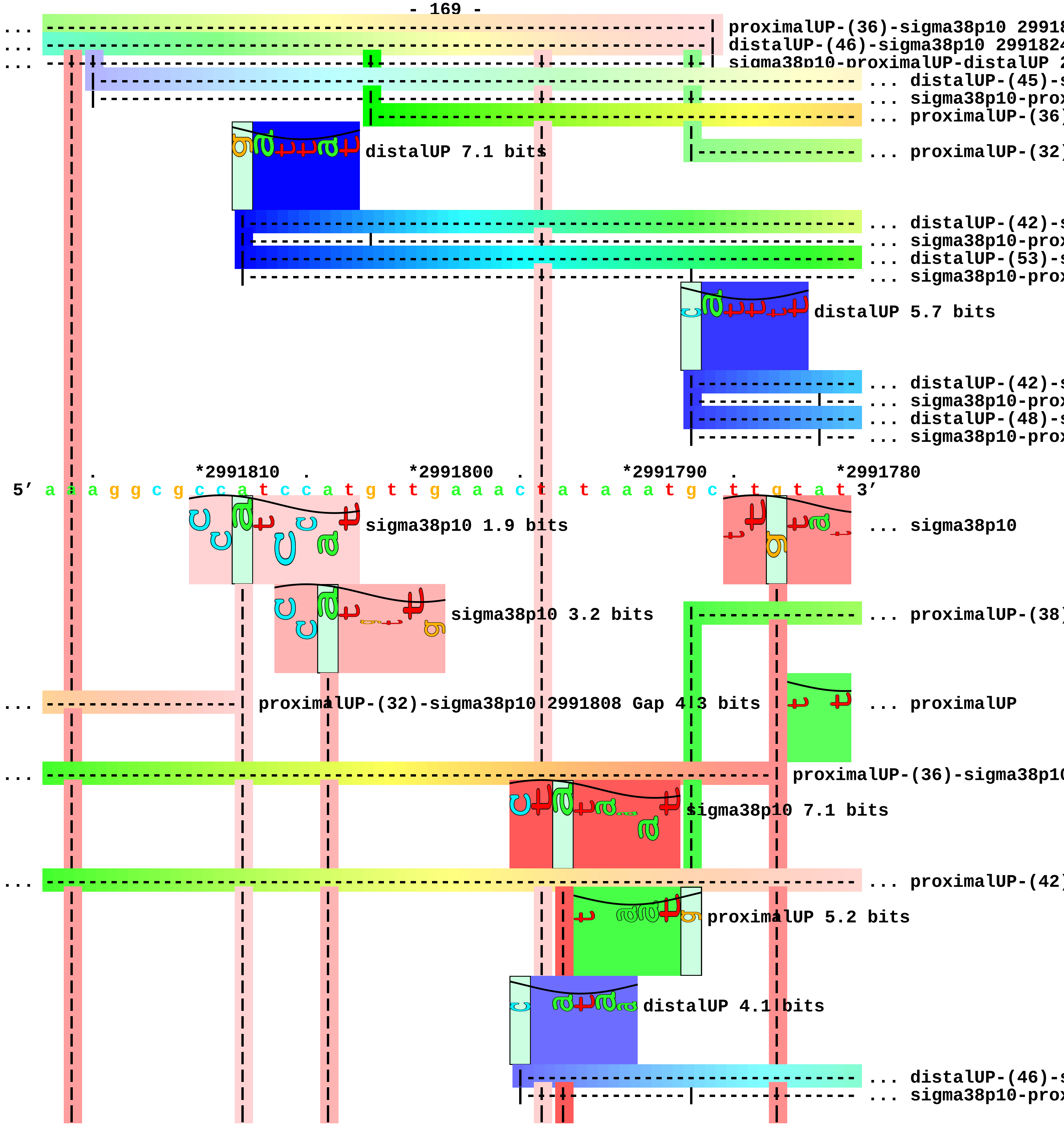
proximalUP-(36)-sigma38p10 29918
distalUP-(46)-sigma38p10 2991824
sigma38p10-proximalUP-distalUP 2
... distalUP-(45)-s
... sigma38p10-prox
... proximalUP-(36)
distalUP 7.1 bits
... proximalUP-(32)
... distalUP-(42)-s
... sigma38p10-prox
... distalUP-(53)-s
... sigma38p10-prox
distalUP 5.7 bits
... distalUP-(42)-s
... sigma38p10-prox
... distalUP-(48)-s
... sigma38p10-prox
*2991810
*2991800
*2991790
*2991780
5′ a a a g g c g c c a t c c a t g t t g a a a c t a t a a a t g c t t g t a t 3′
sigma38p10 1.9 bits
... sigma38p10
sigma38p10 3.2 bits
... proximalUP-(38)
... proximalUP-(32)-sigma38p10 2991808 Gap 4 3 bits
... proximalUP
... proximalUP-(36)-sigma38p10
sigma38p10 7.1 bits
... proximalUP-(42)
proximalUP 5.2 bits
distalUP 4.1 bits
... distalUP-(46)-s
... sigma38p10-prox

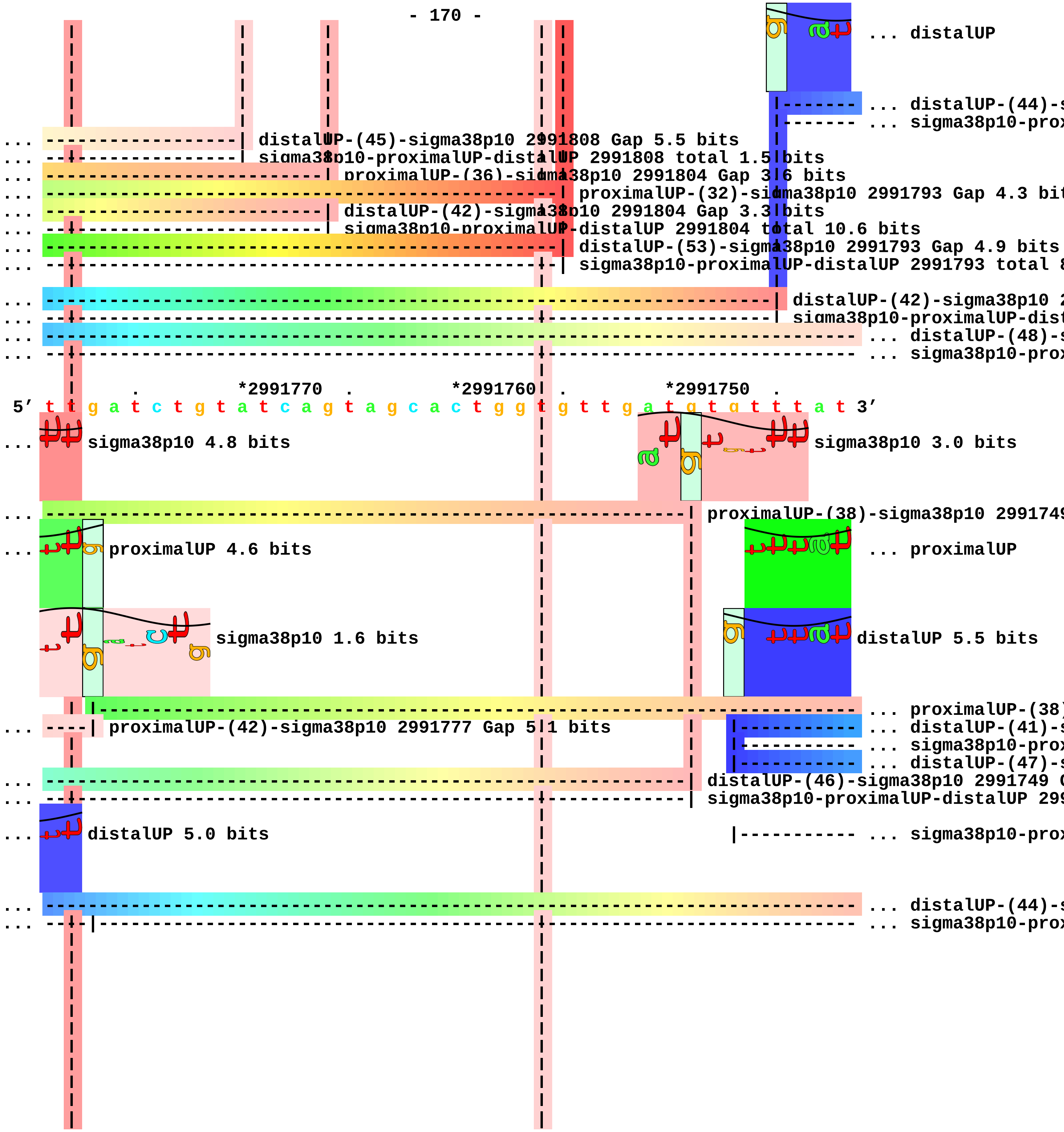
... distalUP
... distalUP-(44)-s
... sigma38p10-prox
... distalUP-(45)-sigma38p10 2991808 Gap 5.5 bits
... sigma38p10-proximalUP-distalUP 2991808 total 1.5 bits
... proximalUP-(36)-sigma38p10 2991804 Gap 3.6 bits
... proximalUP-(32)-sigma38p10 2991793 Gap 4.3 bit
... distalUP-(42)-sigma38p10 2991804 Gap 3.3 bits
... sigma38p10-proximalUP-distalUP 2991804 total 10.6 bits
... distalUP-(53)-sigma38p10 2991793 Gap 4.9 bits
... sigma38p10-proximalUP-distalUP 2991793 total 8
... distalUP-(42)-sigma38p10 2
... sigma38p10-proximalUP-dist
... distalUP-(48)-s
... sigma38p10-prox
*2991770
*2991760
*2991750
5′ t t g a t c t g t a t c a g t a g c a c t g g t g t t g a t g t g t t t a t 3′
... sigma38p10 4.8 bits
sigma38p10 3.0 bits
... proximalUP-(38)-sigma38p10 299174
... proximalUP 4.6 bits
... proximalUP
sigma38p10 1.6 bits
distalUP 5.5 bits
... proximalUP-(38)
... proximalUP-(42)-sigma38p10 2991777 Gap 5.1 bits
... distalUP-(41)-s
... sigma38p10-prox
... distalUP-(47)-s
... distalUP-(46)-sigma38p10 2991749 G
... sigma38p10-proximalUP-distalUP 299
... distalUP 5.0 bits
... sigma38p10-prox
... distalUP-(44)-s
... sigma38p10-prox

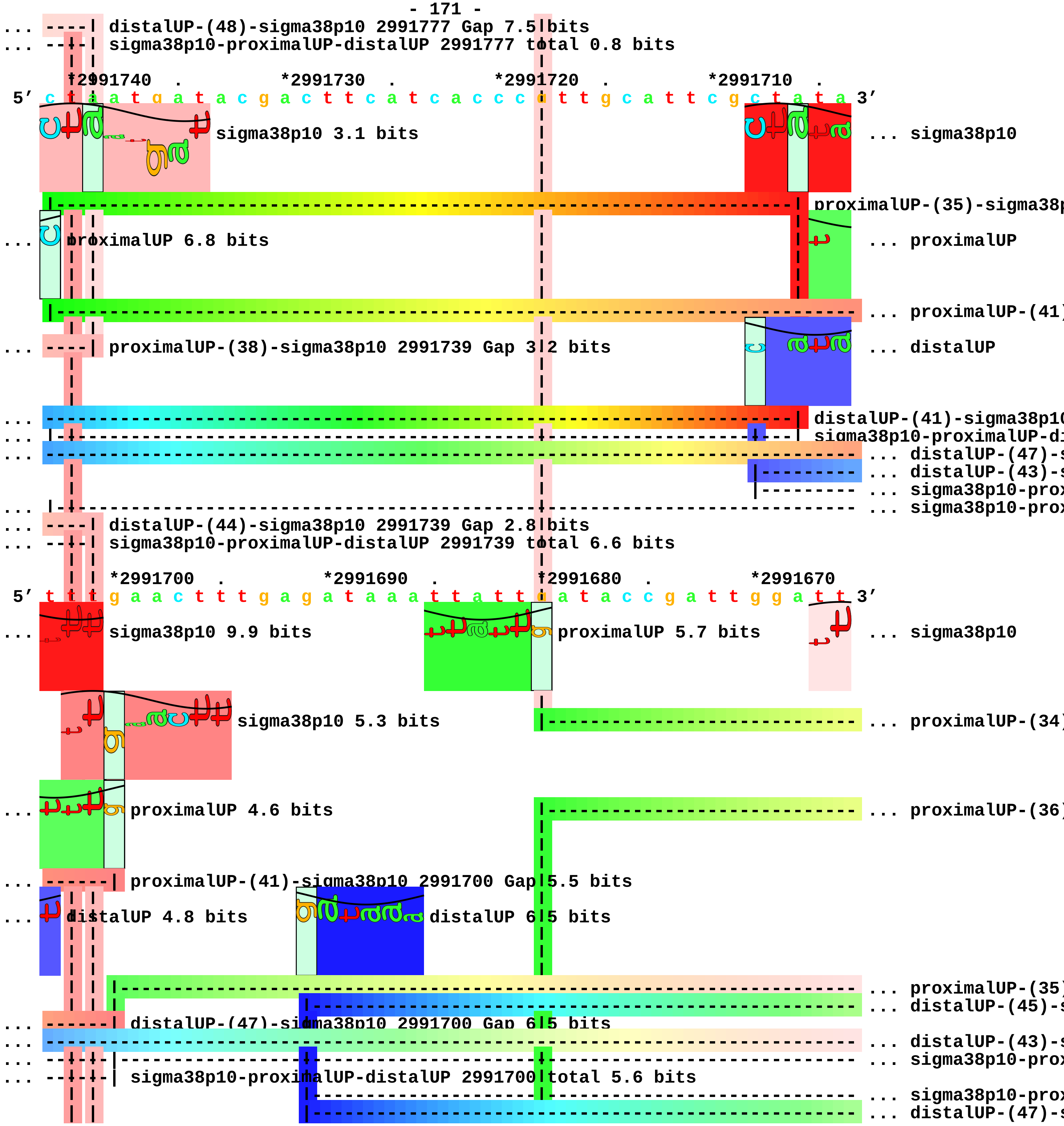
distalUP-(48)-sigma38p10 2991777 Gap 7.5 bits
sigma38p10-proximalUP-distalUP 2991777 total 0.8 bits
*2991740 . *2991730 . *2991720 . *2991710 .
5′ c t a a t g a t a c g a c t t c a t c a c c c a t t g c a t t c g c t a t a 3′
sigma38p10 3.1 bits
... sigma38p10
proximalUP-(35)-sigma38p
... proximalUP 6.8 bits
... proximalUP
... proximalUP-(41)
... proximalUP-(38)-sigma38p10 2991739 Gap 3.2 bits
... distalUP
distalUP-(41)-sigma38p10
sigma38p10-proximalUP-di
... distalUP-(47)-s
... distalUP-(43)-s
... sigma38p10-prox
... sigma38p10-prox
... distalUP-(44)-sigma38p10 2991739 Gap 2.8 bits
... sigma38p10-proximalUP-distalUP 2991739 total 6.6 bits
*2991700 . *2991690 . *2991680 . *2991670
5′ t t t g a a c t t t g a g a t a a a t t a t t a a t a c c g a t t g g a t t 3′
... sigma38p10 9.9 bits
proximalUP 5.7 bits
... sigma38p10
sigma38p10 5.3 bits
... proximalUP-(34)
... proximalUP 4.6 bits
... proximalUP-(36)
... proximalUP-(41)-sigma38p10 2991700 Gap 5.5 bits
... distalUP 4.8 bits
distalUP 6.5 bits
... proximalUP-(35)
... distalUP-(45)-s
... distalUP-(47)-sigma38p10 2991700 Gap 6.5 bits
... distalUP-(43)-s
... sigma38p10-prox
... sigma38p10-proximalUP-distalUP 2991700 total 5.6 bits
... sigma38p10-prox
... distalUP-(47)-s

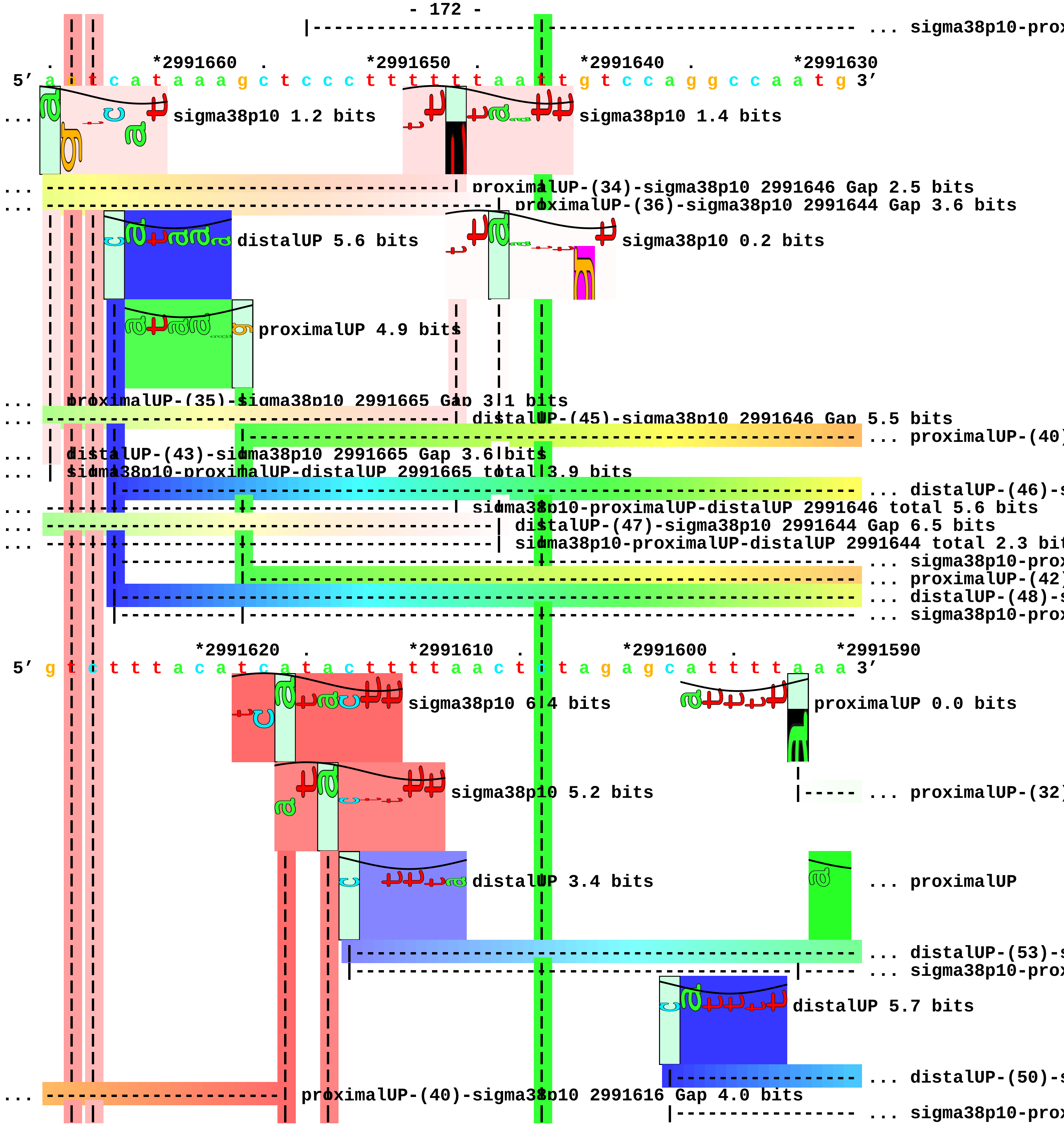
... sigma38p10-prox
*2991660 . *2991650 . *2991640 . *2991630
5′ a a t c a t a a a g c t c c c t t t t t t a a t t g t c c a g g c c a a t g 3′
... sigma38p10 1.2 bits
sigma38p10 1.4 bits
... proximalUP-(34)-sigma38p10 2991646 Gap 2.5 bits
... proximalUP-(36)-sigma38p10 2991644 Gap 3.6 bits
distalUP 5.6 bits
sigma38p10 0.2 bits
proximalUP 4.9 bits
... proximalUP-(35)-sigma38p10 2991665 Gap 3.1 bits
... distalUP-(45)-sigma38p10 2991646 Gap 5.5 bits
... proximalUP-(40)
... distalUP-(43)-sigma38p10 2991665 Gap 3.6 bits
... sigma38p10-proximalUP-distalUP 2991665 total 3.9 bits
... distalUP-(46)-s
... sigma38p10-proximalUP-distalUP 2991646 total 5.6 bits
... distalUP-(47)-sigma38p10 2991644 Gap 6.5 bits
... sigma38p10-proximalUP-distalUP 2991644 total 2.3 bit
... sigma38p10-prox
... proximalUP-(42)
... distalUP-(48)-s
... sigma38p10-prox
*2991620 . *2991610 . *2991600 . *2991590
5′ g t c t t t a c a t c a t a c t t t t t a a c t c t a g a g c a t t t t a a a 3′
sigma38p10 6.4 bits
proximalUP 0.0 bits
sigma38p10 5.2 bits
... proximalUP-(32)
distalUP 3.4 bits
... proximalUP
... distalUP-(53)-s
... sigma38p10-prox
distalUP 5.7 bits
... distalUP-(50)-s
... proximalUP-(40)-sigma38p10 2991616 Gap 4.0 bits
... sigma38p10-prox

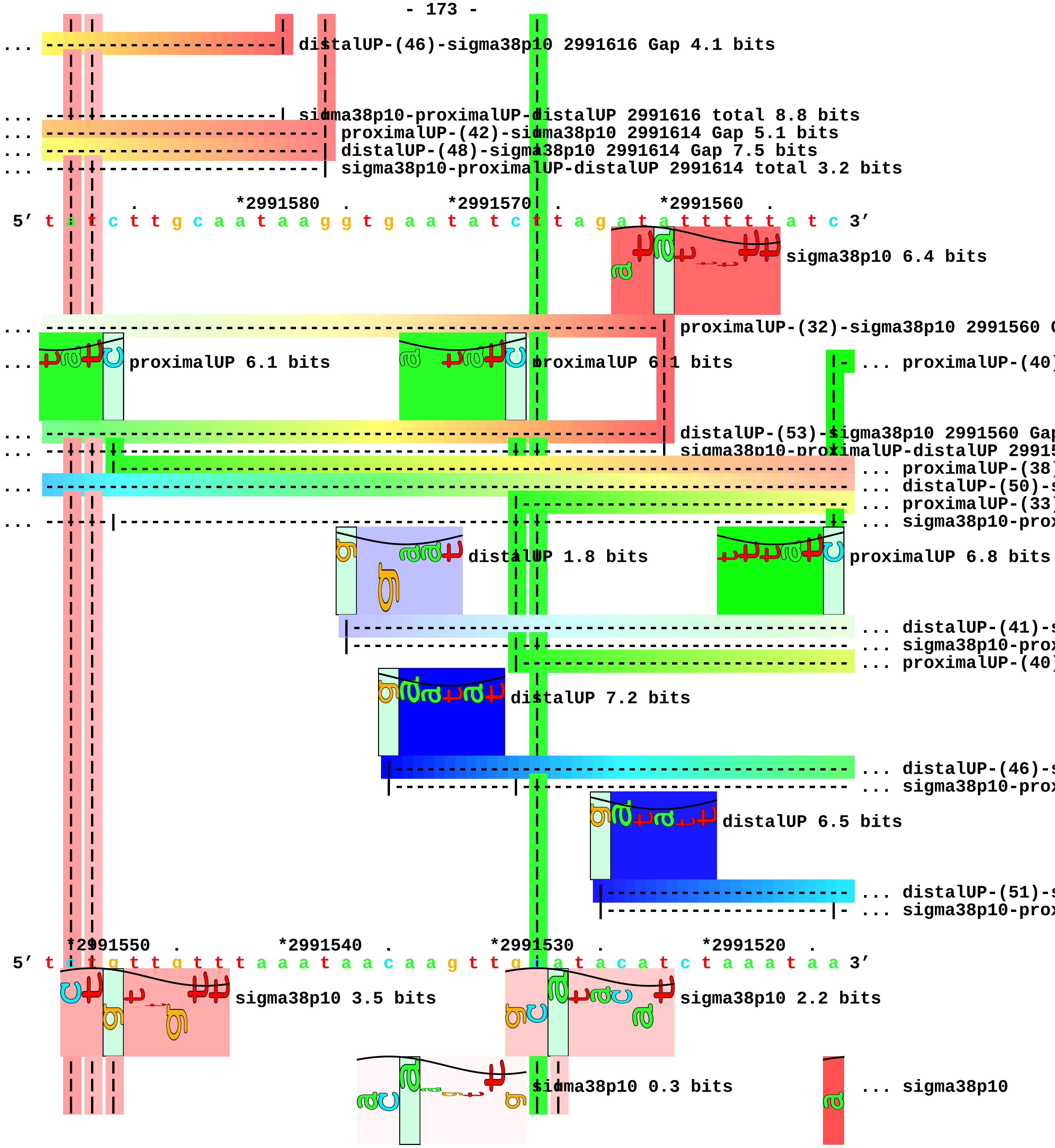
distalUP-(46)-sigma38p10 2991616 Gap 4.1 bits
sigma38p10-proximalUP-distalUP 2991616 total 8.8 bits
proximalUP-(42)-sigma38p10 2991614 Gap 5.1 bits
distalUP-(48)-sigma38p10 2991614 Gap 7.5 bits
sigma38p10-proximalUP-distalUP 2991614 total 3.2 bits
*2991580 . *2991570 . *2991560 .
5′ t a t c t t g c a a t a a g g t g a a t a t c t t a g a t a t t t t t t a t c 3′
sigma38p10 6.4 bits
proximalUP-(32)-sigma38p10 2991560 G
proximalUP 6.1 bits
proximalUP 6.1 bits
... proximalUP-(40)
distalUP-(53)-sigma38p10 2991560 Gap
sigma38p10-proximalUP-distalUP 29915
... proximalUP-(38)
... distalUP-(50)-s
... proximalUP-(33)
... sigma38p10-prox
distalUP 1.8 bits
proximalUP 6.8 bits
... distalUP-(41)-s
... sigma38p10-prox
... proximalUP-(40)
distalUP 7.2 bits
... distalUP-(46)-s
... sigma38p10-prox
distalUP 6.5 bits
... distalUP-(51)-s
... sigma38p10-prox
*2991550 . *2991540 . *2991530 . *2991520 .
5′ t c t g t t g t t t a a a t a a c a a g t t g c a t a c a t c t a a a t a a 3′
sigma38p10 3.5 bits
sigma38p10 2.2 bits
sigma38p10 0.3 bits
... sigma38p10

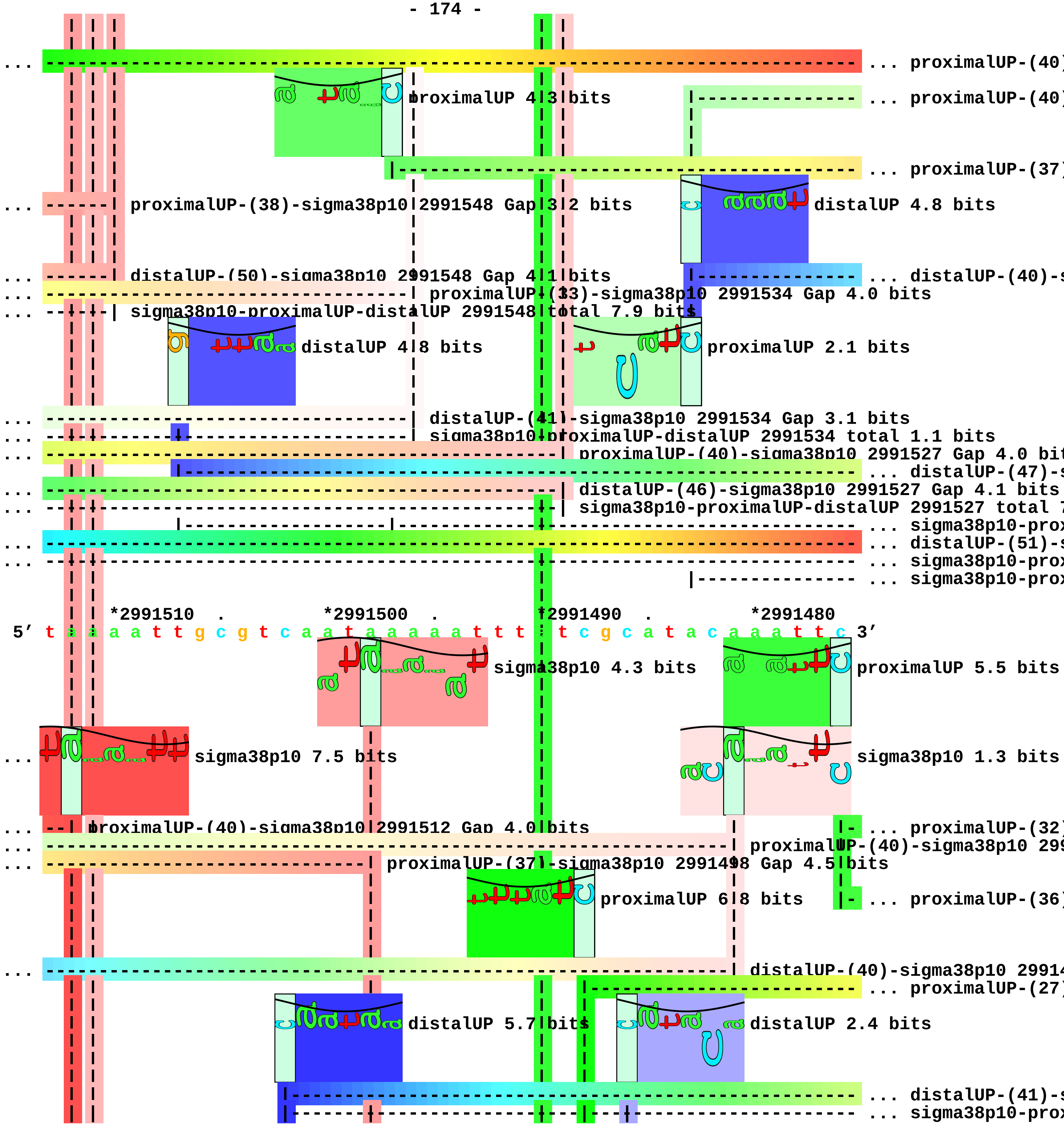
... proximalUP-(40)
proximalUP 4.3 bits
... proximalUP-(40)
... proximalUP-(37)
... proximalUP-(38)-sigma38p10 2991548 Gap 3.2 bits
distalUP 4.8 bits
... distalUP-(50)-sigma38p10 2991548 Gap 4.1 bits
... distalUP-(40)-s
... proximalUP-(33)-sigma38p10 2991534 Gap 4.0 bits
... sigma38p10-proximalUP-distalUP 2991548 total 7.9 bits
distalUP 4.8 bits
proximalUP 2.1 bits
... distalUP-(41)-sigma38p10 2991534 Gap 3.1 bits
... sigma38p10-proximalUP-distalUP 2991534 total 1.1 bits
... proximalUP-(40)-sigma38p10 2991527 Gap 4.0 bit
... distalUP-(47)-s
... distalUP-(46)-sigma38p10 2991527 Gap 4.1 bits
... sigma38p10-proximalUP-distalUP 2991527 total 7
... sigma38p10-prox
... distalUP-(51)-s
... sigma38p10-prox
... sigma38p10-prox
*2991510 . *2991500 . *2991490 . *2991480
5′ t a a a a t t g c g t c a a t a a a a a t t t t t c g c a t a c a a a t t c 3′
sigma38p10 4.3 bits
proximalUP 5.5 bits
... sigma38p10 7.5 bits
sigma38p10 1.3 bits
... proximalUP-(40)-sigma38p10 2991512 Gap 4.0 bits
... proximalUP-(32)
... proximalUP-(40)-sigma38p10 299
... proximalUP-(37)-sigma38p10 2991498 Gap 4.5 bits
proximalUP 6.8 bits
... proximalUP-(36)
... distalUP-(40)-sigma38p10 2991
... proximalUP-(27)
distalUP 5.7 bits
distalUP 2.4 bits
... distalUP-(41)-s
... sigma38p10-prox

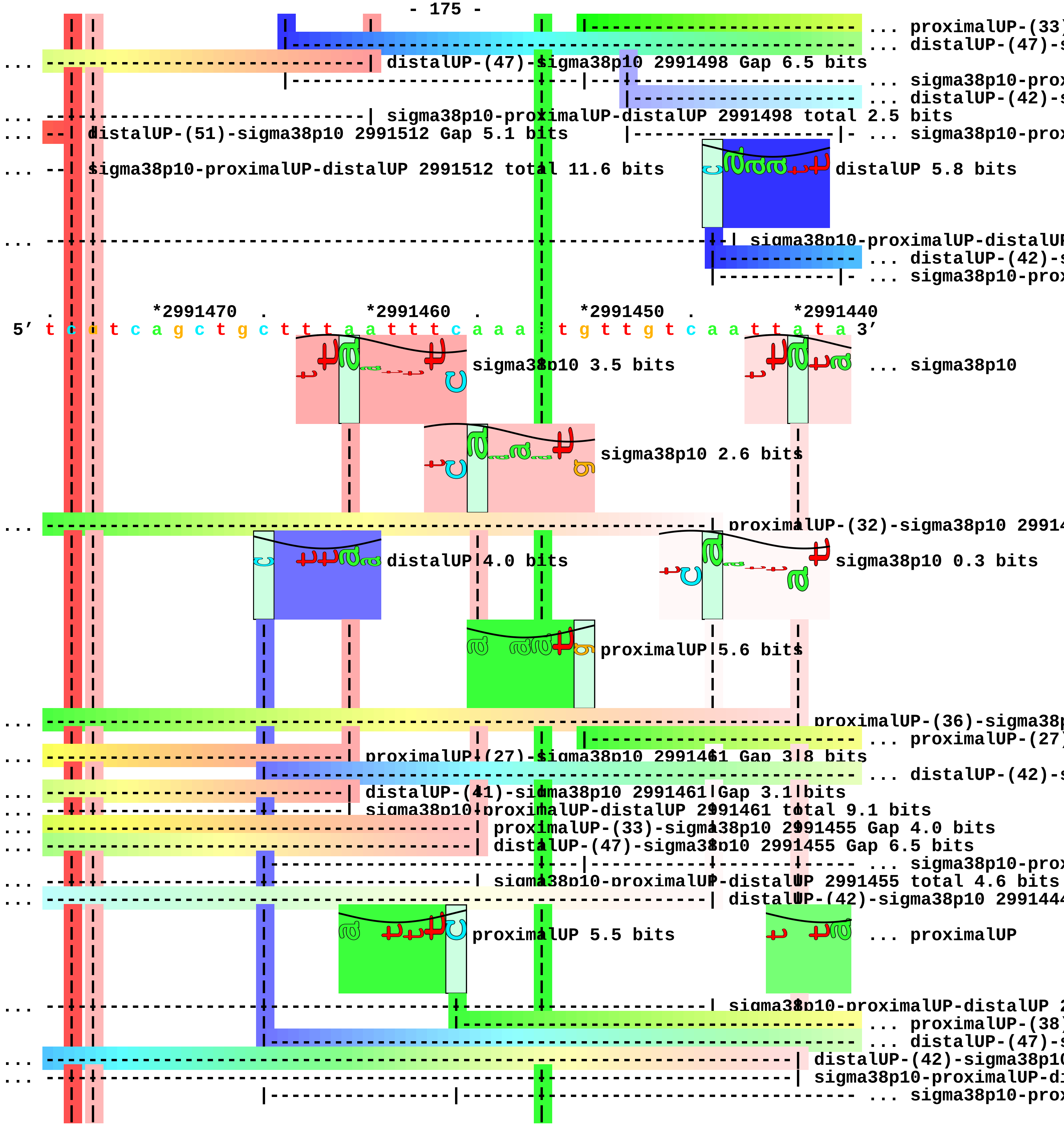
... proximalUP-(33)
... distalUP-(47)-s
... distalUP-(47)-sigma38p10 2991498 Gap 6.5 bits
... sigma38p10-prox
... distalUP-(42)-s
... sigma38p10-proximalUP-distalUP 2991498 total 2.5 bits
... distalUP-(51)-sigma38p10 2991512 Gap 5.1 bits
... sigma38p10-prox
... sigma38p10-proximalUP-distalUP 2991512 total 11.6 bits
distalUP 5.8 bits
sigma38p10-proximalUP-distalUP
... distalUP-(42)-s
... sigma38p10-prox
*2991470
*2991460
*2991450
*2991440
5′ t c c t c a g c t g c t t t a a t t t c a a a a t g t t g t c a a t t a t a 3′
sigma38p10 3.5 bits
... sigma38p10
sigma38p10 2.6 bits
... proximalUP-(32)-sigma38p10 29914
distalUP 4.0 bits
sigma38p10 0.3 bits
proximalUP 5.6 bits
... proximalUP-(36)-sigma38p
... proximalUP-(27)
... proximalUP-(27)-sigma38p10 2991461 Gap 3.8 bits
... distalUP-(42)-s
... distalUP-(41)-sigma38p10 2991461 Gap 3.1 bits
... sigma38p10-proximalUP-distalUP 2991461 total 9.1 bits
... proximalUP-(33)-sigma38p10 2991455 Gap 4.0 bits
... distalUP-(47)-sigma38p10 2991455 Gap 6.5 bits
... sigma38p10-prox
... sigma38p10-proximalUP-distalUP 2991455 total 4.6 bits
... distalUP-(42)-sigma38p10 29914
proximalUP 5.5 bits
... proximalUP
... sigma38p10-proximalUP-distalUP 2
... proximalUP-(38)
... distalUP-(47)-s
... distalUP-(42)-sigma38p10
... sigma38p10-proximalUP-di
... sigma38p10-prox

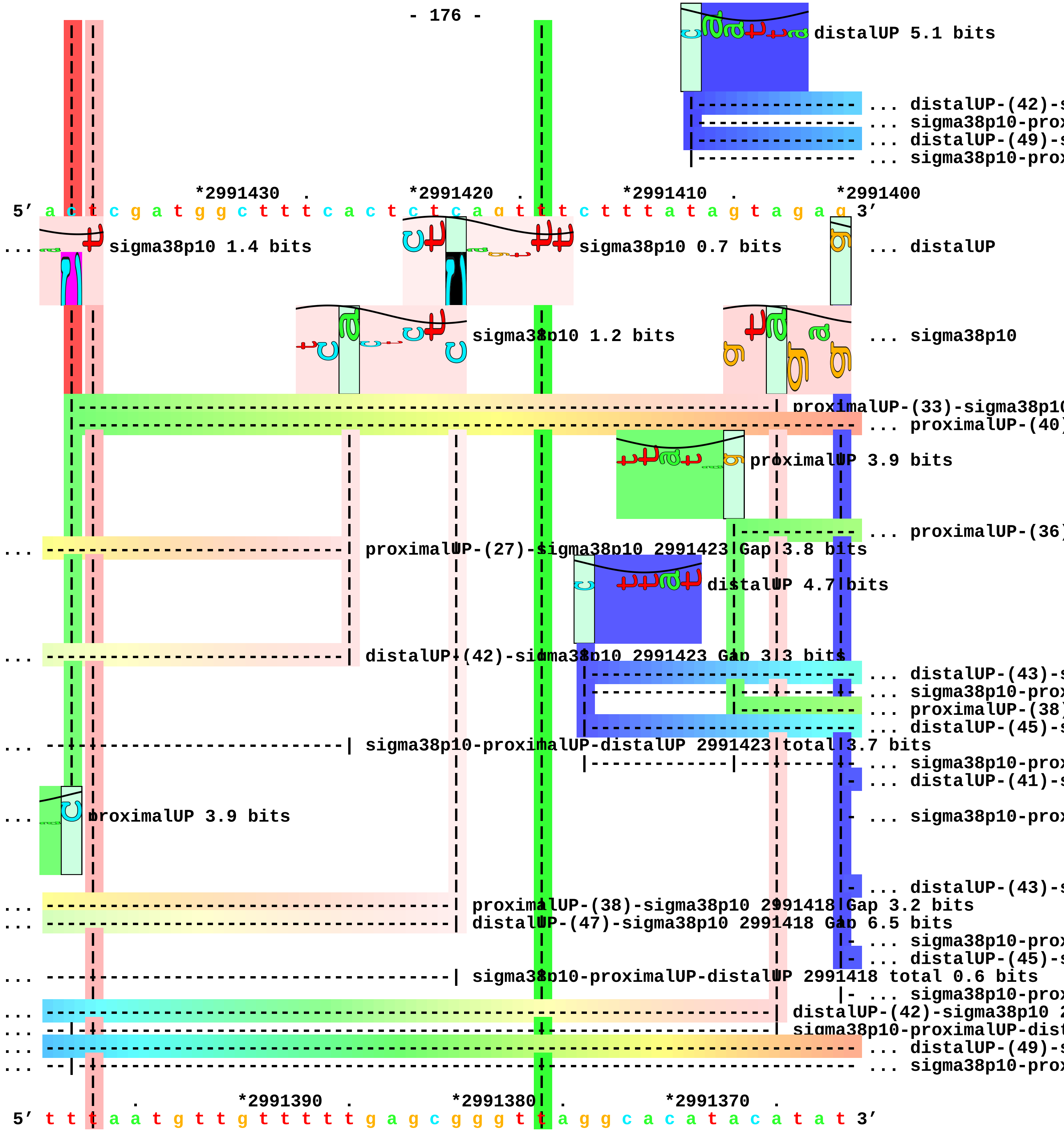
distalUP 5.1 bits
... distalUP-(42)-s
... sigma38p10-prox
... distalUP-(49)-s
... sigma38p10-prox
*2991430 . *2991420 . *2991410 . *2991400
5′ a c t c g a t g g c t t t c a c t c t c a g t t t t c t t t a t a g t a g a g 3′
... sigma38p10 1.4 bits
sigma38p10 0.7 bits
... distalUP
sigma38p10 1.2 bits
... sigma38p10
proximalUP-(33)-sigma38p10
... proximalUP-(40)
proximalUP 3.9 bits
... proximalUP-(36)
... proximalUP-(27)-sigma38p10 2991423 Gap 3.8 bits
distalUP 4.7 bits
... distalUP-(42)-sigma38p10 2991423 Gap 3.3 bits
... distalUP-(43)-s
... sigma38p10-prox
... proximalUP-(38)
... distalUP-(45)-s
... sigma38p10-proximalUP-distalUP 2991423 total 3.7 bits
... sigma38p10-prox
... distalUP-(41)-s
... proximalUP 3.9 bits
... sigma38p10-prox
... distalUP-(43)-s
... proximalUP-(38)-sigma38p10 2991418 Gap 3.2 bits
... distalUP-(47)-sigma38p10 2991418 Gap 6.5 bits
... sigma38p10-prox
... distalUP-(45)-s
... sigma38p10-proximalUP-distalUP 2991418 total 0.6 bits
... sigma38p10-prox
... distalUP-(42)-sigma38p10 2
... sigma38p10-proximalUP-dist
... distalUP-(49)-s
... sigma38p10-prox
. *2991390 . *2991380 . *2991370 .
5′ t t t a a t g t t g t t t t t g a g c g g g t t a g g c a c a t a c a t a t 3′

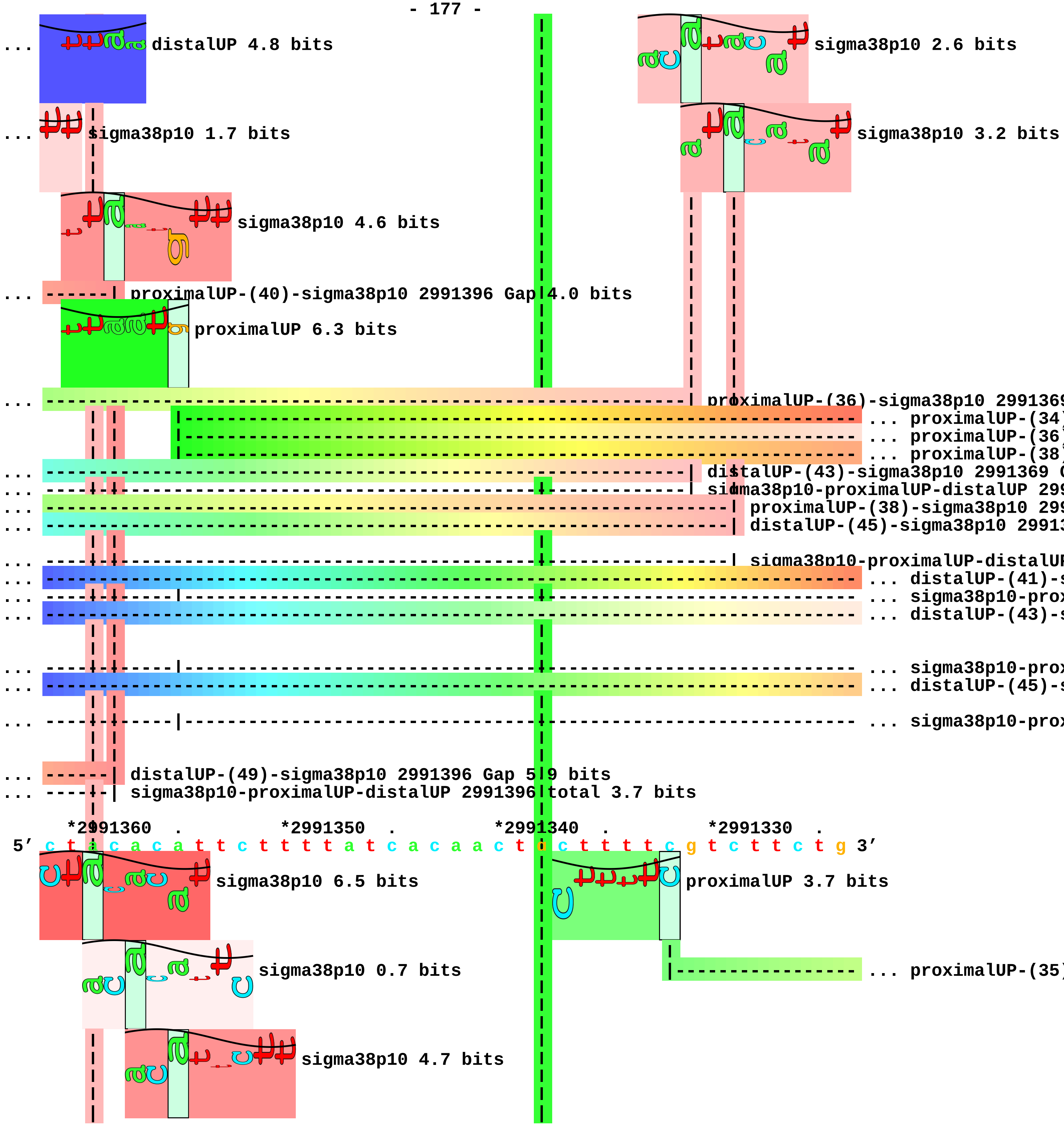
distalUP 4.8 bits
sigma38p10 2.6 bits
sigma38p10 1.7 bits
sigma38p10 3.2 bits
sigma38p10 4.6 bits
proximalUP-(40)-sigma38p10 2991396 Gap 4.0 bits
proximalUP 6.3 bits
proximalUP-(36)-sigma38p10 299136
proximalUP-(34
proximalUP-(36
proximalUP-(38
distalUP-(43)-sigma38p10 2991369
sigma38p10-proximalUP-distalUP 299
proximalUP-(38)-sigma38p10 299
distalUP-(45)-sigma38p10 29913
sigma38p10-proximalUP-distalUP
distalUP-(41)-s
sigma38p10-prox
distalUP-(43)-s
sigma38p10-prox
distalUP-(45)-s
sigma38p10-prox
distalUP-(49)-sigma38p10 2991396 Gap 5.9 bits
sigma38p10-proximalUP-distalUP 2991396 total 3.7 bits
*2991360 . *2991350 . *2991340 . *2991330 .
5' c t a c a c a t t c t t t t a t c a c a a c t g c t t t t c g t c t t c t g 3'
sigma38p10 6.5 bits
proximalUP 3.7 bits
sigma38p10 0.7 bits
proximalUP-(35
sigma38p10 4.7 bits

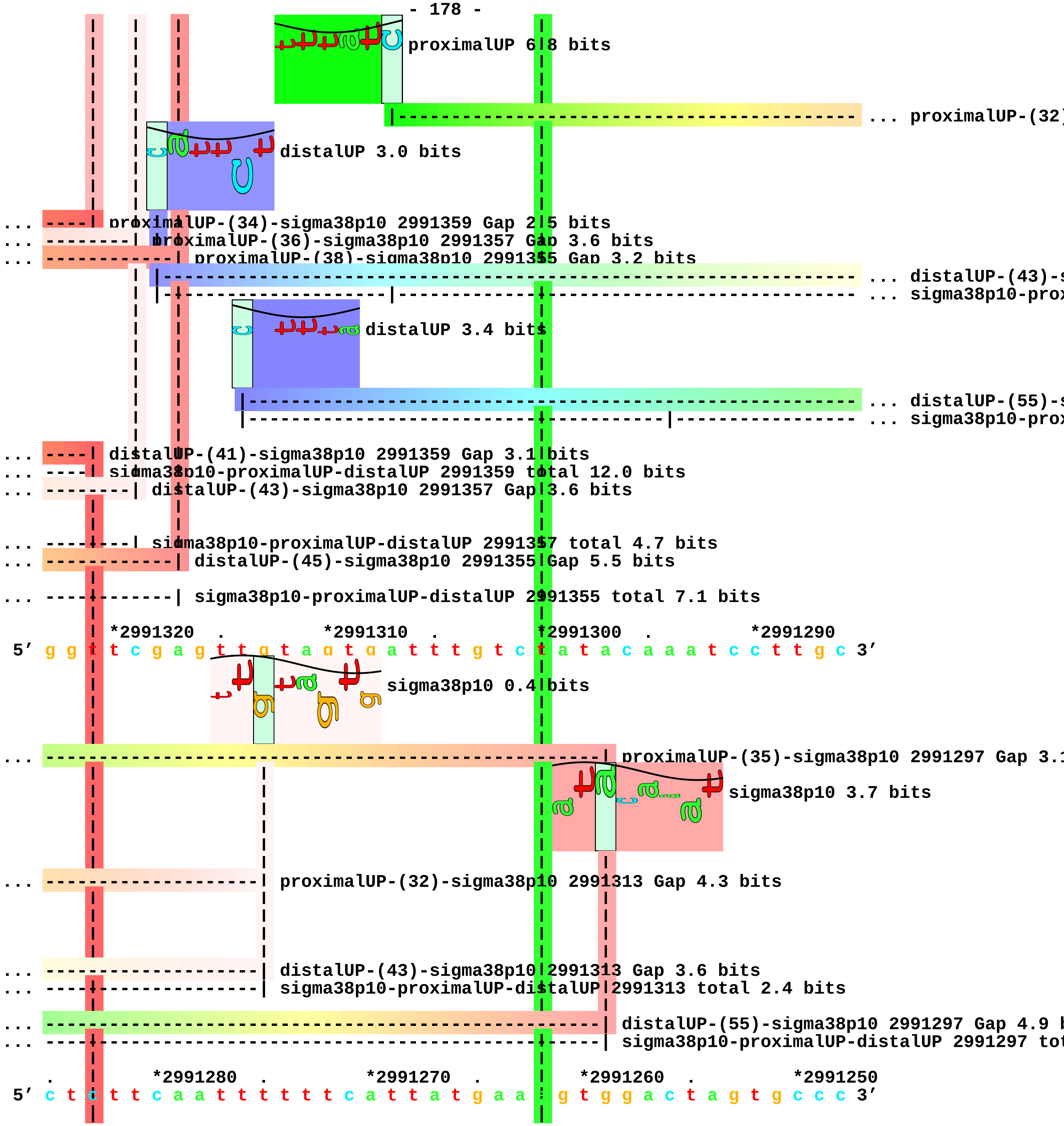
proximalUP 6.8 bits
... proximalUP-(32)
distalUP 3.0 bits
... proximalUP-(34)-sigma38p10 2991359 Gap 2.5 bits
... proximalUP-(36)-sigma38p10 2991357 Gap 3.6 bits
... proximalUP-(38)-sigma38p10 2991355 Gap 3.2 bits
... distalUP-(43)-s
... sigma38p10-prox
distalUP 3.4 bits
... distalUP-(55)-s
... sigma38p10-prox
... distalUP-(41)-sigma38p10 2991359 Gap 3.1 bits
... sigma38p10-proximalUP-distalUP 2991359 total 12.0 bits
... distalUP-(43)-sigma38p10 2991357 Gap 3.6 bits
... sigma38p10-proximalUP-distalUP 2991357 total 4.7 bits
... distalUP-(45)-sigma38p10 2991355 Gap 5.5 bits
... sigma38p10-proximalUP-distalUP 2991355 total 7.1 bits
*2991320 . *2991310 . *2991300 . *2991290
5' g g t t c g a g t t g t a g t g a t t t g t c t a t a c a a a t c c t t g c 3'
sigma38p10 0.4 bits
... proximalUP-(35)-sigma38p10 2991297 Gap 3.1
sigma38p10 3.7 bits
... proximalUP-(32)-sigma38p10 2991313 Gap 4.3 bits
... distalUP-(43)-sigma38p10 2991313 Gap 3.6 bits
... sigma38p10-proximalUP-distalUP 2991313 total 2.4 bits
... distalUP-(55)-sigma38p10 2991297 Gap 4.9 b
... sigma38p10-proximalUP-distalUP 2991297 tot
. *2991280 . *2991270 . *2991260 . *2991250
5' c t c t t c a a t t t t t t c a t t a t g a a a g t g g a c t a g t g c c c 3'

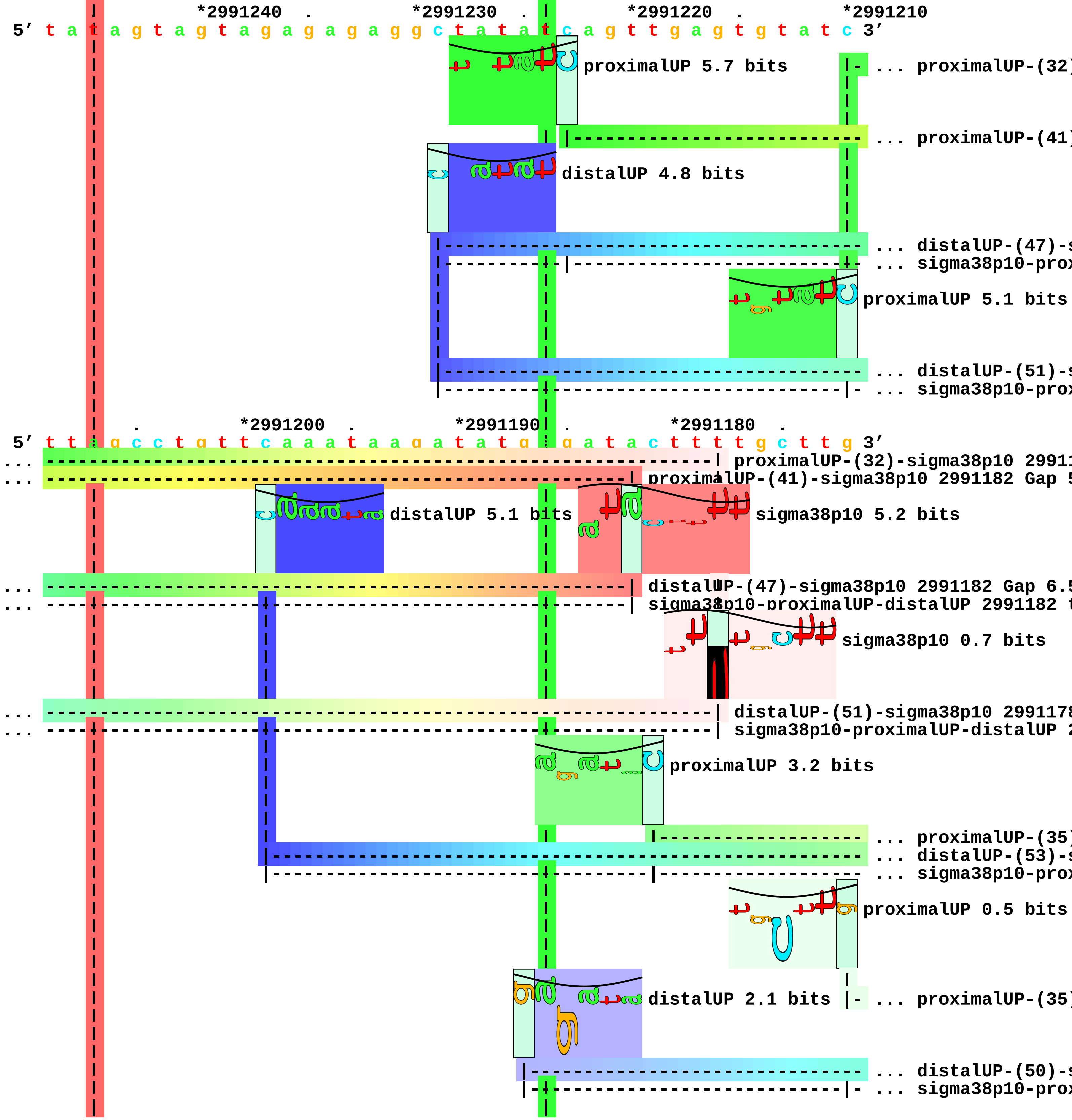
*2991240 . *2991230 . *2991220 . *2991210
5′ t a t a g t a g t a g a g a g g c t a t a t c a g t t g a g t g t a t c 3′
proximalUP 5.7 bits
... proximalUP-(32)
... proximalUP-(41)
distalUP 4.8 bits
... distalUP-(47)-s
... sigma38p10-prox
proximalUP 5.1 bits
... distalUP-(51)-s
... sigma38p10-prox
. *2991200 . *2991190 . *2991180 .
5′ t t a g c c t g t t c a a a t a a g a t a t g t g a t a c t t t t g c t t g 3′
proximalUP-(32)-sigma38p10 29911
proximalUP-(41)-sigma38p10 2991182 Gap 5
distalUP 5.1 bits
sigma38p10 5.2 bits
distalUP-(47)-sigma38p10 2991182 Gap 6.5
sigma38p10-proximalUP-distalUP 2991182 t
sigma38p10 0.7 bits
distalUP-(51)-sigma38p10 2991178
sigma38p10-proximalUP-distalUP 2
proximalUP 3.2 bits
... proximalUP-(35)
... distalUP-(53)-s
... sigma38p10-prox
proximalUP 0.5 bits
distalUP 2.1 bits
... proximalUP-(35)
... distalUP-(50)-s
... sigma38p10-prox

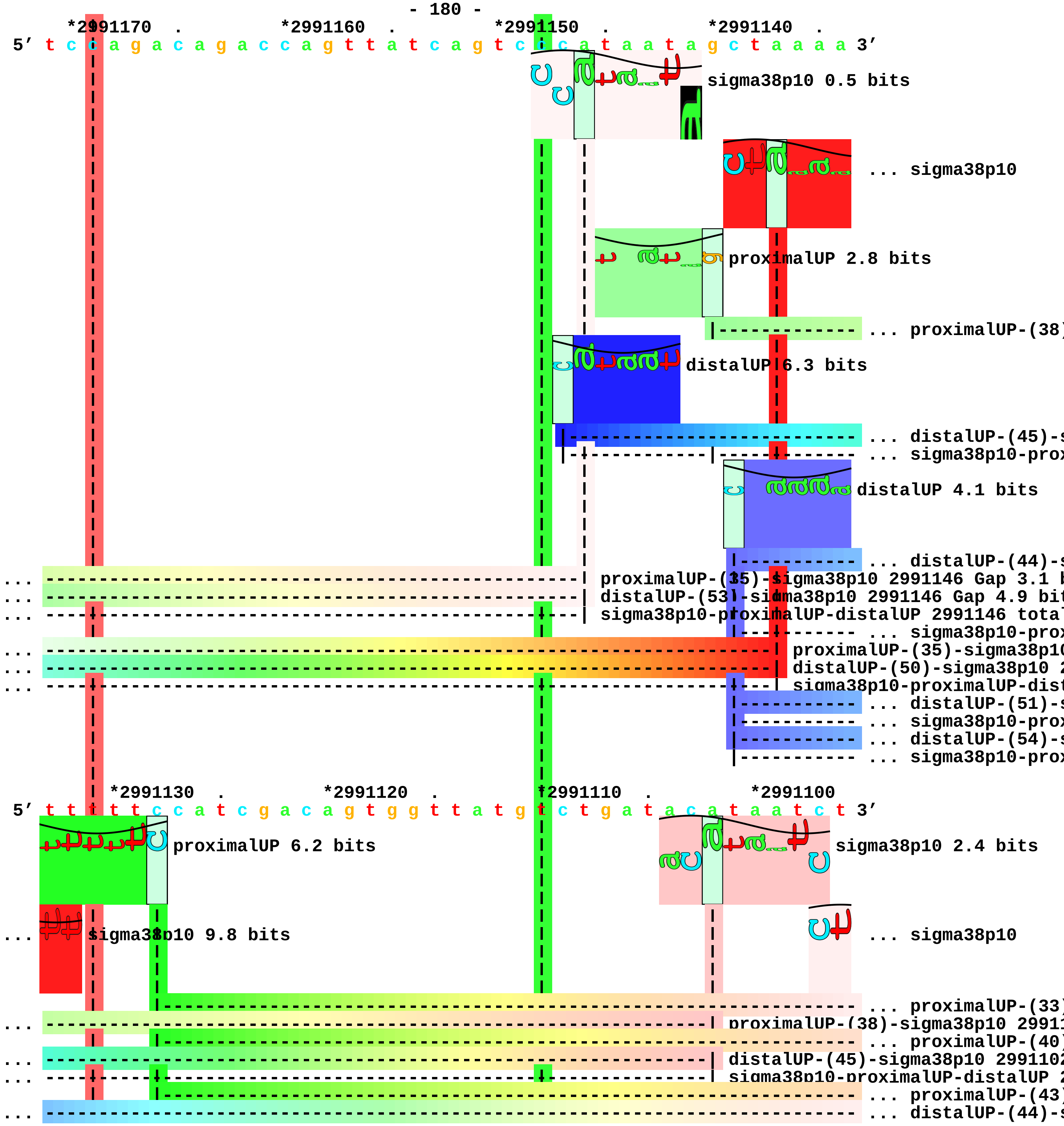
*2991170 . *2991160 . *2991150 . *2991140 .
5′ t c c a g a c a g a c c a g t t a t c a g t c c c a t a a t a g c t a a a a 3′
sigma38p10 0.5 bits
... sigma38p10
proximalUP 2.8 bits
... proximalUP-(38)
distalUP 6.3 bits
... distalUP-(45)-s
... sigma38p10-prox
distalUP 4.1 bits
... distalUP-(44)-s
... proximalUP-(35)-sigma38p10 2991146 Gap 3.1 b
... distalUP-(53)-sigma38p10 2991146 Gap 4.9 bit
... sigma38p10-proximalUP-distalUP 2991146 total
... sigma38p10-prox
... proximalUP-(35)-sigma38p10
... distalUP-(50)-sigma38p10 2
... sigma38p10-proximalUP-dist
... distalUP-(51)-s
... sigma38p10-prox
... distalUP-(54)-s
... sigma38p10-prox
*2991130 . *2991120 . *2991110 . *2991100
5′ t t t t t c c a t c g a c a g t g g t t a t g t c t g a t a c a t a a t c t 3′
proximalUP 6.2 bits
sigma38p10 2.4 bits
... sigma38p10 9.8 bits
... sigma38p10
... proximalUP-(33)
... proximalUP-(38)-sigma38p10 29911
... proximalUP-(40)
... distalUP-(45)-sigma38p10 2991102
... sigma38p10-proximalUP-distalUP 2
... proximalUP-(43)
... distalUP-(44)-s

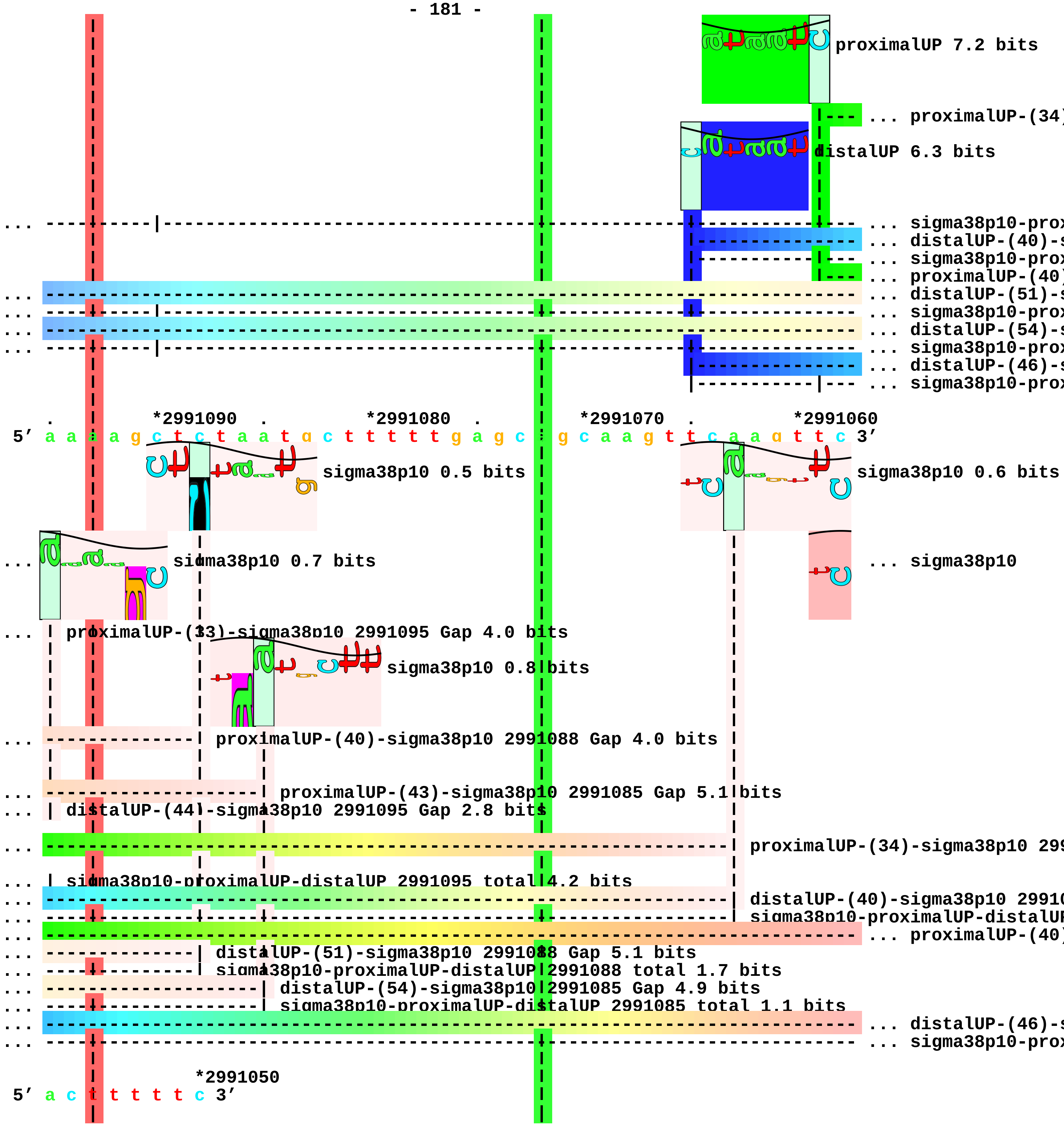
proximalUP 7.2 bits
... proximalUP-(34)
distalUP 6.3 bits
... sigma38p10-prox
... distalUP-(40)-s
... sigma38p10-prox
... proximalUP-(40)
... distalUP-(51)-s
... sigma38p10-prox
... distalUP-(54)-s
... sigma38p10-prox
... distalUP-(46)-s
... sigma38p10-prox
*2991090 *2991080 *2991070 *2991060
5′ a a a a g c t c t a a t g c t t t t t g a g c t g c a a g t t c a a g t t c 3′
sigma38p10 0.5 bits
sigma38p10 0.6 bits
sigma38p10 0.7 bits
... sigma38p10
proximalUP-(33)-sigma38p10 2991095 Gap 4.0 bits
sigma38p10 0.8 bits
proximalUP-(40)-sigma38p10 2991088 Gap 4.0 bits
proximalUP-(43)-sigma38p10 2991085 Gap 5.1 bits
distalUP-(44)-sigma38p10 2991095 Gap 2.8 bits
proximalUP-(34)-sigma38p10 299
sigma38p10-proximalUP-distalUP 2991095 total 4.2 bits
distalUP-(40)-sigma38p10 2991
sigma38p10-proximalUP-distalUP
... proximalUP-(40)
distalUP-(51)-sigma38p10 2991088 Gap 5.1 bits
sigma38p10-proximalUP-distalUP 2991088 total 1.7 bits
distalUP-(54)-sigma38p10 2991085 Gap 4.9 bits
sigma38p10-proximalUP-distalUP 2991085 total 1.1 bits
... distalUP-(46)-s
... sigma38p10-prox
*2991050
5′ a c t t t t t c 3′

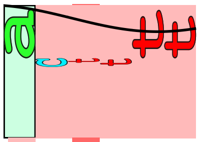

... sigma38p10 2.9 bits

... | proximalUP-(40)-sigma38p10 2991057 Gap 4.0 bits

... | distalUP-(46)-sigma38p10 2991057 Gap 4.1 bits
... | sigma38p10-proximalUP-distalUP 2991057 total 8.3 bits

piece 42, NC_000913, sibC, config: linear, direction: -, begin: 3055002, end: 3054742

```
      *3055000   .         *3054990   .         *3054980   .         *3054970   .
5' c a c c g a a g c g a g g g c t t g a a g g a g a a g g g t t a t g a t g c 3'
 T timestamp:_2018Sep20_11-35-24

          *3054960   .         *3054950   .         *3054940   .         *3054930
5' g a c t t g t c a t c a t a c t g a t t g t a c t g t t a c t c a t a a g t 3'

    .         *3054920   .         *3054910   .         *3054900   .         *3054890
5' t t c a g c g c t t a t t a a c a g t c a g t c t c a g g g g a g g a g c a 3'
```

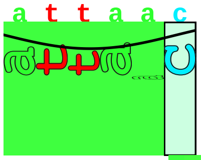

proximalUP 5.4 bits

|----------------------------------------------- ... proximalUP-(33)

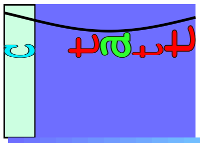

distalUP 4.1 bits

|--------------------------------------------------------------- ... distalUP-(41)-s
|---------------|----------------------------------------------- ... sigma38p10-prox

```
      .         *3054880   .         *3054870   .         *3054860   .
5' a t c c t c c c t t a c c c t t a c t c a c t a a a t t a g g t c a a a g a 3'
```

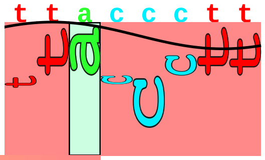

sigma38p10 5.1 bits

... --------------------| proximalUP-(33)-sigma38p10 3054878 Gap 4.0 bits
... --------------------| distalUP-(41)-sigma38p10 3054878 Gap 3.1 bits
... --------------------| sigma38p10-proximalUP-distalUP 3054878 total 7.5 bits

```
   *3054850   .         *3054840   .         *3054830   .         *3054820   .
5' a t c a a c g a t g t c a a t c a g g g c g a t g c g g g t g t a t c g c c 3'

      *3054810   .         *3054800   .         *3054790   .         *3054780   .
5' c t t a c c a c t c c c a g a c t t t c g a c g g t g t a a c c a c c g c a 3'

          *3054770   .         *3054760   .         *3054750   .
5' g g a a g a g g g a t a t c c c a c t c t t c a a c g g g g a g t 3'
```

piece 43, NC_000913, scpA, config: linear, direction: -, begin: 3058799, end: 3058550

```
              .         *3058790  .         *3058780  .         *3058770  .
5' c c a g c a a a a a a a a a t g c a t t c a g a c g t c g t t t t t g c c g 3'
```

proximalUP 5.6 bits

... proximalUP-(38

distalUP 5.7 bits

... distalUP-(48)-s
... sigma38p10-prox
... proximalUP-(43)
... distalUP-(53)-s
... sigma38p10-prox

T timestamp:_2018Sep20_11-35-24

```
    *3058760  .         *3058750  .         *3058740  .         *3058730  .
5' c c t g a g a t t t g t c t a t t t t g t g a t g t g a t t c a a g g a a t 3'
```

sigma38p10 6.2 bits

... proximalUP-(33)

... proximalUP-(38)-sigma38p10 3058747 Gap 3.2 bits

sigma38p10 1.5 bits

... distalUP-(48)-sigma38p10 3058747 Gap 7.5 bits
... sigma38p10-proximalUP-distalUP 3058747 total 6.8 bits
... proximalUP-(43)-sigma38p10 3058742 Gap 5.1 bits
... distalUP-(53)-sigma38p10 3058742 Gap 4.9 bits
... sigma38p10-proximalUP-distalUP 3058742 total 2.8 bits

proximalUP 4.5 bits

distalUP 4.2 bits

... distalUP-(51)-s
... sigma38p10-prox

```
      *3058720  .         *3058710  .         *3058700  .         *3058690
5' a c a a a a a g a a g t g a g g a c a a c g c c t g a t a t g t g c t a t c 3'
```

... proximalUP-(33)-sigma38p10 3058698 Gap 4.0 b

proximalUP 3.8 bits

sigma38p10 2.7 bits

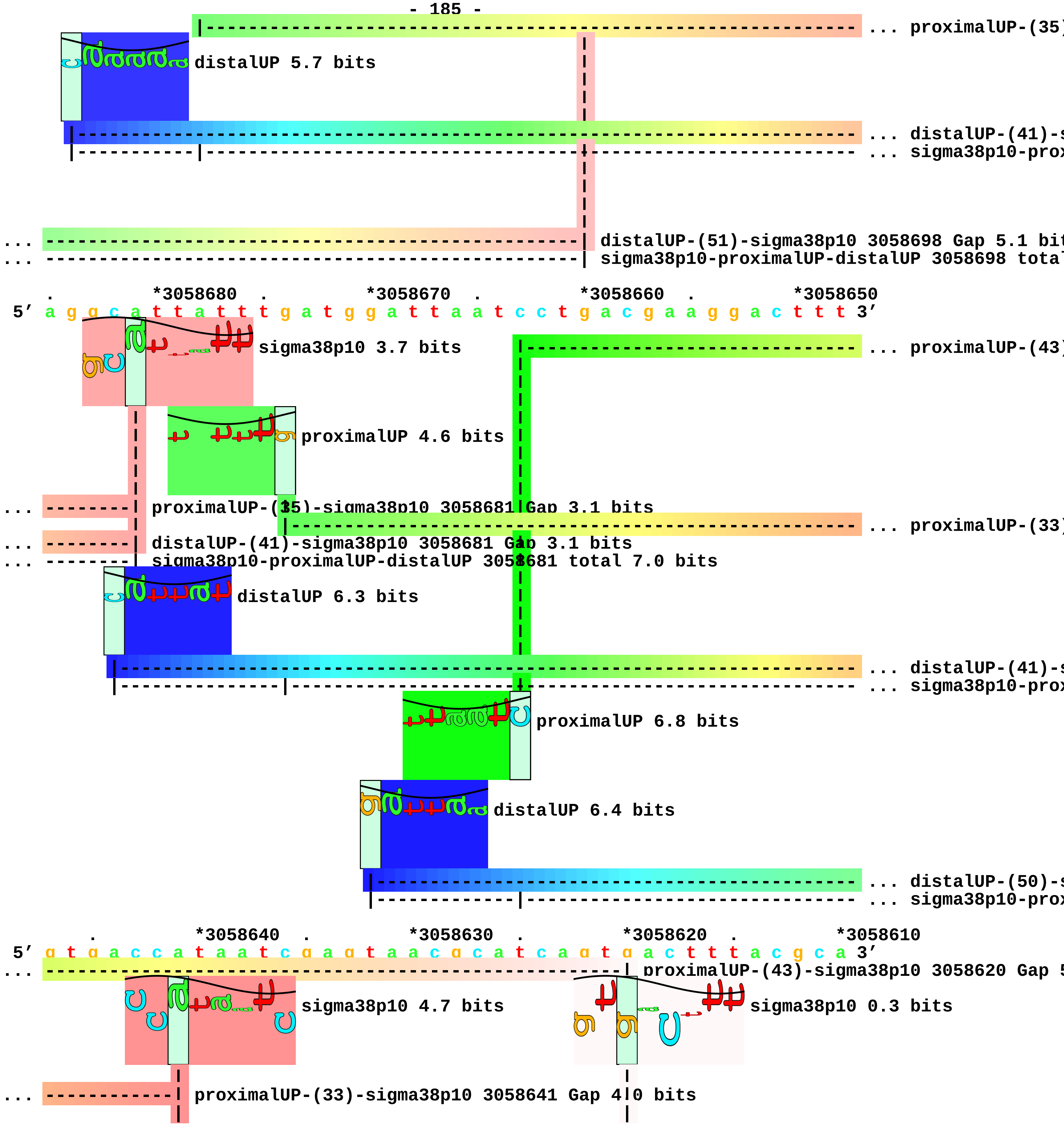
... proximalUP-(35)
distalUP 5.7 bits
... distalUP-(41)-s
... sigma38p10-prox
... distalUP-(51)-sigma38p10 3058698 Gap 5.1 bit
... sigma38p10-proximalUP-distalUP 3058698 total
*3058680
*3058670
*3058660
*3058650
5' a g g c a t t a t t t g a t g g a t t a a t c c t g a c g a a g g a c t t t 3'
sigma38p10 3.7 bits
... proximalUP-(43)
proximalUP 4.6 bits
... proximalUP-(35)-sigma38p10 3058681 Gap 3.1 bits
... proximalUP-(33)
... distalUP-(41)-sigma38p10 3058681 Gap 3.1 bits
... sigma38p10-proximalUP-distalUP 3058681 total 7.0 bits
distalUP 6.3 bits
... distalUP-(41)-s
... sigma38p10-prox
proximalUP 6.8 bits
distalUP 6.4 bits
... distalUP-(50)-s
... sigma38p10-prox
*3058640
*3058630
*3058620
*3058610
5' g t g a c c a t a a t c g a g t a a c g c a t c a g t g a c t t t a c g c a 3'
... proximalUP-(43)-sigma38p10 3058620 Gap 5
sigma38p10 4.7 bits
sigma38p10 0.3 bits
... proximalUP-(33)-sigma38p10 3058641 Gap 4.0 bits

```
                 |                                        |
                 |                                        |
... ------------| distalUP-(41)-sigma38p10 3058641 Gap 3.1|bits
... ------------| sigma38p10-proximalUP-distalUP 3058641 total 8.4 bits
                                                          |
                                                          |
... ------------------------------------------------------| distalUP-(50)-sigma38p10 3058620 Gap 4.1
... ------------------------------------------------------| sigma38p10-proximalUP-distalUP 3058620 t

         .         *3058600  .         *3058590  .         *3058580  .
 5' t c a t g c g g c t t t c a g g a g c a a a g c g g t g c c a g t a g a g c 3'

      *3058570  .         *3058560  .         *3058550
 5' a t c c g t c g t t g a a a t a g c c c a g 3'
```

piece 44, NC_000913, yggE, config: linear, direction: +, begin: 3066000, end: 3066199

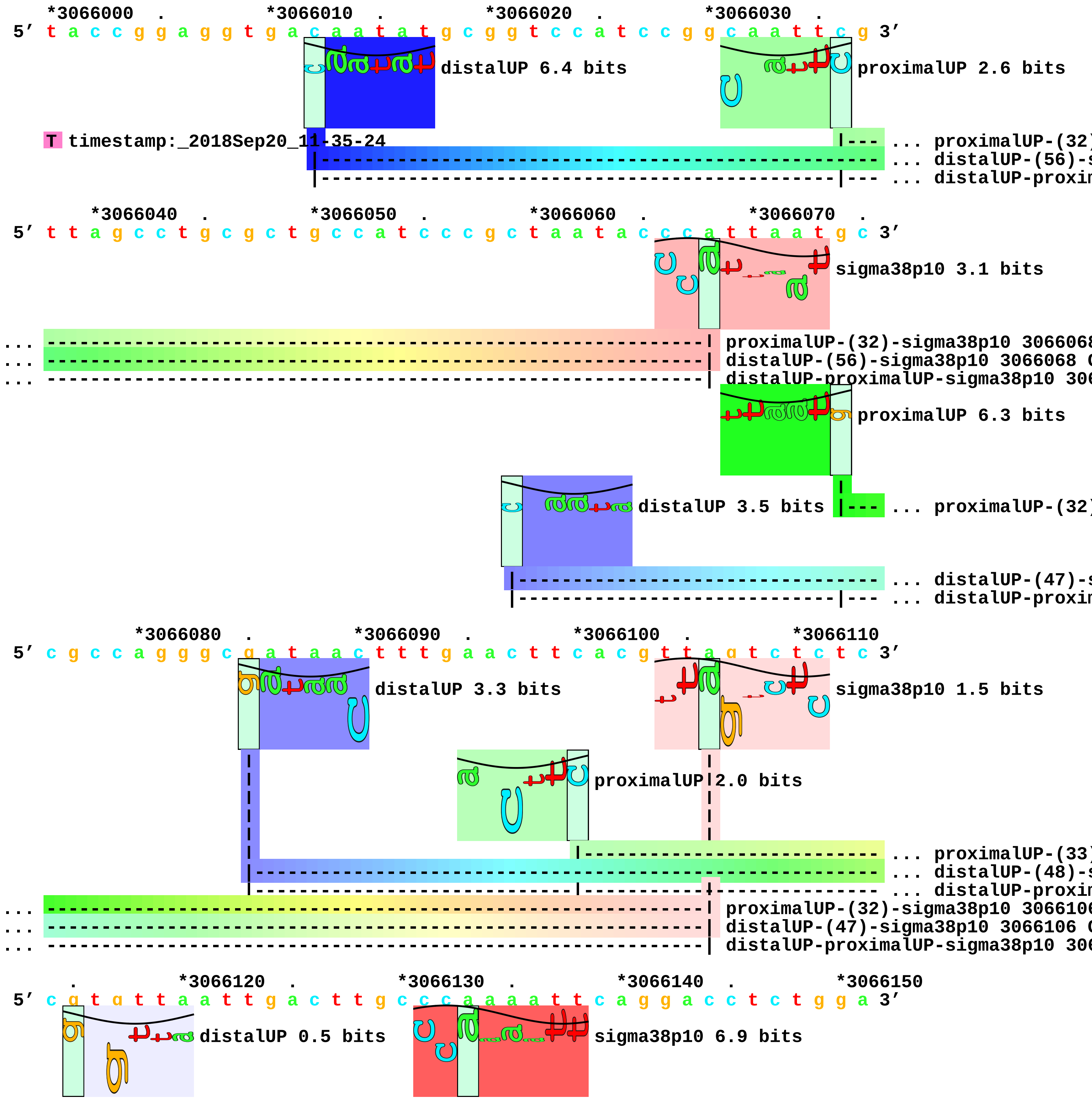

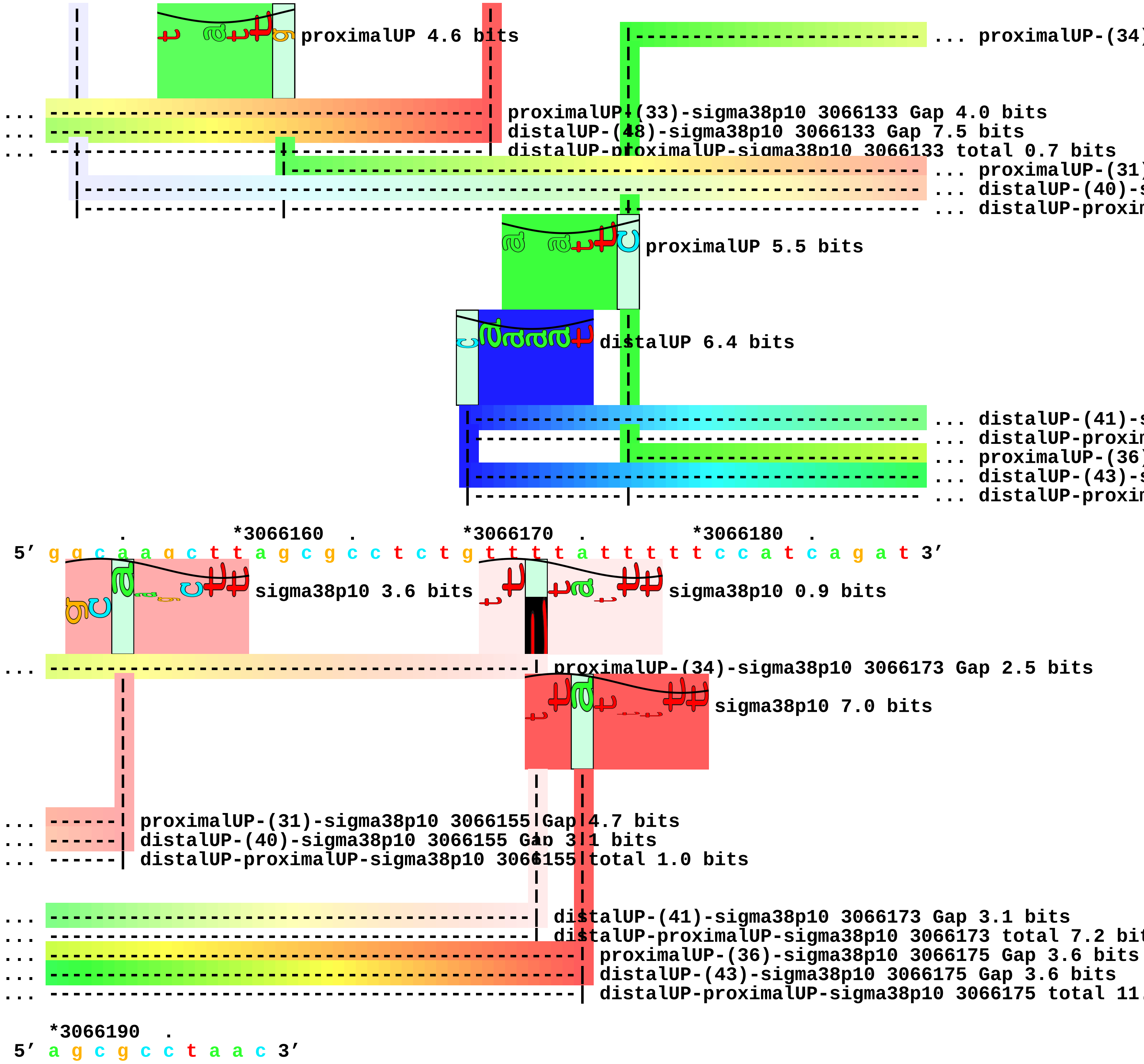
proximalUP 4.6 bits
... proximalUP-(34)
proximalUP-(33)-sigma38p10 3066133 Gap 4.0 bits
distalUP-(48)-sigma38p10 3066133 Gap 7.5 bits
distalUP-proximalUP-sigma38p10 3066133 total 0.7 bits
... proximalUP-(31)
... distalUP-(40)-s
... distalUP-proxin
proximalUP 5.5 bits
distalUP 6.4 bits
... distalUP-(41)-s
... distalUP-proxin
... proximalUP-(36)
... distalUP-(43)-s
... distalUP-proxin
*3066160 . *3066170 . *3066180 .
5′ g g c a a g c t t a g c g c c t c t g t t t t a t t t t t c c a t c a g a t 3′
sigma38p10 3.6 bits
sigma38p10 0.9 bits
proximalUP-(34)-sigma38p10 3066173 Gap 2.5 bits
sigma38p10 7.0 bits
proximalUP-(31)-sigma38p10 3066155 Gap 4.7 bits
distalUP-(40)-sigma38p10 3066155 Gap 3.1 bits
distalUP-proximalUP-sigma38p10 3066155 total 1.0 bits
distalUP-(41)-sigma38p10 3066173 Gap 3.1 bits
distalUP-proximalUP-sigma38p10 3066173 total 7.2 bit
proximalUP-(36)-sigma38p10 3066175 Gap 3.6 bits
distalUP-(43)-sigma38p10 3066175 Gap 3.6 bits
distalUP-proximalUP-sigma38p10 3066175 total 11.
*3066190 .
5′ a g c g c c t a a c 3′

piece 45, NC_000913, ygjR, config: linear, direction: -, begin: 3235431, end: 3235183

```
   *3235430  .         *3235420  .         *3235410  .         *3235400  .
5' g g a a t a t a c g g c g g t t a a c t t g t a t t t a c c g c t c t c a t 3'
 T timestamp:_2018Sep20_11-35-24

       *3235390  .         *3235380  .         *3235370  .         *3235360
5' g g g c g g c c t c g a c g a a c t g g c g a g t g a t c c a g t t c g t a 3'

 .         *3235350  .         *3235340  .         *3235330  .         *3235320
5' c c a a t c a c a g c g a a a c g t a t c a t a a c g c g t g g a t a c t c 3'

     .         *3235310  .         *3235300  .         *3235290  .         *3235280
5' c a t a a a g g g a t t c t g t c g t c a a t g t a g c a c g c c t t t t t 3'
```

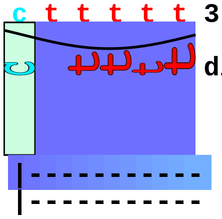


```
       .         *3235270  .         *3235260  .         *3235250  .
5' c a t c t t c g a c g t g c g c a g a t a c g c a t t a t t t c t g c g c c 3'
```

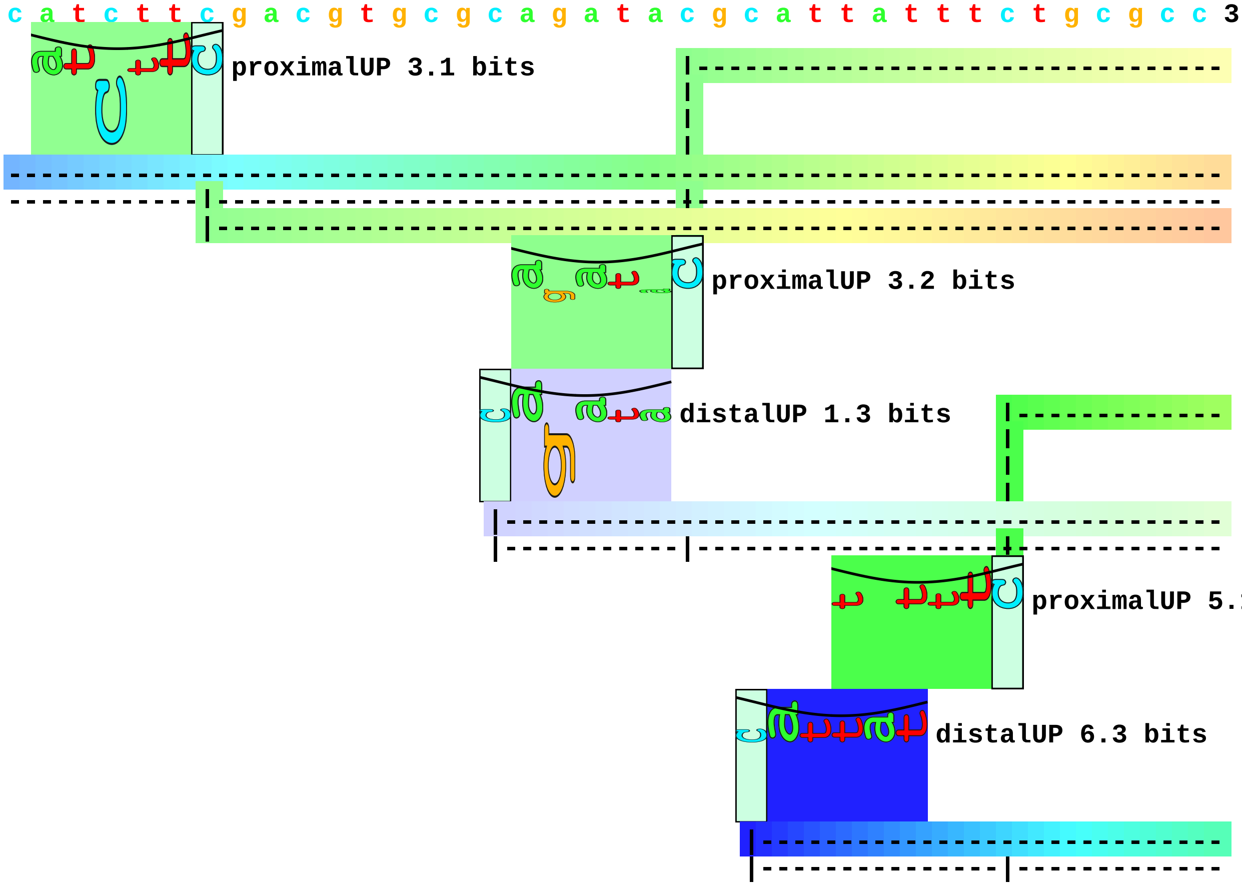


```
   *3235240  .         *3235230  .         *3235220  .         *3235210  .
5' g g a c a g c c a t g c t c a c t a a a c g g c c c a a t c a t g a c c g a 3'
... ----------------------------------| proximalUP-(34)-sigma38p10 3235224 Gap 2.5 bits
... ----------------| distalUP-(52)-sigma38p10 3235233 Gap 5.5 bits
```

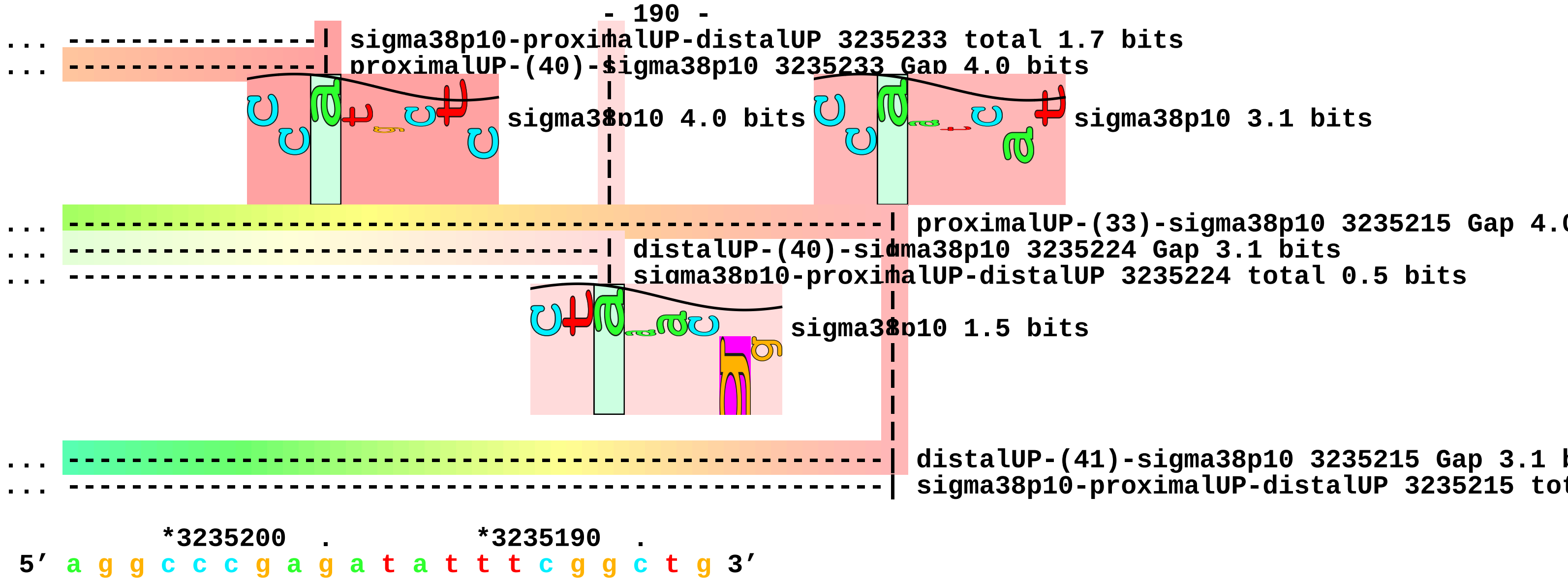
... sigma38p10-proximalUP-distalUP 3235233 total 1.7 bits
... proximalUP-(40)-sigma38p10 3235233 Gap 4.0 bits
sigma38p10 4.0 bits
sigma38p10 3.1 bits
... proximalUP-(33)-sigma38p10 3235215 Gap 4.
... distalUP-(40)-sigma38p10 3235224 Gap 3.1 bits
... sigma38p10-proximalUP-distalUP 3235224 total 0.5 bits
sigma38p10 1.5 bits
... distalUP-(41)-sigma38p10 3235215 Gap 3.1
... sigma38p10-proximalUP-distalUP 3235215 tot
*3235200 .
*3235190 .
5' a g g c c c g a g a t a t t t c g g c t g 3'

piece 46, NC_000913, rpoH, config: linear, direction: +, begin: 3598900, end: 3599149

*3598900 . *3598910 . *3598920 . *3598930 .
5' t c a g a c c g t g a t t t t a t c c a c a a g t t c a a t g c a a g c t t 3'

proximalUP 6.8 bits

T timestamp:_2018Sep20 11-35-24

... proximalUP-(32)

distalUP 6.5 bits

... distalUP-(40)-s
... distalUP-proxim
... proximalUP-(38)
... distalUP-(46)-s
... distalUP-proxim

*3598940 . *3598950 . *3598960 . *3598970 .
5' g t g a a t a a a t t a c g c a c a a a a t g t g a c a t a g a g a t g a a 3'

sigma38p10 0.7 bits

... proximalUP-(32)-sigma38p10 3598949 Gap 4.3 bits

sigma38p10 2.1 bits

... distalUP-(40)-sigma38p10 3598949 Gap 3.1 bits
... distalUP-proximalUP-sigma38p10 3598949 total 6.6 bits
... proximalUP-(38)-sigma38p10 3598955 Gap 3.2 bits
... distalUP-(46)-sigma38p10 3598955 Gap 4.1 bits
... distalUP-proximalUP-sigma38p10 3598955 total 8.0 bits

*3598980 . *3598990 . *3599000 . *3599010
5' a t a c c g g g a a g a g a c a a c g g g g t c t c t t t c c c t g c t a c 3'

. *3599020 . *3599030 . *3599040 . *3599050
5' g g a a c c c a t t g c a g g g a a a g a g t a t a a c a c g c t t t t a t 3'

proximalUP 3.9 bits

... proximalUP-(32)

distalUP 3.4 bits

... proximalUP

... distalUP-(40)-s
... distalUP-proxim

distalUP 3.4 bits

|------------- ... distalUP-(50)-s
|------------- ... distalUP-proxin

. *3599060 . *3599070 . *3599080 .
5′ t a t t c a g g c g t a a a g t g g c g t a a a t g t t g t a c c g t g g c 3′

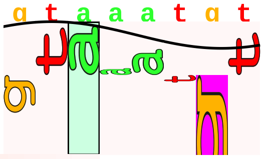
sigma38p10 0.4 bits

... ----------------------------------------| proximalUP-(32)-sigma38p10 3599073 Gap 4.3 bits

... 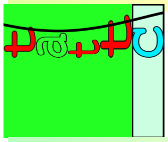
proximalUP 6.2 bits

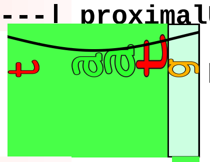
proximalUP 5.2 bits

... ----------------------------------------| distalUP-(40)-sigma38p10 3599073 Gap 3.1 bits
... --------|-------------------------------| distalUP-proximalUP-sigma38p10 3599073 total 0.2 bit
|----------------------------------------------------------- ... proximalUP-(39)
... ---------------------------------------------------------------------- ... distalUP-(50)-s
... --------|----------------------------------------|------------------------ ... distalUP-proxin
|------------------------- ... proximalUP-(34)

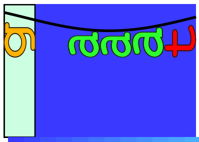
distalUP 5.6 bits

|------------------------------------- ... distalUP-(40)-s
|-----------|------------------------- ... distalUP-proxin

*3599090 . *3599100 . *3599110 . *3599120 .
5′ a a g c c a c g c t g c c a c c c a g c c a a t c a t c g a g c a t a c c a 3′

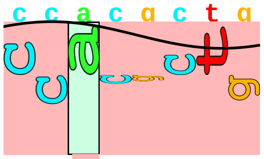
sigma38p10 3.1 bits

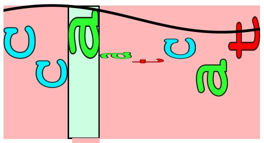
sigma38p10 3.1 bits

... ----------| proximalUP-(39)-sigma38p10 3599095 Gap 7.5 bits
... ----------| distalUP-(50)-sigma38p10 3599095 Gap 4.1 bits
... ----------| distalUP-proximalUP-sigma38p10 3599095 total 1.1 bits
... ----------------------------------------| proximalUP-(34)-sigma38p10 3599111 Gap 2.5 bits
... ----------------------------------------| distalUP-(40)-sigma38p10 3599111 Gap 3.1 bits
... ----------------------------------------| distalUP-proximalUP-sigma38p10 3599111 total 8.3 bit

*3599130 . *3599140 .
5′ g c a g c a a t a g c a g g c a t t c a t c 3′

piece 47, NC_000913, uspB, config: linear, direction: +, begin: 3637700, end: 3637999

```
   *3637700  .         *3637710  .         *3637720  .         *3637730  .
5' c a a a c g a c a c a t a a a g c c c a a a a t a a t g c g a c g g t g c t 3'
   T timestamp:_2018Sep20_11-35-24

     *3637740  .         *3637750  .         *3637760  .         *3637770  .
5' t a t c a t a c c t c c t c c c c g g c g a c c t g c c c g c g g a g t t c 3'

       *3637780  .         *3637790  .         *3637800  .         *3637810
5' c a c c c c g g g g c t a c c g c t c c c g a t a c g c t g c c a a t c a g 3'

   .         *3637820  .         *3637830  .         *3637840  .         *3637850
5' t t a a c a c c a g g t c c t g g a g a a a c c g c t t t t g t g g t g a c 3'
                                              proximalUP 3.8 bits
                                              |------------------------------ ... proximalUP-(34)
                                              distalUP 2.7 bits
                                            |------------------------------------ ... distalUP-(40)-s
                                            |-----------|------------------------ ... distalUP-proxim

         .         *3637860  .         *3637870  .         *3637880  .
5' c a a c a t a c g a g c g g c t c t a t a g a t a g t g t a g g a g a t c a 3'
                                              sigma38p10 4.8 bits
... ------------------------------------| proximalUP-(34)-sigma38p10 3637870 Gap 2.5 bits
                                              proximalUP 2.7 bits
... ------------------------------------| distalUP-(40)-sigma38p10 3637870 Gap 3.1 bits
... ------------------------------------| distalUP-proximalUP-sigma38p10 3637870 total 5.8 bits
                                                        |------------------------ ... proximalUP-(40)
                                              distalUP 2.1 bits
                                                  |------------------------------ ... distalUP-(44)-s
                                                  |-------|---------------------- ... distalUP-proxim

   *3637890  .         *3637900  .         *3637910  .         *3637920  .
5' g g t t g t t t t t t t t c c a g a a g g t t a a c c a c t a t c a a t a t 3'
                              proximalUP 6.2 bits                   sigma38p10 2.3 bits
                              |------------------------------------------- ... proximalUP-(35)
```

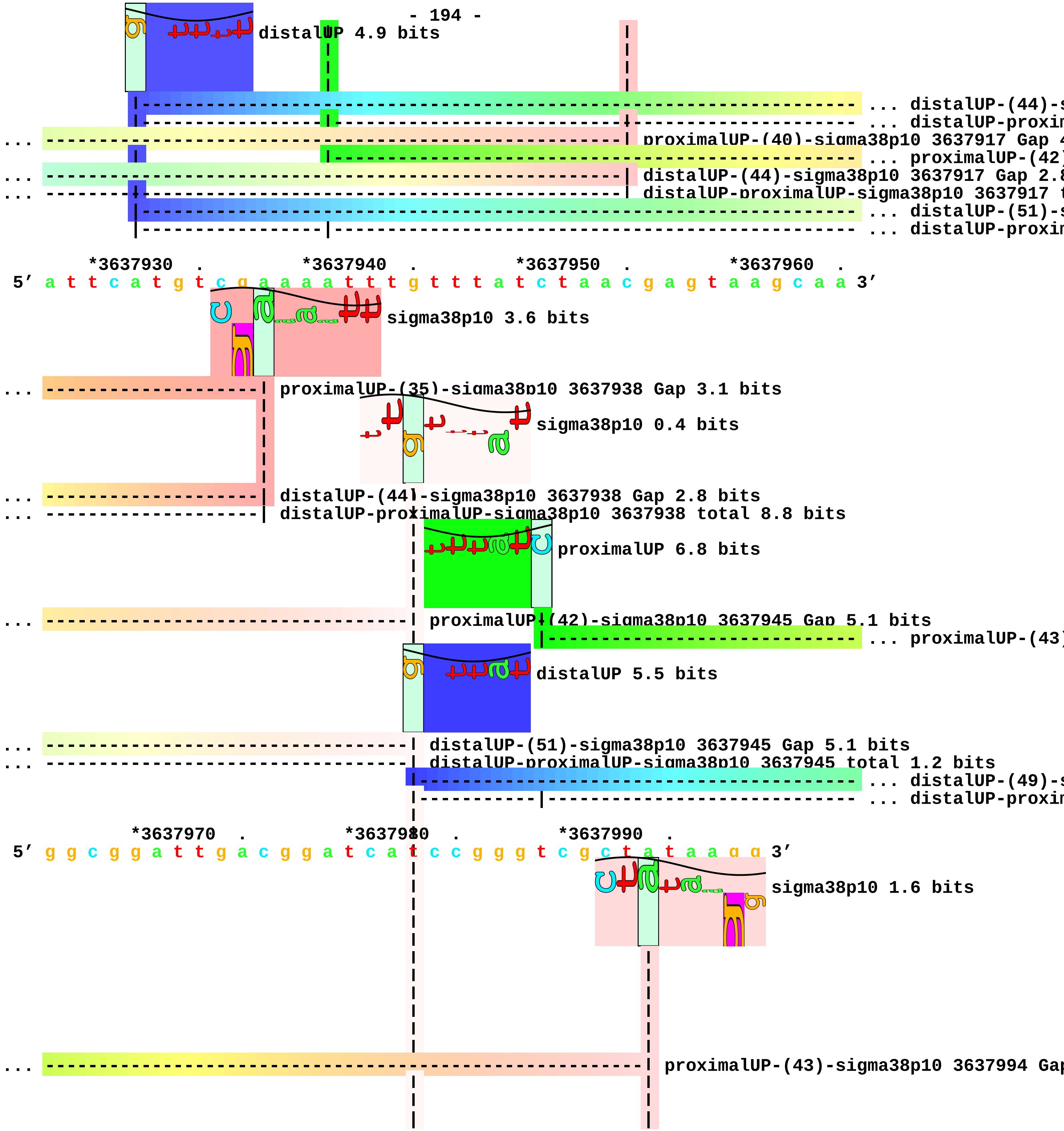
distalUP 4.9 bits
... distalUP-(44)-s
... distalUP-proxim
... proximalUP-(40)-sigma38p10 3637917 Gap 4
... proximalUP-(42)
... distalUP-(44)-sigma38p10 3637917 Gap 2.8
... distalUP-proximalUP-sigma38p10 3637917 t
... distalUP-(51)-s
... distalUP-proxim
*3637930 . *3637940 . *3637950 . *3637960 .
5' a t t c a t g t c g a a a a t t t g t t t a t c t a a c g a g t a a g c a a 3'
sigma38p10 3.6 bits
... proximalUP-(35)-sigma38p10 3637938 Gap 3.1 bits
sigma38p10 0.4 bits
... distalUP-(44)-sigma38p10 3637938 Gap 2.8 bits
... distalUP-proximalUP-sigma38p10 3637938 total 8.8 bits
proximalUP 6.8 bits
... proximalUP-(42)-sigma38p10 3637945 Gap 5.1 bits
... proximalUP-(43)
distalUP 5.5 bits
... distalUP-(51)-sigma38p10 3637945 Gap 5.1 bits
... distalUP-proximalUP-sigma38p10 3637945 total 1.2 bits
... distalUP-(49)-s
... distalUP-proxim
*3637970 . *3637980 . *3637990 .
5' g g c g g a t t g a c g g a t c a t c c g g g t c g c t a t a a g g 3'
sigma38p10 1.6 bits
... proximalUP-(43)-sigma38p10 3637994 Gap

```
... --------------------------------------------------------| distalUP-(49)-sigma38p10 3637994 Gap 5
... --------------------------------------------------------| distalUP-proximalUP-sigma38p10 3637994
                                    |
                                    |
                                    |
```

piece 48, NC_000913, proK, config: linear, direction: +, begin: 3706700, end: 3707049

```
   *3706700   .         *3706710   .         *3706720   .         *3706730   .
5' g c t g c g c c a a t c a c c g a a t g c g g g g c g c a t c t t a c t g c 3'
   T timestamp:_2018Sep20_11-35-24

       *3706740   .         *3706750   .         *3706760   .         *3706770   .
5' g c a g a t a c g c c c t c g t c a a t c c c t t a a t a g c a a a a a t g c 3'

           *3706780   .         *3706790   .         *3706800   .         *3706810
5' c t t t t g a t c g g c g a g a a a g t c a g c a g g c c g c t t a g t t a 3'

     .         *3706820   .         *3706830   .         *3706840   .         *3706850
5' g c c g c t g c c t c t t t t g c c t g c g g g a t g t g a c a c c a g t t 3'

         .         *3706860   .         *3706870   .         *3706880   .
5' g t t g t t t t c g t t a a t t c c a c c a t c c g g t g a a g t a t a g c 3'
                                        proximalUP 5.1 bits
                                         |------------------------------------- ... proximalUP-(33)
                                   distalUP 5.5 bits
                        |--------------------------------------------------------- ... distalUP-(40)-s
                        |-------------|------------------------------------------- ... distalUP-proxim

   *3706890   .         *3706900   .         *3706910   .         *3706920   .
5' c a a g a c a a c c c a t g a t g g t a t c g t a c a g c t c t a c g t g a 3'
                                        sigma38p10 3.7 bits
... ----------------------| proximalUP-(33)-sigma38p10 3706901 Gap 4.0 bits
                          |
... ----------------------| distalUP-(40)-sigma38p10 3706901 Gap 3.1 bits
... ----------------------| distalUP-proximalUP-sigma38p10 3706901 total 7.2 bits

       *3706930   .         *3706940   .         *3706950   .         *3706960   .
5' c g g c g t g g c a c t t t c a t g t c g g c t t c t t t t t c a g c t g 3'
                                                              proximalUP 6.2 bits
                                                              |----------- ... proximalUP-(37)
                                                          distalUP 4.1 bits
                                                 |-------------------------- ... distalUP-(44)-s
                                                 |-------------|----------- ... distalUP-proxim
                                                 |             |----------- ... proximalUP-(39)
                                                 |-------------------------- ... distalUP-(46)-s
                                                 |-------------|----------- ... distalUP-proxim
```

|-----------  ... proximalUP-(41)
|------------------------  ... distalUP-(48)-s
|-------------|-----------  ... distalUP-proxim

*3706970 . *3706980 . *3706990 . *3707000

5' c g c a a a c g c c t g c g c a t t g g c c g g a t t t t c c a g a t a t t 3'

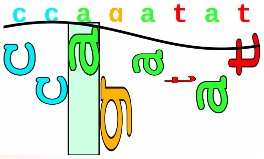

sigma38p10 0.1 bits

... --------------------------------------------------------------| proximalUP-(37)-sigma38p10 37069

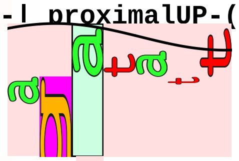

... sigma38p10

... --------------------------------------------------------------| distalUP-(44)-sigma38p10 370699

... --------------------------------------------------------------| distalUP-proximalUP-sigma38p10 3

... ------------------------------------------------------------------| proximalUP-(39)-sigma38p10 3

... ------------------------------------------------------------------| distalUP-(46)-sigma38p10 370

... ------------------------------------------------------------------| distalUP-proximalUP-sigma38p

... ----------------------------------------------------------------------| proximalUP-(41)-sigma38p

... ----------------------------------------------------------------------| distalUP-(48)-sigma38p10

... ----------------------------------------------------------------------| distalUP-proximalUP-sigm

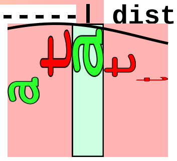

... sigma38p10

. *3707010 . *3707020 . *3707030 . *3707040

5' t a t c t g a c a t c c a g a c c a t c a t c g g t a c g c g g a a c t g c 3'

... 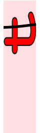 sigma38p10 1.4 bits

... 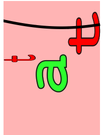 sigma38p10 3.2 bits

.

5' t c c g g c g g 3'

piece 49, NC_000913, yjdC, config: linear, direction: +, begin: 4361237, end: 4361482

```
        *4361240  .         *4361250  .         *4361260  .         *4361270
5' t c c a c a c g t t c a g c a a c c a t c t c c a g c g t g g t g t t g g c 3'
   T timestamp:_2018Sep20_11-35-24

   .         *4361280  .         *4361290  .         *4361300  .         *4361310
5' a a t c c c t t g t a a t t c t a a t a a t t t c a g g g c t t c t c c c a 3'
                                        proximalUP 5.1 bits
                                                                              ... proximalUP-(34)
                              distalUP 5.0 bits                               ... proximalUP-(36)
                                                                              ... distalUP-(40)-s
                                                                              ... distalUP-proxin
                                                 proximalUP 5.5 bits
                                                                              ... distalUP-(52)-s
                                                                              ... distalUP-proxin

       .         *4361320  .         *4361330  .         *4361340  .         *4361350
5' g t a c a t c t t c a c g t t g c a c g c t a t t t t c c t c c g t c t t t 3'
                                   sigma38p10 0.3 bits
... -------------------| proximalUP-(34)-sigma38p10 4361323 Gap 2.5 bits
... ----------------------------------------| proximalUP-(36)-sigma38p10 4361335 Gap 3.6 bits
... -------------------| distalUP-(40)-sigma38p10 4361323 Gap 3.1 bits
... -------------------| distalUP-proximalUP-sigma38p10 4361323 total 4.9 bits
                                                      sigma38p10 5.9 bits
... ----------------------------------------| distalUP-(52)-sigma38p10 4361335 Gap 5.5 bits
... ----------------------------------------| distalUP-proximalUP-sigma38p10 4361335 total 7.4 b

         .         *4361360  .         *4361370  .         *4361380  .
5' c c c a c t g c a a g t g t c g t t c a c g g t t g g c g a t c g c g c a a 3'

     *4361390  .         *4361400  .         *4361410  .         *4361420  .
5' a t g t g c g c t g a a g g t t t c a g c a t c c a t a a a g c c c g t g a 3'

        *4361430  .         *4361440  .         *4361450  .         *4361460
5' c g c g t g c t t g t g g a t g c t c c t g g c c t t g t c c g t c a a a a 3'

   .         *4361470  .         *4361480
5' a a g a g a a t t g t c g g t a g g 3'
```

piece 50, NC_000913, ytfJ, config: linear, direction: +, begin: 4436950, end: 4437399

```
   *4436950  .         *4436960  .         *4436970  .         *4436980  .
5' a a a c a t c c c t g a a c c c g g a a t c g c g t c g t c g g t g t t a a 3'
   T timestamp:_2018Sep20_11-35-24

     *4436990  .         *4437000  .         *4437010  .         *4437020  .
5' c a a t g g t g g t g g t c t g g t a a c g a t c g t g c g g t a a c t t c 3'

       *4437030  .         *4437040  .         *4437050  .         *4437060
5' g c t g a t t t a a t c g c t t c a a t c a g c g t c g c g t t t t t c t c 3'
                                                              proximalUP 6.2 bits
                     distalUP 2.5 bits                                ... proximalUP-(27)
                                                                      ... distalUP-(46)-s
                                                                      ... distalUP-proxim
                                                                      ... proximalUP
                                                   distalUP 4.9 bits
                                                                      ... distalUP-(47)-s
                                                                      ... distalUP-proxim
                                                                      ... proximalUP-(43)
                                                                      ... distalUP-(49)-s
                                                                      ... distalUP-proxim

    .         *4437070  .         *4437080  .         *4437090  .         *4437100
5' t t t t g c a g a g g t g c g a c c a g c a a t a t g t t g c a g t a c t c 3'
                                                 sigma38p10 3.8 bits
... ----------------------------------------| proximalUP-(27)-sigma38p10 4437088 Gap 3.8 bit
... ----------------------------------------| distalUP-(46)-sigma38p10 4437088 Gap 4.1 bits
... ----------------------------------------| distalUP-proximalUP-sigma38p10 4437088 total 4
...       proximalUP 3.2 bits              distalUP 6.4 bits         ... sigma38p10
       |-------------------------------------------------------------- ... proximalUP-(34)
... ------------------------------------------------------------------ ... distalUP-(47)-s
... -------|-----------------------|---------------------------------- ... distalUP-proxim
... ------------------------------------------------------------------ ... proximalUP-(43)
... ------------------------------------------------------------------ ... distalUP-(49)-s
... -------------------------------|---------------------------------- ... distalUP-proxim
```

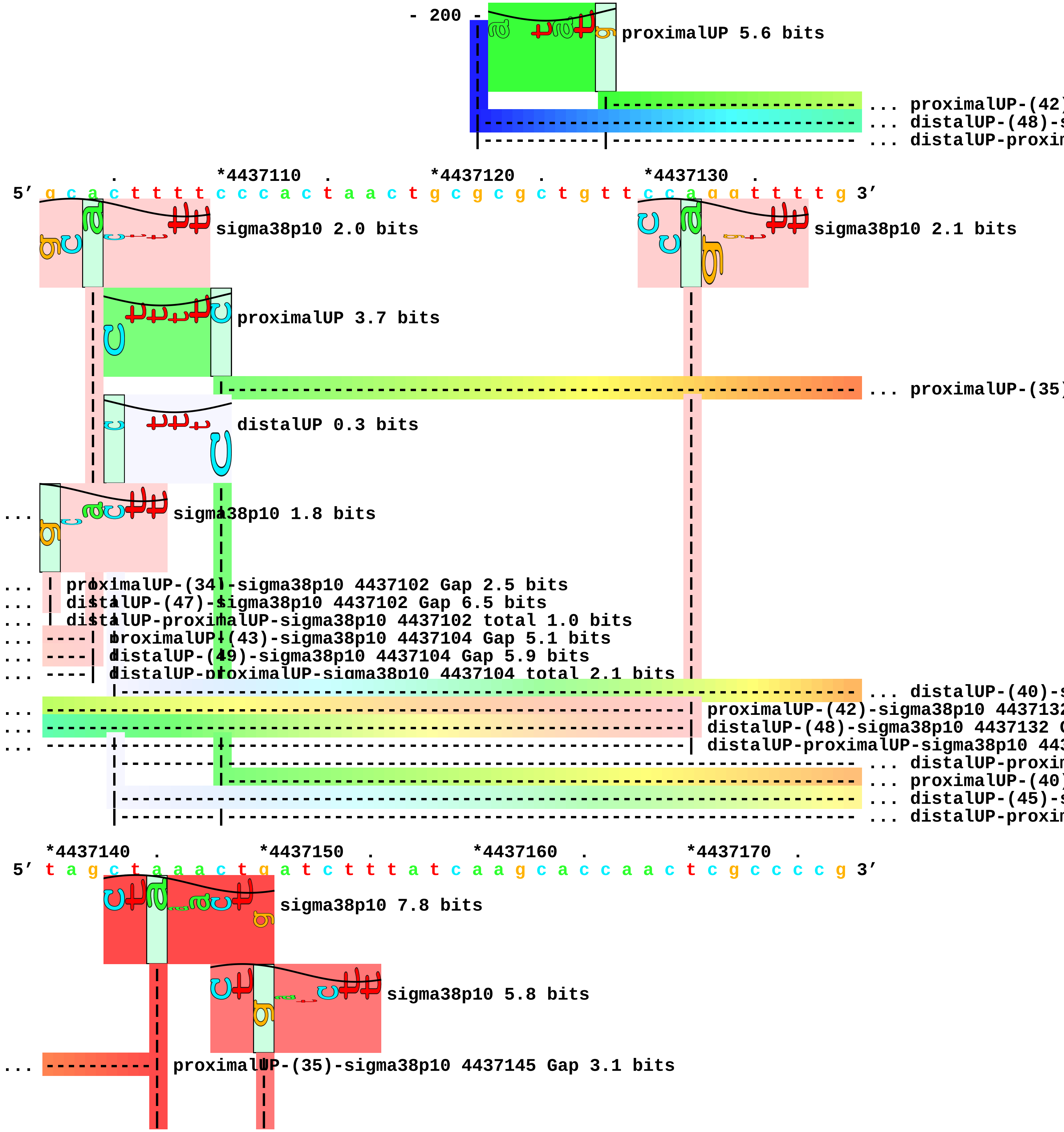

- 200 -
proximalUP 5.6 bits
... proximalUP-(42)
... distalUP-(48)-s
... distalUP-proxim
. *4437110 . *4437120 . *4437130 .
5′ g c a c t t t t c c c a c t a a c t g c g c g c t g t t c c a g g t t t t g 3′
sigma38p10 2.0 bits
sigma38p10 2.1 bits
proximalUP 3.7 bits
... proximalUP-(35)
distalUP 0.3 bits
... sigma38p10 1.8 bits
... proximalUP-(34)-sigma38p10 4437102 Gap 2.5 bits
... distalUP-(47)-sigma38p10 4437102 Gap 6.5 bits
... distalUP-proximalUP-sigma38p10 4437102 total 1.0 bits
... proximalUP-(43)-sigma38p10 4437104 Gap 5.1 bits
... distalUP-(49)-sigma38p10 4437104 Gap 5.9 bits
... distalUP-proximalUP-sigma38p10 4437104 total 2.1 bits
... distalUP-(40)-s
... proximalUP-(42)-sigma38p10 4437132
... distalUP-(48)-sigma38p10 4437132 G
... distalUP-proximalUP-sigma38p10 443
... distalUP-proxim
... proximalUP-(40)
... distalUP-(45)-s
... distalUP-proxim
*4437140 . *4437150 . *4437160 . *4437170 .
5′ t a g c t a a a c t g a t c t t t a t c a a g c a c c a a c t c g c c c c g 3′
sigma38p10 7.8 bits
sigma38p10 5.8 bits
... proximalUP-(35)-sigma38p10 4437145 Gap 3.1 bits

... ----------| distalUP-(40)-sigma38p10 4437145 Gap 3.1 bits

... ----------| distalUP-proximalUP-sigma38p10 4437145 total 5.6 bits
... --------------------| proximalUP-(40)-sigma38p10 4437150 Gap 4.0 bits
... --------------------| distalUP-(45)-sigma38p10 4437150 Gap 5.5 bits
... --------------------| distalUP-proximalUP-sigma38p10 4437150 total 0.3 bits

*4437180 . *4437190 . *4437200 . *4437210 .
5' a t c g g t a a t g c c a a t c g g c g g c a c t c g c t g a c c g g t t t 3'

*4437220 . *4437230 . *4437240 . *4437250
5' c g a a c t g a t g t g c g g a a g c c a t c a t c g g c a a c a g c a g g 3'

. *4437260 . *4437270 . *4437280 . *4437290
5' c a g g t g a g t g c c a g a a t c t t g c g t a g g g t c a t g g t g t t 3'

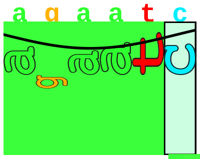

proximalUP 5.5 bits

|---------------------------------------- ... proximalUP-(34)

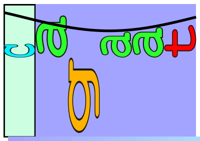

distalUP 2.6 bits

|------------------------------------------------------------ ... distalUP-(40)-s
|-----------|----------------------------------------------- ... distalUP-proxim

. *4437300 . *4437310 . *4437320 .
5' t c c t t c t t a t g a t a t g c a g g t g a t c c g a c c a c t t g g g t 3'

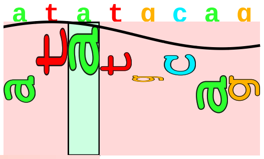

sigma38p10 1.7 bits

... -------------------------| proximalUP-(34)-sigma38p10 4437305 Gap 2.5 bits
... -------------------------| distalUP-(40)-sigma38p10 4437305 Gap 3.1 bits
... -------------------------| distalUP-proximalUP-sigma38p10 4437305 total 4.2 bits

*4437330 . *4437340 . *4437350 . *4437360 .
5' c t g a g t t t t a a t c a t a t g t g c t a t t t a t c g a a t t t c c c 3'

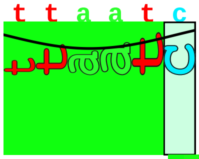

proximalUP 6.8 bits

|------------------------------------------------------ ... proximalUP-(32)

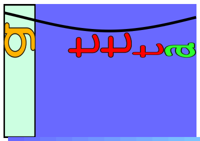

distalUP 4.2 bits

|------------------------------------------------------------------ ... distalUP-(40)-s

|---------------|--------------------------------------------------- ... distalUP-proxin

*4437370 . *4437380 . *4437390 .

5′ g c a a g t g t g a t g c c a g t t t g c g g t c a a g c g c a 3′

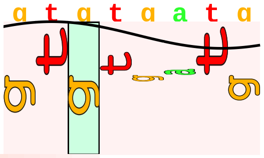

sigma38p10 0.6 bits

... ------------| proximalUP-(32)-sigma38p10 4437374 Gap 4.3 bits

... ------------| distalUP-(40)-sigma38p10 4437374 Gap 3.1 bits

... ------------| distalUP-proximalUP-sigma38p10 4437374 total 4.2 bits
