## Supplement_for_sigma38_paper-directory for "*Escherichia coli σ*^38^ promoters use two UP elements instead of a −35 element: resolution of a paradox and discovery that *σ*^38^ transcribes ribosomal promoters": Peano.Landini2015-Table1-pos-map.pdf

piece 1, NC_000913, hepA, config: linear, direction: -, begin: 63588, end: 63350

```
           .         *63580    .         *63570    .         *63560    .
5' a t g g a c c a c c a a c g g c c c g g a g c c g c t g g a c t a c c a a c 3'
   T timestamp:_2018Sep20_11-34-53

   *63550    .         *63540    .         *63530    .         *63520    .
5' g t t c a c c a c t g g a t t a c g a a c a c t a t c t g a c c g c c a g 3'

     *63510    .         *63500    .         *63490    .         *63480    .
5' c t a c a a c c c g t g g c g g a g g g a a t a c t c c c t t t t a t t g a 3'
                                                                   proximalUP 5.7 bits
                                                                   |--- ... proximalUP-(41)
                                                              distalUP 3.4 bits
                                                              |------------------- ... distalUP-(49)-s
                                                              |---------------|--- ... sigma38p10-prox

       *63470    .         *63460    .         *63450    .         *63440
5' g g a t a a t t t t g c t a c a c t t a t g a c c g g g c a a c t t g g g c 3'
                                                                          ... sigma38p10
... ----------------------------------------------------------------------- ... proximalUP-(41)
                                                                 proximalUP 1.5 bits
... ----------------------------------------------------------------------- ... distalUP-(49)-s
... -------------------------------------------------------------------|------- ... sigma38p10-prox
                                                                       |------- ... proximalUP-(35)
                                                              distalUP 1.0 bits
                                                              |------------------- ... distalUP-(41)-s
                                                              |-----------|------- ... sigma38p10-prox
                                                              |------------------- ... distalUP-(44)-s
                                                              |------------------- ... sigma38p10-prox

     .         *63430    .         *63420    .         *63410    .         *63400
5' t a t t t t g a g c a a a a a a a a g a g t t c g c c a g a t a c c a t t t 3'
... sigma38p10 6.2 bits                                         sigma38p10 4.0 bits
... --| proximalUP-(41)-sigma38p10 63435 Gap 5.5 bits           ... sigma38p10
```

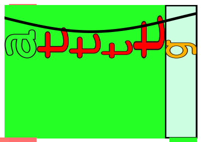

proximalUP 6.1 bits

```
... --| distalUP-(49)-sigma38p10 63435 Gap 5.9 bits
... --| sigma38p10-proximalUP-distalUP 63435 total 3.9 bits
... ---------------------------------------------------------------| proximalUP-(35)-sigma38p10 63405
            |---------------------------------------------------------| proximalUP-(28)-sigma38p10
... ---------------------------------------------------------------| distalUP-(41)-sigma38p10 63405 (
... ---------------------------------------------------------------| sigma38p10-proximalUP-distalUP 6
... ---------------------------------------------------------------------| distalUP-(44)-sigma38p10 6
... ------------|---------------------------------------------------------| sigma38p10-proximalUP-dist

             .         *63390    .         *63380    .         *63370    .
 5' t g a t g c g t g a c g a a t g c t t t g c c a t c c a g t a c c a t a g c 3'
```

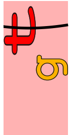

... sigma38p10 3.3 bits

```
    *63360    .         *63350
 5' g c c c t t t c c a t 3'
```

piece 2, NC_000913, lpxC, config: linear, direction: +, begin: 106386, end: 106666

```
            *106390    .          *106400    .          *106410    .          *106420
5' a t g c g c c g c a a a c t g c g a a a g a g c c g g a t t a t c t g g a t 3'

   T timestamp:_2018Sep20_11-34-53                       proximalUP 7.2 bits

                                    distalUP 1.6 bits              ... proximalUP-(28)

                                                                   ... distalUP-(44)-s
                                                                   ... distalUP-proxin
                                                                   ... proximalUP-(37)
                                                 distalUP 7.1 bits

                                                                   ... distalUP-(43)-s
                                                                   ... distalUP-proxin

   .          *106430    .          *106440    .          *106450    .          *106460
5' a t c c c a g c a t t c c t g c g t a a g c a a g c t g a t t a a g a a t t 3'
                                                   sigma38p10 1.0 bits

... ------------------------------------------| proximalUP-(28)-sigma38p10 106446 Gap 5.5 bits
... ------------------------------------------| distalUP-(44)-sigma38p10 106446 Gap 2.8 bits
... ------------------------------------------| distalUP-proximalUP-sigma38p10 106446 total 1.5 bi
... ---------------------------------------------------------| proximalUP-(37)-sigma38p10 10645
                                                                   sigma38p10 3.6 bits

... ---------------------------------------------------------| distalUP-(43)-sigma38p10 106455
... ---------------------------------------------------------| distalUP-proximalUP-sigma38p10 1
                                                        proximalUP 4.9 bits

                                                                   ... proximalUP-(38)
                                                        distalUP 6.4 bits

                                                                   ... distalUP-(44)-s
                                                                   ... distalUP-proxin

   .          *106470    .          *106480    .          *106490    .
5' g a c t g g a a t t t g g g t t t c g a g g c t c t t t g t g c t a a a c t 3'
                                                                   ... sigma38p10
```

```
... ---------------------------------------------------------------------| proximalUP-(38)-sigma38p10 1
... ---------------------------------------------------------------------| distalUP-(44)-sigma38p10 106
... ---------------------------------------------------------------------| distalUP-proximalUP-sigma38p

     *106500    .         *106510    .         *106520    .         *106530    .
 5' g g c c c g c c g a a t g t a t a g t a c a c t t c g g t t g g a t a g g t 3'
... sigma38p10 7.8 bits                                                    ... distalUP

                                                                          |--- ... distalUP-(42)-s
                                                                          |--- ... distalUP-proxin

         *106540    .         *106550    .         *106560    .         *106570    .
 5' a a t t t g g c g a g a t a a t a c g a t g a t c a a a c a a a g g a c a c 3'
... distalUP 5.0 bits

... ---------------------------------------------------------------------------- ... distalUP-(42)-s
... ----------|----------------------------------------------------------------- ... distalUP-proxin
      proximalUP 5.0 bits

            |---------------------------------------------------------------- ... proximalUP-(35)

             *106580    .         *106590    .         *106600    .         *106610
 5' t t a a a c g t a t c g t t c a g g c g a c g g g t g t c g g t t t a c a t 3'
      sigma38p10 2.4 bits

... ----| distalUP-(42)-sigma38p10 106578 Gap 3.3 bits
... ----| distalUP-proximalUP-sigma38p10 106578 total 6.0 bits
        |
... ----| proximalUP-(35)-sigma38p10 106578 Gap 3.1 bits

     .         *106620    .         *106630    .         *106640    .         *106650
 5' a c c g g c a a g a a a g t c a c c c t g a c g t t a c g c c c t g c g c c 3'

         .         *106660    .
 5' g g c c a a c a c c g g g g t 3'
```

piece 3, NC_000913, thrW, config: linear, direction: +, begin: 261990, end: 262252

*261990 . *262000 . *262010 . *262020 .
5′ g g t g a t g c a a a a g t a g c c a t t t g a t t c a c a a g g c c a t t 3′
T timestamp:_2018Sep20_11-34-53

*262030 . *262040 . *262050 . *262060 .
5′ g a c g c a t c g c c c g g t t a g t t t t a a c c t t g t c c a c c g t g 3′
proximalUP 5.0 bits
... proximalUP-(32)
distalUP 0.5 bits
... distalUP-(43)-s
... distalUP-proxim

*262070 . *262080 . *262090 . *262100
5′ a t t c a c g t t c g t g a a c a t g t c c t t t c a g g g c c g a t a t a 3′
sigma38p10 2.4 bits
... proximalUP-(32)-sigma38p10 262084 Gap 4.3 bits
... distalUP-(43)-sigma38p10 262084 Gap 3.6 bits
... distalUP-proximalUP-sigma38p10 262084 total 0.1 bits

. *262110 . *262120 . *262130 . *262140
5′ g c t c a g t t g g t a g a g c a g c g c a t t c g t a a t g c g a a g g t 3′

. *262150 . *262160 . *262170 .
5′ c g t a g g t t c g a c t c c t a t t a t c g g c a c c a t t a a a a t c a 3′
proximalUP 7.2 bits
... proximalUP-(37)
... proximalUP-(36)
distalUP 3.5 bits
proximalUP 6.1 bits
... distalUP-(43)-s
... distalUP-proxim
... proximalUP-(39)
... distalUP-(46)-s
... distalUP-proxim
distalUP 5.6 bits
... distalUP-(46)-s

|----------------|--- ... distalUP-proxin

*262180 . *262190 . *262200 . *262210 .
5' a a t t g t t a c g t a a g a t c t t a t c a t t c t c c c a c c a a a a a a 3'
... ----------------------------------------------------------------------| proximalUP-(37)-sigma38p
... --------------------------------------| proximalUP-(36)-sigma38p10 262199 Gap 3.6 bits

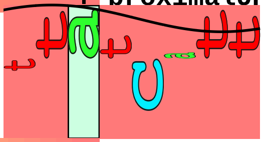

sigma38p10 5.7 bits

... --------------------------------------| distalUP-(43)-sigma38p10 262199 Gap 3.6 bits
... --------------------------------------| distalUP-proximalUP-sigma38p10 262199 total 9.3 bits
... --------------------------------------------| proximalUP-(39)-sigma38p10 262202 Gap 7.5 bits
... --------------------------------------------| distalUP-(46)-sigma38p10 262202 Gap 4.1 bits
... --------------------------------------------| distalUP-proximalUP-sigma38p10 262202 total 1.5 bi

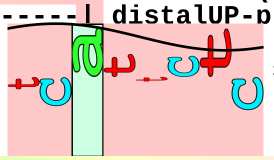

sigma38p10 2.4 bits

... ----------------------------------------------------------------------| distalUP-(46)-sigma38p10
... ----------------------------------------------------------------------| distalUP-proximalUP-sigm

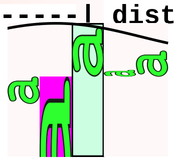

... sigma38p10

*262220 . *262230 . *262240 . *262250
5' a t t a t c t t a a t g t a a c a g c t g g t g t a a g t a a a t t c 3'

...

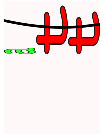

sigma38p10 0.3 bits

piece 4, NC_000913, insEF-2, config: linear, direction: -, begin: 392399, end: 392200

```
              .         *392390   .         *392380   .         *392370   .
5' c g g t c g t t c a c c a g a t c a g c g t t c a g a a c g t a a t c g c t 3'
   T timestamp:_2018Sep20_11-34-53

     *392360   .         *392350   .         *392340   .         *392330   .
5' g c t a t g g a t a t c g a a c a c a c c a g a t a c t a c g t t a c c g t 3'

         *392320   .         *392310   .         *392300   .         *392290
5' t a t g g t c a g c g a t g t t g c c g c t g g t c a g t t c c a g t t t g 3'

   .         *392280   .         *392270   .         *392260   .         *392250
5' t c a g t a c g g a a a g c t t c a g t a t c g a a c t g a t c a a c g t t 3'

       .         *392240   .         *392230   .         *392220   .         *392210
5' c a c a t c c a a t a c a c c g c c g t t a g t t a c g g t g a t a g t a t 3'

           .         *392200
5' t g g c g t a c a g 3'
```

piece 5, NC_000913, yaiA, config: linear, direction: +, begin: 406050, end: 406249

```
   *406050    .         *406060    .         *406070    .         *406080    .
5' c g c t a t a t c g c g a a g t t g c g c a t a t t a t c a t c g a c g c a 3'

 T timestamp:_2018Sep20_11-34-53                                       proximalUP 7.2 bits
                                                          |------------------- ... proximalUP-(37)
                                                     distalUP 5.7 bits
                                         |------------------------------------ ... distalUP-(45)-s
                                         |---------------|-------------------- ... distalUP-proxin

     *406090    .         *406100    .         *406110    .         *406120    .
5' a c a a a c g a a c c c a g c c a g g t g a t t t c t g a a a t t c g c a g 3'
                                                                     sigma38p10 6.4 bits
... ------------------------------------------------------| proximalUP-(37)-sigma38p10 406115 Gap 4.
                                                                     proximalUP 5.5 bits
... ------------------------------------------------------| distalUP-(45)-sigma38p10 406115 Gap 5.5
... ------------------------------------------------------| distalUP-proximalUP-sigma38p10 406115 to
                                                              |--------- ... proximalUP-(31)
                                                     distalUP 2.7 bits
                                         |------------------------------------ ... distalUP-(44)-s
                                         |-------------------------|---------- ... distalUP-proxin

         *406130    .         *406140    .         *406150    .         *406160
5' c g c c c t g g c a c a g a c g a t c a a t t g t t g a t t t t c g a g c g 3'
                                                                   sigma38p10 3.0 bits
... ----------------------------------------------------| proximalUP-(31)-sigma38p10 406152 Gap 4.7
... ----------------------------------------------------| distalUP-(44)-sigma38p10 406152 Gap 2.8 bi
... ----------------------------------------------------| distalUP-proximalUP-sigma38p10 406152 tota

   .         *406170    .         *406180    .         *406190    .         *406200
5' c c t a t a c t t a a c g t t c a t c c c g t g a a a t a a g g a a g a a c 3'
```

```
proximalUP 3.8 bits
|---------------- ... proximalUP-(43)
distalUP 5.9 bits
|------------------------------ ... distalUP-(50)-s
|-------------|--------------- ... distalUP-proxim

           .        *406210    .         *406220    .         *406230    .
5' g a t g c c a a c g a a a c c g c c t t a t c c t c g t g a a g c a t a t a 3'
                                                                   ... sigma38p10
... ----------------------------------------------------------------------| proximalUP-(43)-sigma38p
                                                                          |
... ----------------------------------------------------------------------| distalUP-(50)-sigma38p10
... ----------------------------------------------------------------------| distalUP-proximalUP-sigm

   *406240    .
5' t a g t g a c g a t 3'
... sigma38p10 1.8 bits
```

piece 6, NC_000913, yajO, config: linear, direction: -, begin: 437519, end: 437279

```
               .         *437510    .         *437500    .         *437490    .
5' a t g c t t a a g a a a t t a t t c a t a g a c t c t a a a t a a t t c g a 3'
   T timestamp:_2018Sep20_11-34-53

     *437480    .         *437470    .         *437460    .         *437450    .
5' g t t g c a g g a a g g c g g c a a a c g a g t g a a g c c c c a g g a g c 3'

       *437440    .         *437430    .         *437420    .         *437410
5' t t a c a t a a g t a a g t g a c t g g g g t g a a c g a a t g c a g c c g 3'

   .         *437400    .         *437390    .         *437380    .         *437370
5' c a g c a c a t g c a a c t t g a a g t a t g a c g a g t a t a g c a g g a 3'

     .         *437360    .         *437350    .         *437340    .         *437330
5' g t g g c a g c a t g c a a t a c a a c c c c t t a g g a a a a a c c g a c 3'
```

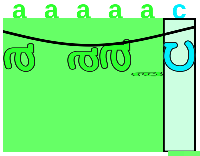

proximalUP 4.3 bits

|--------- ... proximalUP-(37)

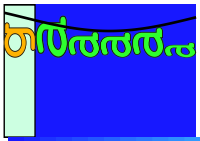

distalUP 6.5 bits

|---------------------- ... distalUP-(43)-s

|-----------|--------- ... sigma38p10-prox

```
               .         *437320    .         *437310    .         *437300    .
5' c t t c g c g t t t c c c g a c t t t g c c t c g g c t g t a t g a c c t t 3'
```

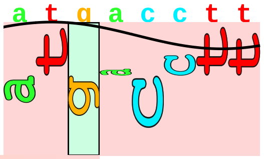

sigma38p10 1.8 bits

... ----------------------------------------------------------------| proximalUP-(37)-sigma38p10 437

... ----------------------------------------------------------------| distalUP-(43)-sigma38p10 43729

... ----------------------------------------------------------------| sigma38p10-proximalUP-distalUP

```
     *437290    .         *437280
5' t g g c g a g c c a g a t 3'
```

piece 7, NC_000913, tomB, config: linear, direction: -, begin: 480165, end: 479870

. *480160 . *480150 . *480140 . *480130
5′ t g t t a t g a a t a g t t c g a g c a a a c t g c t t t t a c c t g c t g 3′
T timestamp:_2018Sep20_11-34-53

. *480120 . *480110 . *480100 . *480090
5′ c g g g t t a g t g c t a g t a t g a a a a a g t g a g t c c t g t c c c g 3′

proximalUP 5.0 bits
... proximalUP-(35)
distalUP 0.5 bits
proximalUP 3.8 bits
... distalUP-(49)-s
... sigma38p10-prox
... proximalUP-(37)
... distalUP-(51)-s
... sigma38p10-prox
... proximalUP-(41)
distalUP 6.5 bits
... distalUP-(47)-s
... sigma38p10-prox

. *480080 . *480070 . *480060 .
5′ c t t c c t t c c t a a t t g t a a t t t t t c g t a a t a a t g c g a t g 3′

sigma38p10 5.3 bits
... proximalUP-(27)
... proximalUP-(35)-sigma38p10 480075 Gap 3.1 bits
sigma38p10 5.4 bits
... proximalUP-(33)
... distalUP-(49)-sigma38p10 480075 Gap 5.9 bits
... sigma38p10-proximalUP-distalUP 480075 total 1.9 bits
... proximalUP-(37)-sigma38p10 480073 Gap 4.5 bits
... distalUP-(51)-sigma38p10 480073 Gap 5.1 bits
... sigma38p10-proximalUP-distalUP 480073 total 1.4 bits
... proximalUP-(41)-sigma38p10 480063 Gap 5.5
sigma38p10 2.7 bits
... distalUP-(47)-sigma38p10 480063 Gap 6.5 bi
... sigma38p10-proximalUP-distalUP 480063 tota
proximalUP 6.7 bits

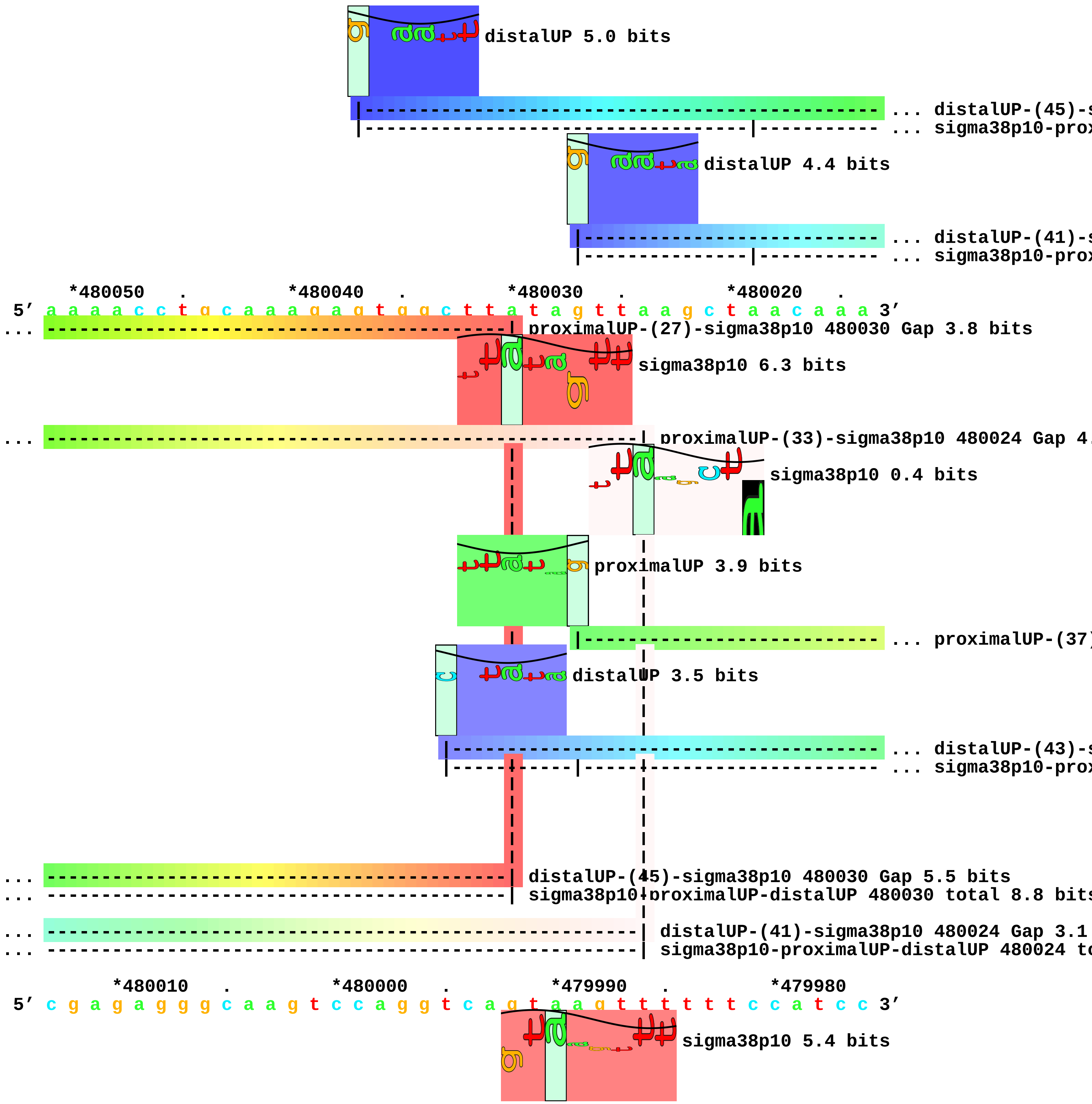
distalUP 5.0 bits
... distalUP-(45)-s
... sigma38p10-prox
distalUP 4.4 bits
... distalUP-(41)-s
... sigma38p10-prox
*480050 . *480040 . *480030 . *480020 .
5′ a a a a c c t g c a a a g a g t g g c t t a t a g t t a a g c t a a c a a a 3′
... proximalUP-(27)-sigma38p10 480030 Gap 3.8 bits
sigma38p10 6.3 bits
... proximalUP-(33)-sigma38p10 480024 Gap 4.
sigma38p10 0.4 bits
proximalUP 3.9 bits
... proximalUP-(37)
distalUP 3.5 bits
... distalUP-(43)-s
... sigma38p10-prox
... distalUP-(45)-sigma38p10 480030 Gap 5.5 bits
... sigma38p10-proximalUP-distalUP 480030 total 8.8 bits
... distalUP-(41)-sigma38p10 480024 Gap 3.1
... sigma38p10-proximalUP-distalUP 480024 to
*480010 . *480000 . *479990 . *479980
5′ c g a g a g g g c a a g t c c a g g t c a g t a a g t t t t t t c c a t c c 3′
sigma38p10 5.4 bits

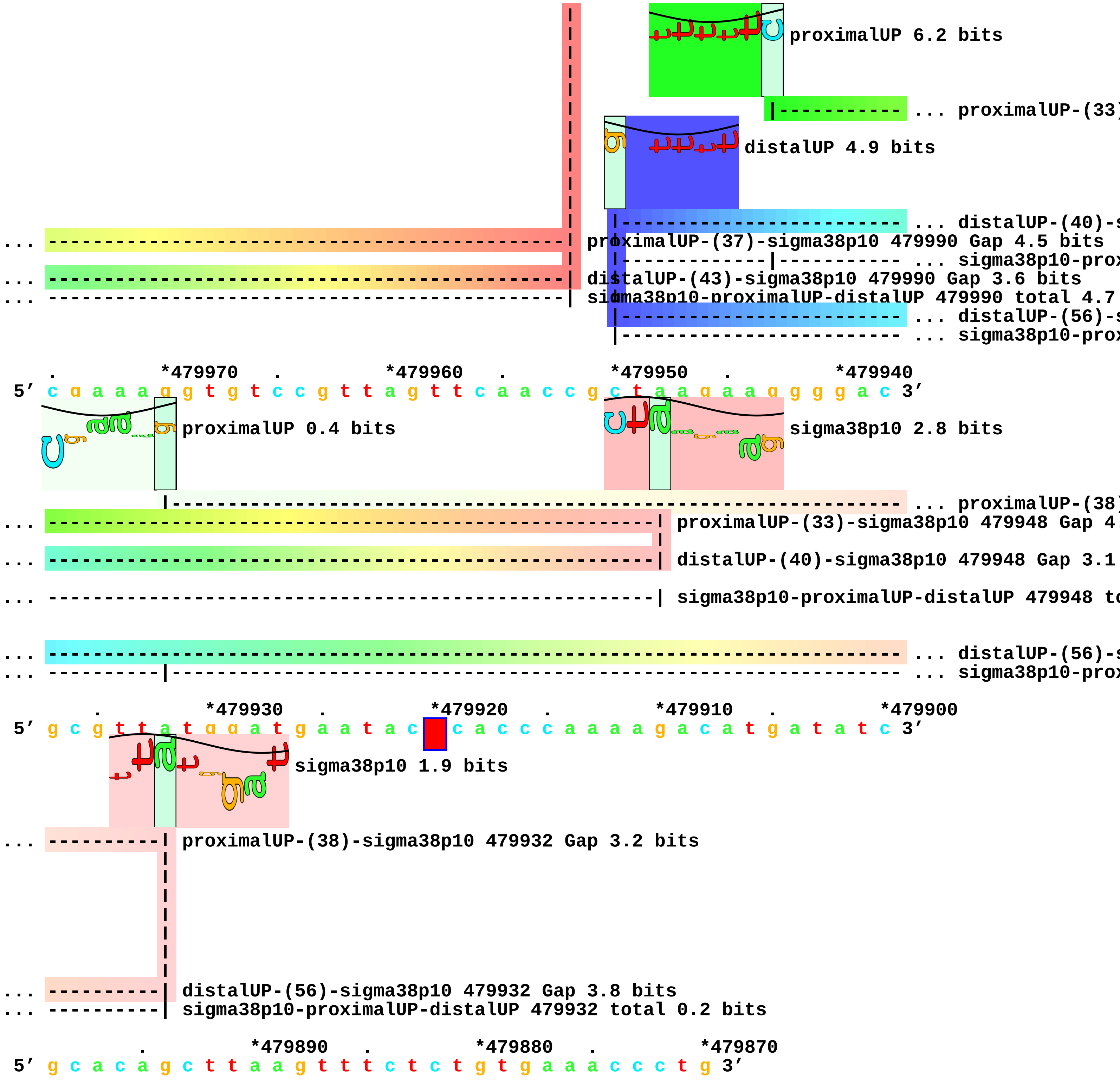
proximalUP 6.2 bits
... proximalUP-(33)
distalUP 4.9 bits
... distalUP-(40)-s
... proximalUP-(37)-sigma38p10 479990 Gap 4.5 bits
... sigma38p10-prox
... distalUP-(43)-sigma38p10 479990 Gap 3.6 bits
... sigma38p10-proximalUP-distalUP 479990 total 4.7
... distalUP-(56)-s
... sigma38p10-prox
. *479970 . *479960 . *479950 . *479940
5′ c g a a a g g t g t c c g t t a g t t c a a c c g c t a a g a a g g g g a c 3′
proximalUP 0.4 bits
sigma38p10 2.8 bits
... proximalUP-(38)
... proximalUP-(33)-sigma38p10 479948 Gap 4.
... distalUP-(40)-sigma38p10 479948 Gap 3.1
... sigma38p10-proximalUP-distalUP 479948 to
... distalUP-(56)-s
... sigma38p10-prox
. *479930 . *479920 . *479910 . *479900
5′ g c g t t a t g g a t g a a t a c c a c c c a a a a g a c a t g a t a t c 3′
sigma38p10 1.9 bits
... proximalUP-(38)-sigma38p10 479932 Gap 3.2 bits
... distalUP-(56)-sigma38p10 479932 Gap 3.8 bits
... sigma38p10-proximalUP-distalUP 479932 total 0.2 bits
. *479890 . *479880 . *479870
5′ g c a c a g c t t a a g t t t c t c t g t g a a a c c c t g 3′

piece 8, NC_000913, insH-2, config: linear, direction: -, begin: 575149, end: 574800

```
           .         *575140   .         *575130   .         *575120   .
5' t a a a g g t a a g g a t c t t g g a g t a t g g g g c g a t c a g g a c t 3'
   T timestamp:_2018Sep20_11-34-53

   *575110   .         *575100   .         *575090   .         *575080   .
5' t a g t c a a a t a t g a t g a t g t a g g t g c a a c c t a t t a c t t c 3'
```

proximalUP 5.6 bits

proximalUP 3.3 bits

... proximalUP-(40)[illegible]

distalUP 5.1 bits

... proximalUP-(34)[illegible]

... distalUP-(47)-[illegible]

... sigma38p10-pro[illegible]

... proximalUP-(42)[illegible]

... distalUP-(49)-[illegible]

... sigma38p10-pro[illegible]

distalUP 3.5 bits

... distalUP-(40)-[illegible]

... sigma38p10-pro[illegible]

```
     *575070   .         *575060   .         *575050   .         *575040
5' a a c a a a a a t a t g t c t a c t t t c g t t g a t t a c a a a a t c a a 3'
```

sigma38p10 1.0 bits

... proximalUP-(40)-sigma38p10 575060 Gap 4.0 bits

... proximalUP-(34)-sigma38p10 575043 [illegible]

... distalUP-(47)-sigma38p10 575060 Gap 6.5 bits

... sigma38p10-proximalUP-distalUP 575060 total 1.2 bits

... proximalUP-(42)-sigma38p10 575058 Gap 5.1 bits

... distalUP-(49)-sigma38p10 575058 Gap 5.9 bits

... sigma38p10-proximalUP-distalUP 575058 total 4.4 bits

sigma38p10 4.7 bits

... distalUP-(40)-sigma38p10 575043 Ga[illegible]

... sigma38p10-proximalUP-distalUP 575[illegible]

proximalUP 5.6 bits

sigma38p10 1.8 bits

... proximalUP-(37)[illegible]

distalUP 5.7 bits

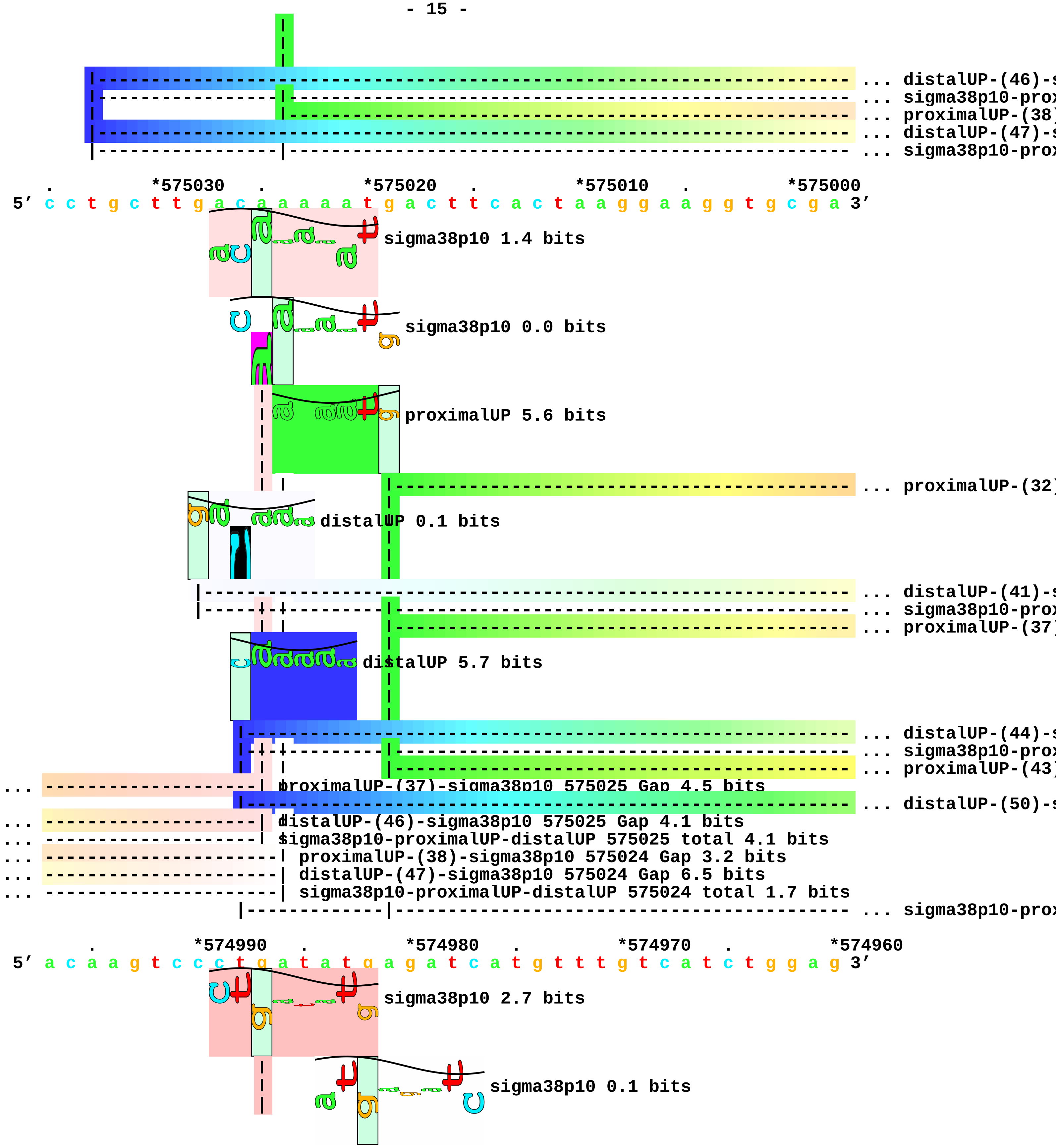

... distalUP-(46)-s
... sigma38p10-prox
... proximalUP-(38)
... distalUP-(47)-s
... sigma38p10-prox
*575030
*575020
*575010
*575000
5′ c c t g c t t g a c a a a a a t g a c t t c a c t a a g g a a g g t g c g a 3′
sigma38p10 1.4 bits
sigma38p10 0.0 bits
proximalUP 5.6 bits
... proximalUP-(32)
distalUP 0.1 bits
... distalUP-(41)-s
... sigma38p10-prox
... proximalUP-(37)
distalUP 5.7 bits
... distalUP-(44)-s
... sigma38p10-prox
... proximalUP-(43)
... proximalUP-(37)-sigma38p10 575025 Gap 4.5 bits
... distalUP-(50)-s
... distalUP-(46)-sigma38p10 575025 Gap 4.1 bits
... sigma38p10-proximalUP-distalUP 575025 total 4.1 bits
... proximalUP-(38)-sigma38p10 575024 Gap 3.2 bits
... distalUP-(47)-sigma38p10 575024 Gap 6.5 bits
... sigma38p10-proximalUP-distalUP 575024 total 1.7 bits
... sigma38p10-prox
*574990
*574980
*574970
*574960
5′ a c a a g t c c c t g a t a t g a g a t c a t g t t t g t c a t c t g g a g 3′
sigma38p10 2.7 bits
sigma38p10 0.1 bits

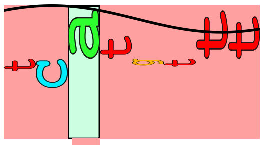

sigma38p10 4.1 bits

```
... --------------------| proximalUP-(32)-sigma38p10 574987 Gap 4.3 bits
... --------------------| distalUP-(41)-sigma38p10 574987 Gap 3.1 bits
... --------------------| sigma38p10-proximalUP-distalUP 574987 total 1.0 bits
... ------------------------------| proximalUP-(37)-sigma38p10 574982 Gap 4.5 bits
... ------------------------------| distalUP-(44)-sigma38p10 574982 Gap 2.8 bits
... ------------------------------| sigma38p10-proximalUP-distalUP 574982 total 4.1 bits
... -----------------------------------------| proximalUP-(43)-sigma38p10 574976 Gap 5.1 bits
... -----------------------------------------| distalUP-(50)-sigma38p10 574976 Gap 4.1 bits

... -----------------------------------------| sigma38p10-proximalUP-distalUP 574976 total 6.1 bits

           .         *574950   .         *574940   .         *574930   .
5' c c a t a g a a c a g g g t t c a t c a t g a g t c a t c a a c t t a c c t 3'

    *574920   .         *574910   .         *574900   .         *574890   .
5' t c g c c g a c a g t g a a t t c a g c a g t a a g c g c c g t c a g a c c 3'

      *574880   .         *574870   .         *574860   .         *574850
5' a g a a a a g a g a t t t t c t t g t c c c g c a t g g a g c a g a t t c t 3'

   .         *574840   .         *574830   .         *574820   .         *574810
5' g c c a t g g c a a a a c a t g g t g g a a g t c a t c g a g c c g t t t t 3'

     .         *574800
5' a c c c c a a g 3'
```

piece 9, NC_000913, ybiI, config: linear, direction: -, begin: 837899, end: 837500

```
         .         *837890    .         *837880    .         *837870    .
5' a t g a t c c g c g a t g a t a a a a a g c g c g c c a t g c t g t t t g a 3'
                                                proximalUP 3.8 bits
   T timestamp:_2018Sep20_11-34-53             |-------------------------------- ... proximalUP-(42)
                                          distalUP 6.5 bits
                              |----------------------------------------------------- ... distalUP-(50)-s
                              |---------------|------------------------------------- ... sigma38p10-prox

    *837860    .         *837850    .         *837840    .         *837830    .
5' a c t g g a c a a c a a a t t c a g t c g c t g a a a a g c c g c t a c g 3'
                                                          sigma38p10 0.6 bits
... -------------------------------------------------| proximalUP-(42)-sigma38p10 837837 Gap 5.1 bits
... -------------------------------------------------| distalUP-(50)-sigma38p10 837837 Gap 4.1 bits
... -------------------------------------------------| sigma38p10-proximalUP-distalUP 837837 total 1.

        *837820    .         *837810    .         *837800    .         *837790
5' g c g a a a g t g a a g a g a t c c t g t c g c t g c t t a a t c t t t a t 3'
                                                          proximalUP 6.8 bits
                                                                |----------- ... proximalUP-(38)
                                                        distalUP 4.7 bits
                                                          |----------------------- ... distalUP-(44)-s
                                                          |-----------|----------- ... sigma38p10-prox
                                                                                   ... proximalUP
                                                                      distalUP 4.7 bits
                                                                |----------- ... distalUP-(44)-s
                                                                |----------- ... sigma38p10-prox

 .         *837780    .         *837770    .         *837760    .         *837750
5' c a t a a t c t g c t g c g g g a a t g g t c g g a g a t c t g a t g c t t 3'
```

sigma38p10 1.8 bits
... ---------------------------------------------------------------| proximalUP-(38)-sigma38p10 837
... sigma38p10
... ---------------------------------------------------------------| distalUP-(44)-sigma38p10 83775
... ---------------------------------------------------------------| sigma38p10-proximalUP-distalUP
... proximalUP 6.8 bits
|------------------------------------------------------------------- ... proximalUP-(38)
... -------------------------------------------------------------------- ... distalUP-(44)-s
... |------------------------------------------------------------------- ... sigma38p10-prox

. *837740 . *837730 . *837720 . *837710
5′ g a a c t g c c c g a g t t c g t c c c t g g a t t a g c t a c a c t t a a 3′
[---------#---------------- ... ybiI

... sigma38p10 2.7 bits
... | proximalUP-(38)-sigma38p10 837747 Gap 3.2 bits
... | distalUP-(44)-sigma38p10 837747 Gap 2.8 bits
... | sigma38p10-proximalUP-distalUP 837747 total 8.1 bits

. *837700 . *837690 . *837680 .
5′ c t g t a c a a g t a t t g a t a t g g g g a g g t c g a t a t g g c a t c 3′
... ---------------------> ybiI

*837670 . *837660 . *837650 . *837640 .
5′ c g g t t g g g c t a a c g a t g a c g c c g t c a a c g a a c a g a t c a 3′
... distalUP
|--------- ... distalUP-(43)-s
|--------- ... sigma38p10-prox

*837630 . *837620 . *837610 . *837600
5′ a c a g t a c a a t t g a a g a t g c g a t t g c c c g c g c t c g g g g t 3′
... distalUP 1.7 bits
... sigma38p10
... -------------------------------------------------------------------- ... distalUP-(43)-s
... -----------------------|-------------------------------------------- ... sigma38p10-prox
proximalUP 2.1 bits

|------------------------------------------------------- ... proximalUP-(27)

. *837590 . *837580 . *837570 . *837560

5' g a a a t t c c g c g c g g c g a a a g c c t g g a t g a a t g t g a a g a 3'

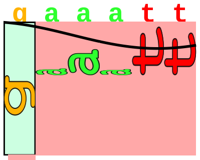

... sigma38p10 3.7 bits

... | distalUP-(43)-sigma38p10 837595 Gap 3.6 bits

... | sigma38p10-proximalUP-distalUP 837595 total 0.2 bits

... | proximalUP-(27)-sigma38p10 837595 Gap 3.8 bits

. *837550 . *837540 . *837530 . *837520

5' g t g c g g t g c c c c c a t c c c g c a g g c c c g t c g g g a a g c c a 3'

. *837510 . *837500

5' t t c c t g g c g t g c g c t t a t g t 3'

piece 10, NC_000913, dps, config: linear, direction: -, begin: 848399, end: 848000

```
             .          *848390   .          *848380   .          *848370   .
5′ c a g a a t t t c t c g c t a c t t t t c c t c t a c a c c g t c t t t a t 3′
```

proximalUP 3.7 bits

distalUP 6.6 bits

... proximalUP-(38)

... distalUP-(56)-s

... sigma38p10-prox

T timestamp:_2018Sep20_11-34-53

```
    *848360   .          *848350   .          *848340   .          *848330   .
5′ a t a t c g a a t t a t g c a a a a g c a t a t t t a t t c c g a a a a t t 3′
```

sigma38p10 4.3 bits

... proximalUP-(38)-sigma38p10 848341 Gap 3.2 bits

... distalUP-(56)-sigma38p10 848341 Gap 3.8 bits

... sigma38p10-proximalUP-distalUP 848341 total 7.6 bits

distalUP 5.9 bits

proximalUP 6.2 bits

... proximalUP-(27)

... distalUP-(51)-s

... sigma38p10-prox

... proximalUP-(34)

distalUP 5.7 bits

... distalUP-(44)-s

... sigma38p10-prox

... proximalUP-(40)

... distalUP-(50)-s

... sigma38p10-prox

... proximalUP

distalUP 7.2 bits

... distalUP-(44)-s

... sigma38p10-prox

```
      *848320   .          *848310   .          *848300   .          *848290
```

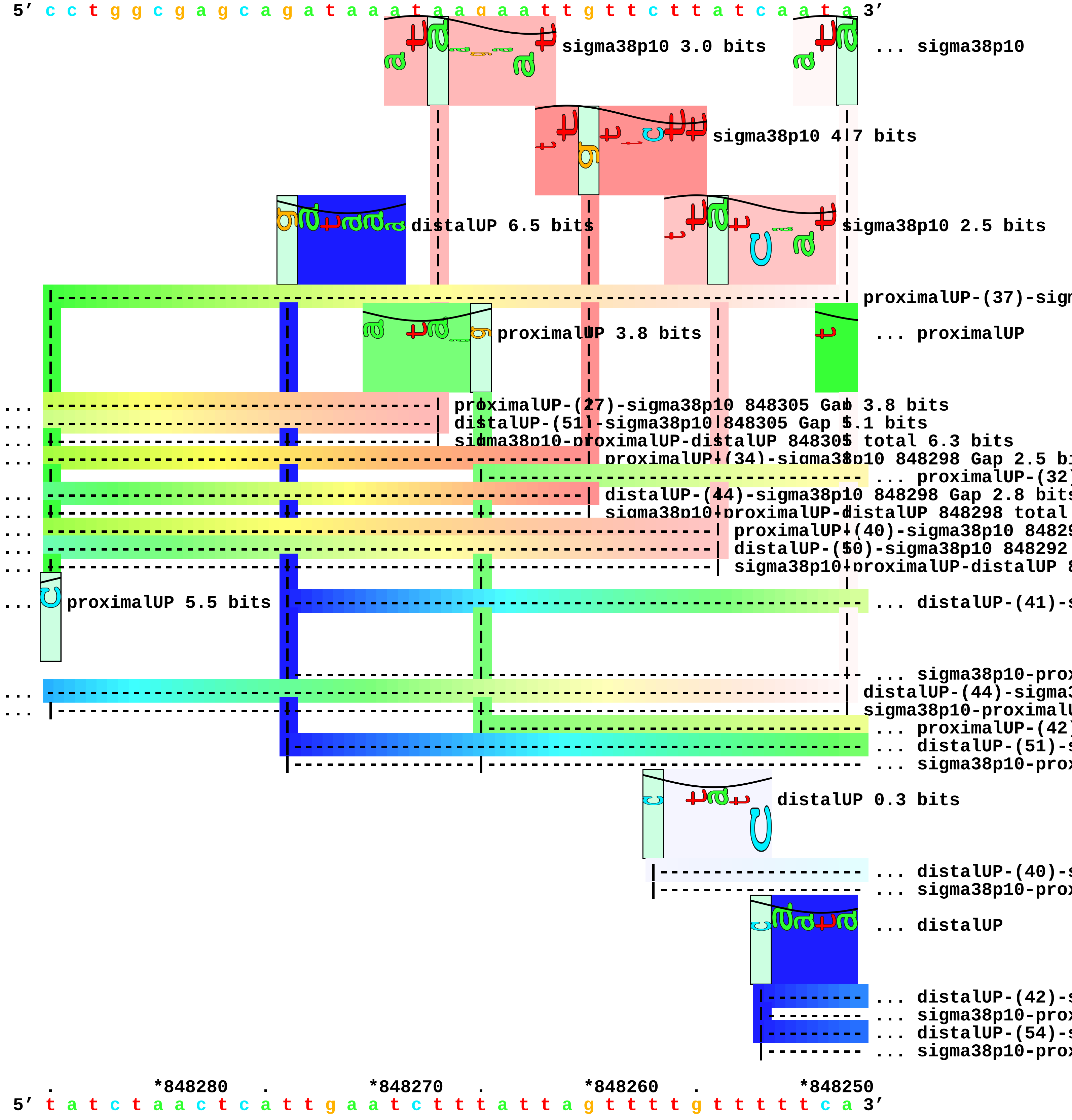
5′ c c t g g c g a g c a g a t a a a t a a g a a t t g t t c t t a t c a a t a 3′
sigma38p10 3.0 bits
... sigma38p10
sigma38p10 4.7 bits
distalUP 6.5 bits
sigma38p10 2.5 bits
proximalUP 3.8 bits
... proximalUP
proximalUP-(27)-sigma38p10 848305 Gap 3.8 bits
distalUP-(51)-sigma38p10 848305 Gap 5.1 bits
sigma38p10-proximalUP-distalUP 848305 total 6.3 bits
proximalUP-(34)-sigma38p10 848298 Gap 2.5 bi
distalUP-(44)-sigma38p10 848298 Gap 2.8 bits
proximalUP 5.5 bits
distalUP 0.3 bits
... distalUP
*848280
*848270
*848260
*848250
5′ t a t c t a a c t c a t t g a a t c t t t a t t a g t t t t g t t t t t c a 3′

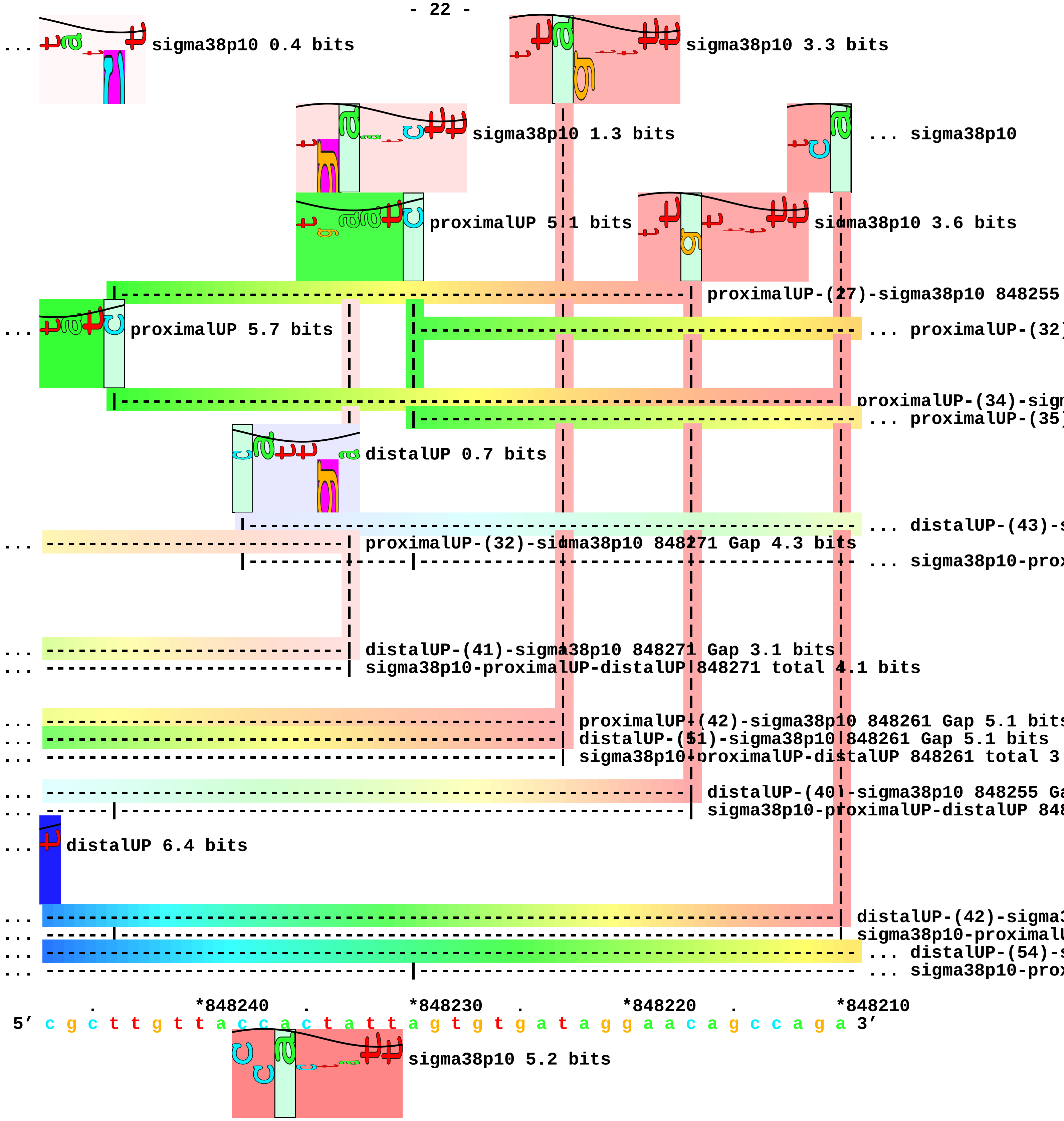
... sigma38p10 0.4 bits
sigma38p10 3.3 bits
sigma38p10 1.3 bits
... sigma38p10
proximalUP 5.1 bits
sigma38p10 3.6 bits
proximalUP-(27)-sigma38p10 848255
... proximalUP 5.7 bits
... proximalUP-(32)
proximalUP-(34)-sign
... proximalUP-(35)
distalUP 0.7 bits
... distalUP-(43)-s
... proximalUP-(32)-sigma38p10 848271 Gap 4.3 bits
... sigma38p10-prox
... distalUP-(41)-sigma38p10 848271 Gap 3.1 bits
... sigma38p10-proximalUP-distalUP 848271 total 4.1 bits
... proximalUP-(42)-sigma38p10 848261 Gap 5.1 bits
... distalUP-(51)-sigma38p10 848261 Gap 5.1 bits
... sigma38p10-proximalUP-distalUP 848261 total 3.
... distalUP-(40)-sigma38p10 848255 Ga
... sigma38p10-proximalUP-distalUP 848
... distalUP 6.4 bits
... distalUP-(42)-sigma3
... sigma38p10-proximalU
... distalUP-(54)-s
... sigma38p10-prox
*848240
*848230
*848220
*848210
5′ c g c t t g t t a c c a c t a t t a g t g t g a t a g g a a c a g c c a g a 3′
sigma38p10 5.2 bits

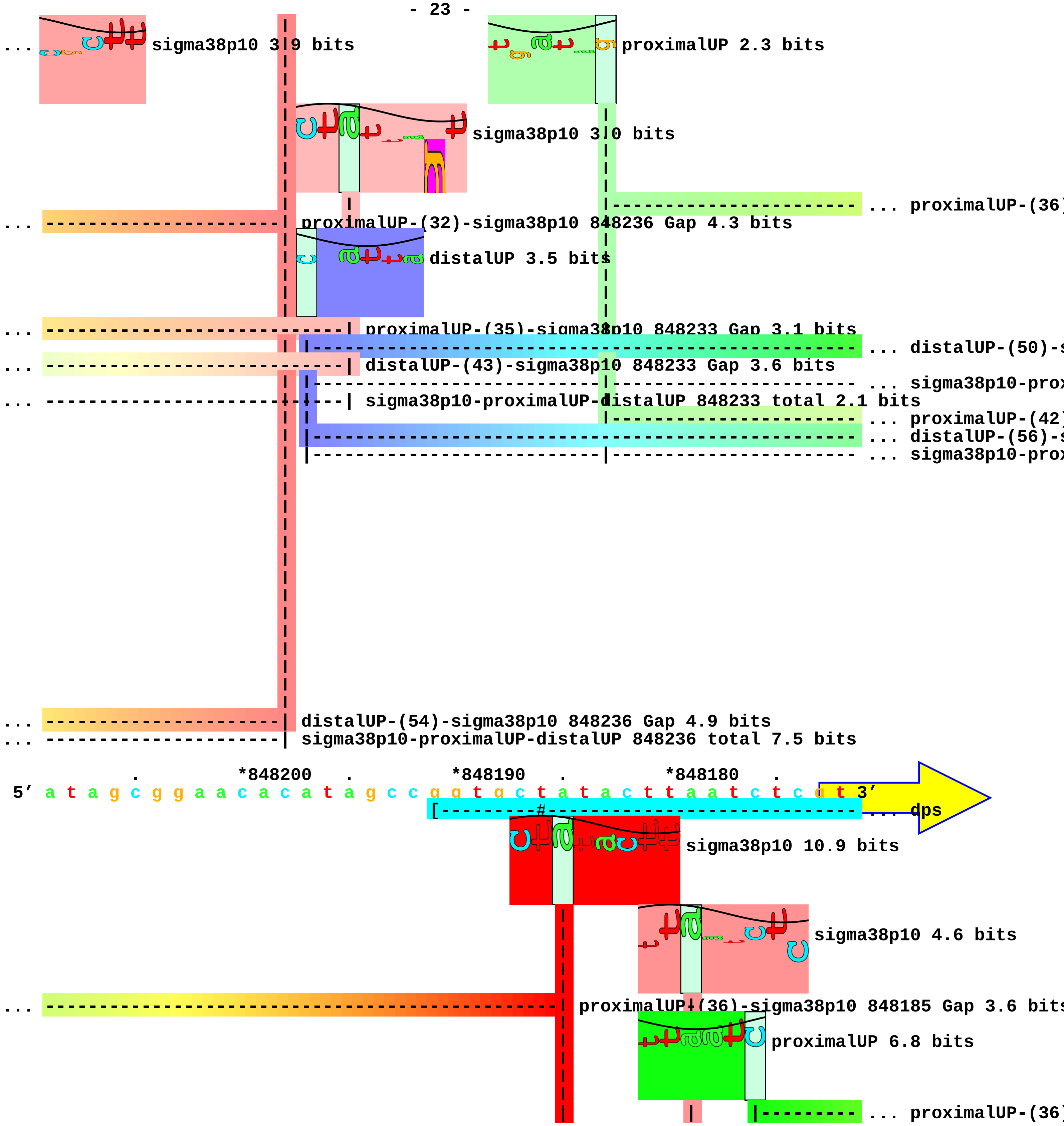
... sigma38p10 3.9 bits
proximalUP 2.3 bits
sigma38p10 3.0 bits
... proximalUP-(36)
... proximalUP-(32)-sigma38p10 848236 Gap 4.3 bits
distalUP 3.5 bits
... proximalUP-(35)-sigma38p10 848233 Gap 3.1 bits
... distalUP-(50)-s
... distalUP-(43)-sigma38p10 848233 Gap 3.6 bits
... sigma38p10-prox
... sigma38p10-proximalUP-distalUP 848233 total 2.1 bits
... proximalUP-(42)
... distalUP-(56)-s
... sigma38p10-prox
... distalUP-(54)-sigma38p10 848236 Gap 4.9 bits
... sigma38p10-proximalUP-distalUP 848236 total 7.5 bits
*848200 *848190 *848180
5' a t a g c g g a a c a c a t a g c c g g t g c t a t a c t t a a t c t c g t 3'
... dps
sigma38p10 10.9 bits
sigma38p10 4.6 bits
... proximalUP-(36)-sigma38p10 848185 Gap 3.6 bits
proximalUP 6.8 bits
... proximalUP-(36)

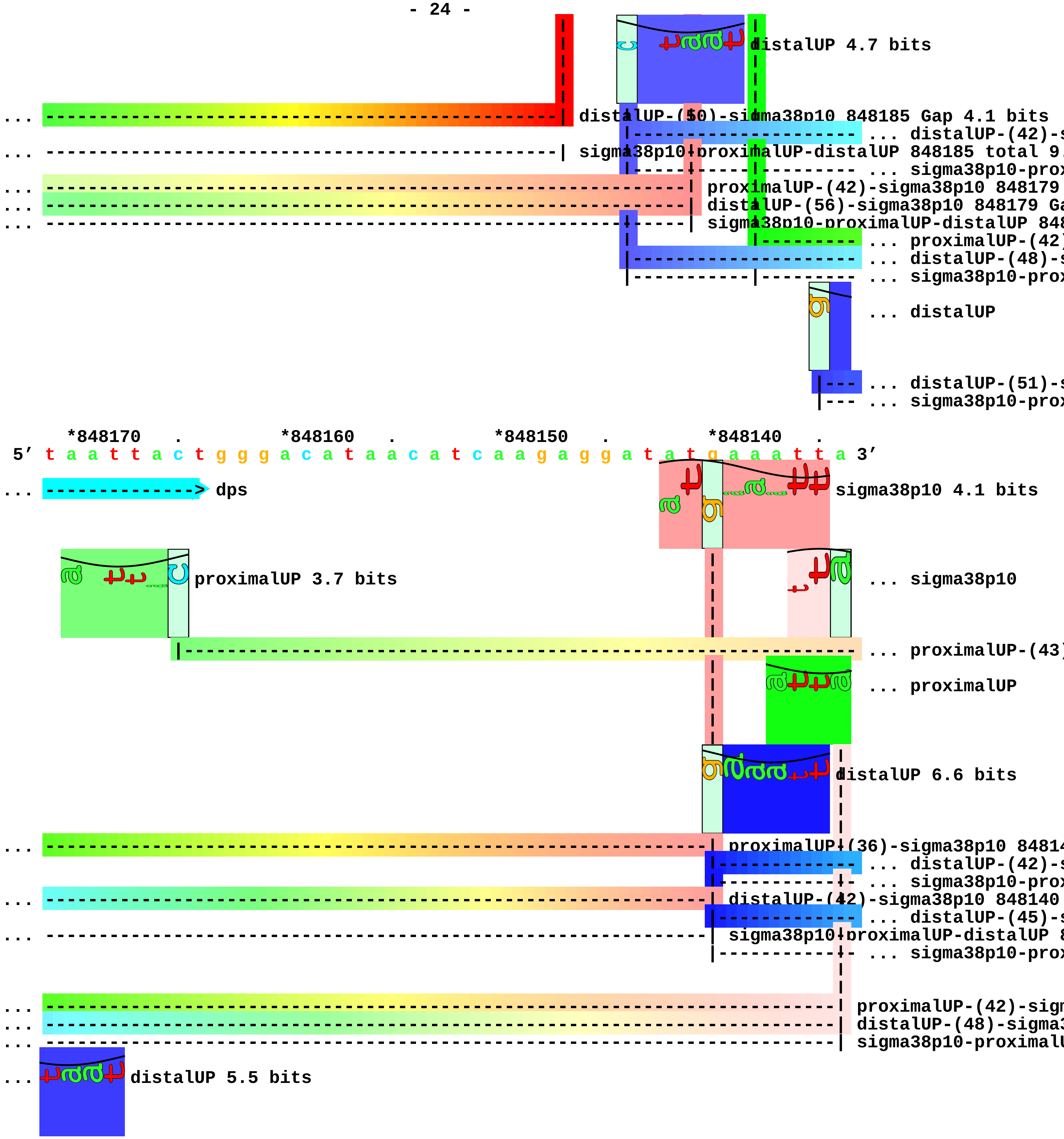
distalUP 4.7 bits
distalUP-(50)-sigma38p10 848185 Gap 4.1 bits
... distalUP-(42)-s
sigma38p10-proximalUP-distalUP 848185 total 9.
... sigma38p10-prox
proximalUP-(42)-sigma38p10 848179
distalUP-(56)-sigma38p10 848179 Ga
sigma38p10-proximalUP-distalUP 848
... proximalUP-(42)
... distalUP-(48)-s
... sigma38p10-prox
... distalUP
... distalUP-(51)-s
... sigma38p10-prox
*848170 . *848160 . *848150 . *848140 .
5' t a a t t a c t g g g a c a t a a c a t c a a g a g g a t a t g a a a t t a 3'
... dps
sigma38p10 4.1 bits
proximalUP 3.7 bits
... sigma38p10
... proximalUP-(43)
... proximalUP
distalUP 6.6 bits
proximalUP-(36)-sigma38p10 84814
... distalUP-(42)-s
... sigma38p10-prox
distalUP-(42)-sigma38p10 848140
... distalUP-(45)-s
sigma38p10-proximalUP-distalUP 8
... sigma38p10-prox
proximalUP-(42)-sign
distalUP-(48)-sigma3
sigma38p10-proximalU
... distalUP 5.5 bits

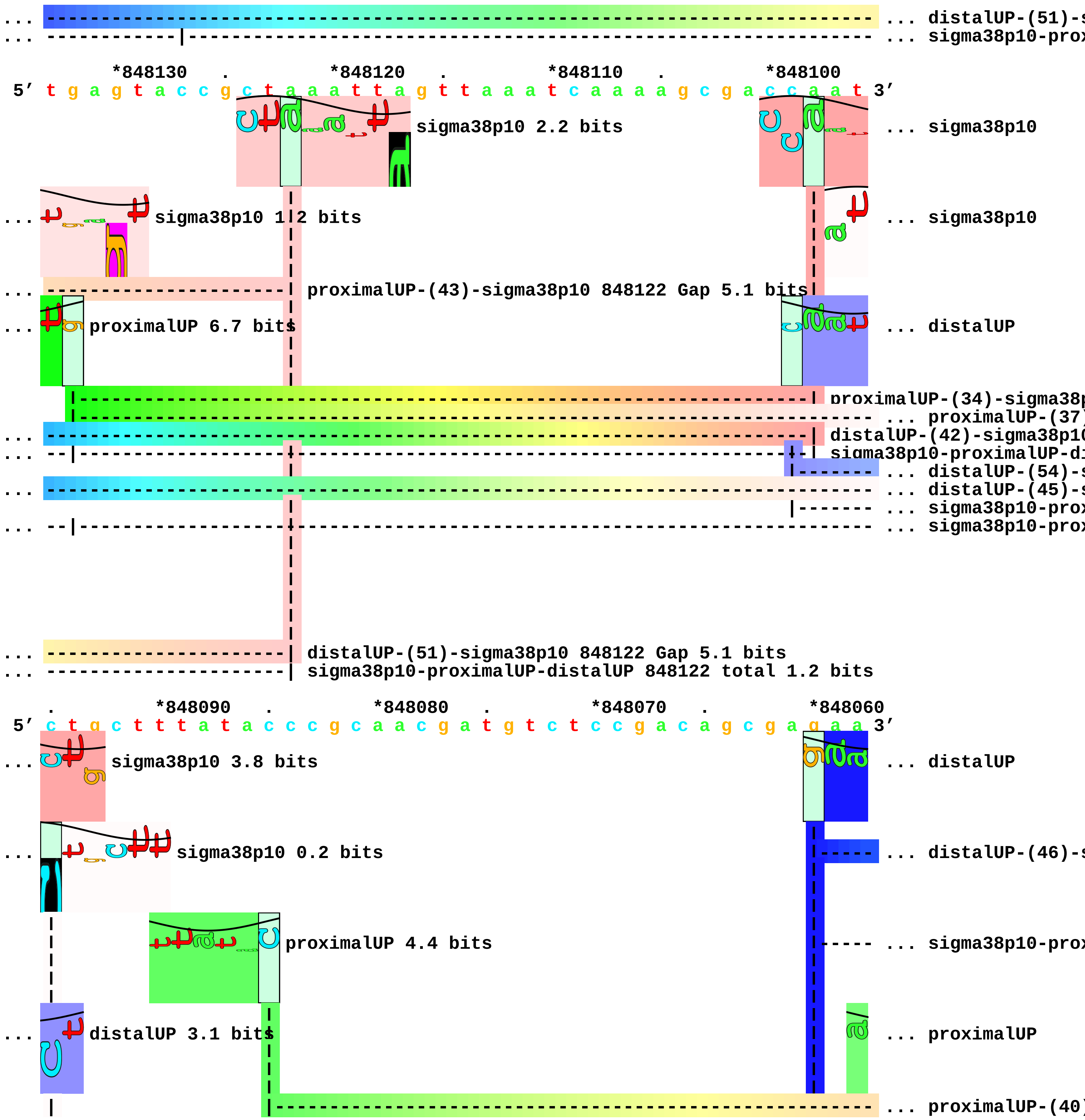
... distalUP-(51)-s
... sigma38p10-prox
*848130 . *848120 . *848110 . *848100
5′ t g a g t a c c g c t a a a t t a g t t a a a t c a a a a g c g a c c a a t 3′
sigma38p10 2.2 bits
... sigma38p10
... sigma38p10 1.2 bits
... sigma38p10
... proximalUP-(43)-sigma38p10 848122 Gap 5.1 bits
... proximalUP 6.7 bits
... distalUP
proximalUP-(34)-sigma38p
... proximalUP-(37)
... distalUP-(42)-sigma38p10
... sigma38p10-proximalUP-di
... distalUP-(54)-s
... distalUP-(45)-s
... sigma38p10-prox
... sigma38p10-prox
... distalUP-(51)-sigma38p10 848122 Gap 5.1 bits
... sigma38p10-proximalUP-distalUP 848122 total 1.2 bits
. *848090 . *848080 . *848070 . *848060
5′ c t g c t t t a t a c c c g c a a c g a t g t c t c c g a c a g c g a g a a 3′
... sigma38p10 3.8 bits
... distalUP
... sigma38p10 0.2 bits
... distalUP-(46)-s
proximalUP 4.4 bits
... sigma38p10-prox
... distalUP 3.1 bits
... proximalUP
... proximalUP-(40)

```
... | proximalUP-(37)-sigma38p10 848095 Gap 4.5 bits                        |----- ... distalUP-(49)-s
    |                                                                       |----- ... sigma38p10-prox
    |
... --------------------------------------------------------------------------- ... distalUP-(54)-s
... | distalUP-(45)-sigma38p10 848095 Gap 5.5 bits
... --------------------|------------------------------------------------------ ... sigma38p10-prox
... | sigma38p10-proximalUP-distalUP 848095 total 3.5 bits

          .         *848050    .         *848040    .         *848030    .         *848020
 5' a a a a g c a a c a g t a g a g t t g c t g a a t c g c c a g g t t a t c c 3'
... distalUP 6.5 bits
... --------------------------------------------------------------------------- ... distalUP-(46)-s
... ----------------|---------------------------------------------------------- ... sigma38p10-prox
... proximalUP 3.8 bits
... ------------------------| proximalUP-(40)-sigma38p10 848045 Gap 4.0 bits
... --------------------------------------------------------------------------- ... distalUP-(49)-s
... --------------------------------------------------------------------------- ... sigma38p10-prox
    sigma38p10 1.7 bits
... ------------------------| distalUP-(54)-sigma38p10 848045 Gap 4.9 bits
    proximalUP 0.2 bits
... --------------------|-------| sigma38p10-proximalUP-distalUP 848045 total 0.4 bits
                        |------------------------------------------------------ ... proximalUP-(35)
... |---------------------------------------------------------------------------- ... proximalUP-(42)

          .         *848010    .         *848000
 5' a g t t t a t t g a t c t t t c t t t g 3'
    sigma38p10 2.0 bits
... ----------| distalUP-(46)-sigma38p10 848014 Gap 4.1 bits
... ----------| sigma38p10-proximalUP-distalUP 848014 total 1.5 bits
    sigma38p10 4.1 bits
... ----------------| distalUP-(49)-sigma38p10 848011 Gap 5.9 bits
... ----------------| sigma38p10-proximalUP-distalUP 848011 total 3.5 bits
... ----------| proximalUP-(35)-sigma38p10 848014 Gap 3.1 bits
```

... ----------------| proximalUP-(42)-sigma38p10 848011 Gap 5.1 bits piece 11, NC_000913, ymgC, config: linear, direction: +, begin: 1215850, end: 1216449

```
    *1215850   .         *1215860   .         *1215870   .         *1215880   .
5' g a t a t g t a a c t t c t a a t t a t t a a g t a t a a g t t t a t a g a 3'
```

proximalUP 3.3 bits ... proximalUP-(30)

... proximalUP-(43)

distalUP 4.2 bits

proximalUP 3.9 bits

... distalUP-(40)-s

... proximalUP-dist

T timestamp:_2018Sep20_11-34-53

proximalUP 4.9 bits ... proximalUP-(40)

... distalUP-(41)-s

... distalUP-proxim

... distalUP

... distalUP-(40)-s

... proximalUP-dist

... proximalUP-(42)

... distalUP-(42)-s

... proximalUP-dist

distalUP 5.5 bits

... distalUP-(56)-s

... distalUP-proxim

```
      *1215890   .         *1215900   .         *1215910   .         *1215920   .
5' a a a c t c a t t c a t c t t a t t t t t g t c t g t c g c t t t a g a c t 3'
```

... proximalUP-(30)-sigma38p10 1215903 Gap 5.9 bits

... proximalUP-(43)-sigma38p10 1215902 Gap 5.1 bits

sigma38p10 0.8 bits ... sigma38p10

... distalUP-(40)-sigma38p10 1215902 Gap 3.1 bits

... proximalUP-distalUP-sigma38p10 1215902 total 0.1 bits

... proximalUP-(40)

... distalUP-(41)-sigma38p10 1215903 Gap 3.1 bits

... distalUP-proximalUP-sigma38p10 1215903 total 7.1 bits

... distalUP 3.4 bits

sigma38p10 7.0 bits

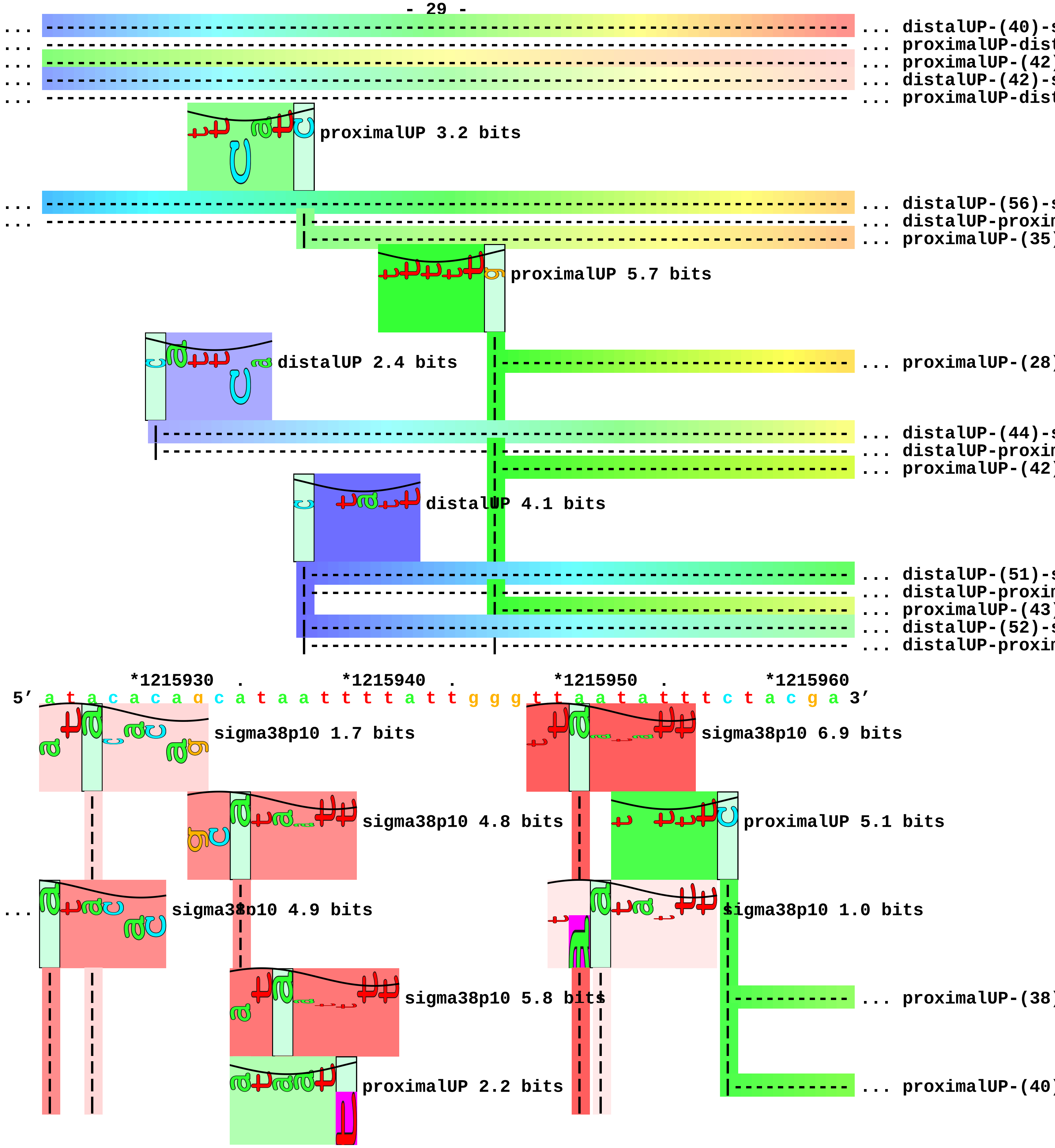
... distalUP-(40)-s
... proximalUP-dist
... proximalUP-(42)
... distalUP-(42)-s
... proximalUP-dist
proximalUP 3.2 bits
... distalUP-(56)-s
... distalUP-proxim
... proximalUP-(35)
proximalUP 5.7 bits
distalUP 2.4 bits
... proximalUP-(28)
... distalUP-(44)-s
... distalUP-proxim
... proximalUP-(42)
distalUP 4.1 bits
... distalUP-(51)-s
... distalUP-proxim
... proximalUP-(43)
... distalUP-(52)-s
... distalUP-proxim
*1215930 . *1215940 . *1215950 . *1215960
5' a t a c a c a g c a t a a t t t t a t t g g g t t a a t a t t t c t a c g a 3'
sigma38p10 1.7 bits
sigma38p10 6.9 bits
sigma38p10 4.8 bits
proximalUP 5.1 bits
sigma38p10 4.9 bits
sigma38p10 1.0 bits
sigma38p10 5.8 bits
... proximalUP-(38)
proximalUP 2.2 bits
... proximalUP-(40)

```
... | proximalUP-(40)-sigma38p10 1215926 Gap 4.0 bits
                                                                  ... proximalUP-(35)
      distalUP 6.3 bits
                                                                  ... distalUP-(41)-s
... | distalUP-(40)-sigma38p10 1215926 Gap 3.1 bits
... | proximalUP-distalUP-sigma38p10 1215926 total 5.1 bits
... ----| proximalUP-(42)-sigma38p10 1215928 Gap 5.1 bits
... ----| distalUP-(42)-sigma38p10 1215928 Gap 3.3 bits
... ----| proximalUP-distalUP-sigma38p10 1215928 total 0.6 bits
                                                                  ... distalUP-proxim
... ------------------| distalUP-(56)-sigma38p10 1215935 Gap 3.8 bits
... ------------------| distalUP-proximalUP-sigma38p10 1215935 total 6.7 bits
... ------------------| proximalUP-(35)-sigma38p10 1215935 Gap 3.1 bits
      distalUP 5.5 bits
... ----------------------| proximalUP-(28)-sigma38p10 1215937 Gap 5.5 bits
... ----------------------| distalUP-(44)-sigma38p10 1215937 Gap 2.8 bits
... ----------------------| distalUP-proximalUP-sigma38p10 1215937 total 5.6 bits
... ---------------------------------------| proximalUP-(42)-sigma38p10 1215951 Gap 5.1 b
                                                                  ... distalUP-(48)-s
... ---------------------------------------| distalUP-(51)-sigma38p10 1215951 Gap 5.1 bit
... ---------------------------------------| distalUP-proximalUP-sigma38p10 1215951 total
... -----------------------------------------| proximalUP-(43)-sigma38p10 1215952 Gap 5.1
... -----------------------------------------| distalUP-(52)-sigma38p10 1215952 Gap 5.5 b
... -----------------------------------------| distalUP-proximalUP-sigma38p10 1215952 tot
                                                                  ... distalUP-proxim
                                                                  ... distalUP-(50)-s
                                                                  ... distalUP-proxim

      .        *1215970   .          *1215980   .          *1215990   .          *1216000
5' g g c t g a c a t g a a t a a t t c a a t c c c a g a g a g a t t t a t t t 3'
      sigma38p10 0.5 bits                                   sigma38p10 0.3 bits
      proximalUP 5.1 bits                                   ... sigma38p10
                                                                  ... proximalUP-(32)
... --------------------------------------------------| proximalUP-(38)-sigma38p10 121
... ----------------------------------------------------| proximalUP-(40)-sigma38p10
      distalUP 6.5 bits                                     ... proximalUP
... ----------------------| proximalUP-(35)-sigma38p10 1215975 Gap 3.1 bits
                                                                  ... distalUP-(40)-s
... ----------------------| distalUP-(41)-sigma38p10 1215975 Gap 3.1 bits
                                                                  ... distalUP-proxim
```

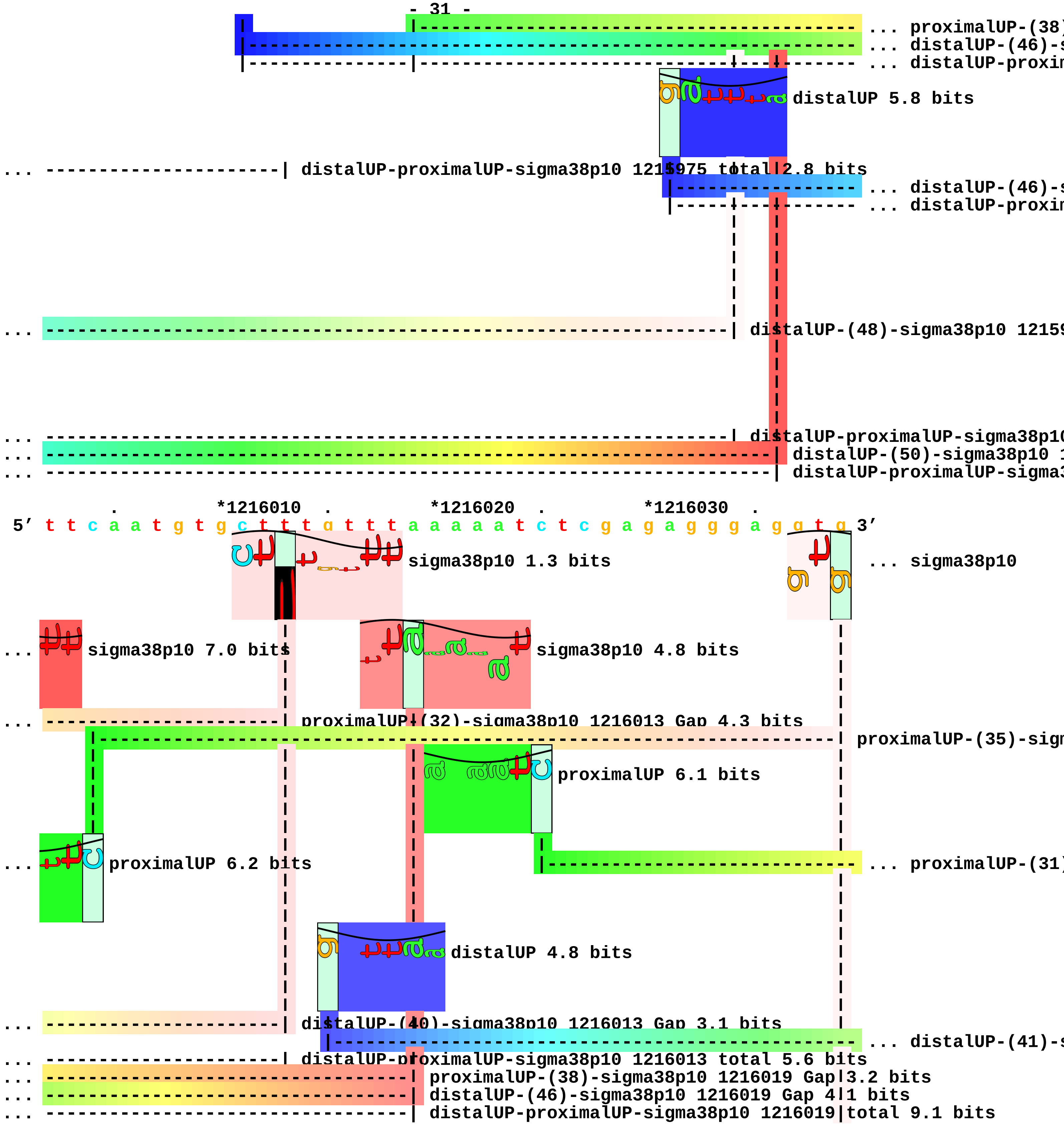
... proximalUP-(38)
... distalUP-(46)-s
... distalUP-proxim
distalUP 5.8 bits
... ---------------------| distalUP-proximalUP-sigma38p10 1215975 total 2.8 bits
... distalUP-(46)-s
... distalUP-proxim
... distalUP-(48)-sigma38p10 1215
... distalUP-proximalUP-sigma38p10
... distalUP-(50)-sigma38p10 1
... distalUP-proximalUP-sigma3
*1216010
*1216020
*1216030
5′ t t c a a t g t g c t t t g t t t a a a a a t c t c g a g a g g g a g g t g 3′
sigma38p10 1.3 bits
... sigma38p10
... sigma38p10 7.0 bits
sigma38p10 4.8 bits
... proximalUP-(32)-sigma38p10 1216013 Gap 4.3 bits
proximalUP-(35)-sigm
proximalUP 6.1 bits
... proximalUP 6.2 bits
... proximalUP-(31)
distalUP 4.8 bits
... distalUP-(40)-sigma38p10 1216013 Gap 3.1 bits
... distalUP-(41)-s
... distalUP-proximalUP-sigma38p10 1216013 total 5.6 bits
... proximalUP-(38)-sigma38p10 1216019 Gap 3.2 bits
... distalUP-(46)-sigma38p10 1216019 Gap 4.1 bits
... distalUP-proximalUP-sigma38p10 1216019 total 9.1 bits

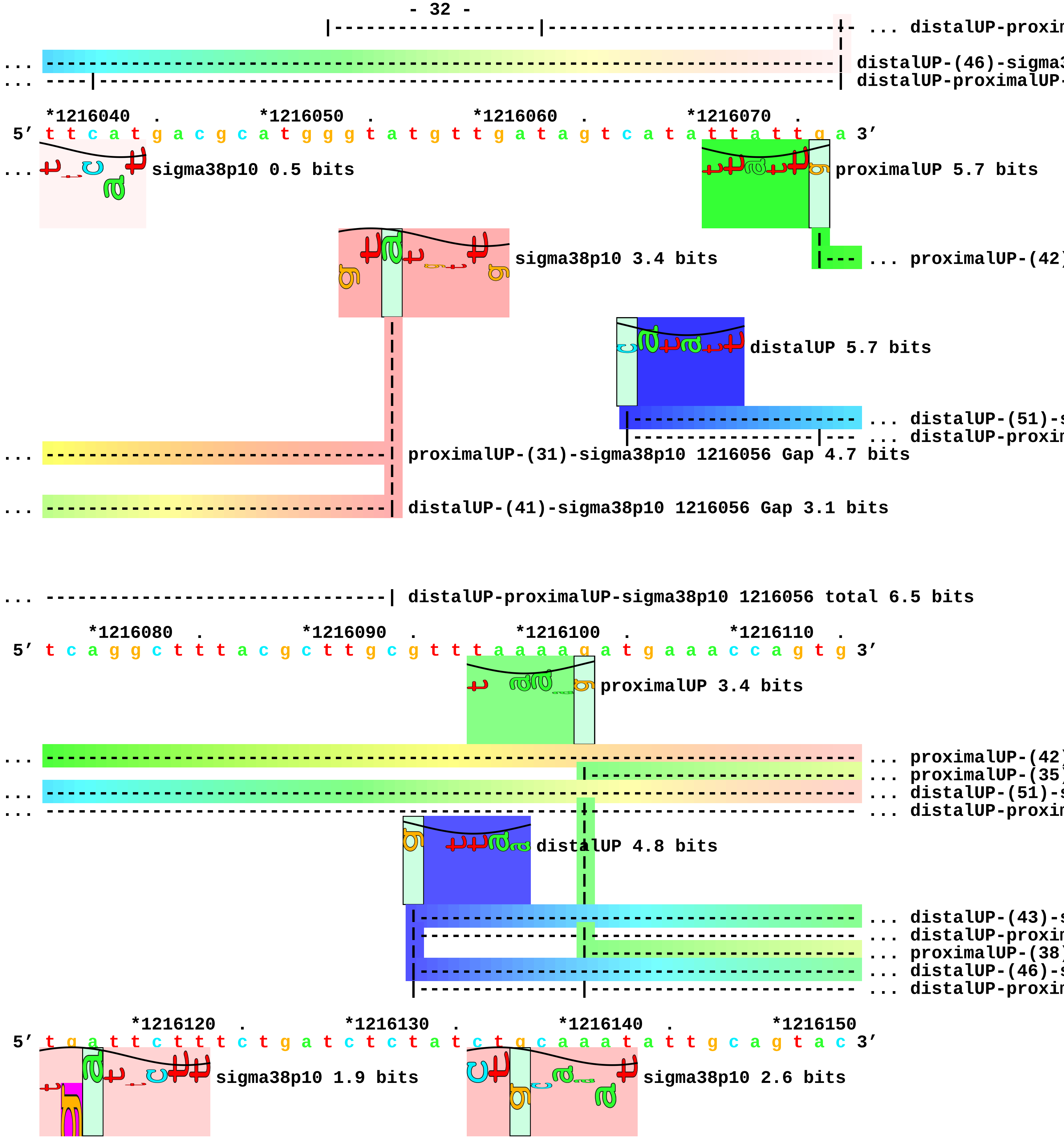
... distalUP-proxin
... distalUP-(46)-sigma3
... distalUP-proximalUP-
*1216040 . *1216050 . *1216060 . *1216070 .
5′ t t c a t g a c g c a t g g g t a t g t t g a t a g t c a t a t t a t t g a 3′
... sigma38p10 0.5 bits
proximalUP 5.7 bits
sigma38p10 3.4 bits
... proximalUP-(42)
distalUP 5.7 bits
... distalUP-(51)-s
... distalUP-proxin
... proximalUP-(31)-sigma38p10 1216056 Gap 4.7 bits
... distalUP-(41)-sigma38p10 1216056 Gap 3.1 bits
... distalUP-proximalUP-sigma38p10 1216056 total 6.5 bits
*1216080 . *1216090 . *1216100 . *1216110 .
5′ t c a g g c t t t a c g c t t g c g t t t a a a a g a t g a a a c c a g t g 3′
proximalUP 3.4 bits
... proximalUP-(42)
... proximalUP-(35)
... distalUP-(51)-s
... distalUP-proxin
distalUP 4.8 bits
... distalUP-(43)-s
... distalUP-proxin
... proximalUP-(38)
... distalUP-(46)-s
... distalUP-proxin
*1216120 . *1216130 . *1216140 . *1216150
5′ t g a t t c t t t c t g a t c t c t a t c t g c a a a t a t t g c a g t a c 3′
sigma38p10 1.9 bits
sigma38p10 2.6 bits

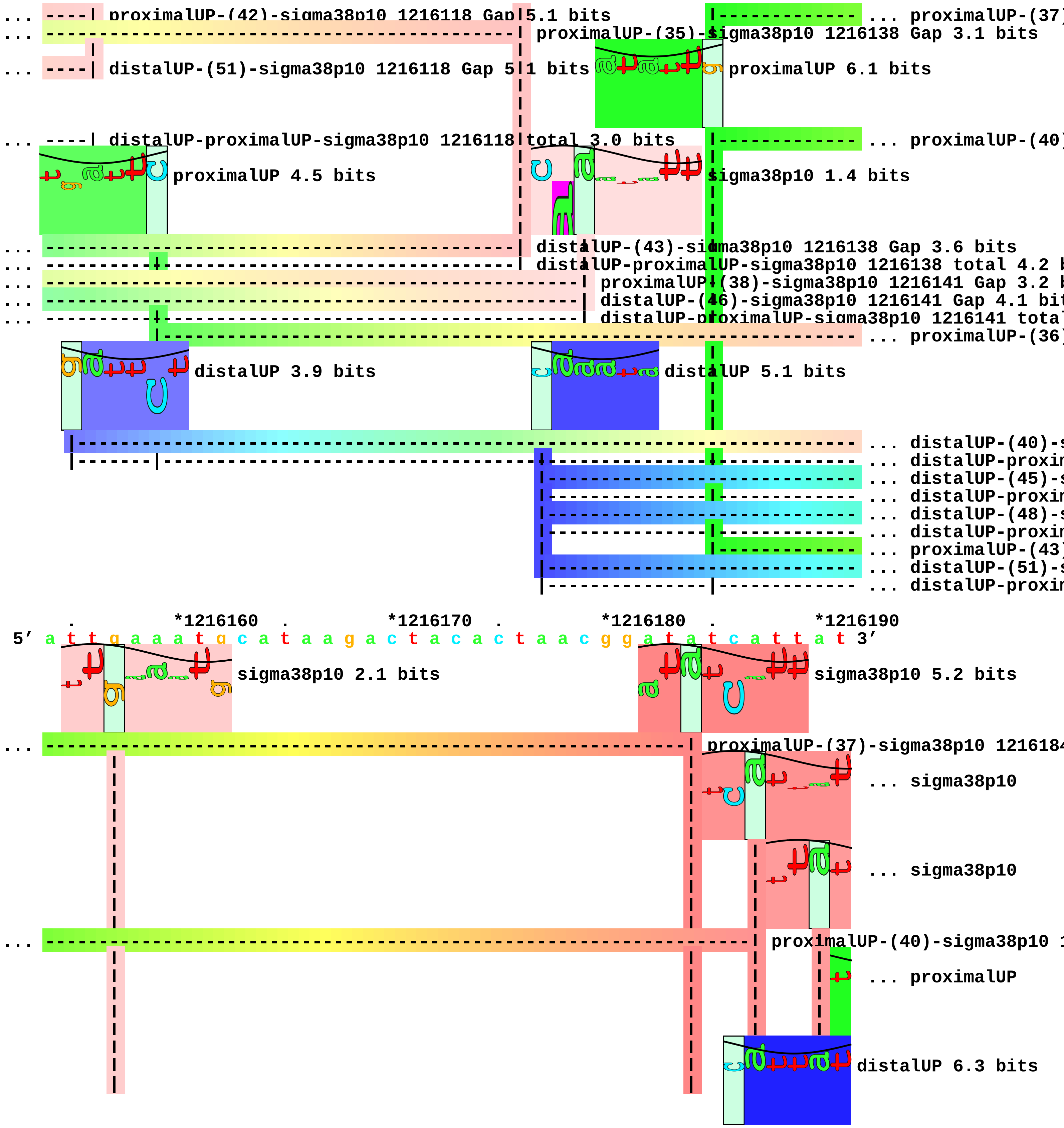
... proximalUP-(42)-sigma38p10 1216118 Gap 5.1 bits
... proximalUP-(37)
... proximalUP-(35)-sigma38p10 1216138 Gap 3.1 bits
... distalUP-(51)-sigma38p10 1216118 Gap 5.1 bits
proximalUP 6.1 bits
... distalUP-proximalUP-sigma38p10 1216118 total 3.0 bits
... proximalUP-(40)
proximalUP 4.5 bits
sigma38p10 1.4 bits
distalUP-(43)-sigma38p10 1216138 Gap 3.6 bits
distalUP-proximalUP-sigma38p10 1216138 total 4.2 b
proximalUP-(38)-sigma38p10 1216141 Gap 3.2 b
distalUP-(46)-sigma38p10 1216141 Gap 4.1 bit
distalUP-proximalUP-sigma38p10 1216141 tota
... proximalUP-(36)
distalUP 3.9 bits
distalUP 5.1 bits
... distalUP-(40)-s
... distalUP-proxim
... distalUP-(45)-s
... distalUP-proxim
... distalUP-(48)-s
... distalUP-proxim
... proximalUP-(43)
... distalUP-(51)-s
... distalUP-proxim
. *1216160 . *1216170 . *1216180 . *1216190
5′ a t t g a a a t g c a t a a g a c t a c a c t a a c g g a t a t c a t t a t 3′
sigma38p10 2.1 bits
sigma38p10 5.2 bits
proximalUP-(37)-sigma38p10 121618
... sigma38p10
... sigma38p10
proximalUP-(40)-sigma38p10 1
... proximalUP
distalUP 6.3 bits

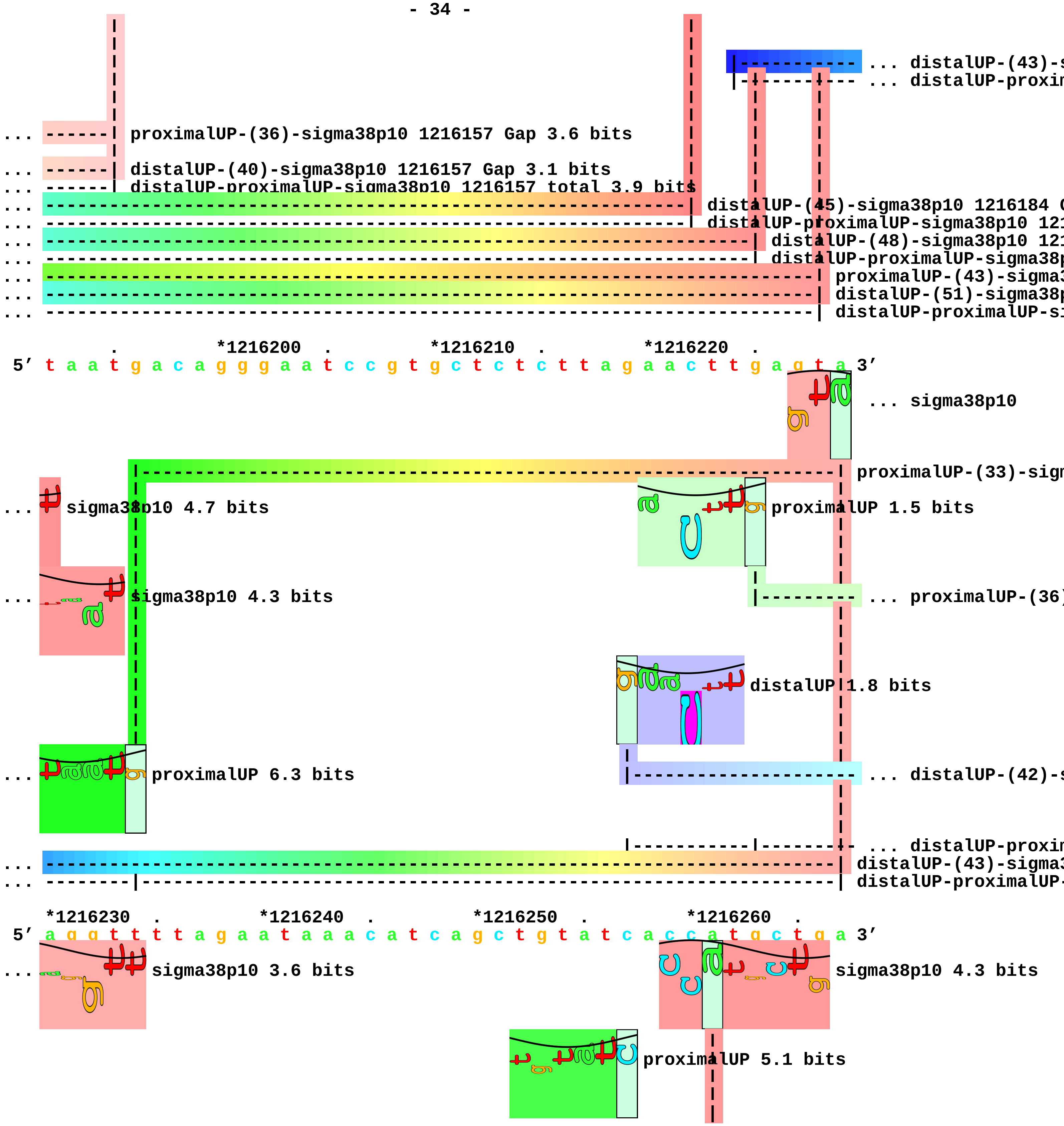
... distalUP-(43)-s
... distalUP-proxim
... proximalUP-(36)-sigma38p10 1216157 Gap 3.6 bits
... distalUP-(40)-sigma38p10 1216157 Gap 3.1 bits
... distalUP-proximalUP-sigma38p10 1216157 total 3.9 bits
... distalUP-(45)-sigma38p10 1216184 G
... distalUP-proximalUP-sigma38p10 121
... distalUP-(48)-sigma38p10 121
... distalUP-proximalUP-sigma38p
... proximalUP-(43)-sigma3
... distalUP-(51)-sigma38p
... distalUP-proximalUP-si
*1216200
*1216210
*1216220
5′ t a a t g a c a g g g a a t c c g t g c t c t c t t a g a a c t t g a g t a 3′
... sigma38p10
proximalUP-(33)-sigm
... sigma38p10 4.7 bits
proximalUP 1.5 bits
... sigma38p10 4.3 bits
... proximalUP-(36)
distalUP 1.8 bits
... proximalUP 6.3 bits
... distalUP-(42)-s
... distalUP-proxim
... distalUP-(43)-sigma3
... distalUP-proximalUP-
*1216230
*1216240
*1216250
*1216260
5′ a g g t t t t t a g a a t a a a c a t c a g c t g t a t c a c c a t g c t g a 3′
... sigma38p10 3.6 bits
sigma38p10 4.3 bits
proximalUP 5.1 bits

```
distalUP 6.5 bits                                  ... proximalUP-(35)
... proximalUP-(36)-sigma38p10 12162
... distalUP-(54)-s
... distalUP-(42)-sigma38p10 1216261
... distalUP-proximalUP-sigma38p10 1
... distalUP-proxim

        *1216270  .       *1216280  .       *1216290  .       *1216300  .
5' t g c a a a g t g a g c a t t t c a g g c g t t a t g c t t t c t t a t t a 3'
sigma38p10 8.0 bits
distalUP 4.1 bits
... proximalUP-(35)-sigma38p10 1216292 Gap 3.1 bit
... distalUP-(51)-s
... distalUP-(54)-sigma38p10 1216292 Gap 4.9 bits
... distalUP-proxim
... distalUP-proximalUP-sigma38p10 1216292 total 1

        *1216310  .       *1216320  .       *1216330  .       *1216340
5' t g t c c g c a a t a t c a g g t g t c a a g a a t g g a g a g t t c t c g 3'
proximalUP 6.1 bits                                ... proximalUP-(31)
... proximalUP-(32)
distalUP 6.4 bits   proximalUP 5.0 bits
... distalUP-(51)-s
... distalUP-(51)-s
... distalUP-proxim
... distalUP-proxim

  .     *1216350  .       *1216360  .       *1216370  .       *1216380
5' c t c t c c a t t c t t g a c g c c t g a t a t c c c g c c t a a c t t a t 3'
... proximalUP-(31)-sigma38p10 1216363 Gap 4.7 bits
... proximalUP-(32)-sigma38p10 1216350 Gap 4.3 bits      ... proximalUP
sigma38p10 6.9 bits                                       ... distalUP
... distalUP-(51)-sigma38p10 1216350 Gap 5.1 bits         ... distalUP-(48)-s
```

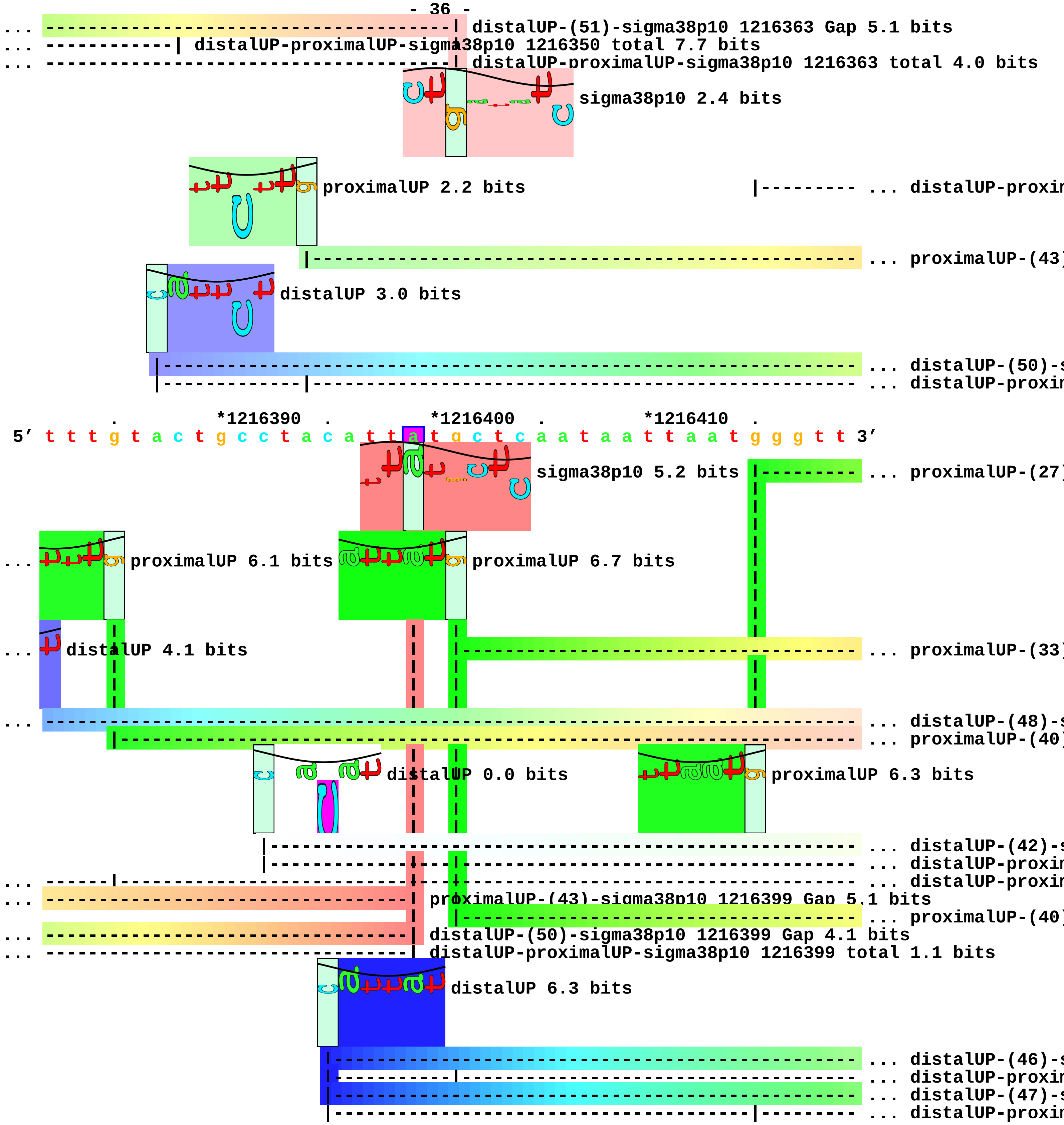
distalUP-(51)-sigma38p10 1216363 Gap 5.1 bits
distalUP-proximalUP-sigma38p10 1216350 total 7.7 bits
distalUP-proximalUP-sigma38p10 1216363 total 4.0 bits
sigma38p10 2.4 bits
proximalUP 2.2 bits
distalUP-proxin
proximalUP-(43)
distalUP 3.0 bits
distalUP-(50)-s
distalUP-proxin
*1216390
*1216400
*1216410
5′ t t t g t a c t g c c t a c a t t a t g c t c a a t a a t t a a t g g g t t 3′
sigma38p10 5.2 bits
proximalUP-(27)
proximalUP 6.1 bits
proximalUP 6.7 bits
distalUP 4.1 bits
proximalUP-(33)
distalUP-(48)-s
proximalUP-(40)
distalUP 0.0 bits
proximalUP 6.3 bits
distalUP-(42)-s
distalUP-proxin
distalUP-proxin
proximalUP-(43)-sigma38p10 1216399 Gap 5.1 bits
proximalUP-(40)
distalUP-(50)-sigma38p10 1216399 Gap 4.1 bits
distalUP-proximalUP-sigma38p10 1216399 total 1.1 bits
distalUP 6.3 bits
distalUP-(46)-s
distalUP-proxin
distalUP-(47)-s
distalUP-proxin

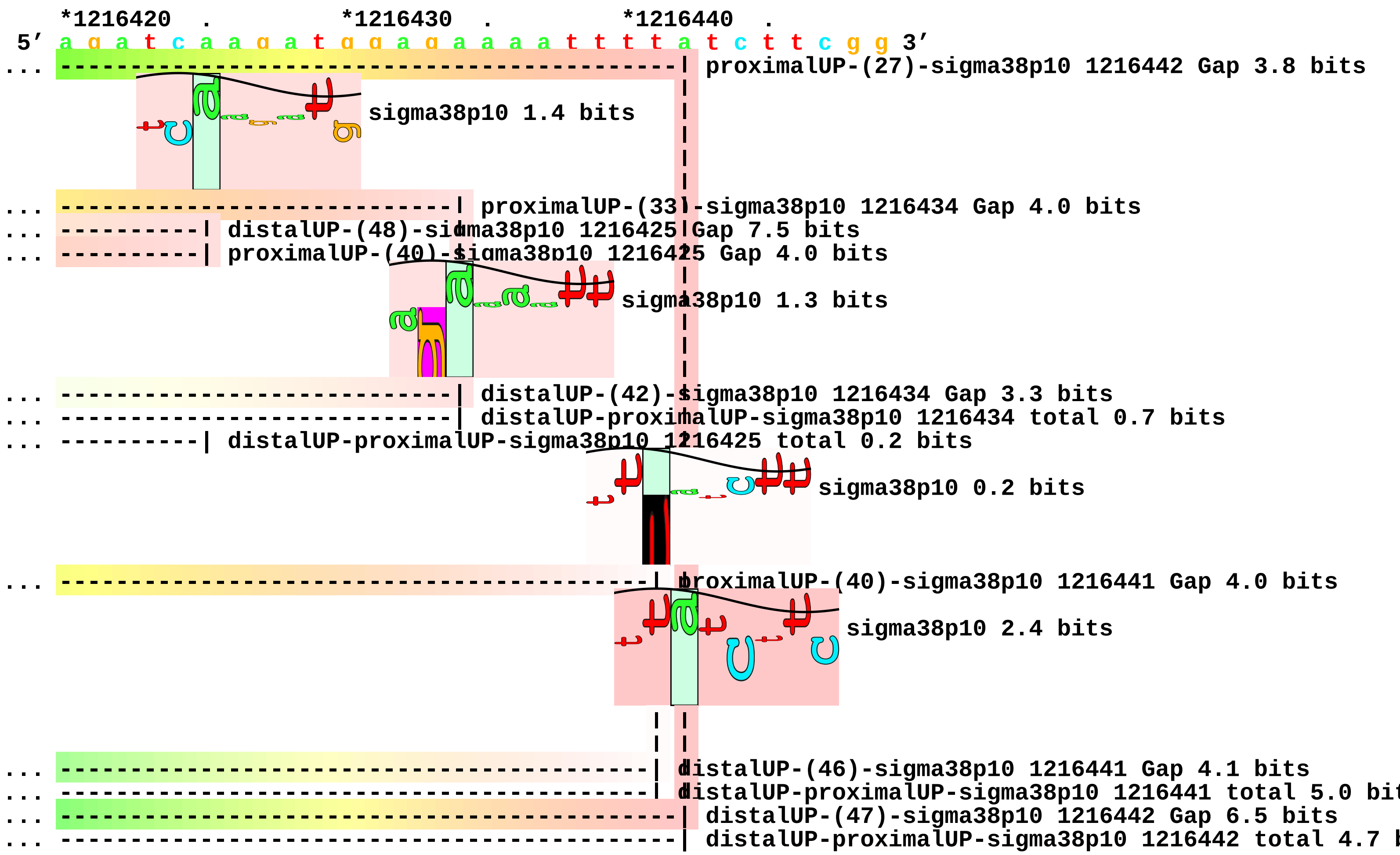
*1216420 . *1216430 . *1216440 .
5′ a g a t c a a g a t g g a g a a a a t t t t a t c t t c g g 3′
proximalUP-(27)-sigma38p10 1216442 Gap 3.8 bits
sigma38p10 1.4 bits
proximalUP-(33)-sigma38p10 1216434 Gap 4.0 bits
distalUP-(48)-sigma38p10 1216425 Gap 7.5 bits
proximalUP-(40)-sigma38p10 1216425 Gap 4.0 bits
sigma38p10 1.3 bits
distalUP-(42)-sigma38p10 1216434 Gap 3.3 bits
distalUP-proximalUP-sigma38p10 1216434 total 0.7 bits
distalUP-proximalUP-sigma38p10 1216425 total 0.2 bits
sigma38p10 0.2 bits
proximalUP-(40)-sigma38p10 1216441 Gap 4.0 bits
sigma38p10 2.4 bits
distalUP-(46)-sigma38p10 1216441 Gap 4.1 bits
distalUP-proximalUP-sigma38p10 1216441 total 5.0 bit
distalUP-(47)-sigma38p10 1216442 Gap 6.5 bits
distalUP-proximalUP-sigma38p10 1216442 total 4.7

piece 12, NC_000913, ycgH, config: linear, direction: +, begin: 1219350, end: 1219999

```
   *1219350  .         *1219360  .         *1219370  .         *1219380  .
5' t t g t t a c c t c t g g g g a a a a t g c a g t a g g t g t t c t t g c a 3'
T timestamp:_2018Sep20_11-34-53

     *1219390  .         *1219400  .         *1219410  .         *1219420  .
5' t g t t c a a g t c c t g g a g a g t c t c g a a c a t g t g t c g a t g c 3'

       *1219430  .         *1219440  .         *1219450  .         *1219460
5' t g t a g a t g a t g a a g t t a g t g a t t c t a a c a g t t a c g a a g 3'
                                          proximalUP 4.5 bits          ... distalUP
                                          |---------------------------- ... proximalUP-(34)
                                          distalUP 1.2 bits          |- ... distalUP-(54)-s
                                          |-------------------------------- ... distalUP-(40)-s
                                          |-----------|-------------------- ... distalUP-proxim
                                                                       |- ... distalUP-proxim

    .         *1219470  .         *1219480  .         *1219490  .         *1219500
5' t t a t t a g c c g t g c t g a t t t a a a a a t g a a t g g t g g t t c c 3'
... distalUP 4.9 bits           sigma38p10 4.8 bits
... -------------------------------------| proximalUP-(34)-sigma38p10 1219483 Gap 2.5 bits
... ---------------------------------------------------------------------- ... distalUP-(54)-s
... -------------------------------------| distalUP-(40)-sigma38p10 1219483 Gap 3.1 bits
... -------------------------------------| distalUP-proximalUP-sigma38p10 1219483 total 5.0 bits
... -------------------------------------------------|------------------------ ... distalUP-proxim
                                          proximalUP 5.6 bits
                                                 |-------------------------- ... proximalUP-(28)
                                                 |-------------------------- ... proximalUP-(38)
                                   distalUP 5.8 bits
                              |--------------------------------------------- ... distalUP-(49)-s
                              |---------------------|------------------------ ... distalUP-proxim

      .         *1219510  .         *1219520  .         *1219530  .
5' a t a a c a a c t a a t g c a t t a a t a g c t a t g g t g c t t a t g c 3'
                                sigma38p10 0.5 bits
                                |
```

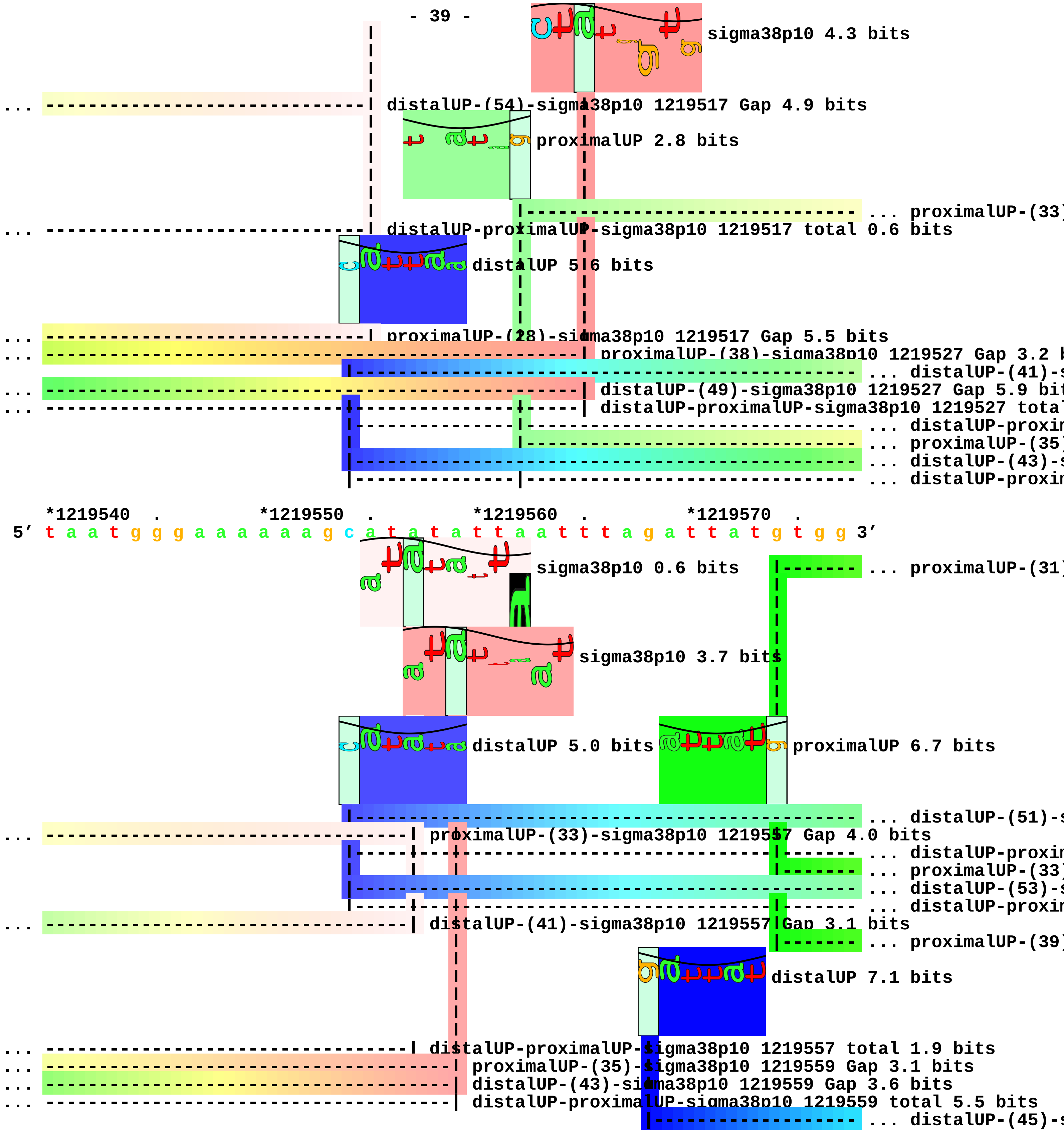
sigma38p10 4.3 bits
... distalUP-(54)-sigma38p10 1219517 Gap 4.9 bits
proximalUP 2.8 bits
... proximalUP-(33)
... distalUP-proximalUP-sigma38p10 1219517 total 0.6 bits
distalUP 5.6 bits
... proximalUP-(28)-sigma38p10 1219517 Gap 5.5 bits
... proximalUP-(38)-sigma38p10 1219527 Gap 3.2 b
... distalUP-(41)-s
... distalUP-(49)-sigma38p10 1219527 Gap 5.9 bit
... distalUP-proximalUP-sigma38p10 1219527 tota
... distalUP-proxim
... proximalUP-(35)
... distalUP-(43)-s
... distalUP-proxim
*1219540 . *1219550 . *1219560 . *1219570 .
5′ t a a t g g g a a a a a a g c a t a t a t t a a t t t a g a t t a t g t g g 3′
sigma38p10 0.6 bits
... proximalUP-(31)
sigma38p10 3.7 bits
distalUP 5.0 bits
proximalUP 6.7 bits
... distalUP-(51)-s
... proximalUP-(33)-sigma38p10 1219557 Gap 4.0 bits
... distalUP-proxim
... proximalUP-(33)
... distalUP-(53)-s
... distalUP-proxim
... distalUP-(41)-sigma38p10 1219557 Gap 3.1 bits
... proximalUP-(39)
distalUP 7.1 bits
... distalUP-proximalUP-sigma38p10 1219557 total 1.9 bits
... proximalUP-(35)-sigma38p10 1219559 Gap 3.1 bits
... distalUP-(43)-sigma38p10 1219559 Gap 3.6 bits
... distalUP-proximalUP-sigma38p10 1219559 total 5.5 bits
... distalUP-(45)-s

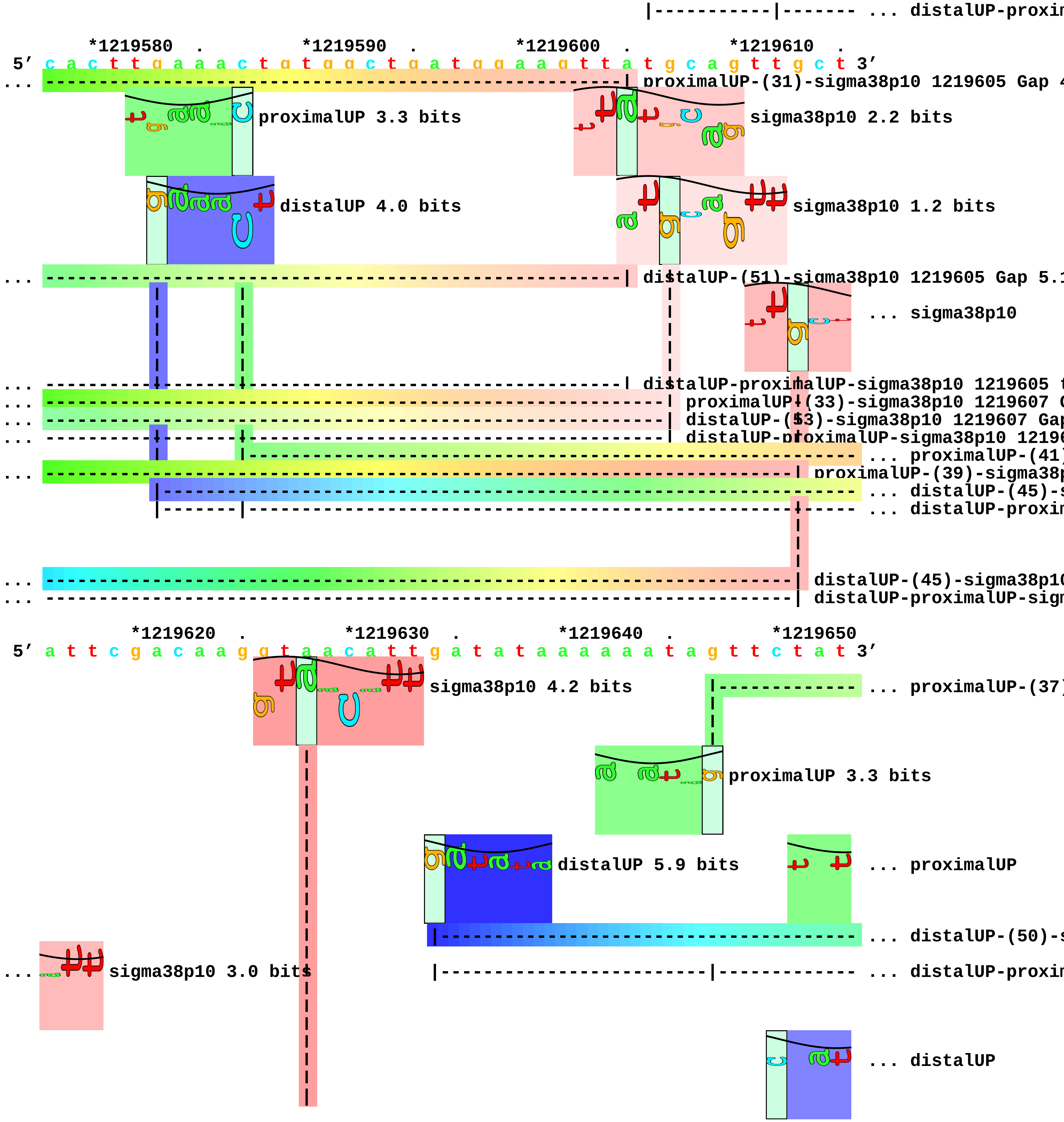
... distalUP-proxi
*1219580 . *1219590 . *1219600 . *1219610 .
5′ c a c t t g a a a c t g t g g c t g a t g g a a g t t a t g c a g t t g c t 3′
... proximalUP-(31)-sigma38p10 1219605 Gap 4
proximalUP 3.3 bits
sigma38p10 2.2 bits
distalUP 4.0 bits
sigma38p10 1.2 bits
... distalUP-(51)-sigma38p10 1219605 Gap 5.1
... sigma38p10
... distalUP-proximalUP-sigma38p10 1219605
... proximalUP-(33)-sigma38p10 1219607 G
... distalUP-(53)-sigma38p10 1219607 Gap
... distalUP-proximalUP-sigma38p10 1219
... proximalUP-(41)
... proximalUP-(39)-sigma38p
... distalUP-(45)-s
... distalUP-proxim
... distalUP-(45)-sigma38p10
... distalUP-proximalUP-sig
*1219620 . *1219630 . *1219640 . *1219650
5′ a t t c g a c a a g g t a a c a t t g a t a t a a a a a a t a g t t c t a t 3′
sigma38p10 4.2 bits
... proximalUP-(37)
proximalUP 3.3 bits
distalUP 5.9 bits
... proximalUP
... distalUP-(50)-s
... sigma38p10 3.0 bits
... distalUP-proxim
... distalUP

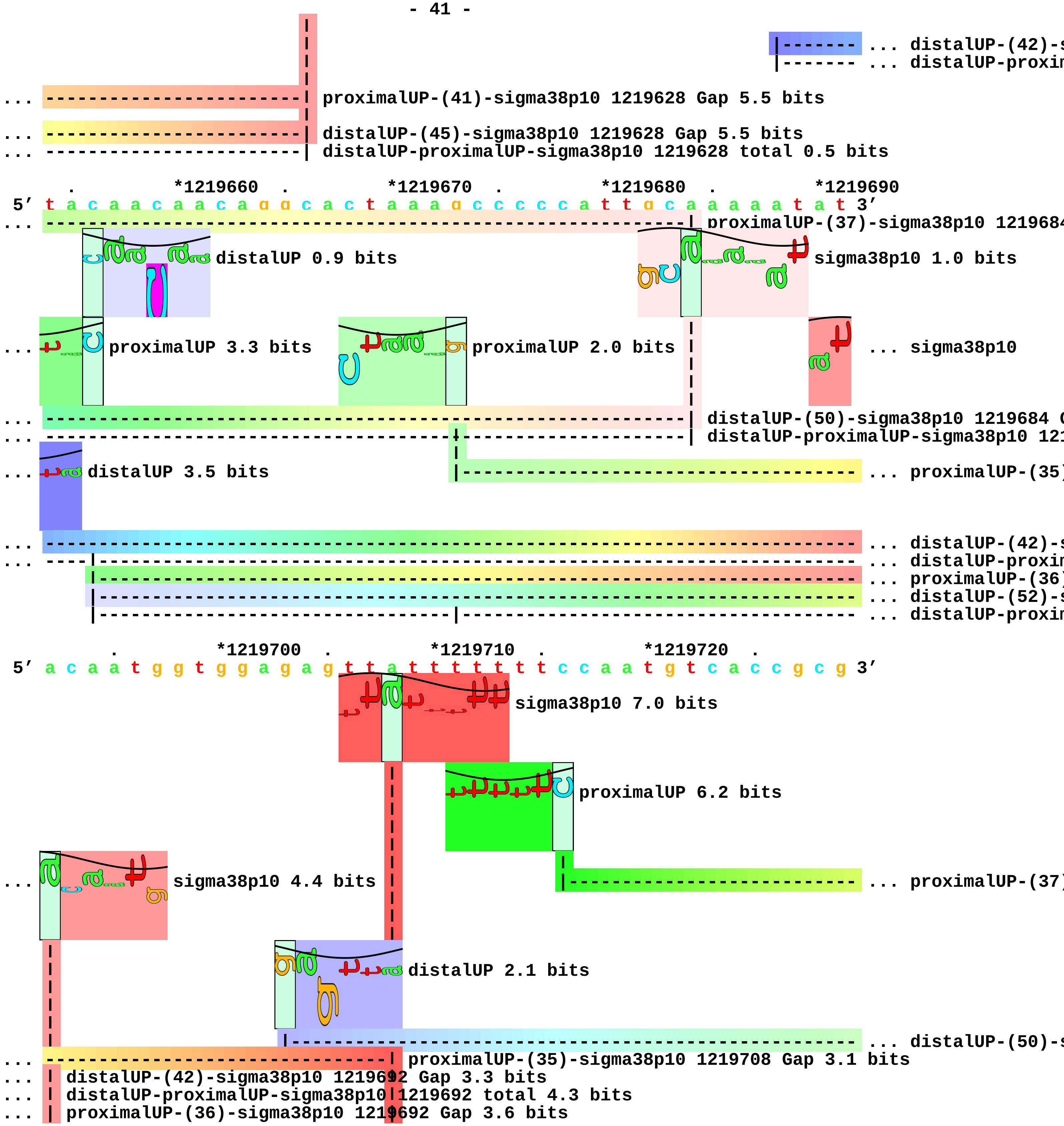
... distalUP-(42)-s
... distalUP-proxim
proximalUP-(41)-sigma38p10 1219628 Gap 5.5 bits
distalUP-(45)-sigma38p10 1219628 Gap 5.5 bits
distalUP-proximalUP-sigma38p10 1219628 total 0.5 bits
*1219660
*1219670
*1219680
*1219690
5′ t a c a a c a a c a g g c a c t a a a g c c c c c a t t g c a a a a a a t a t 3′
proximalUP-(37)-sigma38p10 121968
distalUP 0.9 bits
sigma38p10 1.0 bits
proximalUP 3.3 bits
proximalUP 2.0 bits
... sigma38p10
distalUP-(50)-sigma38p10 1219684 G
distalUP-proximalUP-sigma38p10 121
distalUP 3.5 bits
... proximalUP-(35)
... distalUP-(42)-s
... distalUP-proxim
... proximalUP-(36)
... distalUP-(52)-s
... distalUP-proxim
*1219700
*1219710
*1219720
5′ a c a a t g g t g g a g a g t t a t t t t t t t c c a a t g t c a c c g c g 3′
sigma38p10 7.0 bits
proximalUP 6.2 bits
sigma38p10 4.4 bits
... proximalUP-(37)
distalUP 2.1 bits
... distalUP-(50)-s
proximalUP-(35)-sigma38p10 1219708 Gap 3.1 bits
distalUP-(42)-sigma38p10 1219692 Gap 3.3 bits
distalUP-proximalUP-sigma38p10 1219692 total 4.3 bits
proximalUP-(36)-sigma38p10 1219692 Gap 3.6 bits

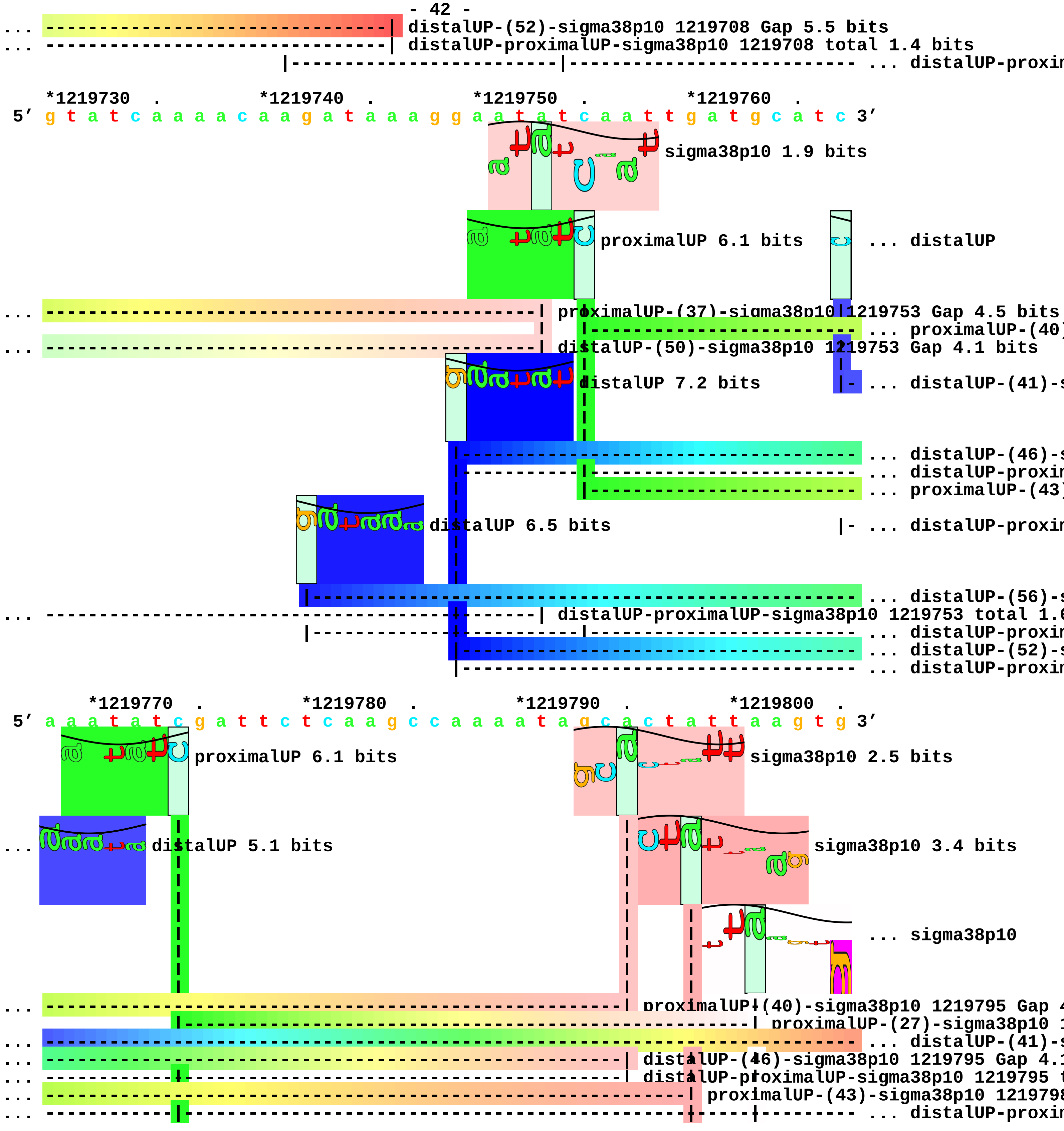

distalUP-(52)-sigma38p10 1219708 Gap 5.5 bits
distalUP-proximalUP-sigma38p10 1219708 total 1.4 bits
... distalUP-proxin
*1219730 . *1219740 . *1219750 . *1219760 .
5′ g t a t c a a a a c a a g a t a a a g g a a t a t c a a t t g a t g c a t c 3′
sigma38p10 1.9 bits
proximalUP 6.1 bits
... distalUP
proximalUP-(37)-sigma38p10 1219753 Gap 4.5 bits
... proximalUP-(40)
distalUP-(50)-sigma38p10 1219753 Gap 4.1 bits
distalUP 7.2 bits
... distalUP-(41)-s
... distalUP-(46)-s
... distalUP-proxin
... proximalUP-(43)
distalUP 6.5 bits
... distalUP-proxin
... distalUP-(56)-s
distalUP-proximalUP-sigma38p10 1219753 total 1.6
... distalUP-proxin
... distalUP-(52)-s
... distalUP-proxin
*1219770 . *1219780 . *1219790 . *1219800 .
5′ a a a t a t c g a t t c t c a a g c c a a a a t a g c a c t a t t a a g t g 3′
proximalUP 6.1 bits
sigma38p10 2.5 bits
distalUP 5.1 bits
sigma38p10 3.4 bits
... sigma38p10
proximalUP-(40)-sigma38p10 1219795 Gap 4
proximalUP-(27)-sigma38p10 1
... distalUP-(41)-s
distalUP-(46)-sigma38p10 1219795 Gap 4.1
distalUP-proximalUP-sigma38p10 1219795 t
proximalUP-(43)-sigma38p10 1219798
... distalUP-proxin

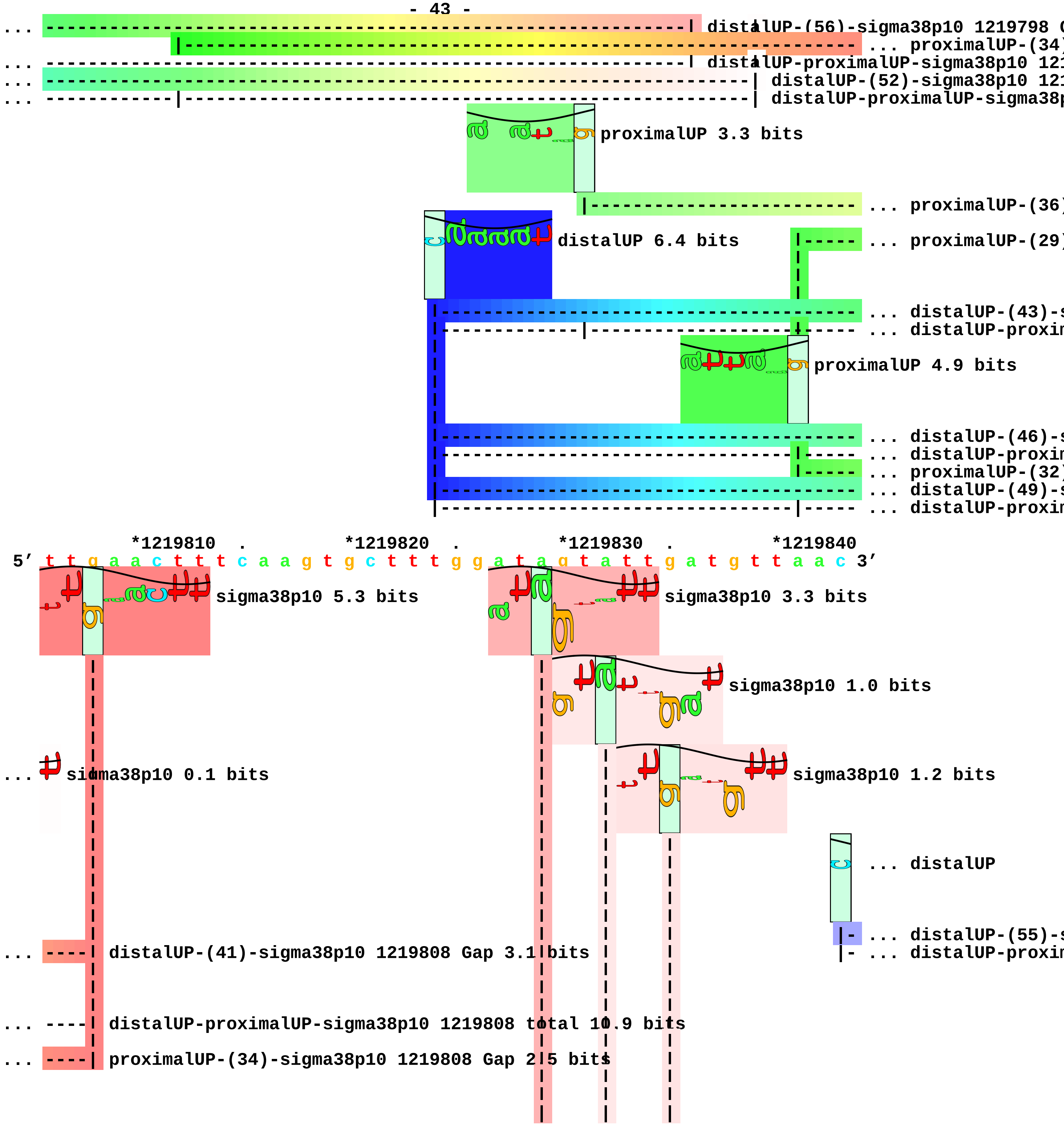
distalUP-(56)-sigma38p10 1219798 (
proximalUP-(34)
distalUP-proximalUP-sigma38p10 121
distalUP-(52)-sigma38p10 121
distalUP-proximalUP-sigma38p
proximalUP 3.3 bits
proximalUP-(36)
distalUP 6.4 bits
proximalUP-(29)
distalUP-(43)-s
distalUP-proxim
proximalUP 4.9 bits
distalUP-(46)-s
distalUP-proxim
proximalUP-(32)
distalUP-(49)-s
distalUP-proxim
*1219810
*1219820
*1219830
*1219840
5′ t t g a a c t t t c a a g t g c t t t g g a t a g t a t t g a t g t t a a c 3′
sigma38p10 5.3 bits
sigma38p10 3.3 bits
sigma38p10 1.0 bits
sigma38p10 0.1 bits
sigma38p10 1.2 bits
distalUP
distalUP-(55)-s
distalUP-proxim
distalUP-(41)-sigma38p10 1219808 Gap 3.1 bits
distalUP-proximalUP-sigma38p10 1219808 total 10.9 bits
proximalUP-(34)-sigma38p10 1219808 Gap 2.5 bits

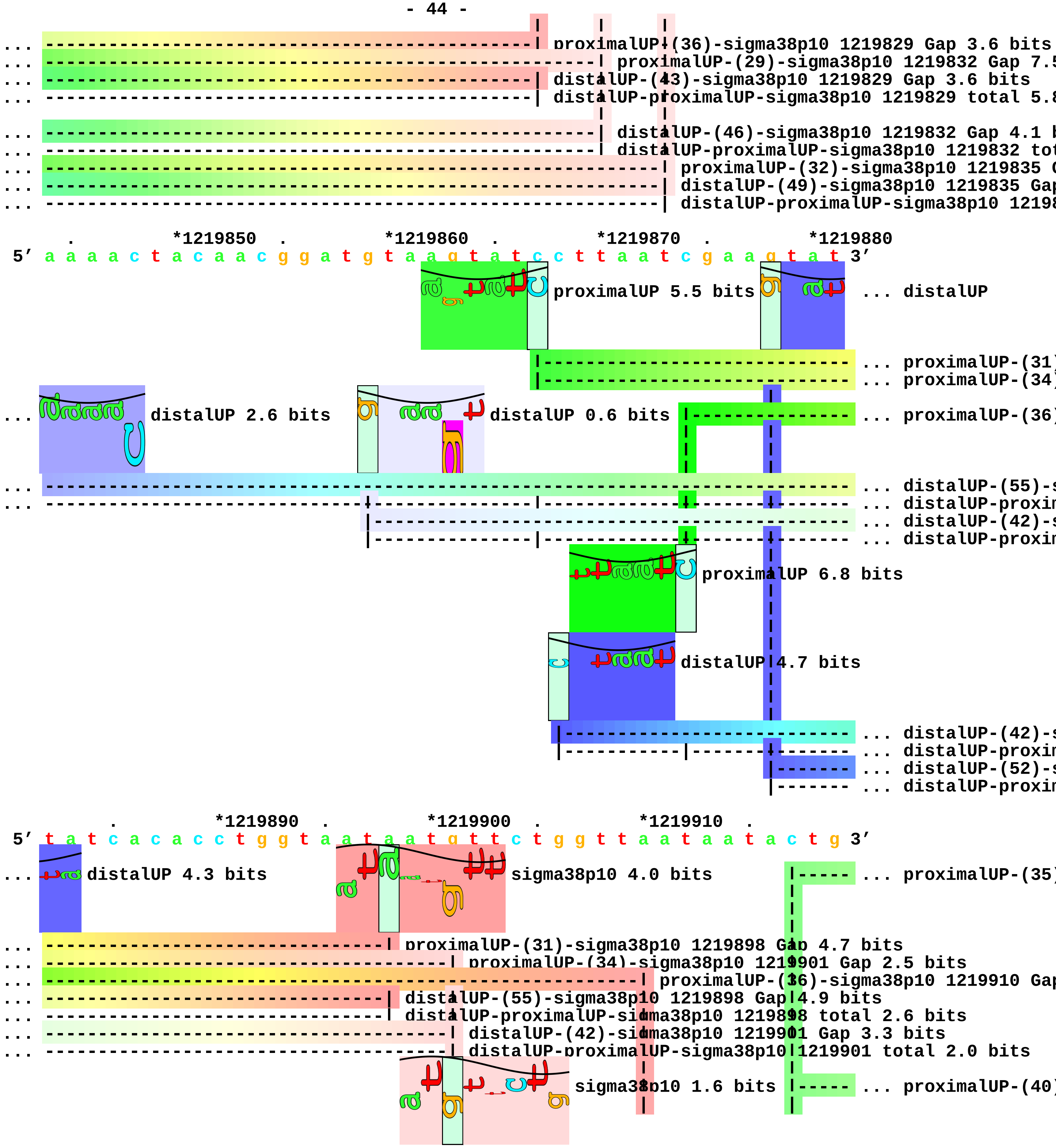
... proximalUP-(36)-sigma38p10 1219829 Gap 3.6 bits
... proximalUP-(29)-sigma38p10 1219832 Gap 7.5
... distalUP-(43)-sigma38p10 1219829 Gap 3.6 bits
... distalUP-proximalUP-sigma38p10 1219829 total 5.8
... distalUP-(46)-sigma38p10 1219832 Gap 4.1 b
... distalUP-proximalUP-sigma38p10 1219832 tot
... proximalUP-(32)-sigma38p10 1219835 G
... distalUP-(49)-sigma38p10 1219835 Gap
... distalUP-proximalUP-sigma38p10 12198
*1219850
*1219860
*1219870
*1219880
5′ a a a a c t a c a a c g g a t g t a a g t a t c c t t a a t c g a a g t a t 3′
proximalUP 5.5 bits
... distalUP
... proximalUP-(31)
... proximalUP-(34)
distalUP 2.6 bits
distalUP 0.6 bits
... proximalUP-(36)
... distalUP-(55)-s
... distalUP-proxim
... distalUP-(42)-s
... distalUP-proxim
proximalUP 6.8 bits
distalUP 4.7 bits
... distalUP-(42)-s
... distalUP-proxim
... distalUP-(52)-s
... distalUP-proxim
*1219890
*1219900
*1219910
5′ t a t c a c a c c t g g t a a t a a t g t t c t g g t t a a t a a t a c t g 3′
... distalUP 4.3 bits
sigma38p10 4.0 bits
... proximalUP-(35)
... proximalUP-(31)-sigma38p10 1219898 Gap 4.7 bits
... proximalUP-(34)-sigma38p10 1219901 Gap 2.5 bits
... proximalUP-(36)-sigma38p10 1219910 Gap
... distalUP-(55)-sigma38p10 1219898 Gap 4.9 bits
... distalUP-proximalUP-sigma38p10 1219898 total 2.6 bits
... distalUP-(42)-sigma38p10 1219901 Gap 3.3 bits
... distalUP-proximalUP-sigma38p10 1219901 total 2.0 bits
sigma38p10 1.6 bits
... proximalUP-(40)

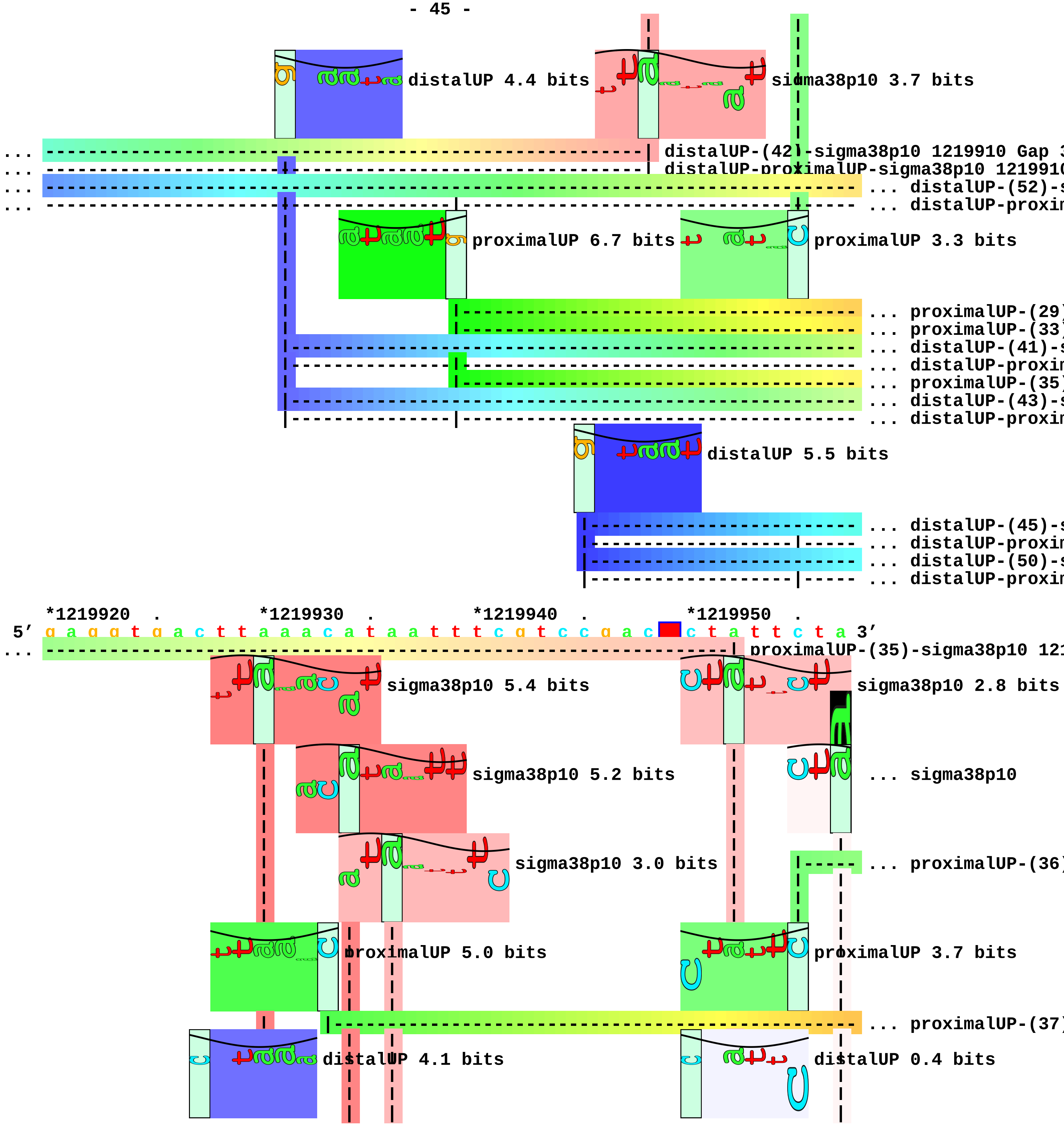
distalUP 4.4 bits
sigma38p10 3.7 bits
distalUP-(42)-sigma38p10 1219910 Gap 3
distalUP-proximalUP-sigma38p10 1219910
... distalUP-(52)-s
... distalUP-proxim
proximalUP 6.7 bits
proximalUP 3.3 bits
... proximalUP-(29)
... proximalUP-(33)
... distalUP-(41)-s
... distalUP-proxim
... proximalUP-(35)
... distalUP-(43)-s
... distalUP-proxim
distalUP 5.5 bits
... distalUP-(45)-s
... distalUP-proxim
... distalUP-(50)-s
... distalUP-proxim
*1219920 . *1219930 . *1219940 . *1219950 .
5′ g a g g t g a c t t a a a c a t a a t t t c g t c c g a c c t a t t c t a 3′
proximalUP-(35)-sigma38p10 121
sigma38p10 5.4 bits
sigma38p10 2.8 bits
sigma38p10 5.2 bits
... sigma38p10
sigma38p10 3.0 bits
... proximalUP-(36)
proximalUP 5.0 bits
proximalUP 3.7 bits
... proximalUP-(37)
distalUP 4.1 bits
distalUP 0.4 bits

```
                 |---------------------------------------------------------- ... distalUP-(43)-s
... ----------------------------------------------------------------------| proximalUP-(40)-sigm
                 |---------|---------------------------------------|-----|- ... distalUP-proxim
                                                                   |-----------  ... distalUP-(41)-s
                                                                   |----|--- ... distalUP-proxim
... --------------------| distalUP-(52)-sigma38p10 1219930 Gap 5.5 bits
... --------------------| distalUP-proximalUP-sigma38p10 1219930 total 3.5 bits

... --------------------| proximalUP-(29)-sigma38p10 1219930 Gap 7.5 bits
... ---------------------------| proximalUP-(33)-sigma38p10 1219934 Gap 4.0 bits
... ---------------------------| distalUP-(41)-sigma38p10 1219934 Gap 3.1 bits
... ---------------------------| distalUP-proximalUP-sigma38p10 1219934 total 9.1 bits
... -------------------------------| proximalUP-(35)-sigma38p10 1219936 Gap 3.1 bits
... -------------------------------| distalUP-(43)-sigma38p10 1219936 Gap 3.6 bits
... -------------------------------| distalUP-proximalUP-sigma38p10 1219936 total 7.4 bits

... ----------------------------------------------------------------| distalUP-(45)-sigma38p10 1219
... ----------------------------------------------------------------| distalUP-proximalUP-sigma38p10
... ----------------------------------------------------------------------| distalUP-(50)-sigma3
... ----------------------------------------------------------------------| distalUP-proximalUP-

     *1219960  .         *1219970  .         *1219980  .         *1219990  .
5' a a t g g a g c g a c t a a a c t c g t c a g c g g c a c a a c c a c g c t 3'
                                     sigma38p10 7.5 bits                      ... sigma38p10

... sigma38p10 0.5 bits

... ------------------------------------------------------------------| proximalUP-(36)-sigma38p10 1
... -------------------------| proximalUP-(37)-sigma38p10 1219970 Gap 4.5 bits
... -------------------------| distalUP-(43)-sigma38p10 1219970 Gap 3.6 bits
... -------------------------| distalUP-proximalUP-sigma38p10 1219970 total 8.5 bits
... ------------------------------------------------------------------| distalUP-(41)-sigma38p10 12
... ------------------------------------------------------------------| distalUP-proximalUP-sigma38p

5' g a a g 3'
... sigma38p10 3.1 bits
```

piece 13, NC_000913, ycgB, config: linear, direction: -, begin: 1236576, end: 1236370

```
     .         *1236570  .         *1236560  .         *1236550  .         *1236540
5' a a t a a g a a a t c g c a a a g a a a a a a t t c t c c g g c g a c g c 3'

   T timestamp:_2018Sep20_11-34-53                       proximalUP 5.5 bits
                                                                                  ... proximalUP-(31)
                                             distalUP 6.5 bits
                                                                                  ... distalUP-(41)-s
                                                                                  ... sigma38p10-prox
                                                                                  ... proximalUP-(37)
                                                                                  ... distalUP-(47)-s
                                                                                  ... sigma38p10-prox

          .         *1236530  .         *1236520  .         *1236510  .
5' g g t t t t t a a c a g g t g t t c t a t g c t t g a a a t g a g g t g c t 3'
                                                       sigma38p10 9.7 bits
...                                            proximalUP-(31)-sigma38p10 1236519 Gap 4.7 bits
                                                                 sigma38p10 2.1 bits
...                                            distalUP-(41)-sigma38p10 1236519 Gap 3.1 bits
...                                            sigma38p10-proximalUP-distalUP 1236519 total 14.0 bits
...                                                        proximalUP-(37)-sigma38p10 1236513 Gap 4.5 b
...                                                        distalUP-(47)-sigma38p10 1236513 Gap 6.5 bit
...                                                        sigma38p10-proximalUP-distalUP 1236513 tota

   *1236500  .         *1236490  .         *1236480  .         *1236470  .
5' c c a c a c a g a g a g c g c c g a c a a c a a t g a g g t g c g c g t a t 3'

       *1236460  .         *1236450  .         *1236440  .         *1236430  .
5' g g c g a c g a t c g a t t c t a t g a a t a a g g a c a c c a c a c g t t 3'

           *1236420  .         *1236410  .         *1236400  .         *1236390
5' t g a g c g a t g g a c c c g a c t g g a c g t t c g a c c t g c t g g a t 3'

     .         *1236380  .         *1236370
5' g t t t a t c t g g c a g a g a t 3'
```

piece 14, NC_000913, osmB, config: linear, direction: -, begin: 1341530, end: 1341254

*1341530 . *1341520 . *1341510 . *1341500 .
5′ a t t a a c a c t c t c a a g a t a t a a a t t a t t a t c a g c g a t a 3′

T timestamp:_2018Sep20_11-34-53

... distalUP
... distalUP-(51)-s
... sigma38p10-prox

*1341490 . *1341480 . *1341470 . *1341460 .
5′ t a a c a g g a a g t c a t t a t c a c c t g c g t g a t a t a a c c c t g 3′

... distalUP 5.9 bits
proximalUP 7.2 bits
proximalUP 3.9 bits
... distalUP-(51)-s
... sigma38p10-prox
... proximalUP-(30)
... proximalUP-(42)
distalUP 6.3 bits
... proximalUP-(28)
... distalUP-(48)-s
... sigma38p10-prox
... distalUP-(50)-s
... sigma38p10-prox
... proximalUP-(37)
distalUP 5.9 bits
... distalUP-(44)-s
... sigma38p10-prox
... proximalUP-(43)
... distalUP-(50)-s
... sigma38p10-prox

*1341450 . *1341440 . *1341430 . *1341420
5′ c g c g c g a g c a g a t t t c a c g g a a t a a t t t c a c c a g a c t t 3′

sigma38p10 0.6 bits
sigma38p10 4.4 bits
... distalUP-(51)-sigma38p10 1341445 Gap 5.1 bits
... sigma38p10
... sigma38p10-proximalUP-distalUP 1341445 total 2.7 bits
... proximalUP-(30)-sigma38p10 1341445 Gap 5.9 bits
... proximalUP

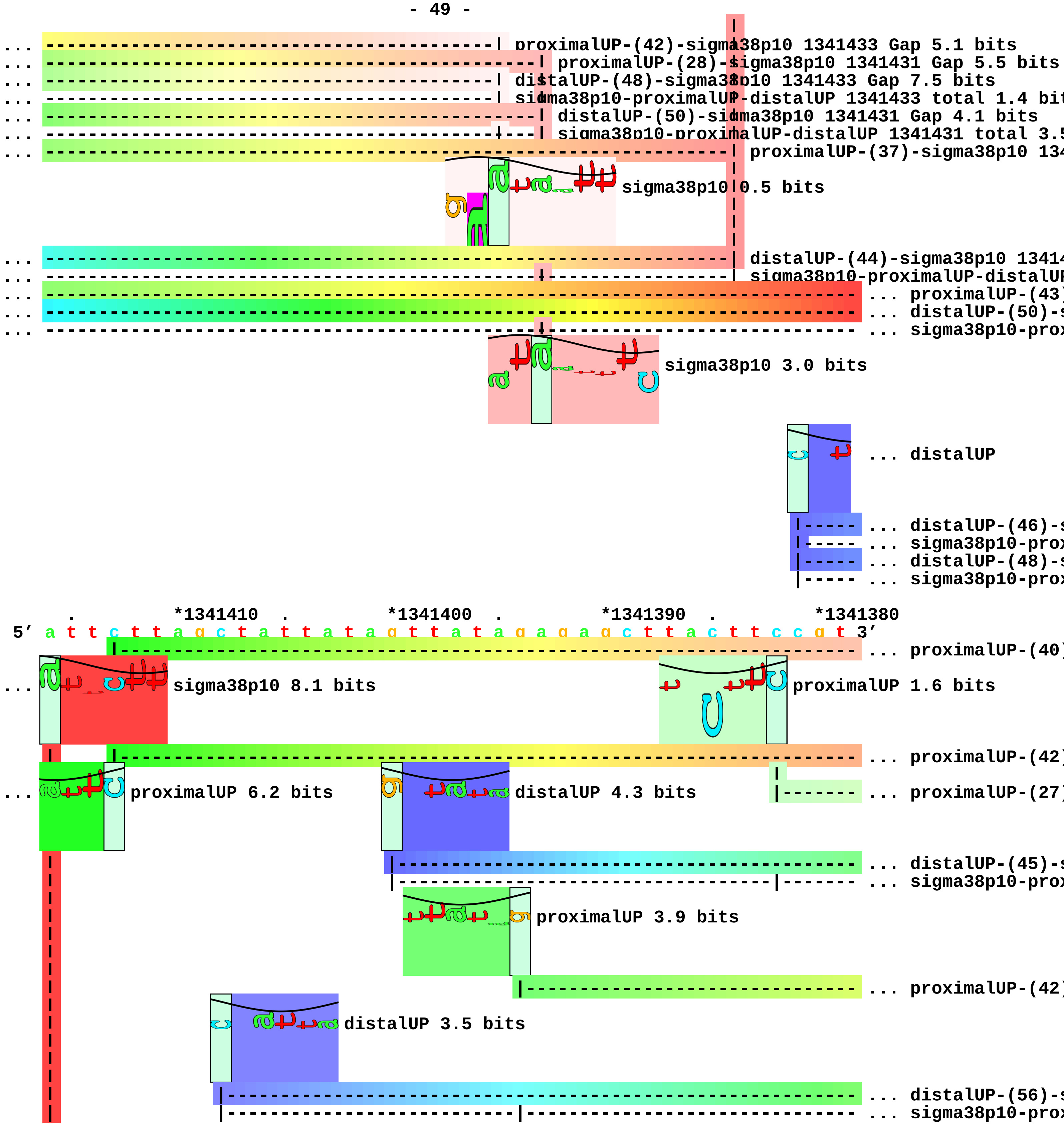
proximalUP-(42)-sigma38p10 1341433 Gap 5.1 bits
proximalUP-(28)-sigma38p10 1341431 Gap 5.5 bits
distalUP-(48)-sigma38p10 1341433 Gap 7.5 bits
sigma38p10-proximalUP-distalUP 1341433 total 1.4 bit
distalUP-(50)-sigma38p10 1341431 Gap 4.1 bits
sigma38p10-proximalUP-distalUP 1341431 total 3.5
proximalUP-(37)-sigma38p10 134
sigma38p10 0.5 bits
distalUP-(44)-sigma38p10 13414
sigma38p10-proximalUP-distalUP
proximalUP-(43)
distalUP-(50)-s
sigma38p10-prox
sigma38p10 3.0 bits
distalUP
distalUP-(46)-s
sigma38p10-prox
distalUP-(48)-s
sigma38p10-prox
*1341410
*1341400
*1341390
*1341380
5′ a t t c t t a g c t a t t a t a g t t a t a g a g a g c t t a c t t c c g t 3′
proximalUP-(40)
sigma38p10 8.1 bits
proximalUP 1.6 bits
proximalUP-(42)
proximalUP 6.2 bits
distalUP 4.3 bits
proximalUP-(27)
distalUP-(45)-s
sigma38p10-prox
proximalUP 3.9 bits
proximalUP-(42)
distalUP 3.5 bits
distalUP-(56)-s
sigma38p10-prox

```
... | proximalUP-(43)-sigma38p10 1341416 Gap 5.1 bits
... | distalUP-(50)-sigma38p10 1341416 Gap 4.1 bits
... | sigma38p10-proximalUP-distalUP 1341416 total 8.5 bits

... atat distalUP 4.1 bits

... ---------------------------------------------------------------------------- ... distalUP-(46)-s
... ------|--------------------------------------------------------------------- ... sigma38p10-prox
... ---------------------------------------------------------------------------- ... distalUP-(48)-s
... ------|--------------------------------------------------------------------- ... sigma38p10-prox

            .         *1341370  .         *1341360  .         *1341350  .
 5' g a a t c a t a a a t t c a g g a g a g a g t a t t a t g t t t g t a a c g 3'
... ----------| proximalUP-(40)-sigma38p10 1341373 Gap 4.0 bits
                sigma38p10 2.6 bits                     sigma38p10 4.0 bits
... --------------| proximalUP-(42)-sigma38p10 1341371 Gap 5.1 bits
... ----------------------------------------------| proximalUP-(27)-sigma38p10 1341355 Gap 3.8 bits
... ----------------------------------------------| distalUP-(45)-sigma38p10 1341355 Gap 5.5 bits
... ----------------------------------------------| sigma38p10-proximalUP-distalUP 1341355 total 0.
                    sigma38p10 4.1 bits                      sigma38p10 6.9 bits
... ----------------------------------------------------------| proximalUP-(42)-sigma38p10 1341352 Gap 5.
                    distalUP 5.6 bits                        proximalUP 6.7 bits
... ----------------------------------------------------------| distalUP-(56)-sigma38p10 1341352 Gap 3.8
... ----------------------------------------------------------| sigma38p10-proximalUP-distalUP 1341352 tot
                                                               |-------------------- ... proximalUP-(31)
        |------------------------------------------------------------------------- ... distalUP-(55)-s
        |------------------------------------------------------|-------------------- ... sigma38p10-prox
                                                               |-------------------- ... proximalUP-(43)
                                                   distalUP 4.3 bits
                                                 |-------------------------------- ... distalUP-(50)-s
                                                 |-------------|------------------ ... sigma38p10-prox
... ----------| distalUP-(46)-sigma38p10 1341373 Gap 4.1 bits
... ----------| sigma38p10-proximalUP-distalUP 1341373 total 4.7 bits
... --------------| distalUP-(48)-sigma38p10 1341371 Gap 7.5 bits
... --------------| sigma38p10-proximalUP-distalUP 1341371 total 1.8 bits

   *1341340  .         *1341330  .         *1341320  .         *1341310  .
 5' a g c a a a a a a a t g a c c g c g g c t g t t c t g g c a a t t a c t t t 3'
```

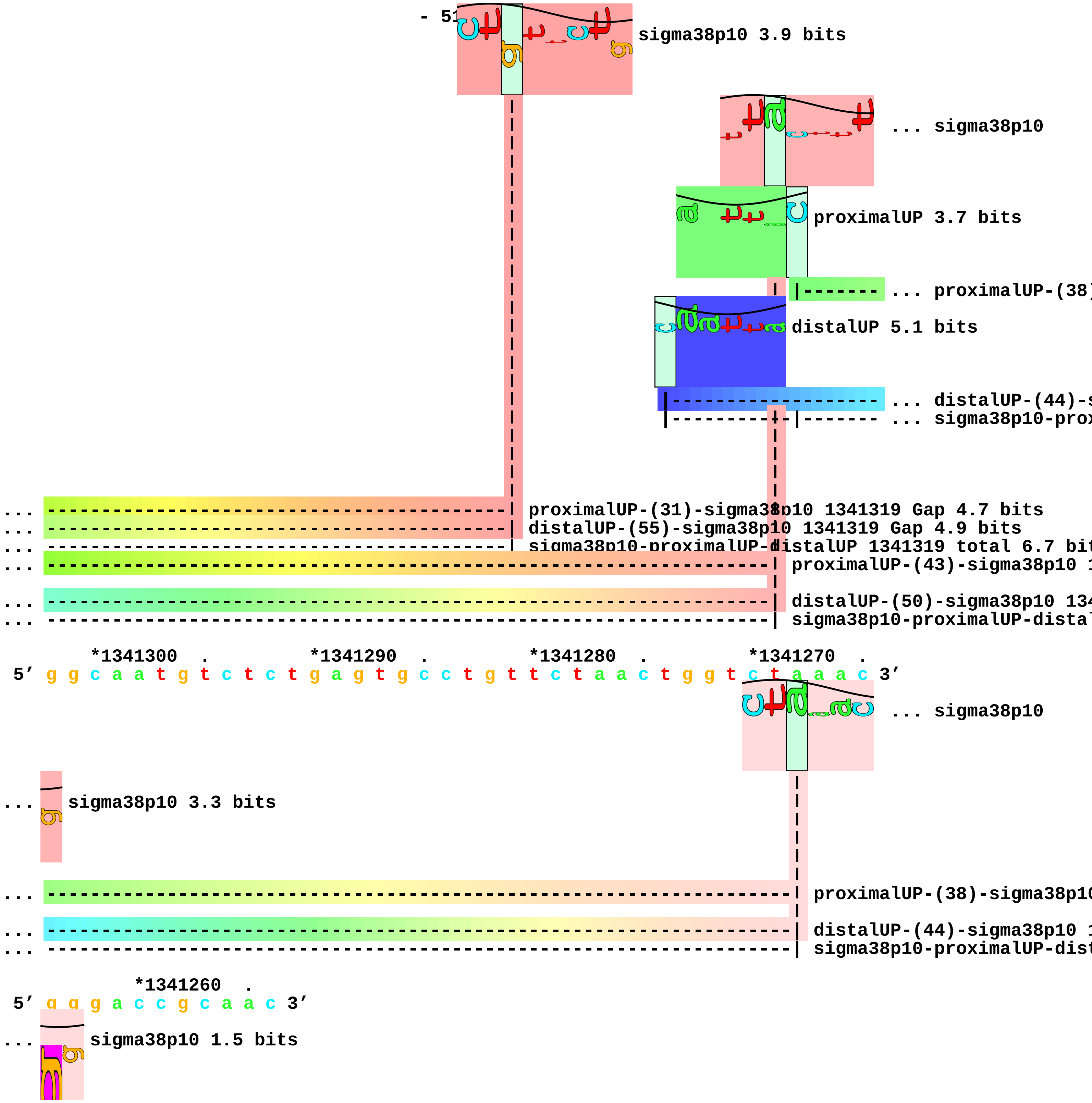

- 51
sigma38p10 3.9 bits
... sigma38p10
proximalUP 3.7 bits
... proximalUP-(38)
distalUP 5.1 bits
... distalUP-(44)-s
... sigma38p10-prox
... proximalUP-(31)-sigma38p10 1341319 Gap 4.7 bits
... distalUP-(55)-sigma38p10 1341319 Gap 4.9 bits
... sigma38p10-proximalUP-distalUP 1341319 total 6.7 bit
... proximalUP-(43)-sigma38p10 1
... distalUP-(50)-sigma38p10 134
... sigma38p10-proximalUP-distal
*1341300 . *1341290 . *1341280 . *1341270 .
5′ g g c a a t g t c t c t g a g t g c c t g t t c t a a c t g g t c t a a a c 3′
... sigma38p10
... sigma38p10 3.3 bits
... proximalUP-(38)-sigma38p10
... distalUP-(44)-sigma38p10 1
... sigma38p10-proximalUP-dist
*1341260 .
5′ g g g a c c g c a a c 3′
... sigma38p10 1.5 bits

piece 15, NC_000913, tfaR, config: linear, direction: +, begin: 1430200, end: 1430599

```
   *1430200  .         *1430210  .         *1430220  .         *1430230  .
5' g a t t g t c t g t t g t g c a t a a t c a a a a c t a t g c a a c a t c a 3'
   T timestamp:_2018Sep20_11-34-53

       *1430240  .         *1430250  .         *1430260  .         *1430270  .
5' t c t g c t g g c g c a c a t a c c c a c t c a c t g t c c g g c a c t g c 3'

           *1430280  .         *1430290  .         *1430300  .         *1430310
5' t g c a a g c g c a g g t g c a c a c g c g c a t a c t g t c g g t a t t g 3'

   .         *1430320  .         *1430330  .         *1430340  .         *1430350
5' g t g c t c a t a c g c a c t c c g t t g c g a t t g g t t c a c a t g g a 3'

       .         *1430360  .         *1430370  .         *1430380  .
5' c a c a c c a t c a c c g t t a a c g c t g c t g g t a a c g c g g a a a a 3'

   *1430390  .         *1430400  .         *1430410  .         *1430420  .
5' c a c c g t c a a a a a c a t c g c a t t t a a c t a t a t t g t g a g g c 3'
                                                 distalUP 4.8 bits
                                                  |--------------------------- ... distalUP-(56)-s
                                                  |--------------------------- ... distalUP-proxin

       *1430430  .         *1430440  .         *1430450  .         *1430460  .
5' t t g c a t a a t g g c a t t c a g a a t g a g t g a a c a a c c a c g g a 3'
                   proximalUP 6.7 bits
... ------------------------------------------------------------------------- ... distalUP-(56)-s
... ------------------|------------------------------------------------------ ... distalUP-proxin
                      |------------------------------------------------------ ... proximalUP-(33)
                      |------------------------------------------------------ ... proximalUP-(34)
                  distalUP 6.3 bits
      |---------------------------------------------------------------------- ... distalUP-(40)-s
      |-----------|---------------------------------------------------------- ... distalUP-proxin
                  |---------------------------------------------------------- ... proximalUP-(41)
      |---------------------------------------------------------------------- ... distalUP-(47)-s
      |-----------|---------------------------------------------------------- ... distalUP-proxin
                  |---------------------------------------------------------- ... proximalUP-(43)
      |---------------------------------------------------------------------- ... distalUP-(49)-s
      |-----------|---------------------------------------------------------- ... distalUP-proxin

           *1430470  .         *1430480  .         *1430490  .         *1430500
5' c c a t a a a a a t t t a t a a t c t g c t g g c c g g a a c t a a t g a a 3'
                   sigma38p10 4.3 bits
... --------| distalUP-(56)-sigma38p10 1430470 Gap 3.8 bits
```

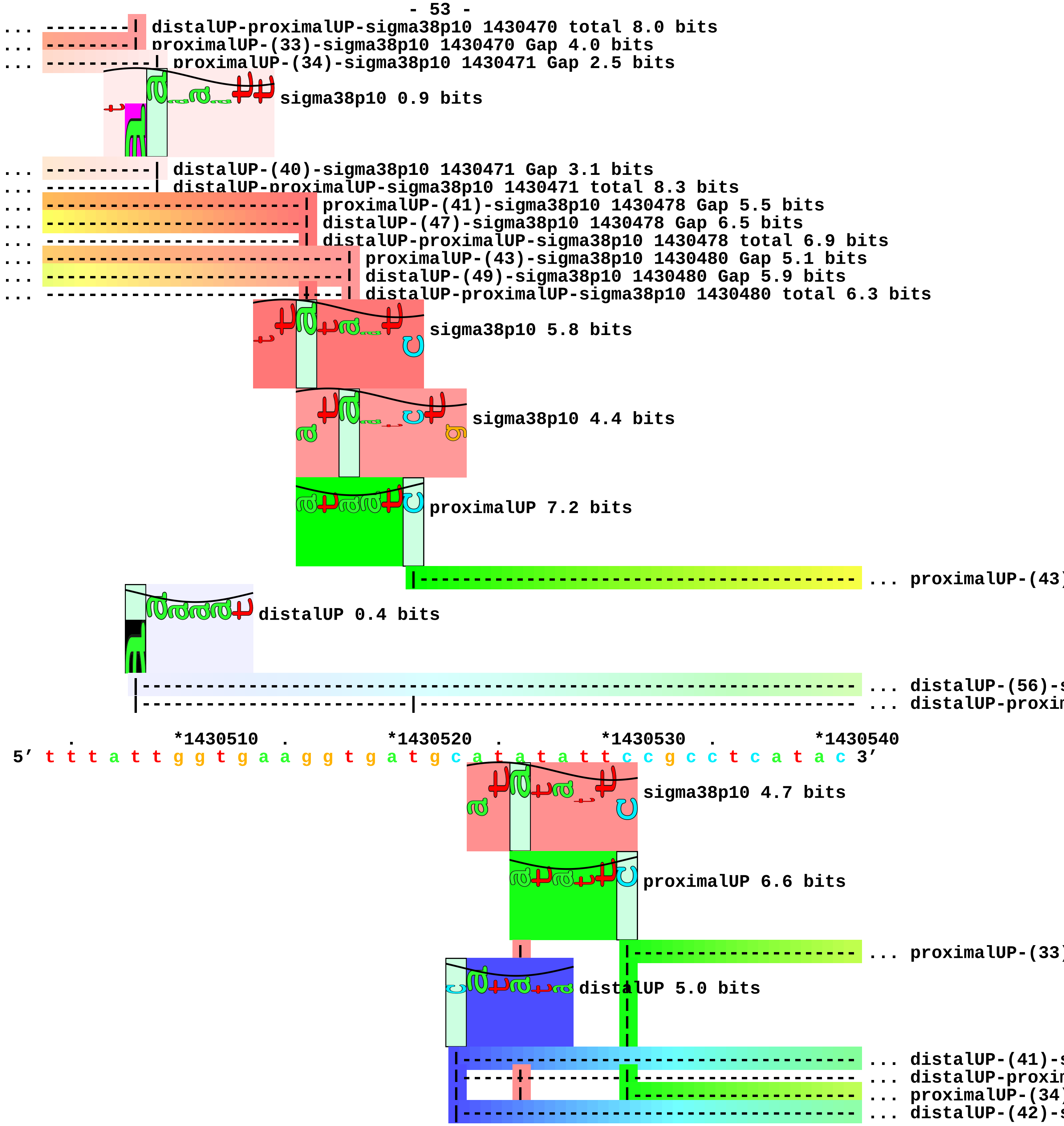
distalUP-proximalUP-sigma38p10 1430470 total 8.0 bits
proximalUP-(33)-sigma38p10 1430470 Gap 4.0 bits
proximalUP-(34)-sigma38p10 1430471 Gap 2.5 bits
sigma38p10 0.9 bits
distalUP-(40)-sigma38p10 1430471 Gap 3.1 bits
distalUP-proximalUP-sigma38p10 1430471 total 8.3 bits
proximalUP-(41)-sigma38p10 1430478 Gap 5.5 bits
distalUP-(47)-sigma38p10 1430478 Gap 6.5 bits
distalUP-proximalUP-sigma38p10 1430478 total 6.9 bits
proximalUP-(43)-sigma38p10 1430480 Gap 5.1 bits
distalUP-(49)-sigma38p10 1430480 Gap 5.9 bits
distalUP-proximalUP-sigma38p10 1430480 total 6.3 bits
sigma38p10 5.8 bits
sigma38p10 4.4 bits
proximalUP 7.2 bits
... proximalUP-(43)
distalUP 0.4 bits
... distalUP-(56)-
... distalUP-proxi
*1430510
*1430520
*1430530
*1430540
5′ t t t a t t g g t g a a g g t g a t g c a t a t a t t c c g c c t c a t a c 3′
sigma38p10 4.7 bits
proximalUP 6.6 bits
... proximalUP-(33)
distalUP 5.0 bits
... distalUP-(41)-
... distalUP-proxi
... proximalUP-(34)
... distalUP-(42)-

```
|-----+---------|-------------------- ... distalUP-proxin

... -------------------------------------------| proximalUP-(43)-sigma38p10 1430526 Gap 5.1 bits
... -------------------------------------------| distalUP-(56)-sigma38p10 1430526 Gap 3.8 bits
... -------------------------------------------| distalUP-proximalUP-sigma38p10 1430526 total 3.5 b

             .       *1430550   .         *1430560  .         *1430570  .
 5' a g g t c t g c c a g c a a a c a g t a c c g a t a t t g c a c c g c c a g 3'
```

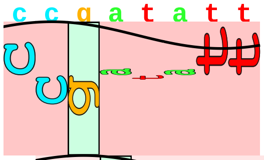

sigma38p10 2.4 bits

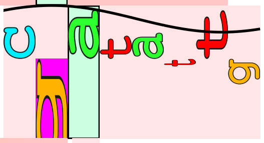

sigma38p10 1.1 bits

```
... -------------------------------------------| proximalUP-(33)-sigma38p10 1430564 Gap 4.0 bits
... -------------------------------------------| distalUP-(41)-sigma38p10 1430564 Gap 3.1 bits
... -------------------------------------------| distalUP-proximalUP-sigma38p10 1430564 total 6.9 b
... ---------------------------------------------| proximalUP-(34)-sigma38p10 1430565 Gap 2.5 bits
... ---------------------------------------------| distalUP-(42)-sigma38p10 1430565 Gap 3.3 bits
... ---------------------------------------------| distalUP-proximalUP-sigma38p10 1430565 total 7.0

    *1430580  .         *1430590  .
 5' a t a t t c c g g c t g g c t t c g t g 3'
```

piece 16, NC_000913, ydcS, config: linear, direction: +, begin: 1509476, end: 1509747

```
              *1509480  .         *1509490  .         *1509500  .         *1509510
5' t c g a a t a g c a g t c a t a t t t c c t a a c t c c t t g a c t a t a c 3'
 T timestamp:_2018Sep20_11-34-53

     .         *1509520  .         *1509530  .         *1509540  .         *1509550
5' t c c a g a a g a t a a c c t t a c a g a c g g c a t a a t g c g c g g t a 3'
                                  proximalUP 0.2 bits
                                 |------------------------------------------------ ... proximalUP-(40)
                                  distalUP 3.3 bits
                    |------------------------------------------------------------- ... distalUP-(46)-s
                    |------------|------------------------------------------------ ... distalUP-proxi

         .         *1509560  .         *1509570  .         *1509580  .
5' g c t c a c a a c c t g a a t a a a t t t t c t c a g g g g c g a a g g t g 3'
                                  sigma38p10 7.0 bits
... ------------------------------| proximalUP-(40)-sigma38p10 1509567 Gap 4.0 bits
                                       proximalUP 6.6 bits
... ------------------------------| distalUP-(46)-sigma38p10 1509567 Gap 4.1 bits
... ------------------------------| distalUP-proximalUP-sigma38p10 1509567 total 2.3 bits
                                          |-------------------------------- ... proximalUP-(35)
                                  distalUP 6.5 bits
                            |----------------------------------------------------- ... distalUP-(46)-s
                            |---------------------|------------------------------- ... distalUP-proxi

   *1509590  .         *1509600  .         *1509610  .         *1509620  .
5' t g c c t g c a a g c c g c c g t c t a t g g t t a a a c a a g g a g a t a 3'
                                        sigma38p10 6.8 bits
                                                   proximalUP 5.0 bits
                                                              |------------------- ... proximalUP-(37)
                                                   distalUP 4.9 bits
```

```
                                          |------------------------------- ... distalUP-(43)-s
... --------------------------------------| proximalUP-(35)-sigma38p10 1509609 Gap 3.1 bits
                                          |-----------|------------------- ... distalUP-proxim
... --------------------------------------| distalUP-(46)-sigma38p10 1509609 Gap 4.1 bits
... --------------------------------------| distalUP-proximalUP-sigma38p10 1509609 total 12.8 bits

       *1509630   .         *1509640   .         *1509650   .         *1509660   .
5' t t t t t a c g g c a c g g c g g c t g a a c a a t t a a t t a c g a c a g 3'
```

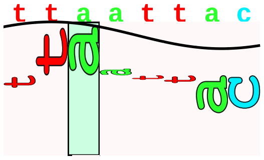

sigma38p10 0.3 bits

```
... ------------------------------------------------------| proximalUP-(37)-sigma38p10 1509655 Gap 4
... ------------------------------------------------------| distalUP-(43)-sigma38p10 1509655 Gap 3.6
... ------------------------------------------------------| distalUP-proximalUP-sigma38p10 1509655 t

         *1509670   .         *1509680   .         *1509690   .         *1509700
5' g a g t a a g a c c t t a t g a g c a a g a c a t t t g c c c g c a g c a g 3'

     .         *1509710   .         *1509720   .         *1509730   .         *1509740
5' c c t g t g t g c g c t c a g c a t g a c a a t a a t g a c c g c t c a c g 3'

       .
5' c c g c c g 3'
```

piece 17, NC_000913, ansP, config: linear, direction: -, begin: 1524249, end: 1523950

```
          .         *1524240  .         *1524230  .         *1524220  .
5′ a g a t t a a g t g t a g g a c a g t a c a c a g g t t a a c c a g c a a a 3′
```

proximalUP 4.9 bits
... proximalUP-(38)
distalUP 6.4 bits
... distalUP-(44)-s
... sigma38p10-prox
... proximalUP-(41)
... distalUP-(47)-s
... sigma38p10-prox

T timestamp:_2018Sep20_11-34-53

```
      *1524210  .         *1524200  .         *1524190  .         *1524180  .
5′ a g g a t t t t g t c a t t t t t t c g t c a g t t g t a t t a a t c c c c t 3′
```

sigma38p10 2.4 bits
proximalUP 6.8 bits
... proximalUP-(38)-sigma38p10 1524204 Gap 3.2 bits
... proximalUP-(37)
sigma38p10 4.1 bits
... distalUP-(44)-sigma38p10 1524204 Gap 2.8 bits
... sigma38p10-proximalUP-distalUP 1524204 total 7.7 bits
... proximalUP-(41)-sigma38p10 1524201 Gap 5.5 bits
... distalUP-(47)-sigma38p10 1524201 Gap 6.5 bits
... sigma38p10-proximalUP-distalUP 1524201 total 3.6 bits
distalUP 4.3 bits
... distalUP-(45)-s
... sigma38p10-prox

```
        *1524170  .         *1524160  .         *1524150  .         *1524140
5′ t c a g g a t t g t a a g g g g a t g t t t t g c a t c g t t t a t c g a t 3′
```

distalUP 0.1 bits
sigma38p10 0.2 bits
... proximalUP-(37)-sigma38p10 152
proximalUP 6.8 bits

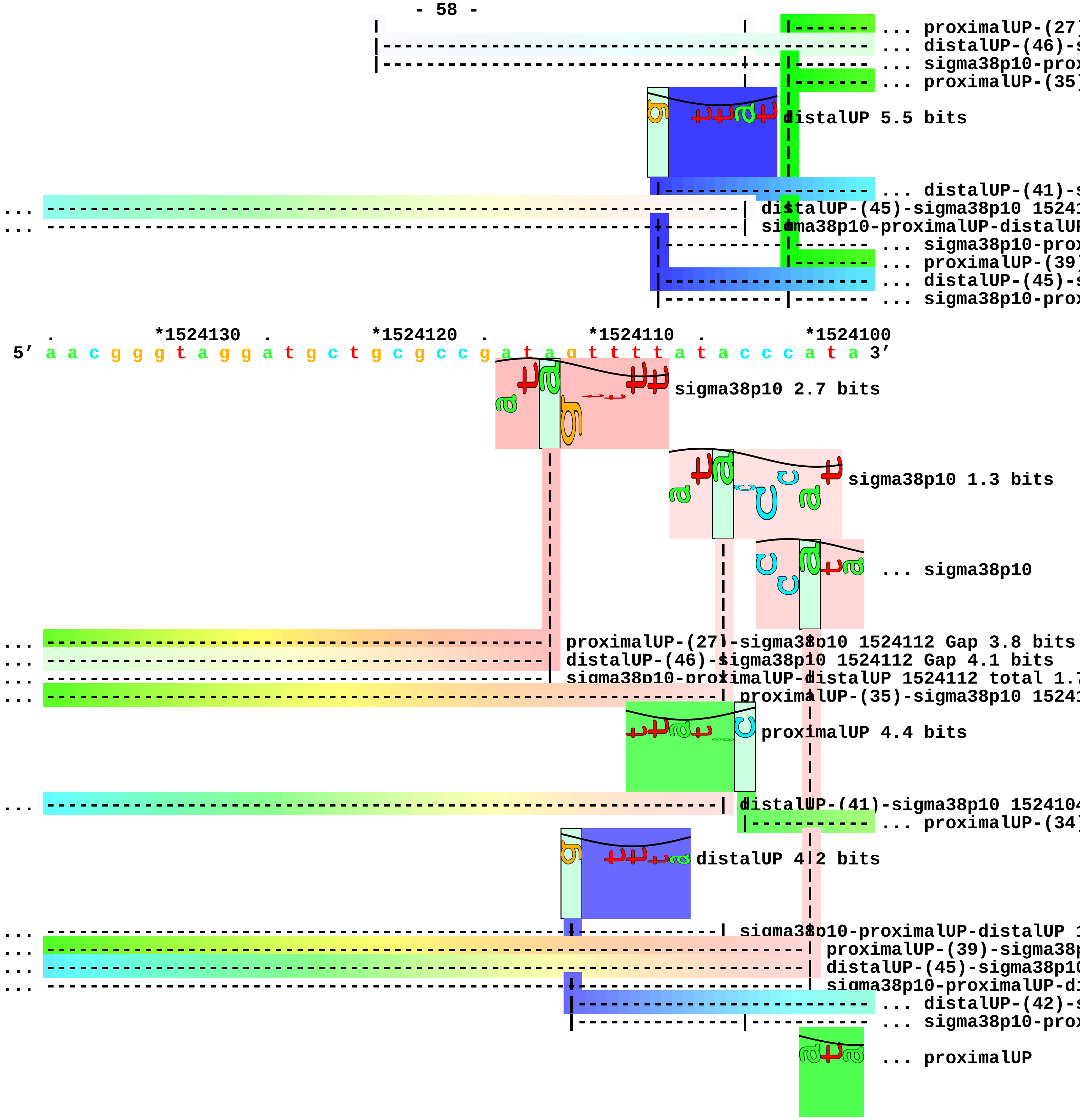

... proximalUP-(27)
... distalUP-(46)-s
... sigma38p10-prox
... proximalUP-(35)
distalUP 5.5 bits
... distalUP-(41)-s
... distalUP-(45)-sigma38p10 15241
... sigma38p10-proximalUP-distalUP
... sigma38p10-prox
... proximalUP-(39)
... distalUP-(45)-s
... sigma38p10-prox
. *1524130 . *1524120 . *1524110 . *1524100
5′ a a c g g g t a g g a t g c t g c g c c g a t a g t t t t a t a c c c a t a 3′
sigma38p10 2.7 bits
sigma38p10 1.3 bits
... sigma38p10
... proximalUP-(27)-sigma38p10 1524112 Gap 3.8 bits
... distalUP-(46)-sigma38p10 1524112 Gap 4.1 bits
... sigma38p10-proximalUP-distalUP 1524112 total 1.7
... proximalUP-(35)-sigma38p10 15241
proximalUP 4.4 bits
... distalUP-(41)-sigma38p10 152410
... proximalUP-(34)
distalUP 4.2 bits
... sigma38p10-proximalUP-distalUP 1
... proximalUP-(39)-sigma38p
... distalUP-(45)-sigma38p10
... sigma38p10-proximalUP-di
... distalUP-(42)-s
... sigma38p10-prox
... proximalUP

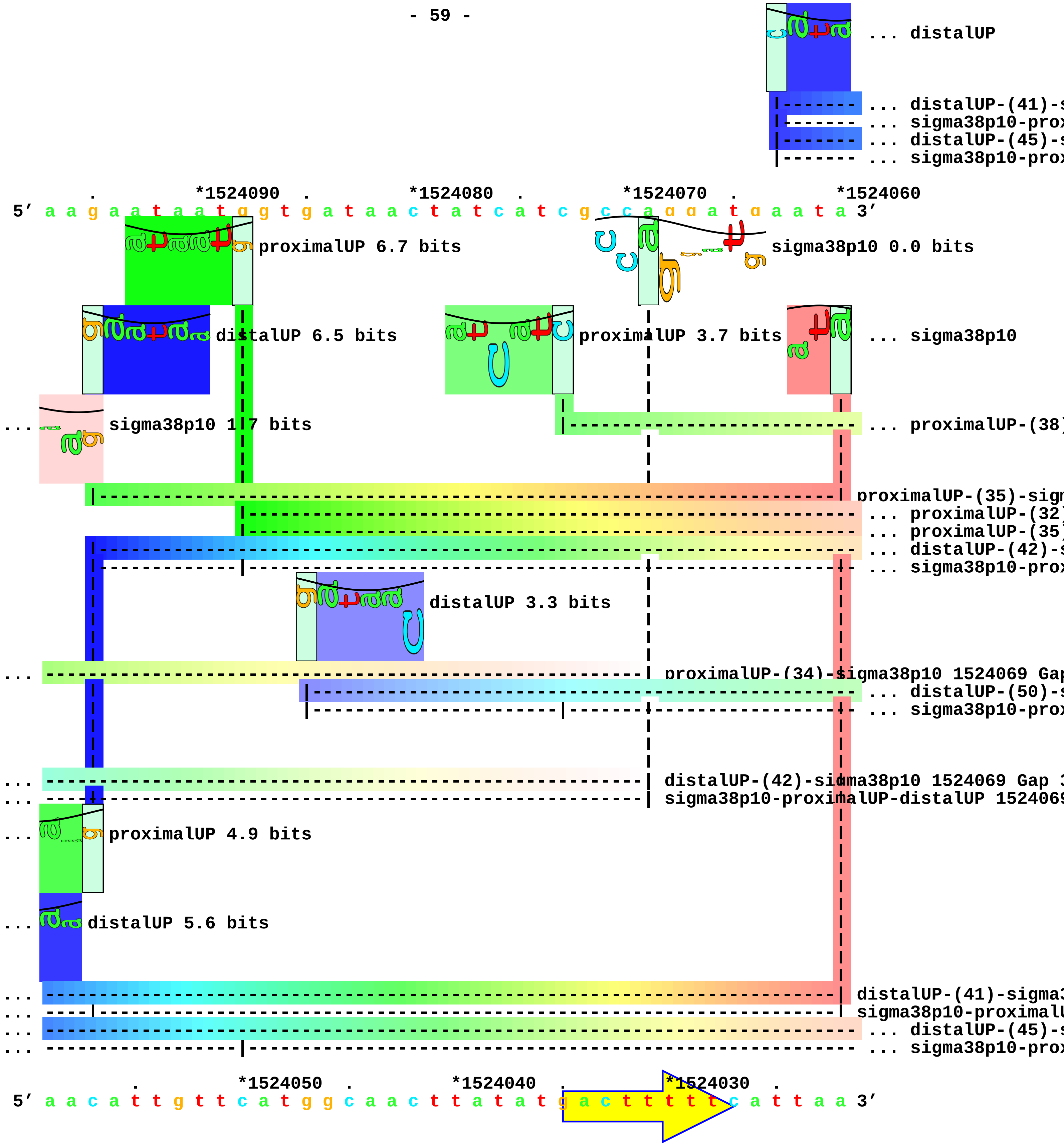
... distalUP
... distalUP-(41)-s
... sigma38p10-prox
... distalUP-(45)-s
... sigma38p10-prox
. *1524090 . *1524080 . *1524070 . *1524060
5' a a g a a t a a t g g t g a t a a c t a t c a t c g c c a g g a t g a a t a 3'
proximalUP 6.7 bits
sigma38p10 0.0 bits
distalUP 6.5 bits
proximalUP 3.7 bits
... sigma38p10
sigma38p10 1.7 bits
... proximalUP-(38)
proximalUP-(35)-sign
... proximalUP-(32)
... proximalUP-(35)
... distalUP-(42)-s
... sigma38p10-prox
distalUP 3.3 bits
proximalUP-(34)-sigma38p10 1524069 Gap
... distalUP-(50)-s
... sigma38p10-prox
distalUP-(42)-sigma38p10 1524069 Gap 3
sigma38p10-proximalUP-distalUP 152406
proximalUP 4.9 bits
distalUP 5.6 bits
distalUP-(41)-sigma3
sigma38p10-proximalU
... distalUP-(45)-s
... sigma38p10-prox
. *1524050 . *1524040 . *1524030 .
5' a a c a t t g t t c a t g g c a a c t t a t a t g a c t t t t t c a t t a a 3'

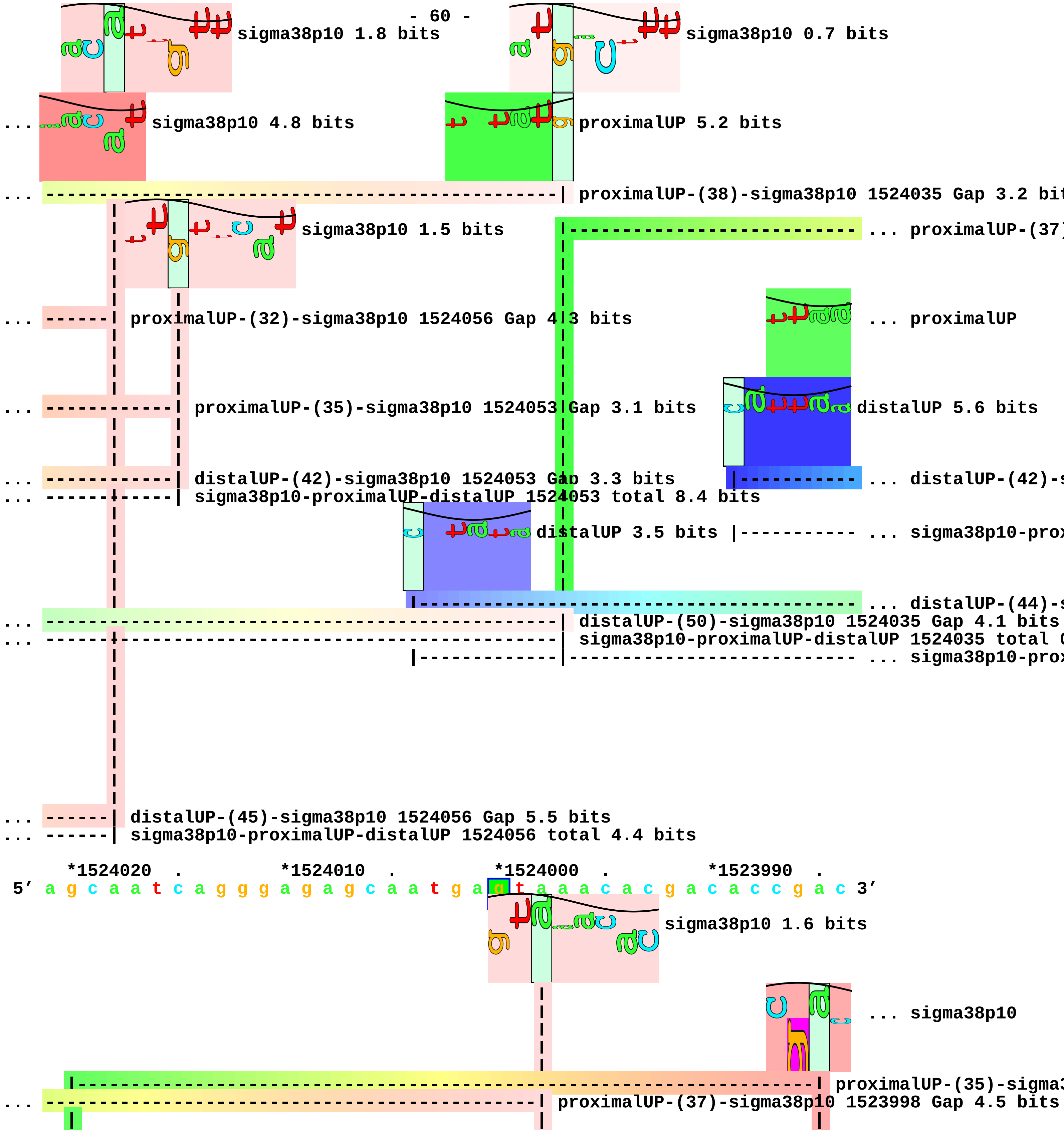
sigma38p10 1.8 bits
sigma38p10 0.7 bits
... sigma38p10 4.8 bits
proximalUP 5.2 bits
... proximalUP-(38)-sigma38p10 1524035 Gap 3.2 bit
sigma38p10 1.5 bits
... proximalUP-(37)
... proximalUP-(32)-sigma38p10 1524056 Gap 4.3 bits
... proximalUP
... proximalUP-(35)-sigma38p10 1524053 Gap 3.1 bits
distalUP 5.6 bits
... distalUP-(42)-sigma38p10 1524053 Gap 3.3 bits
... distalUP-(42)-s
... sigma38p10-proximalUP-distalUP 1524053 total 8.4 bits
distalUP 3.5 bits
... sigma38p10-prox
... distalUP-(44)-s
... distalUP-(50)-sigma38p10 1524035 Gap 4.1 bits
... sigma38p10-proximalUP-distalUP 1524035 total
... sigma38p10-prox
... distalUP-(45)-sigma38p10 1524056 Gap 5.5 bits
... sigma38p10-proximalUP-distalUP 1524056 total 4.4 bits
*1524020 . *1524010 . *1524000 . *1523990 .
5' a g c a a t c a g g g a g a g c a a t g a g t a a a c a c g a c a c c g a c 3'
sigma38p10 1.6 bits
... sigma38p10
proximalUP-(35)-sigma3
... proximalUP-(37)-sigma38p10 1523998 Gap 4.5 bits

```
... proximalUP 4.5 bits

... -----------------------------------------------------------------------| distalUP-(42)-sigma38p

... --|--------------------------------------------------------------------| sigma38p10-proximalUP-
... ----------------------------------------------| distalUP-(44)-sigma38p10 1523998 Gap 2.8 bits

... ----------------------------------------------| sigma38p10-proximalUP-distalUP 1523998 total 2.9

      *1523980  .         *1523970  .         *1523960  .         *1523950
 5' a c t t c a g a t c a a c a c g c c g c g a a a c g c c g c t g g c 3'

... sigma38p10 3.5 bits
```

piece 18, NC_000913, uxaB, config: linear, direction: -, begin: 1608999, end: 1608650

```
            .          *1608990   .          *1608980   .          *1608970   .
5' c t g t a t c g c t c g a a g a a a g a t g g g c g c a a c c g t a c c a g 3'
                 proximalUP 5.1 bits
                                                                                  ... proximalUP-(36)
               distalUP 1.0 bits
                                                                                  ... distalUP-(42)-s
                                                                                  ... sigma38p10-prox
T timestamp:_2018Sep20_11-34-53

   *1608960   .          *1608950   .          *1608940   .          *1608930   .
5' c a c t a t g c g t t a c g g t g a g g a a g t t g t c t g a c a t c t t t 3'
                 sigma38p10 3.5 bits                                              ... proximalUP
... --------| proximalUP-(36)-sigma38p10 1608957 Gap 3.6 bits                     ... distalUP
                                                                                  ... distalUP-(46)-s
... --------| distalUP-(42)-sigma38p10 1608957 Gap 3.3 bits                       ... sigma38p10-prox
... --------| sigma38p10-proximalUP-distalUP 1608957 total 2.7 bits

       *1608920   .          *1608910   .          *1608900   .          *1608890
5' a t t c a t t a c g c g c g a t t t g a a t g c a a t g t g t t c t a c c a 3'
...        proximalUP 6.2 bits
...    distalUP 4.7 bits
... ----------------------------------------------------------------------------- ... distalUP-(46)-s
... ----------------------------------------------------------------------------- ... sigma38p10-prox
      |-------------------------------------------------------------------------- ... proximalUP-(39)

 .         *1608880   .          *1608870   .          *1608860   .          *1608850
5' c t t t a t g a t c c a g g a a a a g g c g t t t t t c c t c g a t t a a a 3'
                 sigma38p10 4.6 bits                   proximalUP 6.2 bits
               proximalUP 6.3 bits                       |----------------------- ... proximalUP-(27)
```

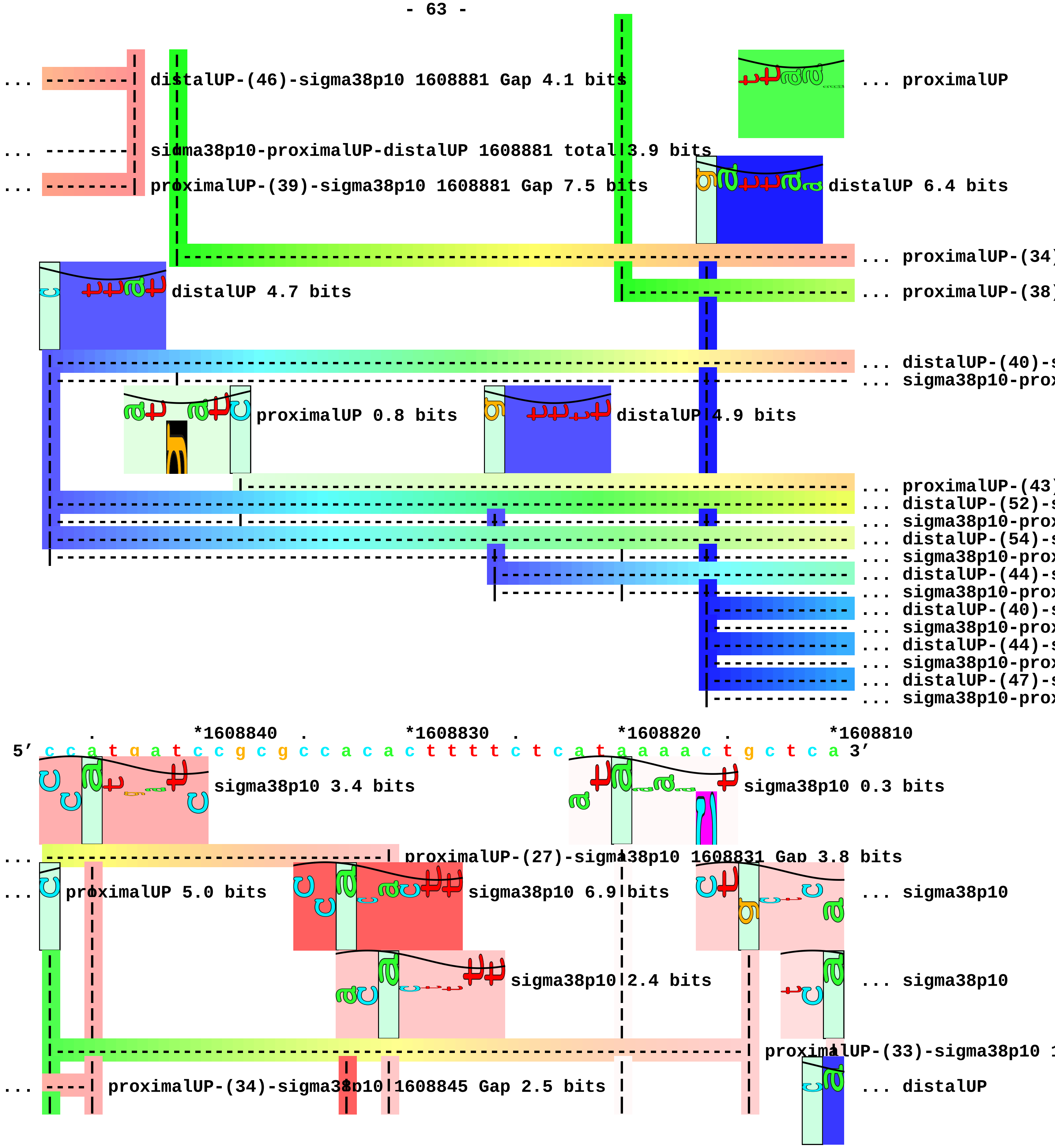
distalUP-(46)-sigma38p10 1608881 Gap 4.1 bits
... proximalUP
sigma38p10-proximalUP-distalUP 1608881 total 3.9 bits
proximalUP-(39)-sigma38p10 1608881 Gap 7.5 bits
distalUP 6.4 bits
... proximalUP-(34)
distalUP 4.7 bits
... proximalUP-(38)
... distalUP-(40)-
... sigma38p10-prox
proximalUP 0.8 bits
distalUP 4.9 bits
... proximalUP-(43)
... distalUP-(52)-
... sigma38p10-prox
... distalUP-(54)-
... sigma38p10-prox
... distalUP-(44)-
... sigma38p10-prox
... distalUP-(40)-
... sigma38p10-prox
... distalUP-(44)-
... sigma38p10-prox
... distalUP-(47)-
... sigma38p10-prox
*1608840 *1608830 *1608820 *1608810
5' c c a t g a t c c g c g c c a c a c t t t t c t c a t a a a a c t g c t c a 3'
sigma38p10 3.4 bits
sigma38p10 0.3 bits
proximalUP-(27)-sigma38p10 1608831 Gap 3.8 bits
proximalUP 5.0 bits
sigma38p10 6.9 bits
... sigma38p10
sigma38p10 2.4 bits
... sigma38p10
proximalUP-(33)-sigma38p10
proximalUP-(34)-sigma38p10 1608845 Gap 2.5 bits
... distalUP

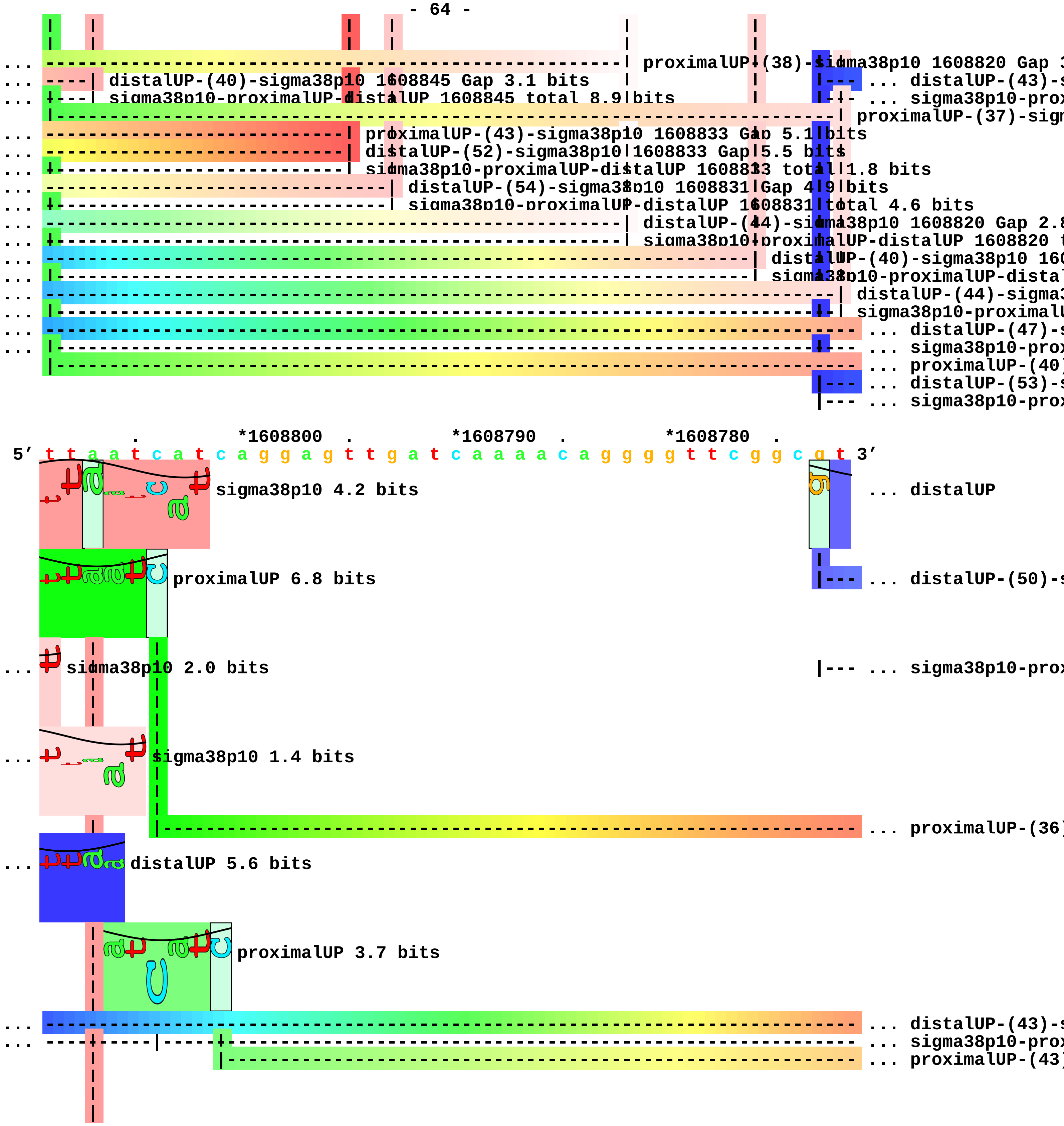
proximalUP-(38)-sigma38p10 1608820 Gap 3
distalUP-(43)-s
distalUP-(40)-sigma38p10 1608845 Gap 3.1 bits
sigma38p10-proximalUP-distalUP 1608845 total 8.9 bits
sigma38p10-prox
proximalUP-(37)-sigm
proximalUP-(43)-sigma38p10 1608833 Gap 5.1 bits
distalUP-(52)-sigma38p10 1608833 Gap 5.5 bits
sigma38p10-proximalUP-distalUP 1608833 total 1.8 bits
distalUP-(54)-sigma38p10 1608831 Gap 4.9 bits
sigma38p10-proximalUP-distalUP 1608831 total 4.6 bits
distalUP-(44)-sigma38p10 1608820 Gap 2.8
sigma38p10-proximalUP-distalUP 1608820 t
distalUP-(40)-sigma38p10 160
sigma38p10-proximalUP-distal
distalUP-(44)-sigma3
sigma38p10-proximalU
distalUP-(47)-s
sigma38p10-prox
proximalUP-(40)
distalUP-(53)-s
sigma38p10-prox
*1608800
*1608790
*1608780
5′ t t a a t c a t c a g g a g t t g a t c a a a a c a g g g g t t c g g c g t 3′
sigma38p10 4.2 bits
distalUP
proximalUP 6.8 bits
distalUP-(50)-s
sigma38p10 2.0 bits
sigma38p10-prox
sigma38p10 1.4 bits
proximalUP-(36)
distalUP 5.6 bits
proximalUP 3.7 bits
distalUP-(43)-s
sigma38p10-prox
proximalUP-(43)

```
... ----| distalUP-(47)-sigma38p10 1608807 Gap 6.5 bits
... ----| sigma38p10-proximalUP-distalUP 1608807 total 5.2 bits
... ----| proximalUP-(40)-sigma38p10 1608807 Gap 4.0 bits
... ----------------------------------------------------------------------------- ... distalUP-(53)-s
... ---------------|------------------------------------------------------------- ... sigma38p10-prox

     *1608770  .         *1608760  .         *1608750  .         *1608740  .
 5' a t t a a a g t t t g c t a a c c t g t c g t a c c a a t t c a a a c a g g 3'
... distalUP 4.3 bits
... ----------------------------------------------------------------------------- ... distalUP-(50)-s
... -------------------|--------------------------------------------------------- ... sigma38p10-prox
sigma38p10 5.7 bits
... ------| proximalUP-(36)-sigma38p10 1608768 Gap 3.6 bits
sigma38p10 4.8 bits
proximalUP 4.5 bits
... ------| distalUP-(43)-sigma38p10 1608768 Gap 3.6 bits
... ------| sigma38p10-proximalUP-distalUP 1608768 total 11.0 bits
... -------------------------| proximalUP-(43)-sigma38p10 1608758 Gap 5.1 bits
|----------------------------------------------------- ... proximalUP-(38)
... -------------------------| distalUP-(53)-sigma38p10 1608758 Gap 4.9 bits
... -------------------------| sigma38p10-proximalUP-distalUP 1608758 total 4.1 bits

         *1608730  .         *1608720  .         *1608710  .         *1608700
 5' t t g t a t g a c t a a t c a g a a g g g a a c c c a t t g t g a a a a c a 3'
```

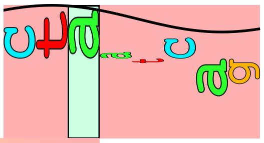

- 66 -
sigma38p10 3.4 bits

... --------------------| distalUP-(50)-sigma38p10 1608723 Gap 4.1 bits
... --------------------| sigma38p10-proximalUP-distalUP 1608723 total 4.8 bits

... --------------------| proximalUP-(38)-sigma38p10 1608723 Gap 3.2 bits

```
    .         *1608690  .         *1608680  .         *1608670  .         *1608660
5' c t a a a t c g t c g c g a t t t t c c c g g t g c a c a g t a t c c a g a 3'

      .         *1608650
5' a c g t a t c a 3'
```

piece 19, NC_000913, ydgA, config: linear, direction: +, begin: 1687694, end: 1687957

```
     .         *1687700  .         *1687710  .         *1687720  .         *1687730
5' a t c a g c g t t t a t t g c c g c c a a c g a a t c a c c g g t g a c t g 3'
 T timestamp:_2018Sep20_11-34-53

       .         *1687740  .         *1687750  .         *1687760  .
5' t c a a a g g c c a c g g c c g t t t a g c g c g t g t t t a c a a c a a g 3'
                                                              distalUP 0.9 bits
                                                               ... distalUP-(44)-s
                                                               ... distalUP-proxin

   *1687770  .         *1687780  .         *1687790  .         *1687800  .
5' c t g t a a g a g c t t a c t g a a a a a a t t a a c a t c t c t t g c t a 3'
     proximalUP 2.8 bits                                          ... sigma38p10
...                                                               distalUP-(44)-sigma3
...                                                               distalUP-proximalUP-
                                                                  proximalUP-(31)-sign

     *1687810  .         *1687820  .         *1687830  .         *1687840  .
5' a g c t t a t t a a a g g c t t a t a a c a c c t t c a g g c g g c c a g t 3'
... sigma38p10 9.1 bits       proximalUP 3.9 bits
          proximalUP 4.5 bits
                                                               ... proximalUP-(33)
        distalUP 4.1 bits                                      ... proximalUP-(35)
                                                               ... distalUP-(42)-s
                                                               ... distalUP-proxin
                    distalUP 3.5 bits
                                                               ... distalUP-(42)-s
                                                               ... distalUP-proxin

       *1687850  .         *1687860  .         *1687870  .         *1687880
5' c c g c c t g a t t t c a t t t t a t g g a t a a t c a t t a t g a a t a a 3'
             sigma38p10 1.9 bits                               ... proximalUP-(34)
```

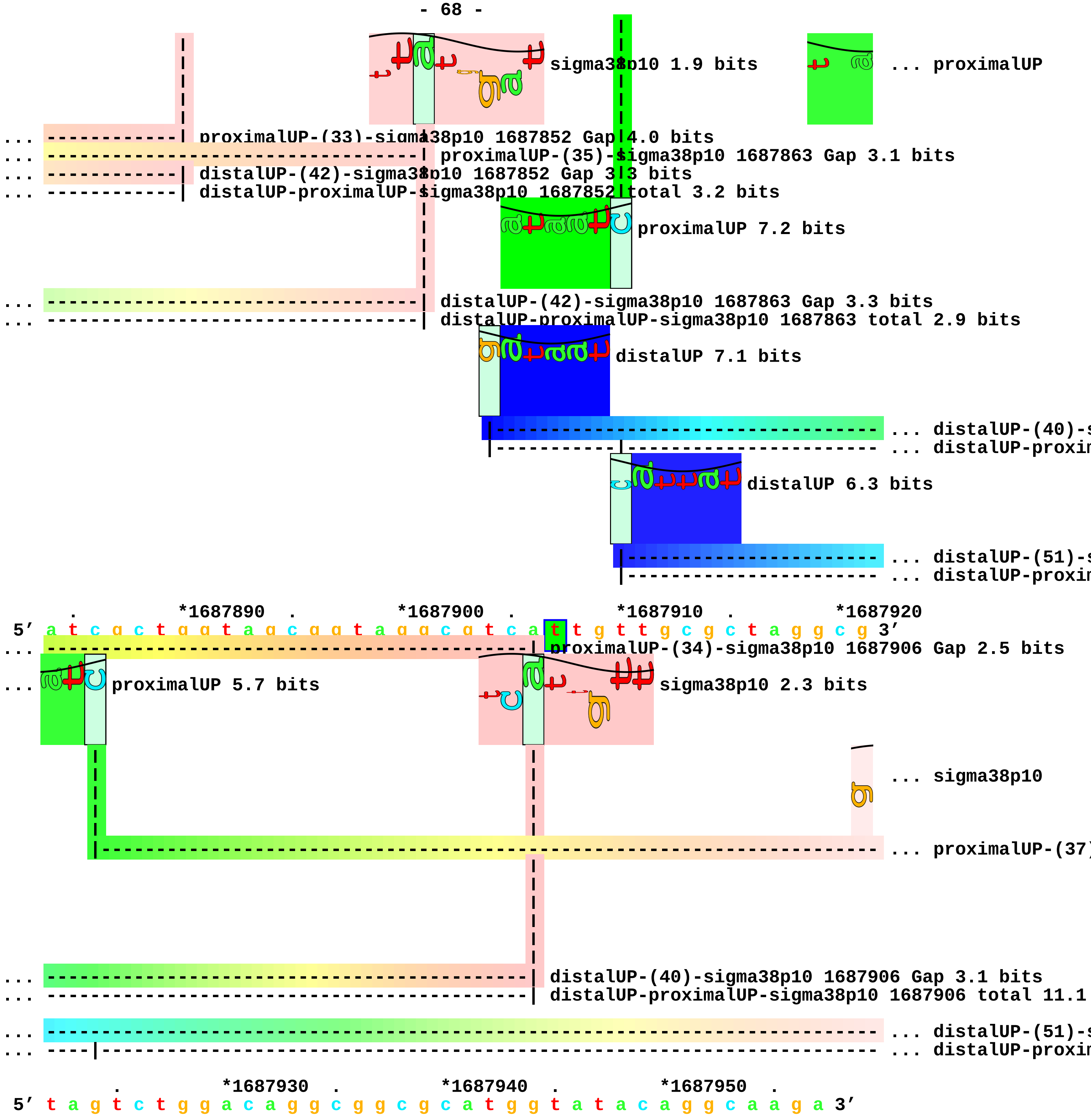
sigma38p10 1.9 bits
... proximalUP
... proximalUP-(33)-sigma38p10 1687852 Gap 4.0 bits
... proximalUP-(35)-sigma38p10 1687863 Gap 3.1 bits
... distalUP-(42)-sigma38p10 1687852 Gap 3.3 bits
... distalUP-proximalUP-sigma38p10 1687852 total 3.2 bits
proximalUP 7.2 bits
... distalUP-(42)-sigma38p10 1687863 Gap 3.3 bits
... distalUP-proximalUP-sigma38p10 1687863 total 2.9 bits
distalUP 7.1 bits
... distalUP-(40)-s
... distalUP-proxim
distalUP 6.3 bits
... distalUP-(51)-s
... distalUP-proxim
*1687890
*1687900
*1687910
*1687920
5′ a t c g c t g g t a g c g g t a g g c g t c a t t g t t g c g c t a g g c g 3′
... proximalUP-(34)-sigma38p10 1687906 Gap 2.5 bits
... proximalUP 5.7 bits
sigma38p10 2.3 bits
... sigma38p10
... proximalUP-(37)
... distalUP-(40)-sigma38p10 1687906 Gap 3.1 bits
... distalUP-proximalUP-sigma38p10 1687906 total 11.1
... distalUP-(51)-s
... distalUP-proxim
*1687930
*1687940
*1687950
5′ t a g t c t g g a c a g g c g g c g c a t g g t a t a c a g g c a a g a 3′

... 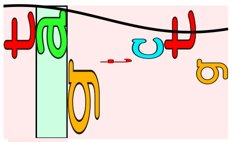 sigma38p10 0.9 bits

... --| proximalUP-(37)-sigma38p10 1687923 Gap 4.5 bits

... --| distalUP-(51)-sigma38p10 1687923 Gap 5.1 bits
... --| distalUP-proximalUP-sigma38p10 1687923 total 3.2 bits

piece 20, NC_000913, lpp, config: linear, direction: +, begin: 1755300, end: 1755549

```
   *1755300   .          *1755310   .          *1755320   .          *1755330   .
5' a t c t a a a a c c c a g c g t t c g a t g c t t c t t t g a g c g a a c g 3'
   T timestamp:_2018Sep20_11-34-53

       *1755340   .          *1755350   .          *1755360   .          *1755370   .
5' a t c a a a a a t a a g t g c c t t c c c a t c a a a a a a a t a t t c t c 3'
                                proximalUP 3.8 bits
                                                                        ... proximalUP-(35)
                 distalUP 5.7 bits
                                                                        ... distalUP-(44)-s
                                                                        ... distalUP-proxim

           *1755380   .          *1755390   .          *1755400   .          *1755410
5' a a c a t a a a a a a c t t t g t g t a a t a c t t g t a a c g c t a c a t 3'
                  sigma38p10 0.9 bits                        proximalUP 3.3 bits
... ----------------| proximalUP-(35)-sigma38p10 1755384 Gap 3.1 bits
                          distalUP 1.1 bits                            ... proximalUP-(32)
... ----------------| distalUP-(44)-sigma38p10 1755384 Gap 2.8 bits
... ----------------| distalUP-proximalUP-sigma38p10 1755384 total 4.5 bits
                                 distalUP 4.4 bits
                                                                        ... distalUP-(45)-s
                                                                        ... distalUP-proxim
                                                                        ... proximalUP-(35)
                                                                        ... distalUP-(50)-s
                                                                        ... distalUP-proxim

      .          *1755420   .          *1755430   .          *1755440   .          *1755450
5' g g a g a t t a a c t c a a t c t a g a g g g t a t t a a t a a t g a a a g 3'
              proximalUP 5.4 bits                         sigma38p10 3.3 bits
             distalUP 6.4 bits                                  sigma38p10 3.7 bits
... ---------------------------------------------| proximalUP-(32)-sigma38p10 1755438 Gap 4.3 bit
```

... proximalUP-(34)
... distalUP-(40)-s
... distalUP-proxim
... distalUP-(45)-sigma38p10 1755438 Gap 5.5 bits
... distalUP-proximalUP-sigma38p10 1755438 total 1
... proximalUP-(35)-sigma38p10 1755441 Gap 3
... distalUP-(50)-sigma38p10 1755441 Gap 4.1
... distalUP-proximalUP-sigma38p10 1755441 t

proximalUP 6.7 bits

... proximalUP-(43)

distalUP 4.3 bits

... distalUP-(54)-s
... distalUP-proxim

. *1755460 . *1755470 . *1755480 .
5′ c t a c t a a a c t g g t a c t g g g c g c g g t a a t c c t g g g t t c t 3′

sigma38p10 7.8 bits ... sigma38p10

... proximalUP-(34)-sigma38p10 1755457 Gap 2.5 bits
... distalUP-(40)-sigma38p10 1755457 Gap 3.1 bits
... distalUP-proximalUP-sigma38p10 1755457 total 14.1 bits

... ... proximalUP-(43)
... ... distalUP-(54)-s
... ... distalUP-proxim

*1755490 . *1755500 . *1755510 . *1755520 .
5′ a c t c t g c t g g c a g g t t g c t c c a g c a a c g c t a a a a t c g a 3′

... sigma38p10 6.0 bits

```
    |
... | proximalUP-(43)-sigma38p10 1755490 Gap 5.1 bits
    |
... | distalUP-(54)-sigma38p10 1755490 Gap 4.9 bits
... | distalUP-proximalUP-sigma38p10 1755490 total 7.0 bits

         *1755530  .         *1755540  .
 5' t c a g c t g t c t t c t g a c g t t c a g 3'
```

piece 21, NC_000913, ynhG, config: linear, direction: -, begin: 1756935, end: 1756770

. *1756930 . *1756920 . *1756910 . *1756900
5′ t t c g c g c a a t t c g c g c c a a a g c c g c t g c a c t t a g c t a a 3′
T timestamp:_2018Sep20_11-34-53

. *1756890 . *1756880 . *1756870 . *1756860
5′ a c t a a a a g g a c a g c t t t c a t c c g g t t g g c t c a c a t t g c 3′

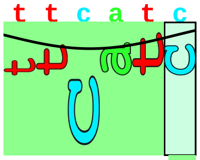
proximalUP 3.2 bits

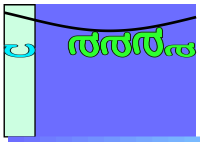
distalUP 4.1 bits

--- ... proximalUP-(27)
--- ... distalUP-(46)-s
--- ... sigma38p10-prox
--- ... proximalUP-(34)
--- ... distalUP-(53)-s
--- ... sigma38p10-prox

. *1756850 . *1756840 . *1756830 .
5′ c g g a t g g t c a a t g t t t a c a c t g a c c t g c a c a g g a g t g t 3′

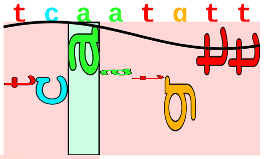
sigma38p10 1.7 bits

... ---| proximalUP-(27)-sigma38p10 1756850 Gap 3.8 bits
... ---| distalUP-(46)-sigma38p10 1756850 Gap 4.1 bits
... ---| sigma38p10-proximalUP-distalUP 1756850 total 1.2 bits
... ---| proximalUP-(34)-sigma38p10 1756843 Gap 2.5 bits
... ---| distalUP-(53)-sigma38p10 1756843 Gap 4.9 bits
... ---| sigma38p10-proximalUP-distalUP 1756843 total 5.5 bits

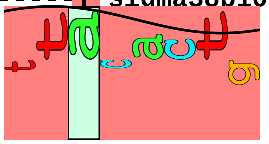
sigma38p10 5.5 bits

*1756820 . *1756810 . *1756800 . *1756790 .
5′ t t g g t t t c a g g t g a a c a t a a g c g c c a t c c g a c g t t t c 3′

*1756780 . *1756770
5′ a g c g t g a g t c t c c g 3′

piece 22, NC_000913, sdaA, config: linear, direction: +, begin: 1894613, end: 1894946

. *1894620 . *1894630 . *1894640 . *1894650
5′ c g c c c a g g c a t t a c a t c t g g g t c g t t a t c a c c c t t t a g 3′

T timestamp:_2018Sep20_11-34-53

... distalUP
... distalUP-(55)-s
... distalUP-proxin

. *1894660 . *1894670 . *1894680 .
5′ a t a t c t a c c g c c g t g g t g a t t c a c a t c g g g t a t g g c t g 3′

... distalUP 2.7 bits
proximalUP 4.5 bits
... ... distalUP-(55)-s
... ... distalUP-proxin
... proximalUP-(33)

*1894690 . *1894700 . *1894710 . *1894720 .
5′ t c c t g g t a c g a a c a g t a t t t t g t a t g g g g a a t g a c c g c 3′

sigma38p10 3.5 bits
... distalUP-(55)-sigma38p10 1894705 Gap 4.9 bits
... distalUP-proximalUP-sigma38p10 1894705 total 1.8 bits
... proximalUP-(33)-sigma38p10 1894705 Gap 4.0 bits
proximalUP 6.1 bits
... proximalUP-(41)
distalUP 2.6 bits
... distalUP-(50)-s
... distalUP-proxin
... proximalUP-(43)
distalUP 5.0 bits
... distalUP-(50)-s
... distalUP-proxin

*1894730 . *1894740 . *1894750 . *1894760
5′ a g g c a t a a t t c g t g a g c t g g c g c t g c a a a t t g g t g t g a 3′

proximalUP 5.1 bits
sigma38p10 2.6 bits

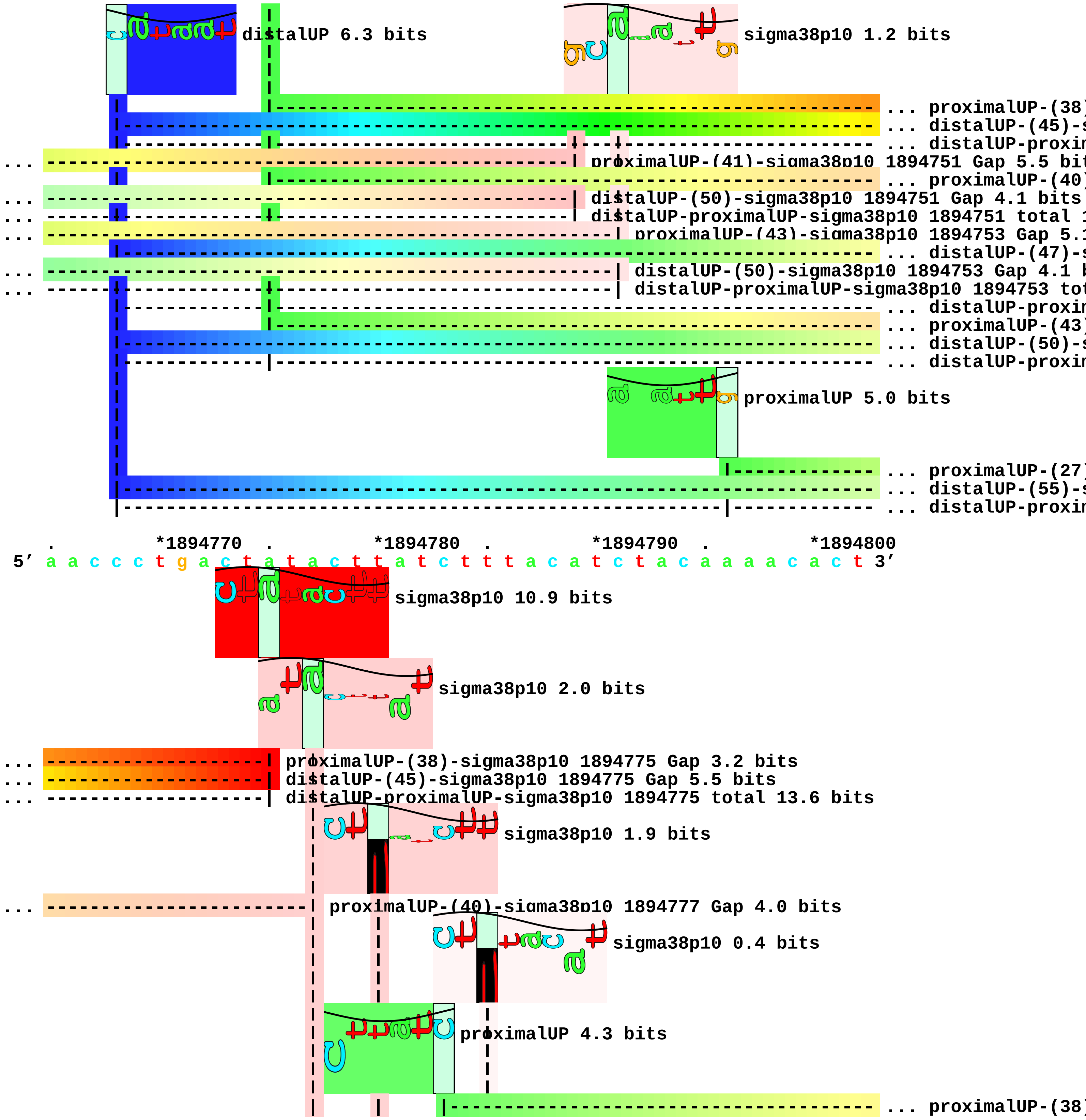
distalUP 6.3 bits
sigma38p10 1.2 bits
... proximalUP-(38)
... distalUP-(45)-s
... distalUP-proxim
... proximalUP-(41)-sigma38p10 1894751 Gap 5.5 bit
... proximalUP-(40)
... distalUP-(50)-sigma38p10 1894751 Gap 4.1 bits
... distalUP-proximalUP-sigma38p10 1894751 total 1
... proximalUP-(43)-sigma38p10 1894753 Gap 5.1
... distalUP-(47)-s
... distalUP-(50)-sigma38p10 1894753 Gap 4.1 b
... distalUP-proximalUP-sigma38p10 1894753 tot
... distalUP-proxim
... proximalUP-(43)
... distalUP-(50)-s
... distalUP-proxim
proximalUP 5.0 bits
... proximalUP-(27)
... distalUP-(55)-s
... distalUP-proxim
. *1894770 . *1894780 . *1894790 . *1894800
5′ a a c c c t g a c t a t a c t t a t c t t t a c a t c t a c a a a a c a c t 3′
sigma38p10 10.9 bits
sigma38p10 2.0 bits
... proximalUP-(38)-sigma38p10 1894775 Gap 3.2 bits
... distalUP-(45)-sigma38p10 1894775 Gap 5.5 bits
... distalUP-proximalUP-sigma38p10 1894775 total 13.6 bits
sigma38p10 1.9 bits
... proximalUP-(40)-sigma38p10 1894777 Gap 4.0 bits
sigma38p10 0.4 bits
proximalUP 4.3 bits
... proximalUP-(38)

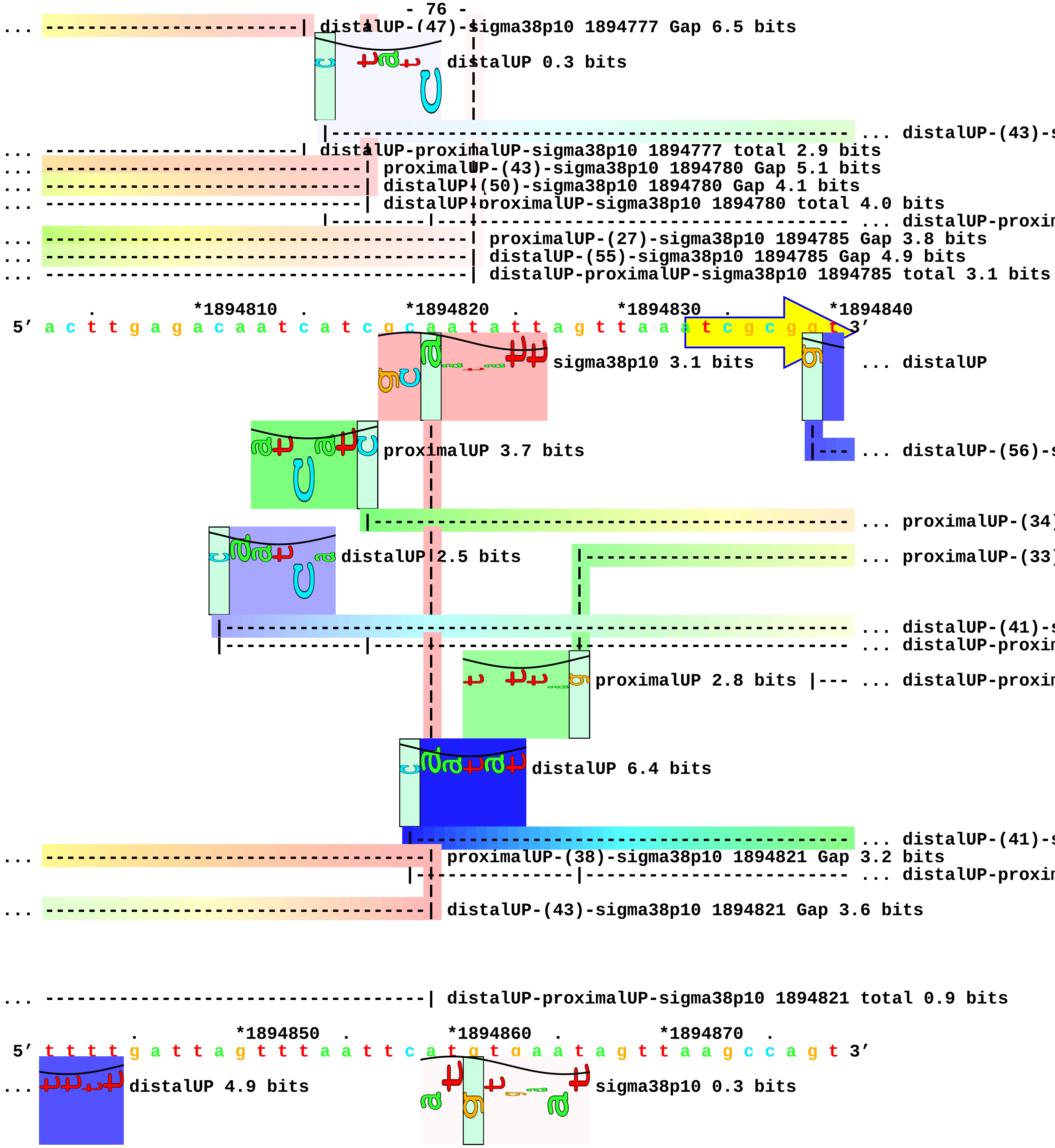
distalUP-(47)-sigma38p10 1894777 Gap 6.5 bits
distalUP 0.3 bits
... distalUP-(43)-s
distalUP-proximalUP-sigma38p10 1894777 total 2.9 bits
proximalUP-(43)-sigma38p10 1894780 Gap 5.1 bits
distalUP-(50)-sigma38p10 1894780 Gap 4.1 bits
distalUP-proximalUP-sigma38p10 1894780 total 4.0 bits
... distalUP-proxim
proximalUP-(27)-sigma38p10 1894785 Gap 3.8 bits
distalUP-(55)-sigma38p10 1894785 Gap 4.9 bits
distalUP-proximalUP-sigma38p10 1894785 total 3.1 bits
*1894810
*1894820
*1894830
*1894840
5′ a c t t g a g a c a a t c a t c g c a a t a t t a g t t a a a t c g c g g t 3′
sigma38p10 3.1 bits
... distalUP
proximalUP 3.7 bits
... distalUP-(56)-s
... proximalUP-(34)
distalUP 2.5 bits
... proximalUP-(33)
... distalUP-(41)-s
... distalUP-proxim
proximalUP 2.8 bits
... distalUP-proxim
distalUP 6.4 bits
... distalUP-(41)-s
proximalUP-(38)-sigma38p10 1894821 Gap 3.2 bits
... distalUP-proxim
distalUP-(43)-sigma38p10 1894821 Gap 3.6 bits
distalUP-proximalUP-sigma38p10 1894821 total 0.9 bits
*1894850
*1894860
*1894870
5′ t t t t g a t t a g t t t a a t t c a t g t g a a t a g t t a a g c c a g t 3′
distalUP 4.9 bits
sigma38p10 0.3 bits

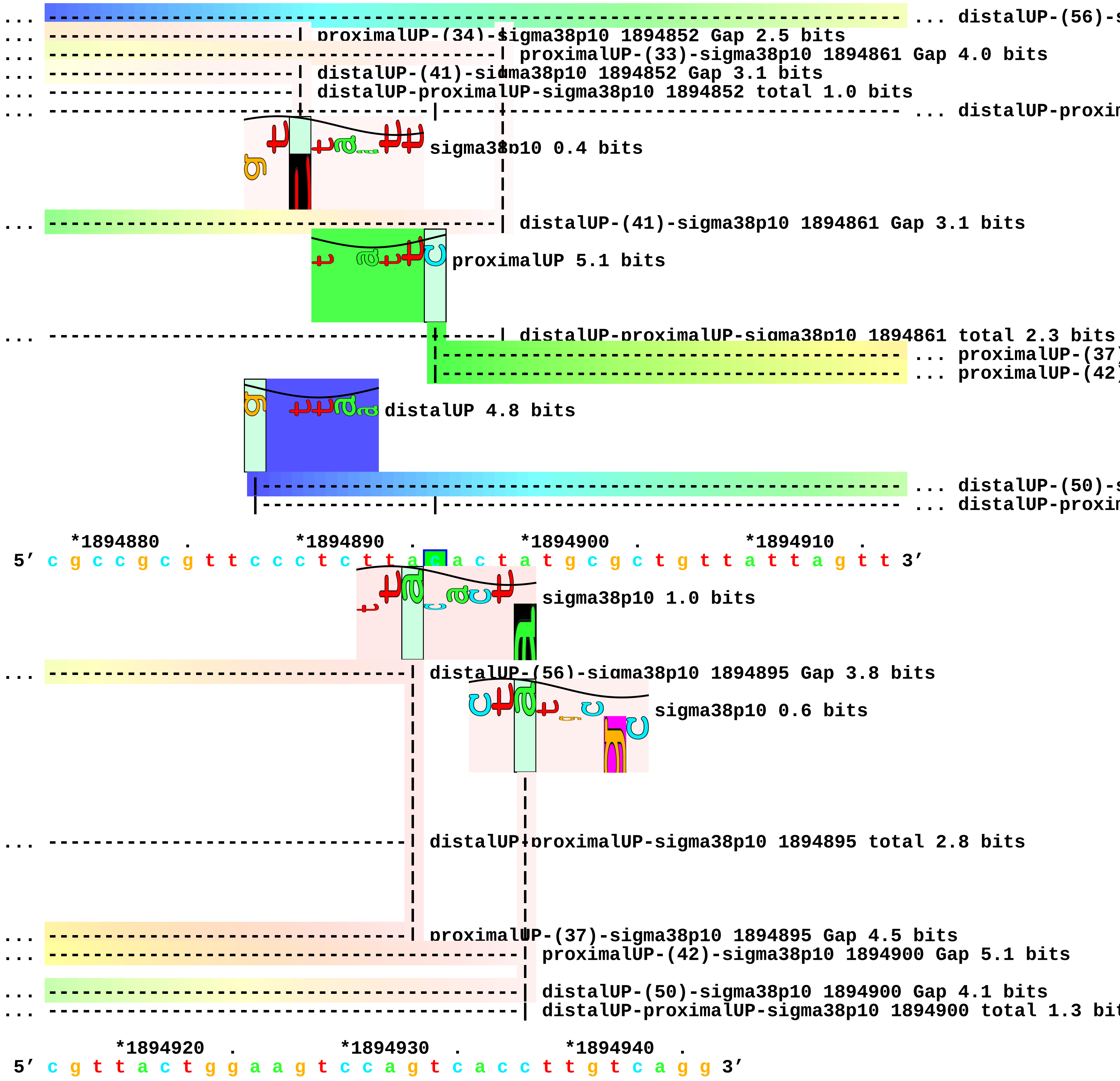
... distalUP-(56)-s
... proximalUP-(34)-sigma38p10 1894852 Gap 2.5 bits
... proximalUP-(33)-sigma38p10 1894861 Gap 4.0 bits
... distalUP-(41)-sigma38p10 1894852 Gap 3.1 bits
... distalUP-proximalUP-sigma38p10 1894852 total 1.0 bits
... distalUP-proxin
sigma38p10 0.4 bits
... distalUP-(41)-sigma38p10 1894861 Gap 3.1 bits
proximalUP 5.1 bits
... distalUP-proximalUP-sigma38p10 1894861 total 2.3 bits
... proximalUP-(37)
... proximalUP-(42)
distalUP 4.8 bits
... distalUP-(50)-s
... distalUP-proxin
*1894880 . *1894890 . *1894900 . *1894910 .
5′ c g c c g c g t t c c c t c t t a c a c t a t g c g c t g t t a t t a g t t 3′
sigma38p10 1.0 bits
... distalUP-(56)-sigma38p10 1894895 Gap 3.8 bits
sigma38p10 0.6 bits
... distalUP-proximalUP-sigma38p10 1894895 total 2.8 bits
... proximalUP-(37)-sigma38p10 1894895 Gap 4.5 bits
... proximalUP-(42)-sigma38p10 1894900 Gap 5.1 bits
... distalUP-(50)-sigma38p10 1894900 Gap 4.1 bits
... distalUP-proximalUP-sigma38p10 1894900 total 1.3 bit
*1894920 . *1894930 . *1894940 .
5′ c g t t a c t g g a a g t c c a g t c a c c t t g t c a g g 3′

piece 23, NC_000913, yobF, config: linear, direction: -, begin: 1905834, end: 1905497

*1905830 . *1905820 . *1905810 . *1905800
5' g a a c t a t g t t a g a a a a t a c g c c a g t t t a a g t a t c t g c c 3'
proximalUP 3.7 bits ... proximalUP-(28)
T timestamp:_2018Sep20_11-34-53 ... proximalUP-(35)
distalUP 7.2 bits
... distalUP-(42)-s
... sigma38p10-prox
proximalUP 4.5 bits
... distalUP-(46)-s
... sigma38p10-prox

. *1905790 . *1905780 . *1905770 . *1905760
5' t g a a c t g g c a a g g t t a a g c a c a a t g a t a t a t c g g c g c g 3'
... proximalUP-(28)-sigma38p10 1905777 Gap 5.5 bits
... proximalUP-(35)-sigma38p10 1905781 Gap 3.1 bits
sigma38p10 1.3 bits
... distalUP-(42)-sigma38p10 1905781 Gap 3.3 bits
... sigma38p10-proximalUP-distalUP 1905781 total 5.9 bits
sigma38p10 1.1 bits
... distalUP-(46)-sigma38p10 1905777 Gap 4.1 bits
... sigma38p10-proximalUP-distalUP 1905777 total 3.2 bits

. *1905750 . *1905740 . *1905730 .
5' t a t t c c g t t g c a t a a g t g t g c a a a a a a a g t g g a a g a c g 3'
proximalUP 3.8 bits
... proximalUP-(38)
distalUP 5.7 bits ... proximalUP
... distalUP-(46)-s
... sigma38p10-prox
... distalUP-(52)-s
... sigma38p10-prox

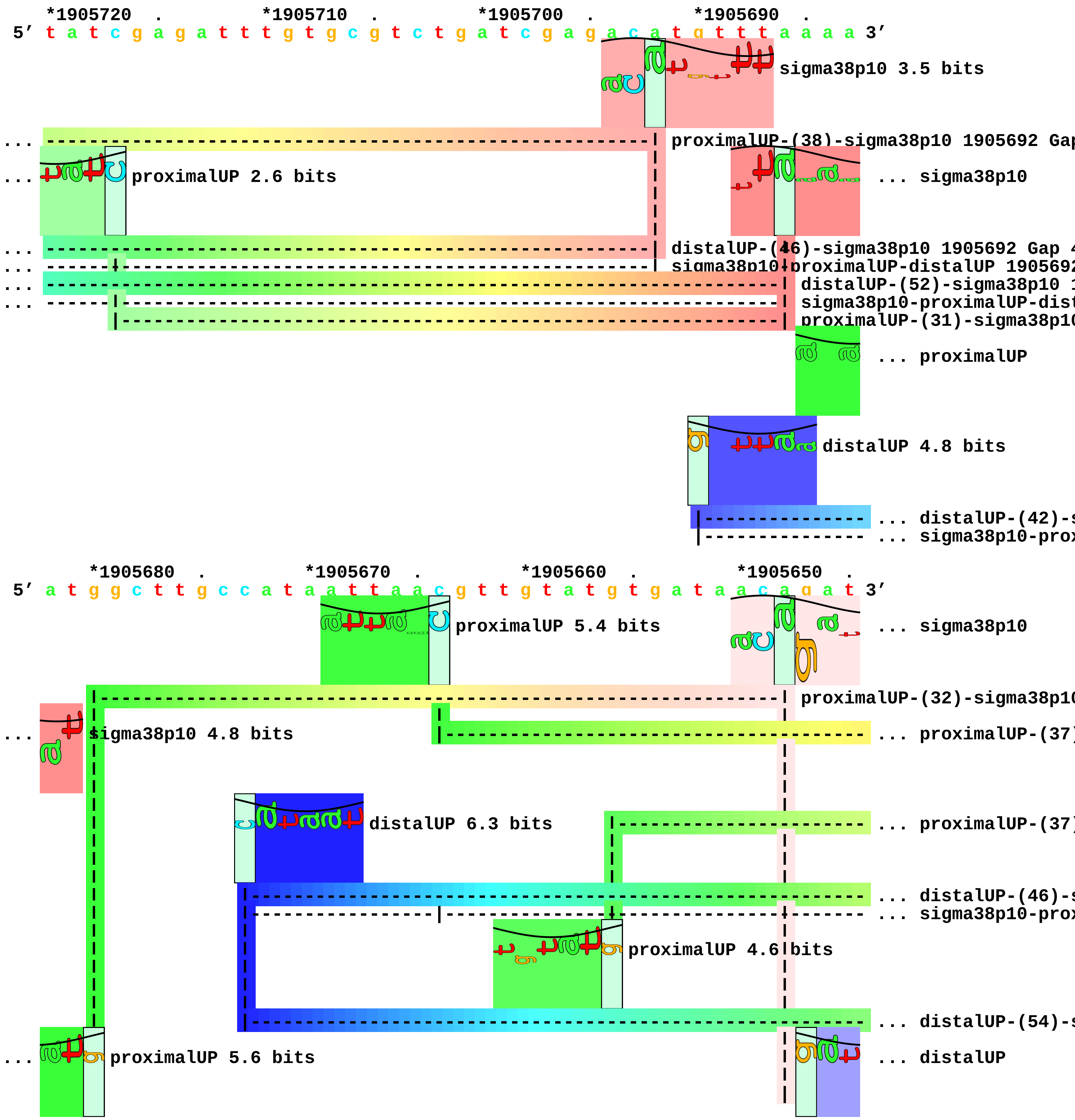
*1905720 . *1905710 . *1905700 . *1905690 .
5′ t a t c g a g a t t t g t g c g t c t g a t c g a g a c a t g t t t a a a a 3′
sigma38p10 3.5 bits
proximalUP-(38)-sigma38p10 1905692 Gap
proximalUP 2.6 bits
... sigma38p10
distalUP-(46)-sigma38p10 1905692 Gap 4
sigma38p10-proximalUP-distalUP 190569
distalUP-(52)-sigma38p10 1
sigma38p10-proximalUP-dist
proximalUP-(31)-sigma38p10
... proximalUP
distalUP 4.8 bits
... distalUP-(42)-s
... sigma38p10-prox
*1905680 . *1905670 . *1905660 . *1905650 .
5′ a t g g c t t g c c a t a a t t a a c g t t g t a t g t g a t a a c a g a t 3′
proximalUP 5.4 bits
... sigma38p10
proximalUP-(32)-sigma38p10
sigma38p10 4.8 bits
... proximalUP-(37)
distalUP 6.3 bits
... proximalUP-(37)
... distalUP-(46)-s
... sigma38p10-prox
proximalUP 4.6 bits
... distalUP-(54)-s
proximalUP 5.6 bits
... distalUP

```
                       |-----------------------------------|-----------------|--|----- ... sigma38p10-prox
... -----------------------------------------------------------------------|  distalUP-(42)-sigma38p10 1
... ----|--------------------------------------------------------------------|  sigma38p10-proximalUP-dist
                                                                               |----- ... distalUP-(41)-s
                                                                               |----- ... sigma38p10-prox

            *1905640   .            *1905630   .            *1905620   .            *1905610
5' t t c g g g t t a a a c g a g g t a c a g t t c t g t t t a t g t g t g g c 3'
... sigma38p10 1.0 bits                         sigma38p10 4.2 bits                    ... sigma38p10
                  proximalUP 5.0 bits                         sigma38p10 2.1 bits
... -----------------------------------| proximalUP-(37)-sigma38p10 1905627 Gap 4.5 bits
... ------------------------------------------------------| proximalUP-(37)-sigma38p10 1905619 Gap 4.5 b
... -----------------------------------| distalUP-(46)-sigma38p10 1905627 Gap 4.1 bits
... -----------------------------------| sigma38p10-proximalUP-distalUP 1905627 total 7.3 bits
                      |-------------------------------------------------------- ... proximalUP-(27)
... ------------------------------------------------------| distalUP-(54)-sigma38p10 1905619 Gap 4.9 bit
... distalUP 2.7 bits

... ------------------------------------------------------| sigma38p10-proximalUP-distalUP 1905619 total

... ----------------------------------------------------------------------- ... distalUP-(41)-s
... ---------------------|------------------------------------------------- ... sigma38p10-prox

      .        *1905600   .            *1905590   .            *1905580   .            *1905570
5' a t t t t c a g t a a a g a a g t c c t g a g t a a a c a c g t t g a c g t 3'
... sigma38p10 0.3 bits

... | proximalUP-(27)-sigma38p10 1905606 Gap 3.8 bits

... | distalUP-(41)-sigma38p10 1905606 Gap 3.1 bits
... | sigma38p10-proximalUP-distalUP 1905606 total 1.1 bits

         .        *1905560   .            *1905550   .            *1905540   .
5' t g a a t a c c g c t t c t c t g c c g a g c c t t a t a t t g g t g c c t 3'
                                                           proximalUP 6.1 bits
```

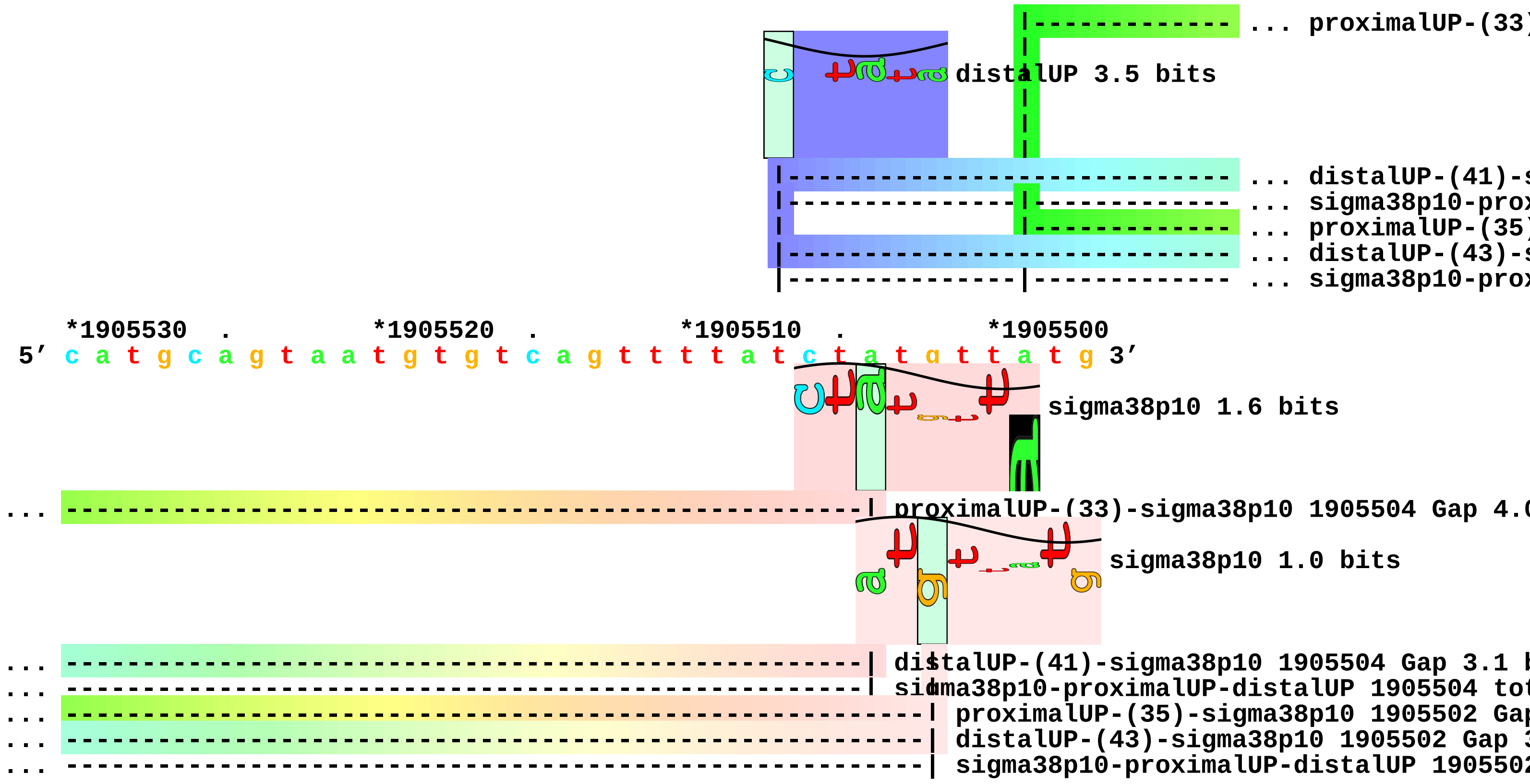

... proximalUP-(33)
distalUP 3.5 bits
... distalUP-(41)-s
... sigma38p10-prox
... proximalUP-(35)
... distalUP-(43)-s
... sigma38p10-prox
*1905530 . *1905520 . *1905510 . *1905500
5' c a t g c a g t a a t g t g t c a g t t t t a t c t a t g t t a t g 3'
sigma38p10 1.6 bits
... proximalUP-(33)-sigma38p10 1905504 Gap 4.0
sigma38p10 1.0 bits
... distalUP-(41)-sigma38p10 1905504 Gap 3.1 b
... sigma38p10-proximalUP-distalUP 1905504 tot
... proximalUP-(35)-sigma38p10 1905502 Gap
... distalUP-(43)-sigma38p10 1905502 Gap 3
... sigma38p10-proximalUP-distalUP 1905502

piece 24, NC_000913, yebW, config: linear, direction: +, begin: 1919983, end: 1920253

. *1919990 . *1920000 . *1920010 . *1920020

5' c c t t t a c c g t a c c c g t t a c g a t c a a c a a t c t g a t g c c t 3'

T timestamp:_2018Sep20_11-34-53

. *1920030 . *1920040 . *1920050 .

5' g g a t t a t g c g t c [illegible] t g c c t g a t c c a a a a a g a a c c c g t c g 3'

proximalUP 6.7 bits

... proximalUP-(43)

distalUP 7.1 bits

... distalUP-(49)-s

... distalUP-proxim

*1920060 . *1920070 . *1920080 . *1920090 .

5' g c a t g g c g g g t t a t t t g t c c t g g t t a t t c c c c c g t t g t 3'

sigma38p10 0.8 bits

... distalUP

... proximalUP-(43)-sigma38p10 1920071 Gap 5.1 bits

proximalUP 4.6 bits

... distalUP-(47)-s

... distalUP-(49)-sigma38p10 1920071 Gap 5.9 bits ... distalUP-proxim

... distalUP-proximalUP-sigma38p10 1920071 total 3.5 bits

... proximalUP-(34)

distalUP 4.9 bits

... distalUP-(41)-s

... distalUP-proxim

distalUP 4.9 bits

... distalUP-(48)-s

... distalUP-proxim

*1920100 . *1920110 . *1920120 . *1920130

5' a a a a t c t c t c c t a a a c t t a a c g g t a c g g c a c c a c a c t t 3'

... distalUP 5.0 bits

sigma38p10 6.9 bits

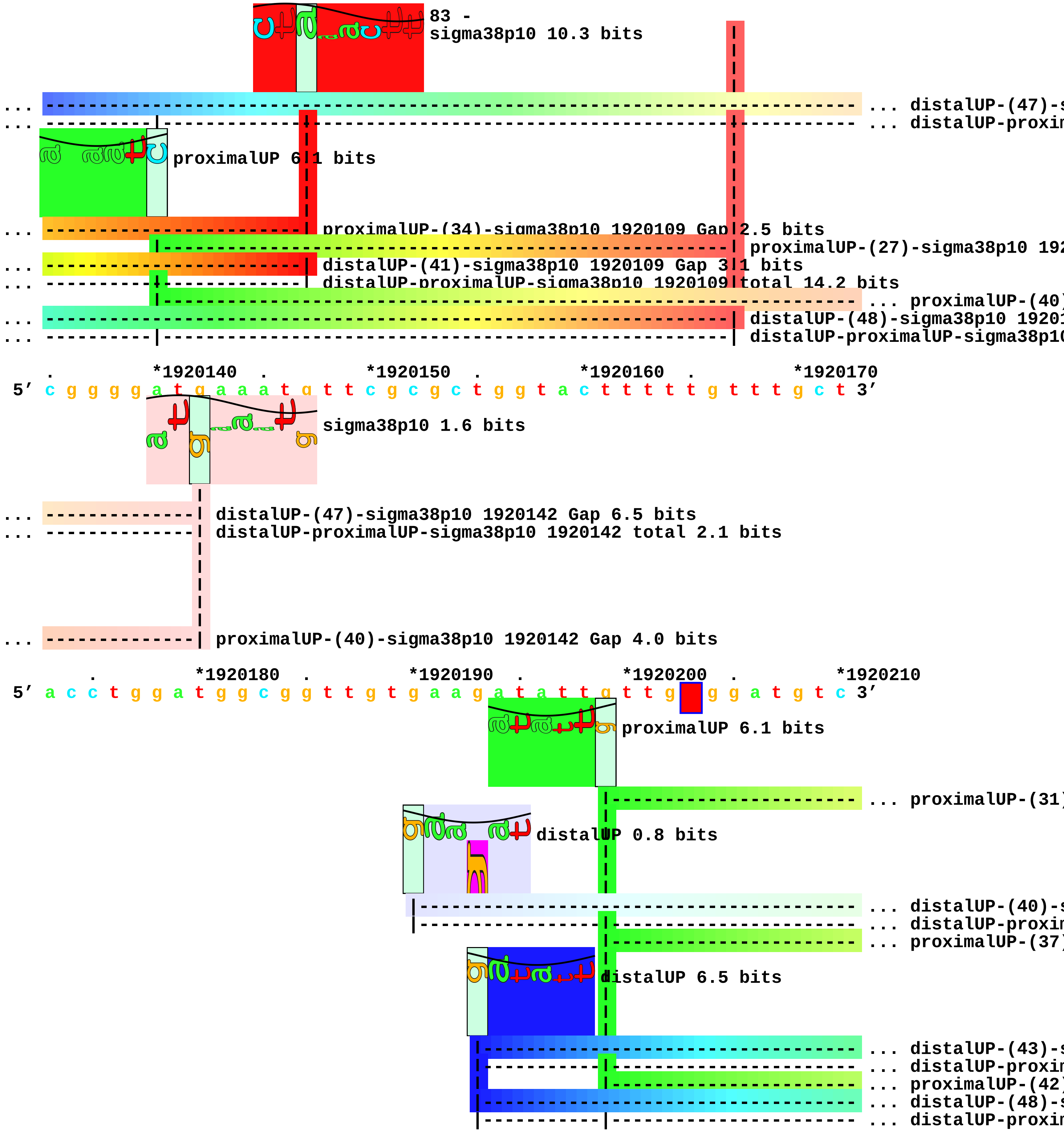
83 -
sigma38p10 10.3 bits
... distalUP-(47)-s
... distalUP-proxin
proximalUP 6.1 bits
... proximalUP-(34)-sigma38p10 1920109 Gap 2.5 bits
proximalUP-(27)-sigma38p10 192
... distalUP-(41)-sigma38p10 1920109 Gap 3.1 bits
... distalUP-proximalUP-sigma38p10 1920109 total 14.2 bits
... proximalUP-(40)
... distalUP-(48)-sigma38p10 1920
... distalUP-proximalUP-sigma38p10
*1920140 *1920150 *1920160 *1920170
5′ c g g g g a t g a a a t g t t c g c g c t g g t a c t t t t t g t t t g c t 3′
sigma38p10 1.6 bits
... distalUP-(47)-sigma38p10 1920142 Gap 6.5 bits
... distalUP-proximalUP-sigma38p10 1920142 total 2.1 bits
... proximalUP-(40)-sigma38p10 1920142 Gap 4.0 bits
*1920180 *1920190 *1920200 *1920210
5′ a c c t g g a t g g c g g t t g t g a a g a t a t t g t t g g g a t g t c 3′
proximalUP 6.1 bits
... proximalUP-(31)
distalUP 0.8 bits
... distalUP-(40)-s
... distalUP-proxin
... proximalUP-(37)
distalUP 6.5 bits
... distalUP-(43)-s
... distalUP-proxin
... proximalUP-(42)
... distalUP-(48)-s
... distalUP-proxin

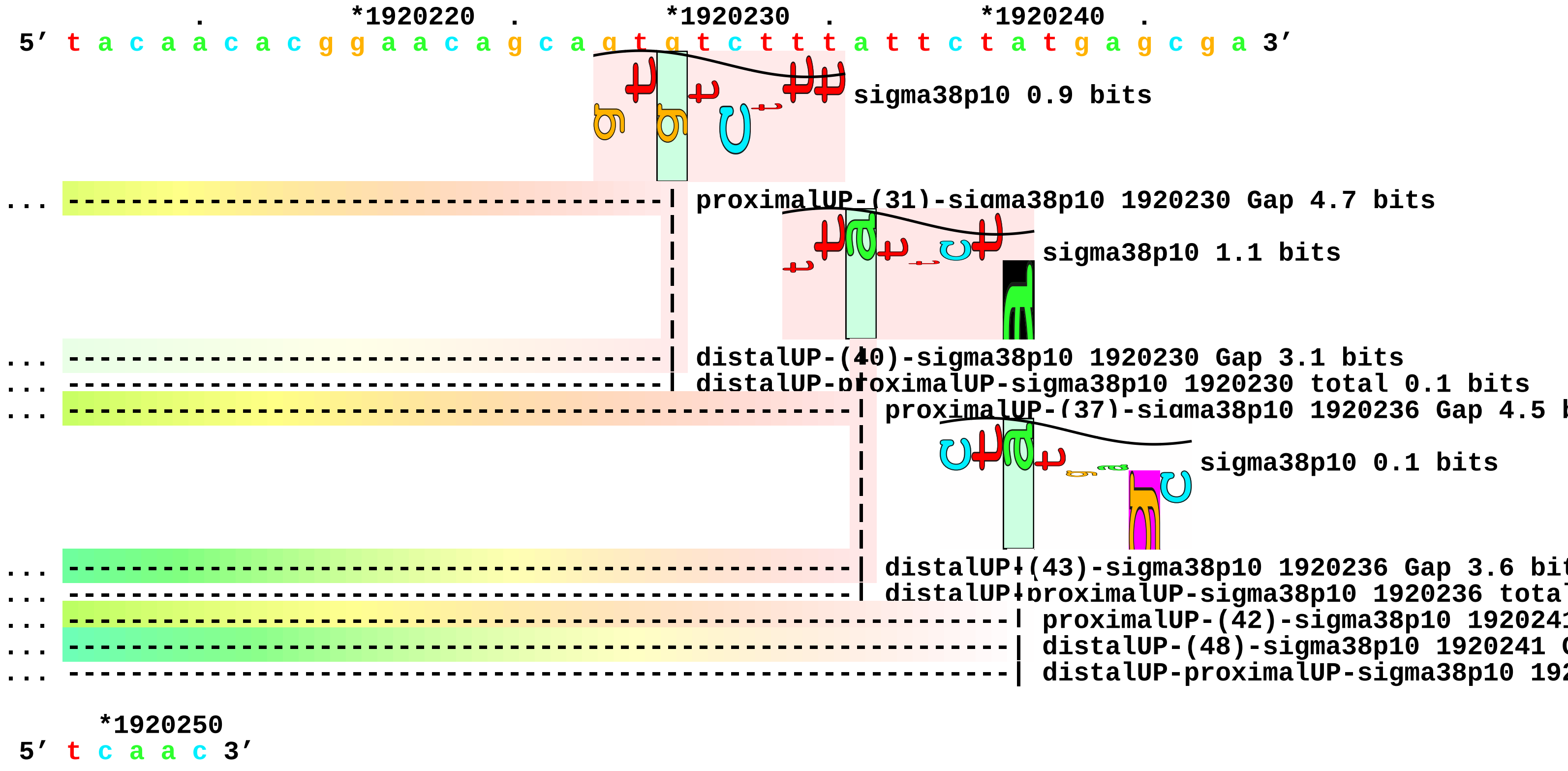
*1920220
*1920230
*1920240
5′ t a c a a c a c g g a a c a g c a g t g t c t t t a t t c t a t g a g c g a 3′
sigma38p10 0.9 bits
proximalUP-(31)-sigma38p10 1920230 Gap 4.7 bits
sigma38p10 1.1 bits
distalUP-(40)-sigma38p10 1920230 Gap 3.1 bits
distalUP-proximalUP-sigma38p10 1920230 total 0.1 bits
proximalUP-(37)-sigma38p10 1920236 Gap 4.5 b
sigma38p10 0.1 bits
distalUP-(43)-sigma38p10 1920236 Gap 3.6 bit
distalUP-proximalUP-sigma38p10 1920236 tota
proximalUP-(42)-sigma38p10 192024
distalUP-(48)-sigma38p10 1920241 (
distalUP-proximalUP-sigma38p10 192
*1920250
5′ t c a a c 3′

piece 25, NC_000913, ryeB, config: linear, direction: -, begin: 1921349, end: 1921100

. *1921340 . *1921330 . *1921320 .
5′ c c t c a g t t c t t t c a c c a g a a c g g g c g g t t t t t a a c a t t 3′

T timestamp:_2018Sep20_11-34-53

proximalUP 5.0 bits

|------- ... proximalUP-(36)

distalUP 4.9 bits

|------------------------- ... distalUP-(44)-s
|---------------|------- ... sigma38p10-prox

*1921310 . *1921300 . *1921290 . *1921280 .
5′ t c a g c t g a t g a c c a c c a c g c t t t t t a t t g a c c a t t t t g 3′

sigma38p10 3.3 bits

... ----------------------------------------------------------------| proximalUP-(36)-sigma38p10 192

proximalUP 6.1 bits

... ----------------------------------------------------------------| distalUP-(44)-sigma38p10 19212
... ----------------------------------------------------------------| sigma38p10-proximalUP-distalUP

distalUP 4.1 bits

|- ... proximalUP-(33)

|------------------------------------- ... distalUP-(51)-s
|-------------------------------------|- ... sigma38p10-prox

*1921270 . *1921260 . *1921250 . *1921240
5′ c a c g c a a a c t g g a a a a c c t g g c g t c g t c a t c t a t t c t t 3′

sigma38p10 9.8 bits

proximalUP 3.7 bits

|----- ... proximalUP-(36)

distalUP 0.3 bits

|---------------------- ... distalUP-(44)-s
... ----------------------------------------------------------------| proximalUP-(33)-sigma38p10 192

```
... ----------------------------------------------------------------| distalUP-(51)-sigma38p10 19212
... ----------------------------------------------------------------| sigma38p10-proximalUP-distalUP
                                                          |---------------|----- ... sigma38p10-prox

     .         *1921230   .         *1921220   .         *1921210   .         *1921200
5' a a a g g g c a a g g c a a c t a a g c c t g c a t t a a t g c c a a c t t 3'
```

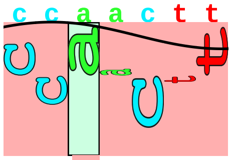

```
                                                                                  ... sigma38p10

... ------------------------------------------------------------------| proximalUP-(36)-sigma38p10 1
... ------------------------------------------------------------------| distalUP-(44)-sigma38p10 192

... ------------------------------------------------------------------| sigma38p10-proximalUP-dista

         .         *1921190   .         *1921180   .         *1921170   .         *1921160
5' t t a g c g c a c g g c t c t c t c c c a a g a g c c a t t t c c c t g g a 3'
```

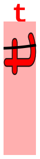

```
... sigma38p10 3.4 bits

             .         *1921150   .         *1921140   .         *1921130   .
5' c c g a a t a c a g g a a t c g t g t t c g g t c t c t t t t t a t c t g t 3'

      *1921120   .         *1921110   .         *1921100
5' t a a a a g c c a g a a g c a t t t c c t t 3'
```

piece 26, NC_000913, yodC, config: linear, direction: -, begin: 2026555, end: 2026334

```
   .         *2026550   .         *2026540   .         *2026530   .         *2026520
5' a t c t g a t t c t t g g a t t t a t c a t c g g c g g c a t c g t g g t g 3'
```

T timestamp:_2018Sep20_11-34-53

```
     .         *2026510   .         *2026500   .         *2026490   .         *2026480
5' c t g g g a g t g a a a g c c g t a g a g a a a a t g c g c g g t c a g g c 3'
```

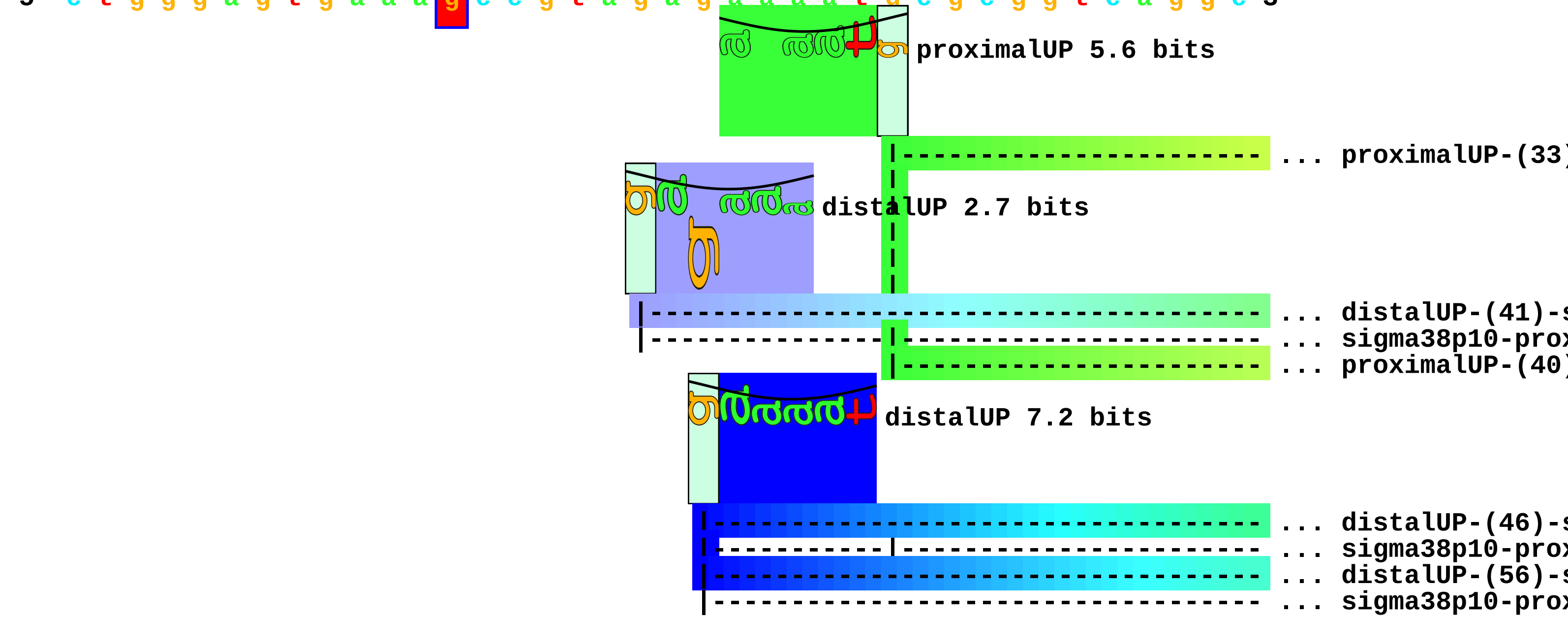


```
         .         *2026470   .         *2026460   .         *2026450   .
5' a c a t t a a t a c t c g c a c t g g c t a t t a t g c a a a a t t c g c c 3'
```

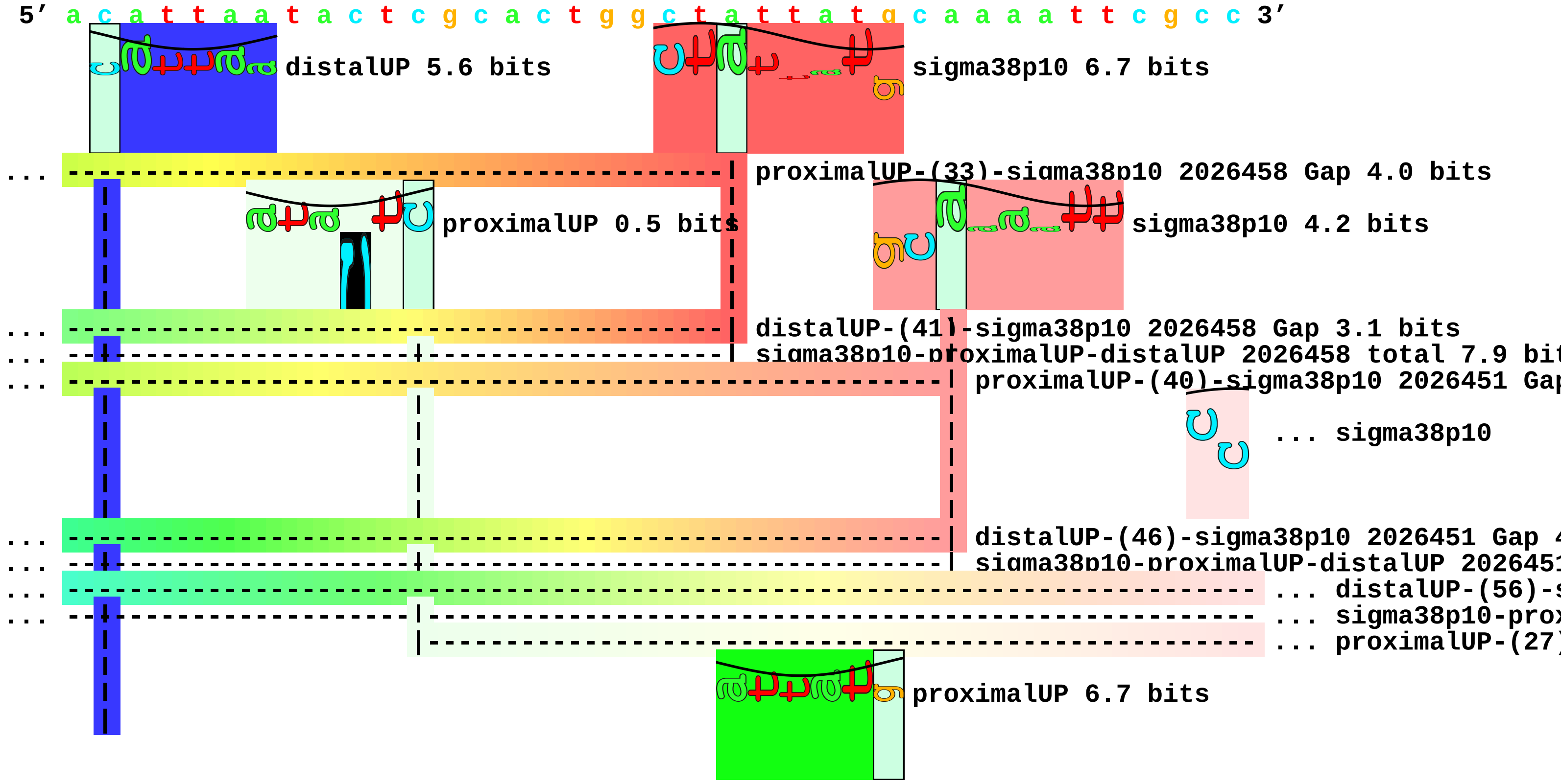

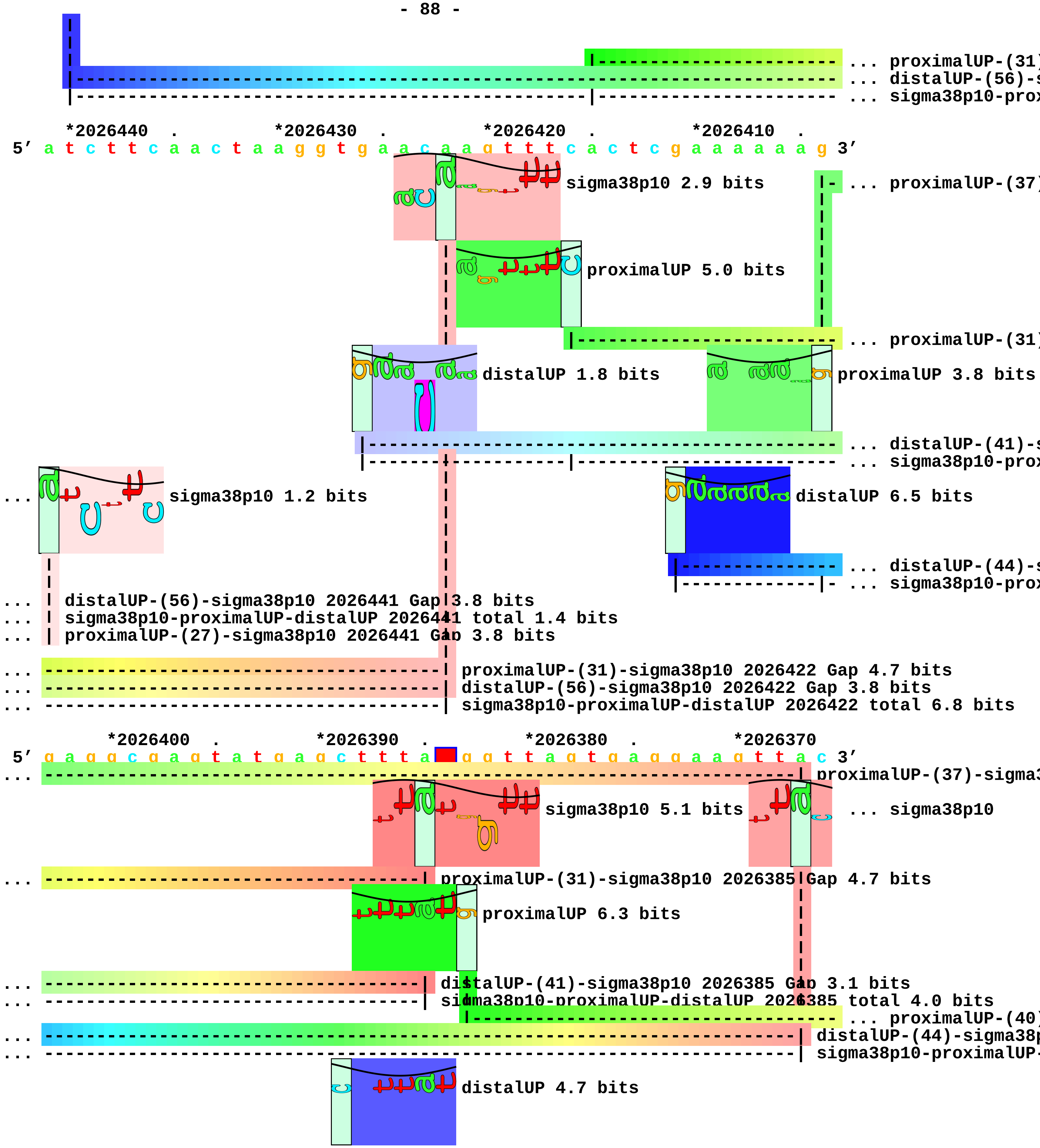
... proximalUP-(31)
... distalUP-(56)-s
... sigma38p10-prox
*2026440 . *2026430 . *2026420 . *2026410 .
5′ a t c t t c a a c t a a g g t g a a c a a g t t t c a c t c g a a a a a a g 3′
sigma38p10 2.9 bits
... proximalUP-(37)
proximalUP 5.0 bits
... proximalUP-(31)
distalUP 1.8 bits
proximalUP 3.8 bits
... distalUP-(41)-s
... sigma38p10-prox
... sigma38p10 1.2 bits
distalUP 6.5 bits
... distalUP-(44)-s
... sigma38p10-prox
... distalUP-(56)-sigma38p10 2026441 Gap 3.8 bits
... sigma38p10-proximalUP-distalUP 2026441 total 1.4 bits
... proximalUP-(27)-sigma38p10 2026441 Gap 3.8 bits
... proximalUP-(31)-sigma38p10 2026422 Gap 4.7 bits
... distalUP-(56)-sigma38p10 2026422 Gap 3.8 bits
... sigma38p10-proximalUP-distalUP 2026422 total 6.8 bits
*2026400 . *2026390 . *2026380 . *2026370
5′ g a g g c g a g t a t g a g c t t t a g g t t a g t g a g g a a g t t a c 3′
... proximalUP-(37)-sigma3
sigma38p10 5.1 bits
... sigma38p10
... proximalUP-(31)-sigma38p10 2026385 Gap 4.7 bits
proximalUP 6.3 bits
... distalUP-(41)-sigma38p10 2026385 Gap 3.1 bits
... sigma38p10-proximalUP-distalUP 2026385 total 4.0 bits
... proximalUP-(40)
... distalUP-(44)-sigma38p
... sigma38p10-proximalUP-
distalUP 4.7 bits

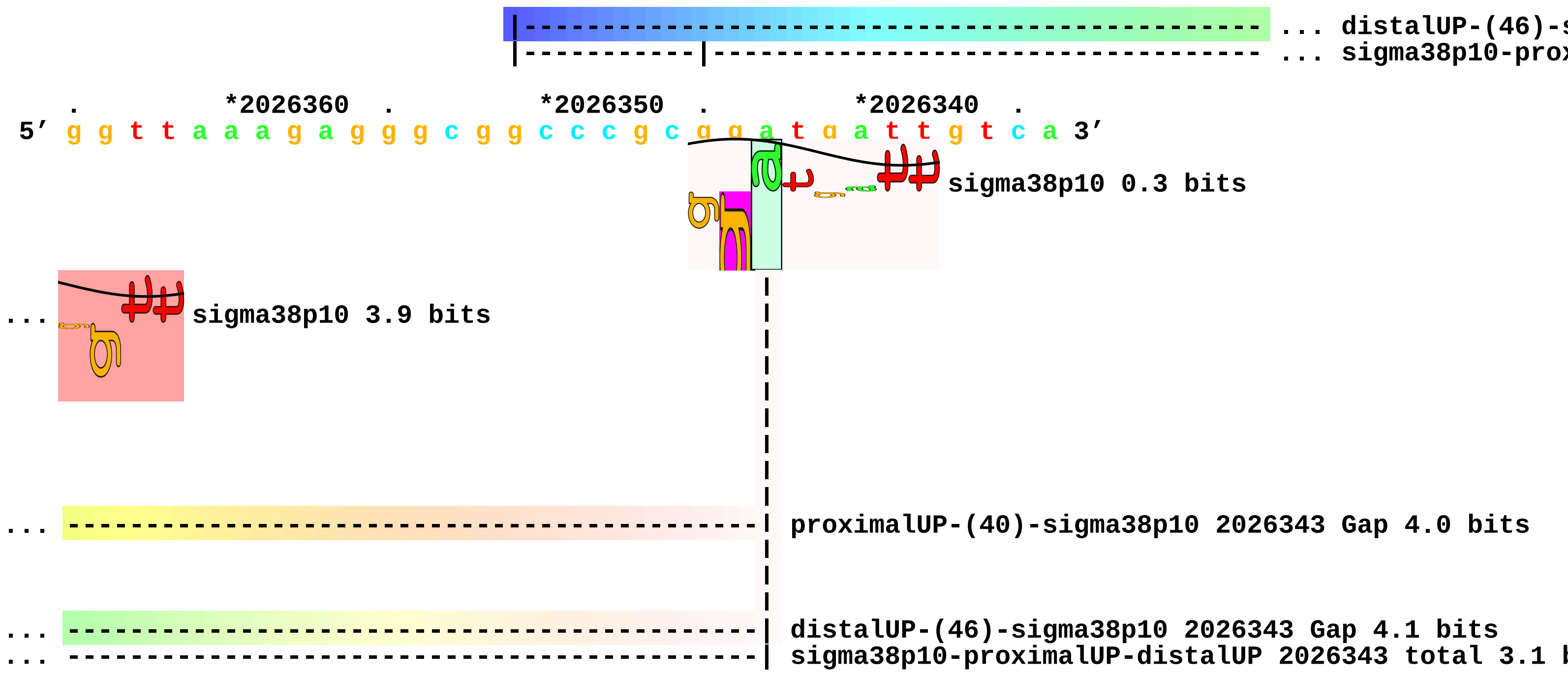
... distalUP-(46)-s
... sigma38p10-prox
*2026360
*2026350
*2026340
5′ g g t t a a a g a g g g c g g c c c g c g g a t g a t t g t c a 3′
sigma38p10 0.3 bits
... sigma38p10 3.9 bits
... proximalUP-(40)-sigma38p10 2026343 Gap 4.0 bits
... distalUP-(46)-sigma38p10 2026343 Gap 4.1 bits
... sigma38p10-proximalUP-distalUP 2026343 total 3.1 b

piece 27, NC_000913, erfK, config: linear, direction: -, begin: 2061534, end: 2061211

```
             *2061530   .          *2061520   .          *2061510   .          *2061500
5' a g t g g c g c a g c t c t g g c g a t g c c c a t c a t c g a a g c t g c 3'

   T timestamp:_2018Sep20_11-34-53                                proximalUP 3.7 bits

                                                                  |---------------- ... proximalUP-(34)
                                                                distalUP 1.5 bits

                                                          |------------------------------ ... distalUP-(40)-s
                                                          |-----------|------------------ ... sigma38p10-prox
                                                          |------------------------------ ... distalUP-(55)-s
                                                          |------------------------------ ... sigma38p10-prox

    .         *2061490   .          *2061480   .          *2061470   .          *2061460
5' c t g t g c g a t a t a c a a c a a c a t g g g c g a a c t t g c t g c c a 3'
                          proximalUP 4.9 bits                 sigma38p10 1.4 bits
... -----------------------------------------------------| proximalUP-(34)-sigma38p10 2061471 Gap 2.5 b
                        |------------------------------------------------ ... proximalUP-(28)
... -----------------------------------------------------| distalUP-(40)-sigma38p10 2061471 Gap 3.1 bit
... -----------------------------------------------------| sigma38p10-proximalUP-distalUP 2061471 tota
... --------------------------------------------------------------------- ... distalUP-(55)-s
... -----------------------|--------------------------------------------- ... sigma38p10-prox

        .         *2061450   .          *2061440   .          *2061430   .
5' g t a a t a t t g t t c t a c c g g g a a t a c g a c t t c t g a t t t g 3'
                  sigma38p10 6.0 bits

                    proximalUP 6.1 bits

... ----| proximalUP|(28)-sigma38p10 2061456 Gap 5.5 bits
                  |------------------------------------------------------------ ... proximalUP-(37)
                distalUP 4.4 bits

... ----| distalUP-(55)-sigma38p10 2061456 Gap 4.9 bits
... |---| sigma38p10-proximalUP-distalUP 2061456 total 2.0 bits
    |---------------------------------------------------------------------- ... distalUP-(45)-s
    |---------------|------------------------------------------------------ ... sigma38p10-prox

    *2061420   .          *2061410   .          *2061400   .          *2061390   .
5' a a c a g t t a a a a a c g a a a a g t c t g a t a a a a g c t a t a c t t 3'
```

sigma38p10 2.0 bits

proximalUP 4.3 bits

... proximalUP-(33

... proximalUP-(37)-sigma38p10 2061413 Gap 4.5 bits

distalUP 4.9 bits

... distalUP-(41)-s

... sigma38p10-prox

... distalUP-(45)-sigma38p10 2061413 Gap 5.5 bits

... sigma38p10-proximalUP-distalUP 2061413 total 2.5 bits

*2061380 . *2061370 . *2061360 . *2061350 .

5′ a a c g g a t a g c t t t c g c g a c a t a g g a a a g g g a c a t g a t g 3′

sigma38p10 1.0 bits

... proximalUP-(33)-sigma38p10 2061375 Gap 4.0 bits

... distalUP-(41)-sigma38p10 2061375 Gap 3.1 bits

... sigma38p10-proximalUP-distalUP 2061375 total 3.0 bits

*2061340 . *2061330 . *2061320 . *2061310

5′ c g t c g t g t a a a t a t t c t t t g c t c a t t t g c t c t g c t t t t 3′

proximalUP 6.6 bits

... proximalUP-(31

distalUP 5.6 bits

... distalUP-(40)-s

... sigma38p10-prox

... proximalUP-(36

... distalUP-(45)-s

... sigma38p10-prox

. *2061300 . *2061290 . *2061280 . *2061270

5′ t g c c a g c c a t a c t a g c c t g g c g g t a a c t t a t c c a t t a c 3′

sigma38p10 1.0 bits

... proximalUP-(31)-sigma38p10 2061298 Gap 4.7 bits

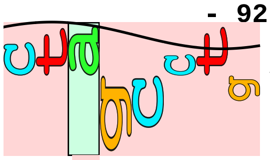


sigma38p10 1.7 bits

```
... ----------------| distalUP-(40)-sigma38p10 2061298 Gap 3.1 bits
... ----------------| sigma38p10-proximalUP-distalUP 2061298 total 5.5 bits
... -------------------------| proximalUP-(36)-sigma38p10 2061293 Gap 3.6 bits
... -------------------------| distalUP-(45)-sigma38p10 2061293 Gap 5.5 bits
... -------------------------| sigma38p10-proximalUP-distalUP 2061293 total 4.9 bits

          .         *2061260  .         *2061250  .         *2061240  .
 5' c t c c a g a g g g t a g c c g t t t a g t g g g g c a g t c g t t t a c t 3'

    *2061230  .         *2061220  .
 5' g t a a c t g t t c c t g a t c a c a a 3'
```

piece 28, NC_000913, wbbI, config: linear, direction: -, begin: 2104249, end: 2103800

```
            .         *2104240  .         *2104230  .         *2104220  .
5' g t c t c g a a a t t a c a t a t c t g t g a a a a t a t t t t a t t c a t 3'
```

proximalUP 3.7 bits

proximalUP 1.8 bits

... proximalUP-(35)

distalUP 6.6 bits

... proximalUP-(27)

... distalUP-(42)-s

... sigma38p10-prox

... proximalUP-(37)

... distalUP-(44)-s

... sigma38p10-prox

proximalUP 3.2 bits

proximalUP 6.2 bits

T timestamp: 2018Sep20 11-34-53

... proximalUP-(36)

... distalUP-(48)-s

... sigma38p10-prox

distalUP 7.2 bits

... distalUP-(40)-s

... sigma38p10-prox

... proximalUP-(33)

... distalUP-(47)-s

... sigma38p10-prox

... distalUP

... distalUP-(53)-s

... sigma38p10-prox

```
     *2104210  .         *2104200  .         *2104190  .         *2104180  .
5' a t t t t a t t t a t a c c t t t t c t t t c a t t t t t t a t c a t g a a 3'
```

sigma38p10 2.0 bits

sigma38p10 3.2 bits

... proximalUP-(35)-sigma38p10 2104202 Gap 3.1 bits

... proximalUP-(27)-sigma38p10 2104188 Gap 3.8 bits

... distalUP-(42)-sigma38p10 2104202 Gap 3.3 bits

... sigma38p10-proximalUP-distalUP 2104202 total 6.0 bits

... proximalUP-(37)-sigma38p10 2104200 Gap 4.5 bits

... distalUP-(44)-sigma38p10 2104200 Gap 2.8 bits

... sigma38p10-proximalUP-distalUP 2104200 total 6.5 bits

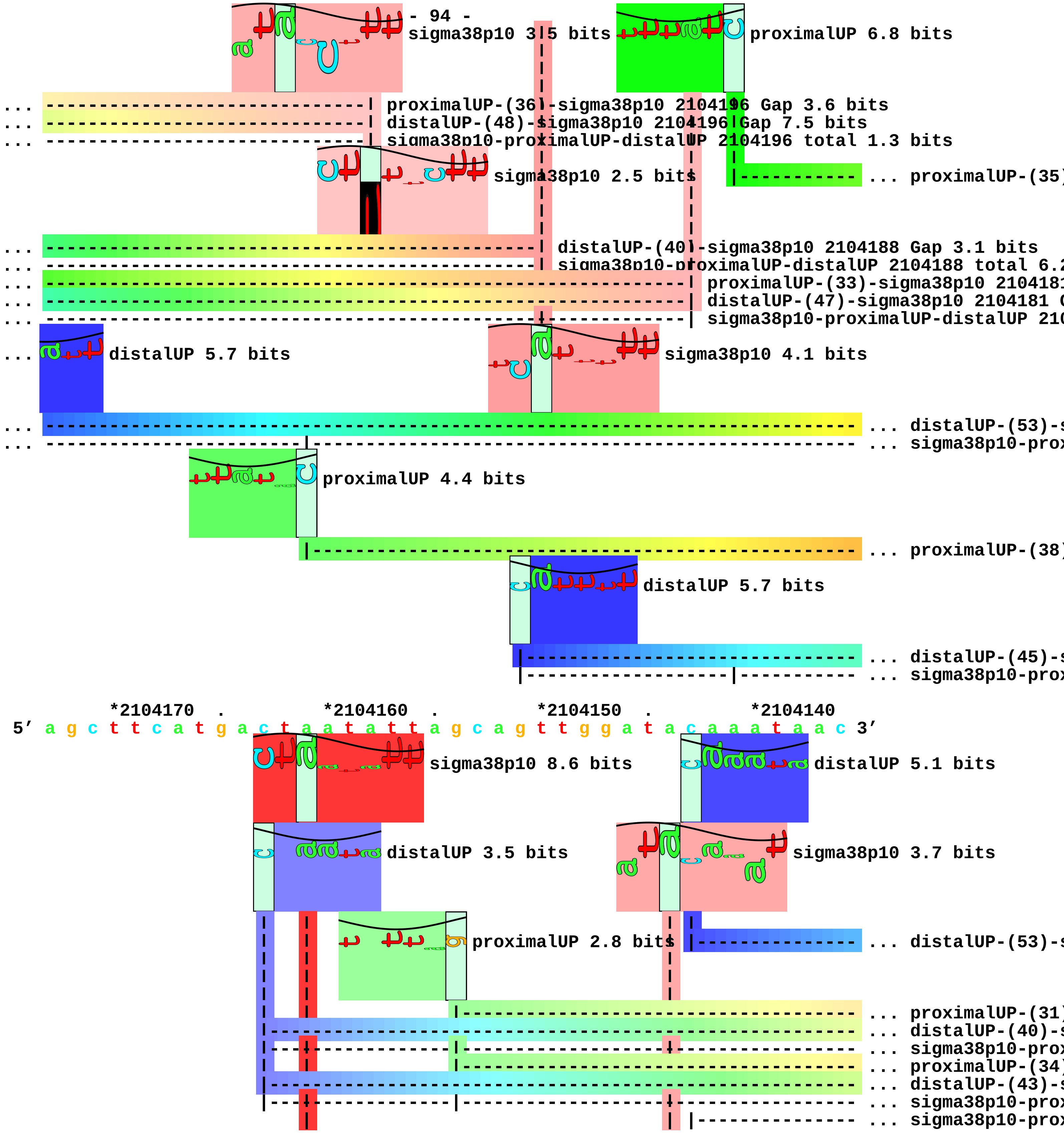
sigma38p10 3.5 bits
proximalUP 6.8 bits
proximalUP-(36)-sigma38p10 2104196 Gap 3.6 bits
distalUP-(48)-sigma38p10 2104196 Gap 7.5 bits
sigma38p10-proximalUP-distalUP 2104196 total 1.3 bits
sigma38p10 2.5 bits
proximalUP-(35)
distalUP-(40)-sigma38p10 2104188 Gap 3.1 bits
sigma38p10-proximalUP-distalUP 2104188 total 6.2
proximalUP-(33)-sigma38p10 2104181
distalUP-(47)-sigma38p10 2104181 G
sigma38p10-proximalUP-distalUP 210
distalUP 5.7 bits
sigma38p10 4.1 bits
distalUP-(53)-s
sigma38p10-prox
proximalUP 4.4 bits
proximalUP-(38)
distalUP 5.7 bits
distalUP-(45)-s
sigma38p10-prox
*2104170 . *2104160 . *2104150 . *2104140
5′ a g c t t c a t g a c t a a t a t t a g c a g t t g g a t a c a a a t a a c 3′
sigma38p10 8.6 bits
distalUP 5.1 bits
distalUP 3.5 bits
sigma38p10 3.7 bits
proximalUP 2.8 bits
distalUP-(53)-s
proximalUP-(31)
distalUP-(40)-s
sigma38p10-prox
proximalUP-(34)
distalUP-(43)-s
sigma38p10-prox
sigma38p10-prox

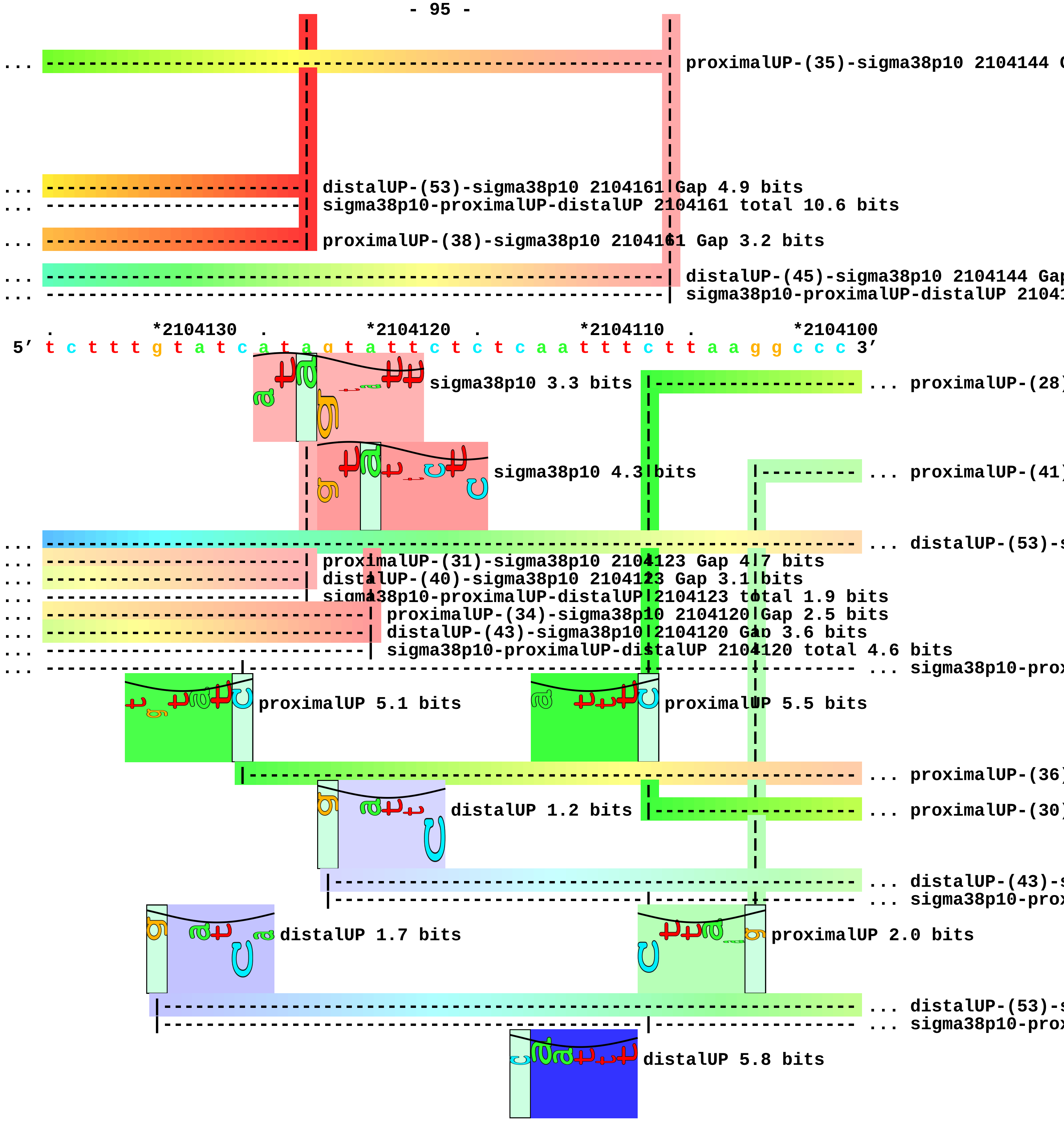
proximalUP-(35)-sigma38p10 2104144 G
distalUP-(53)-sigma38p10 2104161 Gap 4.9 bits
sigma38p10-proximalUP-distalUP 2104161 total 10.6 bits
proximalUP-(38)-sigma38p10 2104161 Gap 3.2 bits
distalUP-(45)-sigma38p10 2104144 Gap
sigma38p10-proximalUP-distalUP 21041
*2104130
*2104120
*2104110
*2104100
5' t c t t t g t a t c a t a g t a t t c t c t c a a t t t c t t a a g g c c c 3'
sigma38p10 3.3 bits
proximalUP-(28)
sigma38p10 4.3 bits
proximalUP-(41)
distalUP-(53)-s
proximalUP-(31)-sigma38p10 2104123 Gap 4.7 bits
distalUP-(40)-sigma38p10 2104123 Gap 3.1 bits
sigma38p10-proximalUP-distalUP 2104123 total 1.9 bits
proximalUP-(34)-sigma38p10 2104120 Gap 2.5 bits
distalUP-(43)-sigma38p10 2104120 Gap 3.6 bits
sigma38p10-proximalUP-distalUP 2104120 total 4.6 bits
sigma38p10-prox
proximalUP 5.1 bits
proximalUP 5.5 bits
proximalUP-(36)
distalUP 1.2 bits
proximalUP-(30)
distalUP-(43)-s
sigma38p10-prox
distalUP 1.7 bits
proximalUP 2.0 bits
distalUP-(53)-s
sigma38p10-prox
distalUP 5.8 bits

... distalUP-(52)-s
... sigma38p10-prox

. *2104090 . *2104080 . *2104070 . *2104060

5′ a g a a a a t a a a g t g a a a a t g t a t t t t t t g a a t g a t t t a a 3′

... proximalUP-(28)-sigma38p10 2104079 Gap 5.5 bits
... proximalUP-(41)-sigma3
... distalUP-(53)-sigma38p10 2104090 Gap 4.9 bits
... sigma38p10
sigma38p10 2.6 bits
... proximalUP
sigma38p10 4.2 bits
sigma38p10 6.0 bits
proximalUP 5.6 bits
distalUP 5.8 bits
... proximalUP-(35)
distalUP 7.2 bits
... distalUP-(48)-s
... sigma38p10-proximalUP-distalUP 2104090 total 4.4 bits
... distalUP-(41)-s
... proximalUP-(36)-sigma38p10 2104090 Gap 3.6 bits
... proximalUP-(30)-sigma38p10 2104077 Gap 5.9 bits
... distalUP-(43)-sigma38p10 2104079 Gap 3.6 bits
... sigma38p10-proximalUP-distalUP 2104079 total 1.9 bits
... sigma38p10-prox
... distalUP-(53)-sigma38p10 2104077 Gap 4.9 bits
... sigma38p10-proximalUP-distalUP 2104077 total 2.5 bits
... proximalUP-(40)
... distalUP-(52)-sigma38p
... sigma38p10-proximalUP-
... distalUP-(46)-s
... sigma38p10-prox
... sigma38p10-prox
... distalUP-(53)-s
... sigma38p10-prox

. *2104050 . *2104040 . *2104030 .

5′ a t t t c t c t a g a c g c g a t g c t g g a t t t a a a g c a a g a a a a 3′

sigma38p10 1.0 bits

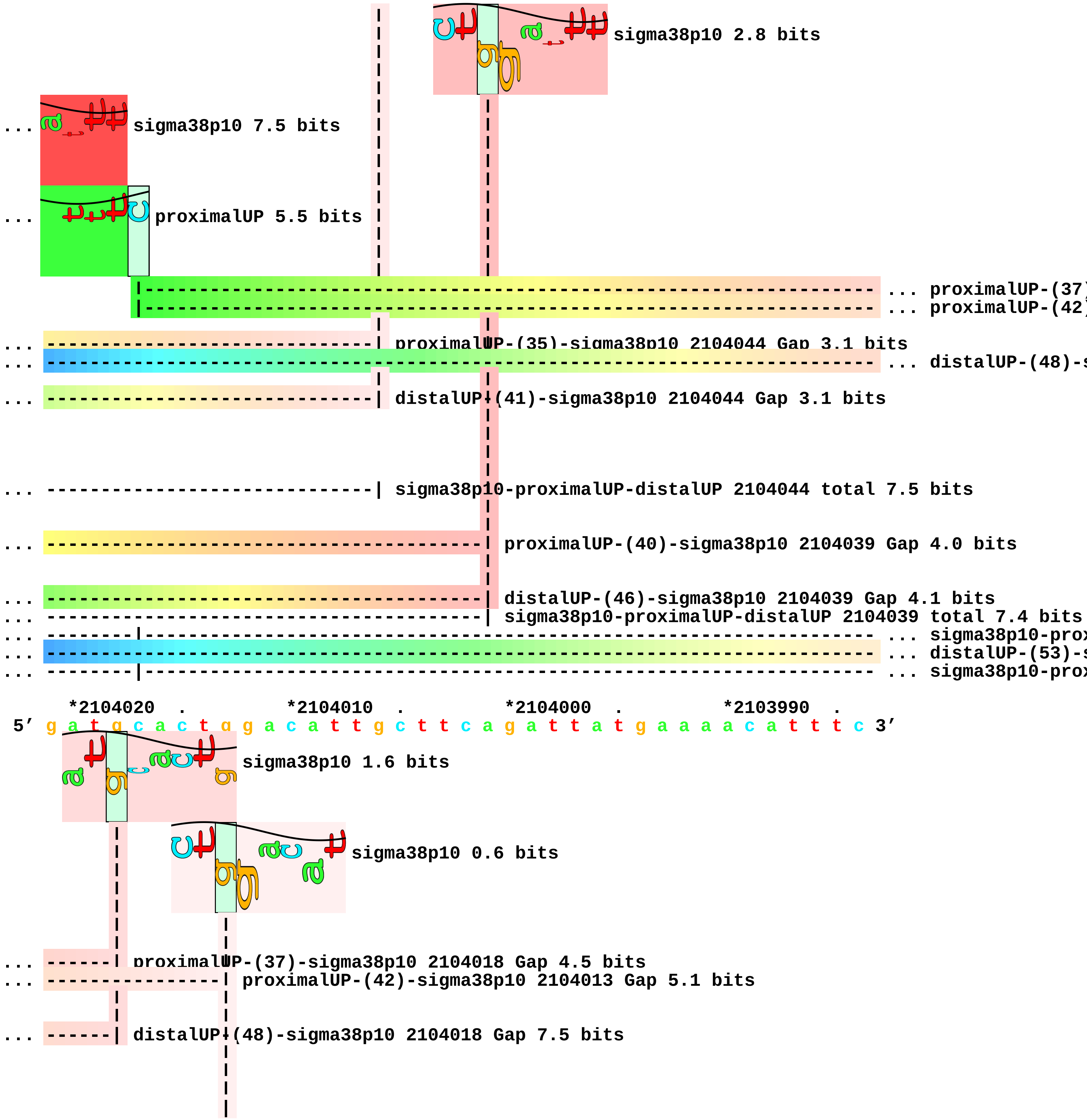
sigma38p10 2.8 bits
... sigma38p10 7.5 bits
... proximalUP 5.5 bits
... proximalUP-(37
... proximalUP-(42
... ------------------------------| proximalUP-(35)-sigma38p10 2104044 Gap 3.1 bits
... distalUP-(48)-s
... ------------------------------| distalUP-(41)-sigma38p10 2104044 Gap 3.1 bits
... ------------------------------| sigma38p10-proximalUP-distalUP 2104044 total 7.5 bits
... ----------------------------------------| proximalUP-(40)-sigma38p10 2104039 Gap 4.0 bits
... ----------------------------------------| distalUP-(46)-sigma38p10 2104039 Gap 4.1 bits
... ----------------------------------------| sigma38p10-proximalUP-distalUP 2104039 total 7.4 bits
... sigma38p10-prox
... distalUP-(53)-s
... sigma38p10-prox
*2104020 . *2104010 . *2104000 . *2103990 .
5′ g a t g c a c t g g a c a t t g c t t c a g a t t a t g a a a a c a t t t c 3′
sigma38p10 1.6 bits
sigma38p10 0.6 bits
... ------| proximalUP-(37)-sigma38p10 2104018 Gap 4.5 bits
... ----------------| proximalUP-(42)-sigma38p10 2104013 Gap 5.1 bits
... ------| distalUP-(48)-sigma38p10 2104018 Gap 7.5 bits

```
... ------| sigma38p10-proximalUP-distalUP 2104018 total 1.0 bits
... ----------------| distalUP-(53)-sigma38p10 2104013 Gap 4.9 bits
... ----------------| sigma38p10-proximalUP-distalUP 2104013 total 2.0 bits

       *2103980    .         *2103970    .         *2103960    .         *2103950
5' t g t t g t t a a c a t t c c t c t a t g g g g t g g a g t a g t c c a g a 3'

    .         *2103940    .         *2103930    .         *2103920    .         *2103910
5' g a a t t a t t a g t t c t g t t a a g c t t a g t a c a t t t c t c t g c 3'
                                   proximalUP 2.8 bits
                   |--------------------------------------------------------- ... proximalUP-(35)
                   distalUP 5.9 bits
   |------------------------------------------------------------------------- ... distalUP-(44)-s
   |-----------------|------------------------------------------------------- ... sigma38p10-prox
                     |------------------------------------------------------- ... proximalUP-(41)
   |------------------------------------------------------------------------- ... distalUP-(50)-s
   |-----------------|------------------------------------------------------- ... sigma38p10-prox

        .         *2103900    .         *2103890    .         *2103880    .         *2103870
5' g g t c t t g a a a a t a a a g a t g t t t t a a t t t t c a a t t t c c c 3'
                      sigma38p10 1.5 bits                     proximalUP 6.6 bits
... ------------| proximalUP-(35)-sigma38p10 2103901 Gap 3.1 bits     |----- ... proximalUP-(33)
                             sigma38p10 1.9 bits                  |---------------- ... proximalUP-(31)
... ------------| distalUP-(44)-sigma38p10 2103901 Gap 2.8 bits       proximalUP 5.5 bits
... ------------| sigma38p10-proximalUP-distalUP 2103901 total 4.3 bits
... ------------------------| proximalUP-(41)-sigma38p10 2103895 Gap 5.5 bits
... ------------------------| distalUP-(50)-sigma38p10 2103895 Gap 4.1 bits
... ------------------------| sigma38p10-proximalUP-distalUP 2103895 total 1.1 bits
                            proximalUP 4.9 bits                  |---------------- ... proximalUP-(33)
```

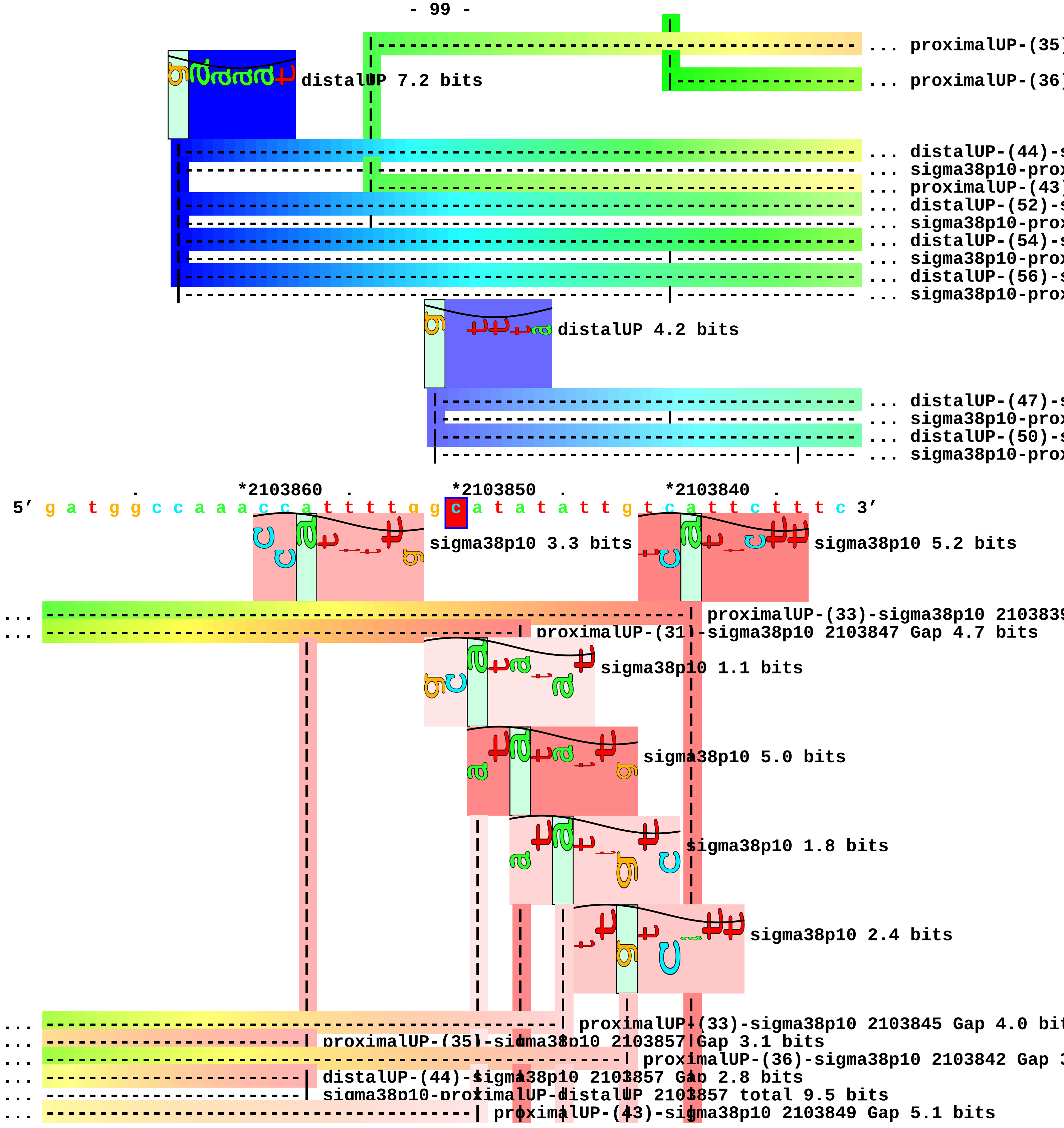
... proximalUP-(35)
... proximalUP-(36)
distalUP 7.2 bits
... distalUP-(44)-s
... sigma38p10-prox
... proximalUP-(43)
... distalUP-(52)-s
... sigma38p10-prox
... distalUP-(54)-s
... sigma38p10-prox
... distalUP-(56)-s
... sigma38p10-prox
distalUP 4.2 bits
... distalUP-(47)-s
... sigma38p10-prox
... distalUP-(50)-s
... sigma38p10-prox
*2103860
*2103850
*2103840
5′ g a t g g c c a a a c c a t t t t g g c a t a t a t t g t c a t t c t t t c 3′
sigma38p10 3.3 bits
sigma38p10 5.2 bits
... proximalUP-(33)-sigma38p10 2103839
... proximalUP-(31)-sigma38p10 2103847 Gap 4.7 bits
sigma38p10 1.1 bits
sigma38p10 5.0 bits
sigma38p10 1.8 bits
sigma38p10 2.4 bits
... proximalUP-(33)-sigma38p10 2103845 Gap 4.0 bit
... proximalUP-(35)-sigma38p10 2103857 Gap 3.1 bits
... proximalUP-(36)-sigma38p10 2103842 Gap 3
... distalUP-(44)-sigma38p10 2103857 Gap 2.8 bits
... sigma38p10-proximalUP-distalUP 2103857 total 9.5 bits
... proximalUP-(43)-sigma38p10 2103849 Gap 5.1 bits

```
                                         - 100 -
... ----------------------------------------| distalUP-(52)-sigma38p10 2103849 Gap 5.5 bits
... ----------------------------------------| sigma38p10-proximalUP-distalUP 2103849 total 2.6 bits
... --------------------------------------------| distalUP-(54)-sigma38p10 2103847 Gap 4.9 bits
... --------------------------------------------| sigma38p10-proximalUP-distalUP 2103847 total 9.3 b
... ------------------------------------------------| distalUP-(56)-sigma38p10 2103845 Gap 3.8 bits
... ------------------------------------------------| sigma38p10-proximalUP-distalUP 2103845 total 7
                                                          |     |
... ------------------------------------------------------| distalUP-(47)-sigma38p10 2103842 Gap 6.5
... ------------------------------------------------------| sigma38p10-proximalUP-distalUP 2103842 t
... ------------------------------------------------------------| distalUP-(50)-sigma38p10 2103839 G
... ------------------------------------------------------------| sigma38p10-proximalUP-distalUP 210

      *2103830  .         *2103820  .         *2103810  .         *2103800
 5' a c c g c c t t c t a a a a t t t a g a a t a g t a c c t c t g 3'
```

piece 29, NC_000913, wbbH, config: linear, direction: -, begin: 2105649, end: 2104500

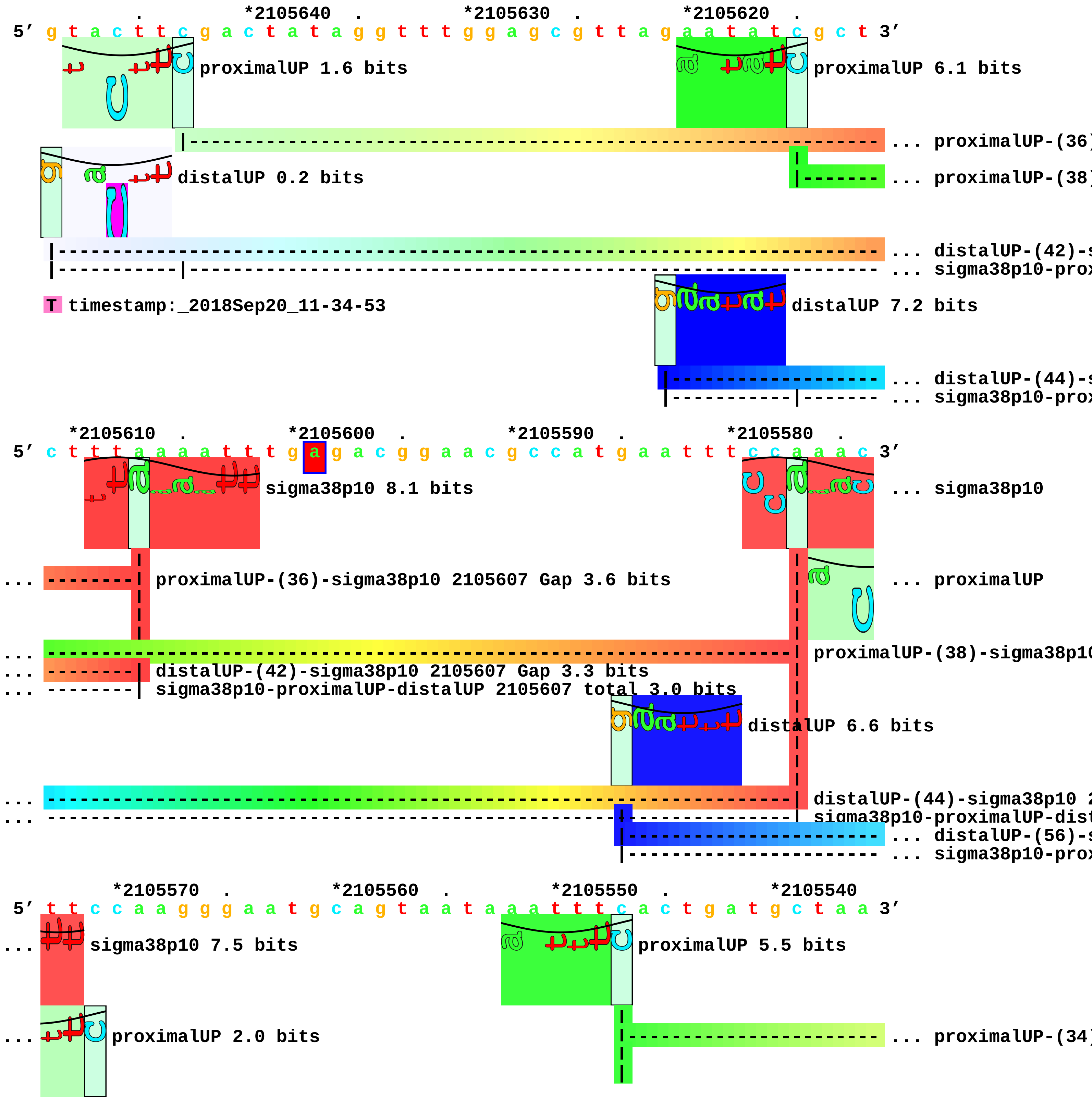

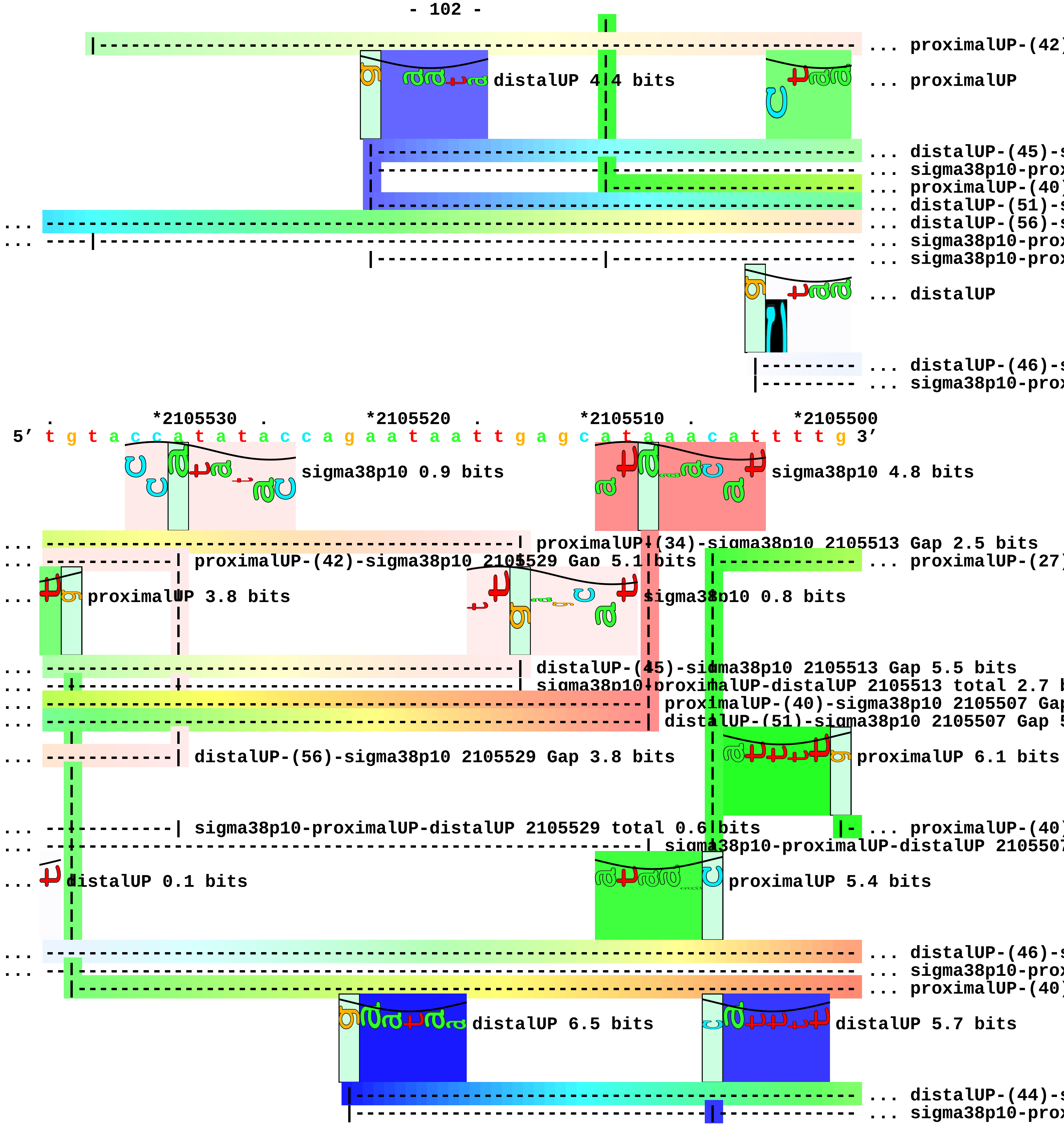
... proximalUP-(42)
distalUP 4.4 bits
... proximalUP
... distalUP-(45)-s
... sigma38p10-prox
... proximalUP-(40)
... distalUP-(51)-s
... distalUP-(56)-s
... sigma38p10-prox
... sigma38p10-prox
... distalUP
... distalUP-(46)-s
... sigma38p10-prox
*2105530
*2105520
*2105510
*2105500
5′ t g t a c c a t a t a c c a g a a t a a t t g a g c a t a a a c a t t t t g 3′
sigma38p10 0.9 bits
sigma38p10 4.8 bits
proximalUP-(34)-sigma38p10 2105513 Gap 2.5 bits
proximalUP-(42)-sigma38p10 2105529 Gap 5.1 bits
... proximalUP-(27)
proximalUP 3.8 bits
sigma38p10 0.8 bits
distalUP-(45)-sigma38p10 2105513 Gap 5.5 bits
sigma38p10-proximalUP-distalUP 2105513 total 2.7 b
proximalUP-(40)-sigma38p10 2105507 Gap
distalUP-(51)-sigma38p10 2105507 Gap 5
distalUP-(56)-sigma38p10 2105529 Gap 3.8 bits
proximalUP 6.1 bits
sigma38p10-proximalUP-distalUP 2105529 total 0.6 bits
... proximalUP-(40)
sigma38p10-proximalUP-distalUP 210550
distalUP 0.1 bits
proximalUP 5.4 bits
... distalUP-(46)-s
... sigma38p10-prox
... proximalUP-(40)
distalUP 6.5 bits
distalUP 5.7 bits
... distalUP-(44)-s
... sigma38p10-prox

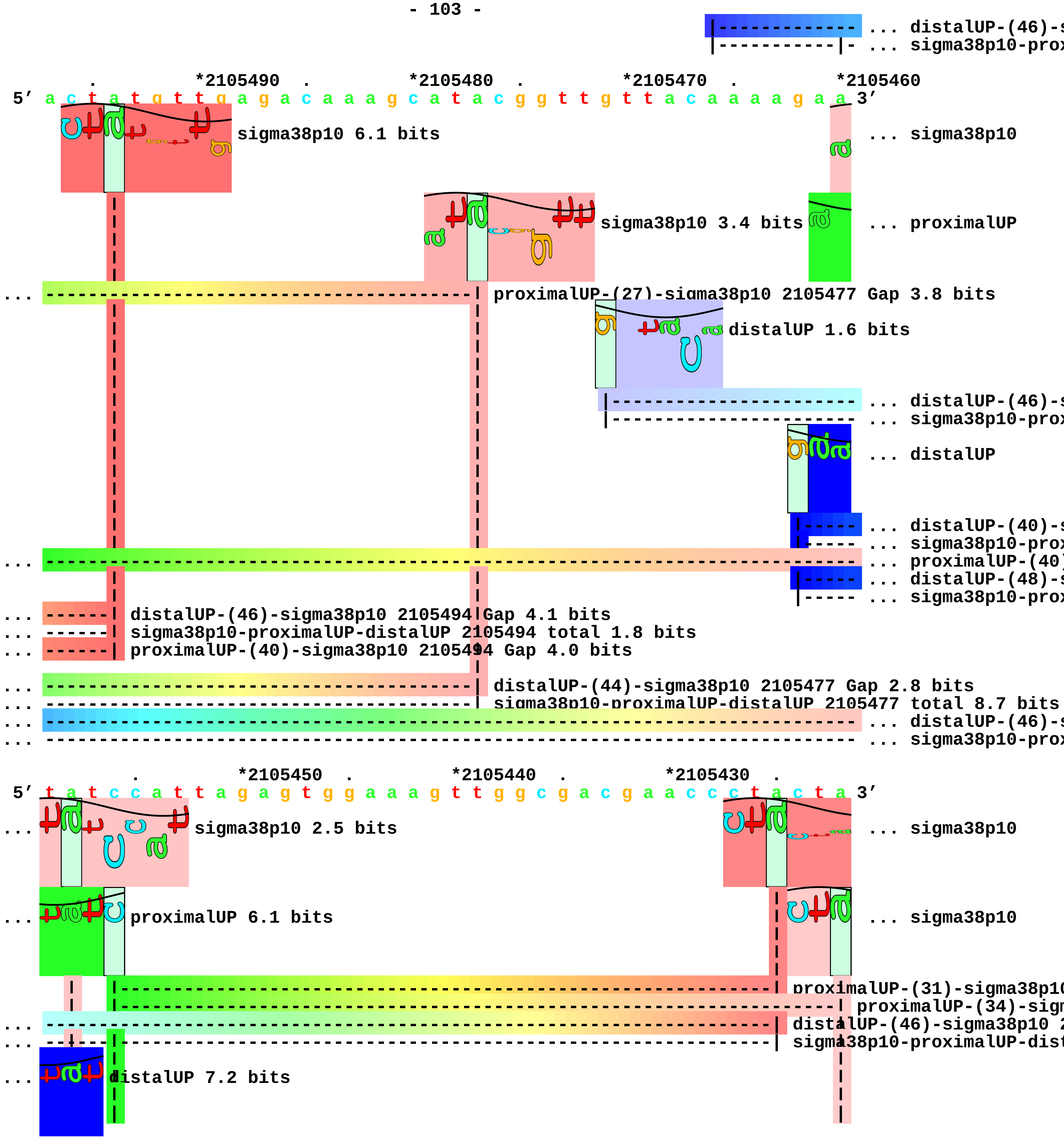
... distalUP-(46)-s
... sigma38p10-prox
*2105490
*2105480
*2105470
*2105460
5′ a c t a t g t t g a g a c a a a g c a t a c g g t t g t t a c a a a a g a a 3′
sigma38p10 6.1 bits
... sigma38p10
sigma38p10 3.4 bits
... proximalUP
... proximalUP-(27)-sigma38p10 2105477 Gap 3.8 bits
distalUP 1.6 bits
... distalUP-(46)-s
... sigma38p10-prox
... distalUP
... distalUP-(40)-s
... sigma38p10-prox
... proximalUP-(40)
... distalUP-(48)-s
... sigma38p10-prox
... distalUP-(46)-sigma38p10 2105494 Gap 4.1 bits
... sigma38p10-proximalUP-distalUP 2105494 total 1.8 bits
... proximalUP-(40)-sigma38p10 2105494 Gap 4.0 bits
... distalUP-(44)-sigma38p10 2105477 Gap 2.8 bits
... sigma38p10-proximalUP-distalUP 2105477 total 8.7 bits
... distalUP-(46)-s
... sigma38p10-prox
*2105450
*2105440
*2105430
5′ t a t c c a t t a g a g t g g a a a g t t g g c g a c g a a c c c t a c t a 3′
... sigma38p10 2.5 bits
... sigma38p10
... proximalUP 6.1 bits
... sigma38p10
proximalUP-(31)-sigma38p10
proximalUP-(34)-sigm
... distalUP-(46)-sigma38p10 2
... sigma38p10-proximalUP-dist
... distalUP 7.2 bits

... ---------------------------------------------------------------------| distalUP-(40)-sigma3
... --|-----------------------------------------------------------------| sigma38p10-proximalU
... --| proximalUP-(40)-sigma38p10 2105458 Gap 4.0 bits
... ------------------------------------------------------------------------ ... distalUP-(48)-s
... ------------------------------------------------------------------------ ... sigma38p10-prox
|------------------------------------------------------------------ ... proximalUP-(42)
... --| distalUP-(46)-sigma38p10 2105458 Gap 4.1 bits
... --| sigma38p10-proximalUP-distalUP 2105458 total 6.2 bits

*2105420 . *2105410 . *2105400 . *2105390 .
5' t c c a g t t a a t g a t a a t a a a a a c a t g g a g c t t t t t a a g a 3'

... sigma38p10 5.2 bits
proximalUP 4.5 bits
... sigma38p10 2.2 bits
|--- ... proximalUP-(37)
sigma38p10 1.4 bits
distalUP 4.1 bits
|------------------- ... distalUP-(45)-s
|---------------|--- ... sigma38p10-prox

... --------------| distalUP-(48)-sigma38p10 2105414 Gap 7.5 bits
... --------------| sigma38p10-proximalUP-distalUP 2105414 total 2.0 bits
... --------------| proximalUP-(42)-sigma38p10 2105414 Gap 5.1 bits

*2105380 . *2105370 . *2105360 . *2105350
5' a a t a t a g a g a g t t a g c t a g c a g a g a a g a c a a g g t t a t a 3'

... sigma38p10
... ---------------------------------------------------------------------| proximalUP-(37)-sigma38p
... proximalUP
... ---------------------------------------------------------------------| distalUP-(45)-sigma38p10
... ---------------------------------------------------------------------| sigma38p10-proximalUP-di
distalUP 4.3 bits

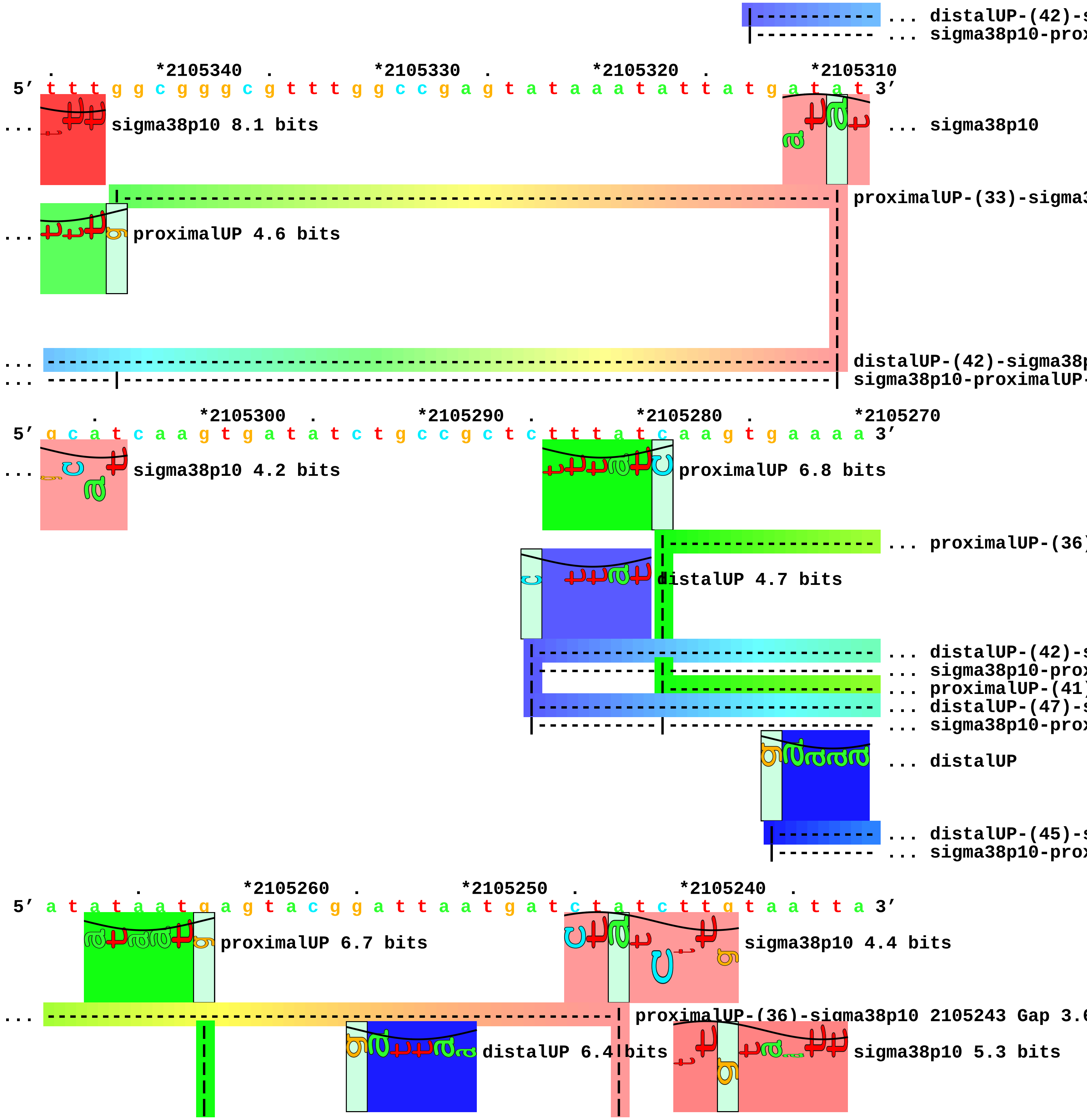
... distalUP-(42)-s
... sigma38p10-prox
*2105340 *2105330 *2105320 *2105310
5′ t t t g g c g g g c g t t t g g c c g a g t a t a a a t a t t a t g a t a t 3′
... sigma38p10 8.1 bits
... sigma38p10
proximalUP-(33)-sigma3
... proximalUP 4.6 bits
... distalUP-(42)-sigma38p
... sigma38p10-proximalUP-
*2105300 *2105290 *2105280 *2105270
5′ g c a t c a a g t g a t a t c t g c c g c t c t t t a t c a a g t g a a a a a 3′
... sigma38p10 4.2 bits
proximalUP 6.8 bits
... proximalUP-(36)
distalUP 4.7 bits
... distalUP-(42)-s
... sigma38p10-prox
... proximalUP-(41)
... distalUP-(47)-s
... sigma38p10-prox
... distalUP
... distalUP-(45)-s
... sigma38p10-prox
*2105260 *2105250 *2105240
5′ a t a t a a t g a g t a c g g a t t a a t g a t c t a t c t t g t a a t t a 3′
proximalUP 6.7 bits
sigma38p10 4.4 bits
... proximalUP-(36)-sigma38p10 2105243 Gap 3.6
distalUP 6.4 bits
sigma38p10 5.3 bits

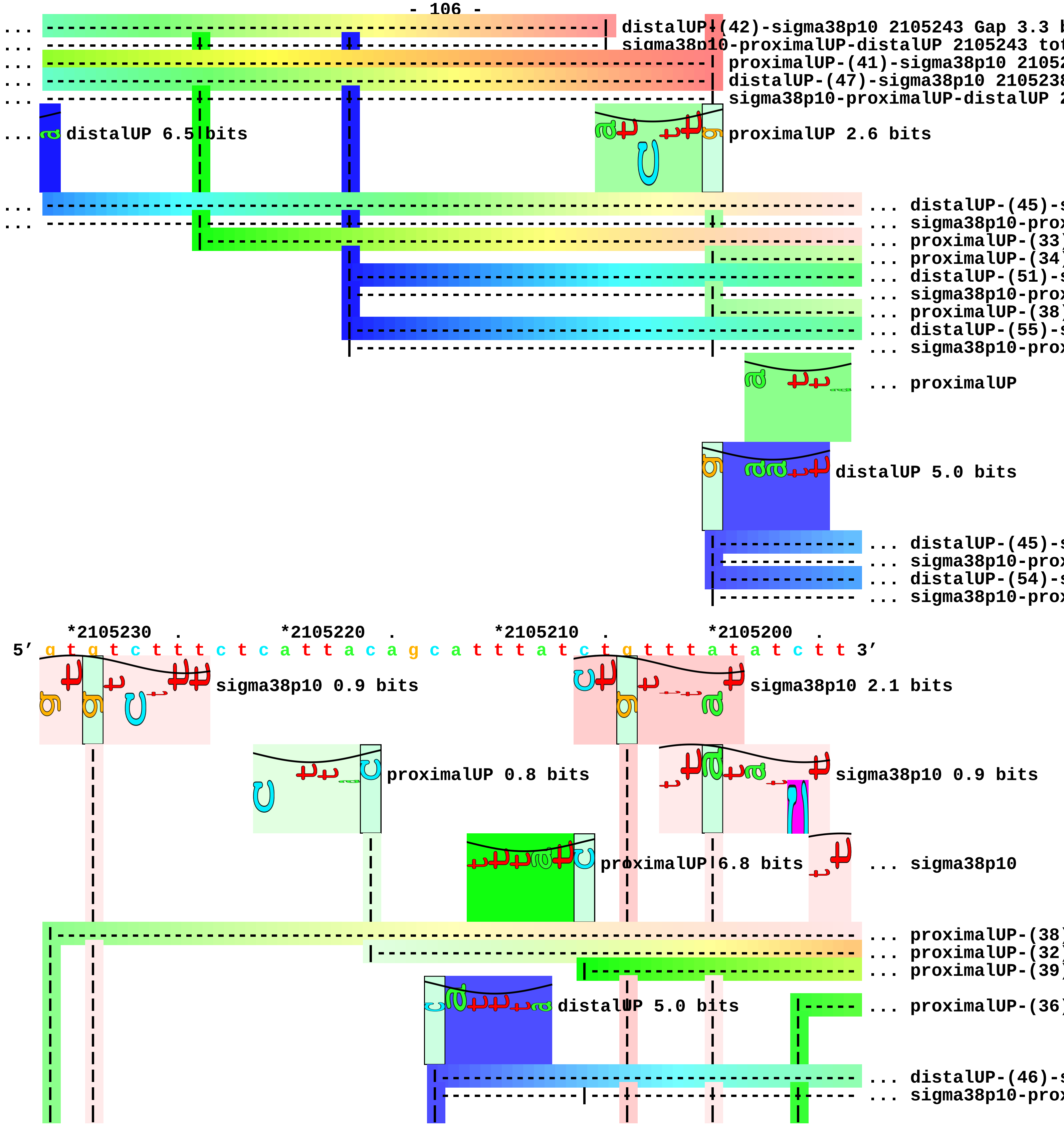
distalUP-(42)-sigma38p10 2105243 Gap 3.3 b
sigma38p10-proximalUP-distalUP 2105243 tot
proximalUP-(41)-sigma38p10 21052
distalUP-(47)-sigma38p10 2105238
sigma38p10-proximalUP-distalUP 2
distalUP 6.5 bits
proximalUP 2.6 bits
... distalUP-(45)-s
... sigma38p10-prox
... proximalUP-(33)
... proximalUP-(34)
... distalUP-(51)-s
... sigma38p10-prox
... proximalUP-(38)
... distalUP-(55)-s
... sigma38p10-prox
... proximalUP
distalUP 5.0 bits
... distalUP-(45)-s
... sigma38p10-prox
... distalUP-(54)-s
... sigma38p10-prox
*2105230 . *2105220 . *2105210 . *2105200 .
5′ g t g t c t t t c t c a t t a c a g c a t t t a t c t g t t t a t a t c t t 3′
sigma38p10 0.9 bits
sigma38p10 2.1 bits
proximalUP 0.8 bits
sigma38p10 0.9 bits
proximalUP 6.8 bits
... sigma38p10
... proximalUP-(38)
... proximalUP-(32)
... proximalUP-(39)
distalUP 5.0 bits
... proximalUP-(36)
... distalUP-(46)-s
... sigma38p10-prox

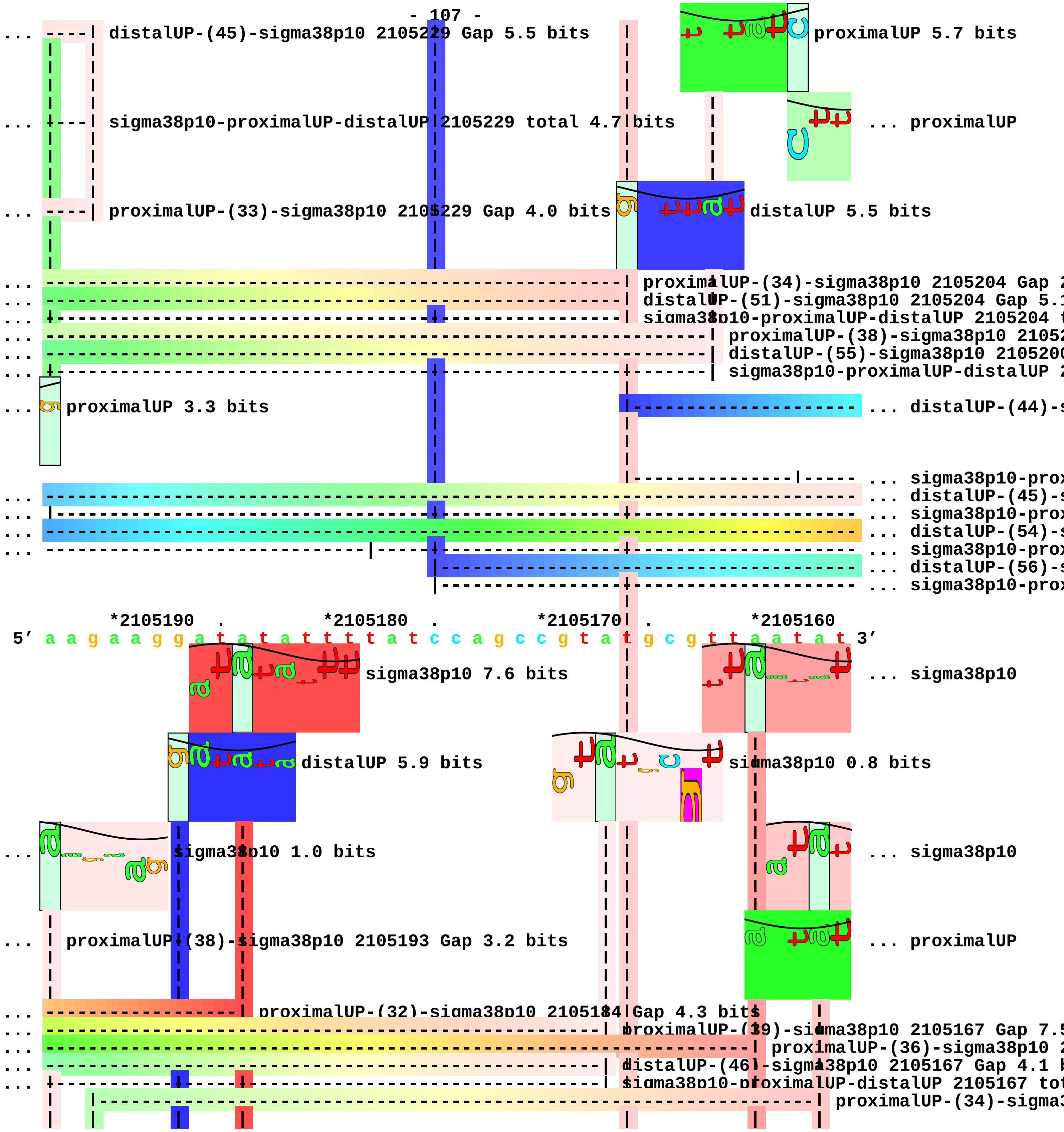
distalUP-(45)-sigma38p10 2105229 Gap 5.5 bits
proximalUP 5.7 bits
sigma38p10-proximalUP-distalUP 2105229 total 4.7 bits
... proximalUP
proximalUP-(33)-sigma38p10 2105229 Gap 4.0 bits
distalUP 5.5 bits
proximalUP-(34)-sigma38p10 2105204 Gap
distalUP-(51)-sigma38p10 2105204 Gap 5.
sigma38p10-proximalUP-distalUP 2105204
proximalUP-(38)-sigma38p10 2105
distalUP-(55)-sigma38p10 210520
sigma38p10-proximalUP-distalUP
proximalUP 3.3 bits
... distalUP-(44)-
... sigma38p10-prox
... distalUP-(45)-
... sigma38p10-prox
... distalUP-(54)-
... sigma38p10-prox
... distalUP-(56)-
... sigma38p10-prox
*2105190 . *2105180 . *2105170 . *2105160
5′ a a g a a g g a t a t a t t t t a t c c a g c c g t a t g c g t t a a t a t 3′
sigma38p10 7.6 bits
... sigma38p10
distalUP 5.9 bits
sigma38p10 0.8 bits
sigma38p10 1.0 bits
... sigma38p10
proximalUP-(38)-sigma38p10 2105193 Gap 3.2 bits
... proximalUP
proximalUP-(32)-sigma38p10 2105184 Gap 4.3 bits
proximalUP-(39)-sigma38p10 2105167 Gap 7.5
proximalUP-(36)-sigma38p10 2
distalUP-(46)-sigma38p10 2105167 Gap 4.1 b
sigma38p10-proximalUP-distalUP 2105167 tot
proximalUP-(34)-sigma3

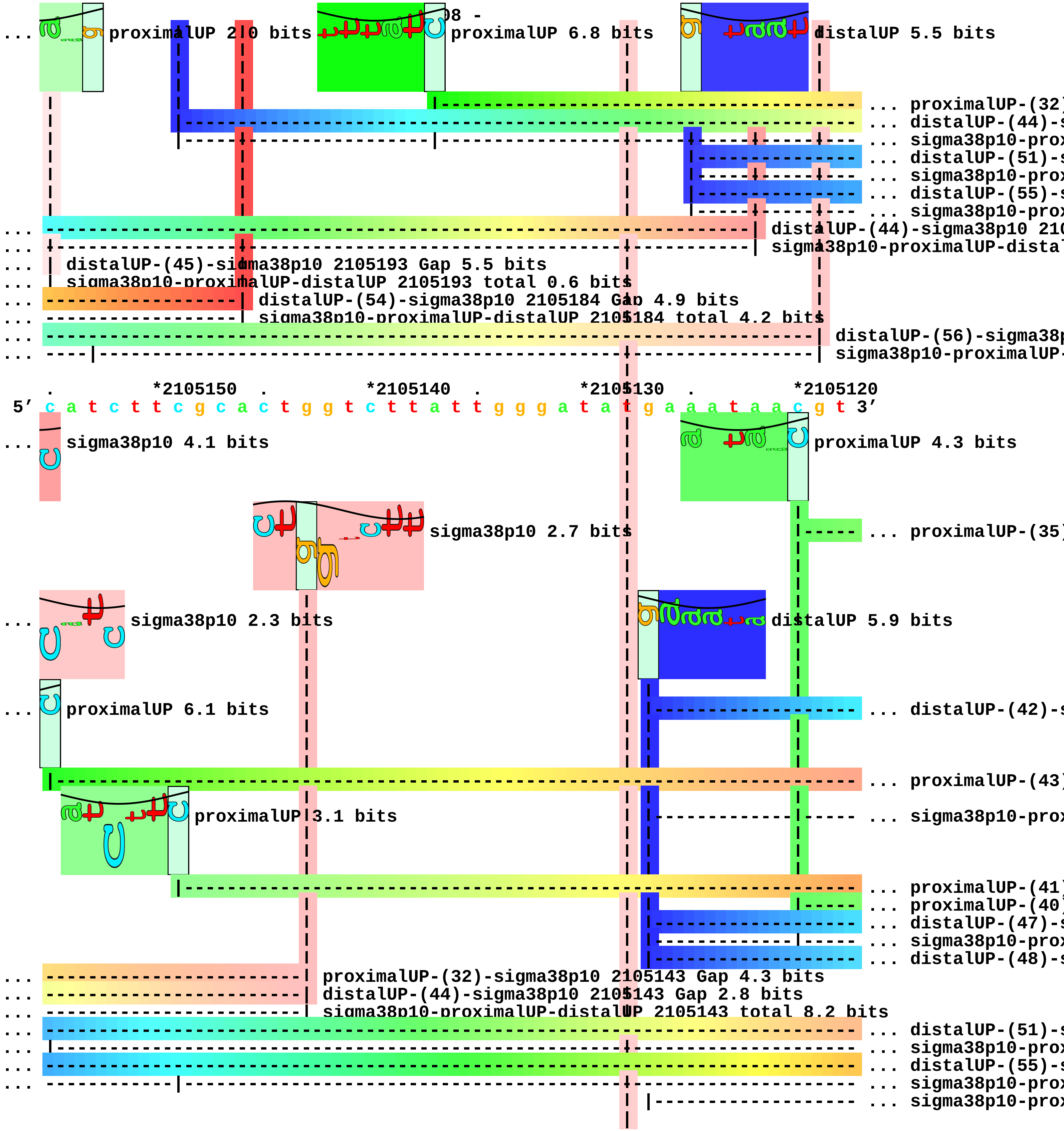
proximalUP 2.0 bits
08 -
proximalUP 6.8 bits
distalUP 5.5 bits
... proximalUP-(32)
... distalUP-(44)-s
... sigma38p10-prox
... distalUP-(51)-s
... sigma38p10-prox
... distalUP-(55)-s
... sigma38p10-prox
distalUP-(44)-sigma38p10 210
sigma38p10-proximalUP-distal
distalUP-(45)-sigma38p10 2105193 Gap 5.5 bits
sigma38p10-proximalUP-distalUP 2105193 total 0.6 bits
distalUP-(54)-sigma38p10 2105184 Gap 4.9 bits
sigma38p10-proximalUP-distalUP 2105184 total 4.2 bits
distalUP-(56)-sigma38p
sigma38p10-proximalUP-
*2105150
*2105140
*2105130
*2105120
5' c a t c t t c g c a c t g g t c t t a t t g g g a t a t g a a a t a a c g t 3'
sigma38p10 4.1 bits
proximalUP 4.3 bits
sigma38p10 2.7 bits
... proximalUP-(35)
sigma38p10 2.3 bits
distalUP 5.9 bits
proximalUP 6.1 bits
... distalUP-(42)-s
... proximalUP-(43)
proximalUP 3.1 bits
... sigma38p10-prox
... proximalUP-(41)
... proximalUP-(40)
... distalUP-(47)-s
... sigma38p10-prox
... distalUP-(48)-s
proximalUP-(32)-sigma38p10 2105143 Gap 4.3 bits
distalUP-(44)-sigma38p10 2105143 Gap 2.8 bits
sigma38p10-proximalUP-distalUP 2105143 total 8.2 bits
... distalUP-(51)-s
... sigma38p10-prox
... distalUP-(55)-s
... sigma38p10-prox
... sigma38p10-prox

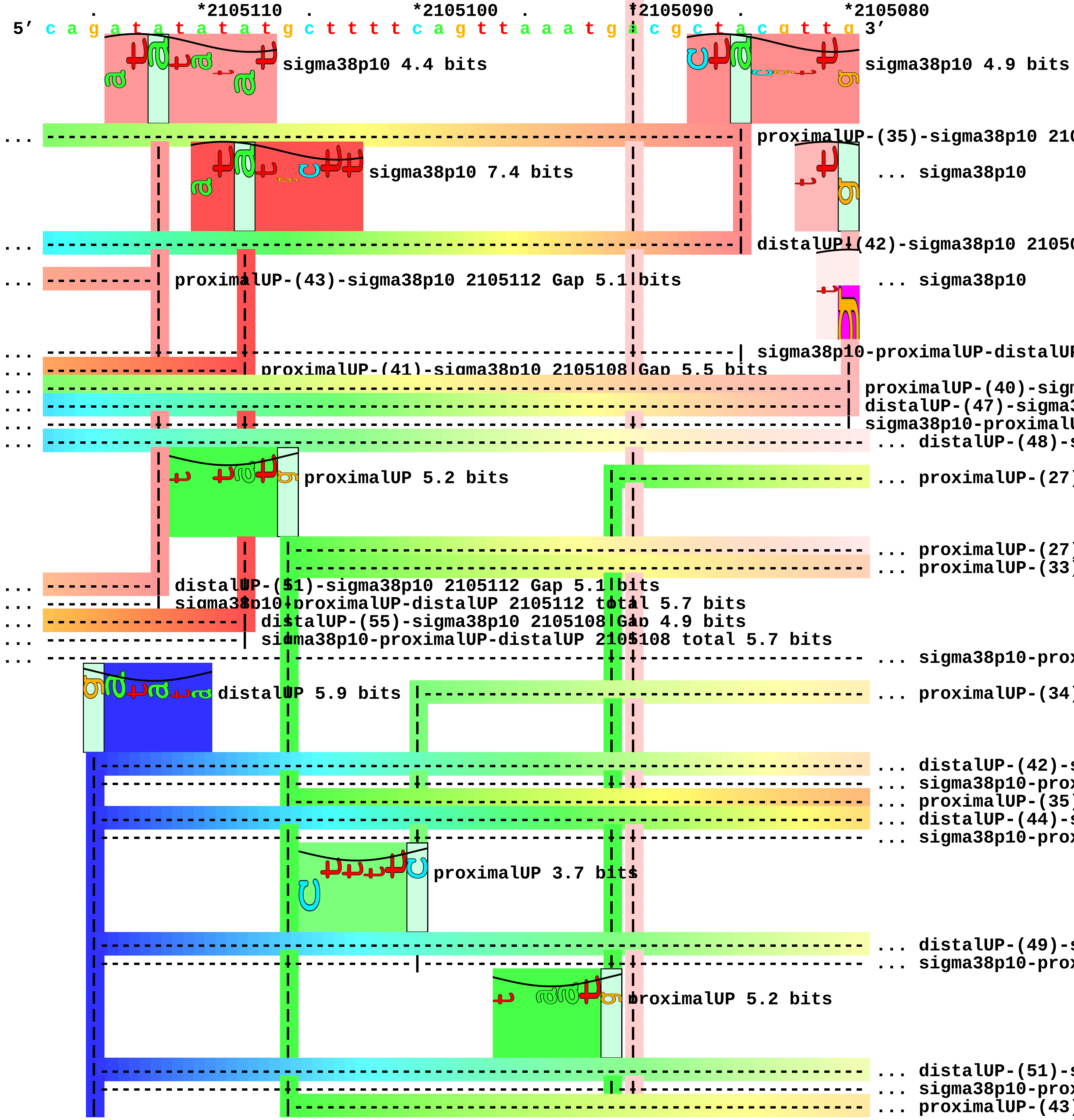
*2105110
*2105100
*2105090
*2105080
5' c a g a t a t a t a t g c t t t t c a g t t a a a t g a c g c t a c g t t g 3'
sigma38p10 4.4 bits
sigma38p10 4.9 bits
proximalUP-(35)-sigma38p10 210
sigma38p10 7.4 bits
... sigma38p10
distalUP-(42)-sigma38p10 2105
proximalUP-(43)-sigma38p10 2105112 Gap 5.1 bits
... sigma38p10
sigma38p10-proximalUP-distalUP
proximalUP-(41)-sigma38p10 2105108 Gap 5.5 bits
proximalUP-(40)-sign
distalUP-(47)-sigma3
sigma38p10-proximalU
... distalUP-(48)-s
proximalUP 5.2 bits
... proximalUP-(27)
... proximalUP-(27)
... proximalUP-(33)
distalUP-(51)-sigma38p10 2105112 Gap 5.1 bits
sigma38p10-proximalUP-distalUP 2105112 total 5.7 bits
distalUP-(55)-sigma38p10 2105108 Gap 4.9 bits
sigma38p10-proximalUP-distalUP 2105108 total 5.7 bits
... sigma38p10-prox
distalUP 5.9 bits
... proximalUP-(34)
... distalUP-(42)-s
... sigma38p10-prox
... proximalUP-(35)
... distalUP-(44)-s
... sigma38p10-prox
proximalUP 3.7 bits
... distalUP-(49)-s
... sigma38p10-prox
proximalUP 5.2 bits
... distalUP-(51)-s
... sigma38p10-prox
... proximalUP-(43)

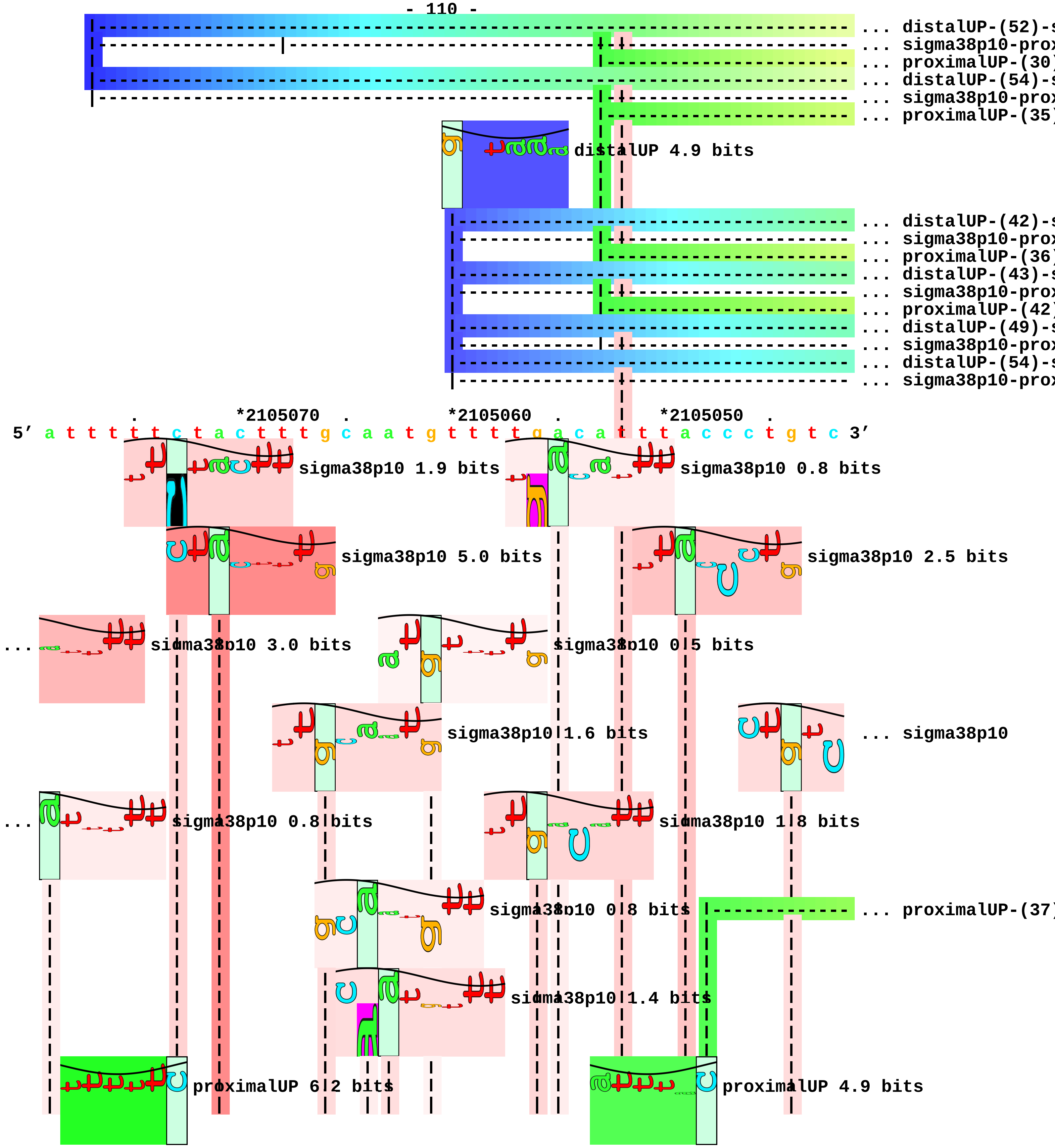
... distalUP-(52)-s
... sigma38p10-prox
... proximalUP-(30)
... distalUP-(54)-s
... sigma38p10-prox
... proximalUP-(35)
distalUP 4.9 bits
... distalUP-(42)-s
... sigma38p10-prox
... proximalUP-(36)
... distalUP-(43)-s
... sigma38p10-prox
... proximalUP-(42)
... distalUP-(49)-s
... sigma38p10-prox
... distalUP-(54)-s
... sigma38p10-prox
*2105070
*2105060
*2105050
5′ a t t t t t c t a c t t t g c a a t g t t t t g a c a t t t a c c c t g t c 3′
sigma38p10 1.9 bits
sigma38p10 0.8 bits
sigma38p10 5.0 bits
sigma38p10 2.5 bits
sigma38p10 3.0 bits
sigma38p10 0.5 bits
sigma38p10 1.6 bits
... sigma38p10
sigma38p10 0.8 bits
sigma38p10 1.8 bits
sigma38p10 0.8 bits
... proximalUP-(37)
sigma38p10 1.4 bits
proximalUP 6.2 bits
proximalUP 4.9 bits

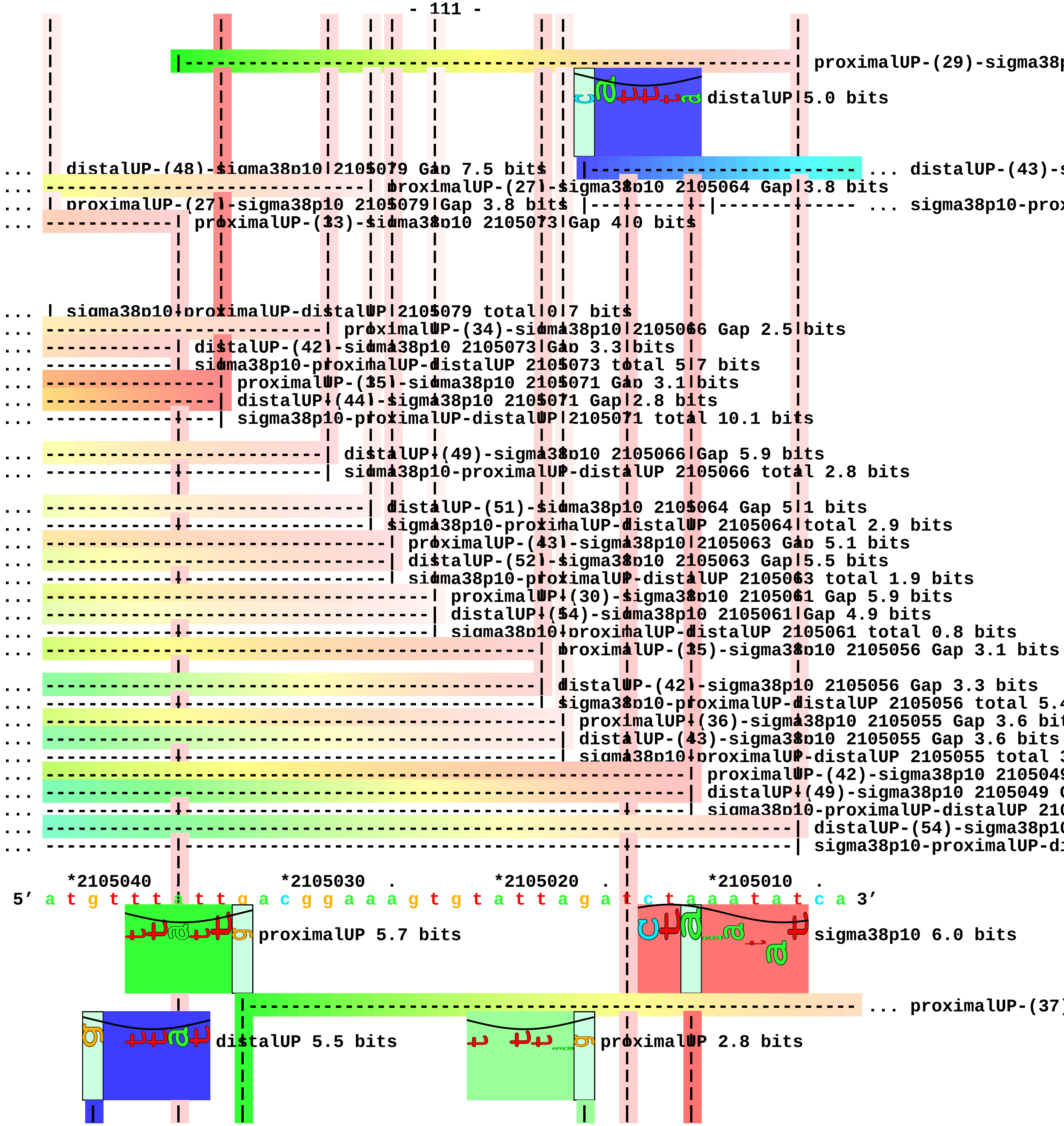
proximalUP-(29)-sigma38p
distalUP 5.0 bits
... distalUP-(48)-sigma38p10 2105079 Gap 7.5 bits
... distalUP-(43)-s
... proximalUP-(27)-sigma38p10 2105064 Gap 3.8 bits
... proximalUP-(27)-sigma38p10 2105079 Gap 3.8 bits
... sigma38p10-prox
... proximalUP-(33)-sigma38p10 2105073 Gap 4.0 bits
... sigma38p10-proximalUP-distalUP 2105079 total 0.7 bits
... proximalUP-(34)-sigma38p10 2105066 Gap 2.5 bits
... distalUP-(42)-sigma38p10 2105073 Gap 3.3 bits
... sigma38p10-proximalUP-distalUP 2105073 total 5.7 bits
... proximalUP-(35)-sigma38p10 2105071 Gap 3.1 bits
... distalUP-(44)-sigma38p10 2105071 Gap 2.8 bits
... sigma38p10-proximalUP-distalUP 2105071 total 10.1 bits
... distalUP-(49)-sigma38p10 2105066 Gap 5.9 bits
... sigma38p10-proximalUP-distalUP 2105066 total 2.8 bits
... distalUP-(51)-sigma38p10 2105064 Gap 5.1 bits
... sigma38p10-proximalUP-distalUP 2105064 total 2.9 bits
... proximalUP-(43)-sigma38p10 2105063 Gap 5.1 bits
... distalUP-(52)-sigma38p10 2105063 Gap 5.5 bits
... sigma38p10-proximalUP-distalUP 2105063 total 1.9 bits
... proximalUP-(30)-sigma38p10 2105061 Gap 5.9 bits
... distalUP-(54)-sigma38p10 2105061 Gap 4.9 bits
... sigma38p10-proximalUP-distalUP 2105061 total 0.8 bits
... proximalUP-(35)-sigma38p10 2105056 Gap 3.1 bits
... distalUP-(42)-sigma38p10 2105056 Gap 3.3 bits
... sigma38p10-proximalUP-distalUP 2105056 total 5.4
... proximalUP-(36)-sigma38p10 2105055 Gap 3.6 bit
... distalUP-(43)-sigma38p10 2105055 Gap 3.6 bits
... sigma38p10-proximalUP-distalUP 2105055 total 3
... proximalUP-(42)-sigma38p10 2105049
... distalUP-(49)-sigma38p10 2105049 G
... sigma38p10-proximalUP-distalUP 210
... distalUP-(54)-sigma38p10
... sigma38p10-proximalUP-di
*2105040 *2105030 . *2105020 . *2105010 .
5′ a t g t t t a t t g a c g g a a a g t g t a t t a g a t c t a a a t a t c a 3′
proximalUP 5.7 bits
sigma38p10 6.0 bits
... proximalUP-(37)
distalUP 5.5 bits
proximalUP 2.8 bits

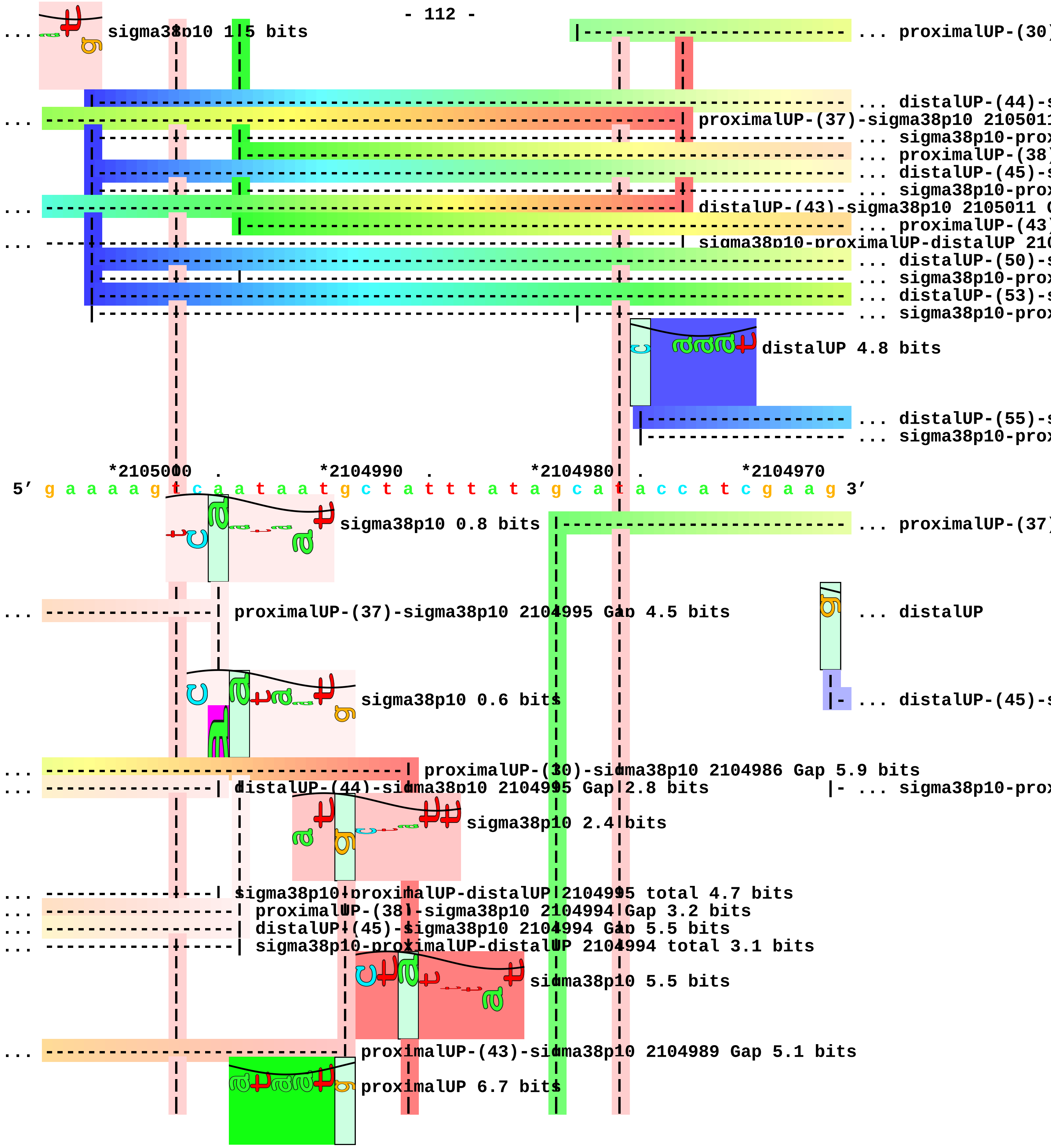
sigma38p10 1.5 bits
... proximalUP-(30)
... distalUP-(44)-s
... proximalUP-(37)-sigma38p10 2105011
... sigma38p10-prox
... proximalUP-(38)
... distalUP-(45)-s
... sigma38p10-prox
... distalUP-(43)-sigma38p10 2105011 G
... proximalUP-(43)
... sigma38p10-proximalUP-distalUP 210
... distalUP-(50)-s
... sigma38p10-prox
... distalUP-(53)-s
... sigma38p10-prox
distalUP 4.8 bits
... distalUP-(55)-s
... sigma38p10-prox
*2105000 . *2104990 . *2104980 . *2104970
5′ g a a a a g t c a a t a a t g c t a t t t a t a g c a t a c c a t c g a a g 3′
sigma38p10 0.8 bits
... proximalUP-(37)
... proximalUP-(37)-sigma38p10 2104995 Gap 4.5 bits
... distalUP
sigma38p10 0.6 bits
... distalUP-(45)-s
... proximalUP-(30)-sigma38p10 2104986 Gap 5.9 bits
... distalUP-(44)-sigma38p10 2104995 Gap 2.8 bits
... sigma38p10-prox
sigma38p10 2.4 bits
... sigma38p10-proximalUP-distalUP 2104995 total 4.7 bits
... proximalUP-(38)-sigma38p10 2104994 Gap 3.2 bits
... distalUP-(45)-sigma38p10 2104994 Gap 5.5 bits
... sigma38p10-proximalUP-distalUP 2104994 total 3.1 bits
sigma38p10 5.5 bits
... proximalUP-(43)-sigma38p10 2104989 Gap 5.1 bits
proximalUP 6.7 bits

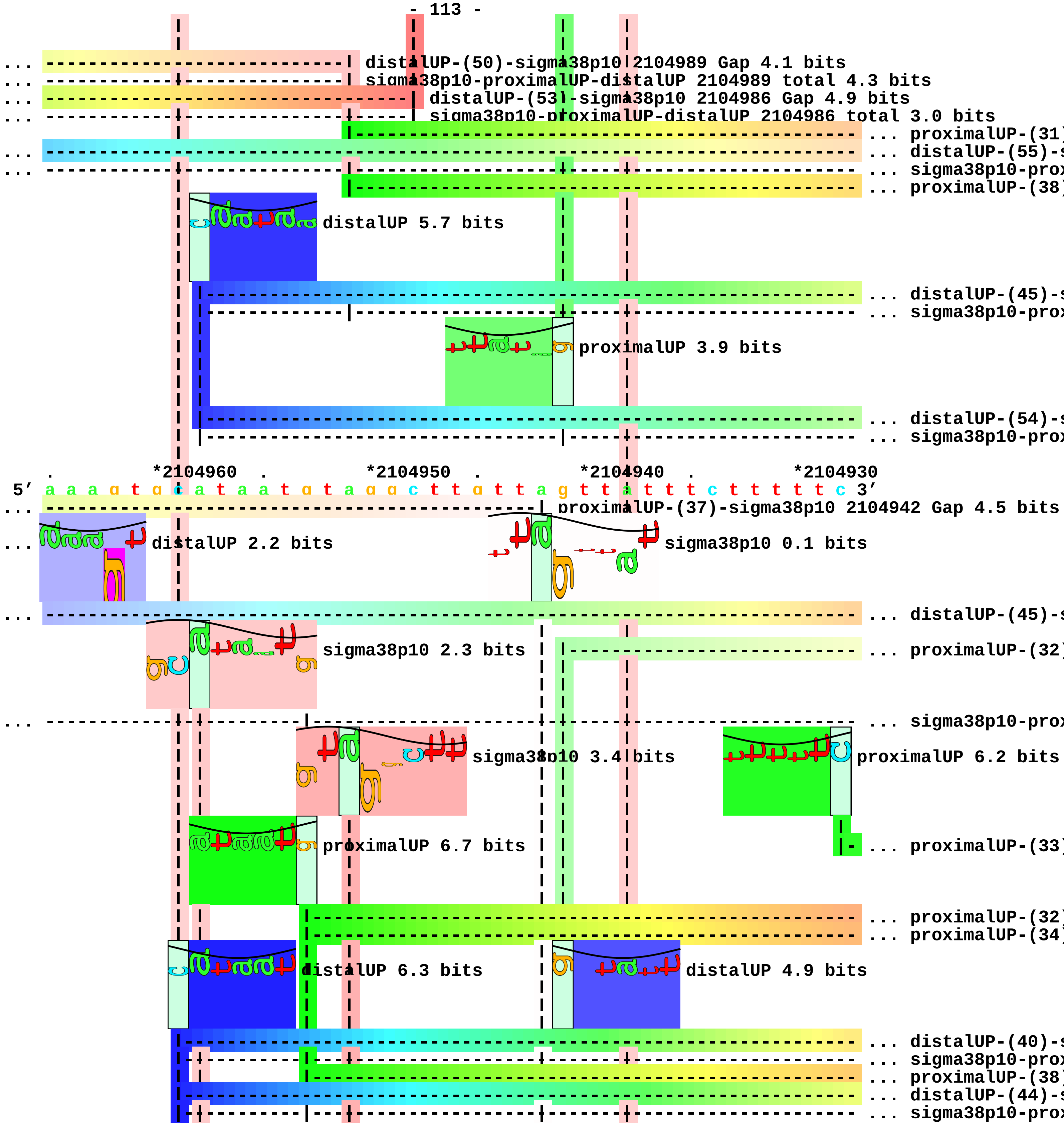
distalUP-(50)-sigma38p10 2104989 Gap 4.1 bits
sigma38p10-proximalUP-distalUP 2104989 total 4.3 bits
distalUP-(53)-sigma38p10 2104986 Gap 4.9 bits
sigma38p10-proximalUP-distalUP 2104986 total 3.0 bits
proximalUP-(31)
distalUP-(55)-s
sigma38p10-prox
proximalUP-(38)
distalUP 5.7 bits
distalUP-(45)-s
sigma38p10-prox
proximalUP 3.9 bits
distalUP-(54)-s
sigma38p10-prox
*2104960
*2104950
*2104940
*2104930
5′ a a a g t g c a t a a t g t a g g c t t g t t a g t t a t t t c t t t t t c 3′
proximalUP-(37)-sigma38p10 2104942 Gap 4.5 bits
distalUP 2.2 bits
sigma38p10 0.1 bits
distalUP-(45)-s
sigma38p10 2.3 bits
proximalUP-(32)
sigma38p10-prox
sigma38p10 3.4 bits
proximalUP 6.2 bits
proximalUP 6.7 bits
proximalUP-(33)
proximalUP-(32)
proximalUP-(34)
distalUP 6.3 bits
distalUP 4.9 bits
distalUP-(40)-s
sigma38p10-prox
proximalUP-(38)
distalUP-(44)-s
sigma38p10-prox

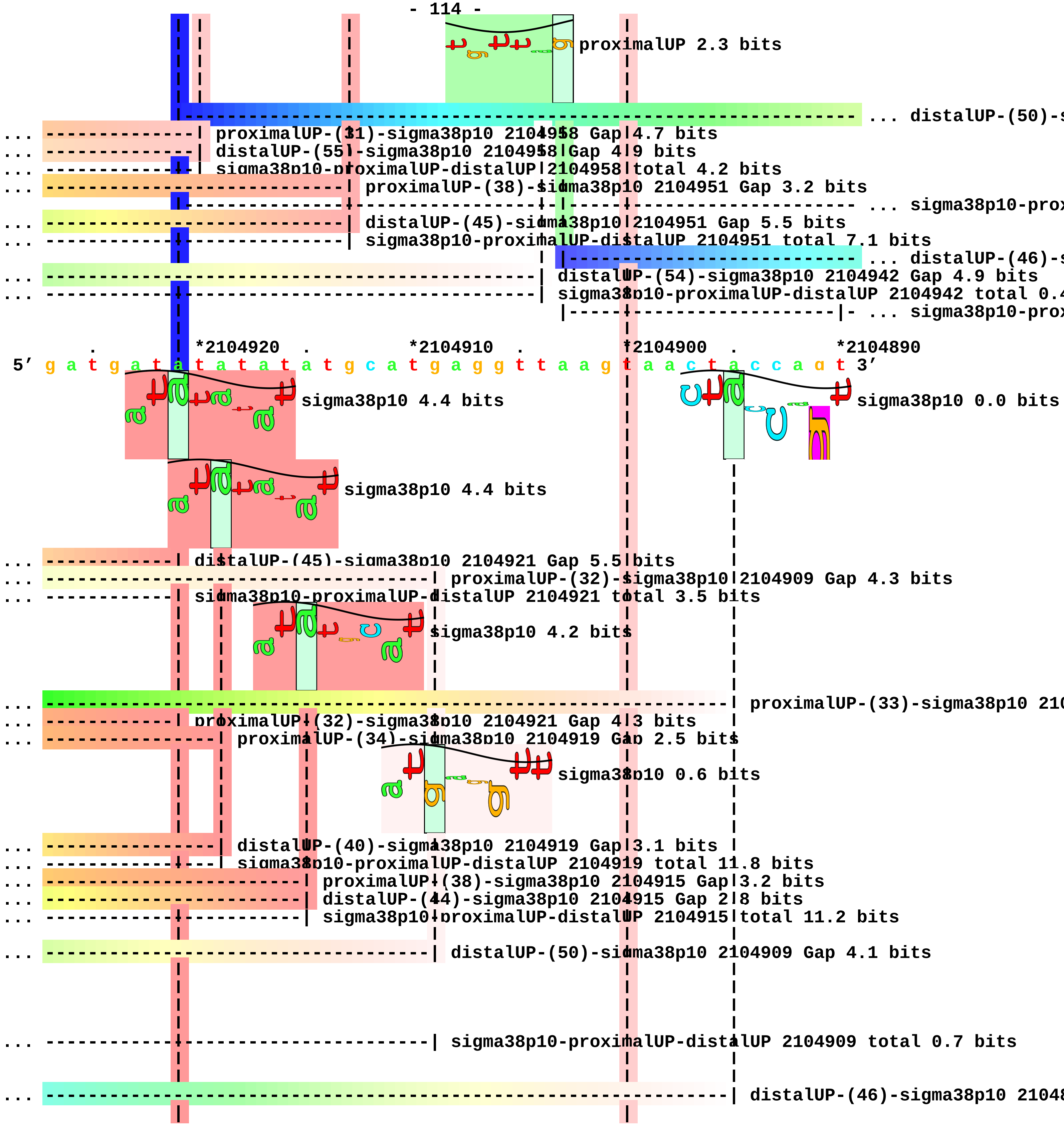
proximalUP 2.3 bits
... distalUP-(50)-s
proximalUP-(31)-sigma38p10 2104958 Gap 4.7 bits
distalUP-(55)-sigma38p10 2104958 Gap 4.9 bits
sigma38p10-proximalUP-distalUP 2104958 total 4.2 bits
proximalUP-(38)-sigma38p10 2104951 Gap 3.2 bits
... sigma38p10-prox
distalUP-(45)-sigma38p10 2104951 Gap 5.5 bits
sigma38p10-proximalUP-distalUP 2104951 total 7.1 bits
... distalUP-(46)-s
distalUP-(54)-sigma38p10 2104942 Gap 4.9 bits
sigma38p10-proximalUP-distalUP 2104942 total 0.4
... sigma38p10-prox
*2104920
*2104910
*2104900
*2104890
5′ g a t g a t a t a t a t a t g c a t g a g g t t a a g t a a c t a c c a g t 3′
sigma38p10 4.4 bits
sigma38p10 0.0 bits
sigma38p10 4.4 bits
distalUP-(45)-sigma38p10 2104921 Gap 5.5 bits
proximalUP-(32)-sigma38p10 2104909 Gap 4.3 bits
sigma38p10-proximalUP-distalUP 2104921 total 3.5 bits
sigma38p10 4.2 bits
proximalUP-(33)-sigma38p10 210
proximalUP-(32)-sigma38p10 2104921 Gap 4.3 bits
proximalUP-(34)-sigma38p10 2104919 Gap 2.5 bits
sigma38p10 0.6 bits
distalUP-(40)-sigma38p10 2104919 Gap 3.1 bits
sigma38p10-proximalUP-distalUP 2104919 total 11.8 bits
proximalUP-(38)-sigma38p10 2104915 Gap 3.2 bits
distalUP-(44)-sigma38p10 2104915 Gap 2.8 bits
sigma38p10-proximalUP-distalUP 2104915 total 11.2 bits
distalUP-(50)-sigma38p10 2104909 Gap 4.1 bits
sigma38p10-proximalUP-distalUP 2104909 total 0.7 bits
distalUP-(46)-sigma38p10 21048

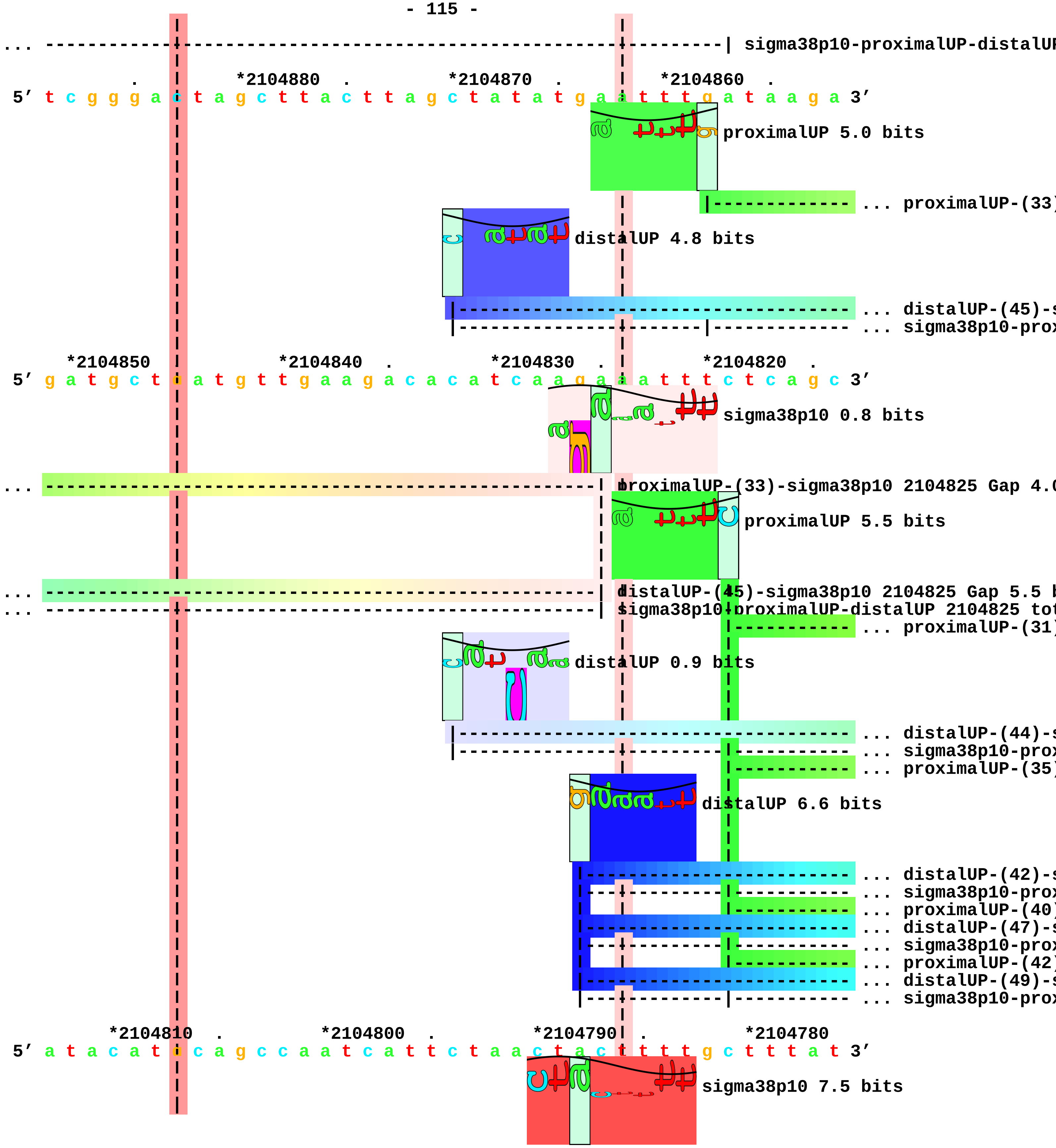
... ---------------------------------------------------------| sigma38p10-proximalUP-distalUP
*2104880 . *2104870 . *2104860 .
5' t c g g g a c t a g c t t a c t t a g c t a t a t g a a t t t g a t a a g a 3'
proximalUP 5.0 bits
... proximalUP-(33)
distalUP 4.8 bits
... distalUP-(45)-s
... sigma38p10-prox
*2104850 *2104840 . *2104830 . *2104820 .
5' g a t g c t c a t g t t g a a g a c a c a t c a a g a a a t t t c t c a g c 3'
sigma38p10 0.8 bits
... proximalUP-(33)-sigma38p10 2104825 Gap 4.0
proximalUP 5.5 bits
... distalUP-(45)-sigma38p10 2104825 Gap 5.5 b
... sigma38p10-proximalUP-distalUP 2104825 tot
... proximalUP-(31)
distalUP 0.9 bits
... distalUP-(44)-s
... sigma38p10-prox
... proximalUP-(35)
distalUP 6.6 bits
... distalUP-(42)-s
... sigma38p10-prox
... proximalUP-(40)
... distalUP-(47)-s
... sigma38p10-prox
... proximalUP-(42)
... distalUP-(49)-s
... sigma38p10-prox
*2104810 . *2104800 . *2104790 . *2104780
5' a t a c a t c c a g c c a a t c a t t c t a a c t a c t t t t g c t t t a t 3'
sigma38p10 7.5 bits

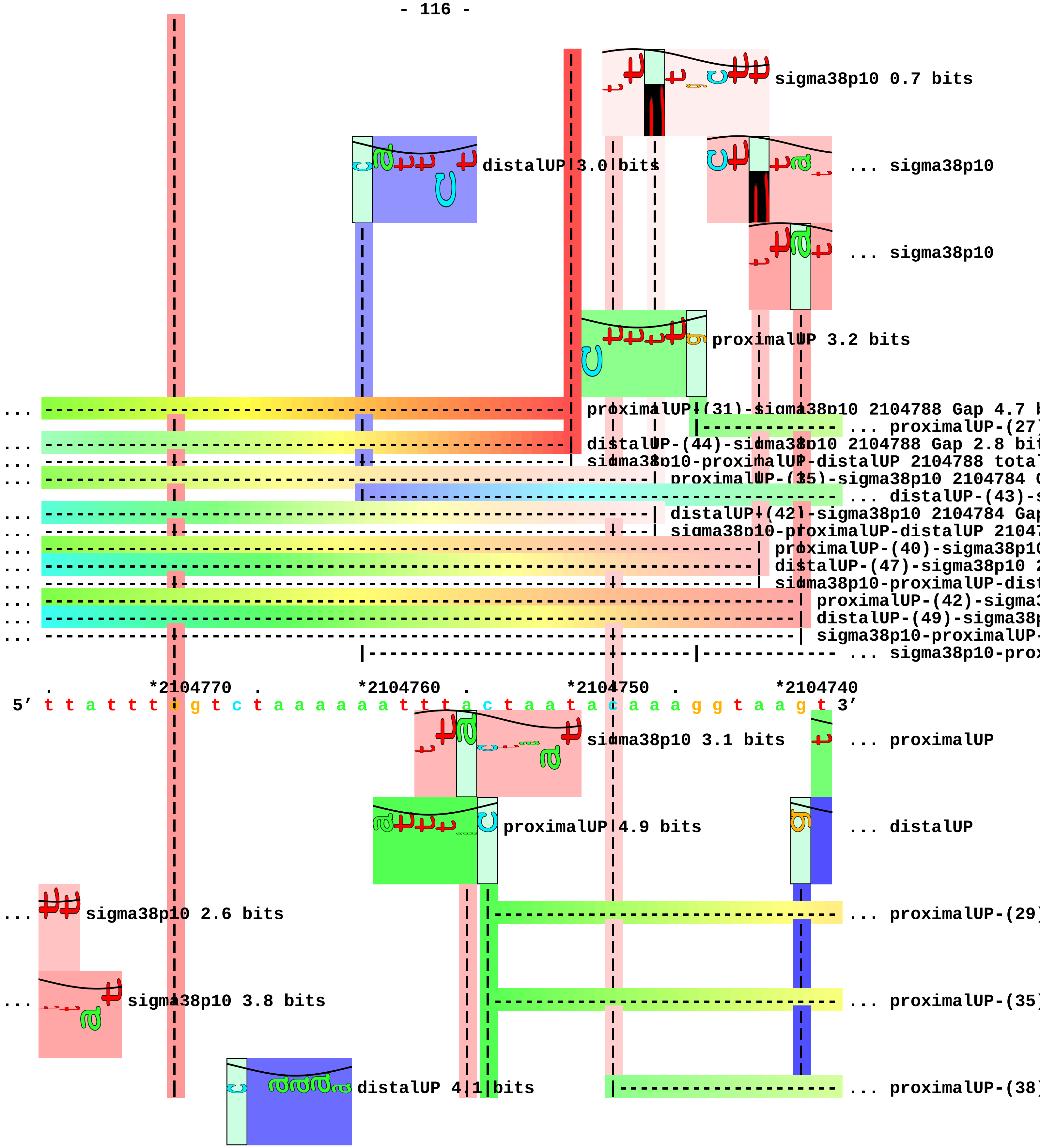
sigma38p10 0.7 bits
distalUP 3.0 bits
... sigma38p10
... sigma38p10
proximalUP 3.2 bits
proximalUP-(31)-sigma38p10 2104788 Gap 4.7 b
... proximalUP-(27)
distalUP-(44)-sigma38p10 2104788 Gap 2.8 bit
sigma38p10-proximalUP-distalUP 2104788 total
proximalUP-(35)-sigma38p10 2104784 G
... distalUP-(43)-s
distalUP-(42)-sigma38p10 2104784 Gap
sigma38p10-proximalUP-distalUP 21047
proximalUP-(40)-sigma38p10
distalUP-(47)-sigma38p10 2
sigma38p10-proximalUP-dist
proximalUP-(42)-sigma3
distalUP-(49)-sigma38p
sigma38p10-proximalUP-
... sigma38p10-prox
*2104770
*2104760
*2104750
*2104740
5′ t t a t t t g g t c t a a a a a a t t t a c t a a t a c a a a g g t a a g t 3′
sigma38p10 3.1 bits
... proximalUP
proximalUP 4.9 bits
... distalUP
... sigma38p10 2.6 bits
... proximalUP-(29)
... sigma38p10 3.8 bits
... proximalUP-(35)
distalUP 4.1 bits
... proximalUP-(38)

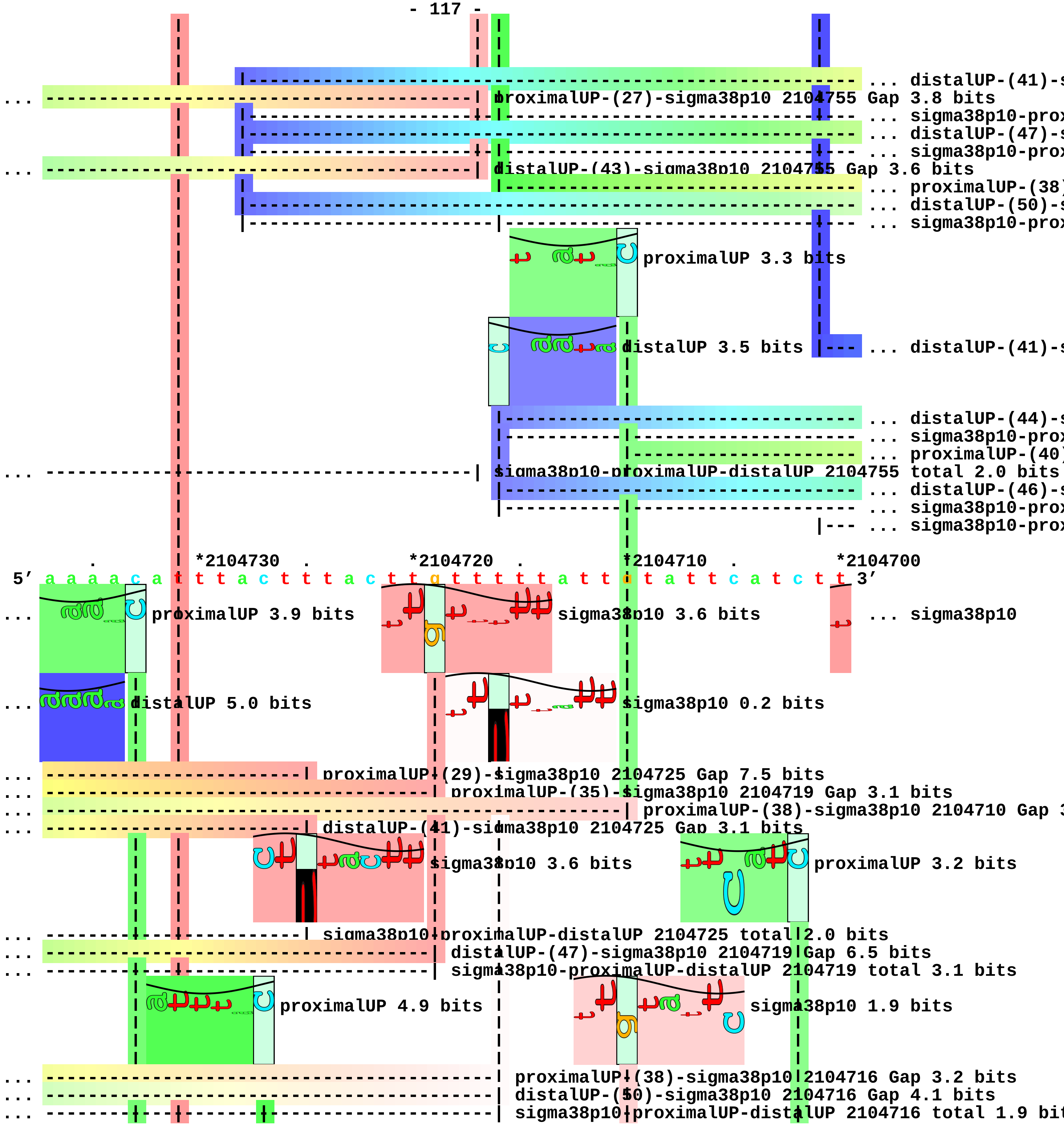
... distalUP-(41)-s
... proximalUP-(27)-sigma38p10 2104755 Gap 3.8 bits
... sigma38p10-prox
... distalUP-(47)-s
... sigma38p10-prox
... distalUP-(43)-sigma38p10 2104755 Gap 3.6 bits
... proximalUP-(38)
... distalUP-(50)-s
... sigma38p10-prox
proximalUP 3.3 bits
distalUP 3.5 bits
... distalUP-(41)-s
... distalUP-(44)-s
... sigma38p10-prox
... proximalUP-(40)
... sigma38p10-proximalUP-distalUP 2104755 total 2.0 bits
... distalUP-(46)-s
... sigma38p10-prox
... sigma38p10-prox
*2104730 *2104720 *2104710 *2104700
5′ a a a a c a t t t a c t t t a c t t g t t t t t a t t g t a t t c a t c t t 3′
... proximalUP 3.9 bits
sigma38p10 3.6 bits
... sigma38p10
... distalUP 5.0 bits
sigma38p10 0.2 bits
... proximalUP-(29)-sigma38p10 2104725 Gap 7.5 bits
... proximalUP-(35)-sigma38p10 2104719 Gap 3.1 bits
... proximalUP-(38)-sigma38p10 2104710 Gap [illegible]
... distalUP-(41)-sigma38p10 2104725 Gap 3.1 bits
sigma38p10 3.6 bits
proximalUP 3.2 bits
... sigma38p10-proximalUP-distalUP 2104725 total 2.0 bits
... distalUP-(47)-sigma38p10 2104719 Gap 6.5 bits
... sigma38p10-proximalUP-distalUP 2104719 total 3.1 bits
proximalUP 4.9 bits
sigma38p10 1.9 bits
... proximalUP-(38)-sigma38p10 2104716 Gap 3.2 bits
... distalUP-(50)-sigma38p10 2104716 Gap 4.1 bits
... sigma38p10-proximalUP-distalUP 2104716 total 1.9 bi[illegible]

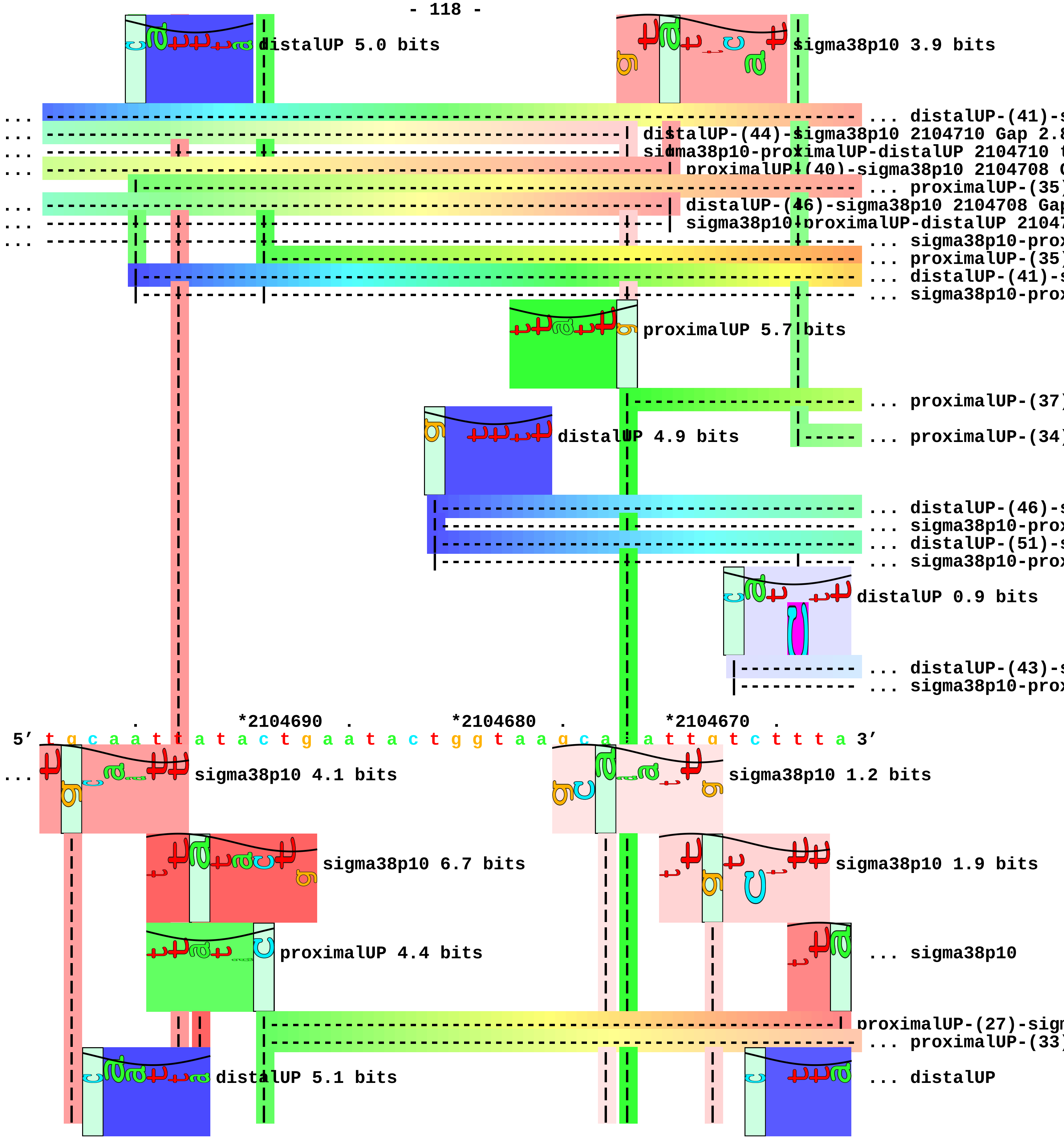
distalUP 5.0 bits
sigma38p10 3.9 bits
... distalUP-(41)-s
distalUP-(44)-sigma38p10 2104710 Gap 2.8
sigma38p10-proximalUP-distalUP 2104710 t
proximalUP-(40)-sigma38p10 2104708 G
... proximalUP-(35)
distalUP-(46)-sigma38p10 2104708 Gap
sigma38p10-proximalUP-distalUP 2104
... sigma38p10-prox
... proximalUP-(35)
... distalUP-(41)-
... sigma38p10-prox
proximalUP 5.7 bits
... proximalUP-(37)
distalUP 4.9 bits
... proximalUP-(34)
... distalUP-(46)-s
... sigma38p10-prox
... distalUP-(51)-s
... sigma38p10-prox
distalUP 0.9 bits
... distalUP-(43)-s
... sigma38p10-prox
*2104690
*2104680
*2104670
5′ t g c a a t t a t a c t g a a t a c t g g t a a g c a t a t t g t c t t t a 3′
sigma38p10 4.1 bits
sigma38p10 1.2 bits
sigma38p10 6.7 bits
sigma38p10 1.9 bits
proximalUP 4.4 bits
... sigma38p10
proximalUP-(27)-sigm
... proximalUP-(33)
distalUP 5.1 bits
... distalUP

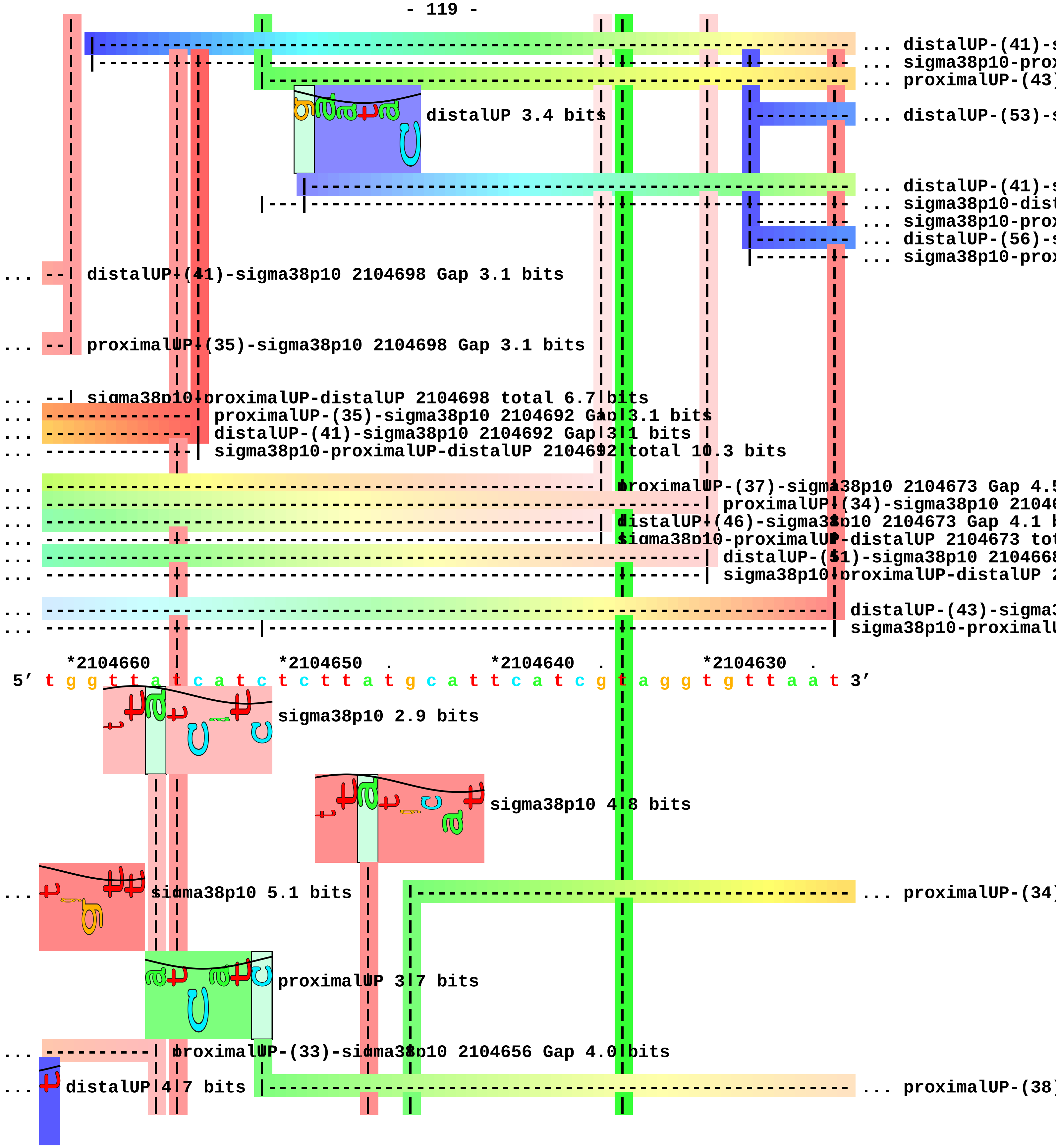
... distalUP-(41)-s
... sigma38p10-prox
... proximalUP-(43)
distalUP 3.4 bits
... distalUP-(53)-s
... distalUP-(41)-s
... sigma38p10-dist
... sigma38p10-prox
... distalUP-(56)-s
... sigma38p10-prox
... distalUP-(41)-sigma38p10 2104698 Gap 3.1 bits
... proximalUP-(35)-sigma38p10 2104698 Gap 3.1 bits
... sigma38p10-proximalUP-distalUP 2104698 total 6.7 bits
... proximalUP-(35)-sigma38p10 2104692 Gap 3.1 bits
... distalUP-(41)-sigma38p10 2104692 Gap 3.1 bits
... sigma38p10-proximalUP-distalUP 2104692 total 10.3 bits
proximalUP-(37)-sigma38p10 2104673 Gap 4.5
proximalUP-(34)-sigma38p10 21046
distalUP-(46)-sigma38p10 2104673 Gap 4.1 b
sigma38p10-proximalUP-distalUP 2104673 tot
distalUP-(51)-sigma38p10 210466
sigma38p10-proximalUP-distalUP 2
distalUP-(43)-sigma3
sigma38p10-proximalU
*2104660 *2104650 . *2104640 . *2104630 .
5′ t g g t t a t c a t c t c t t a t g c a t t c a t c g t a g g t g t t a a t 3′
sigma38p10 2.9 bits
sigma38p10 4.8 bits
... sigma38p10 5.1 bits
... proximalUP-(34)
proximalUP 3.7 bits
... proximalUP-(33)-sigma38p10 2104656 Gap 4.0 bits
... distalUP 4.7 bits
... proximalUP-(38)

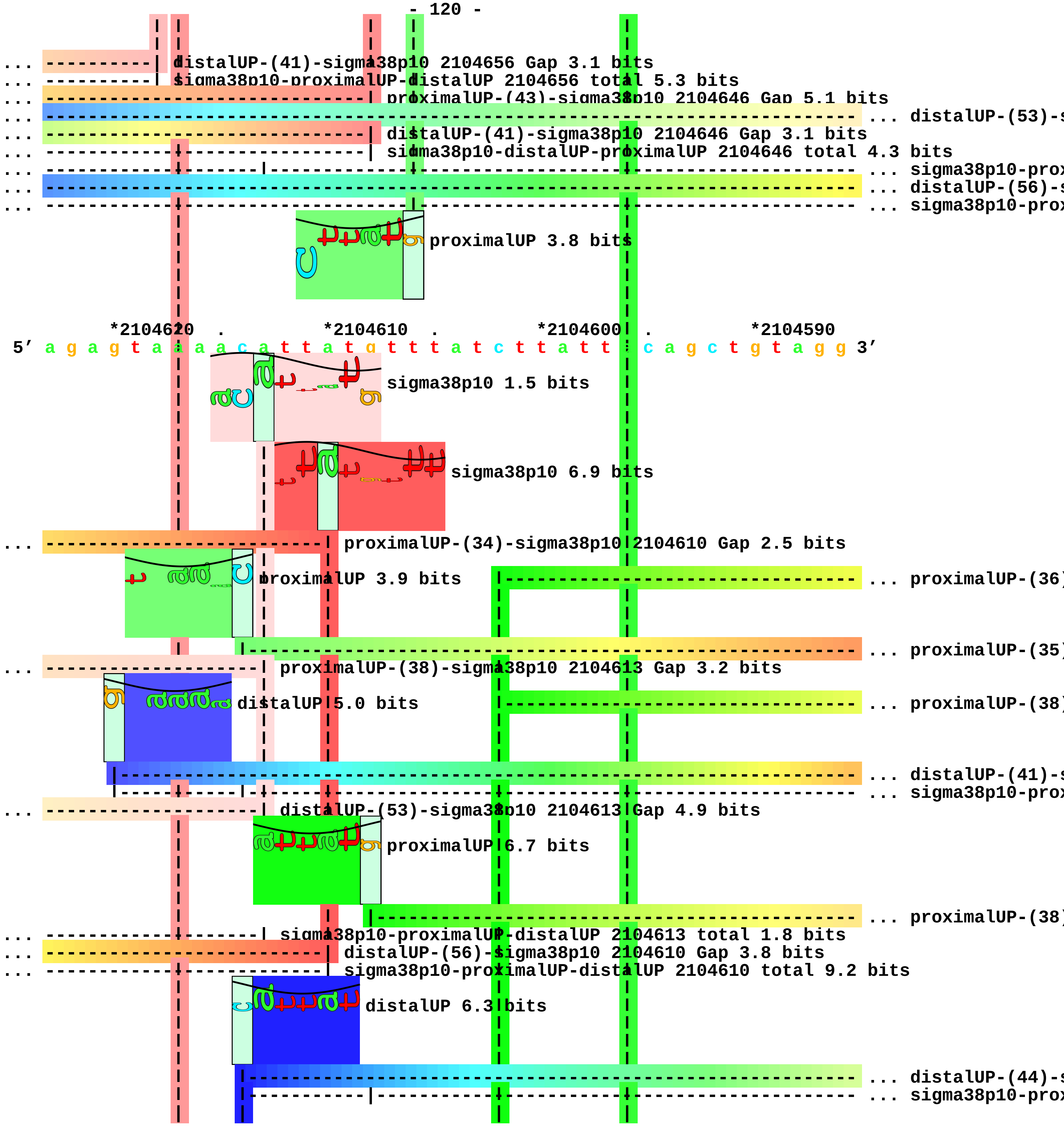
distalUP-(41)-sigma38p10 2104656 Gap 3.1 bits
sigma38p10-proximalUP-distalUP 2104656 total 5.3 bits
proximalUP-(43)-sigma38p10 2104646 Gap 5.1 bits
... distalUP-(53)-s
distalUP-(41)-sigma38p10 2104646 Gap 3.1 bits
sigma38p10-distalUP-proximalUP 2104646 total 4.3 bits
... sigma38p10-prox
... distalUP-(56)-s
... sigma38p10-prox
proximalUP 3.8 bits
*2104620 . *2104610 . *2104600 . *2104590
5' a g a g t a a a a c a t t a t g t t t a t c t t a t t t c a g c t g t a g g 3'
sigma38p10 1.5 bits
sigma38p10 6.9 bits
proximalUP-(34)-sigma38p10 2104610 Gap 2.5 bits
proximalUP 3.9 bits
... proximalUP-(36)
... proximalUP-(35)
proximalUP-(38)-sigma38p10 2104613 Gap 3.2 bits
distalUP 5.0 bits
... proximalUP-(38)
... distalUP-(41)-s
... sigma38p10-prox
distalUP-(53)-sigma38p10 2104613 Gap 4.9 bits
proximalUP 6.7 bits
... proximalUP-(38)
sigma38p10-proximalUP-distalUP 2104613 total 1.8 bits
distalUP-(56)-sigma38p10 2104610 Gap 3.8 bits
sigma38p10-proximalUP-distalUP 2104610 total 9.2 bits
distalUP 6.3 bits
... distalUP-(44)-s
... sigma38p10-prox

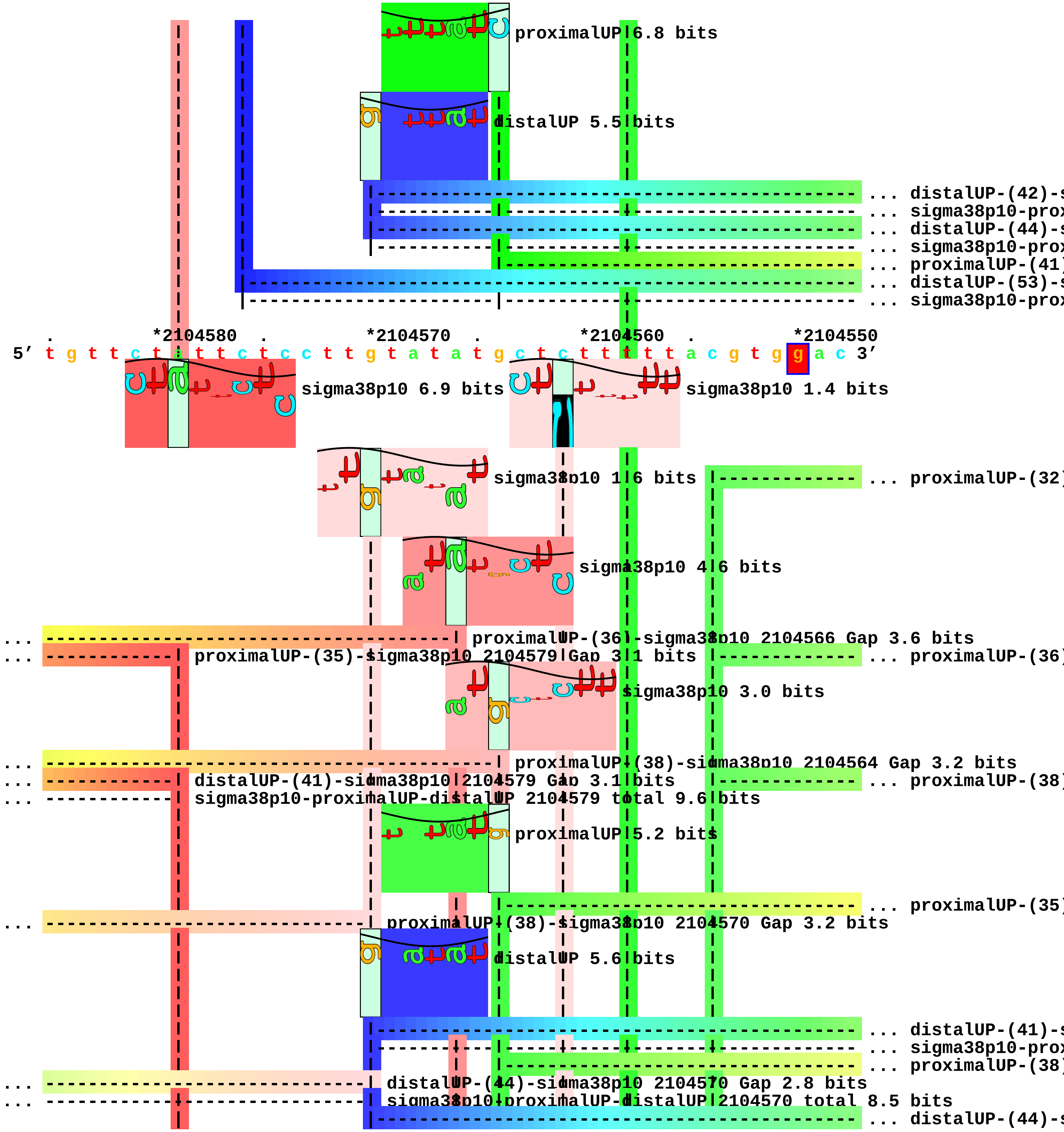

proximalUP 6.8 bits
distalUP 5.5 bits
... distalUP-(42)-s
... sigma38p10-prox
... distalUP-(44)-s
... sigma38p10-prox
... proximalUP-(41)
... distalUP-(53)-s
... sigma38p10-prox
*2104580
*2104570
*2104560
*2104550
5′ t g t t c t a t t c t c c t t g t a t a t g c t c t t t t t a c g t g g a c 3′
sigma38p10 6.9 bits
sigma38p10 1.4 bits
sigma38p10 1.6 bits
... proximalUP-(32)
sigma38p10 4.6 bits
proximalUP-(36)-sigma38p10 2104566 Gap 3.6 bits
proximalUP-(35)-sigma38p10 2104579 Gap 3.1 bits
... proximalUP-(36)
sigma38p10 3.0 bits
proximalUP-(38)-sigma38p10 2104564 Gap 3.2 bits
distalUP-(41)-sigma38p10 2104579 Gap 3.1 bits
... proximalUP-(38)
sigma38p10-proximalUP-distalUP 2104579 total 9.6 bits
proximalUP 5.2 bits
... proximalUP-(35)
proximalUP-(38)-sigma38p10 2104570 Gap 3.2 bits
distalUP 5.6 bits
... distalUP-(41)-s
... sigma38p10-prox
... proximalUP-(38)
distalUP-(44)-sigma38p10 2104570 Gap 2.8 bits
sigma38p10-proximalUP-distalUP 2104570 total 8.5 bits
... distalUP-(44)-s

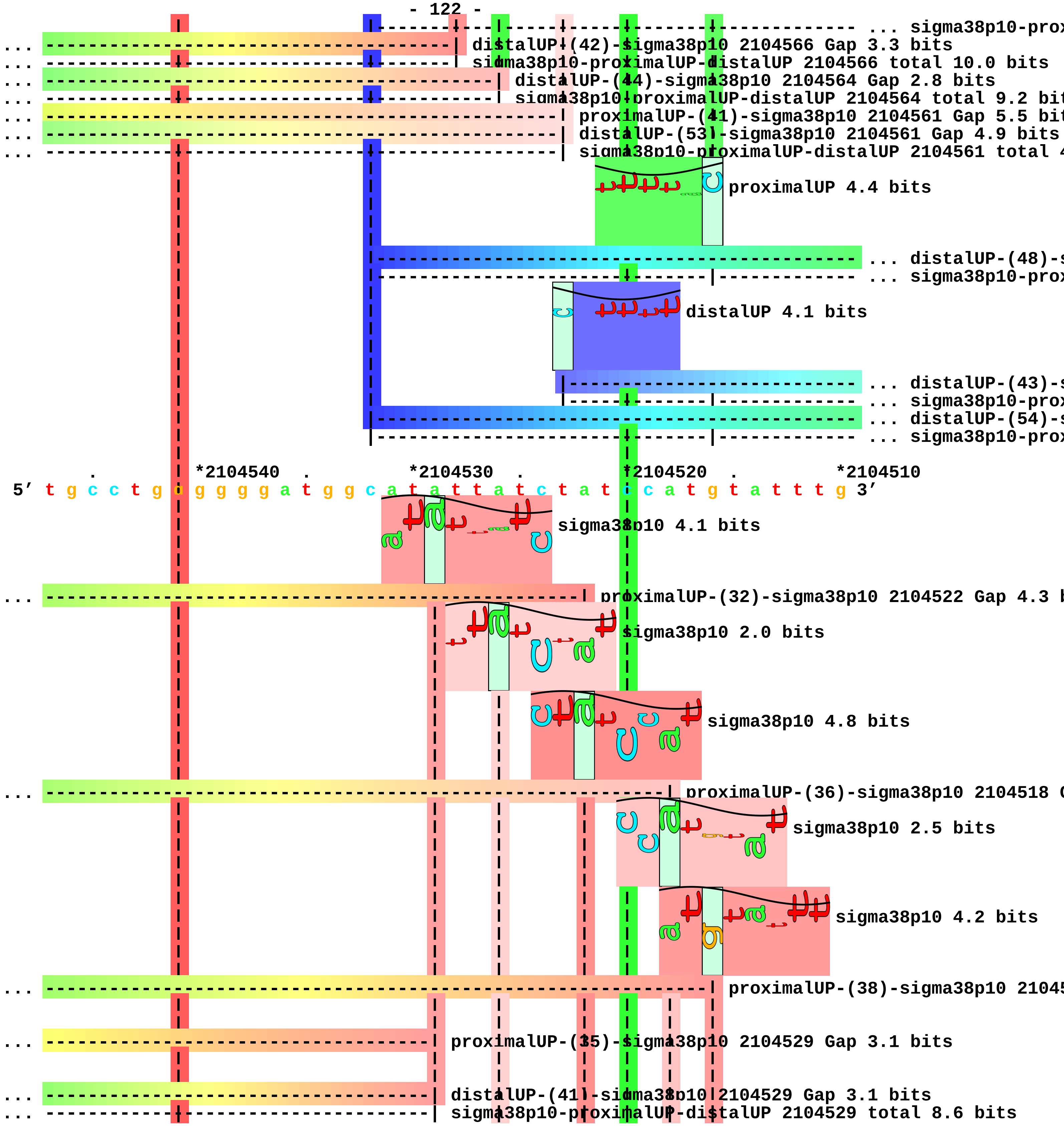
... sigma38p10-prox
... distalUP-(42)-sigma38p10 2104566 Gap 3.3 bits
... sigma38p10-proximalUP-distalUP 2104566 total 10.0 bits
... distalUP-(44)-sigma38p10 2104564 Gap 2.8 bits
... sigma38p10-proximalUP-distalUP 2104564 total 9.2 bit
... proximalUP-(41)-sigma38p10 2104561 Gap 5.5 bit
... distalUP-(53)-sigma38p10 2104561 Gap 4.9 bits
... sigma38p10-proximalUP-distalUP 2104561 total 4
proximalUP 4.4 bits
... distalUP-(48)-s
... sigma38p10-prox
distalUP 4.1 bits
... distalUP-(43)-s
... sigma38p10-prox
... distalUP-(54)-s
... sigma38p10-prox
*2104540
*2104530
*2104520
*2104510
5′ t g c c t g g g g g g a t g g c a t a t t a t c t a t c c a t g t a t t t g 3′
sigma38p10 4.1 bits
... proximalUP-(32)-sigma38p10 2104522 Gap 4.3 b
sigma38p10 2.0 bits
sigma38p10 4.8 bits
... proximalUP-(36)-sigma38p10 2104518 G
sigma38p10 2.5 bits
sigma38p10 4.2 bits
... proximalUP-(38)-sigma38p10 21045
... proximalUP-(35)-sigma38p10 2104529 Gap 3.1 bits
... distalUP-(41)-sigma38p10 2104529 Gap 3.1 bits
... sigma38p10-proximalUP-distalUP 2104529 total 8.6 bits

```
... ------------------------------------------| proximalUP-(38)-sigma38p10 2104526 Gap 3.2 bits

... ------------------------------------------| distalUP-(44)-sigma38p10 2104526 Gap 2.8 bits
... ------------------------------------------| sigma38p10-proximalUP-distalUP 2104526 total 6.7 bit

... --------------------------------------------------| distalUP-(48)-sigma38p10 2104522 Gap 7.5 bit
... --------------------------------------------------| sigma38p10-proximalUP-distalUP 2104522 tota

... ----------------------------------------------------------| distalUP-(43)-sigma38p10 2104518 Gap
... ----------------------------------------------------------| sigma38p10-proximalUP-distalUP 21045
... --------------------------------------------------------------| distalUP-(54)-sigma38p10 210451
... --------------------------------------------------------------| sigma38p10-proximalUP-distalUP 2

        .         *2104500
 5' g t c a g c c c t a 3'
```

piece 30, NC_000913, yehE, config: linear, direction: -, begin: 2190999, end: 2190750

. *2190990 . *2190980 . *2190970 .

5′ t g a a c a c t t a t t t a a t a c t g c g a a t a t t c t g c g g a c c t 3′

proximalUP 3.3 bits ... proximalUP-(36)

T timestamp: 2018Sep20 11-34-53 ... proximalUP-(35)

distalUP 4.1 bits proximalUP 6.6 bits

... distalUP-(46)-s

... sigma38p10-prox

distalUP 7.2 bits

... distalUP-(43)-s

... sigma38p10-prox

... proximalUP-(40)

... distalUP-(47)-s

... sigma38p10-prox

*2190960 . *2190950 . *2190940 . *2190930 .

5′ t c g t c c g a a a c t c t a c a a a g t c g a t t g a t c a t a c a g g c 3′

... proximalUP-(36)-sigma38p10 2190935 Gap 3.6

... proximalUP-(35)-sigma38p10 2190947 Gap 3.1 bits

sigma38p10 3.4 bits

... distalUP-(46)-sigma38p10 2190947 Gap 4.1 bits

... sigma38p10-proximalUP-distalUP 2190947 total 3.6 bits

sigma38p10 0.9 bits

... distalUP-(43)-sigma38p10 2190935 Gap 3.6 b

... sigma38p10-proximalUP-distalUP 2190935 tot

... proximalUP-(40)-sigma38p10 219093

... distalUP-(47)-sigma38p10 2190931 G

... sigma38p10-proximalUP-distalUP 219

sigma38p10 0.6 bits

distalUP 2.4 bits

... distalUP-(55)-s

... sigma38p10-prox

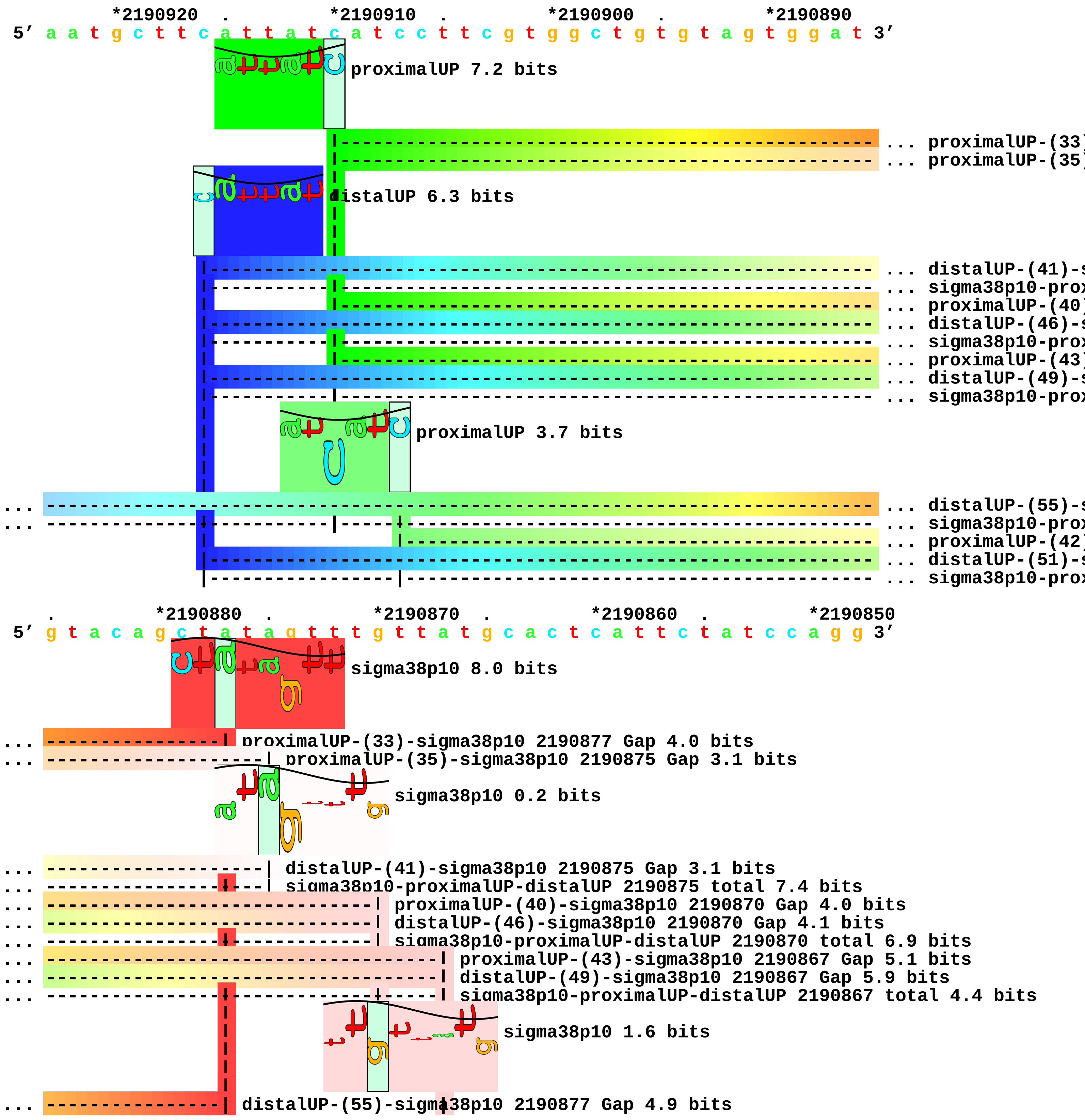
*2190920 . *2190910 . *2190900 . *2190890
5′ a a t g c t t c a t t a t c a t c c t t c g t g g c t g t g t a g t g g a t 3′
proximalUP 7.2 bits
... proximalUP-(33)
... proximalUP-(35)
distalUP 6.3 bits
... distalUP-(41)-s
... sigma38p10-prox
... proximalUP-(40)
... distalUP-(46)-s
... sigma38p10-prox
... proximalUP-(43)
... distalUP-(49)-s
... sigma38p10-prox
proximalUP 3.7 bits
... distalUP-(55)-s
... sigma38p10-prox
... proximalUP-(42)
... distalUP-(51)-s
... sigma38p10-prox
. *2190880 . *2190870 . *2190860 . *2190850
5′ g t a c a g c t a t a g t t t g t t a t g c a c t c a t t c t a t c c a g g 3′
sigma38p10 8.0 bits
proximalUP-(33)-sigma38p10 2190877 Gap 4.0 bits
proximalUP-(35)-sigma38p10 2190875 Gap 3.1 bits
sigma38p10 0.2 bits
distalUP-(41)-sigma38p10 2190875 Gap 3.1 bits
sigma38p10-proximalUP-distalUP 2190875 total 7.4 bits
proximalUP-(40)-sigma38p10 2190870 Gap 4.0 bits
distalUP-(46)-sigma38p10 2190870 Gap 4.1 bits
sigma38p10-proximalUP-distalUP 2190870 total 6.9 bits
proximalUP-(43)-sigma38p10 2190867 Gap 5.1 bits
distalUP-(49)-sigma38p10 2190867 Gap 5.9 bits
sigma38p10-proximalUP-distalUP 2190867 total 4.4 bits
sigma38p10 1.6 bits
distalUP-(55)-sigma38p10 2190877 Gap 4.9 bits

```
... ----------------| sigma38p10-proximalUP-distalUP 2190877 total 8.8 bits
... ----------------------------------------| proximalUP-(42)-sigma38p10 2190865 Gap 5.1 bits
... ----------------------------------------| distalUP-(51)-sigma38p10 2190865 Gap 5.1 bits
... ------------------------------|-----| sigma38p10-proximalUP-distalUP 2190865 total 0.9 bits
                                          sigma38p10 1.9 bits
                                              sigma38p10 1.3 bits

     .         *2190840  .         *2190830  .         *2190820  .         *2190810
5' c a t a a c c t t t a c t a c a g g g a t a a c g t t c a a t g a a t a a g 3'
                                                                proximalUP 3.8 bits
                                          distalUP 3.3 bits              |
                                                                         |- ... proximalUP-(32)
                                         |------------------------------------ ... distalUP-(51)-s
                                         |------------------------------------|- ... sigma38p10-prox

         .         *2190800  .         *2190790  .         *2190780  .
5' t a t t g g c t g _ c a g g a a t t a t t t t t c t t g c c t a t g g a t t 3'
                                                            sigma38p10 3.6 bits
... -------------------------------------------------------------| proximalUP-(32)-sigma38p10 21907
... -------------------------------------------------------------| distalUP-(51)-sigma38p10 2190778
... -------------------------------------------------------------| sigma38p10-proximalUP-distalUP 2

   *2190770  .         *2190760  .         *2190750
5' g g c t t c t c c c g c a t t c t c t t c a 3'
```

piece 31, NC_000913, yohF, config: linear, direction: -, begin: 2225440, end: 2225229

```
   *2225440   .          *2225430   .          *2225420   .          *2225410   .
5' c t g c g t g a g c g g g c c a a t g g c c t g t t a t t g c a a g g g c a 3'
   T timestamp:_2018Sep20_11-34-53

       *2225400   .          *2225390   .          *2225380   .          *2225370   .
5' g t g g c t g g a t g c c t c c a t t c a a c t c a c t g g t g c g t t g g 3'

           *2225360   .          *2225350   .          *2225340   .          *2225330
5' g c g g g g g g t a c a a a c g c t g a t g a t a t g c g c c t t c t a t a 3'

   .          *2225320   .          *2225310   .          *2225300   .          *2225290
5' c t t a a c g t t t a t t c a g c g t t a a g t g g a g a a c t c g a t g g 3'
                                        proximalUP 6.2 bits
                                                            ... proximalUP-(33
                              distalUP 5.5 bits
                                                            ... distalUP-(40)-s
                                                            ... sigma38p10-prox
                                                            ... proximalUP-(36
                                                            ... distalUP-(43)-s
                                                            ... sigma38p10-prox

       .          *2225280   .          *2225270   .          *2225260   .
5' c a c a g g t t g c g a t t a t t a c c g c c t c c g a t t c g g g g a t c 3'
                                        sigma38p10 2.9 bits
... ----------------| proximalUP-(33)-sigma38p10 2225280 Gap 4.0 bits
                                        sigma38p10 3.0 bits
... ----------------| distalUP-(40)-sigma38p10 2225280 Gap 3.1 bits
... ----------------| sigma38p10-proximalUP-distalUP 2225280 total 7.5 bits
... ---------------------| proximalUP-(36)-sigma38p10 2225277 Gap 3.6 bits
... ---------------------| distalUP-(43)-sigma38p10 2225277 Gap 3.6 bits
... ---------------------| sigma38p10-proximalUP-distalUP 2225277 total 7.6 bits
                                        proximalUP 3.3 bits
                                                            ... proximalUP-(34
                                        distalUP 7.1 bits
                                                            ... distalUP-(42)-s
```

|---------------|------------------------------------- ... sigma38p10-prox

*2225250 . *2225240 . *2225230

5′ g g c a a a g a g t g c g c g t t a t t a c 3′

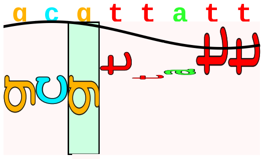

sigma38p10 0.3 bits

... ----------------------------| proximalUP-(34)-sigma38p10 2225236 Gap 2.5 bits

... ----------------------------| distalUP-(42)-sigma38p10 2225236 Gap 3.3 bits

... ----------------------------| sigma38p10-proximalUP-distalUP 2225236 total 5.0 bits piece 32, NC_000913, tfaS, config: linear, direction: +, begin: 2468627, end: 2468932

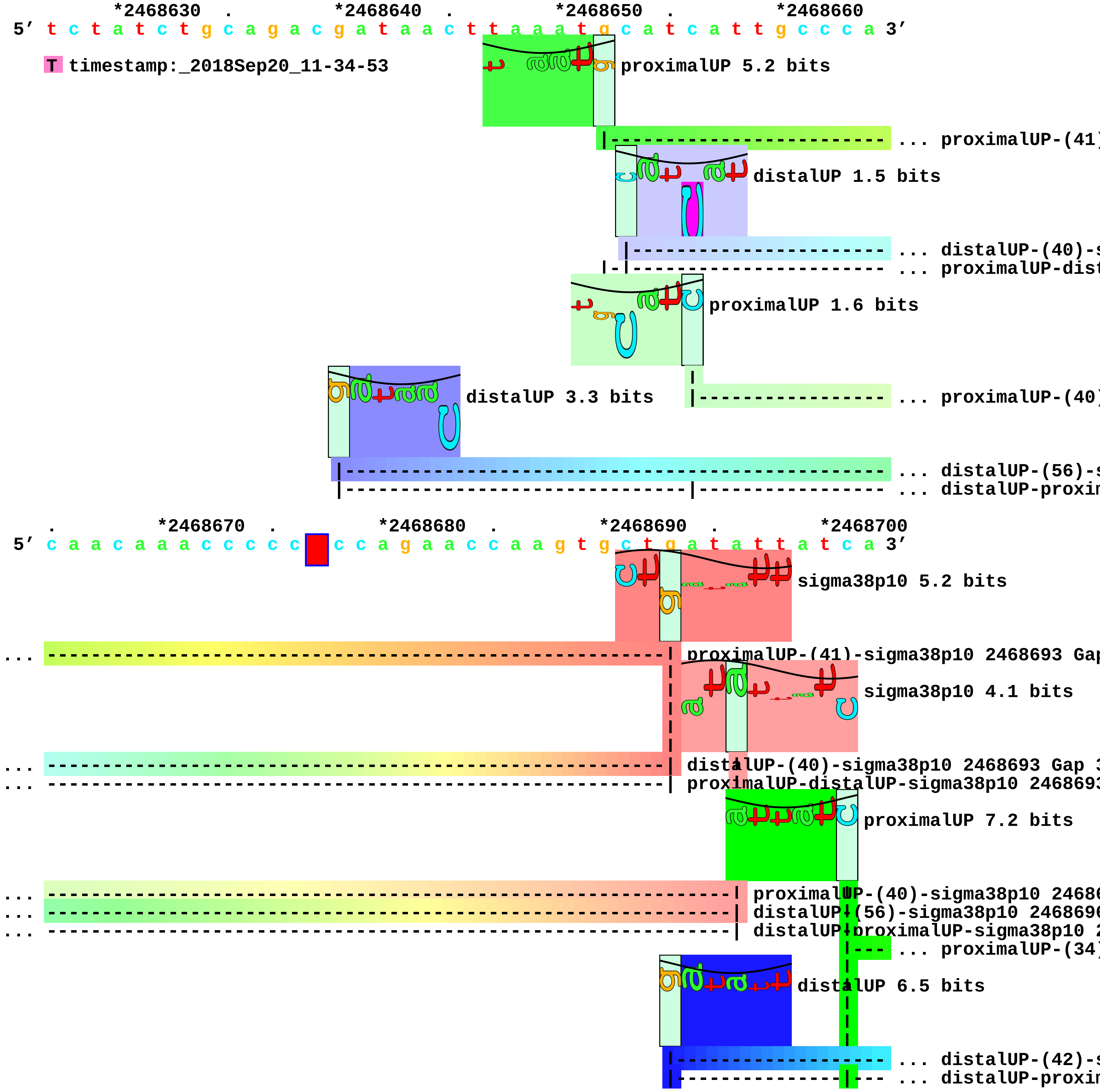

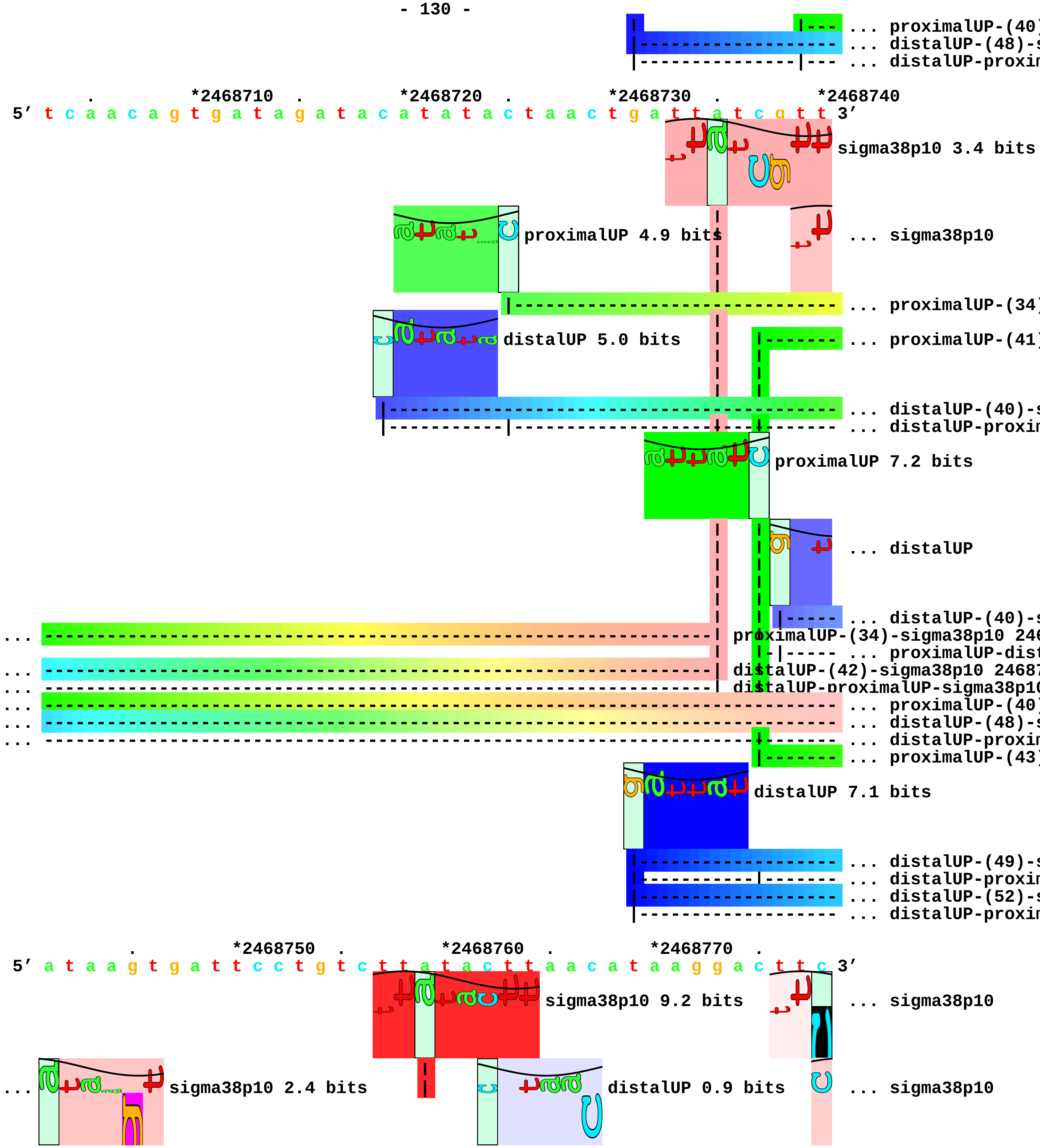
... proximalUP-(40)
... distalUP-(48)-s
... distalUP-proxim
. *2468710 . *2468720 . *2468730 . *2468740
5′ t c a a c a g t g a t a g a t a c a t a t a c t a a c t g a t t a t c g t t 3′
sigma38p10 3.4 bits
proximalUP 4.9 bits
... sigma38p10
... proximalUP-(34)
distalUP 5.0 bits
... proximalUP-(41)
... distalUP-(40)-s
... distalUP-proxim
proximalUP 7.2 bits
... distalUP
... distalUP-(40)-s
... proximalUP-(34)-sigma38p10 246
... proximalUP-dist
... distalUP-(42)-sigma38p10 2468
... distalUP-proximalUP-sigma38p10
... proximalUP-(40)
... distalUP-(48)-s
... distalUP-proxim
... proximalUP-(43)
distalUP 7.1 bits
... distalUP-(49)-s
... distalUP-proxim
... distalUP-(52)-s
... distalUP-proxim
. *2468750 . *2468760 . *2468770 .
5′ a t a a g t g a t t c c t g t c t t a t a c t t a a c a t a a g g a c t t c 3′
sigma38p10 9.2 bits
... sigma38p10
... sigma38p10 2.4 bits
distalUP 0.9 bits
... sigma38p10

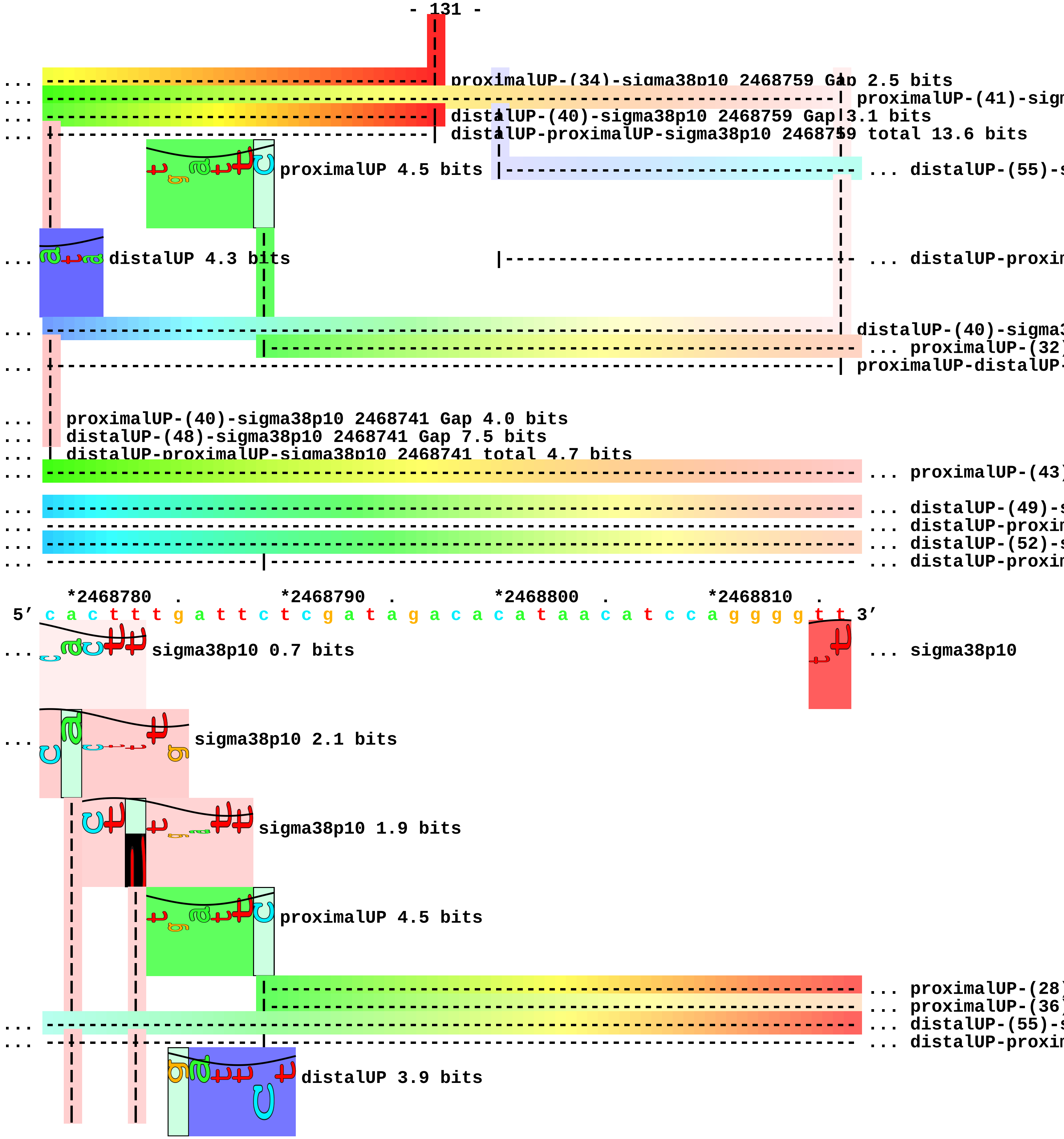
proximalUP-(34)-sigma38p10 2468759 Gap 2.5 bits
distalUP-(40)-sigma38p10 2468759 Gap 3.1 bits
distalUP-proximalUP-sigma38p10 2468759 total 13.6 bits
proximalUP 4.5 bits
distalUP 4.3 bits
proximalUP-(40)-sigma38p10 2468741 Gap 4.0 bits
distalUP-(48)-sigma38p10 2468741 Gap 7.5 bits
distalUP-proximalUP-sigma38p10 2468741 total 4.7 bits
*2468780 . *2468790 . *2468800 . *2468810 .
5′ c a c t t t g a t t c t c g a t a g a c a c a t a a c a t c c a g g g g t t 3′
sigma38p10 0.7 bits
sigma38p10
sigma38p10 2.1 bits
sigma38p10 1.9 bits
proximalUP 4.5 bits
distalUP 3.9 bits

... --------| proximalUP-(32)-sigma38p10 2468783 Gap 4.3 bits
|------------------------------------------------------------------ ... distalUP-(40)-s
|--------|---------------------------------------------------------- ... distalUP-proxin

... --| proximalUP-(43)-sigma38p10 2468780 Gap 5.1 bits

... --| distalUP-(49)-sigma38p10 2468780 Gap 5.9 bits
... --| distalUP-proximalUP-sigma38p10 2468780 total 5.4 bits
... --------| distalUP-(52)-sigma38p10 2468783 Gap 5.5 bits
... --------| distalUP-proximalUP-sigma38p10 2468783 total 3.7 bits

*2468820 . *2468830 . *2468840 . *2468850
5′ a t g t t t g t a t g c g t c c c g c g a c t a t c g c c c c a t t a a c g 3′

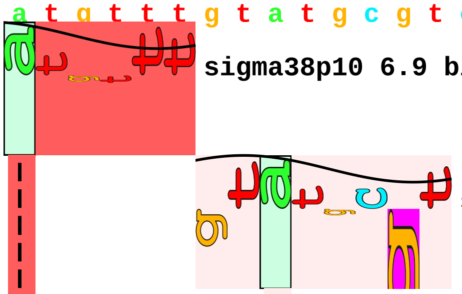

... sigma38p10 6.9 bits

sigma38p10 0.8 bits

... | proximalUP-(28)-sigma38p10 2468817 Gap 5.5 bits
... ----------------| proximalUP-(36)-sigma38p10 2468825 Gap 3.6 bits
... | distalUP-(55)-sigma38p10 2468817 Gap 4.9 bits
... | distalUP-proximalUP-sigma38p10 2468817 total 2.0 bits

... ----------------| distalUP-(40)-sigma38p10 2468825 Gap 3.1 bits
... ----------------| distalUP-proximalUP-sigma38p10 2468825 total 2.6 bits

. *2468860 . *2468870 . *2468880 . *2468890
5′ c c a t a c g a t a a a t g g g a t g g t g a g a a a g g g t g a c g g a 3′

. *2468900 . *2468910 . *2468920 . *2468930
5′ t a c c g a g g c a c a g c a t a g c g t c g c a g t a g a t g c a g c a g 3′

5′ a a 3′

piece 33, NC_000913, csiE, config: linear, direction: +, begin: 2663314, end: 2663551

```
     .          *2663320  .         *2663330  .         *2663340  .         *2663350
5' t g c t a a t c c a c t g g t c a g a a c a g t t t a a g a t g a g a a a a 3'
```

distalUP 0.4 bits

proximalUP 4.5 bits

T timestamp: 2018Sep20 11-34-53

... proximalUP-(27)

... distalUP-(53)-s

... distalUP-proxir

... proximalUP-(37)

distalUP 4.8 bits

... distalUP-(43)-s

... distalUP-proxir

... proximalUP

distalUP 2.7 bits

... distalUP-(40)-s

... distalUP-proxir

```
     .          *2663360  .         *2663370  .         *2663380  .
5' a t t c t g t g a c g c t t g c c a a c a t t t c t g a t g a t t a g c a t 3'
```

sigma38p10 4.0 bits

... proximalUP-(36)

... proximalUP-(27)-sigma38p10 2663369 Gap 3.8 bits

... distalUP-(53)-sigma38p10 2663369 Gap 4.9 bits

... distalUP-proximalUP-sigma38p10 2663369 total 0.2 bits

... proximalUP-(37)-sigma38p10 2663379 Gap 4

sigma38p10 1.3 bits

... distalUP-(43)-sigma38p10 2663379 Gap 3.6

... distalUP-proximalUP-sigma38p10 2663379 t

... proximalUP 5.5 bits

distalUP 1.6 bits

... sigma38p10

proximalUP-(30)-sigma38p10 2

... distalUP-(40)-sigma38p10 266

... distalUP-proximalUP-sigma38p

proximalUP 2.6 bits

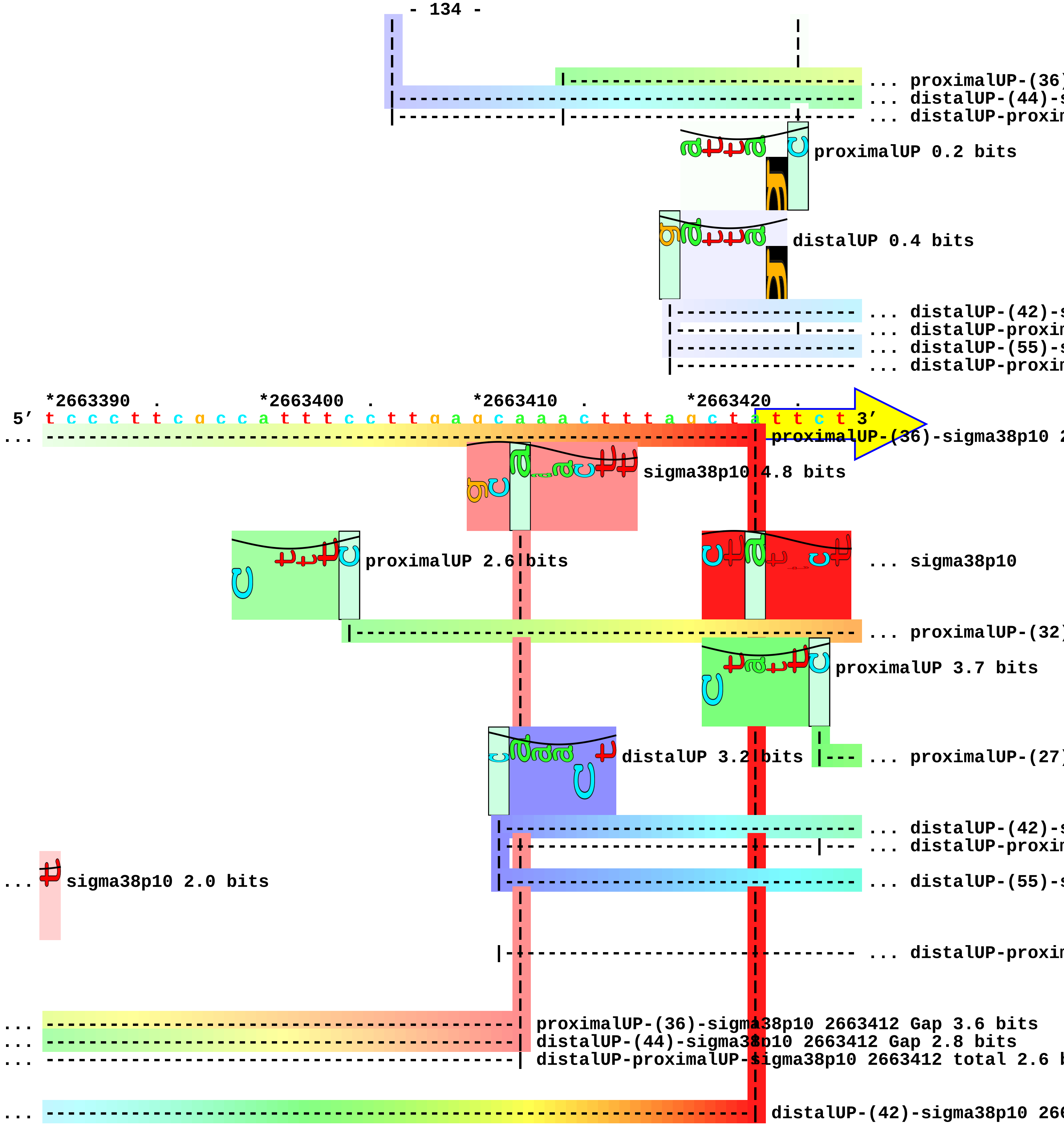
... proximalUP-(36)
... distalUP-(44)-s
... distalUP-proxim
proximalUP 0.2 bits
distalUP 0.4 bits
... distalUP-(42)-s
... distalUP-proxim
... distalUP-(55)-s
... distalUP-proxim
*2663390 . *2663400 . *2663410 . *2663420 .
5′ t c c c t t c g c c a t t t c c t t g a g c a a a c t t t a g c t a t t c t 3′
proximalUP-(36)-sigma38p10 2
sigma38p10 4.8 bits
proximalUP 2.6 bits
... sigma38p10
... proximalUP-(32)
proximalUP 3.7 bits
distalUP 3.2 bits
... proximalUP-(27)
... distalUP-(42)-s
... distalUP-proxim
sigma38p10 2.0 bits
... distalUP-(55)-s
... distalUP-proxim
proximalUP-(36)-sigma38p10 2663412 Gap 3.6 bits
distalUP-(44)-sigma38p10 2663412 Gap 2.8 bits
distalUP-proximalUP-sigma38p10 2663412 total 2.6 b
distalUP-(42)-sigma38p10 266

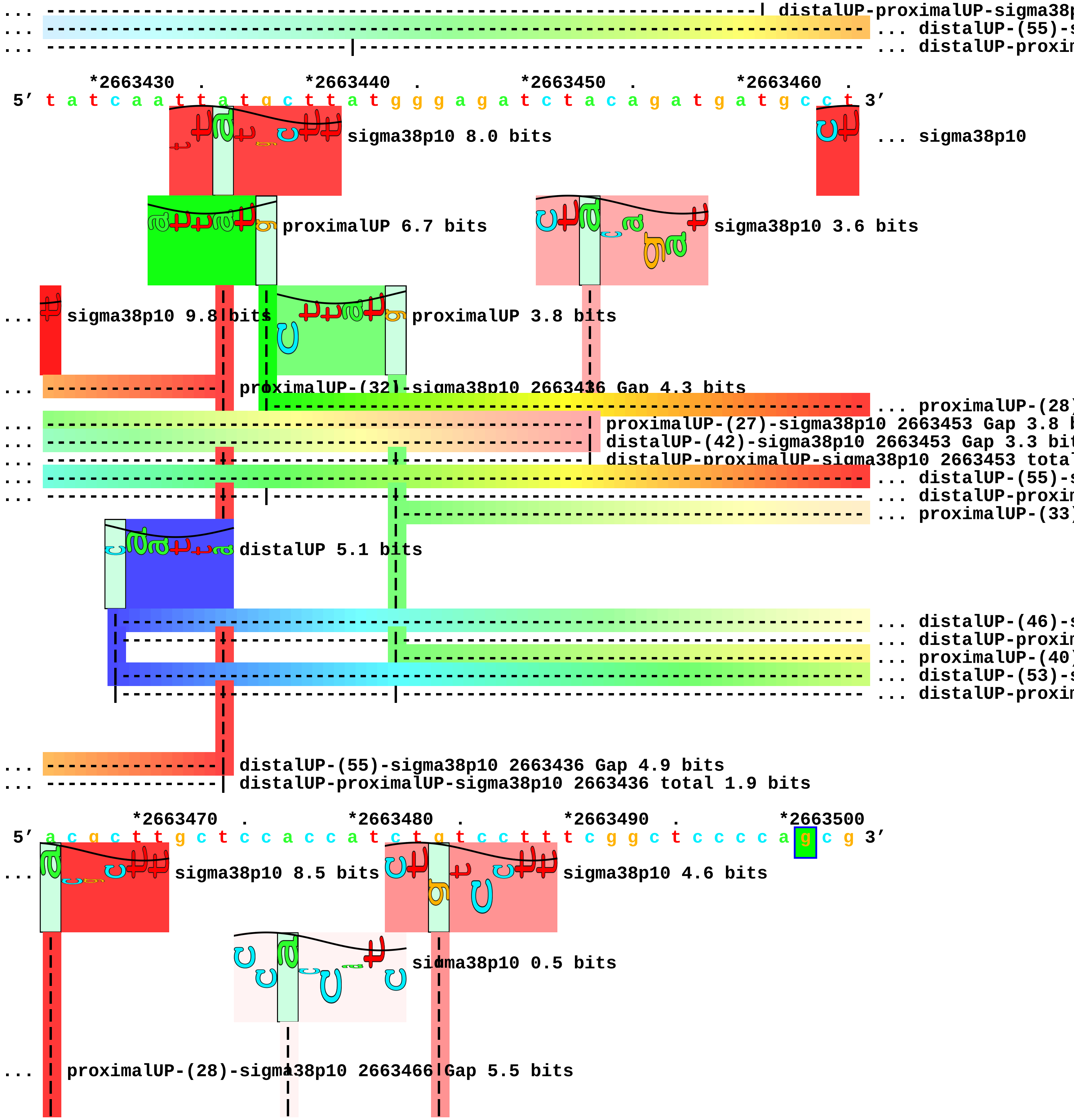
... ----------------------------------------------------------------------| distalUP-proximalUP-sigma38p
... ---------------------------------------------------------------------------- ... distalUP-(55)-s
... ---------------------------|------------------------------------------------ ... distalUP-proxin
*2663430 . *2663440 . *2663450 . *2663460 .
5′ t a t c a a t t a t g c t t a t g g g a g a t c t a c a g a t g a t g c c t 3′
sigma38p10 8.0 bits
... sigma38p10
proximalUP 6.7 bits
sigma38p10 3.6 bits
... sigma38p10 9.8 bits
proximalUP 3.8 bits
... ----------------| proximalUP-(32)-sigma38p10 2663436 Gap 4.3 bits
... proximalUP-(28)
... proximalUP-(27)-sigma38p10 2663453 Gap 3.8 b
... distalUP-(42)-sigma38p10 2663453 Gap 3.3 bit
... distalUP-proximalUP-sigma38p10 2663453 tota
... distalUP-(55)-s
... distalUP-proxin
... proximalUP-(33)
distalUP 5.1 bits
... distalUP-(46)-s
... distalUP-proxin
... proximalUP-(40)
... distalUP-(53)-s
... distalUP-proxin
... ----------------| distalUP-(55)-sigma38p10 2663436 Gap 4.9 bits
... ----------------| distalUP-proximalUP-sigma38p10 2663436 total 1.9 bits
*2663470 . *2663480 . *2663490 . *2663500
5′ a c g c t t g c t c c a c c a t c t g t c c t t t c g g c t c c c c a g c g 3′
... sigma38p10 8.5 bits
sigma38p10 4.6 bits
sigma38p10 0.5 bits
... proximalUP-(28)-sigma38p10 2663466 Gap 5.5 bits

```
    |                      |             |
... | distalUP-(55)-sigma38p10 2663466 Gap 4.9 bits
... | distalUP-proximalUP-sigma38p10 2663466 total 8.0 bits
... ----------------------| proximalUP-(33)-sigma38p10 2663477 Gap 4.0 bits
                          |              |
... ----------------------| distalUP-(46)-sigma38p10 2663477 Gap 4.1 bits
... ----------------------| distalUP-proximalUP-sigma38p10 2663477 total 1.3 bits
... ------------------------------------| proximalUP-(40)-sigma38p10 2663484 Gap 4.0 bits
... ------------------------------------| distalUP-(53)-sigma38p10 2663484 Gap 4.9 bits
... ------------------------------------| distalUP-proximalUP-sigma38p10 2663484 total 4.7 bits

      .         *2663510 .         *2663520  .         *2663530  .         *2663540
 5' c c g c t g c c a g a t c t t g c t g a c g c t c t t t c a g c c g g g g t 3'

          .         *2663550
 5' t g a c c g c c a c 3'
```

piece 34, NC_000913, raiA, config: linear, direction: +, begin: 2734860, end: 2735131

```
   *2734860   .         *2734870   .         *2734880   .         *2734890   .
5' t g a a a a a g t a g c g a a a a t c a t c g c c g c a a a c a g c a g c a 3'
   T timestamp:_2018Sep20_11-34-53

       *2734900   .         *2734910   .         *2734920   .         *2734930   .
5' a t a c a t a a c a g a a a c c t g a a a c a c a a a a c g g c a g c c c t 3'
            proximalUP 3.8 bits
                                                    ... proximalUP-(33
      distalUP 2.5 bits                             ... proximalUP-(34)
                                                    ... distalUP-(44)-s
                                                    ... distalUP-proxim
                   proximalUP 3.3 bits
                 distalUP 0.3 bits
                                                    ... distalUP-(40)-s
                                                    ... distalUP-proxim

         *2734940   .         *2734950   .         *2734960   .         *2734970
5' t g a g c t g c c g t t t t t t t a t t c t g t c a g t t g t g a a a c t g 3'
            sigma38p10 2.5 bits
... ------------------| proximalUP-(33)-sigma38p10 2734945 Gap 4.0 bits
... ----------------------------------| proximalUP-(34)-sigma38p10 2734953 Gap 2.5 bits
... ------------------| distalUP-(44)-sigma38p10 2734945 Gap 2.8 bits
... ------------------| distalUP-proximalUP-sigma38p10 2734945 total 1.9 bits
                         sigma38p10 5.5 bits
                       proximalUP 6.2 bits
... ----------------------------------| dist[a]lUP-(40)-sigma38p10 2734953 Gap 3.1 bits
... ----------------------------------| dist[a]lUP-proximalUP-sigma38p10 2734953 total 3.7 bits
                                         |--------------------------- ... proximalUP-(34)
            distalUP 4.9 bits
```

... distalUP-(45)-s
... distalUP-proxim

. *2734980 . *2734990 . *2735000 . *2735010
5′ a a g c g a t t t a g t c g c t a t c g a t c t c a t c a a a t a t g g c t 3′

sigma38p10 1.9 bits

... proximalUP-(34)-sigma38p10 2734990 Gap 2.5 bits
... distalUP-(45)-sigma38p10 2734990 Gap 5.5 bits
... distalUP-proximalUP-sigma38p10 2734990 total 5.0 bits

. *2735020 . *2735030 . *2735040 .
5′ c g c t t t g a g a t a t t c c t c a a g t a a a a a a a c a t c t c t t c 3′

*2735050 . *2735060 . *2735070 . *2735080 .
5′ c t g c g a t t t c t c a c a a a a a a g a t t c g t t g a c a a a a a g t 3′

proximalUP 5.0 bits
... proximalUP-(27)
... proximalUP-(37)
distalUP 5.7 bits
proximalUP 3.8 bits
... distalUP-(48)-s
... distalUP-proxim
... distalUP-(50)-s
... distalUP-proxim
... proximalUP-(41)
... distalUP-(52)-s
... distalUP-proxim

*2735090 . *2735100 . *2735110 . *2735120 .
5′ g a c a a a a t t a t g a g a t t t t c a t c a c a c a t t t t g a c a t c 3′
... proximalUP-(27)-sigma38p10 2735113 Gap 3.8 b
... proximalUP-(37)-sigma38p10 2735111 Gap 4.5 bits

sigma38p10 1.9 bits

... distalUP-(48)-sigma38p10 2735111 Gap 7.5 bits
... distalUP-proximalUP-sigma38p10 2735111 total 0.7
... distalUP-(50)-sigma38p10 2735113 Gap 4.1 bit
... distalUP-proximalUP-sigma38p10 2735113 total

... ------------------------------------------------------------| proximalUP-(41)-sigma38p10 2735115 Gap 5
... ------------------------------------------------------------| distalUP-(52)-sigma38p10 2735115 Gap 5.5
... ------------------------------------------------------------| distalUP-proximalUP-sigma38p10 2735115 t

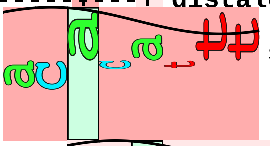

sigma38p10 3.5 bits

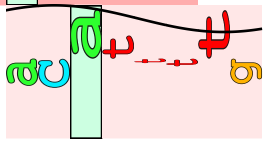

sigma38p10 1.0 bits

*2735130
5’ a g g a a c 3’

piece 35, NC_000913, ssrA, config: linear, direction: +, begin: 2753452, end: 2753757

```
        .         *2753460  .         *2753470  .         *2753480  .
5' g c g c g a a t g a a c a t c t t a t t g g c t a t c a c a t c c g a c a c 3'
 T timestamp:_2018Sep20_11-34-53

 *2753490  .         *2753500  .         *2753510  .         *2753520  .
5' a a a t g t t g c c a t c c c a t t g c t t a a t c g a a t a a a a a t c a 3'

   *2753530  .         *2753540  .         *2753550  .         *2753560  .
5' g g c t a c a t g g g t g c t a a a t c t t t a a c g a t a a c g c c a t t 3'
                                            proximalUP 5.7 bits
                                                     |---------------------------------- ... proximalUP-(38)
                                        distalUP 4.8 bits
                                        |------------------------------------------------ ... distalUP-(44)-s
                                        |-------------|---------------------------------- ... distalUP-proxin
                                                              proximalUP 5.0 bits
                                                                     |-------------------- ... proximalUP-(43)
                                                                        distalUP 3.3 bits
                                                                       |------------------ ... distalUP-(42)-s
                                                                     |-|------------------ ... proximalUP-dist

     *2753570  .         *2753580  .         *2753590  .         *2753600
5' g a g g c t g g t c a t g g c g c t c a t a a a t c t g g t a t a c t t a c 3'
                                                    sigma38p10 2.6 bits
... --------------------------------------| proximalUP-(38)-sigma38p10 2753585 Gap 3.2 bits
                                                                   sigma38p10 8.3 bits
... --------------------------------------| distalUP-(44)-sigma38p10 2753585 Gap 2.8 bits
... --------------------------------------| distalUP-proximalUP-sigma38p10 2753585 total 7.0 bits
... ------------------------------------------------------------| proximalUP-(43)-sigma38p10 2753596
... ------------------------------------------------------------| distalUP-(42)-sigma38p10 2753596 G
... ------------------------------------------------------------| proximalUP-distalUP-sigma38p10 275

 .         *2753610  .         *2753620  .         *2753630  .         *2753640
5' c t t t a c a c a t t g g g g c t g a t t c t g g a t t c g a c g g g a t t 3'
```

```
        .         *2753650  .         *2753660  .         *2753670  .
5' t g c g a a a c c c a a g g t g c a t g c c g a g g g g c g g t t g g c c t 3'

  *2753680  .         *2753690  .         *2753700  .         *2753710  .
5' c g t a a a a a g c c g c a a a a a a t a g t c g c a a a c g a c g a a a a 3'
                                                   proximalUP 3.3 bits
                                                     ... proximalUP-(34)
                      distalUP 5.7 bits
                                                     ... distalUP-(43)-s
                                                     ... distalUP-proxin
                                                     ... proximalUP-(37)
                                                     ... distalUP-(46)-s
                                                     ... distalUP-proxin

    *2753720  .         *2753730  .         *2753740  .         *2753750  .
5' c t a c g c t t t a g c a g c t t a a t a a c c t g c t t a g a g c c c t c 3'
                                    sigma38p10 0.8 bits
... ------------------------------------| proximalUP-(34)-sigma38p10 2753735 Gap 2.5 bits
                                           sigma38p10 2.6 bits
... ------------------------------------| distalUP-(43)-sigma38p10 2753735 Gap 3.6 bits
... ------------------------------------| distalUP-proximalUP-sigma38p10 2753735 total 3.8 bits
... ------------------------------------| proximalUP-(37)-sigma38p10 2753738 Gap 4.5 bits
... ------------------------------------| distalUP-(46)-sigma38p10 2753738 Gap 4.1 bits
... ------------------------------------| distalUP-proximalUP-sigma38p10 2753738 total 2.9 bits

5' t c 3'
```

piece 36, NC_000913, yfjJ, config: linear, direction: +, begin: 2758250, end: 2759049

*2758250 . *2758260 . *2758270 . *2758280 .
5′ t t t c t t c c t g c a t c a a g a a a a a t a t t a t c t t a c a a t t t 3′
T timestamp:_2018Sep20_11-34-53

*2758290 . *2758300 . *2758310 . *2758320 .
5′ g g g c c a a a t a a a a t t a g a a a t c g t g a c a a g g c a a a t g a 3′

proximalUP 4.4 bits
proximalUP 2.7 bits
... proximalUP-(29[illegible]
distalUP 5.1 bits
... proximalUP-(35[illegible]
... distalUP-(41)-[illegible]
... distalUP-proxi[illegible]
... proximalUP-(31[illegible]
... distalUP-(43)-[illegible]
... distalUP-proxi[illegible]
... proximalUP-(36[illegible]
... distalUP-(48)-[illegible]
... distalUP-proxi[illegible]
... proximalUP-(37[illegible]
... distalUP-(49)-[illegible]
... distalUP-proxi[illegible]
distalUP 2.8 bits
... distalUP
... distalUP-(55)-[illegible]
... distalUP-proxi[illegible]
... distalUP-(41)-[illegible]
... distalUP-proxi[illegible]

*2758330 . *2758340 . *2758350 . *2758360
5′ a t t a a t t a g a a t c t t a a t t t c a c a a a a c a a a a t a t t t a 3′

sigma38p10 2.1 bits
... sigma38p10
... proximalUP-(29)-sigma38p10 2758333 Gap 7.5 bits
... sigma38p10
... proximalUP-(35)-sigma38p10 [illegible]
... distalUP-(41)-sigma38p10 2758333 Gap 3.1 bits
... proximalUP-(31[illegible]
... distalUP-proximalUP-sigma38p10 2758333 total 1.0 bits
... proximalUP
... proximalUP-(31)-sigma38p10 2758335 Gap 4.7 bits
... distalUP-(43)-sigma38p10 2758335 Gap 3.6 bits

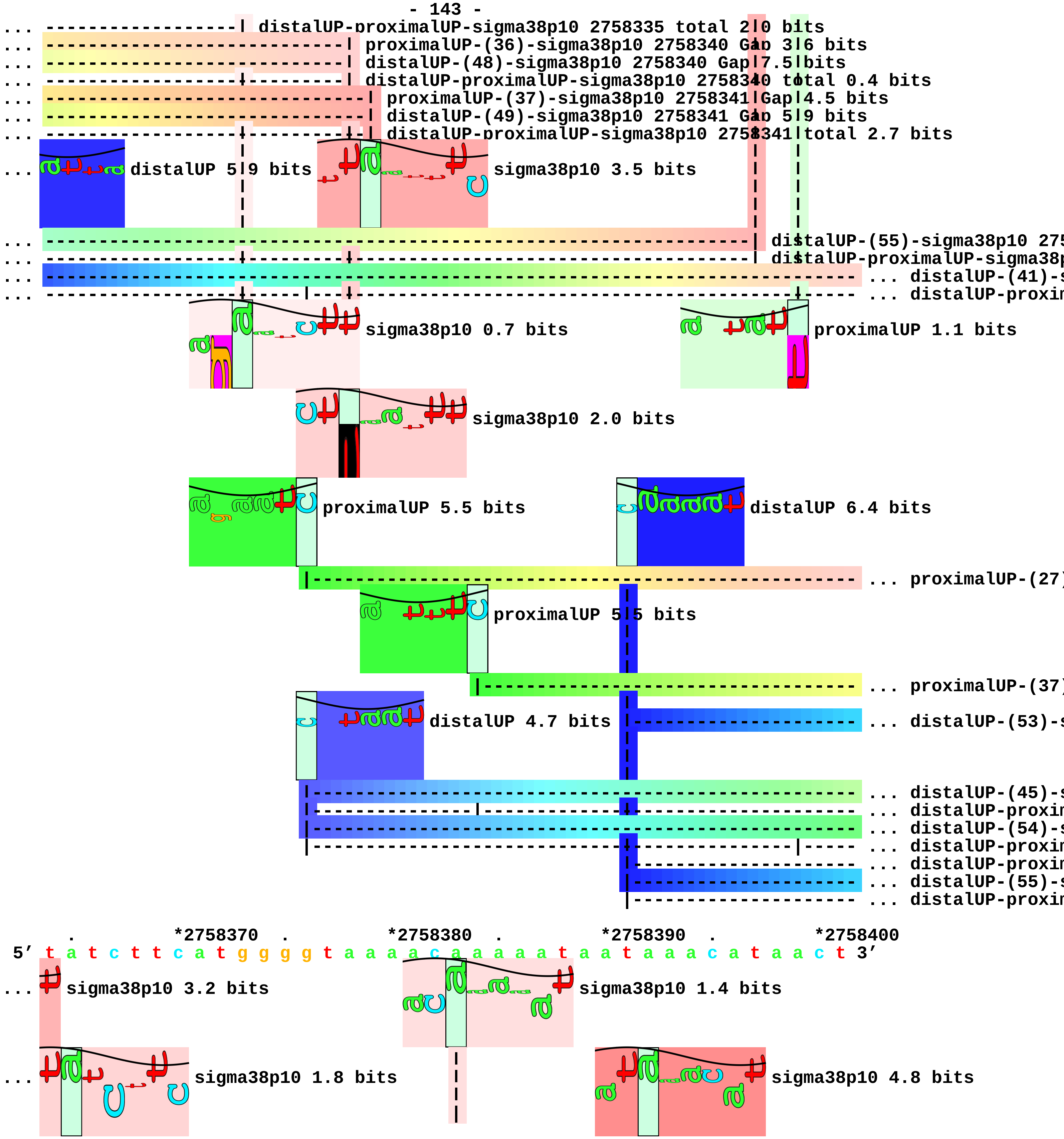
distalUP-proximalUP-sigma38p10 2758335 total 2.0 bits
proximalUP-(36)-sigma38p10 2758340 Gap 3.6 bits
distalUP-(48)-sigma38p10 2758340 Gap 7.5 bits
distalUP-proximalUP-sigma38p10 2758340 total 0.4 bits
proximalUP-(37)-sigma38p10 2758341 Gap 4.5 bits
distalUP-(49)-sigma38p10 2758341 Gap 5.9 bits
distalUP-proximalUP-sigma38p10 2758341 total 2.7 bits
distalUP 5.9 bits
sigma38p10 3.5 bits
distalUP-(55)-sigma38p10 275
distalUP-proximalUP-sigma38p
distalUP-(41)-s
distalUP-proxim
sigma38p10 0.7 bits
proximalUP 1.1 bits
sigma38p10 2.0 bits
proximalUP 5.5 bits
distalUP 6.4 bits
proximalUP-(27)
proximalUP 5.5 bits
proximalUP-(37)
distalUP 4.7 bits
distalUP-(53)-s
distalUP-(45)-s
distalUP-proxim
distalUP-(54)-s
distalUP-proxim
distalUP-proxim
distalUP-(55)-s
distalUP-proxim
*2758370
*2758380
*2758390
*2758400
5’ t a t c t t c a t g g g g t a a a a c a a a a a t a a t a a a c a t a a c t 3’
sigma38p10 3.2 bits
sigma38p10 1.4 bits
sigma38p10 1.8 bits
sigma38p10 4.8 bits

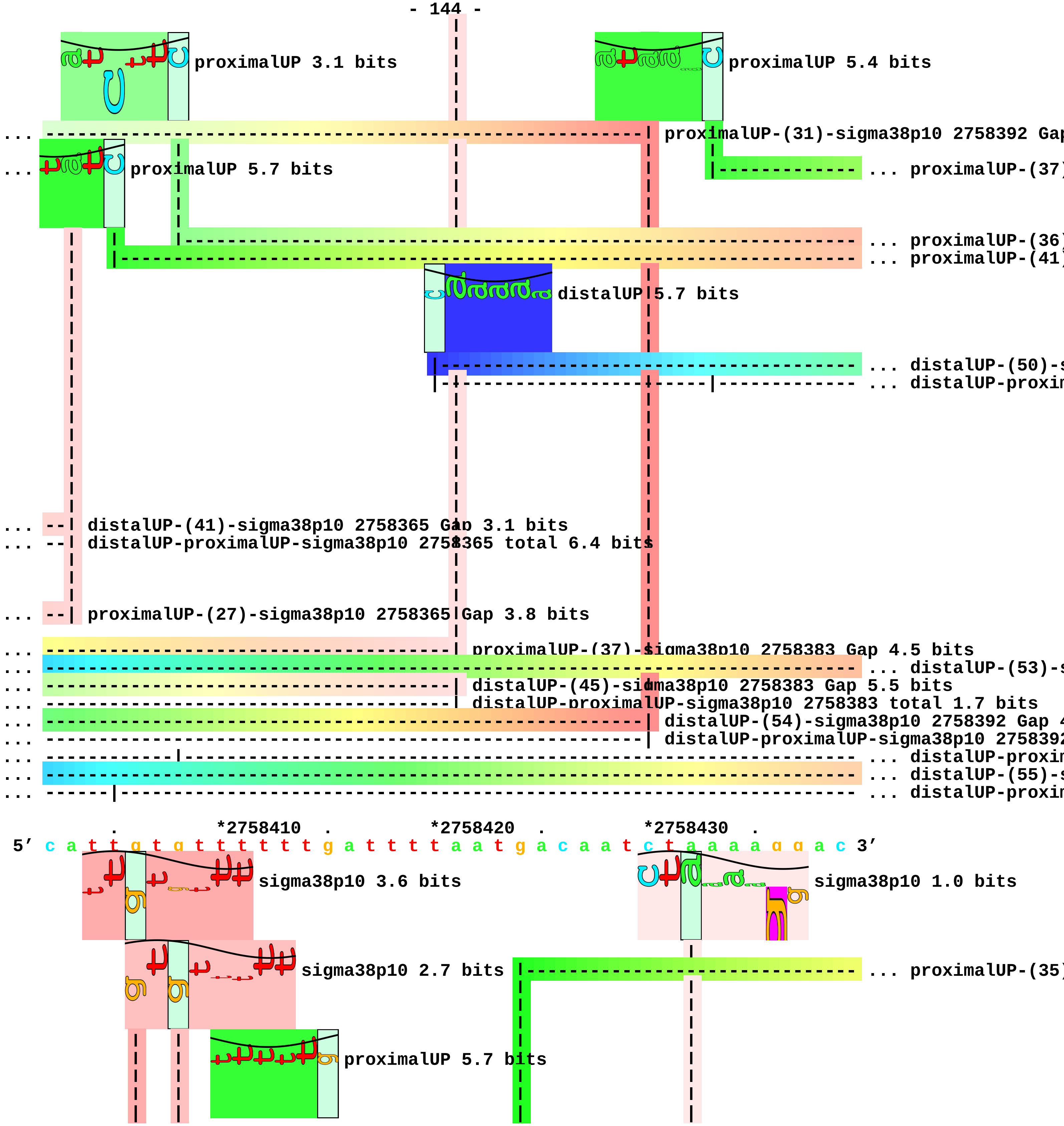
proximalUP 3.1 bits
proximalUP 5.4 bits
... proximalUP-(31)-sigma38p10 2758392 Gap
... proximalUP 5.7 bits
... proximalUP-(37)
... proximalUP-(36)
... proximalUP-(41)
distalUP 5.7 bits
... distalUP-(50)-s
... distalUP-proxin
... distalUP-(41)-sigma38p10 2758365 Gap 3.1 bits
... distalUP-proximalUP-sigma38p10 2758365 total 6.4 bits
... proximalUP-(27)-sigma38p10 2758365 Gap 3.8 bits
... proximalUP-(37)-sigma38p10 2758383 Gap 4.5 bits
... distalUP-(53)-s
... distalUP-(45)-sigma38p10 2758383 Gap 5.5 bits
... distalUP-proximalUP-sigma38p10 2758383 total 1.7 bits
... distalUP-(54)-sigma38p10 2758392 Gap 4
... distalUP-proximalUP-sigma38p10 2758392
... distalUP-proxin
... distalUP-(55)-s
... distalUP-proxin
*2758410
*2758420
*2758430
5′ c a t t g t g t t t t t t g a t t t t a a t g a c a a t c t a a a a g g a c 3′
sigma38p10 3.6 bits
sigma38p10 1.0 bits
sigma38p10 2.7 bits
... proximalUP-(35)
proximalUP 5.7 bits

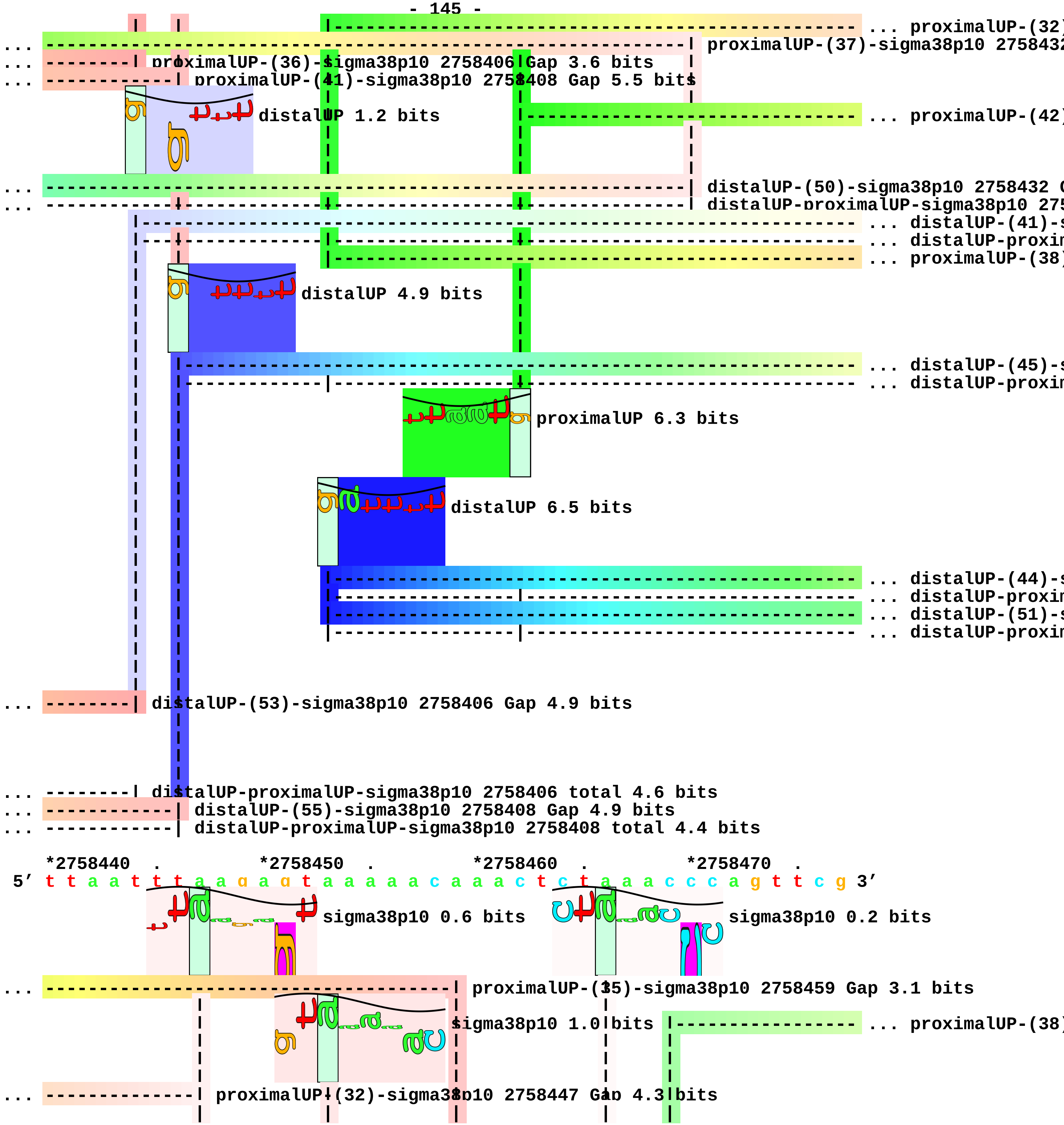
... proximalUP-(32)
... proximalUP-(37)-sigma38p10 2758432
... proximalUP-(36)-sigma38p10 2758406 Gap 3.6 bits
... proximalUP-(41)-sigma38p10 2758408 Gap 5.5 bits
distalUP 1.2 bits
... proximalUP-(42)
... distalUP-(50)-sigma38p10 2758432
... distalUP-proximalUP-sigma38p10 275
... distalUP-(41)-
... distalUP-proxim
... proximalUP-(38)
distalUP 4.9 bits
... distalUP-(45)-
... distalUP-proxim
proximalUP 6.3 bits
distalUP 6.5 bits
... distalUP-(44)-
... distalUP-proxim
... distalUP-(51)-
... distalUP-proxim
... distalUP-(53)-sigma38p10 2758406 Gap 4.9 bits
... distalUP-proximalUP-sigma38p10 2758406 total 4.6 bits
... distalUP-(55)-sigma38p10 2758408 Gap 4.9 bits
... distalUP-proximalUP-sigma38p10 2758408 total 4.4 bits
*2758440 . *2758450 . *2758460 . *2758470 .
5’ t t a a t t t a a g a g t a a a a a c a a a c t c t a a a c c c a g t t c g 3’
sigma38p10 0.6 bits
sigma38p10 0.2 bits
... proximalUP-(35)-sigma38p10 2758459 Gap 3.1 bits
sigma38p10 1.0 bits
... proximalUP-(38)
... proximalUP-(32)-sigma38p10 2758447 Gap 4.3 bits

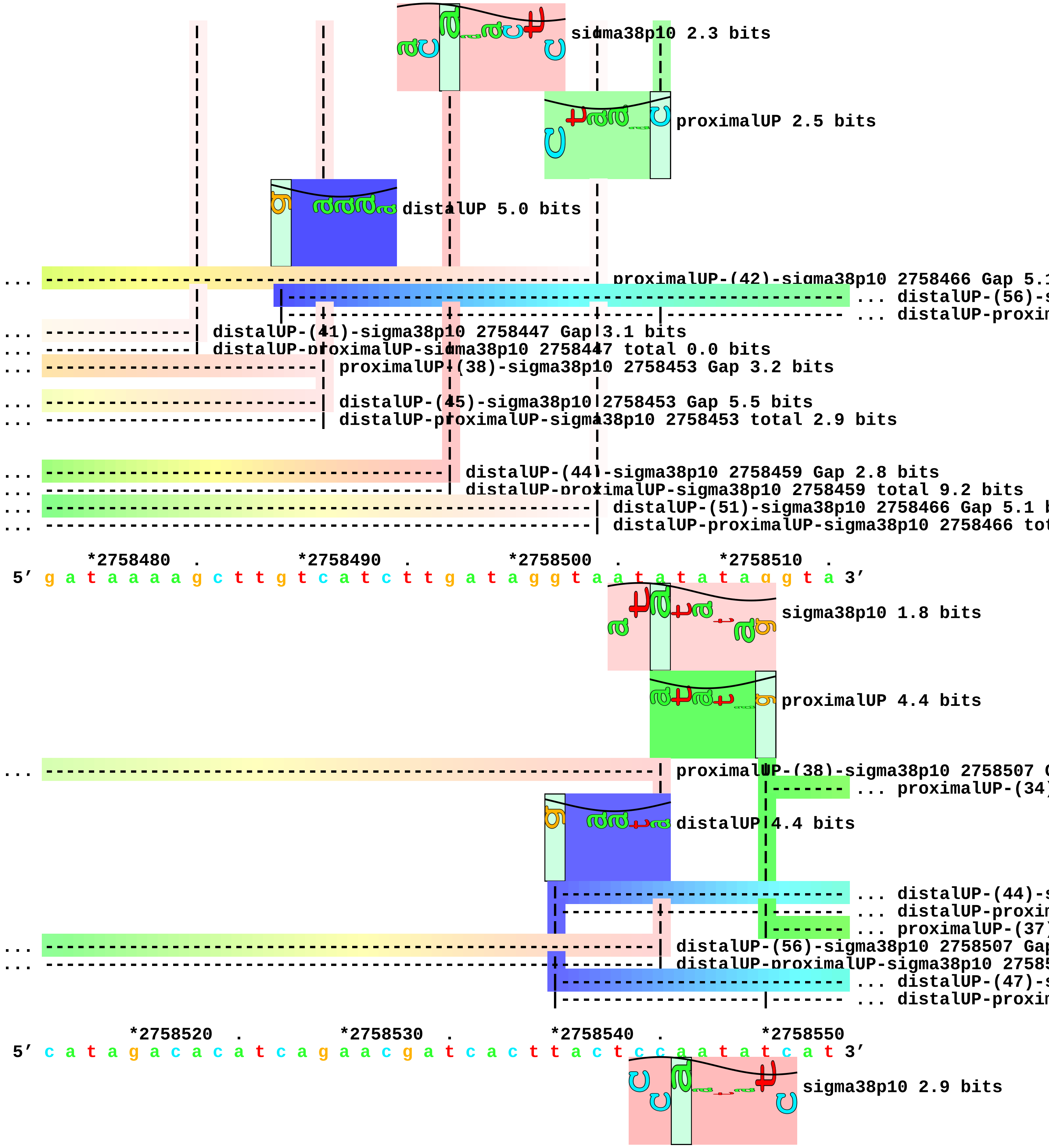

sigma38p10 2.3 bits
proximalUP 2.5 bits
distalUP 5.0 bits
... proximalUP-(42)-sigma38p10 2758466 Gap 5.1
... distalUP-(56)-s
... distalUP-proxim
... distalUP-(41)-sigma38p10 2758447 Gap 3.1 bits
... distalUP-proximalUP-sigma38p10 2758447 total 0.0 bits
... proximalUP-(38)-sigma38p10 2758453 Gap 3.2 bits
... distalUP-(45)-sigma38p10 2758453 Gap 5.5 bits
... distalUP-proximalUP-sigma38p10 2758453 total 2.9 bits
... distalUP-(44)-sigma38p10 2758459 Gap 2.8 bits
... distalUP-proximalUP-sigma38p10 2758459 total 9.2 bits
... distalUP-(51)-sigma38p10 2758466 Gap 5.1 b
... distalUP-proximalUP-sigma38p10 2758466 tot
*2758480 . *2758490 . *2758500 . *2758510 .
5' g a t a a a a g c t t g t c a t c t t g a t a g g t a a t a t a t a g g t a 3'
sigma38p10 1.8 bits
proximalUP 4.4 bits
... proximalUP-(38)-sigma38p10 2758507 G
... proximalUP-(34)
distalUP 4.4 bits
... distalUP-(44)-s
... distalUP-proxim
... proximalUP-(37)
... distalUP-(56)-sigma38p10 2758507 Gap
... distalUP-proximalUP-sigma38p10 27585
... distalUP-(47)-s
... distalUP-proxim
*2758520 . *2758530 . *2758540 . *2758550
5' c a t a g a c a c a t c a g a a c g a t c a c t t a c t c c a a t a t c a t 3'
sigma38p10 2.9 bits

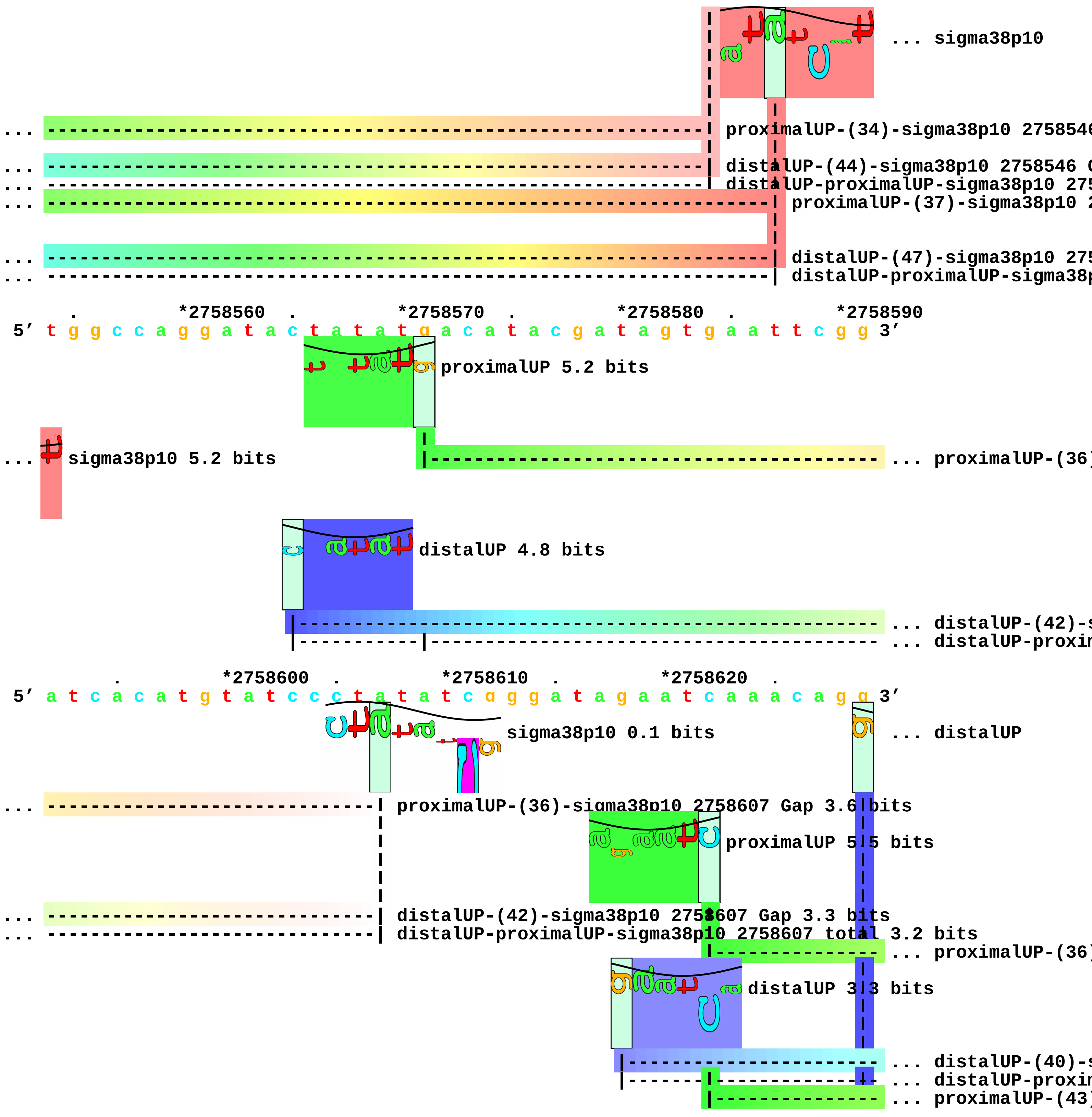
... sigma38p10
... proximalUP-(34)-sigma38p10 275854
... distalUP-(44)-sigma38p10 2758546
... distalUP-proximalUP-sigma38p10 275
... proximalUP-(37)-sigma38p10 2
... distalUP-(47)-sigma38p10 275
... distalUP-proximalUP-sigma38p
*2758560
*2758570
*2758580
*2758590
5' t g g c c a g g a t a c t a t a t g a c a t a c g a t a g t g a a t t c g g 3'
proximalUP 5.2 bits
... sigma38p10 5.2 bits
... proximalUP-(36)
distalUP 4.8 bits
... distalUP-(42)-s
... distalUP-proxin
*2758600
*2758610
*2758620
5' a t c a c a t g t a t c c c t a t a t c g g g a t a g a a t c a a a c a g g 3'
sigma38p10 0.1 bits
... distalUP
... proximalUP-(36)-sigma38p10 2758607 Gap 3.6 bits
proximalUP 5.5 bits
... distalUP-(42)-sigma38p10 2758607 Gap 3.3 bits
... distalUP-proximalUP-sigma38p10 2758607 total 3.2 bits
... proximalUP-(36)
distalUP 3.3 bits
... distalUP-(40)-s
... distalUP-proxin
... proximalUP-(43)

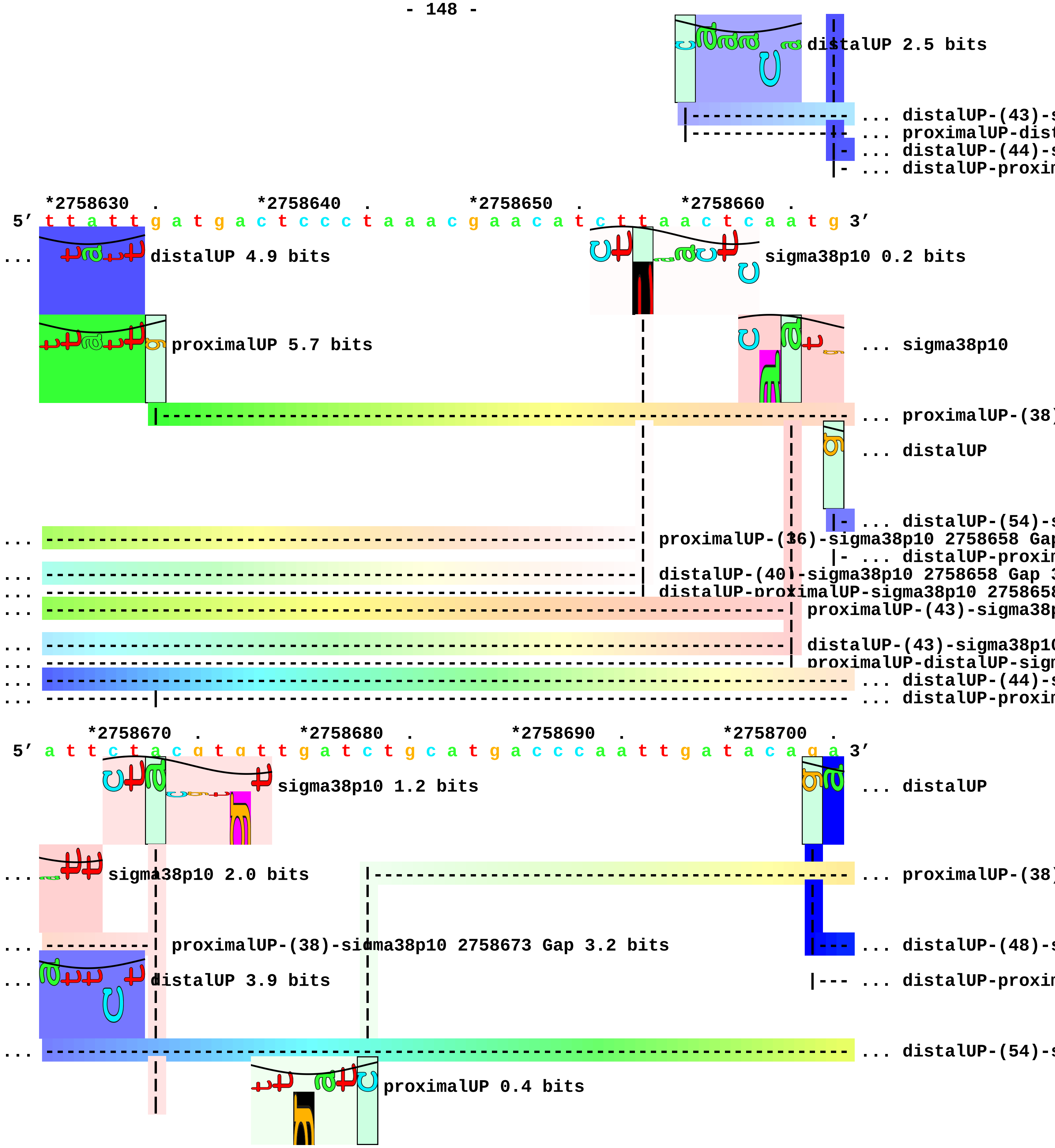
distalUP 2.5 bits
... distalUP-(43)-s
... proximalUP-dist
... distalUP-(44)-s
... distalUP-proxim
*2758630 . *2758640 . *2758650 . *2758660 .
5′ t t a t t g a t g a c t c c c t a a a c g a a c a t c t t a a c t c a a t g 3′
... distalUP 4.9 bits
sigma38p10 0.2 bits
proximalUP 5.7 bits
... sigma38p10
... proximalUP-(38)
... distalUP
... distalUP-(54)-s
... proximalUP-(36)-sigma38p10 2758658 Gap
... distalUP-proxim
... distalUP-(40)-sigma38p10 2758658 Gap 3
... distalUP-proximalUP-sigma38p10 275865
... proximalUP-(43)-sigma38p
... distalUP-(43)-sigma38p10
... proximalUP-distalUP-sigm
... distalUP-(44)-s
... distalUP-proxim
*2758670 . *2758680 . *2758690 . *2758700 .
5′ a t t c t a c g t g t t g a t c t g c a t g a c c c a a t t g a t a c a g a 3′
sigma38p10 1.2 bits
... distalUP
... sigma38p10 2.0 bits
... proximalUP-(38)
... proximalUP-(38)-sigma38p10 2758673 Gap 3.2 bits
... distalUP-(48)-s
... distalUP 3.9 bits
... distalUP-proxim
... distalUP-(54)-s
proximalUP 0.4 bits

... ----------|---------------------------------------------------- ... distalUP-proxin

... ----------| distalUP-(44)-sigma38p10 2758673 Gap 2.8 bits
... ----------| distalUP-proximalUP-sigma38p10 2758673 total 5.8 bits

*2758710 . *2758720 . *2758730 . *2758740
5' a a a t a t g g a t a a c c c a t t c t t t c a a c c c a g g g t t g a c t 3'
... distalUP 7.2 bits sigma38p10 6.9 bits
... ------------------------------| proximalUP-(38)-sigma38p10 2758721 Gap 3.2 bits
... -------------------------------------------------------------------------- ... distalUP-(48)-s
... ------------|-----------------|------------------------------------------- ... distalUP-proxin
... ------------------------------| distalUP-(54)-sigma38p10 2758721 Gap 4.9 bits
proximalUP 5.6 bits
... ------------|-----------------| distalUP-proximalUP-sigma38p10 2758721 total 3.1 bits
|------------------------------------------------------------- ... proximalUP-(40)

. *2758750 . *2758760 . *2758770 . *2758780
5' c t g g t g c t a t a t c t c g c t t t a c c a g t g c g t t a a a a g c a 3'
sigma38p10 2.6 bits proximalUP 3.4 bits
proximalUP 5.7 bits |----- ... proximalUP-(27)
... ----------------| distalUP-(48)-sigma38p10 2758752 Gap 7.5 bits
... ----------------| distalUP-proximalUP-sigma38p10 2758752 total 3.9 bits
|--------------------------------------------------- ... proximalUP-(41)
distalUP 4.8 bits
|------------------------------------------------------------- ... distalUP-(47)-s
... ----------------| proximalUP-(40)-sigma38p10 2758752 Gap 4.0 bits
|-----------|------------------------------------------------- ... distalUP-proxin
|------------------------------------------------------------- ... distalUP-(56)-s
|--------------------------------------------------------|----- ... distalUP-proxin

. *2758790 . *2758800 . *2758810 .
5' a a g c t t a a a c a t g a t a a g c a t a t t a a a a c t c a a c g g a a 3'
sigma38p10 3.6 bits

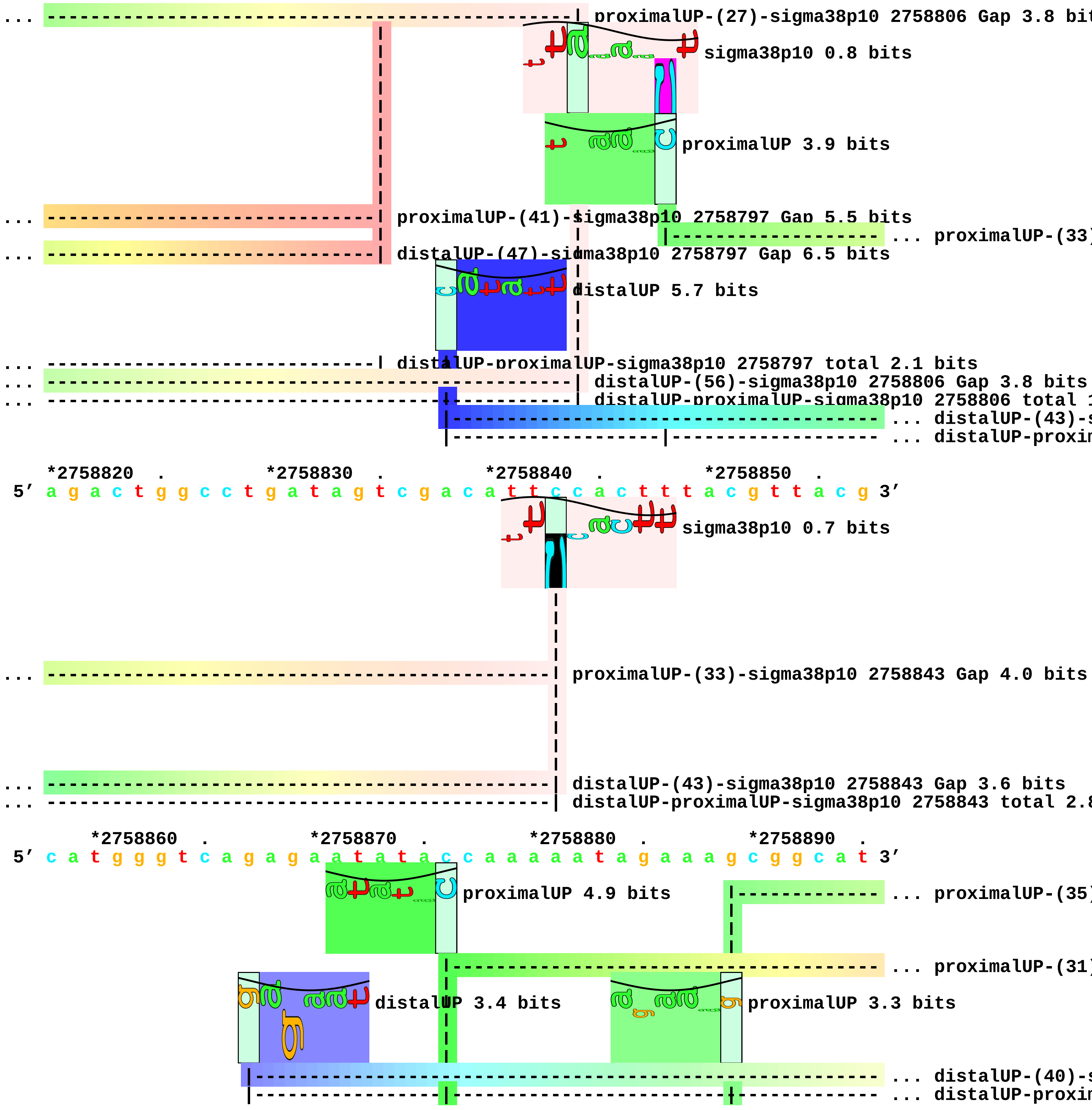
... proximalUP-(27)-sigma38p10 2758806 Gap 3.8 bit
sigma38p10 0.8 bits
proximalUP 3.9 bits
... proximalUP-(41)-sigma38p10 2758797 Gap 5.5 bits
... proximalUP-(33)
... distalUP-(47)-sigma38p10 2758797 Gap 6.5 bits
distalUP 5.7 bits
... distalUP-proximalUP-sigma38p10 2758797 total 2.1 bits
... distalUP-(56)-sigma38p10 2758806 Gap 3.8 bits
... distalUP-proximalUP-sigma38p10 2758806 total 1
... distalUP-(43)-s
... distalUP-proxin
*2758820 . *2758830 . *2758840 . *2758850 .
5′ a g a c t g g c c t g a t a g t c g a c a t t c c a c t t t a c g t t a c g 3′
sigma38p10 0.7 bits
... proximalUP-(33)-sigma38p10 2758843 Gap 4.0 bits
... distalUP-(43)-sigma38p10 2758843 Gap 3.6 bits
... distalUP-proximalUP-sigma38p10 2758843 total 2.8
*2758860 . *2758870 . *2758880 . *2758890 .
5′ c a t g g g t c a g a g a a t a t a c c a a a a a t a g a a a g c g g c a t 3′
proximalUP 4.9 bits
... proximalUP-(35)
... proximalUP-(31)
distalUP 3.4 bits
proximalUP 3.3 bits
... distalUP-(40)-s
... distalUP-proxin

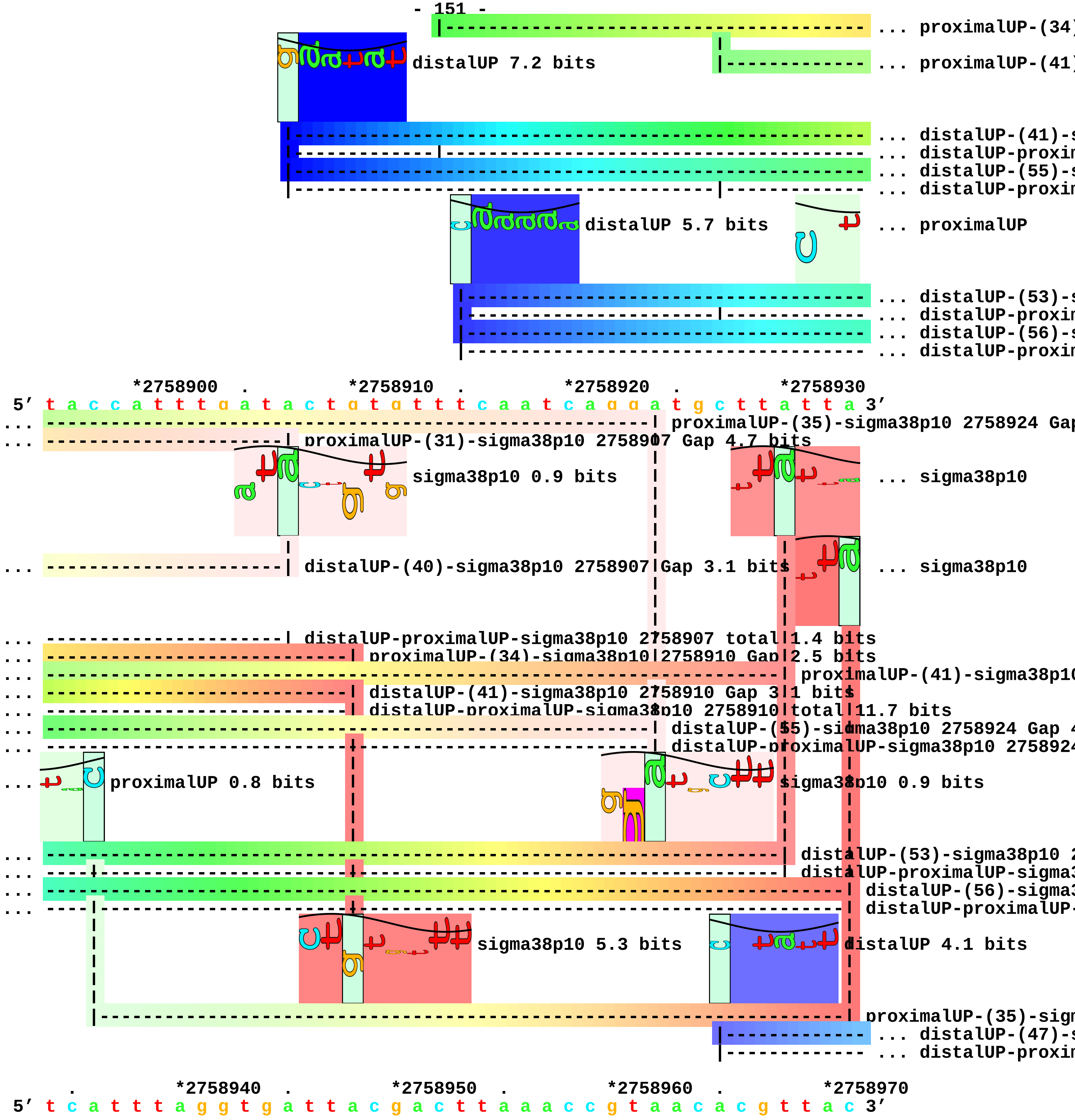
... proximalUP-(34)
... proximalUP-(41)
distalUP 7.2 bits
... distalUP-(41)-s
... distalUP-proxim
... distalUP-(55)-s
... distalUP-proxim
distalUP 5.7 bits
... proximalUP
... distalUP-(53)-s
... distalUP-proxim
... distalUP-(56)-s
... distalUP-proxim
*2758900 . *2758910 . *2758920 . *2758930
5' t a c c a t t t g a t a c t g t g t t t c a a t c a g g a t g c t t a t t a 3'
... proximalUP-(35)-sigma38p10 2758924 Gap
... proximalUP-(31)-sigma38p10 2758907 Gap 4.7 bits
sigma38p10 0.9 bits
... sigma38p10
... distalUP-(40)-sigma38p10 2758907 Gap 3.1 bits
... sigma38p10
... distalUP-proximalUP-sigma38p10 2758907 total 1.4 bits
... proximalUP-(34)-sigma38p10 2758910 Gap 2.5 bits
... proximalUP-(41)-sigma38p10
... distalUP-(41)-sigma38p10 2758910 Gap 3.1 bits
... distalUP-proximalUP-sigma38p10 2758910 total 11.7 bits
... distalUP-(55)-sigma38p10 2758924 Gap 4
... distalUP-proximalUP-sigma38p10 275892
... proximalUP 0.8 bits
sigma38p10 0.9 bits
... distalUP-(53)-sigma38p10 2
... distalUP-proximalUP-sigma3
... distalUP-(56)-sigma3
... distalUP-proximalUP-
sigma38p10 5.3 bits
distalUP 4.1 bits
proximalUP-(35)-sigm
... distalUP-(47)-s
... distalUP-proxim
. *2758940 . *2758950 . *2758960 . *2758970
5' t c a t t t a g g t g a t t a c g a c t t a a a c c g t a a c a c g t t a c 3'

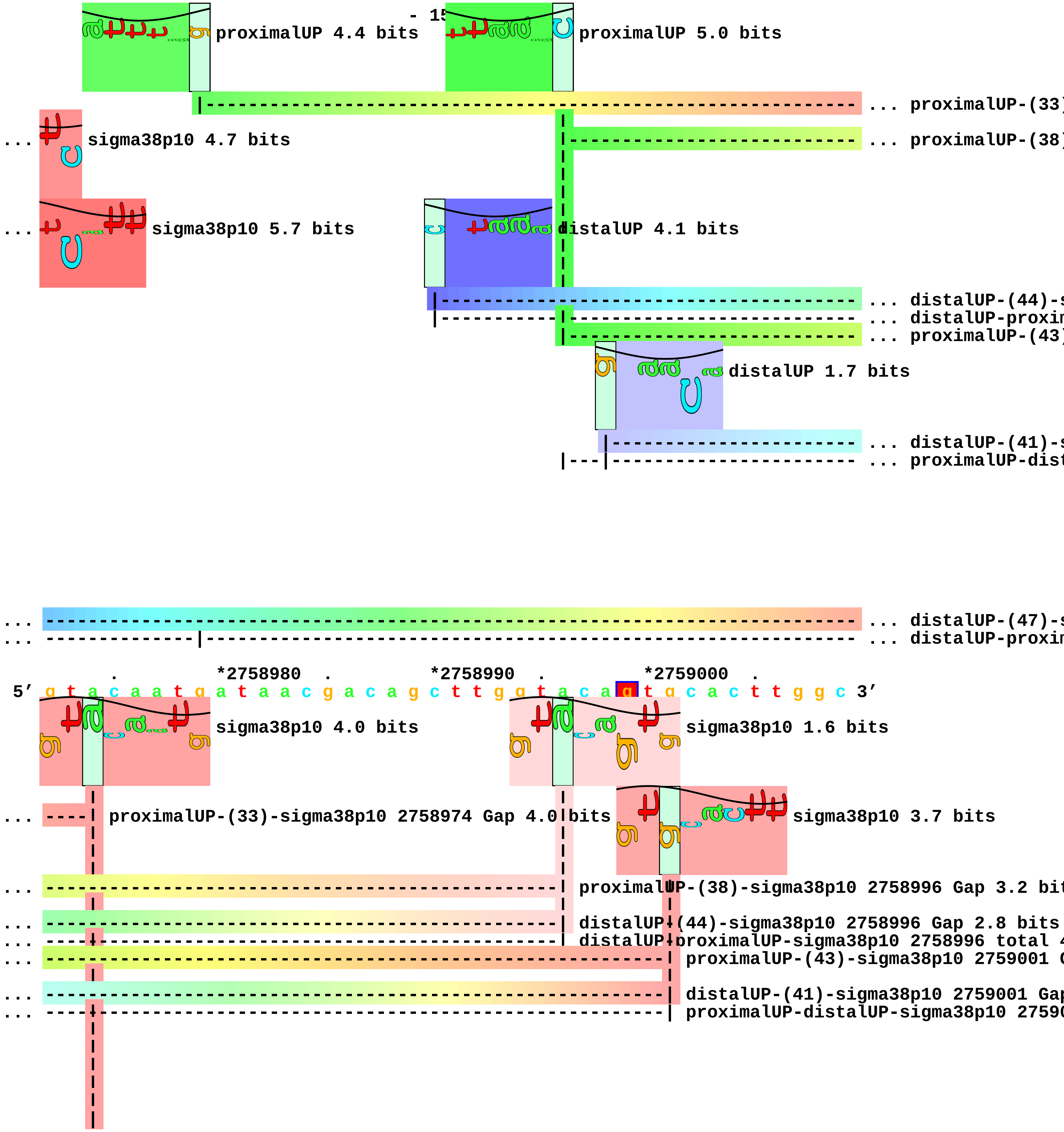

- 15
proximalUP 4.4 bits
proximalUP 5.0 bits
... proximalUP-(33)
... sigma38p10 4.7 bits
... proximalUP-(38)
... sigma38p10 5.7 bits
distalUP 4.1 bits
... distalUP-(44)-s
... distalUP-proxim
... proximalUP-(43)
distalUP 1.7 bits
... distalUP-(41)-s
... proximalUP-dist
... distalUP-(47)-s
... distalUP-proxim
. *2758980 . *2758990 . *2759000 .
5′ g t a c a a t g a t a a c g a c a g c t t g g t a c a g t g c a c t t g g c 3′
sigma38p10 4.0 bits
sigma38p10 1.6 bits
... proximalUP-(33)-sigma38p10 2758974 Gap 4.0 bits
sigma38p10 3.7 bits
... proximalUP-(38)-sigma38p10 2758996 Gap 3.2 bit
... distalUP-(44)-sigma38p10 2758996 Gap 2.8 bits
... distalUP-proximalUP-sigma38p10 2758996 total 4
... proximalUP-(43)-sigma38p10 2759001 G
... distalUP-(41)-sigma38p10 2759001 Gap
... proximalUP-distalUP-sigma38p10 27590

```
        |
        |
... ----| distalUP-(47)-sigma38p10 2758974 Gap 6.5 bits
... ----| distalUP-proximalUP-sigma38p10 2758974 total 1.9 bits

    *2759010  .         *2759020  .         *2759030  .         *2759040  .
 5' a t c c c t a t a g a t a g c t c g g g g a a g t t a g t t a a t t a c c c 3'


 5' g c 3'
```

piece 37, NC_000913, alaE, config: linear, direction: +, begin: 2797050, end: 2797299

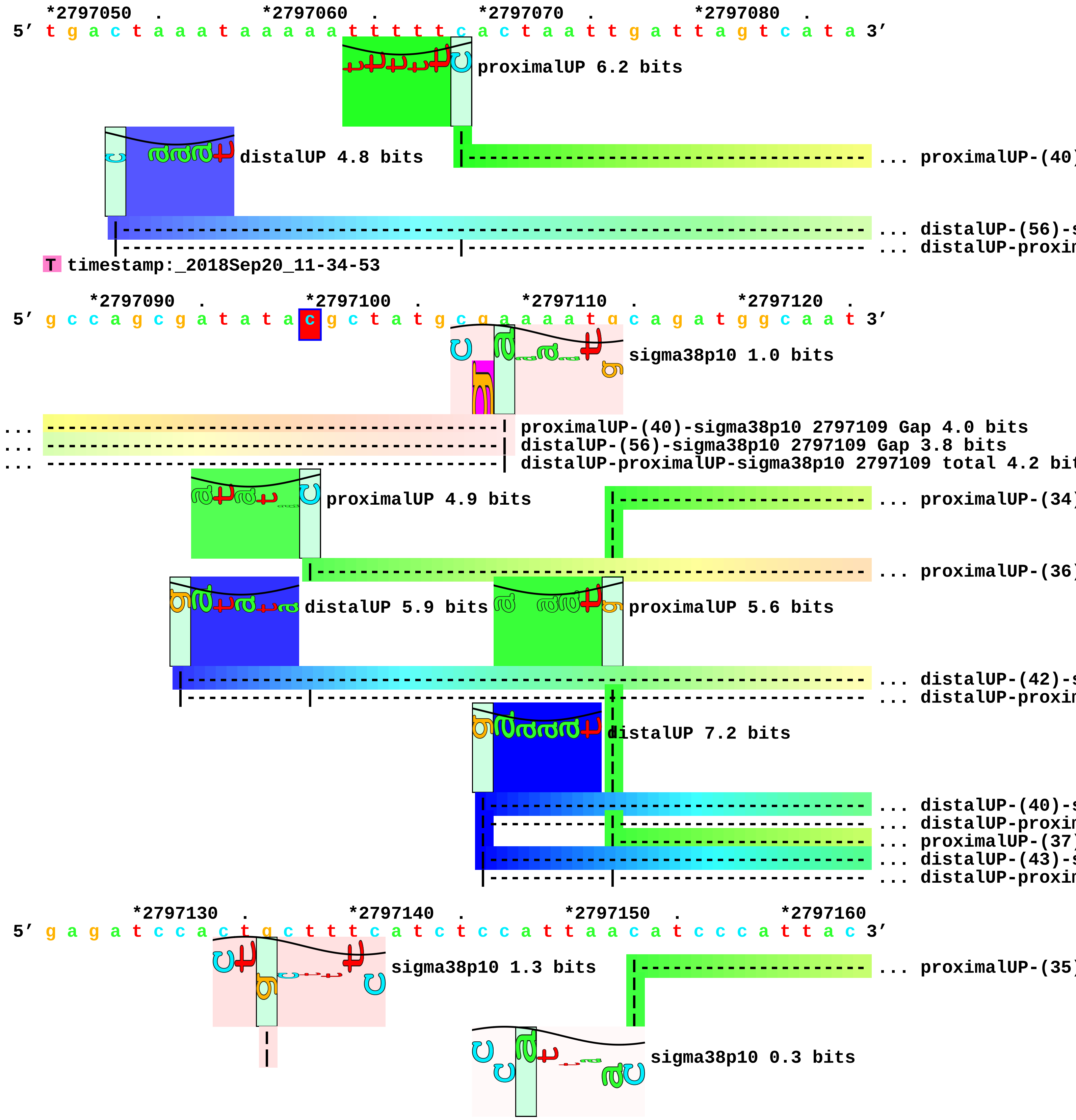

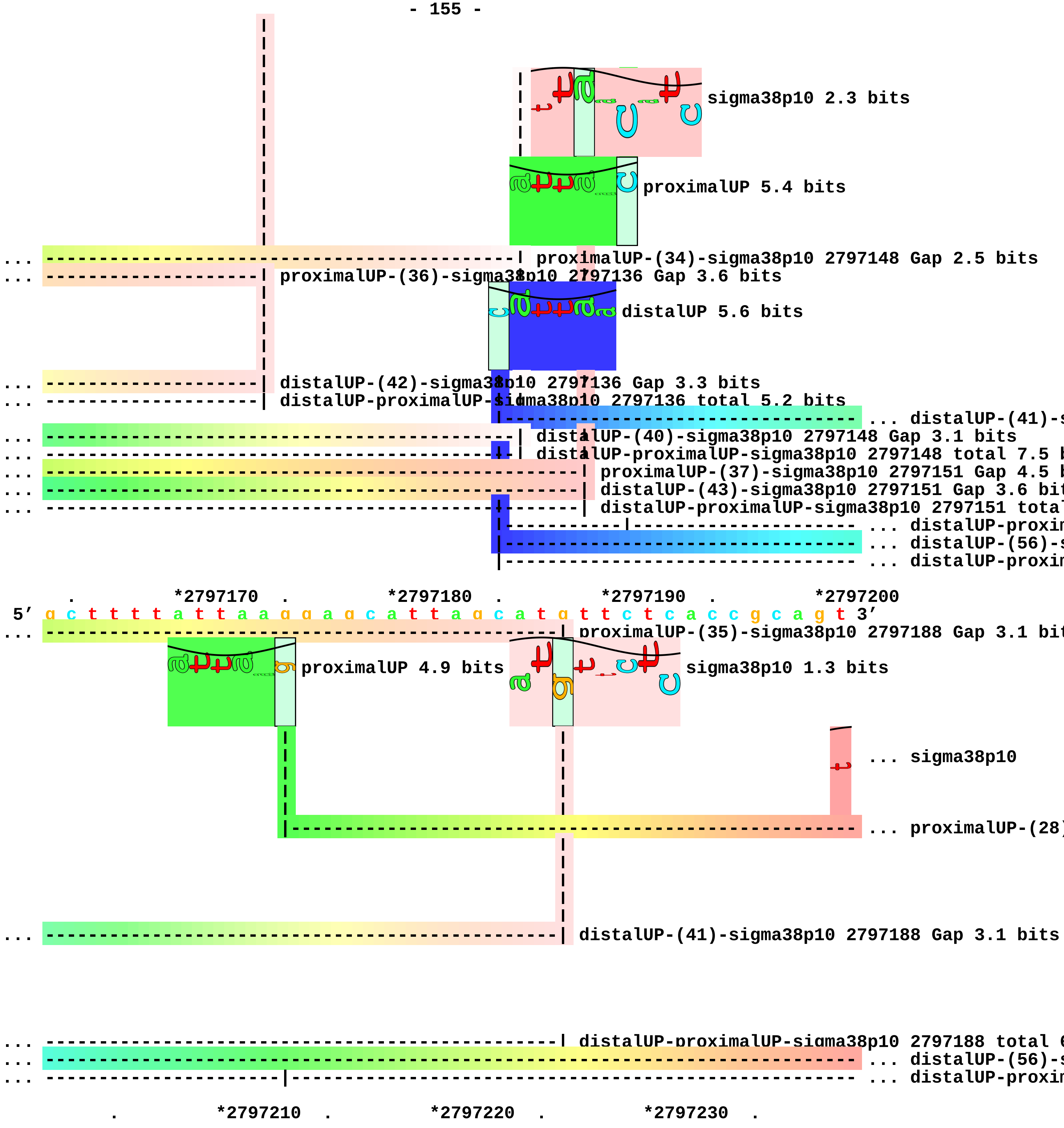
sigma38p10 2.3 bits
proximalUP 5.4 bits
... proximalUP-(34)-sigma38p10 2797148 Gap 2.5 bits
... proximalUP-(36)-sigma38p10 2797136 Gap 3.6 bits
distalUP 5.6 bits
... distalUP-(42)-sigma38p10 2797136 Gap 3.3 bits
... distalUP-proximalUP-sigma38p10 2797136 total 5.2 bits
... distalUP-(41)-s
... distalUP-(40)-sigma38p10 2797148 Gap 3.1 bits
... distalUP-proximalUP-sigma38p10 2797148 total 7.5 b
... proximalUP-(37)-sigma38p10 2797151 Gap 4.5 b
... distalUP-(43)-sigma38p10 2797151 Gap 3.6 bit
... distalUP-proximalUP-sigma38p10 2797151 total
... distalUP-proxin
... distalUP-(56)-s
... distalUP-proxin
. *2797170 . *2797180 . *2797190 . *2797200
5' g c t t t t a t t a a g g a g c a t t a g c a t g t t c t c a c c g c a g t 3'
... proximalUP-(35)-sigma38p10 2797188 Gap 3.1 bit
proximalUP 4.9 bits
sigma38p10 1.3 bits
... sigma38p10
... proximalUP-(28)
... distalUP-(41)-sigma38p10 2797188 Gap 3.1 bits
... distalUP-proximalUP-sigma38p10 2797188 total 6
... distalUP-(56)-s
... distalUP-proxin
. *2797210 . *2797220 . *2797230 .

5′ c a c g c t t g c g t c a t g c a g t t g c a g a t a c g t t c g c g a t g 3′

... 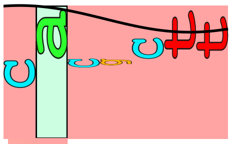 sigma38p10 3.9 bits

... --| proximalUP-(28)-sigma38p10 2797203 Gap 5.5 bits

... --| distalUP-(56)-sigma38p10 2797203 Gap 3.8 bits
... --| distalUP-proximalUP-sigma38p10 2797203 total 5.3 bits

*2797240 . *2797250 . *2797260 . *2797270 .
5′ g t t g t t t a c g t t c t g t c g t g a a c a t g t g t a t t g a a g t 3′

*2797280 . *2797290 .
5′ t t t c c t c t c c g g a a t g a g c t t c 3′

piece 38, NC_000913, csrA, config: linear, direction: -, begin: 2817445, end: 2817177

```
   .         *2817440  .         *2817430  .         *2817420  .         *2817410
5' g t t a g c c a g t g t g a a a g g c t g g g t c a g c g c g a a a t t g c 3'
   T timestamp:_2018Sep20_11-34-53

     .         *2817400  .         *2817390  .         *2817380  .         *2817370
5' a a t a a t a t a a g c g t c a g g c a a t g c c g t g g a c t c g c t t c 3'

       .         *2817360  .         *2817350  .         *2817340  .
5' a c g g c a t t c g c a t t a a c g c t a t c g a c a a c g a t a a a g t c 3'
                                proximalUP 5.4 bits                  proximalUP 4.9 bits
                                                                     ... proximalUP-(35)
                                                                     ... proximalUP-(27)
                      distalUP 5.6 bits
                                                                     ... distalUP-(41)-s
                                                                     ... sigma38p10-prox
                                                                     ... distalUP-(52)-s
                                                                     ... sigma38p10-prox

    *2817330  .         *2817320  .         *2817310  .         *2817300  .
5' a g g t t g a a g t t g t g t a t a t c g g c t a a a c t t a g g t t t a a 3'
                              sigma38p10 0.6 bits                    ... proximalUP
... ------------------------| proximalUP-(35)-sigma38p10 2817318 Gap 3.1 bits
... ---------------------------------------------| proximalUP-(27)-sigma38p10 2817307 Gap 3.8 bit
... ------------------------| distalUP-(41)-sigma38p10 2817318 Gap 3.1 bits
... ------------------------| sigma38p10-proximalUP-distalUP 2817318 total 5.4 bits
... ---------------------------------------------| distalUP-(52)-sigma38p10 2817307 Gap 5.5 bits
... ---------------------------------------------| sigma38p10-proximalUP-distalUP 2817307 total 1
                                        sigma38p10 10.3 bits
                      distalUP 5.6 bits                              distalUP 4.8 bits
                                                                     ... distalUP-(53)-s
                                                                     ... sigma38p10-prox
                                                                     ... distalUP-(55)-s
                                                                     ... sigma38p10-prox
                                                                     ... distalUP-(45)-s
                                                                     ... sigma38p10-prox
                                                                     ... distalUP-(55)-s
                                                                     ... sigma38p10-prox

         *2817290  .         *2817280  .         *2817270  .         *2817260
5' c a g a a t g t a a t g c c a t g a c t g c t t a g a t g t a a t g t g t t 3'
```

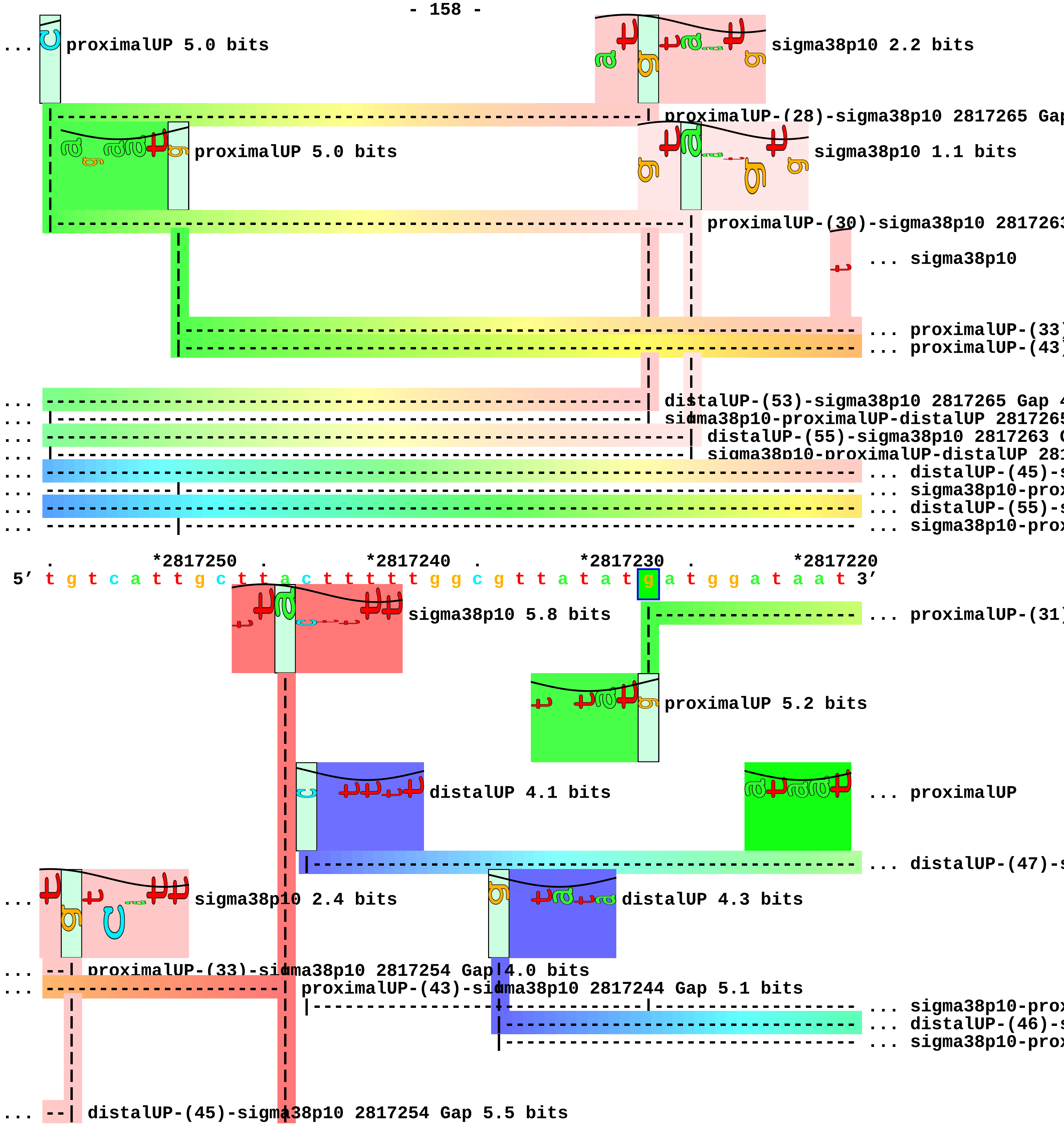

proximalUP 5.0 bits
sigma38p10 2.2 bits
proximalUP-(28)-sigma38p10 2817265 Gap
proximalUP 5.0 bits
sigma38p10 1.1 bits
proximalUP-(30)-sigma38p10 281726
... sigma38p10
... proximalUP-(33)
... proximalUP-(43)
distalUP-(53)-sigma38p10 2817265 Gap
sigma38p10-proximalUP-distalUP 281726
distalUP-(55)-sigma38p10 2817263
sigma38p10-proximalUP-distalUP 281
... distalUP-(45)-
... sigma38p10-prox
... distalUP-(55)-
... sigma38p10-prox
*2817250 *2817240 *2817230 *2817220
5′ t g t c a t t g c t t a c t t t t t g g c g t t a t a t g a t g g a t a a t 3′
sigma38p10 5.8 bits
... proximalUP-(31)
proximalUP 5.2 bits
distalUP 4.1 bits
... proximalUP
... distalUP-(47)-
sigma38p10 2.4 bits
distalUP 4.3 bits
proximalUP-(33)-sigma38p10 2817254 Gap 4.0 bits
proximalUP-(43)-sigma38p10 2817244 Gap 5.1 bits
... sigma38p10-prox
... distalUP-(46)-
... sigma38p10-prox
distalUP-(45)-sigma38p10 2817254 Gap 5.5 bits

```
... --| sigma38p10-proximalUP-distalUP 2817254 total 2.8 bits
... ----------------------| distalUP-(55)-sigma38p10 2817244 Gap 4.9 bits
... ----------------------| sigma38p10-proximalUP-distalUP 2817244 total 5.7 bits

         .         *2817210  .         *2817200  .         *2817190  .         *2817180
 5' g c c g g g a t a c a g a g a g a c c c g a c t c t t t t a a t c t t t c a 3'
... ------------------------------------------| proximalUP-(31)-sigma38p10 2817196 Gap 4.7 bits
```

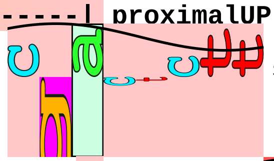

sigma38p10 2.4 bits

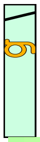

... proximalUP 6.7 bits

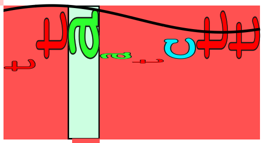

sigma38p10 7.5 bits

```
... ------------------------------------------| distalUP-(47)-sigma38p10 2817196 Gap 6.5 bits
    |---------------------------------------------------------| proximalUP-(29)-sigma38p10 2817188 G

... ------------------------------------------| sigma38p10-proximalUP-distalUP 2817196 total 0.5 bit
... ----------------------------------------------------------| distalUP-(46)-sigma38p10 2817188 Gap
... |---------------------------------------------------------| sigma38p10-proximalUP-distalUP 28171

 5' a g g 3'
```

piece 39, NC_000913, ygdH, config: linear, direction: +, begin: 2924202, end: 2924420

. *2924210 . *2924220 . *2924230 .
5′ g a c t g g t t c g g c a a t a a a t t t t t t c t c a a t t t t g c g t g 3′
T timestamp:_2018Sep20_11-34-53

*2924240 . *2924250 . *2924260 . *2924270 .
5′ c t g g a t t c a c g c a g a a g g t t g t g a a a g g t c a t c a g g c a 3′

*2924280 . *2924290 . *2924300 . *2924310 .
5′ g g g c t a t t g t a a t c a a a g g g a a t g a c g a t a t t c g t c c c 3′
distalUP 6.5 bits
... distalUP-(56)-s
... distalUP-proxim

*2924320 . *2924330 . *2924340 . *2924350
5′ a t a a g g a g t t t t t c t t g a t t a c a c a t a t t a g c c c g c t t 3′
proximalUP 6.2 bits
proximalUP 0.2 bits
... distalUP-(56)-s
... distalUP-proxim
... proximalUP-(31)
... proximalUP-(37)
distalUP 4.9 bits
... proximalUP-(43)
... distalUP-(43)-s
... distalUP-proxim
distalUP 5.7 bits
... distalUP-(51)-s
... distalUP-proxim

. *2924360 . *2924370 . *2924380 . *2924390
5′ g g c t c c a t g g a t a t g t g t c g c a g c t g g a a g t g g a t a t 3′
sigma38p10 0.7 bits
... sigma38p10
... distalUP-(56)-sigma38p10 2924360 Gap 3.8 bits
... distalUP-proximalUP-sigma38p10 2924360 total 5.0 bits
... proximalUP-(31)-sigma38p10 2924360 Gap 4.7 bits
... proximalUP-(37)-sigma38p10 2924366 Gap 4.5 bits
... proximalUP-(43)-sigma3
... distalUP-(43)-sigma38p10 2924366 Gap 3.6 bits
... distalUP-proximalUP-sigma38p10 2924366 total 6.9 bits
sigma38p10 3.8 bits

```
... ------------------------------------------------------------------------| distalUP-(51)-sigma38p
... ------------------------------------------------------------------------| distalUP-proximalUP-si

            .         *2924400   .         *2924410   .         *2924420
 5′ g c t t a a a c g c a c c g c c a g c a g c g a c c t c t 3′
```

... 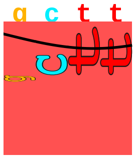 sigma38p10 7.4 bits piece 40, NC_000913, omrA, config: linear, direction: -, begin: 2974328, end: 2974103

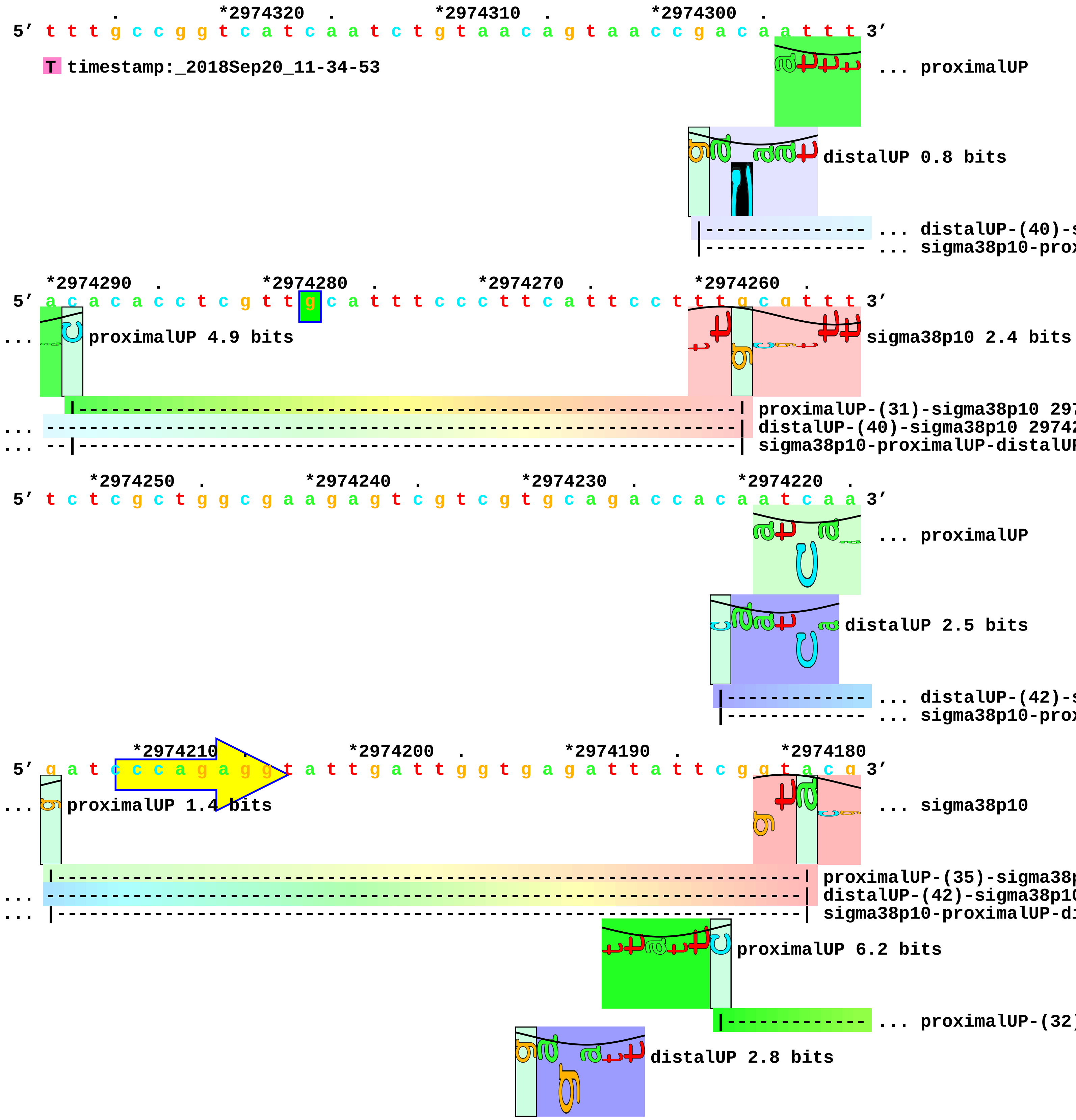

```
                                                |---------------------------- ... distalUP-(41)-s
                                                |----------------|------------- ... sigma38p10-prox

      .         *2974170  .         *2974160  .         *2974150  .         *2974140
5' c t c t t c g t a c c c t g t c t c t t g c a c c a a c c t g c g c g g a t 3'
...   sigma38p10 3.0 bits                                          sigma38p10 2.0 bits

... --------------------------------------------------| proximalUP-(32)-sigma38p10 2974151 Gap 4.3 b
... --------------------------------------------------| distalUP-(41)-sigma38p10 2974151 Gap 3.1 bit
... --------------------------------------------------| sigma38p10-proximalUP-distalUP 2974151 tota

        .         *2974130  .         *2974120  .         *2974110  .
5' g c g c a g g t t t t t t t c g c a c c t a a t t t a c t g t c g c t 3'
```

piece 41, NC_000913, ygeI, config: linear, direction: +, begin: 2991050, end: 2992349

*2991050 . *2991060 . *2991070 . *2991080 .
5′ g a a a a a g t g a a c t t g a a c t t g c t g c t c a a a a a g c a t t a 3′

proximalUP 1.5 bits
... proximalUP-(37)
distalUP 6.5 bits
... distalUP-(51)-s
... distalUP-proxi
T timestamp:_2018Sep20_11-34-53

*2991090 . *2991100 . *2991110 . *2991120 .
5′ g a g c t t t t a g a t [illegible] a t g t a t c a g a c a t a a c c a c t g t c g a 3′

sigma38p10 3.7 bits
... proximalUP-(37)-sigma38p10 2991101 Gap 4.5 bits
proximalUP 5.1 bits
... distalUP-(51)-sigma38p10 2991101 Gap 5.1 bits
... distalUP-proximalUP-sigma38p10 2991101 total 2.1 bits
... proximalUP-(35)
distalUP 7.1 bits
... distalUP-(45)-s
... distalUP-proxi

*2991130 . *2991140 . *2991150 . *2991160
5′ t g g a a a a a t t t t a g c t a t t a t g g g a c t g a t a a c t g g t c 3′

sigma38p10 6.7 bits
... proximalUP-(35)-sigma38p10 2991142 Gap 3.1 bits
... distalUP-(45)-sigma38p10 2991142 Gap 5.5 bits
... distalUP-proximalUP-sigma38p10 2991142 total 10.3 bits

. *2991170 . *2991180 . *2991190 . *2991200
5′ t g t c t g g a c a a g c a a a a g t a t c t c a t a t c t t a t t t g a a 3′

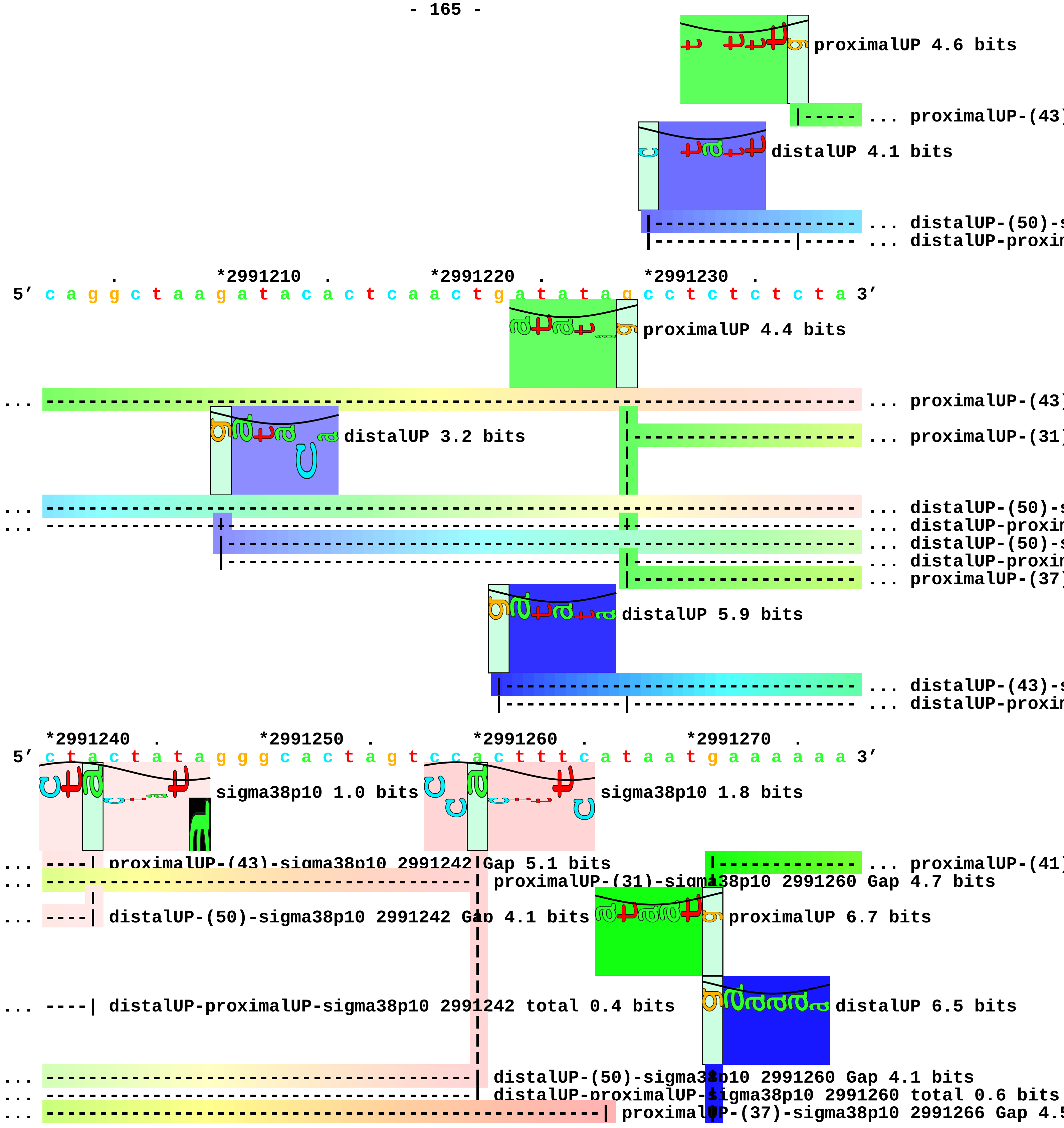
proximalUP 4.6 bits
... proximalUP-(43)
distalUP 4.1 bits
... distalUP-(50)-s
... distalUP-proxim
. *2991210 . *2991220 . *2991230 .
5′ c a g g c t a a g a t a c a c t c a a c t g a t a t a g c c t c t c t c t a 3′
proximalUP 4.4 bits
... proximalUP-(43)
distalUP 3.2 bits
... proximalUP-(31)
... distalUP-(50)-s
... distalUP-proxim
... distalUP-(50)-s
... distalUP-proxim
... proximalUP-(37)
distalUP 5.9 bits
... distalUP-(43)-s
... distalUP-proxim
*2991240 . *2991250 . *2991260 . *2991270 .
5′ c t a c t a t a g g g c a c t a g t c c a c t t t c a t a a t g a a a a a a 3′
sigma38p10 1.0 bits
sigma38p10 1.8 bits
... ----| proximalUP-(43)-sigma38p10 2991242 Gap 5.1 bits
... proximalUP-(41)
... proximalUP-(31)-sigma38p10 2991260 Gap 4.7 bits
... ----| distalUP-(50)-sigma38p10 2991242 Gap 4.1 bits
proximalUP 6.7 bits
... ----| distalUP-proximalUP-sigma38p10 2991242 total 0.4 bits
distalUP 6.5 bits
... distalUP-(50)-sigma38p10 2991260 Gap 4.1 bits
... distalUP-proximalUP-sigma38p10 2991260 total 0.6 bits
... proximalUP-(37)-sigma38p10 2991266 Gap 4.5

sigma38p10 3.2 bits

... ---------------------------------------------------------| distalUP-(43)-sigma38p10 2991266 Gap 3.6 b
... ---------------------------------------------------------| distalUP-proximalUP-sigma38p10 2991266 tot
|------------- ... distalUP-(41)-s
|------------- ... proximalUP-dist

*2991280 . *2991290 . *2991300 . *2991310 .
5' t t g a a g a g g c a a g g a t t t g t a t a g a c a a a t c a c t a c a a 3'

... sigma38p10

... ---------------------------------------------------------------| proximalUP-(41)-sigma38p10

... ---------------------------------------------------------------| distalUP-(41)-sigma38p10 2
... ---------------------------------------------------------------| proximalUP-distalUP-sigma3

*2991320 . *2991330 . *2991340 . *2991350
5' c t c g a a c c c a g a a g a c g a a a a g c a g t t g t g a t a a a a g a 3'

... sigma38p10 1.9 bits ... proximalUP

distalUP 6.5 bits

|----------------- ... distalUP-(46)-s
|----------------- ... distalUP-proxin
|----------------- ... distalUP-(52)-s
|----------------- ... distalUP-proxin

. *2991360 . *2991370 . *2991380 . *2991390
5' a t g t g t a g a t a t g t a t g t g c c t a a c c c g c t c a a a a a c a 3'

... proximalUP 5.0 bits ... sigma38p10

|--------------------------------------------------------------| proximalUP-(35)-sigm
... ---------------------------------------------------------------| distalUP-(46)-sigma3
... ----|----------------------------------------------------------| distalUP-proximalUP-
... ---------------------------------------------------------------- ... distalUP-(52)-s
... -----------------------------|---------------------------------- ... distalUP-proxin

proximalUP 4.6 bits

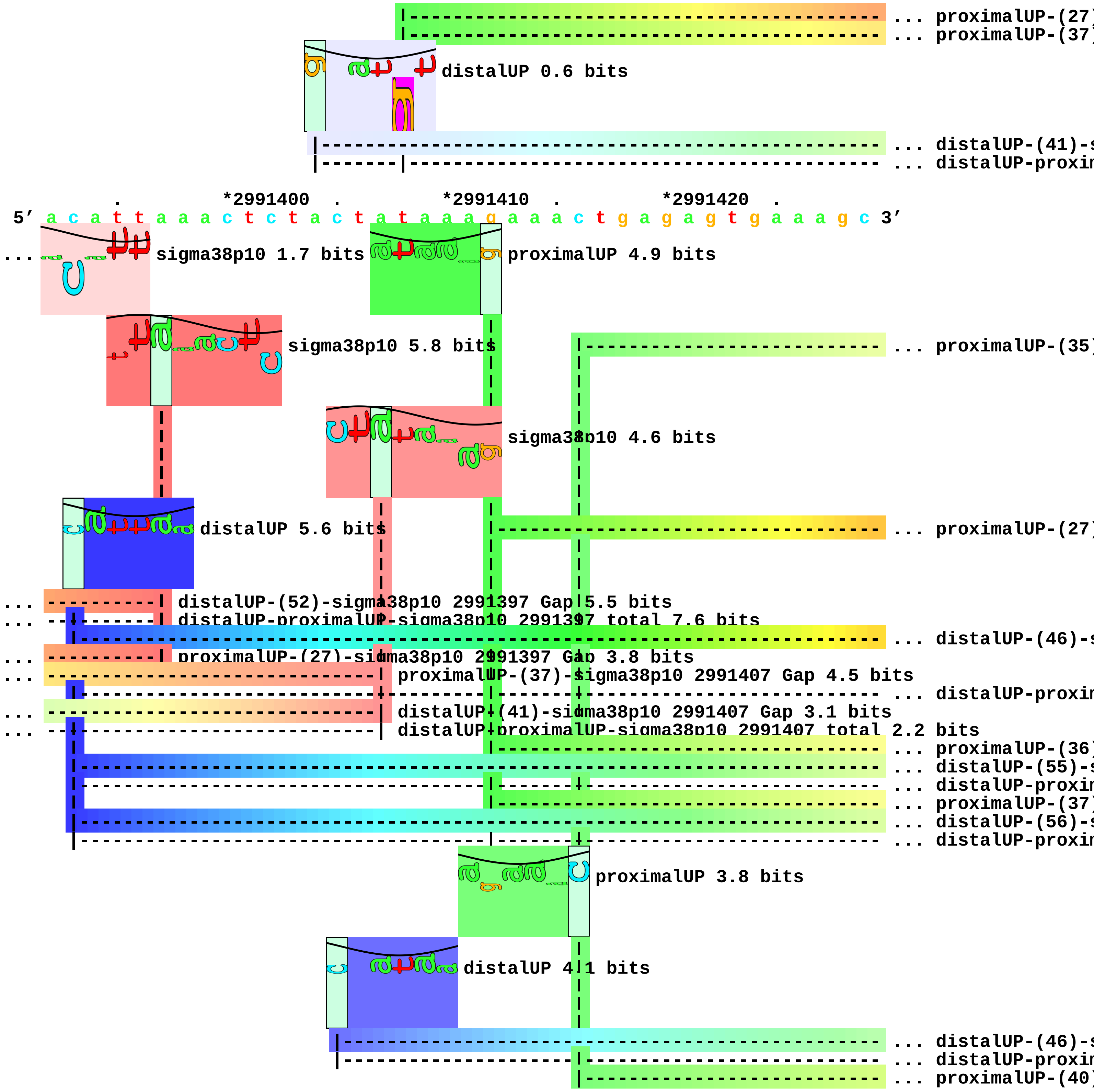
... proximalUP-(27)
... proximalUP-(37)
distalUP 0.6 bits
... distalUP-(41)-s
... distalUP-proxim
*2991400
*2991410
*2991420
5' a c a t t a a a c t c t a c t a t a a a g a a a c t g a g a g t g a a a g c 3'
... sigma38p10 1.7 bits
proximalUP 4.9 bits
sigma38p10 5.8 bits
... proximalUP-(35)
sigma38p10 4.6 bits
distalUP 5.6 bits
... proximalUP-(27)
... distalUP-(52)-sigma38p10 2991397 Gap 5.5 bits
... distalUP-proximalUP-sigma38p10 2991397 total 7.6 bits
... distalUP-(46)-s
... proximalUP-(27)-sigma38p10 2991397 Gap 3.8 bits
... proximalUP-(37)-sigma38p10 2991407 Gap 4.5 bits
... distalUP-proxim
... distalUP-(41)-sigma38p10 2991407 Gap 3.1 bits
... distalUP-proximalUP-sigma38p10 2991407 total 2.2 bits
... proximalUP-(36)
... distalUP-(55)-s
... distalUP-proxim
... proximalUP-(37)
... distalUP-(56)-s
... distalUP-proxim
proximalUP 3.8 bits
distalUP 4.1 bits
... distalUP-(46)-s
... distalUP-proxim
... proximalUP-(40)

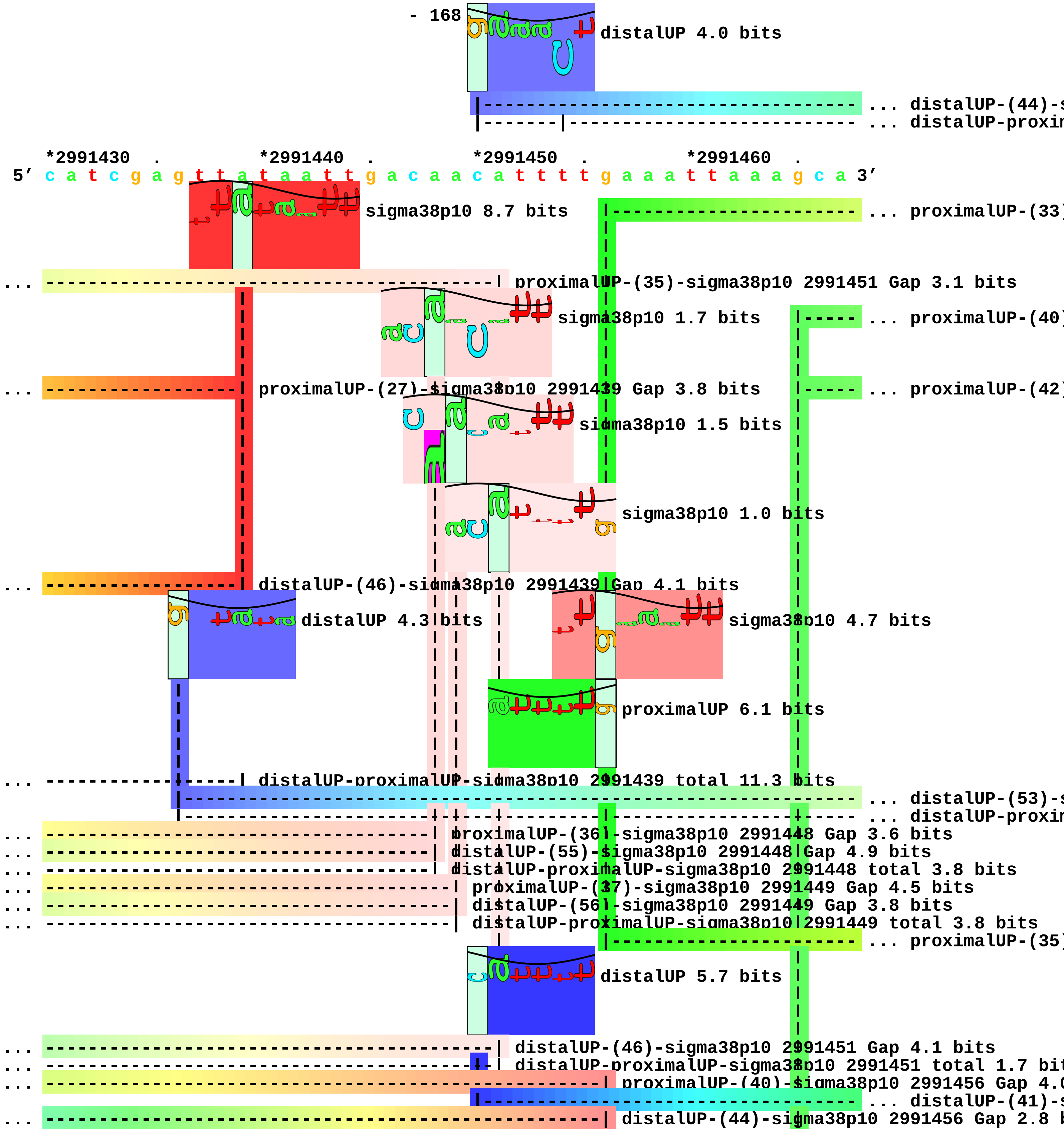
distalUP 4.0 bits
... distalUP-(44)-s
... distalUP-proxin
*2991430 .
*2991440 .
*2991450 .
*2991460 .
5' c a t c g a g t t a t a a t t g a c a a c a t t t t g a a a t t a a a g c a 3'
sigma38p10 8.7 bits
... proximalUP-(33)
... proximalUP-(35)-sigma38p10 2991451 Gap 3.1 bits
sigma38p10 1.7 bits
... proximalUP-(40)
... proximalUP-(27)-sigma38p10 2991439 Gap 3.8 bits
... proximalUP-(42)
sigma38p10 1.5 bits
sigma38p10 1.0 bits
... distalUP-(46)-sigma38p10 2991439 Gap 4.1 bits
distalUP 4.3 bits
sigma38p10 4.7 bits
proximalUP 6.1 bits
... distalUP-proximalUP-sigma38p10 2991439 total 11.3 bits
... distalUP-(53)-s
... distalUP-proxin
... proximalUP-(36)-sigma38p10 2991448 Gap 3.6 bits
... distalUP-(55)-sigma38p10 2991448 Gap 4.9 bits
... distalUP-proximalUP-sigma38p10 2991448 total 3.8 bits
... proximalUP-(37)-sigma38p10 2991449 Gap 4.5 bits
... distalUP-(56)-sigma38p10 2991449 Gap 3.8 bits
... distalUP-proximalUP-sigma38p10 2991449 total 3.8 bits
... proximalUP-(35)
distalUP 5.7 bits
... distalUP-(46)-sigma38p10 2991451 Gap 4.1 bits
... distalUP-proximalUP-sigma38p10 2991451 total 1.7 bit
... proximalUP-(40)-sigma38p10 2991456 Gap 4.0
... distalUP-(41)-s
... distalUP-(44)-sigma38p10 2991456 Gap 2.8 b

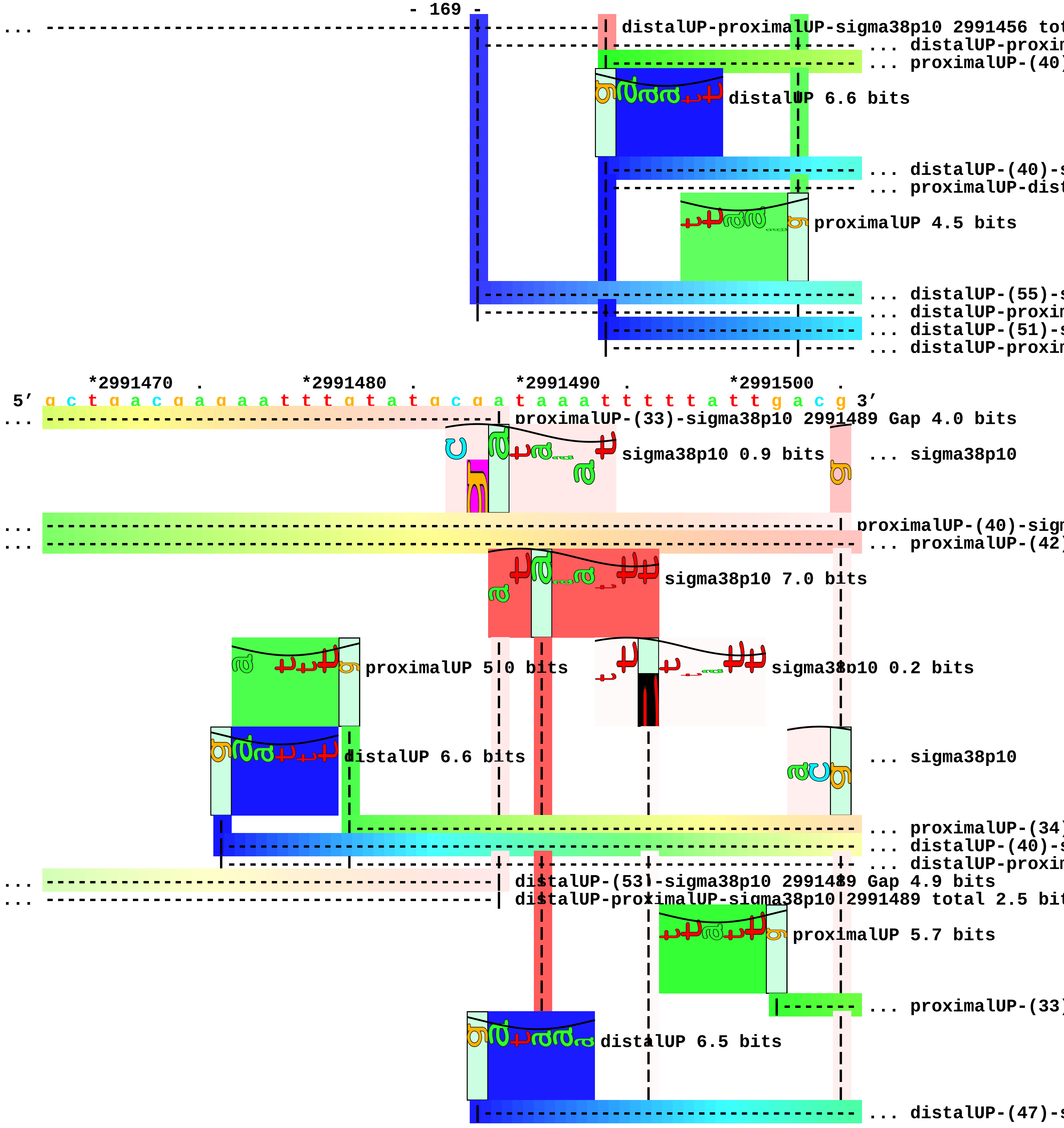

... distalUP-proximalUP-sigma38p10 2991456 tot
... distalUP-proxim
... proximalUP-(40)
distalUP 6.6 bits
... distalUP-(40)-s
... proximalUP-dist
proximalUP 4.5 bits
... distalUP-(55)-s
... distalUP-proxim
... distalUP-(51)-s
... distalUP-proxim
*2991470 . *2991480 . *2991490 . *2991500 .
5′ g c t g a c g a g a a t t t g t a t g c g a t a a a t t t t t a t t g a c g 3′
... proximalUP-(33)-sigma38p10 2991489 Gap 4.0 bits
sigma38p10 0.9 bits
... sigma38p10
... proximalUP-(40)-sign
... proximalUP-(42)
sigma38p10 7.0 bits
proximalUP 5.0 bits
sigma38p10 0.2 bits
distalUP 6.6 bits
... sigma38p10
... proximalUP-(34)
... distalUP-(40)-s
... distalUP-proxim
... distalUP-(53)-sigma38p10 2991489 Gap 4.9 bits
... distalUP-proximalUP-sigma38p10 2991489 total 2.5 bit
proximalUP 5.7 bits
... proximalUP-(33)
distalUP 6.5 bits
... distalUP-(47)-s

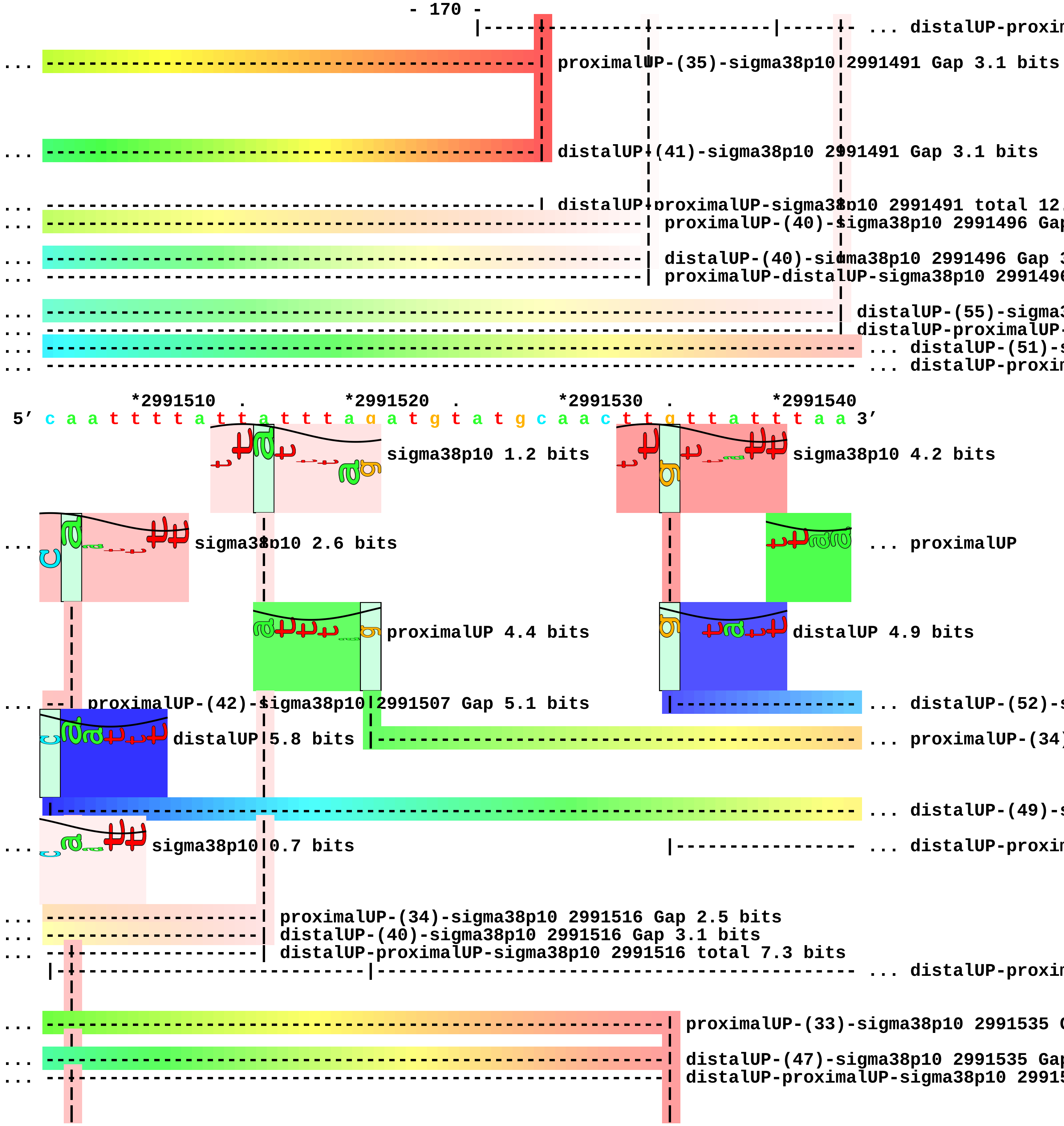
... distalUP-proxim
proximalUP-(35)-sigma38p10 2991491 Gap 3.1 bits
distalUP-(41)-sigma38p10 2991491 Gap 3.1 bits
distalUP-proximalUP-sigma38p10 2991491 total 12.
proximalUP-(40)-sigma38p10 2991496 Gap
distalUP-(40)-sigma38p10 2991496 Gap 3
proximalUP-distalUP-sigma38p10 299149
distalUP-(55)-sigma3
distalUP-proximalUP-
... distalUP-(51)-s
... distalUP-proxim
*2991510 . *2991520 . *2991530 . *2991540
5′ c a a t t t t a t t a t t t a g a t g t a t g c a a c t t g t t a t t t a a 3′
sigma38p10 1.2 bits
sigma38p10 4.2 bits
sigma38p10 2.6 bits
... proximalUP
proximalUP 4.4 bits
distalUP 4.9 bits
proximalUP-(42)-sigma38p10 2991507 Gap 5.1 bits
... distalUP-(52)-s
distalUP 5.8 bits
... proximalUP-(34)
... distalUP-(49)-s
sigma38p10 0.7 bits
... distalUP-proxim
proximalUP-(34)-sigma38p10 2991516 Gap 2.5 bits
distalUP-(40)-sigma38p10 2991516 Gap 3.1 bits
distalUP-proximalUP-sigma38p10 2991516 total 7.3 bits
... distalUP-proxim
proximalUP-(33)-sigma38p10 2991535 G
distalUP-(47)-sigma38p10 2991535 Gap
distalUP-proximalUP-sigma38p10 29915

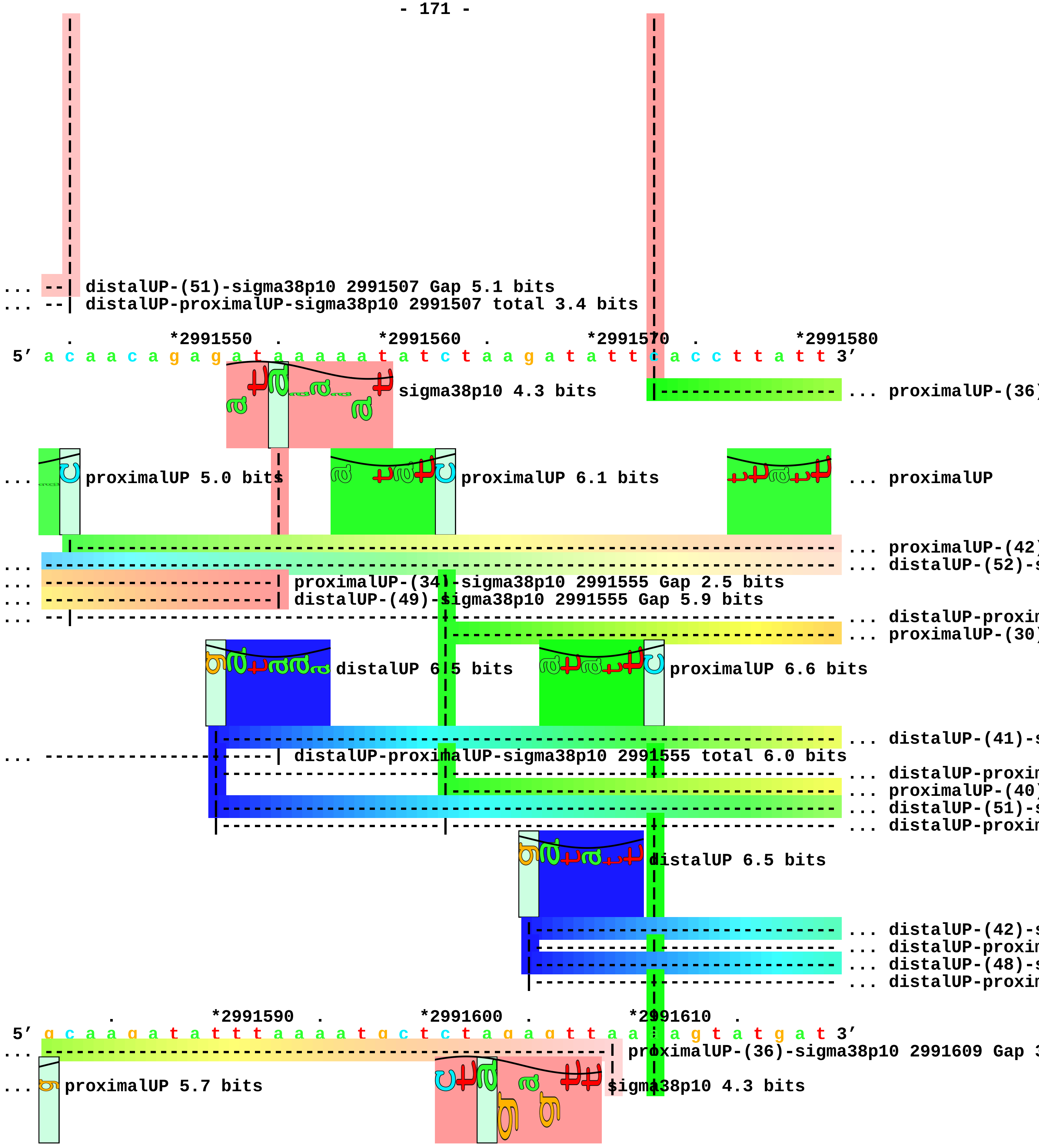
... --| distalUP-(51)-sigma38p10 2991507 Gap 5.1 bits
... --| distalUP-proximalUP-sigma38p10 2991507 total 3.4 bits
. *2991550 . *2991560 . *2991570 . *2991580
5' a c a a c a g a g a t a a a a a a t a t c t a a g a t a t t c a c c t t a t t 3'
sigma38p10 4.3 bits
... proximalUP-(36)
... proximalUP 5.0 bits
proximalUP 6.1 bits
... proximalUP
... proximalUP-(42)
... distalUP-(52)-s
... proximalUP-(34)-sigma38p10 2991555 Gap 2.5 bits
... distalUP-(49)-sigma38p10 2991555 Gap 5.9 bits
... distalUP-proxim
... proximalUP-(30)
distalUP 6.5 bits
proximalUP 6.6 bits
... distalUP-(41)-s
... distalUP-proximalUP-sigma38p10 2991555 total 6.0 bits
... distalUP-proxim
... proximalUP-(40)
... distalUP-(51)-s
... distalUP-proxim
distalUP 6.5 bits
... distalUP-(42)-s
... distalUP-proxim
... distalUP-(48)-s
... distalUP-proxim
. *2991590 . *2991600 . *2991610 .
5' g c a a g a t a t t t a a a a t g c t c t a g a g t t a a t a g t a t g a t 3'
... proximalUP-(36)-sigma38p10 2991609 Gap 3
... proximalUP 5.7 bits
sigma38p10 4.3 bits

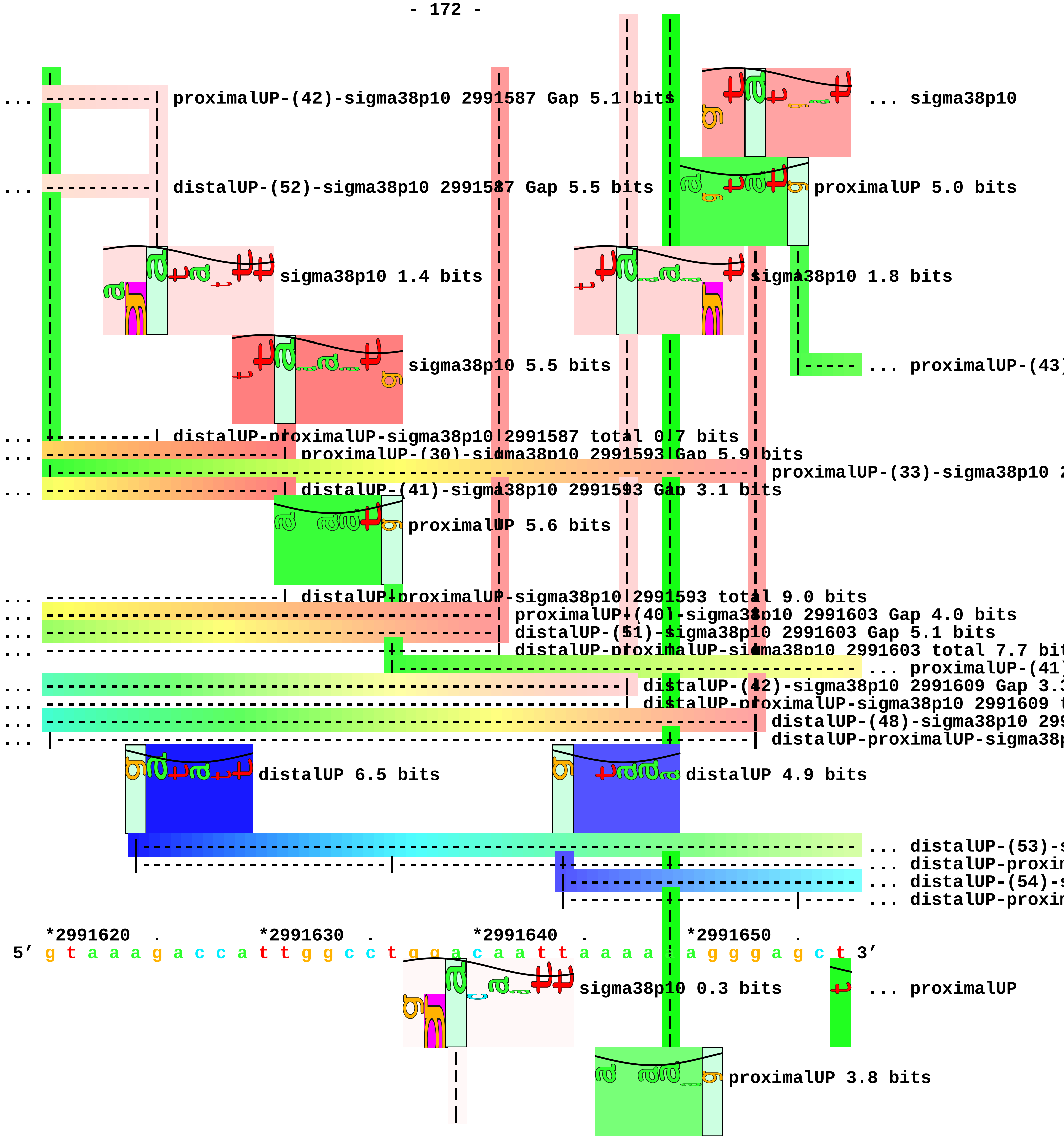
... proximalUP-(42)-sigma38p10 2991587 Gap 5.1 bits
... sigma38p10
... distalUP-(52)-sigma38p10 2991587 Gap 5.5 bits
proximalUP 5.0 bits
sigma38p10 1.4 bits
sigma38p10 1.8 bits
sigma38p10 5.5 bits
... proximalUP-(43)
... distalUP-proximalUP-sigma38p10 2991587 total 0.7 bits
... proximalUP-(30)-sigma38p10 2991593 Gap 5.9 bits
proximalUP-(33)-sigma38p10 2
... distalUP-(41)-sigma38p10 2991593 Gap 3.1 bits
proximalUP 5.6 bits
... distalUP-proximalUP-sigma38p10 2991593 total 9.0 bits
... proximalUP-(40)-sigma38p10 2991603 Gap 4.0 bits
... distalUP-(51)-sigma38p10 2991603 Gap 5.1 bits
... distalUP-proximalUP-sigma38p10 2991603 total 7.7 bit
... proximalUP-(41)
... distalUP-(42)-sigma38p10 2991609 Gap 3.
... distalUP-proximalUP-sigma38p10 2991609
... distalUP-(48)-sigma38p10 299
... distalUP-proximalUP-sigma38
distalUP 6.5 bits
distalUP 4.9 bits
... distalUP-(53)-
... distalUP-proxim
... distalUP-(54)-
... distalUP-proxim
*2991620 . *2991630 . *2991640 . *2991650 .
5′ g t a a a g a c c a t t g g c c t g g a c a a t t a a a a t a g g g a g c t 3′
sigma38p10 0.3 bits
... proximalUP
proximalUP 3.8 bits

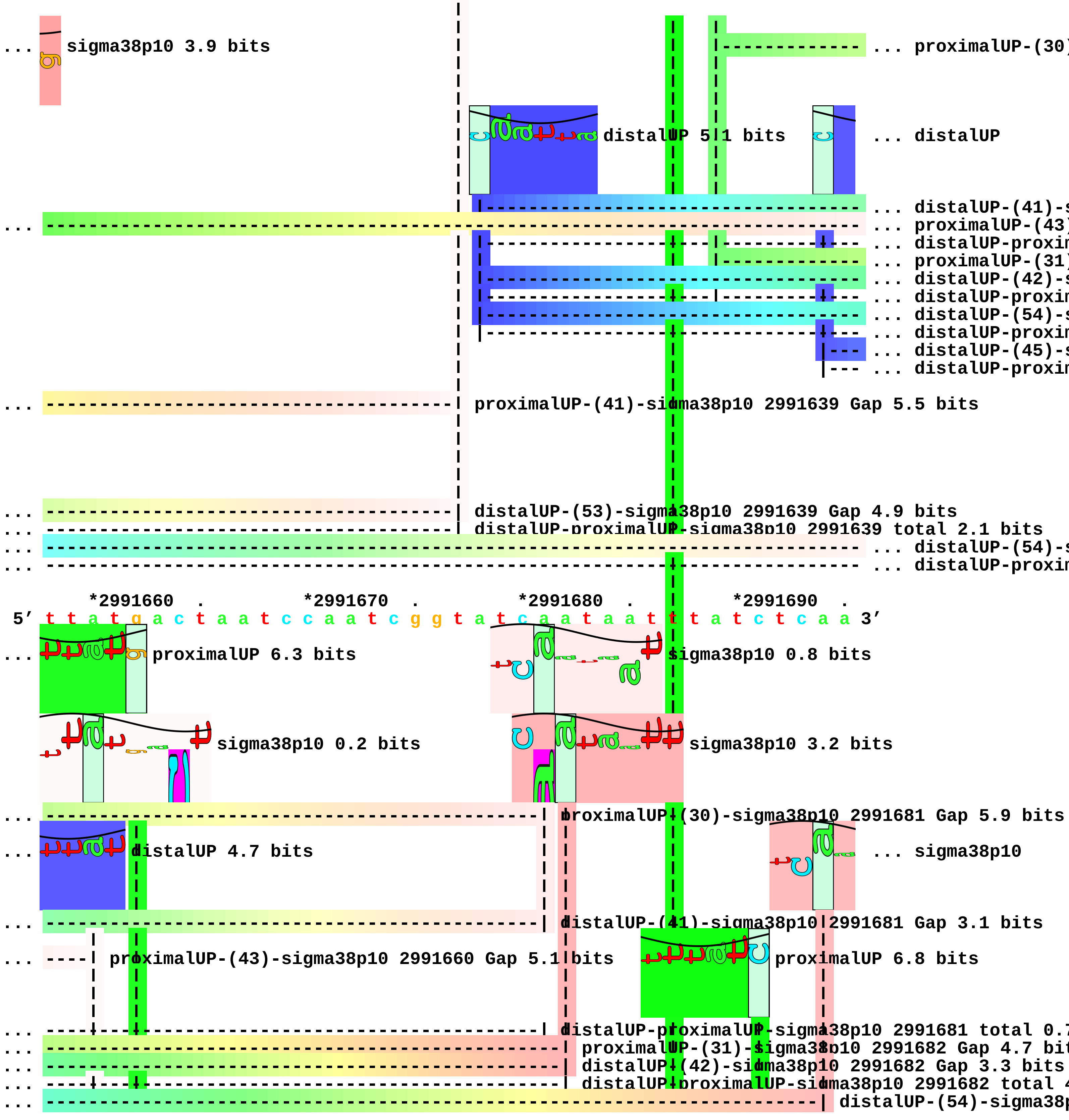
sigma38p10 3.9 bits
proximalUP-(30)
distalUP 5.1 bits
distalUP
distalUP-(41)-s
proximalUP-(43)
distalUP-proxin
proximalUP-(31)
distalUP-(42)-s
distalUP-proxin
distalUP-(54)-s
distalUP-proxin
distalUP-(45)-s
distalUP-proxin
proximalUP-(41)-sigma38p10 2991639 Gap 5.5 bits
distalUP-(53)-sigma38p10 2991639 Gap 4.9 bits
distalUP-proximalUP-sigma38p10 2991639 total 2.1 bits
distalUP-(54)-s
distalUP-proxin
*2991660 . *2991670 . *2991680 . *2991690 .
5′ t t a t g a c t a a t c c a a t c g g t a t c a a t a a t t t a t c t c a a 3′
proximalUP 6.3 bits
sigma38p10 0.8 bits
sigma38p10 0.2 bits
sigma38p10 3.2 bits
proximalUP-(30)-sigma38p10 2991681 Gap 5.9 bits
distalUP 4.7 bits
sigma38p10
distalUP-(41)-sigma38p10 2991681 Gap 3.1 bits
proximalUP-(43)-sigma38p10 2991660 Gap 5.1 bits
proximalUP 6.8 bits
distalUP-proximalUP-sigma38p10 2991681 total 0.7
proximalUP-(31)-sigma38p10 2991682 Gap 4.7 bit
distalUP-(42)-sigma38p10 2991682 Gap 3.3 bits
distalUP-proximalUP-sigma38p10 2991682 total 4
distalUP-(54)-sigma38p

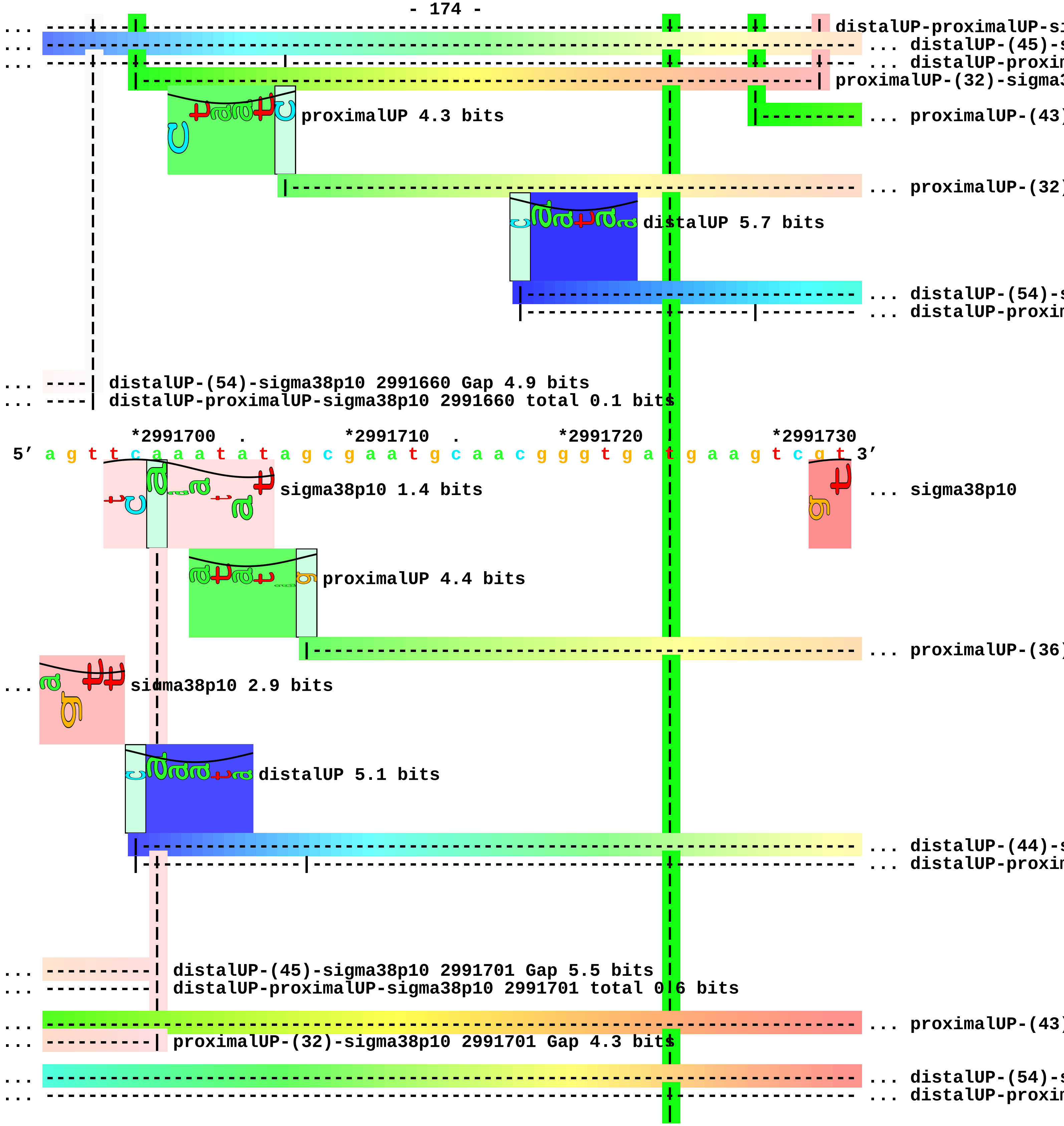
distalUP-proximalUP-si
... distalUP-(45)-s
... distalUP-proxim
proximalUP-(32)-sigma3
proximalUP 4.3 bits
... proximalUP-(43)
... proximalUP-(32)
distalUP 5.7 bits
... distalUP-(54)-s
... distalUP-proxim
distalUP-(54)-sigma38p10 2991660 Gap 4.9 bits
distalUP-proximalUP-sigma38p10 2991660 total 0.1 bits
*2991700
*2991710
*2991720
*2991730
5' a g t t c a a a t a t a g c g a a t g c a a c g g g t g a t g a a g t c g t 3'
sigma38p10 1.4 bits
... sigma38p10
proximalUP 4.4 bits
... proximalUP-(36)
sigma38p10 2.9 bits
distalUP 5.1 bits
... distalUP-(44)-s
... distalUP-proxim
distalUP-(45)-sigma38p10 2991701 Gap 5.5 bits
distalUP-proximalUP-sigma38p10 2991701 total 0.6 bits
... proximalUP-(43)
proximalUP-(32)-sigma38p10 2991701 Gap 4.3 bits
... distalUP-(54)-s
... distalUP-proxim

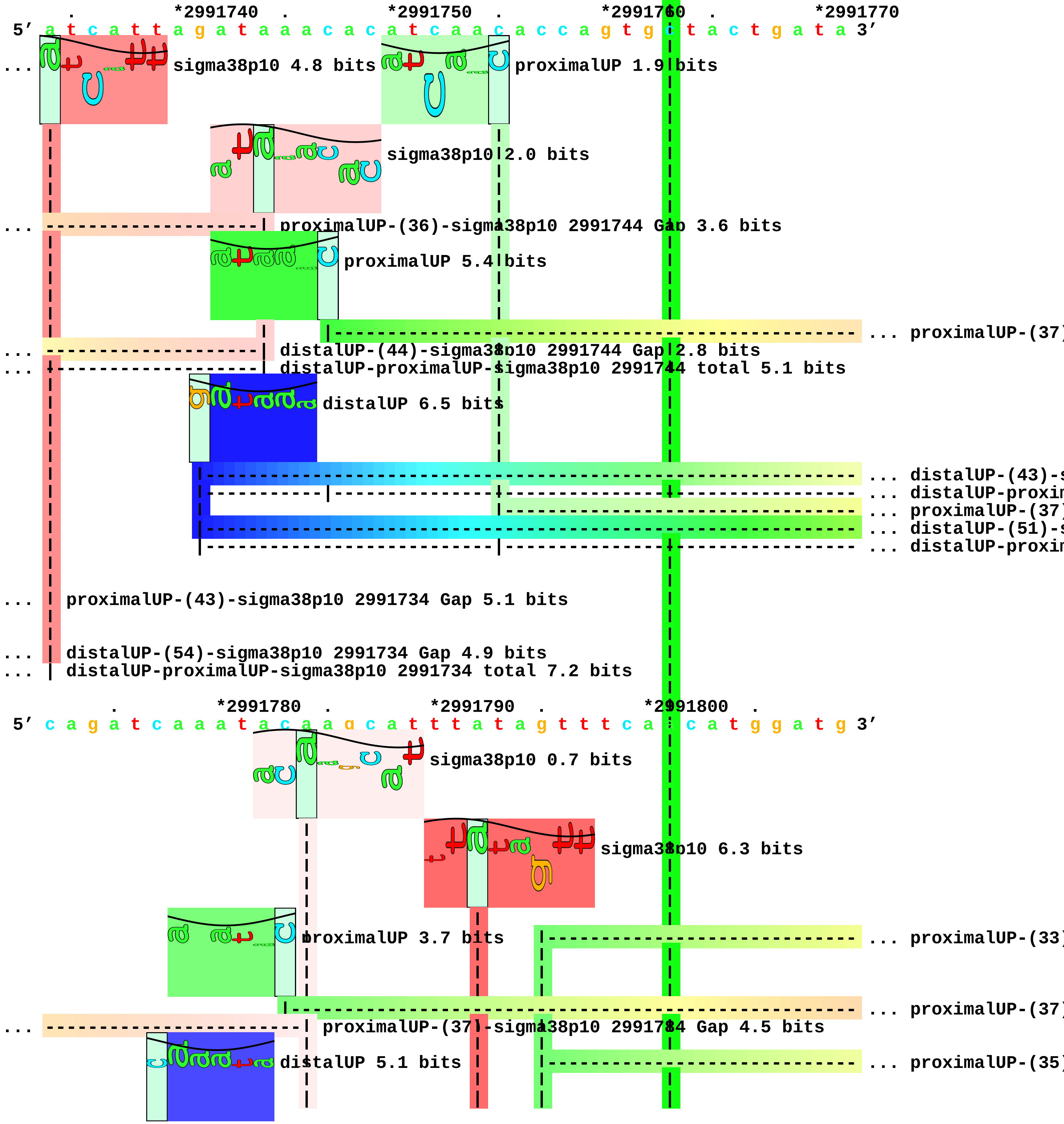

. *2991740 . *2991750 . *2991760 . *2991770
5′ a t c a t t a g a t a a a c a c a t c a a c a c c a g t g c t a c t g a t a 3′
... sigma38p10 4.8 bits
proximalUP 1.9 bits
sigma38p10 2.0 bits
... proximalUP-(36)-sigma38p10 2991744 Gap 3.6 bits
proximalUP 5.4 bits
... proximalUP-(37)
... distalUP-(44)-sigma38p10 2991744 Gap 2.8 bits
... distalUP-proximalUP-sigma38p10 2991744 total 5.1 bits
distalUP 6.5 bits
... distalUP-(43)-s
... distalUP-proxin
... proximalUP-(37)
... distalUP-(51)-s
... distalUP-proxin
... proximalUP-(43)-sigma38p10 2991734 Gap 5.1 bits
... distalUP-(54)-sigma38p10 2991734 Gap 4.9 bits
... distalUP-proximalUP-sigma38p10 2991734 total 7.2 bits
. *2991780 . *2991790 . *2991800 .
5′ c a g a t c a a a t a c a a g c a t t t a t a g t t t c a c c a t g g a t g 3′
sigma38p10 0.7 bits
sigma38p10 6.3 bits
proximalUP 3.7 bits
... proximalUP-(33)
... proximalUP-(37)
... proximalUP-(37)-sigma38p10 2991784 Gap 4.5 bits
distalUP 5.1 bits
... proximalUP-(35)

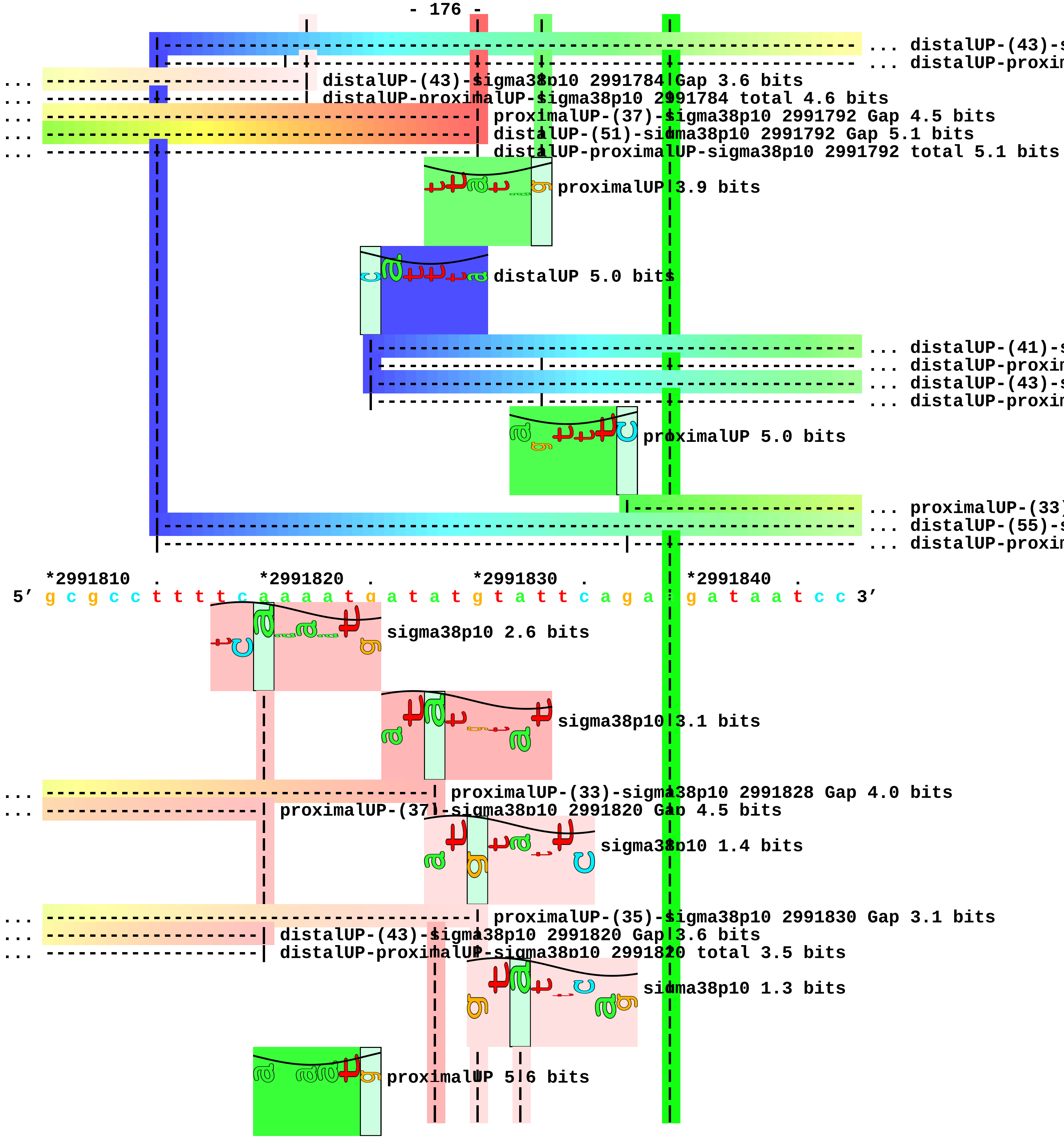
... distalUP-(43)-s
... distalUP-proxim
distalUP-(43)-sigma38p10 2991784 Gap 3.6 bits
distalUP-proximalUP-sigma38p10 2991784 total 4.6 bits
proximalUP-(37)-sigma38p10 2991792 Gap 4.5 bits
distalUP-(51)-sigma38p10 2991792 Gap 5.1 bits
distalUP-proximalUP-sigma38p10 2991792 total 5.1 bits
proximalUP 3.9 bits
distalUP 5.0 bits
... distalUP-(41)-s
... distalUP-proxim
... distalUP-(43)-s
... distalUP-proxim
proximalUP 5.0 bits
... proximalUP-(33)
... distalUP-(55)-s
... distalUP-proxim
*2991810 . *2991820 . *2991830 . *2991840 .
5′ g c g c c t t t t c a a a a t g a t a t g t a t t c a g a t g a t a a t c c 3′
sigma38p10 2.6 bits
sigma38p10 3.1 bits
proximalUP-(33)-sigma38p10 2991828 Gap 4.0 bits
proximalUP-(37)-sigma38p10 2991820 Gap 4.5 bits
sigma38p10 1.4 bits
proximalUP-(35)-sigma38p10 2991830 Gap 3.1 bits
distalUP-(43)-sigma38p10 2991820 Gap 3.6 bits
distalUP-proximalUP-sigma38p10 2991820 total 3.5 bits
sigma38p10 1.3 bits
proximalUP 5.6 bits

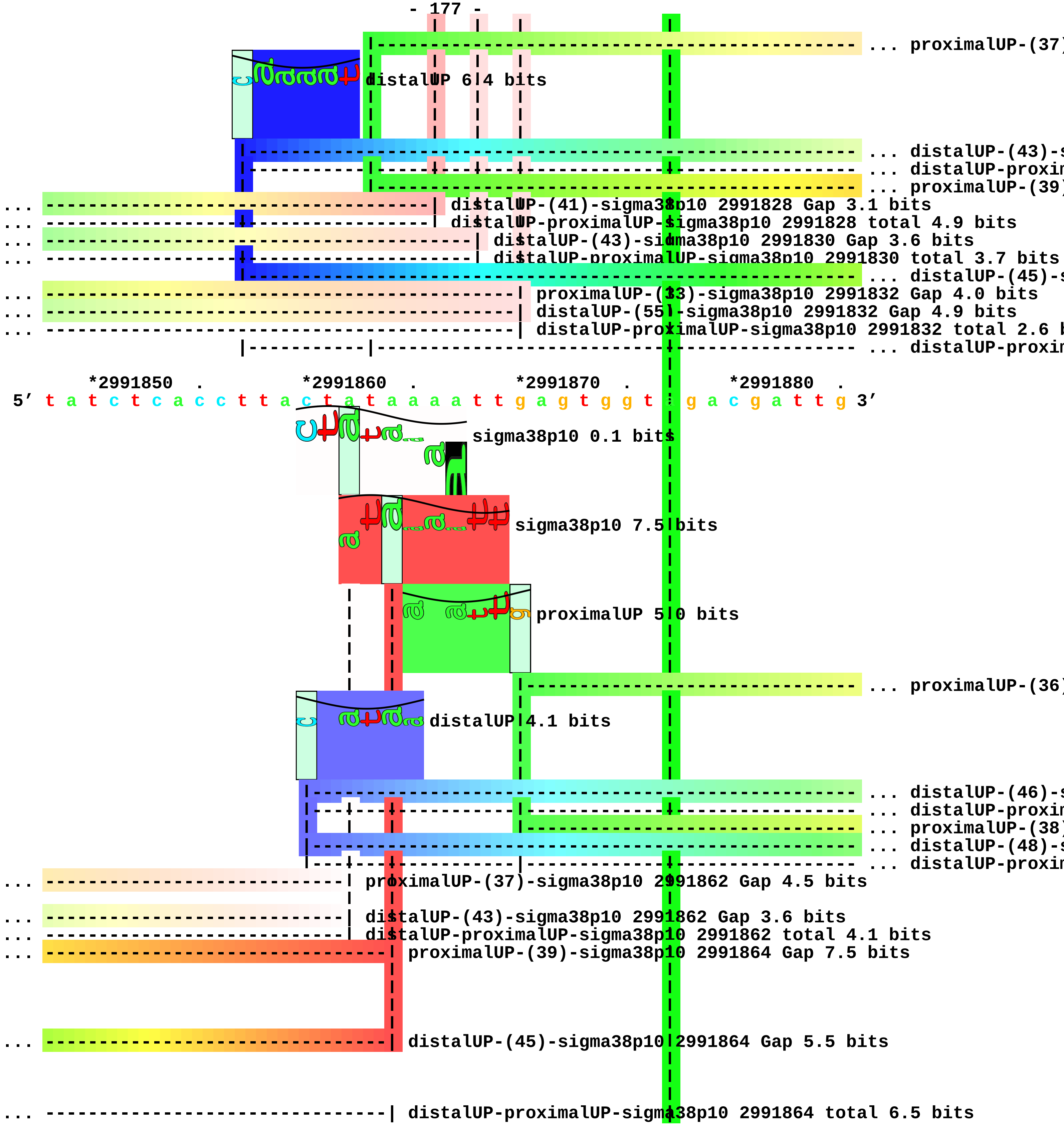
... proximalUP-(37)
distalUP 6.4 bits
... distalUP-(43)-s
... distalUP-proxim
... proximalUP-(39)
... distalUP-(41)-sigma38p10 2991828 Gap 3.1 bits
... distalUP-proximalUP-sigma38p10 2991828 total 4.9 bits
... distalUP-(43)-sigma38p10 2991830 Gap 3.6 bits
... distalUP-proximalUP-sigma38p10 2991830 total 3.7 bits
... distalUP-(45)-s
... proximalUP-(33)-sigma38p10 2991832 Gap 4.0 bits
... distalUP-(55)-sigma38p10 2991832 Gap 4.9 bits
... distalUP-proximalUP-sigma38p10 2991832 total 2.6 b
... distalUP-proxim
*2991850 . *2991860 . *2991870 . *2991880 .
5′ t a t c t c a c c t t a c t a t a a a a t t g a g t g g t t g a c g a t t g 3′
sigma38p10 0.1 bits
sigma38p10 7.5 bits
proximalUP 5.0 bits
... proximalUP-(36)
distalUP 4.1 bits
... distalUP-(46)-s
... distalUP-proxim
... proximalUP-(38)
... distalUP-(48)-s
... distalUP-proxim
... proximalUP-(37)-sigma38p10 2991862 Gap 4.5 bits
... distalUP-(43)-sigma38p10 2991862 Gap 3.6 bits
... distalUP-proximalUP-sigma38p10 2991862 total 4.1 bits
... proximalUP-(39)-sigma38p10 2991864 Gap 7.5 bits
... distalUP-(45)-sigma38p10 2991864 Gap 5.5 bits
... distalUP-proximalUP-sigma38p10 2991864 total 6.5 bits

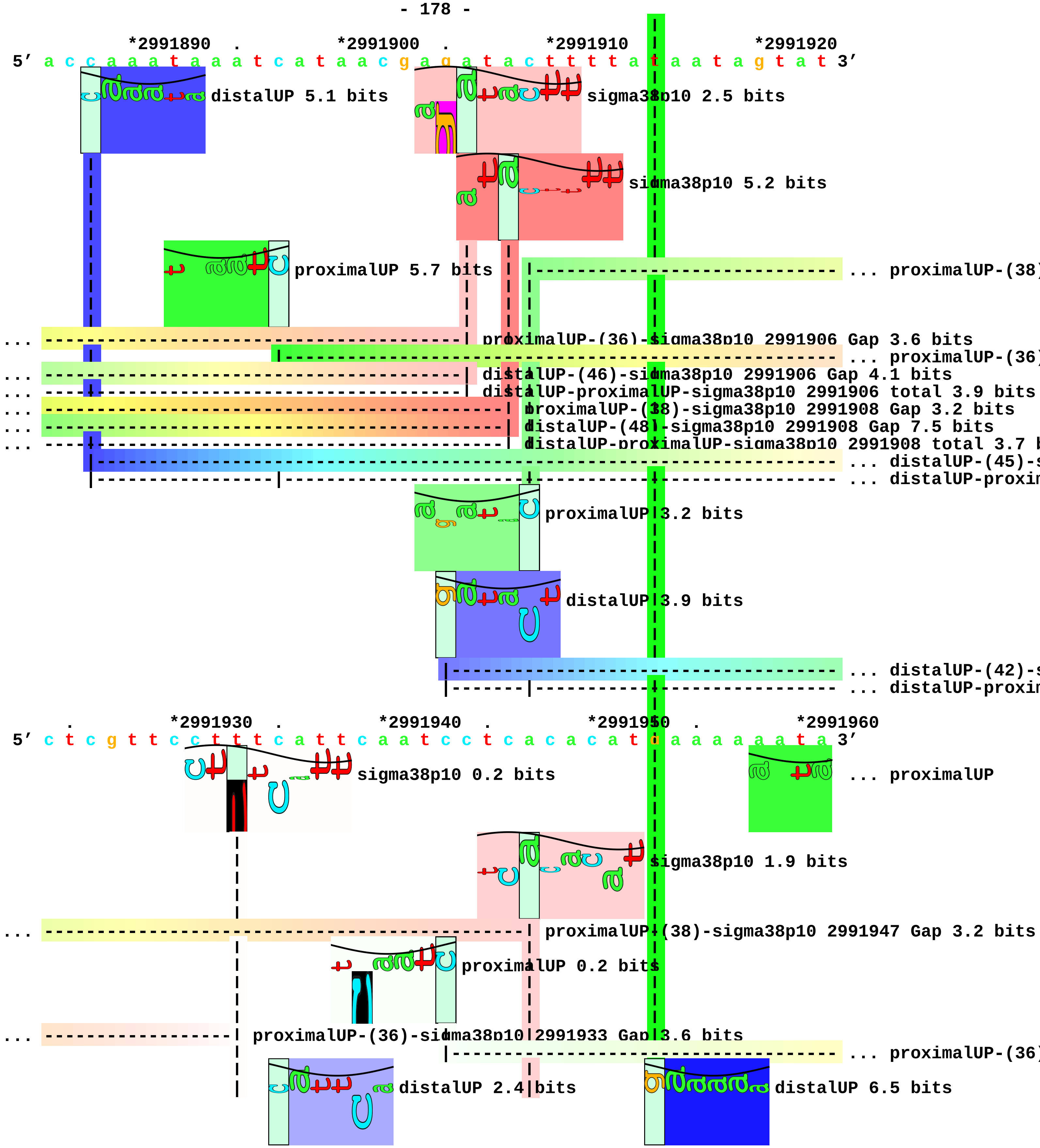
*2991890 . *2991900 . *2991910 *2991920
5′ a c c a a a t a a a t c a t a a c g a g a t a c t t t t a t a a t a g t a t 3′
distalUP 5.1 bits
sigma38p10 2.5 bits
sigma38p10 5.2 bits
proximalUP 5.7 bits
... proximalUP-(38)
... proximalUP-(36)-sigma38p10 2991906 Gap 3.6 bits
... proximalUP-(36)
... distalUP-(46)-sigma38p10 2991906 Gap 4.1 bits
... distalUP-proximalUP-sigma38p10 2991906 total 3.9 bits
... proximalUP-(38)-sigma38p10 2991908 Gap 3.2 bits
... distalUP-(48)-sigma38p10 2991908 Gap 7.5 bits
... distalUP-proximalUP-sigma38p10 2991908 total 3.7 b
... distalUP-(45)-s
... distalUP-proxin
proximalUP 3.2 bits
distalUP 3.9 bits
... distalUP-(42)-s
... distalUP-proxin
. *2991930 . *2991940 . *2991950 . *2991960
5′ c t c g t t c c t t t c a t t c a a t c c t c a c a c a t g a a a a a a t a 3′
sigma38p10 0.2 bits
... proximalUP
sigma38p10 1.9 bits
... proximalUP-(38)-sigma38p10 2991947 Gap 3.2 bits
proximalUP 0.2 bits
... proximalUP-(36)-sigma38p10 2991933 Gap 3.6 bits
... proximalUP-(36)
distalUP 2.4 bits
distalUP 6.5 bits

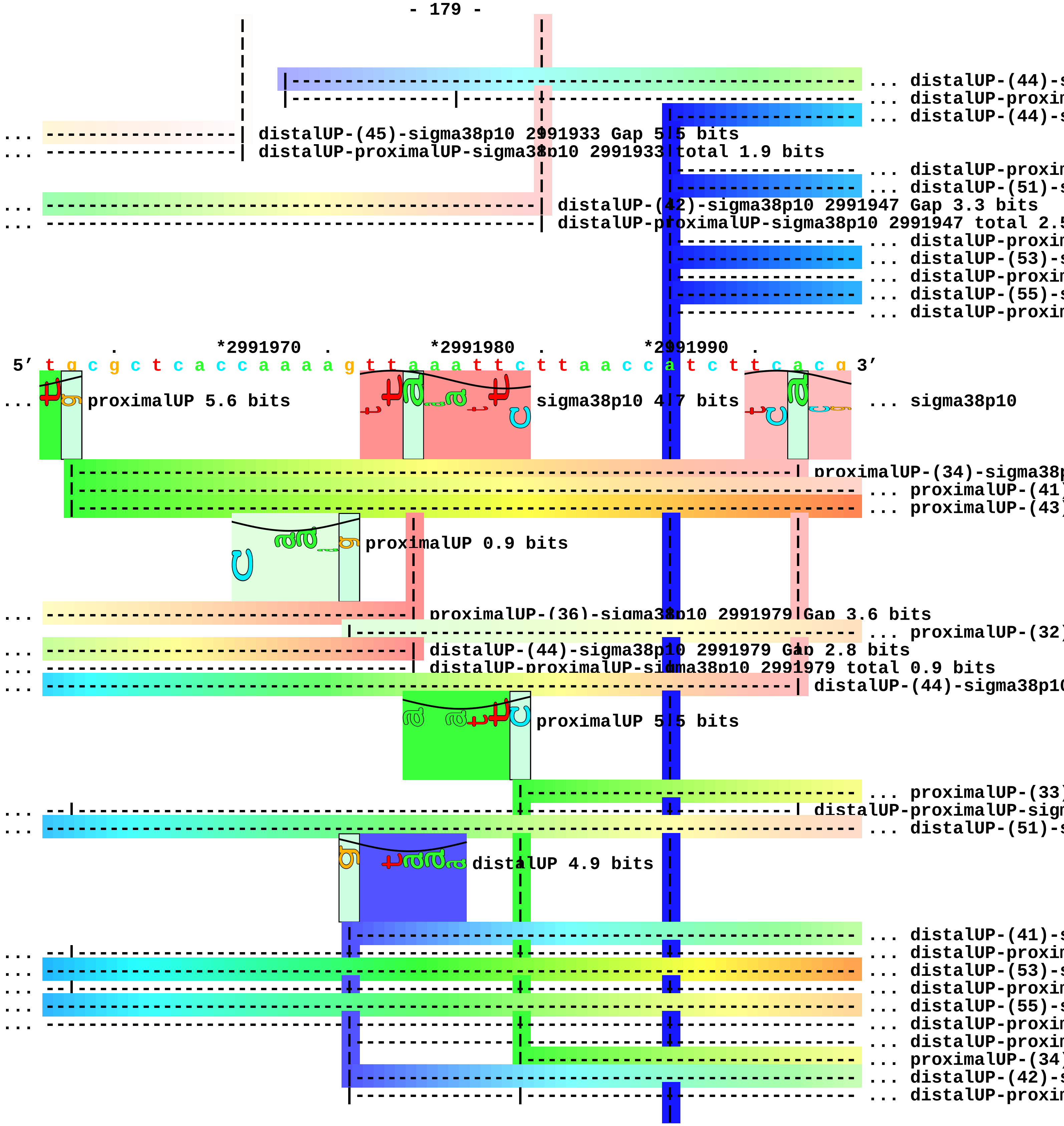

... distalUP-(44)-s
... distalUP-proxin
... distalUP-(44)-s
... distalUP-(45)-sigma38p10 2991933 Gap 5.5 bits
... distalUP-proximalUP-sigma38p10 2991933 total 1.9 bits
... distalUP-proxin
... distalUP-(51)-s
... distalUP-(42)-sigma38p10 2991947 Gap 3.3 bits
... distalUP-proximalUP-sigma38p10 2991947 total 2.5
... distalUP-proxin
... distalUP-(53)-s
... distalUP-proxin
... distalUP-(55)-s
... distalUP-proxin
*2991970
*2991980
*2991990
5′ t g c g c t c a c c a a a a g t t a a a t t c t t a a c c a t c t t c a c g 3′
... proximalUP 5.6 bits
sigma38p10 4.7 bits
... sigma38p10
proximalUP-(34)-sigma38p
... proximalUP-(41)
... proximalUP-(43)
proximalUP 0.9 bits
... proximalUP-(36)-sigma38p10 2991979 Gap 3.6 bits
... proximalUP-(32)
... distalUP-(44)-sigma38p10 2991979 Gap 2.8 bits
... distalUP-proximalUP-sigma38p10 2991979 total 0.9 bits
... distalUP-(44)-sigma38p10
proximalUP 5.5 bits
... proximalUP-(33)
... distalUP-proximalUP-sigm
... distalUP-(51)-s
distalUP 4.9 bits
... distalUP-(41)-s
... distalUP-proxin
... distalUP-(53)-s
... distalUP-proxin
... distalUP-(55)-s
... distalUP-proxin
... distalUP-proxin
... proximalUP-(34)
... distalUP-(42)-s
... distalUP-proxin

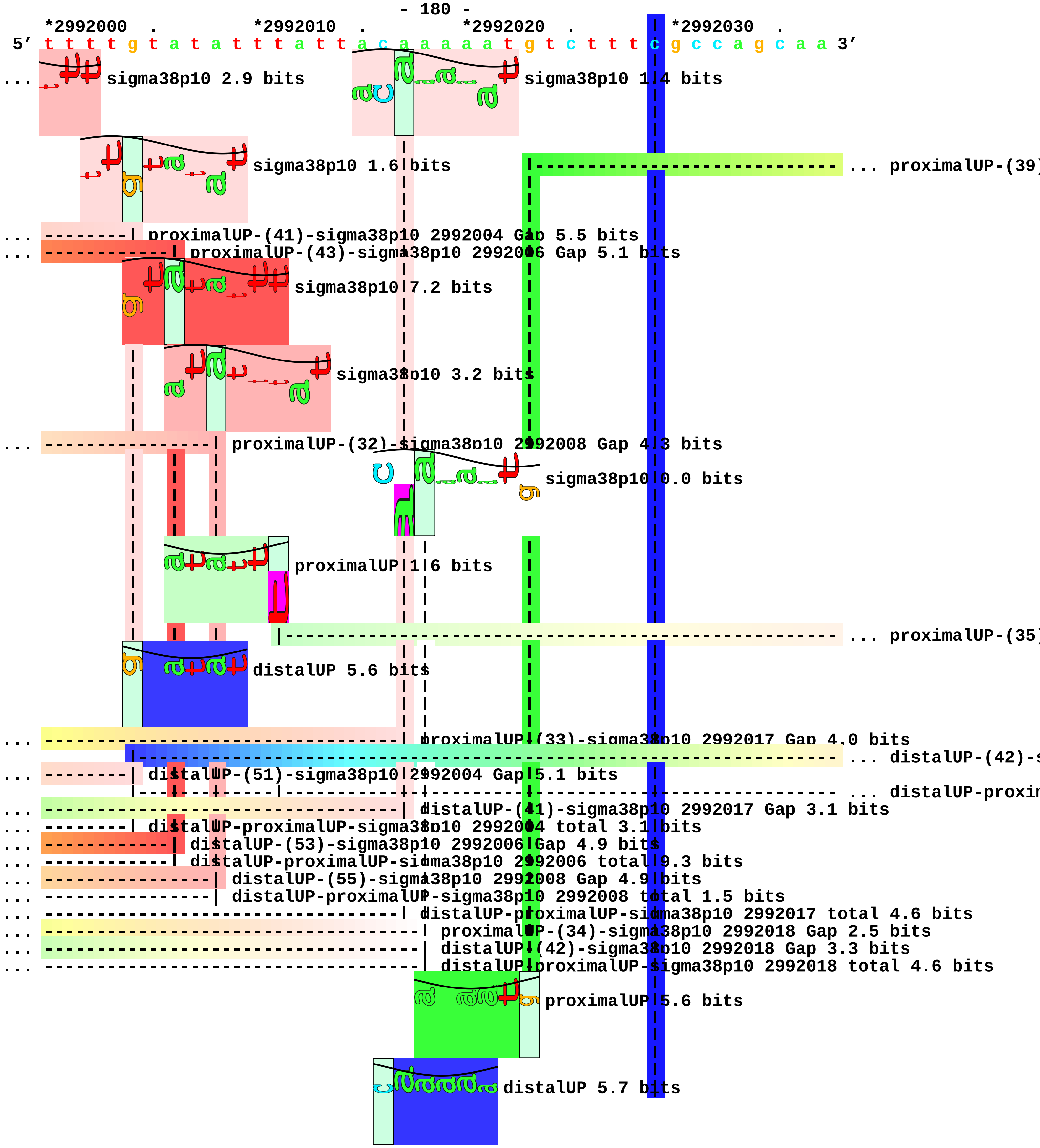
*2992000 . *2992010 . *2992020 . *2992030 .
5′ t t t t g t a t a t t t a t t t a c a a a a a a t g t c t t t c g c c a g c a a 3′
sigma38p10 2.9 bits
sigma38p10 1.4 bits
sigma38p10 1.6 bits
... proximalUP-(39)
... proximalUP-(41)-sigma38p10 2992004 Gap 5.5 bits
... proximalUP-(43)-sigma38p10 2992006 Gap 5.1 bits
sigma38p10 7.2 bits
sigma38p10 3.2 bits
... proximalUP-(32)-sigma38p10 2992008 Gap 4.3 bits
sigma38p10 0.0 bits
proximalUP 1.6 bits
... proximalUP-(35)
distalUP 5.6 bits
... proximalUP-(33)-sigma38p10 2992017 Gap 4.0 bits
... distalUP-(42)-s
... distalUP-(51)-sigma38p10 2992004 Gap 5.1 bits
... distalUP-proxim
... distalUP-(41)-sigma38p10 2992017 Gap 3.1 bits
... distalUP-proximalUP-sigma38p10 2992004 total 3.1 bits
... distalUP-(53)-sigma38p10 2992006 Gap 4.9 bits
... distalUP-proximalUP-sigma38p10 2992006 total 9.3 bits
... distalUP-(55)-sigma38p10 2992008 Gap 4.9 bits
... distalUP-proximalUP-sigma38p10 2992008 total 1.5 bits
... distalUP-proximalUP-sigma38p10 2992017 total 4.6 bits
... proximalUP-(34)-sigma38p10 2992018 Gap 2.5 bits
... distalUP-(42)-sigma38p10 2992018 Gap 3.3 bits
... distalUP-proximalUP-sigma38p10 2992018 total 4.6 bits
proximalUP 5.6 bits
distalUP 5.7 bits

```
                                                         ... distalUP-(46)-s
                                                         ... distalUP-proxim

      *2992040  .         *2992050  .         *2992060  .         *2992070  .
5′ c t c t t g c t c t a a t g a a g c a g g a a c a a t g t t c a g a a t c g 3′
                         sigma38p10 0.5 bits

... ------------------------------------------------| proximalUP-(39)-sigma38p10 2992062 Gap 7.5 bit
                                                       sigma38p10 1.2 bits

... ----------------| proximalUP-(35)-sigma38p10 2992046 Gap 3.1 bits

... ----------------| distalUP-(42)-sigma38p10 2992046 Gap 3.3 bits
... ----------------| distalUP-proximalUP-sigma38p10 2992046 total 1.4 bits

... ------------------------------------------------| distalUP-(46)-sigma38p10 2992062 Gap 4.1 bits
... ------------------------------------------------| distalUP-proximalUP-sigma38p10 2992062 total (

          *2992080  .         *2992090  .         *2992100  .         *2992110
5′ a a c c a a a t t t a a t t a a a g c t a t c g c t c t t g t t g a a a g c 3′
                                                         ... distalUP

                                                       |- ... distalUP-(43)-s
                                                       |- ... distalUP-proxim

    .         *2992120  .         *2992130  .         *2992140  .         *2992150
5′ a a t c t a a a a a a g g a t a g c a t a g g c a a g a a c a g a g a t a a 3′
...        distalUP 3.1 bits
```

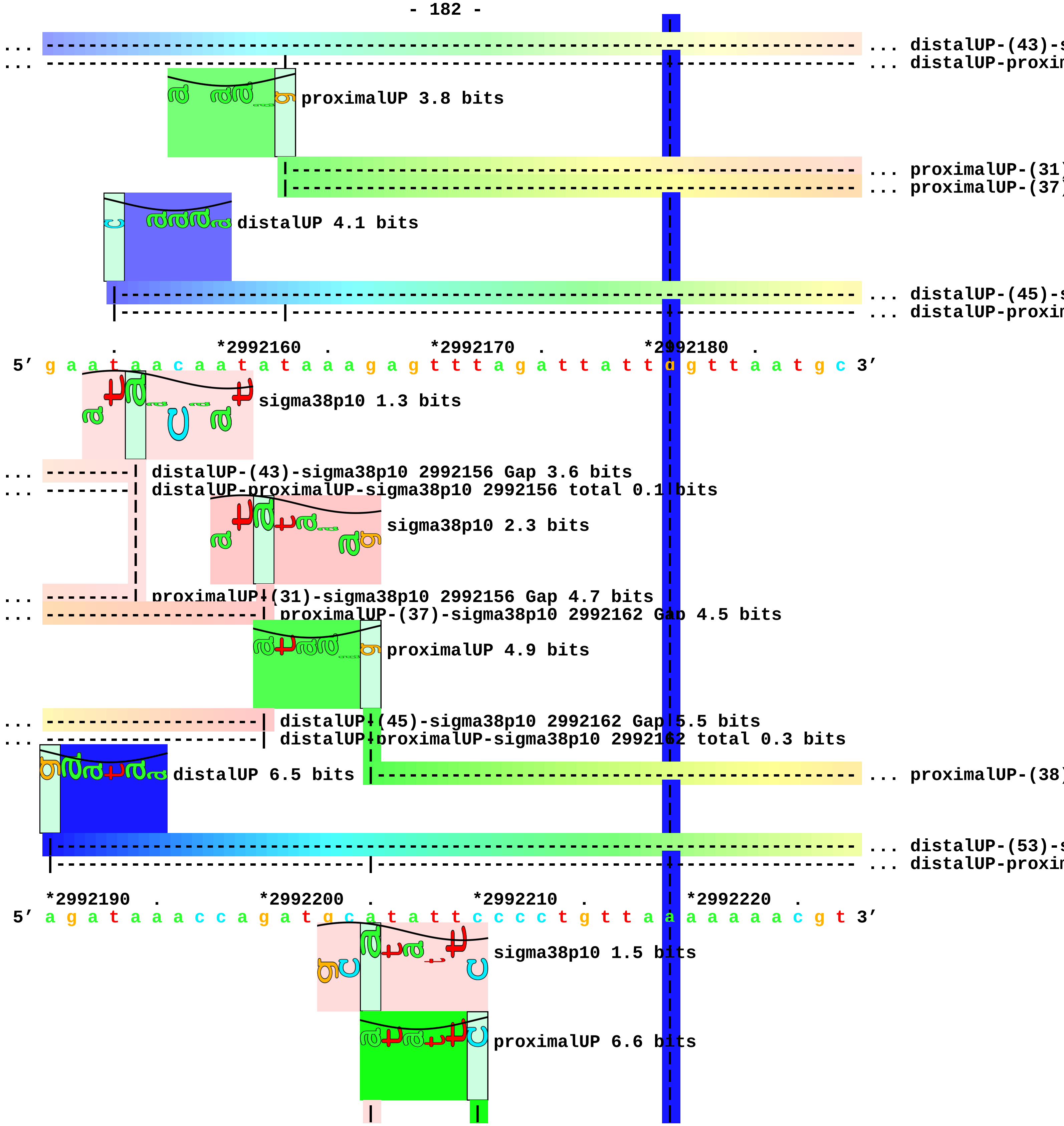

... distalUP-(43)-s
... distalUP-proxin
proximalUP 3.8 bits
... proximalUP-(31)
... proximalUP-(37)
distalUP 4.1 bits
... distalUP-(45)-s
... distalUP-proxin
*2992160
*2992170
*2992180
5′ g a a t a a c a a t a t a a a g a g t t t a g a t t a t t a g t t a a t g c 3′
sigma38p10 1.3 bits
distalUP-(43)-sigma38p10 2992156 Gap 3.6 bits
distalUP-proximalUP-sigma38p10 2992156 total 0.1 bits
sigma38p10 2.3 bits
proximalUP-(31)-sigma38p10 2992156 Gap 4.7 bits
proximalUP-(37)-sigma38p10 2992162 Gap 4.5 bits
proximalUP 4.9 bits
distalUP-(45)-sigma38p10 2992162 Gap 5.5 bits
distalUP-proximalUP-sigma38p10 2992162 total 0.3 bits
distalUP 6.5 bits
... proximalUP-(38)
... distalUP-(53)-s
... distalUP-proxin
*2992190
*2992200
*2992210
*2992220
5′ a g a t a a a c c a g a t g c a t a t t c c c c t g t t a a a a a a a c g t 3′
sigma38p10 1.5 bits
proximalUP 6.6 bits

distalUP 6.5 bits

|----------------------------------- ... proximalUP-(37)

|--------------------------------------------------------------------------- ... distalUP-(56)-s
|-----------------------------|----------|------------------------------------ ... distalUP-proxim

... -----------------------------| proximalUP-(38)-sigma38p10 2992205 Gap 3.2 bits
... -----------------------------| distalUP-(53)-sigma38p10 2992205 Gap 4.9 bits
... -----------------------------| distalUP-proximalUP-sigma38p10 2992205 total 4.8 bits

*2992230 . *2992240 . *2992250 . *2992260 .
5' g g a a t a a t a a a a g a t g a a c g a g a t t t g t t a g a t a a c c c 3'

distalUP 6.5 bits

sigma38p10 0.0 bits

proximalUP 3.4 bits

proximalUP 2.3 bits

... ------------------------------------------| proximalUP-(37)-sigma38p10 2992247 Gap 4.5 bits
... ------------------------------------------| distalUP-(56)-sigma38p10 2992247 Gap 3.8 bits
... --------------------------|---------------| distalUP-proximalUP-sigma38p10 2992247 total 4.9 bits
|----------------------------------------------------------- ... proximalUP-(32)
|------------------------------------------------------------------------- ... distalUP-(43)-s
|---------------------|-------------------------------------------|-------------- ... distalUP-proxim
|-------------- ... proximalUP-(33)

distalUP 2.8 bits

|------------------------------------------ ... distalUP-(44)-s
|-----------------------|-------------- ... distalUP-proxim

*2992270 . *2992280 . *2992290 . *2992300
5' t t g t c t g a a c a t t a a a a t a g g a a c c g a g a t t t t a t a t a 3'

sigma38p10 3.7 bits

sigma38p10 2.3 bits

... ------------| proximalUP-(32)-sigma38p10 2992272 Gap 4.3 bits
... ------------| distalUP-(43)-sigma38p10 2992272 Gap 3.6 bits
... ------------| distalUP-proximalUP-sigma38p10 2992272 total 5.8 bits
... --------------------------------------------------| proximalUP-(33)-sigma38p10 2992291 Gap 4.0 b

... --------------------------------------------------| distalUP-(44)-sigma38p10 2992291 Gap 2.8 bit
... --------------------------------------------------| distalUP-proximalUP-sigma38p10 2992291 tota

. *2992310 . *2992320 . *2992330 . *2992340
5' a c c a t t t t t c a c g t t g t g g a g t a a c g t g g c a a t g c c t g 3'

5′ g g g a c t t a 3′

piece 42, NC_000913, sibC, config: linear, direction: +, begin: 3054742, end: 3055002

```
        .         *3054750  .         *3054760  .         *3054770  .
5' a c t c c c c g t t g a a g a g t g g g a t a t c c c t c t t c c t g c g g 3'
   T timestamp:_2018Sep20_11-34-53

   *3054780  .         *3054790  .         *3054800  .         *3054810  .
5' t g g t t a c a c c g t c g a a a g t c t g g g a g t g g t a a g g g c g a 3'

     *3054820  .         *3054830  .         *3054840  .         *3054850  .
5' t a c a c c c g c a t c g c c c t g a t t g a c a t c g t t g a t t c t t t 3'

       *3054860  .         *3054870  .         *3054880  .         *3054890
5' g a c c t a a t t t a g t g a g t a a g g g t a a g g g a g g a t t g c t c 3'
                                             proximalUP 3.3 bits
                                             |------------------------------------- ... proximalUP-(36)
                                       distalUP 2.7 bits
                                  |-------------------------------------------------- ... distalUP-(42)-s
                                  |-----------|-------------------------------------- ... distalUP-proxim

    .         *3054900  .         *3054910  .         *3054920  .         *3054930
5' c t c c c c t g a g a c t g a c t g t t a a t a a g c g c t g a a a c t t a 3'
                                                   sigma38p10 2.6 bits
... ----------------------------------| proximalUP-(36)-sigma38p10 3054911 Gap 3.6 bits
                                                   proximalUP 3.8 bits
... ----------------------------------| distalUP-(42)-sigma38p10 3054911 Gap 3.3 bits
... ----------------------------------| distalUP-proximalUP-sigma38p10 3054911 total 1.8 bits
                                                         |-------------------------- ... proximalUP-(33)
                                                   distalUP 5.5 bits
                                        |-------------------------------------------- ... distalUP-(41)-s
                                        |---------------|---------------------------- ... distalUP-proxim

        .         *3054940  .         *3054950  .         *3054960  .
5' t g a g t a a c a g t a c a a t c a g t a t g a t g a c a a g t c g c a t c 3'
                                                   sigma38p10 3.9 bits
```

```
                                        |
                                        |
... ----------------------------------------| proximalUP-(33)-sigma38p10 3054952 Gap 4.0 bits
                                        |
... ----------------------------------------| distalUP-(41)-sigma38p10 3054952 Gap 3.1 bits
... ----------------------------------------| distalUP-proximalUP-sigma38p10 3054952 total 6.1 bits

    *3054970  .         *3054980  .         *3054990  .         *3055000
 5' a t a a c c c t t c t c c t t c a a g c c c t c g c t t c g g t g 3'
```

piece 43, NC_000913, scpA, config: linear, direction: +, begin: 3058550, end: 3058799

*3058550 . *3058560 . *3058570 . *3058580 .
5′ c t g g g c t a t t t c a a c g a c g g a t g c t c t a c t g g c a c c g c 3′
T timestamp:_2018Sep20_11-34-53

*3058590 . *3058600 . *3058610 . *3058620 .
5′ t t t g c t c c t g a a a g c c g c a t g a t g c g t a a a g t c a c t g a 3′

*3058630 . *3058640 . *3058650 . *3058660
5′ t g c g t t a c t c g a t t a t g g t c a c a a a g t c c t t c g t c a g g 3′
... distalUP
... distalUP-(43)-s
... distalUP-proxin

. *3058670 . *3058680 . *3058690 . *3058700
5′ a t t a a t c c a t c a a a t a a t g c c t g a t a g c a c a t a t c a g g 3′
... distalUP 6.4 bits ... proximalUP-(33)
... ... distalUP-(43)-s
... ... distalUP-proxin
proximalUP 6.8 bits ... proximalUP-(40)
... proximalUP-(36)
proximalUP 6.7 bits
distalUP 5.1 bits
... distalUP-(41)-s
... distalUP-proxin
... distalUP-(48)-s
... distalUP-proxin

. *3058710 . *3058720 . *3058730 .
5′ c g t t g t c c t c a c t t c t t t t t g t a t t c c t t g a a t c a c a t 3′
... proximalUP-(33)-sigma38p10 3058715 Gap 4.0 bits
... distalUP-(43)-sigma38p10 3058706 Gap 3.6 bits
... distalUP-proximalUP-sigma38p10 3058706 total 6.2 bits
... proximalUP-(40)-sigma38p10 3058722 Gap 4.0 bits
... proximalUP-(36)-sigma38p10 3058706 Gap 3.6 bits
sigma38p10 0.1 bits

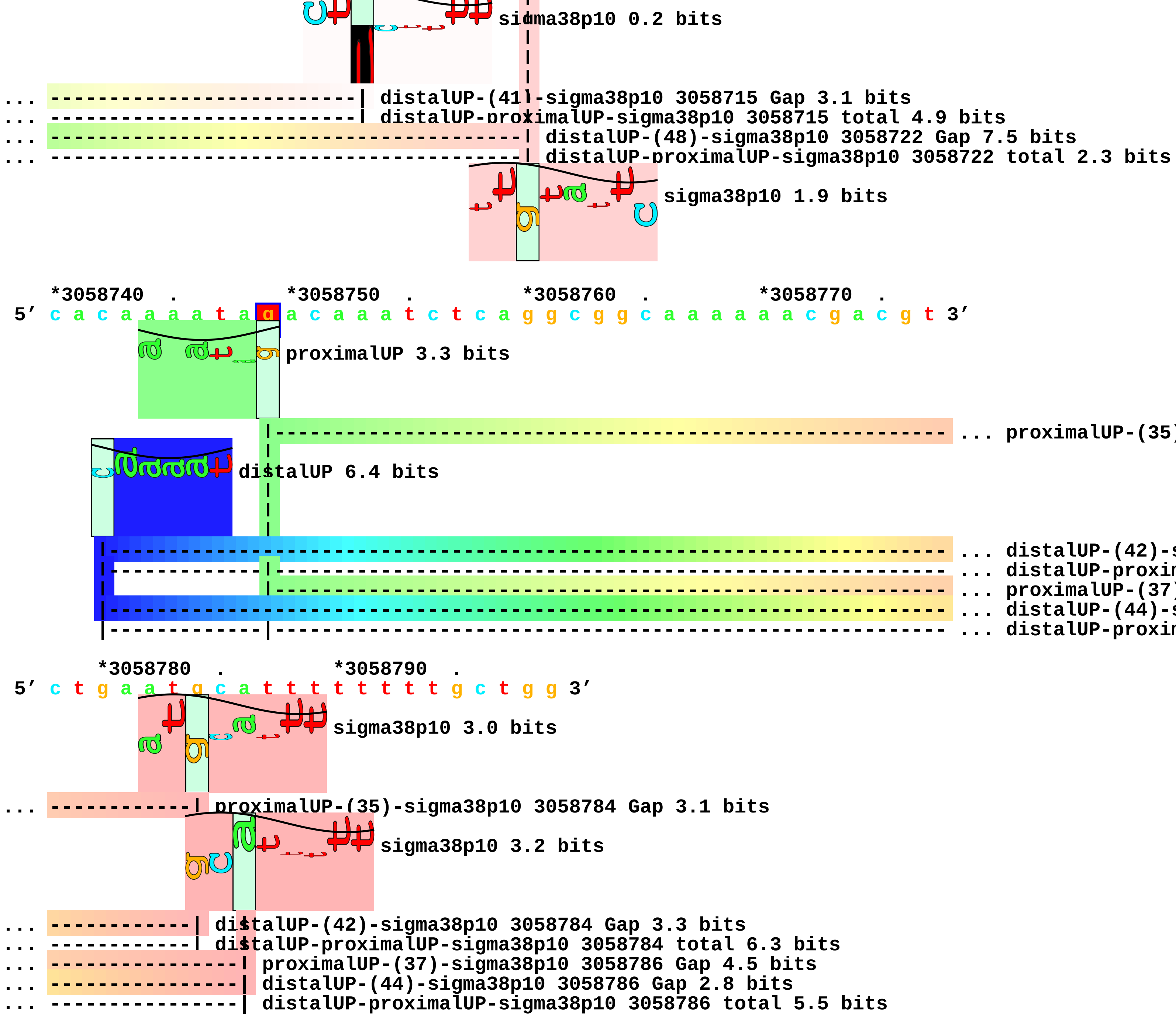
sigma38p10 0.2 bits
... distalUP-(41)-sigma38p10 3058715 Gap 3.1 bits
... distalUP-proximalUP-sigma38p10 3058715 total 4.9 bits
... distalUP-(48)-sigma38p10 3058722 Gap 7.5 bits
... distalUP-proximalUP-sigma38p10 3058722 total 2.3 bits
sigma38p10 1.9 bits
*3058740 . *3058750 . *3058760 . *3058770 .
5′ c a c a a a a t a g a c a a a t c t c a g g c g g c a a a a a a c g a c g t 3′
proximalUP 3.3 bits
... proximalUP-(35)
distalUP 6.4 bits
... distalUP-(42)-s
... distalUP-proxim
... proximalUP-(37)
... distalUP-(44)-s
... distalUP-proxim
*3058780 . *3058790 .
5′ c t g a a t g c a t t t t t t t t g c t g g 3′
sigma38p10 3.0 bits
... proximalUP-(35)-sigma38p10 3058784 Gap 3.1 bits
sigma38p10 3.2 bits
... distalUP-(42)-sigma38p10 3058784 Gap 3.3 bits
... distalUP-proximalUP-sigma38p10 3058784 total 6.3 bits
... proximalUP-(37)-sigma38p10 3058786 Gap 4.5 bits
... distalUP-(44)-sigma38p10 3058786 Gap 2.8 bits
... distalUP-proximalUP-sigma38p10 3058786 total 5.5 bits

piece 44, NC_000913, yggE, config: linear, direction: -, begin: 3066199, end: 3066000

```
              .          *3066190   .          *3066180   .          *3066170   .
5' g t t a g g c g c t a t c t g a t g g a a a a a t a a a a c a g a g g c g c 3'

   T timestamp:_2018Sep20_11-34-53                            proximalUP 3.9 bits

                                                                       ... proximalUP-(31)
                                          distalUP 6.5 bits

                                                                       ... distalUP-(42)-s
                                                                       ... sigma38p10-prox
                                                                       ... proximalUP-(32)
                                                                       ... distalUP-(43)-s
                                                                       ... sigma38p10-prox

     *3066160   .          *3066150   .          *3066140   .          *3066130   .
5' t a a g c t t g c c t c c c a g a g g t c c t g a a t t t t g g g c a a g t c 3'
                                                         sigma38p10 5.9 bits

... ------------------------------------------| proximalUP-(31)-sigma38p10 3066139 Gap 4.7 bits
                                                           sigma38p10 0.2 bits

... ------------------------------------------| distalUP-(42)-sigma38p10 3066139 Gap 3.3 bits
... ------------------------------------------| sigma38p10-proximalUP-distalUP 3066139 total 8.4 b
... --------------------------------------------| proximalUP-(32)-sigma38p10 3066138 Gap 4.3 bits
... --------------------------------------------| distalUP-(43)-sigma38p10 3066138 Gap 3.6 bits
... --------------------------------------------| sigma38p10-proximalUP-distalUP 3066138 total 2.8
                                                              proximalUP 6.1 bits

                                                                       ... proximalUP-(32)
                                                      distalUP 1.0 bits

                                                                       ... distalUP-(41)-s
                                                                       ... sigma38p10-prox
                                                                       ... proximalUP-(39)
                                                         distalUP 6.6 bits

                                                                       ... distalUP-(46)-s
                                                                       ... sigma38p10-prox
```

*3066120 . *3066110 . *3066100 . *3066090
5′ a a t t a a c a c g g a g a g a c t a a c g t g a a g t t c a a a g t t a t 3′

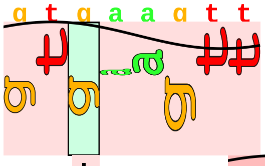
sigma38p10 1.4 bits

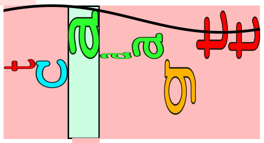
sigma38p10 2.9 bits

... ----------------------------------------------| proximalUP-(32)-sigma38p10 3066100 Gap 4.3 bits
... ----------------------------------------------| distalUP-(41)-sigma38p10 3066100 Gap 3.1 bits
... ----------------------------------------------| sigma38p10-proximalUP-distalUP 3066100 total 1.1
... ------------------------------------------------------------| proximalUP-(39)-sigma38p10 306609
... ------------------------------------------------------------| distalUP-(46)-sigma38p10 3066093 C
... ------------------------------------------------------------| sigma38p10-proximalUP-distalUP 306

. *3066080 . *3066070 . *3066060 . *3066050
5′ c g c c c t g g c g g c a t t a a t g g g t a t t a g c g g g a t g g c a g 3′

. *3066040 . *3066030 . *3066020 . *3066010
5′ c g c a g g c t a a c g a a t t g c c g g a t g g a c c g c a t a t t g t c 3′

. *3066000
5′ a c c t c c g g t a 3′

piece 45, NC_000913, ygjR, config: linear, direction: +, begin: 3235183, end: 3235431

```
         .         *3235190  .         *3235200  .         *3235210  .         *3235220
5' c a g c c g a a a t a t c t c g g g c c t t c g g t c a t g a t t g g g c c 3'
   T timestamp:_2018Sep20_11-34-53

         .         *3235230  .         *3235240  .         *3235250  .
5' g t t t a g t g a g c a t g g c t g t c c g g c g c a g a a a t a a t g c g 3'
                                                            proximalUP 6.7 bits
                                                            |----- ... proximalUP-(37)
                                                distalUP 5.9 bits
                                                |---------------------- ... distalUP-(45)-s
                                                |---------------|----- ... distalUP-proxin

   *3235260  .         *3235270  .         *3235280  .         *3235290  .
5' t a t c t g c g c a c g t c g a a g a t g a a a a a g g c g t g c t a c a t 3'
                                                                    ... sigma38p10
... ------------------------------------------------------------| proximalUP-(37)-sigma38p10
                                                                    ... distalUP
... ------------------------------------------------------------| distalUP-(45)-sigma38p10 3
... ------------------------------------------------------------| distalUP-proximalUP-sigma3
                                                             |----- ... distalUP-(44)-s
                                                             |----- ... distalUP-proxin

      *3235300  .         *3235310  .         *3235320  .         *3235330
5' t g a c g a c a g a a t c c c t t t a t g g a g t a t c c a c g c g t t a t 3'
... sigma38p10 6.1 bits |-------------------------------------------- ... proximalUP-(29)
                  proximalUP 5.5 bits
... distalUP 0.7 bits          proximalUP 6.3 bits
                                |----------------------------------- ... proximalUP-(31)
                  distalUP 2.6 bits
```

```
... --------------------------------------------------------------------------- ... distalUP-(44)-s
... ------------|-----------|---------------------------------------------------- ... distalUP-proxim
            |---------------------------------------------------------------- ... distalUP-(45)-s
            |---------------------------|------------------------------------ ... distalUP-proxim

    .         *3235340  .         *3235350  .         *3235360  .         *3235370
 5' g a t a c g t t t c g c t g t g a t t g g t a c g a a c t g g a t c a c t c 3'
... ------| proximalUP-(29)-sigma38p10 3235338 Gap 7.5 bits
```

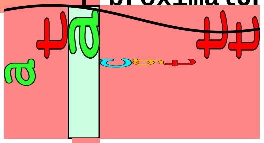

sigma38p10 5.2 bits

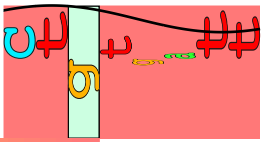

sigma38p10 5.8 bits

```
... --------------------------| proximalUP-(31)-sigma38p10 3235348 Gap 4.7 bits

... ------| distalUP-(44)-sigma38p10 3235338 Gap 2.8 bits
... ------| distalUP-proximalUP-sigma38p10 3235338 total 1.1 bits
... --------------------------| distalUP-(45)-sigma38p10 3235348 Gap 5.5 bits
... --------------------------| distalUP-proximalUP-sigma38p10 3235348 total 4.5 bits

        .         *3235380  .         *3235390  .         *3235400  .         *3235410
 5' g c c a g t t c g t c g a g g c c g c c c a t g a g a g c g g t a a a t a c 3'

            .         *3235420  .         *3235430
 5' a a g t t a a c c g c c g t a t a t t c c 3'
```

piece 46, NC_000913, rpoH, config: linear, direction: -, begin: 3599149, end: 3598900

```
            .         *3599140  .         *3599130  .         *3599120  .
5' g a t g a a t g c c t g c t a t t g c t g c t g g t a t g c t c g a t g a t 3'
  T timestamp:_2018Sep20_11-34-53

  *3599110  .         *3599100  .         *3599090  .         *3599080  .
5' t g g c t g g g t g g c a g c g t g g c t t g c c a c g g t a c a a c a t t 3'
                                                             ... proximalUP
                                                             distalUP 1.6 bits
                                                             ... distalUP-(42)-s
                                                             ... sigma38p10-prox
                                                             ... distalUP
                                                             ... distalUP-(41)-s
                                                             ... sigma38p10-prox
                                                             ... distalUP-(52)-s
                                                             ... sigma38p10-prox

    *3599070  .         *3599060  .         *3599050  .         *3599040
5' t a c g c c a c t t t a c g c c t g a a t a a t a a a a g c g t g t t a t a 3'
... proximalUP 4.9 bits                                      ... sigma38p10
                                                             proximalUP-(33)-sigma38p
...                                                          distalUP-(42)-sigma38p10
...                                                          sigma38p10-proximalUP-di
... distalUP 5.0 bits                                        ... sigma38p10
...                                                          distalUP-(41)-sigma3
...                                                          sigma38p10-proximalU
...                                                          ... distalUP-(52)-s
...                                                          ... sigma38p10-prox
                                                             proximalUP-(35)-sign
          proximalUP 1.9 bits
                                                             ... proximalUP-(36)

.         *3599030  .         *3599020  .         *3599010  .         *3599000
5' c t c t t t c c c t g c a a t g g g t t c c g t a g c a g g g a a a g a g a 3'
... sigma38p10 6.4 bits
```

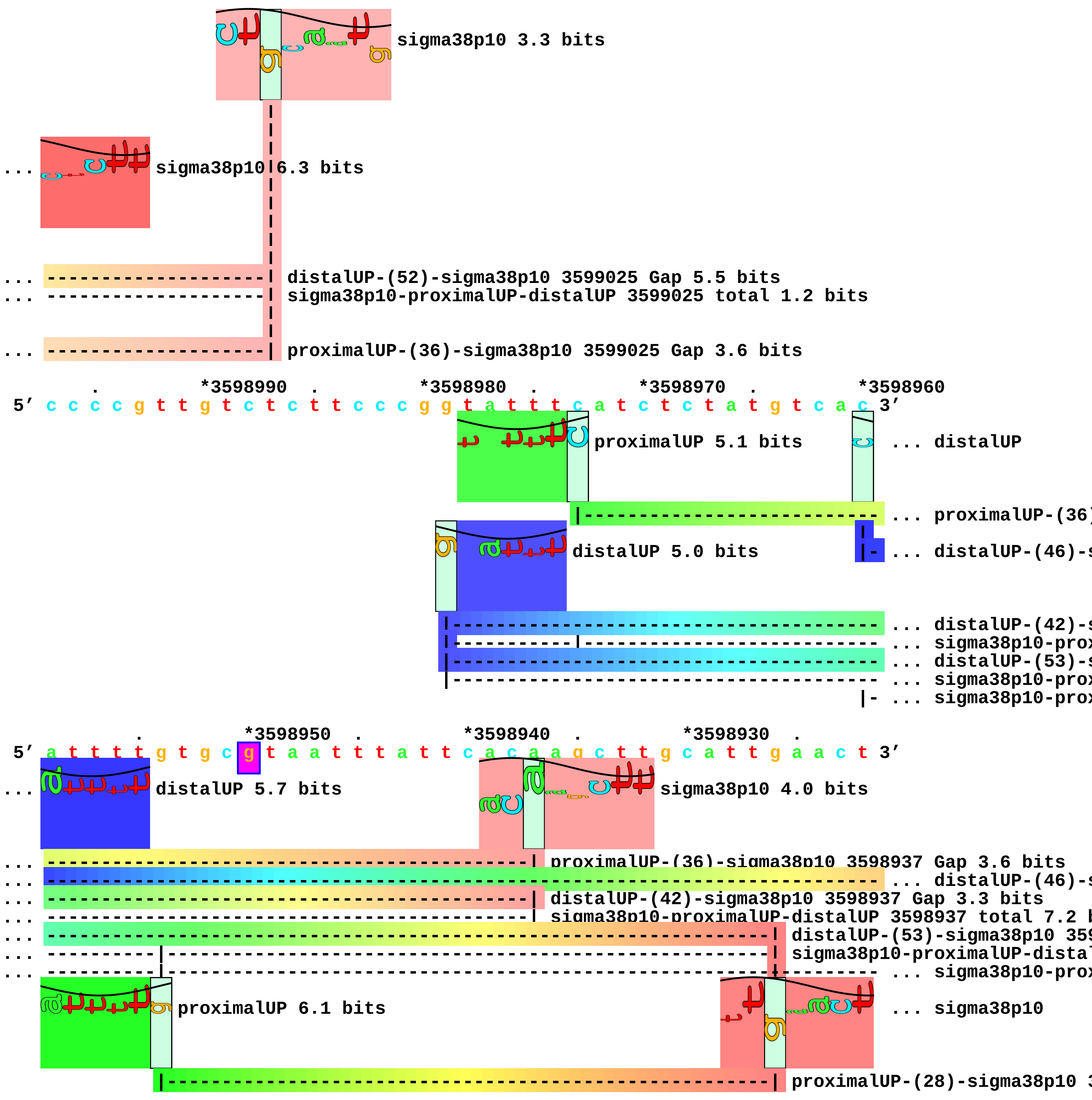
sigma38p10 3.3 bits
... sigma38p10 6.3 bits
... distalUP-(52)-sigma38p10 3599025 Gap 5.5 bits
... sigma38p10-proximalUP-distalUP 3599025 total 1.2 bits
... proximalUP-(36)-sigma38p10 3599025 Gap 3.6 bits
. *3598990 . *3598980 . *3598970 . *3598960
5′ c c c c g t t g t c t c t t c c c g g t a t t t c a t c t c t a t g t c a c 3′
proximalUP 5.1 bits
... distalUP
... proximalUP-(36)
distalUP 5.0 bits
... distalUP-(46)-s
... distalUP-(42)-s
... sigma38p10-prox
... distalUP-(53)-s
... sigma38p10-prox
... sigma38p10-prox
. *3598950 . *3598940 . *3598930 .
5′ a t t t t g t g c g t a a t t t a t t c a c a a g c t t g c a t t g a a c t 3′
... distalUP 5.7 bits
sigma38p10 4.0 bits
... proximalUP-(36)-sigma38p10 3598937 Gap 3.6 bits
... distalUP-(46)-s
... distalUP-(42)-sigma38p10 3598937 Gap 3.3 bits
... sigma38p10-proximalUP-distalUP 3598937 total 7.2 b
... distalUP-(53)-sigma38p10 359
... sigma38p10-proximalUP-dista
... sigma38p10-prox
proximalUP 6.1 bits
... sigma38p10
proximalUP-(28)-sigma38p10 3

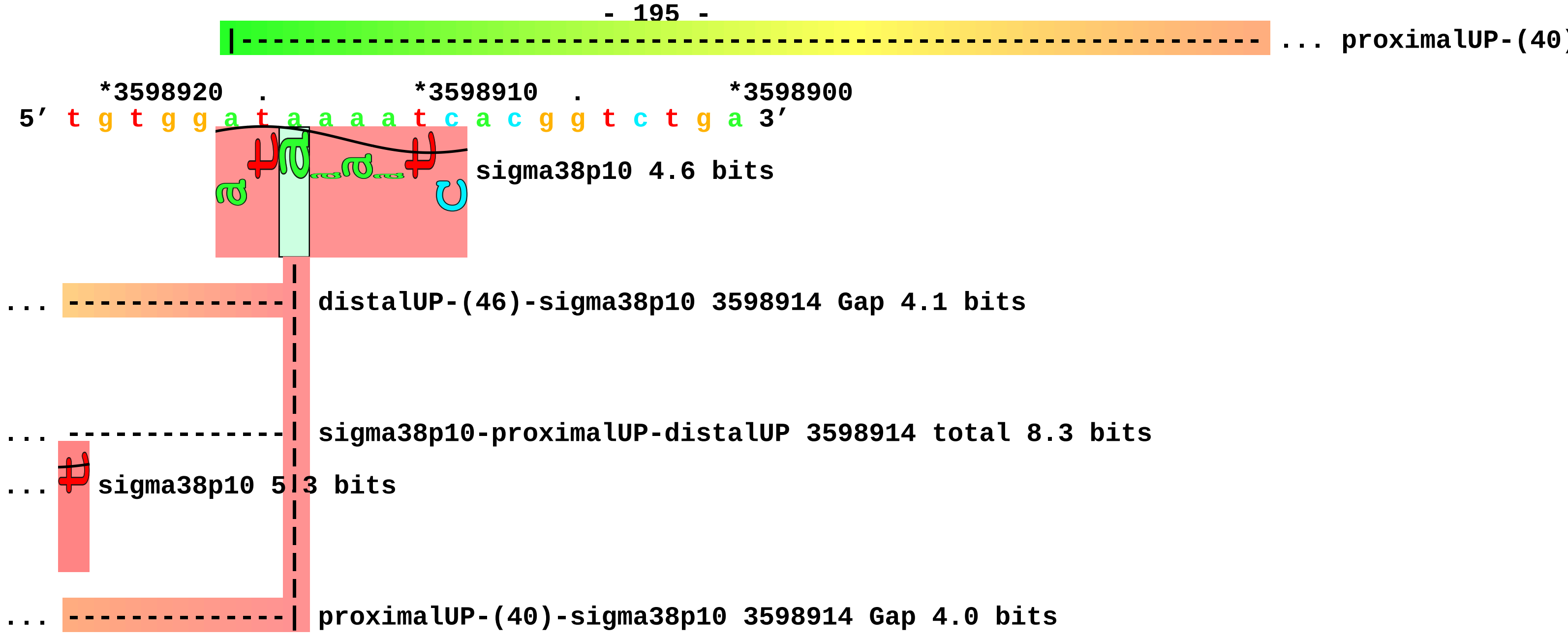
... proximalUP-(40)
*3598920 . *3598910 . *3598900
5′ t g t g g a t a a a a t c a c g g t c t g a 3′
sigma38p10 4.6 bits
... distalUP-(46)-sigma38p10 3598914 Gap 4.1 bits
... sigma38p10-proximalUP-distalUP 3598914 total 8.3 bits
... sigma38p10 5.3 bits
... proximalUP-(40)-sigma38p10 3598914 Gap 4.0 bits

piece 47, NC_000913, uspB, config: linear, direction: -, begin: 3637999, end: 3637700

. *3637990 . *3637980 . *3637970 .
5′ c c t t a t a g c g a c c c g g a t g a t c c g t c a a t c c g c c t t g c 3′

distalUP 3.5 bits
proximalUP 0.8 bits
... distalUP

T timestamp: 2018Sep20 11-34-53
... proximalUP-(34)
... distalUP-(54)-s
... sigma38p10-prox
... distalUP-(50)-s
... sigma38p10-prox

*3637960 . *3637950 . *3637940 . *3637930 .
5′ t t a c t c g t t a g a t a a a c a a a t t t t c g a c a t g a a t a t a t 3′

... distalUP 1.5 bits
sigma38p10 4.1 bits
... proximalUP

... proximalUP-(34)-sigma38p10 3637944 Gap 2.5 bits
... distalUP-(54)-sigma38p10 3637944 Gap 4.9 bits
... sigma38p10-proximalUP-distalUP 3637944 total 1.0 bits
... ... distalUP-(50)-s
... ... sigma38p10-prox

proximalUP 5.4 bits
distalUP 7.2 bits

... proximalUP-(33)
... distalUP-(43)-s
... sigma38p10-prox
... distalUP-(51)-s
... sigma38p10-prox
... distalUP-(53)-s
... sigma38p10-prox
... distalUP-(56)-s
... sigma38p10-prox

*3637920 . *3637910 . *3637900 . *3637890
5′ t g a t a g t g g t t a a c c t t c t g g a a a a a a a a c a a c c t g a t 3′

... proximalUP 6.1 bits
... sigma38p10

sigma38p10 5.7 bits
... proximalUP-(37)

proximalUP-(34)-sigma38p
... proximalUP-(42)
... distalUP-(50)-sigma38p10 3637912 Gap 4.1 bits
... sigma38p10-proximalUP-distalUP 3637912 total 4.4 bits

proximalUP 2.3 bits
proximalUP 4.3 bits

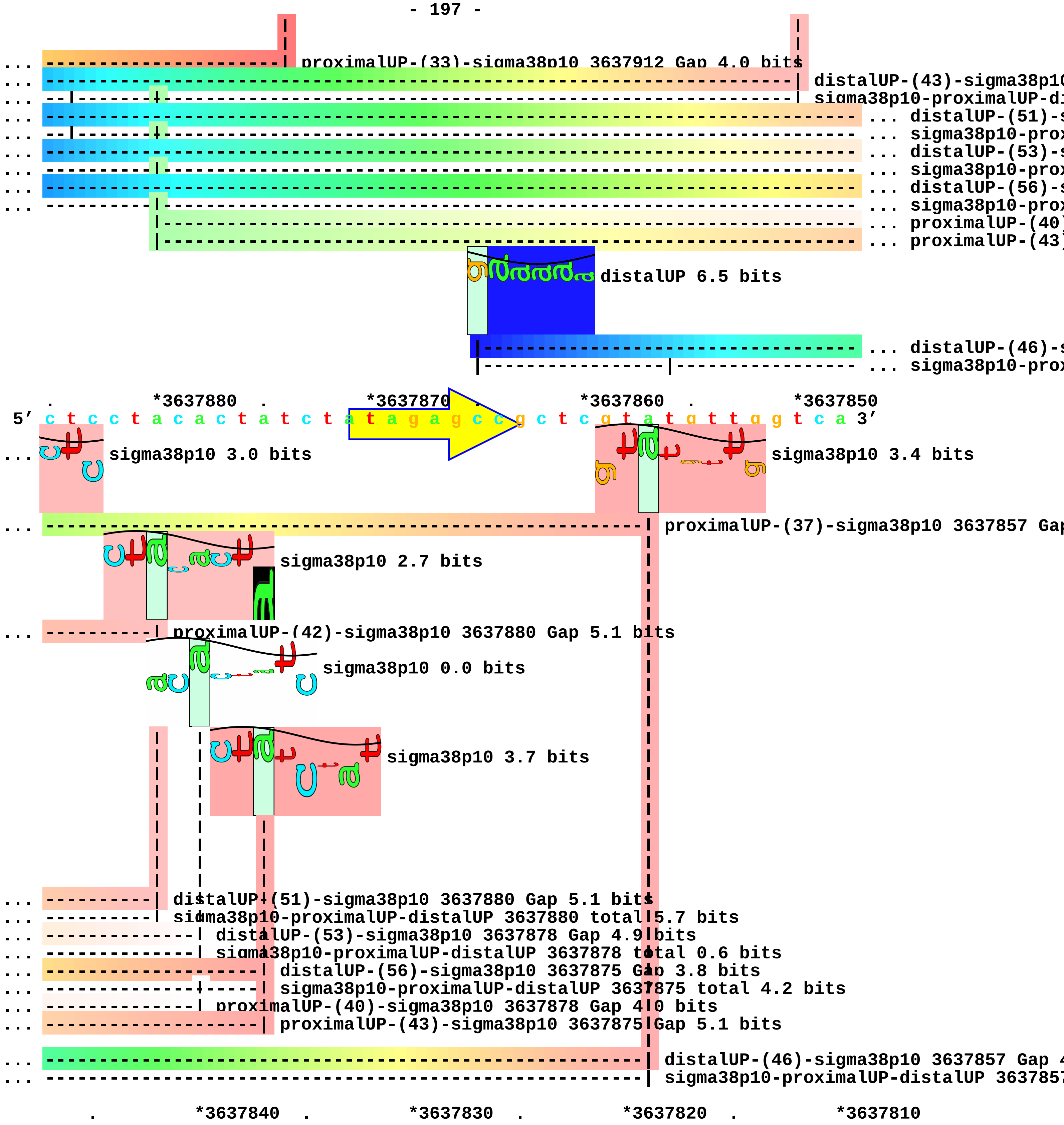
... proximalUP-(33)-sigma38p10 3637912 Gap 4.0 bits
... distalUP-(43)-sigma38p10
... sigma38p10-proximalUP-di
... distalUP-(51)-s
... sigma38p10-prox
... distalUP-(53)-s
... sigma38p10-prox
... distalUP-(56)-s
... sigma38p10-prox
... proximalUP-(40)
... proximalUP-(43)
distalUP 6.5 bits
... distalUP-(46)-s
... sigma38p10-prox
. *3637880 . *3637870 . *3637860 . *3637850
5′ c t c c t a c a c t a t c t a t a g a g c c g c t c g t a t g t t g g t c a 3′
sigma38p10 3.0 bits
sigma38p10 3.4 bits
... proximalUP-(37)-sigma38p10 3637857 Gap
sigma38p10 2.7 bits
... proximalUP-(42)-sigma38p10 3637880 Gap 5.1 bits
sigma38p10 0.0 bits
sigma38p10 3.7 bits
... distalUP-(51)-sigma38p10 3637880 Gap 5.1 bits
... sigma38p10-proximalUP-distalUP 3637880 total 5.7 bits
... distalUP-(53)-sigma38p10 3637878 Gap 4.9 bits
... sigma38p10-proximalUP-distalUP 3637878 total 0.6 bits
... distalUP-(56)-sigma38p10 3637875 Gap 3.8 bits
... sigma38p10-proximalUP-distalUP 3637875 total 4.2 bits
... proximalUP-(40)-sigma38p10 3637878 Gap 4.0 bits
... proximalUP-(43)-sigma38p10 3637875 Gap 5.1 bits
... distalUP-(46)-sigma38p10 3637857 Gap 4
... sigma38p10-proximalUP-distalUP 363785
. *3637840 . *3637830 . *3637820 . *3637810

5’ c c a c a a a a g c g g t t t c t c c a g g a c c t g g t g t t a a c t g a 3’

. *3637800 . *3637790 . *3637780 .

5’ t t g g c a g c g t a t c g g g a g c g g t a g c c c c g g g g t g g a a c 3’

*3637770 . *3637760 . *3637750 . *3637740 .

5’ t c c g c g g g c a g g t c g c c g g g g a g g a g g t a t g a t a a g c a 3’

*3637730 . *3637720 . *3637710 . *3637700

5’ c c g t c g c a t t a t t t t g g g c t t t a t g t g t c g t t t g 3’ piece 48, NC_000913, proK, config: linear, direction: -, begin: 3707049, end: 3706700

```
           .         *3707040   .         *3707030   .         *3707020   .
5' c c g c c g g a g c a g t t c c g c g t a c c g a t g a t g g t c t g g a t 3'
T timestamp:_2018Sep20_11-34-53

  *3707010   .         *3707000   .         *3706990   .         *3706980   .
5' g t c a g a t a a a t a t c t g g a a a a t c c g g c c a a t g c g c a g g 3'

      *3706970   .         *3706960   .         *3706950   .         *3706940
5' c g t t t g c g c a g c t g a a a a a a g a a g c c g a c a t g a a a g t g 3'
                                distalUP 6.5 bits               proximalUP 2.8 bits
                                                                  |----- ... proximalUP-(33)
            |---------------------------------------------------------- ... distalUP-(55)-s
            |------------------------------------------|----- ... sigma38p10-prox

 .         *3706930   .         *3706920   .         *3706910   .         *3706900
5' c c a c g c c g t c a c g t a g a g c t g t a c g a t a c c a t c a t g g g 3'
                                                            sigma38p10 2.0 bits
... ------------------------------------------------------------| proximalUP-(33)-sigma38p10 3706905
... ------------------------------------------------------------| distalUP-(55)-sigma38p10 3706905 0
... ------------------------------------------------------------| sigma38p10-proximalUP-distalUP 370

    .         *3706890   .         *3706880   .         *3706870   .         *3706860
5' t t g t c t t g g c t a t a c t t c a c c g g a t g g t g g a a t t a a c g 3'

        .         *3706850   .         *3706840   .         *3706830   .
5' a a a a c a a c a a c t g g t g t c a c a t c c c g c a g g c a a a a g a g 3'

  *3706820   .         *3706810   .         *3706800   .         *3706790   .
5' g c a g c g g c t a a c t a a g c g g c c t g c t g a c t t t c t c g c c g 3'

      *3706780   .         *3706770   .         *3706760   .         *3706750
5' a t c a a a a g g c a t t t t g c t a t t a a g g g a t t g a c g a g g g c 3'
                              proximalUP 6.1 bits
                 |-------------------------------------------- ... proximalUP-(36)
                           distalUP 5.7 bits
         |------------------------------------------------------ ... distalUP-(42)-s
         |-----------|------------------------------------------ ... sigma38p10-prox

 .         *3706740   .         *3706730   .         *3706720   .         *3706710
5' g t a t c t g c g c a g t a a g a t g c g c c c c g c a t t c g g t g a t t 3'
                                      sigma38p10 3.3 bits
```

```
... -------------------------| proximalUP-(36)-sigma38p10 3706732 Gap 3.6 bits
                             |
... -------------------------| distalUP-(42)-sigma38p10 3706732 Gap 3.3 bits
... -------------------------| sigma38p10-proximalUP-distalUP 3706732 total 8.3 bits

        .         *3706700
 5' g g c g c a g c 3'
```

piece 49, NC_000913, yjdC, config: linear, direction: -, begin: 4361482, end: 4361237

```
      *4361480   .         *4361470   .         *4361460   .         *4361450   .
5' c c t a c c g a c a a t t c t c t t t t t t g a c g g a c a a g g c c a g g 3'
 T timestamp:_2018Sep20_11-34-53

          *4361440   .         *4361430   .         *4361420   .         *4361410
5' a g c a t c c a c a a g c a c g c g t c a c g g g c t t t a t g g a t g c t 3'
                                                                          ... proximalUP
                                                      distalUP 4.7 bits
                                                      |------------------------- ... distalUP-(55)-s
                                                      |------------------------- ... sigma38p10-pro

    .         *4361400   .         *4361390   .         *4361380   .         *4361370
5' g a a a c c t t c a g c g c a c a t t t g c g c g a t c g c c a a c c g t g 3'
... proximalUP 3.3 bits
      |---------------------------------------------------------- ... proximalUP-(38)
... ---------------------------------------------------------------- ... distalUP-(55)-s
... --------|---------------------------------------------------------- ... sigma38p10-pro

      .         *4361360   .         *4361350   .         *4361340   .
5' a a c g a c a c t t g c a g t g g g a a a g a c g g a g g a a a a t a g c g 3'
          sigma38p10 3.5 bits                         proximalUP 3.3 bits
... --------| proximalUP-(38)-sigma38p10 4361364 Gap 3.2 bits       |----- ... proximalUP-(34)
... --------| distalUP-(55)-sigma38p10 4361364 Gap 4.9 bits   distalUP 7.2 bits

... --------| sigma38p10-proximalUP-distalUP 4361364 total 3.5 bits |----- ... proximalUP-(43)
                                                 |-------------------- ... distalUP-(41)-s
                                                 |------------|----- ... sigma38p10-pro
                                                 |-------------------- ... distalUP-(50)-s
                                                 |------------|----- ... sigma38p10-pro

   *4361330   .         *4361320   .         *4361310   .         *4361300   .
5' t g c a a c g t g a a g a t g t a c t g g g a g a a g c c c t g a a a t t a 3'
                                                          sigma38p10 6.4 bits
... ------------------------------------------------------| proximalUP-(34)-sigma38p10 43612
... ---------------------------------------------------------------- ... proximalUP-(43)
```

```
... ----------------------------------------------------------------| distalUP-(41)-sigma38p10 4361299
... ----------------------------------------------------------------| sigma38p10-proximalUP-distalUP 4
... ------------------------------------------------------------------------------------ ... distalUP-(50)-s
... ------------------------------------------------------------------------------------ ... sigma38p10-prox

        *4361290  .         *4361280  .         *4361270  .         *4361260  .
5' t t a g a a t t a c a a g g g a t t g c c a a c a c c a c g c t g g a g a t 3'
                    sigma38p10 5.0 bits
                        proximalUP 3.7 bits
                         |------------------------------------------------------------ ... proximalUP-(36)
... ----| proximalUP-(43)-sigma38p10 4361290 Gap 5.1 bits
                      distalUP 5.9 bits
            |------------------------------------------------------------------------ ... distalUP-(42)-s
... ----| distalUP-(50)-sigma38p10 4361290 Gap 4.1 bits
... ----| sigma38p10-proximalUP-distalUP 4361290 total 6.1 bits
            |-----------|------------------------------------------------------------ ... sigma38p10-prox

            *4361250  .         *4361240
5' g g t t g c t g a a c g t g t g g a 3'
                          sigma38p10 0.7 bits

... --------------| proximalUP-(36)-sigma38p10 4361247 Gap 3.6 bits

... --------------| distalUP-(42)-sigma38p10 4361247 Gap 3.3 bits

... --------------| sigma38p10-proximalUP-distalUP 4361247 total 3.5 bits
```

piece 50, NC_000913, ytfJ, config: linear, direction: -, begin: 4437399, end: 4436950

. *4437390 . *4437380 . *4437370 .
5′ t g c g c t t g a c c g c a a a c t g g c a t c a c a c t t g c g g g a a a 3′
T timestamp:_2018Sep20_11-34-53

*4437360 . *4437350 . *4437340 . *4437330 .
5′ t t c g a t a a a t a g c a c a t a t g a t t a a a c t c a g a c c c a a 3′

proximalUP 2.7 bits
distalUP 6.5 bits
|----------------------------------- ... proximalUP-(34)
|---------------------------------------------------------------------- ... distalUP-(50)-s
|--------------------------------|---------------------------------- ... sigma38p10-prox
proximalUP 3.9 bits
|---------------------- ... proximalUP-(35)
distalUP 6.4 bits
|------------------------------------------ ... distalUP-(43)-s
|---------------|---------------------- ... sigma38p10-prox

*4437320 . *4437310 . *4437300 . *4437290
5′ g t g g t c g g a t c a c c t g c a t a t c a t a a g a a g g a a a c a c c 3′

sigma38p10 2.1 bits
... --------------------------------| proximalUP-(34)-sigma38p10 4437308 Gap 2.5 bits
... --------------------------------| distalUP-(50)-sigma38p10 4437308 Gap 4.1 bits
... --------------------------------| sigma38p10-proximalUP-distalUP 4437308 total 4.6 bits
sigma38p10 0.5 bits
... ------------------------------------------------| proximalUP-(35)-sigma38p10 4437299 Gap 3.1 bit
distalUP 3.3 bits
... ------------------------------------------------| distalUP-(43)-sigma38p10 4437299 Gap 3.6 bits
... ------------------------------------------------| sigma38p10-proximalUP-distalUP 4437299 total 4
|--------------- ... distalUP-(53)-s
|--------------- ... sigma38p10-prox

. *4437280 . *4437270 . *4437260 . *4437250

5′ a t g a c c c t a c g c a a g a t t c t g g c a c t c a c c t g c c t g c t 3′

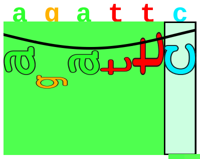
proximalUP 5.0 bits

|------------------------------------- ... proximalUP-(27)

... ---------------------------------------------------------------------------- ... distalUP-(53)-s
... ------------------------------------|-------------------------------------- ... sigma38p10-prox

. *4437240 . *4437230 . *4437220 . *4437210
5′ g t t g c c g a t g a t g g c t t c c g c a c a t c a g t t c g a a a c c g 3′

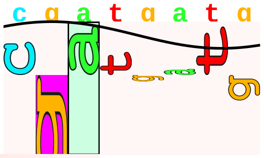
sigma38p10 0.4 bits

... --------------| proximalUP-(27)-sigma38p10 4437240 Gap 3.8 bits

... --------------| distalUP-(53)-sigma38p10 4437240 Gap 4.9 bits
... --------------| sigma38p10-proximalUP-distalUP 4437240 total 0.0 bits

. *4437200 . *4437190 . *4437180 .
5′ g t c a g c g a g t g c c g c c g a t t g g c a t t a c c g a t c g g g g c 3′

*4437170 . *4437160 . *4437150 . *4437140 .
5′ g a g t t g g t g c t t g a t a a a g a t c a g t t t a g c t a c a a a a c 3′

*4437130 . *4437120 . *4437110 . *4437100
5′ c t g g a a c a g c g c g c a g t t a g t g g g a a a a g t g c g a g t a c 3′

. *4437090 . *4437080 . *4437070 . *4437060
5′ t g c a a c a t a t t g c t g g t c g c a c c t c t g c a a a a g a g a a a 3′

. *4437050 . *4437040 . *4437030 . *4437020
5′ a a c g c g a c g c t g a t t g a a g c g a t t a a a t c a g c g a a g t t 3′

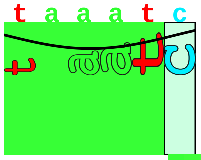
proximalUP 5.7 bits

|-------------------- ... proximalUP-(35)

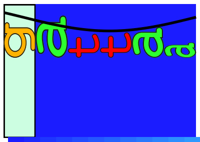
distalUP 6.4 bits

|------------------------------------ ... distalUP-(43)-s
|---------------|-------------------- ... sigma38p10-prox

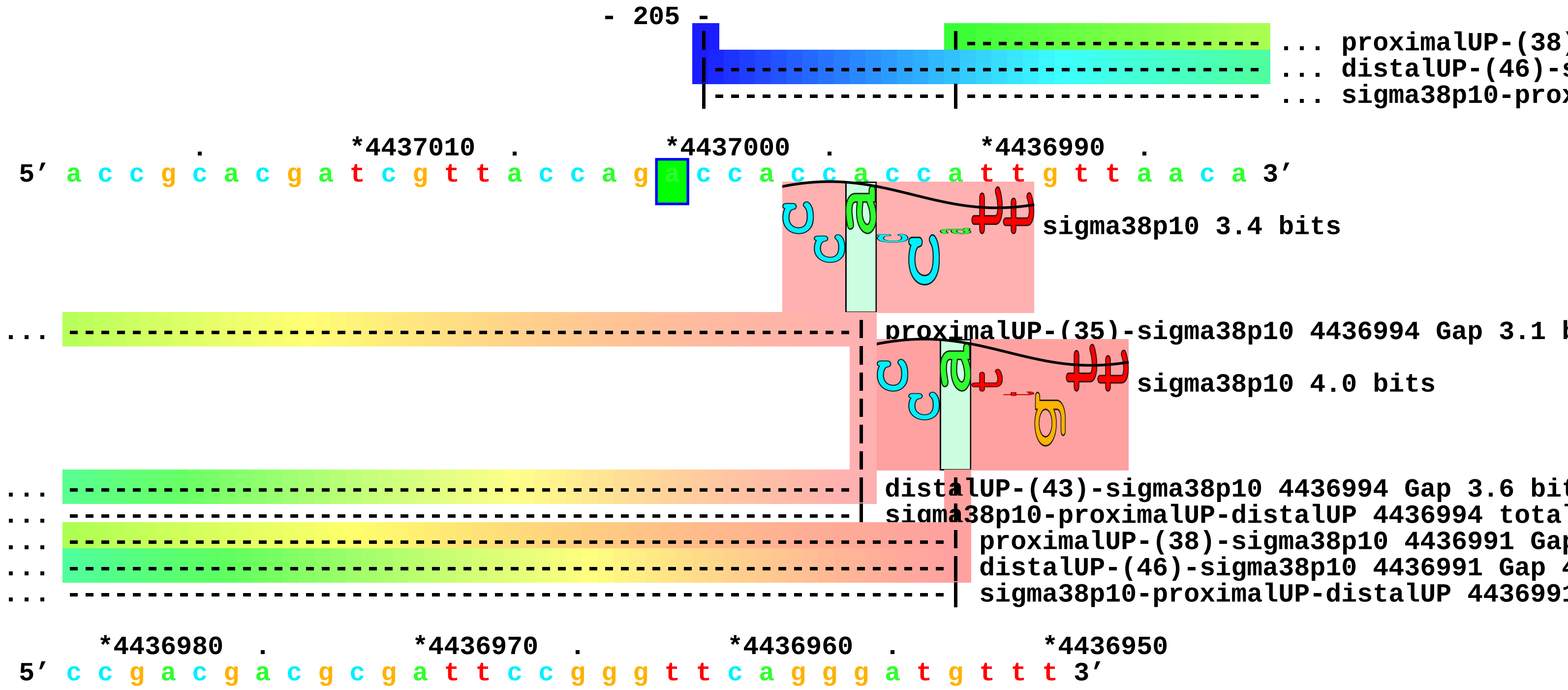
- 205 -
... proximalUP-(38)
... distalUP-(46)-s
... sigma38p10-prox
. *4437010 . *4437000 . *4436990 .
5′ a c c g c a c g a t c g t t a c c a g a c c a c c a c c a t t g t t a a c a 3′
sigma38p10 3.4 bits
... proximalUP-(35)-sigma38p10 4436994 Gap 3.1 b
sigma38p10 4.0 bits
... distalUP-(43)-sigma38p10 4436994 Gap 3.6 bit
... sigma38p10-proximalUP-distalUP 4436994 total
... proximalUP-(38)-sigma38p10 4436991 Gap
... distalUP-(46)-sigma38p10 4436991 Gap 4
... sigma38p10-proximalUP-distalUP 443699
*4436980 . *4436970 . *4436960 . *4436950
5′ c c g a c g a c g c g a t t c c g g g t t c a g g g a t g t t t 3′
