## Supplement_for_sigma38_paper-directory for "*Escherichia coli σ*^38^ promoters use two UP elements instead of a −35 element: resolution of a paradox and discovery that *σ*^38^ transcribes ribosomal promoters": Peano.Landini2015-Table2-neg-map.pdf

piece 1, NC_000913, ychH-pth, config: linear, direction: -, begin: 1258249, end: 1257700

. *1258240 . *1258230 . *1258220 .
5′ t a g t g g c g a a c c c a c c a g t a a c g g t c g c a c a c c t g t t c 3′
T timestamp:_2018Sep20_11-36-13

*1258210 . *1258200 . *1258190 . *1258180 .
5′ a t g c c c g c c a a c a c g g g c a c c c g c c a g c c a g a g a a t g g 3′

*1258170 . *1258160 . *1258150 . *1258140
5′ c a c c g a c g a a a a t a c t t a g c a c t g c a c c a t g t g c g a a a 3′

proximalUP 3.7 bits
... proximalUP-(38)
distalUP 7.2 bits
... distalUP-(45)-s
... sigma38p10-prox

. *1258130 . *1258120 . *1258110 . *1258100
5′ t a c t g g g g c a t a t t a a a c t g t g g t a a c t g g t t g a g g a t 3′

sigma38p10 6.1 bits
... proximalUP-(38)-sigma38p10 1258121 Gap 3.2 bits
proximalUP 5.0 bits
... distalUP-(45)-sigma38p10 1258121 Gap 5.5 bits
... sigma38p10-proximalUP-distalUP 1258121 total 8.3 bits
... proximalUP-(40)
distalUP 5.7 bits
... distalUP-(49)-s
... sigma38p10-prox

. *1258090 . *1258080 . *1258070 . *1258060
5′ t g a a t a c c c c a c g c c g a c c a c c a t t a c c a c c a g a c c c a 3′

sigma38p10 3.4 bits
... proximalUP-(40)-sigma38p10 1258078 Gap 4.0 bits
... distalUP-(49)-sigma38p10 1258078 Gap 5.9 bits

... ------------------------------------| sigma38p10-proximalUP-distalUP 1258078 total 4.2 bits

. *1258050 . *1258040 . *1258030 .
5’ a c c c c a t g a g c a c g t t a c c g a g t a a c g a a g c g t t t t t g 3’

*1258020 . *1258010 . *1258000 . *1257990 .
5’ c g t t t c a t a t g t c a c c t c c g g a a c t t t c t g g g t t g t a a 3’

*1257980 . *1257970 . *1257960 . *1257950
5’ c a g g g a a t a c c c c t c t t c c t t a t g t g t a a a g t a t a g a c 3’

. *1257940 . *1257930 . *1257920 . *1257910
5’ a a c a c c c a g t g g g t t a t g t g c g g g c g t g a t c a c a a t t a 3’

... proximalUP

distalUP 5.1 bits

|----------- ... distalUP-(44)-s
|----------- ... sigma38p10-prox

. *1257900 . *1257890 . *1257880 . *1257870
5’ c a a c c c t t a t t t c a a c a a a a c t t t a c a a a t a a a c g c c t 3’

... proximalUP 3.7 bits ... sigma38p10

|----------------------------------------------------------------------- ... proximalUP-(38)
... ------------------------------------------------------------------------ ... distalUP-(44)-s
... |----------------------------------------------------------------------- ... sigma38p10-prox

proximalUP 5.1 bits

|------------------------------------------------------ ... proximalUP-(35)

distalUP 4.1 bits

|------------------------------------------------------------------ ... distalUP-(42)-s
|------------- ---------------------------------------------------- ... sigma38p10-prox
|---------------------------------------------------- ... proximalUP-(43)
|------------------------------------------------------------------ ... distalUP-(50)-s
|-------------|---------------------------------------------------- ... sigma38p10-prox

. *1257860 . *1257850 . *1257840 .
5’ g a a c a c c t t g a c t t t c t a a t a g t c c a c t c a g g c a a t t a 3’

... sigma38p10 0.9 bits

... | proximalUP-(38)-sigma38p10 1257869 Gap 3.2 bits
... | distalUP-(44)-sigma38p10 1257869 Gap 2.8 bits

```
... | sigma38p10-proximalUP-distalUP 1257869 total 3.7 bits
                                                sigma38p10 1.2 bits
... ------------------| proximalUP-(35)-sigma38p10 1257860 Gap 3.1 bits
                                                sigma38p10 2.4 bits
... ------------------| distalUP-(42)-sigma38p10 1257860 Gap 3.3 bits
... ------------------| sigma38p10-proximalUP-distalUP 1257860 total 4.1 bits
... ----------------------------------| proximalUP-(43)-sigma38p10 1257852 Gap 5.1 bits
... ----------------------------------| distalUP-(50)-sigma38p10 1257852 Gap 4.1 bits
... ----------------------------------| sigma38p10-proximalUP-distalUP 1257852 total 2.3 bits

     *1257830  .          *1257820  .          *1257810  .          *1257800  .
5' t t a c g t t a t c t g c a t t t g a t a c g c a g t t t t t t g t c a a g 3'
                                                          proximalUP 5.7 bits
                                                          |----------- ... proximalUP-(38)
                                                          distalUP 4.9 bits
                                                        |-----------------------  ... distalUP-(45)-s
                                                        |---------|-----------  ... sigma38p10-prox
                                                                  |-----------  ... proximalUP-(42)
                                                        |-----------------------  ... distalUP-(49)-s
                                                        |---------|-----------  ... sigma38p10-prox

         *1257790  .          *1257780  .          *1257770  .          *1257760
5' c g g g g c c g c a a c c a g t a a a c t a c g c g c c a g t t a t g t a c 3'
                                                                  sigma38p10 0.9 bits
... ----------------------------------------------------------------| proximalUP-(38)-sigma38p10 125
                                                                   ... sigma38p10
... ----------------------------------------------------------------| distalUP-(45)-sigma38p10 12577
... ----------------------------------------------------------------| sigma38p10-proximalUP-distalUP
... -----------------------------------------------------------------------| proximalUP-(42)-sigma3
... -----------------------------------------------------------------------| distalUP-(49)-sigma38p
... -----------------------------------------------------------------------| sigma38p10-proximalUP-

   .         *1257750  .          *1257740  .          *1257730  .          *1257720
5' a c a c t c a g g a c a a a a a a a c g t g a c g a t t a a a t t g a t t g 3'

... sigma38p10 1.0 bits
```

```
          .         *1257710  .         *1257700
5′ t c g g c c t g g c g a a c c c c g 3′
```

piece 2, NC_000913, rssA-ychJ, config: linear, direction: -, begin: 1288449, end: 1288200

. *1288440 . *1288430 . *1288420 .
5′ c t g g a a t g c a a t c g t a g c c a c a t c g c t a c t g c t c g g t t 3′
T timestamp:_2018Sep20_11-36-13

*1288410 . *1288400 . *1288390 . *1288380 .
5′ a a c a t a g t g c t a c t c a g g c g g t t a a t a c g c c g t t a t t g 3′
proximalUP 3.3 bits
... proximalUP-(35)
distalUP 5.5 bits
... distalUP-(42)-s
... sigma38p10-prox

*1288370 . *1288360 . *1288350 . *1288340
5′ t t t c c a g g g a g a t t a t t t g t g t c t c a g c t t t g t c c c t g 3′
sigma38p10 1.8 bits
... proximalUP-(35)-sigma38p10 1288349 Gap 3.1 bit
distalUP 7.1 bits
proximalUP 0.1 bits
... distalUP-(42)-sigma38p10 1288349 Gap 3.3 bits
... sigma38p10-proximalUP-distalUP 1288349 total 4
... proximalUP-(29)
... distalUP-(52)-s
... sigma38p10-prox

. *1288330 . *1288320 . *1288310 . *1288300
5′ t g g t a g t g c t g t c g a g t a t a g c c t a t g t t g c c a c c c t t 3′
sigma38p10 6.1 bits
... proximalUP-(29)-sigma38p10 1288311 Gap 7.5 bit
... distalUP-(52)-sigma38p10 1288311 Gap 5.5 bits
... sigma38p10-proximalUP-distalUP 1288311 total 6

. *1288290 . *1288280 . *1288270 . *1288260
5′ a t g t g t c t g g t g a a a a g g t t g c a c c t g a t c c a g a a c a t 3′

. *1288250 . *1288240 . *1288230 .
5′ c t c a t g c g t t c g c g t t a c t g c g c t t t t g t g a t g c a a g a 3′

```
    *1288220  .         *1288210  .         *1288200
5' c g c a g a t t a t t t a a t a a a g a c c 3'
```

piece 3, NC_000913, ydbJ-ydbK, config: linear, direction: -, begin: 1439099, end: 1438750

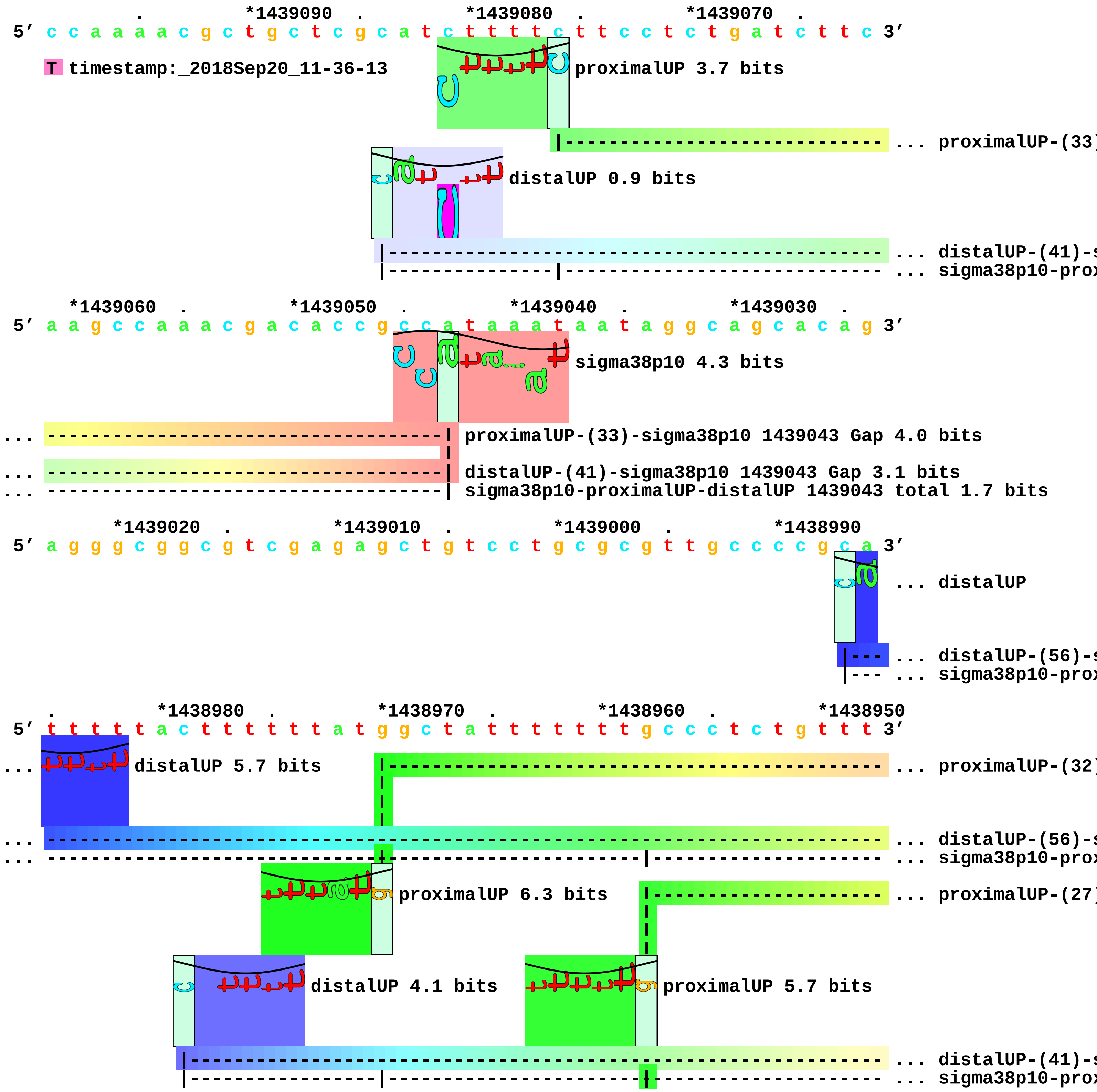

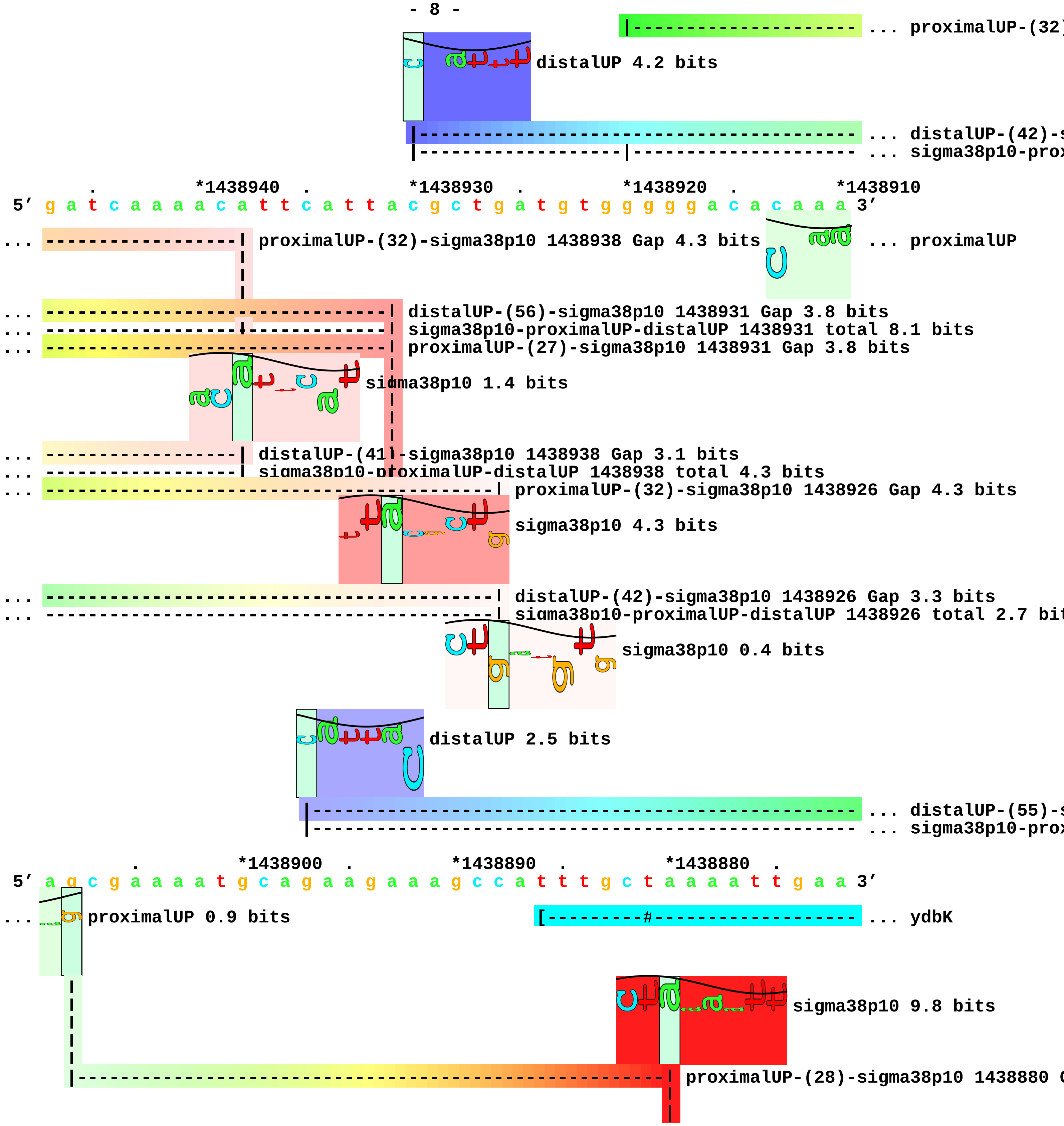
... proximalUP-(32)
distalUP 4.2 bits
... distalUP-(42)-s
... sigma38p10-prox
. *1438940 . *1438930 . *1438920 . *1438910
5′ g a t c a a a a c a t t c a t t a c g c t g a t g t g g g g g a c a c a a a 3′
... proximalUP-(32)-sigma38p10 1438938 Gap 4.3 bits
... proximalUP
... distalUP-(56)-sigma38p10 1438931 Gap 3.8 bits
... sigma38p10-proximalUP-distalUP 1438931 total 8.1 bits
... proximalUP-(27)-sigma38p10 1438931 Gap 3.8 bits
sigma38p10 1.4 bits
... distalUP-(41)-sigma38p10 1438938 Gap 3.1 bits
... sigma38p10-proximalUP-distalUP 1438938 total 4.3 bits
... proximalUP-(32)-sigma38p10 1438926 Gap 4.3 bits
sigma38p10 4.3 bits
... distalUP-(42)-sigma38p10 1438926 Gap 3.3 bits
... sigma38p10-proximalUP-distalUP 1438926 total 2.7 bit
sigma38p10 0.4 bits
distalUP 2.5 bits
... distalUP-(55)-s
... sigma38p10-prox
. *1438900 . *1438890 . *1438880 .
5′ a g c g a a a a t g c a g a a g a a a g c c a t t t g c t a a a a t t g a a 3′
... proximalUP 0.9 bits
[---------#------------------- ... ydbK
sigma38p10 9.8 bits
proximalUP-(28)-sigma38p10 1438880 G

```
... ----------------------------------------------------------| distalUP-(55)-sigma38p10 1438880 Gap
... --|-------------------------------------------------------| sigma38p10-proximalUP-distalUP 14388

     *1438870  .         *1438860  .         *1438850  .         *1438840  .
 5' a g a t t a c t a c t g g g c g c g c a g c a a t t t c g t g c g c c c c t 3'
... ----------------------> ydbK                            proximalUP 5.5 bits
                                                        |-------------------- ... proximalUP-(36)
                                                       distalUP 5.8 bits
                                                 |----------------------------- ... distalUP-(42)-s
                                                 |-----------|----------------- ... sigma38p10-prox
                                                             |----------------- ... proximalUP-(42)
                                                 |----------------------------- ... distalUP-(48)-s
                                                 |-----------|----------------- ... sigma38p10-prox

       *1438830  .         *1438820  .         *1438810  .         *1438800
 5' c a t t t g c g c a a t g t a a g g g t g t c a t a t g a t t a c t a t t g 3'
                                                             sigma38p10 6.9 bits
... ---------------------------------------------------------| proximalUP-(36)-sigma38p10 1438808 Gap 3.6 b
                                                                  sigma38p10 6.3 bits
... ---------------------------------------------------------| distalUP-(42)-sigma38p10 1438808 Gap 3.3 bit
... ---------------------------------------------------------| sigma38p10-proximalUP-distalUP 1438808 tota
... ---------------------------------------------------------------------| proximalUP-(42)-sigma38p10 14388
... ---------------------------------------------------------------------| distalUP-(48)-sigma38p10 143880
... ---------------------------------------------------------------------| sigma38p10-proximalUP-distalUP 1

    .        *1438790  .         *1438780  .         *1438770  .         *1438760
 5' a c g g t a a t g g c g c g g t t g c t t c g g t c g c a t t t c g c a c c 3'

      .        *1438750
 5' a g t g a a g t 3'
```

piece 4, NC_000913, cybB-gapC1, config: linear, direction: -, begin: 1488999, end: 1488600

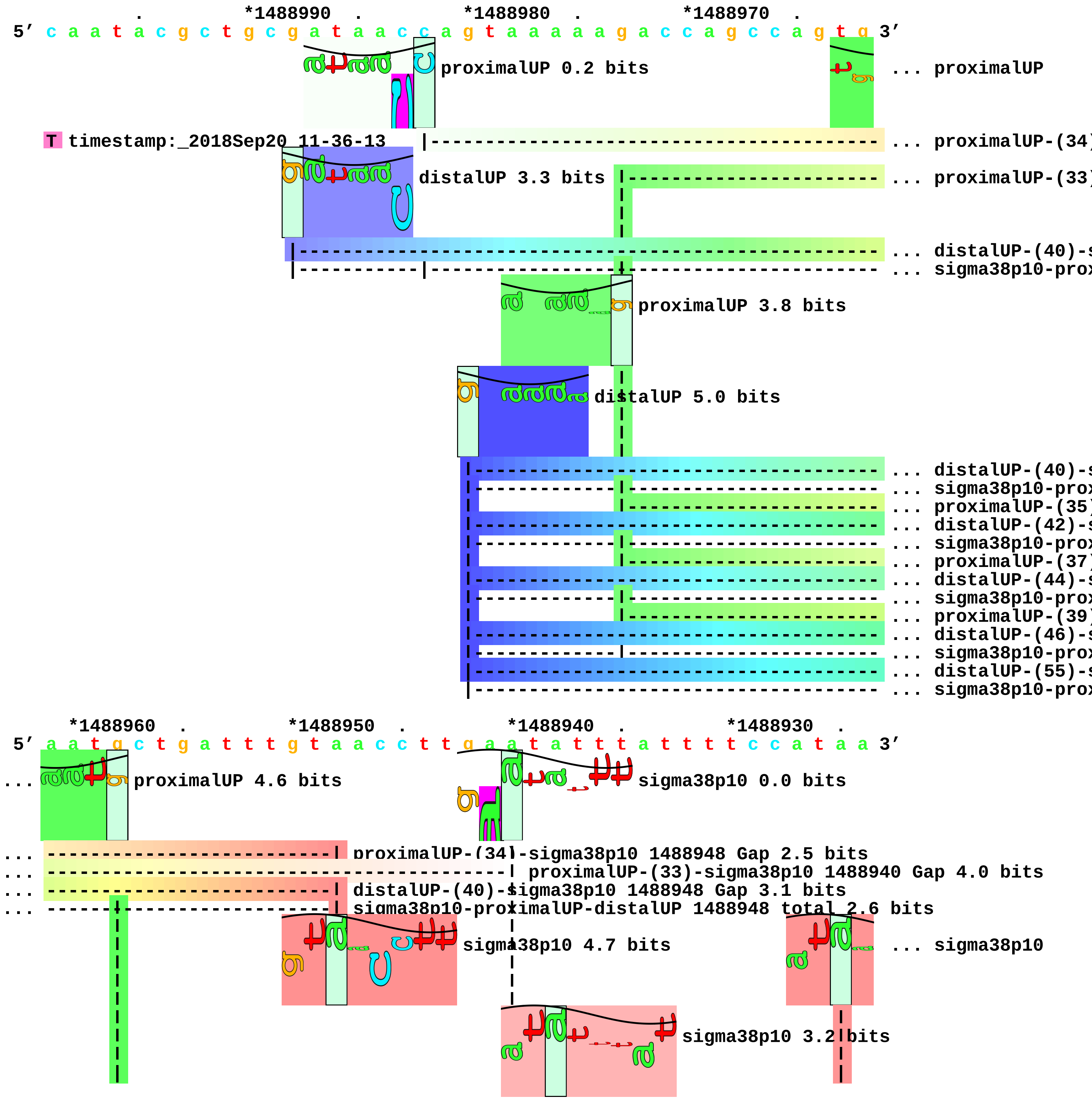

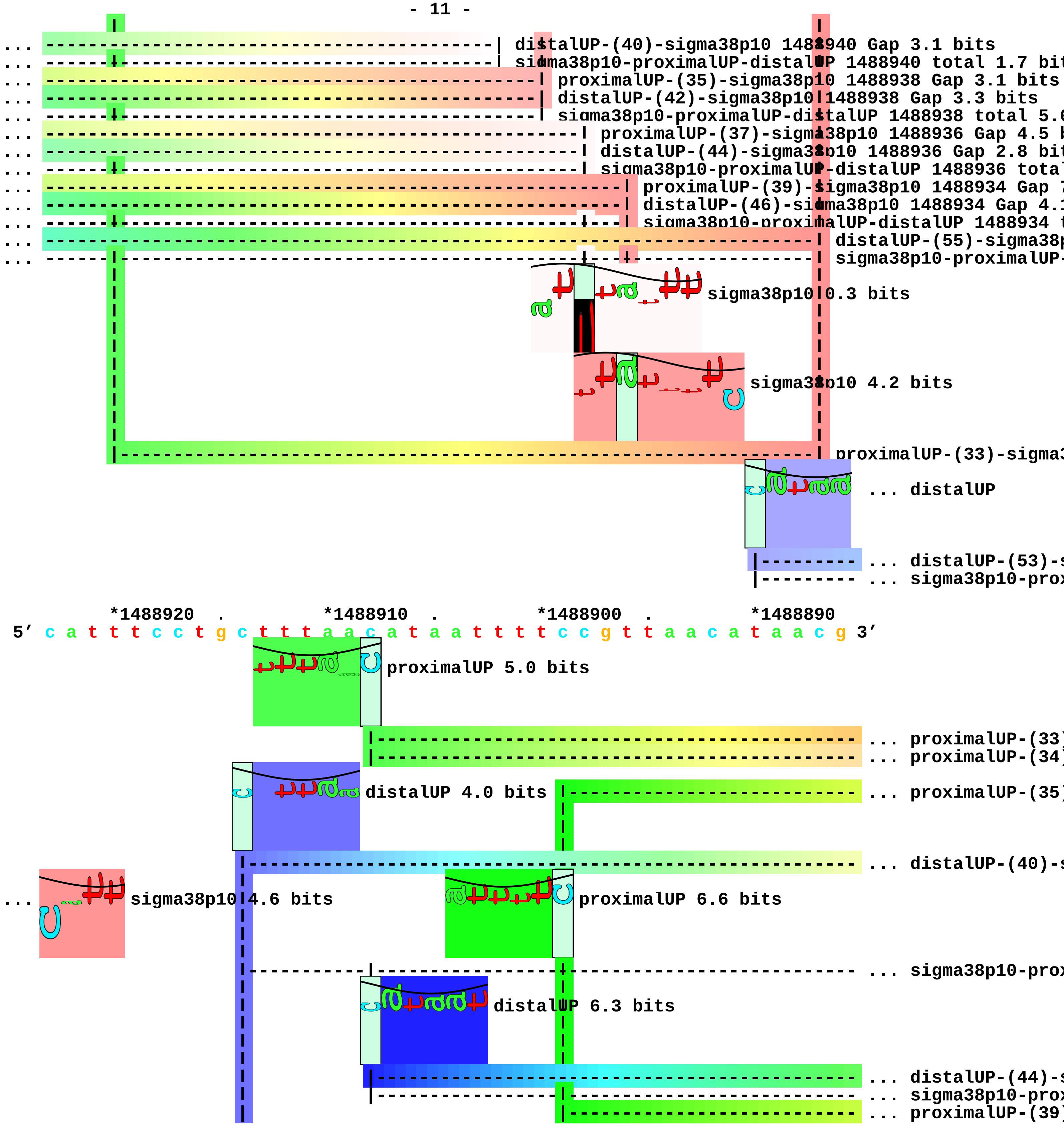

distalUP-(40)-sigma38p10 1488940 Gap 3.1 bits
sigma38p10-proximalUP-distalUP 1488940 total 1.7 bit
proximalUP-(35)-sigma38p10 1488938 Gap 3.1 bits
distalUP-(42)-sigma38p10 1488938 Gap 3.3 bits
sigma38p10-proximalUP-distalUP 1488938 total 5.
proximalUP-(37)-sigma38p10 1488936 Gap 4.5 b
distalUP-(44)-sigma38p10 1488936 Gap 2.8 bit
sigma38p10-proximalUP-distalUP 1488936 total
proximalUP-(39)-sigma38p10 1488934 Gap
distalUP-(46)-sigma38p10 1488934 Gap 4.
sigma38p10-proximalUP-distalUP 1488934
distalUP-(55)-sigma38p
sigma38p10-proximalUP-
sigma38p10 0.3 bits
sigma38p10 4.2 bits
proximalUP-(33)-sigma
... distalUP
... distalUP-(53)-
... sigma38p10-prox
*1488920 . *1488910 . *1488900 . *1488890
5′ c a t t t c c t g c t t t a a c a t a a t t t t c c g t t a a c a t a a c g 3′
proximalUP 5.0 bits
... proximalUP-(33)
... proximalUP-(34)
distalUP 4.0 bits
... proximalUP-(35)
... distalUP-(40)-
... sigma38p10 4.6 bits
proximalUP 6.6 bits
... sigma38p10-prox
distalUP 6.3 bits
... distalUP-(44)-
... sigma38p10-prox
... proximalUP-(39)

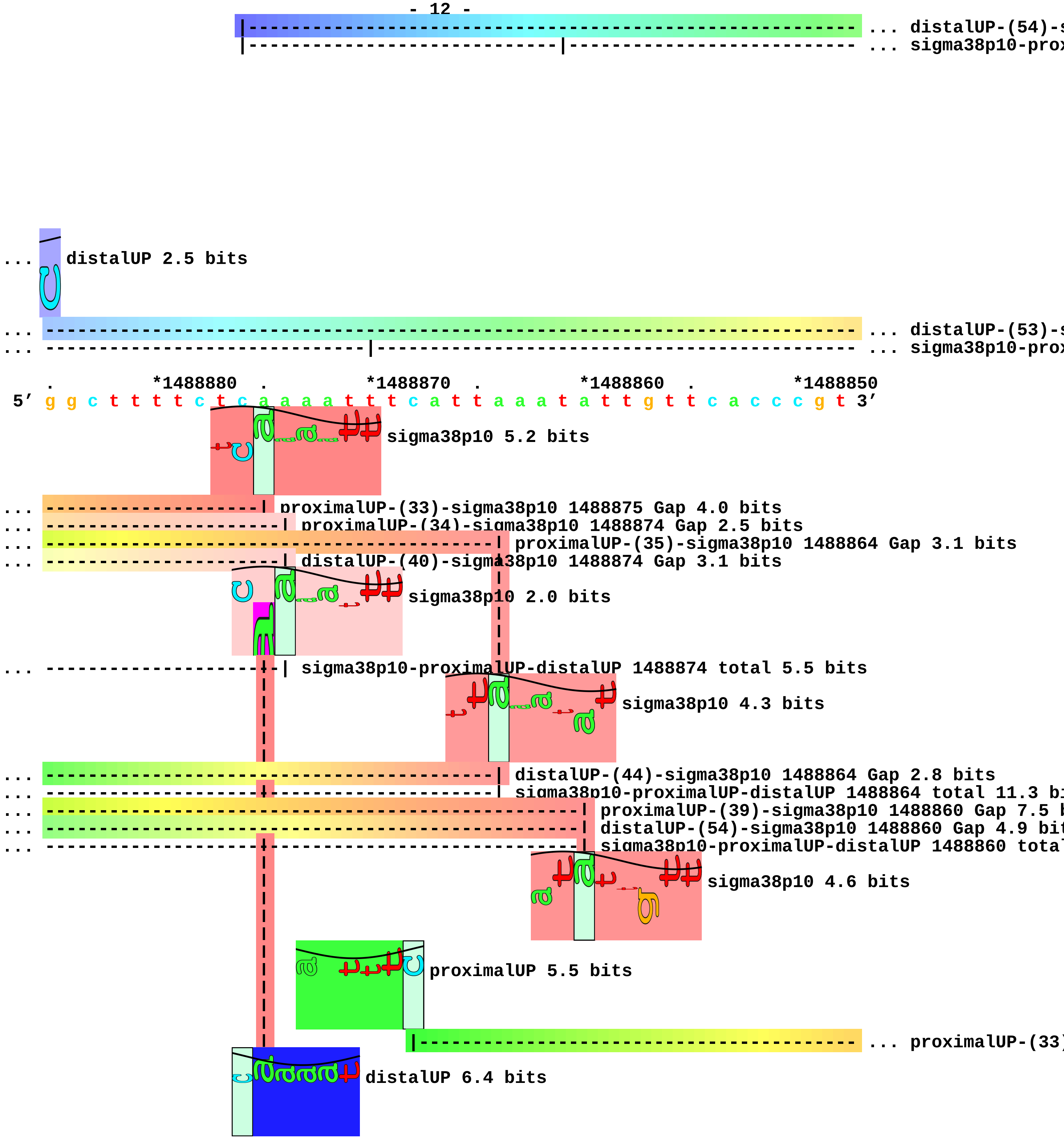
... distalUP-(54)-s
... sigma38p10-prox
... distalUP 2.5 bits
... distalUP-(53)-s
... sigma38p10-prox
*1488880
*1488870
*1488860
*1488850
5′ g g c t t t t c t c a a a a t t t c a t t a a a t a t t g t t c a c c c g t 3′
sigma38p10 5.2 bits
... proximalUP-(33)-sigma38p10 1488875 Gap 4.0 bits
... proximalUP-(34)-sigma38p10 1488874 Gap 2.5 bits
... proximalUP-(35)-sigma38p10 1488864 Gap 3.1 bits
... distalUP-(40)-sigma38p10 1488874 Gap 3.1 bits
sigma38p10 2.0 bits
... sigma38p10-proximalUP-distalUP 1488874 total 5.5 bits
sigma38p10 4.3 bits
... distalUP-(44)-sigma38p10 1488864 Gap 2.8 bits
... sigma38p10-proximalUP-distalUP 1488864 total 11.3 bi
... proximalUP-(39)-sigma38p10 1488860 Gap 7.5 b
... distalUP-(54)-sigma38p10 1488860 Gap 4.9 bit
... sigma38p10-proximalUP-distalUP 1488860 tota
sigma38p10 4.6 bits
proximalUP 5.5 bits
... proximalUP-(33)
distalUP 6.4 bits

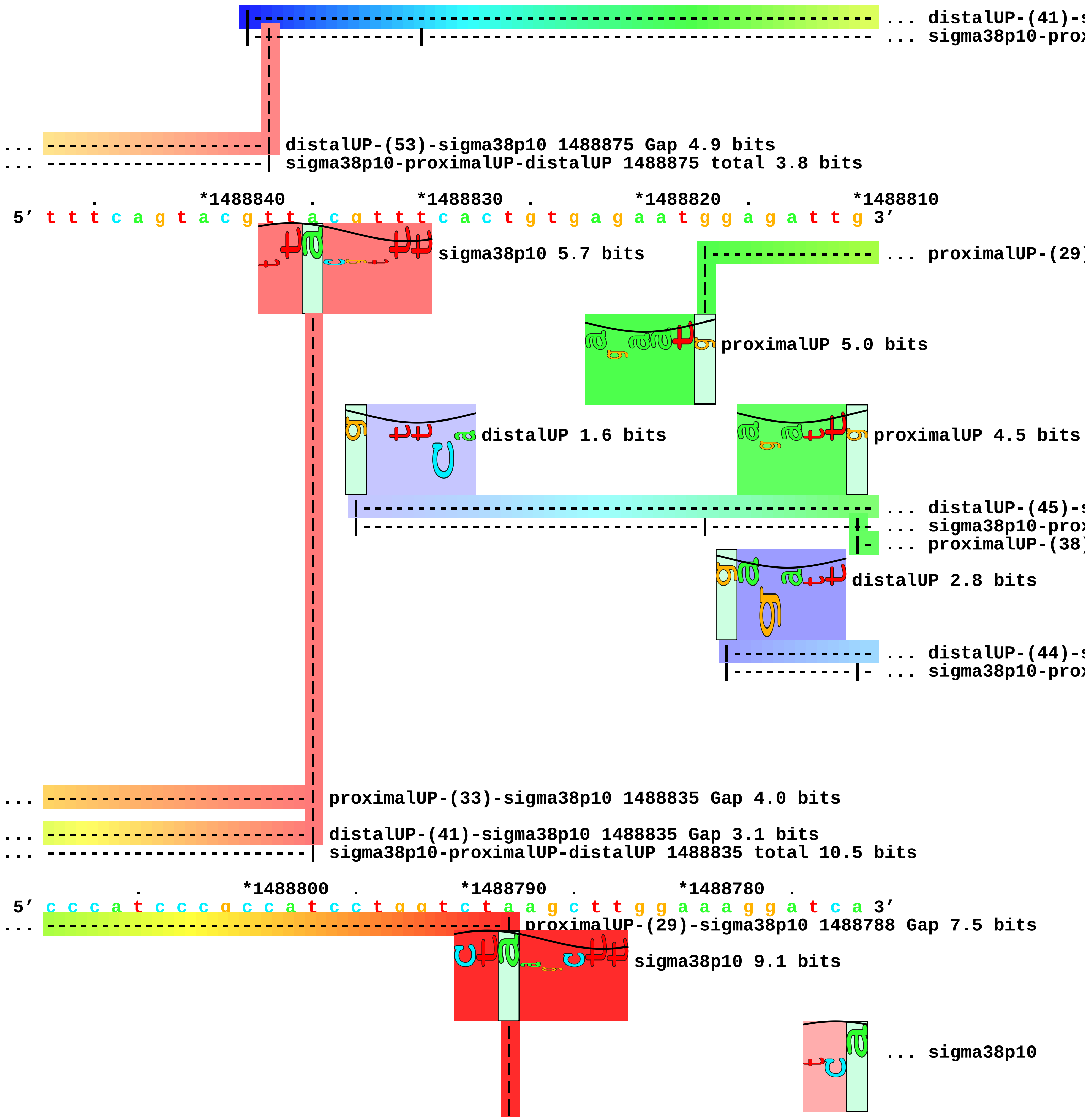
... distalUP-(41)-s
... sigma38p10-prox
... distalUP-(53)-sigma38p10 1488875 Gap 4.9 bits
... sigma38p10-proximalUP-distalUP 1488875 total 3.8 bits
. *1488840 . *1488830 . *1488820 . *1488810
5′ t t t c a g t a c g t t a c g t t t c a c t g t g a g a a t g g a g a t t g 3′
sigma38p10 5.7 bits
... proximalUP-(29)
proximalUP 5.0 bits
distalUP 1.6 bits
proximalUP 4.5 bits
... distalUP-(45)-s
... sigma38p10-prox
... proximalUP-(38)
distalUP 2.8 bits
... distalUP-(44)-s
... sigma38p10-prox
... proximalUP-(33)-sigma38p10 1488835 Gap 4.0 bits
... distalUP-(41)-sigma38p10 1488835 Gap 3.1 bits
... sigma38p10-proximalUP-distalUP 1488835 total 10.5 bits
. *1488800 . *1488790 . *1488780 .
5′ c c c a t c c c g c c a t c c t g g t c t a a g c t t g g a a a g g a t c a 3′
... proximalUP-(29)-sigma38p10 1488788 Gap 7.5 bits
sigma38p10 9.1 bits
... sigma38p10

... ----------------------------------------| distalUP-(45)-sigma38p10 1488788 Gap 5.5 bits
... ----------------------------------------| sigma38p10-proximalUP-distalUP|1488788 total 2.8 bit
... ----------------------------------------------------------------------------| proximalUP-(38)-sigm
... distalUP

... ----------------------------------------------------------------------------| distalUP-(44)-sigma3
... ----------------------------------------------------------------------------| sigma38p10-proximalU
|--- ... distalUP-(42)-s
|--- ... sigma38p10-prox

*1488770 . *1488760 . *1488750 . *1488740 .
5′ a t t t t c a t c c g a a c g t t c c t g a c a g g a g a a a a c c a t g a 3′

proximalUP 6.6 bits

|------------------------------------------------------------ ... proximalUP-(35)

... sigma38p10 3.5 bits

... distalUP 5.8 bits

... ---------------------------------------------------------------------------- ... distalUP-(42)-s
... ----------|----------------------------------------------------------------- ... sigma38p10-prox

*1488730 . *1488720 . *1488710 . *1488700
5′ g t a a a g t t g g t a t t a a c g g t t t t g g t c g t a t c g g t c g a 3′

sigma38p10 4.8 bits

... ----| proximalUP-(35)-sigma38p10 1488731 Gap 3.1 bits

... ----| distalUP-(42)-sigma38p10 1488731 Gap 3.3 bits
... ----| sigma38p10-proximalUP-distalUP 1488731 total 10.8 bits

. *1488690 . *1488680 . *1488670 . *1488660
5′ c t g g t g t t g g g t c g a t t a c t t g a a g t c a a a a g c a a c a t 3′

. *1488650 . *1488640 . *1488630 . *1488620
5′ a g a c g t t g t c g c t a t t a a t g a t c t c a c t t c c c c a t a a a 3′

```
               .          *1488610  .         *1488600
5' t t c t c g c c t a c c t g c t g a a a 3'
```

piece 5, NC_000913, nadE-osmE, config: linear, direction: -, begin: 1820399, end: 1820200

```
             .         *1820390  .         *1820380  .         *1820370  .
5' g g t g g t t a a a a c g t c a t a c a g g a a t c c t c t g a a a g c g g 3'
   T timestamp:_2018Sep20_11-36-13

   *1820360  .         *1820350  .         *1820340  .         *1820330  .
5' a g c c g t t t c g t t c a c g g g c t t g a a a a a g c g c c c a a t g t 3'

     *1820320  .         *1820310  .         *1820300  .         *1820290
5' a t t c c a g g c t t a t c t a a c a c g c t g a t a a a c a a g a g g a c 3'

   .         *1820280  .         *1820270  .         *1820260  .         *1820250
5' g g a a t a t g a a c a a g a a t a t g g c a g g a a t t c t g a g t g c a 3'
```

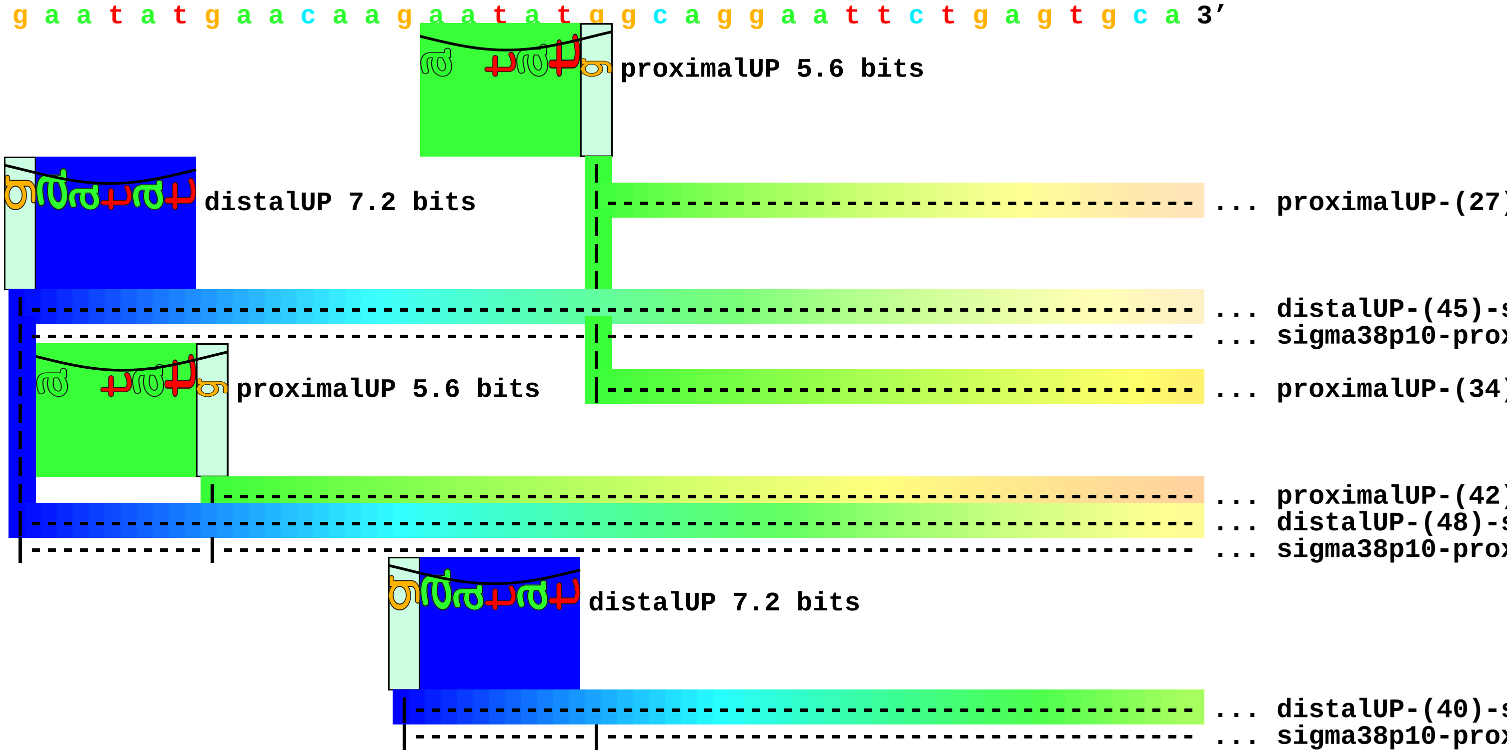


```
       .         *1820240  .         *1820230  .         *1820220  .         *1820210
5' g c g g c g g t a t t a a c c a t g c t g g c g g g t t g t a c g g c t t a 3'
```

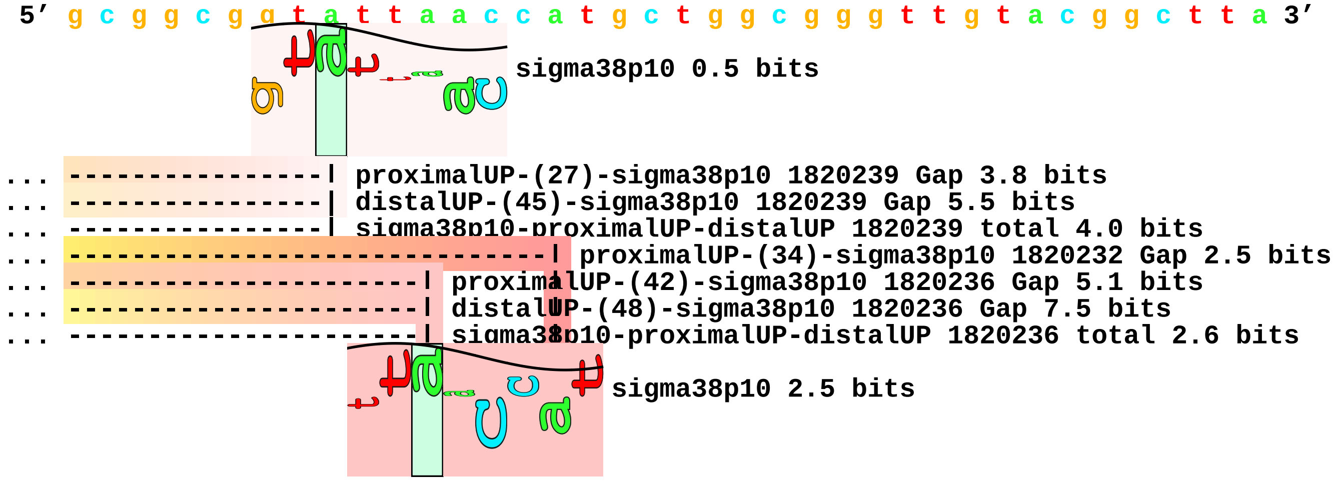

... ------------------------------| distalUP-(40)-sigma38p10 1820232 Gap 3.1 bits

... ------------------------------| sigma38p10-proximalUP-distalUP 1820232 total 11.5 bits

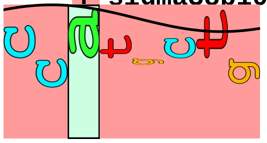

sigma38p10 4.3 bits

. *1820200

5′ t g a t c g t a c c 3′

piece 6, NC_000913, yodD-dsrB, config: linear, direction: -, begin: 2023199, end: 2022800

. *2023190 . *2023180 . *2023170 .

5′ c c a g t t c g a a g g c a t c c g c c a a t t c a c g c c a t g t a t g a 3′

T timestamp:_2018Sep20_11-36-13

... proximalUP

... distalUP

... distalUP-(46)-s
... sigma38p10-prox

*2023160 . *2023150 . *2023140 . *2023130 .

5′ t a t t c a c g t c c a t c g a c g c t a a c g t g t t c a g a a t c g t t 3′

... proximalUP 6.6 bits

... distalUP 6.5 bits

... ... distalUP-(46)-s
... ... sigma38p10-prox
... proximalUP-(40)

*2023120 . *2023110 . *2023100 . *2023090

5′ c t g g c t g c g a t g a a c t t c g c t t t c g c t a a t t t c a t t g a 3′

sigma38p10 2.0 bits

proximalUP 5.5 bits

... proximalUP-(37)
... proximalUP-(43)

... distalUP-(46)-sigma38p10 2023117 Gap 4.1 bits
... sigma38p10-proximalUP-distalUP 2023117 total 7.0 bits
... proximalUP-(40)-sigma38p10 2023117 Gap 4.0 bits

distalUP 4.2 bits

... distalUP-(44)-s
... sigma38p10-prox
... distalUP-(50)-s
... sigma38p10-prox

. *2023080 . *2023070 . *2023060 . *2023050

5′ t c g c c g c c a g c a g g g c a t c g a c a t c g a c g c t g a c c t c a 3′

sigma38p10 1.2 bits

... ----------------------------------------------------------------| proximalUP-(37)-sigma38p10 20230
... ----------------------------------------------------------------| proximalUP-(43)-sigm

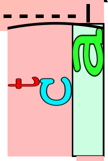

... sigma38p10

... ----------------------------------------------------------------| distalUP-(44)-sigma38p10 202305
... ----------------------------------------------------------------| sigma38p10-proximalUP-distalUP 2
... ----------------------------------------------------------------| distalUP-(50)-sigma3
... ----------------------------------------------------------------| sigma38p10-proximalU

. *2023040 . *2023030 . *2023020 . *2023010
5′ c g t t t t g c g g t a t c g c t g t a c t c t t t t g c g g t t t t c a t 3′

... 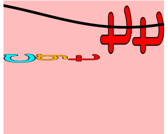 sigma38p10 2.9 bits

. *2023000 . *2022990 . *2022980 .
5′ c a t c t t t c c t c g c a a c c g t t a c t c t g t t g t t g c c g c c a 3′

*2022970 . *2022960 . *2022950 . *2022940 .
5′ g t c g t t a a g t c c g c c t g g c a t c t c t t a a g t a t a g a c c g 3′

*2022930 . *2022920 . *2022910 . *2022900
5′ t c g a g a a a a t c a g c a c t c g c g c g c g c c t g g c g c t g c c a 3′

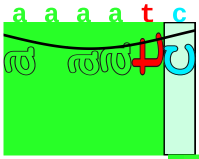 proximalUP 6.1 bits

------------------------------------------------------ ... proximalUP-(35)

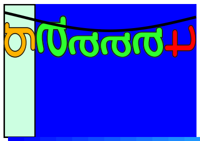 distalUP 7.2 bits

------------------------------------------------------ ... distalUP-(41)-s
------------------------------------------------------ ... sigma38p10-prox
------------------------------------------------------ ... proximalUP-(36)
------------------------------------------------------ ... distalUP-(42)-s
------------------------------------------------------ ... sigma38p10-prox

. *2022890 . *2022880 . *2022870 . *2022860
5′ g a a a a a c g a a a t t g t t c t a c a c t g g c a c a a a g c c a c a g 3′

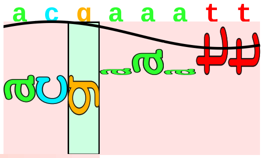 sigma38p10 1.3 bits

... -------------| proximalUP-(35)-sigma38p10 2022888 Gap 3.1 bits

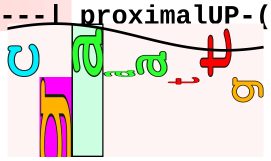 sigma38p10 0.5 bits

```
... --------------| distalUP-(41)-sigma38p10 2022888 Gap 3.1 bits
... --------------| sigma38p10-proximalUP-distalUP 2022888 total 8.3 bits
... ----------------| proximalUP-(36)-sigma38p10 2022887 Gap 3.6 bits
... ----------------| distalUP-(42)-sigma38p10 2022887 Gap 3.3 bits
... ----------------| sigma38p10-proximalUP-distalUP 2022887 total 6.9 bits

          .         *2022850  .         *2022840  .         *2022830  .         *2022820
 5' g a g g a a a a c g a t g a a g g t g a a t g a t c g g g t a a c a g t c a 3'

              .         *2022810  .         *2022800
 5' a a a c g g a t g g c g g t c c g c g t 3'
```

piece 7, NC_000913, lpxP-yfdY, config: linear, direction: -, begin: 2493599, end: 2493400

. *2493590 . *2493580 . *2493570 .
5′ t g g t t g g g c c a t c a t g t a c g a c a a t g g t t c c t g t a a a g 3′
T timestamp:_2018Sep20_11-36-13

*2493560 . *2493550 . *2493540 . *2493530 .
5′ a g t g t a a a g a c t a g t c a g g a c a t a a c c a c g a g g a g c a t 3′
... proximalUP

*2493520 . *2493510 . *2493500 . *2493490
5′ g a a t g c g g t a t c t g t a c t t t a t c a c t c g c t g a c t g c t t 3′
... proximalUP 4.6 bits proximalUP 6.8 bits
... proximalUP-(43)
distalUP 2.4 bits ... proximalUP-(29)
... distalUP-(40)-s
... sigma38p10-dist
... distalUP-(44)-s
... sigma38p10-prox
... proximalUP-(32)
distalUP 0.2 bits
... distalUP-(41)-s
... sigma38p10-prox

. *2493480 . *2493470 . *2493460 . *2493450
5′ t c a g c t a c t g a t t a t t a a t c g t g c a a a g t a a a t a t t t t 3′
sigma38p10 1.5 bits ... proximalUP
... proximalUP-(43)-sigma38p10 2493476 Gap 5.1 bits ... proximalUP
... proximalUP-(29)-sigma38p10 2493472 Gap 7.5 bits
... distalUP-(40)-sigma38p10 2493476 Gap 3.1 bits
... sigma38p10-distalUP-proximalUP 2493476 total 0.3 bits
... distalUP-(44)-sigma38p10 2493472 Gap 2.8 bits
... sigma38p10-proximalUP-distalUP 2493472 total 3.1 bits
... proximalUP-(32)-sigma38p10 2493469 Gap 4.3 bits
sigma38p10 4.3 bits

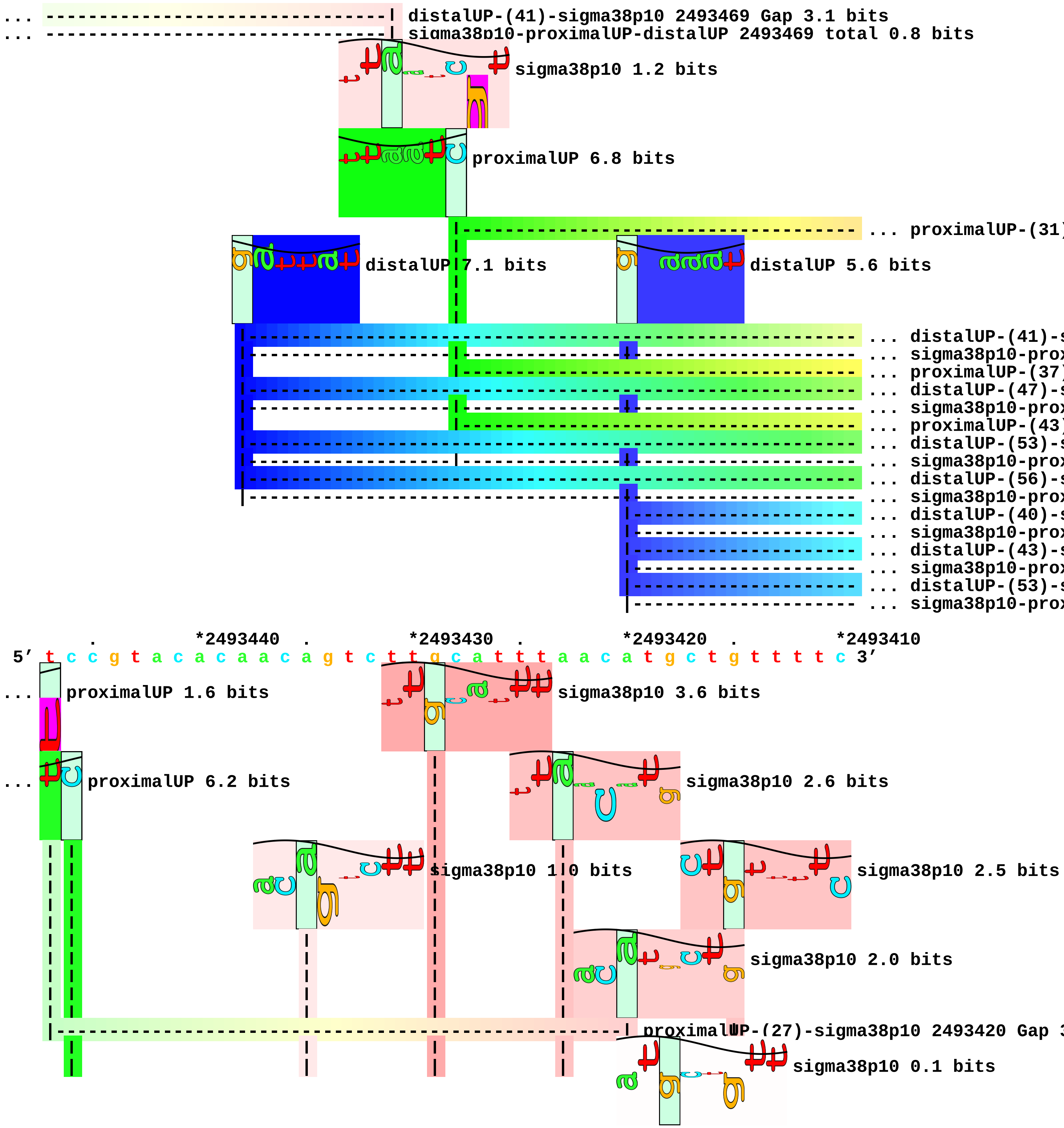
distalUP-(41)-sigma38p10 2493469 Gap 3.1 bits
sigma38p10-proximalUP-distalUP 2493469 total 0.8 bits
sigma38p10 1.2 bits
proximalUP 6.8 bits
distalUP 7.1 bits
distalUP 5.6 bits
*2493440
*2493430
*2493420
*2493410
5′ t c c g t a c a c a a c a g t c t t g c a t t t a a c a t g c t g t t t t c 3′
proximalUP 1.6 bits
sigma38p10 3.6 bits
proximalUP 6.2 bits
sigma38p10 2.6 bits
sigma38p10 1.0 bits
sigma38p10 2.5 bits
sigma38p10 2.0 bits
sigma38p10 0.1 bits

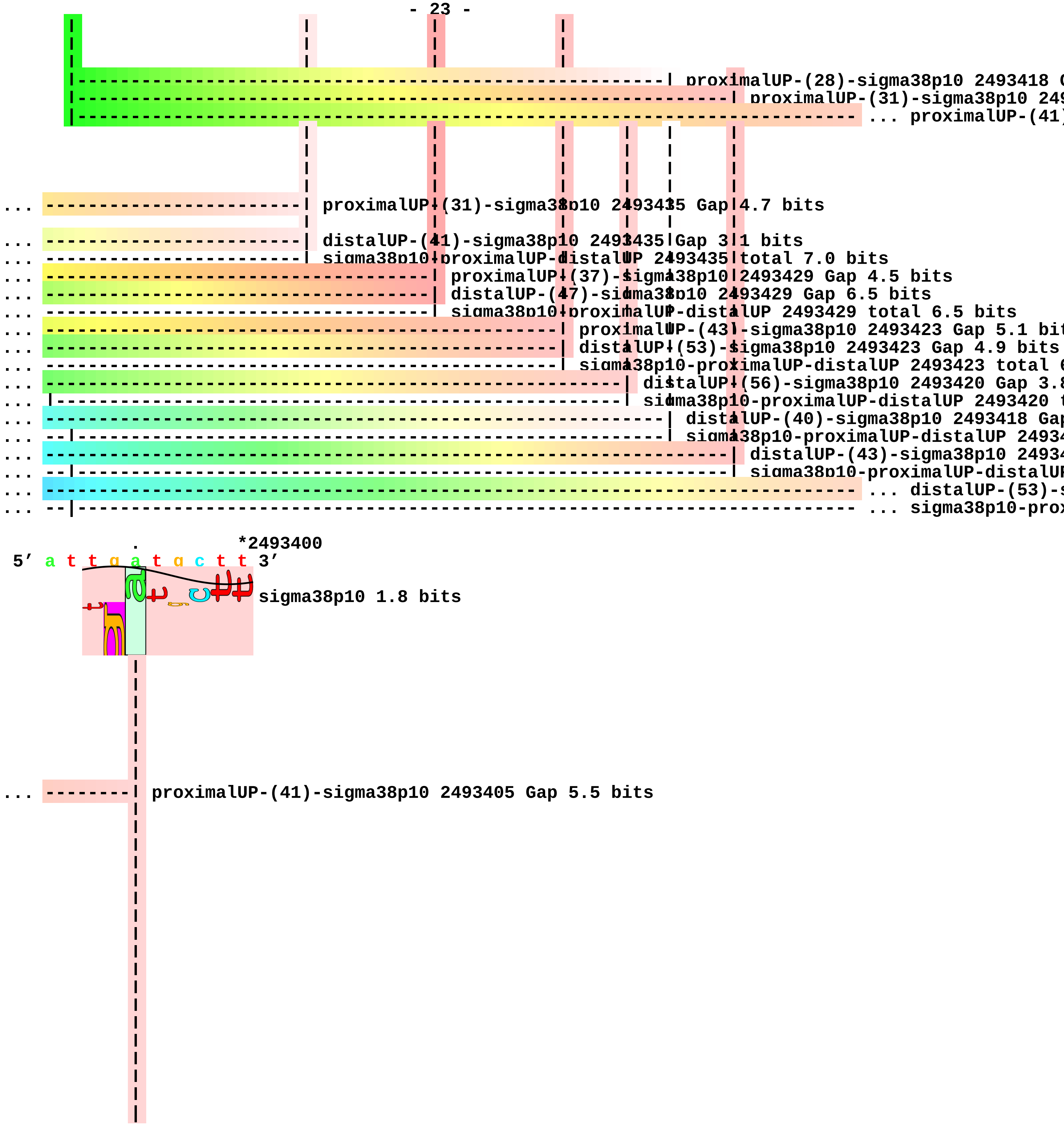
proximalUP-(28)-sigma38p10 2493418
proximalUP-(31)-sigma38p10
... proximalUP-(41)
... proximalUP-(31)-sigma38p10 2493435 Gap 4.7 bits
... distalUP-(41)-sigma38p10 2493435 Gap 3.1 bits
... sigma38p10-proximalUP-distalUP 2493435 total 7.0 bits
... proximalUP-(37)-sigma38p10 2493429 Gap 4.5 bits
... distalUP-(47)-sigma38p10 2493429 Gap 6.5 bits
... sigma38p10-proximalUP-distalUP 2493429 total 6.5 bits
... proximalUP-(43)-sigma38p10 2493423 Gap 5.1 bit
... distalUP-(53)-sigma38p10 2493423 Gap 4.9 bits
... sigma38p10-proximalUP-distalUP 2493423 total
... distalUP-(56)-sigma38p10 2493420 Gap 3.
... sigma38p10-proximalUP-distalUP 2493420
... distalUP-(40)-sigma38p10 2493418 Ga
... sigma38p10-proximalUP-distalUP 24934
... distalUP-(43)-sigma38p10 24934
... sigma38p10-proximalUP-distalU
... distalUP-(53)-
... sigma38p10-pro
*2493400
5′ a t t g a t g c t t 3′
sigma38p10 1.8 bits
... proximalUP-(41)-sigma38p10 2493405 Gap 5.5 bits

```
                  |
                  |
... --------| distalUP-(53)-sigma38p10 2493405 Gap 4.9 bits
... --------| sigma38p10-proximalUP-distalUP 2493405 total 3.3 bits
```

piece 8, NC_000913, yfgG-yfgF, config: linear, direction: -, begin: 2627449, end: 2627050

. *2627440 . *2627430 . *2627420 .
5′ g c t c t g a t g g t g c t g c c a g g c a c c g a c g g a c g a g t a g a 3′
T timestamp:_2018Sep20_11-36-13

*2627410 . *2627400 . *2627390 . *2627380 .
5′ t a a a a c g g c c a a a g a a g a a g a t a a a g c t g a t g a g c a g t 3′

*2627370 . *2627360 . *2627350 . *2627340
5′ a c g a t a c g g g t c a t g c g a c t g t t a a a t c g g t g t c g t t t 3′

. *2627330 . *2627320 . *2627310 . *2627300
5′ t c g c a t a c t g g t t g c c t g a c t c a c a a a a g g t t c c t t g a 3′

. *2627290 . *2627280 . *2627270 . *2627260
5′ a g t a t g t c c c a c g c t g t g g a c g g t a c t t a c a t t a a g g c 3′

. *2627250 . *2627240 . *2627230 .
5′ a c a a c a g g a c a a a a t g g t c a a t t c t t c g t t a t g t a a a a 3′

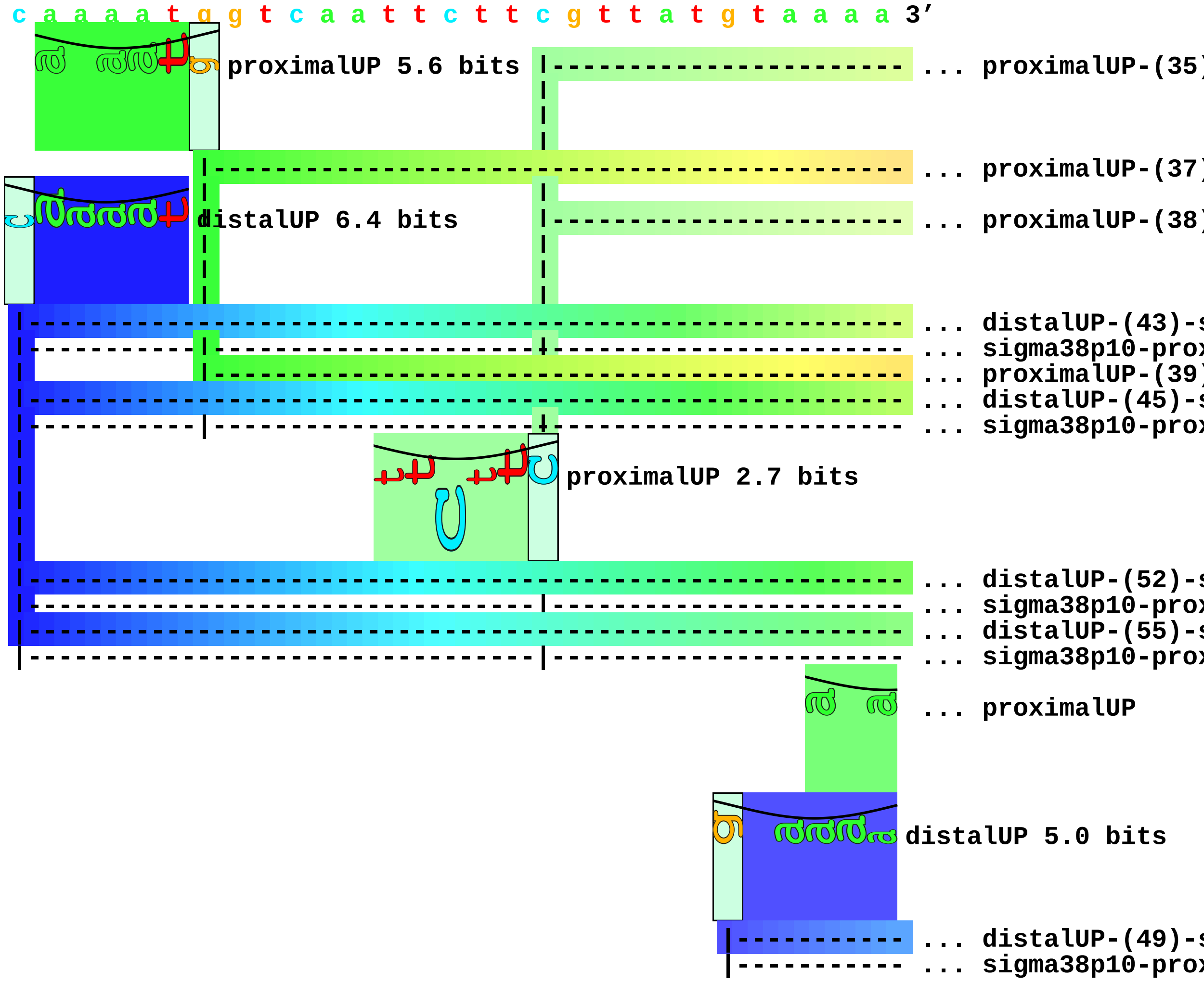


*2627220 . *2627210 . *2627200 . *2627190 .
5′ a a g c g t c a g t g g a t a c a t a t t t t a a t g t t a t g g a a g t t 3′
... proximalUP-(35)-sigma38p10 2627198 Gap 3.1 bits

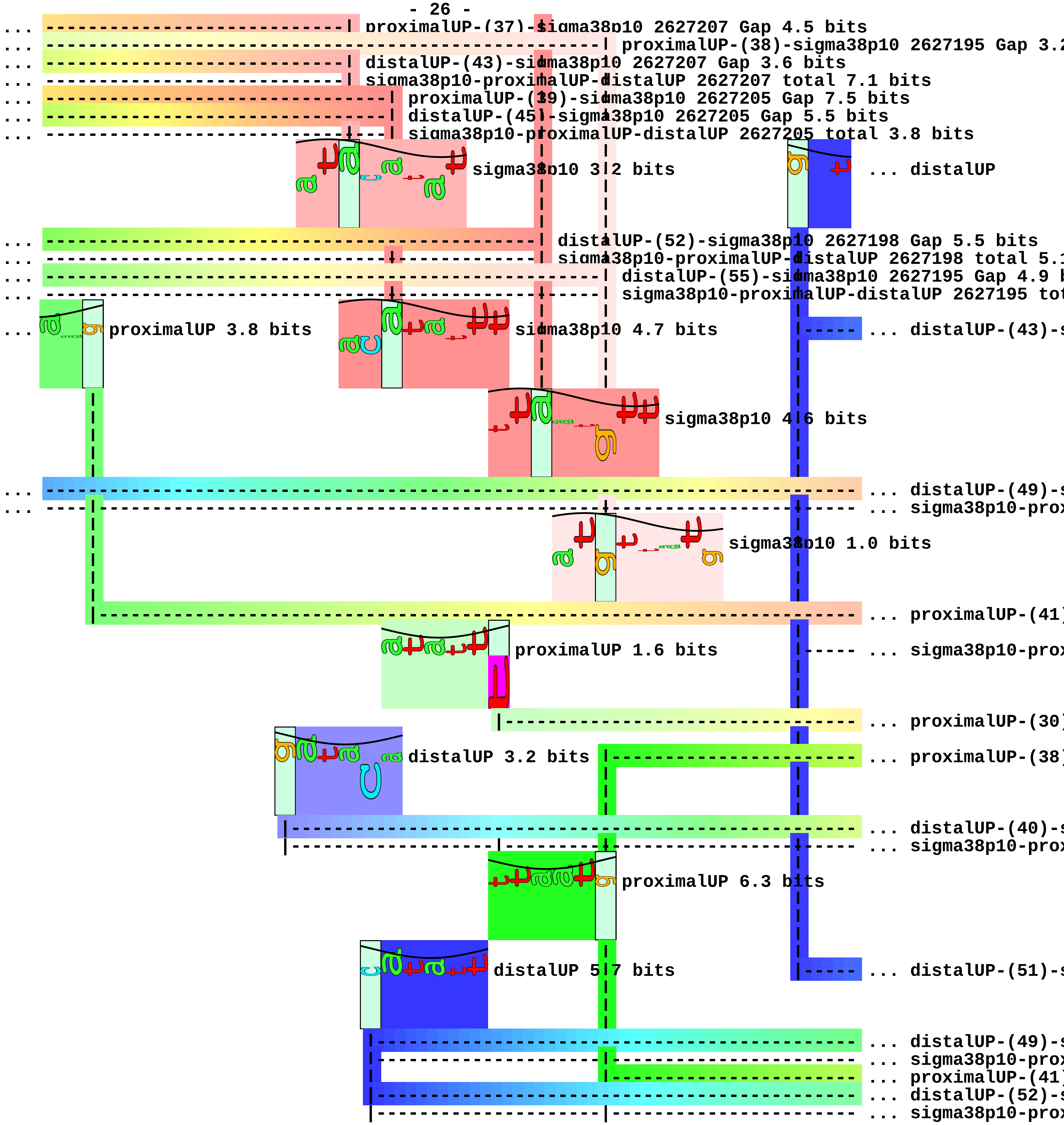
proximalUP-(37)-sigma38p10 2627207 Gap 4.5 bits
proximalUP-(38)-sigma38p10 2627195 Gap 3.2
distalUP-(43)-sigma38p10 2627207 Gap 3.6 bits
sigma38p10-proximalUP-distalUP 2627207 total 7.1 bits
proximalUP-(39)-sigma38p10 2627205 Gap 7.5 bits
distalUP-(45)-sigma38p10 2627205 Gap 5.5 bits
sigma38p10-proximalUP-distalUP 2627205 total 3.8 bits
sigma38p10 3.2 bits
... distalUP
distalUP-(52)-sigma38p10 2627198 Gap 5.5 bits
sigma38p10-proximalUP-distalUP 2627198 total 5.1
distalUP-(55)-sigma38p10 2627195 Gap 4.9 b
sigma38p10-proximalUP-distalUP 2627195 tot
proximalUP 3.8 bits
sigma38p10 4.7 bits
... distalUP-(43)-s
sigma38p10 4.6 bits
... distalUP-(49)-s
... sigma38p10-prox
sigma38p10 1.0 bits
... proximalUP-(41)
proximalUP 1.6 bits
... sigma38p10-prox
... proximalUP-(30)
distalUP 3.2 bits
... proximalUP-(38)
... distalUP-(40)-s
... sigma38p10-prox
proximalUP 6.3 bits
distalUP 5.7 bits
... distalUP-(51)-s
... distalUP-(49)-s
... sigma38p10-prox
... proximalUP-(41)
... distalUP-(52)-s
... sigma38p10-prox

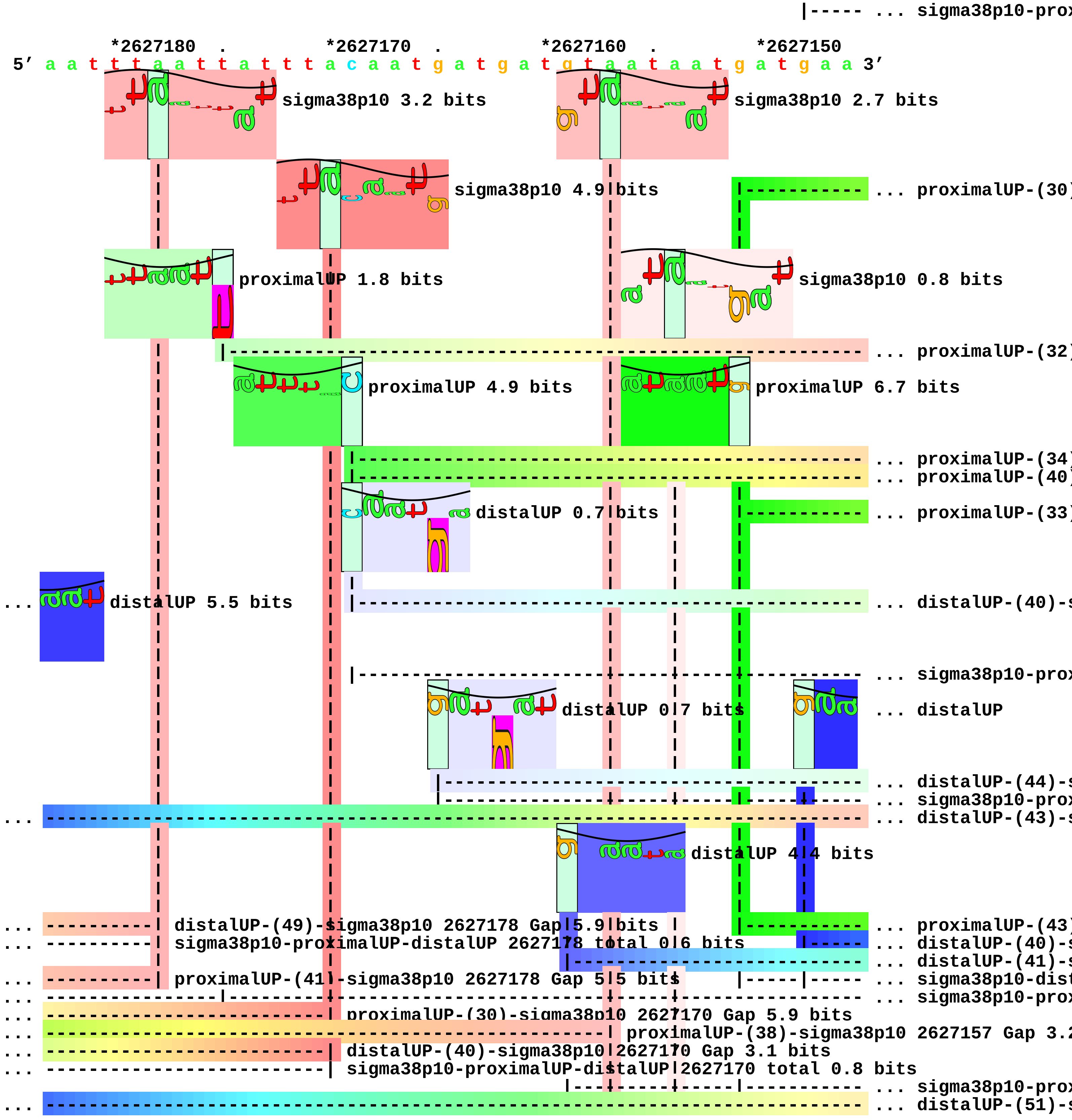
|----- ... sigma38p10-prox
*2627180 . *2627170 . *2627160 . *2627150
5' a a t t t a a t t a t t t a c a a t g a t g a t g t a a t a a t g a t g a a 3'
sigma38p10 3.2 bits
sigma38p10 2.7 bits
sigma38p10 4.9 bits
... proximalUP-(30)
proximalUP 1.8 bits
sigma38p10 0.8 bits
... proximalUP-(32)
proximalUP 4.9 bits
proximalUP 6.7 bits
... proximalUP-(34)
... proximalUP-(40)
distalUP 0.7 bits
... proximalUP-(33)
... distalUP 5.5 bits
... distalUP-(40)-s
... sigma38p10-prox
distalUP 0.7 bits
... distalUP
... distalUP-(44)-s
... sigma38p10-prox
... distalUP-(43)-s
distalUP 4.4 bits
... distalUP-(49)-sigma38p10 2627178 Gap 5.9 bits
... sigma38p10-proximalUP-distalUP 2627178 total 0.6 bits
... proximalUP-(41)-sigma38p10 2627178 Gap 5.5 bits
... proximalUP-(43)
... distalUP-(40)-s
... distalUP-(41)-s
... sigma38p10-dist
... sigma38p10-prox
... proximalUP-(30)-sigma38p10 2627170 Gap 5.9 bits
... proximalUP-(38)-sigma38p10 2627157 Gap 3.2
... distalUP-(40)-sigma38p10 2627170 Gap 3.1 bits
... sigma38p10-proximalUP-distalUP 2627170 total 0.8 bits
... sigma38p10-prox
... distalUP-(51)-s

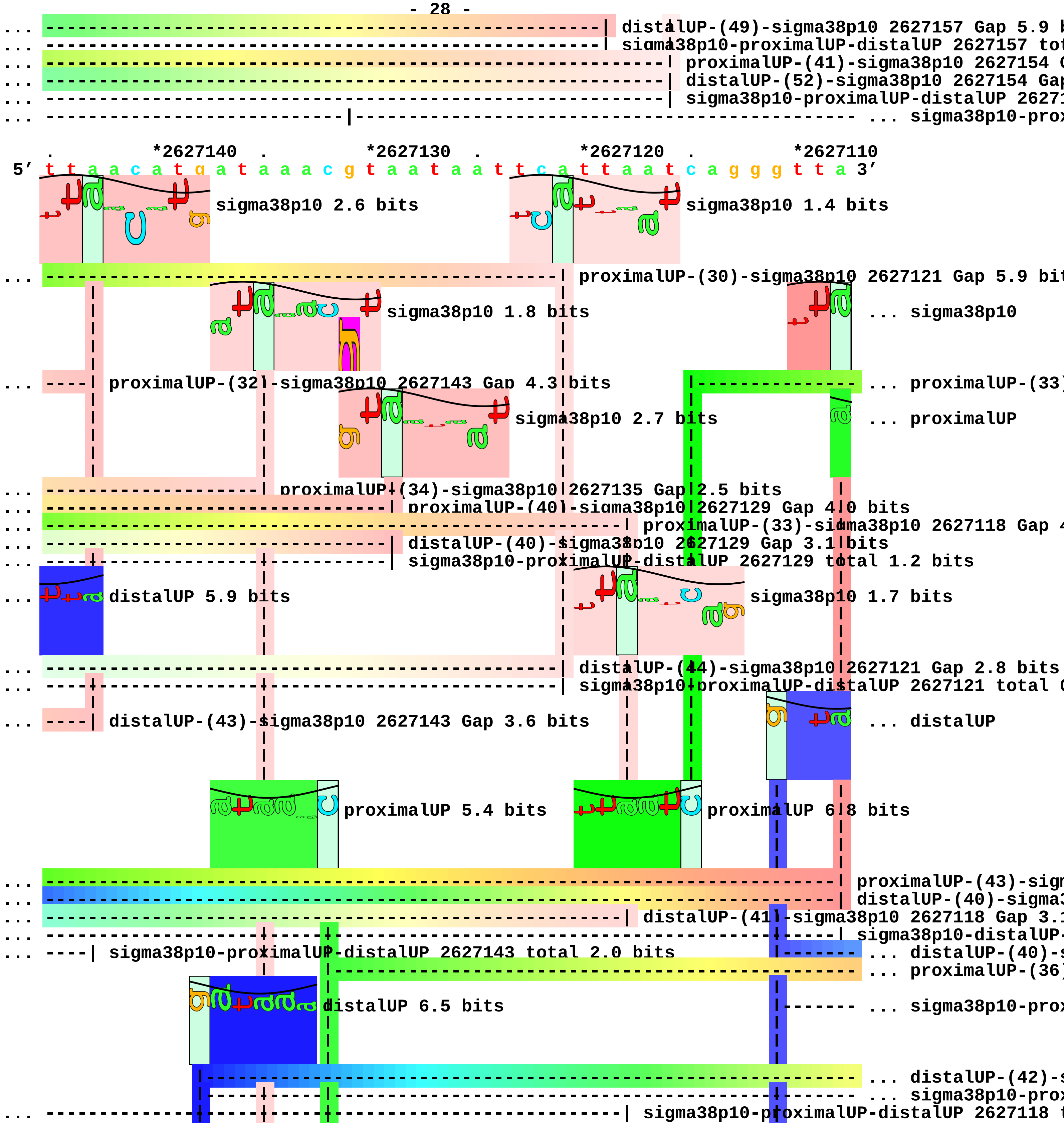
distalUP-(49)-sigma38p10 2627157 Gap 5.9 b
sigma38p10-proximalUP-distalUP 2627157 tot
proximalUP-(41)-sigma38p10 2627154 G
distalUP-(52)-sigma38p10 2627154 Gap
sigma38p10-proximalUP-distalUP 26271
... sigma38p10-prox
*2627140
*2627130
*2627120
*2627110
5′ t t a a c a t g a t a a a c g t a a t a a t t c a t t a a t c a g g g t t a 3′
sigma38p10 2.6 bits
sigma38p10 1.4 bits
proximalUP-(30)-sigma38p10 2627121 Gap 5.9 bit
sigma38p10 1.8 bits
... sigma38p10
proximalUP-(32)-sigma38p10 2627143 Gap 4.3 bits
... proximalUP-(33)
sigma38p10 2.7 bits
... proximalUP
proximalUP-(34)-sigma38p10 2627135 Gap 2.5 bits
proximalUP-(40)-sigma38p10 2627129 Gap 4.0 bits
proximalUP-(33)-sigma38p10 2627118 Gap 4
distalUP-(40)-sigma38p10 2627129 Gap 3.1 bits
sigma38p10-proximalUP-distalUP 2627129 total 1.2 bits
distalUP 5.9 bits
sigma38p10 1.7 bits
distalUP-(44)-sigma38p10 2627121 Gap 2.8 bits
sigma38p10-proximalUP-distalUP 2627121 total 0
distalUP-(43)-sigma38p10 2627143 Gap 3.6 bits
... distalUP
proximalUP 5.4 bits
proximalUP 6.8 bits
proximalUP-(43)-sigm
distalUP-(40)-sigma3
distalUP-(41)-sigma38p10 2627118 Gap 3.1
sigma38p10-distalUP-
sigma38p10-proximalUP-distalUP 2627143 total 2.0 bits
... distalUP-(40)-s
... proximalUP-(36)
distalUP 6.5 bits
... sigma38p10-prox
... distalUP-(42)-s
... sigma38p10-prox
sigma38p10-proximalUP-distalUP 2627118 t

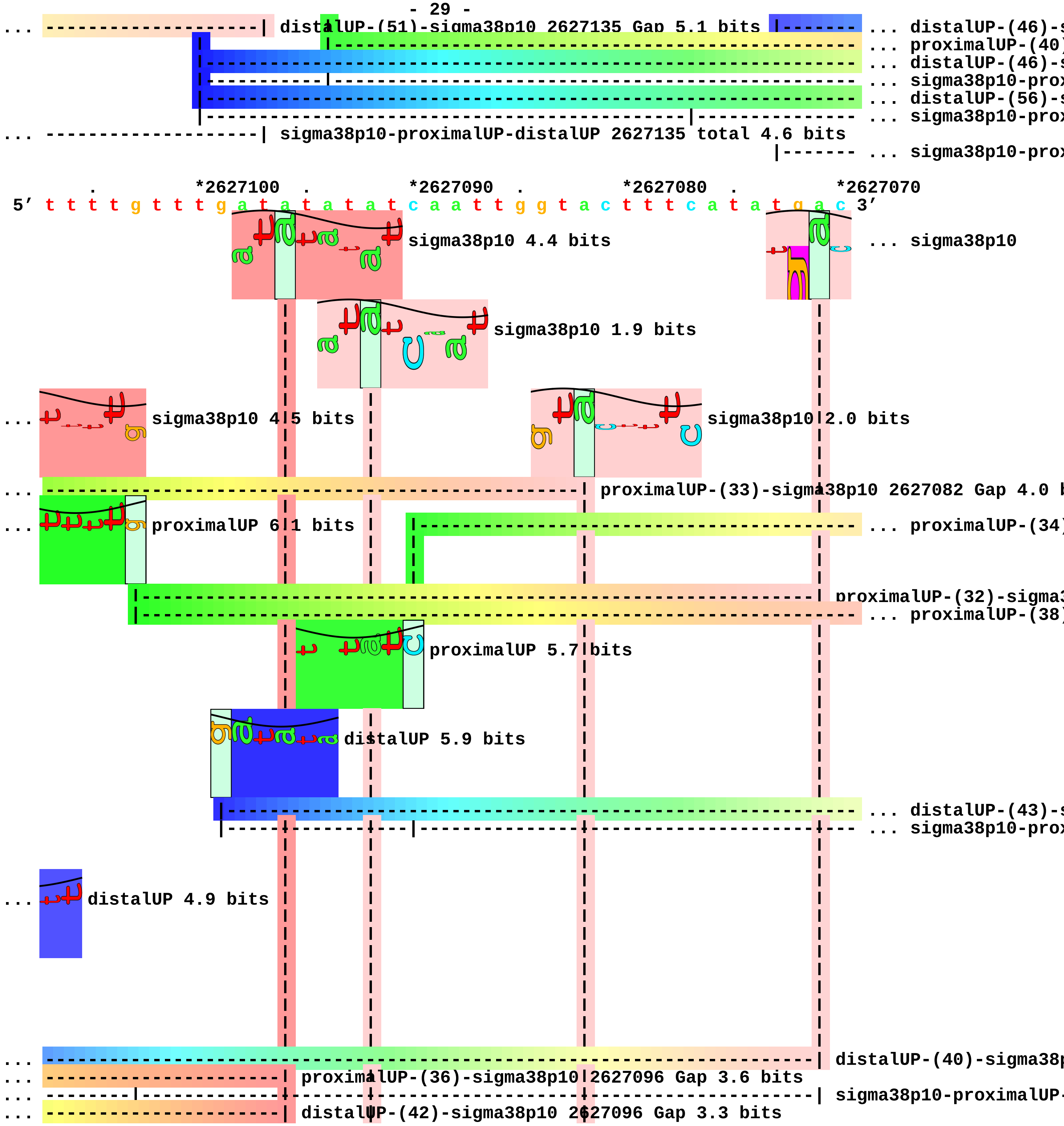

distalUP-(51)-sigma38p10 2627135 Gap 5.1 bits
... distalUP-(46)-s
... proximalUP-(40)
... distalUP-(46)-s
... sigma38p10-prox
... distalUP-(56)-s
... sigma38p10-prox
sigma38p10-proximalUP-distalUP 2627135 total 4.6 bits
... sigma38p10-prox
*2627100
*2627090
*2627080
*2627070
5′ t t t t g t t t g a t a t a t a t c a a t t g g t a c t t t c a t a t g a c 3′
sigma38p10 4.4 bits
... sigma38p10
sigma38p10 1.9 bits
sigma38p10 4.5 bits
sigma38p10 2.0 bits
proximalUP-(33)-sigma38p10 2627082 Gap 4.0 b
proximalUP 6.1 bits
... proximalUP-(34)
proximalUP-(32)-sigma3
... proximalUP-(38)
proximalUP 5.7 bits
distalUP 5.9 bits
... distalUP-(43)-s
... sigma38p10-prox
distalUP 4.9 bits
distalUP-(40)-sigma38p
proximalUP-(36)-sigma38p10 2627096 Gap 3.6 bits
sigma38p10-proximalUP-
distalUP-(42)-sigma38p10 2627096 Gap 3.3 bits

... ---------------------| sigma38p10-proximalUP-distalUP 2627096 total 9.4 bits

... ----------------------------------------------------------------------------- ... distalUP-(46)-s
... ------------------------------| proximalUP-(40)-sigma38p10 2627092 Gap 4.0 bits
... ------------------------------| distalUP-(46)-sigma38p10 2627092 Gap 4.1 bits
... ------------------------------| sigma38p10-proximalUP-distalUP 2627092 total 5.7 bits
... --------------------------------------------------| distalUP-(56)-sigma38p10 2627082 Gap 3.8 bit
... --------------------------------------------------| sigma38p10-proximalUP-distalUP 2627082 tota

... --------|------------------------------------------------------------------ ... sigma38p10-prox

. *2627060 . *2627050

5′ a c t t a g a a t c a a t g c t c t c t 3′

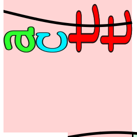

... sigma38p10 1.8 bits

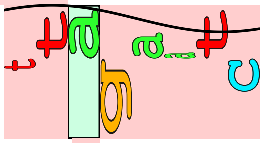

sigma38p10 2.1 bits

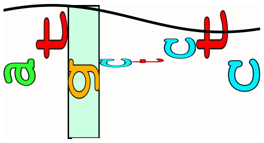

sigma38p10 0.1 bits

... --------------------------| proximalUP-(34)-sigma38p10 2627056 Gap 2.5 bits

... --------| proximalUP-(38)-sigma38p10 2627065 Gap 3.2 bits

... --------------------------| distalUP-(43)-sigma38p10 2627056 Gap 3.6 bits
... --------------------------| sigma38p10-proximalUP-distalUP 2627056 total 5.6 bits

... --------| distalUP-(46)-sigma38p10 2627065 Gap 4.1 bits

... --------| sigma38p10-proximalUP-distalUP 2627065 total 5.8 bits piece 9, NC_000913, ygcG-queE, config: linear, direction: -, begin: 2903699, end: 2903300

. *2903690 . *2903680 . *2903670 .
5′ t g a c g g c g g c a t c g g t a a t t t t a t a t t c g t t a t c a t t a 3′

T timestamp:_2018Sep20_11-36-13

proximalUP 6.6 bits
... proximalUP-(31)
distalUP 5.0 bits
... proximalUP
... distalUP-(44)-s
... sigma38p10-prox
... proximalUP-(35)
... distalUP-(48)-s
... sigma38p10-prox
... proximalUP-(37)
... distalUP-(50)-s
... sigma38p10-prox
... distalUP
... distalUP-(44)-s
... sigma38p10-prox
... distalUP-(52)-s
... sigma38p10-prox

*2903660 . *2903650 . *2903640 . *2903630 .
5′ a a a a c t a a t a c a g a a a a a a t a a a t a t a a t t t t g g t g a a 3′

sigma38p10 3.7 bits
... sigma38p10
... proximalUP-(31)-sigma38p10 2903641 Gap 4.7 bits
... proximalUP 4.3 bits
sigma38p10 8.1 bits
... distalUP-(44)-sigma38p10 2903641 Gap 2.8 bits
... sigma38p10-proximalUP-distalUP 2903641 total 7.9 bits
... proximalUP-(35)-sigma38p10 2903637 Gap 3.1 bit
... distalUP-(48)-sigma38p10 2903637 Gap 7.5 bits
... sigma38p10-proximalUP-distalUP 2903637 total 9
... proximalUP-(37)-sigma38p10 2903635 Gap 4.5
... distalUP-(50)-sigma38p10 2903635 Gap 4.1 b
... sigma38p10-proximalUP-distalUP 2903635 tot
... distalUP 5.6 bits
sigma38p10 5.8 bits
... ... distalUP-(44)-s
... ... sigma38p10-prox

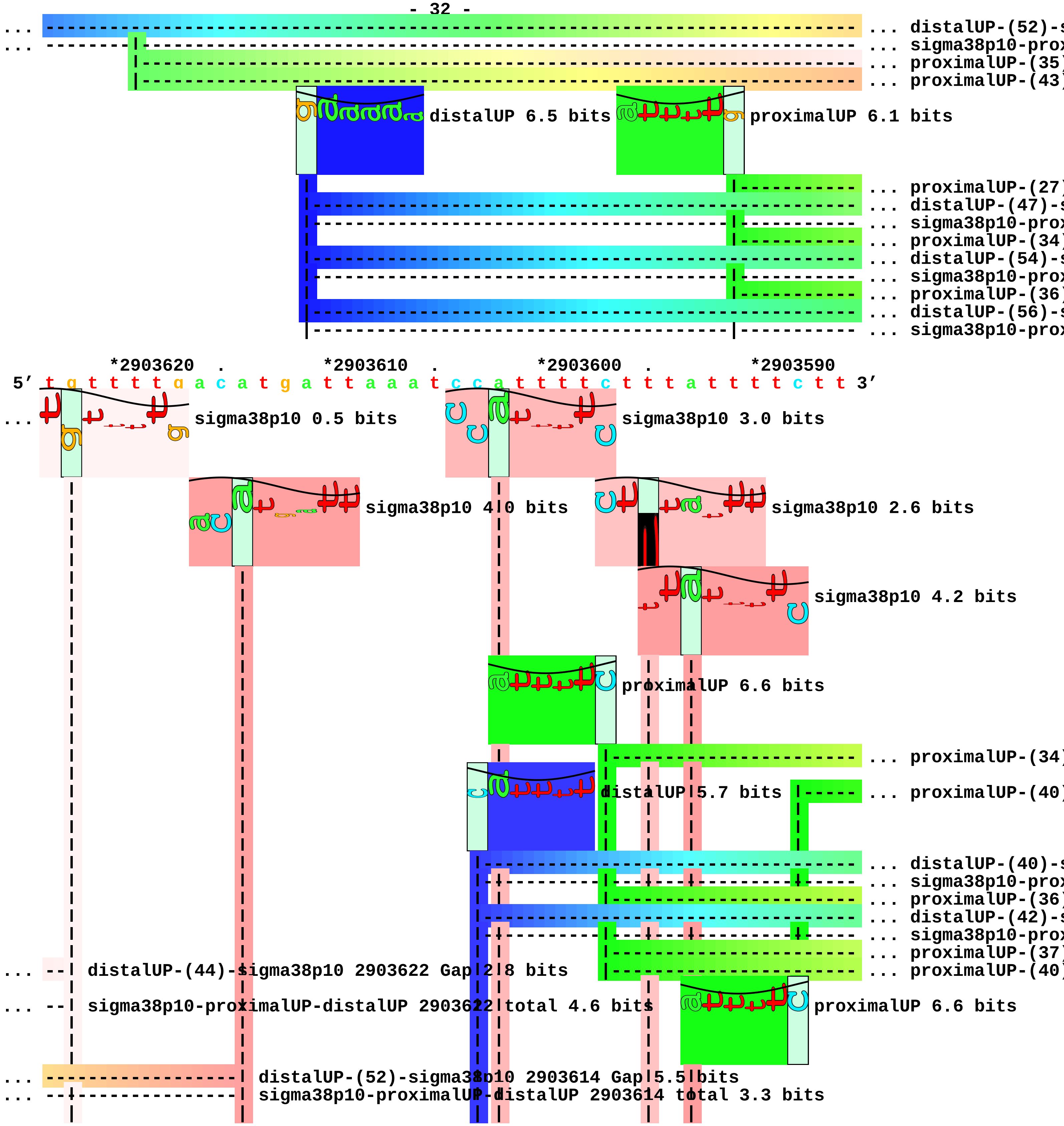
... distalUP-(52)-s
... sigma38p10-prox
... proximalUP-(35)
... proximalUP-(43)
distalUP 6.5 bits
proximalUP 6.1 bits
... proximalUP-(27)
... distalUP-(47)-s
... sigma38p10-prox
... proximalUP-(34)
... distalUP-(54)-s
... sigma38p10-prox
... proximalUP-(36)
... distalUP-(56)-s
... sigma38p10-prox
*2903620 . *2903610 . *2903600 . *2903590
5′ t g t t t t g a c a t g a t t a a a t c c a t t t t c t t t a t t t t t c t t 3′
sigma38p10 0.5 bits
sigma38p10 3.0 bits
sigma38p10 4.0 bits
sigma38p10 2.6 bits
sigma38p10 4.2 bits
proximalUP 6.6 bits
... proximalUP-(34)
distalUP 5.7 bits
... proximalUP-(40)
... distalUP-(40)-s
... sigma38p10-prox
... proximalUP-(36)
... distalUP-(42)-s
... sigma38p10-prox
... proximalUP-(37)
... proximalUP-(40)
distalUP-(44)-sigma38p10 2903622 Gap 2.8 bits
sigma38p10-proximalUP-distalUP 2903622 total 4.6 bits
proximalUP 6.6 bits
distalUP-(52)-sigma38p10 2903614 Gap 5.5 bits
sigma38p10-proximalUP-distalUP 2903614 total 3.3 bits

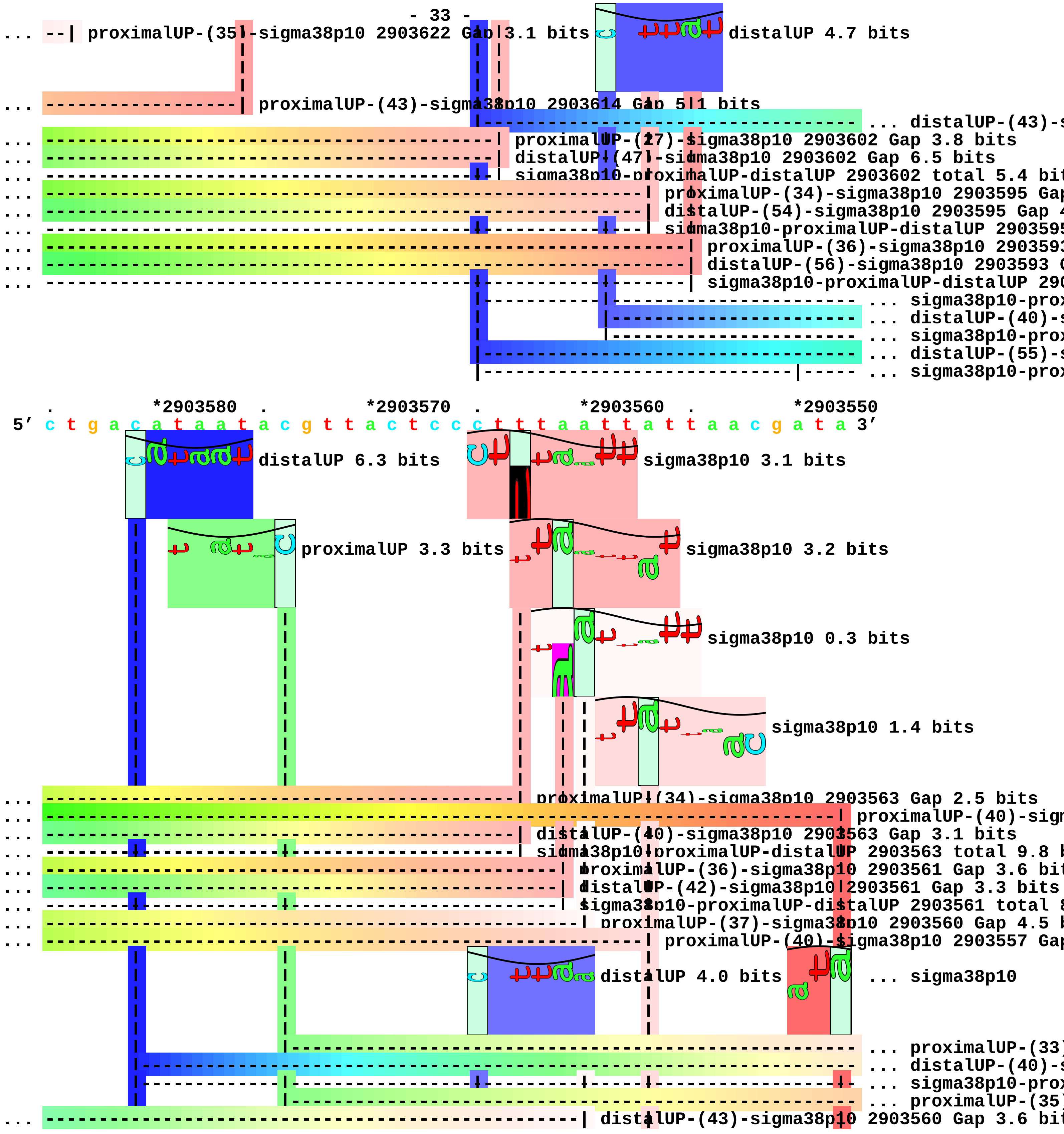
... --| proximalUP-(35)-sigma38p10 2903622 Gap 3.1 bits
distalUP 4.7 bits
... -----------------| proximalUP-(43)-sigma38p10 2903614 Gap 5.1 bits
... distalUP-(43)-s
... proximalUP-(27)-sigma38p10 2903602 Gap 3.8 bits
... distalUP-(47)-sigma38p10 2903602 Gap 6.5 bits
... sigma38p10-proximalUP-distalUP 2903602 total 5.4 bit
... proximalUP-(34)-sigma38p10 2903595 Gap
... distalUP-(54)-sigma38p10 2903595 Gap 4
... sigma38p10-proximalUP-distalUP 290359
... proximalUP-(36)-sigma38p10 290359
... distalUP-(56)-sigma38p10 2903593 G
... sigma38p10-proximalUP-distalUP 290
... sigma38p10-prox
... distalUP-(40)-s
... sigma38p10-prox
... distalUP-(55)-s
... sigma38p10-prox
. *2903580 . *2903570 . *2903560 . *2903550
5′ c t g a c a t a a t a c g t t a c t c c c t t t a a t t a t t a a c g a t a 3′
distalUP 6.3 bits
sigma38p10 3.1 bits
proximalUP 3.3 bits
sigma38p10 3.2 bits
sigma38p10 0.3 bits
sigma38p10 1.4 bits
... proximalUP-(34)-sigma38p10 2903563 Gap 2.5 bits
... proximalUP-(40)-sign
... distalUP-(40)-sigma38p10 2903563 Gap 3.1 bits
... sigma38p10-proximalUP-distalUP 2903563 total 9.8 b
... proximalUP-(36)-sigma38p10 2903561 Gap 3.6 bit
... distalUP-(42)-sigma38p10 2903561 Gap 3.3 bits
... sigma38p10-proximalUP-distalUP 2903561 total 8
... proximalUP-(37)-sigma38p10 2903560 Gap 4.5 b
... proximalUP-(40)-sigma38p10 2903557 Gap
distalUP 4.0 bits
... sigma38p10
... proximalUP-(33)
... distalUP-(40)-s
... sigma38p10-prox
... proximalUP-(35)
... distalUP-(43)-sigma38p10 2903560 Gap 3.6 bit

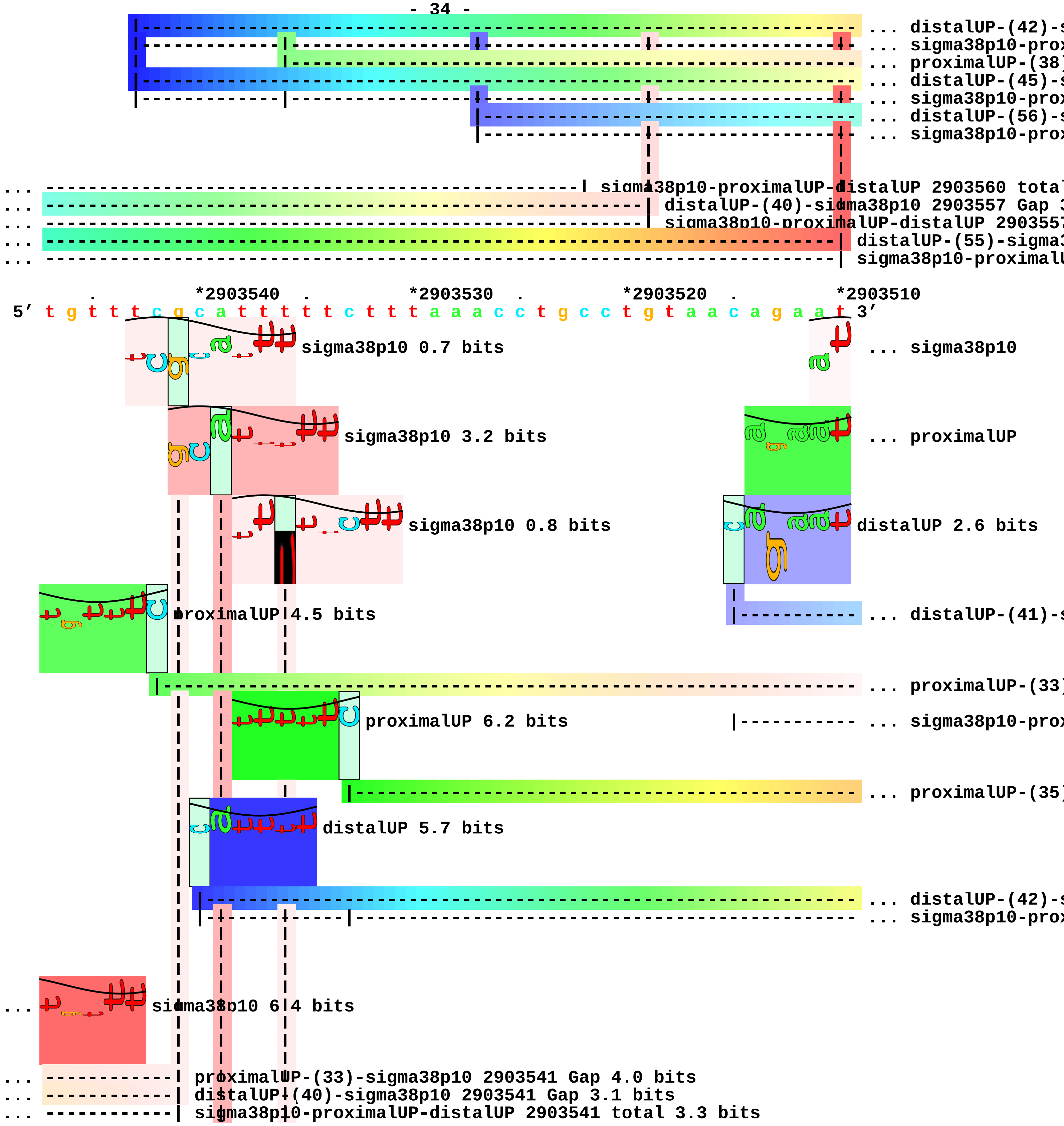
... distalUP-(42)-s
... sigma38p10-prox
... proximalUP-(38)
... distalUP-(45)-s
... sigma38p10-prox
... distalUP-(56)-s
... sigma38p10-prox
sigma38p10-proximalUP-distalUP 2903560 total
distalUP-(40)-sigma38p10 2903557 Gap 3
sigma38p10-proximalUP-distalUP 290355
distalUP-(55)-sigma3
sigma38p10-proximalU
*2903540
*2903530
*2903520
*2903510
5′ t g t t t c g c a t t t t t c t t t a a a c c t g c c t g t a a c a g a a t 3′
sigma38p10 0.7 bits
... sigma38p10
sigma38p10 3.2 bits
... proximalUP
sigma38p10 0.8 bits
distalUP 2.6 bits
proximalUP 4.5 bits
... distalUP-(41)-s
... proximalUP-(33)
proximalUP 6.2 bits
... sigma38p10-prox
... proximalUP-(35)
distalUP 5.7 bits
... distalUP-(42)-s
... sigma38p10-prox
sigma38p10 6.4 bits
proximalUP-(33)-sigma38p10 2903541 Gap 4.0 bits
distalUP-(40)-sigma38p10 2903541 Gap 3.1 bits
sigma38p10-proximalUP-distalUP 2903541 total 3.3 bits

```
... ----------------| proximalUP-(35)-sigma38p10 2903539 Gap 3.1 bits
                    |
... ----------------| distalUP-(42)-sigma38p10 2903539 Gap 3.3 bits
... ----------------| sigma38p10-proximalUP-distalUP 2903539 total 6.4 bits
... ----------------------| proximalUP-(38)-sigma38p10 2903536 Gap 3.2 bits
... ----------------------| distalUP-(45)-sigma38p10 2903536 Gap 5.5 bits
... ----------------------| sigma38p10-proximalUP-distalUP 2903536 total 1.7 bits
... ------------------------------------------------------------------------------ ... distalUP-(56)-s
... ----------|------------------------------------------------------------------- ... sigma38p10-prox

              .          *2903500  .          *2903490  .          *2903480  .
 5' g t t a a t a a g a c a t g a t t g t t g t g g c a g g a t c t g c a a t a 3'
... sigma38p10 0.4 bits                                                        ... sigma38p10
... proximalUP 5.0 bits                proximalUP 4.0 bits
                                       sigma38p10 4.0 bits
... ------------------------------------------------------------------------| distalUP-(41)-sigma38p10
... | proximalUP-(33)-sigma38p10 2903509 Gap 4.0 bits
... |-----------------------------------------------------------------------| sigma38p10-proximalUP-di
... ----------------------| proximalUP-(35)-sigma38p10 2903498 Gap 3.1 bits
... |-----------------------------------------------------------------------| proximalUP-(35)-sigma38p
... ----------------------| distalUP-(42)-sigma38p10 2903498 Gap 3.3 bits
... |---------------------| sigma38p10-proximalUP-distalUP 2903498 total 9.5 bits
                      proximalUP 3.8 bits
                      |----------------------------------------------------------- ... proximalUP-(33)
    distalUP 5.5 bits                          |------------------------------------ ... proximalUP-(33)
... |------------------------------------------------------------------------------ ... distalUP-(41)-s
... |------------------------------------------------------------------------------ ... sigma38p10-prox
... |------------------------------------------------------------------------------ ... proximalUP-(34)
... |------------------------------------------------------------------------------ ... distalUP-(42)-s
... |---------------------|-------------------------------------------------------- ... sigma38p10-prox
... |------------------------------------------------------------------------------ ... distalUP-(50)-s
... |------------------------------|----------------------------------------------- ... sigma38p10-prox
... | distalUP-(56)-sigma38p10 2903509 Gap 3.8 bits
... | sigma38p10-proximalUP-distalUP 2903509 total 1.2 bits

      *2903470  .          *2903460  .          *2903450  .          *2903440  .
 5' a t t a a c a a t t a t c c a c t t c a c a g a g a a t g c t a t g c a g t 3'
```

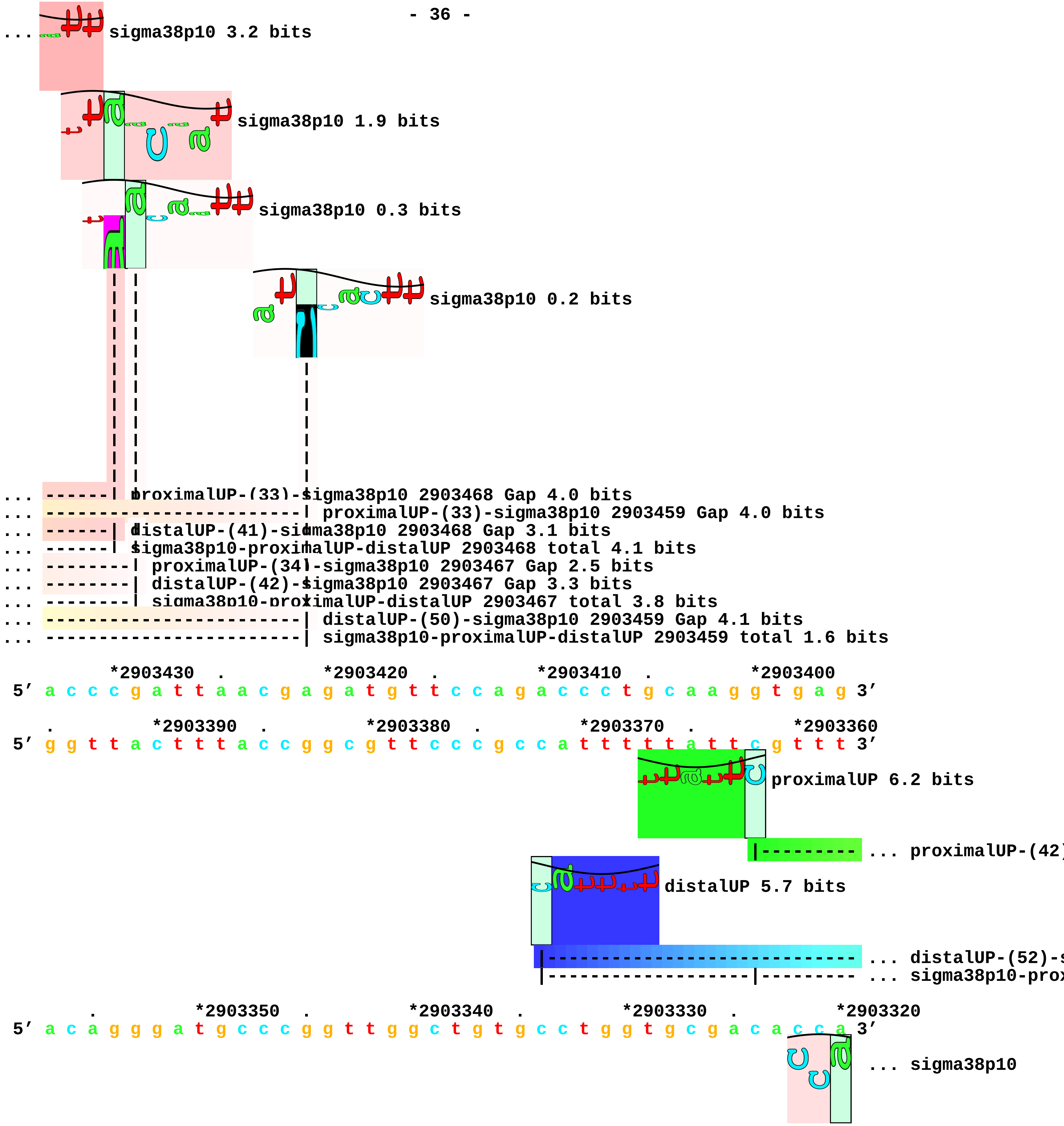
... sigma38p10 3.2 bits
sigma38p10 1.9 bits
sigma38p10 0.3 bits
sigma38p10 0.2 bits
... ------| proximalUP-(33)-sigma38p10 2903468 Gap 4.0 bits
... -----------------------| proximalUP-(33)-sigma38p10 2903459 Gap 4.0 bits
... ------| distalUP-(41)-sigma38p10 2903468 Gap 3.1 bits
... ------| sigma38p10-proximalUP-distalUP 2903468 total 4.1 bits
... --------| proximalUP-(34)-sigma38p10 2903467 Gap 2.5 bits
... --------| distalUP-(42)-sigma38p10 2903467 Gap 3.3 bits
... --------| sigma38p10-proximalUP-distalUP 2903467 total 3.8 bits
... -----------------------| distalUP-(50)-sigma38p10 2903459 Gap 4.1 bits
... -----------------------| sigma38p10-proximalUP-distalUP 2903459 total 1.6 bits
*2903430 . *2903420 . *2903410 . *2903400
5′ a c c c g a t t a a c g a g a t g t t c c a g a c c c t g c a a g g t g a g 3′
. *2903390 . *2903380 . *2903370 . *2903360
5′ g g t t a c t t t a c c g g c g t t c c c g c c a t t t t t a t t c g t t t 3′
proximalUP 6.2 bits
|--------- ... proximalUP-(42)
distalUP 5.7 bits
|------------------------------ ... distalUP-(52)-s
|-------------------|--------- ... sigma38p10-prox
. *2903350 . *2903340 . *2903330 . *2903320
5′ a c a g g g a t g c c c g g t t g g c t g t g c c t g g t g c g a c a [illegible] [illegible] [illegible] 3′
... sigma38p10

```
... ----------------------------------------------------------------------| proximalUP-(42)-sigm
                                                                          |
... ----------------------------------------------------------------------| distalUP-(52)-sigma3
... ----------------------------------------------------------------------| sigma38p10-proximalU

             .         *2903310   .         *2903300
 5' a a c a c a c c t g g g a a a a g c t t 3'
```

... 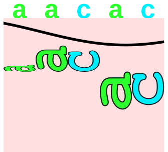 sigma38p10 1.4 bits piece 10, NC_000913, tisB-istR-1-istR-2, config: linear, direction: -, begin: 3851449, end: 385105

. *3851440 . *3851430 . *3851420 .

5′ g c g g g g t t a a g g c g c t g a c g g c a c c a c c c g t t t c a g c c 3′

T timestamp:_2018Sep20_11-36-13

*3851410 . *3851400 . *3851390 . *3851380 .

5′ a g g a c t t a g t g c g c c g g g t a a c g a t g c g g c c c c c g c a t 3′

... proximalUP

... distalUP

... distalUP-(41)-s

... sigma38p10-prox

*3851370 . *3851360 . *3851350 . *3851340

5′ a a c a c a t t g c g t a c a g t g a t a t t a t a g a a g t t t a c t g t 3′

... proximalUP 1.4 bits distalUP 6.5 bits ... sigma38p10

... distalUP 2.5 bits proximalUP 3.9 bits

... distalUP-(41)-s

... sigma38p10-prox

... proximalUP-(36)

... proximalUP-(43)

... distalUP-(52)-s

... sigma38p10-prox

... distalUP

... distalUP-(40)-s

... sigma38p10-prox

... distalUP

... distalUP-(45)-s

... sigma38p10-prox

. *3851330 . *3851320 . *3851310 . *3851300

5′ a t a a a t a a a c a g t a t a t t t g g a c a a a a c g c a a a c t g t 3′

... sigma38p10 4.5 bits sigma38p10 2.2 bits

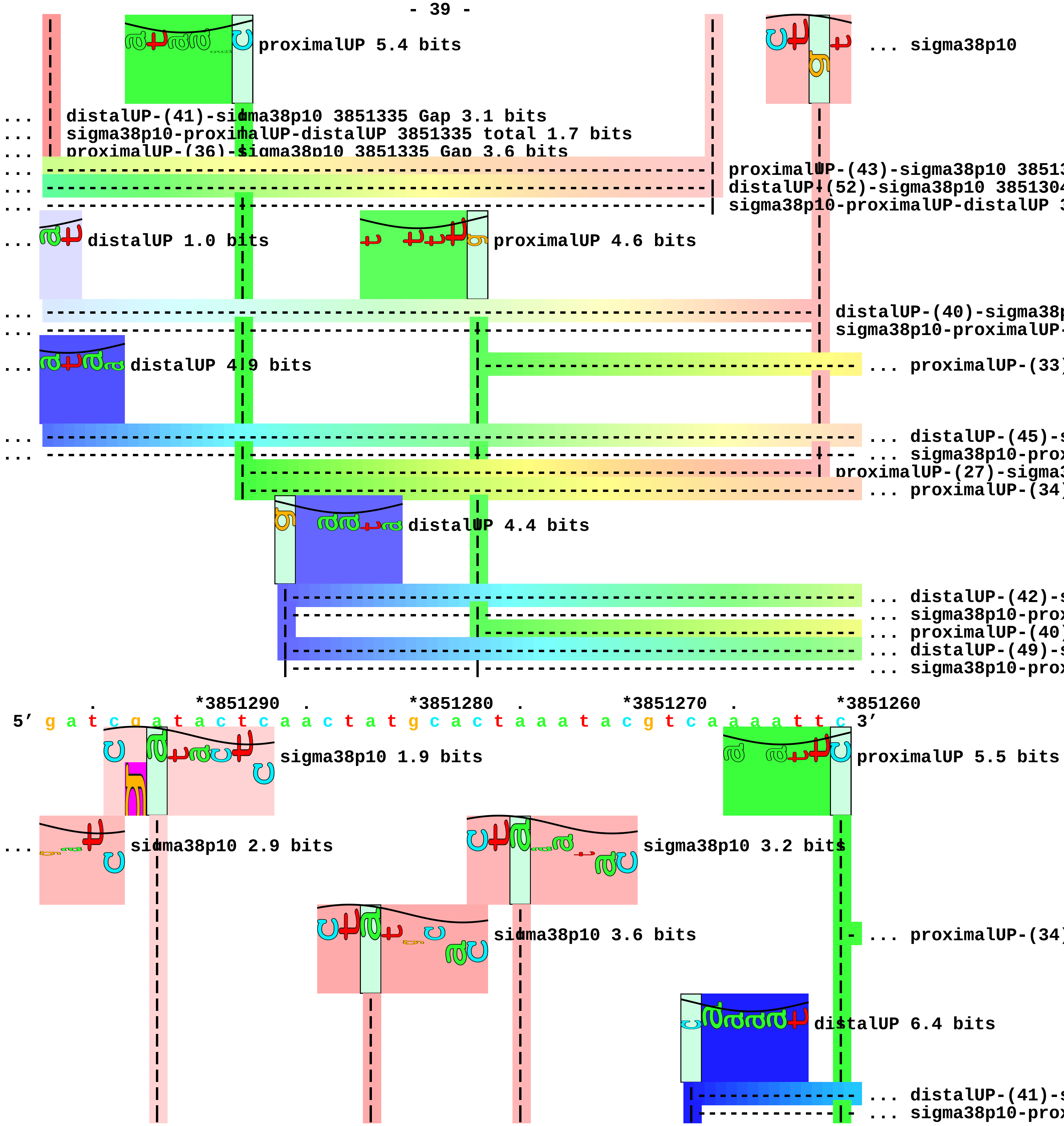
proximalUP 5.4 bits
... sigma38p10
... distalUP-(41)-sigma38p10 3851335 Gap 3.1 bits
... sigma38p10-proximalUP-distalUP 3851335 total 1.7 bits
... proximalUP-(36)-sigma38p10 3851335 Gap 3.6 bits
proximalUP-(43)-sigma38p10 38513
distalUP-(52)-sigma38p10 385130
sigma38p10-proximalUP-distalUP 3
... distalUP 1.0 bits
proximalUP 4.6 bits
distalUP-(40)-sigma38p
sigma38p10-proximalUP-
... distalUP 4.9 bits
... proximalUP-(33)
... distalUP-(45)-s
... sigma38p10-prox
proximalUP-(27)-sigma3
... proximalUP-(34)
distalUP 4.4 bits
... distalUP-(42)-s
... sigma38p10-prox
... proximalUP-(40)
... distalUP-(49)-s
... sigma38p10-prox
. *3851290 . *3851280 . *3851270 . *3851260
5’ g a t c g a t a c t c a a c t a t g c a c t a a a t a c g t c a a a a t t c 3’
sigma38p10 1.9 bits
proximalUP 5.5 bits
... sigma38p10 2.9 bits
sigma38p10 3.2 bits
sigma38p10 3.6 bits
... proximalUP-(34)
distalUP 6.4 bits
... distalUP-(41)-s
... sigma38p10-prox

```
                                                                                  |            |- ... proximalUP-(36)
                                                                                  |--------------- ... distalUP-(43)-s
                                                                                  |------------|- ... sigma38p10-prox
... ------------------------------| proximalUP-(33)-sigma38p10 3851282 Gap 4.0 bits
... ----------| distalUP-(45)-sigma38p10 3851292 Gap 5.5 bits
... ----------| sigma38p10-proximalUP-distalUP 3851292 total 4.3 bits
... ----------| proximalUP-(34)-sigma38p10 3851292 Gap 2.5 bits
... ------------------------------| distalUP-(42)-sigma38p10 3851282 Gap 3.3 bits
... ------------------------------| sigma38p10-proximalUP-distalUP 3851282 total 5.3 bits
... ---------------------------------------------| proximalUP-(40)-sigma38p10 3851275 Gap 4.0 bits
... ---------------------------------------------| distalUP-(49)-sigma38p10 3851275 Gap 5.9 bits
... ---------------------------------------------| sigma38p10-proximalUP-distalUP 3851275 total 2.2 b

              .         *3851250   .         *3851240   .         *3851230   .
5' g t g c c g a a a t t g c g c g t t c t g c g c g g a a c a c g t a t a c t 3'
                                                                            ... sigma38p10
                                                                            ... sigma38p10
... -----------------------------------------------------------------------| proximalUP-(34)-sigma38p10 3
... -----------------------------------------------------------------------| distalUP-(41)-sigma38p10 385
... -----------------------------------------------------------------------| sigma38p10-proximalUP-dista
... ---------------------------------------------------------------------------| proximalUP-(36)-sigma38p
... ---------------------------------------------------------------------------| distalUP-(43)-sigma38p10
... ---------------------------------------------------------------------------| sigma38p10-proximalUP-di

     *3851220   .         *3851210   .         *3851200   .         *3851190   .
5' t t c a g t g t t g a c a t a a t a c a g t g t g c t t t g c g g t t a c c 3'
... sigma38p10 8.3 bits
... sigma38p10 2.4 bits

       *3851180   .         *3851170   .         *3851160   .         *3851150
5' a g c c g c a g g c g a c t g a c g a a a c c t c g c t c c g g c g g g g t 3'
                                                                            ... distalUP
                                                                        |--- ... distalUP-(44)-s
                                                                        |--- ... sigma38p10-prox
                                                                        |--- ... distalUP-(45)-s
                                                                        |--- ... sigma38p10-prox
```

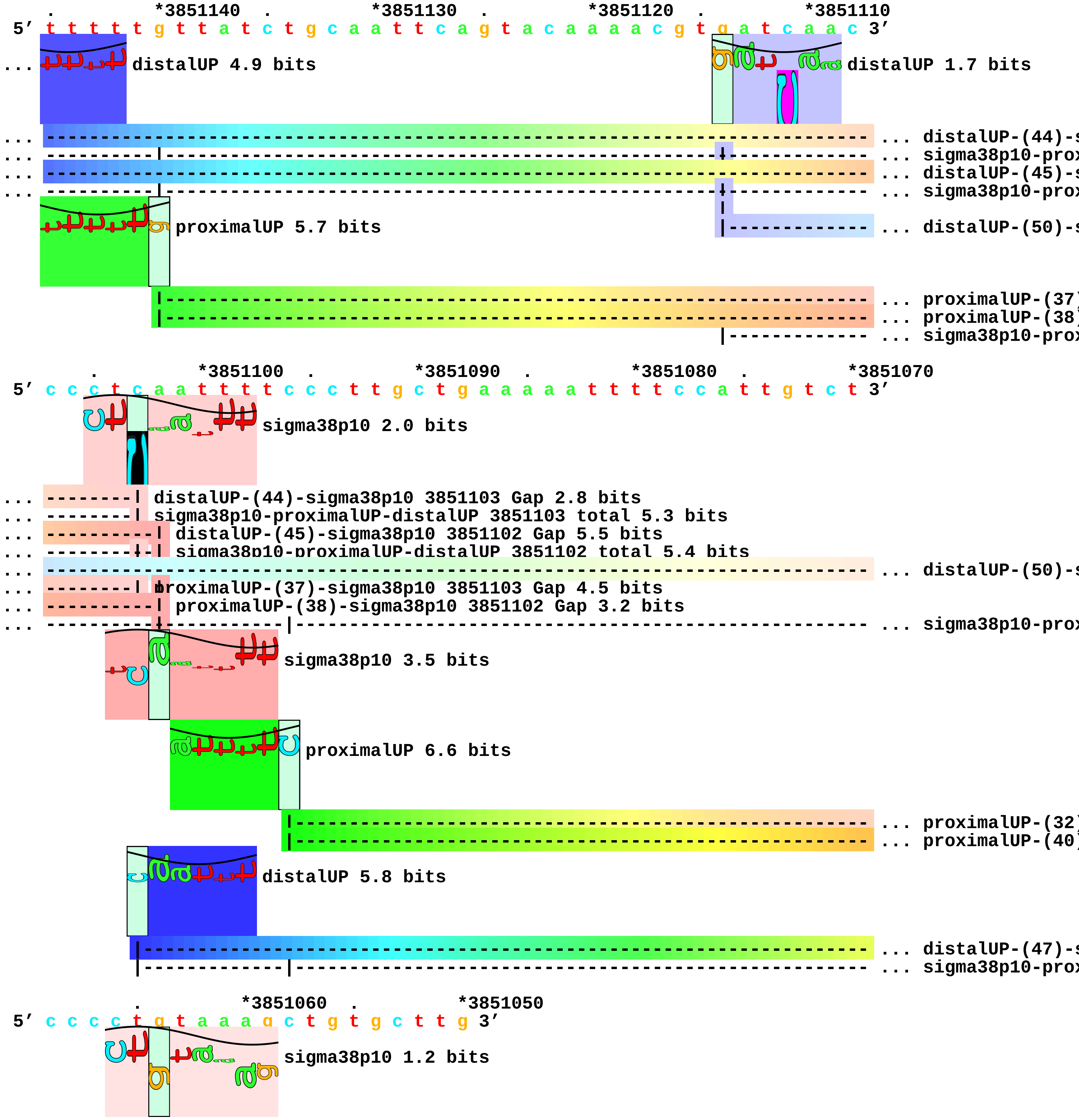
. *3851140 . *3851130 . *3851120 . *3851110
5' t t t t t g t t a t c t g c a a t t c a g t a c a a a a c g t g a t c a a c 3'
... distalUP 4.9 bits
distalUP 1.7 bits
... distalUP-(44)-s
... sigma38p10-prox
... distalUP-(45)-s
... sigma38p10-prox
proximalUP 5.7 bits
... distalUP-(50)-s
... proximalUP-(37)
... proximalUP-(38)
... sigma38p10-prox
. *3851100 . *3851090 . *3851080 . *3851070
5' c c c t c a a t t t t c c c t t g c t g a a a a a t t t t c c a t t g t c t 3'
sigma38p10 2.0 bits
... distalUP-(44)-sigma38p10 3851103 Gap 2.8 bits
... sigma38p10-proximalUP-distalUP 3851103 total 5.3 bits
... distalUP-(45)-sigma38p10 3851102 Gap 5.5 bits
... sigma38p10-proximalUP-distalUP 3851102 total 5.4 bits
... distalUP-(50)-s
... proximalUP-(37)-sigma38p10 3851103 Gap 4.5 bits
... proximalUP-(38)-sigma38p10 3851102 Gap 3.2 bits
... sigma38p10-prox
sigma38p10 3.5 bits
proximalUP 6.6 bits
... proximalUP-(32)
... proximalUP-(40)
distalUP 5.8 bits
... distalUP-(47)-s
... sigma38p10-prox
. *3851060 . *3851050
5' c c c c t g t a a a g c t g t g c t t g 3'
sigma38p10 1.2 bits

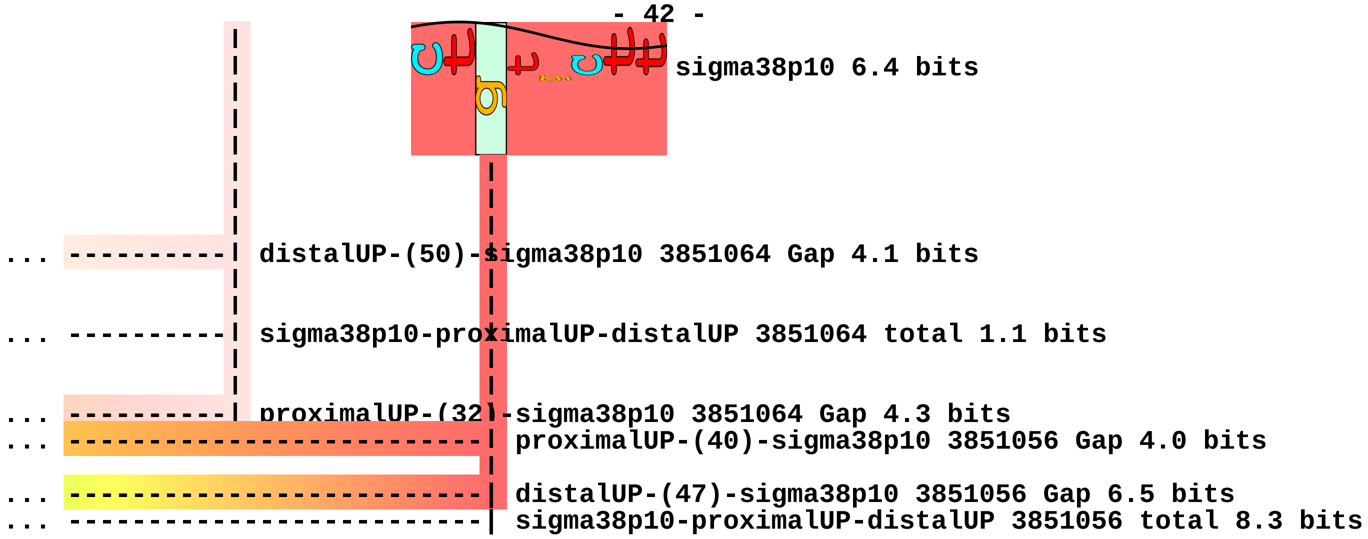
sigma38p10 6.4 bits
distalUP-(50)-sigma38p10 3851064 Gap 4.1 bits
sigma38p10-proximalUP-distalUP 3851064 total 1.1 bits
proximalUP-(32)-sigma38p10 3851064 Gap 4.3 bits
proximalUP-(40)-sigma38p10 3851056 Gap 4.0 bits
distalUP-(47)-sigma38p10 3851056 Gap 6.5 bits
sigma38p10-proximalUP-distalUP 3851056 total 8.3 bits

piece 11, NC_000913, rpmE-priA, config: linear, direction: -, begin: 4125099, end: 4124800

. *4125090 . *4125080 . *4125070 .
5′ t t a c g t t a c c g c a a g a g c a g c t a g c a g t a a t t t c t t c g 3′
T timestamp:_2018Sep20_11-36-13

*4125060 . *4125050 . *4125040 . *4125030 .
5′ t a t t t c g g g t g a a t a t c t t t t t t c a t g g g a a a a c c t c g 3′

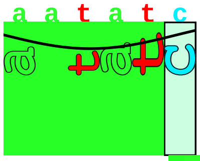

proximalUP 6.1 bits

|---------------------------------------- ... proximalUP-(37)

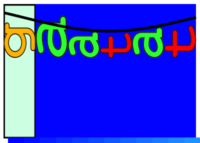

distalUP 7.2 bits

|------------------------------------------------------ ... distalUP-(43)-s
|-----------|----------------------------------------- ... sigma38p10-prox

*4125020 . *4125010 . *4125000 . *4124990
5′ g t t t a a g g c c g c g t c g c t c t t c c a g c c c t a a c g c c a g a 3′

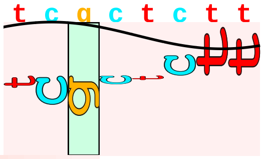

sigma38p10 0.7 bits

... ------------------------------| proximalUP-(37)-sigma38p10 4125008 Gap 4.5 bits
... ------------------------------| distalUP-(43)-sigma38p10 4125008 Gap 3.6 bits
... ------------------------------| sigma38p10-proximalUP-distalUP 4125008 total 5.9 bits

. *4124980 . *4124970 . *4124960 . *4124950
5′ c a c c a c g c g a t g t t a a a a g t a t a g c t t c a a t a c g a t c a 3′

. *4124940 . *4124930 . *4124920 . *4124910
5′ t t t c g t a c g a a g g c g c g a a a t c a t a c a g a a a t t a a c c a 3′

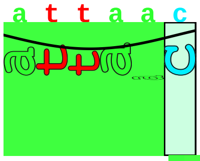

proximalUP 5.4 bits

|----- ... proximalUP-(43)

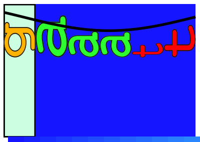

distalUP 6.6 bits

|---------------------- ... distalUP-(51)-s
|---------------|----- ... sigma38p10-prox

. *4124900 . *4124890 . *4124880 .
5′ g c g t a t g c a a a c t g a t c c g c a c t c t t c t a c g g c a a t g t 3′

... ------------------------------------------------------------------------ ... proximalUP-(43)
... ------------------------------------------------------------------------ ... distalUP-(51)-s
... ------------------------------------------------------------------------ ... sigma38p10-prox

```
     *4124870  .         *4124860  .         *4124850  .         *4124840  .
5' g t a t a c t a a c c c a c c g a a t t t c a a g t c a g g a t g a t g c t 3'
```

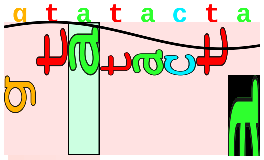

sigma38p10 1.2 bits

```
... ----| proximalUP-(43)-sigma38p10 4124869 Gap 5.1 bits
        |
... ----| distalUP-(51)-sigma38p10 4124869 Gap 5.1 bits
... ----| sigma38p10-proximalUP-distalUP 4124869 total 3.0 bits
```

```
         *4124830  .         *4124820  .         *4124810  .         *4124800
5' a t g c c c g t t g c c c a c g t t g c c t t g c c c g t t c c g c 3'
```

piece 12, NC_000913, yjfP-bsmA, config: linear, direction: -, begin: 4414949, end: 4414600

```
             .         *4414940   .         *4414930   .         *4414920   .
5' a c a t g a a g a a a t c c t t a g c a c a a g t t t c t g c t g g c g g t 3'
```

T timestamp:_2018Sep20_11-36-13

proximalUP 5.0 bits

... proximalUP-(39)

distalUP 2.3 bits

... distalUP-(43)-s

... sigma38p10-prox

... proximalUP

... distalUP

... distalUP-(41)-s

... sigma38p10-prox

```
   *4414910   .         *4414900   .         *4414890   .         *4414880   .
5' a t t a g t t c a t a a t g t t g a g a t g t g g g t t a c g c t t t c g t 3'
```

[---------#----------------------------------------> bsmA

... sigma38p10

... proximalUP-(39)-sigma38p10 4414883 Gap

proximalUP 6.7 bits

sigma38p10 6.8 bits

... distalUP-(43)-sigma38p10 4414883 Gap 3

... sigma38p10-proximalUP-distalUP 441488

... proximalUP 2.8 bits

... distalUP 4.3 bits

... proximalUP-(39)

... distalUP-(41)-s

... sigma38p10-prox

... proximalUP-(35)

distalUP 6.3 bits

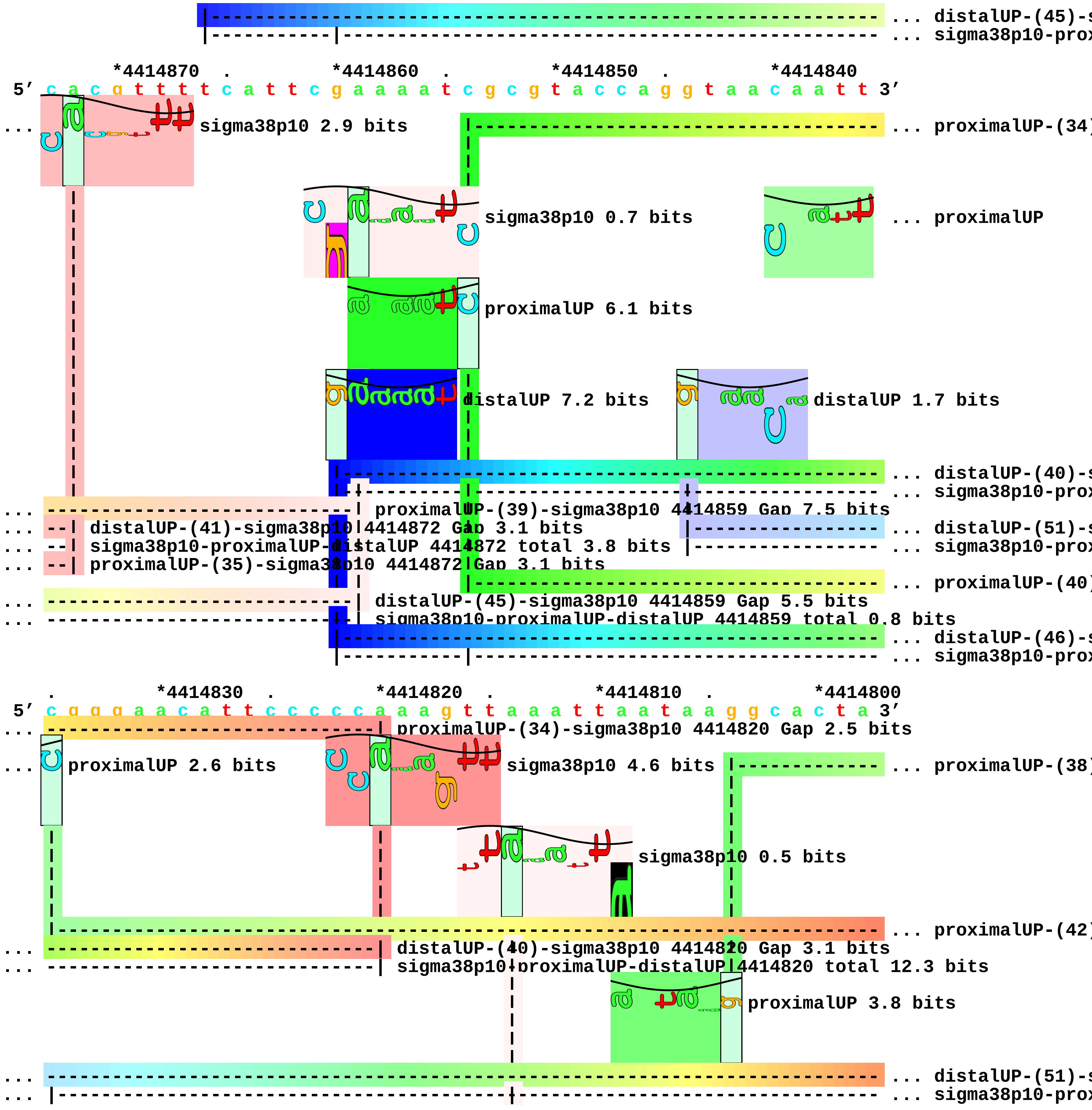
... distalUP-(45)-s
... sigma38p10-prox
*4414870 . *4414860 . *4414850 . *4414840
5′ c a c g t t t t c a t t c g a a a a t c g c g t a c c a g g t a a c a a t t 3′
sigma38p10 2.9 bits
... proximalUP-(34)
sigma38p10 0.7 bits
... proximalUP
proximalUP 6.1 bits
distalUP 7.2 bits
distalUP 1.7 bits
... distalUP-(40)-s
... sigma38p10-prox
proximalUP-(39)-sigma38p10 4414859 Gap 7.5 bits
distalUP-(41)-sigma38p10 4414872 Gap 3.1 bits
... distalUP-(51)-s
sigma38p10-proximalUP-distalUP 4414872 total 3.8 bits
... sigma38p10-prox
proximalUP-(35)-sigma38p10 4414872 Gap 3.1 bits
... proximalUP-(40)
distalUP-(45)-sigma38p10 4414859 Gap 5.5 bits
sigma38p10-proximalUP-distalUP 4414859 total 0.8 bits
... distalUP-(46)-s
... sigma38p10-prox
. *4414830 . *4414820 . *4414810 . *4414800
5′ c g g g a a c a t t c c c c c a a a g t t a a a t t a a t a a g g c a c t a 3′
proximalUP-(34)-sigma38p10 4414820 Gap 2.5 bits
proximalUP 2.6 bits
sigma38p10 4.6 bits
... proximalUP-(38)
sigma38p10 0.5 bits
... proximalUP-(42)
distalUP-(40)-sigma38p10 4414820 Gap 3.1 bits
sigma38p10-proximalUP-distalUP 4414820 total 12.3 bits
proximalUP 3.8 bits
... distalUP-(51)-s
... sigma38p10-prox

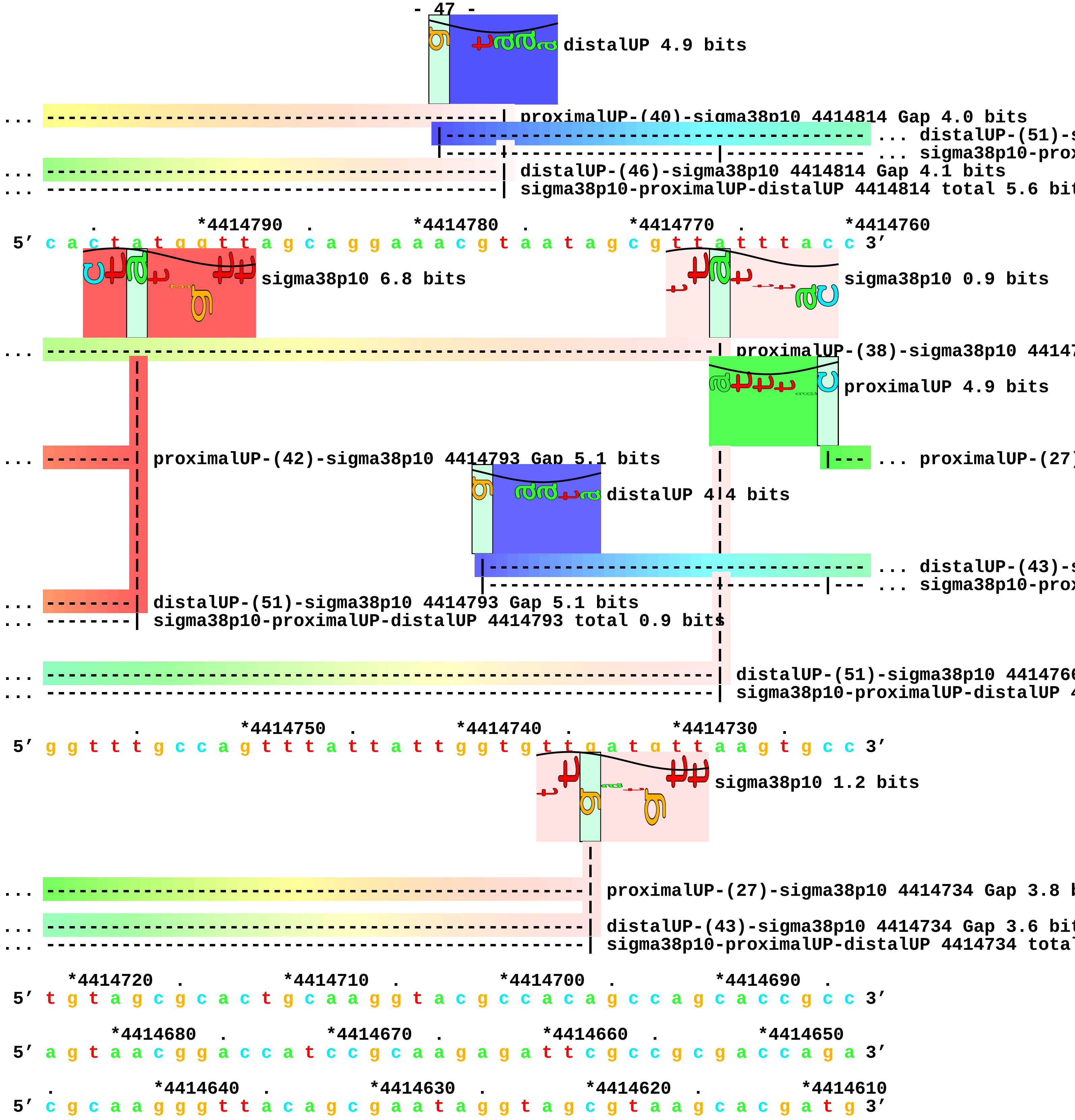
distalUP 4.9 bits
... ----------------------------------------| proximalUP-(40)-sigma38p10 4414814 Gap 4.0 bits
|----------------------------------------- ... distalUP-(51)-s
|------------------------|------------- ... sigma38p10-prox
... ----------------------------------------| distalUP-(46)-sigma38p10 4414814 Gap 4.1 bits
... ----------------------------------------| sigma38p10-proximalUP-distalUP 4414814 total 5.6 bit
. *4414790 . *4414780 . *4414770 . *4414760
5′ c a c t a t g g t t a g c a g g a a a c g t a a t a g c g t t a t t t a c c 3′
sigma38p10 6.8 bits
sigma38p10 0.9 bits
... -------------------------------------------------------------| proximalUP-(38)-sigma38p10 44147
proximalUP 4.9 bits
... --------| proximalUP-(42)-sigma38p10 4414793 Gap 5.1 bits
|--- ... proximalUP-(27)
distalUP 4.4 bits
|------------------------------------- ... distalUP-(43)-s
|-------------------------------|--- ... sigma38p10-prox
... --------| distalUP-(51)-sigma38p10 4414793 Gap 5.1 bits
... --------| sigma38p10-proximalUP-distalUP 4414793 total 0.9 bits
... -------------------------------------------------------------| distalUP-(51)-sigma38p10 4414760
... -------------------------------------------------------------| sigma38p10-proximalUP-distalUP 4
. *4414750 . *4414740 . *4414730 .
5′ g g t t t g c c a g t t t a t t a t t g g t g t t g a t g t t a a g t g c c 3′
sigma38p10 1.2 bits
... --------------------------------------------------| proximalUP-(27)-sigma38p10 4414734 Gap 3.8 b
... --------------------------------------------------| distalUP-(43)-sigma38p10 4414734 Gap 3.6 bit
... --------------------------------------------------| sigma38p10-proximalUP-distalUP 4414734 total
*4414720 . *4414710 . *4414700 . *4414690 .
5′ t g t a g c g c a c t g c a a g g t a c g c c a c a g c c a g c a c c g c c 3′
*4414680 . *4414670 . *4414660 . *4414650
5′ a g t a a c g g a c c a t c c g c a a g a g a t t c g c c g c g a c c a g a 3′
. *4414640 . *4414630 . *4414620 . *4414610
5′ c g c a a g g g t t a c a g c g a a t a g g t a g c g t a a g c a c g a t g 3′

```
       .         *4414600
5’ g t t c g g g g 3’
```

piece 13, NC_000913, cysQ-cpdB, config: linear, direction: -, begin: 4434799, end: 4434350

```
              .          *4434790  .          *4434780  .          *4434770  .
5' g c t g g c a t a c t t g a t c t a a c a t t t c t c c a c c t c g t c t c 3'
```

T timestamp:_2018Sep20_11-36-13

proximalUP 2.6 bits

|-------------------------- ... proximalUP-(32)

distalUP 1.1 bits

|---------------------------------------------------- ... distalUP-(44)-

|----------------------|------------------------- ... sigma38p10-prox

```
    *4434760  .          *4434750  .          *4434740  .          *4434730  .
5' t g t g a g c g g t g t t a a c t t a t t g t t t t a c t t a t a c c c t a 3'
```

sigma38p10 5.2 bits

... ----------------------------------| proximalUP-(32)-sigma38p10 4434743 Gap 4.3 bits

proximalUP 4.4 bits

... ----------------------------------| distalUP-(44)-sigma38p10 4434743 Gap 2.8 bits

... ----------------------------------| sigma38p10-proximalUP-distalUP 4434743 total 1.8 bits

|--------- ... proximalUP-(37)

distalUP 3.5 bits

|--------------------- ... distalUP-(43)-

|-----------|--------- ... sigma38p10-prox

|--------------------- ... distalUP-(46)-

|--------------------- ... sigma38p10-prox

```
      *4434720  .          *4434710  .          *4434700  .          *4434690
5' t c g t t a a t g a a t g c g c c a a c t g t g a t a g t g t c a t c a t t 3'
```

proximalUP 6.3 bits

sigma38p10 2.9 bits

distalUP 5.5 bits

... sigma38p10

|------------------------------------------------------| proximalUP-(27)-sigma38p

|------------------------------------------------------- ... proximalUP-(40)

|------------------------------------------------------- ... distalUP-(46)-

... ------------------------------------------------------| proximalUP-(37)-sigma38p10 443

|-----------|-------------------------------------------|-----|----- ... sigma38p10-prox

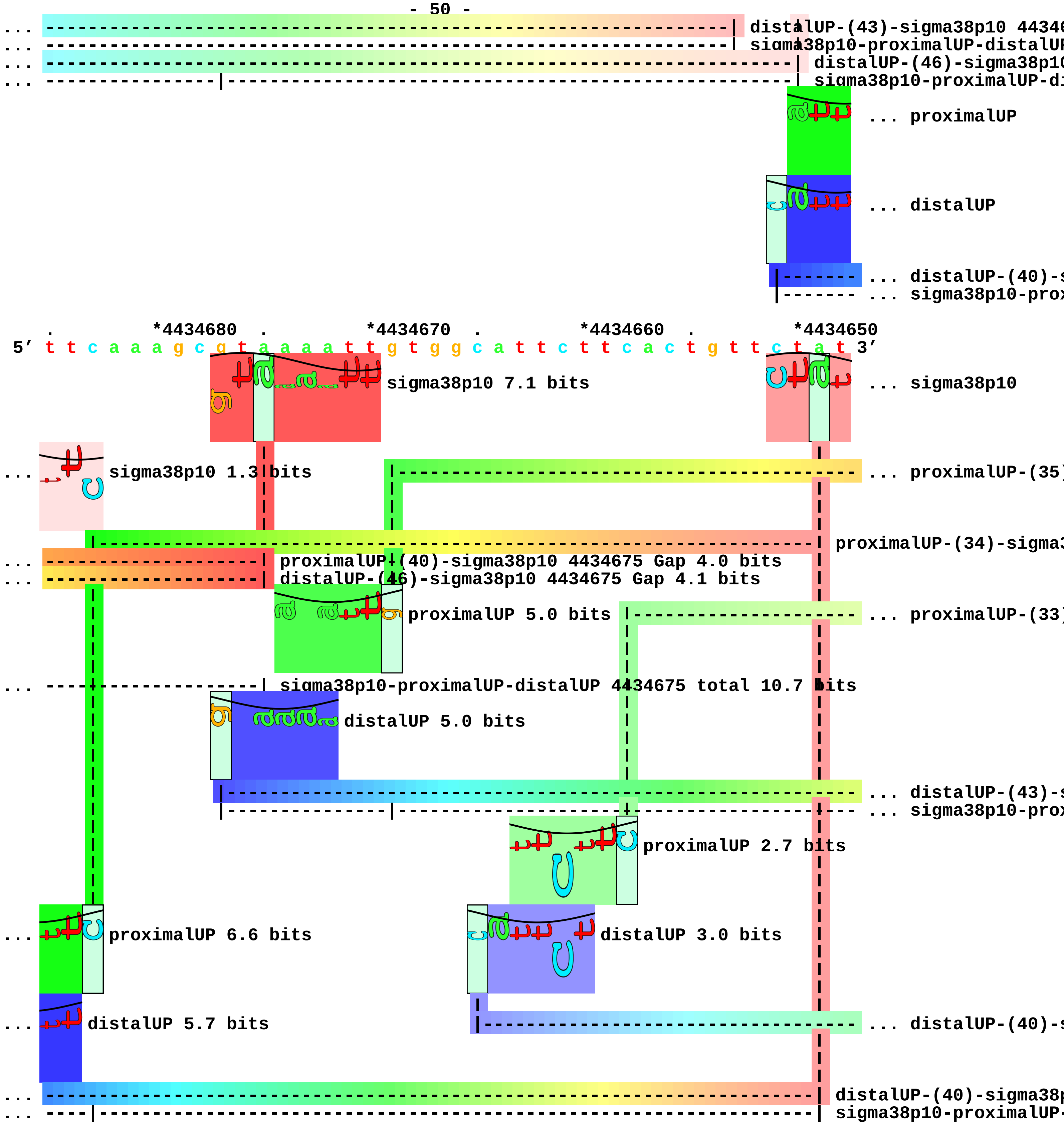
distalUP-(43)-sigma38p10 4434
sigma38p10-proximalUP-distalUP
distalUP-(46)-sigma38p10
sigma38p10-proximalUP-di
... proximalUP
... distalUP
... distalUP-(40)-s
... sigma38p10-prox
*4434680
*4434670
*4434660
*4434650
5′ t t c a a a g c g t a a a a t t g t g g c a t t c t t c a c t g t t c t a t 3′
sigma38p10 7.1 bits
... sigma38p10
sigma38p10 1.3 bits
... proximalUP-(35)
proximalUP-(34)-sigma3
proximalUP-(40)-sigma38p10 4434675 Gap 4.0 bits
distalUP-(46)-sigma38p10 4434675 Gap 4.1 bits
proximalUP 5.0 bits
... proximalUP-(33)
sigma38p10-proximalUP-distalUP 4434675 total 10.7 bits
distalUP 5.0 bits
... distalUP-(43)-s
... sigma38p10-prox
proximalUP 2.7 bits
proximalUP 6.6 bits
distalUP 3.0 bits
distalUP 5.7 bits
... distalUP-(40)-s
distalUP-(40)-sigma38p
sigma38p10-proximalUP-

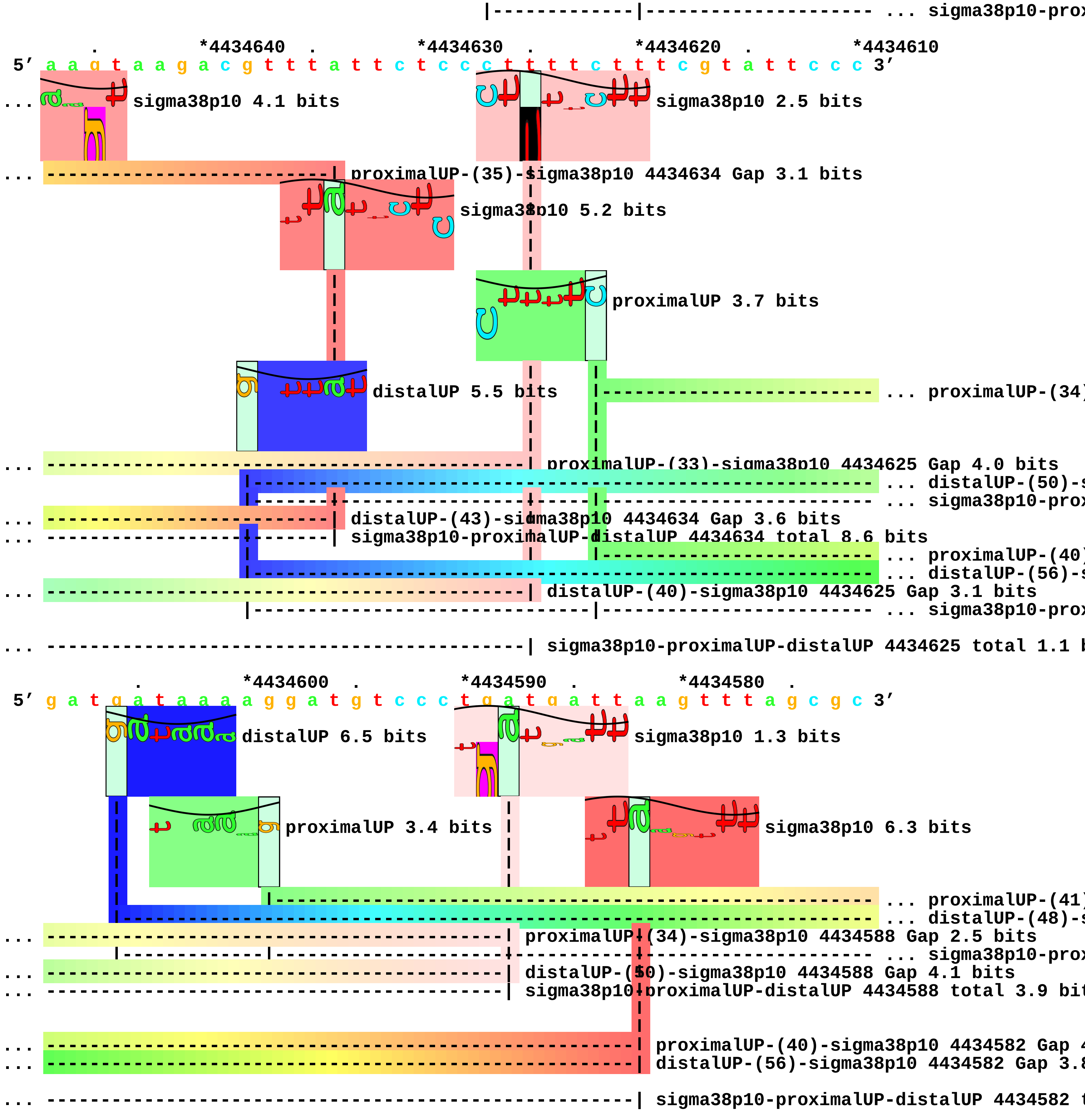
... sigma38p10-prox
*4434640
*4434630
*4434620
*4434610
5′ a a g t a a g a c g t t t a t t c t c c c t t t t c t t t c g t a t t c c c 3′
sigma38p10 4.1 bits
sigma38p10 2.5 bits
proximalUP-(35)-sigma38p10 4434634 Gap 3.1 bits
sigma38p10 5.2 bits
proximalUP 3.7 bits
distalUP 5.5 bits
... proximalUP-(34)
proximalUP-(33)-sigma38p10 4434625 Gap 4.0 bits
... distalUP-(50)-s
... sigma38p10-prox
distalUP-(43)-sigma38p10 4434634 Gap 3.6 bits
sigma38p10-proximalUP-distalUP 4434634 total 8.6 bits
... proximalUP-(40)
... distalUP-(56)-s
distalUP-(40)-sigma38p10 4434625 Gap 3.1 bits
... sigma38p10-prox
sigma38p10-proximalUP-distalUP 4434625 total 1.1 b
*4434600
*4434590
*4434580
5′ g a t g a t a a a a g g a t g t c c c t g a t g a t t a a g t t t a g c g c 3′
distalUP 6.5 bits
sigma38p10 1.3 bits
proximalUP 3.4 bits
sigma38p10 6.3 bits
... proximalUP-(41)
... distalUP-(48)-s
proximalUP-(34)-sigma38p10 4434588 Gap 2.5 bits
... sigma38p10-prox
distalUP-(50)-sigma38p10 4434588 Gap 4.1 bits
sigma38p10-proximalUP-distalUP 4434588 total 3.9 bit
proximalUP-(40)-sigma38p10 4434582 Gap 4
distalUP-(56)-sigma38p10 4434582 Gap 3.8
sigma38p10-proximalUP-distalUP 4434582 t

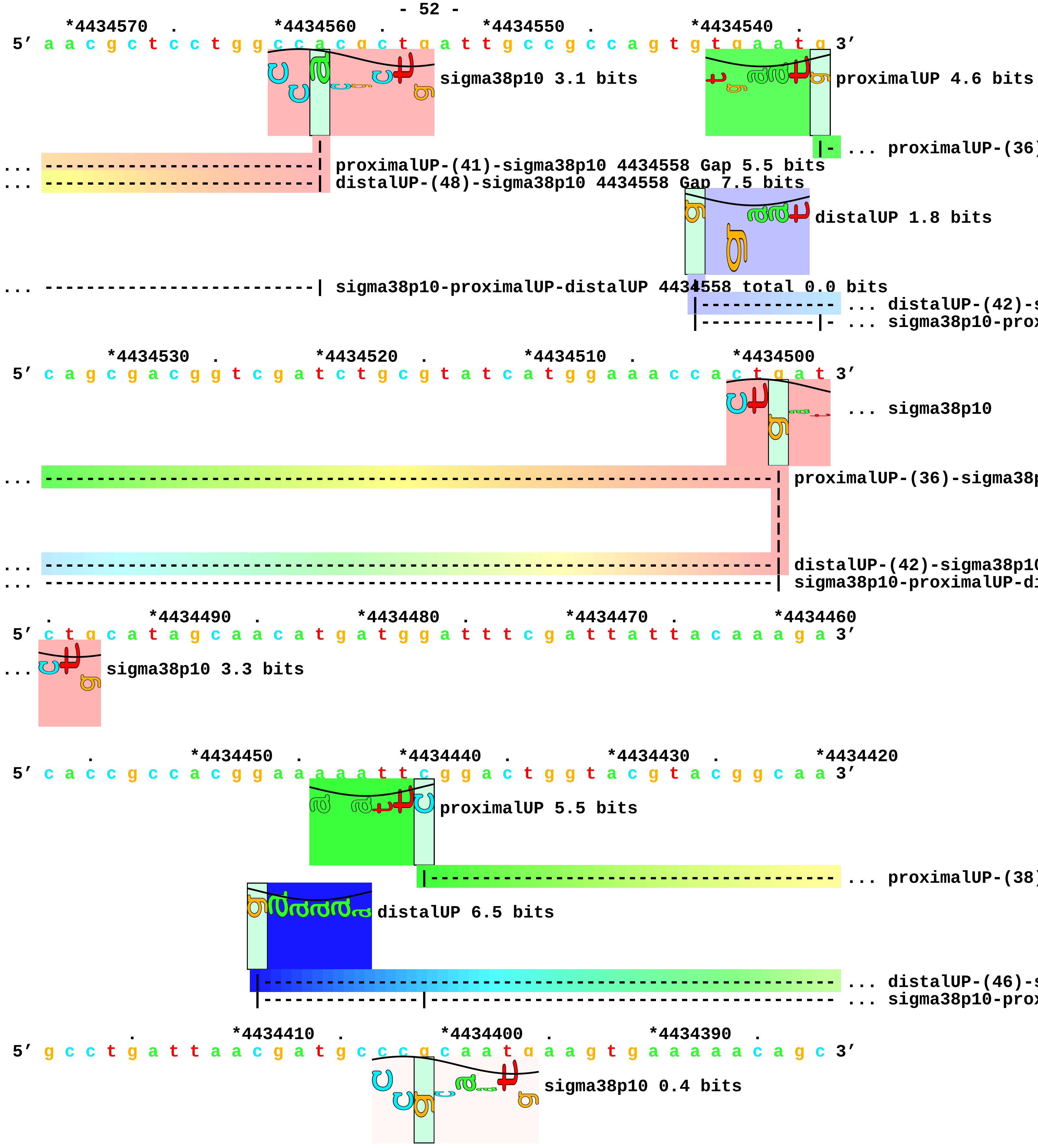
*4434570 . *4434560 . *4434550 . *4434540 .
5' a a c g c t c c t g g c c a c g c t g a t t g c c g c c a g t g t g a a t g 3'
sigma38p10 3.1 bits
proximalUP 4.6 bits
... proximalUP-(36)
... proximalUP-(41)-sigma38p10 4434558 Gap 5.5 bits
... distalUP-(48)-sigma38p10 4434558 Gap 7.5 bits
distalUP 1.8 bits
... sigma38p10-proximalUP-distalUP 4434558 total 0.0 bits
... distalUP-(42)-s
... sigma38p10-prox
*4434530 . *4434520 . *4434510 . *4434500
5' c a g c g a c g g t c g a t c t g c g t a t c a t g g a a a c c a c t g a t 3'
... sigma38p10
... proximalUP-(36)-sigma38p
... distalUP-(42)-sigma38p10
... sigma38p10-proximalUP-di
. *4434490 . *4434480 . *4434470 . *4434460
5' c t g c a t a g c a a c a t g a t g g a t t t c g a t t a t t a c a a a g a 3'
... sigma38p10 3.3 bits
. *4434450 . *4434440 . *4434430 . *4434420
5' c a c c g c c a c g g a a a a a t t c g g a c t g g t a c g t a c g g c a a 3'
proximalUP 5.5 bits
... proximalUP-(38)
distalUP 6.5 bits
... distalUP-(46)-s
... sigma38p10-prox
. *4434410 . *4434400 . *4434390 .
5' g c c t g a t t a a c g a t g c c c g c a a t g a a g t g a a a a a c a g c 3'
sigma38p10 0.4 bits

```
... ------------------------------------| proximalUP-(38)-sigma38p10 4434401 Gap 3.2 bits
                                        |
... ------------------------------------| distalUP-(46)-sigma38p10 4434401 Gap 4.1 bits
... ------------------------------------| sigma38p10-proximalUP-distalUP 4434401 total 5.1 bits

     *4434380  .         *4434370  .         *4434360  .         *4434350
 5' g t a c t g g t t g a t a a c g g c g a t t t g a t t c a g g g 3'
```
