## Supplement_for_sigma38_paper-directory for "*Escherichia coli σ*^38^ promoters use two UP elements instead of a −35 element: resolution of a paradox and discovery that *σ*^38^ transcribes ribosomal promoters": Peano.Landini2015-Table2-pos-map.pdf

piece 1, NC_000913, ychH-pth, config: linear, direction: +, begin: 1257700, end: 1258249

```
   *1257700  .         *1257710  .         *1257720  .         *1257730  .
5' c g g g g t t c g c c a g g c c g a c a a t c a a t t t a a t c g t c a c g 3'
   T timestamp:_2018Sep20_11-36-03

     *1257740  .         *1257750  .         *1257760  .         *1257770  .
5' t t t t t t t g t c c t g a g t g t g t a c a t a a c t g g c g c g t a g t 3'

       *1257780  .         *1257790  .         *1257800  .         *1257810
5' t t a c t g g t t g c g g c c c c g c t t g a c a a a a a a c t g c g t a t 3'

   .         *1257820  .         *1257830  .         *1257840  .         *1257850
5' c a a a t g c a g a t a a c g t a a t a a t t g c c t g a g t g g a c t a t 3'
```

proximalUP 4.6 bits ... proximalUP

... proximalUP-(41

distalUP 4.4 bits

... distalUP-(50)-s

... distalUP-proxim

... distalUP-(56)-s

... distalUP-proxim

```
         .         *1257860  .         *1257870  .         *1257880  .
5' t a g a a a g t c a a g g t g t t c a g g c g t t t a t t t g t a a a g t t 3'
```

... proximalUP 2.8 bits

sigma38p10 0.8 bits

... proximalUP-(41)-sigma38p10 1257878 Gap 5.5

distalUP 2.2 bits

sigma38p10 4.8 bits

... distalUP-(50)-sigma38p10 1257878 Gap 4.1 b

... distalUP-proximalUP-sigma38p10 1257878 tot

... distalUP-(56)-sigma38p10 12578

... distalUP-proximalUP-sigma38p10

proximalUP-(30)-sigma38p10 125

proximalUP 1.8 bits

... proximalUP-(27

... distalUP-(53)-s

... distalUP-proxim

proximalUP 4.6 bits

... proximalUP-(43

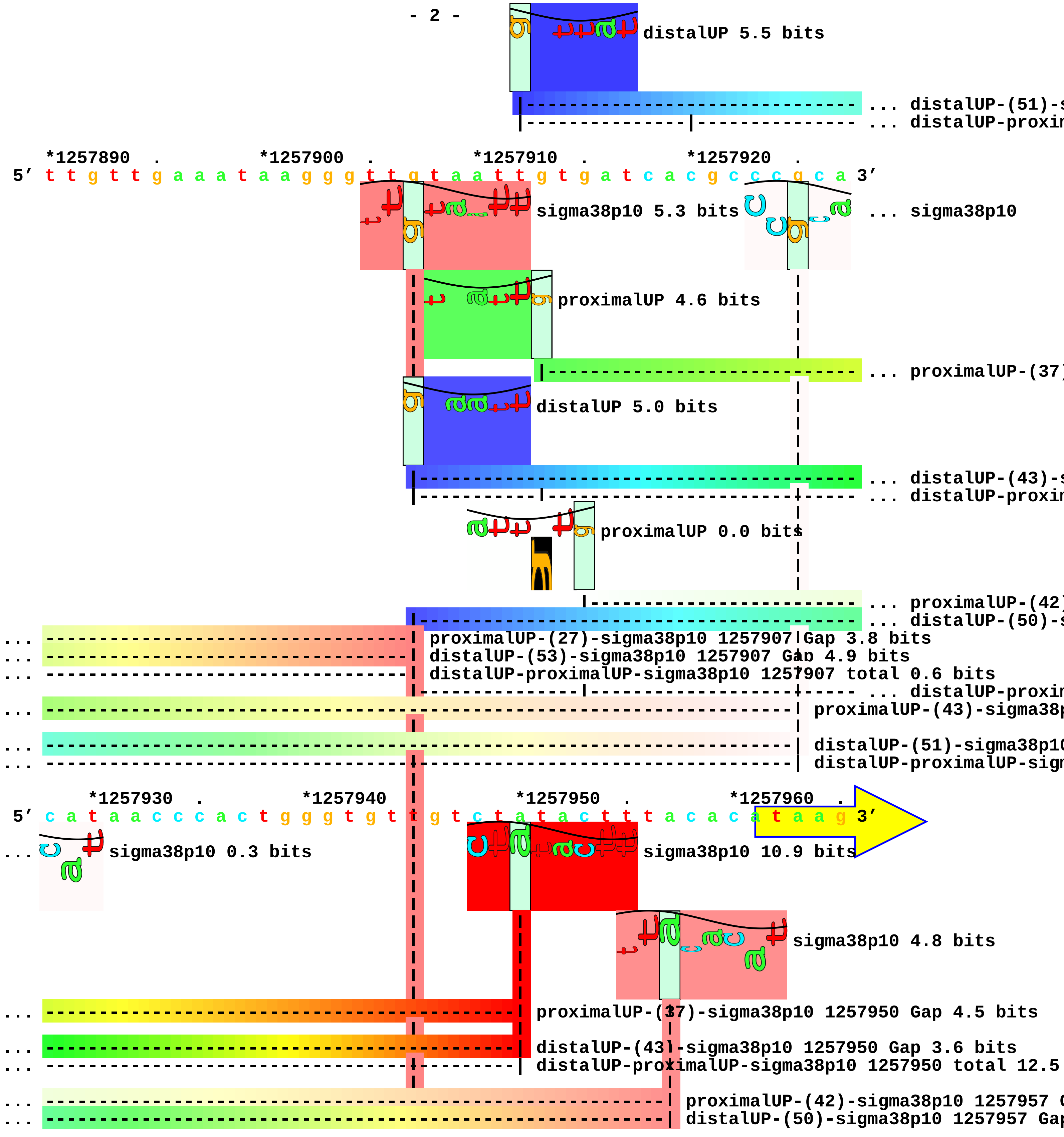
distalUP 5.5 bits
... distalUP-(51)-s
... distalUP-proxim
*1257890 . *1257900 . *1257910 . *1257920 .
5′ t t g t t g a a a t a a g g g t t g t a a t t g t g a t c a c g c c c g c a 3′
sigma38p10 5.3 bits
... sigma38p10
proximalUP 4.6 bits
... proximalUP-(37)
distalUP 5.0 bits
... distalUP-(43)-s
... distalUP-proxim
proximalUP 0.0 bits
... proximalUP-(42)
... distalUP-(50)-s
... proximalUP-(27)-sigma38p10 1257907 Gap 3.8 bits
... distalUP-(53)-sigma38p10 1257907 Gap 4.9 bits
... distalUP-proximalUP-sigma38p10 1257907 total 0.6 bits
... distalUP-proxim
... proximalUP-(43)-sigma38p
... distalUP-(51)-sigma38p10
... distalUP-proximalUP-sign
*1257930 . *1257940 . *1257950 . *1257960 .
5′ c a t a a c c c a c t g g g t g t t g t c t a t a c t t t a c a c a t a a g 3′
... sigma38p10 0.3 bits
sigma38p10 10.9 bits
sigma38p10 4.8 bits
... proximalUP-(37)-sigma38p10 1257950 Gap 4.5 bits
... distalUP-(43)-sigma38p10 1257950 Gap 3.6 bits
... distalUP-proximalUP-sigma38p10 1257950 total 12.5
... proximalUP-(42)-sigma38p10 1257957 G
... distalUP-(50)-sigma38p10 1257957 Gap

... ---------------------------------------------------------------------| distalUP-proximalUP-sigma38p10 12579

*1257970 . *1257980 . *1257990 . *1258000
5′ g a a g a g g g g t a t t c c c t g t t a c a a c c c a g a a a g t t c c g 3′

. *1258010 . *1258020 . *1258030 . *1258040
5′ g a g g t g a c a t a t g a a a c g c a a a a a c g c t t c g t t a c t c g 3′

. *1258050 . *1258060 . *1258070 .
5′ g t a a c g t g c t c a t g g g g t t g g g t c t g g t g g t a a t g g t g 3′

*1258080 . *1258090 . *1258100 . *1258110 .
5′ g t c g g c g t g g g g t a t t c a a t c c t c a a c c a g t t a c c a c a 3′

... distalUP

|--- ... distalUP-(41)-s
|--- ... distalUP-proxin

*1258120 . *1258130 . *1258140 . *1258150 .
5′ g t t t a a t a t g c c c c a g t a t t t c g c a c a t g g t g c a g t g c 3′

... distalUP 2.0 bits proximalUP 5.1 bits ... sigma38p10

... ------------------------------------------------------------------------ ... distalUP-(41)-s
... ------------------------------------------------------------------------ ... distalUP-proxin

proximalUP 5.6 bits |--------------------------------- ... proximalUP-(34)

|------------------------------------------------------------ ... proximalUP-(30)
|------------------------------------------------------------ ... proximalUP-(34)

distalUP 4.8 bits distalUP 5.0 bits

|------------------------------------------------------------------------ ... distalUP-(43)-s
|------------------|------------------------------------------------------ ... distalUP-proxin
|------------------------------------------------------------ ... distalUP-(40)-s
|------------|------------------------------------------------ ... distalUP-proxin
|------------------------------------------------ ... proximalUP-(37)
|------------------------------------------------------------ ... distalUP-(43)-s
|------------|------------------------------------------------ ... distalUP-proxin

*1258160 . *1258170 . *1258180 . *1258190
5′ t a a g t a t t t t c g t c g g t g c c a t t c t c t g g c t g g c g g g t 3′

... sigma38p10 4.8 bits sigma38p10 4.1 bits

... --| distalUP-(41)-sigma38p10 1258157 Gap 3.1 bits
... --| distalUP-proximalUP-sigma38p10 1258157 total 3.3 bits

```
... ------------------------------------| proximalUP-(34)-sigma38p10 1258173 Gap 2.5 bits
... --| proximalUP-(30)-sigma38p10 1258157 Gap 5.9 bits
... ----------| proximalUP-(34)-sigma38p10 1258161 Gap 2.5 bits
```

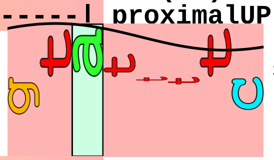

sigma38p10 3.2 bits

```
... ----------| distalUP-(43)-sigma38p10 1258161 Gap 3.6 bits
... ----------| distalUP-proximalUP-sigma38p10 1258161 total 7.6 bits
... ------------------------------------| distalUP-(40)-sigma38p10 1258173 Gap 3.1 bits
... ------------------------------------| distalUP-proximalUP-sigma38p10 1258173 total 4.8 bits
... ------------------------------------------| proximalUP-(37)-sigma38p10 1258176 Gap 4.5 bits
... ------------------------------------------| distalUP-(43)-sigma38p10 1258176 Gap 3.6 bits
... ------------------------------------------| distalUP-proximalUP-sigma38p10 1258176 total 6.1 bits
```

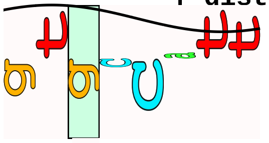

sigma38p10 0.2 bits

```
      .         *1258200   .         *1258210   .         *1258220   .         *1258230
5' g c c c g t g t t g g c g g g c a t g a a c a g g t g t g c g a c c g t t a 3'

          .         *1258240   .
5' c t g g t g g g t t c g c c a c t a 3'
```

```
piece 2, NC_000913, rssA-ychJ, config: linear, direction: +, begin: 1288200, end: 1288449

   *1288200   .         *1288210   .         *1288220   .         *1288230   .
5' g g t c t t t a t t a a a t a a t c t g c g t c t t g c a t c a c a a a a g 3'
                                                proximalUP 7.2 bits

T timestamp: 2018Sep20_11-36-03        |-------------------------------------- ... proximalUP-(40)
                  distalUP 4.7 bits

        |-------------------------------------------------------------------- ... distalUP-(54)-s
        |--------------------------|--------------------------------------- ... distalUP-proxin

     *1288240   .         *1288250   .         *1288260   .         *1288270   .
5' c g c a g t a a c g c g a a c g c a t g a g a t g t t c t g g a t c a g g t 3'
                                                       sigma38p10 0.4 bits
... ---------------------------------------| proximalUP-(40)-sigma38p10 1288257 Gap 4.0 bits
                                           |
... ---------------------------------------| distalUP-(54)-sigma38p10 1288257 Gap 4.9 bits
... ---------------------------------------| distalUP-proximalUP-sigma38p10 1288257 total 3.3 bits

       *1288280   .         *1288290   .         *1288300   .         *1288310
5' g c a a c c t t t t c a c c a g a c a c a t a a g g g t g g c a a c a t a g 3'

   .         *1288320   .         *1288330   .         *1288340   .         *1288350
5' g c t a t a c t c g a c a g c a c t a c c a c a g g g a c a a a g c t g a g 3'

     .         *1288360   .         *1288370   .         *1288380   .
5' a c a c a a a t a a t c t c c c t g g a a a c a a t a a c g g c g t a t t a 3'
                  distalUP 5.1 bits                         proximalUP 4.3 bits

        |                                                   |------------------ ... proximalUP-(27)
        |-------------------------------------------------------------------- ... distalUP-(52)-s
        |---------------------------------------------------|------------------ ... distalUP-proxin

   *1288390   .         *1288400   .         *1288410   .         *1288420   .
5' a c c g c c t g a g t a g c a c t a t g t t a a c c g a g c a g t a g c g a 3'
                                                    sigma38p10 1.6 bits

... -----------------------------------| proximalUP-(27)-sigma38p10 1288407 Gap 3.8 bits
... -----------------------------------| distalUP-(52)-sigma38p10 1288407 Gap 5.5 bits
... -----------------------------------| distalUP-proximalUP-sigma38p10 1288407 total 1.8 bits

     *1288430   .         *1288440   .
5' t g t g g c t a c g a t t g c a t t c c a g 3'
```

piece 3, NC_000913, ydbJ-ydbK, config: linear, direction: +, begin: 1438750, end: 1439099

```
   *1438750  .         *1438760  .         *1438770  .         *1438780  .
5' a c t t c a c t g g t g c g a a a t g c g a c c g a a g c a a c c g c g c c 3'
   T timestamp:_2018Sep20_11-36-03

     *1438790  .         *1438800  .         *1438810  .         *1438820  .
5' a t t a c c g t c a a t a g t a a t c a t a t g a c a c c c t t a c a t t g 3'

       *1438830  .         *1438840  .         *1438850  .         *1438860
5' c g c a a a t g a g g g g c g c a c g a a a t t t g c t g c g c g c c c a g t 3'
              proximalUP 2.7 bits        distalUP 6.6 bits    <------- ... ydbK
            |------------------------------------------------------- ... proximalUP-(34)
        distalUP 0.2 bits             |------------------------------ ... distalUP-(55)-s
   |------------------------------------------------------------------ ... distalUP-(40)-s
   |-----------|------------------------------------------------------ ... distalUP-proxim
                                      |------------------------------ ... distalUP-proxim

   .         *1438870  .         *1438880  .         *1438890  .         *1438900
5' a g t a a t c t t t c a a t t t t a g c a a a t g g c t t t c t t c t g c a 3'
... ----------------------------------#---------] ydbK                        ... sigma38p10
... ------| proximalUP-(34)-sigma38p10 1438867 Gap 2.5 bits
... -----------------------------------------------------------------------| distalUP-(55)-sigma38p10
... ------| distalUP-(40)-sigma38p10 1438867 Gap 3.1 bits
... ------| distalUP-proximalUP-sigma38p10 1438867 total 3.9 bits
... ------------|------------------------------------------------------------| distalUP-proximalUP-sigm
     sigma38p10 6.5 bits
     proximalUP 0.7 bits
          |-----------------------------------------------------------------| proximalUP-(29)-sigma38p

   .         *1438910  .         *1438920  .         *1438930  .
5' t t t t c g c t t t t g t g t c c c c c a c a t c a g c g t a a t g a a t g 3'
... sigma38p10 5.3 bits                                       proximalUP 4.6 bits
                                                            |- ... proximalUP-(42)
                                                               ... distalUP
```

```
                                                                   |--------- ... distalUP-(46)-s
                                                                   |-------|- ... distalUP-proxim

    *1438940  .         *1438950  .         *1438960  .         *1438970  .
 5' t t t t g a t c a a a c a g a g g g c a a a a a a t a g c c a t a a a a a 3'
                                                                 distalUP 5.6 bits
... ---------------------------------------------------------------------------- ... proximalUP-(42)
...    distalUP 2.2 bits                                          |--------------- ... distalUP-(56)-s

... ---------------------------------------------------------------------------- ... distalUP-(46)-s
... ---------------------------------------------------------------------------- ... distalUP-proxim
                                                                  |--------------- ... distalUP-proxim

      *1438980  .         *1438990  .         *1439000  .         *1439010  .
 5' a g t a a a a a t g c g g g g c a a c g c g c a g g a c a g c t c t c g a c 3'
                          sigma38p10 3.9 bits
... ------| proximalUP-(42)-sigma38p10 1438981 Gap 5.1 bits
... ---------------------------------------------------------------------------- ... distalUP-(56)-s
... ------| distalUP-(46)-sigma38p10 1438981 Gap 4.1 bits
... ------| distalUP-proximalUP-sigma38p10 1438981 total 1.4 bits
... -----------------|---------------------------------------------------------- ... distalUP-proxim
                            proximalUP 5.6 bits
                  |------------------------------------------------------------- ... proximalUP-(39)

          *1439020  .         *1439030  .         *1439040  .         *1439050
 5' g c c g c c c t c t g t g c t g c c t a t t a t t t a t g g c g g t g t c g 3'
                                       sigma38p10 3.8 bits

... --------------------|  distalUP-(56)-sigma38p10 1439026 Gap 3.8 bits

... --------------------|  distalUP-proximalUP-sigma38p10 1439026 total 3.8 bits
... --------------------|  proximalUP-(39)-sigma38p10 1439026 Gap 7.5 bits

   .         *1439060  .         *1439070  .         *1439080  .         *1439090
 5' t t t g g c t t g a a g a t c a g a g g a a g a a a a g a t g c g a g c a g 3'

         .
 5' c g t t t t g g 3'
```

piece 4, NC_000913, cybB-gapC1, config: linear, direction: +, begin: 1488600, end: 1488999

```
   *1488600   .          *1488610   .          *1488620   .          *1488630   .
5' t t t c a g c a g g t a g g c g a g a a t t t a t g g g g a a g t g a g a t 3'
```

T timestamp:_2018Sep20_11-36-03

proximalUP 6.3 bits

... proximalUP-(33)

distalUP 6.6 bits

... distalUP-(41)-s
... distalUP-proxim
... proximalUP-(38)
... distalUP-(46)-s
... distalUP-proxim

```
     *1488640   .          *1488650   .          *1488660   .          *1488670   .
5' c a t t a a t a g c g a c a a c g t c t a t g t t g c t t t t g a c t t c a 3'
```

sigma38p10 6.1 bits

... proximalUP-(33)-sigma38p10 1488658 Gap 4.0 bits

sigma38p10 2.4 bits

... distalUP-(41)-sigma38p10 1488658 Gap 3.1 bits
... distalUP-proximalUP-sigma38p10 1488658 total 11.8 bits
... proximalUP-(38)-sigma38p10 1488663 Gap 3.2 b
... distalUP-(46)-sigma38p10 1488663 Gap 4.1 bit
... distalUP-proximalUP-sigma38p10 1488663 tota

```
       *1488680   .          *1488690   .          *1488700   .          *1488710
5' a g t a a t c g a c c c a a c a c c a g t c g a c c g a t a c g a c c a a a 3'
```

```
   .          *1488720   .          *1488730   .          *1488740   .          *1488750
5' a c c g t t a a t a c c a a c t t t a c t c a t g g t t t t c t c c t g t c 3'
```

proximalUP 1.9 bits

... proximalUP-(37)

distalUP 5.5 bits

... proximalUP-(34)

... distalUP-(50)-s
... distalUP-proxim

proximalUP 0.1 bits

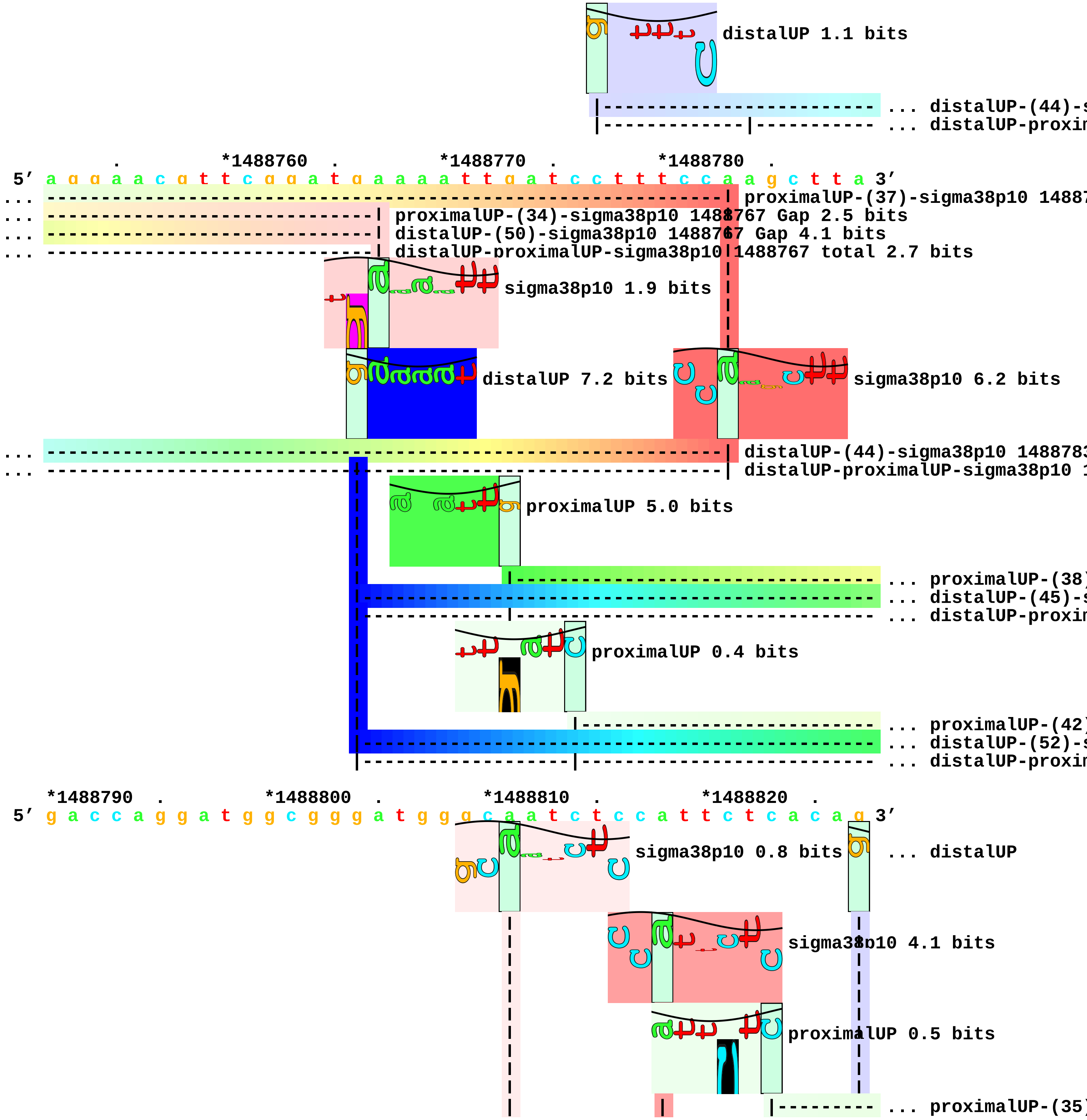
distalUP 1.1 bits
... distalUP-(44)-s
... distalUP-proxim
. *1488760 . *1488770 . *1488780 .
5′ a g g a a c g t t c g g a t g a a a a t t g a t c c t t t c c a a g c t t a 3′
proximalUP-(37)-sigma38p10 14887
proximalUP-(34)-sigma38p10 1488767 Gap 2.5 bits
distalUP-(50)-sigma38p10 1488767 Gap 4.1 bits
distalUP-proximalUP-sigma38p10 1488767 total 2.7 bits
sigma38p10 1.9 bits
distalUP 7.2 bits
sigma38p10 6.2 bits
distalUP-(44)-sigma38p10 148878
distalUP-proximalUP-sigma38p10 1
proximalUP 5.0 bits
... proximalUP-(38)
... distalUP-(45)-s
... distalUP-proxim
proximalUP 0.4 bits
... proximalUP-(42)
... distalUP-(52)-s
... distalUP-proxim
*1488790 . *1488800 . *1488810 . *1488820 .
5′ g a c c a g g a t g g c g g g a t g g g c a a t c t c c a t t c t c a c a g 3′
sigma38p10 0.8 bits
... distalUP
sigma38p10 4.1 bits
proximalUP 0.5 bits
... proximalUP-(35)

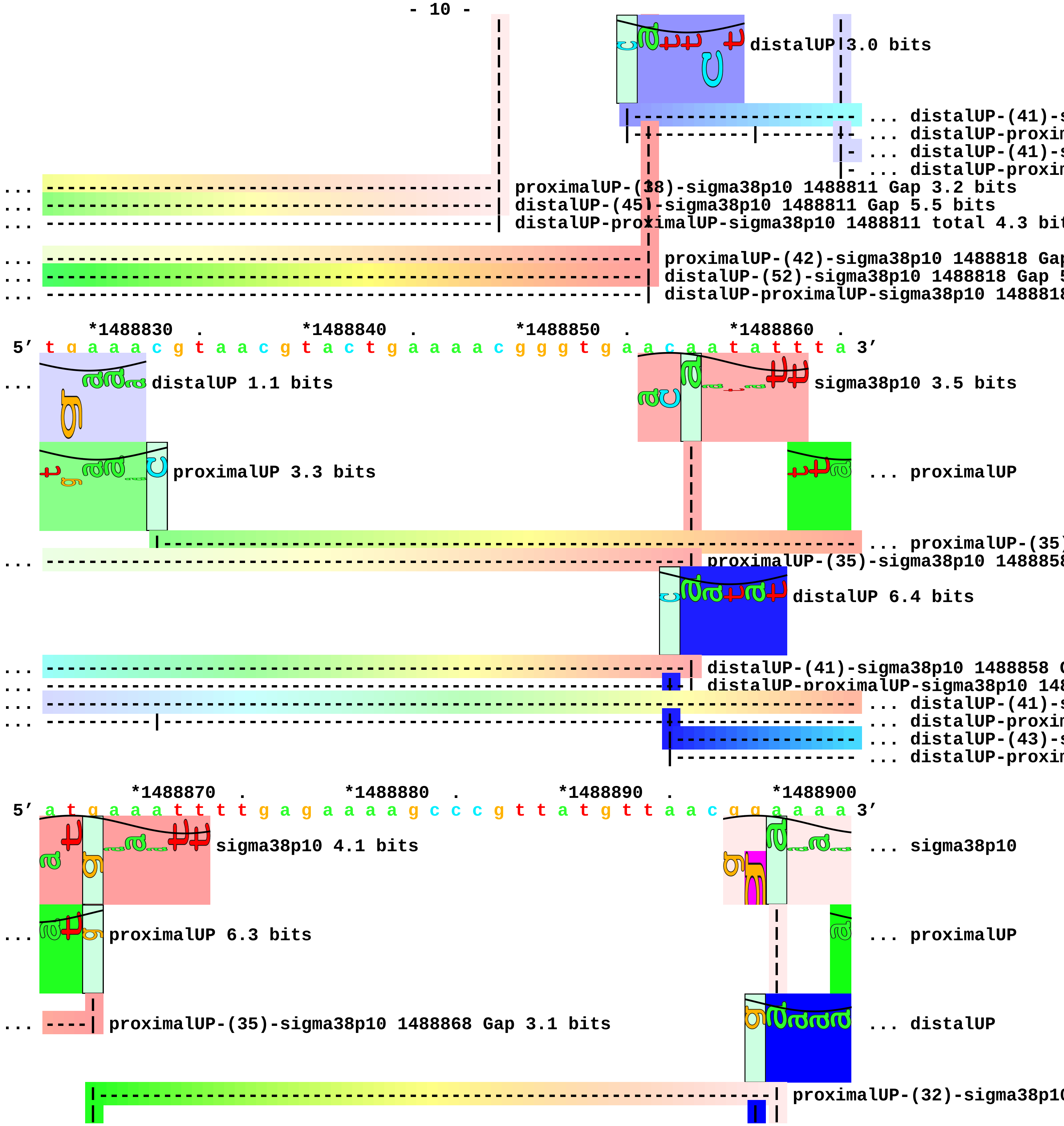
distalUP 3.0 bits
... distalUP-(41)-s
... distalUP-proxim
... distalUP-(41)-s
... distalUP-proxim
... proximalUP-(38)-sigma38p10 1488811 Gap 3.2 bits
... distalUP-(45)-sigma38p10 1488811 Gap 5.5 bits
... distalUP-proximalUP-sigma38p10 1488811 total 4.3 bit
... proximalUP-(42)-sigma38p10 1488818 Gap
... distalUP-(52)-sigma38p10 1488818 Gap 5
... distalUP-proximalUP-sigma38p10 1488818
*1488830 . *1488840 . *1488850 . *1488860 .
5′ t g a a a c g t a a c g t a c t g a a a a c g g g t g a a c a a t a t t t a 3′
... distalUP 1.1 bits
sigma38p10 3.5 bits
proximalUP 3.3 bits
... proximalUP
... proximalUP-(35)
... proximalUP-(35)-sigma38p10 148885
distalUP 6.4 bits
... distalUP-(41)-sigma38p10 1488858 G
... distalUP-proximalUP-sigma38p10 148
... distalUP-(41)-s
... distalUP-proxim
... distalUP-(43)-s
... distalUP-proxim
*1488870 . *1488880 . *1488890 . *1488900
5′ a t g a a a t t t t g a g a a a a g c c c g t t a t g t t a a c g g a a a a 3′
sigma38p10 4.1 bits
... sigma38p10
... proximalUP 6.3 bits
... proximalUP
... proximalUP-(35)-sigma38p10 1488868 Gap 3.1 bits
... distalUP
proximalUP-(32)-sigma38p10

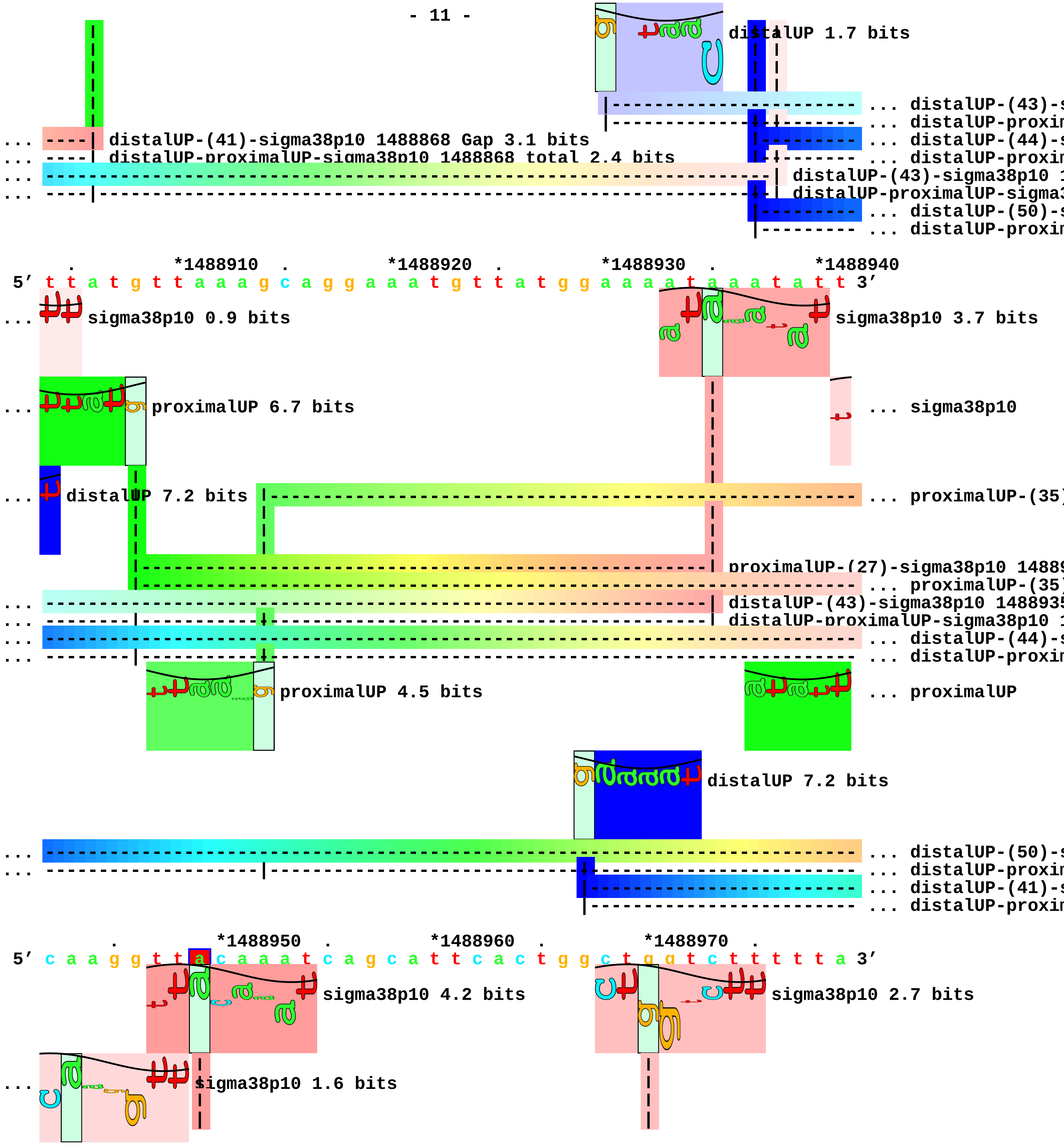
distalUP 1.7 bits
... distalUP-(43)-s
... distalUP-proxin
... distalUP-(41)-sigma38p10 1488868 Gap 3.1 bits
... distalUP-proximalUP-sigma38p10 1488868 total 2.4 bits
... distalUP-(44)-s
... distalUP-proxin
distalUP-(43)-sigma38p10 1
distalUP-proximalUP-sigma3
... distalUP-(50)-s
... distalUP-proxin
*1488910
*1488920
*1488930
*1488940
5′ t t a t g t t a a a g c a g g a a a t g t t a t g g a a a a t a a a t a t t 3′
... sigma38p10 0.9 bits
sigma38p10 3.7 bits
... proximalUP 6.7 bits
... sigma38p10
... distalUP 7.2 bits
... proximalUP-(35)
proximalUP-(27)-sigma38p10 1488
... proximalUP-(35)
distalUP-(43)-sigma38p10 148893
distalUP-proximalUP-sigma38p10 1
... distalUP-(44)-s
... distalUP-proxin
proximalUP 4.5 bits
... proximalUP
distalUP 7.2 bits
... distalUP-(50)-s
... distalUP-proxin
... distalUP-(41)-s
... distalUP-proxin
*1488950
*1488960
*1488970
5′ c a a g g t t a c a a a t c a g c a t t c a c t g g c t g g t c t t t t t a 3′
sigma38p10 4.2 bits
sigma38p10 2.7 bits
... sigma38p10 1.6 bits

```
... --------------| proximalUP-(35)-sigma38p10 1488949 Gap 3.1 bits
    |-------------------------------------------------------| proximalUP-(28)-sigma38p10 1488970 Gap
... --| proximalUP-(35)-sigma38p10 1488943 Gap 3.1 bits
... --| distalUP-(44)-sigma38p10 1488943 Gap 2.8 bits
... +-| distalUP-proximalUP-sigma38p10 1488943 total 9.6 bits
... C proximalUP 6.6 bits
... --------------| distalUP-(50)-sigma38p10 1488949 Gap 4.1 bits
... --------------| distalUP-proximalUP-sigma38p10 1488949 total 8.7 bits
... --------------------------------------------------------| distalUP-(41)-sigma38p10 1488970 Gap 3
... |-------------------------------------------------------| distalUP-proximalUP-sigma38p10 148897

    *1488980  .         *1488990  .
 5' c t g g t t a t c g c a g c g t a t t g 3'
```

piece 5, NC_000913, nadE-osmE, config: linear, direction: +, begin: 1820200, end: 1820399

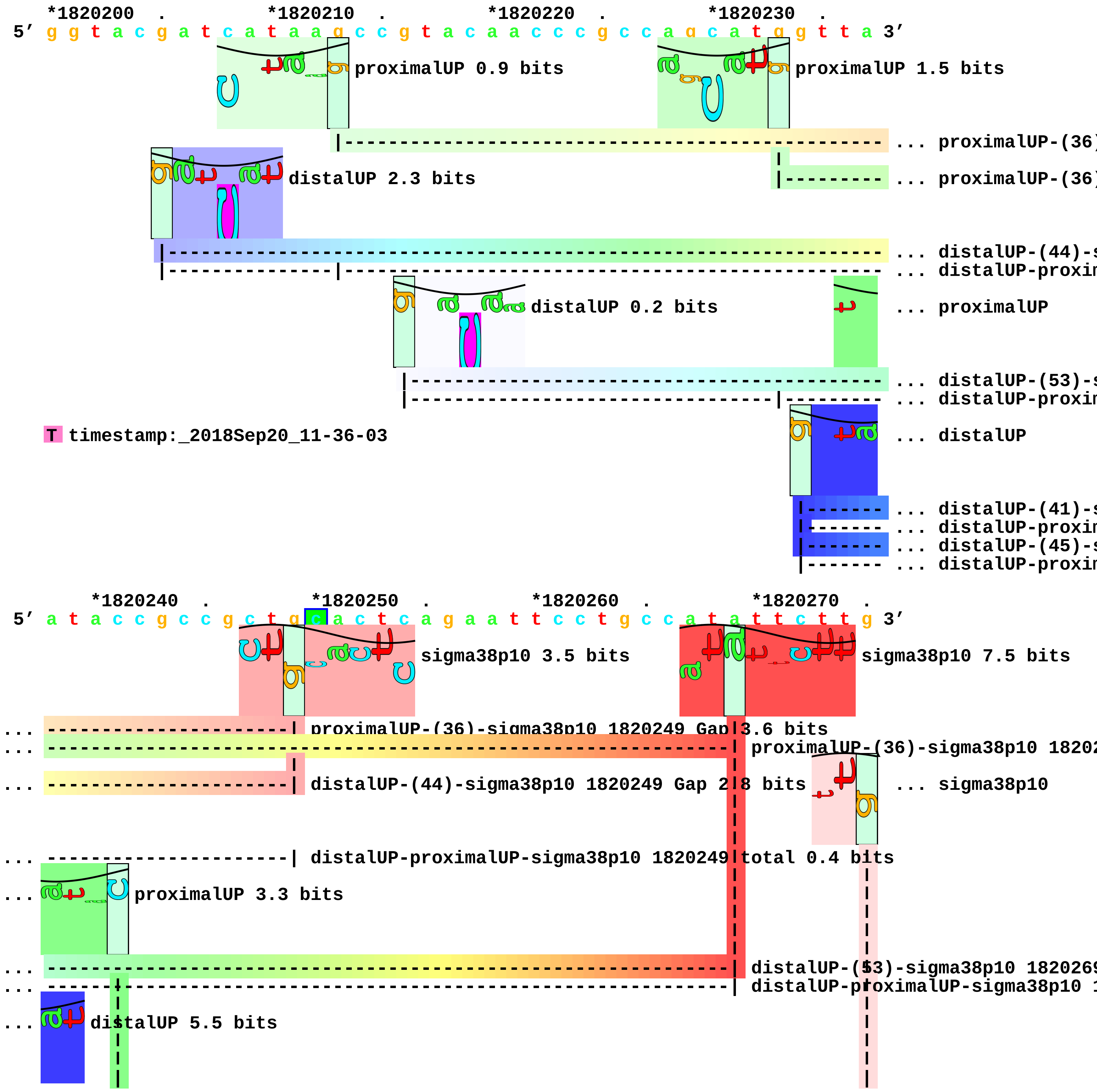

```
... -----------------------------------------------------------------------| distalUP-(41)-sigma3
... ------|----------------------------------------------------------------| distalUP-proximalUP-
... ------------------------------------------------------------------------ ... distalUP-(45)-s
... ------|------------------------------------------------------------------ ... distalUP-proxin
      |----------------------------------------------------------------| proximalUP-(34)-sign
      |----------------------------------------------------------------- ... proximalUP-(38)

        *1820280  .         *1820290  .         *1820300  .         *1820310
 5' t t c a t a t t c c g t c c t c t t g t t t a t c a g c g t g t t a g a t a 3'
                      sigma38p10 2.4 bits

                      proximalUP 6.6 bits

                |-------------------------------------------------------- ... proximalUP-(40)
...             sigma38p10 1.5 bits

                distalUP 5.7 bits

      |------------------------------------------------------------------ ... distalUP-(46)-s
      |---------|------------------------------------------------------- ... distalUP-proxin

... ------| distalUP-(45)-sigma38p10 1820279 Gap 5.5 bits
... ------| distalUP-proximalUP-sigma38p10 1820279 total 2.6 bits

... ------| proximalUP-(38)-sigma38p10 1820279 Gap 3.2 bits

   .         *1820320  .         *1820330  .         *1820340  .         *1820350
 5' a g c c t g g a a t a c a t t g g g c g c t t t t t c a a g c c c g t g a a 3'
                      sigma38p10 3.8 bits

... --------------------| proximalUP-(40)-sigma38p10 1820324 Gap 4.0 bits

... --------------------| distalUP-(46)-sigma38p10 1820324 Gap 4.1 bits
... --------------------| distalUP-proximalUP-sigma38p10 1820324 total 8.0 bits

      .         *1820360  .         *1820370  .         *1820380  .
 5' c g a a a c g g c t c c g c t t t c a g a g g a t t c c t g t a t g a c g t 3'

   *1820390  .
 5' t t t a a c c a c c 3'
```

piece 6, NC_000913, yodD-dsrB, config: linear, direction: +, begin: 2022800, end: 2023199

*2022800 . *2022810 . *2022820 . *2022830 .
5′ a c g c g g a c c g c c a t c c g t t t t g a c t g t t a c c c g a t c a t 3′
T timestamp:_2018Sep20_11-36-03

*2022840 . *2022850 . *2022860 . *2022870 .
5′ t c a c c t t c a t c g t t t c c t c c t g t g g c t t t g t g c c a g t 3′

*2022880 . *2022890 . *2022900 . *2022910
5′ g t a g a a c a a t t t c g t t t t t c t g g c a g c g c c a g g c g c g c 3′

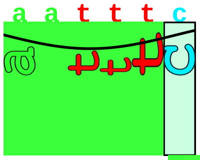

proximalUP 5.5 bits

... proximalUP-(35)[illegible]

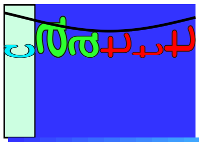

distalUP 5.8 bits

... distalUP-(41)-[illegible]
... distalUP-proxi[illegible]

. *2022920 . *2022930 . *2022940 . *2022950
5′ g c g a g t g c t g a t t t t c t c g a c g g t c t a t a c t t a a g a g a g a 3′

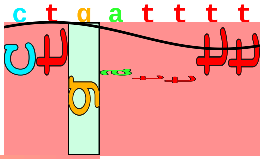

sigma38p10 4.7 bits

... proximalUP-(35)-sigma38p10 2022923 Gap 3.1 bits
... distalUP-(41)-sigma38p10 2022923 Gap 3.1 bits
... distalUP-proximalUP-sigma38p10 2022923 total 9.8 bits

. *2022960 . *2022970 . *2022980 .
5′ t g c c a g g c g g a c t t a a c g a c t g g c g g c a a c a a c a g a g t 3′

*2022990 . *2023000 . *2023010 . *2023020 .
5′ a a c g g t t g c g a g g a a a g a t g a t g a a a a c c g c a a a a g a g 3′

*2023030 . *2023040 . *2023050 . *2023060 .
5′ t a c a g c g a t a c c g c a a a a c g t g a g g t c a g c g t c g a t g t 3′

*2023070 . *2023080 . *2023090 . *2023100
5′ c g a t g c c c t g c t g g c g g c g a t c a a t g a a a t t a g c g a a a 3′

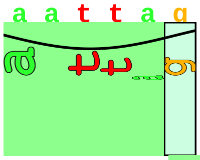

proximalUP 3.3 bits

... proximalUP-(35)[illegible]

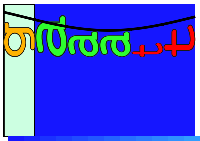

distalUP 6.6 bits

... distalUP-(42)-[illegible]
... distalUP-proxi[illegible]

```
   .         *2023110  .         *2023120  .         *2023130  .         *2023140
5' g c g a a g t t c a t c g c a g c c a g a a c g a t t c t g a a c a c g t t 3'
```

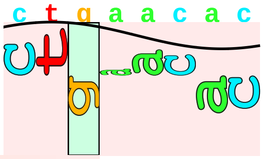

sigma38p10 0.9 bits

```
... ----------------------------------------------------------| proximalUP-(35)-sigma38p10 2023133 G
                                                              |
... ----------------------------------------------------------| distalUP-(42)-sigma38p10 2023133 Gap
... ----------------------------------------------------------| distalUP-proximalUP-sigma38p10 20231
```

```
         .         *2023150  .         *2023160  .         *2023170  .
5' a g c g t c g a t g g a c g t g a a t a t c a t a c a t g g c g t g a a t t 3'
```

```
   *2023180  .         *2023190  .
5' g g c g g a t g c c t t c g a a c t g g 3'
```

piece 7, NC_000913, lpxP-yfdY, config: linear, direction: +, begin: 2493400, end: 2493599

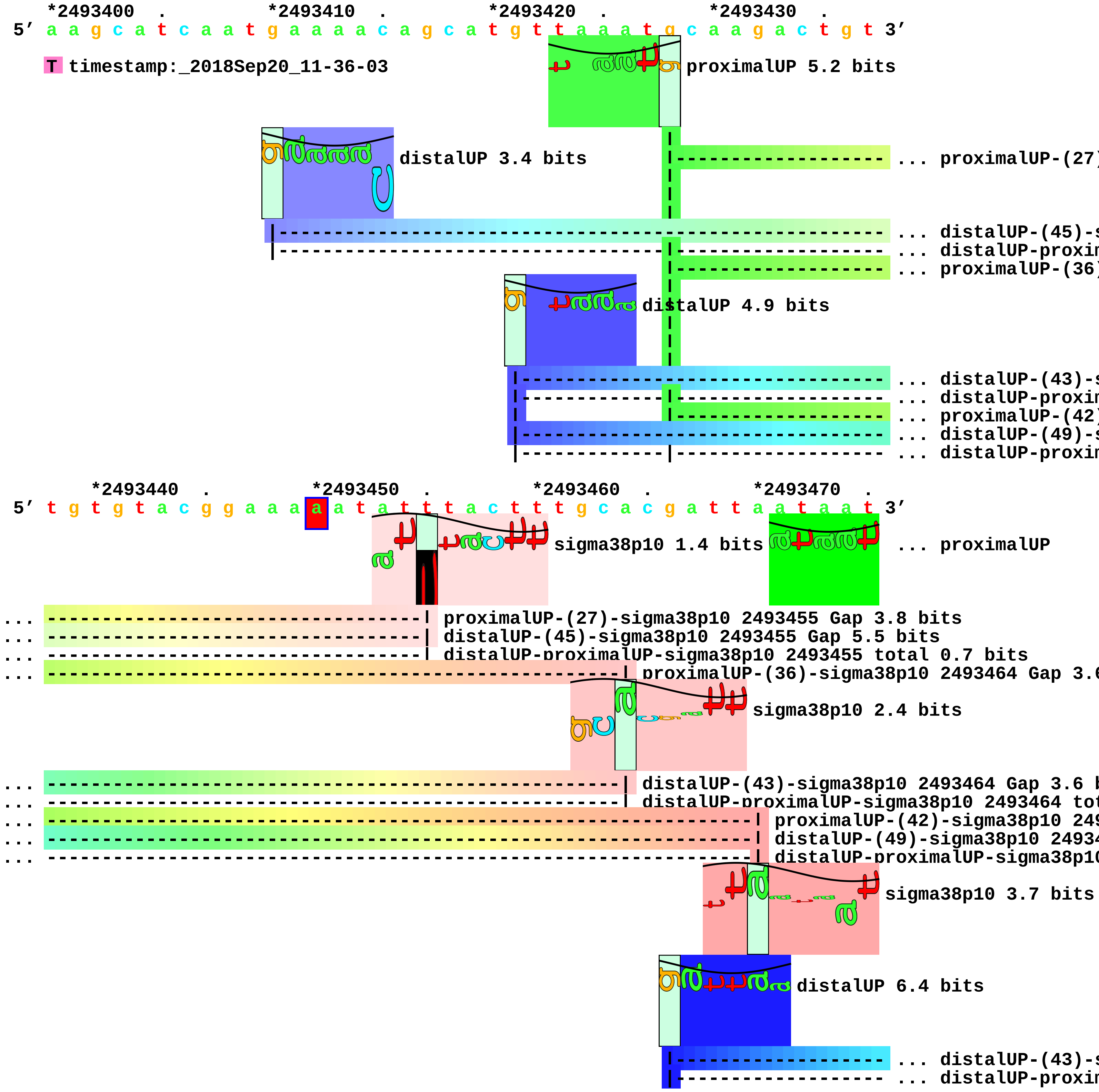

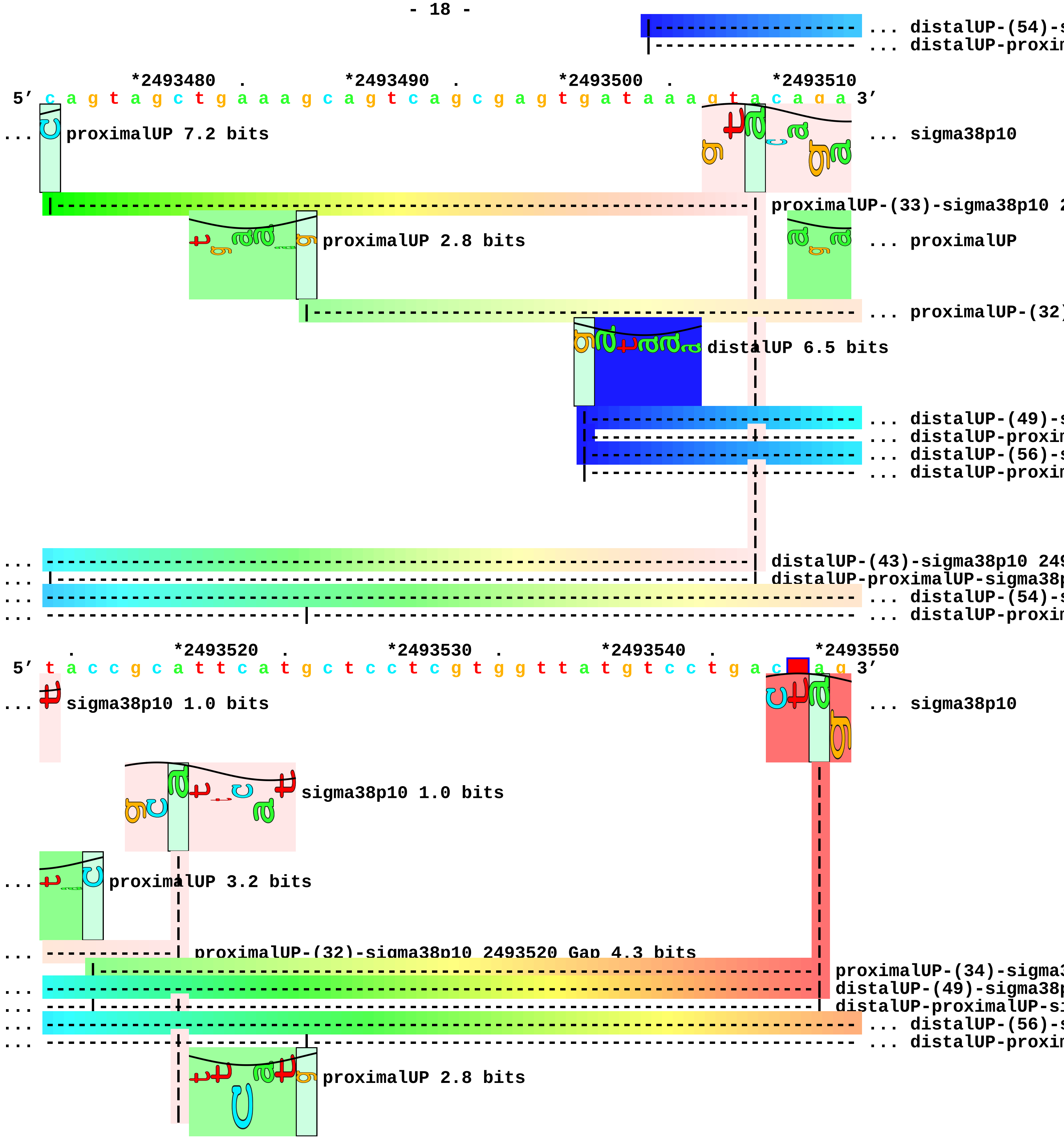
... distalUP-(54)-s
... distalUP-proxim
*2493480 . *2493490 . *2493500 . *2493510
5′ c a g t a g c t g a a a g c a g t c a g c g a g t g a t a a a g t a c a g a 3′
... proximalUP 7.2 bits
... sigma38p10
proximalUP-(33)-sigma38p10 2
proximalUP 2.8 bits
... proximalUP
... proximalUP-(32)
distalUP 6.5 bits
... distalUP-(49)-s
... distalUP-proxim
... distalUP-(56)-s
... distalUP-proxim
... distalUP-(43)-sigma38p10 249
... distalUP-proximalUP-sigma38p
... distalUP-(54)-s
... distalUP-proxim
. *2493520 . *2493530 . *2493540 . *2493550
5′ t a c c g c a t t c a t g c t c c t c g t g g t t a t g t c c t g a c a g 3′
... sigma38p10 1.0 bits
... sigma38p10
sigma38p10 1.0 bits
... proximalUP 3.2 bits
... proximalUP-(32)-sigma38p10 2493520 Gap 4.3 bits
proximalUP-(34)-sigma3
... distalUP-(49)-sigma38p
... distalUP-proximalUP-si
... distalUP-(56)-s
... distalUP-proxim
proximalUP 2.8 bits

```
                  |-------------------------------------------------------------- ... proximalUP-(31)
                  |-------------------------------------------------------------- ... proximalUP-(33)
            distalUP 2.4 bits
      |------------------------------------------------------------------------ ... distalUP-(40)-s
      |--------------|--------------------------------------------------------- ... distalUP-proxim
... ------------| distalUP-(54)-sigma38p10 2493520 Gap 4.9 bits
... ------------| distalUP-proximalUP-sigma38p10 2493520 total 1.1 bits

             .         *2493560   .         *2493570   .         *2493580   .
 5' t c t t t a c a c t c t t t a c a g g a a c c a t t g t c g t a c a t g a t 3'
...         sigma38p10 6.1 bits
            sigma38p10 5.2 bits
            sigma38p10 3.5 bits
... ----------| distalUP-(56)-sigma38p10 2493557 Gap 3.8 bits
... ----------| distalUP-proximalUP-sigma38p10 2493557 total 6.0 bits
... ----------| proximalUP-(31)-sigma38p10 2493557 Gap 4.7 bits
... --------------| proximalUP-(33)-sigma38p10 2493559 Gap 4.0 bits
... --------------| distalUP-(40)-sigma38p10 2493559 Gap 3.1 bits
... --------------| distalUP-proximalUP-sigma38p10 2493559 total 1.5 bits

   *2493590   .
 5' g g c c c a a c c a 3'
```

piece 8, NC_000913, yfgG-yfgF, config: linear, direction: +, begin: 2627050, end: 2627449

```
   *2627050   .          *2627060   .          *2627070   .          *2627080   .
5' a g a g a g c a t t g a t t c t a a g t g t c a t a t g a a a g t a c c a a 3'
   T timestamp:_2018Sep20_11-36-03

       *2627090   .          *2627100   .          *2627110   .          *2627120   .
5' t t g a t a t a t a t c a a a c a a a a t a a c c c t g a t t a a t g a a t 3'
```

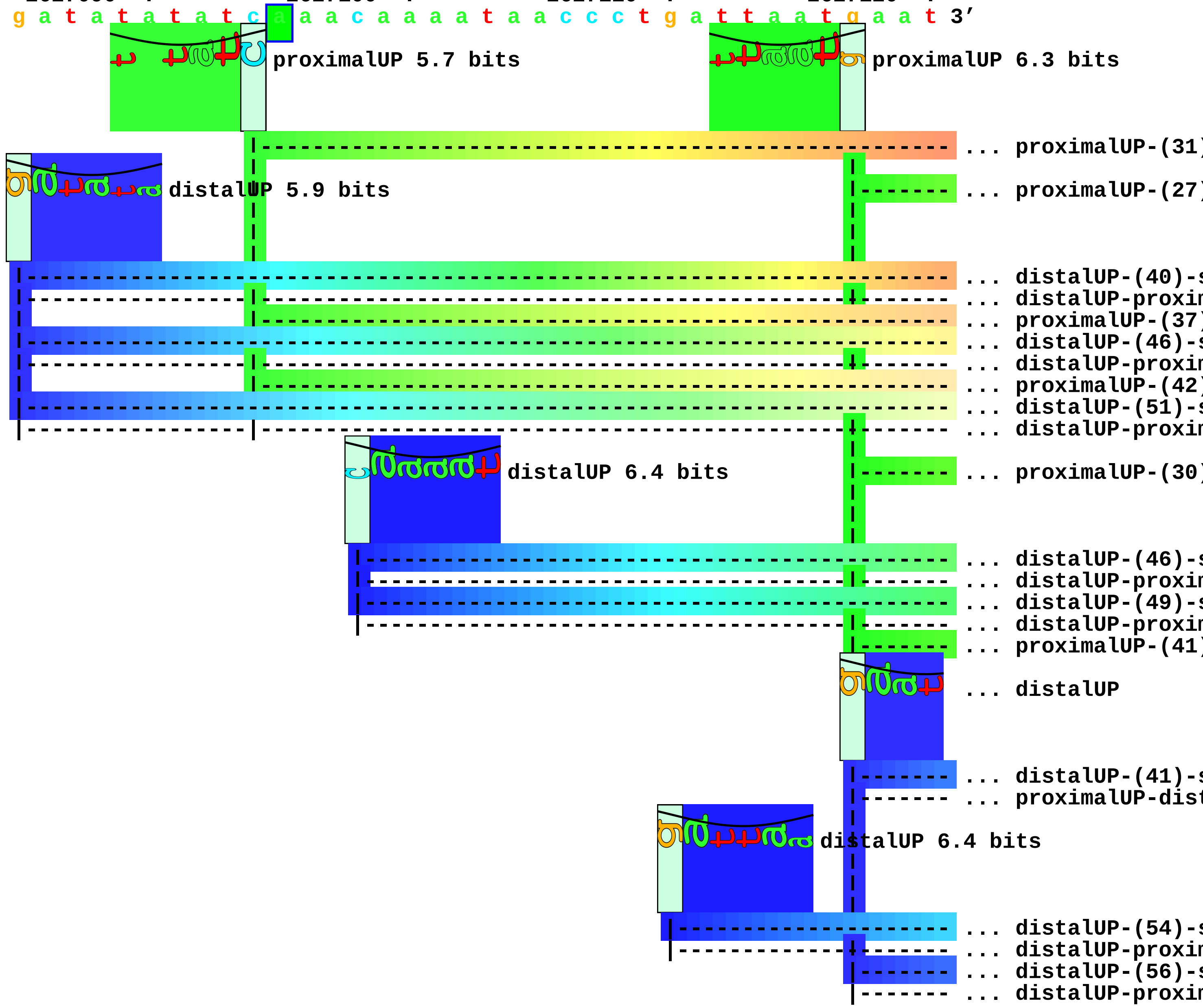


```
           *2627130   .          *2627140   .          *2627150   .          *2627160
5' t a t t a c g t t t a t c a t g t t a a t t c a t c a t t a t t a c a t c a 3'
```

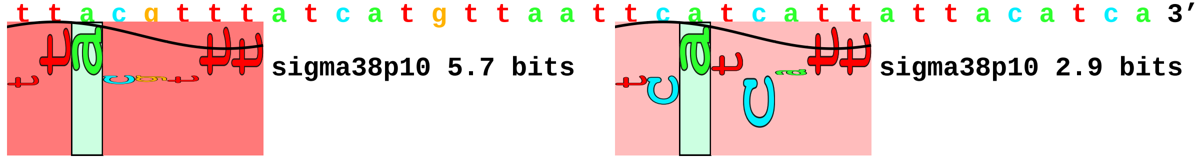

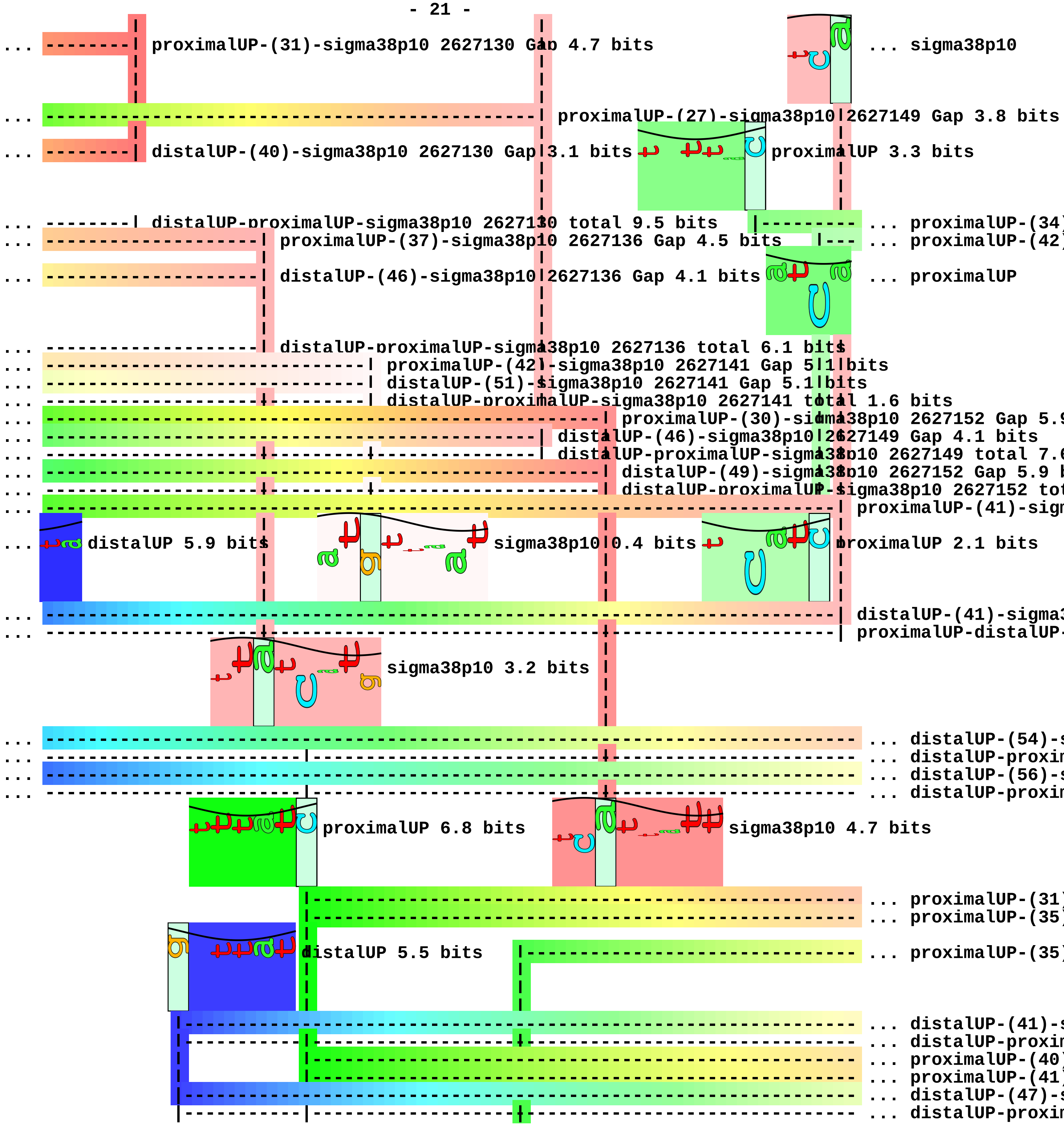
... proximalUP-(31)-sigma38p10 2627130 Gap 4.7 bits
... sigma38p10
... proximalUP-(27)-sigma38p10 2627149 Gap 3.8 bits
... distalUP-(40)-sigma38p10 2627130 Gap 3.1 bits
proximalUP 3.3 bits
... distalUP-proximalUP-sigma38p10 2627130 total 9.5 bits
... proximalUP-(37)-sigma38p10 2627136 Gap 4.5 bits
... distalUP-(46)-sigma38p10 2627136 Gap 4.1 bits
... proximalUP
... distalUP-proximalUP-sigma38p10 2627136 total 6.1 bits
... proximalUP-(42)-sigma38p10 2627141 Gap 5.1 bits
... distalUP-(51)-sigma38p10 2627141 Gap 5.1 bits
... distalUP-proximalUP-sigma38p10 2627141 total 1.6 bits
... distalUP-(46)-sigma38p10 2627149 Gap 4.1 bits
... distalUP 5.9 bits
sigma38p10 0.4 bits
proximalUP 2.1 bits
sigma38p10 3.2 bits
proximalUP 6.8 bits
sigma38p10 4.7 bits
distalUP 5.5 bits

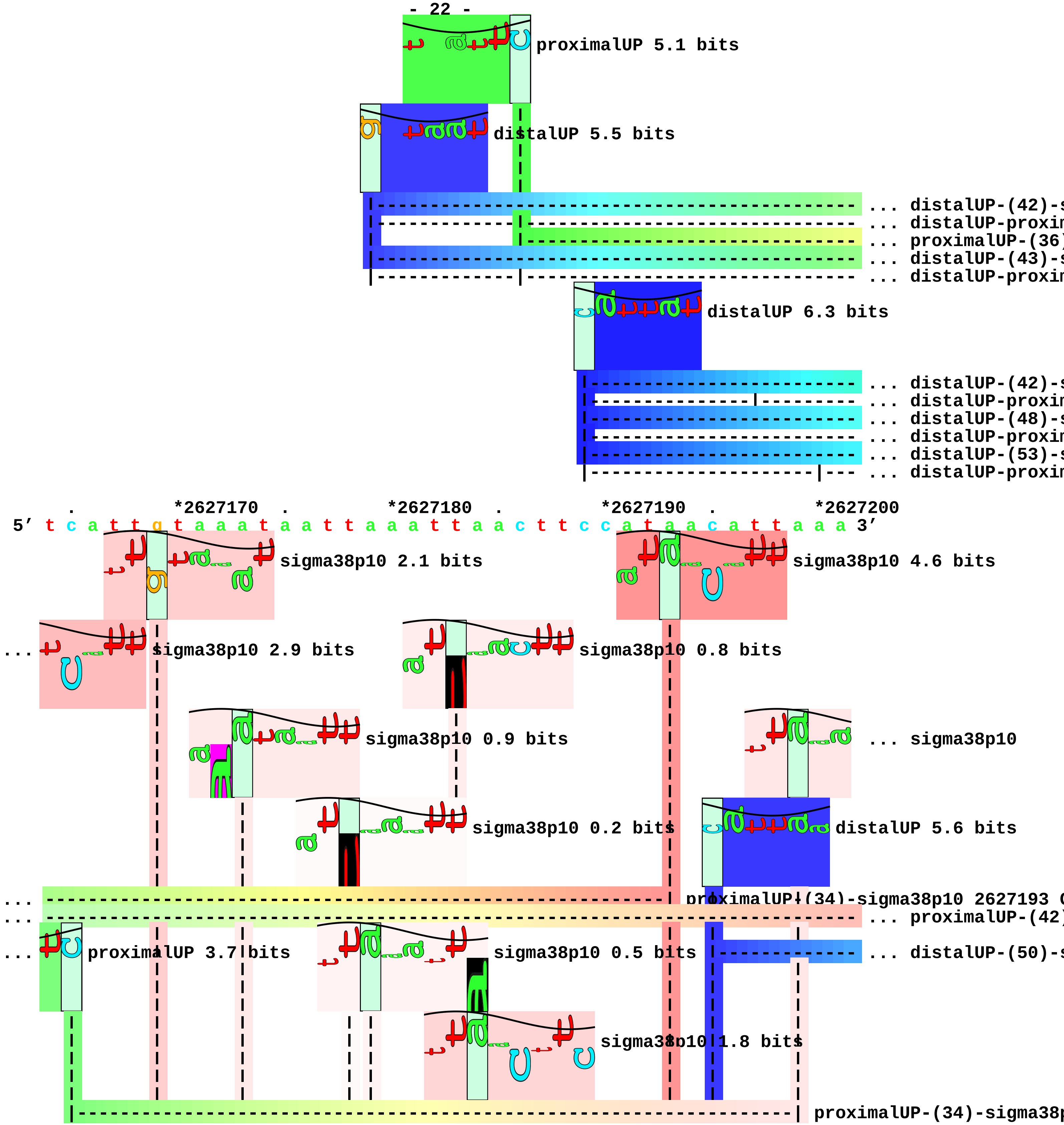
proximalUP 5.1 bits
distalUP 5.5 bits
... distalUP-(42)-s
... distalUP-proxin
... proximalUP-(36)
... distalUP-(43)-s
... distalUP-proxin
distalUP 6.3 bits
... distalUP-(42)-s
... distalUP-proxin
... distalUP-(48)-s
... distalUP-proxin
... distalUP-(53)-s
... distalUP-proxin
*2627170
*2627180
*2627190
*2627200
5′ t c a t t g t a a a t a a t t a a a t t a a c t t c c a t a a c a t t a a a 3′
sigma38p10 2.1 bits
sigma38p10 4.6 bits
sigma38p10 2.9 bits
sigma38p10 0.8 bits
sigma38p10 0.9 bits
... sigma38p10
sigma38p10 0.2 bits
distalUP 5.6 bits
proximalUP-(34)-sigma38p10 2627193 (
... proximalUP-(42)
proximalUP 3.7 bits
sigma38p10 0.5 bits
... distalUP-(50)-s
sigma38p10 1.8 bits
proximalUP-(34)-sigma38p

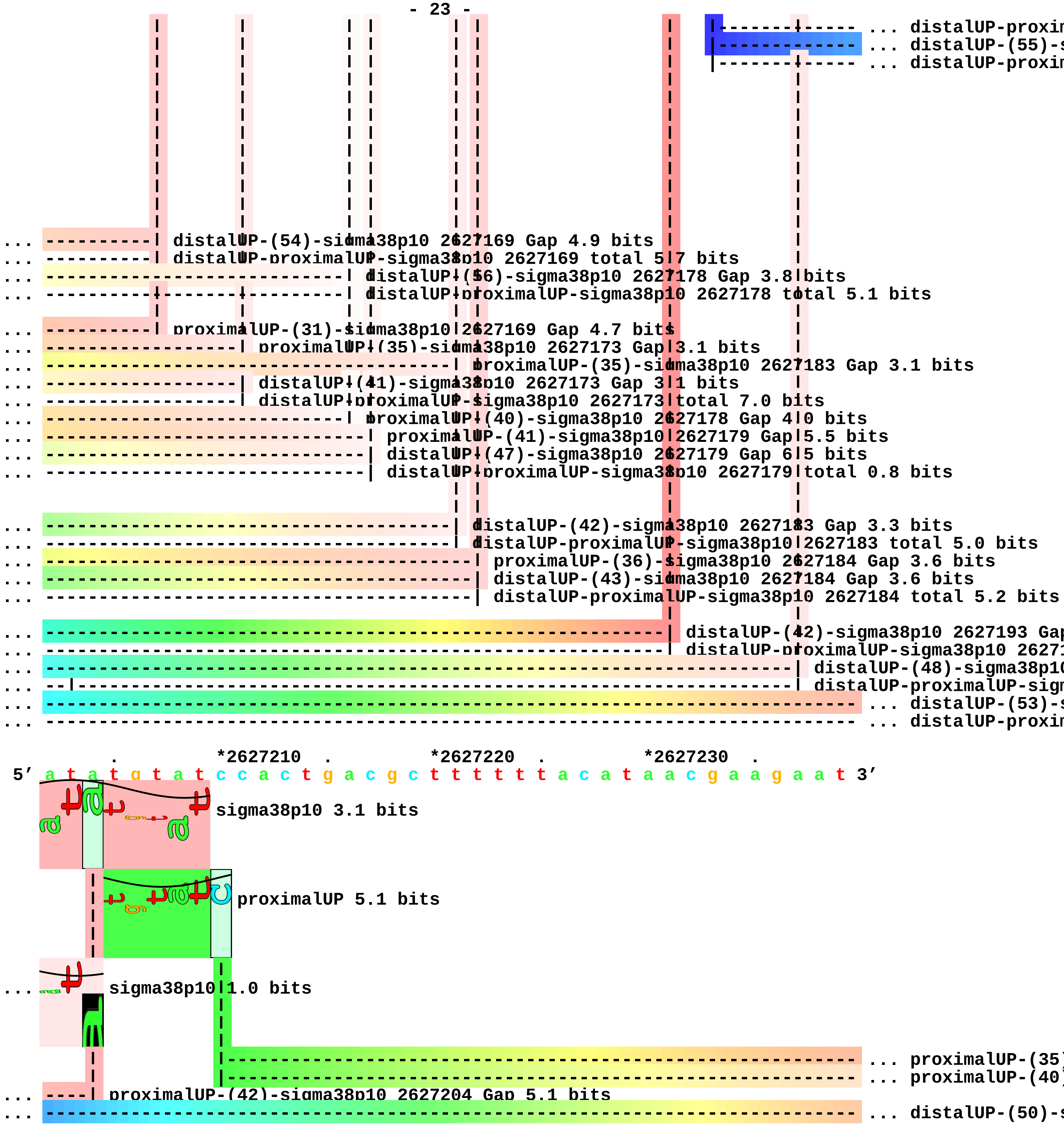
... distalUP-proxin
... distalUP-(55)-s
... distalUP-proxin
... distalUP-(54)-sigma38p10 2627169 Gap 4.9 bits
... distalUP-proximalUP-sigma38p10 2627169 total 5.7 bits
... distalUP-(56)-sigma38p10 2627178 Gap 3.8 bits
... distalUP-proximalUP-sigma38p10 2627178 total 5.1 bits
... proximalUP-(31)-sigma38p10 2627169 Gap 4.7 bits
... proximalUP-(35)-sigma38p10 2627173 Gap 3.1 bits
... proximalUP-(35)-sigma38p10 2627183 Gap 3.1 bits
... distalUP-(41)-sigma38p10 2627173 Gap 3.1 bits
... distalUP-proximalUP-sigma38p10 2627173 total 7.0 bits
... proximalUP-(40)-sigma38p10 2627178 Gap 4.0 bits
... proximalUP-(41)-sigma38p10 2627179 Gap 5.5 bits
... distalUP-(47)-sigma38p10 2627179 Gap 6.5 bits
... distalUP-proximalUP-sigma38p10 2627179 total 0.8 bits
... distalUP-(42)-sigma38p10 2627183 Gap 3.3 bits
... distalUP-proximalUP-sigma38p10 2627183 total 5.0 bits
... proximalUP-(36)-sigma38p10 2627184 Gap 3.6 bits
... distalUP-(43)-sigma38p10 2627184 Gap 3.6 bits
... distalUP-proximalUP-sigma38p10 2627184 total 5.2 bits
... distalUP-(42)-sigma38p10 2627193 Gap
... distalUP-proximalUP-sigma38p10 26271
... distalUP-(48)-sigma38p10
... distalUP-proximalUP-sigm
... distalUP-(53)-s
... distalUP-proxin
. *2627210 . *2627220 . *2627230 .
5′ a t a t g t a t c c a c t g a c g c t t t t t t a c a t a a c g a a g a a t 3′
sigma38p10 3.1 bits
proximalUP 5.1 bits
... sigma38p10 1.0 bits
... proximalUP-(35)
... proximalUP-(40)
... proximalUP-(42)-sigma38p10 2627204 Gap 5.1 bits
... distalUP-(50)-s

```
... ---------------|--------------------------------------------------------- ... distalUP-proxim
... -------------------------------------------------------------------------- ... distalUP-(55)-s
... ---------------|--------------------------------------------------------- ... distalUP-proxim

... ----| distalUP-(53)-sigma38p10 2627204 Gap 4.9 bits
... ----| distalUP-proximalUP-sigma38p10 2627204 total 1.5 bits

   *2627240  .         *2627250  .         *2627260  .         *2627270  .
5' t g a c c a t t t t g t c c t g t t g t g c c t t a a t g t a a g t a c c g 3'
```

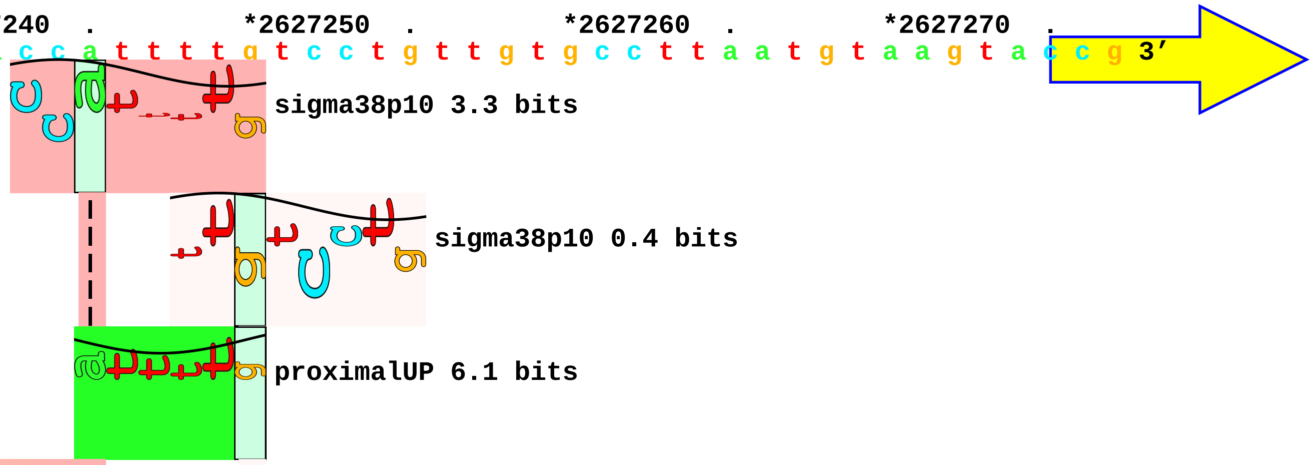


```
... ----------| proximalUP-(35)-sigma38p10 2627245 Gap 3.1 bits
... --------------------| proximalUP-(40)-sigma38p10 2627250 Gap 4.0 bits
```

```
                             |--------------------------------------------------------------- ... proximalUP-(43)
... ----------| distalUP-(50)-sigma38p10 2627245 Gap 4.1 bits
                             distalUP 5.7 bits
               |------------------------------------------------------------------------- ... distalUP-(49)-s
... ----------| distalUP-proximalUP-sigma38p10 2627245 total 6.8 bits
... --------------------| distalUP-(55)-sigma38p10 2627250 Gap 4.9 bits
... --------------------| distalUP-proximalUP-sigma38p10 2627250 total 2.2 bits
               |-----------|------------------------------------------------------------ ... distalUP-proxim

        *2627280   .         *2627290   .         *2627300   .         *2627310   .
5' t c c a c a g c g t g g g a c a t a c t t c a a g g a a c c t t t t g t g a 3'
                                                  sigma38p10 5.8 bits

... ------------------------------| proximalUP-(43)-sigma38p10 2627293 Gap 5.1 bits

... ------------------------------| distalUP-(49)-sigma38p10 2627293 Gap 5.9 bits

... ------------------------------| distalUP-proximalUP-sigma38p10 2627293 total 6.6 bits

            *2627320   .         *2627330   .         *2627340   .         *2627350
5' g t c a g g c a a c c a g t a t g c g a a a a c g a c a c c g a t t t a a c 3'
                                                                   proximalUP 5.0 bits
                                                                     |- ... proximalUP-(36)
                                                                distalUP 5.8 bits
                                                               |--------------- ... distalUP-(43)-s
                                                               |-------------|- ... distalUP-proxim
                                                                             |- ... proximalUP-(37)
                                                               |--------------- ... distalUP-(44)-s
                                                               |-------------|- ... distalUP-proxim
                                                                             |- ... proximalUP-(42)
                                                               |--------------- ... distalUP-(49)-s
                                                               |-------------|- ... distalUP-proxim

   .         *2627360   .         *2627370   .         *2627380   .         *2627390
5' a g t c g c a t g a c c c g t a t c g t a c t g c t c a t c a g c t t t a t 3'
                                                                            ... sigma38p10
```

```
... ------------------------------------------------------------------------| proximalUP-(36)-sigma38p
                                                                              ... sigma38p10

... ------------------------------------------------------------------------| distalUP-(43)-sigma38p10
... ------------------------------------------------------------------------| distalUP-proximalUP-sign
... --------------------------------------------------------------------------| proximalUP-(37)-sigma3
... --------------------------------------------------------------------------| distalUP-(44)-sigma38p
... --------------------------------------------------------------------------| distalUP-proximalUP-si
... ------------------------------------------------------------------------------ ... proximalUP-(42)
... ------------------------------------------------------------------------------ ... distalUP-(49)-s
... ------------------------------------------------------------------------------ ... distalUP-proxin

            .         *2627400   .         *2627410   .         *2627420   .
 5' c t t c t t c t t t g g c c g t t t t a t c t a c t c g t c c g t c g g t g 3'
... sigma38p10 0.2 bits

       sigma38p10 0.8 bits

... sigma38p10 2.4 bits

... ------| proximalUP-(42)-sigma38p10 2627395 Gap 5.1 bits
... ------| distalUP-(49)-sigma38p10 2627395 Gap 5.9 bits
... ------| distalUP-proximalUP-sigma38p10 2627395 total 0.6 bits

    *2627430   .         *2627440   .
 5' c c t g g c a g c a c c a t c a g a g c 3'
```

piece 9, NC_000913, ygcG-queE, config: linear, direction: +, begin: 2903300, end: 2903699

```
   *2903300  .         *2903310  .         *2903320  .         *2903330  .
5' a a g c t t t t c c c a g g t g t g t t t g g t g t c g c a c c a g g c a c 3'
   T timestamp:_2018Sep20_11-36-03

     *2903340  .         *2903350  .         *2903360  .         *2903370  .
5' a g c c a a c c g g g c a t c c c t g t a a a c g a a t a a a a a t g g c g 3'

       *2903380  .         *2903390  .         *2903400  .         *2903410
5' g g a a c g c c g g t a a a g t a a c c c t c a c c t t g c a g g g t c t g 3'

   .         *2903420  .         *2903430  .         *2903440  .         *2903450
5' g a a c a t c t c g t t a a t c g g g t a c t g c a t a g c a t t c t c t g 3'
                                 proximalUP 6.8 bits              distalUP 3.0 bits
   distalUP 2.4 bits  |---------------------------------------- ... proximalUP-(31)
   |------------------------------------------------------------ ... distalUP-(46)-s
   |------------------------------------------------------------ ... distalUP-proxim
                      |---------------------------------------- ... proximalUP-(37)
            distalUP 5.5 bits                       |---------- ... distalUP-(50)-s
            |-------------------------------------------------- ... distalUP-(43)-s
            |-----------|-------------------------------------- ... distalUP-proxim
                        |-------------------------------------- ... proximalUP-(40)
            |-------------------------------------------------- ... distalUP-(46)-s
            |-----------|-------------------------------------- ... distalUP-proxim
                        |-------------------------------------- ... proximalUP-(41)
            |-------------------------------------------------- ... distalUP-(47)-s
            |-----------|-------------------------------------- ... distalUP-proxim
                                                    |---------- ... distalUP-proxim

       .         *2903460  .         *2903470  .         *2903480  .
5' t g a a g t g g a t a a t t g t t a a t t a t t g c a g a t c c t g c c a c 3'
                  sigma38p10 1.5 bits  |------------------------ ... proximalUP-(31)
... ----------------| proximalUP-(31)-sigma38p10 2903460 Gap 4.7 bits
... ----------------| distalUP-(46)-sigma38p10 2903460 Gap 4.1 bits
... ----------------| distalUP-proximalUP-sigma38p10 2903460 total 1.9 bits
... ----------------------------| proximalUP-(37)-sigma38p10 2903466 Gap 4.5 bits
... ------------------------------------------------------------ ... distalUP-(50)-s
... ----------------------------| distalUP-(43)-sigma38p10 2903466 Gap 3.6 bits
... ----------------------------| distalUP-proximalUP-sigma38p10 2903466 total 5.2 bits
... ----------------------------------| proximalUP-(40)-sigma38p10 2903469 Gap 4.0 bits
... ----------------------------------| distalUP-(46)-sigma38p10 2903469 Gap 4.1 bits
... ----------------------------|-----| distalUP-proximalUP-sigma38p10 2903469 total 7.3 bits
... ------------------------------------| proximalUP-(41)-sigma38p10 2903470 Gap 5.5 bits
... ------------------------------------| distalUP-(47)-sigma38p10 2903470 Gap 6.5 bits
```

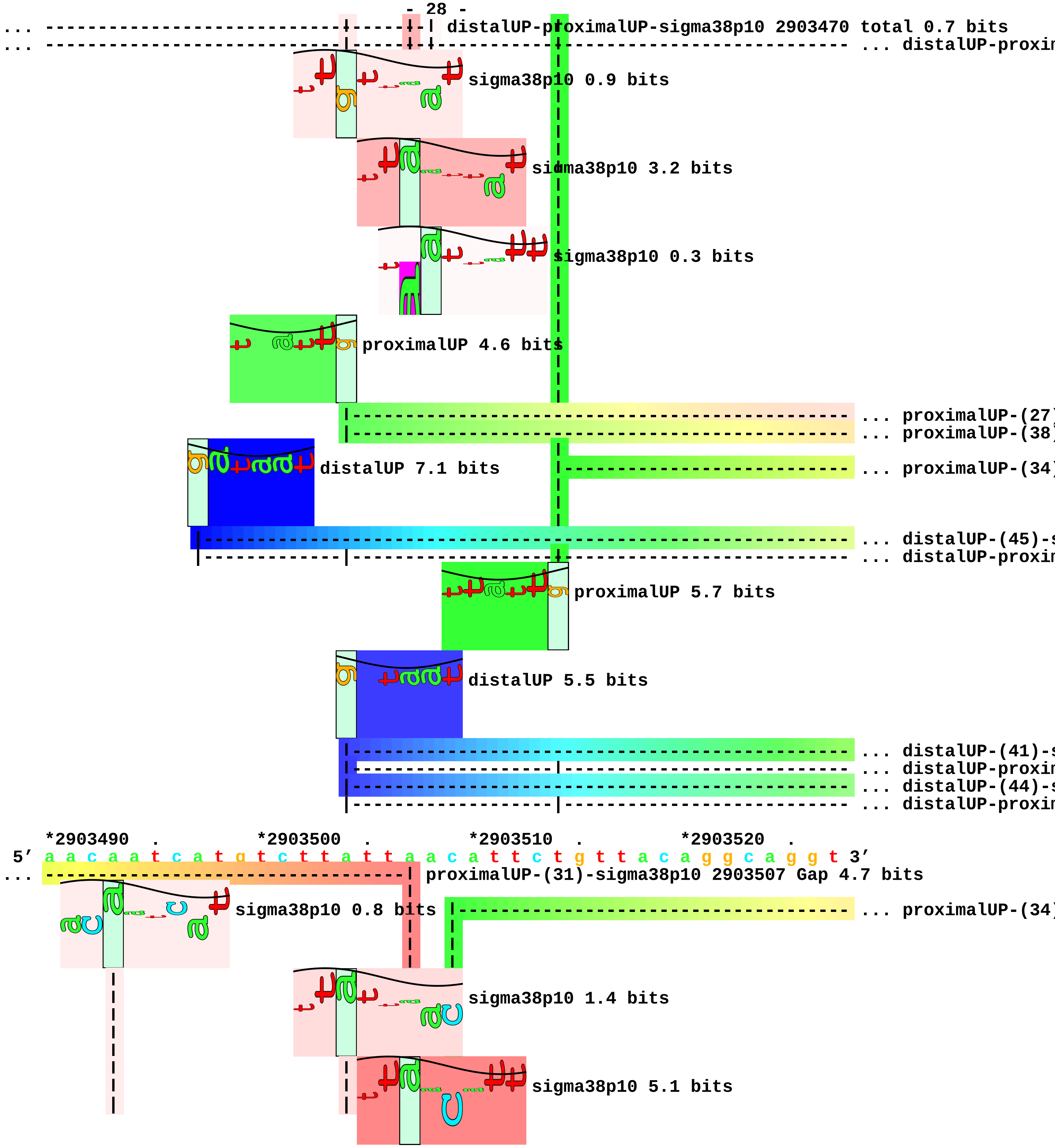

... distalUP-proximalUP-sigma38p10 2903470 total 0.7 bits
... distalUP-proxin
sigma38p10 0.9 bits
sigma38p10 3.2 bits
sigma38p10 0.3 bits
proximalUP 4.6 bits
... proximalUP-(27)
... proximalUP-(38)
distalUP 7.1 bits
... proximalUP-(34)
... distalUP-(45)-s
... distalUP-proxin
proximalUP 5.7 bits
distalUP 5.5 bits
... distalUP-(41)-s
... distalUP-proxin
... distalUP-(44)-s
... distalUP-proxin
*2903490 . *2903500 . *2903510 . *2903520 .
5' a a c a a t c a t g t c t t a t t a a c a t t c t g t t a c a g g c a g g t 3'
... proximalUP-(31)-sigma38p10 2903507 Gap 4.7 bits
sigma38p10 0.8 bits
... proximalUP-(34)
sigma38p10 1.4 bits
sigma38p10 5.1 bits

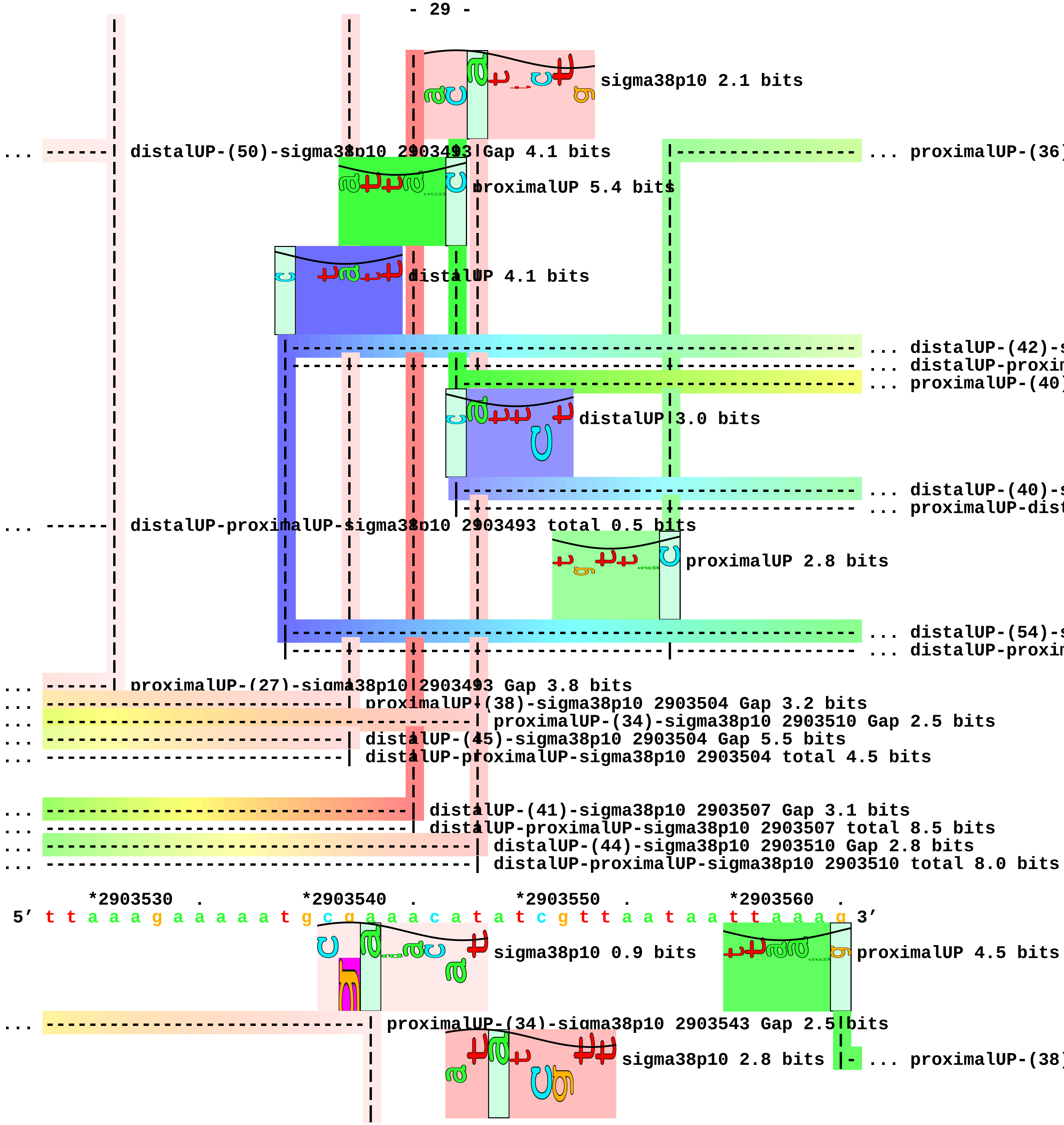
sigma38p10 2.1 bits
... ------ distalUP-(50)-sigma38p10 2903493 Gap 4.1 bits
... proximalUP-(36)
proximalUP 5.4 bits
distalUP 4.1 bits
... distalUP-(42)-s
... distalUP-proxim
... proximalUP-(40)
distalUP 3.0 bits
... distalUP-(40)-s
... proximalUP-dist
... ------ distalUP-proximalUP-sigma38p10 2903493 total 0.5 bits
proximalUP 2.8 bits
... distalUP-(54)-s
... distalUP-proxim
... ------ proximalUP-(27)-sigma38p10 2903493 Gap 3.8 bits
... proximalUP-(38)-sigma38p10 2903504 Gap 3.2 bits
... proximalUP-(34)-sigma38p10 2903510 Gap 2.5 bits
... distalUP-(45)-sigma38p10 2903504 Gap 5.5 bits
... distalUP-proximalUP-sigma38p10 2903504 total 4.5 bits
... distalUP-(41)-sigma38p10 2903507 Gap 3.1 bits
... distalUP-proximalUP-sigma38p10 2903507 total 8.5 bits
... distalUP-(44)-sigma38p10 2903510 Gap 2.8 bits
... distalUP-proximalUP-sigma38p10 2903510 total 8.0 bits
*2903530 . *2903540 . *2903550 . *2903560 .
5′ t t a a a g a a a a a t g c g a a a c a t a t c g t t a a t a a t t a a a g 3′
sigma38p10 0.9 bits
proximalUP 4.5 bits
... proximalUP-(34)-sigma38p10 2903543 Gap 2.5 bits
sigma38p10 2.8 bits
... proximalUP-(38)

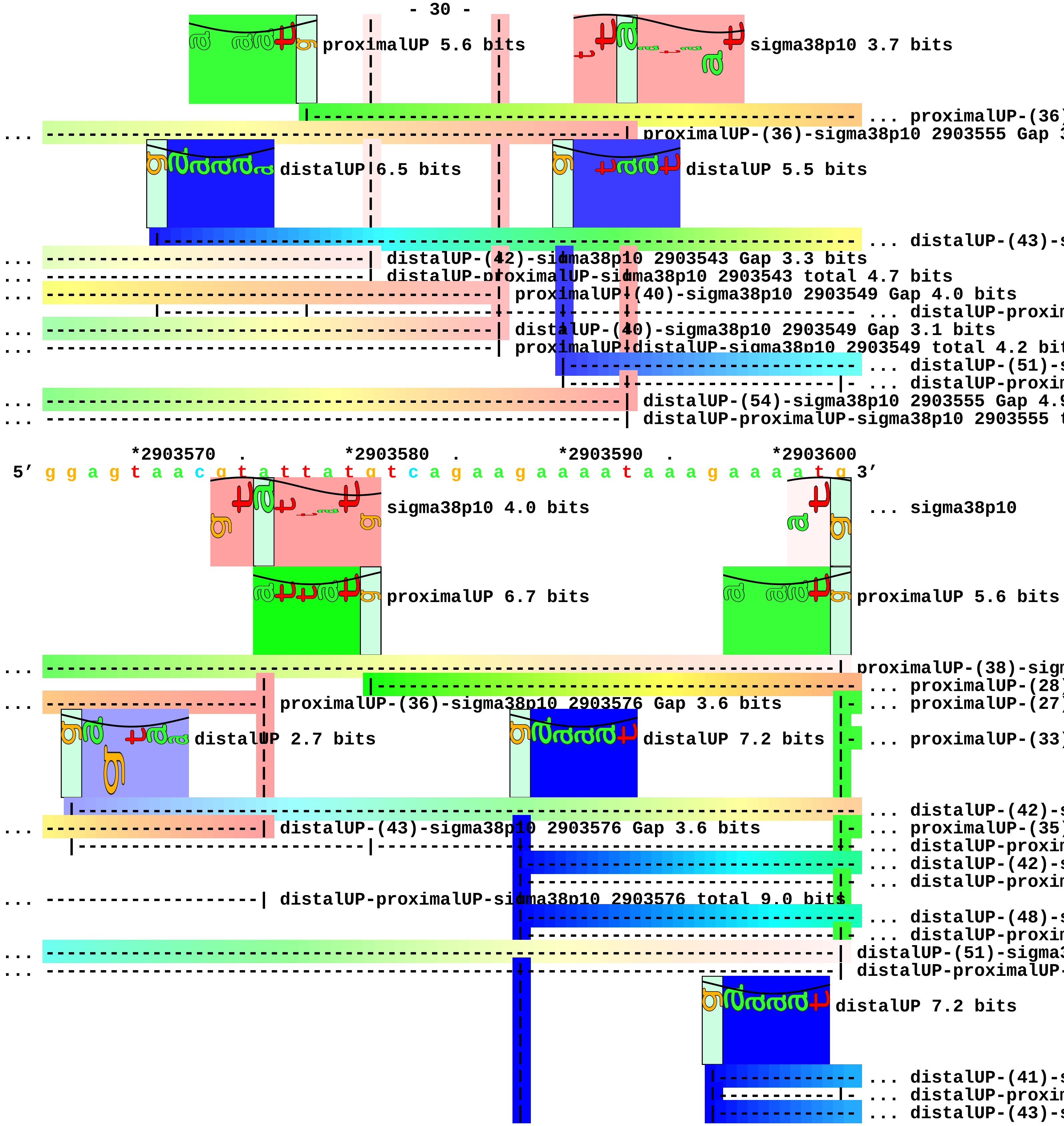
proximalUP 5.6 bits
sigma38p10 3.7 bits
... proximalUP-(36)
... proximalUP-(36)-sigma38p10 2903555 Gap 3
distalUP 6.5 bits
distalUP 5.5 bits
... distalUP-(43)-s
... distalUP-(42)-sigma38p10 2903543 Gap 3.3 bits
... distalUP-proximalUP-sigma38p10 2903543 total 4.7 bits
... proximalUP-(40)-sigma38p10 2903549 Gap 4.0 bits
... distalUP-proxim
... distalUP-(40)-sigma38p10 2903549 Gap 3.1 bits
... proximalUP-distalUP-sigma38p10 2903549 total 4.2 bit
... distalUP-(51)-s
... distalUP-proxim
... distalUP-(54)-sigma38p10 2903555 Gap 4.9
... distalUP-proximalUP-sigma38p10 2903555 t
*2903570 . *2903580 . *2903590 . *2903600
5' g g a g t a a c g t a t t a t g t c a g a a g a a a a t a a a g a a a a t g 3'
sigma38p10 4.0 bits
... sigma38p10
proximalUP 6.7 bits
proximalUP 5.6 bits
... proximalUP-(38)-sigm
... proximalUP-(28)
... proximalUP-(36)-sigma38p10 2903576 Gap 3.6 bits
... proximalUP-(27)
distalUP 2.7 bits
distalUP 7.2 bits
... proximalUP-(33)
... distalUP-(42)-s
... distalUP-(43)-sigma38p10 2903576 Gap 3.6 bits
... proximalUP-(35)
... distalUP-proxim
... distalUP-(42)-s
... distalUP-proxim
... distalUP-proximalUP-sigma38p10 2903576 total 9.0 bits
... distalUP-(48)-s
... distalUP-proxim
... distalUP-(51)-sigma3
... distalUP-proximalUP-
distalUP 7.2 bits
... distalUP-(41)-s
... distalUP-proxim
... distalUP-(43)-s

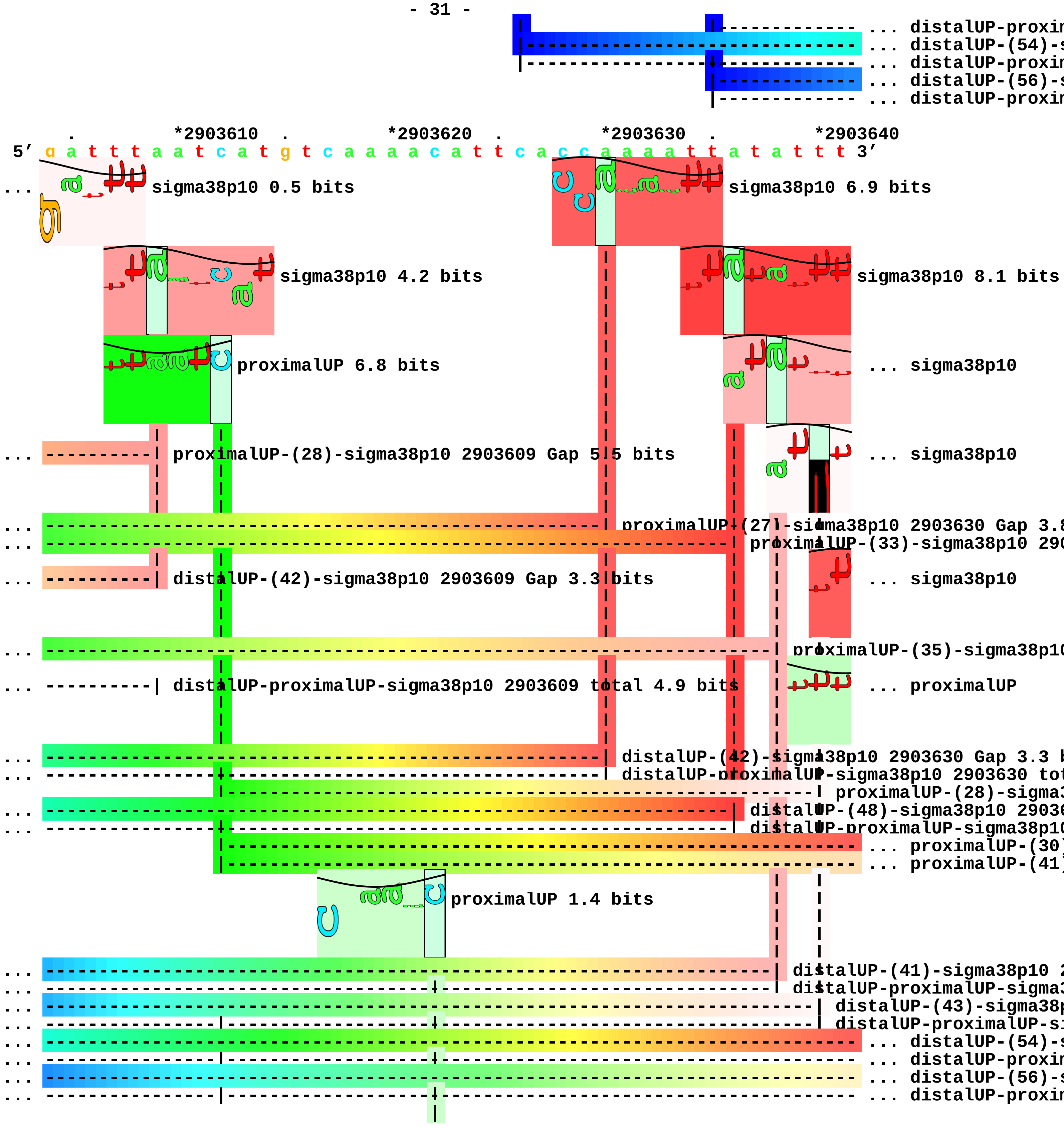
... distalUP-proxim
... distalUP-(54)-s
... distalUP-proxim
... distalUP-(56)-s
... distalUP-proxim
*2903610
*2903620
*2903630
*2903640
5′ g a t t t a a t c a t g t c a a a a c a t t c a c c a a a a t t a t a t t t 3′
sigma38p10 0.5 bits
sigma38p10 6.9 bits
sigma38p10 4.2 bits
sigma38p10 8.1 bits
proximalUP 6.8 bits
... sigma38p10
proximalUP-(28)-sigma38p10 2903609 Gap 5.5 bits
... sigma38p10
proximalUP-(27)-sigma38p10 2903630 Gap 3.8
proximalUP-(33)-sigma38p10 290
distalUP-(42)-sigma38p10 2903609 Gap 3.3 bits
... sigma38p10
proximalUP-(35)-sigma38p10
distalUP-proximalUP-sigma38p10 2903609 total 4.9 bits
... proximalUP
distalUP-(42)-sigma38p10 2903630 Gap 3.3 b
distalUP-proximalUP-sigma38p10 2903630 tot
proximalUP-(28)-sigma3
distalUP-(48)-sigma38p10 29036
distalUP-proximalUP-sigma38p10
... proximalUP-(30)
... proximalUP-(41)
proximalUP 1.4 bits
distalUP-(41)-sigma38p10 2
distalUP-proximalUP-sigma3
distalUP-(43)-sigma38p
distalUP-proximalUP-si
... distalUP-(54)-s
... distalUP-proxim
... distalUP-(56)-s
... distalUP-proxim

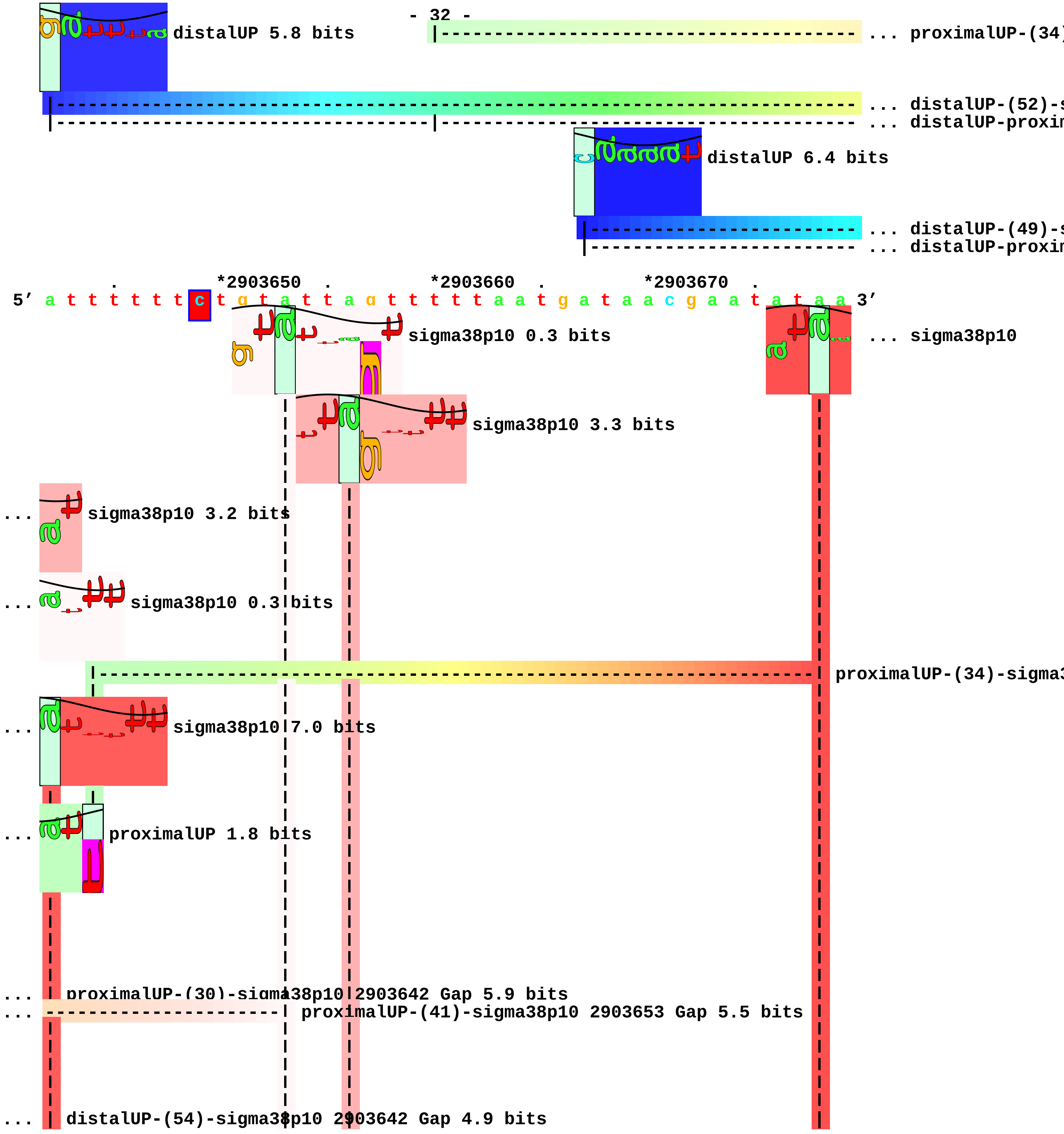
distalUP 5.8 bits
... proximalUP-(34)
... distalUP-(52)-s
... distalUP-proxim
distalUP 6.4 bits
... distalUP-(49)-s
... distalUP-proxim
*2903650
*2903660
*2903670
5′ a t t t t t t c t g t a t t a g t t t t t a a t g a t a a c g a a t a t a a 3′
sigma38p10 0.3 bits
... sigma38p10
sigma38p10 3.3 bits
... sigma38p10 3.2 bits
... sigma38p10 0.3 bits
proximalUP-(34)-sigma3
... sigma38p10 7.0 bits
... proximalUP 1.8 bits
... proximalUP-(30)-sigma38p10 2903642 Gap 5.9 bits
... proximalUP-(41)-sigma38p10 2903653 Gap 5.5 bits
... distalUP-(54)-sigma38p10 2903642 Gap 4.9 bits

```
... | distalUP-proximalUP-sigma38p10 2903642 total 10.2 bits
... ----------------------| distalUP-(56)-sigma38p10 2903653 Gap 3.8 bits
... ----------------------| distalUP-proximalUP-sigma38p10 2903653 total 5.1 bits
... ----------------------------| proximalUP-(34)-sigma38p10 2903656 Gap 2.5 bits
... ----------------------------| distalUP-(52)-sigma38p10 2903656 Gap 5.5 bits
... ----------------------------| distalUP-proximalUP-sigma38p10 2903656 total 2.6 bits

... -------------------------------------------------------------------------| distalUP-(49)-sigma38p
... ----|--------------------------------------------------------------------| distalUP-proximalUP-si

    *2903680  .         *2903690  .
 5' a a t t a c c g a t g c c g c c g t c a 3'
```

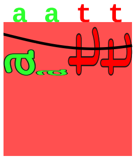

```
... sigma38p10 7.5 bits
```

piece 10, NC_000913, tisB-istR-1-istR-2, config: linear, direction: +, begin: 3851050, end: 385144

```
    *3851050   .         *3851060   .         *3851070   .         *3851080   .
5′ c a a g c a c a g c t t t a c a g g g g a g a c a a t g g a a a a t t t t t 3′
```

T timestamp:_2018Sep20_11-36-03

... proximalUP

distalUP 0.8 bits

... distalUP-(44)-s
... distalUP-proxi

distalUP 7.2 bits

... distalUP-(44)-s
... distalUP-proxi
... distalUP-(49)-s
... distalUP-proxi

```
    *3851090   .         *3851100   .         *3851110   .         *3851120   .
5′ c a g c a a g g g a a a a t t g a g g g g t t g a t c a c g t t t t g t a c 3′
```

... proximalUP 6.2 bits

sigma38p10 2.9 bits

proximalUP-(27)-sigma38p10 3851115 Gap 3
... distalUP-(44)-sigma38p10 3851115 Gap 2.8
... distalUP-proximalUP-sigma38p10 3851115 t

... sigma38p10

... distalUP-(44)-sigma38p10 3
... distalUP-proximalUP-sigma3
... ... distalUP-(49)-s
... ... distalUP-proxi
proximalUP-(34)-sigma38p10

... sigma38p10

... proximalUP-(39)

```
        *3851130   .         *3851140   .         *3851150   .         *3851160
5′ t g a a t t g c a g a t a a c a a a a a c c c c g c c g g a g c g a g g t 3′
```

... sigma38p10 3.3 bits

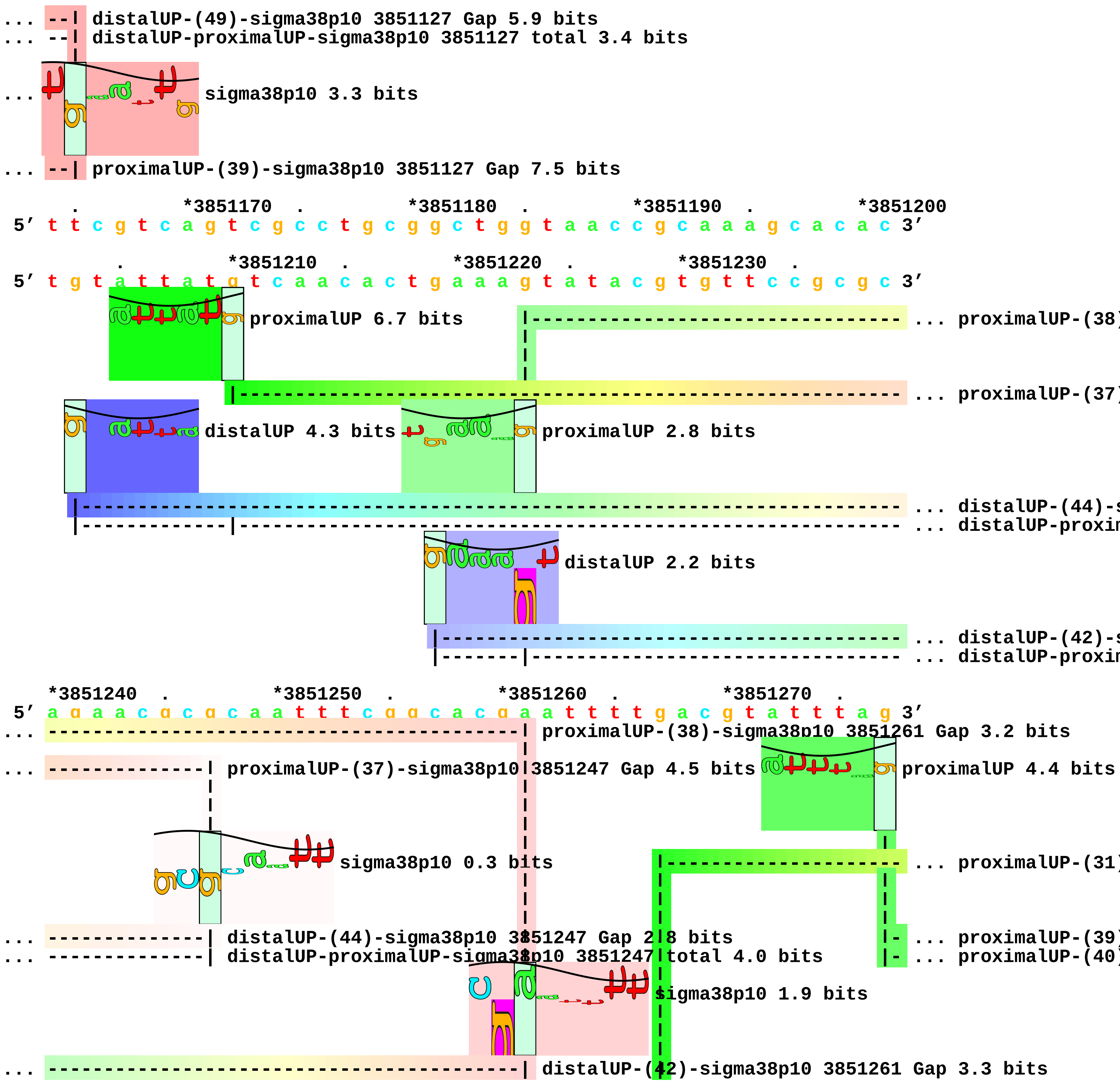
... --| distalUP-(49)-sigma38p10 3851127 Gap 5.9 bits
... --| distalUP-proximalUP-sigma38p10 3851127 total 3.4 bits
... sigma38p10 3.3 bits
... --| proximalUP-(39)-sigma38p10 3851127 Gap 7.5 bits
. *3851170 . *3851180 . *3851190 . *3851200
5' t t c g t c a g t c g c c t g c g g c t g g t a a c c g c a a a g c a c a c 3'
. *3851210 . *3851220 . *3851230 .
5' t g t a t t a t g t c a a c a c t g a a a g t a t a c g t g t t c c g c g c 3'
proximalUP 6.7 bits
... proximalUP-(38
... proximalUP-(37
distalUP 4.3 bits
proximalUP 2.8 bits
... distalUP-(44)-s
... distalUP-proxim
distalUP 2.2 bits
... distalUP-(42)-s
... distalUP-proxim
*3851240 . *3851250 . *3851260 . *3851270 .
5' a g a a c g c g c a a t t t c g g c a c g a a t t t t g a c g t a t t t a g 3'
... proximalUP-(38)-sigma38p10 3851261 Gap 3.2 bits
... proximalUP-(37)-sigma38p10 3851247 Gap 4.5 bits
proximalUP 4.4 bits
sigma38p10 0.3 bits
... proximalUP-(31
... distalUP-(44)-sigma38p10 3851247 Gap 2.8 bits
... distalUP-proximalUP-sigma38p10 3851247 total 4.0 bits
... proximalUP-(39
... proximalUP-(40
sigma38p10 1.9 bits
... distalUP-(42)-sigma38p10 3851261 Gap 3.3 bits

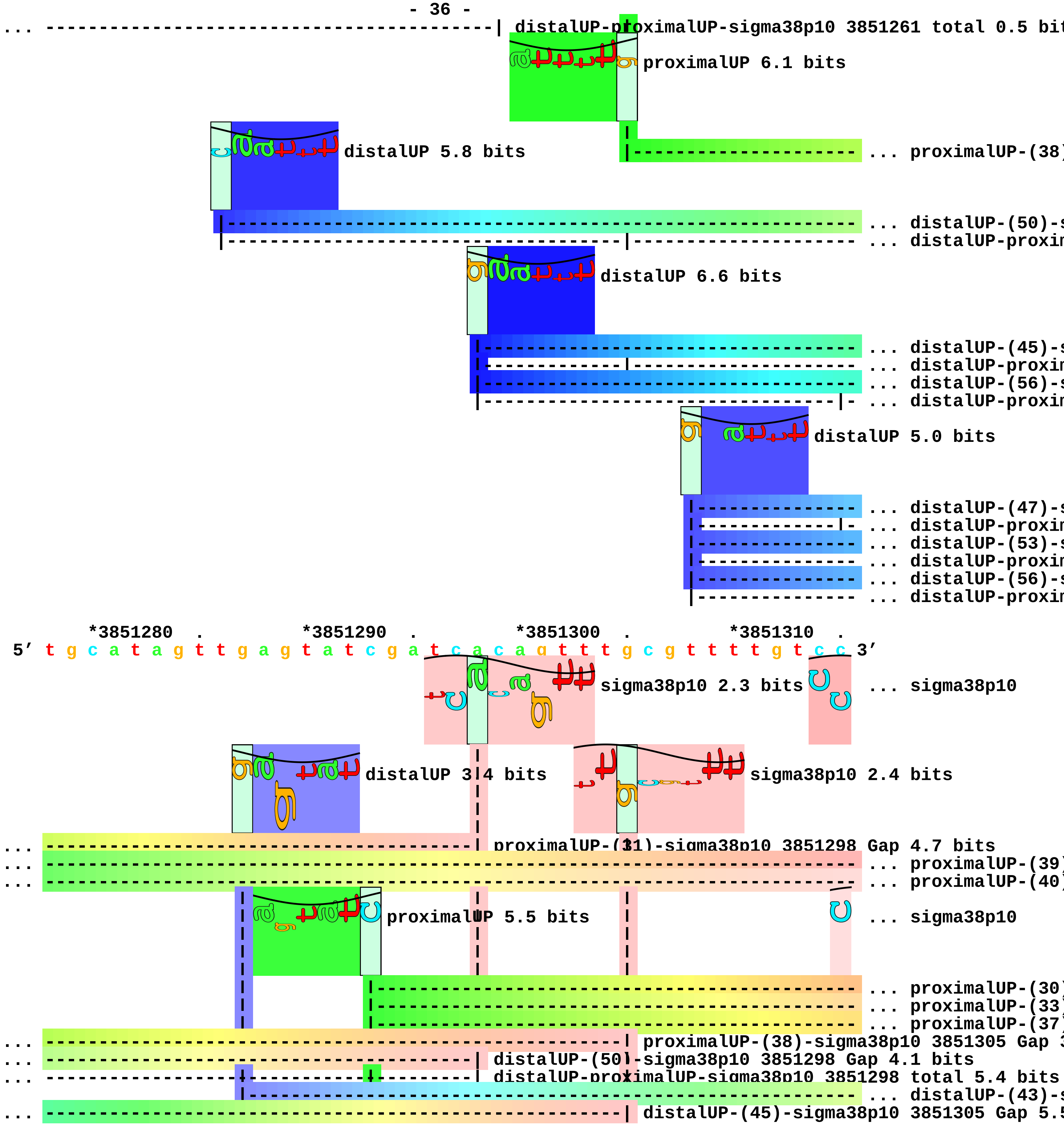
distalUP-proximalUP-sigma38p10 3851261 total 0.5 bit
proximalUP 6.1 bits
distalUP 5.8 bits
... proximalUP-(38)
... distalUP-(50)-s
... distalUP-proxim
distalUP 6.6 bits
... distalUP-(45)-s
... distalUP-proxim
... distalUP-(56)-s
... distalUP-proxim
distalUP 5.0 bits
... distalUP-(47)-s
... distalUP-proxim
... distalUP-(53)-s
... distalUP-proxim
... distalUP-(56)-s
... distalUP-proxim
*3851280 . *3851290 . *3851300 . *3851310 .
5′ t g c a t a g t t g a g t a t c g a t c a c a g t t t g c g t t t t g t c c 3′
sigma38p10 2.3 bits
... sigma38p10
distalUP 3.4 bits
sigma38p10 2.4 bits
proximalUP-(31)-sigma38p10 3851298 Gap 4.7 bits
... proximalUP-(39)
... proximalUP-(40)
proximalUP 5.5 bits
... sigma38p10
... proximalUP-(30)
... proximalUP-(33)
... proximalUP-(37)
proximalUP-(38)-sigma38p10 3851305 Gap 3
distalUP-(50)-sigma38p10 3851298 Gap 4.1 bits
distalUP-proximalUP-sigma38p10 3851298 total 5.4 bits
... distalUP-(43)-s
distalUP-(45)-sigma38p10 3851305 Gap 5.5

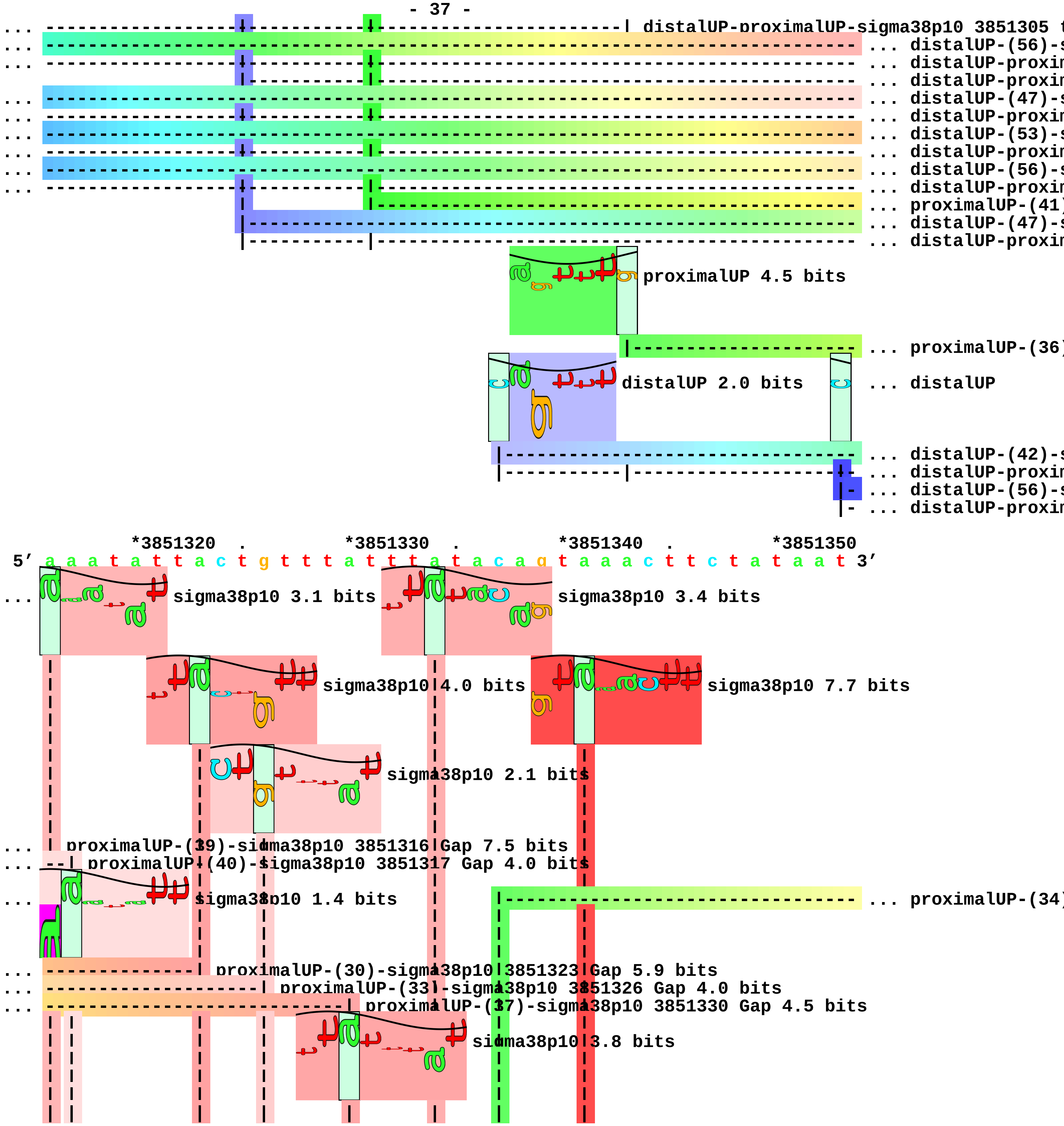
distalUP-proximalUP-sigma38p10 3851305 t
distalUP-(56)-s
distalUP-proxim
distalUP-proxim
distalUP-(47)-s
distalUP-proxim
distalUP-(53)-s
distalUP-proxim
distalUP-(56)-s
distalUP-proxim
proximalUP-(41)
distalUP-(47)-s
distalUP-proxim
proximalUP 4.5 bits
proximalUP-(36)
distalUP 2.0 bits
distalUP
distalUP-(42)-s
distalUP-proxim
distalUP-(56)-s
distalUP-proxim
*3851320
*3851330
*3851340
*3851350
5′ a a a t a t t a c t g t t t a t t t a t a c a g t a a a c t t c t a t a a t 3′
sigma38p10 3.1 bits
sigma38p10 3.4 bits
sigma38p10 4.0 bits
sigma38p10 7.7 bits
sigma38p10 2.1 bits
proximalUP-(39)-sigma38p10 3851316 Gap 7.5 bits
proximalUP-(40)-sigma38p10 3851317 Gap 4.0 bits
sigma38p10 1.4 bits
proximalUP-(34)
proximalUP-(30)-sigma38p10 3851323 Gap 5.9 bits
proximalUP-(33)-sigma38p10 3851326 Gap 4.0 bits
proximalUP-(37)-sigma38p10 3851330 Gap 4.5 bits
sigma38p10 3.8 bits

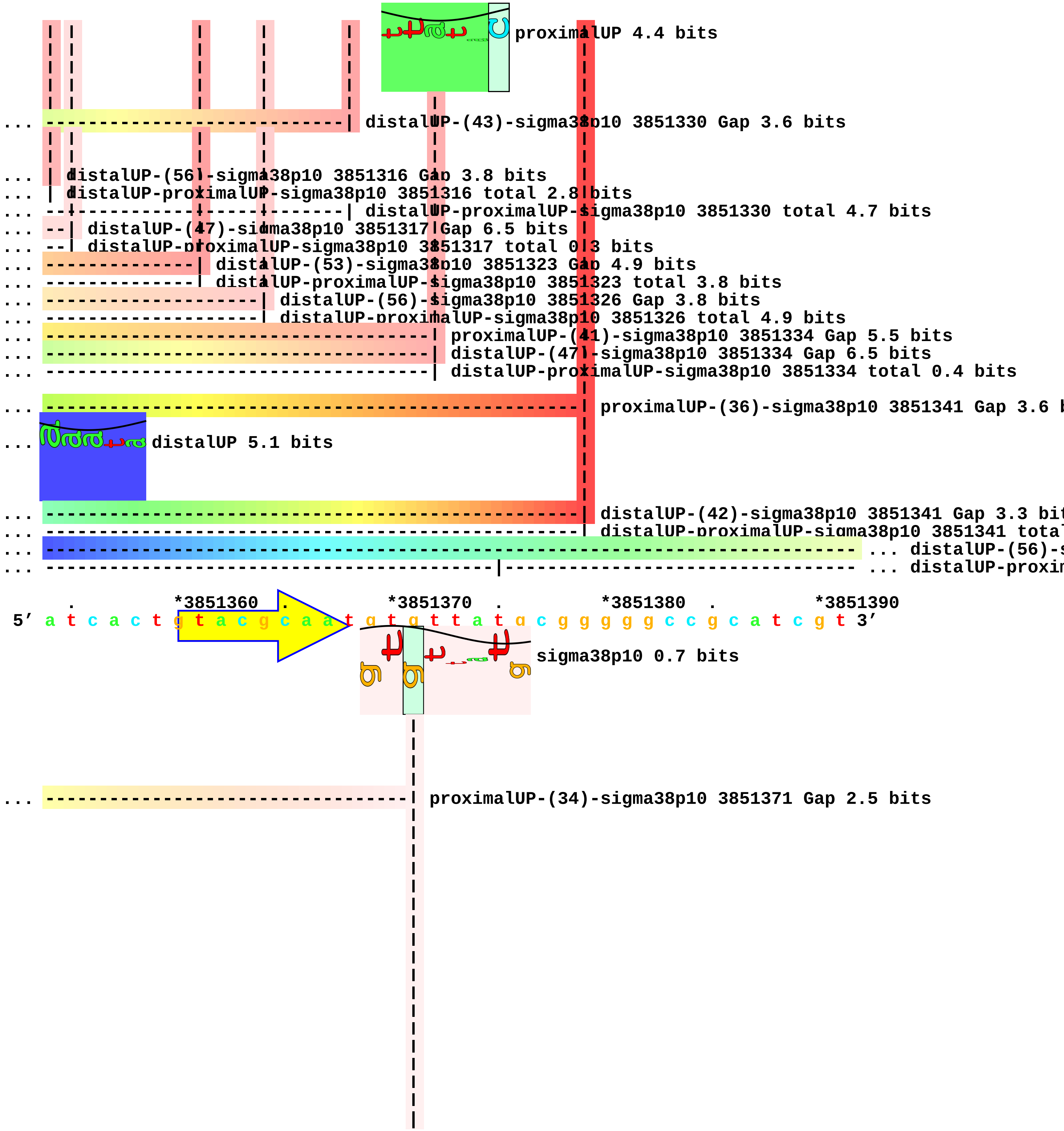
proximalUP 4.4 bits
distalUP-(43)-sigma38p10 3851330 Gap 3.6 bits
distalUP-(56)-sigma38p10 3851316 Gap 3.8 bits
distalUP-proximalUP-sigma38p10 3851316 total 2.8 bits
distalUP-proximalUP-sigma38p10 3851330 total 4.7 bits
distalUP-(47)-sigma38p10 3851317 Gap 6.5 bits
distalUP-proximalUP-sigma38p10 3851317 total 0.3 bits
distalUP-(53)-sigma38p10 3851323 Gap 4.9 bits
distalUP-proximalUP-sigma38p10 3851323 total 3.8 bits
distalUP-(56)-sigma38p10 3851326 Gap 3.8 bits
distalUP-proximalUP-sigma38p10 3851326 total 4.9 bits
proximalUP-(41)-sigma38p10 3851334 Gap 5.5 bits
distalUP-(47)-sigma38p10 3851334 Gap 6.5 bits
distalUP-proximalUP-sigma38p10 3851334 total 0.4 bits
proximalUP-(36)-sigma38p10 3851341 Gap 3.6 b
distalUP 5.1 bits
distalUP-(42)-sigma38p10 3851341 Gap 3.3 bit
distalUP-proximalUP-sigma38p10 3851341 total
... distalUP-(56)-s
... distalUP-proxim
*3851360
*3851370
*3851380
*3851390
5′ a t c a c t g t a c g c a a t g t g t t a t g c g g g g g c c g c a t c g t 3′
sigma38p10 0.7 bits
proximalUP-(34)-sigma38p10 3851371 Gap 2.5 bits

```
... ----------------------------------| distalUP-(56)-sigma38p10 3851371 Gap 3.8 bits
... ----------------------------------| distalUP-proximalUP-sigma38p10 3851371 total 4.0 bits

           .         *3851400  .         *3851410  .         *3851420  .
 5' t a c c c g g c g c a c t a a g t c c t g g c t g a a a c g g g t g g t g c 3'

    *3851430  .         *3851440  .
 5' c g t c a g c g c c t t a a c c c c g c 3'
```

piece 11, NC_000913, rpmE-priA, config: linear, direction: +, begin: 4124800, end: 4125099

```
   *4124800  .         *4124810  .         *4124820  .         *4124830  .
5' g c g g a a c g g g c a a g g c a a c g t g g g c a a c g g g c a t a g c a 3'
   T timestamp:_2018Sep20_11-36-03

     *4124840  .         *4124850  .         *4124860  .         *4124870  .
5' t c a t c c t g a c t t g a a a t t c g g t g g g t t a g t a t a c a c a t 3'

       *4124880  .         *4124890  .         *4124900  .         *4124910
5' t g c c g t a g a a g a g t g c g g a t c a g t t t g c a t a c g c t g g t 3'
                                                                     ... distalUP

                                                                     |--- ... distalUP-(52)-s
                                                                     |--- ... distalUP-proxim
                                                                     |--- ... distalUP-(54)-s
                                                                     |--- ... distalUP-proxim

    .         *4124920  .         *4124930  .         *4124940  .         *4124950
5' t a a t t t c t g t a t g a t t t c g c g c c t t c g t a c g a a a t g a t 3'
... distalUP 5.5 bits

... ---------------------------------------------------------------------- ... distalUP-(52)-s
... ------------------------|--------------------------------------------- ... distalUP-proxim
... ---------------------------------------------------------------------- ... distalUP-(54)-s
... ------------------------|--------------------------------------------- ... distalUP-proxim
              proximalUP 4.6 bits

                        |--------------------------------------------- ... proximalUP-(38)
                        |--------------------------------------------- ... proximalUP-(40)

      .         *4124960  .         *4124970  .         *4124980  .
5' c g t a t t g a a g c t a t a c t t t t a a c a t c g c g t g g t g t c t g 3'
                 sigma38p10 10.9 bits

... ------------------------| distalUP-(52)-sigma38p10 4124964 Gap 5.5 bits
... ------------------------| distalUP-proximalUP-sigma38p10 4124964 total 12.4 bits
... ----------------------------| distalUP-(54)-sigma38p10 4124966 Gap 4.9 bits
... ----------------------------| distalUP-proximalUP-sigma38p10 4124966 total 6.5 bits
                   sigma38p10 5.2 bits

... ------------------------| proximalUP-(38)-sigma38p10 4124964 Gap 3.2 bits
... ----------------------------| proximalUP-(40)-sigma38p10 4124966 Gap 4.0 bits

   *4124990  .         *4125000  .         *4125010  .         *4125020  .
5' g c g t t a g g g c t g g a a g a g c g a c g c g g c c t t a a a c c g a g 3'
```

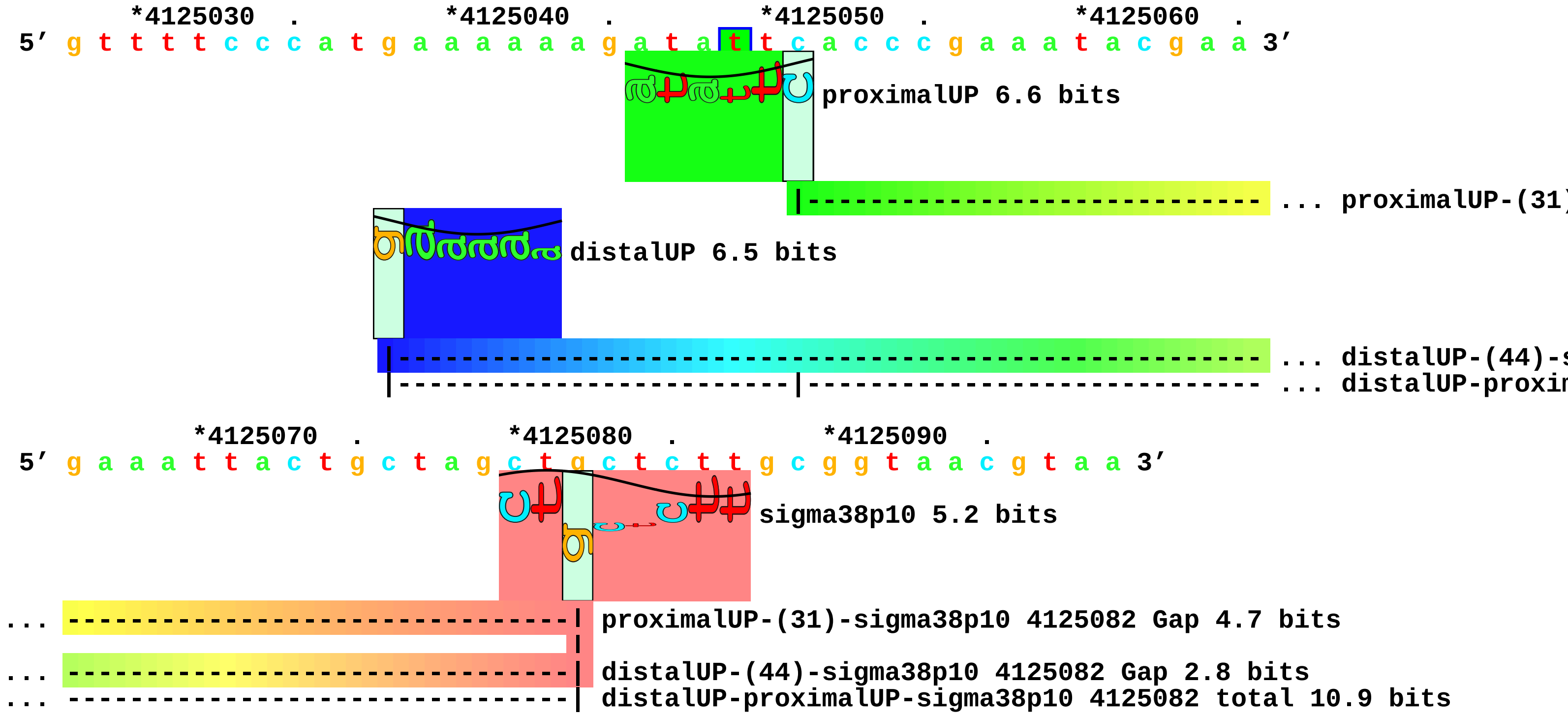
*4125030 . *4125040 . *4125050 . *4125060 .
5′ g t t t t c c c a t g a a a a a a g a t a t t c a c c c g a a a t a c g a a 3′
proximalUP 6.6 bits
... proximalUP-(31)
distalUP 6.5 bits
... distalUP-(44)-s
... distalUP-proxi
*4125070 . *4125080 . *4125090 .
5′ g a a a t t a c t g c t a g c t g c t c t t g c g g t a a c g t a a 3′
sigma38p10 5.2 bits
... proximalUP-(31)-sigma38p10 4125082 Gap 4.7 bits
... distalUP-(44)-sigma38p10 4125082 Gap 2.8 bits
... distalUP-proximalUP-sigma38p10 4125082 total 10.9 bits

piece 12, NC_000913, yjfP-bsmA, config: linear, direction: +, begin: 4414600, end: 4414949

```
   *4414600  .         *4414610  .         *4414620  .         *4414630  .
5' c c c c g a a c c a t c g t g c t t a c g c t a c c t a t t c g c t g t a a 3'
   T timestamp:_2018Sep20_11-36-03

     *4414640  .         *4414650  .         *4414660  .         *4414670  .
5' c c c t t g c g t c t g g t c g c g g c g a a t c t c t t g c g g a t g g t 3'

       *4414680  .         *4414690  .         *4414700  .         *4414710
5' c c g t t a c t g g c g g t g c t g g c t g t g g c g t a c c t t g c a g t 3'

   .         *4414720  .         *4414730  .         *4414740  .         *4414750
5' g c g c t a c a g g c a c t t a a c a t c a a c a c c a a t a a t a a a c t 3'
                                                                 proximalUP 5.4 bits
                                                                 ... proximalUP-(38)
                                             distalUP 5.7 bits
                                                                 ... distalUP-(48)-s
                                                                 ... distalUP-proxim
                                                                 ... proximalUP-(42)
                                                                 ... distalUP-(52)-s
                                                                 ... distalUP-proxim

     .         *4414760  .         *4414770  .         *4414780  .
5' g g c a a a c c g g t a a a t a a c g c t a t t a c g t t t c c t g c t a a 3'
                 proximalUP 4.3 bits                             ... sigma38p10
...                                                              proximalUP-(38)-sigma3
                                                                 ... proximalUP-(38)
...                                                              distalUP-(48)-sigma38p
...                                                              distalUP-proximalUP-si
...                                                              ... proximalUP-(42)
...                                                              ... distalUP-(52)-s
...                                                              ... distalUP-proxim
            distalUP 5.6 bits                                    ... proximalUP-(37)
                                                                 ... distalUP-(46)-s
                                                                 ... distalUP-proxim
                             proximalUP 3.3 bits
                                                                 ... distalUP-(53)-s
                                                                 ... distalUP-proxim
                                                                 ... proximalUP-(38)
                                                                 ... distalUP-(54)-s
```

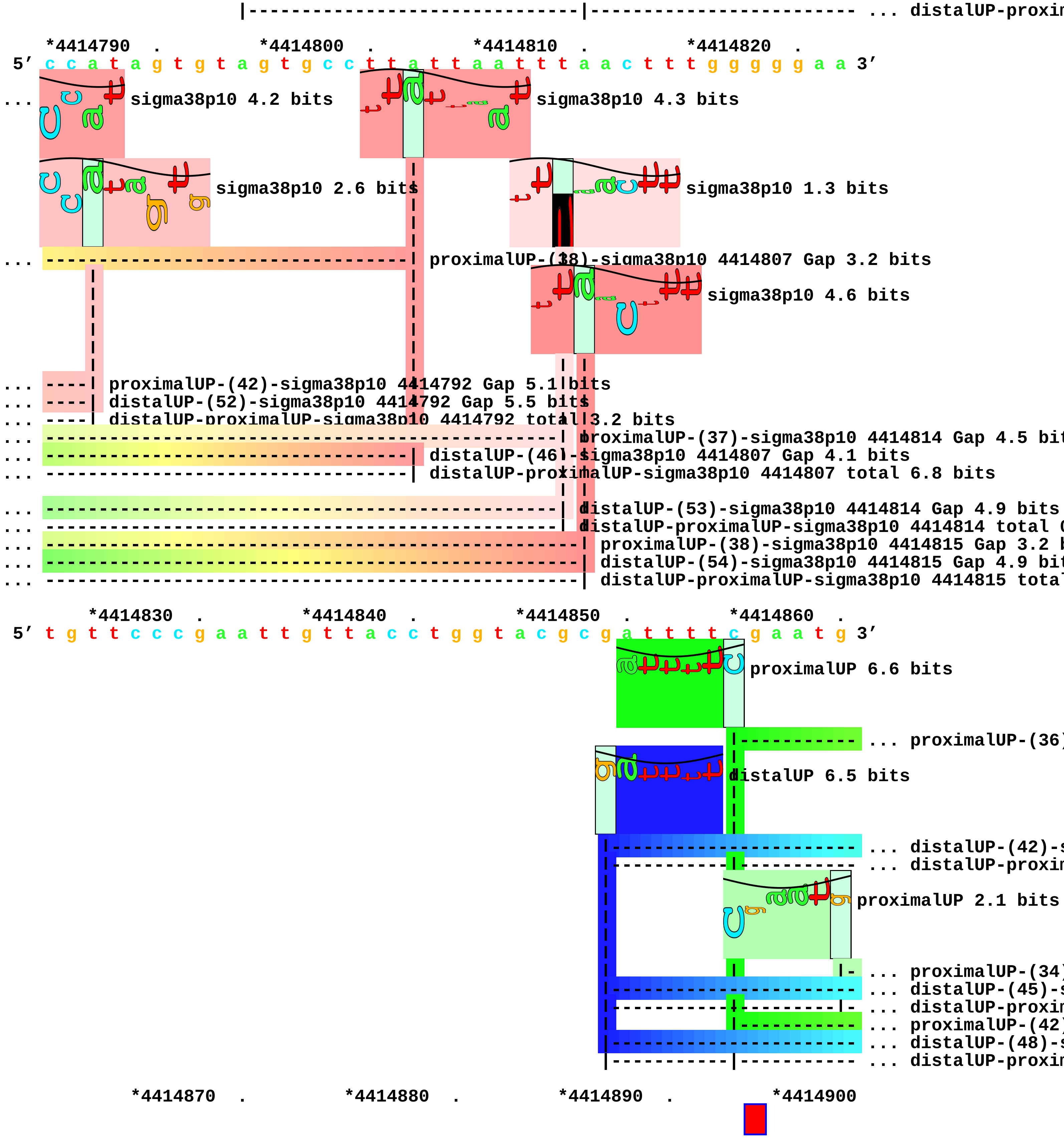
... distalUP-proxim
*4414790 . *4414800 . *4414810 . *4414820 .
5′ c c a t a g t g t a g t g c c t t a t t a a t t t a a c t t t g g g g g a a 3′
sigma38p10 4.2 bits
sigma38p10 4.3 bits
sigma38p10 2.6 bits
sigma38p10 1.3 bits
proximalUP-(38)-sigma38p10 4414807 Gap 3.2 bits
sigma38p10 4.6 bits
proximalUP-(42)-sigma38p10 4414792 Gap 5.1 bits
distalUP-(52)-sigma38p10 4414792 Gap 5.5 bits
distalUP-proximalUP-sigma38p10 4414792 total 3.2 bits
proximalUP-(37)-sigma38p10 4414814 Gap 4.5 bit
distalUP-(46)-sigma38p10 4414807 Gap 4.1 bits
distalUP-proximalUP-sigma38p10 4414807 total 6.8 bits
distalUP-(53)-sigma38p10 4414814 Gap 4.9 bits
distalUP-proximalUP-sigma38p10 4414814 total
proximalUP-(38)-sigma38p10 4414815 Gap 3.2 b
distalUP-(54)-sigma38p10 4414815 Gap 4.9 bit
distalUP-proximalUP-sigma38p10 4414815 total
*4414830 . *4414840 . *4414850 . *4414860 .
5′ t g t t c c c g a a t t g t t a c c t g g t a c g c g a t t t t c g a a t g 3′
proximalUP 6.6 bits
... proximalUP-(36)
distalUP 6.5 bits
... distalUP-(42)-
... distalUP-proxim
proximalUP 2.1 bits
... proximalUP-(34)
... distalUP-(45)-
... distalUP-proxim
... proximalUP-(42)
... distalUP-(48)-
... distalUP-proxim
*4414870 . *4414880 . *4414890 . *4414900

5′ a a a a c g t g a c g a a a g c g t a a c c c a c a t c t c a a c a t t a t 3′
<-------------------------------------- ... bsmA
... ------------------------------------------------------------| proximalUP-(36)-sigma38p10 4414896
sigma38p10 2.2 bits
... ------------------------------------------------------------| distalUP-(42)-sigma38p10 4414896 0
... ------------------------------------------------------------| distalUP-proximalUP-sigma38p10 441
... sigma38p10
... ------------------------------------------------------------------| proximalUP-(34)-sigma38p10 4
... ------------------------------------------------------------------| distalUP-(45)-sigma38p10 441
... ------------------------------------------------------------------| distalUP-proximalUP-sigma38p
... ------------------------------------------------------------------------| proximalUP-(42)-sigma3
... ------------------------------------------------------------------------| distalUP-(48)-sigma38p
... ------------------------------------------------------------------------| distalUP-proximalUP-si
... sigma38p10

. *4414910 . *4414920 . *4414930 . *4414940
5′ g a a c t a a t a c c g c c a g c a g a a a c t t g t g c t a a g g a t t t 3′
... #---------] bsmA

... sigma38p10 1.5 bits

... sigma38p10 1.4 bits

.
5′ c t t c a t g t 3′

piece 13, NC_000913, cysQ-cpdB, config: linear, direction: +, begin: 4434350, end: 4434799

```
   *4434350  .         *4434360  .         *4434370  .         *4434380  .
5′ c c c t g a a t c a a a t c g c c g t t a t c a a c c a g t a c g c t g t t 3′
   T timestamp:_2018Sep20_11-36-03

     *4434390  .         *4434400  .         *4434410  .         *4434420  .
5′ t t t c a c t t c a t t g c g g g c a t c g t t a a t c a g g c t t g c c g 3′
                                               proximalUP 6.8 bits
                                                            ... proximalUP-(42)
                                             distalUP 5.5 bits
                                                            ... distalUP-(48)-s
                                                            ... distalUP-proxim

       *4434430  .         *4434440  .         *4434450  .         *4434460
5′ t a c g t a c c a g t c c g a a t t t t t c c g t g g c g g t g t c t t t g 3′
                               distalUP 6.6 bits              sigma38p10 0.9 bits
...                                                    proximalUP-(42)-sigma38p10 44344
                                     proximalUP 6.2 bits
...                                                    distalUP-(48)-sigma38p10 4434457
...                                                    distalUP-proximalUP-sigma38p10 4
                                                            ... proximalUP-(32)
                                                            ... distalUP-(40)-s
                                                            ... distalUP-proxim
                                                            ... proximalUP-(35)
                                                            ... distalUP-(43)-s
                                                            ... distalUP-proxim
                                                            ... proximalUP-(40)
                                                            ... distalUP-(48)-s
                                                            ... distalUP-proxim
                                                            ... proximalUP-(43)
                                                            ... distalUP-(51)-s
                                                            ... distalUP-proxim

    .         *4434470  .         *4434480  .         *4434490  .         *4434500
5′ t a a t a a t c g a a a t c c a t c a t g t t g c t a t g c a g a t c a g t 3′
                                       sigma38p10 2.0 bits
                                          sigma38p10 1.5 bits
```

```
proximalUP 7.2 bits                          sigma38p10 0.4 bits
distalUP 2.8 bits                            sigma38p10 3.9 bits
                                                              ... proximalUP-(43)
... ---------------------------------| proximalUP-(32)-sigma38p10 4434479 Gap 4.3 bits
... ---------------------------------| distalUP-(40)-sigma38p10 4434479 Gap 3.1 bits
... -------------------------------- | distalUP-proximalUP-sigma38p10 4434479 total 7.4 bits
... ------------------------------------| proximalUP-(35)-sigma38p10 4434482 Gap 3.1 bits
... ------------------------------------| distalUP-(43)-sigma38p10 4434482 Gap 3.6 bits
... ------------------------------------| distalUP-proximalUP-sigma38p10 4434482 total 7.7 bits
... ------------------------------------------| proximalUP-(40)-sigma38p10 4434487 Gap 4.0 bits
... ------------------------------------------| distalUP-(48)-sigma38p10 4434487 Gap 7.5 bits
... ------------------------------------------| distalUP-proximalUP-sigma38p10 4434487 total 1.7
... ---------------------------------------------| proximalUP-(43)-sigma38p10 4434490 Gap 5.1
... ---------------------------------------------| distalUP-(51)-sigma38p10 4434490 Gap 5.1 b
... ---------------------------------------------| distalUP-proximalUP-sigma38p10 4434490 tot
          |------------------------------------------------ ... distalUP-(42)-s
        |-|------------------------------------------------ ... proximalUP-dist

       .      *4434510  .       *4434520  .       *4434530  .
5' g g t t t c c a t g a t a c g c a g a t c g a c c g t c g c t g c a t t c a 3'
                                   sigma38p10 0.5 bits

... ------------------------| proximalUP-(43)-sigma38p10 4434514 Gap 5.1 bits

... ------------------------| distalUP-(42)-sigma38p10 4434514 Gap 3.3 bits
... ------------------------| proximalUP-distalUP-sigma38p10 4434514 total 2.0 bits

   *4434540  .       *4434550  .       *4434560  .       *4434570  .
5' c a c t g g c g g c a a t c a g c g t g g c c a g g a g c g t t g c g c t a 3'

     *4434580  .       *4434590  .       *4434600  .       *4434610  .
5' a a c t t a a t c a t c a g g g a c a t c c t t t t a t c a t c g g g a a t 3'
       distalUP 4.7 bits                          proximalUP 6.8 bits
```

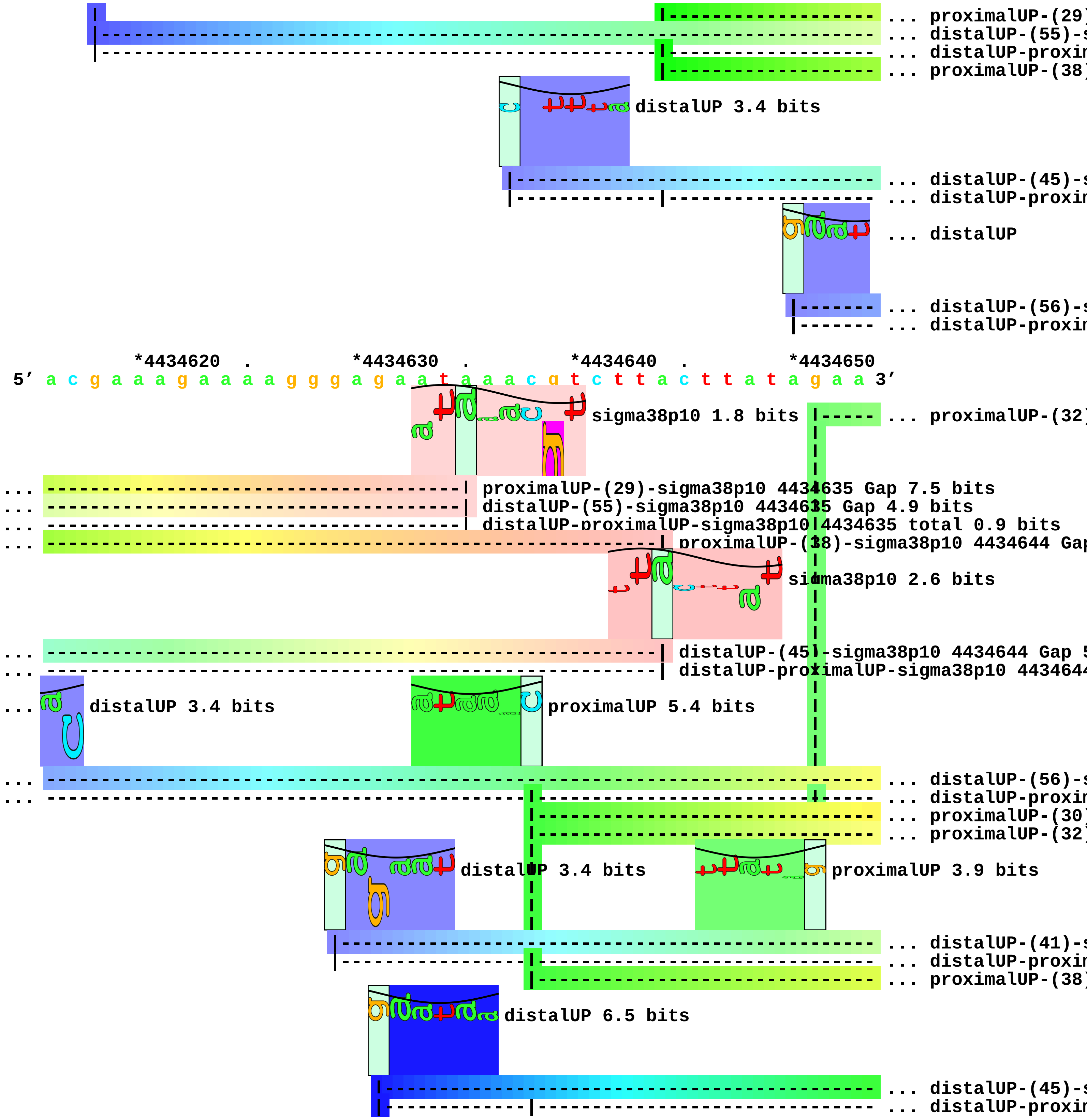
... proximalUP-(29)
... distalUP-(55)-s
... distalUP-proxin
... proximalUP-(38)
distalUP 3.4 bits
... distalUP-(45)-s
... distalUP-proxin
... distalUP
... distalUP-(56)-s
... distalUP-proxin
*4434620 . *4434630 . *4434640 . *4434650
5′ a c g a a a g a a a a g g g a g a a t a a a c g t c t t a c t t a t a g a a 3′
sigma38p10 1.8 bits
... proximalUP-(32)
... proximalUP-(29)-sigma38p10 4434635 Gap 7.5 bits
... distalUP-(55)-sigma38p10 4434635 Gap 4.9 bits
... distalUP-proximalUP-sigma38p10 4434635 total 0.9 bits
... proximalUP-(38)-sigma38p10 4434644 Gap
sigma38p10 2.6 bits
... distalUP-(45)-sigma38p10 4434644 Gap 5
... distalUP-proximalUP-sigma38p10 443464
... distalUP 3.4 bits
proximalUP 5.4 bits
... distalUP-(56)-s
... distalUP-proxin
... proximalUP-(30)
... proximalUP-(32)
distalUP 3.4 bits
proximalUP 3.9 bits
... distalUP-(41)-s
... distalUP-proxin
... proximalUP-(38)
distalUP 6.5 bits
... distalUP-(45)-s
... distalUP-proxin

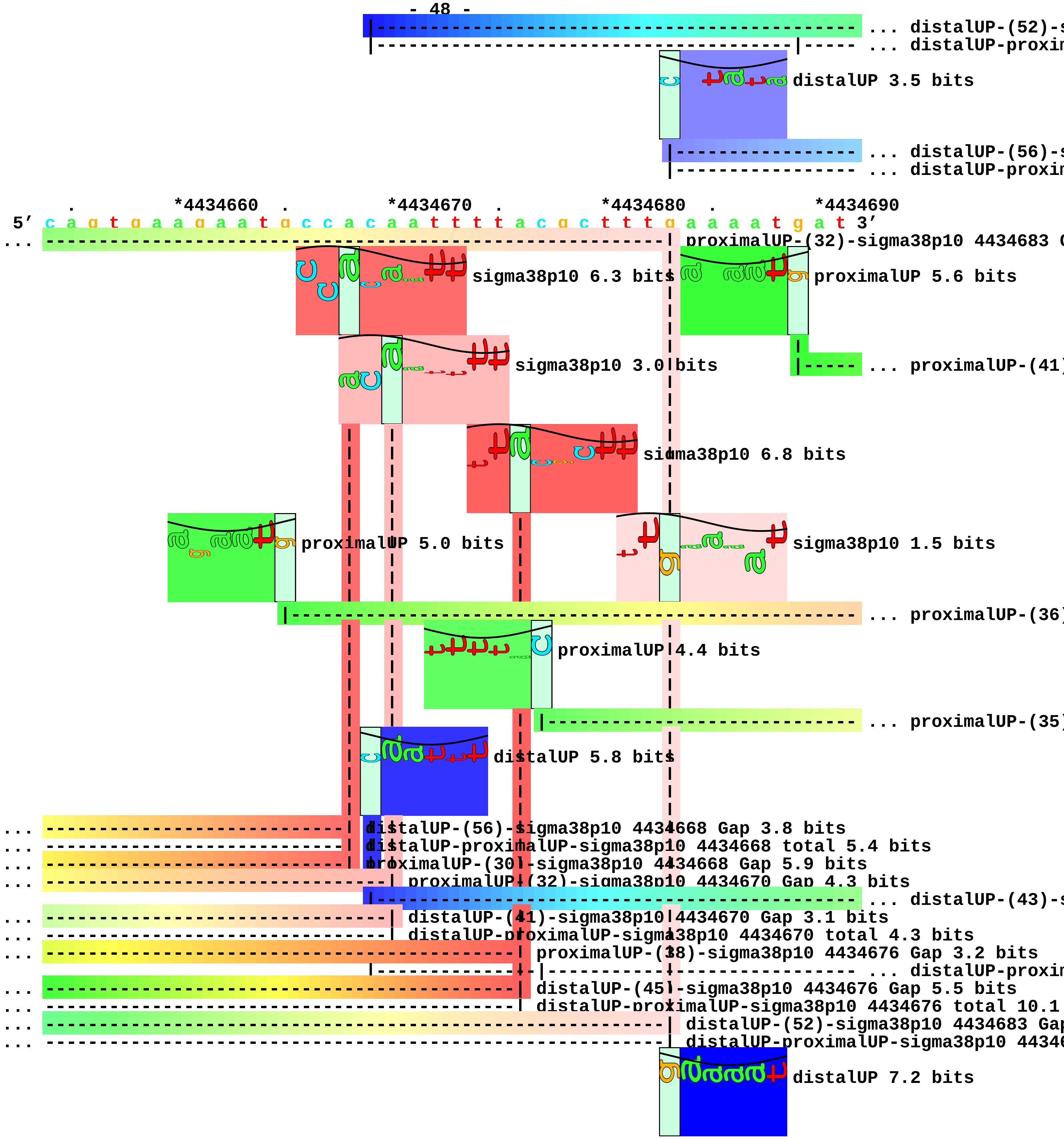
... distalUP-(52)-s
... distalUP-proxim
distalUP 3.5 bits
... distalUP-(56)-s
... distalUP-proxim
. *4434660 . *4434670 . *4434680 . *4434690
5′ c a g t g a a g a a t g c c a c a a t t t t a c g c t t t g a a a a t g a t 3′
... proximalUP-(32)-sigma38p10 4434683 G
sigma38p10 6.3 bits
proximalUP 5.6 bits
sigma38p10 3.0 bits
... proximalUP-(41)
sigma38p10 6.8 bits
proximalUP 5.0 bits
sigma38p10 1.5 bits
... proximalUP-(36)
proximalUP 4.4 bits
... proximalUP-(35)
distalUP 5.8 bits
... distalUP-(56)-sigma38p10 4434668 Gap 3.8 bits
... distalUP-proximalUP-sigma38p10 4434668 total 5.4 bits
... proximalUP-(30)-sigma38p10 4434668 Gap 5.9 bits
... proximalUP-(32)-sigma38p10 4434670 Gap 4.3 bits
... distalUP-(43)-s
... distalUP-(41)-sigma38p10 4434670 Gap 3.1 bits
... distalUP-proximalUP-sigma38p10 4434670 total 4.3 bits
... proximalUP-(38)-sigma38p10 4434676 Gap 3.2 bits
... distalUP-proxim
... distalUP-(45)-sigma38p10 4434676 Gap 5.5 bits
... distalUP-proximalUP-sigma38p10 4434676 total 10.1
... distalUP-(52)-sigma38p10 4434683 Gap
... distalUP-proximalUP-sigma38p10 44346
distalUP 7.2 bits

```
... ----------------------------------------------------------------------------- ... distalUP-(56)-s
... ---------------------|----------------------------------------------------- ... distalUP-proxim
                                                              |---------------- ... distalUP-(47)-s
                                                              |-----------|----- ... distalUP-proxim

              .         *4434700  .         *4434710  .         *4434720  .
5' g a c a c t a t c a c a g t t g g c g c a t t c a t t a a c g a t a g g g t 3'
                                     sigma38p10 2.3 bits                          ... sigma38p10

                                                       sigma38p10 1.0 bits

... ----------------------------------------------------------------------------- ... proximalUP-(41)

... ------------------| proximalUP-(36)-sigma38p10 4434701 Gap 3.6 bits
... --------------------------------------| proximalUP-(35)-sigma38p10 4434712 Gap 3.1 bits

... --------------------------------------| distalUP-(43)-sigma38p10 4434712 Gap 3.6 bits

... ------------------|-------------------| distalUP-proximalUP-sigma38p10 4434712 total 4.6 bits

... ------------------| distalUP-(56)-sigma38p10 4434701 Gap 3.8 bits
... ------------------| distalUP-proximalUP-sigma38p10 4434701 total 3.4 bits
... ----------------------------------------------------------------------------- ... distalUP-(47)-s
... ----------------------------------------------------------------------------- ... distalUP-proxim

 *4434730  .         *4434740  .         *4434750  .         *4434760  .
5' a t a a g t a a a a c a a t a a g t t a a c a c c g c t c a c a g a g a c g 3'
... sigma38p10 1.5 bits

... | proximalUP-(41)-sigma38p10 4434730 Gap 5.5 bits
```

```
    |
    |
    |
    |
    |
    |
    |
    |
    |
    |
    |
    |
    |
... | distalUP-(47)-sigma38p10 4434730 Gap 6.5 bits
... | distalUP-proximalUP-sigma38p10 4434730 total 2.3 bits

        *4434770  .         *4434780  .         *4434790  .
 5' a g g t g g a g a a a t g t t a g a t c a a g t a t g c c a g c 3'
```
