## Supplement_for_sigma38_paper-directory for "*Escherichia coli σ*^38^ promoters use two UP elements instead of a −35 element: resolution of a paradox and discovery that *σ*^38^ transcribes ribosomal promoters": Peano.Landini2015-Table34.pdf

piece 1, NC_000913, p10-dependent-ansP-P1-1524035-gttcatggcaacttatatga, config: linear, direction: -, begin: 1524154, end: 1523994

*1524150 . *1524140 . *1524130 . *1524120

5' t t t t g c a t c g t t t a t c g a t a a c g g g t a g g a t g c t g c g c 3'

T timestamp:_2018Sep20_11-34-26

p35 1.2 bits

proximalUP 6.8 bits

p35 1.4 bits

... proximalUP-(35)-sigma38p10 1524104 Gap

distalUP 5.5 bits

... distalUP-(41)-sigma38p10 1524104 Gap

... sigma38p10-proximalUP-distalUP 1524104 total 7.4

p35 6.2 bits

p35 3.3 bits

p35 0.9 bits

. *1524110 . *1524100 . *1524090 . *1524080

5' c g a t a g t t t t a t a c c c a t a a a g a a t a a t g g t g a t a a c t 3'

sigma38p10 1.3 bits

... proximalUP-(32)-sigma38p10 1524056 Gap

proximalUP 4.4 bits

... proximalUP-(35)-sigma38p10 1524053 Gap

... proximalUP-(35)-sigma38p10 1524104 Gap 3.1 bits ... p35

... proximalUP-(34)-sigma38p10 1524069 Gap

... distalUP-(41)-sigma38p10 1524104 Gap 3.1 bits

... sigma38p10-proximalUP-distalUP 1524104 total 7.4 bits

distalUP 4.2 bits

... proximalUP-(35)-sigma38p10 1524060 Gap

... distalUP-(42)-sigma38p10 1524069 Gap

... sigma38p10-proximalUP-distalUP 1524069 total 2.9

p35 0.7 bits

proximalUP 4.9 bits

distalUP 5.6 bits

... distalUP-(41)-sigma38p10 1524060 Gap

... sigma38p10-proximalUP-distalUP 1524060 total 9.2

p35 4.8 bits

proximalUP 6.7 bits

... distalUP-(45)-sigma38p10 1524056 Gap

... sigma38p10-proximalUP-distalUP 1524056 total 4.4

p35 4.7 bits

distalUP 6.5 bits

p35 0.4 bits

... distalUP-(42)-sigma38p10 1524053 Gap

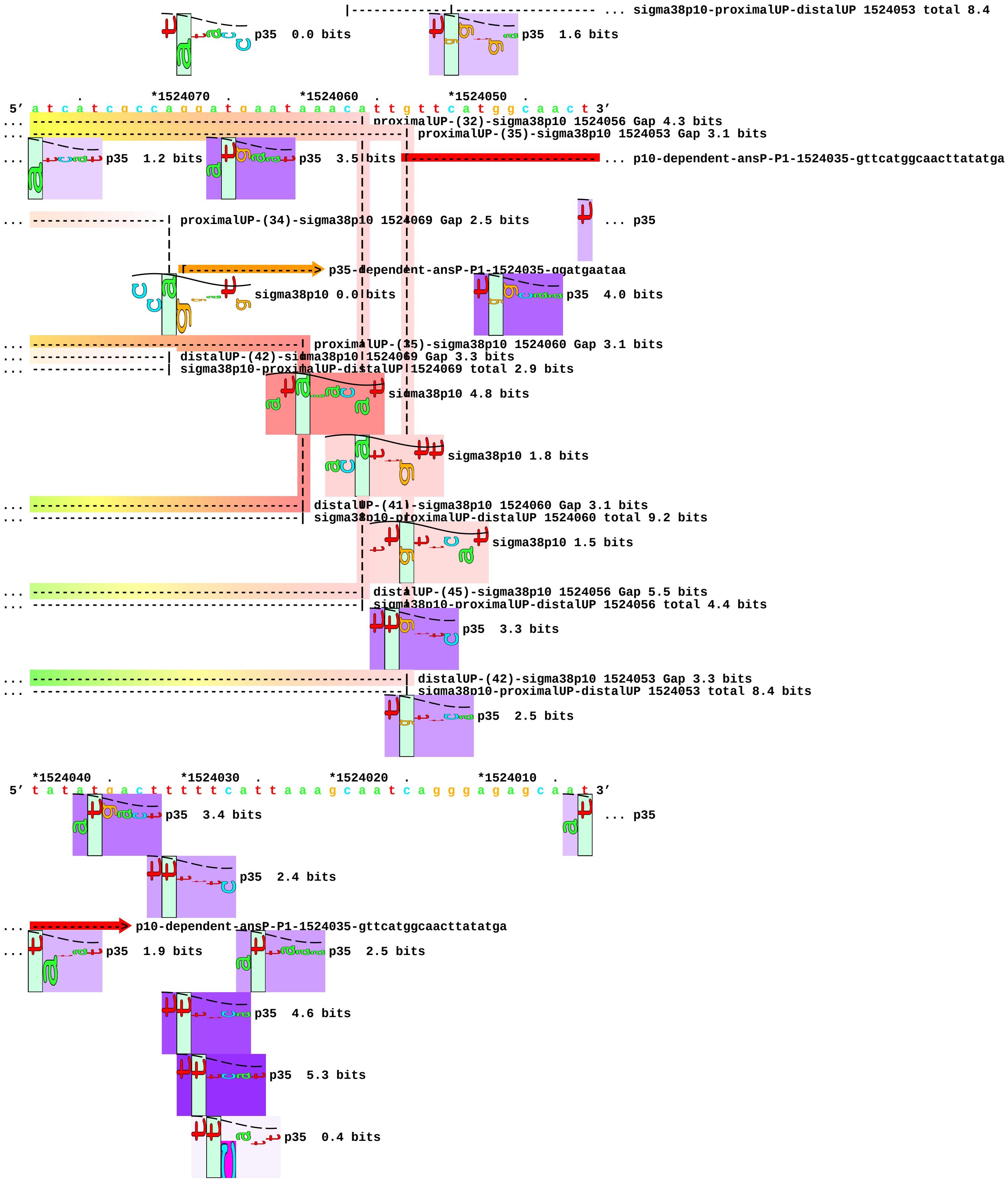
... sigma38p10-proximalUP-distalUP 1524053 total 8.4
p35 0.0 bits
p35 1.6 bits
*1524070 . *1524060 . *1524050 .
5' a t c a t c g c c a g g a t g a a t a a a c a t t g t t c a t g g c a a c t 3'
... proximalUP-(32)-sigma38p10 1524056 Gap 4.3 bits
... proximalUP-(35)-sigma38p10 1524053 Gap 3.1 bits
... p35 1.2 bits
p35 3.5 bits
... p10-dependent-ansP-P1-1524035-gttcatggcaacttatatga
... proximalUP-(34)-sigma38p10 1524069 Gap 2.5 bits
... p35
p35-dependent-ansP-P1-1524035-ggatgaataa
sigma38p10 0.0 bits
p35 4.0 bits
... proximalUP-(35)-sigma38p10 1524060 Gap 3.1 bits
... distalUP-(42)-sigma38p10 1524069 Gap 3.3 bits
... sigma38p10-proximalUP-distalUP 1524069 total 2.9 bits
sigma38p10 4.8 bits
sigma38p10 1.8 bits
... distalUP-(41)-sigma38p10 1524060 Gap 3.1 bits
... sigma38p10-proximalUP-distalUP 1524060 total 9.2 bits
sigma38p10 1.5 bits
... distalUP-(45)-sigma38p10 1524056 Gap 5.5 bits
... sigma38p10-proximalUP-distalUP 1524056 total 4.4 bits
p35 3.3 bits
... distalUP-(42)-sigma38p10 1524053 Gap 3.3 bits
... sigma38p10-proximalUP-distalUP 1524053 total 8.4 bits
p35 2.5 bits
*1524040 . *1524030 . *1524020 . *1524010 .
5' t a t a t g a c t t t t t c a t t a a a g c a a t c a g g g a g a g c a a t 3'
p35 3.4 bits
... p35
p35 2.4 bits
... p10-dependent-ansP-P1-1524035-gttcatggcaacttatatga
... p35 1.9 bits
p35 2.5 bits
p35 4.6 bits
p35 5.3 bits
p35 0.4 bits

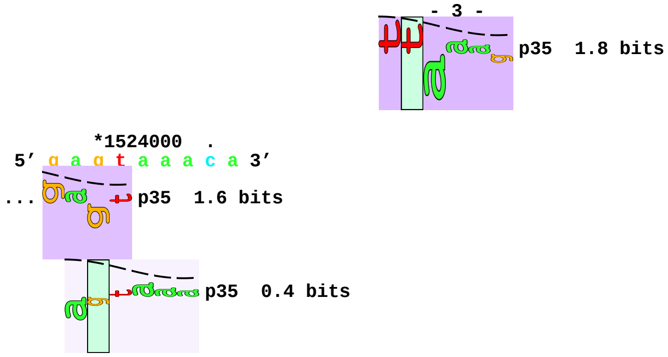
- 3 -
p35 1.8 bits
*1524000 .
5′ g a g t a a a c a 3′
... p35 1.6 bits
p35 0.4 bits

piece 2, NC_000913, p10-dependent-cpdB-4434652-aattgtggcattcttcactg, config: linear, direction: -, begin: 4434774, end: 4434614

*4434770 . *4434760 . *4434750 . *4434740

5' t c c a c c t c g t c t c t g t g a g c g g t g t t a a c t t a t t g t t t 3'

T timestamp:_2018Sep20_11-34-26

p35 2.6 bits ... p35

p35 2.8 bits ... p35

p35 0.0 bits

p35 4.8 bits

... p35

... p35

. *4434730 . *4434720 . *4434710 . *4434700

5' t a c t t t a t a c c c t a t c g t t a a t g a a t g c g c c a a c t g t g a 3'

... p35 3.2 bits proximalUP 6.3 bits

... p35 1.6 bits ... proximalUP-(27)-sigma38p10 4434688 Gap

proximalUP 4.4 bits ... proximalUP-(40)-sigma38p10 4434675 Gap

... proximalUP-(37)-sigma38p10 4434691 Gap

... p35 1.8 bits distalUP 5.5 bits

... p35 5.7 bits ... distalUP-(46)-sigma38p10 4434675 Gap

distalUP 3.5 bits p35 3.5 bits

... distalUP-(43)-sigma38p10 4434691 Gap
... sigma38p10-proximalUP-distalUP 4434691 total 2.7
... distalUP-(46)-sigma38p10 4434688 Gap
... sigma38p10-proximalUP-distalUP 4434688 total 3.1
... sigma38p10-proximalUP-distalUP 4434675 total 10.7

p35 0.4 bits p35 0.9 bits

p35 0.0 bits

. *4434690 . *4434680 . *4434670 .

5' t a g t g t c a t c a t t t t c a a a g c g t a a a a t t g t g g c a t t c 3'

sigma38p10 2.9 bits ... p10-dependent-cpdB-4434652-aattgtggcattcttcactg

... proximalUP-(27)-sigma38p10 4434688 Gap 3.8 bits
... proximalUP-(40)-sigma38p10 4434675 Gap 4.0 bits
... proximalUP-(37)-sigma38p10 4434691 Gap 4.5 bits ... p35

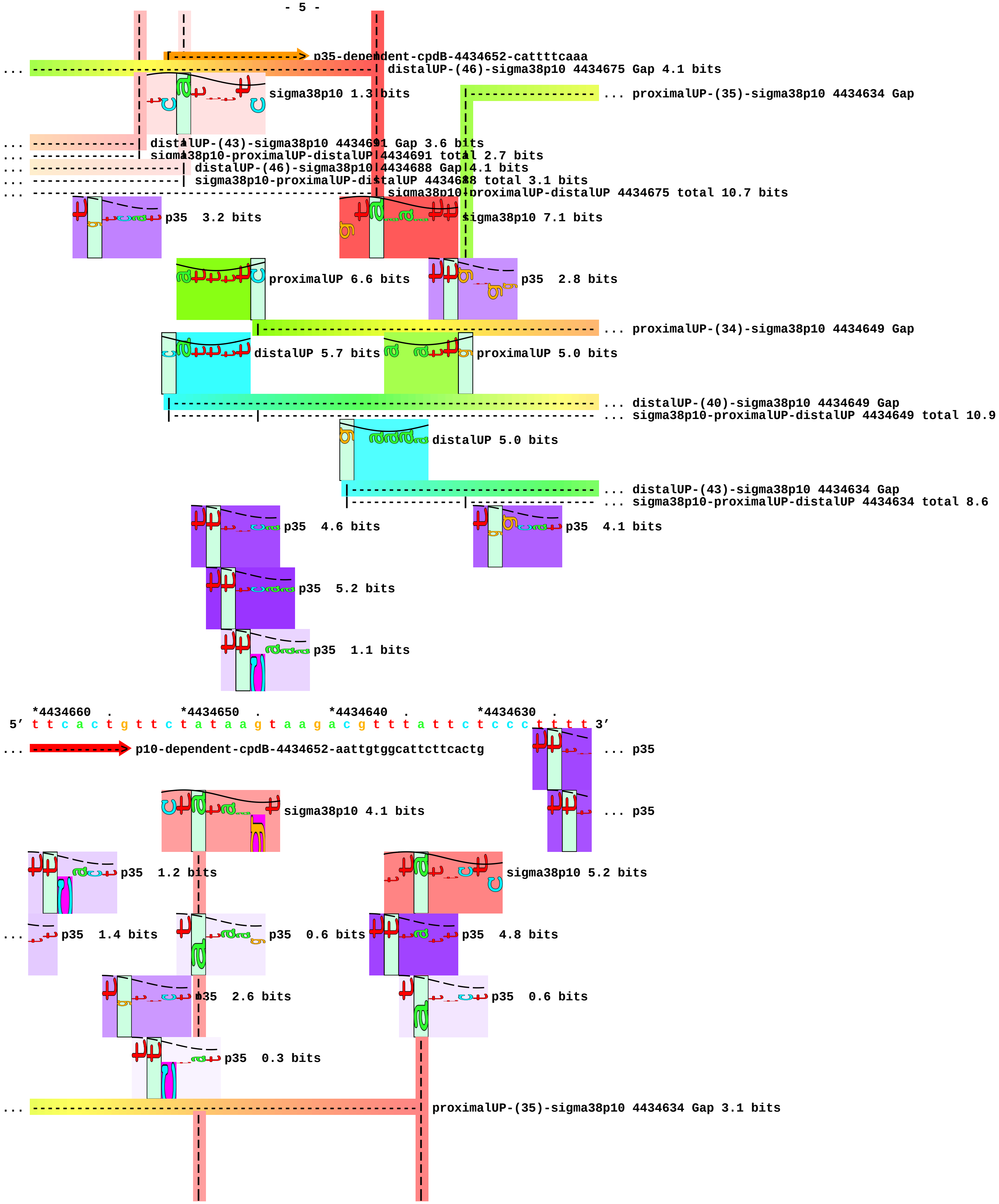
p35-dependent-cpdB-4434652-cattttcaaa
... distalUP-(46)-sigma38p10 4434675 Gap 4.1 bits
sigma38p10 1.3 bits
... proximalUP-(35)-sigma38p10 4434634 Gap
... distalUP-(43)-sigma38p10 4434691 Gap 3.6 bits
... sigma38p10-proximalUP-distalUP 4434691 total 2.7 bits
... distalUP-(46)-sigma38p10 4434688 Gap 4.1 bits
... sigma38p10-proximalUP-distalUP 4434688 total 3.1 bits
... sigma38p10-proximalUP-distalUP 4434675 total 10.7 bits
p35 3.2 bits
sigma38p10 7.1 bits
proximalUP 6.6 bits
p35 2.8 bits
... proximalUP-(34)-sigma38p10 4434649 Gap
distalUP 5.7 bits
proximalUP 5.0 bits
... distalUP-(40)-sigma38p10 4434649 Gap
... sigma38p10-proximalUP-distalUP 4434649 total 10.9
distalUP 5.0 bits
... distalUP-(43)-sigma38p10 4434634 Gap
... sigma38p10-proximalUP-distalUP 4434634 total 8.6
p35 4.6 bits
p35 4.1 bits
p35 5.2 bits
p35 1.1 bits
*4434660 . *4434650 . *4434640 . *4434630 .
5′ t t c a c t g t t c t a t a a g t a a g a c g t t t a t t c t c c c t t t t 3′
... p10-dependent-cpdB-4434652-aattgtggcattcttcactg
... p35
sigma38p10 4.1 bits
... p35
p35 1.2 bits
sigma38p10 5.2 bits
... p35 1.4 bits
p35 0.6 bits
p35 4.8 bits
p35 2.6 bits
p35 0.6 bits
p35 0.3 bits
... proximalUP-(35)-sigma38p10 4434634 Gap 3.1 bits

```
... ----------------------| proximalUP-(34)-sigma38p10 4434649 Gap 2.5 bits
                          |                             |
... ----------------------| distalUP-(40)-sigma38p10 4434649 Gap 3.1 bits
... ----------------------| sigma38p10-proximalUP-distalUP 4434649 total 10.9 bits
                                                        |
... ---------------------------------------------------| distalUP-(43)-sigma38p10 4434634 Gap 3.6 bits
... ---------------------------------------------------| sigma38p10-proximalUP-distalUP 4434634 total 8.6 bits

          *4434620  .
 5' c t t t c g t a t 3'
...           p35  4.8 bits

...             p35  4.5 bits

                        p35  3.5 bits
```

piece 3, NC_000913, p10-dependent-csiE-2663423-tccttgagcaaactttagct, config: linear, direction: +, begin: 2663283, end: 2663443

. *2663290 . *2663300 . *2663310 . *2663320
5’ c g a t a a t c a t t c a c a a a a c c a c c t t a a g a c a t g c t a a t 3’

T timestamp:_2018Sep20_11-34-26

proximalUP 2.0 bits

distalUP 7.1 bits

... proximalUP-(30)-sigma38p10 2663340 Gap

... distalUP-(56)-sigma38p10 2663340 Gap
... distalUP-proximalUP-sigma38p10 2663340 total 3.8

p35 0.7 bits
p35 0.8 bits
p35 1.1 bits
p35 2.1 bits

. *2663330 . *2663340 . *2663350 .
5’ c c a c t g g t c a g a a c a g t t t a a g a t g a g a a a a a t t c t g t 3’

p35 3.4 bits
sigma38p10 4.3 bits

... proximalUP-(30)-sigma38p10 2663340 Gap 5.9 bits
... distalUP-(56)-sigma38p10 2663340 Gap 3.8 bits
... distalUP-proximalUP-sigma38p10 2663340 total 3.8 bits

proximalUP 4.5 bits

... proximalUP-(37)-sigma38p10 2663379 Gap

distalUP 4.8 bits
p35 0.4 bits

... distalUP-(43)-sigma38p10 2663379 Gap
... distalUP-proximalUP-sigma38p10 2663379 total 2.6

p35 4.7 bits
p35 0.8 bits
p35 1.5 bits

*2663360 . *2663370 . *2663380 . *2663390 .
5’ g a c g c t t g c c a a c a t t t c t g a t g a t t a g c a t t c c c t t c 3’

proximalUP 2.6 bits [-----------t----> p35-dependent-csiE-2663423-cattcctttc.6

p35 6.1 bits
sigma38p10 1.3 bits

... proximalUP-(36)-sigma38p10 2663412 Gap

distalUP 1.6 bits
... proximalUP-(36)-sigma38p10 2663423 Gap

... distalUP-(44)-sigma38p10 2663412 Gap
... proximalUP-(37)-sigma38p10 2663379 Gap 4.5 bits
... distalUP-proximalUP-sigma38p10 2663412 total 2.6
... distalUP-(43)-sigma38p10 2663379 Gap 3.6 bits
... distalUP-proximalUP-sigma38p10 2663379 total 2.6 bits

proximalUP 0.2 bits

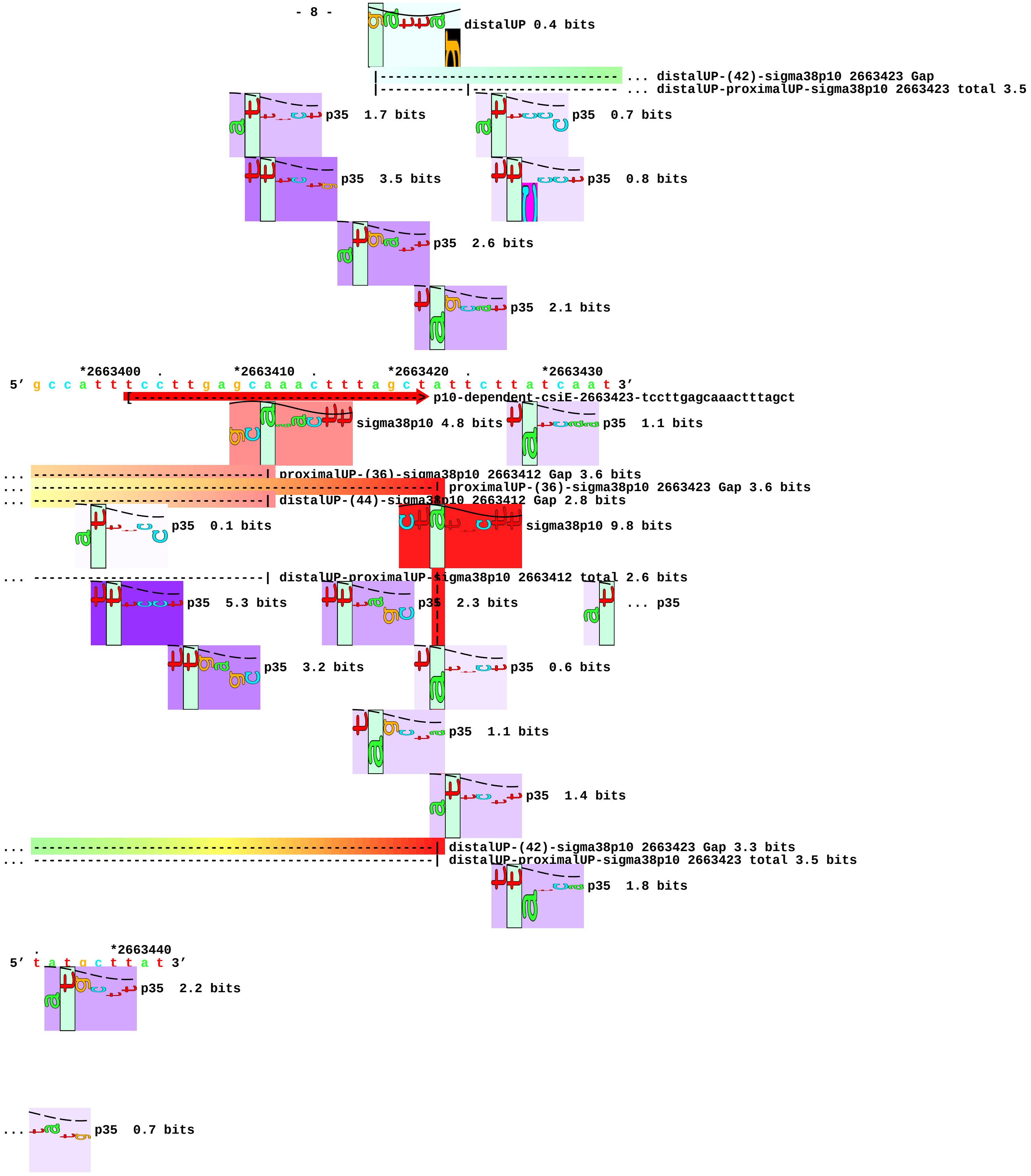
distalUP 0.4 bits
... distalUP-(42)-sigma38p10 2663423 Gap
... distalUP-proximalUP-sigma38p10 2663423 total 3.5
p35 1.7 bits
p35 0.7 bits
p35 3.5 bits
p35 0.8 bits
p35 2.6 bits
p35 2.1 bits
*2663400 . *2663410 . *2663420 . *2663430
5' g c c a t t t c c t t g a g c a a a c t t t a g c t a t t c t t a t c a a t 3'
p10-dependent-csiE-2663423-tccttgagcaaactttagct
sigma38p10 4.8 bits
p35 1.1 bits
... proximalUP-(36)-sigma38p10 2663412 Gap 3.6 bits
... proximalUP-(36)-sigma38p10 2663423 Gap 3.6 bits
... distalUP-(44)-sigma38p10 2663412 Gap 2.8 bits
p35 0.1 bits
sigma38p10 9.8 bits
... distalUP-proximalUP-sigma38p10 2663412 total 2.6 bits
p35 5.3 bits
p35 2.3 bits
... p35
p35 3.2 bits
p35 0.6 bits
p35 1.1 bits
p35 1.4 bits
... distalUP-(42)-sigma38p10 2663423 Gap 3.3 bits
... distalUP-proximalUP-sigma38p10 2663423 total 3.5 bits
p35 1.8 bits
. *2663440
5' t a t g c t t a t 3'
p35 2.2 bits
... p35 0.7 bits

piece 4, NC_000913, p10-dependent-csrA-2817295-atatcggctaaacttaggtt, config: linear, direction: -, begin: 2817417, end: 2817257

. *2817410 . *2817400 . *2817390 . *2817380
5' g c g a a a t t g c a a t a a t a t a a g c g t c a g g c a a t g c c g t g 3'

T timestamp:_2018Sep20_11-34-26

p35 0.9 bits

p35 6.1 bits

p35 2.1 bits

p35 0.6 bits

... p35

. *2817370 . *2817360 . *2817350 .
5' g a c t c g c t t c a c g g c a t t c g c a t t a a c g c t a t c g a c a a 3'

p35 0.2 bits

proximalUP 5.4 bits

... proximalUP-(35)-sigma38p10 2817318 Gap

... p35 4.4 bits

distalUP 5.6 bits

... distalUP-(41)-sigma38p10 2817318 Gap
... sigma38p10-proximalUP-distalUP 2817318 total 5.4
... distalUP-(52)-sigma38p10 2817307 Gap
... sigma38p10-proximalUP-distalUP 2817307 total 11.6

p35 1.0 bits

p35 1.8 bits

*2817340 . *2817330 . *2817320 . *2817310 .
5' c g a t a a a g t c a g g t t g a a g t t g t g t a t a t c g g c t a a a c 3'

proximalUP 4.9 bits

... p10-dependent-csrA-2817295-atatcggctaaacttaggtt

... proximalUP-(35)-sigma38p10 2817318 Gap 3.1 bits
p35-dependent-csrA-2817295-gtcaggttga
... distalUP-(41)-sigma38p10 2817318 Gap 3.1 bits
... sigma38p10-proximalUP-distalUP 2817318 total 5.4 bits
... distalUP-(52)-sigma38p10 2817307 Gap 5.5 bits
... sigma38p10-proximalUP-distalUP 2817307 total 11.6 bits

sigma38p10 0.6 bits

p35 5.5 bits

... sigma38p10

proximalUP-(27)-sigma38p10 2817307 Gap 3.8 bits

distalUP 5.6 bits

... distalUP-(53)-sigma38p10 2817265 Gap
... sigma38p10-proximalUP-distalUP 2817265 total 2.4

p35 3.8 bits

p35 2.6 bits

*2817300 . *2817290 . *2817280 . *2817270
5' t t a g g t t t a a c a g a a t g t a a t g c c a t g a c t g c t t a g a t 3'

... p10-dependent-csrA-2817295-atatcggctaaacttaggtt

... sigma38p10

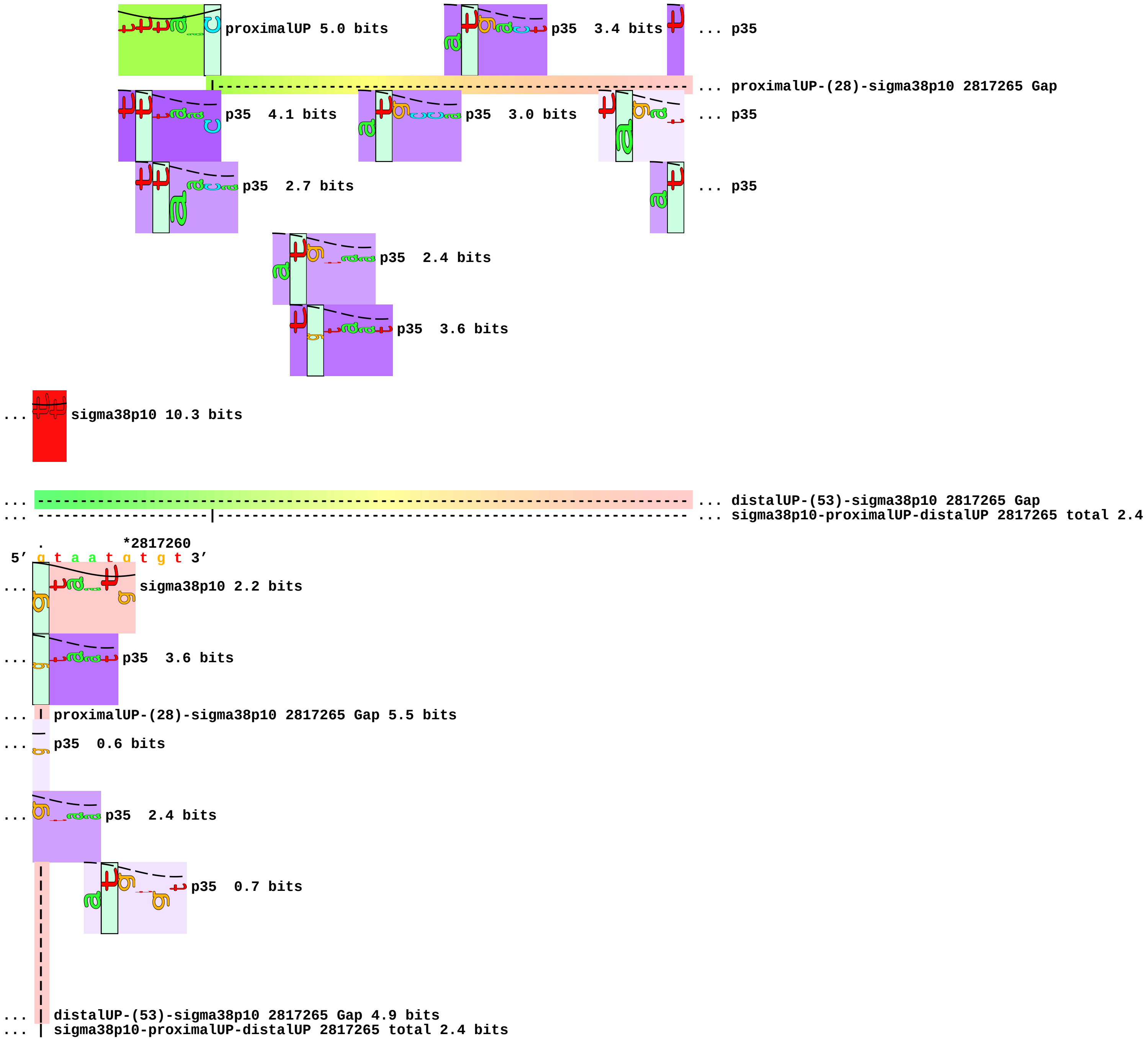
proximalUP 5.0 bits
p35 3.4 bits
... p35
... proximalUP-(28)-sigma38p10 2817265 Gap
p35 4.1 bits
p35 3.0 bits
... p35
p35 2.7 bits
... p35
p35 2.4 bits
p35 3.6 bits
... sigma38p10 10.3 bits
... distalUP-(53)-sigma38p10 2817265 Gap
... sigma38p10-proximalUP-distalUP 2817265 total 2.4
*2817260
5' g t a a t g t g t 3'
... sigma38p10 2.2 bits
... p35 3.6 bits
... proximalUP-(28)-sigma38p10 2817265 Gap 5.5 bits
... p35 0.6 bits
... p35 2.4 bits
p35 0.7 bits
... distalUP-(53)-sigma38p10 2817265 Gap 4.9 bits
... sigma38p10-proximalUP-distalUP 2817265 total 2.4 bits

piece 5, NC_000913, p10-dependent-dps-848173-gccggtgctatacttaatct, config: linear, direction: -, begin: 848295, end: 848135

```
   .          *848290    .          *848280    .          *848270    .          *848260
5' c t t a t c a a t a t a t c t a a c t c a t t g a a t c t t t a t t a g t t 3'
```

T timestamp:_2018Sep20_11-34-26

proximalUP 5.1 bits

proximalUP 5.7 bits

... proximalUP-(32)-sigma38p10 848236 Gap

... proximalUP-(27)-sigma38p10 848255 Gap

distalUP 0.3 bits

... proximalUP-(35)-sigma38p10 848233 Gap

... distalUP-(40)-sigma38p10 848255 Gap
... sigma38p10-proximalUP-distalUP 848255 total 2.7
... proximalUP-(34)-sigma38p10 848248 Gap

distalUP 6.4 bits

p35 6.5 bits

... p35

... distalUP-(42)-sigma38p10 848248 Gap
... sigma38p10-proximalUP-distalUP 848248 total 10.2
... distalUP-(54)-sigma38p10 848236 Gap
... sigma38p10-proximalUP-distalUP 848236 total 7.5

p35 1.8 bits

distalUP 0.7 bits

p35 1.1 bits

... distalUP-(43)-sigma38p10 848233 Gap

... sigma38p10-proximalUP-distalUP 848233 total 2.1

p35 0.2 bits

p35 4.8 bits

p35 0.9 bits

p35 0.7 bits

... p35

```
   .          *848250    .          *848240    .          *848230    .          *848220
5' t t g t t t t t c a c g c t t g t t a c c a c t a t t a g t g t g a t a g g 3'
```

sigma38p10 3.6 bits

sigma38p10 5.2 bits

... proximalUP-(32)-sigma38p10 848236 Gap 4.3 bits

... proximalUP-(27)-sigma38p10 848255 Gap 3.8 bits

proximalUP 2.3 bits

... proximalUP-(35)-sigma38p10 848233 Gap 3.1 bits

... distalUP-(40)-sigma38p10 848255 Gap 3.1 bits

... proximalUP-(36)-sigma38p10 848185 Gap

... sigma38p10-proximalUP-distalUP 848255 total 2.7 bits

... proximalUP-(34)-sigma38p10 848248 Gap 2.5 bits

... p35 2.9 bits

sigma38p10 3.0 bits

... distalUP-(42)-sigma38p10 848248 Gap 3.3 bits
... sigma38p10-proximalUP-distalUP 848248 total 10.2 bits
... distalUP-(54)-sigma38p10 848236 Gap 4.9 bits
... sigma38p10-proximalUP-distalUP 848236 total 7.5 bits

sigma38p10 3.9 bits

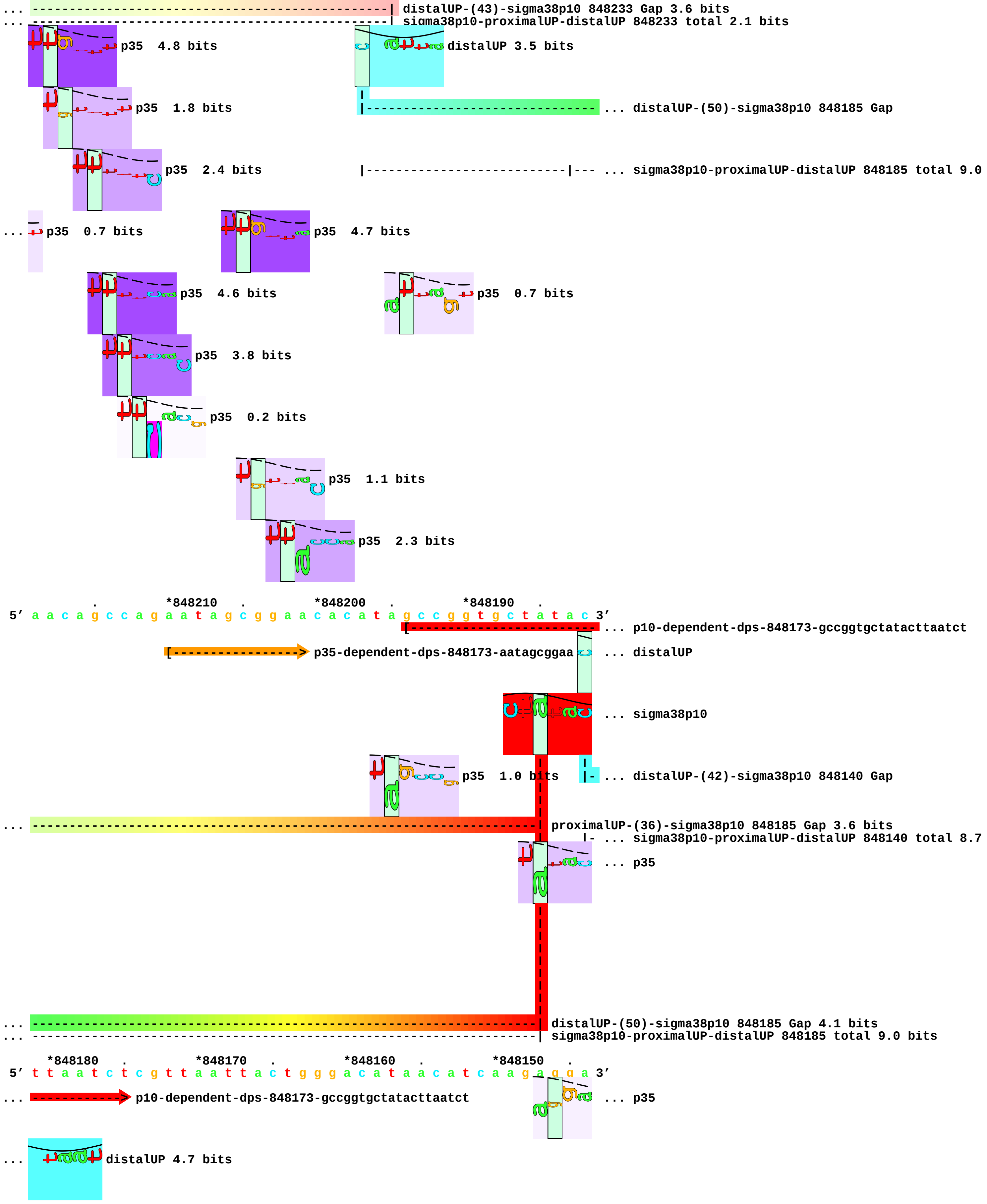
... distalUP-(43)-sigma38p10 848233 Gap 3.6 bits
... sigma38p10-proximalUP-distalUP 848233 total 2.1 bits
p35 4.8 bits
distalUP 3.5 bits
p35 1.8 bits
... distalUP-(50)-sigma38p10 848185 Gap
p35 2.4 bits
... sigma38p10-proximalUP-distalUP 848185 total 9.0
... p35 0.7 bits
p35 4.7 bits
p35 4.6 bits
p35 0.7 bits
p35 3.8 bits
p35 0.2 bits
p35 1.1 bits
p35 2.3 bits
. *848210 . *848200 . *848190 .
5' a a c a g c c a g a a t a g c g g a a c a c a t a g c c g g t g c t a t a c 3'
... p10-dependent-dps-848173-gccggtgctatacttaatct
p35-dependent-dps-848173-aatagcggaa
... distalUP
... sigma38p10
p35 1.0 bits
... distalUP-(42)-sigma38p10 848140 Gap
... proximalUP-(36)-sigma38p10 848185 Gap 3.6 bits
... sigma38p10-proximalUP-distalUP 848140 total 8.7
... p35
... distalUP-(50)-sigma38p10 848185 Gap 4.1 bits
... sigma38p10-proximalUP-distalUP 848185 total 9.0 bits
*848180 . *848170 . *848160 . *848150 .
5' t t a a t c t c g t t a a t t a c t g g g a c a t a a c a t c a a g a g g a 3'
... p10-dependent-dps-848173-gccggtgctatacttaatct
... p35
... distalUP 4.7 bits

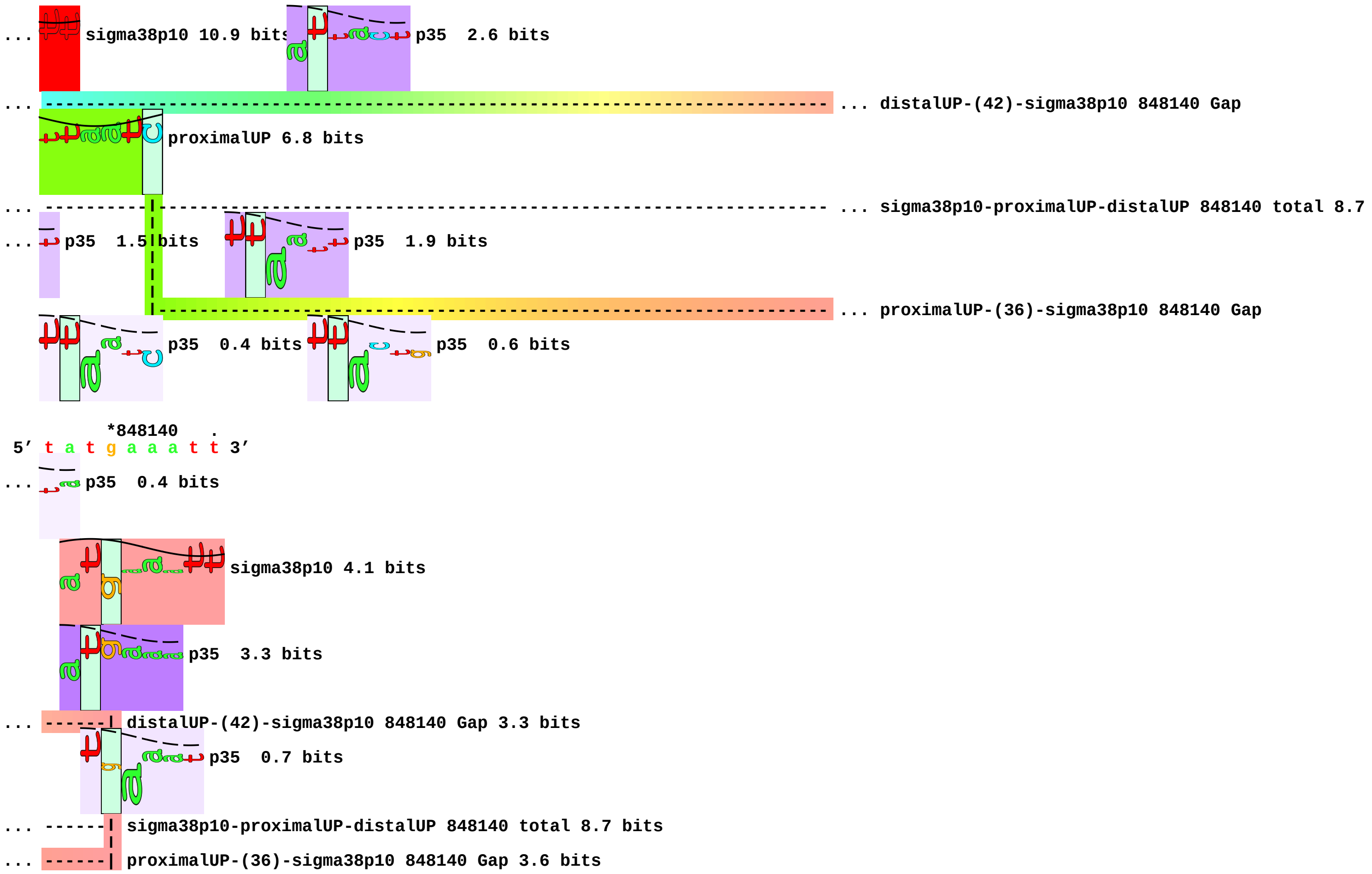
... sigma38p10 10.9 bits
p35 2.6 bits
... distalUP-(42)-sigma38p10 848140 Gap
proximalUP 6.8 bits
... sigma38p10-proximalUP-distalUP 848140 total 8.7
... p35 1.5 bits
p35 1.9 bits
... proximalUP-(36)-sigma38p10 848140 Gap
p35 0.4 bits
p35 0.6 bits
*848140 .
5' t a t g a a a t t 3'
... p35 0.4 bits
sigma38p10 4.1 bits
p35 3.3 bits
... distalUP-(42)-sigma38p10 848140 Gap 3.3 bits
p35 0.7 bits
... sigma38p10-proximalUP-distalUP 848140 total 8.7 bits
... proximalUP-(36)-sigma38p10 848140 Gap 3.6 bits

piece 6, NC_000913, p10-dependent-omrA-2974211-gtgcagaccacaatcaagat, config: linear, direction: -, begin: 2974332, end: 2974172

```
    *2974330  .         *2974320  .         *2974310  .         *2974300  .
5' t t t t t t t g c c g g t c a t c a a t c t g t a a c a g t a a c c g a c a 3'
```

T timestamp:_2018Sep20_11-34-26

p35 2.0 bits

p35 3.9 bits

p35 3.9 bits

p35 2.9 bits

p35 1.4 bits

p35 5.2 bits

```
      *2974290  .         *2974280  .         *2974270  .         *2974260
5' a t t t a c a c a c c t c g t t g c a t t t c c c t t c a t t c c t t g c 3'
```

p35 0.1 bits

p35 6.2 bits

p35 2.2 bits

p35 5.5 bits

p35 0.1 bits

... p35

p35 0.9 bits

p35 3.8 bits

p35 0.8 bits

p35 0.4 bits

```
   .        *2974250  .         *2974240  .         *2974230  .         *2974220
5' g t t t t c t c g c t g g c g a a g a g t c g t c g t g c a g a c c a c a a 3'
```

p35 4.8 bits

[-------------------------- ... p10-dependent-omrA-2974211-gtgcagaccacaatcaagat

... p35 4.4 bits

p35 2.1 bits

[-------------------> p35-dependent-omrA-2974211-cgctggcgaa

p35 2.9 bits

```
     .        *2974210  .         *2974200  .         *2974190  .
5' t c a a g a t c c c a g a g g t a t t g a t t g g t g a g a t t a t t c g g 3'
```

... -------------> p10-dependent-omrA-2974211-gtgcagaccacaatcaagat

p35 0.4 bits

p35 1.7 bits

p35 5.7 bits

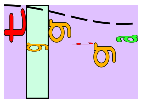

p35 1.6 bits

*2974180 .
5′ t a c g c t c t t 3′

piece 7, NC_000913, p10-dependent-osmB-1341393-ttcttagctattatagttat, config: linear, direction: -, begin: 1341516, end: 1341356

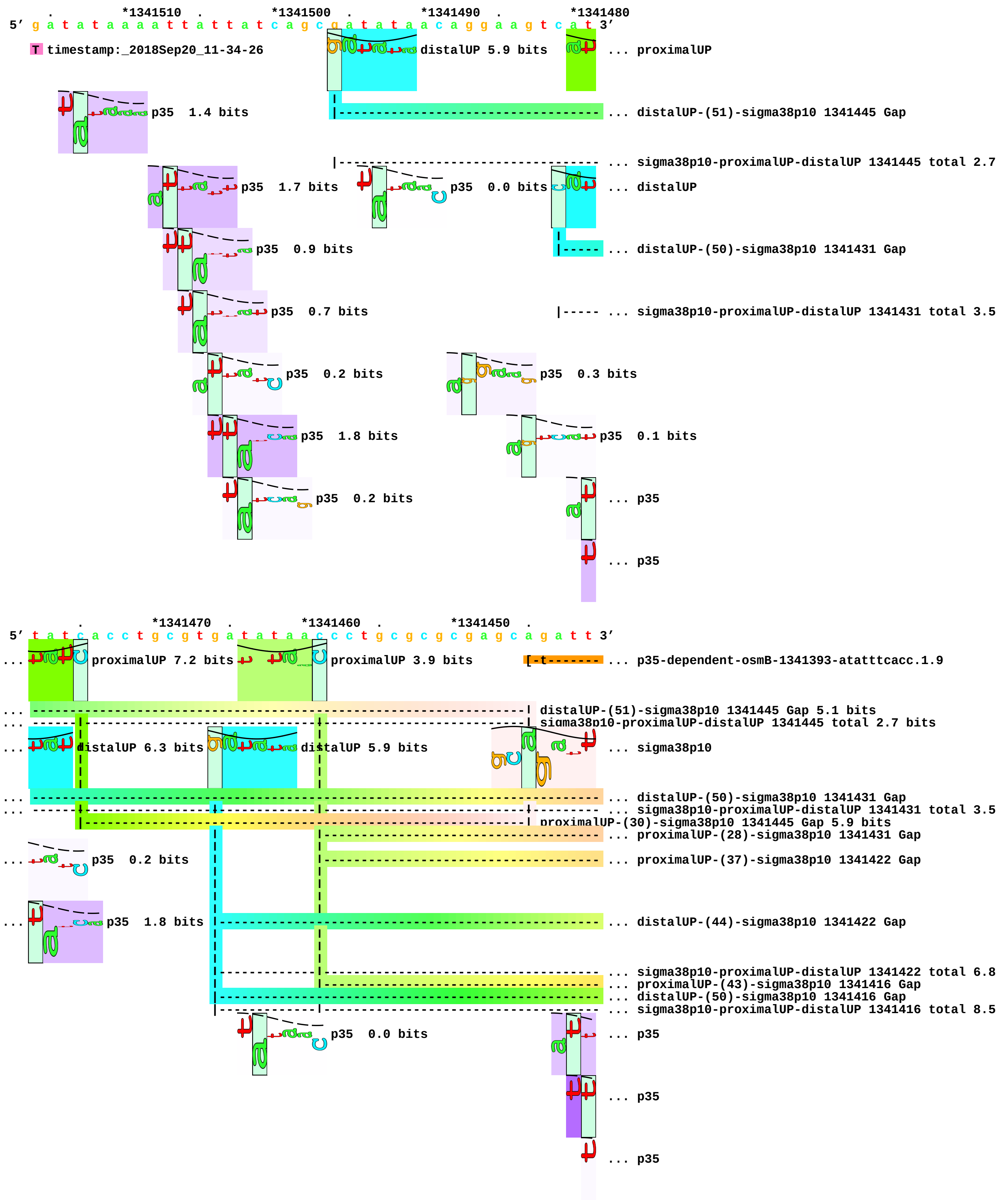

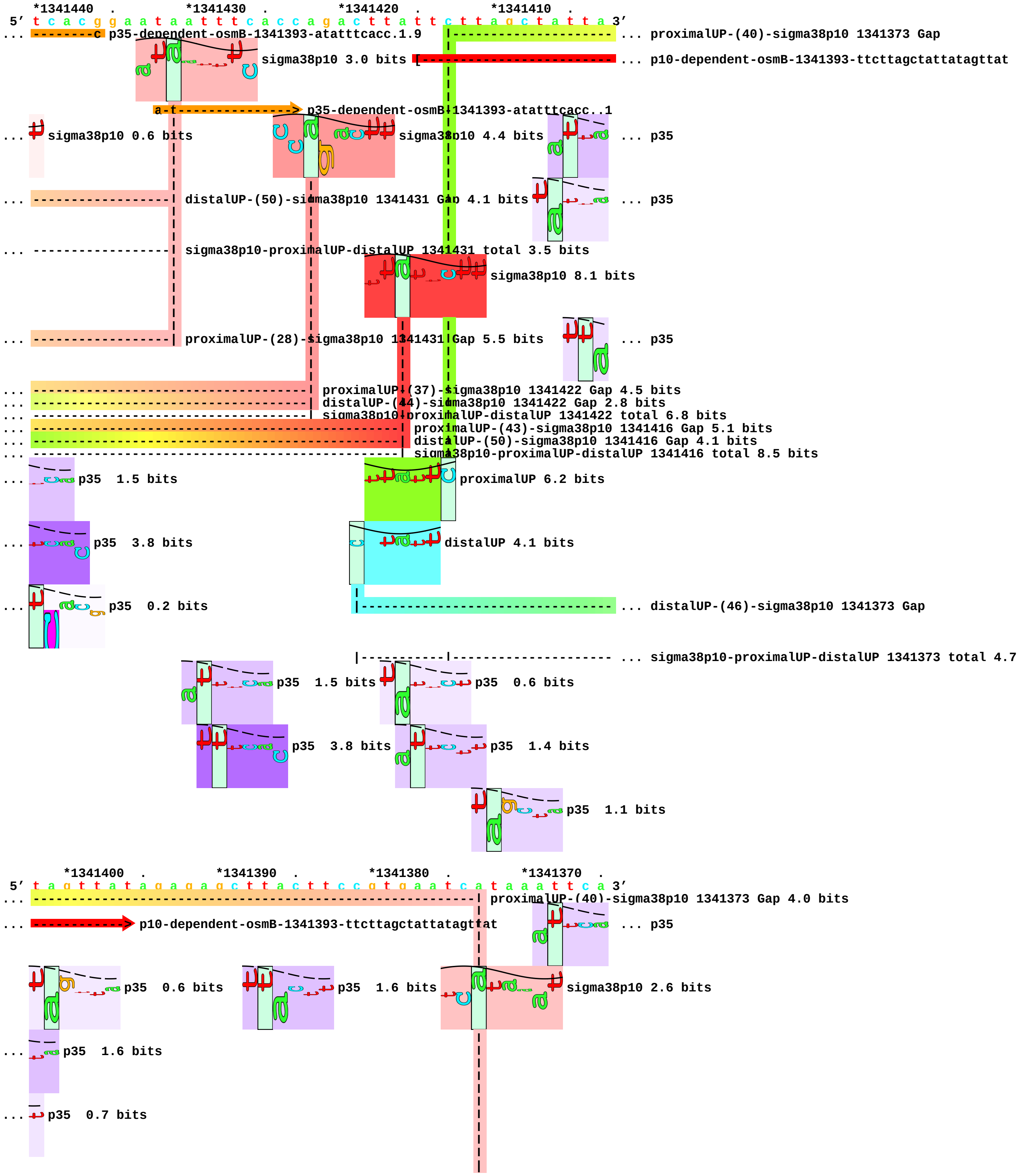
*1341440 . *1341430 . *1341420 . *1341410 .
5' t c a c g g a a t a a t t t c a c c a g a c t t a t t c t t a g c t a t t a 3'
p35-dependent-osmB-1341393-atatttcacc.1.9
... proximalUP-(40)-sigma38p10 1341373 Gap
sigma38p10 3.0 bits
... p10-dependent-osmB-1341393-ttcttagctattatagttat
p35-dependent-osmB-1341393-atatttcacc..1
... sigma38p10 0.6 bits
sigma38p10 4.4 bits
... p35
... distalUP-(50)-sigma38p10 1341431 Gap 4.1 bits
... p35
... sigma38p10-proximalUP-distalUP 1341431 total 3.5 bits
sigma38p10 8.1 bits
... proximalUP-(28)-sigma38p10 1341431 Gap 5.5 bits
... p35
... proximalUP-(37)-sigma38p10 1341422 Gap 4.5 bits
... distalUP-(44)-sigma38p10 1341422 Gap 2.8 bits
... sigma38p10-proximalUP-distalUP 1341422 total 6.8 bits
... proximalUP-(43)-sigma38p10 1341416 Gap 5.1 bits
... distalUP-(50)-sigma38p10 1341416 Gap 4.1 bits
... sigma38p10-proximalUP-distalUP 1341416 total 8.5 bits
... p35 1.5 bits
proximalUP 6.2 bits
... p35 3.8 bits
distalUP 4.1 bits
... p35 0.2 bits
... distalUP-(46)-sigma38p10 1341373 Gap
... sigma38p10-proximalUP-distalUP 1341373 total 4.7
p35 1.5 bits
p35 0.6 bits
p35 3.8 bits
p35 1.4 bits
p35 1.1 bits
*1341400 . *1341390 . *1341380 . *1341370 .
5' t a g t t a t a g a g a g c t t a c t t c c g t g a a t c a t a a a t t c a 3'
... proximalUP-(40)-sigma38p10 1341373 Gap 4.0 bits
... p10-dependent-osmB-1341393-ttcttagctattatagttat
... p35
p35 0.6 bits
p35 1.6 bits
sigma38p10 2.6 bits
... p35 1.6 bits
... p35 0.7 bits

p35 0.9 bits

p35 0.5 bits

... p35 0.9 bits

... ------------------------------------------------------------| distalUP-(46)-sigma38p10 1341373 Gap 4.1 bits
... ------------------------------------------------------------| sigma38p10-proximalUP-distalUP 1341373 total 4.7 bits

*1341360
5’ g g a g a g a g t 3’

... p35 1.2 bits

piece 8, NC_000913, p10-dependent-osmE-1820307-atgtattccaggcttatcta, config: linear, direction: -, begin: 1820428, end: 1820268

```
         .         *1820420  .         *1820410  .         *1820400  .
5' c t g a a a c g a c a a g c g a c a g c g g g c t g a a t g g t g g t t a a 3'
```

T timestamp:_2018Sep20_11-34-26

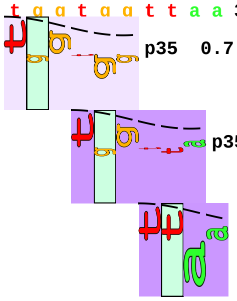
p35 0.7 bits
p35 2.6 bits
... p35

```
 *1820390  .         *1820380  .         *1820370  .         *1820360  .
5' a a c g t c a t a c a g g a a t c c t c t g a a a g c g g a g c c g t t t c 3'
```

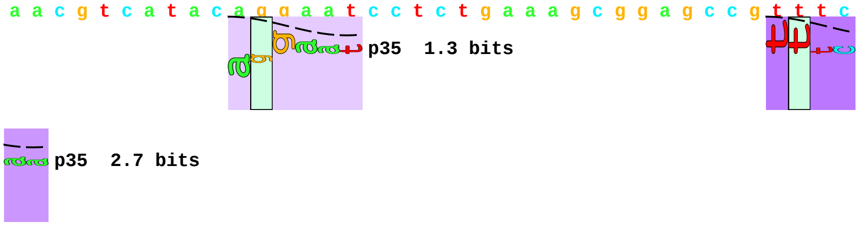
p35 1.3 bits
... p35
... p35 2.7 bits

```
     *1820350  .         *1820340  .         *1820330  .         *1820320  .
5' g t t c a c g g g c t t g a a a a a g c g c c c a a t g t a t t c c a g g c 3'
```

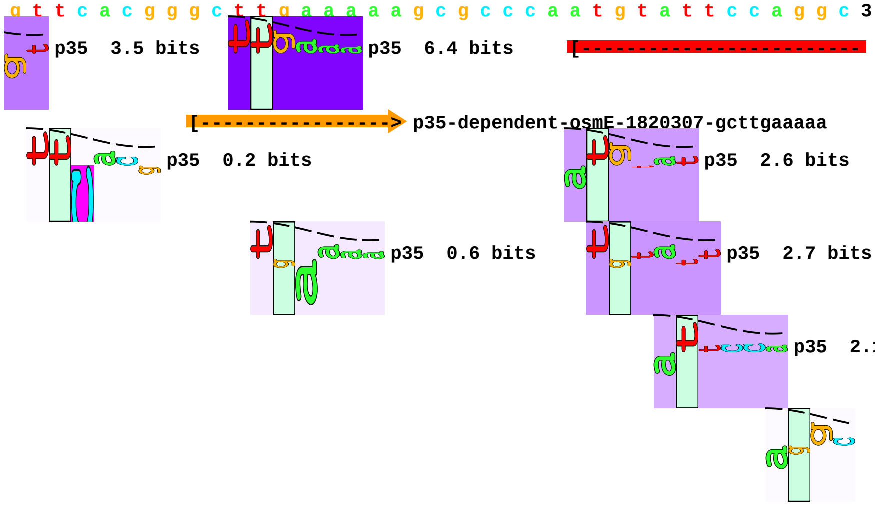
... p35 3.5 bits
p35 6.4 bits
[-------------------------- ... p10-dependent-osmE-1820307-atgtattccaggcttatcta
[-------------------> p35-dependent-osmE-1820307-gcttgaaaaa
p35 0.2 bits
p35 2.6 bits
p35 0.6 bits
p35 2.7 bits
p35 2.1 bits
... p35

```
         *1820310  .         *1820300  .         *1820290  .         *1820280
5' t t a t c t a a c a c g c t g a t a a a c a a g a g g a c g g a a t a t g a 3'
```

... -------------> p10-dependent-osmE-1820307-atgtattccaggcttatcta

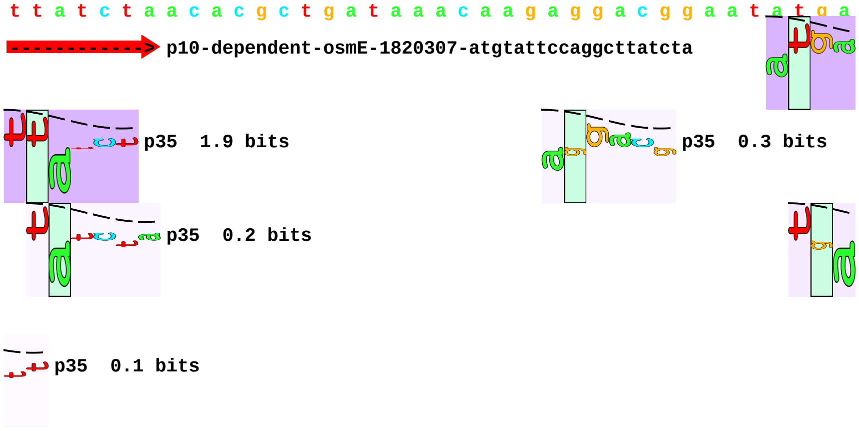
... p35
p35 1.9 bits
p35 0.3 bits
p35 0.2 bits
... p35
... p35 0.1 bits

```
     .         *1820270
5' a c a a g a a t a 3'
```

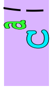
... p35 1.9 bits

... 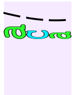 p35 0.5 bits piece 9, NC_000913, p10-dependent-rssA-1288329-acataggctatactcgacag, config: linear, direction: +, begin: 1288188, end: 1288348

```
     *1288190  .         *1288200  .         *1288210  .         *1288220  .
5' a g a g g g a t g c c a g g t c t t t a t t a a a t a a t c t g c g t c t t 3'
```

T timestamp:_2018Sep20_11-34-26

proximalUP 7.2 bits

p35 3.0 bits

... proximalUP-(40)-sigma38p10 1288257 Gap

distalUP 4.7 bits

... p35

... distalUP-(54)-sigma38p10 1288257 Gap

... distalUP-proximalUP-sigma38p10 1288257 total 3.3

p35 0.4 bits

p35 4.8 bits

p35 0.9 bits

p35 0.6 bits

p35 2.5 bits

p35 2.8 bits

```
         *1288230  .         *1288240  .         *1288250  .         *1288260
5' g c a t c a c a a a a g c g c a g t a a c g c g a a c g c a t g a g a t g t 3'
```

sigma38p10 0.4 bits

... proximalUP-(40)-sigma38p10 1288257 Gap 4.0 bits

... p35 6.2 bits

p35 1.5 bits

... distalUP-(54)-sigma38p10 1288257 Gap 4.9 bits

... distalUP-proximalUP-sigma38p10 1288257 total 3.3 bits

... p35

... p35

```
   .         *1288270  .         *1288280  .         *1288290  .         *1288300
5' t c t g g a t c a g g t g c a a c c t t t t c a c c a g a c a c a t a a g g 3'
```

p35 2.1 bits

[-----------------> p35-dependent-rssA-1288329-gacacataag

p35 4.6 bits

p35 3.8 bits

... 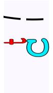 p35 0.2 bits

... 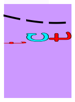 p35 2.6 bits

```
               .          *1288310  .          *1288320  .          *1288330  .
5' g t g g c a a c a t a g g c t a t a c t c g a c a g c a c t a c c a c a g g 3'
```

p10-dependent-rssA-1288329-acataggctatactcgacag

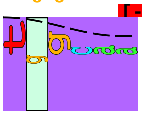 p35 4.0 bits 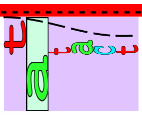 p35 1.5 bits

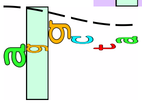 p35 0.0 bits

```
   *1288340  .
5' g a c a a a g c t 3'
```

piece 10, NC_000913, p10-dependent-sibC-3054873-ctttgacctaatttagtgag, config: linear, direction: +, begin: 3054732, end: 3054892

. *3054740 . *3054750 . *3054760 .
5' t g g t g g a a a a a c t c c c c g t t g a a g a g t g g g a t a t c c c t 3'

T timestamp:_2018Sep20_11-34-26

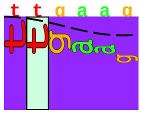
p35 5.5 bits

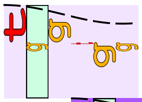
p35 0.7 bits

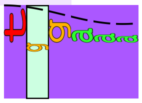
p35 4.3 bits

*3054770 . *3054780 . *3054790 . *3054800 .
5' c t t c c t g c g g t g g t t a c a c c g t c g a a a g t c t g g g a g t g 3'

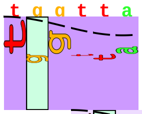
p35 2.6 bits

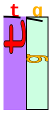
... p35

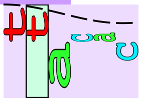
p35 0.9 bits

*3054810 . *3054820 . *3054830 . *3054840 .
5' g t a a g g g c g a t a c a c c c g c a t c g c c c t g a t t g a c a t c g 3'

... 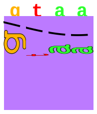
p35 3.4 bits

[------------------> p35-dependent-sibC-3054873-gattgacatc

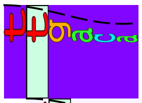
p35 6.4 bits

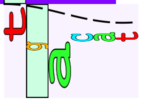
p35 0.3 bits

*3054850 . *3054860 . *3054870 . *3054880
5' t t g a t t c t t t g a c c t a a t t t a g t g a g t a a g g g t a a g g g 3'

[------------------------------------------> p10-dependent-sibC-3054873-ctttgacctaatttagtgag

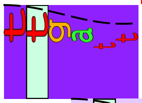
p35 5.7 bits

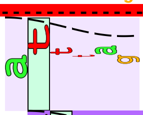
p35 0.7 bits

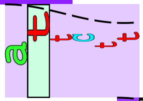
p35 1.4 bits

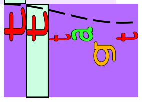
p35 3.8 bits

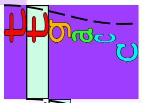
p35 5.0 bits

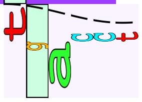
p35 0.3 bits

. *3054890
5' a g g a t t g c t 3'

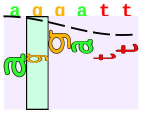
p35 0.5 bits

piece 11, NC_000913, p10-dependent-uspB-3637871-tgatctcctacactatctat, config: linear, direction: -, begin: 3637990, end: 3637830

*3637990 . *3637980 . *3637970 . *3637960 .
5' g a c c c g g a t g a t c c g t c a a t c c g c c t t g c t t a c t c g t t 3'

T timestamp:_2018Sep20_11-34-26

distalUP 1.5 bits

p35 1.1 bits

... distalUP-(50)-sigma38p10 3637912 Gap

... sigma38p10-proximalUP-distalUP 3637912 total 4.4

p35 5.3 bits

p35 0.0 bits

... p35

*3637950 . *3637940 . *3637930 . *3637920 .
5' a g a t a a a c a a a t t t t c g a c a t g a a t a t a t t g a t a g t g g 3'

proximalUP 5.4 bits

proximalUP 6.1 bits

... ... distalUP-(50)-sigma38p10 3637912 Gap
... ... sigma38p10-proximalUP-distalUP 3637912 total 4.4
... proximalUP-(33)-sigma38p10 3637912 Gap

p35 3.7 bits

... proximalUP-(34)-sigma38p10 3637888 Gap

... p35 1.5 bits

distal|UP 7.2 bits

... distalUP-(43)-sigma38p10 3637888 Gap
... sigma38p10-proximalUP-distalUP 3637888 total 10.3

p35 3|4 bits

... proximalUP-(42)-sigma38p10 3637880 Gap

... distalUP-(51)-sigma38p10 3637880 Gap
... sigma38p10-proximalUP-distalUP 3637880 total 5.7

proximalUP 2.3 bits

p35 3.5 bits

... proximalUP-(43)-sigma38p10 3637875 Gap

... distalUP-(56)-sigma38p10 3637875 Gap
... sigma38p10-proximalUP-distalUP 3637875 total 4.2

p35 0.7 bits

p35 5.6 bits

... p35

*3637910 . *3637900 . *3637890 . *3637880
5' t t a a c c t t c t g g a a a a a a a a c a a a c c t g a t c t c c t a c a c 3'

sigma38p10 5.7 bits

[ ... p10-dependent-uspB-3637871-tgatctcctacactatctat

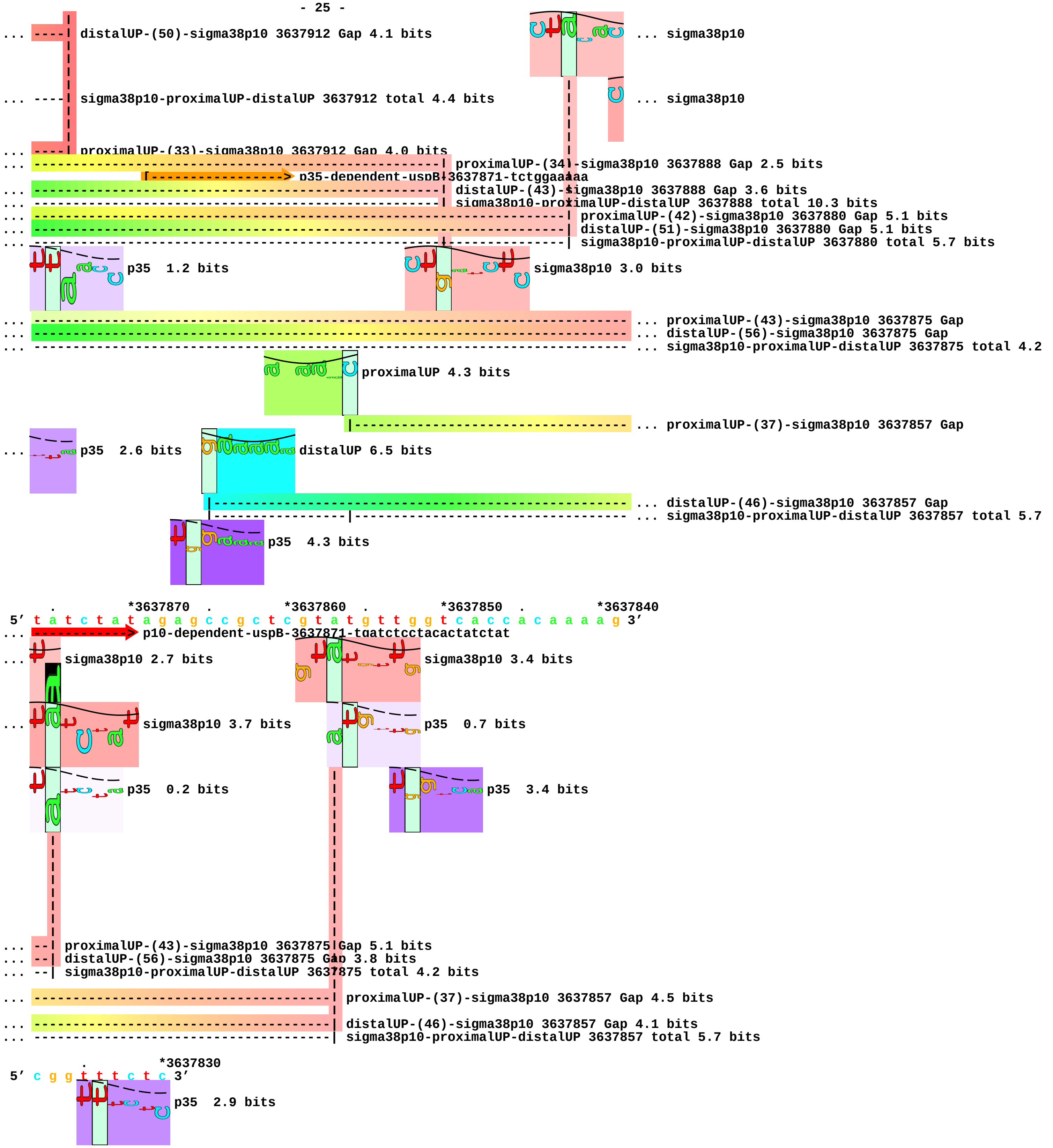

... ----| distalUP-(50)-sigma38p10 3637912 Gap 4.1 bits
... sigma38p10
... ----| sigma38p10-proximalUP-distalUP 3637912 total 4.4 bits
... sigma38p10
... ----| proximalUP-(33)-sigma38p10 3637912 Gap 4.0 bits
... proximalUP-(34)-sigma38p10 3637888 Gap 2.5 bits
p35-dependent-uspB-3637871-tctggaaaaa
... distalUP-(43)-sigma38p10 3637888 Gap 3.6 bits
... sigma38p10-proximalUP-distalUP 3637888 total 10.3 bits
... proximalUP-(42)-sigma38p10 3637880 Gap 5.1 bits
... distalUP-(51)-sigma38p10 3637880 Gap 5.1 bits
... sigma38p10-proximalUP-distalUP 3637880 total 5.7 bits
p35 1.2 bits
sigma38p10 3.0 bits
... proximalUP-(43)-sigma38p10 3637875 Gap
... distalUP-(56)-sigma38p10 3637875 Gap
... sigma38p10-proximalUP-distalUP 3637875 total 4.2
proximalUP 4.3 bits
... proximalUP-(37)-sigma38p10 3637857 Gap
... p35 2.6 bits
distalUP 6.5 bits
... distalUP-(46)-sigma38p10 3637857 Gap
... sigma38p10-proximalUP-distalUP 3637857 total 5.7
p35 4.3 bits
. *3637870 . *3637860 . *3637850 . *3637840
5′ t a t c t a t a g a g c c g c t c g t a t g t t g g t c a c c a c a a a a g 3′
... p10-dependent-uspB-3637871-tgatctcctacactatctat
... sigma38p10 2.7 bits
sigma38p10 3.4 bits
... sigma38p10 3.7 bits
p35 0.7 bits
p35 0.2 bits
p35 3.4 bits
... --| proximalUP-(43)-sigma38p10 3637875 Gap 5.1 bits
... --| distalUP-(56)-sigma38p10 3637875 Gap 3.8 bits
... --| sigma38p10-proximalUP-distalUP 3637875 total 4.2 bits
... proximalUP-(37)-sigma38p10 3637857 Gap 4.5 bits
... distalUP-(46)-sigma38p10 3637857 Gap 4.1 bits
... sigma38p10-proximalUP-distalUP 3637857 total 5.7 bits
. *3637830
5′ c g g t t t c t c 3′
p35 2.9 bits

piece 12, NC_000913, p10-dependent-ybiI-837707-ggattagctacacttaactg, config: linear, direction: -, begin: 837827, end: 837667

```
           .         *837820   .         *837810   .         *837800   .         *837790
5' t a c g g c g a a a g t g a a g a g a t c c t g t c g c t g c t t a a t c t 3'
T timestamp:_2018Sep20_11-34-26
                                                                       proximalUP 6.8 bits
                                                                       |--- ... proximalUP-(38)-sigma38p10 837753 Gap
                                                                       distalUP 4.7 bits
                                                                       |-------------- ... distalUP-(44)-sigma38p10 837753 Gap
                                                                       |-----------|--- ... sigma38p10-proximalUP-distalUP 837753 total 7.2
                                                                       ... proximalUP
                                                                       ... distalUP
                                                                       |--- ... distalUP-(44)-sigma38p10 837747 Gap
                                                                       |--- ... sigma38p10-proximalUP-distalUP 837747 total 8.1
                                                                       p35  0.4 bits
                                                                       ... p35

             .         *837780   .         *837770   .         *837760   .
5' t t a t c a t a a t c t g c t g c g g g a a t g g t c g g a g a t c t g a t 3'
p35  1.8 bits                                                          ... sigma38p10
... -------------------------------------------------------------| proximalUP-(38)-sigma38p10 837753 Gap 3.2 bits
    |---------------------------------------------------------------- ... proximalUP-(38)-sigma38p10 837747 Gap
... -------------------------------------------------------------| distalUP-(44)-sigma38p10 837753 Gap 2.8 bits
... -------------------------------------------------------------| sigma38p10-proximalUP-distalUP 837753 total 7.2 bits
... proximalUP 6.8 bits                 p35  2.5 bits              ... p35
... distalUP 4.7 bits
... ----------------------------------------------------------------- ... distalUP-(44)-sigma38p10 837747 Gap
... --------|-------------------------------------------------------- ... sigma38p10-proximalUP-distalUP 837747 total 8.1
    p35  1.2 bits
... p35  3.3 bits

     *837750   .         *837740   .         *837730   .         *837720   .
5' g c t t g a a c t g c c c g a g t t c g t c c c t g g a t t a g c t a c a c 3'
... sigma38p10 1.8 bits                     [-------------------------- ... p10-dependent-ybiI-837707-ggattagctacacttaactg
              [------------------> p35-dependent-ybiI-837707-tacccgagtt
... --------| proximalUP-(38)-sigma38p10 837747 Gap 3.2 bits          p35  1.1 bits
    sigma38p10 2.7 bits                     p35  3.6 bits
    p35  5.0 bits
```

```
...                 p35   2.2 bits

                          p35   0.7 bits
... --------| distalUP-(44)-sigma38p10 837747 Gap 2.8 bits
... --------| sigma38p10-proximalUP-distalUP 837747 total 8.1 bits

        *837710    .         *837700    .         *837690    .         *837680
 5' t t a a c t g t a c a a g t a t t g a t a t g g g g a g g t c g a t a t g g 3'
... -------------> p10-dependent-ybiI-837707-ggattagctacacttaactg          ... p35

                  p35   2.8 bits                  p35   5.6 bits
                        p35   3.4 bits

    .         *837670
 5' c a t c c g g t t 3'
...       p35   4.1 bits
```

piece 13, NC_000913, p10-dependent-ycgB-1236508-caggtgttctatgcttgaaa, config: linear, direction: -, begin: 1236630, end: 1236470

```
   *1236630  .         *1236620  .         *1236610  .         *1236600  .
5' c c c t t t t a t c c a g c a t a a a a t t c c g t t c a g a a g c g g a 3'
T timestamp:_2018Sep20_11-34-26                     proximalUP 5.5 bits          ... p35
               proximalUP 6.8 bits
                                                                                 ... proximalUP-(31)-sigma38p10 1236576 Gap
                                                                                 ... proximalUP-(36)-sigma38p10 1236584 Gap
          distalUP 4.1 bits                           p35   1.2 bits
                                                                                 ... distalUP-(44)-sigma38p10 1236584 Gap
                                                                                 ... sigma38p10-proximalUP-distalUP 1236584 total 5.0
                              distalUP 5.6 bits
                                                                                 ... distalUP-(40)-sigma38p10 1236576 Gap
                                                                                 ... sigma38p10-proximalUP-distalUP 1236576 total 4.0
            p35   3.8 bits
              p35   4.8 bits
                p35   3.3 bits
                  p35   0.3 bits
                    p35   1.1 bits

       *1236590  .         *1236580  .         *1236570  .         *1236560  .
5' t t a g t g g c a c t c t g a t a a t a a g a a a t c g c a a a g a a a a a 3'
...     p35   0.7 bits                 sigma38p10 0.6 bits                       ... proximalUP
... ---------------------------------| proximalUP-(31)-sigma38p10 1236576 Gap 4.7 bits
... ----------------| proximalUP-(36)-sigma38p10 1236584 Gap 3.6 bits
                      sigma38p10 0.5 bits                       distalUP 6.5 bits
... ----------------| distalUP-(44)-sigma38p10 1236584 Gap 2.8 bits |----------- ... distalUP-(41)-sigma38p10 1236519 Gap
... ----------------| sigma38p10-proximalUP-distalUP 1236584 total 5.0 bits
              p35   2.5 bits                                       |----------- ... sigma38p10-proximalUP-distalUP 1236519 total 14.0
... ---------------------------------| distalUP-(40)-sigma38p10 1236576 Gap 3.1 bits
... ---------------------------------| sigma38p10-proximalUP-distalUP 1236576 total 4.0 bits
                                                                   |----------- ... distalUP-(47)-sigma38p10 1236513 Gap
                                                                   |----------- ... sigma38p10-proximalUP-distalUP 1236513 total 3.3

          *1236550  .         *1236540  .         *1236530  .         *1236520
5' a a t t c t c c g g c g a c g c g g t t t t t a a c a g g t g t t c t a t g 3'
...     proximalUP 5.5 bits                            [------------------------ ... p10-dependent-ycgB-1236508-caggtgttctatgcttgaaa
           [------------------> p35-dependent-ycgB-1236508-cggcgacgcg
                                       p35   3.8 bits                            ... sigma38p10
         |-------------------------------------------------------------| proximalUP-(31)-sigma38p10 1236519 Gap 4.7 bits
```

... ---------------------------------------------------------------------| distalUP-(41)-sigma38p10 1236519 Gap 3.1 bits
|------------------------------------------------------------------- ... proximalUP-(37)-sigma38p10 1236513 Gap
... --------|-----------------------------------------------------------| sigma38p10-proximalUP-distalUP 1236519 total 14.0 bits

p35 4.7 bits ... p35

p35 4.1 bits

... ---------------------------------------------------------------------------- ... distalUP-(47)-sigma38p10 1236513 Gap
... --------|------------------------------------------------------------------- ... sigma38p10-proximalUP-distalUP 1236513 total 3.3

p35 2.7 bits

p35 2.6 bits

p35 0.3 bits

. *1236510 . *1236500 . *1236490 . *1236480

5’ c t t g a a a t g a g g t g c t c c a c a c a g a g a g c g c c g a c a a c 3’

... -------------> p10-dependent-ycgB-1236508-caggtgttctatgcttgaaa

sigma38p10 2.1 bits

... sigma38p10 9.7 bits

p35 6.4 bits

p35 0.7 bits

... ------| proximalUP-(37)-sigma38p10 1236513 Gap 4.5 bits

p35 0.6 bits

... p35 2.2 bits

... ------| distalUP-(47)-sigma38p10 1236513 Gap 6.5 bits
... ------| sigma38p10-proximalUP-distalUP 1236513 total 3.3 bits

. *1236470

5’ a a t g a g g t g 3’

p35 0.6 bits

piece 14, NC_000913, p10-dependent-ychH-1257961-gtgttgtctatactttacac, config: linear, direction: +, begin: 1257821, end: 1257981

```
           .         *1257830   .         *1257840   .         *1257850   .
5' a g a t a a c g t a a t a a t t g c c t g a g t g g a c t a t t a g a a a g 3'
```

T timestamp:_2018Sep20_11-34-26

proximalUP 2.8 bits

distalUP 4.4 bits

... proximalUP-(30)-sigma38p10 1257884 Gap

... distalUP-(56)-sigma38p10 1257884 Gap

... distalUP-proximalUP-sigma38p10 1257884 total 2.3

p35 6.2 bits

p35 0.6 bits

p35 4.4 bits

p35 2.3 bits

```
    *1257860   .         *1257870   .         *1257880   .         *1257890   .
5' t c a a g g t g t t c a g g c g t t t a t t t g t a a a g t t t t g t t g a 3'
```

p35 2.5 bits

sigma38p10 4.8 bits

... proximalUP-(30)-sigma38p10 1257884 Gap 5.9 bits

... distalUP-(56)-sigma38p10 1257884 Gap 3.8 bits

... distalUP-proximalUP-sigma38p10 1257884 total 2.3 bits

p35 4.8 bits

p35 2.9 bits

p35 1.0 bits

p35 3.8 bits

p35 5.5 bits

... p35

p35 3.4 bits

... p35

p35 0.7 bits

```
        *1257900   .         *1257910   .         *1257920   .         *1257930
5' a a t a a g g g t t g t a a t t g t g a t c a c g c c c g c a c a t a a c c 3'
```

proximalUP 4.6 bits [-----------------> p35-dependent-ychH-1257961-cgcacataac

... proximalUP-(37)-sigma38p10 1257950 Gap

distalUP 5.0 bits

... distalUP-(43)-sigma38p10 1257950 Gap

... distalUP-proximalUP-sigma38p10 1257950 total 12.5

p35 5.5 bits

... p35 6.4 bits

p35 3.6 bits

... p35 0.7 bits

p35 3.7 bits

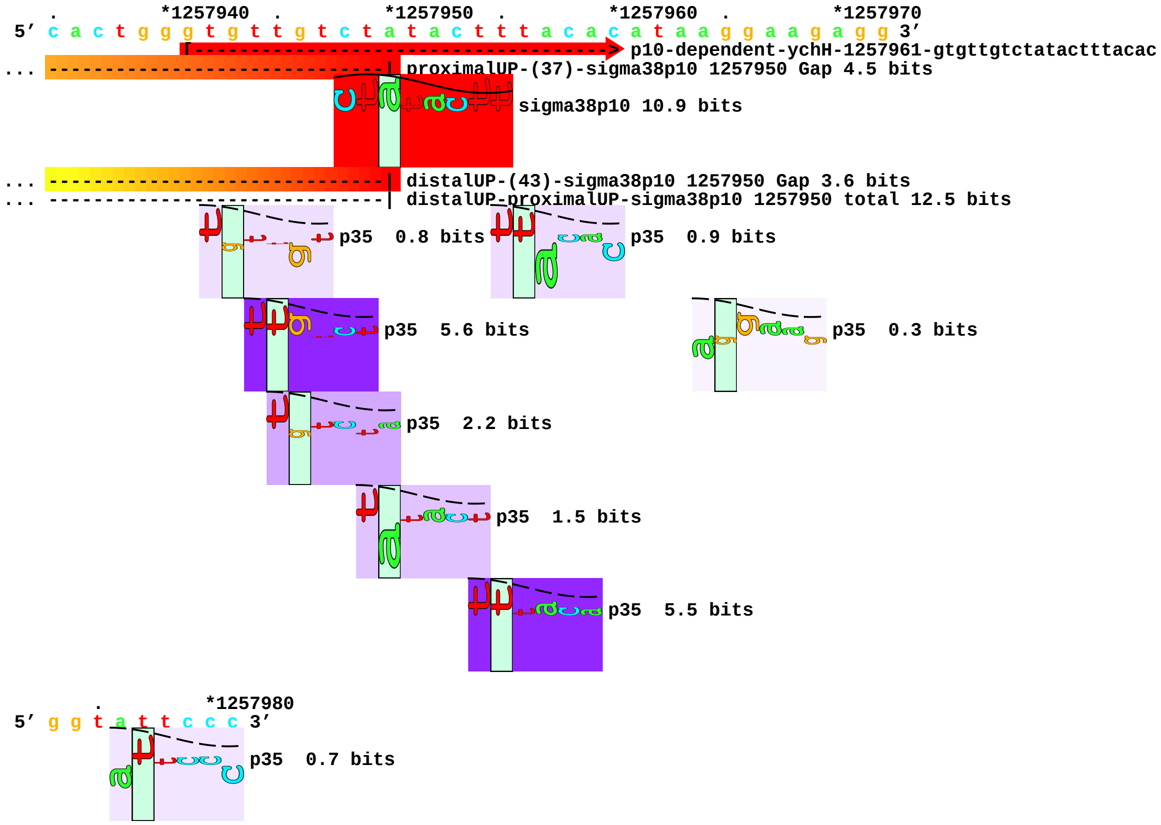
*1257940
*1257950
*1257960
*1257970
5′ c a c t g g g t g t t g t c t a t a c t t t a c a c a t a a g g a a g a g g 3′
p10-dependent-ychH-1257961-gtgttgtctatactttacac
proximalUP-(37)-sigma38p10 1257950 Gap 4.5 bits
sigma38p10 10.9 bits
distalUP-(43)-sigma38p10 1257950 Gap 3.6 bits
distalUP-proximalUP-sigma38p10 1257950 total 12.5 bits
p35 0.8 bits
p35 0.9 bits
p35 5.6 bits
p35 0.3 bits
p35 2.2 bits
p35 1.5 bits
p35 5.5 bits
*1257980
5′ g g t a t t c c c 3′
p35 0.7 bits

piece 15, NC_000913, p10-dependent-ydbK-1438870-ccatttgctaaaattgaaag, config: linear, direction: -, begin: 1438990, end: 1438830

*1438990 . *1438980 . *1438970 . *1438960 .

5' c c g c a t t t t t a c t t t t t t a t g g c t a t t t t t t t g c c c t c 3'

T timestamp:_2018Sep20_11-34-26

... proximalUP-(32)-sigma38p10 1438938 Gap

proximalUP 6.3 bits

... proximalUP-(27)-sigma38p10 1438931 Gap

distalUP 4.1 bits

proximalUP 5.7 bits

... distalUP-(41)-sigma38p10 1438938 Gap

... sigma38p10-proximalUP-distalUP 1438938 total 4.3

distalUP 5.7 bits

p35 3.1 bits

... proximalUP-(32)-sigma38p10 1438926 Gap

... distalUP-(56)-sigma38p10 1438931 Gap

... sigma38p10-proximalUP-distalUP 1438931 total 8.1

p35 0.8 bits

distalUP 4.2 bits

p35 3.8 bits

... distalUP-(42)-sigma38p10 1438926 Gap

p35 3.2 bits

... sigma38p10-proximalUP-distalUP 1438926 total 2.7

p35 5.7 bits

p35 0.8 bits

p35 1.6 bits

p35 3.9 bits

p35 3.9 bits

p35 3.9 bits

p35 3.8 bits

p35 2.9 bits

p35 4.8 bits

p35 1.4 bits

p35 3.8 bits

p35 4.6 bits

*1438950 . *1438940 . *1438930 . *1438920 .

5' t g t t t g a t c a a a a c a t t c a t t a c g c t g a t g t g g g g g a c 3'

... proximalUP-(32)-sigma38p10 1438938 Gap 4.3 bits

... proximalUP-(27)-sigma38p10 1438931 Gap 3.8 bits

sigma38p10 1.4 bits

... distalUP-(41)-sigma38p10 1438938 Gap 3.1 bits

... sigma38p10-proximalUP-distalUP 1438938 total 4.3 bits

... proximalUP-(32)-sigma38p10 1438926 Gap 4.3 bits

... distalUP-(56)-sigma38p10 1438931 Gap 3.8 bits

... sigma38p10-proximalUP-distalUP 1438931 total 8.1 bits

p35 0.8 bits

sigma38p10 4.3 bits

... distalUP-(42)-sigma38p10 1438926 Gap 3.3 bits

... sigma38p10-proximalUP-distalUP 1438926 total 2.7 bits

p35 4.2 bits

sigma38p10 0.4 bits

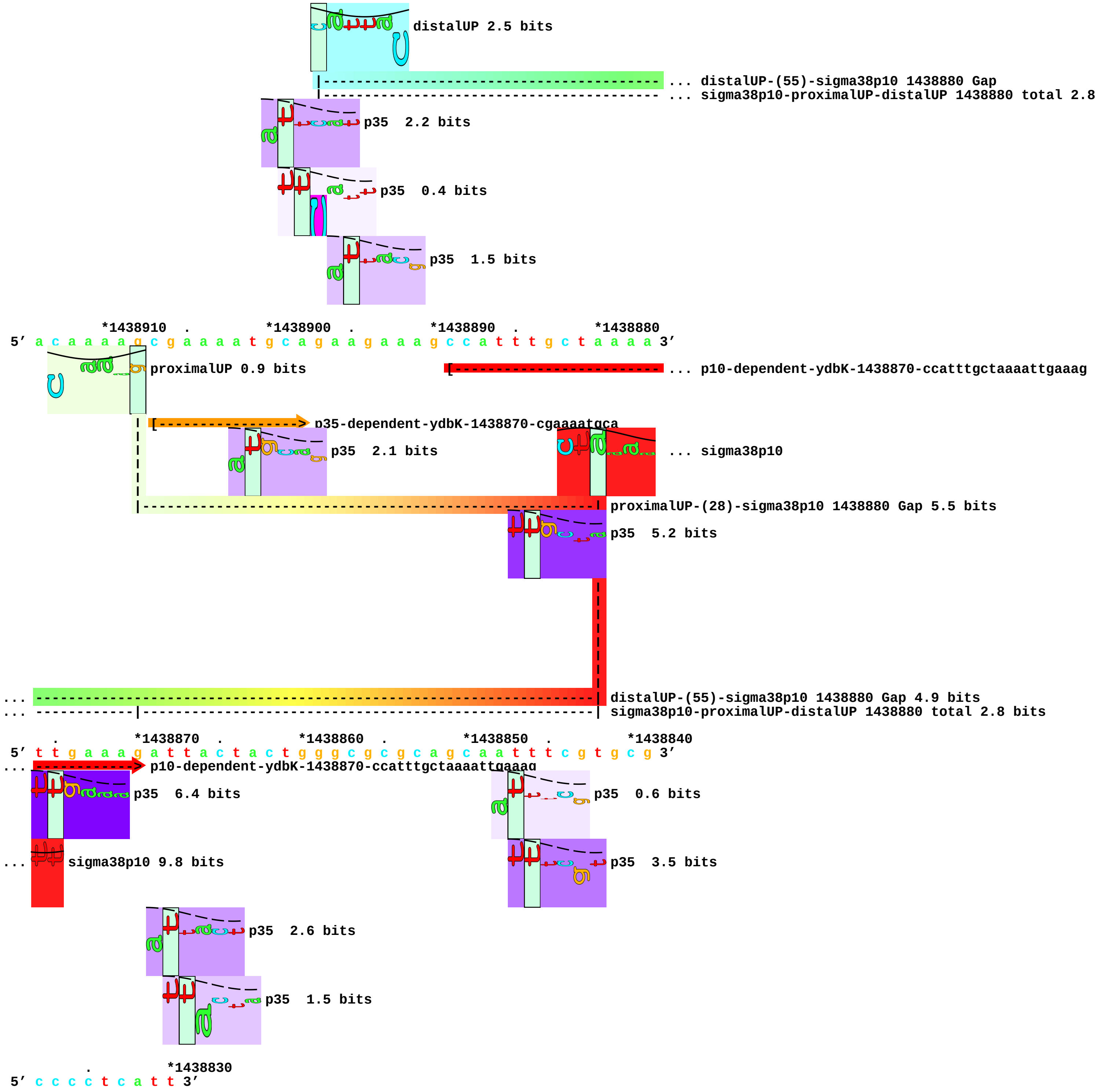
distalUP 2.5 bits
... distalUP-(55)-sigma38p10 1438880 Gap
... sigma38p10-proximalUP-distalUP 1438880 total 2.8
p35 2.2 bits
p35 0.4 bits
p35 1.5 bits
*1438910 . *1438900 . *1438890 . *1438880
5’ a c a a a a g c g a a a a t g c a g a a g a a a g c c a t t t g c t a a a a 3’
proximalUP 0.9 bits
... p10-dependent-ydbK-1438870-ccatttgctaaaattgaaag
p35-dependent-ydbK-1438870-cgaaaatgca
p35 2.1 bits
... sigma38p10
proximalUP-(28)-sigma38p10 1438880 Gap 5.5 bits
p35 5.2 bits
... distalUP-(55)-sigma38p10 1438880 Gap 4.9 bits
... sigma38p10-proximalUP-distalUP 1438880 total 2.8 bits
. *1438870 . *1438860 . *1438850 . *1438840
5’ t t g a a a g a t t a c t a c t g g g c g c g c a g c a a t t t c g t g c g 3’
... p10-dependent-ydbK-1438870-ccatttgctaaaattgaaag
p35 6.4 bits
p35 0.6 bits
... sigma38p10 9.8 bits
p35 3.5 bits
p35 2.6 bits
p35 1.5 bits
. *1438830
5’ c c c c t c a t t 3’

piece 16, NC_000913, p10-dependent-ydcS-1509623-cgccggtctatggttaaaca, config: linear, direction: +, begin: 1509480, end: 1509640

```
   *1509480  .          *1509490  .          *1509500  .          *1509510  .
5' a t a g c a g t c a t a t t t c c t a a c t c c t t g a c t a t a c t c c a 3'
```

T timestamp:_2018Sep20_11-34-26

p35 6.5 bits

p35 1.1 bits

p35 1.5 bits

p35 0.1 bits

p35 0.1 bits

p35 5.3 bits

```
     *1509520  .          *1509530  .          *1509540  .          *1509550  .
5' g a a g a t a a c c t t a c a g a c g g c a t a a t g c g c g g t a g c t c 3'
```

proximalUP 0.2 bits

|--------------------------------------------------------------- ... proximalUP-(40)-sigma38p10 1509567 Gap

distalUP 3.3 bits

|----------------------------------------------------------------------------- ... distalUP-(46)-sigma38p10 1509567 Gap

|-----------|---------------------------------------------------------------- ... distalUP-proximalUP-sigma38p10 1509567 total 2.3

p35 1.4 bits

```
         *1509560  .          *1509570  .          *1509580  .          *1509590
5' a c a a c c t g a a t a a a t t t t c t c a g g g g c g a a g g t g t g c c 3'
```

distalUP 6.5 bits

[-----------------> p35-dependent-ydcS-1509623-cgaaggtgtg

... -----------------------| proximalUP-(40)-sigma38p10 1509567 Gap 4.0 bits

sigma38p10 7.0 bits

... ----------------------| distalUP-(46)-sigma38p10 1509567 Gap 4.1 bits

... ----------------------| distalUP-proximalUP-sigma38p10 1509567 total 2.3 bits

proximalUP 6.6 bits

|---------------------------------------- ... proximalUP-(35)-sigma38p10 1509609 Gap

|---------------------------------------------------------------- ... distalUP-(46)-sigma38p10 1509609 Gap

|---------------------|------------------------------------------ ... distalUP-proximalUP-sigma38p10 1509609 total 12.8

p35 4.8 bits

p35 2.9 bits

p35 0.2 bits

```
   .          *1509600  .          *1509610  .          *1509620  .          *1509630
5' t g c a a g c c g c c g t c t a t g g t t a a a c a a g g a g a t a t t t t 3'
```

[-g-c---g-----------------------------------> p10-dependent-ydcS-1509623-cgccggtctatggttaaaca.1.2.4

sigma38p10 6.8 bits

... p35

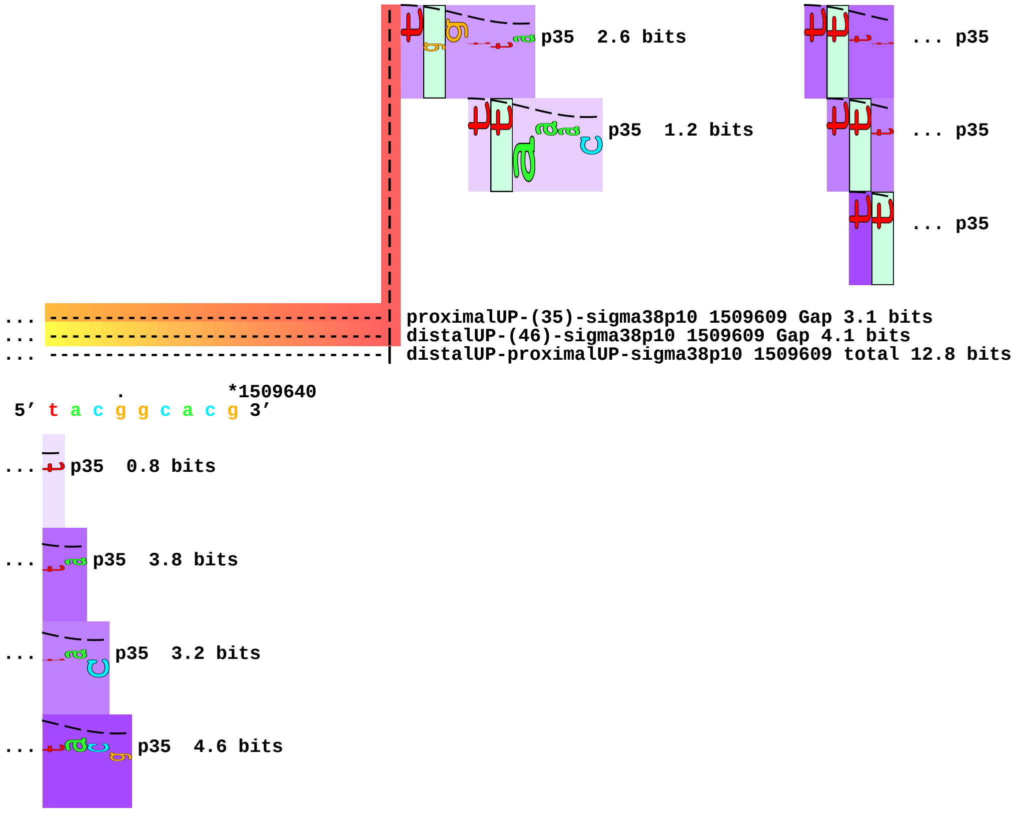
p35 2.6 bits
... p35
p35 1.2 bits
... p35
... p35
... proximalUP-(35)-sigma38p10 1509609 Gap 3.1 bits
... distalUP-(46)-sigma38p10 1509609 Gap 4.1 bits
... distalUP-proximalUP-sigma38p10 1509609 total 12.8 bits
. *1509640
5' t a c g g c a c g 3'
... p35 0.8 bits
... p35 3.8 bits
... p35 3.2 bits
... p35 4.6 bits

piece 17, NC_000913, p10-dependent-ydgA-1687818-tctcttgctaagcttattaa, config: linear, direction: +, begin: 1687678, end: 1687838

*1687680 . *1687690 . *1687700 . *1687710 .

5′ a g c t t a a a c c g g g t g a a t c a g c g t t t a t t g c c g c c a a c 3′

T timestamp:_2018Sep20_11-34-26

p35 4.8 bits

p35 1.2 bits

p35 0.0 bits

p35 5.2 bits

*1687720 . *1687730 . *1687740 . *1687750

5′ g a a t c a c c g g t g a c t g t c a a a g g c c a c g g c c g t t t a g c 3′

p35 3.1 bits

p35 2.3 bits

p35 0.8 bits

. *1687760 . *1687770 . *1687780 . *1687790

5′ g c g t g t t t a c a a c a a g c t g t a a g a g c t t a c t g a a a a a a a 3′

distalUP 0.9 bits

[------------------> p35-dependent-ydgA-1687818-tactgaaaaa

p35 1.7 bits

proximalUP 2.8 bits

... p35

p35 5.5 bits

|---------------------------------- ... proximalUP-(31)-sigma38p10 1687807 Gap

|---------------------------------------------------------- ... distalUP-(44)-sigma38p10 1687807 Gap

|--------------------------|-------------------------------- ... distalUP-proximalUP-sigma38p10 1687807 total 5.4

p35 2.3 bits

p35 0.6 bits

p35 2.6 bits

p35 0.6 bits

. *1687800 . *1687810 . *1687820 .

5′ t t a a c a t c t c t t g c t a a g c t t a t t a a a g g c t t a t a a c a 3′

[-------------------------------------> p10-dependent-ydgA-1687818-tctcttgctaagcttattaa

... p35 1.0 bits

sigma38p10 9.1 bits

p35 0.0 bits

... ---------------------------------| proximalUP-(31)-sigma38p10 1687807 Gap 4.7 bits

... ---------------------------------| distalUP-(44)-sigma38p10 1687807 Gap 2.8 bits

... ---------------------------------| distalUP-proximalUP-sigma38p10 1687807 total 5.4 bits

p35 2.7 bits

p35 0.9 bits

p35 5.2 bits

p35 0.1 bits

p35 0.6 bits

p35 2.5 bits

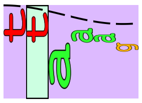

p35 1.8 bits

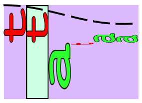

p35 1.8 bits

 *1687830 .
5’ c c t t c a g g c 3’

piece 18, NC_000913, p10-dependent-yggE-3066148-agaggcgctaagcttgcctc, config: linear, direction: -, begin: 3066270, end: 3066110

```
   *3066270   .       *3066260   .       *3066250   .       *3066240   .
5' g g g a t g t g t t a t g t g g t t t a t t g c c t t g c a g c t g g c g a 3'
```

T timestamp:_2018Sep20_11-34-26

p35 4.8 bits
p35 2.1 bits
p35 0.7 bits
p35 0.0 bits
p35 2.7 bits
p35 6.2 bits
p35 0.1 bits
p35 5.2 bits
p35 2.7 bits

```
     *3066230   .       *3066220   .       *3066210   .       *3066200   .
5' g a g a c g g t a t t g c t c a t g c a c a a g c c t t g t t c a g t t a g 3'
```

p35 3.8 bits
proximalUP 2.7 bits
p35 1.6 bits
|- ... proximalUP-(35)-sigma38p10 3066160 Gap
distalUP 1.3 bits
... distalUP-(41)-sigma38p10 3066160 Gap
|-----------|- ... sigma38p10-proximalUP-distalUP 3066160 total 6.9
p35 3.3 bits
p35 2.5 bits

```
         *3066190   .       *3066180   .       *3066170   .       *3066160
5' g c g c t a t c t g a t g g a a a a a t a a a a c a g a g g c g c t a a g c 3'
```

p35 4.3 bits
[---------------------------- ... p10-dependent-yggE-3066148-agaggcgctaagcttgcctc

... ---------------------------------------------------------------------- proximalUP-(35)-sigma38p10 3066160 Gap 3.1 bits
[-----------------> p35-dependent-yggE-3066148-tgatggaaaaa
... ---------------------------------------------------------------------- distalUP-(41)-sigma38p10 3066160 Gap 3.1 bits
... ---------------------------------------------------------------------- sigma38p10-proximalUP-distalUP 3066160 total 6.9 bits
distalUP 6.5 bits
... sigma38p10
proximalUP 3.9 bits
|-------------------------- ... proximalUP-(31)-sigma38p10 3066139 Gap
|------------------------------------------------------ ... distalUP-(42)-sigma38p10 3066139 Gap
|--------------------|------------------------------ ... sigma38p10-proximalUP-distalUP 3066139 total 8.4
|-------------------------- ... proximalUP-(32)-sigma38p10 3066138 Gap
|------------------------------------------------------ ... distalUP-(43)-sigma38p10 3066138 Gap
|--------------------|------------------------------ ... sigma38p10-proximalUP-distalUP 3066138 total 2.8

```
     .     *3066150   .       *3066140   .       *3066130   .       *3066120
5' t t g c c t c c a g a g g t c c t g a a t t t t g g g c a a g t c a a t t a 3'
```

... -------------> p10-dependent-yggE-3066148-agaggcgctaagcttgcctc
... p35

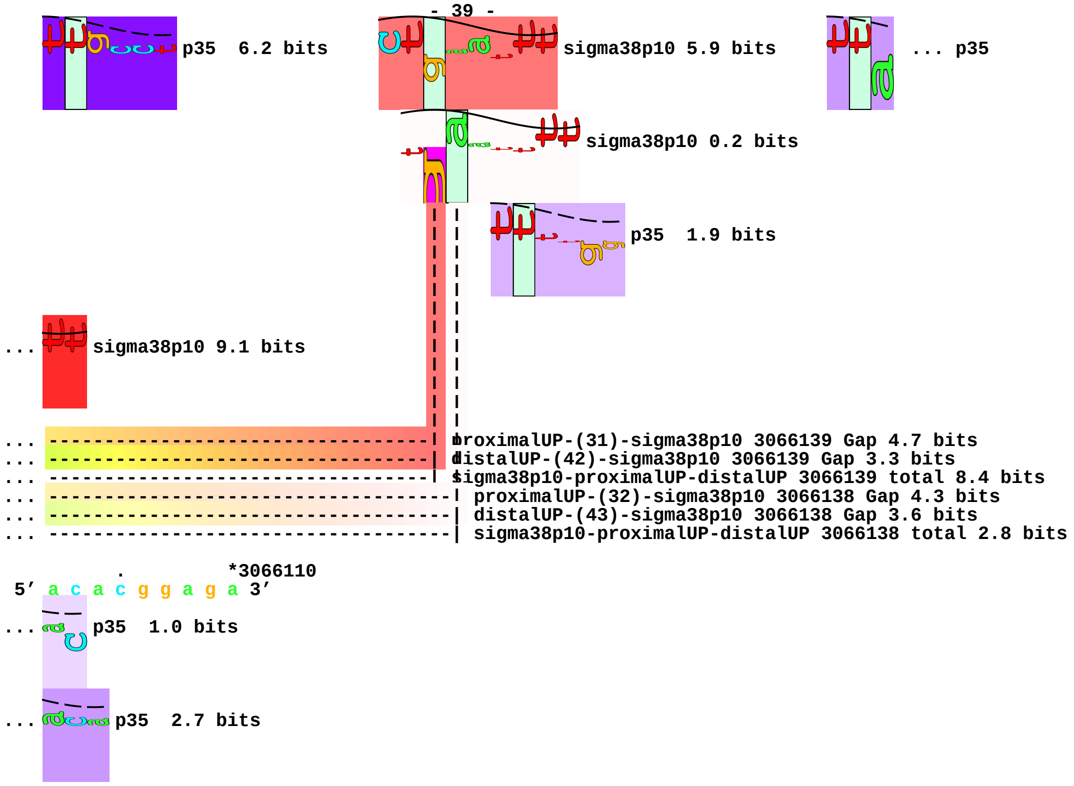
p35 6.2 bits
sigma38p10 5.9 bits
... p35
sigma38p10 0.2 bits
p35 1.9 bits
... sigma38p10 9.1 bits
... proximalUP-(31)-sigma38p10 3066139 Gap 4.7 bits
... distalUP-(42)-sigma38p10 3066139 Gap 3.3 bits
... sigma38p10-proximalUP-distalUP 3066139 total 8.4 bits
... proximalUP-(32)-sigma38p10 3066138 Gap 4.3 bits
... distalUP-(43)-sigma38p10 3066138 Gap 3.6 bits
... sigma38p10-proximalUP-distalUP 3066138 total 2.8 bits
*3066110
5' a c a c g g a g a 3'
... p35 1.0 bits
... p35 2.7 bits

piece 19, NC_000913, p10-dependent-yjdC-4361353-ccgtgaacgacacttgcagt, config: linear, direction: -, begin: 4361474, end: 4361314

*4361470 . *4361460 . *4361450 . *4361440

5' c a a t t c t c t t t t t t g a c g g a c a a g g c c a g g a g c a t c c a 3'

T timestamp:_2018Sep20_11-34-26

p35 0.8 bits

p35 0.3 bits

p35 3.9 bits

p35 2.9 bits

p35 2.8 bits

p35 5.5 bits

. *4361430 . *4361420 . *4361410 . *4361400

5' c a a g c a c g c g t c a c g g g c t t t a t g g a t g c t g a a a c c t t 3'

[---t-------c-g---a p35-dependent-yjdC-4361353-ttttggcgga.2.6.7.9

distalUP 4.7 bits t----- ... p35-dependent-yjdC-4361353-ttttggcgga..3.4.8

p35 3.8 bits <- ... p35-dependent-yjdC-4361353-ttttggcgga.7.4.3.1

proximalUP 3.3 bits

|------- ... proximalUP-(38)-sigma38p10 4361364 Gap

|--------------------------------------------------- ... distalUP-(55)-sigma38p10 4361364 Gap

|--------------------------------|------- ... sigma38p10-proximalUP-distalUP 4361364 total 3.5

p35 2.6 bits

p35 1.2 bits

. *4361390 . *4361380 . *4361370 .

5' c a g c g c a c a t t t g c g c g a t c g c c a a c c g t g a a c g a c a c 3'

p35 2.8 bits [------------------------- ... p10-dependent-yjdC-4361353-ccgtgaacgacacttgcagt

... t-g-------g-> p35-dependent-yjdC-4361353-ttttggcgga..3.4.8 ... sigma38p10

... --c-----c-a---a-] p35-dependent-yjdC-4361353-ttttggcgga.7.4.3.1

t---------g-c-g---> p35-dependent-yjdC-4361353-ttttg[illegible]cgga..5.6.7

... ------------------------------------------------------------| proximalUP-(38)-sigma38p10 4361364 Gap 3.2 bits

... ------------------------------------------------------------| distalUP-(55)-sigma38p10 4361364 Gap 4.9 bits

... ------------------------------------------------------------| sigma38p10-proximalUP-distalUP 4361364 total 3.5 bits

[-----t-g-------g-a p35-dependent-yjdC-4361353-ttttggcgga.3.4.8.9

<-c-------------a-a p35-dependent-yjdC-4361353-ttttggcgga.9.8.1.

*4361360 . *4361350 . *4361340 . *4361330 .

5' t t g c a g t g g g a a a g a c g g a g g a a a a t a g c g t g c a a c g t 3'

... -------------> p10-dependent-yjdC-4361353-ccgtgaacgacacttgcagt

... sigma38p10 3.5 bits p35 1.2 bits

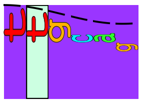
p35 5.2 bits

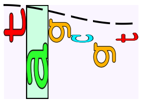
p35 0.3 bits

```
     *4361320  .
5′ g a a g a t g t a 3′
```

piece 20, NC_000913, p10-dependent-ytfJ-4437309-ccaagtggtcggatcacctg, config: linear, direction: -, begin: 4437428, end: 4437268

```
       .         *4437420   .         *4437410   .         *4437400   .
5' g t a t a a g c a t t c a t t t t t c a t a t g a t t t g t g c g c t t g a 3'
T timestamp: 2018Sep20_11-34-26                                      ... proximalUP-(35)-sigma38p10 4437375 Gap
proximalUP 3.4 bits          p35   2.6 bits                          ... p35
                                                                     ... proximalUP-(36)-sigma38p10 4437386 Gap
distalUP 4.9 bits     p35   5.3 bits
                                                                     ... distalUP-(42)-sigma38p10 4437386 Gap
                                                                     ... sigma38p10-proximalUP-distalUP 4437386 total 3.7
proximalUP 6.2 bits
distalUP 5.7 bits     p35   2.3 bits
                                                                     ... distalUP-(42)-sigma38p10 4437375 Gap
                                                                     ... sigma38p10-proximalUP-distalUP 4437375 total 10.7
p35   0.6 bits     p35   4.6 bits
p35   2.2 bits
p35   0.4 bits
p35   0.8 bits
p35   2.4 bits
p35   0.2 bits

 *4437390   .         *4437380   .         *4437370   .         *4437360   .
5' c c g c a a a c t g g c a t c a c a c t t g c g g g a a a t t c g a t a a a 3'
... proximalUP-(35)-sigma38p10 4437375 Gap 3.1 bits
... p35   5.0 bits     sigma38p10 5.2 bits     distalUP 6.5 bits
... proximalUP-(36)-sigma38p10 4437386 Gap 3.6 bits     ... distalUP-(50)-sigma38p10 4437308 Gap
sigma38p10 2.2 bits                                     ... sigma38p10-proximalUP-distalUP 4437308 total 4.6
... distalUP-(42)-sigma38p10 4437386 Gap 3.3 bits     p35   0.3 bits
... sigma38p10-proximalUP-distalUP 4437386 total 3.7 bits
p35   4.1 bits
p35   3.4 bits
... distalUP-(42)-sigma38p10 4437375 Gap 3.3 bits
... sigma38p10-proximalUP-distalUP 4437375 total 10.7 bits

   *4437350   .         *4437340   .         *4437330   .         *4437320   .
```

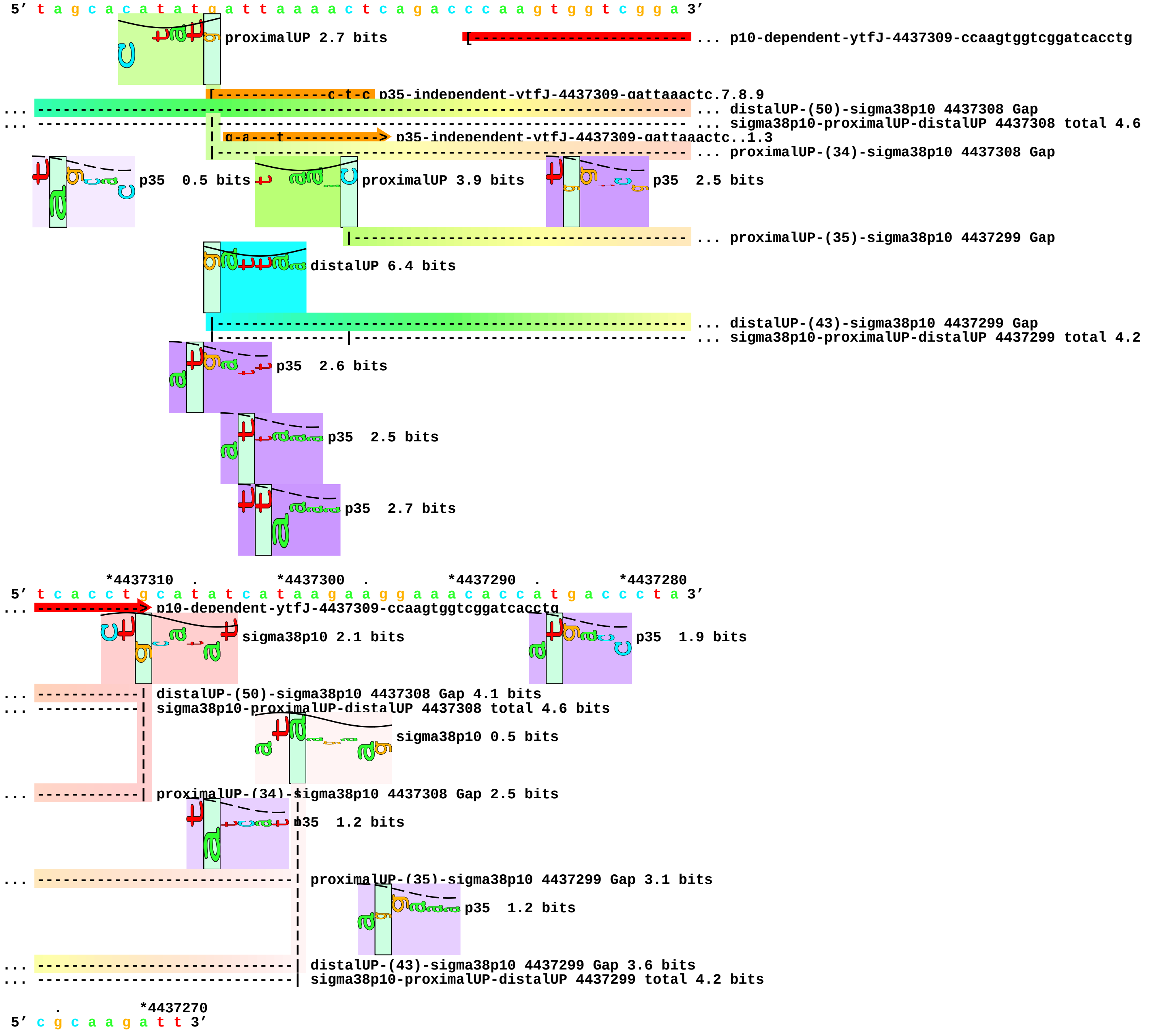

5’ t a g c a c a t a t g a t t a a a a c t c a g a c c c a a g t g g t c g g a 3’
proximalUP 2.7 bits
... p10-dependent-ytfJ-4437309-ccaagtggtcggatcacctg
p35-independent-ytfJ-4437309-gattaaactc.7.8.9
... distalUP-(50)-sigma38p10 4437308 Gap
... sigma38p10-proximalUP-distalUP 4437308 total 4.6
g-a---t p35-independent-ytfJ-4437309-gattaaactc..1.3
... proximalUP-(34)-sigma38p10 4437308 Gap
p35 0.5 bits
proximalUP 3.9 bits
p35 2.5 bits
... proximalUP-(35)-sigma38p10 4437299 Gap
distalUP 6.4 bits
... distalUP-(43)-sigma38p10 4437299 Gap
... sigma38p10-proximalUP-distalUP 4437299 total 4.2
p35 2.6 bits
p35 2.5 bits
p35 2.7 bits
*4437310 . *4437300 . *4437290 . *4437280
5’ t c a c c t g c a t a t c a t a a g a a g g a a a c a c c a t g a c c c t a 3’
p10-dependent-ytfJ-4437309-ccaagtggtcggatcacctg
sigma38p10 2.1 bits
p35 1.9 bits
distalUP-(50)-sigma38p10 4437308 Gap 4.1 bits
sigma38p10-proximalUP-distalUP 4437308 total 4.6 bits
sigma38p10 0.5 bits
proximalUP-(34)-sigma38p10 4437308 Gap 2.5 bits
p35 1.2 bits
proximalUP-(35)-sigma38p10 4437299 Gap 3.1 bits
p35 1.2 bits
distalUP-(43)-sigma38p10 4437299 Gap 3.6 bits
sigma38p10-proximalUP-distalUP 4437299 total 4.2 bits
. *4437270
5’ c g c a a g a t t 3’

piece 21, NC_000913, p10-independent-alaE-2796688-caagattatatattagccat, config: linear, direction: +, begin: 2796549, end: 2796709

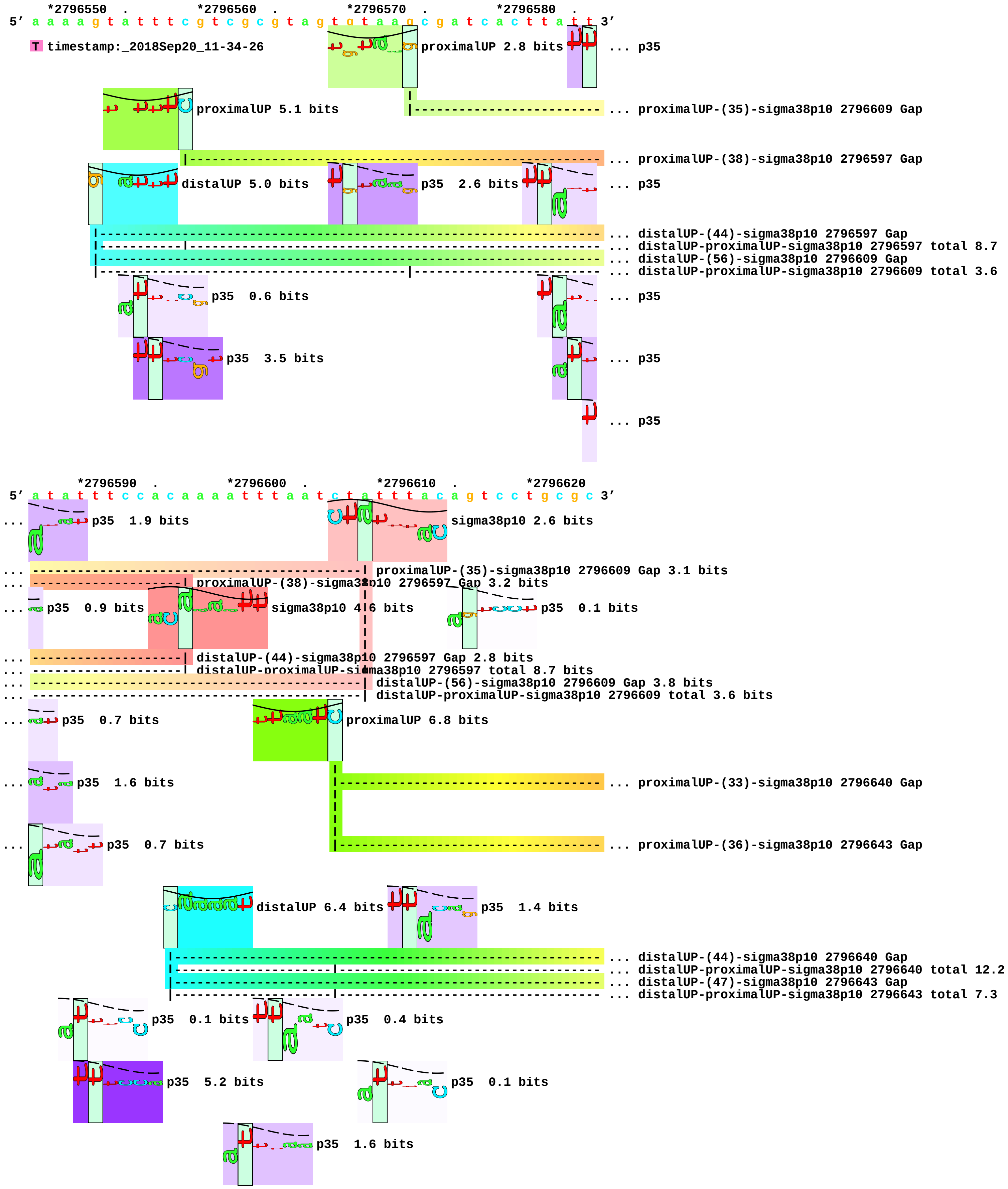

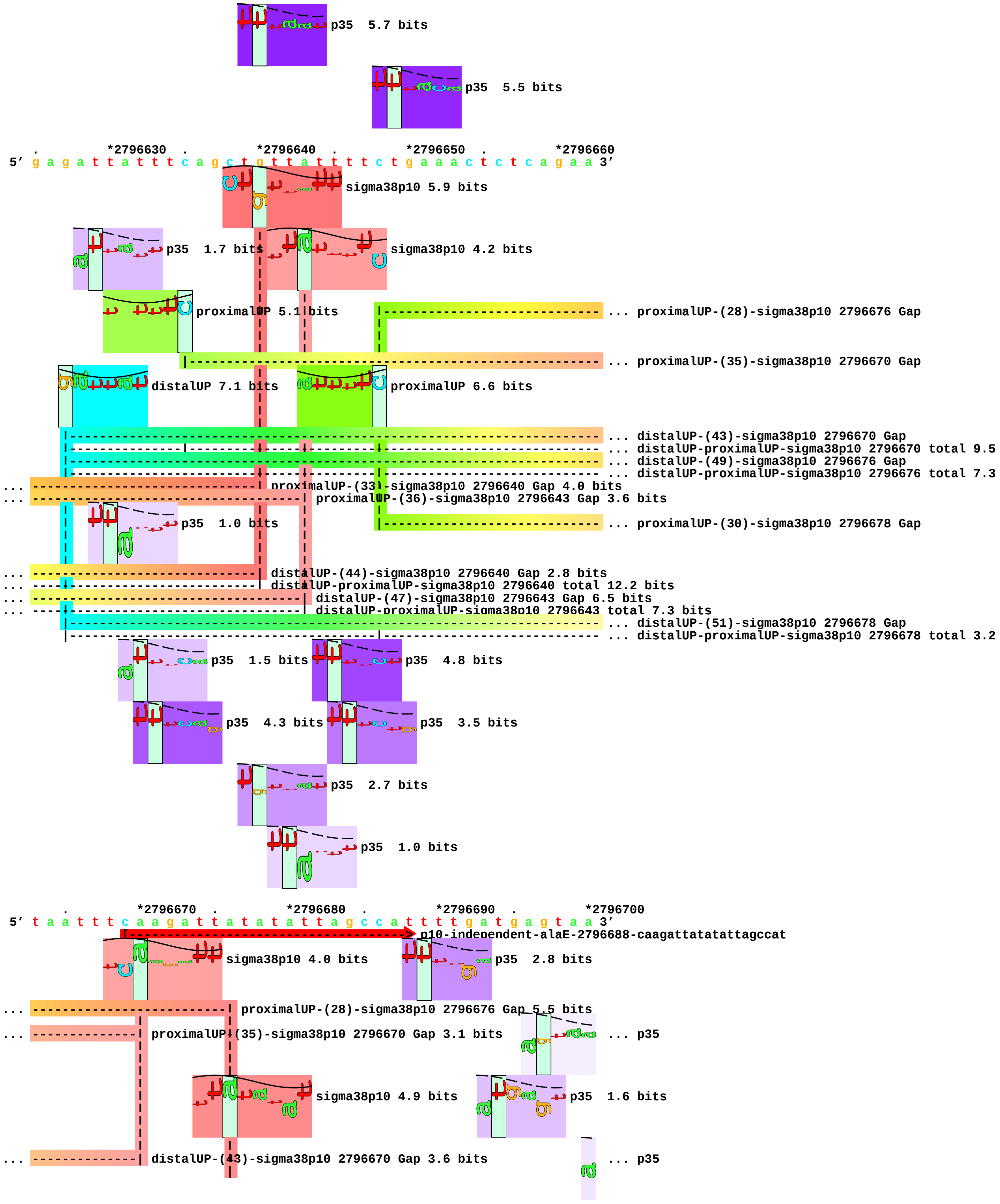
p35 5.7 bits
p35 5.5 bits
. *2796630 . *2796640 . *2796650 . *2796660
5' g a g a t t a t t t c a g c t g t t a t t t t c t g a a a c t c t c a g a a 3'
sigma38p10 5.9 bits
p35 1.7 bits
sigma38p10 4.2 bits
proximalUP 5.1 bits
... proximalUP-(28)-sigma38p10 2796676 Gap
... proximalUP-(35)-sigma38p10 2796670 Gap
distalUP 7.1 bits
proximalUP 6.6 bits
... distalUP-(43)-sigma38p10 2796670 Gap
... distalUP-proximalUP-sigma38p10 2796670 total 9.5
... distalUP-(49)-sigma38p10 2796676 Gap
... distalUP-proximalUP-sigma38p10 2796676 total 7.3
... proximalUP-(33)-sigma38p10 2796640 Gap 4.0 bits
... proximalUP-(36)-sigma38p10 2796643 Gap 3.6 bits
p35 1.0 bits
... proximalUP-(30)-sigma38p10 2796678 Gap
... distalUP-(44)-sigma38p10 2796640 Gap 2.8 bits
... distalUP-proximalUP-sigma38p10 2796640 total 12.2 bits
... distalUP-(47)-sigma38p10 2796643 Gap 6.5 bits
... distalUP-proximalUP-sigma38p10 2796643 total 7.3 bits
... distalUP-(51)-sigma38p10 2796678 Gap
... distalUP-proximalUP-sigma38p10 2796678 total 3.2
p35 1.5 bits
p35 4.8 bits
p35 4.3 bits
p35 3.5 bits
p35 2.7 bits
p35 1.0 bits
. *2796670 . *2796680 . *2796690 . *2796700
5' t a a t t t c a a g a t t a t a t a t t a g c c a t t t t g a t g a g t a a 3'
p10-independent-alaE-2796688-caagattatatattagccat
sigma38p10 4.0 bits
p35 2.8 bits
... proximalUP-(28)-sigma38p10 2796676 Gap 5.5 bits
... proximalUP-(35)-sigma38p10 2796670 Gap 3.1 bits
... p35
sigma38p10 4.9 bits
p35 1.6 bits
... distalUP-(43)-sigma38p10 2796670 Gap 3.6 bits
... p35

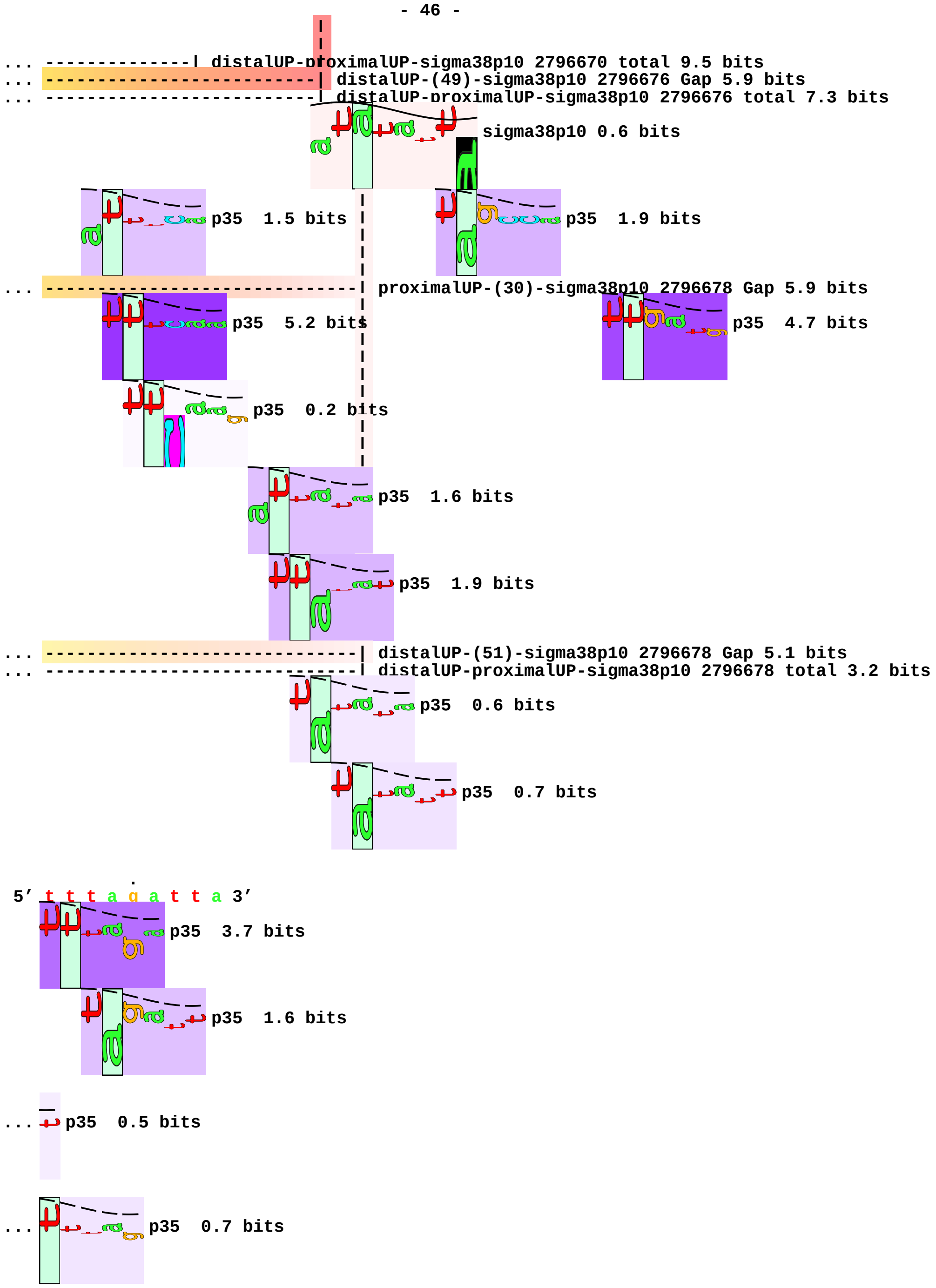
distalUP-proximalUP-sigma38p10 2796670 total 9.5 bits
distalUP-(49)-sigma38p10 2796676 Gap 5.9 bits
distalUP-proximalUP-sigma38p10 2796676 total 7.3 bits
sigma38p10 0.6 bits
p35 1.5 bits
p35 1.9 bits
proximalUP-(30)-sigma38p10 2796678 Gap 5.9 bits
p35 5.2 bits
p35 4.7 bits
p35 0.2 bits
p35 1.6 bits
p35 1.9 bits
distalUP-(51)-sigma38p10 2796678 Gap 5.1 bits
distalUP-proximalUP-sigma38p10 2796678 total 3.2 bits
p35 0.6 bits
p35 0.7 bits
5' t t t a g a t t a 3'
p35 3.7 bits
p35 1.6 bits
p35 0.5 bits
p35 0.7 bits

piece 22, NC_000913, p10-independent-lpp-1755407-actttgtgtaatacttgtaa, config: linear, direction: +, begin: 1755266, end: 1755426

```
            *1755270  .         *1755280  .         *1755290  .         *1755300
5' c g a t g g a a g c a t c c t g t t t t c t c t c a a t t t t t t t a t c t 3'
```

T timestamp:_2018Sep20_11-34-26 p35 4.8 bits ... proximalUP

p35 3.4 bits distalUP 5.8 bits

... distalUP-(51)-sigma38p10 1755341 Gap
... distalUP-proximalUP-sigma38p10 1755341 total 2.5

p35 1.8 bits p35 3.9 bits

p35 2.9 bits p35 4.8 bits

p35 0.3 bits p35 3.3 bits

p35 0.8 bits

p35 3.9 bits

p35 3.8 bits

p35 1.9 bits

... p35

```
   .         *1755310  .         *1755320  .         *1755330  .         *1755340
5' a a a a c c c a g c g t t c g a t g c t t c t t t g a g c g a a c g a t c a 3'
```

... proximalUP 3.9 bits p35 2.2 bits ... sigma38p10

proximalUP-(33)-sigma38p10 1755341 Gap 4.0 bits
... distalUP-(51)-sigma38p10 1755341 Gap 5.1 bits
... distalUP-proximalUP-sigma38p10 1755341 total 2.5 bits

p35 3.2 bits ... distalUP

... distalUP-(44)-sigma38p10 1755384 Gap
... distalUP-proximalUP-sigma38p10 1755384 total 4.5

... p35 0.2 bits

```
        .         *1755350  .         *1755360  .         *1755370  .
5' a a a a t a a g t g c c t t c c c a t c a a a a a a a t a t t c t c a a c a 3'
```

... sigma38p10 2.0 bits [----------------> p35-independent-lpp-1755407-tattctcaac

proximalUP 3.8 bits p35 0.6 bits

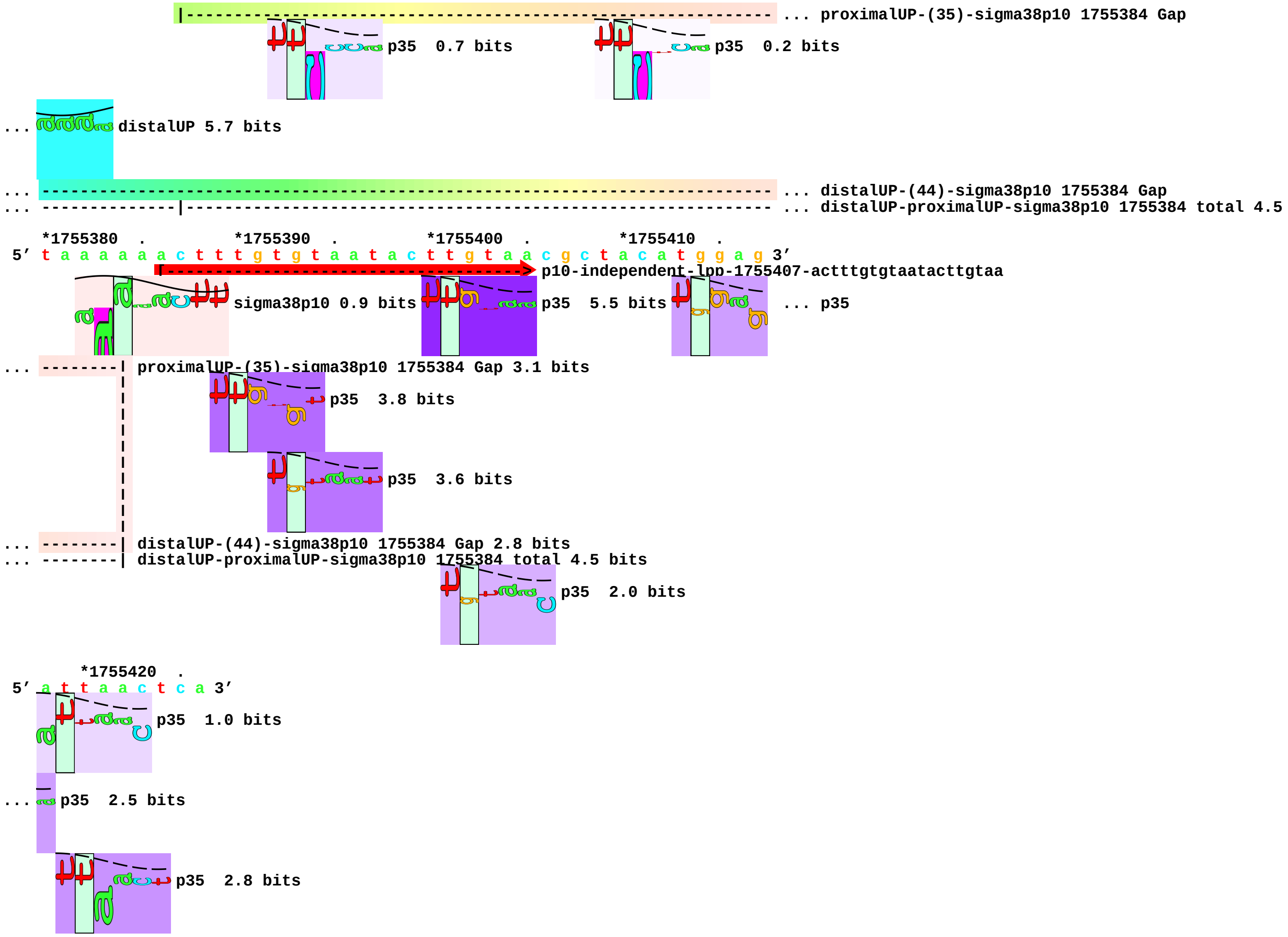
|------------------------------------------------------------ ... proximalUP-(35)-sigma38p10 1755384 Gap
p35 0.7 bits
p35 0.2 bits
... distalUP 5.7 bits
... ---------------------------------------------------------------------- ... distalUP-(44)-sigma38p10 1755384 Gap
... -------------|------------------------------------------------------- ... distalUP-proximalUP-sigma38p10 1755384 total 4.5
*1755380 . *1755390 . *1755400 . *1755410 .
5' t a a a a a a c t t t g t g t a a t a c t t g t a a c g c t a c a t g g a g 3'
p10-independent-lpp-1755407-actttgtgtaatacttgtaa
sigma38p10 0.9 bits
p35 5.5 bits
... p35
... --------| proximalUP-(35)-sigma38p10 1755384 Gap 3.1 bits
p35 3.8 bits
p35 3.6 bits
... --------| distalUP-(44)-sigma38p10 1755384 Gap 2.8 bits
... --------| distalUP-proximalUP-sigma38p10 1755384 total 4.5 bits
p35 2.0 bits
*1755420 .
5' a t t a a c t c a 3'
p35 1.0 bits
... p35 2.5 bits
p35 2.8 bits

piece 23, NC_000913, p10-independent-lpxC-106530-atgtatagtacacttcggtt, config: linear, direction: +, begin: 106390, end: 106550

*106390 . *106400 . *106410 . *106420 .
5' g c c g c a a a c t g c g a a a g a g c c g g a t t a t c t g g a t a t c c 3'

T timestamp:_2018Sep20_11-34-26

proximalUP 7.2 bits
... proximalUP-(37)-sigma38p10 106455 Gap
distalUP 7.1 bits
... distalUP-(43)-sigma38p10 106455 Gap
... distalUP-proximalUP-sigma38p10 106455 total 9.9
p35 0.2 bits
p35 1.9 bits
p35 3.5 bits

*106430 . *106440 . *106450 . *106460 .
5' c a g c a t t c c t g c g t a a g c a a g c t g a t t a a g a a t t g a c t 3'

p35 2.2 bits
sigma38p10 3.6 bits
... proximalUP-(37)-sigma38p10 106455 Gap 4.5 bits
proximalUP 4.9 bits
... distalUP-(43)-sigma38p10 106455 Gap 3.6 bits
... distalUP-proximalUP-sigma38p10 106455 total 9.9 bits
... proximalUP-(38)-sigma38p10 106495 Gap
distalUP 6.4 bits
... distalUP-(44)-sigma38p10 106495 Gap
... distalUP-proximalUP-sigma38p10 106495 total 13.1
p35 1.6 bits
... p35
p35 0.8 bits
p35 6.5 bits

*106470 . *106480 . *106490 . *106500
5' g g a a t t t g g g t t t c g a g g c t c t t t g t g c t a a a c t g g c c 3'

p35 3.4 bits
[------------------> p35-independent-lpxC-106530-gctaaactgg
sigma38p10 7.8 bits
p35 2.3 bits
... p35

- 50 -

... ----------------------------------------------------------| proximalUP-(38)-sigma38p10 106495 Gap 3.2 bits

... ----------------------------------------------------------| distalUP-(44)-sigma38p10 106495 Gap 2.8 bits

... ----------------------------------------------------------| distalUP-proximalUP-sigma38p10 106495 total 13.1 bits

... p35 4.4 bits

. *106510 . *106520 . *106530 . *106540

5’ c g c c g a a t g t a t a g t a c a c t t c g g t t g g a t a g g t a a t t 3’

p10-independent-lpxC-106530-atgtatagtacacttcggtt

p35 2.6 bits

p35 3.5 bits

p35 2.6 bits

p35 0.3 bits

... p35 2.5 bits

p35 0.0 bits

p35 0.3 bits

. *106550

5’ t g g c g a g a t 3’

p35 2.1 bits

piece 24, NC_000913, p10-independent-sdaA-1894833-aatcatcgcaatattagtta, config: linear, direction: +, begin: 1894692, end: 1894852

. *1894700 . *1894710 . *1894720 .
5' t g g t a c g a a c a g t a t t t t g t a t g g g g a a t g a c c g c a g g 3'
I timestamp:_2018Sep20_11-34-26
|------------------------------------------- ... proximalUP-(43)-sigma38p10 1894753 Gap
p35 2.0 bits
proximalUP 6.1 bits
... p35
distalUP 5.0 bits
p35 1.9 bits
|---------------------------------------------------------------- ... distalUP-(50)-sigma38p10 1894753 Gap
|-------------|-------------------------------------------------- ... distalUP-proximalUP-sigma38p10 1894753 total 3.0
p35 2.9 bits
p35 5.7 bits
p35 1.7 bits

*1894730 . *1894740 . *1894750 . *1894760 .
5' c a t a a t t c g t g a g c t g g c g c t g c a a a t t g g t g t g a a a c 3'
... ---------------------------------------------| proximalUP-(43)-sigma38p10 1894753 Gap 5.1 bits
... p35 1.0 bits
sigma38p10 1.2 bits
proximalUP 5.1 bits
proximalUP 5.0 bits
... ---------------------------------------------| distalUP-(50)-sigma38p10 1894753 Gap 4.1 bits
... ---------------------------------------------| distalUP-proximalUP-sigma38p10 1894753 total 3.0 bits
|---------------------------------------------------------- ... proximalUP-(38)-sigma38p10 1894775 Gap
|------------------- ... proximalUP-(27)-sigma38p10 1894785 Gap
distalUP 6.3 bits
|---------------------------------------------------------------------- ... distalUP-(45)-sigma38p10 1894775 Gap
|------------|-------------------------------------------------------- ... distalUP-proximalUP-sigma38p10 1894775 total 13.6
|-------------------------------------------------------- ... proximalUP-(40)-sigma38p10 1894777 Gap
|---------------------------------------------------------------------- ... distalUP-(47)-sigma38p10 1894777 Gap
|------------|-------------------------------------------------------- ... distalUP-proximalUP-sigma38p10 1894777 total 2.9
|-------------------------------------------------------- ... proximalUP-(43)-sigma38p10 1894780 Gap
|---------------------------------------------------------------------- ... distalUP-(50)-sigma38p10 1894780 Gap
|------------|-------------------------------------------------------- ... distalUP-proximalUP-sigma38p10 1894780 total 4.0
|---------------------------------------------------------------------- ... distalUP-(55)-sigma38p10 1894785 Gap
|-------------------------------------------------|-------------------- ... distalUP-proximalUP-sigma38p10 1894785 total 3.1
p35 0.4 bits
p35 1.7 bits
p35 0.7 bits

*1894770 . *1894780 . *1894790 . *1894800 .
5' c c t g a c t a t a c t t a t c t t t a c a t c t a c a a a a c a c t a c t 3'
sigma38p10 10.9 bits
[----------------> p35-independent-sdaA-1894833-aaaacactac
sigma38p10 2.0 bits
... p35
sigma38p10 1.9 bits
sigma38p10 0.4 bits

p35 1.5 bits

```
... ---------------| proximalUP-(38)-sigma38p10 1894775 Gap 3.2 bits
... -------------------------------------| proximalUP-(27)-sigma38p10 1894785 Gap 3.8 bits
... ---------------| distalUP-(45)-sigma38p10 1894775 Gap 5.5 bits
... ---------------| distalUP-proximalUP-sigma38p10 1894775 total 13.6 bits
... -------------------| proximalUP-(40)-sigma38p10 1894777 Gap 4.0 bits
... -------------------| distalUP-(47)-sigma38p10 1894777 Gap 6.5 bits
... -------------------| distalUP-proximalUP-sigma38p10 1894777 total 2.9 bits
... -------------------------| proximalUP-(43)-sigma38p10 1894780 Gap 5.1 bits
... -------------------------| distalUP-(50)-sigma38p10 1894780 Gap 4.1 bits
... -------------------------| distalUP-proximalUP-sigma38p10 1894780 total 4.0 bits
... -------------------------------------| distalUP-(55)-sigma38p10 1894785 Gap 4.9 bits
... -------------------------------------| distalUP-proximalUP-sigma38p10 1894785 total 3.1 bits
```

p35 1.9 bits

p35 0.4 bits

p35 5.5 bits

p35 2.4 bits

```
          *1894810   .         *1894820   .         *1894830   .         *1894840
5' t g a g a c a a t c a t c g c a a t a t t a g t t a a a t c g c g g t t t t 3'
              [---------------------------------------> p10-independent-sdaA-1894833-aatcatcgcaatattagtta
```

... p35 4.6 bits

p35 0.7 bits

... p35

p35 0.6 bits

... p35

p35 2.8 bits

... p35

```
        .        *1894850
5' t g a t t a g t t 3'
```

p35 0.7 bits

... p35 2.9 bits

... p35 2.8 bits

... p35 5.7 bits

piece 25, NC_000913, p10-independent-ssrA-2753608-aaatctggtatacttacctt, config: linear, direction: +, begin: 2753467, end: 2753627

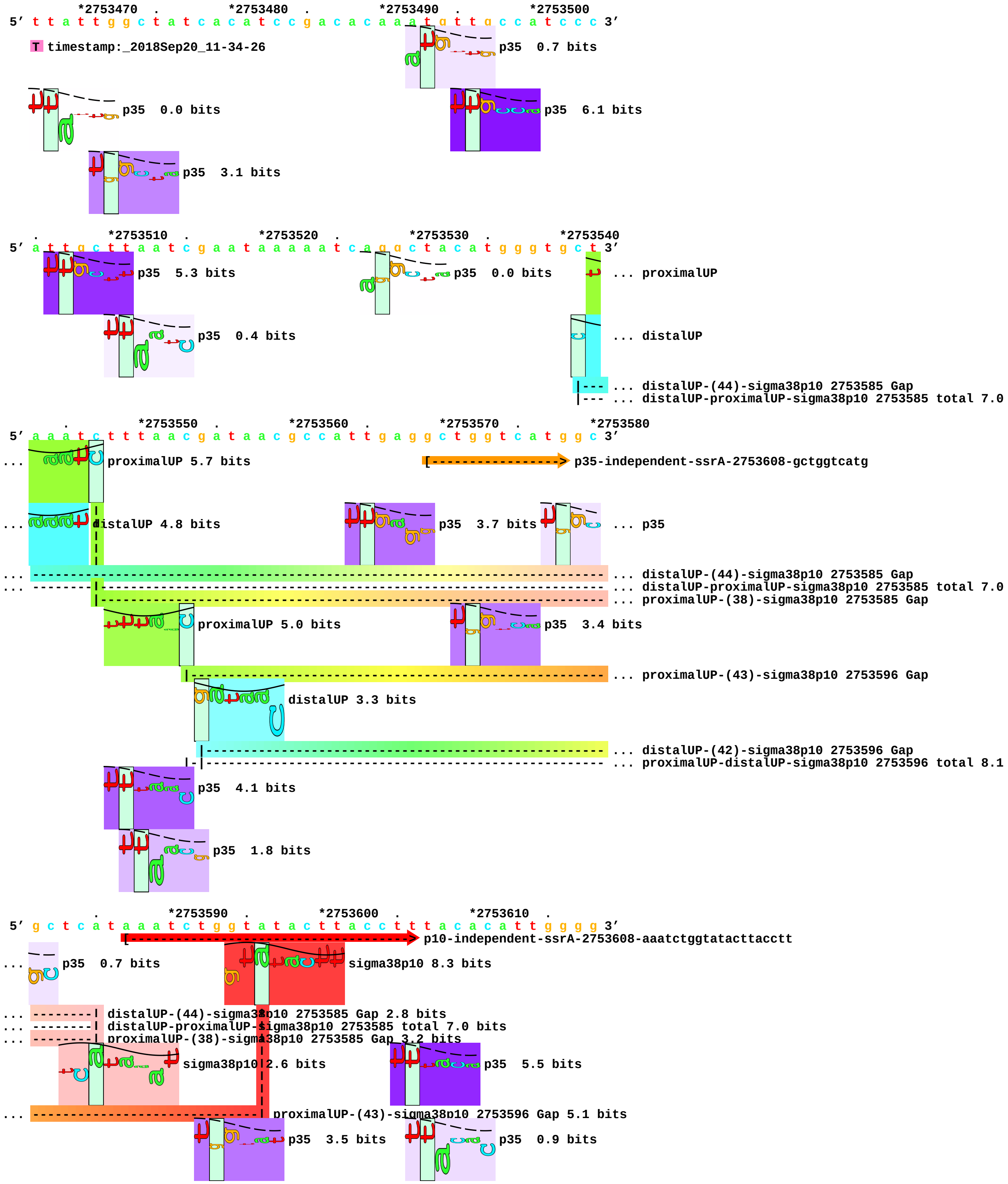

```
... ------------------------------| distalUP-(42)-sigma38p10 2753596 Gap 3.3 bits
... ------------------------------- proximalUP-distalUP-sigma38p10 2753596 total 8.1 bits
```

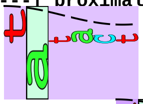

p35 1.5 bits

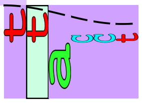

p35 2.4 bits

```
      *2753620  .
5′ c t g a t t c t g 3′
```

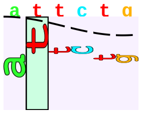

p35 0.4 bits

piece 26, NC_000913, p10-independent-uxaB-1608744-aagtttgctaacctgtcgta, config: linear, direction: -, begin: 1608868, end: 1608708

. *1608860 . *1608850 . *1608840 .

5' a g g c g t t t t t c c t c g a t t a a a c c a t g a t c c g c g c c a c a 3'

T timestamp:_2018Sep20_11-34-26

proximalUP 5.0 bits

proximalUP 6.2 bits ... proximalUP-(33)-sigma38p10 1608814 Gap

... proximalUP-(38)-sigma38p10 1608820 Gap

distalUP 4.9 bits ... proximalUP-(37)-sigma38p10 1608810 Gap

... distalUP-(44)-sigma38p10 1608820 Gap

... sigma38p10-proximalUP-distalUP 1608820 total 5.3

distalUP 6.4 bits

... distalUP-(40)-sigma38p10 1608814 Gap

... sigma38p10-proximalUP-distalUP 1608814 total 6.4

... distalUP-(44)-sigma38p10 1608810 Gap

... sigma38p10-proximalUP-distalUP 1608810 total 5.6

p35 2.4 bits ... proximalUP-(40)-sigma38p10 1608807 Gap

... distalUP-(47)-sigma38p10 1608807 Gap

... sigma38p10-proximalUP-distalUP 1608807 total 5.2

p35 3.2 bits

p35 1.1 bits

p35 5.3 bits

p35 2.5 bits

p35 1.2 bits

*1608830 . *1608820 . *1608810 . *1608800 .

5' c t t t t c t c a t a a a a c t g c t c a t t a a t c a t c a g g a g t t g 3'

sigma38p10 0.3 bits ... proximalUP-(36)-sigma38p10 1608768 Gap

... proximalUP-(33)-sigma38p10 1608814 Gap 4.0 bits

... proximalUP-(38)-sigma38p10 1608820 Gap 3.2 bits ... p35

... proximalUP-(37)-sigma38p10 1608810 Gap 4.5 bits

... distalUP-(44)-sigma38p10 1608820 Gap 2.8 bits

... sigma38p10-proximalUP-distalUP 1608820 total 5.3 bits

p35 4.8 bits

sigma38p10 2.0 bits

... distalUP-(40)-sigma38p10 1608814 Gap 3.1 bits

... sigma38p10-proximalUP-distalUP 1608814 total 6.4 bits

... distalUP-(44)-sigma38p10 1608810 Gap 2.8 bits

... sigma38p10-proximalUP-distalUP 1608810 total 5.6 bits

... proximalUP-(40)-sigma38p10 1608807 Gap 4.0 bits

... distalUP-(47)-sigma38p10 1608807 Gap 6.5 bits

... sigma38p10-proximalUP-distalUP 1608807 total 5.2 bits

p35 2.9 bits

sigma38p10 1.4 bits

p35 0.2 bits

sigma38p10 4.2 bits

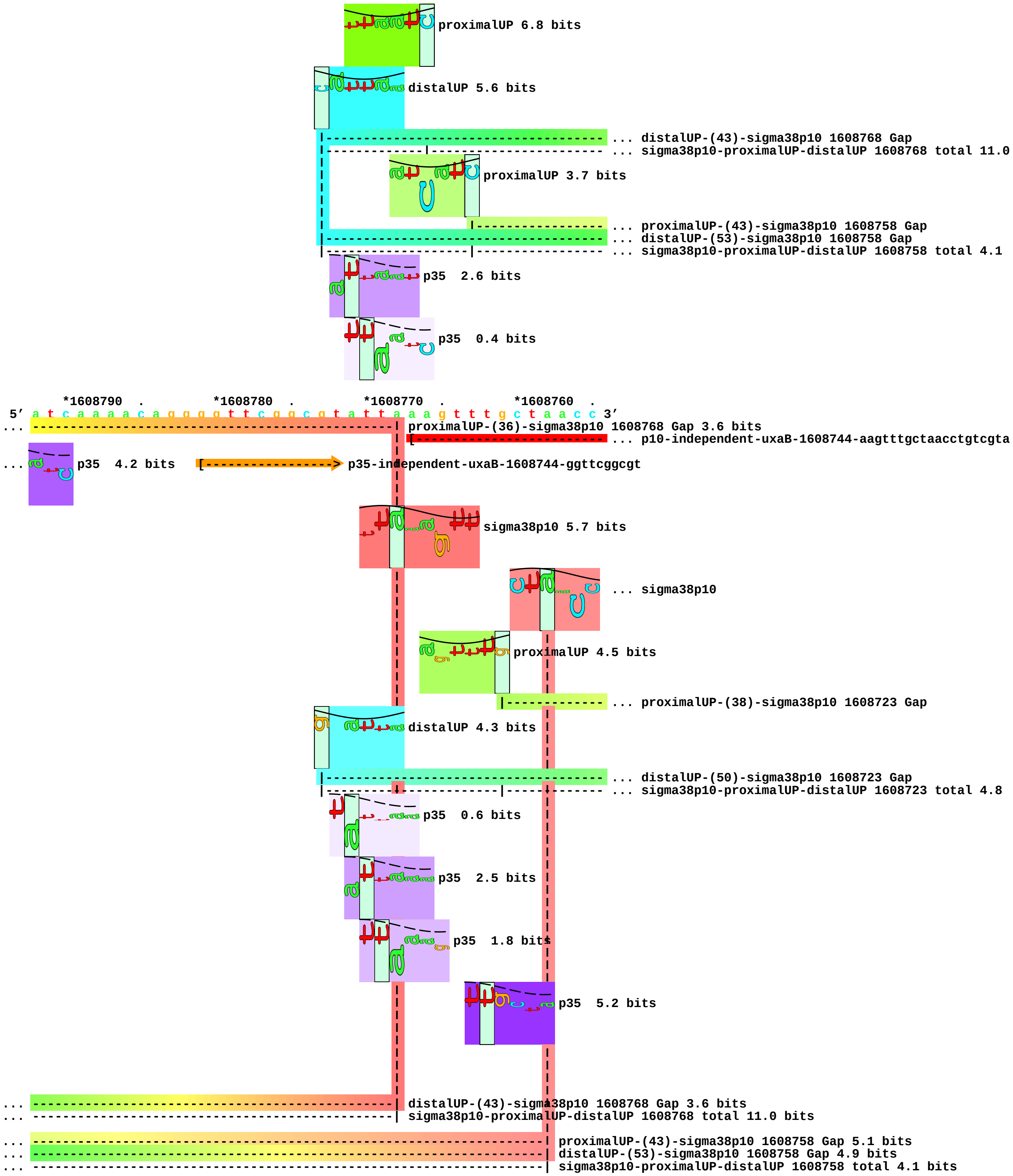
proximalUP 6.8 bits
distalUP 5.6 bits
... distalUP-(43)-sigma38p10 1608768 Gap
... sigma38p10-proximalUP-distalUP 1608768 total 11.0
proximalUP 3.7 bits
... proximalUP-(43)-sigma38p10 1608758 Gap
... distalUP-(53)-sigma38p10 1608758 Gap
... sigma38p10-proximalUP-distalUP 1608758 total 4.1
p35 2.6 bits
p35 0.4 bits
*1608790 . *1608780 . *1608770 . *1608760 .
5' a t c a a a a c a g g g g t t c g g c g t a t t a a a g t t t g c t a a c c 3'
... proximalUP-(36)-sigma38p10 1608768 Gap 3.6 bits
... p10-independent-uxaB-1608744-aagtttgctaacctgtcgta
... p35 4.2 bits
p35-independent-uxaB-1608744-ggttcggcgt
sigma38p10 5.7 bits
... sigma38p10
proximalUP 4.5 bits
... proximalUP-(38)-sigma38p10 1608723 Gap
distalUP 4.3 bits
... distalUP-(50)-sigma38p10 1608723 Gap
... sigma38p10-proximalUP-distalUP 1608723 total 4.8
p35 0.6 bits
p35 2.5 bits
p35 1.8 bits
p35 5.2 bits
... distalUP-(43)-sigma38p10 1608768 Gap 3.6 bits
... sigma38p10-proximalUP-distalUP 1608768 total 11.0 bits
... proximalUP-(43)-sigma38p10 1608758 Gap 5.1 bits
... distalUP-(53)-sigma38p10 1608758 Gap 4.9 bits
... sigma38p10-proximalUP-distalUP 1608758 total 4.1 bits

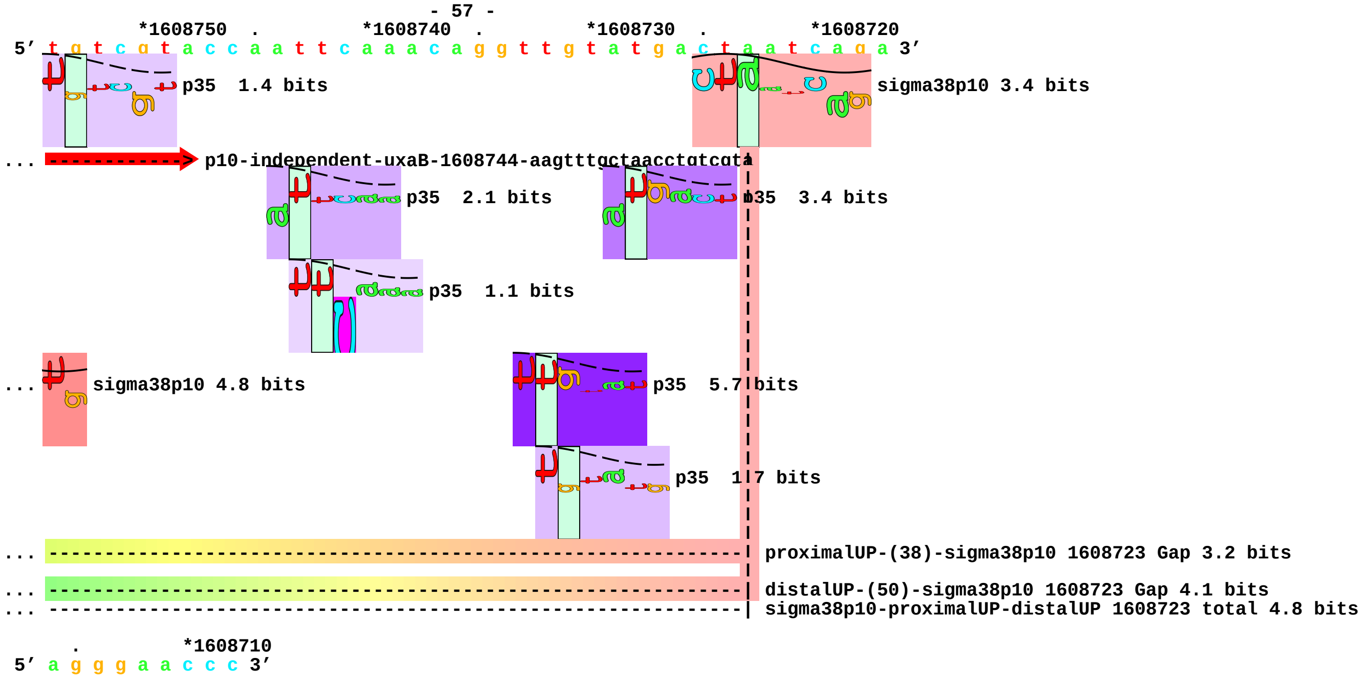
*1608750 . *1608740 . *1608730 . *1608720
5′ t g t c g t a c c a a t t c a a a c a g g t t g t a t g a c t a a t c a g a 3′
p35 1.4 bits
sigma38p10 3.4 bits
... p10-independent-uxaB-1608744-aagtttgctaacctgtcgta
p35 2.1 bits
p35 3.4 bits
p35 1.1 bits
... sigma38p10 4.8 bits
p35 5.7 bits
p35 1.7 bits
... proximalUP-(38)-sigma38p10 1608723 Gap 3.2 bits
... distalUP-(50)-sigma38p10 1608723 Gap 4.1 bits
... sigma38p10-proximalUP-distalUP 1608723 total 4.8 bits
. *1608710
5′ a g g g a a c c c 3′

piece 27, NC_000913, p10-independent-ygjR-3235304-aggcgtgctacattgacgac, config: linear, direction: +, begin: 3235164, end: 3235324

```
  .         *3235170  .         *3235180  .         *3235190  .         *3235200
5' c g c c a a a a t c c t c c a t c g c c a g c c g a a a t a t c t c g g g c 3'
T timestamp: 2018Sep20_11-34-26
proximalUP 6.1 bits
|------------------------------------------------------------ ... proximalUP-(37)-sigma38p10 3235210 Gap
distalUP 6.4 bits
|-------------------------------------------------------------------------- ... distalUP-(43)-sigma38p10 3235210 Gap
|-----------|-------------------------------------------------------------- ... distalUP-proximalUP-sigma38p10 3235210 total 9.0

  .         *3235210  .         *3235220  .         *3235230  .
5' c t t c g g t c a t g a t t g g g c c g t t t a g t g a g c a t g g c t g t 3'
sigma38p10 4.6 bits          p35  2.2 bits
p35  2.6 bits                ... p35
... ----------------| proximalUP-(37)-sigma38p10 3235210 Gap 4.5 bits
p35  3.8 bits
... ----------------| distalUP-(43)-sigma38p10 3235210 Gap 3.6 bits
... ----------------| distalUP-proximalUP-sigma38p10 3235210 total 9.0 bits

 *3235240  .         *3235250  .         *3235260  .         *3235270  .
5' c c g g c g c a g a a a t a a t g c g t a t c t g c g c a c g t c g a a g a 3'
proximalUP 6.7 bits     [----------------- ... p35-independent-ygjR-3235304-cgtcgaagat
... p35  2.2 bits
|----------------------------------------- ... proximalUP-(37)-sigma38p10 3235293 Gap
distalUP 5.9 bits       ... p35
|--------------------------------------------------------- ... distalUP-(45)-sigma38p10 3235293 Gap
|-----------------|---------------------------------------- ... distalUP-proximalUP-sigma38p10 3235293 total 8.8
p35  1.3 bits

     *3235280  .         *3235290  .         *3235300  .         *3235310  .
5' t g a a a a a g g c g t g c t a c a t t g a c g a c g a a t c c c t t t a 3'
... > p35-independent-ygjR-3235304-cgtcgaagat          ... p35
... -----------------------------| proximalUP-(37)-sigma38p10 3235293 Gap 4.5 bits
... p35  3.3 bits       sigma38p10 6.1 bits
... -----------------------------| distalUP-(45)-sigma38p10 3235293 Gap 5.5 bits
... -----------------------------| distalUP-proximalUP-sigma38p10 3235293 total 8.8 bits
[-------------------------> p10-independent-ygjR-3235304-aggcgtgctacattgacgac
p35  0.6 bits           p35  5.5 bits

       *3235320
5' t g g a g t a t c 3'
... p35  3.8 bits
```

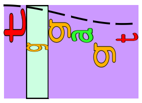

p35 2.6 bits

piece 28, NC_000913, p10-independent-yobF-1905641-gtatgtgataacagatttcg, config: linear, direction: -, begin: 1905761, end: 1905601

```
    *1905760  .         *1905750  .         *1905740  .         *1905730  .
5' g c g t a t t c c g t t g c a t a a g t g t g c a a a a a a a g t g g a a g 3'
```

T timestamp:_2018Sep20_11-34-26

proximalUP 3.8 bits

... proximalUP-(38)-sigma38p10 1905692 Gap

p35 1.2 bits

p35 6.2 bits

distalUP 5.7 bits

... distalUP-(46)-sigma38p10 1905692 Gap
... sigma38p10-proximalUP-distalUP 1905692 total 5.7
... distalUP-(52)-sigma38p10 1905686 Gap
... sigma38p10-proximalUP-distalUP 1905686 total 3.1

p35 3.4 bits

```
      *1905720  .         *1905710  .         *1905700  .         *1905690
5' a c g t a t c g a g a t t t g t g c g t c t g a t c g a g a c a t g t t t a 3'
```

proximalUP 2.6 bits

<---t-----c-----g-] p35-independent-yobF-1905641-gcctggaaca.7.4.1

... proximalUP-(38)-sigma38p10 1905692 Gap 3.2 bits

p35 2.3 bits

sigma38p10 3.5 bits

... distalUP-(46)-sigma38p10 1905692 Gap 4.1 bits
... sigma38p10-proximalUP-distalUP 1905692 total 5.7 bits
... distalUP-(52)-sigma38p10 1905686 Gap 5.5 bits
... sigma38p10-proximalUP-distalUP 1905686 total 3.1 bits

... sigma38p10

proximalUP-(31)-sigma38p10 1905686 Gap 4.7 bits

... distalUP

... distalUP-(42)-sigma38p10 1905648 Gap
... sigma38p10-proximalUP-distalUP 1905648 total 3.9

p35 1.7 bits

p35 1.7 bits

... p35

... p35

```
    .     *1905680  .         *1905670  .         *1905660  .         *1905650
5' a a a a t g g c t t g c c a t a a t t a a c g t t g t a t g t g a t a a c a 3'
```

proximalUP 5.6 bits

[--------------------------- ... p10-independent-yobF-1905641-gtatgtgataacagatttcg

proximalUP 5.4 bits

... sigma38p10

proximalUP-(32)-sigma38p10 1905648 Gap 4.3 bits

p35 3.2 bits

... proximalUP-(37)-sigma38p10 1905627 Gap

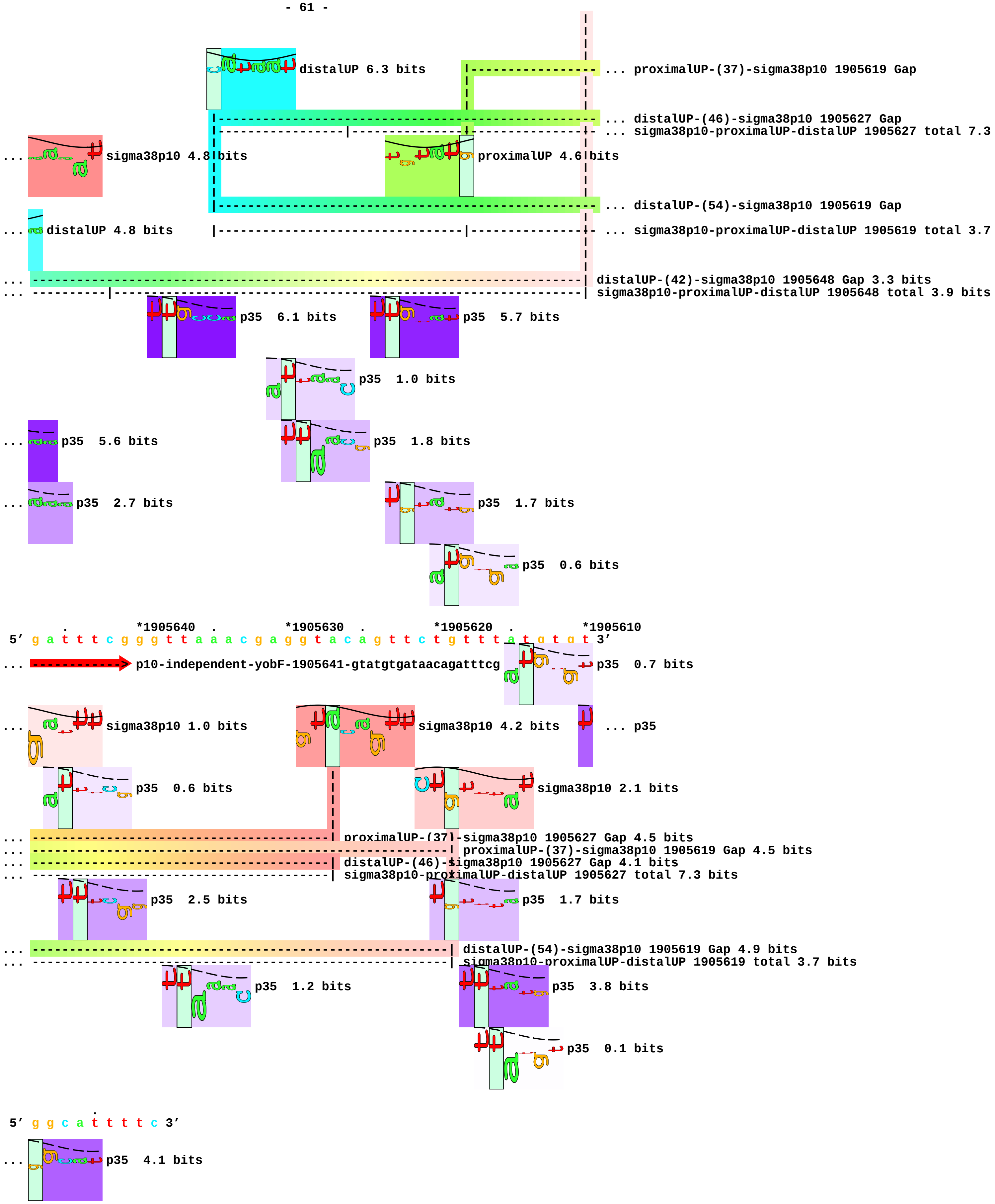
distalUP 6.3 bits
... proximalUP-(37)-sigma38p10 1905619 Gap
... distalUP-(46)-sigma38p10 1905627 Gap
... sigma38p10-proximalUP-distalUP 1905627 total 7.3
... sigma38p10 4.8 bits
proximalUP 4.6 bits
... distalUP-(54)-sigma38p10 1905619 Gap
... distalUP 4.8 bits
... sigma38p10-proximalUP-distalUP 1905619 total 3.7
...
...
distalUP-(42)-sigma38p10 1905648 Gap 3.3 bits
sigma38p10-proximalUP-distalUP 1905648 total 3.9 bits
p35 6.1 bits
p35 5.7 bits
p35 1.0 bits
... p35 5.6 bits
p35 1.8 bits
... p35 2.7 bits
p35 1.7 bits
p35 0.6 bits
. *1905640 . *1905630 . *1905620 . *1905610
5' g a t t t c g g g t t a a a c g a g g t a c a g t t c t g t t t a t g t g t 3'
... p10-independent-yobF-1905641-gtatgtgataacagatttcg
p35 0.7 bits
... sigma38p10 1.0 bits
sigma38p10 4.2 bits
... p35
p35 0.6 bits
sigma38p10 2.1 bits
...
...
...
...
proximalUP-(37)-sigma38p10 1905627 Gap 4.5 bits
proximalUP-(37)-sigma38p10 1905619 Gap 4.5 bits
distalUP-(46)-sigma38p10 1905627 Gap 4.1 bits
sigma38p10-proximalUP-distalUP 1905627 total 7.3 bits
p35 2.5 bits
p35 1.7 bits
...
...
distalUP-(54)-sigma38p10 1905619 Gap 4.9 bits
sigma38p10-proximalUP-distalUP 1905619 total 3.7 bits
p35 1.2 bits
p35 3.8 bits
p35 0.1 bits
.
5' g g c a t t t t c 3'
... p35 4.1 bits
