## Supplement_for_sigma38_paper-directory for "*Escherichia coli σ*^38^ promoters use two UP elements instead of a −35 element: resolution of a paradox and discovery that *σ*^38^ transcribes ribosomal promoters": README.pdf

### README: Supplementary Material for *Escherichia coli* $\sigma^{38}$ promoters use two UP elements instead of a −35 element: resolution of a paradox and discovery that $\sigma^{38}$ transcribes ribosomal promoters

Kevin S. Franco, Zhe Sun, Yixiong Chen, Cedric Cagliero, Yuhong Zuo, Yan Ning Zhou, Mikhail Kashlev, Ding Jun Jin, and Thomas D. Schneider *

version = 1.50 of README.tex 2020 Jan 08

**Glossary of information theory terms:**
`http://alum.mit.edu/www/toms/glossaryframes.html`

1. **README.pdf** contains the name and description of every document within the supplement. When the figures are in the same directory as this document, one can click on the underlined blue text to directly pop up a figure.

2. **File names that have the form 'figure-##.pdf' are the figures from the paper. The legends are in the main text.** We provide these copies of the main figures so that readers can examine their fine details.

| | | | |
|---|---|---|---|
| figure-01.pdf | figure-02.pdf | figure-03.pdf | figure-04.pdf |
| figure-05.pdf | figure-06.pdf | figure-07.pdf | figure-08.pdf |
| figure-09.pdf | figure-10.pdf | figure-11.pdf | figure-12.pdf |
| figure-13.pdf | figure-14.pdf | figure-15.pdf | figure-16.pdf |
| figure-17.pdf | figure-18.pdf | figure-19.pdf | figure-20.pdf |

3. **sigma38coverfigure.pdf** contains our proposed cover figure along with a description of the figure.

4. **sigma38cover.pdf** contains our proposed cover figure without a description.

5. **sigma38cover.eps** contains the eps version of our cover figure.

*National Institutes of Health, National Cancer Institute, Center for Cancer Research, RNA Biology Laboratory, P. O. Box B, Frederick, MD 21702-1201. (301) 846-5581, `http://alum.mit.edu/www/toms/`

6. **169sigma38map.pdf** contains the **lister** map (output of the **lister** program) of the $\sigma^{38}$ model scanned on the known 169 $\sigma^{38}$ genes and shown as sequence walkers. A sequence walker is a graphical representation of a single binding site, with the height of each letter indicating the information content at that position [1, 2]. The colored rectangles 'petals' with sequence walkers inside indicate the kind of site by hue and the strength of the binding site is distinguished by the saturation. The label to the right of each rectangle gives the kind of site and the bits of information of that site. The multi-part flexible sequence walker visualizes the locations of the -10, proximal, and distal UP elements that $\sigma^{38}$ binds to. To indicate the relationship there is a connecting bar that shades from the color of one site petal to the color of the other. The dashed line within the colored bridges has a label to the right that names the two kinds of site, the distance between the two and the gap surprisal for the distance between them. The dashed line labeled 'total' gives the kind of sites, the coordinate of the farthest downstream site, and the total information for the entire site found by the flexible model. Red petal: -10 region, green: proximal UP element region and blue: distal UP element region. The left edge of dark blue rectangles represent the $-10$ locations that were chosen to make the $\sigma^{38}$ model. Transcription starts 6 to 18 bp downstream from the blunt end of the arrow. The pink arrows represent the 78 experimentally proven transcription sites from regulonDB.

   As we were completing our analyses we noticed that two reported promoters, artPp2 and artPp3 are at the same location (see supplement 169sigma38map.pdf; the pages 36 and 37 are identical). The two promoters were found to have the same coordinate in RegulonDB database (Release: 8.2 Date: April-22-2013). Since that time a new dataset has been released ((Release: 9.2 Date: 09-08-2016) and the duplicate coordinate has been fixed. Ideally we would remove one of these, but the base frequencies would change only a small amount, less than 1%, so we do not anticipate that this would alter any of our results significantly. Furthermore, as discussed in the text (see supplement new22map.pdf), there are already 22 new $\sigma^{38}$ sites that could be incorporated into an even more complete model, but adding them is outside the scope of this work.

   In addition, yqaE and ygaU (page 105 of 169sigma38map.pdf) were reported in the RegulonDB database (Release: 8.2 Date: April-22-2013) and Weber *et al.* [3] to have the same initiation base. However in this case our models clearly identify two promoters, so each of these was used separately.

7. **sigma38.78.-10.inst.txt** is the instruction file required by the **delila** program to extract the sequences for the **lister** map. The instructions are for *E. coli* K-12 MG1655 in GenBank Accession NC000913 version 2 and are aligned by the base representing the $-10$ of the 78 $\sigma^{38}$ genes we used to make the first $\sigma^{38}$ model. These 78 $\sigma^{38}$ sites are from RegulonDB dataset: 8.2 Date: April-22-2013. 09-08-2016.

8. **sigma38.78.-10.book.txt** **Delila** was used to extract the sequences from sigma38.78.-10.inst.txt to create this collection of sequences (a 'book' of sequences).

9. **sigma38.169.-10.inst.txt** is the instruction file required by the **delila** program to extract the sequences for the **lister** map. The instructions are for *E. coli* K-12 MG1655 in GenBank

Accession NC000913 version 2 and are aligned by the base representing the -10 of the 169 $\sigma^{38}$ genes.

10. **sigma38.169.-10.book.txt** **Delila** was used to extract the sequences from sigma38.169.-10.inst.txt to create this collection of sequences (a 'book' of sequences).

11. **new22map.pdf** contains the **lister** map (output of the **lister** program) of the $\sigma^{38}$ model scanned on a new version of the RegulonDB database (Release: 9.2 Date: 09-08-2016) and shown as sequence walkers. The $\sigma^{38}$ model located 22 of the 35 new sequences not in our 169 dataset.

12. **File names that begin with 'sigma38.169.' make up the $\sigma^{38}$ model:**

    (a) **sigma38.169.ribl.txt** is a concatenated file containing the individual information weight matrices (ribls) of all elements used in the scan. The ribl weight matrix is constructed using individual information theory and is a model of a binding site. The order of these ribls matches the order of elements in the scanp file and the order of elements in the multiscanp file.

    (b) **sigma38.169.scanp.txt** is a concatenated file containing the **scan** program parameter files (scanp) for all of the elements.

    (c) **sigma38.169.multiscanp.txt** is the parameter file for the **multiscan** program.

    (d) **sigma38.169.histog.txt** concatenated histog files for each of the $\sigma^{38}$ elements. A histog file is a histogram of the distances between the sites. These files need to be in the same order as the multiscanp, scanp, and ribl files. Histog files can be generated by the program **genhis**.

13. **File names that begin with 'sigma38.169.[part]' and end with '.inst.txt' are the instruction files used by the delila program to generate the individual parts of the $\sigma^{38}$ model and the $\sigma^{38}$ combined UP element model:**

    (a) **sigma38.169.-10.inst.txt** is the instruction file used by the **delila** program [4]. The instructions are aligned by the -10 base of the 169 $\sigma^{38}$ genes. This file is the same as 169.inst.txt.

    (b) **sigma38.169.proximalUP.inst.txt** is the instruction file used by the **delila** program. The instructions are aligned by the base representing the proximal UP element of the 169 $\sigma^{38}$ genes.

    (c) **sigma38.169.distalUP.inst.txt** is the instruction file used by the **delila** program. The instructions are aligned by the base representing the distal UP element of the 169 $\sigma^{38}$ genes.

    (d) **sigma38.169.combinedUP.instshiftp.txt** is the parameter file that is used by the **instshift** program in order to shift an inst file to a certain base and range. In this case the parameter file creates the complementary **delila** instruction file. This is done by starting the third line with the letter N. This parameter file is used on the distal instructions to allow for the complementary instructions to align with the proximal instructions in order to make the combined $\sigma^{38}$ instructions.

(e) **sigma38.169.combinedUP.inst.txt** is the instruction file required by the **delila** program. The instructions are aligned by the base representing the proximal element of the 169 $\sigma^{38}$ genes and by the base of the distal element, in the opposite orientation, of the 169 $\sigma^{38}$ genes. These instructions are used in order to make a model of combined UP elements.

14. **rrnBp1-bolAp1-mutation-map.pdf contains the lister map of the *in vitro* constructs, analyzed by our $\sigma^{38}$ model.**

(a) The first page contains sequence walkers for the *rrnB* P1 promoter constructs. rrnB _P1_WT, is the first sequence shown (piece 1). The wild-type sequence is marked as a dashed line labeled rrnBP1_WT. The pink boxes on the ends of each sequence show the wild type sequence before it was replaced by EcoRI and HindIII sites in all plasmid constructs. In subsequent sequences (pieces 2-5) the green boxes on the sequence ends show the EcoRI and HindIII sites. The green boxes in the middle of sequences highlight the mutant sequences. The −10 of the $\sigma^{38}$ model is shown by a chartreuse petal, the proximal UP is shown in aquamarine and the distal is shown in blue. Fis sites are shown as purple petals. The −35 of *rrnB* P1 has a red petal. As described in our paper the most distal Fis site drops from 5.3 to 3.2 bits because the 5′ most base was changed from an A to a C by the creation of the EcoRI site. This analysis predicts:

piece 1: wild type sequence in *E. coli*
piece 2: wild type sequence in plasmid construct
piece 3: −10 mutation blocks $\sigma^{38}$
piece 4: UP region mutation blocks $\sigma^{38}$
piece 5: −35 mutation allows $\sigma^{38}$

These predictions match those shown in the Figure 18.

(b) The second page contains the sequence walkers for the *bolA* P1 promoter constructs. This analysis predicts:

piece 6: wild type sequence in *E. coli*
piece 7: wild type sequence in plasmid construct
piece 8: −10 mutation blocks $\sigma^{38}$
piece 9: UP region mutation blocks $\sigma^{38}$
piece 10: −35 mutation allows $\sigma^{38}$

These predictions match those shown in the Figure 19 except for the −35 case.

15. **File names that end with '.csv' are tables that show the sequences used for the *in vitro* experiments.**

(a) **Supplementary_Table_S1.csv** contains the promoter sequences used.

(b) **Supplementary_Table_S2.csv** contains the primers to construct the plasmids.

16. **File names that end with '.tif' are the original images of the *in vitro* gel experiments. For details see the corresponding figure legends in the main text.**

(a) **EXPTsigma38rrnBP1.tif - Figure 17**

(b) **EXPTsigma38rrnBP1Mutants.tif - Figure 18**

(c) **EXPTsigma38bolAP1Mutants.tif - Figure 19**

17. **The following papers were analyzed using the $\sigma^{38}$ model constructed from 169 sites.**

(a) **"A third recognition element in bacterial promoters: DNA binding by the $\alpha$ subunit of RNA polymerase"**
**Ross *et al.* [5]**

i. **Ross.Gourse1993-Fig7.pdf** Ross *et al.* showed that the UP element comprises a third area of recognition for the promoter. Through mutation and hydroxyl radical footprinting experiments they were able to locate the areas of protection upstream of the −35 in the −60 to −40 region. Their findings show that the UP-element increases promoter activity in the following promoters: *rrnB* P1, *rrnB* P2, *leuV*, and RNA-II. We scanned the sequences with our combined model of the UP element.

A. *rrnB* P1 is known to have an UP element located between −60 and −40 and we found two strong sites. One 7.0 bit site is from −58 to −53 and a 6.1 bit site is from −45 to −40. This promoter is well characterized and is described in our paper.

B. *rrnB* P2 is found to have its promoter activity significantly increased when the sequence extends to −68 rather than when it only extends to −39. Our model supports this finding because we find two strong UP elements located from −50 to −39.

C. RNA-II transcription was found to be dependent on the $\alpha$CTD and our model was able to locate two sites: a 3.0 bit site from −56 to −51 and a 2.6 bit site from −42 to −37.

D. *leuV* is also dependent on the $\alpha$CTD and it is found to have regions upstream from the −35 that increase its transcription. Sequences between −47 and −37 are known to increase activity by a factor of 3 *in vivo* and our model matches these findings because it found strong sites on *leuV* between −48 and −40.

E. They also find that the following promoters and mutations do not depend on the UP-element and therefore have less promoter acitivity: *lacUV5*, RNA-I, P1 ("SUB"), P1 (−41*A*), P1 (−41*B*) and P2 (−39). When our model was scanned on these sequences, it only found two weak sites on *lacUV5* and none on the rest of the sequences.

ii. These results show that our combined UP element model correctly predicts the results of Ross *et al.*

(b) **"Identification of an UP element consensus sequence for bacterial promoters"**
**Estrem *et al.* [6]**

i. **Estrem.Gourse1998-logo.pdf** (constructed from their Figure 2) Estrem *et al.* randomized the region upstream of the natural *E. coli rrnB* P1 promoter and selected sequences that had strong transcription [6]. The corresponding sequence logo has

23.7 ± 0.3 bits which is more than the information required to locate a single site in the *E. coli* genome ($\log_2(2 \times 4639675) = 23.1$ bits) and therefore it is unreasonably large [7].

ii. **Estrem.Gourse1998-rrnB-P1-selection.pdf** (walkers constructed from their Figure 2A; the randomized sequence is marked by a line followed by the 'Upstream Sequence Name' and the 'Relative Activity') We analyzed the individual mutant sequences with our 169 site $\sigma^{38}$ model and found that most of the conservation observed in the logo (−7 to +3) is explained by UP elements at various positions. We do not know why +7 to +9 is conserved as our model does not provide for proximal UP elements in this region, but it may prevent binding of the $\alpha$CTD in ways that reduce transcription. Some of the distal UP elements are predicted to use a C 5′ to the randomized region. This analysis using our two-UP element model suggests that the interpretation of the original randomization experiment is complicated and that its representation as a sequence logo does not directly represent UP element binding since they are not rigidly aligned relative to the rest of the promoter.

iii. **Estrem.Gourse1998-rrnB-P1-selection.pdf** (See only the first page of this document.) Estrem *et al.* performed DNase I footprinting on the 4192 selected variant. (See their section "UP Element Sequences Protected by RNAP and by Purified $\alpha$" and their Figure 4 on their page 9764). We marked the protected (square) and enhanced (triangle) DNA cleavage bases on our walker map. These effects correspond to the zero base of both the distal and proximal UP elements predicted by our model.

(c) **"Bacterial promoter architecture: subsite structure of UP elements and interactions with the carboxy-terminal domain of the RNA polymerase $\alpha$ subunit" Estrem *et al.* [8]**

i. We scanned their Fig.1 sequences with our $\sigma^{70}$ model attached to the proximal and distal UP element models. This model was only able to locate 2 of the sequences. Our model was unable to pick up the proximal locations because the distance of the experimentally selected proximal UP elements were closer to the −10 than the distance our model allows. Our UP element model is based on natural sequences and therefore cannot locate UP elements that do not follow the characteristics of natural occurring $\sigma^{38}$ promoters.

**Estrem.Gourse1999-fig1-map.pdf** If we allowed our UP element model to be closer to the −10 it was able to locate the correct proximal UP element. Allowing a range closer to the −10 did not change the other results described in our paper and therefore we did not modify our $\sigma^{38}$ model. Our model did not locate a distal element on any of the Estrem *et al.* sequences because the distal region in this experiment had a "SUB" sequence that did not contain UP element type sequence. The selected proximal region (light blue rectangle) and the distal region with a "SUB" sequence that does not influence transcription (light red rectangle) are marked above the sequence. Our model also did not locate an UP element on the sequence that was randomized.

ii. **Estrem.Gourse1999-fig3-map.pdf** We also scanned our $\sigma^{70}$ UP model on the selected distal sequences from their Fig.3B and it was able to locate the distal regions on every sequence that was shown to have a distal UP element. The selected distal region (light blue rectangle) and the proximal region with a "SUB" sequence that does not influence transcription (light red rectangle) are marked above the sequence

iii. These results show that our $\sigma^{70}$ UP model correctly predicts the results of Estrem *et al.*

(d) **"Mechanism of specific recognition of the *aidB* promoter by $\sigma^S$-RNA polymerase"** **Lacour *et al.* [9]**

i. **Lacour.Landini2002-map.pdf** We scanned our $\sigma^{70}$ (with UP) and $\sigma^{38}$ models on the mutant sequences which Lacour *et al.* generated. In the natural sequence, we find a 7.5 bit $\sigma^{70}$ site and a 4.7 bit $\sigma^{38}$ site. The black circle at −40 (NC000913.2 coordinate 4412222) and the black triangle at −50 (NC000913.2 coordinate 4412232) represent DNaseI-hypersensitive bands found by Lacour *et al.* The hypersensitive −50 band is located in the middle of the proximal and distal sites located by our $\sigma^{38}$ model.

In the mutant sequence, created by deleting base −13 C (NC000913.2 coordinate 4412259; highlighted pink square on page 1 and cursor symbol on pg.2) $\sigma^{38}$ is unable to locate a site but $\sigma^{70}$ is unaffected, as Lacour *et al.* observed.

Note: Lacour *et al.* reported a G at coordinate 4412248 in the wild type *aidB*, but in GenBank NC000913.2 it is a C.

(e) **"Differential ability of $\sigma^S$ and $\sigma^{70}$ *Escherichia coli* to utilize promoters containing half or full UP-element sites"**
**Typas *et al.* [10]**

i. **Typas.Hengge2005-table1-map.pdf** Some of their primer sequences matched GenBank NC_000913.2 but were different in their tables, so to do the analysis we located and used the corresponding sequences in GenBank. Then we constructed sequence walker maps using their Table 1 and compared our predictions to the data in their Figure 1. Mutational changes by Typas *et al.* are indicated by black arrows. Dashed objects are primers or −10 regions.

A. *rrnB* P1 (our page 1). Their sequence differs from GenBank NC_000913.2 at bases 3939501 and 3939502 where they report a TC, but Genbank reports a CA. Our model predicts that with either sequence variation there is a $\sigma^{38}$ site with an information content of 11.3 bits.

B. *bolA* P1 (our page 2, their figure 1A). Typas *et al.* show that the *bolA* promoter in rpoS+ cells gave low $\beta$-galactosidase activity for the No-UP element construct, slightly higher activity for the Wild-Type and the highest activity for the putative Full-UP construct.
- **Wild-Type (piece 3)**: A 6.4 bit $\sigma^{38}$ site was located.
- **No-UP (piece 4)**: No $\sigma^{38}$ site was located.
- **Full-UP (piece 5)**: A 6.4 bit $\sigma^{38}$ site was located.

The sites found by our model on the Wild-Type and No-UP element constructs match the Typas *et al.* data. Although we do predict a site for the Full-UP element construct, our model did not indicate the observed increase in activity. This was also not explained by Typas *et al.* They thought that for $\sigma^{38}$ binding there must be a −35 and a distal UP and that they were mutating only the distal UP element (No-UP, piece 4). However our model shows that those mutations ruin both the proximal and distal UP sites. Their supposed proximal mutations (piece 5) do not affect either UP element of our model. We suggest that the supposed Full-UP construct may have affected binding by another protein in this *in vivo* experiment. This is the only case of Typas *et al.* data that does not fit our $\sigma^{38}$ model.

C. *csiE* (our page 3, their figure 1C). This sequence has an inconsistency: at base 2663400 Genbank reports an A, but the Typas *et al.* sequence is missing this base. This makes a deletion in their sequence. The base is marked with a pink box on the Wild-Type sequence and with a light green box on the no-UP element and Full-UP element constructs.

   Typas *et al.* found that *csiE* promoter in rpoS+ cells gave very low $\beta$-galactosidase activity for the No-UP construct, slightly higher activity for the Wild-Type and much higher activity for the Full-UP construct.

   - **Wild-type (piece 6)**: A 2.6 bit $\sigma^{38}$ site at 2663412 was found with the −10 matching the −10 in Table 1. Downstream at 2663423 there is an unreported 3.5 bit $\sigma^{38}$ site with a strong −10 of 9.8 bits but weak UP elements. We propose that this site is non-functional because both UP elements show a G base with a black rectangular background. This indicates that G has never been seen before in the original sequences at those positions.
   - **No-UP (piece 7)**: No $\sigma^{38}$ site was located at 2663412 for the −10 in Table 1. The poor $\sigma^{38}$ site at 2663423 was not changed.
   - **Full-UP (piece 8)**: A $\sigma^{38}$ site with an information content of 2.6 bits was located at 2663412. Further downstream the site at 2663423 was found with an information content of 10.1 bits. We also found a site further downstream at 2663429 with an information content of 2.5 bits.

   In summary, the No-UP sequence destroyed the $\sigma^{38}$ site and low transcription was observed. The Wild-type *csiE* sequence (2.6 bits) had low transcription compared to *bolA* (6.4 bits). Our model predicted three sites (2.6, 10.1 and 2.5 bits) on the so-called Full-UP sequence, consistent with the strong observed $\sigma^{38}$ transcription. Therefore our $\sigma^{38}$ model fits these data.

D. *osmY* (our page 4, their figure 1E). Typas *et al.* showed that *osmY* in rpoS+ cells had significant $\beta$-galactosidase activity for the No-UP construct, slightly higher activity for the Wild-Type construct and much higher activity for the Full-UP construct.

   - **Wild-type (piece 9)**: A $\sigma^{38}$ site with an information content of 7.6 bits was found at the reported -10. Further upstream a 4.8 bit site with a different -10 was located. The two promoters share a proximal UP element.

- **No-UP (piece 10)**: The reported -10 no longer has a $\sigma^{38}$ site. However, the unreported $-10$ site found upstream on the Wild-Type sequence is found again, but with a lower information content of 4.0 bits. Although the 3.8 bit proximal UP site was destroyed by the mutations, a slightly weaker 3.2 bit proximal site was found upstream.
- **Full-UP (piece 11)**: The reported site is found again, and the information content remains the same as on Wild-Type. The unreported site found upstream remains and has the same 4.8 bits of information as it does on Wild-Type. A new 3.5 bit site, for which a strong distal UP was created by the mutations, is found further downstream.

The promoter variants for *osmY* are more than 10 times stronger than those of *bolA* and *csiE*, so even the putative No-UP construct has significant $\sigma^{38}$ transcription. Consistent with this, our $\sigma^{38}$ model located one site for the so-called No-UP construct, two sites for the Wild-Type sequence and three sites for the Full-UP construct. In summary, our model shows that the increase in $\beta$-galactosidase activity correlates to increasing numbers of predicted $\sigma^{38}$ sites.

ii. **Typas.Hengge2005-supplement-S1-map.pdf** We analyzed Supplementary Table S1 of Typas *et al.* (MMI_4382_sm_TableS1.doc) using our 169 site $\sigma^{38}$ model. First we mapped their sequences to the *E. coli* genome NC_000913.2, and then scanned them with the sigma38.169 model to produce the PDF.

The results are summarized in the text table Typas.Hengge2005-supplement-S1-map.txt. This table lists both their predictions and our predictions for the number of UP elements of $\sigma^{38}$ promoters.

Typas *et al.* predicted that of 51 $\sigma^{38}$ promoters only 3 would have two UP elements, 18 would have just one UP and 30 would have no UP elements. In contrast, out of the 51 promoters our model predicts 46 tripart $\sigma^{38}$ sites and therefore predicts at least two UP elements for each of these 46.

We believe this occurred because Typas *et al.* used a rigid consensus sequence to predict the UP element region. It has been shown that such models do not make good predictions [11].

(f) **"Selectivity of the *Escherichia coli* RNA polymerase E$\sigma^{38}$ for overlapping promoters and ability to support CRP activation"**
**Kolb *et al.* [12]**

i. **Kolb.Ishihama1995-fig2-map.pdf** Kolb *et al.* showed that both $\sigma^{38}$ and $\sigma^{70}$ are capable of transcribing the *gal*P1 promoter. We scanned our $\sigma^{38}$ and $\sigma^{70}$ (with UP) models on the *gal*P1 promoter sequence from Figure 2 of Kolb *et al.* This identified a 9.6 bit $\sigma^{38}$ site and a 5.6 bit $\sigma^{70}$ site. Kolb *et al.* found that $\sigma^{38}$ transcibes *gal*P1 4.7 fold more than $\sigma^{70}$ transcribes it, consistent with our analysis. In the figure the dashed lines represent the $-10$ locations for both $\sigma^{38}$ and $\sigma^{70}$ as marked by Kolb *et al.*

(g) **"Characterization of the *Escherichia coli* $\sigma^S$ core regulon by Chromatin Immunoprecipitation-sequencing (ChIP-seq) analysis"**

**Peano *et al.* [13]**

The tables were reproduced as machine-readable comma-separated-variable (csv) files here with permission from Stephan Lacour. Table 1 and 2 (**Peano.Landini2015-Table1.csv** and **Peano.Landini2015-Table2.csv**) were from updated versions graciously sent to us by Stephan Lacour. **Peano.Landini2015-Table34-list.csv** was created by combining Tables 3 and 4.

i. **Peano.Landini2015-Table1-pos-map.pdf** corresponds to their Table 1. The sequence range of each peak was extended 50 bases on each end to allow the $\sigma^{38}$ model to overlap the edges. The 169 site $\sigma^{38}$ model was scanned in the given orientation ('pos') so that the walkers point at the designated gene. The rear end of yellow arrows indicate the coordinate of the 'experimentally validated TSS located inside peak'. Small rectangles on the sequence indicate the ends of the defined chip peak; they are green if the region contains a validated peak and red if not. The coordinates of some experimentally validated peaks (*e.g. yaiA*) are not inside the defined peak ranges. If the peak is validated but the coordinate is not inside the given peak range, the peak edge rectangles are marked purple and there is no yellow arrow. The reversed ('neg') orientation scan **Peano.Landini2015-Table1-neg-map.pdf** has the model pointing away from the gene. The number of sites found for the two orientations was nearly identical.

ii. **Peano.Landini2015-Table2-pos-map.pdf** and **Peano.Landini2015-Table2-neg-map.pdf** correspond to their Table 2 in two orientations. The positive orientation means that the $\sigma^{38}$ models point towards the first gene listed; while the negative orientation means that the $\sigma^{38}$ models point towards the second gene listed.

iii. **Peano.Landini2015-Table34.pdf** are scans for their Supplementary Table S3 ('Sequence of experimentally determined −10 promoter regions lying within ChIP-seq peaks') and their Supplementary Table S4 ('Sequence of experimentally determined −35 promoter regions lying within ChIP-seq peaks'). Based on the sequences given in these tables, the proposed −10 regions are marked with red arrows and the proposed −35 regions by orange arrows. When a sequence could not be found in NC_000913.2 the number of allowed mismatches was increased until at least one was found. The base(s) that needed to be changed are indicated inside the arrow. For comparison, the −35 model from [14] is shown in purple.

Of the 104 sequences analyzed from the four tables (not including Peano.Landini2015-Table1-neg-map.pdf since that is a control), with 4 exceptions (insEF-2, omrA, osmE, sibC), we found that each sequence from Peano *et al.* has one or more predicted $\sigma^{38}$ site of at least 2 bits. In some cases, because of the broad regions defined by ChIP-seq, there are many predicted $\sigma^{38}$ sites. The results of these scans are often too complex to interpret in detail but we provide the analysis to help future investigations.

(h) **"Interaction of the C-terminal domain of the *E. coli* RNA polymerase $\alpha$ subunit with the UP element: recognizing the backbone structure in the minor groove surface"**
**Yasuno *et al.* [15] Yasuno.Kyogoku2001-figure1b.pdf**

### References


[1] Schneider,T.D. (1997) Sequence Walkers: a graphical method to display how binding proteins interact with DNA or RNA sequences. *Nucleic Acids Res.,* **25**, 4408–4415 https://doi.org/10.1093/nar/25.21.4408, https://alum.mit.edu/www/toms/papers/walker/, erratum: NAR 26(4): 1135, 1998.

[2] Schneider,T.D. and Spouge,J. (1997) Information content of individual genetic sequences. *J. Theor. Biol.,* **189**, 427–441 https://doi.org/10.1006/jtbi.1997.0540, https://alum.mit.edu/www/toms/papers/ri/.

[3] Weber,H., Polen,T., Heuveling,J., Wendisch,V.F., and Hengge,R. (2005) Genome-wide analysis of the general stress response network in *Escherichia coli*: $\sigma^S$-dependent genes, promoters, and sigma factor selectivity. *J. Bacteriol.,* **187**, 1591–1603 https://doi.org/10.1128/JB.187.5.1591-1603.2005.

[4] Schneider,T.D. and Stephens,R.M. (1990) Sequence Logos: A New Way to Display Consensus Sequences. *Nucleic Acids Res.,* **18**, 6097–6100 https://doi.org/10.1093/nar/18.20.6097, https://alum.mit.edu/www/toms/papers/logopaper/.

[5] Ross,W., Gosink,K.K., Salomon,J., Igarashi,K., Zou,C., Ishihama,A., Severinov,K., and Gourse,R.L. (1993) A third recognition element in bacterial promoters: DNA binding by the $\alpha$ subunit of RNA polymerase. *Science,* **262**, 1407–1413 https://doi.org/10.1126/science.8248780.

[6] Estrem,S.T., Gaal,T., Ross,W., and Gourse,R.L. (1998) Identification of an UP element consensus sequence for bacterial promoters. *Proc. Natl. Acad. Sci. USA,* **95**, 9761–9766 https://doi.org/10.1073/pnas.95.17.9761.

[7] Schneider,T.D., Stormo,G.D., Gold,L., and Ehrenfeucht,A. (1986) Information content of binding sites on nucleotide sequences. *J. Mol. Biol.,* **188**, 415–431 https://doi.org/10.1016/0022-2836(86)90165-8, https://alum.mit.edu/www/toms/papers/schneider1986/.

[8] Estrem,S.T., Ross,W., Gaal,T., Chen,Z.W., Niu,W., Ebright,R.H., and Gourse,R.L. (1999) Bacterial promoter architecture: subsite structure of UP elements and interactions with the carboxy-terminal domain of the RNA polymerase $\alpha$ subunit. *Genes Dev,* **13**, 2134–2147.

[9] Lacour,S., Kolb,A., Boris Zehnder,A.J., and Landini,P. (2002) Mechanism of specific recognition of the *aidB* promoter by $\sigma^S$-RNA polymerase. *Biochem. Biophys. Res. Commun.,* **292**, 922–930 https://doi.org/10.1006/bbrc.2002.6744.

[10] Typas,A. and Hengge,R. (2005) Differential ability of $\sigma^s$ and $\sigma^{70}$ of *Escherichia coli* to utilize promoters containing half or full UP-element sites. *Mol. Microbiol.,* **55**, 250–260 https://doi.org/10.1111/j.1365-2958.2004.04382.x.

[11] Schneider,T.D. (2002) Consensus Sequence Zen. *Applied Bioinformatics,* **1**, 111–119 https://www.ncbi.nlm.nih.gov/pmc/articles/PMC1852464/, https://alum.mit.edu/www/toms/papers/zen/.

[12] Kolb,A., Kotlarz,D., Kusano,S., and Ishihama,A. (1995) Selectivity of the *Escherichia coli* RNA polymerase E$\sigma^{38}$ for overlapping promoters and ability to support CRP activation. *Nucleic Acids Res.,* **23**, 819–826 https://doi.org/10.1093/nar/23.5.819.

[13] Peano,C., Wolf,J., Demol,J., Rossi,E., Petiti,L., De Bellis,G., Geiselmann,J., Egli,T., Lacour,S., and Landini,P. (2015) Characterization of the *Escherichia coli* $\sigma^S$ core regulon by Chromatin Immunoprecipitation-sequencing (ChIP-seq) analysis. *Sci Rep,* **5**, 10469 https://doi.org/10.1038/srep10469.

[14] Shultzaberger,R.K., Chen,Z., Lewis,K.A., and Schneider,T.D. (2007) Anatomy of *Escherichia coli* $\sigma^{70}$ promoters. *Nucleic Acids Res.,* **35**, 771–788 https://doi.org/10.1093/nar/gkl956, https://alum.mit.edu/www/toms/papers/flexprom/.

[15] Yasuno,K., Yamazaki,T., Tanaka,Y., Kodama,T.S., Matsugami,A., Katahira,M., Ishihama,A., and Kyogoku,Y. (2001) Interaction of the C-terminal domain of the *E. coli* RNA polymerase $\alpha$ subunit with the UP element: recognizing the backbone structure in the minor groove surface. *J. Mol. Biol.,* **306**, 213–225 https://doi.org/10.1006/jmbi.2000.4369.
