## Supplement_for_sigma38_paper-directory for "*Escherichia coli σ*^38^ promoters use two UP elements instead of a −35 element: resolution of a paradox and discovery that *σ*^38^ transcribes ribosomal promoters": Ross.Gourse1993-Fig7.pdf

rrnB(P1)

```
            *-60      .         *-50      .         *-40      .         *-30
5′ c g g t c a g a a a a t t a t t t t a a a t t t c c t c t t g t c a g g c c 3′
```

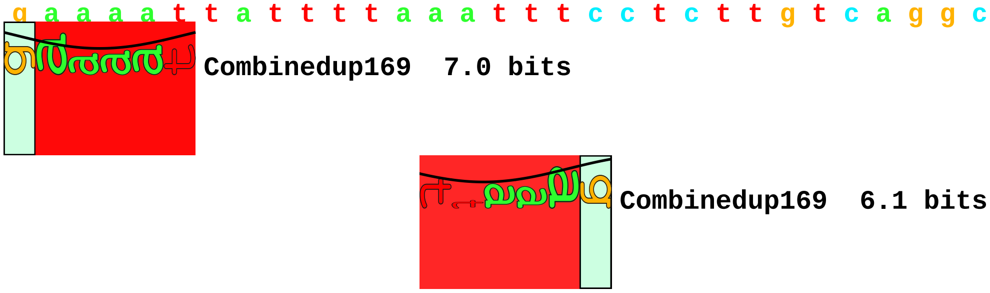


rrnB(P2)

```
            *-60      .         *-50      .         *-40      .         *-30
5′ c t c c g g c a g a g a a a g c a a a a a t a a a t g c t t g a c t c t g t 3′
```

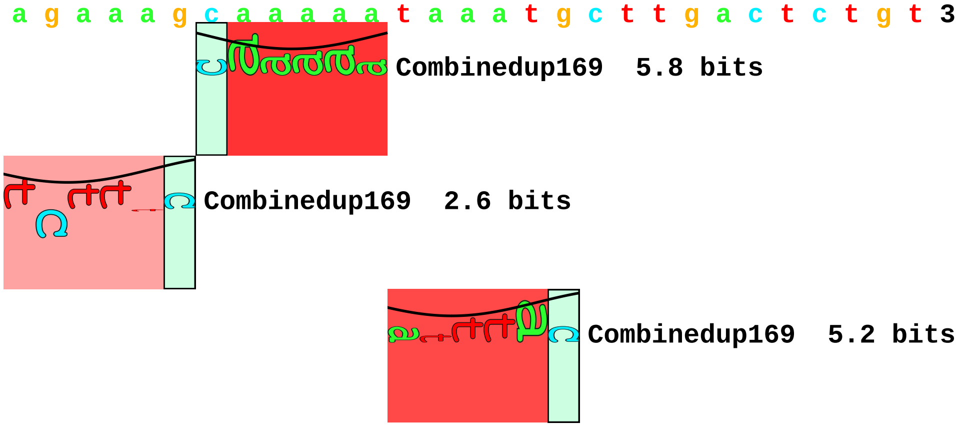


RNA-II

```
            *-60      .         *-50      .         *-40      .         *-30
5′ a c c c c g t a g a a a a g a t c a a a g g a t c t t c t t g a g a t c c t 3′
```

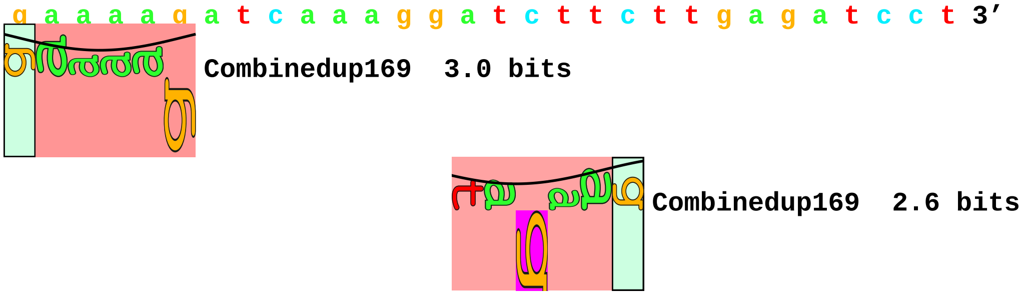


leuV

```
            *-60      .         *-50      .         *-40      .         *-30
5′ c a a t t a c a c c t c t g t c g a t a a t t a a c t a t t g a c g a a a a 3′
```

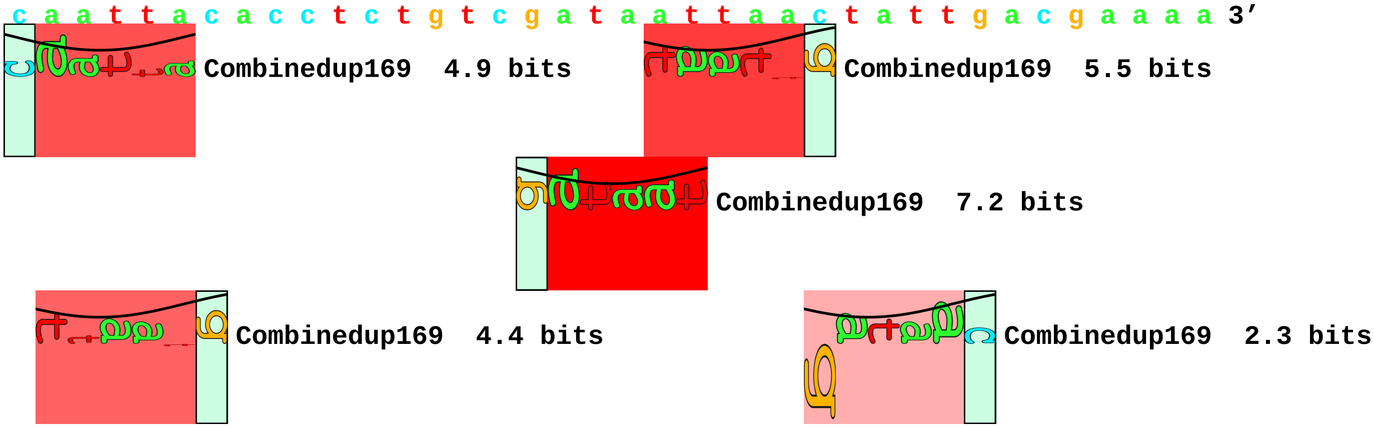

**lacUV5**

```
           *-60      .         *-50      .         *-40      .         *-30
5' g a a t t c t c a c t c a t t a g g c a c c c c a g g c t t t a c a c t t t 3'
```

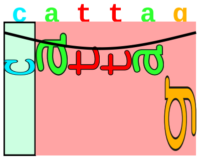

Combinedup169 2.6 bits

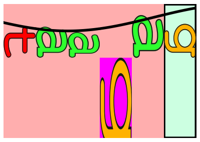

Combinedup169 2.3 bits

**RNA-I**

```
           *-60      .         *-50      .         *-40      .         *-30
5' g a g g t a t g t a g g c g g t g c t a c a g a g t t c t t g a a g t g g t 3'
```

**P1(SUB)**

```
           *-60      .         *-50      .         *-40      .         *-30
5' c g g t c g a c t g c a g t g g t a c c t a g g c c t c t t g t c a g g c c 3'
```

**P1(-41A)**

```
           *-60      .         *-50      .         *-40      .         *-30
5' c c c t t t c g t c t t c a a g a a t t c c a t c c t c t t g t c a g g c c 3'
```

**P1(-41B)**

```
           *-60      .         *-50      .         *-40      .         *-30
5' g g g a t c c t c t a g c c g g a a t t c c a t c c t c t t g t c a g g c c 3'
```

**P2(-39B)**

```
           *-60      .         *-50      .         *-40      .         *-30
5' g c c c t t t c g t c t t c a a g a a t t c c g g t g c t t g a c t c t g t 3'
```
