## Supplement_for_sigma38_paper-directory for "*Escherichia coli σ*^38^ promoters use two UP elements instead of a −35 element: resolution of a paradox and discovery that *σ*^38^ transcribes ribosomal promoters": rrnBp1-bolAp1-mutation-map.pdf

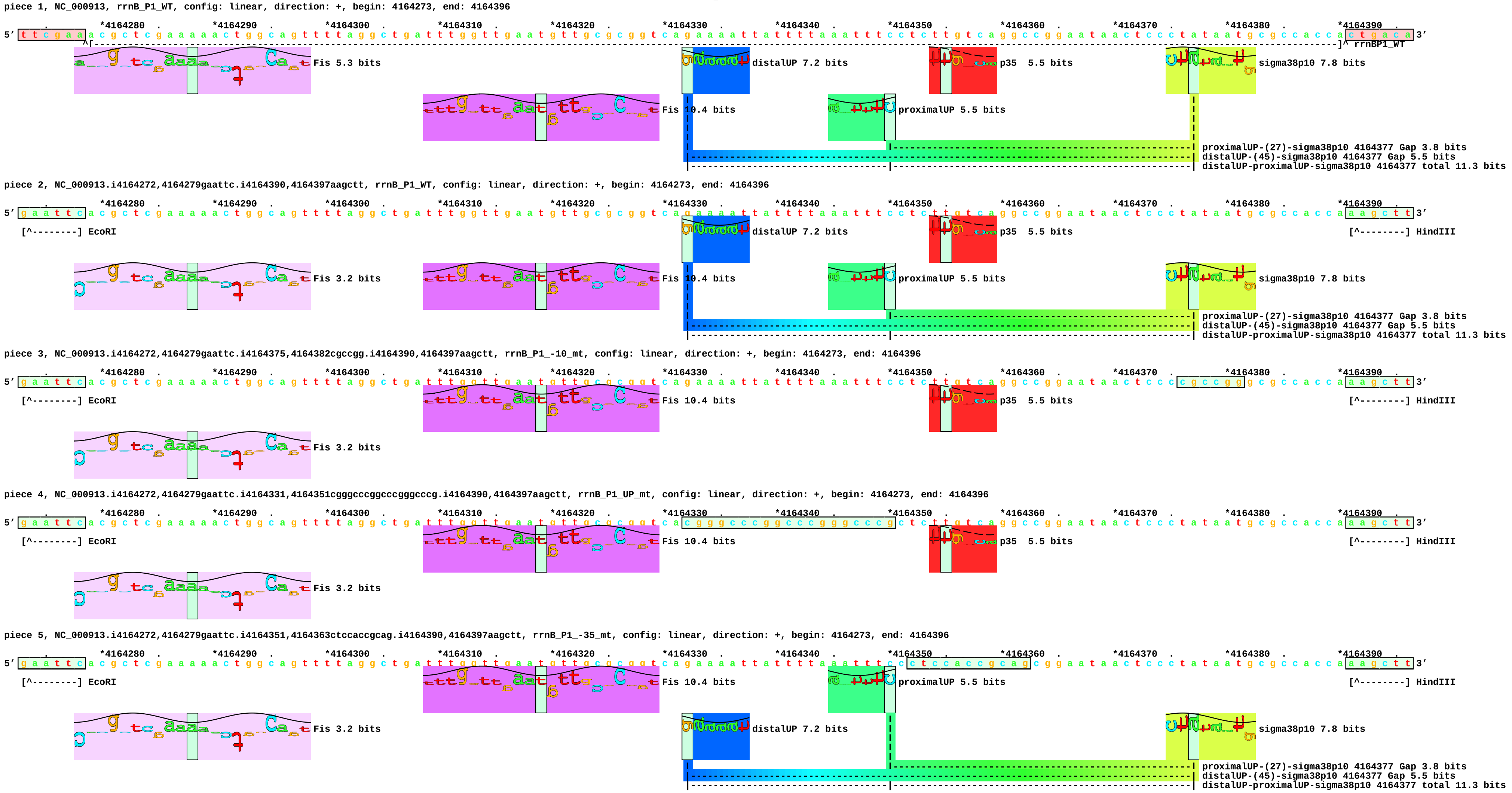
piece 1, NC_000913, rrnB_P1_WT, config: linear, direction: +, begin: 4164273, end: 4164396
5' ttcgaaacgctcgaaaaactggcagttttaggctgattttggttgaatgttgcgcggtcagaaaattatttttaaatttcctcttgtcaggccggaataactccctataatgcgccaccactgaca 3'
rrnBP1_WT
Fis 5.3 bits
Fis 10.4 bits
distalUP 7.2 bits
p35 5.5 bits
proximalUP 5.5 bits
sigma38p10 7.8 bits
proximalUP-(27)-sigma38p10 4164377 Gap 3.8 bits
distalUP-(45)-sigma38p10 4164377 Gap 5.5 bits
distalUP-proximalUP-sigma38p10 4164377 total 11.3 bits
piece 2, NC_000913.i4164272,4164279gaattc.i4164390,4164397aagctt, rrnB_P1_WT, config: linear, direction: +, begin: 4164273, end: 4164396
[^-------] EcoRI
[^-------] HindIII
Fis 3.2 bits
Fis 10.4 bits
distalUP 7.2 bits
p35 5.5 bits
proximalUP 5.5 bits
sigma38p10 7.8 bits
proximalUP-(27)-sigma38p10 4164377 Gap 3.8 bits
distalUP-(45)-sigma38p10 4164377 Gap 5.5 bits
distalUP-proximalUP-sigma38p10 4164377 total 11.3 bits
piece 3, NC_000913.i4164272,4164279gaattc.i4164375,4164382cgccgg.i4164390,4164397aagctt, rrnB_P1_-10_mt, config: linear, direction: +, begin: 4164273, end: 4164396
[^-------] EcoRI
[^-------] HindIII
Fis 10.4 bits
p35 5.5 bits
Fis 3.2 bits
piece 4, NC_000913.i4164272,4164279gaattc.i4164331,4164351cgggcccggcccgggcccg.i4164390,4164397aagctt, rrnB_P1_UP_mt, config: linear, direction: +, begin: 4164273, end: 4164396
[^-------] EcoRI
[^-------] HindIII
Fis 10.4 bits
p35 5.5 bits
Fis 3.2 bits
piece 5, NC_000913.i4164272,4164279gaattc.i4164351,4164363ctccaccgcag.i4164390,4164397aagctt, rrnB_P1_-35_mt, config: linear, direction: +, begin: 4164273, end: 4164396
[^-------] EcoRI
[^-------] HindIII
Fis 10.4 bits
proximalUP 5.5 bits
Fis 3.2 bits
distalUP 7.2 bits
sigma38p10 7.8 bits
proximalUP-(27)-sigma38p10 4164377 Gap 3.8 bits
distalUP-(45)-sigma38p10 4164377 Gap 5.5 bits
distalUP-proximalUP-sigma38p10 4164377 total 11.3 bits

piece 6, NC_000913, bolA_P1_WT, config: linear, direction: +, begin: 453570, end: 453678

```
  *453570   .        *453580   .        *453590   .        *453600   .        *453610   .        *453620   .        *453630   .        *453640   .        *453650   .        *453660   .        *453670   .
5' c a t a c g c t t g a c g g a a a a a a c c a g g a c g a a a c c t a a a t a t t t g t t g t t a a g c t g c a a t g g a a a c g g t a a a a g c g g c t a g t a t t t a a a g g g a t g g a t g a c a t c t c a g c g t t 3'
            ^[------------------------------------------------------------------------------------------------------------------------------------------------------------------------------------]^ bolAP1_WT
```

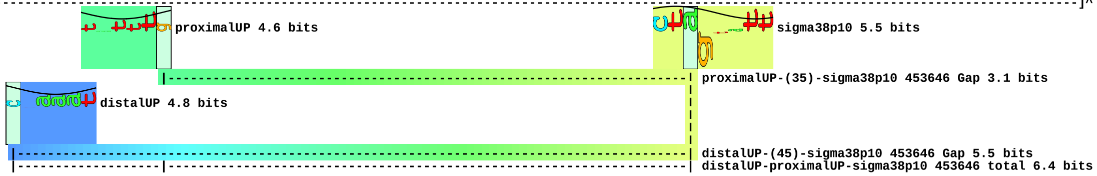


piece 7, NC_000913.i453569,453576gaattc.i453672,453679aagctt, bolA_P1_WT, config: linear, direction: +, begin: 453570, end: 453678

```
  *453570   .        *453580   .        *453590   .        *453600   .        *453610   .        *453620   .        *453630   .        *453640   .        *453650   .        *453660   .        *453670   .
5' g a a t t c c t t g a c g g a a a a a a c c a g g a c g a a a c c t a a a t a t t t g t t g t t a a g c t g c a a t g g a a a c g g t a a a a g c g g c t a g t a t t t a a a g g g a t g g a t g a c a t c t c a a g c t t 3'
   [^--------] EcoRI                                                                                                                                                                          [^--------] HindIII
```

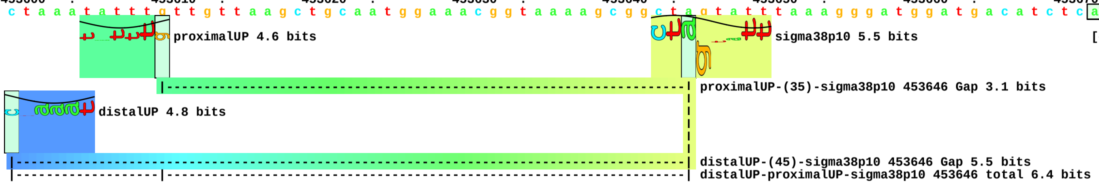


piece 8, NC_000913.i453569,453576gaattc.i453643,453652gacgtaca.i453672,453679aagctt, bolA_P1_-10 mt, config: linear, direction: +, begin: 453570, end: 453678

```
  *453570   .        *453580   .        *453590   .        *453600   .        *453610   .        *453620   .        *453630   .        *453640   .        *453650   .        *453660   .        *453670   .
5' g a a t t c c t t g a c g g a a a a a a c c a g g a c g a a a c c t a a a t a t t t g t t g t t a a g c t g c a a t g g a a a c g g t a a a a g c g g g a c g t a c a t a a a g g g a t g g a t g a c a t c t c a a g c t t 3'
   [^--------] EcoRI                                                                                                                                                                          [^--------] HindIII
```

piece 9, NC_000913.i453569,453576gaattc.i453602,453619cagcagtggtagtagg.i453672,453679aagctt, bolA_P1_up_mt, config: linear, direction: +, begin: 453570, end: 453678

```
  *453570   .        *453580   .        *453590   .        *453600   .        *453610   .        *453620   .        *453630   .        *453640   .        *453650   .        *453660   .        *453670   .
5' g a a t t c c t t g a c g g a a a a a a c c a g g a c g a a a c c t c a g c a g t g g t a g t a g g g c t g c a a t g g a a a c g g t a a a a g c g g c t a g t a t t t a a a g g g a t g g a t g a c a t c t c a a g c t t 3'
   [^--------] EcoRI                                                                                                                                                                          [^--------] HindIII
```

piece 10, NC_000913.i453569,453576gaattc.i453620,453632ctccaccgcag.i453672,453679aagctt, bolA_P1_-35_mt, config: linear, direction: +, begin: 453570, end: 453678

```
  *453570   .        *453580   .        *453590   .        *453600   .        *453610   .        *453620   .        *453630   .        *453640   .        *453650   .        *453660   .        *453670   .
5' g a a t t c c t t g a c g g a a a a a a c c a g g a c g a a a c c t a a a t a t t t g t t g t t a a g c c t c c a c c g c a g c g g t a a a a g c g g c t a g t a t t t a a a g g g a t g g a t g a c a t c t c a a g c t t 3'
   [^--------] EcoRI                                                                                                                                                                          [^--------] HindIII
```

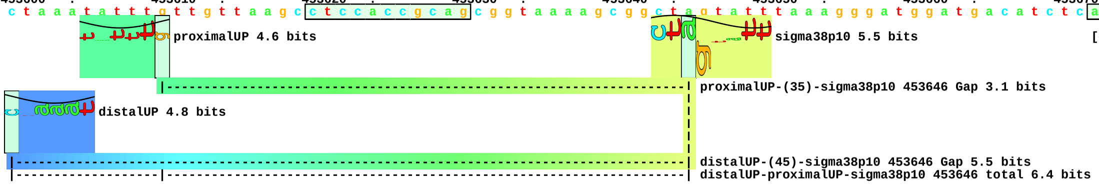
