## Supplement_for_sigma38_paper-directory for "*Escherichia coli σ*^38^ promoters use two UP elements instead of a −35 element: resolution of a paradox and discovery that *σ*^38^ transcribes ribosomal promoters": sigma38.169.combinedUP.instshiftp.txt

1.58      version of instshift that this parameter file is designed for.
+0        number of bases to shift the coordinate system
N -10 +10 rangechange: nrft fromrange torange
n -1  +1  splatter: ns fromrange torange
0.007     frequency of random mutations for "m" splatter
-0.5      timeseed for random mutations
