## Supplement_for_sigma38_paper-directory for "*Escherichia coli σ*^38^ promoters use two UP elements instead of a −35 element: resolution of a paradox and discovery that *σ*^38^ transcribes ribosomal promoters": sigma38.169.histog.txt

* genhis 1.95: generalized histogram data from
*
* parameters:
*            1  is the data column used
*          169  numbers are in the file
*       -43.00  is the minimum number
*       -27.00  is the maximum number
*       -34.91  is the MEAN
*         3.95  is the STANDARD DEVIATION
*         0.30  is the STANDARD ERROR OF THE MEAN (SEM)
*        15.62  is the variance
*         3.66  is the uncertainty in bits
*         4.03  is the computed uncertainty in bits (Shannon p.57)
*
*       -43.00  to        -27.00 is the range of data plotted
*         1.00  is the x-axis interval
*     17     is the number of intervals
*           32 is the maximum y-axis value
*         1.00 is the y-axis scale
*
* gaussian standard plotted
*        17.06 is the maximum Gaussian curve value 
*
* interval  | histogram | histogram
* beginning | number    | frequency
*
       -43.00            5  0.0296 ++*++
       -42.00            5  0.0296 +++*+
       -41.00            4  0.0237 ++++ :
       -40.00           11  0.0651 ++++++++*++
       -39.00            1  0.0059 +         :
       -38.00           19  0.1124 +++++++++++++*+++++
       -37.00            8  0.0473 ++++++++       :
       -36.00           15  0.0888 +++++++++++++++ :
       -35.00           21  0.1243 ++++++++++++++++*++++
       -34.00           32  0.1893 +++++++++++++++*++++++++++++++++
       -33.00           11  0.0651 +++++++++++  :
       -32.00            9  0.0533 +++++++++  :
       -31.00            7  0.0414 +++++++ :
       -30.00            3  0.0178 +++   :
       -29.00            1  0.0059 +   :
       -28.00            4  0.0237 ++*+
       -27.00           13  0.0769 +*+++++++++++
* genhis 1.95: generalized histogram data from
*
* parameters:
*            1  is the data column used
*          169  numbers are in the file
*       -56.00  is the minimum number
*       -40.00  is the maximum number
*       -45.88  is the MEAN
*         5.29  is the STANDARD DEVIATION
*         0.41  is the STANDARD ERROR OF THE MEAN (SEM)
*        28.01  is the variance
*         3.71  is the uncertainty in bits
*         4.45  is the computed uncertainty in bits (Shannon p.57)
*
*       -56.00  to        -40.00 is the range of data plotted
*         1.00  is the x-axis interval
*     17     is the number of intervals
*           25 is the maximum y-axis value
*         1.00 is the y-axis scale
*
* gaussian standard plotted
*        12.74 is the maximum Gaussian curve value 
*
* interval  | histogram | histogram
* beginning | number    | frequency
*
       -56.00           13  0.0769 +*+++++++++++
       -55.00            6  0.0355 ++*+++
       -54.00            6  0.0355 ++++*+
       -53.00            6  0.0355 +++++*
       -52.00            4  0.0237 ++++  :
       -51.00            5  0.0296 +++++   :
       -50.00           10  0.0592 +++++++++*
       -49.00            3  0.0178 +++       :
       -48.00            1  0.0059 +          :
       -47.00            2  0.0118 ++          :
       -46.00           10  0.0592 ++++++++++  :
       -45.00            4  0.0237 ++++       :
       -44.00           25  0.1479 +++++++++++*+++++++++++++
       -43.00           15  0.0888 +++++++++*+++++
       -42.00           18  0.1065 ++++++++*+++++++++
       -41.00           20  0.1183 +++++++*++++++++++++
       -40.00           21  0.1243 +++++*+++++++++++++++
