## Supplement_for_sigma38_paper-directory for "*Escherichia coli σ*^38^ promoters use two UP elements instead of a −35 element: resolution of a paradox and discovery that *σ*^38^ transcribes ribosomal promoters": sigma38.169.multiscanp.txt

1.03    version of multiscan that this parameter file is designed for.
200     maximum range
0     total Ri cutoff
f       f means filter out duplicates of the features
0       essential number cutoff (Ri - gap), if negative use scanp cutoff, no gap.
"sigma38p10" (null)  e s n b o3  parent, essential or non, orient tog, use GS, bar, overlap tog
"proximalUP" (sigma38p10)  e s g b o3  parent, essential or non, orient tog, use GS, bar, overlap tog
"distalUP" (sigma38p10)  e s g b o3  parent, essential or non, orient tog, use GS, bar, overlap tog
.
