## Supplement_for_sigma38_paper-directory for "*Escherichia coli σ*^38^ promoters use two UP elements instead of a −35 element: resolution of a paradox and discovery that *σ*^38^ transcribes ribosomal promoters": sigma38.169.ribl.txt

ribl "sigma38p10"
* Ri 2.82
*
* Ri(b,l) table is from:
** rseq 5.41 rsequence calculated from encoded sequences from:
** encode 1.42; encoding of sequences in
** * 2013/07/02 14:12:51, 2013/04/03 01:02:58, multiscan 3.96
*
* BOOK/INST sequences are from:
* 2013/07/02 14:12:51, 2013/04/03 01:02:58, multiscan 3.96
* title "sigma38 multiscan";
*
* PARAMETERS FOR Ri:
* -2 5 from-to
* 1 column of value file
* -2147483647.000000 2147483647.000000 lowest to highest Ri selected
* -2147483647.000000 2147483647.000000 lowest to highest Value selected
* not printing sequences to sequ
*     printing sequences to rixyin
* -: whole line printed when partial site
*  using Staden's Method: when f(b,l) = 0, replace with f(b,l) = 1/(n+t), t = 2.0
* 
* i alignmenttype first, book, instructions
*    Ri(a,l)    Ri(c,l)    Ri(g,l)    Ri(t,l)        l        a        c        g        t
      -2        5 frombase, tobase
   -1.021362   1.244532  -1.413679  -0.459483       -2       21      101       16       31
   -5.413679  -1.091751  -4.413679   1.776145       -1        1       20        2      146
    1.853107  -5.430653  -1.506789  -5.430653        0      154        0       15        0
    0.258746  -0.326217  -2.828717   0.871723        1       51       34        6       78
    0.978638  -1.954248  -0.243754  -0.165752        2       84       11       36       38
    0.230177   0.795774  -2.091751  -0.284396        3       50       74       10       35
   -1.413679  -5.413679  -4.413679   1.815139        4       16        1        2      150
   -5.430653  -1.243754  -0.954248   1.597548        5        0       18       22      129
*
  6.448546 bits = mean (Rsequence of selected region)
  2.642942 bits = standard deviation
*
 10.932606 bits = Ri of consensus sequence from -2 to 5
-29.977059 bits = Ri of anticonsensus sequence from -2 to 5
*
-10.381182 bits = average Ri for random sequence from -2 to 5
*
169 n, number of sequences used to create the matrix
*
asymmetric symmetry of the matrix
*
-5000.000000 Ri bound: lower bound on Ri
100.000000 Z bound: lower bound on Z
  1.000000 P bound: upper probability
*
h  extreme: char; h or l, the high or low extreme to be defined
-1.0  wavelocation: real; the location in bases of the extreme
 2.0  wavebit: real; the location in bits of the extreme
 0.5  waveamplitude: real; the amplitude of the wave in bits
10.6  wavelength: real; the wave length of the wave in bases
 0.0  0.2  0.0 dashon, dashoff, dashoffset (in bases)
 0.0  thickness: real; the thickness of the cosine wave.  <=0 means default
.

ribl "proximalUP"
* Ri 2.82
*
* Ri(b,l) table is from:
** rseq 5.41 rsequence calculated from encoded sequences from:
** encode 1.42; encoding of sequences in
** * 2016/08/25 10:11:27, 2016/02/04 12:17:57, target
*
* BOOK/INST sequences are from:
* 2016/08/25 10:11:27, 2016/02/04 12:17:57, target
* title 'target';
*
* PARAMETERS FOR Ri:
* -5 0 from-to
* 1 column of value file
* -2147483647.000000 2147483647.000000 lowest to highest Ri selected
* -2147483647.000000 2147483647.000000 lowest to highest Value selected
* not printing sequences to sequ
*     printing sequences to rixyin
* -: whole line printed when partial site
*  using Staden's Method: when f(b,l) = 0, replace with f(b,l) = 1/(n+t), t = 2.0
*
* i alignmenttype first, book, instructions
*    Ri(a,l)    Ri(c,l)    Ri(g,l)    Ri(t,l)        l        a        c        g        t
      -5        0 frombase, tobase
    1.078174  -1.828717  -5.430653   0.652410       -5       90       12        0       67
    0.045752  -5.430653  -0.506789   1.156176       -4       44        0       30       95
    0.926171  -2.606325  -5.430653   0.926171       -3       81        7        0       81
    1.244532  -5.430653  -5.430653   0.673783       -2      101        0        0       68
   -0.165752  -5.413679  -5.430653   1.608688       -1       38        1        0      130
   -5.430653   1.186233   0.694845  -3.828717        0        0       97       69        3
*
  5.134314 bits = mean (Rsequence of selected region)
  1.666666 bits = standard deviation
*
  7.199974 bits = Ri of consensus sequence from -5 to 0
-32.583915 bits = Ri of anticonsensus sequence from -5 to 0
*
-10.542903 bits = average Ri for random sequence from -5 to 0
*
169 n, number of sequences used to create the matrix
*
asymmetric symmetry of the matrix
*
-5000.000000 Ri bound: lower bound on Ri
100.000000 Z bound: lower bound on Z
  1.000000 P bound: upper probability
*
h  extreme: char; h or l, the high or low extreme to be defined
+2.5  wavelocation: real; the location in bases of the extreme
 2.0  wavebit: real; the location in bits of the extreme
 0.5  waveamplitude: real; the amplitude of the wave in bits
10.6  wavelength: real; the wave length of the wave in bases
 0.0  0.2  0.0 dashon, dashoff, dashoffset (in bases)
 0.0  thickness: real; the thickness of the cosine wave.  <=0 means default
.

ribl "distalUP"
* Ri 2.82
*
* Ri(b,l) table is from:
** rseq 5.41 rsequence calculated from encoded sequences from:
** encode 1.42; encoding of sequences in
** * 2016/08/25 10:17:02, 2016/02/04 12:17:57, target
*
* BOOK/INST sequences are from:
* 2016/08/25 10:17:02, 2016/02/04 12:17:57, target
* title 'target';
*
* PARAMETERS FOR Ri:
* 0 5 from-to
* 1 column of value file
* -2147483647.000000 2147483647.000000 lowest to highest Ri selected
* -2147483647.000000 2147483647.000000 lowest to highest Value selected
* not printing sequences to sequ
*     printing sequences to rixyin
* -: whole line printed when partial site
*  using Staden's Method: when f(b,l) = 0, replace with f(b,l) = 1/(n+t), t = 2.0
*
* i alignmenttype first, book, instructions
*    Ri(a,l)    Ri(c,l)    Ri(g,l)    Ri(t,l)        l        a        c        g        t
       0        5 frombase, tobase
   -5.430653   0.517058   1.327788  -5.413679        0        0       61      107        1
    1.575005  -5.430653  -5.430653  -0.021362        1      127        0        0       42
    0.978638  -5.430653  -2.828717   0.890101        2       84        0        6       79
    0.961360  -3.828717  -5.413679   0.943873        3       83        3        1       82
    1.140909  -2.091751  -3.828717   0.540517        4       94       10        3       62
    0.563600  -2.606325  -5.430653   1.215677        5       63        7        0       99
*
  5.261177 bits = mean (Rsequence of selected region)
  1.700187 bits = standard deviation
*
  7.199377 bits = Ri of consensus sequence from 0 to 5
-30.965006 bits = Ri of anticonsensus sequence from 0 to 5
*
-10.632921 bits = average Ri for random sequence from 0 to 5
*
169 n, number of sequences used to create the matrix
*
asymmetric symmetry of the matrix
*
-5000.000000 Ri bound: lower bound on Ri
100.000000 Z bound: lower bound on Z
  1.000000 P bound: upper probability
*
h  extreme: char; h or l, the high or low extreme to be defined
-2.5  wavelocation: real; the location in bases of the extreme
 2.0  wavebit: real; the location in bits of the extreme
 0.5  waveamplitude: real; the amplitude of the wave in bits
10.6  wavelength: real; the wave length of the wave in bases
 0.0  0.2  0.0 dashon, dashoff, dashoffset (in bases)
 0.0  thickness: real; the thickness of the cosine wave.  <=0 means default
.
