## Supplement_for_sigma38_paper-directory for "*Escherichia coli σ*^38^ promoters use two UP elements instead of a −35 element: resolution of a paradox and discovery that *σ*^38^ transcribes ribosomal promoters": sigma38.169.scanp.txt

2.90    version of scan that this parameter file is designed for.
-1      number of seqs to scan 0 = none, positive = that number; negative = all
0.0       information content at or above which to report in the data file.
100     Z score below which to report in the data file
0       probability at or above which to report in the data file.
-2 +5   desired region of the ribl weight matrix to use
1       0: program figures it out; 1: one way scan; 2: two way scan.
define "sigma38p10" " w" "  " "  " -2 +5
 string "@"
 coordinate 4
 string "|Ri="
Ri 1 1  Riwidth Ridecimal: character places for reporting bits to scanfeatures
string " bits"
 string "|Z="
 Z 7 4   z score
 string "|p="
 probability 5 3
.  end of print definitions
DFi     dfi: data, features, inst: files output
n       normalizeRi: n: normal, s: Ri(l)-Ri(0), d: Ri(l)/Ri(0)   
-50 +50 range to make the scaninst file (if made)
2.90    version of scan that this parameter file is designed for.
-1      number of seqs to scan 0 = none, positive = that number; negative = all
0.0       information content at or above which to report in the data file.
100     Z score below which to report in the data file
0       probability at or above which to report in the data file.
-5 +0   desired region of the ribl weight matrix to use
1       0: program figures it out; 1: one way scan; 2: two way scan.
define "proximalUP" " w" "  " "  " -5 +0
 string "@"
 coordinate 4
 string "|Ri="
Ri 1 1  Riwidth Ridecimal: character places for reporting bits to scanfeatures
string " bits"
 string "|Z="
 Z 7 4   z score
 string "|p="
 probability 5 3
.  end of print definitions
DFi     dfi: data, features, inst: files output
n       normalizeRi: n: normal, s: Ri(l)-Ri(0), d: Ri(l)/Ri(0)   
-50 +50 range to make the scaninst file (if made)
2.90    version of scan that this parameter file is designed for.
-1      number of seqs to scan 0 = none, positive = that number; negative = all
0.0       information content at or above which to report in the data file.
100     Z score below which to report in the data file
0       probability at or above which to report in the data file.
-0 +5   desired region of the ribl weight matrix to use
1       0: program figures it out; 1: one way scan; 2: two way scan.
define "distalUP" " w" "  " "  " -0 +5
 string "@"
 coordinate 4
 string "|Ri="
Ri 1 1  Riwidth Ridecimal: character places for reporting bits to scanfeatures
string " bits"
 string "|Z="
 Z 7 4   z score
 string "|p="
 probability 5 3
.  end of print definitions
DFi     dfi: data, features, inst: files output
n       normalizeRi: n: normal, s: Ri(l)-Ri(0), d: Ri(l)/Ri(0)   
-50 +50 range to make the scaninst file (if made)
