## Supplement_for_sigma38_paper-directory for "*Escherichia coli σ*^38^ promoters use two UP elements instead of a −35 element: resolution of a paradox and discovery that *σ*^38^ transcribes ribosomal promoters": sigma38coverfigure.pdf

### COVER FIGURE
### for
### *Escherichia coli* $\sigma^{38}$ promoters use two UP elements instead of a $-35$ element: resolution of a paradox and discovery that $\sigma^{38}$ transcribes ribosomal promoters

Kevin S. Franco, Zhe Sun, Yixiong Chen, Cedric Cagliero, Yuhong Zuo,
Yan Ning Zhou, Mikhail Kashlev, Ding Jun Jin, and Thomas D. Schneider

version = 1.13 of sigma38coverfigure.tex 2018 Oct 02

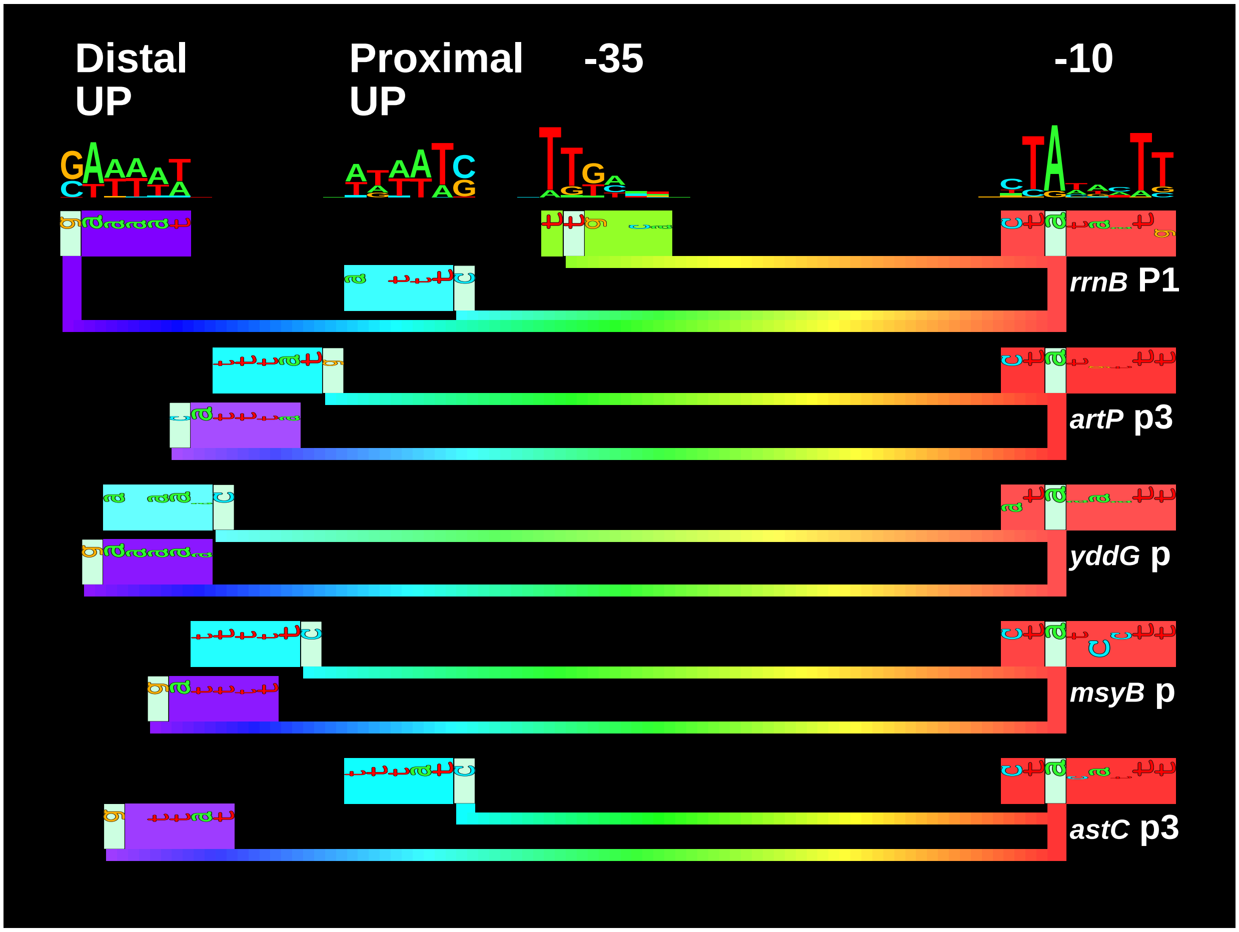


The identity of elements required for stress-responding $\sigma^{38}$ promoters in *Escherichia coli* has remained paradoxical for more than two decades. In contrast, housekeeping $\sigma^{70}$ promoters are well known to have −10, −35 and UP elements, as in the case of the ribosomal promoter *rrnB* P1. These four components are shown by sequence logos at the top of the figure. Below the logos are the four matching components of the *rrnB* P1 promoter of *E. coli* shown as sequence walkers, spaced as they are found in the genome and connected by spectrum-colored bars. Franco *et al.* show that most naturally occurring $\sigma^{38}$ promoters do not have a −35 component; instead they have a −10 and two strong UP elements. Four experimentally verified examples of $\sigma^{38}$ promoters are shown below *rrnB* P1: *artP* p3, *yddG* p, *msyB* p and *astC* p3. Franco *et al.* also found that the ribosomal operon promoter *rrnB* P1 is activated by $\sigma^{38}$ *in vitro*, but the −35 is not used. This has important physiological implications for *E. coli* since it implies that the ribosomal promoters should continue firing during entry to stationary phase.
