## Supplement_for_sigma38_paper-directory for "*Escherichia coli σ*^38^ promoters use two UP elements instead of a −35 element: resolution of a paradox and discovery that *σ*^38^ transcribes ribosomal promoters": Typas.Hengge2005-supplement-S1-map.pdf

piece 1, NC_000913.2, adhE-P1, config: linear, direction: -, begin: 1297722, end: 1297616

```
     *1297720  .        *1297710  .        *1297700  .        *1297690  .
5' c t a c t t a a a a t t g c t a t c a t t c g t t a t t g t t a t c t a g t t g 3'
                                                          [------------------ ... adhE-P1
distal  4.9 bits
                                                                              ... distal-(53)-sigma38p10 1297647 Gap
                                                                              ... sigma38p10-proximal-distal 1297647 total
crp  2.0 bits
crp  1.8 bits

     *1297680  .        *1297670  .        *1297660  .        *1297650  .
5' t g c a a a a c a t g c t a a t g t a g c c a c c a a a t c a t a c t a c a a t 3'
... --------------------------------------------------------------------------- ... adhE-P1
proximal  1.4 bits                                                            ... sigma38p10
...                                                        distal-(53)-sigma38p10 1297647 Gap 4.9 bits
...                                                        sigma38p10-proximal-distal 1297647 total 5.1 bits
                                                           proximal-(28)-sigma38p10 1297647 Gap 5.5 bits
proximal  3.8 bits
                                                                              ... proximal-(40)-sigma38p10 1297626 Gap
distal  2.6 bits
                                                                              ... distal-(54)-sigma38p10 1297626 Gap
                                                                              ... sigma38p10-proximal-distal 1297626 total

     *1297640  .        *1297630  .        *1297620
5' t t a t t a a c t g t t a g c t a t a a t g g c g a a 3'
... ------------] adhE-P1          sigma38p10  7.8 bits
```

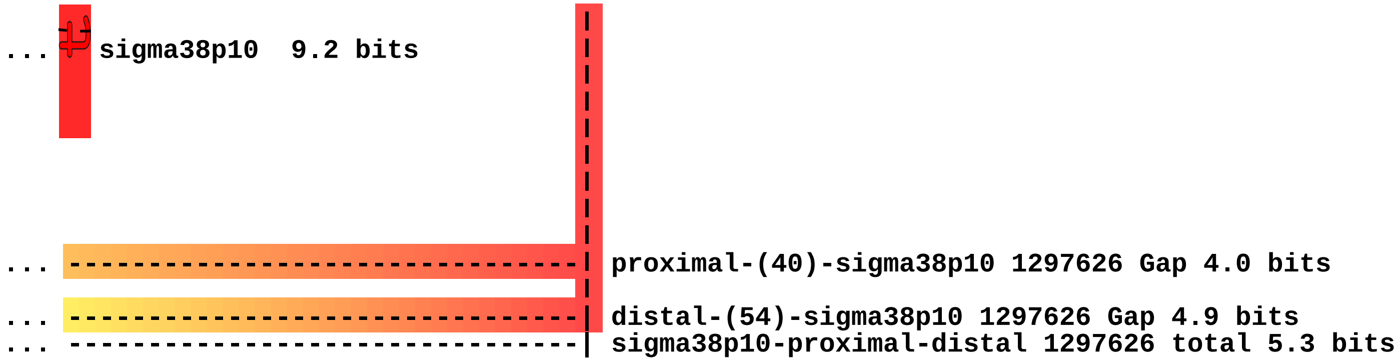

sigma38p10 9.2 bits
proximal-(40)-sigma38p10 1297626 Gap 4.0 bits
distal-(54)-sigma38p10 1297626 Gap 4.9 bits
sigma38p10-proximal-distal 1297626 total 5.3 bits

piece 2, NC_000913.2, osmB-P1, config: linear, direction: -, begin: 1341628, end: 1341524

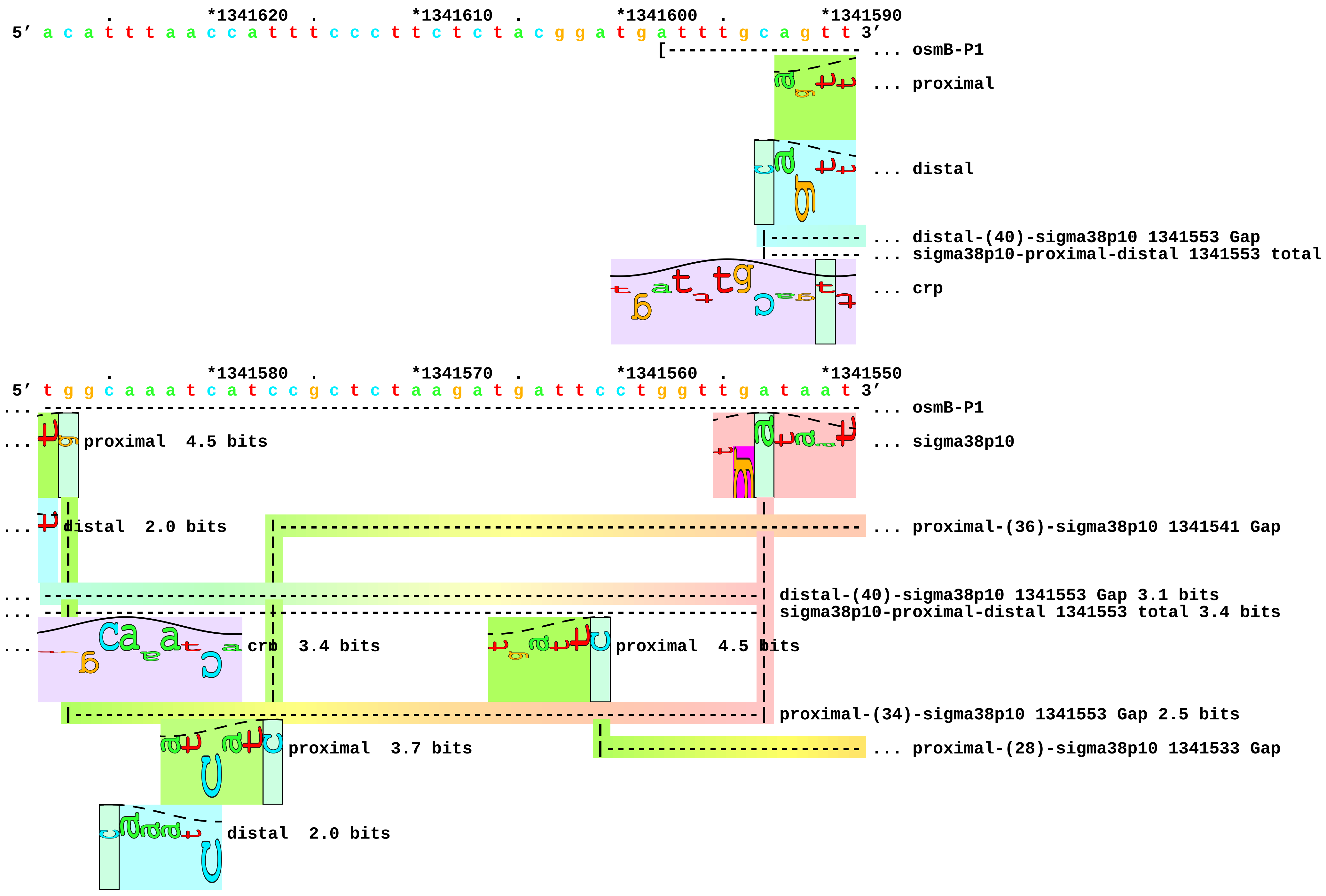

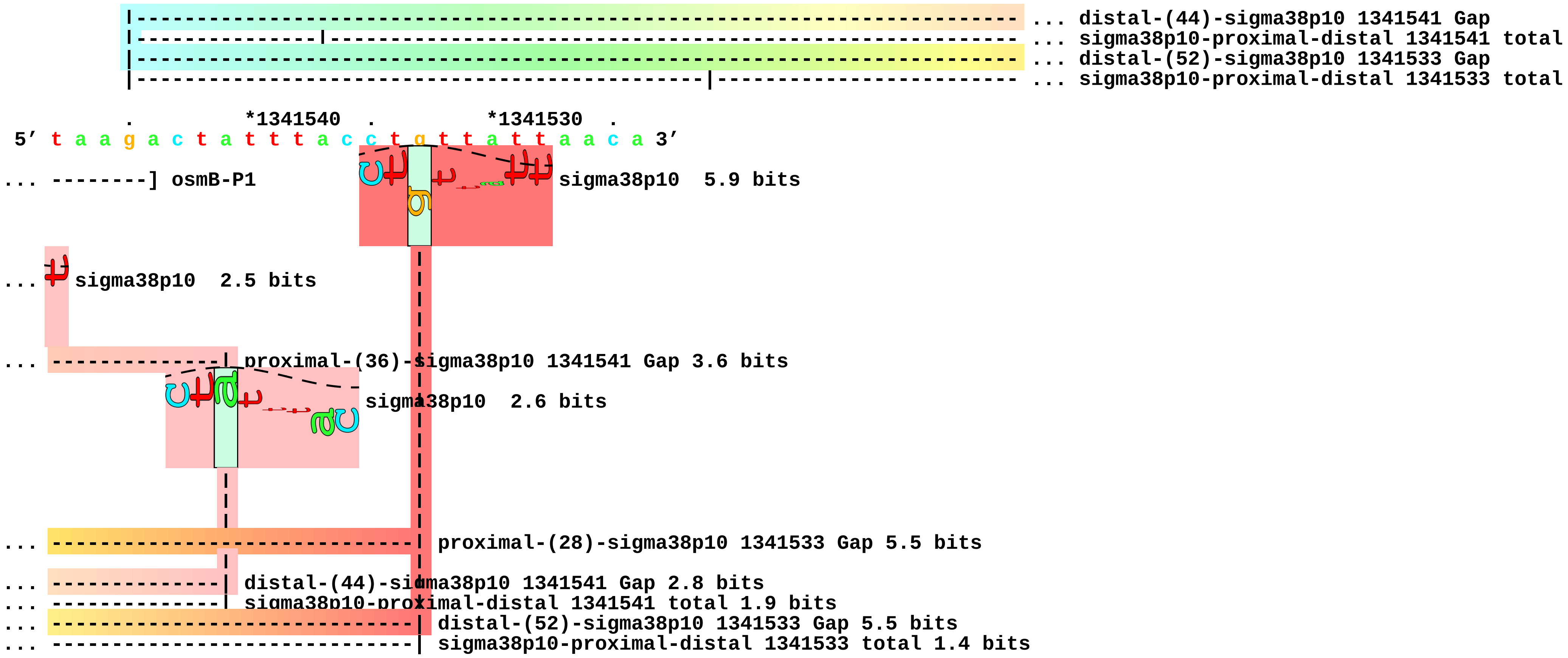
... distal-(44)-sigma38p10 1341541 Gap
... sigma38p10-proximal-distal 1341541 total
... distal-(52)-sigma38p10 1341533 Gap
... sigma38p10-proximal-distal 1341533 total
*1341540
*1341530
5′ t a a g a c t a t t t a c c t g t t a t t a a c a 3′
... --------] osmB-P1
sigma38p10 5.9 bits
... sigma38p10 2.5 bits
... proximal-(36)-sigma38p10 1341541 Gap 3.6 bits
sigma38p10 2.6 bits
... proximal-(28)-sigma38p10 1341533 Gap 5.5 bits
... distal-(44)-sigma38p10 1341541 Gap 2.8 bits
... sigma38p10-proximal-distal 1341541 total 1.9 bits
... distal-(52)-sigma38p10 1341533 Gap 5.5 bits
... sigma38p10-proximal-distal 1341533 total 1.4 bits

piece 3, NC_000913.2, xthA, config: linear, direction: +, begin: 1830338, end: 1830443

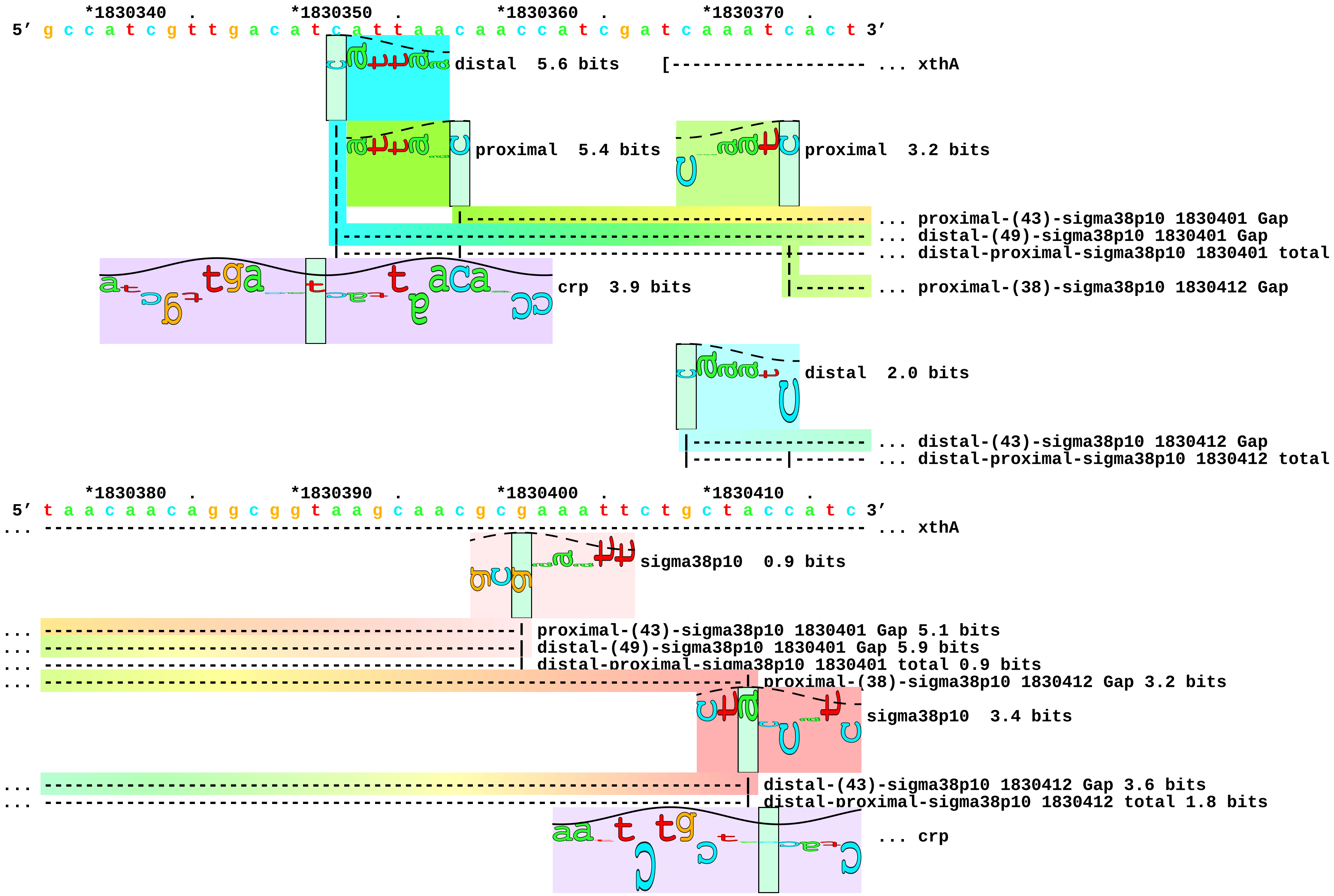

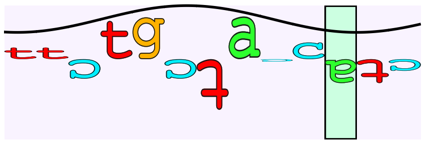

... crp

```
      *1830420   .         *1830430   .         *1830440
5' c a c g c a c t c t t t a t c t g a a t a a a t g g 3'
... ----------] xthA
```

... 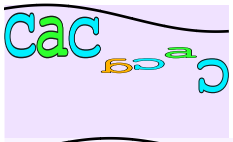 crp  2.9 bits

... 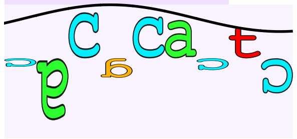 crp  1.2 bits piece 4, NC_000913.2, adhE-P2, config: linear, direction: -, begin: 1297616, end: 1297512

```
      .         *1297610  .         *1297600  .         *1297590  .         *1297580
5' a a a g c g a t g c t g a a a g g t g t c a g c t t t g c a a a a a t t t g a t 3'
                                                                  [---------------- ... adhE-P2
proximal  5.0 bits
crp  1.8 bits
... proximal-(35)-sigma38p10 1297544 Gap
distal  5.7 bits
... distal-(44)-sigma38p10 1297544 Gap
... sigma38p10-proximal-distal 1297544 total
crp  1.9 bits
... crp

      .         *1297570  .         *1297560  .         *1297550  .         *1297540
5' t t g g a t c a c g t a a t c a g t a c c c a g a a g t g a g t a a t c t t g c 3'
... ------------------------------------------------------------------------------ ... adhE-P2
sigma38p10  6.5 bits
... proximal-(35)-sigma38p10 1297544 Gap 3.1 bits
... crp
... distal-(44)-sigma38p10 1297544 Gap 2.8 bits
... sigma38p10-proximal-distal 1297544 total 11.4 bits
... crp
```

... 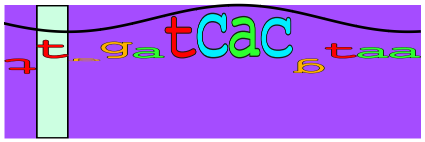 crp 17.2 bits

```
       .          *1297530  .          *1297520  .
 5' t t a c g c c a c c t g g a a g t g a c g c a t t 3'
... --------] adhE-P2
```

... 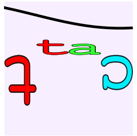 crp  1.6 bits

... 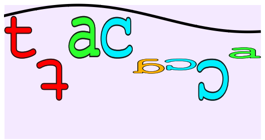 crp  2.2 bits piece 5, NC_000913.2, aidB, config: linear, direction: +, begin: 4412185, end: 4412292

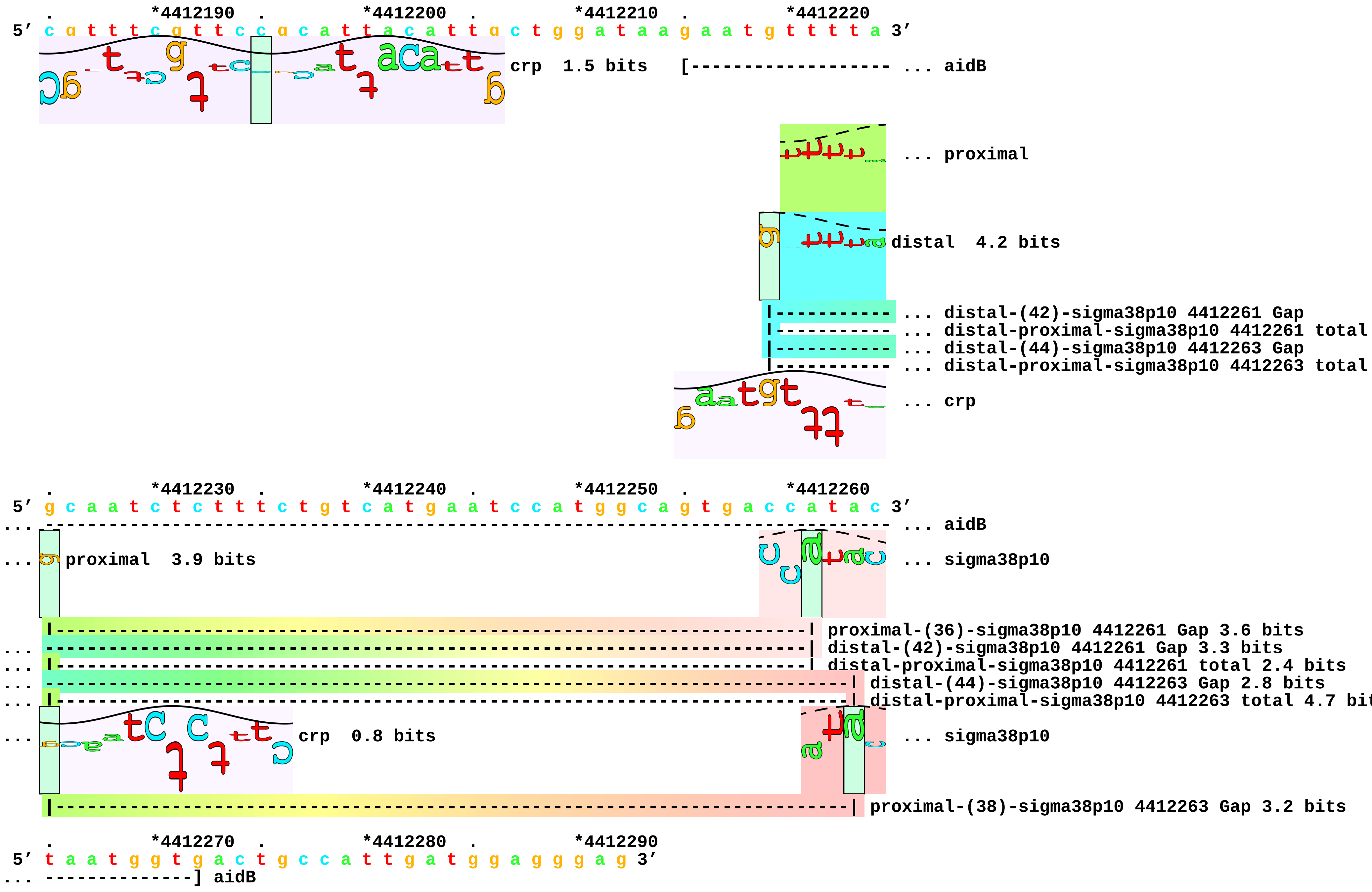

... 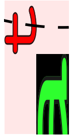 sigma38p10 1.0 bits

... 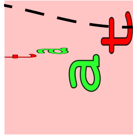 sigma38p10 2.5 bits piece 6, NC_000913.2, appY, config: linear, direction: +, begin: 582812, end: 582921

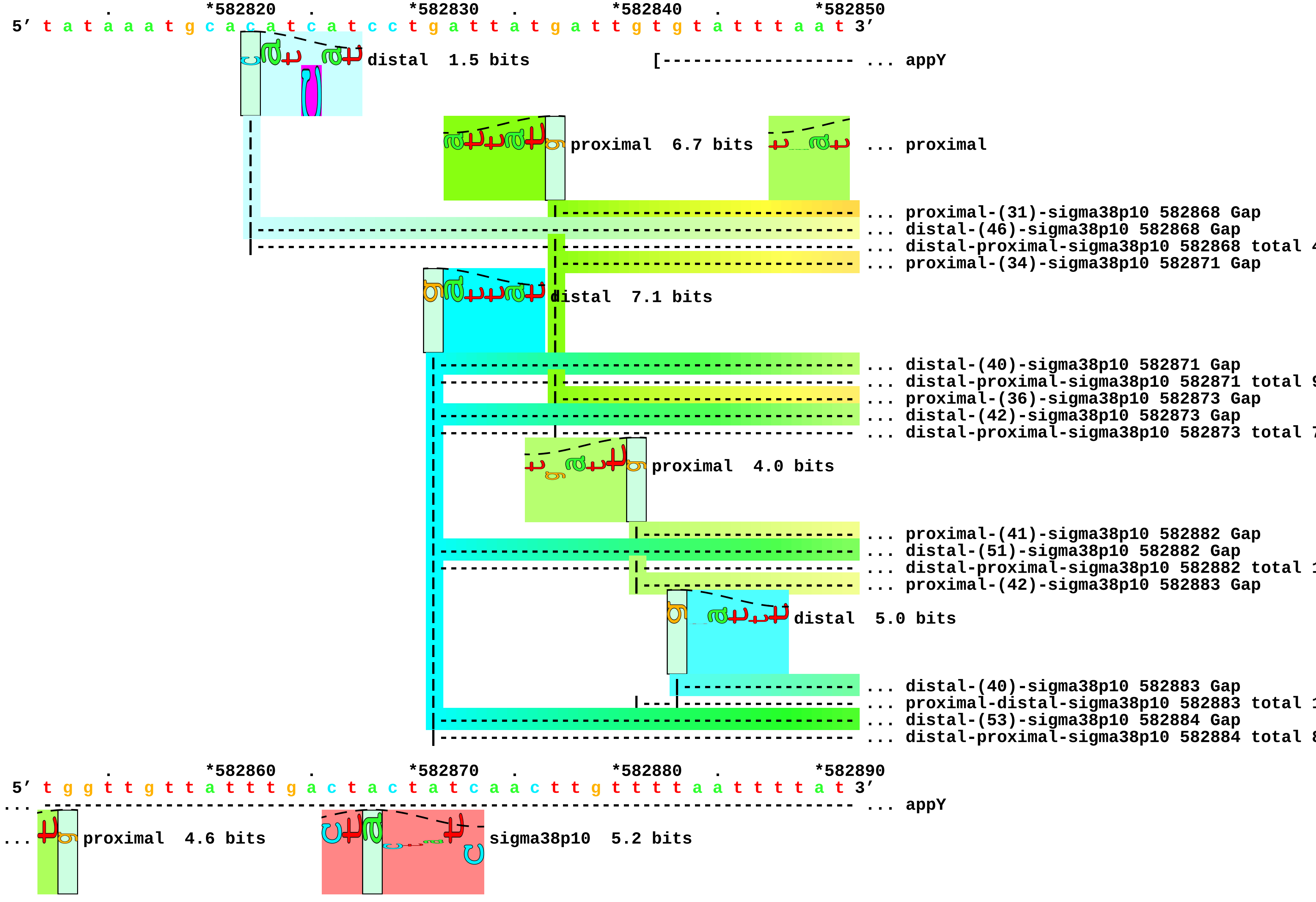

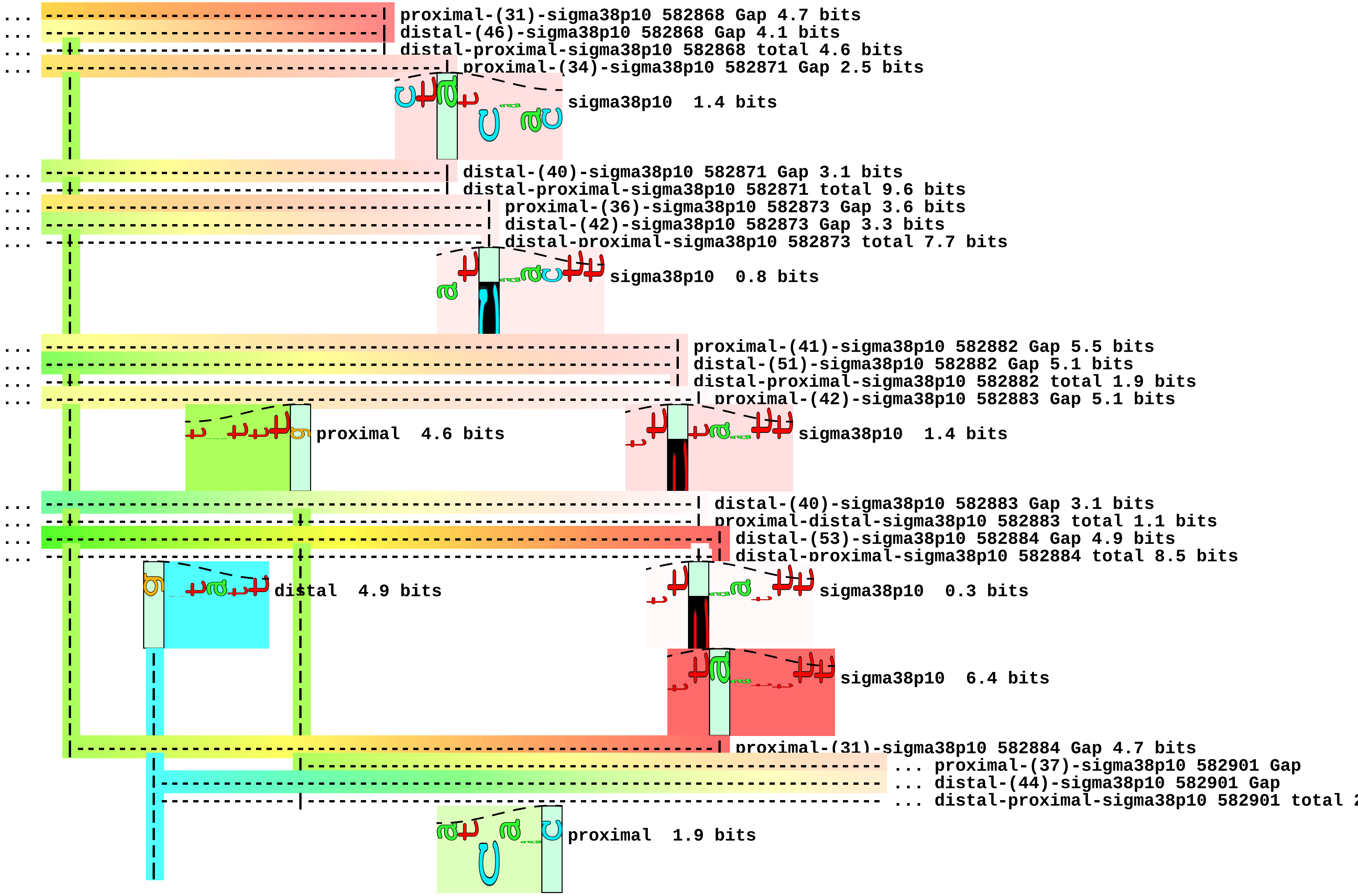
proximal-(31)-sigma38p10 582868 Gap 4.7 bits
distal-(46)-sigma38p10 582868 Gap 4.1 bits
distal-proximal-sigma38p10 582868 total 4.6 bits
proximal-(34)-sigma38p10 582871 Gap 2.5 bits
sigma38p10 1.4 bits
distal-(40)-sigma38p10 582871 Gap 3.1 bits
distal-proximal-sigma38p10 582871 total 9.6 bits
proximal-(36)-sigma38p10 582873 Gap 3.6 bits
distal-(42)-sigma38p10 582873 Gap 3.3 bits
distal-proximal-sigma38p10 582873 total 7.7 bits
sigma38p10 0.8 bits
proximal-(41)-sigma38p10 582882 Gap 5.5 bits
distal-(51)-sigma38p10 582882 Gap 5.1 bits
distal-proximal-sigma38p10 582882 total 1.9 bits
proximal-(42)-sigma38p10 582883 Gap 5.1 bits
proximal 4.6 bits
sigma38p10 1.4 bits
distal-(40)-sigma38p10 582883 Gap 3.1 bits
proximal-distal-sigma38p10 582883 total 1.1 bits
distal-(53)-sigma38p10 582884 Gap 4.9 bits
distal-proximal-sigma38p10 582884 total 8.5 bits
distal 4.9 bits
sigma38p10 0.3 bits
sigma38p10 6.4 bits
proximal-(31)-sigma38p10 582884 Gap 4.7 bits
... proximal-(37)-sigma38p10 582901 Gap
... distal-(44)-sigma38p10 582901 Gap
... distal-proximal-sigma38p10 582901 total
proximal 1.9 bits

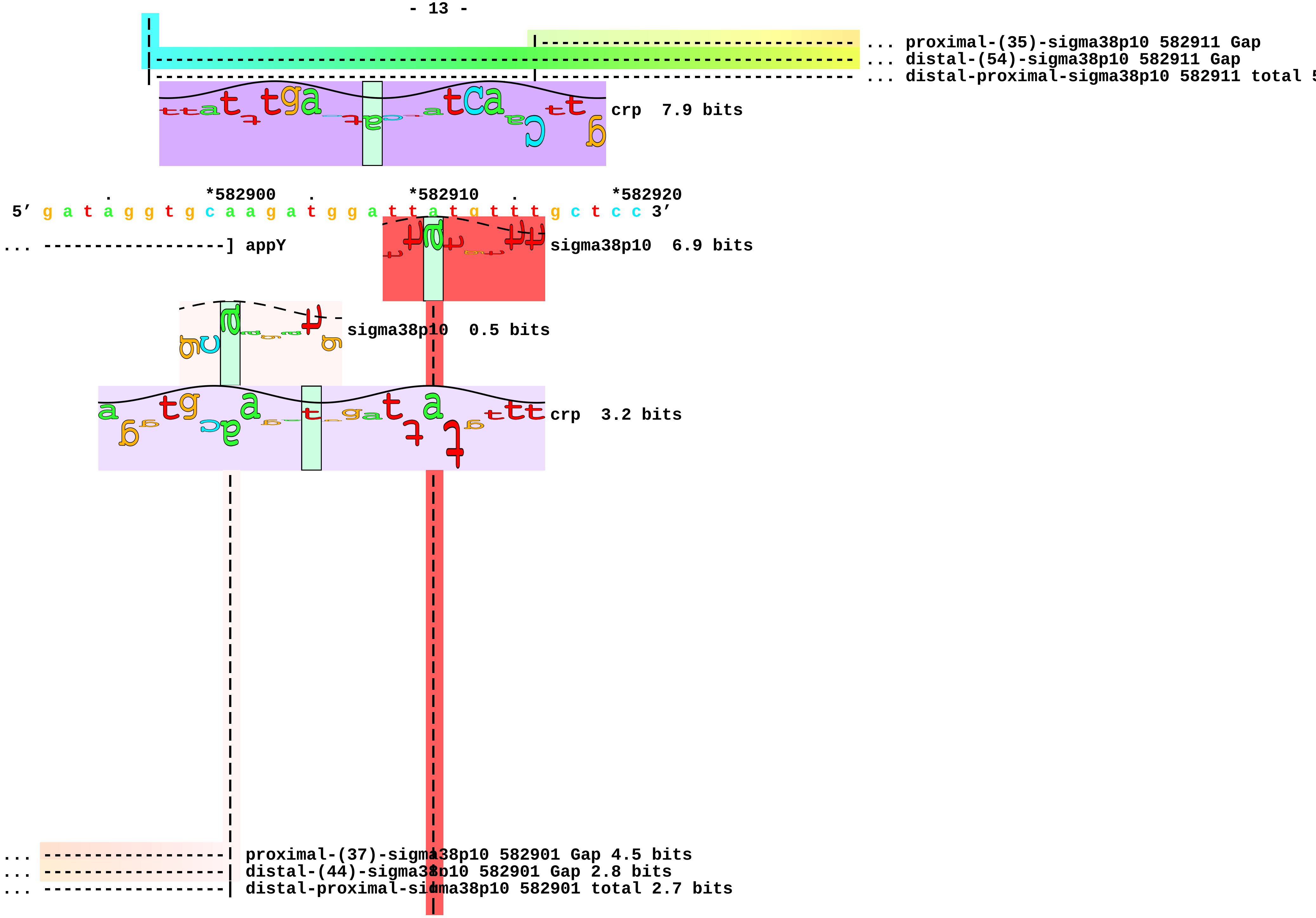
... proximal-(35)-sigma38p10 582911 Gap
... distal-(54)-sigma38p10 582911 Gap
... distal-proximal-sigma38p10 582911 total
crp 7.9 bits
*582900
*582910
*582920
5′ g a t a g g t g c a a g a t g g a t t a t g t t t g c t c c 3′
... ------------------] appY
sigma38p10 6.9 bits
sigma38p10 0.5 bits
crp 3.2 bits
... ------------------| proximal-(37)-sigma38p10 582901 Gap 4.5 bits
... ------------------| distal-(44)-sigma38p10 582901 Gap 2.8 bits
... ------------------| distal-proximal-sigma38p10 582901 total 2.7 bits

```
... --------------------------------------| proximal-(35)-sigma38p10 582911 Gap 3.1 bits
... --------------------------------------| distal-(54)-sigma38p10 582911 Gap 4.9 bits
... --------------------------------------| distal-proximal-sigma38p10 582911 total 5.8 bits
```

piece 7, NC_000913.2, blc, config: linear, direction: -, begin: 4375852, end: 4375748

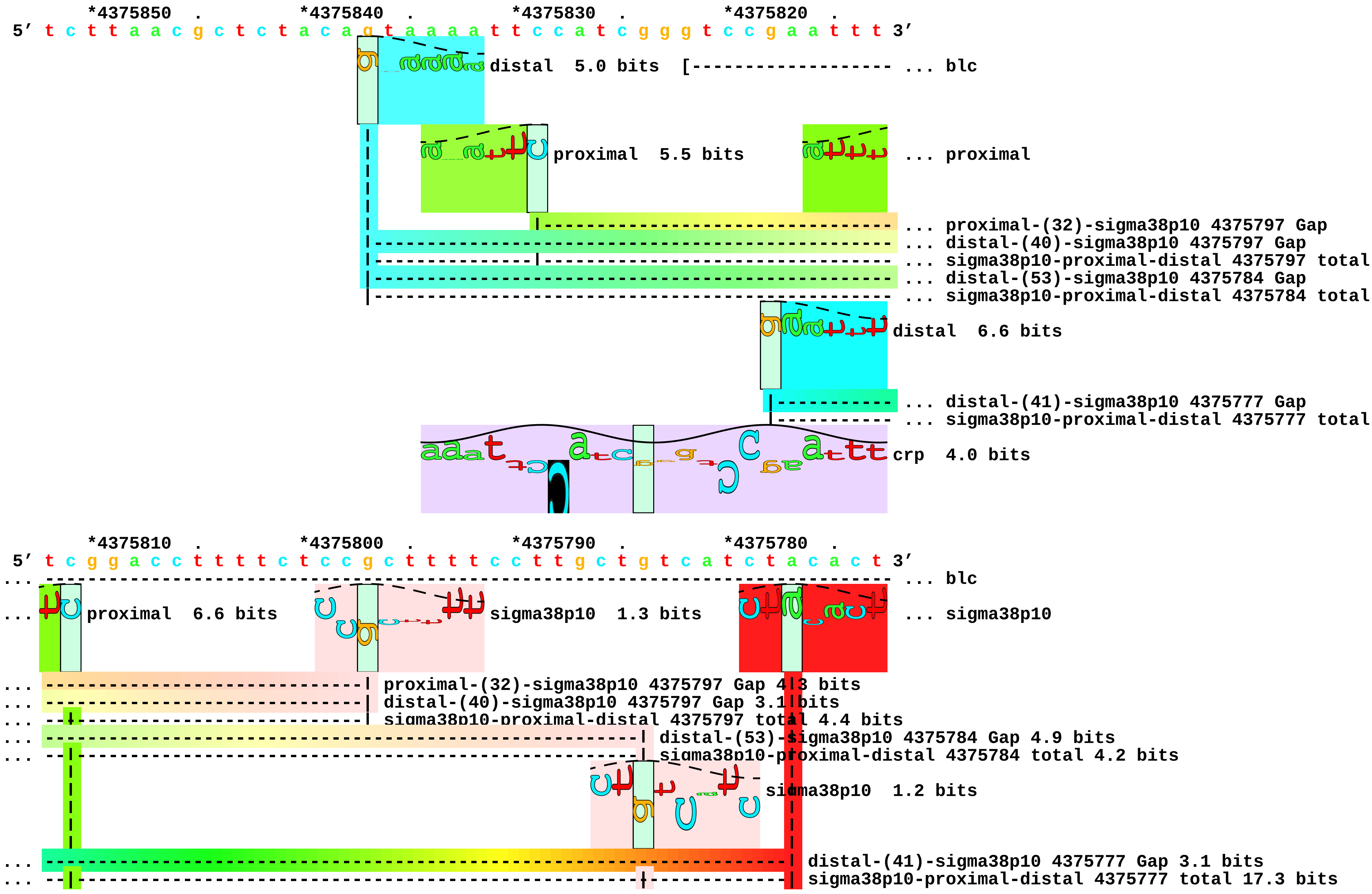

```
|------------------------------------------------------------| proximal-(27)-sigma38p10 4375784 Gap 3.8 bits
|---------------------------------------------------------------| proximal-(34)-sigma38p10 4375777 Gap 2.5 bits
```

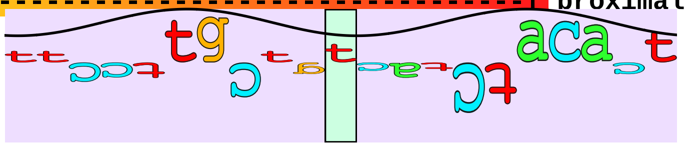

... crp

```
         *4375770  .         *4375760  .         *4375750
5' t a g a a a a a a a c c a g t a a g g a a a c a t 3'
... --------] blc
... t sigma38p10  9.7 bits
... t crp  3.2 bits
```

piece 8, NC_000913.2, bolA-P1, config: linear, direction: +, begin: 453570, end: 453677

```
   *453570    .          *453580    .          *453590    .          *453600    .
5' c a t a c g c t t g a c g g a a a a a c c a g g a c g a a a c c t a a a t a t t 3'
                                                                  g-g---------------- ... bolA-P1.-57.-56
                                                                                      ... proximal
                                                                              distal  4.8 bits
                                                                     |--------------- ... distal-(45)-sigma38p10 453646 Gap
                                                                     |--------------- ... distal-proximal-sigma38p10 453646 total [illegible]

   *453610    .          *453620    .          *453630    .          *453640    .
5' t g t t g t t a a g c t g c a a t g g a a a c g g t a a a a g c g g c t a g t a 3'
... ------------------------------------------------------------------------------- ... bolA-P1.-57.-56
...      proximal  4.6 bits                                                          ... sigma38p10
      |---------------------------------------------------------------------| proximal-(35)-sigma38p10 453646 Gap 3.1 bits
... ------------------------------------------------------------------------| distal-(45)-sigma38p10 453646 Gap 5.5 bits
... --|---------------------------------------------------------------------| distal-proximal-sigma38p10 453646 total 6.4 bits

   *453650    .          *453660    .          *453670    .
5' t t t a a a g g g a t g g a t g a c a t c t c a g c g t 3'
... --------------] bolA-P1.-57.-56
...      sigma38p10  5.5 bits
```

piece 9, NC_000913.2, cbpA, config: linear, direction: -, begin: 1063135, end: 1063033

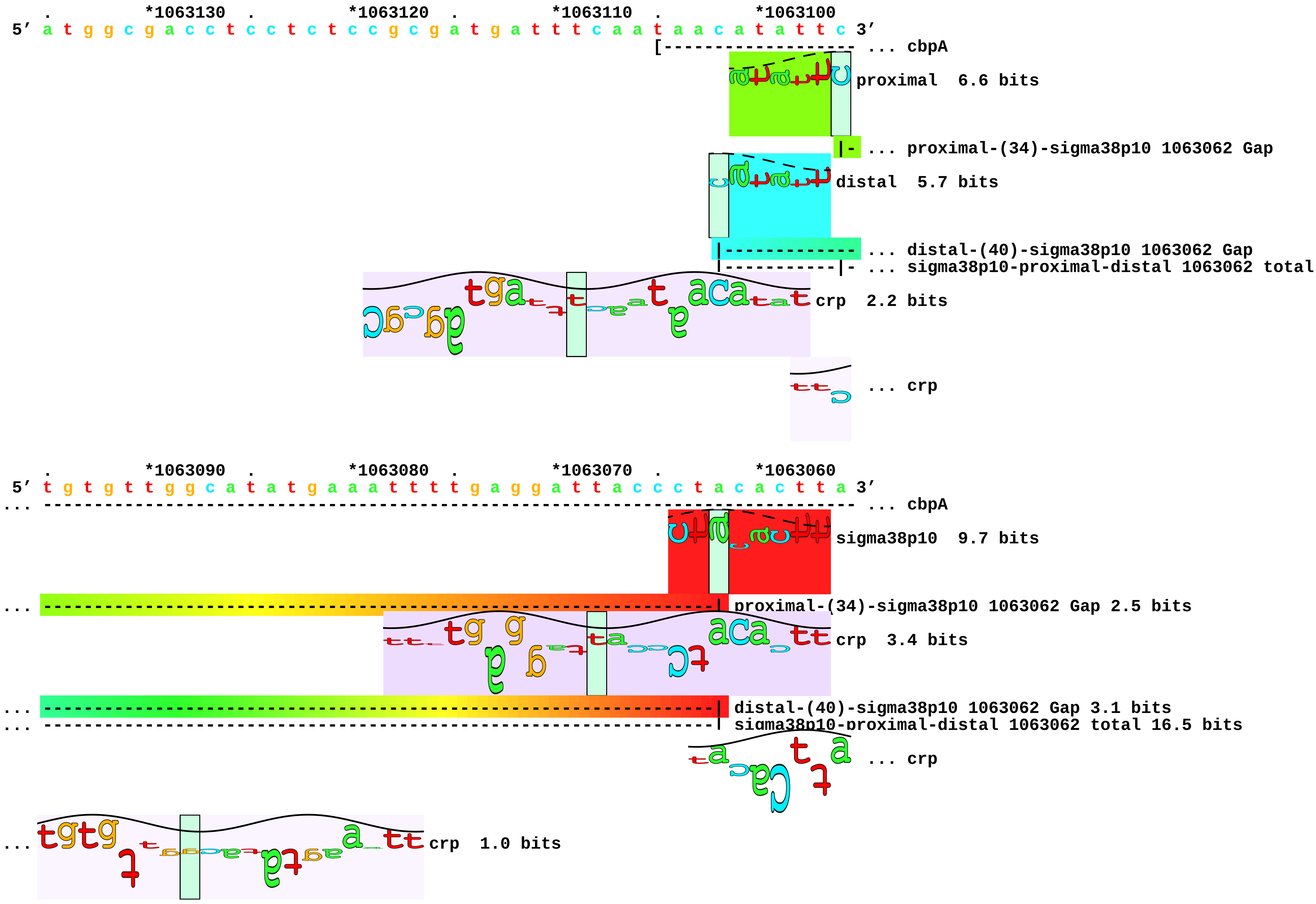

```
     .         *1063050  .         *1063040  .
5' t a g g a g t t a c c t t a c a g g g g t t c 3'
... ----] cbpA
```

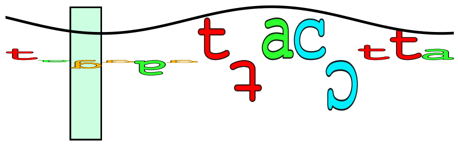

... crp 0.1 bits

piece 10, NC_000913.2, csiD, config: linear, direction: +, begin: 2786864, end: 2786969

. *2786870 . *2786880 . *2786890 . *2786900
5′ t a c t a a t t t g t t g c t t t t g a t c a c a a t a a g a a a a c a a t a t 3′

proximal 3.2 bits [------------------- ... csiD

distal 4.2 bits

|------------------------------------- ... proximal-(37)-sigma38p10 2786919 Gap

|------------------------------------- ... distal-(53)-sigma38p10 2786919 Gap
|------------------------------------- ... distal-proximal-sigma38p10 2786919 total

crp 10.8 bits ... proximal

distal 3.4 bits

|--------------------- ... distal-(41)-sigma38p10 2786934 Gap
|--------------------- ... distal-proximal-sigma38p10 2786934 total

distal 5.7 bits ... crp

|--------------------------------- ... distal-(49)-sigma38p10 2786936 Gap
|--------------------------------- ... distal-proximal-sigma38p10 2786936 total

distal 6.4 bits

|----------- ... distal-(40)-sigma38p10 2786938 Gap
|----------- ... distal-proximal-sigma38p10 2786938 total

crp 1.1 bits

. *2786910 . *2786920 . *2786930 . *2786940
5′ g t c g c t t t t g t g c g c a t t t t t c a g a a a t g t a g a t a t t t t t 3′
... ------------------------------------------------------------------------------ ... csiD
... ------------------------------| proximal-(37)-sigma38p10 2786919 Gap 4.5 bits

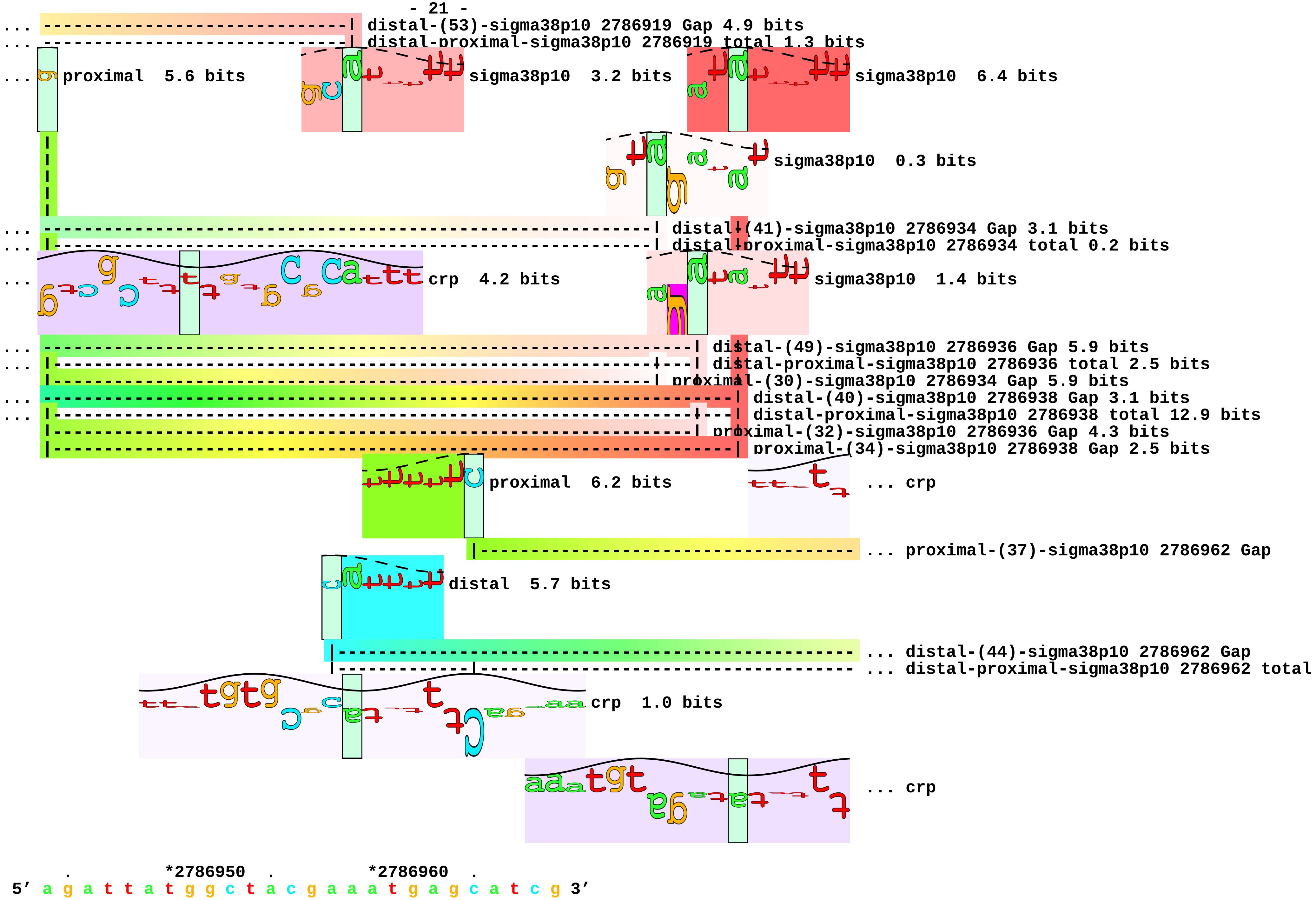
distal-(53)-sigma38p10 2786919 Gap 4.9 bits
distal-proximal-sigma38p10 2786919 total 1.3 bits
proximal 5.6 bits
sigma38p10 3.2 bits
sigma38p10 6.4 bits
sigma38p10 0.3 bits
distal-(41)-sigma38p10 2786934 Gap 3.1 bits
distal-proximal-sigma38p10 2786934 total 0.2 bits
crp 4.2 bits
sigma38p10 1.4 bits
distal-(49)-sigma38p10 2786936 Gap 5.9 bits
distal-proximal-sigma38p10 2786936 total 2.5 bits
proximal-(30)-sigma38p10 2786934 Gap 5.9 bits
distal-(40)-sigma38p10 2786938 Gap 3.1 bits
distal-proximal-sigma38p10 2786938 total 12.9 bits
proximal-(32)-sigma38p10 2786936 Gap 4.3 bits
proximal-(34)-sigma38p10 2786938 Gap 2.5 bits
proximal 6.2 bits
... crp
... proximal-(37)-sigma38p10 2786962 Gap
distal 5.7 bits
... distal-(44)-sigma38p10 2786962 Gap
... distal-proximal-sigma38p10 2786962 total
crp 1.0 bits
... crp
*2786950
*2786960
5′ a g a t t a t g g c t a c g a a a t g a g c a t c g 3′

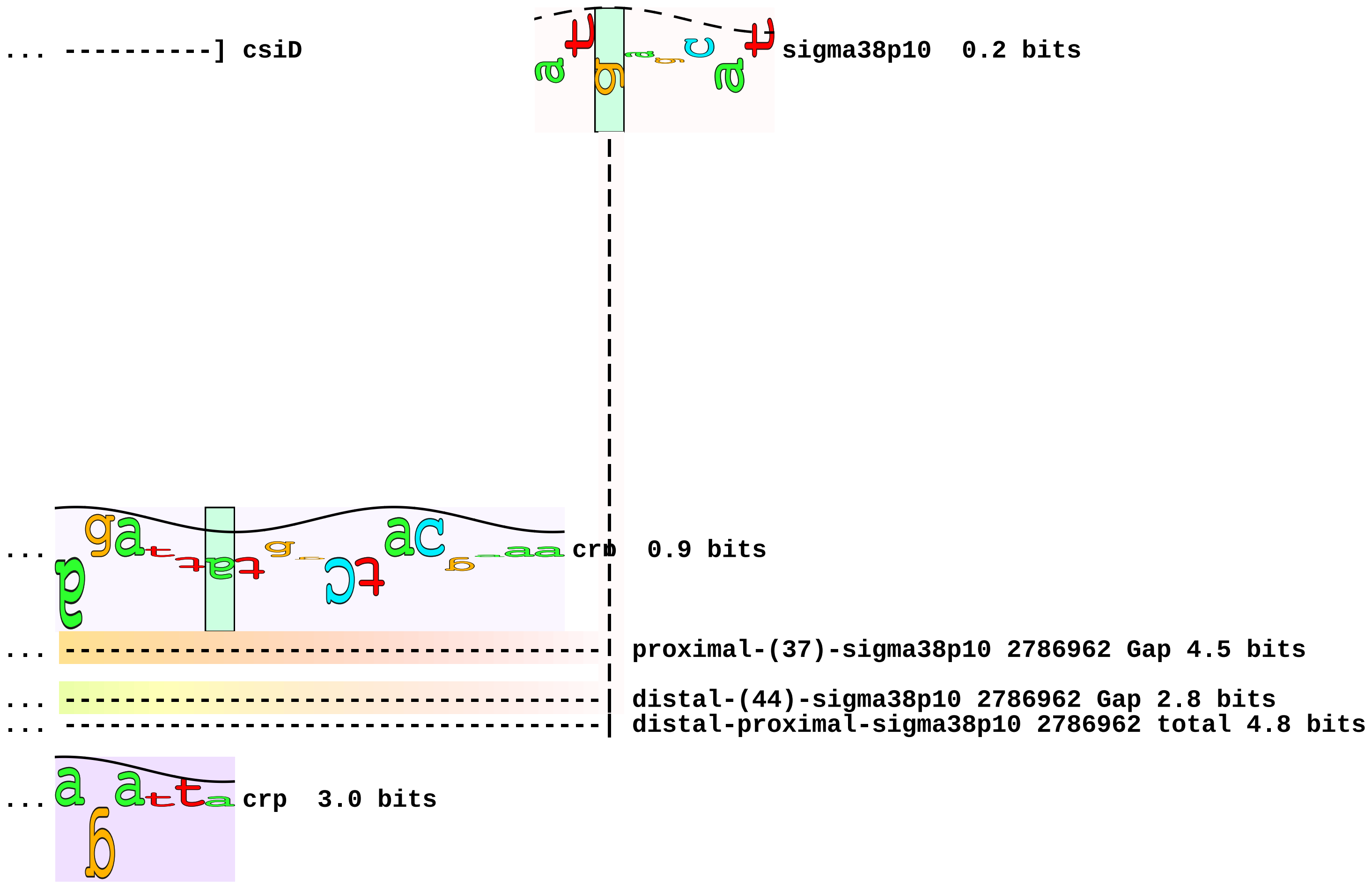

... ----------] csiD
sigma38p10 0.2 bits
crp 0.9 bits
proximal-(37)-sigma38p10 2786962 Gap 4.5 bits
distal-(44)-sigma38p10 2786962 Gap 2.8 bits
distal-proximal-sigma38p10 2786962 total 4.8 bits
crp 3.0 bits

piece 11, NC_000913.2, csiE, config: linear, direction: +, begin: 2663338, end: 2663443

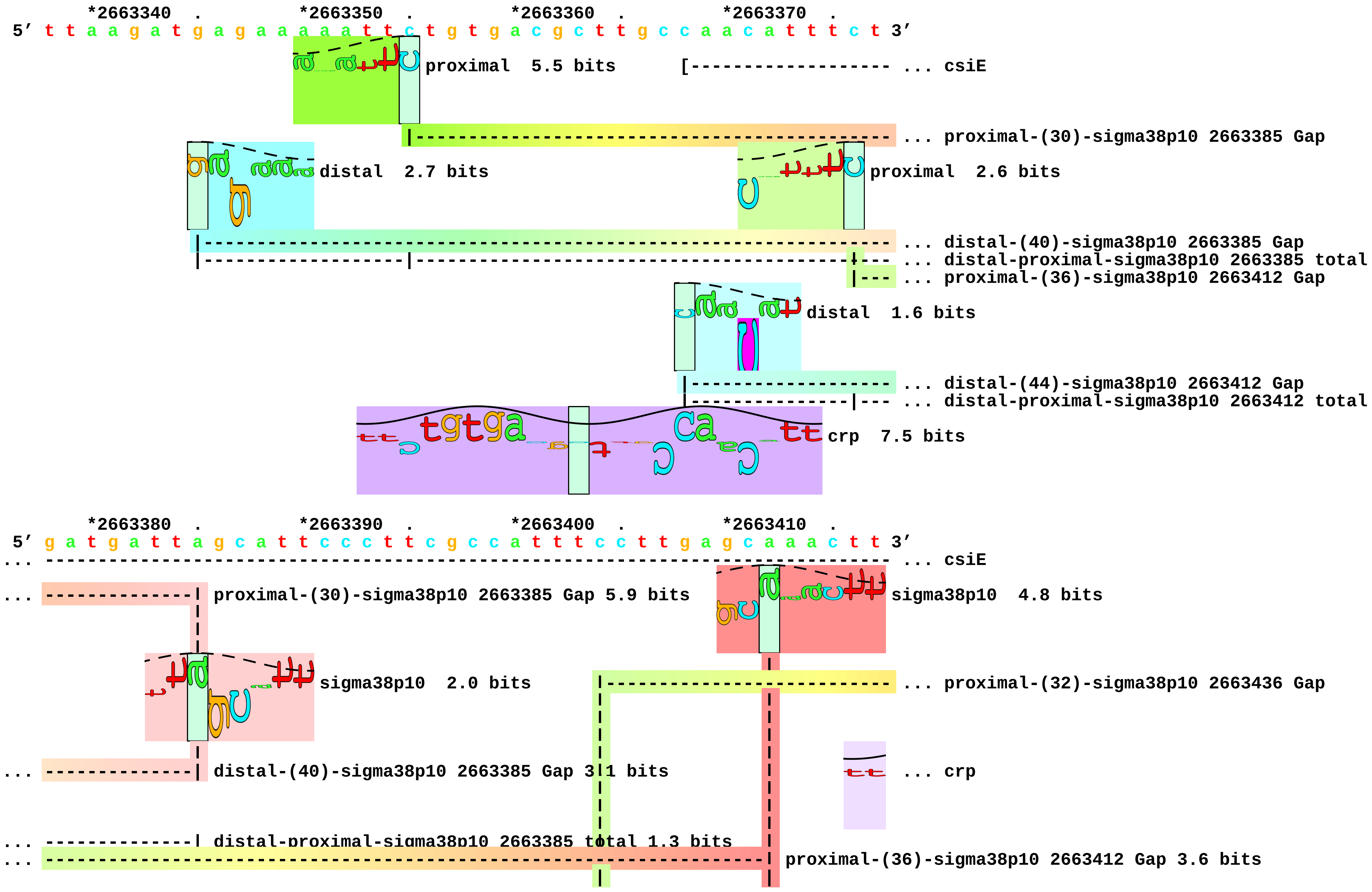

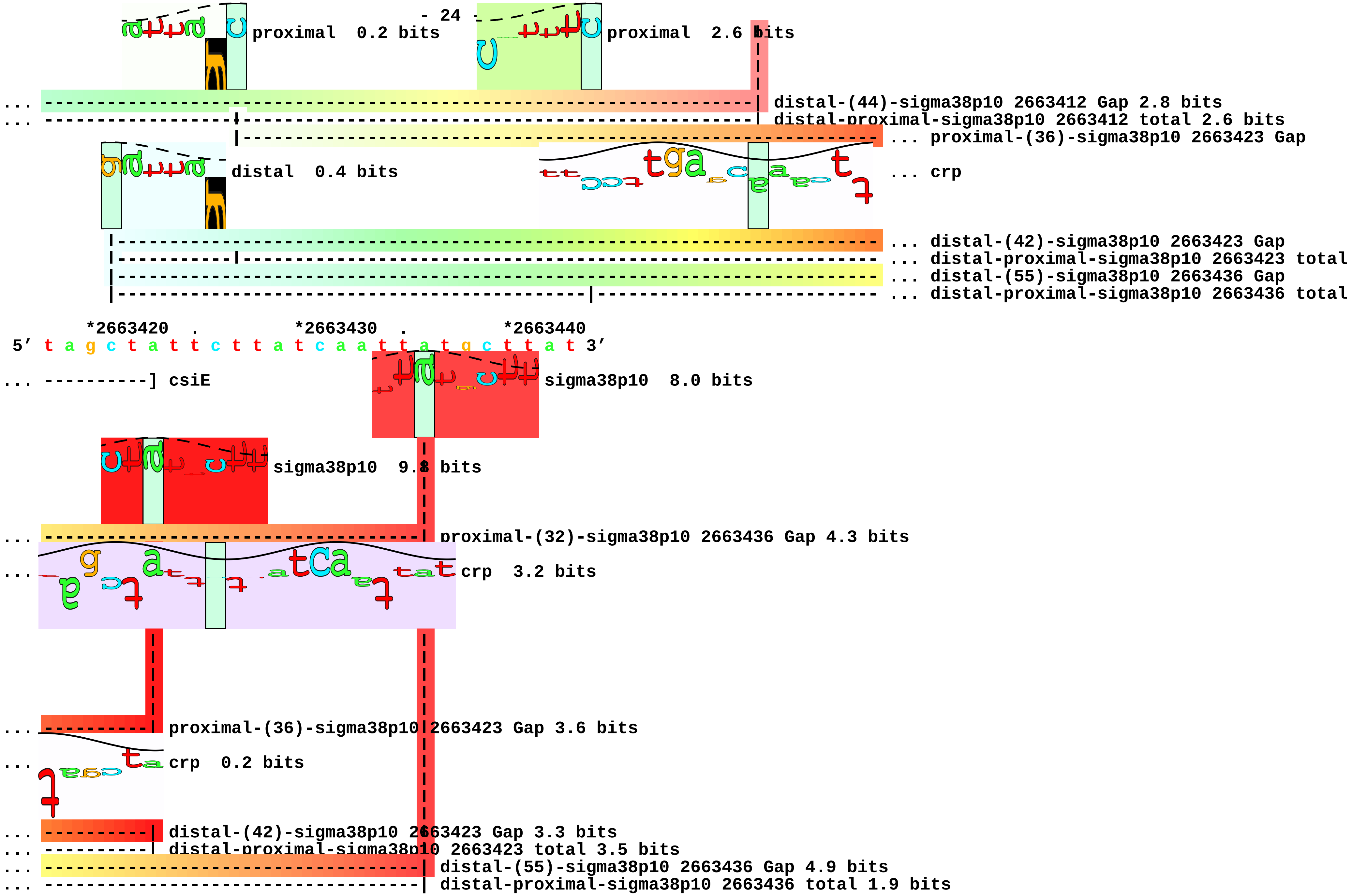
- 24
proximal 0.2 bits
proximal 2.6 bits
... distal-(44)-sigma38p10 2663412 Gap 2.8 bits
... distal-proximal-sigma38p10 2663412 total 2.6 bits
... proximal-(36)-sigma38p10 2663423 Gap
distal 0.4 bits
... crp
... distal-(42)-sigma38p10 2663423 Gap
... distal-proximal-sigma38p10 2663423 total
... distal-(55)-sigma38p10 2663436 Gap
... distal-proximal-sigma38p10 2663436 total
*2663420 . *2663430 . *2663440
5′ t a g c t a t t c t t a t c a a t t a t g c t t a t 3′
... ----------] csiE
sigma38p10 8.0 bits
sigma38p10 9.8 bits
... proximal-(32)-sigma38p10 2663436 Gap 4.3 bits
... crp 3.2 bits
... proximal-(36)-sigma38p10 2663423 Gap 3.6 bits
... crp 0.2 bits
... distal-(42)-sigma38p10 2663423 Gap 3.3 bits
... distal-proximal-sigma38p10 2663423 total 3.5 bits
... distal-(55)-sigma38p10 2663436 Gap 4.9 bits
... distal-proximal-sigma38p10 2663436 total 1.9 bits

piece 12, NC_000913.2, dnaN-P1, config: linear, direction: -, begin: 3880843, end: 3880736

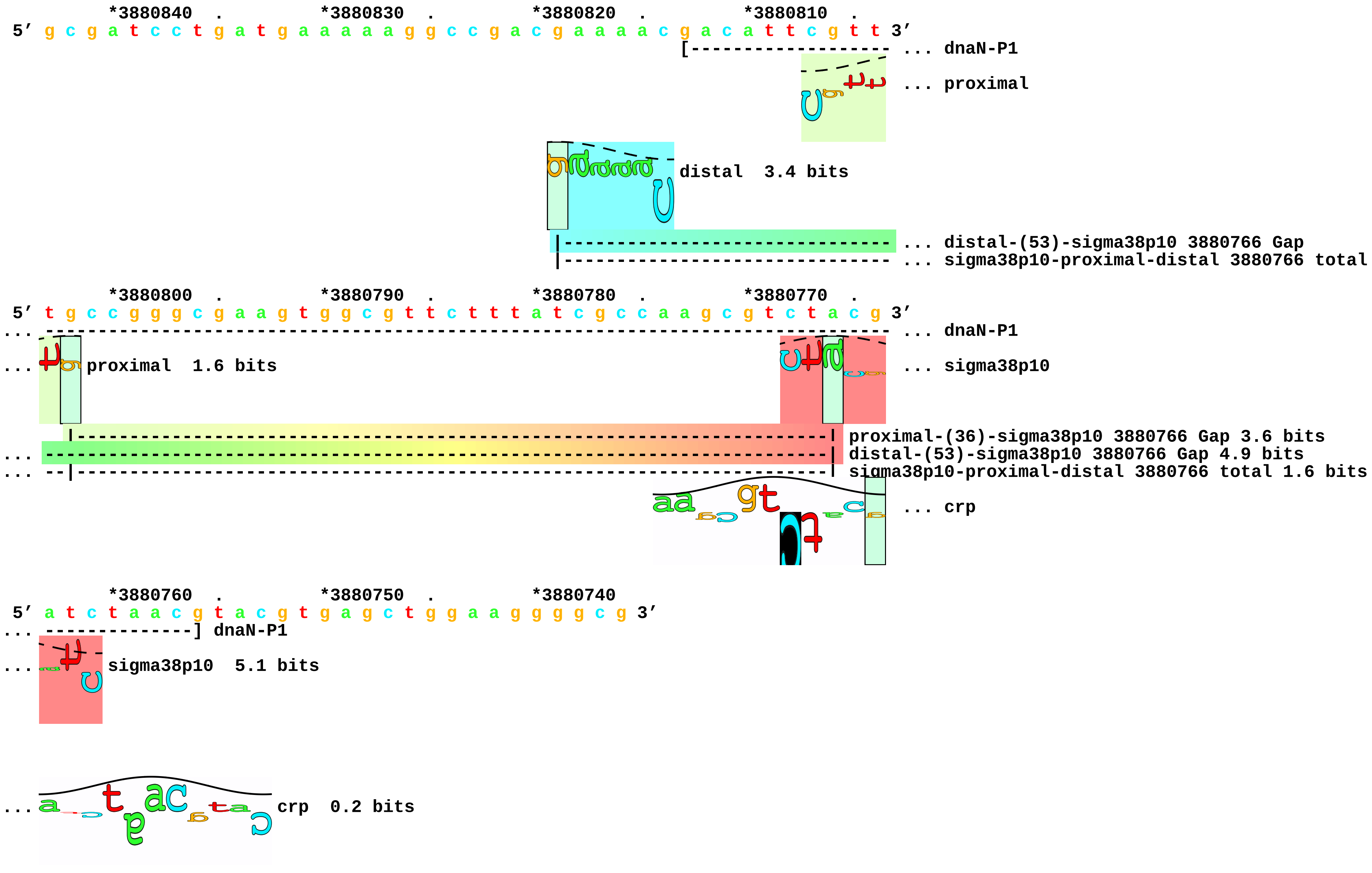

piece 13, NC_000913.2, frdA, config: linear, direction: -, begin: 4380525, end: 4380420

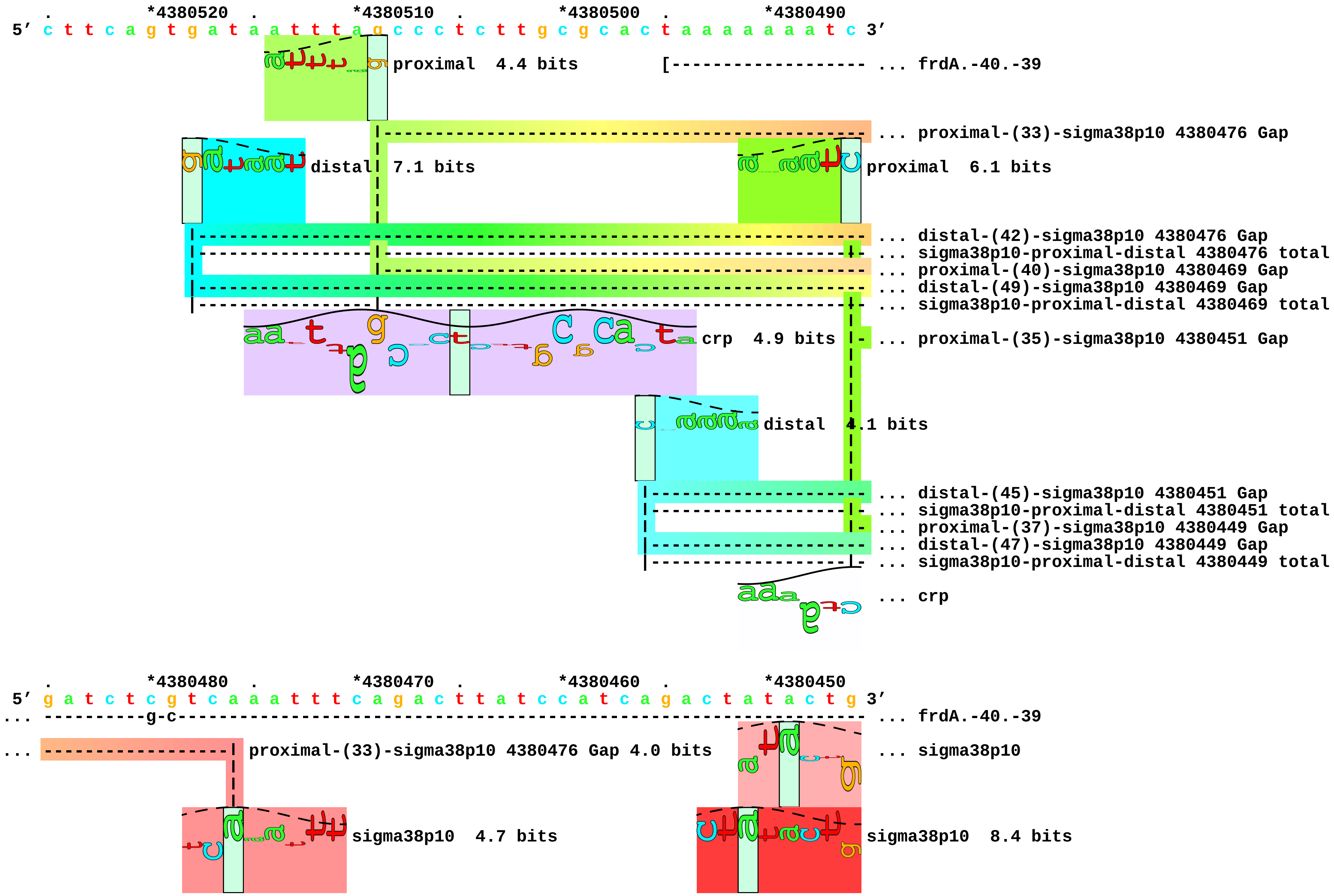

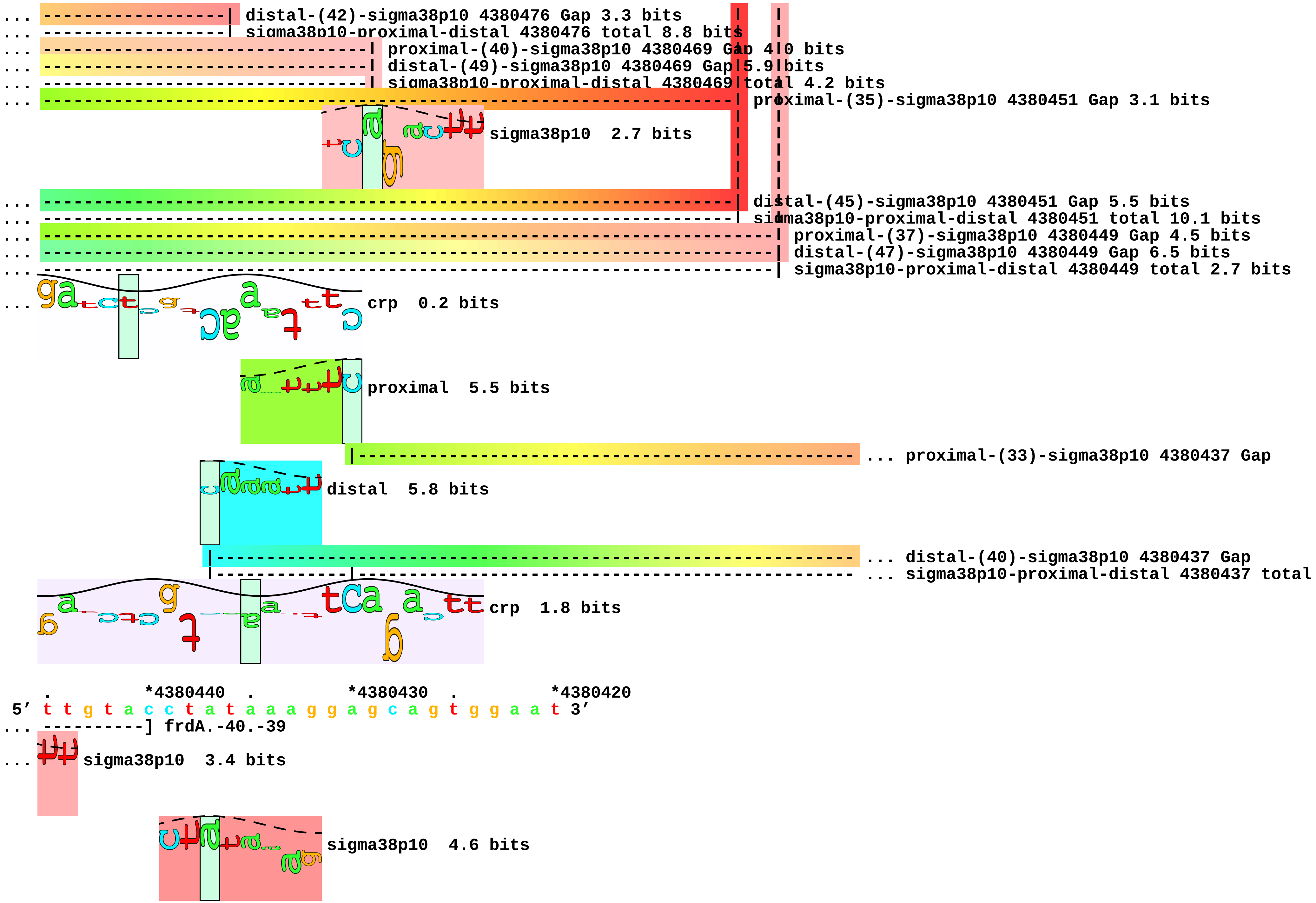
distal-(42)-sigma38p10 4380476 Gap 3.3 bits
sigma38p10-proximal-distal 4380476 total 8.8 bits
proximal-(40)-sigma38p10 4380469 Gap 4.0 bits
distal-(49)-sigma38p10 4380469 Gap 5.9 bits
sigma38p10-proximal-distal 4380469 total 4.2 bits
proximal-(35)-sigma38p10 4380451 Gap 3.1 bits
sigma38p10 2.7 bits
distal-(45)-sigma38p10 4380451 Gap 5.5 bits
sigma38p10-proximal-distal 4380451 total 10.1 bits
proximal-(37)-sigma38p10 4380449 Gap 4.5 bits
distal-(47)-sigma38p10 4380449 Gap 6.5 bits
sigma38p10-proximal-distal 4380449 total 2.7 bits
crp 0.2 bits
proximal 5.5 bits
proximal-(33)-sigma38p10 4380437 Gap
distal 5.8 bits
distal-(40)-sigma38p10 4380437 Gap
sigma38p10-proximal-distal 4380437 total
crp 1.8 bits
*4380440
*4380430
*4380420
5′ t t g t a c c t a t a a a g g a g c a g t g g a a t 3′
] frdA.-40.-39
sigma38p10 3.4 bits
sigma38p10 4.6 bits

```
...  ----------------|  proximal-(33)-sigma38p10 4380437 Gap 4.0 bits
...  ----------------|  distal-(40)-sigma38p10 4380437 Gap 3.1 bits
...  ----------------|  sigma38p10-proximal-distal 4380437 total 8.8 bits
```

piece 14, NC_000913.2, ftsQ-P1, config: linear, direction: +, begin: 102787, end: 102891

```
         *102790    .          *102800    .          *102810    .          *102820    .
5' g c c g g a g t t c a c g g t t g c g a t a c t c g g t g a a g a a a t t t t a 3'
                                                              [------------------- ... ftsQ-P1.-1
                                                                                   ... proximal
                                                                          distal  6.6 bits
                                                                |----------------- ... distal-(43)-sigma38p10 102861 Gap
                                                                |----------------- ... distal-proximal-sigma38p10 102861 total 1
                                                                                   ... crp

         *102830    .          *102840    .          *102850    .          *102860    .
5' c c g t c a a t a c g t a t t c a a c c g t c c g g a a c c t t c t a t g a t t 3'
... ---------------------------------------------------------------------------- ... ftsQ-P1.-1
... proximal  4.4 bits                                                    sigma38p10  9.1 bits
   |------------------------------------------------------------------| proximal-(34)-sigma38p10 102861 Gap 2.5 bits
... ------------------------------------------------------------------| distal-(43)-sigma38p10 102861 Gap 3.6 bits
... |-----------------------------------------------------------------| distal-proximal-sigma38p10 102861 total 14.1 bits
...                           crp  4.7 bits

         *102870    .          *102880    .          *102890
5' a t g a g g c g a a g t a t c t c t c t g a t g a 3'
... ------t-] ftsQ-P1.-1
                                            crp  2.6 bits
```

piece 15, NC_000913.2, gadX, config: linear, direction: -, begin: 3663950, end: 3663843

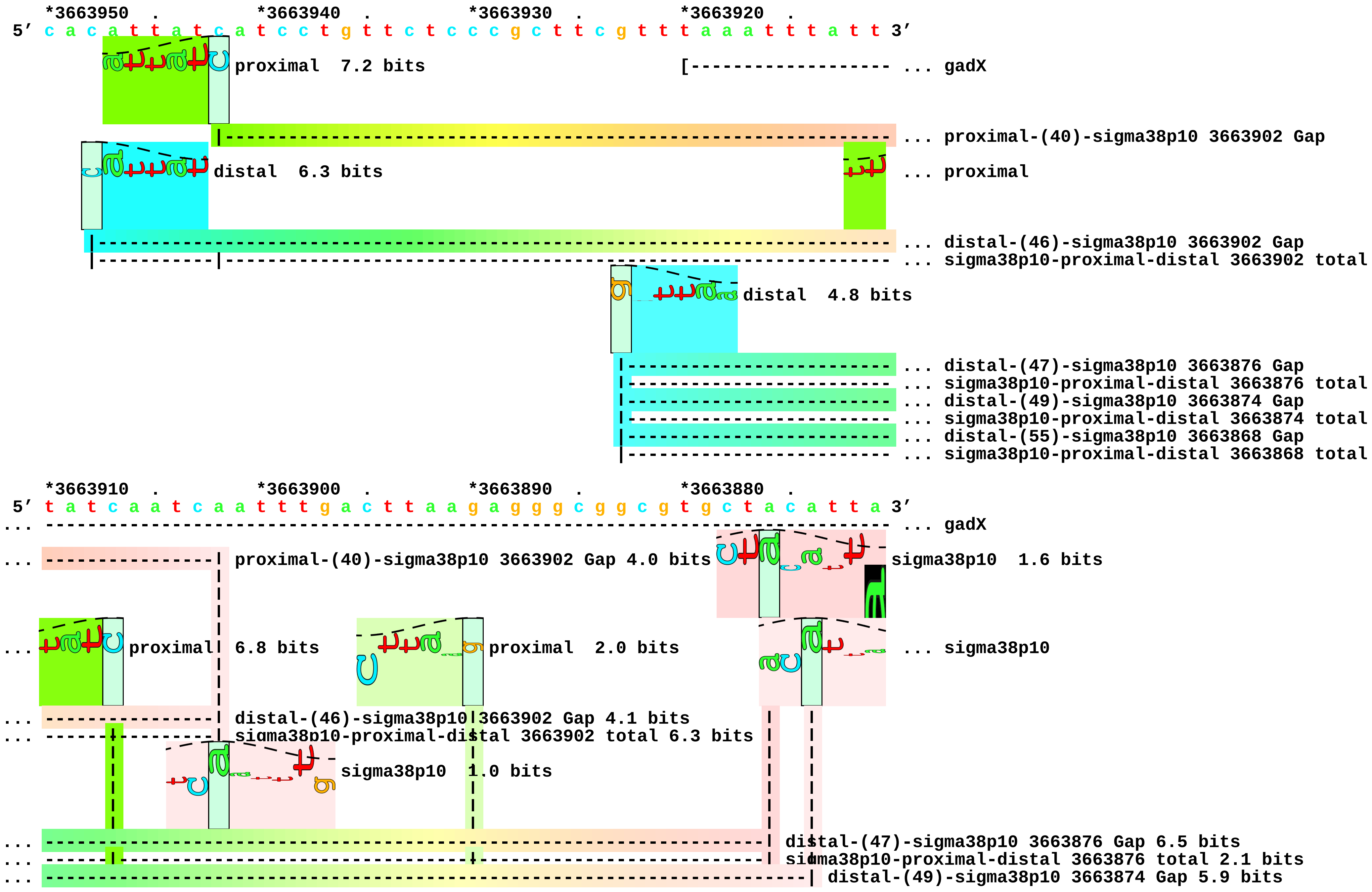

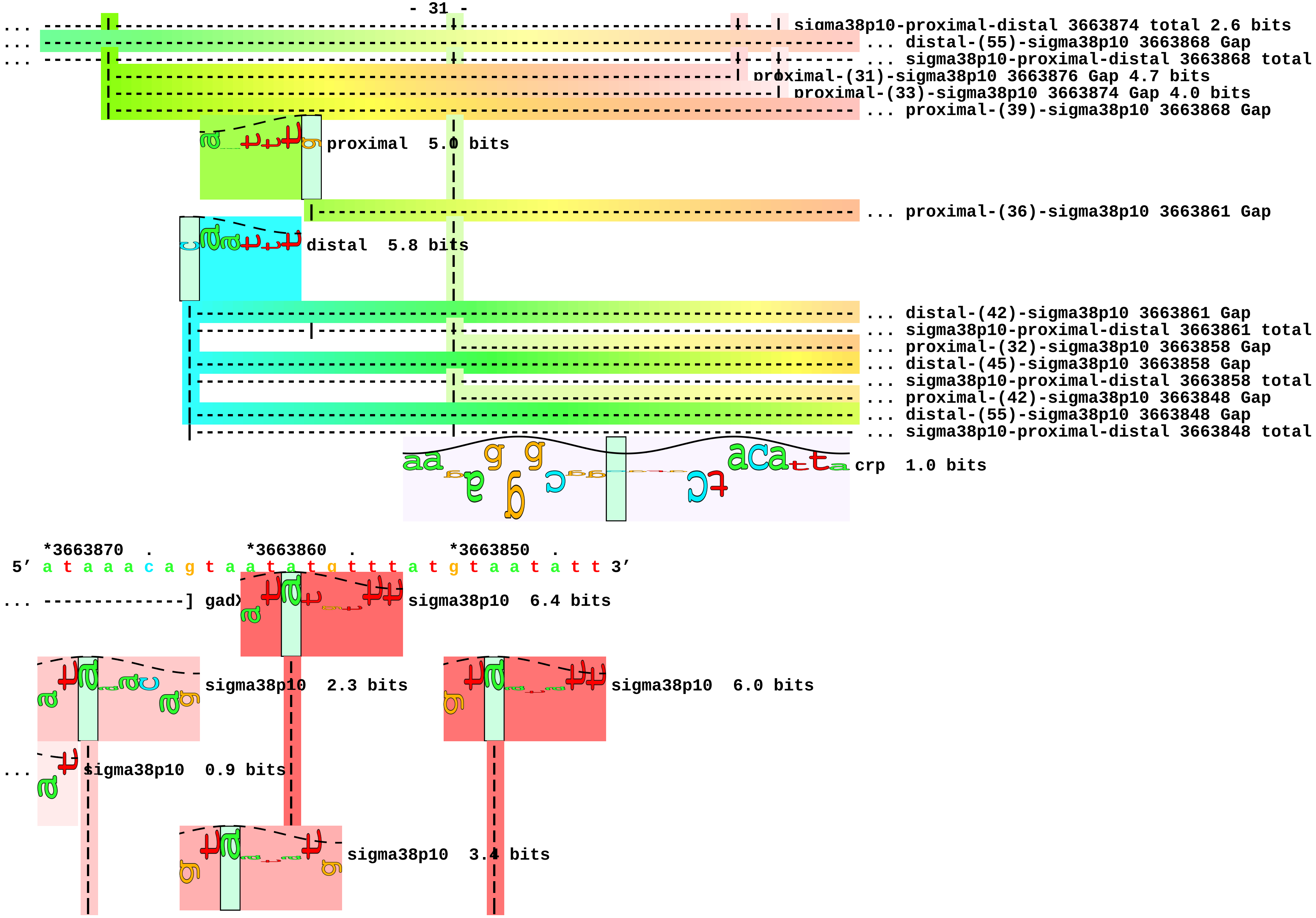
sigma38p10-proximal-distal 3663874 total 2.6 bits
distal-(55)-sigma38p10 3663868 Gap
sigma38p10-proximal-distal 3663868 total
proximal-(31)-sigma38p10 3663876 Gap 4.7 bits
proximal-(33)-sigma38p10 3663874 Gap 4.0 bits
proximal-(39)-sigma38p10 3663868 Gap
proximal 5.0 bits
proximal-(36)-sigma38p10 3663861 Gap
distal 5.8 bits
distal-(42)-sigma38p10 3663861 Gap
sigma38p10-proximal-distal 3663861 total
proximal-(32)-sigma38p10 3663858 Gap
distal-(45)-sigma38p10 3663858 Gap
sigma38p10-proximal-distal 3663858 total
proximal-(42)-sigma38p10 3663848 Gap
distal-(55)-sigma38p10 3663848 Gap
sigma38p10-proximal-distal 3663848 total
crp 1.0 bits
*3663870 . *3663860 . *3663850 .
5′ a t a a a c a g t a a t a t g t t t a t g t a a t a t t 3′
gadX
sigma38p10 6.4 bits
sigma38p10 2.3 bits
sigma38p10 6.0 bits
sigma38p10 0.9 bits
sigma38p10 3.4 bits

```
... ----| distal-(55)-sigma38p10 3663868 Gap 4.9 bits
... ----| sigma38p10-proximal-distal 3663868 total 1.5 bits
... ----| proximal-(39)-sigma38p10 3663868 Gap 7.5 bits
... ------------------| proximal-(36)-sigma38p10 3663861 Gap 3.6 bits
... ------------------| distal-(42)-sigma38p10 3663861 Gap 3.3 bits
... ------------------| sigma38p10-proximal-distal 3663861 total 7.3 bits
... ------------------------| proximal-(32)-sigma38p10 3663858 Gap 4.3 bits
... ------------------------| distal-(45)-sigma38p10 3663858 Gap 5.5 bits
... ------------------------| sigma38p10-proximal-distal 3663858 total 4.4 bits
... --------------------------------------------| proximal-(42)-sigma38p10 3663848 Gap 5.1 bits
... --------------------------------------------| distal-(55)-sigma38p10 3663848 Gap 4.9 bits
... --------------------------------------------| sigma38p10-proximal-distal 3663848 total 3.7 bits
```

piece 16, NC_000913.2, htrE-P2, config: linear, direction: -, begin: 156309, end: 156204

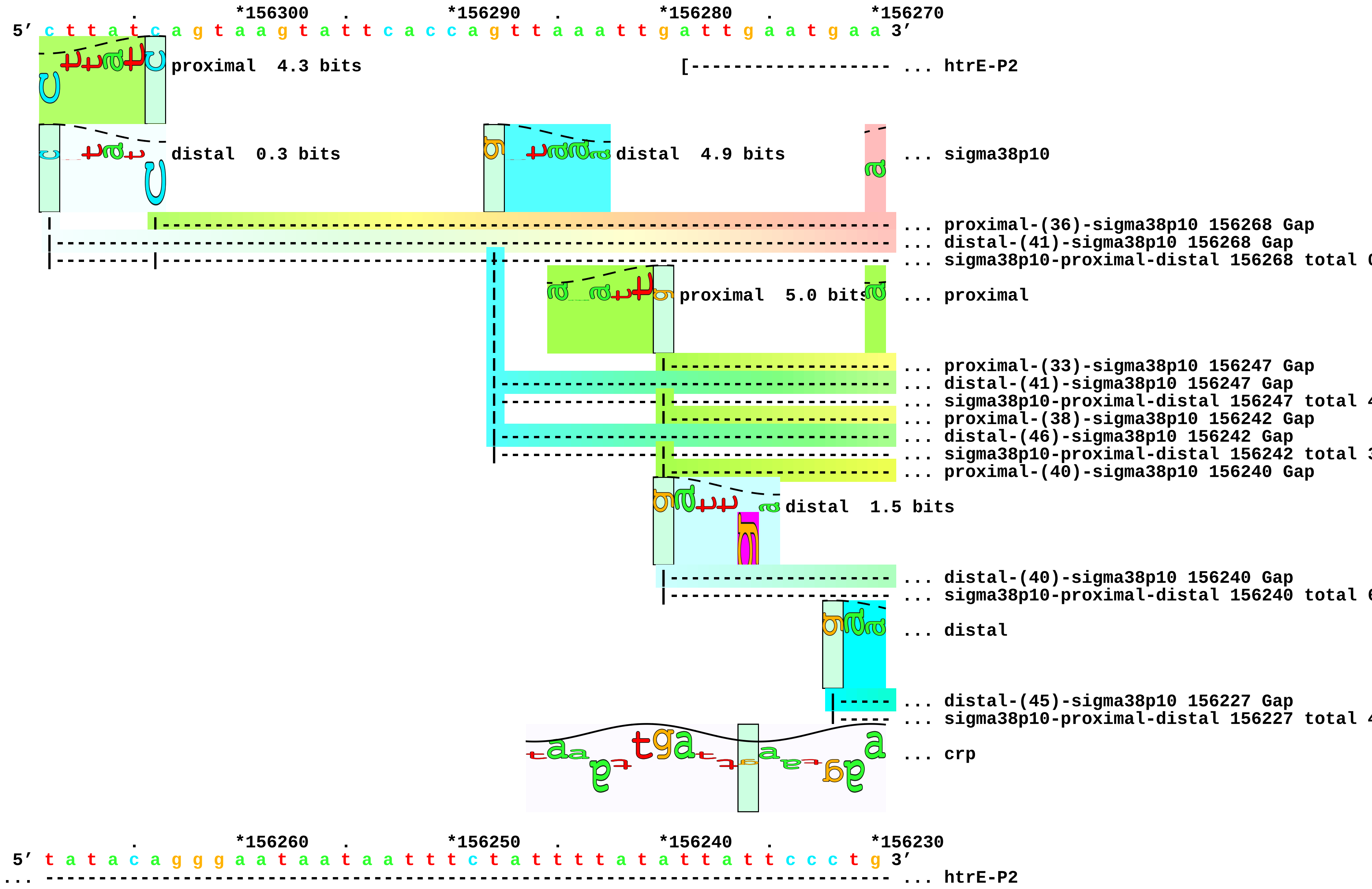

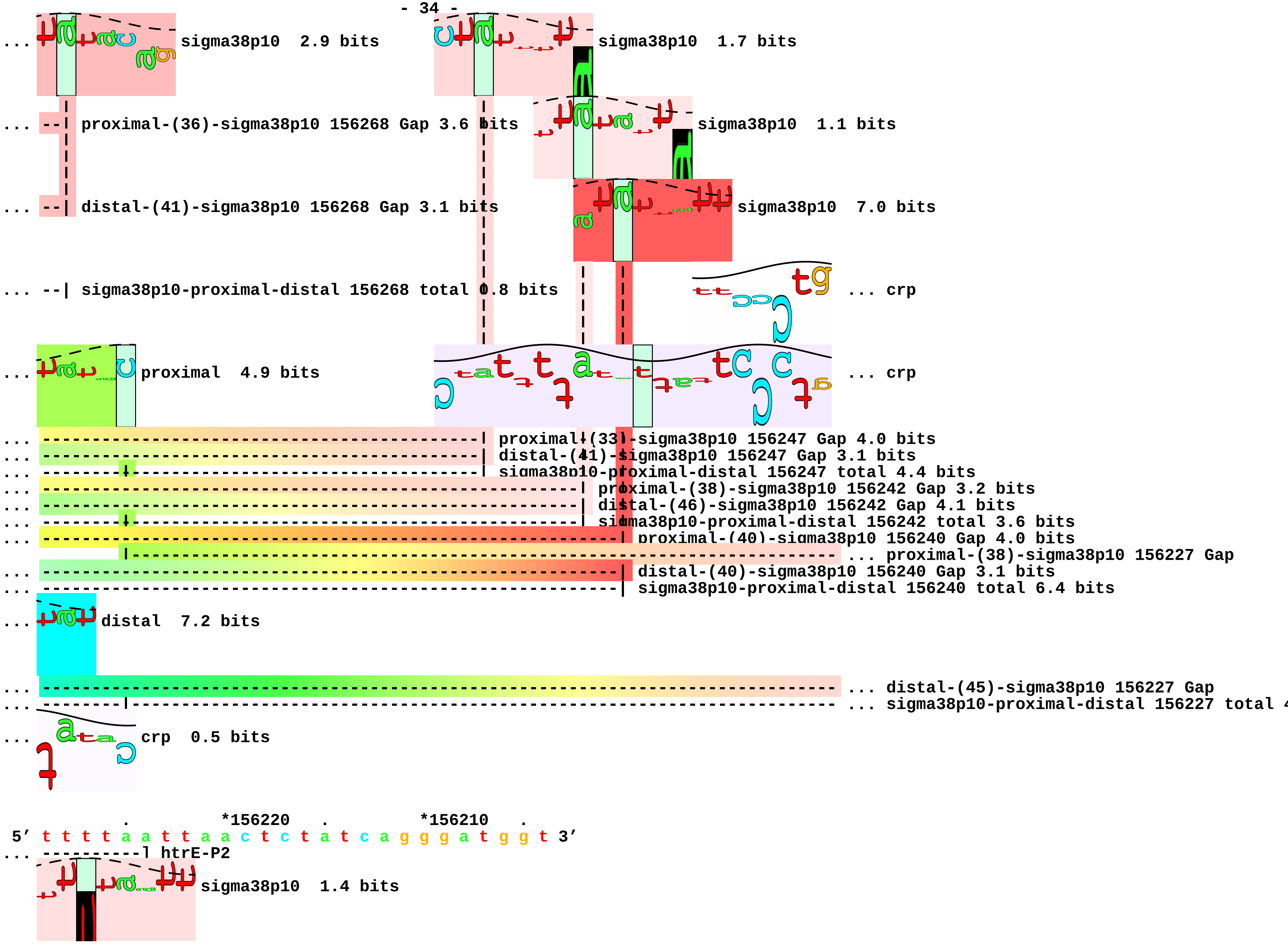
sigma38p10 2.9 bits
sigma38p10 1.7 bits
proximal-(36)-sigma38p10 156268 Gap 3.6 bits
sigma38p10 1.1 bits
distal-(41)-sigma38p10 156268 Gap 3.1 bits
sigma38p10 7.0 bits
sigma38p10-proximal-distal 156268 total 0.8 bits
... crp
proximal 4.9 bits
... crp
proximal-(33)-sigma38p10 156247 Gap 4.0 bits
distal-(41)-sigma38p10 156247 Gap 3.1 bits
sigma38p10-proximal-distal 156247 total 4.4 bits
proximal-(38)-sigma38p10 156242 Gap 3.2 bits
distal-(46)-sigma38p10 156242 Gap 4.1 bits
sigma38p10-proximal-distal 156242 total 3.6 bits
proximal-(40)-sigma38p10 156240 Gap 4.0 bits
... proximal-(38)-sigma38p10 156227 Gap
distal-(40)-sigma38p10 156240 Gap 3.1 bits
sigma38p10-proximal-distal 156240 total 6.4 bits
distal 7.2 bits
... distal-(45)-sigma38p10 156227 Gap
... sigma38p10-proximal-distal 156227 total
crp 0.5 bits
*156220
*156210
5′ t t t t a a t t a a c t c t a t c a g g g a t g g t 3′
htrE-P2
sigma38p10 1.4 bits

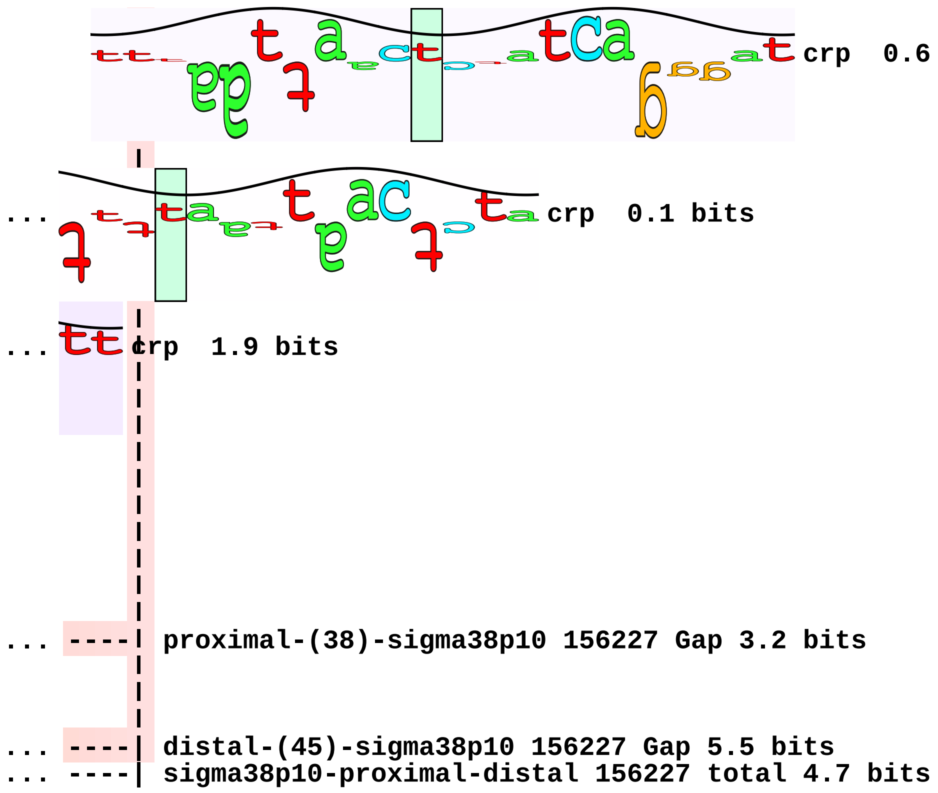
crp 0.6 bits
... crp 0.1 bits
... crp 1.9 bits
... ----| proximal-(38)-sigma38p10 156227 Gap 3.2 bits
... ----| distal-(45)-sigma38p10 156227 Gap 5.5 bits
... ----| sigma38p10-proximal-distal 156227 total 4.7 bits

piece 17, NC_000913.2, mscL, config: linear, direction: +, begin: 3435936, end: 3436042

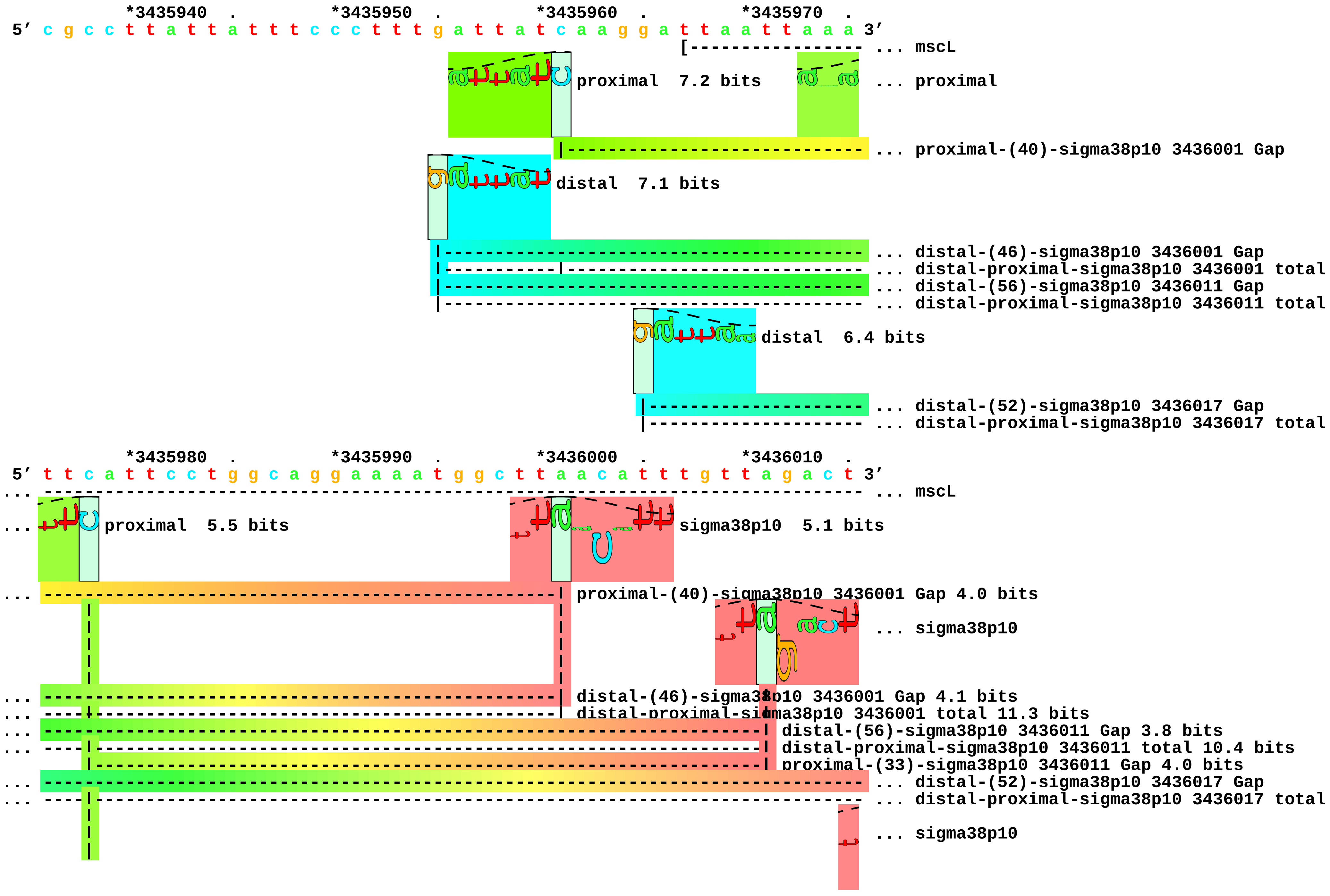

```
                                                                          ... proximal-(39)-sigma38p10 3436017 Gap
                                                                          ... crp

        *3436020  .        *3436030  .        *3436040
5' t a t g g t t g t c g g c t t c a t a g g g a g a a t 3'
... ------------#-] mscL

... sigma38p10  5.5 bits

... --| distal-(52)-sigma38p10 3436017 Gap 5.5 bits
... --| distal-proximal-sigma38p10 3436017 total 4.1 bits
...          sigma38p10  5.1 bits

... --| proximal-(39)-sigma38p10 3436017 Gap 7.5 bits

... crp  0.1 bits
```

piece 18, NC_000913.2, mscS, config: linear, direction: -, begin: 3067979, end: 3067873

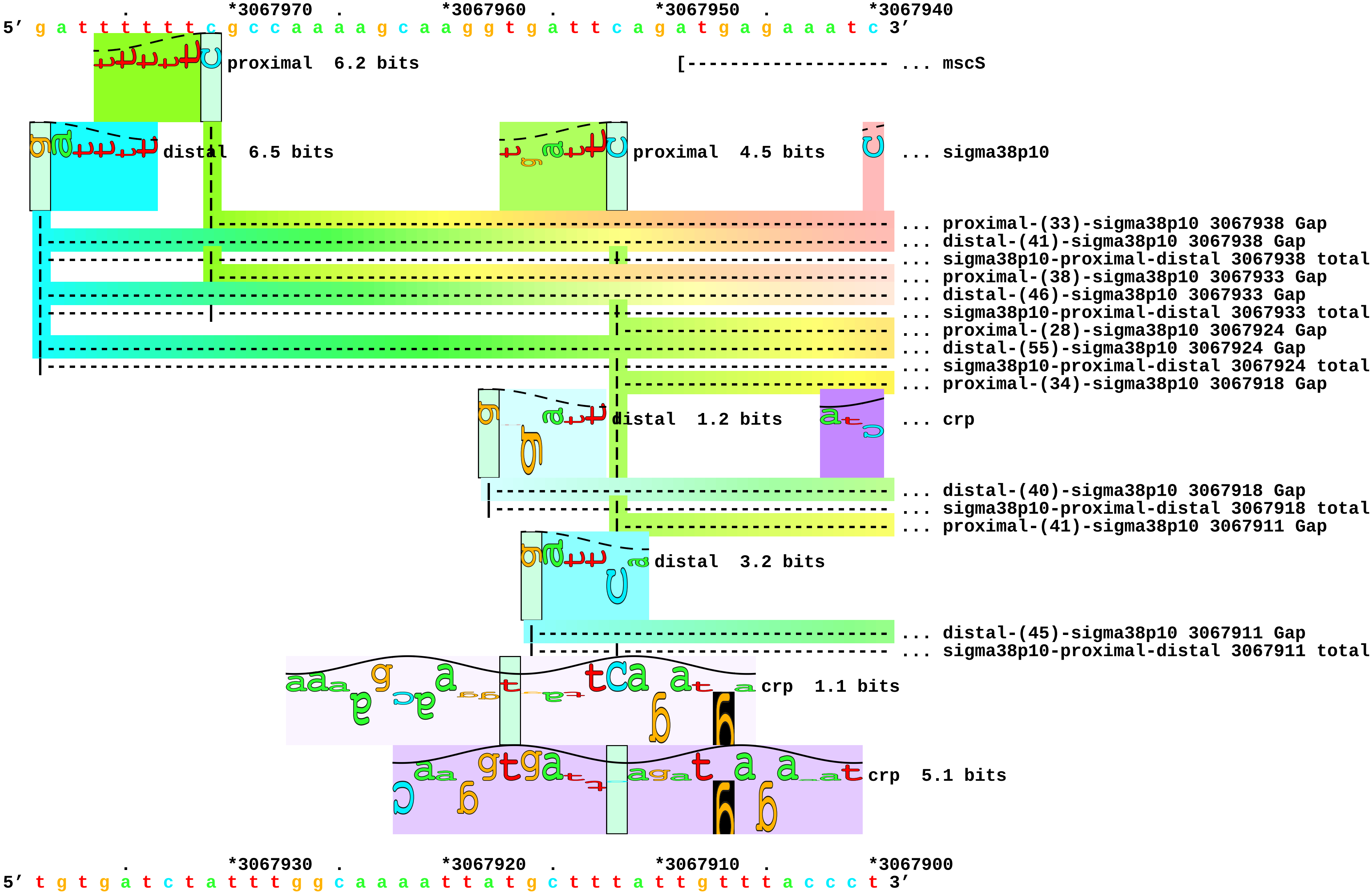

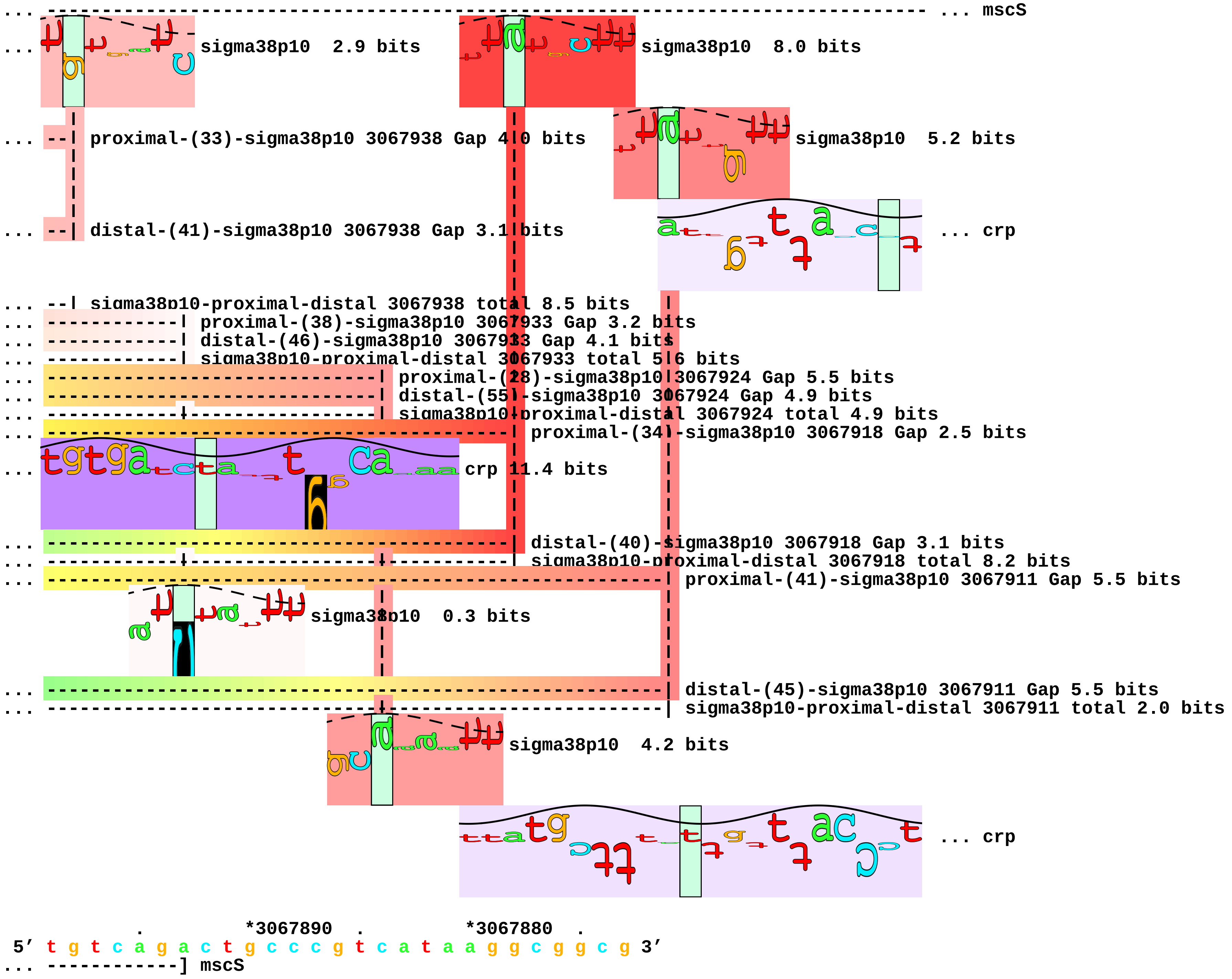
... mscS
sigma38p10 2.9 bits
sigma38p10 8.0 bits
proximal-(33)-sigma38p10 3067938 Gap 4.0 bits
sigma38p10 5.2 bits
distal-(41)-sigma38p10 3067938 Gap 3.1 bits
... crp
sigma38p10-proximal-distal 3067938 total 8.5 bits
proximal-(38)-sigma38p10 3067933 Gap 3.2 bits
distal-(46)-sigma38p10 3067933 Gap 4.1 bits
sigma38p10-proximal-distal 3067933 total 5.6 bits
proximal-(28)-sigma38p10 3067924 Gap 5.5 bits
distal-(55)-sigma38p10 3067924 Gap 4.9 bits
sigma38p10-proximal-distal 3067924 total 4.9 bits
proximal-(34)-sigma38p10 3067918 Gap 2.5 bits
crp 11.4 bits
distal-(40)-sigma38p10 3067918 Gap 3.1 bits
sigma38p10-proximal-distal 3067918 total 8.2 bits
proximal-(41)-sigma38p10 3067911 Gap 5.5 bits
sigma38p10 0.3 bits
distal-(45)-sigma38p10 3067911 Gap 5.5 bits
sigma38p10-proximal-distal 3067911 total 2.0 bits
sigma38p10 4.2 bits
... crp
*3067890
*3067880
5′ t g t c a g a c t g c c c g t c a t a a g g c g g c g 3′
... ------------] mscS

... 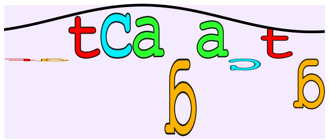 crp 1.9 bits

... 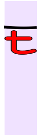 crp 2.8 bits piece 19, NC_000913.2, osmY, config: linear, direction: +, begin: 4609089, end: 4609196

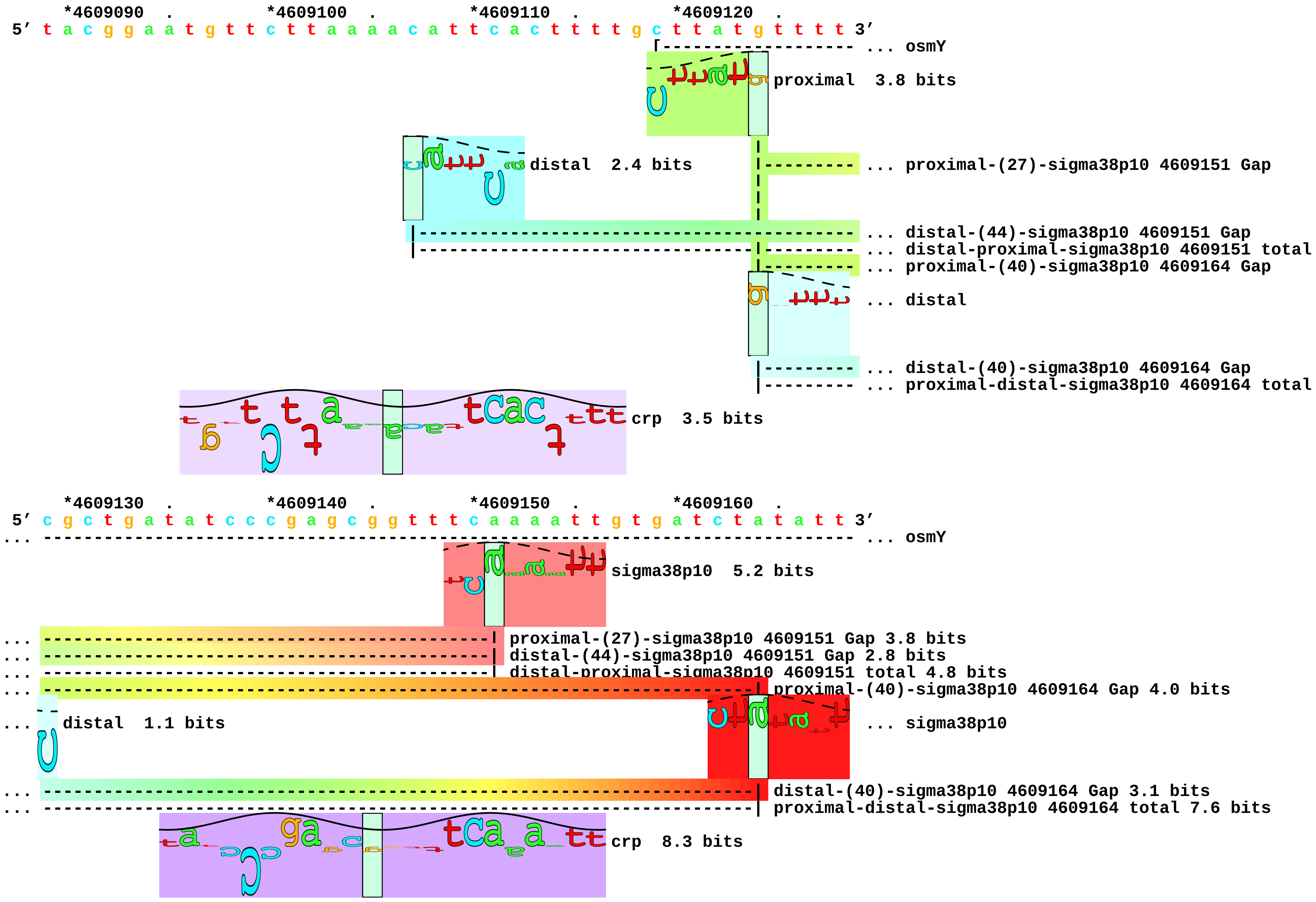

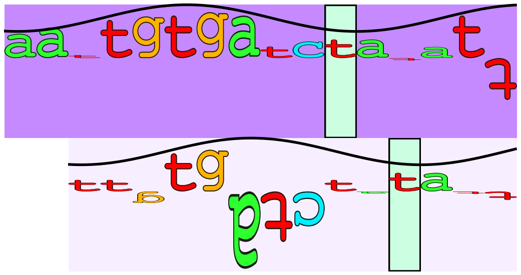

... crp

... crp

```
     *4609170  .         *4609180  .         *4609190  .
5' t a a c a a a g t g a t g a c a t t t c t g a c g g c g 3'
... --------------] osmY
```

... 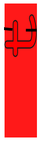 sigma38p10  9.9 bits

... 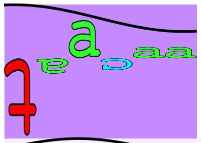 crp 11.2 bits

... 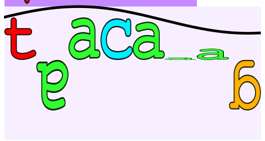 crp  1.6 bits piece 20, NC_000913.2, proP-P2, config: linear, direction: +, begin: 4328344, end: 4328450

```
        .         *4328350  .         *4328360  .         *4328370  .         *4328380
5' t c a t t a a c t g c c c a a t t c a g g c g t c a a c t g g t t t g a t t g t 3'
                                                          [-------------------- ... proP-P2
                                                                  proximal  4.0 bits
                                                                        |--- ... proximal-(37)-sigma38p10 4328419 Gap
                                                                  distal  2.1 bits
                                                                 |----------- ... distal-(41)-sigma38p10 4328419 Gap
                                                                 |--------|--- ... distal-proximal-sigma38p10 4328419 total

        .         *4328390  .         *4328400  .         *4328410  .         *4328420
5' a c a t t c c t t a a c c g g a g g g t g t a a g c a a a c c c g c t a c g c t 3'
... ---------------------------------------------------------------------------- ... proP-P2
                                        distal  1.1 bits                        ... sigma38p10
... ---------------------------------------------------------------------| proximal-(37)-sigma38p10 4328419 Gap 4.5 bits
                                        proximal  2.8 bits                      ... crp
... ---------------------------------------------------------------------| distal-(41)-sigma38p10 4328419 Gap 3.1 bits
... ---------------------------------------------------------------------| distal-proximal-sigma38p10 4328419 total 7.1 bits
                                                  |---------------------------- ... proximal-(36)-sigma38p10 4328444 Gap
                                        |-------------------------------------- ... distal-(42)-sigma38p10 4328444 Gap
                                        |----------|--------------------------- ... distal-proximal-sigma38p10 4328444 total

        .         *4328430  .         *4328440  .         *4328450
5' t g t t a c a g a g a t t g c a t c c t g c a a t t c 3'
... ------------] proP-P2
                                        sigma38p10  5.8 bits
... sigma38p10  8.5 bits
```

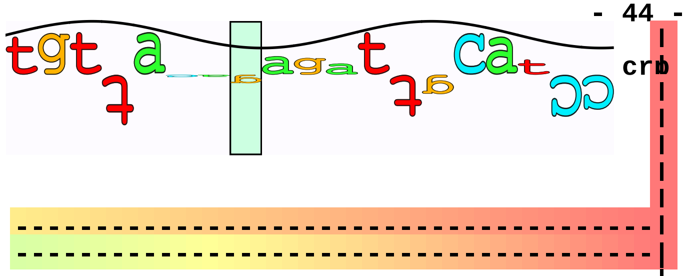


... proximal-(36)-sigma38p10 4328444 Gap 3.6 bits
... distal-(42)-sigma38p10 4328444 Gap 3.3 bits
... distal-proximal-sigma38p10 4328444 total 2.9 bits

piece 21, NC_000913.2, uspB, config: linear, direction: -, begin: 3637955, end: 3637851

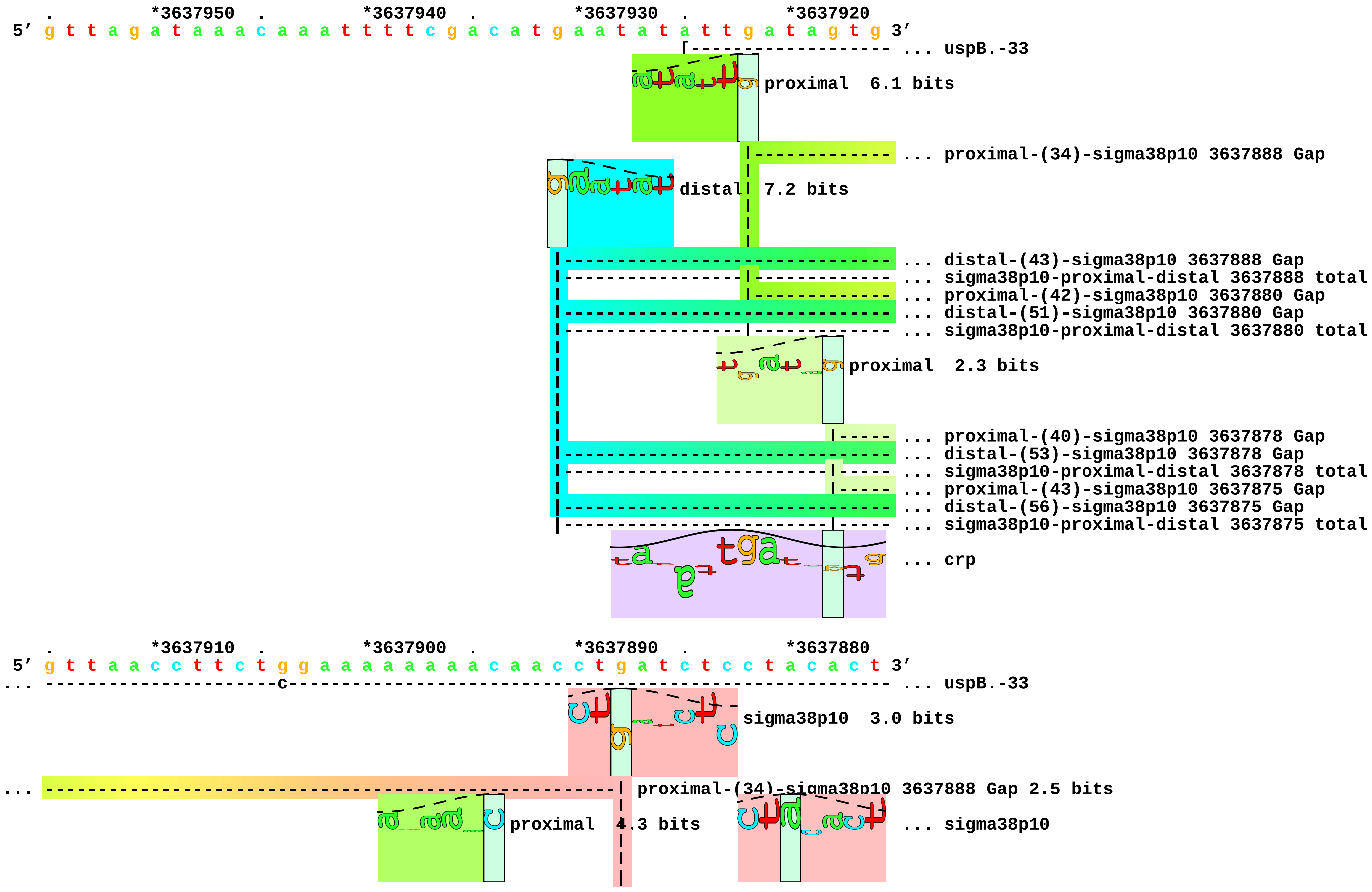

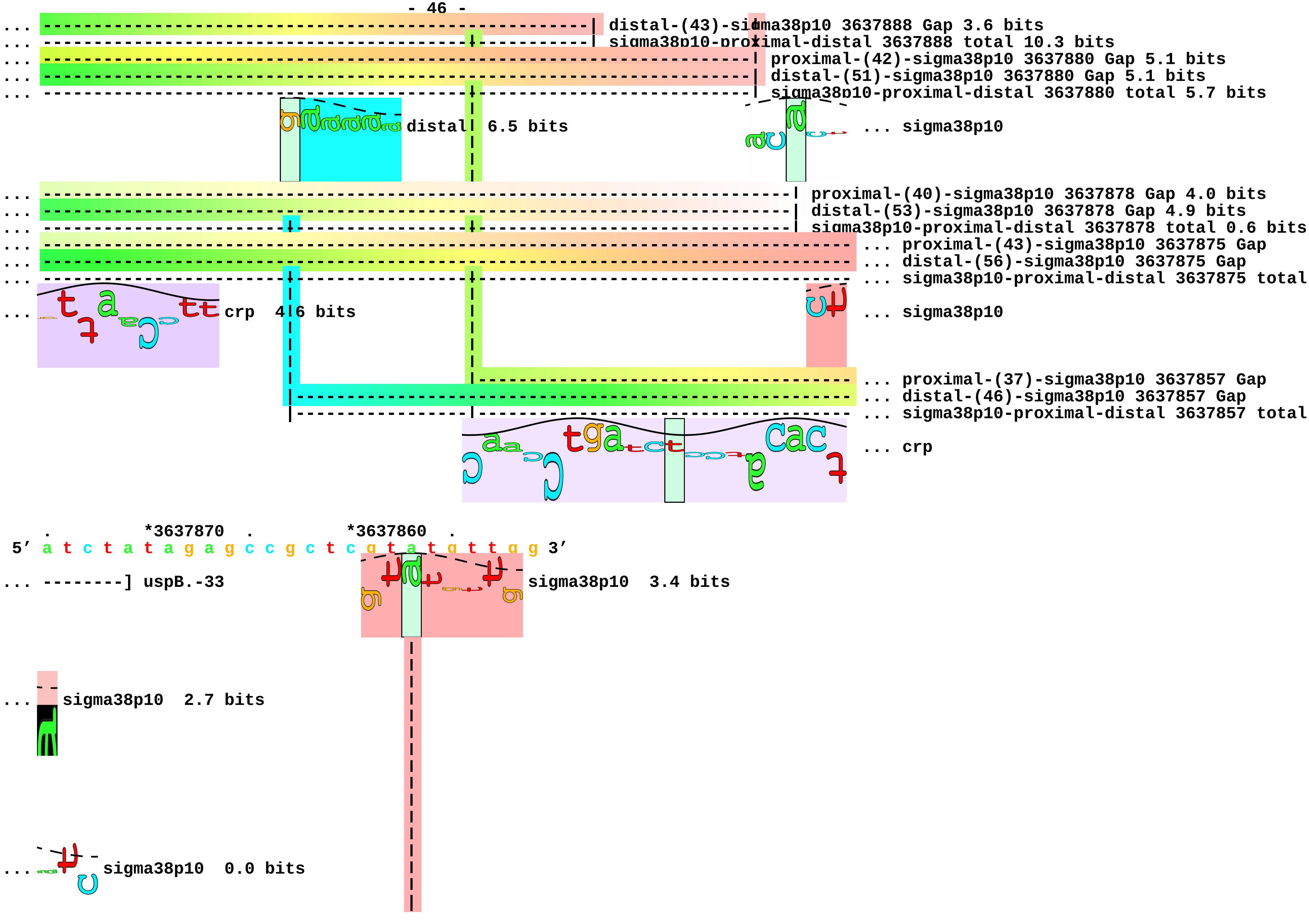
... distal-(43)-sigma38p10 3637888 Gap 3.6 bits
... sigma38p10-proximal-distal 3637888 total 10.3 bits
... proximal-(42)-sigma38p10 3637880 Gap 5.1 bits
... distal-(51)-sigma38p10 3637880 Gap 5.1 bits
... sigma38p10-proximal-distal 3637880 total 5.7 bits
distal 6.5 bits
... sigma38p10
... proximal-(40)-sigma38p10 3637878 Gap 4.0 bits
... distal-(53)-sigma38p10 3637878 Gap 4.9 bits
... sigma38p10-proximal-distal 3637878 total 0.6 bits
... proximal-(43)-sigma38p10 3637875 Gap
... distal-(56)-sigma38p10 3637875 Gap
... sigma38p10-proximal-distal 3637875 total
... crp 4.6 bits
... sigma38p10
... proximal-(37)-sigma38p10 3637857 Gap
... distal-(46)-sigma38p10 3637857 Gap
... sigma38p10-proximal-distal 3637857 total
... crp
. *3637870 . *3637860 .
5′ a t c t a t a g a g c c g c t c g t a t g t t g g 3′
... --------] uspB.-33
sigma38p10 3.4 bits
... sigma38p10 2.7 bits
... sigma38p10 0.0 bits

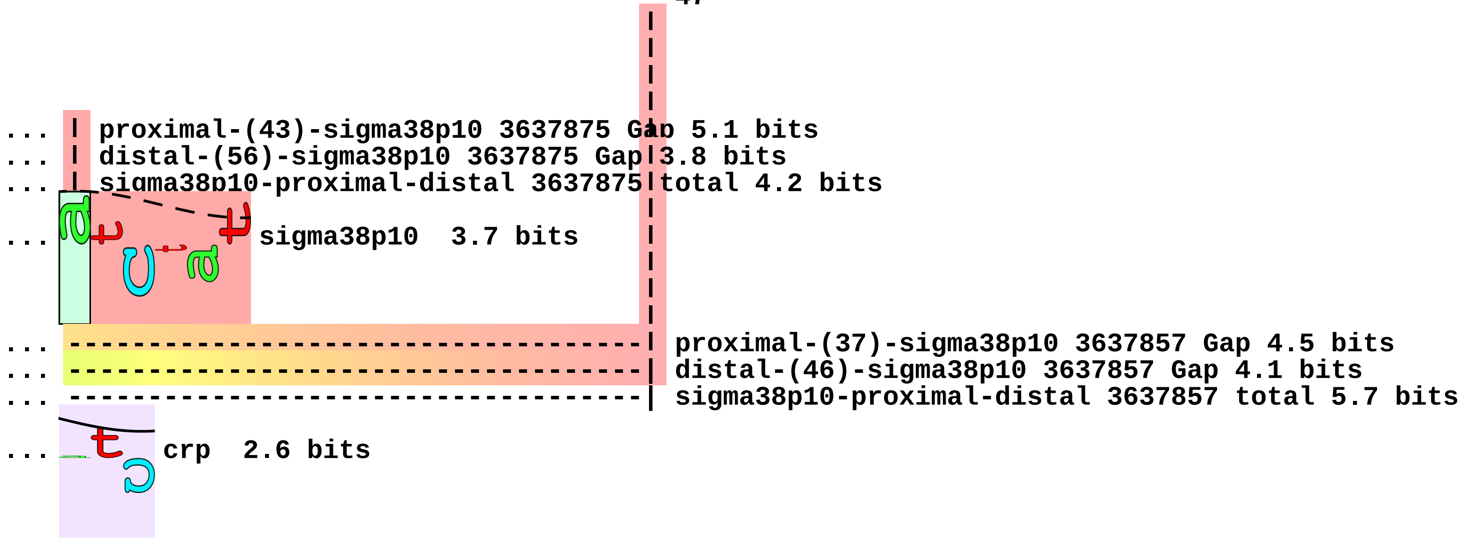
... proximal-(43)-sigma38p10 3637875 Gap 5.1 bits
... distal-(56)-sigma38p10 3637875 Gap 3.8 bits
... sigma38p10-proximal-distal 3637875 total 4.2 bits
... sigma38p10 3.7 bits
... proximal-(37)-sigma38p10 3637857 Gap 4.5 bits
... distal-(46)-sigma38p10 3637857 Gap 4.1 bits
... sigma38p10-proximal-distal 3637857 total 5.7 bits
... crp 2.6 bits

piece 22, NC_000913.2, ada, config: linear, direction: -, begin: 2308536, end: 2308432

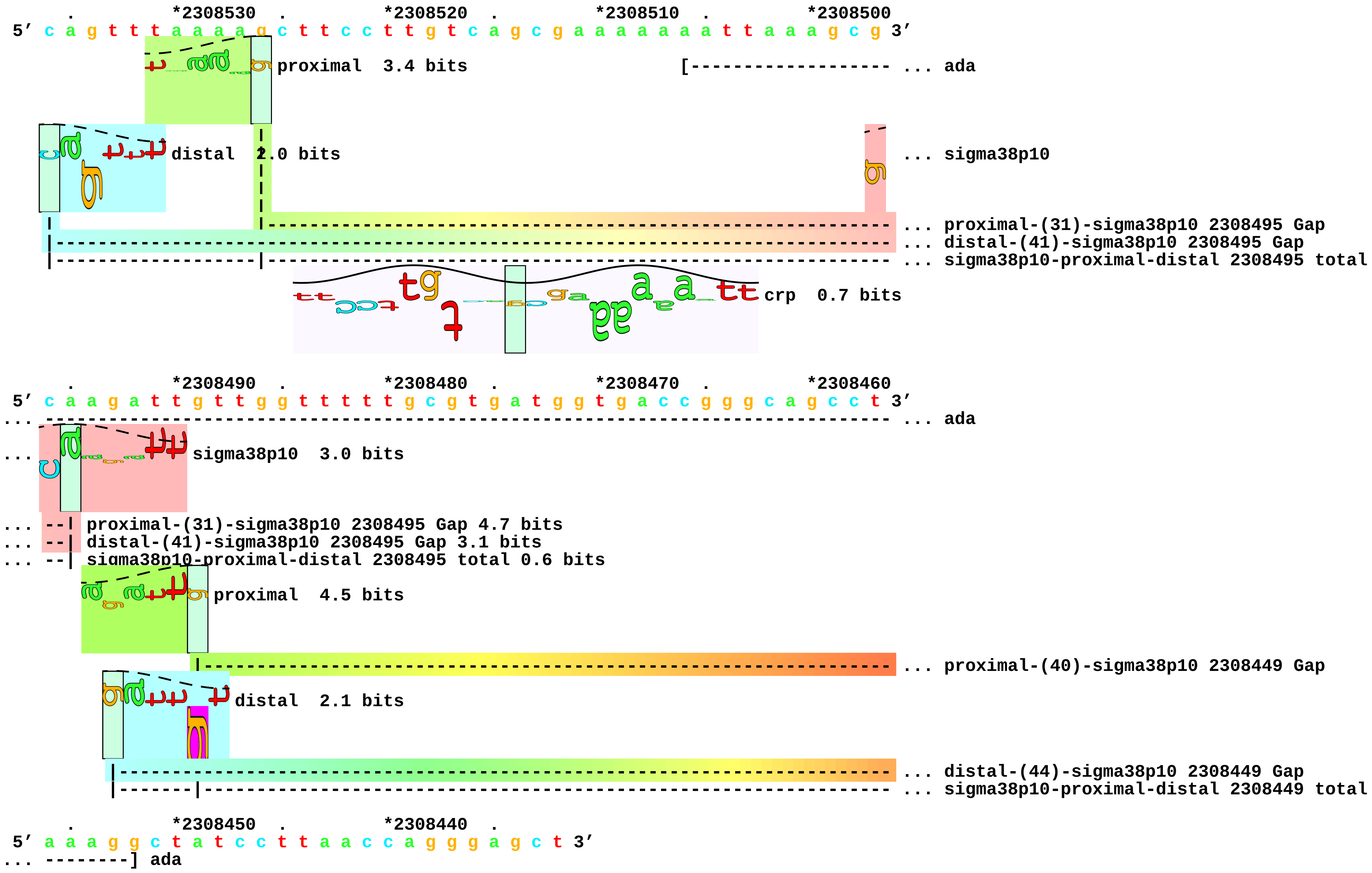

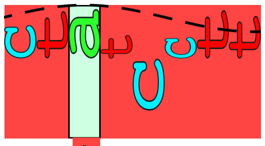

sigma38p10 8.0 bits

... --------------| proximal-(40)-sigma38p10 2308449 Gap 4.0 bits

... --------------| distal-(44)-sigma38p10 2308449 Gap 2.8 bits

... --------------| sigma38p10-proximal-distal 2308449 total 7.8 bits piece 23, NC_000913.2, aldB, config: linear, direction: -, begin: 3754645, end: 3754538

```
   .          *3754640   .          *3754630   .          *3754620   .          *3754610
5' g t a a c c c t t g c c c g t a a a t t c g t g a t a g c t g t c g t a a a g c 3'
```

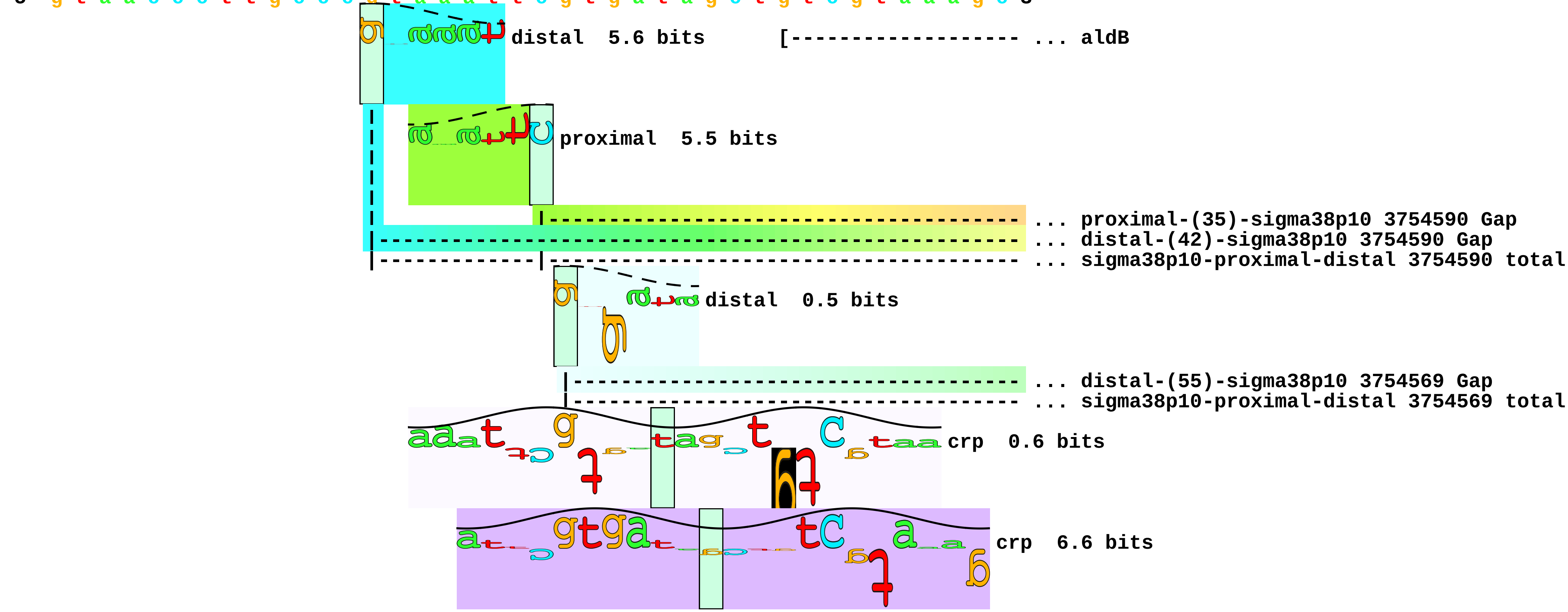


```
   .          *3754600   .          *3754590   .          *3754580   .          *3754570
5' t g t t a c c g a c t g g c g a a g a t t t c g c c a g t c a c g t c t a c c c 3'
```

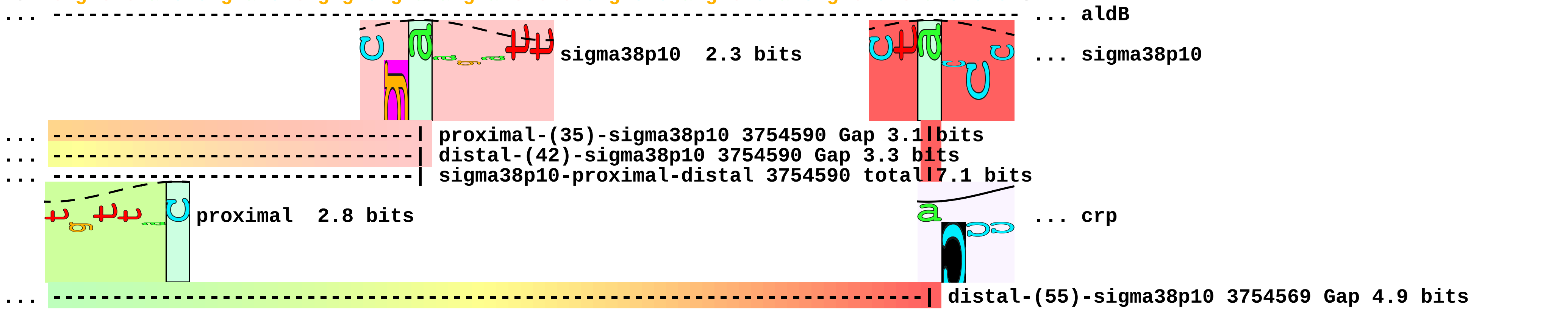

```
... ----------|-------------------------------------------------------------| sigma38p10-proximal-distal 3754569 total 0.6 bits
              |-------------------------------------------------------------| proximal-(31)-sigma38p10 3754569 Gap 4.7 bits

     .         *3754560  .         *3754550  .         *3754540
 5' t t g t t a t a c c t c a c a c c g c a a g g a g a c g 3'
... --------------] aldB
```

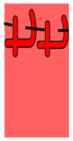

... sigma38p10   6.8 bits

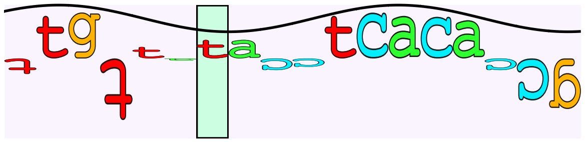

... crp   1.1 bits

```
piece 24, NC_000913.2, appA, config: linear, direction: +, begin: 1039735, end: 1039843

    .         *1039740  .         *1039750  .         *1039760  .         *1039770
5' g g c t g g a c c c g c g t c t g a a a a g t t a a c g a a c g t a g g c c t g 3'
                                                                 [------------------- ... appA

    .         *1039780  .         *1039790  .         *1039800  .         *1039810
5' a t g c g g c g c a t t a g c a t c g c a t c a g g c a a t c a a t a a t g t c 3'
... -------------------------------------------------------------------------------- ... appA

    .         *1039820  .         *1039830  .         *1039840
5' a g a t a t g a a a a g c g g a a a c a t a t c g a t g a 3'
... ----------------] appA
```

```
piece 25, NC_000913.2, appCB, config: linear, direction: +, begin: 1036845, end: 1036953

   .         *1036850  .         *1036860  .         *1036870  .         *1036880
5' c g c t t t c a g t c t t a t g a a t a t c g c a a t c g g c g a a t a c c t c 3'
                                                               [------------------- ... appCB
                                                      crp  4.3 bits

   .         *1036890  .         *1036900  .         *1036910  .         *1036920
5' t g g t c g t a g a g t t t c a g g a t a a a g a g g g a g a t c t a c c a t t 3'
... ---------------------------------------------------------------------------------- ... appCB
                         proximal  5.0 bits                                             ... crp
                                 |---------------------------------------------------- ... proximal-(34)-sigma38p10 1036933 Gap
                       distal  2.8 bits
                     |-------------------------------------------------------------- ... distal-(40)-sigma38p10 1036933 Gap
                     |-----------|-------------------------------------------------- ... distal-proximal-sigma38p10 1036933 total
                                                                                        ... crp

   .         *1036930  .         *1036940  .         *1036950
5' a t c g g g t t a t t t t t c t c t c t t c g c c t a c a 3'
... ----------------] appCB
...                                             crp  0.6 bits
... ----------------| proximal-(34)-sigma38p10 1036933 Gap 2.5 bits
                         sigma38p10  7.0 bits
... ----------------| distal-(40)-sigma38p10 1036933 Gap 3.1 bits
... ----------------| distal-proximal-sigma38p10 1036933 total 9.2 bits

... crp  0.3 bits
```

piece 26, NC_000913.2, cfa-P2, config: linear, direction: +, begin: 1739318, end: 1739423

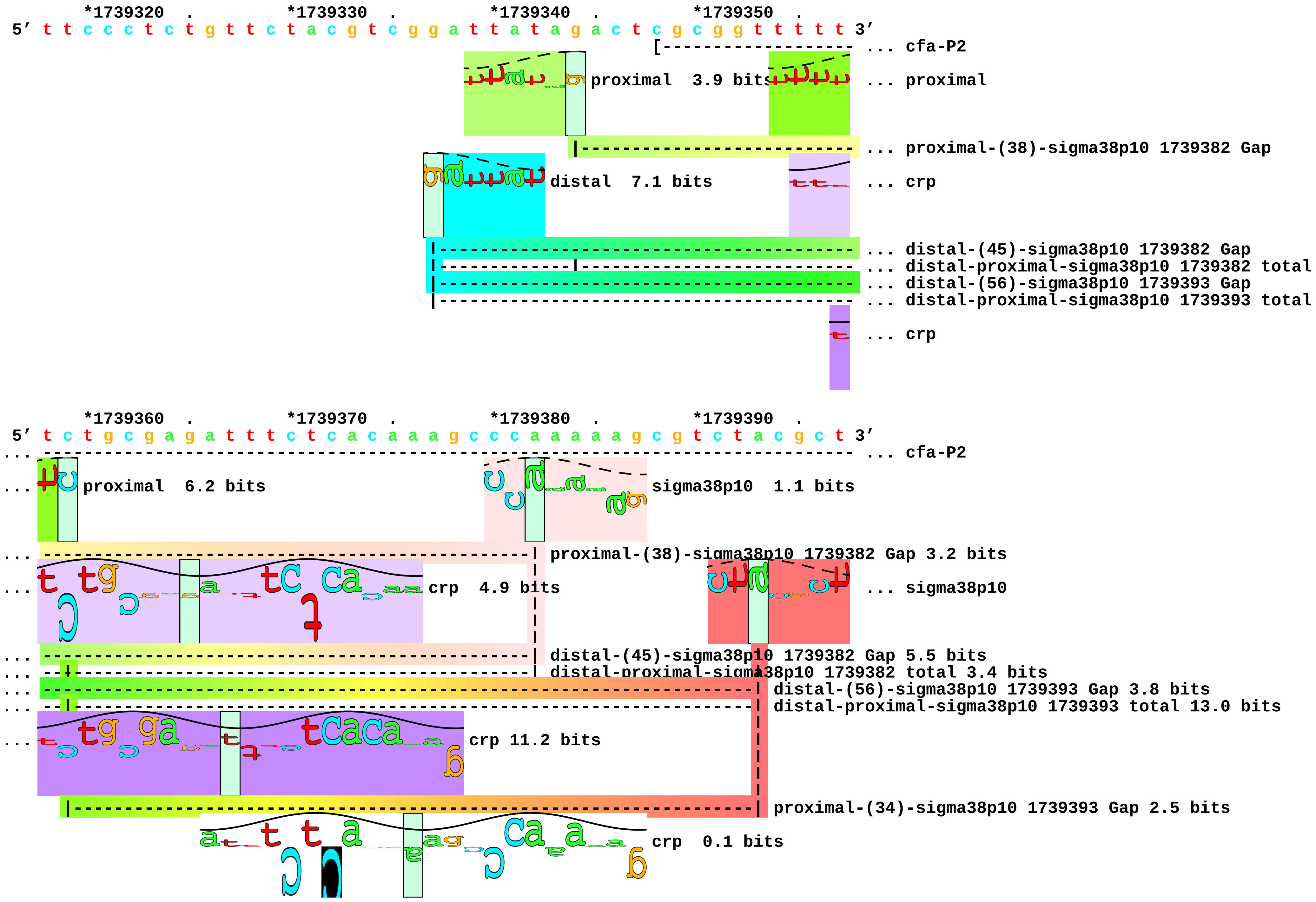

```
         *1739400  .         *1739410  .         *1739420
 5' g t t t t a a g g t t c t g a t c a c c g a c c a g 3'
... ----------] cfa-P2

... g sigma38p10  6.0 bits
```

piece 27, NC_000913.2, cpxRA, config: linear, direction: -, begin: 4103795, end: 4103688

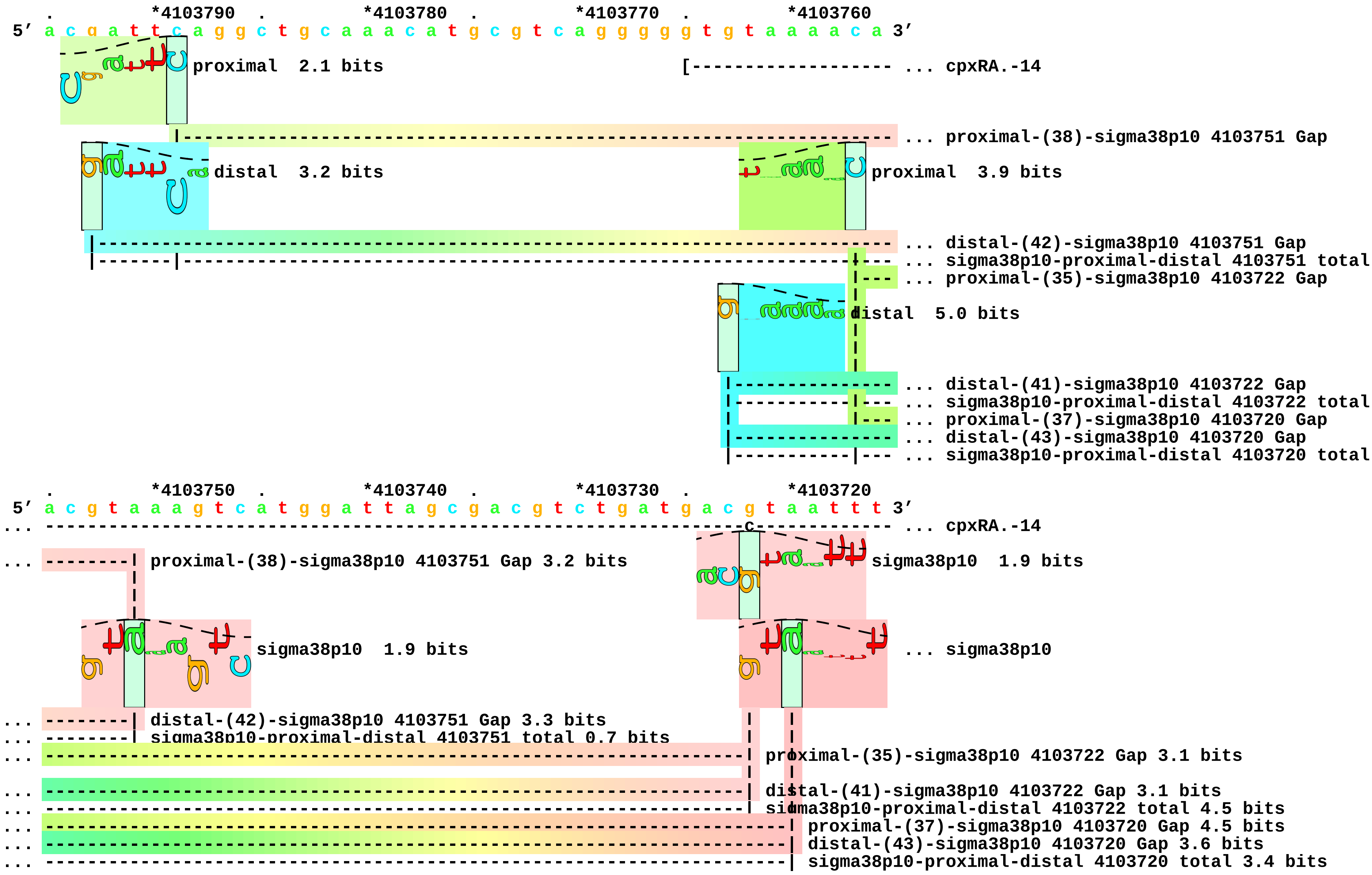

```
    .         *4103710  .         *4103700  .         *4103690
 5' c t g c c t c g g a g g t a t t t a a a c a a t g a a t 3'
... -------------] cpxRA.-14

... -- sigma38p10  2.6 bits
```

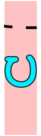

piece 28, NC_000913.2, csgBA, config: linear, direction: +, begin: 1102996, end: 1103102

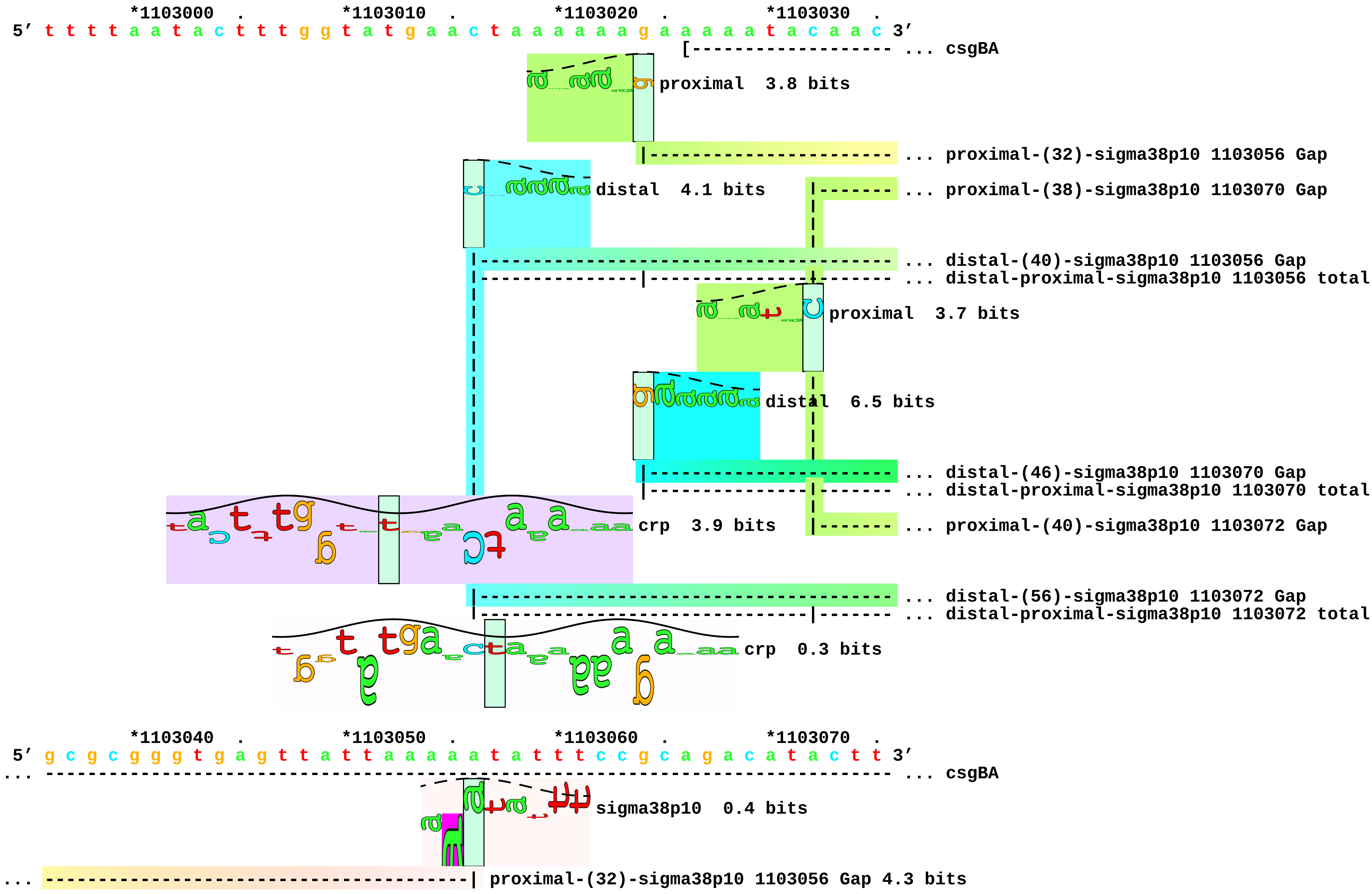

```
... ----------------------------------------------------------------------| proximal-(38)-sigma38p10 1103070 Gap 3.2 bits
... ----------------------------------------| distal-(40)-sigma38p10 1103056 Gap 3.1 bits
... ----------------------------------------| distal-proximal-sigma38p10 1103056 total 1.0 bits
                                                                          sigma38p10   5.8 bits
                                                                          ... sigma38p10
... ----------------------------------------------------------------------| distal-(46)-sigma38p10 1103070 Gap 4.1 bits
... ----------------------------------------------------------------------| distal-proximal-sigma38p10 1103070 total 8.7 bits
... ------------------------------------------------------------------------| proximal-(40)-sigma38p10 1103072 Gap 4.0 bits
... ------------------------------------------------------------------------| distal-(56)-sigma38p10 1103072 Gap 3.8 bits
... ------------------------------------------------------------------------| distal-proximal-sigma38p10 1103072 total 2.5 bits

             *1103080  .         *1103090  .         *1103100
 5' t c c a t c g t a a c g c a g c g t t a a c a a a a t 3'
... ------------] csgBA

... sigma38p10   2.4 bits
```

piece 29, NC_000913.2, dnaN-P2, config: linear, direction: -, begin: 3880698, end: 3880593

```
              .         *3880690  .         *3880680  .         *3880670  .         *3880660
5' g g g c g a t c a c c a t c g a c t t c g t g c g t g a g g c g c t g c g c g a 3'
                                                               [-------------------- ... dnaN-P2
                                                                                     ... crp
```

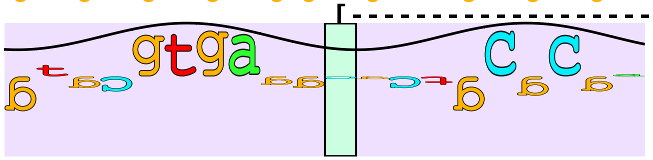

```
              .         *3880650  .         *3880640  .         *3880630  .         *3880620
5' c t t g c t g g c a t t g c a g g a a a a a c t g g t c a c c a t c g a c a a t 3'
... ---------------------------------------------------------------------------------- ... dnaN-P2
...   crp  3.0 bits                                                                 crp  0.7 bits
```

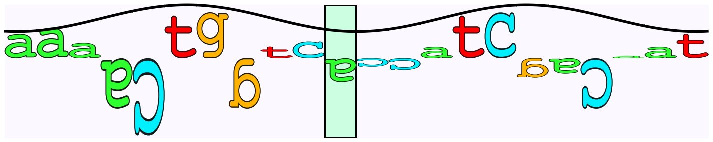

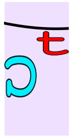

```
              .         *3880610  .         *3880600  .
5' a t t c a g a a g a c g g t g g c g g a g t a c t a 3'
... ----------] dnaN-P2
```

piece 30, NC_000913.2, dnaN-P4, config: linear, direction: -, begin: 3880461, end: 3880354

```
    *3880460  .         *3880450  .         *3880440  .         *3880430  .
5' g c c g t g a c c a c a c g a c g g t g c t t c a t g c c t g c c g t a a g a t 3'
                                                                [------------------- ... dnaN-P4

    *3880420  .         *3880410  .         *3880400  .         *3880390  .
5' c g a g c a g t t g c g t g a a g a g a g c c a c g a t a t c a a a g a a g a t 3'
... ---------------------------------------------------------------------------------- ... dnaN-P4
```

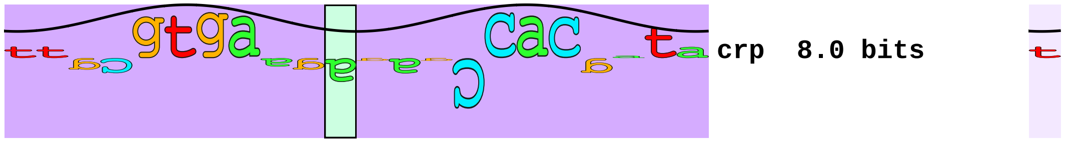

crp 8.0 bits ... crp

```
    *3880380  .         *3880370  .         *3880360  .
5' t t t t c a a a t t t a a t c a g a a c a t t g t c a t 3'
... --------------] dnaN-P4
```

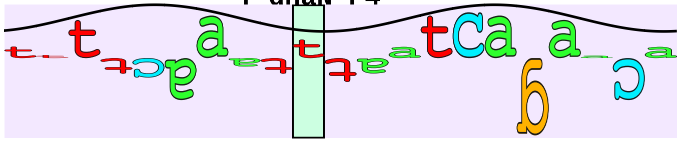

... crp 2.3 bits

piece 31, NC_000913.2, dps, config: linear, direction: -, begin: 848260, end: 848153

```
   *848260   .         *848250   .         *848240   .         *848230   .
5' g t t t t g t t t t t c a c g c t t g t t a c c a c t a t t a g t g t g a t a g 3'
                                                               [----------------- ... dps
                                                                     proximal  2.3 bits
                                                                              |- ... proximal-(36)-sigma38p10 848185 Gap
                                                        distal  3.5 bits|- ... proximal-(42)-sigma38p10 848179 Gap
                                                                                 ... distal-(50)-sigma38p10 848185 Gap
                                                                                 ... sigma38p10-proximal-distal 848185 total 9
                                                                                 ... distal-(56)-sigma38p10 848179 Gap
                                                                                 ... sigma38p10-proximal-distal 848179 total 1
                                                                                 ... crp

   *848220   .         *848210   .         *848200   .         *848190   .
5' g a a c a g c c a g a a t a g c g g a a c a c a t a g c c g g t g c t a t a c t 3'
... ------------------------------------------------------------------------------ ... dps
                                                                                   ... sigma38p10
... -------------------------------------------------------------| proximal-(36)-sigma38p10 848185 Gap 3.6 bits
... ------------------------------------------------------------------------------ ... proximal-(42)-sigma38p10 848179 Gap
... -------------------------------------------------------------| distal-(50)-sigma38p10 848185 Gap 4.1 bits
... -------------------------------------------------------------| sigma38p10-proximal-distal 848185 total 9.0 bits
... ------------------------------------------------------------------------------ ... distal-(56)-sigma38p10 848179 Gap
... ------------------------------------------------------------------------------ ... sigma38p10-proximal-distal 848179 total 1
...                    crp  0.6 bits                                                ... sigma38p10
                                                                                   ... crp

   *848180   .         *848170   .         *848160   .
```

5′ t a a t c t c g t t a a t t a c t g g g a c a t a a c a 3′

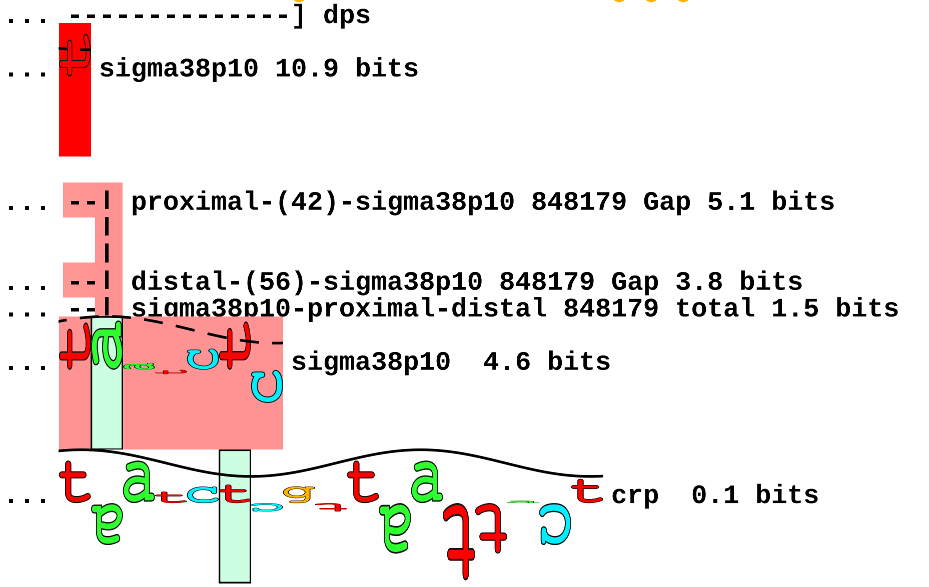

piece 32, NC_000913.2, ecnB, config: linear, direction: +, begin: 4374448, end: 4374553

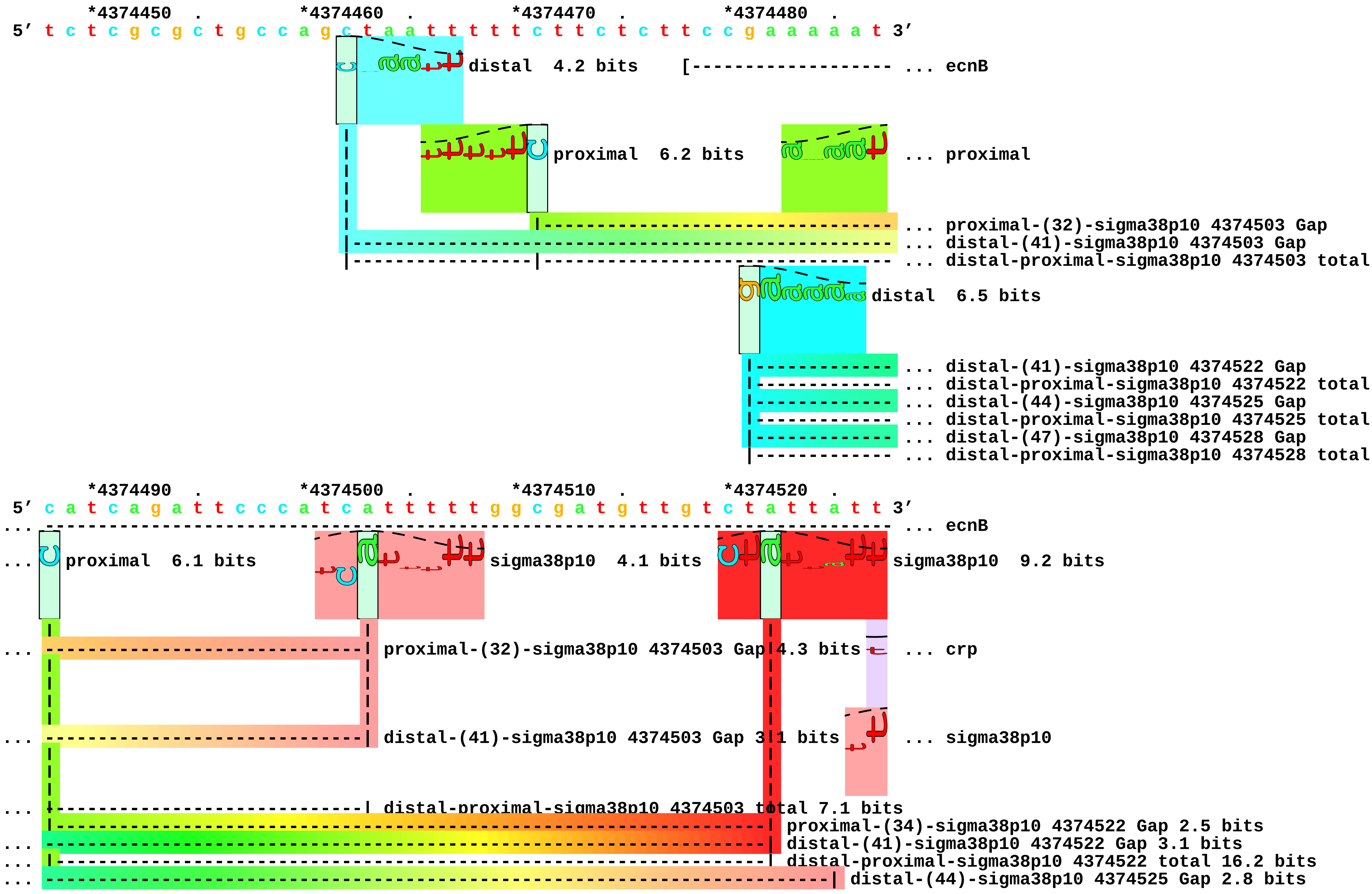

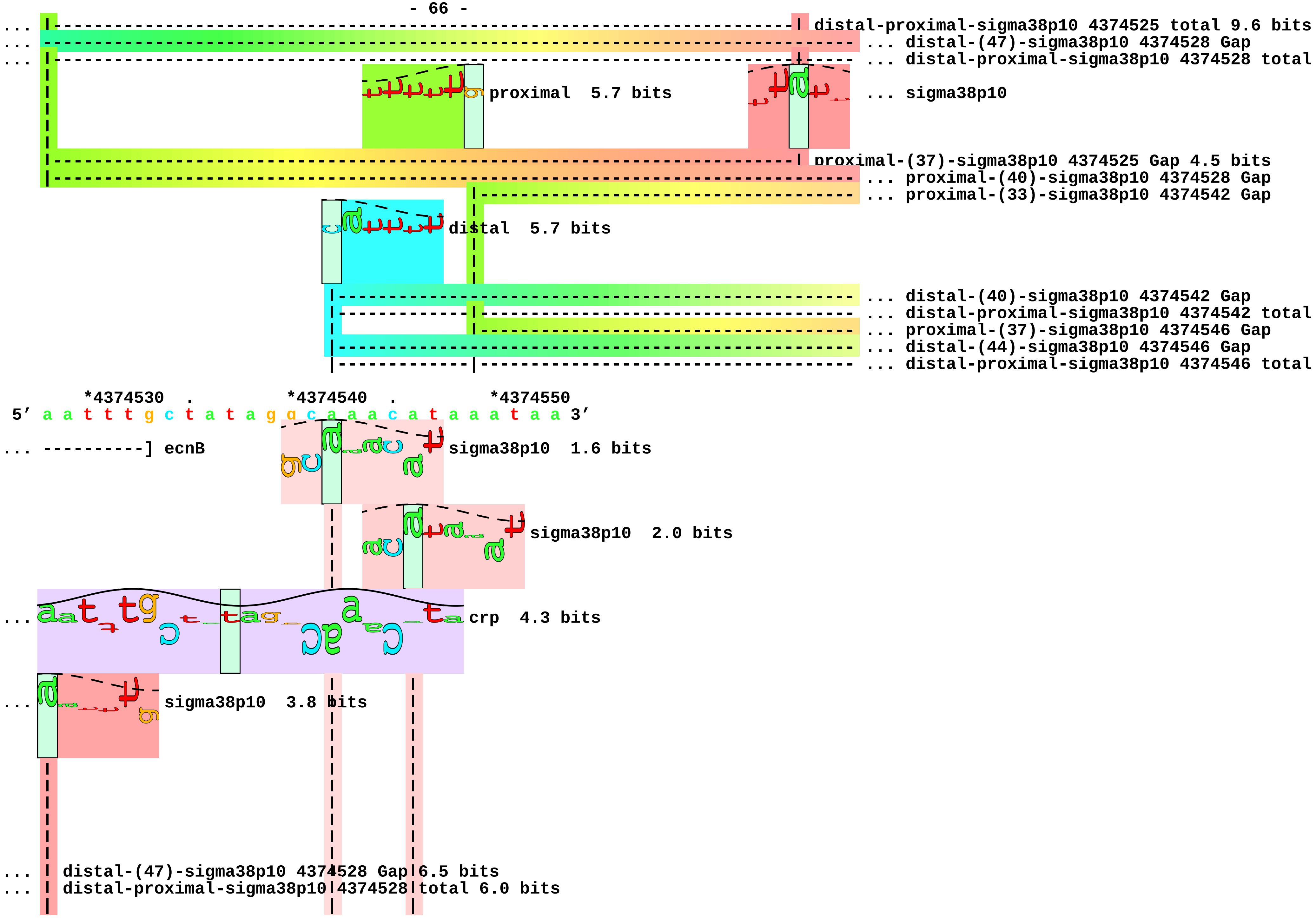
distal-proximal-sigma38p10 4374525 total 9.6 bits
... distal-(47)-sigma38p10 4374528 Gap
... distal-proximal-sigma38p10 4374528 total
proximal 5.7 bits
... sigma38p10
proximal-(37)-sigma38p10 4374525 Gap 4.5 bits
... proximal-(40)-sigma38p10 4374528 Gap
... proximal-(33)-sigma38p10 4374542 Gap
distal 5.7 bits
... distal-(40)-sigma38p10 4374542 Gap
... distal-proximal-sigma38p10 4374542 total
... proximal-(37)-sigma38p10 4374546 Gap
... distal-(44)-sigma38p10 4374546 Gap
... distal-proximal-sigma38p10 4374546 total
*4374530 . *4374540 . *4374550
5' a a t t t g c t a t a g g g c a a a c a t a a a t a a 3'
... ----------] ecnB
sigma38p10 1.6 bits
sigma38p10 2.0 bits
... crp 4.3 bits
... sigma38p10 3.8 bits
... distal-(47)-sigma38p10 4374528 Gap 6.5 bits
... distal-proximal-sigma38p10 4374528 total 6.0 bits

```
...          sigma38p10  4.3 bits
...  | proximal-(40)-sigma38p10 4374528 Gap 4.0 bits
...  ----------------------------| proximal-(33)-sigma38p10 4374542 Gap 4.0 bits
...  ----------------------------| distal-(40)-sigma38p10 4374542 Gap 3.1 bits
...  ----------------------------| distal-proximal-sigma38p10 4374542 total 5.9 bits
...  ------------------------------------| proximal-(37)-sigma38p10 4374546 Gap 4.5 bits
...  ------------------------------------| distal-(44)-sigma38p10 4374546 Gap 2.8 bits
...  ------------------------------------| distal-proximal-sigma38p10 4374546 total 6.1 bits
```

```
piece 33, NC_000913.2, fic, config: linear, direction: -, begin: 3489754, end: 3489650

          *3489750  .         *3489740  .         *3489730  .         *3489720  .
5' t g c c g t a a t g a t t t c t c g c g c g g g c a a t c t t g c c c g c g c t 3'
                                                              [------------------- ... fic.-36.
                                                                 proximal  2.6 bits
                                                               |------------------ ... proximal-(43)-sigma38p10 3489680 Gap
                                                           distal  3.1 bits
                                                       |-------------------------- ... distal-(50)-sigma38p10 3489680 Gap
                                                       |-------------|------------ ... sigma38p10-proximal-distal 3489680 total

          *3489710  .         *3489700  .         *3489690  .         *3489680  .
5' t c t g c t c t c c c g g c g t a a c c c g g a t t t g c c g c t t a t a c t t 3'
... ----------------t------------------------------------------------------------- ... fic.-36.
                                                                         sigma38p10  9.2 bits
... -----------------------------------------------------------------------| proximal-(43)-sigma38p10 3489680 Gap 5.1 bits
... -----------------------------------------------------------------------| distal-(50)-sigma38p10 3489680 Gap 4.1 bits
... -----------------------------------------------------------------------| sigma38p10-proximal-distal 3489680 total 5.7 bits

          *3489670  .         *3489660  .         *3489650
5' g t g g c a a a t g g a c a c g t t c a g g g a g 3'
... --------g fic.-36.
```

piece 34, NC_000913.2, gadA, config: linear, direction: -, begin: 3665717, end: 3665611

```
         .         *3665710  .         *3665700  .         *3665690  .         *3665680
5' a a g t c g t t t t t c t g c t t a g g a t t t t g t t a t t t a a a t t a a g 3'
                                                           [--------------------- ... gadA
                                 distal   6.5 bits                               proximal   4.9 bits
                                                                                  ... proximal-(36)-sigma38p10 3665642 Gap
                                                                                  ... distal-(56)-sigma38p10 3665642 Gap
                                                                                  ... sigma38p10-proximal-distal 3665642 total
                                                                                  ... crp

         .         *3665670  .         *3665660  .         *3665650  .         *3665640
5' c c t g t a a t g c c t t g c t t c c a t t g c g g a t a a a t c c t a c t t t 3'
... ------------------------------------------------------------------------------ ... gadA
                                                                                  ... sigma38p10
... ---------------------------------------------------------------------| proximal-(36)-sigma38p10 3665642 Gap 3.6 bits
... ---------------------------------------------------------------------| distal-(56)-sigma38p10 3665642 Gap 3.8 bits
... ---------------------------------------------------------------------| sigma38p10-proximal-distal 3665642 total 11.6 bits
... crp   5.7 bits                                                        ... crp

         .         *3665630  .         *3665620  .
5' t t t a t t g c c t t c a a a t a a a t t t a a g g a 3'
... -----------] gadA
... sigma38p10   7.5 bits
                                              crp   1.3 bits
```

...
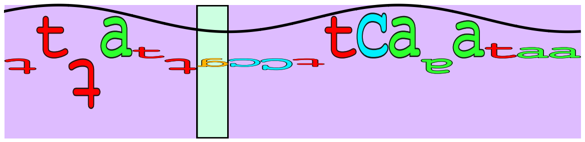
- 70 -
crp 6.5 bits

piece 35, NC_000913.2, gadB, config: linear, direction: -, begin: 1570182, end: 1570077

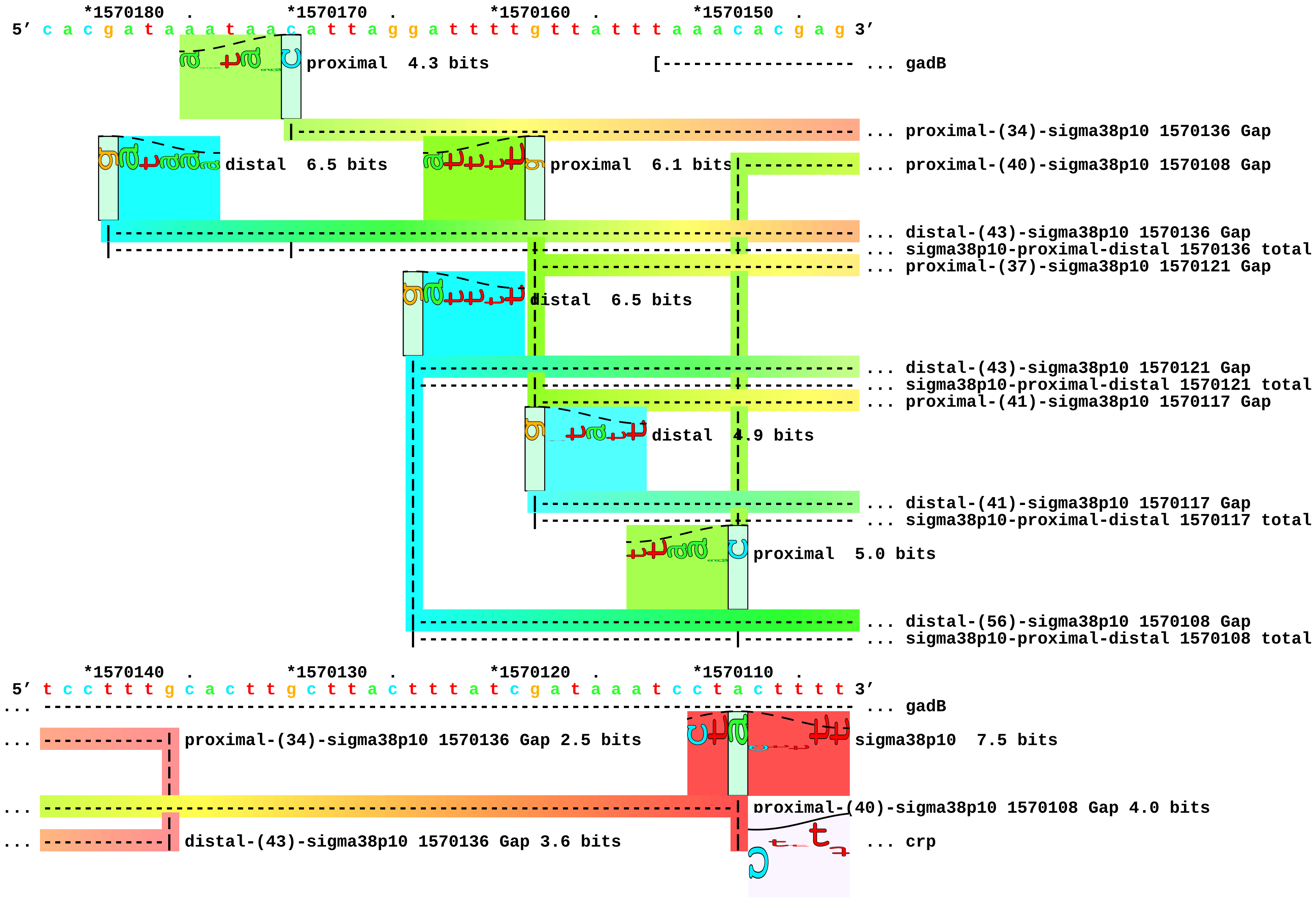

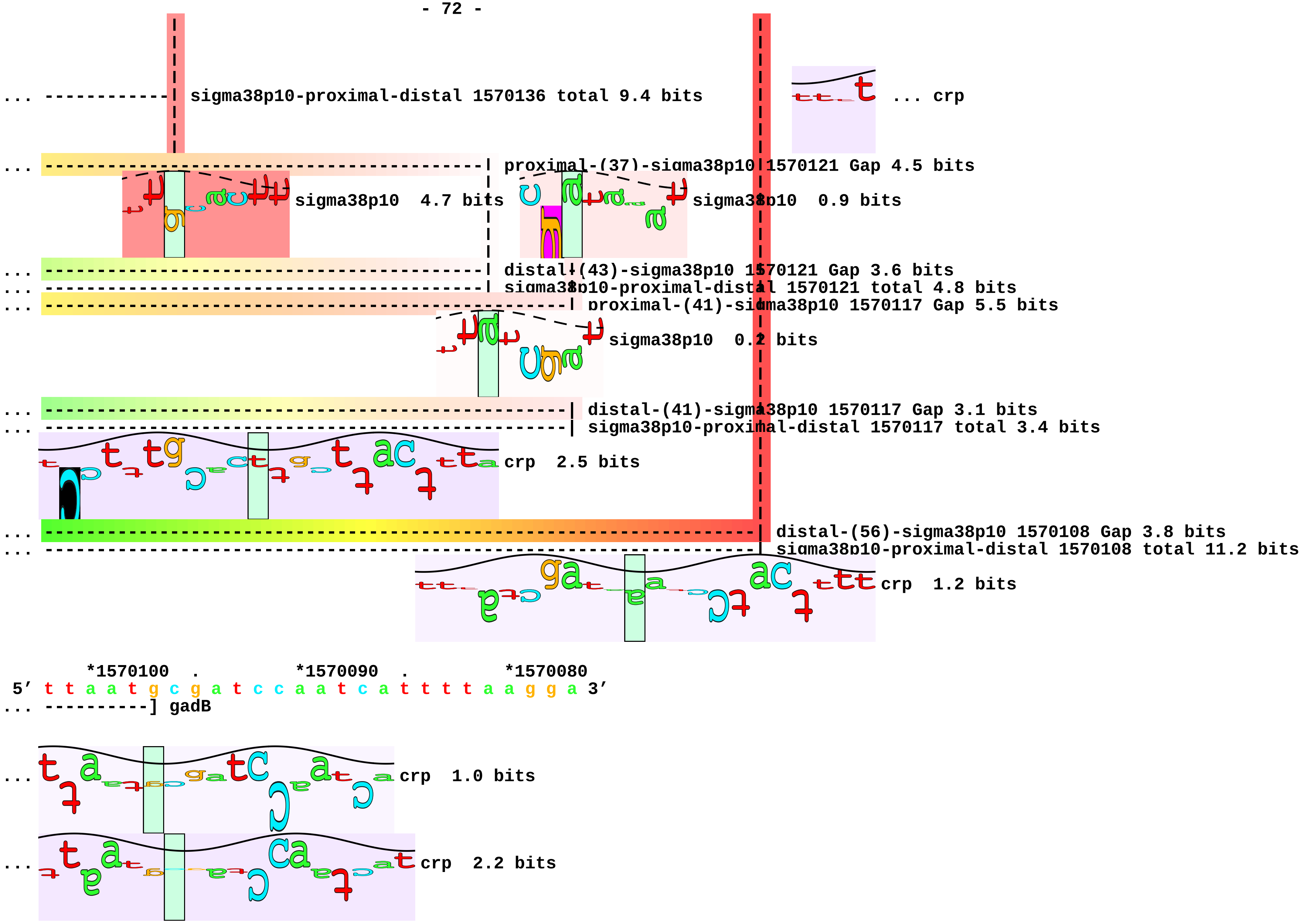
sigma38p10-proximal-distal 1570136 total 9.4 bits
... crp
proximal-(37)-sigma38p10 1570121 Gap 4.5 bits
sigma38p10 4.7 bits
sigma38p10 0.9 bits
distal-(43)-sigma38p10 1570121 Gap 3.6 bits
sigma38p10-proximal-distal 1570121 total 4.8 bits
proximal-(41)-sigma38p10 1570117 Gap 5.5 bits
sigma38p10 0.2 bits
distal-(41)-sigma38p10 1570117 Gap 3.1 bits
sigma38p10-proximal-distal 1570117 total 3.4 bits
crp 2.5 bits
distal-(56)-sigma38p10 1570108 Gap 3.8 bits
sigma38p10-proximal-distal 1570108 total 11.2 bits
crp 1.2 bits
*1570100 . *1570090 . *1570080
5′ t t a a t g c g a t c c a a t c a t t t t a a g g a 3′
gadB
crp 1.0 bits
crp 2.2 bits

piece 36, NC_000913.2, gadC, config: linear, direction: -, begin: 1568670, end: 1568567

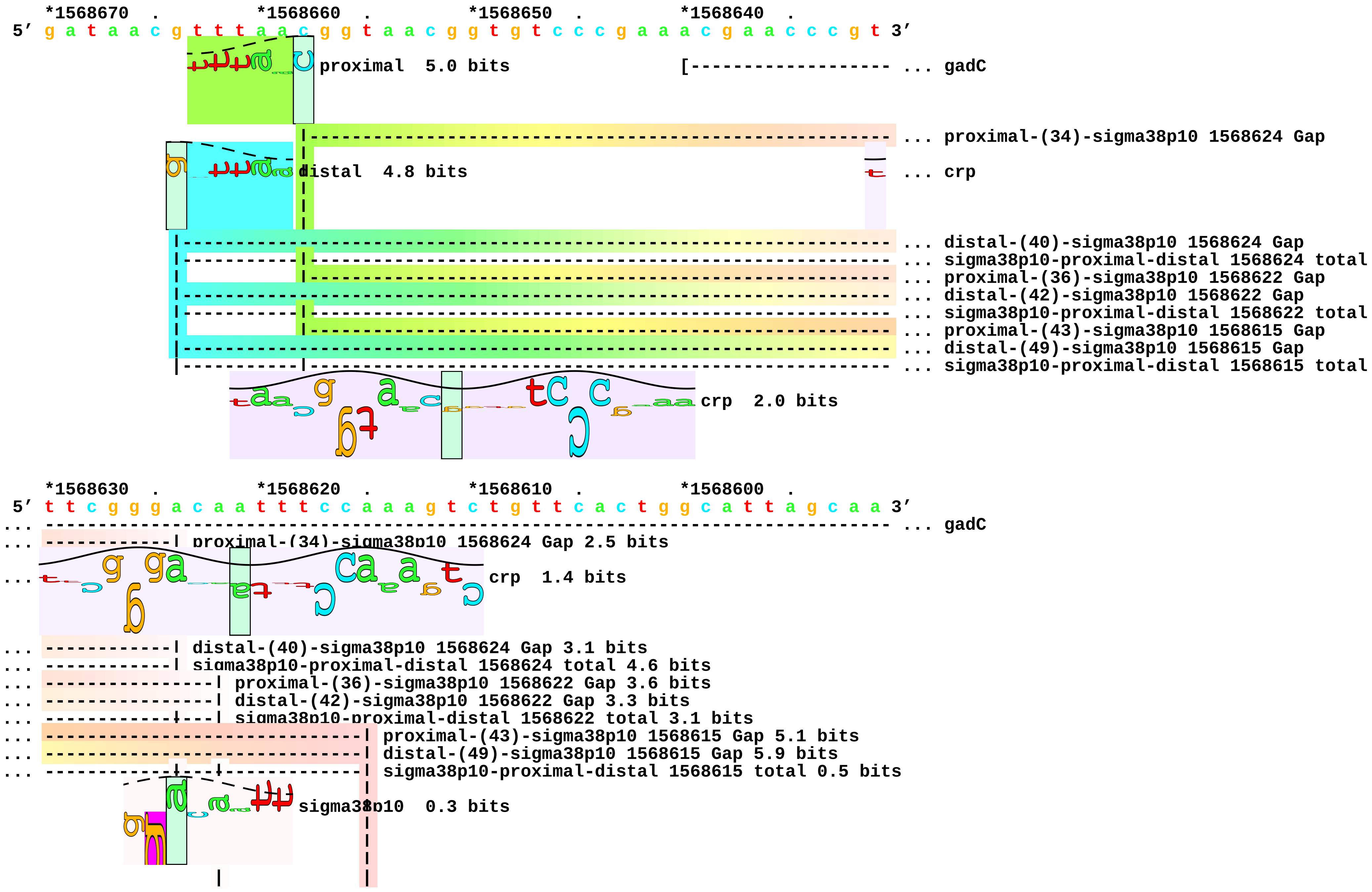

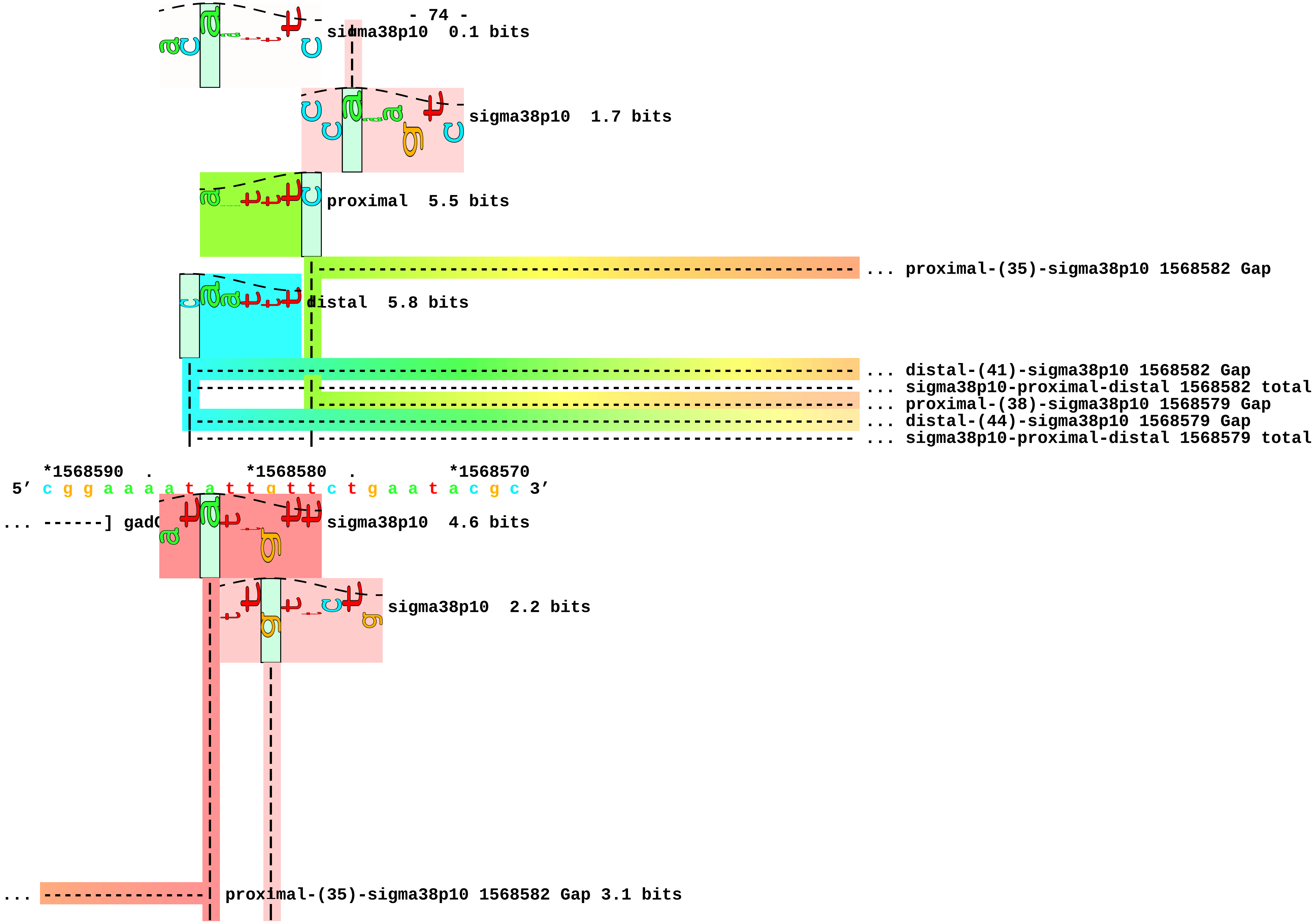

sigma38p10 0.1 bits
sigma38p10 1.7 bits
proximal 5.5 bits
... proximal-(35)-sigma38p10 1568582 Gap
distal 5.8 bits
... distal-(41)-sigma38p10 1568582 Gap
... sigma38p10-proximal-distal 1568582 total
... proximal-(38)-sigma38p10 1568579 Gap
... distal-(44)-sigma38p10 1568579 Gap
... sigma38p10-proximal-distal 1568579 total
*1568590 . *1568580 . *1568570
5′ c g g a a a a t a t t g t t c t g a a t a c g c 3′
... ------] gadC
sigma38p10 4.6 bits
sigma38p10 2.2 bits
... proximal-(35)-sigma38p10 1568582 Gap 3.1 bits

```
... ----------------| distal-(41)-sigma38p10 1568582 Gap 3.1 bits
... ----------------| sigma38p10-proximal-distal 1568582 total 9.7 bits
... ----------------------| proximal-(38)-sigma38p10 1568579 Gap 3.2 bits
... ----------------------| distal-(44)-sigma38p10 1568579 Gap 2.8 bits
... ----------------------| sigma38p10-proximal-distal 1568579 total 7.4 bits
```

piece 37, NC_000913.2, glgS-P2, config: linear, direction: -, begin: 3190111, end: 3190004

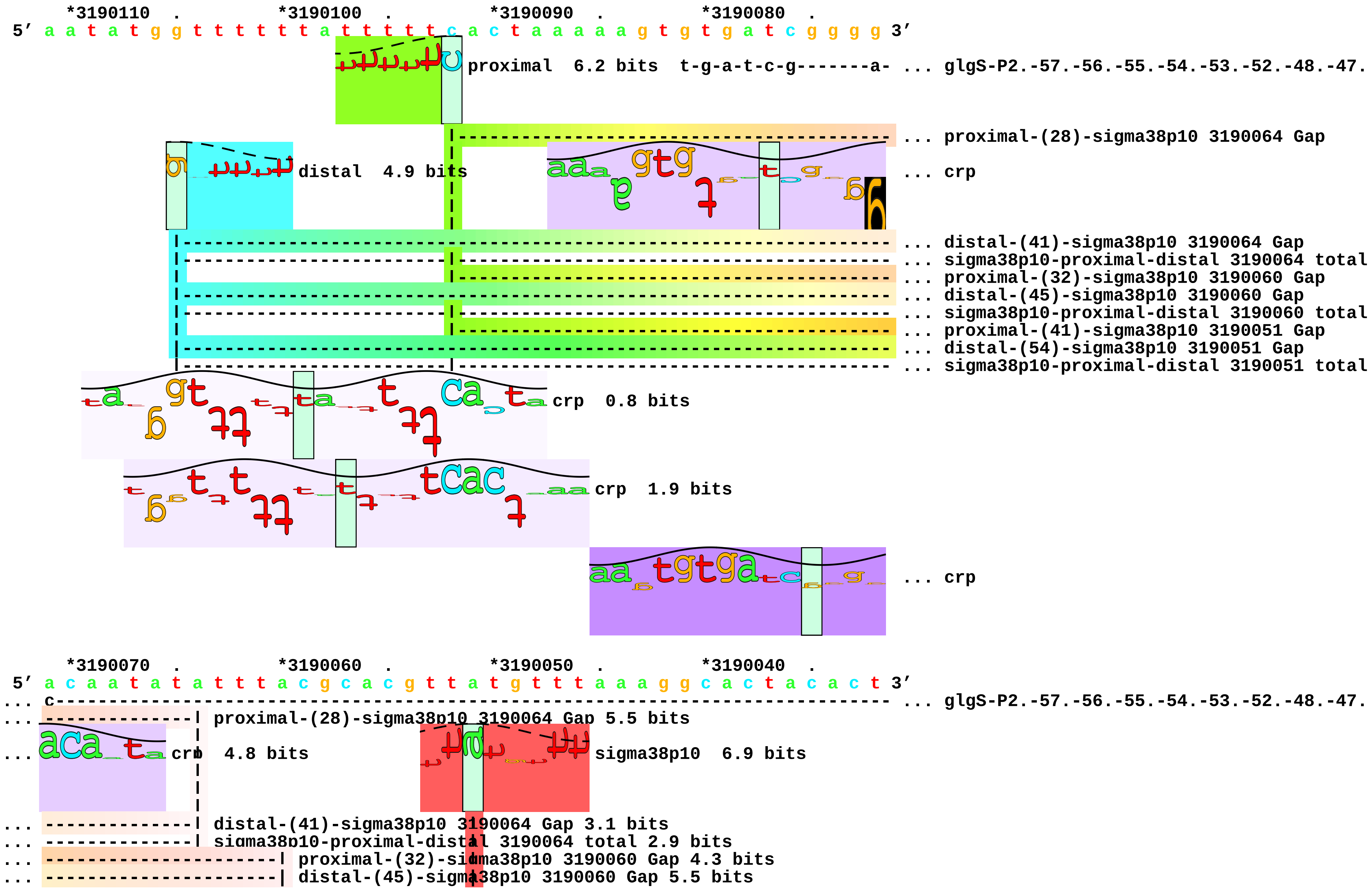

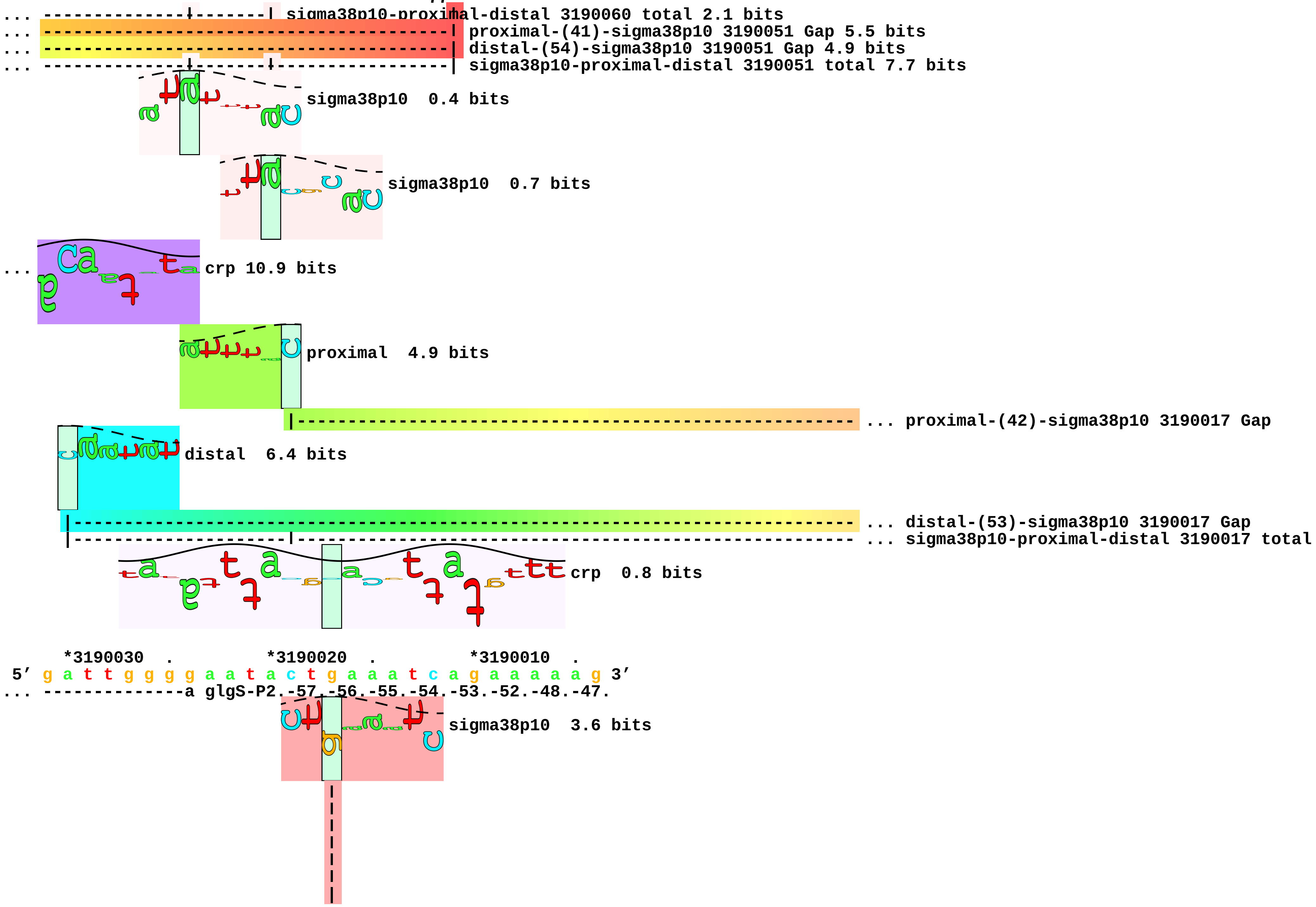
sigma38p10-proximal-distal 3190060 total 2.1 bits
proximal-(41)-sigma38p10 3190051 Gap 5.5 bits
distal-(54)-sigma38p10 3190051 Gap 4.9 bits
sigma38p10-proximal-distal 3190051 total 7.7 bits
sigma38p10 0.4 bits
sigma38p10 0.7 bits
crp 10.9 bits
proximal 4.9 bits
proximal-(42)-sigma38p10 3190017 Gap
distal 6.4 bits
distal-(53)-sigma38p10 3190017 Gap
sigma38p10-proximal-distal 3190017 total
crp 0.8 bits
*3190030 . *3190020 . *3190010 .
5′ g a t t g g g g a a t a c t g a a a t c a g a a a a a g 3′
... --------------a glgS-P2.-57.-56.-55.-54.-53.-52.-48.-47.
sigma38p10 3.6 bits

```
                            |
                            |
                            |
                            |
                            |
... ------------------------| proximal-(42)-sigma38p10 3190017 Gap 5.1 bits
                            |
... ------------------------| distal-(53)-sigma38p10 3190017 Gap 4.9 bits
... ------------------------| sigma38p10-proximal-distal 3190017 total 4.7 bits
```

piece 38, NC_000913.2, gor, config: linear, direction: +, begin: 3644213, end: 3644318

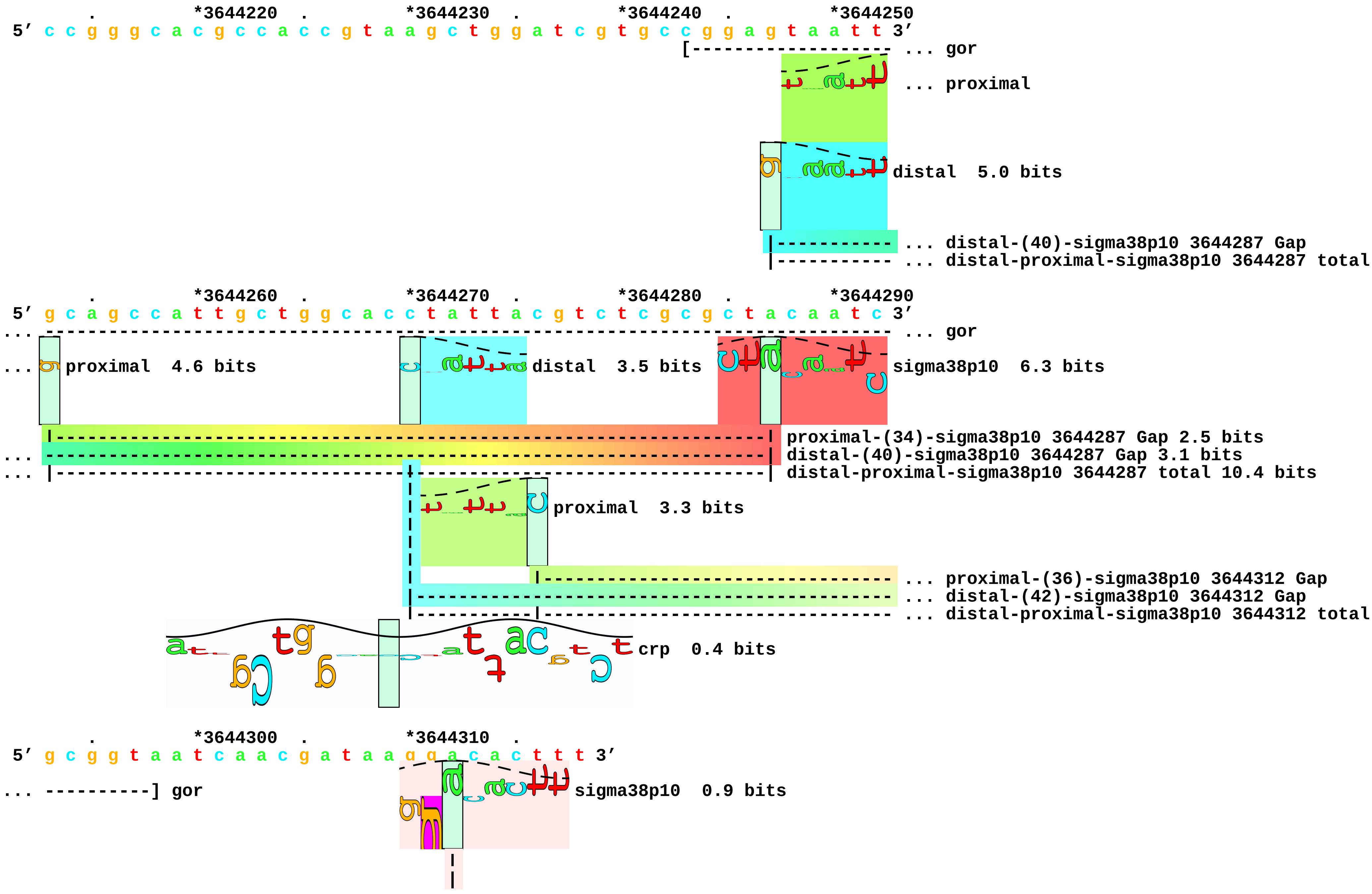

```
                                     |
                                     |
                                     |
... -------------------------------------| proximal-(36)-sigma38p10 3644312 Gap 3.6 bits
... -------------------------------------| distal-(42)-sigma38p10 3644312 Gap 3.3 bits
... -------------------------------------| distal-proximal-sigma38p10 3644312 total 0.9 bits
```

piece 39, NC_000913.2, himA-P4, config: linear, direction: -, begin: 1793830, end: 1793725

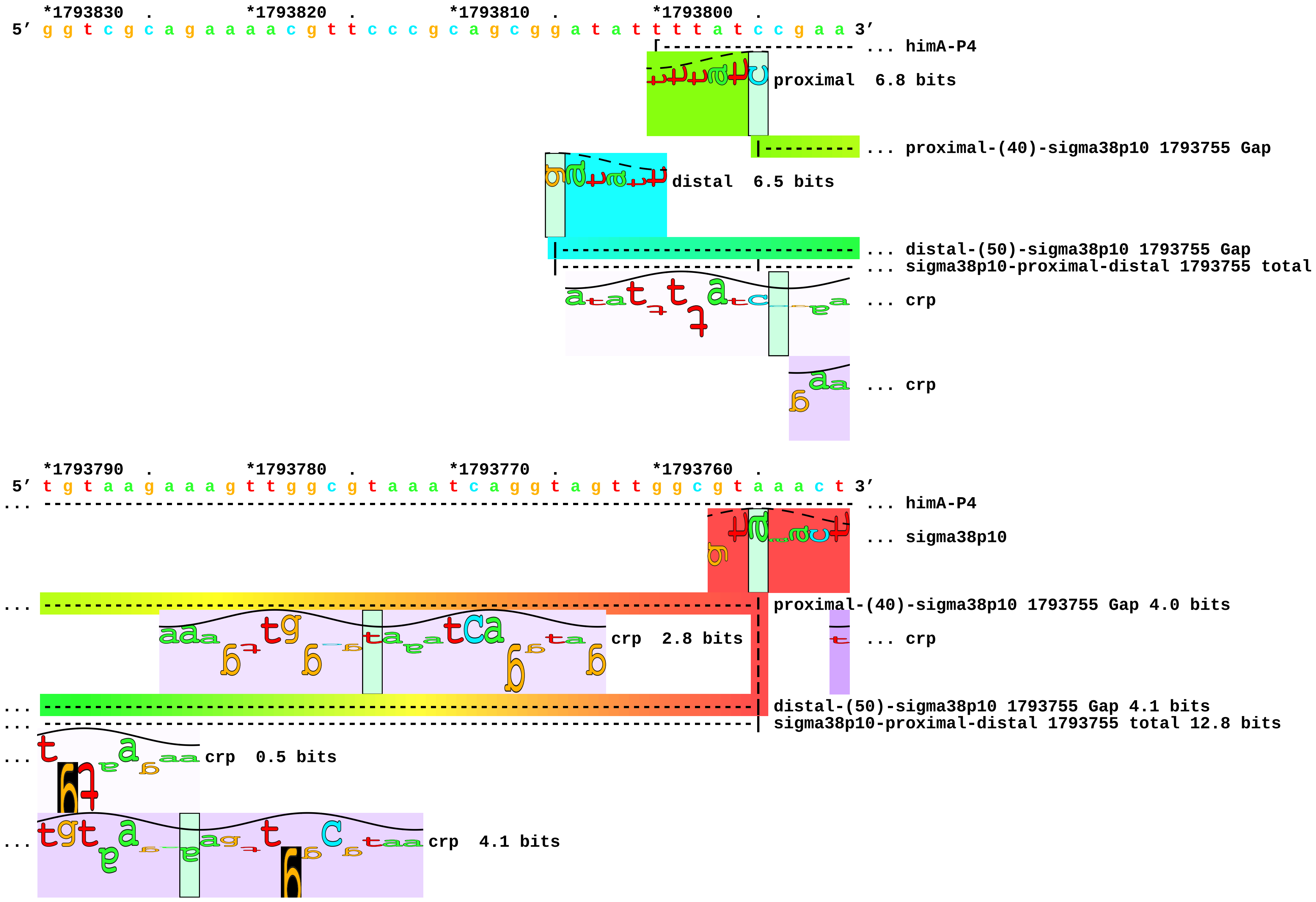

```
    *1793750  .         *1793740  .         *1793730  .
 5' t a t t t g a c g t g t a c c g c g g t a a g g g t 3'
... ----------] himA-P4
```

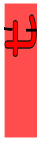

... sigma38p10 7.7 bits

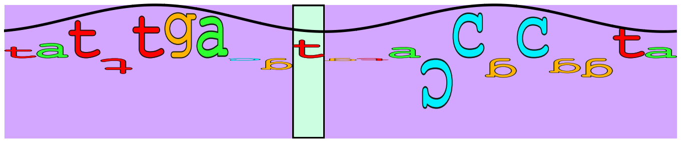

... crp 8.5 bits

piece 40, NC_000913.2, hyaAB, config: linear, direction: +, begin: 1031122, end: 1031227

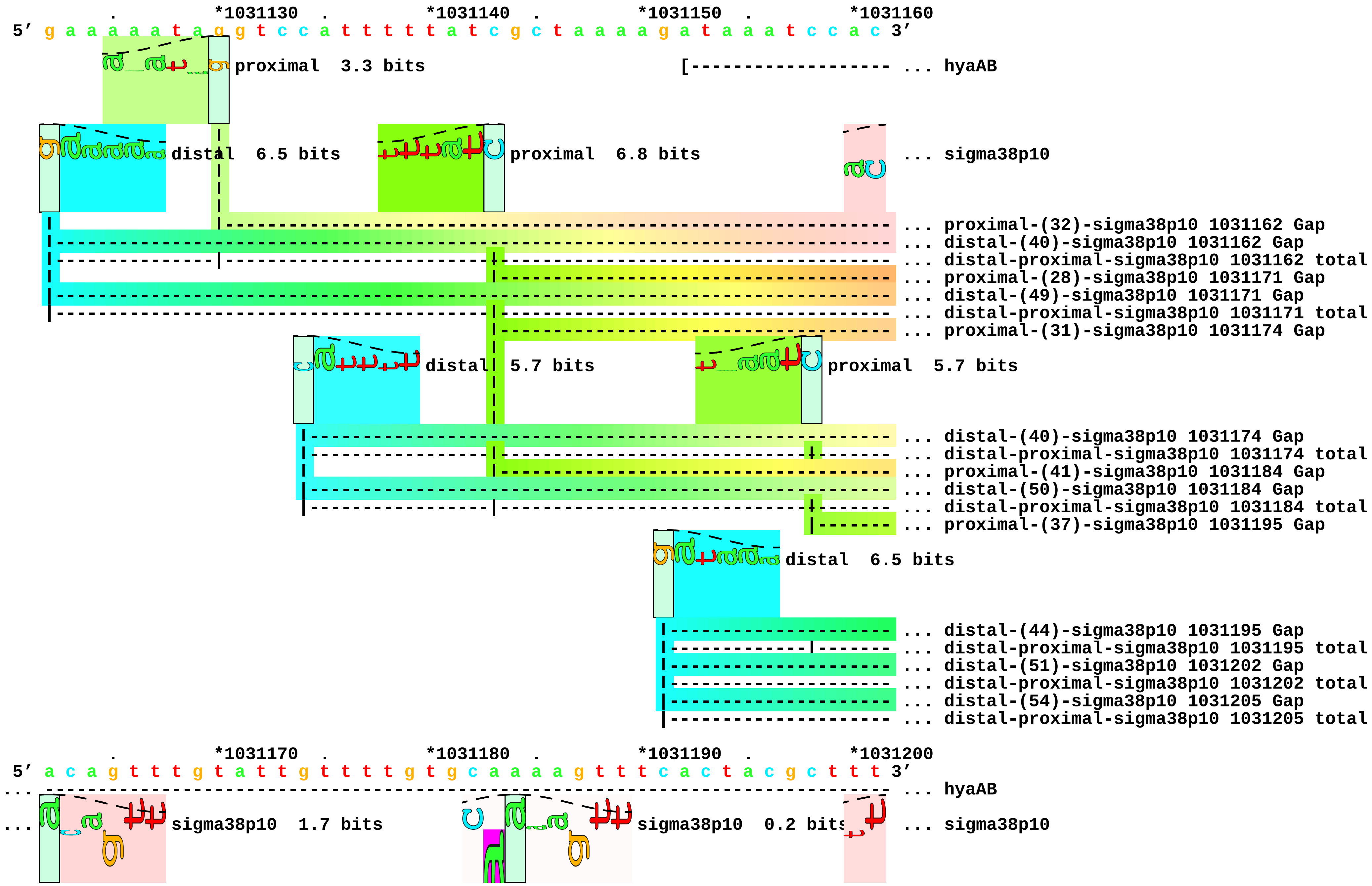

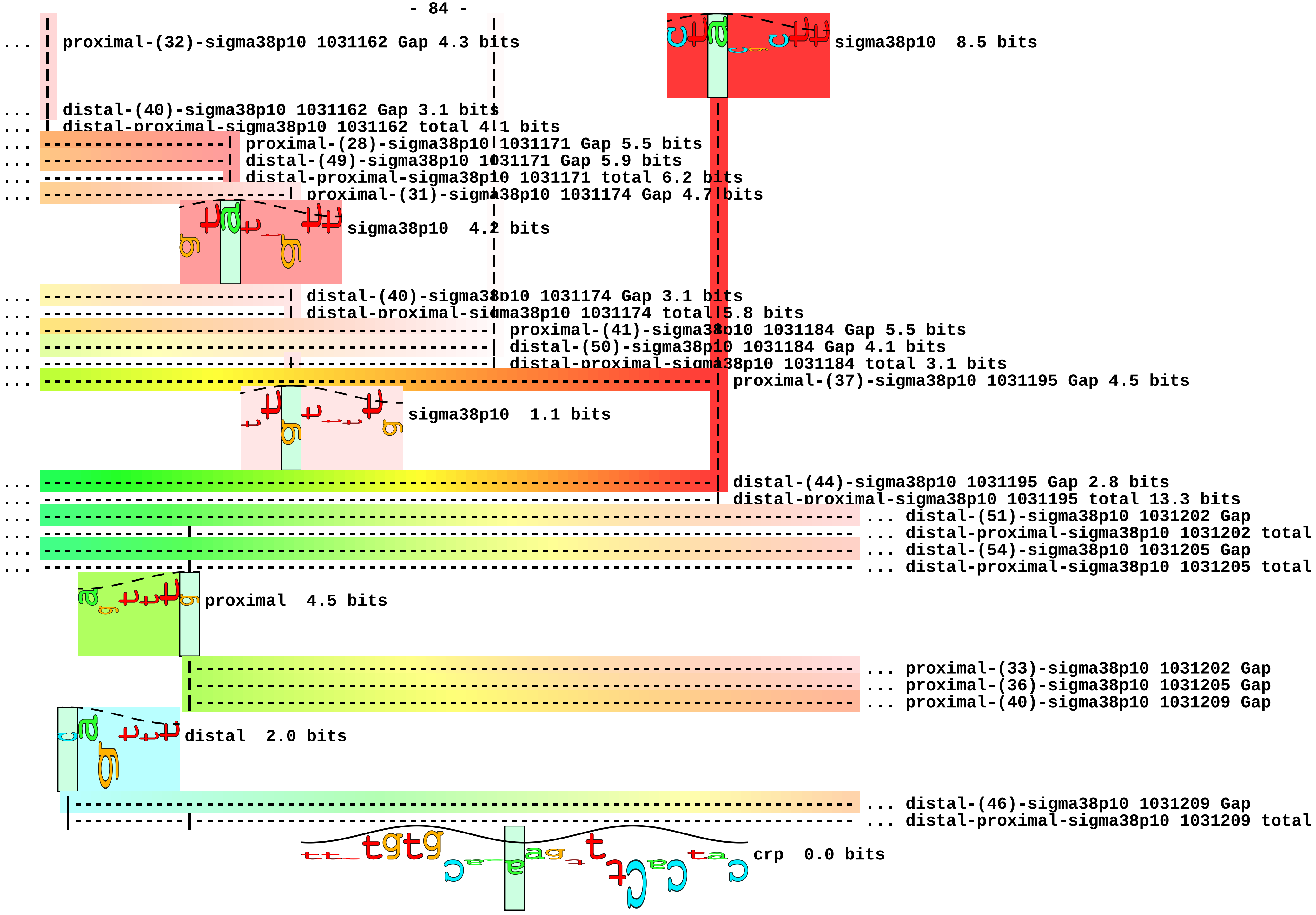
... proximal-(32)-sigma38p10 1031162 Gap 4.3 bits
sigma38p10 8.5 bits
... distal-(40)-sigma38p10 1031162 Gap 3.1 bits
... distal-proximal-sigma38p10 1031162 total 4.1 bits
... proximal-(28)-sigma38p10 1031171 Gap 5.5 bits
... distal-(49)-sigma38p10 1031171 Gap 5.9 bits
... distal-proximal-sigma38p10 1031171 total 6.2 bits
... proximal-(31)-sigma38p10 1031174 Gap 4.7 bits
sigma38p10 4.2 bits
... distal-(40)-sigma38p10 1031174 Gap 3.1 bits
... distal-proximal-sigma38p10 1031174 total 5.8 bits
... proximal-(41)-sigma38p10 1031184 Gap 5.5 bits
... distal-(50)-sigma38p10 1031184 Gap 4.1 bits
... distal-proximal-sigma38p10 1031184 total 3.1 bits
... proximal-(37)-sigma38p10 1031195 Gap 4.5 bits
sigma38p10 1.1 bits
... distal-(44)-sigma38p10 1031195 Gap 2.8 bits
... distal-proximal-sigma38p10 1031195 total 13.3 bits
... ... distal-(51)-sigma38p10 1031202 Gap
... ... distal-proximal-sigma38p10 1031202 total
... ... distal-(54)-sigma38p10 1031205 Gap
... ... distal-proximal-sigma38p10 1031205 total
proximal 4.5 bits
... proximal-(33)-sigma38p10 1031202 Gap
... proximal-(36)-sigma38p10 1031205 Gap
... proximal-(40)-sigma38p10 1031209 Gap
distal 2.0 bits
... distal-(46)-sigma38p10 1031209 Gap
... distal-proximal-sigma38p10 1031209 total
crp 0.0 bits

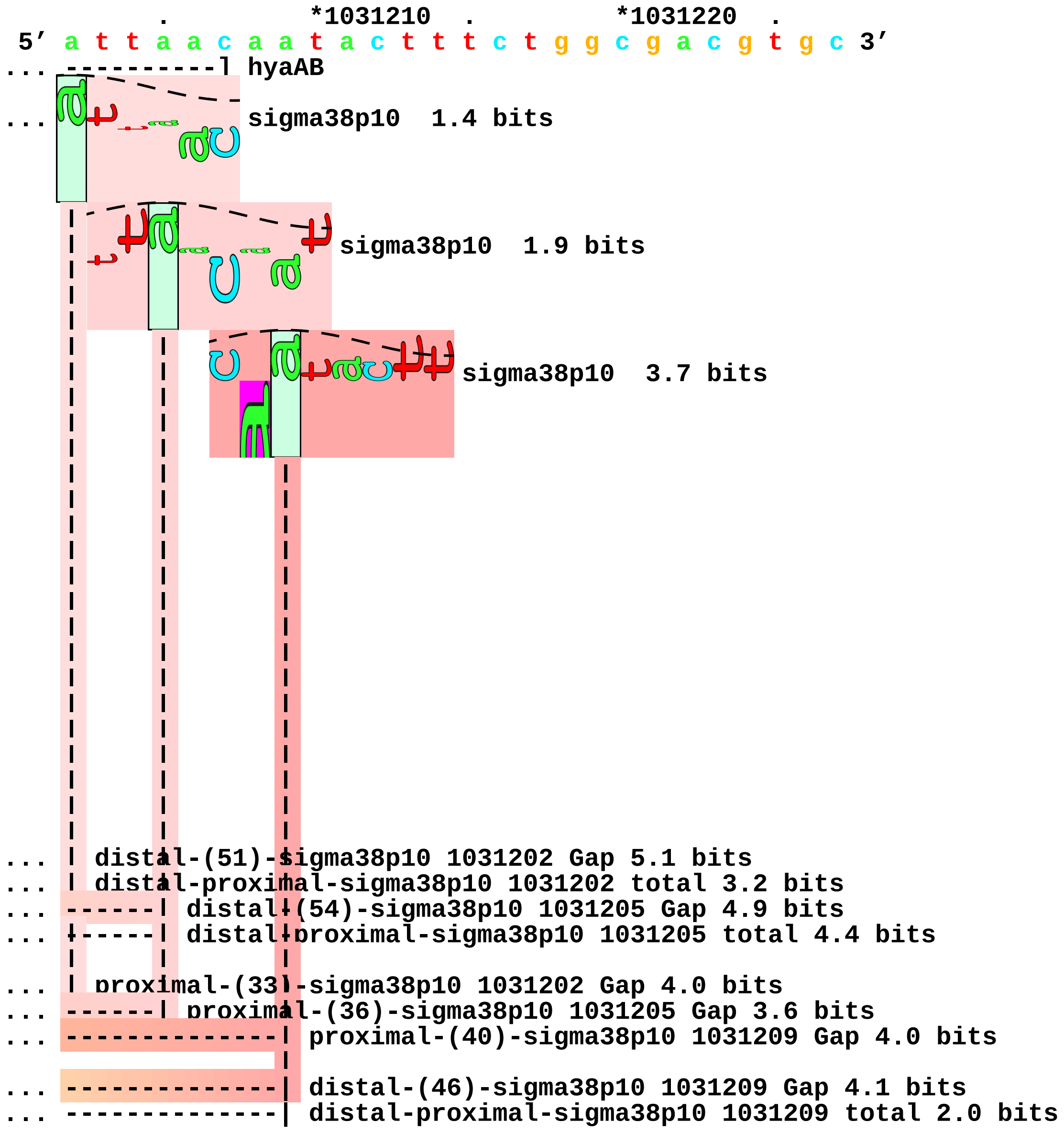
. *1031210 . *1031220 .
5′ a t t a a c a a t a c t t t c t g g c g a c g t g c 3′
... ----------] hyaAB
... sigma38p10 1.4 bits
sigma38p10 1.9 bits
sigma38p10 3.7 bits
... distal-(51)-sigma38p10 1031202 Gap 5.1 bits
... distal-proximal-sigma38p10 1031202 total 3.2 bits
... distal-(54)-sigma38p10 1031205 Gap 4.9 bits
... distal-proximal-sigma38p10 1031205 total 4.4 bits
... proximal-(33)-sigma38p10 1031202 Gap 4.0 bits
... proximal-(36)-sigma38p10 1031205 Gap 3.6 bits
... proximal-(40)-sigma38p10 1031209 Gap 4.0 bits
... distal-(46)-sigma38p10 1031209 Gap 4.1 bits
... distal-proximal-sigma38p10 1031209 total 2.0 bits

piece 41, NC_000913.2, katG, config: linear, direction: +, begin: 4131748, end: 4131855

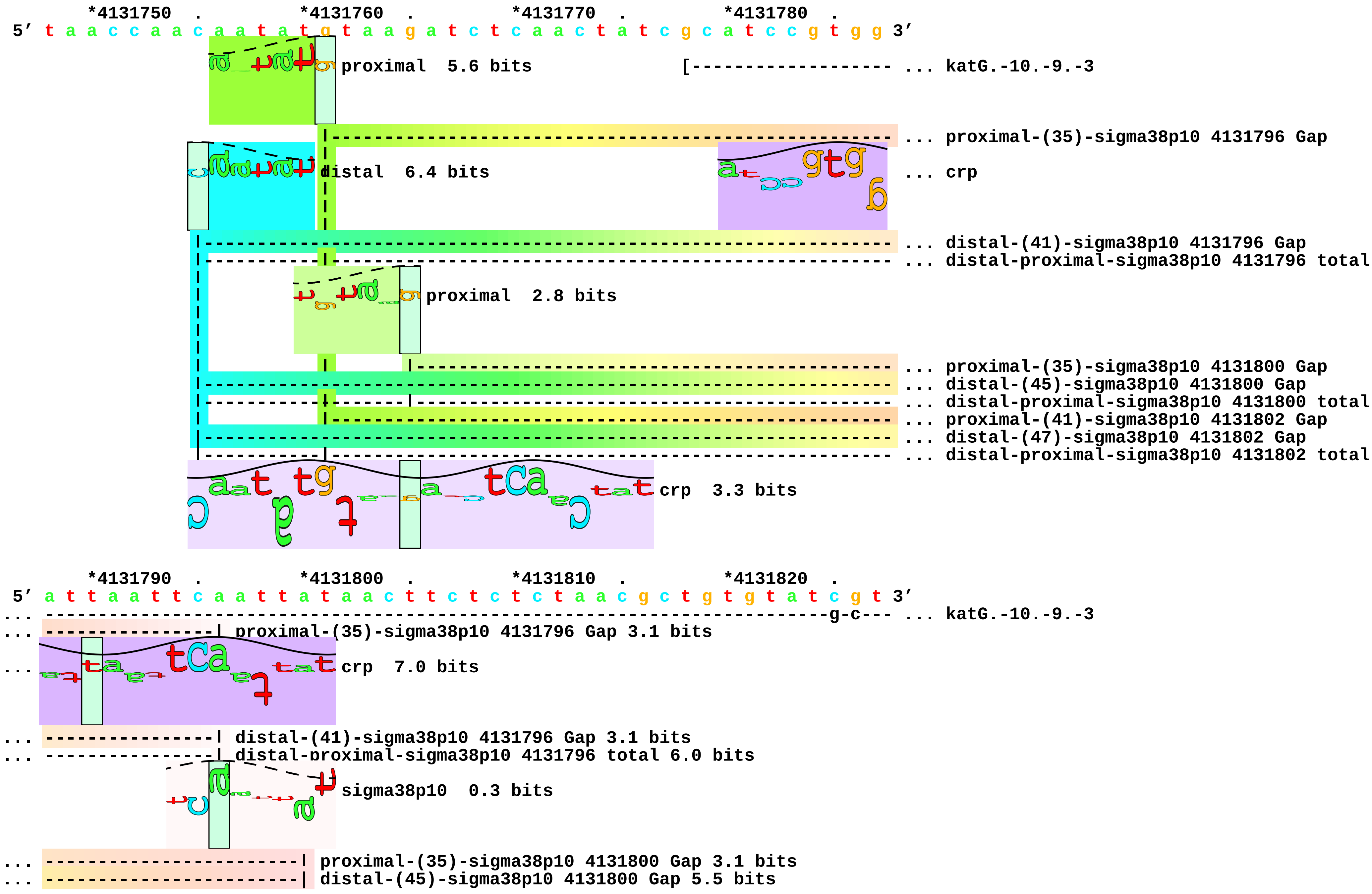

```
... -------------------------| distal-proximal-sigma38p10 4131800 total 2.1 bits
... -----------------------------| proximal-(41)-sigma38p10 4131802 Gap 5.5 bits
... -----------------------------| distal-(47)-sigma38p10 4131802 Gap 6.5 bits
... -------------------------|---| distal-proximal-sigma38p10 4131802 total 1.2 bits
```

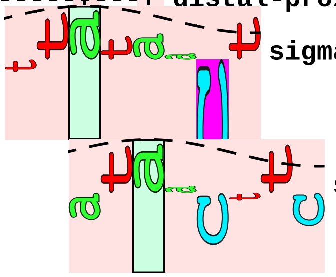

```
                                          sigma38p10  1.4 bits
                                              sigma38p10  1.2 bits

         *4131830  .         *4131840  .         *4131850  .
 5' a a c g g t a a c a c t g t a g a g g g g a g c a c a t 3'
... --------c-----] katG.-10.-9.-3
```

piece 42, NC_000913.2, osmB-P2, config: linear, direction: -, begin: 1341480, end: 1341375

```
   *1341480  .         *1341470  .         *1341460  .         *1341450  .
5' a t t a t c a c c t g c g t g a t a t a a c c c t g c g c g a g c a g a t t 3'
                                         distal  5.9 bits   [------------------- ... osmB-P2
                                           proximal  3.9 bits
                                                                                 ... proximal-(37)-sigma38p10 1341422 Gap
                                                                                 ... distal-(44)-sigma38p10 1341422 Gap
                                                                                 ... sigma38p10-proximal-distal 1341422 total
                                                                                 ... proximal-(43)-sigma38p10 1341416 Gap
                                                                                 ... distal-(50)-sigma38p10 1341416 Gap
                                                                                 ... sigma38p10-proximal-distal 1341416 total
                                                                                 ... crp

   *1341440  .         *1341430  .         *1341420  .         *1341410  .
5' t c a c g g a a t a a t t t c a c c a g a c t t a t t c t t a g c t a t t a t a 3'
... ---------------------------------------------------------------------------- ... osmB-P2
                                          sigma38p10  4.4 bits                   ... crp
... ------------------------------------ proximal-(37)-sigma38p10 1341422 Gap 4.5 bits
... ------------------------------------ distal-(44)-sigma38p10 1341422 Gap 2.8 bits
... ------------------------------------ sigma38p10-proximal-distal 1341422 total 6.8 bits
... ------------------------------------------------ proximal-(43)-sigma38p10 1341416 Gap 5.1 bits
... ------------------------------------------------ distal-(50)-sigma38p10 1341416 Gap 4.1 bits
... ------------------------------------------------ sigma38p10-proximal-distal 1341416 total 8.5 bits
... crp  3.8 bits                                            sigma38p10  8.1 bits

   *1341400  .         *1341390  .         *1341380  .
5' g t t a t a g a g a g c t t a c t t c c g t g a a t 3'
... ----------] osmB-P2
...                                     crp  2.1 bits
```

piece 43, NC_000913.2, osmC-P2, config: linear, direction: +, begin: 1554543, end: 1554650

```
        .         *1554550    .         *1554560    .         *1554570    .         *1554580
5' g c a g a a g t a a t a t t t t c g c c g g a t t t t a t t c g g a a t a t c c 3'
                                                               a-t-t-c-g-g---a-t-a- ... osmC-P2.-57.-56.-55.-54.-53.-52.-50.-49.
                                                                    proximal  6.1 bits
                                                                    --- ... proximal-(34)-sigma38p10 1554615 Gap
                                                                   distal  7.2 bits
                                                               |-------------- ... distal-(40)-sigma38p10 1554615 Gap
                                                               |-----------|--- ... distal-proximal-sigma38p10 1554615 total
                                                                           |--- ... proximal-(36)-sigma38p10 1554617 Gap
                                                               |-------------- ... distal-(42)-sigma38p10 1554617 Gap
                                                               |-----------|--- ... distal-proximal-sigma38p10 1554617 total

        .         *1554590    .         *1554600    .         *1554610    .         *1554620
5' t g c t t a t c c t c g t g c t g t t t c t c a c g t a g t c t a t a a t t t c 3'
... --c-----g-c---t-a------------------------------------------------------------ ... osmC-P2.-57.-56.-55.-54.-53.-52.-50.-49.
                                                                   sigma38p10 10.4 bits
... ------------------------------------------------------------| proximal-(34)-sigma38p10 1554615 Gap 2.5 bits
                                                                      sigma38p10  3.0 bits
... ------------------------------------------------------------| distal-(40)-sigma38p10 1554615 Gap 3.1 bits
... ------------------------------------------------------------| distal-proximal-sigma38p10 1554615 total 18.1 bits
... ----------------------------------------------------------------| proximal-(36)-sigma38p10 1554617 Gap 3.6 bits
... ----------------------------------------------------------------| distal-(42)-sigma38p10 1554617 Gap 3.3 bits
... ----------------------------------------------------------------| distal-proximal-sigma38p10 1554617 total 9.4 bits
                                                          crp  1.6 bits

        .         *1554630    .         *1554640    .         *1554650
5' c t t t t t a a g c c c a c a g g a g a g c a a c a a t 3'
... -------------] osmC-P2.-57.-56.-55.-54.-53.-52.-50.-49.-48.-46.-43.-42.-40.-39
```

piece 44, NC_000913.2, osmE, config: linear, direction: -, begin: 1820392, end: 1820287

```
      *1820390  .         *1820380  .         *1820370  .         *1820360  .
5' a a a a c g t c a t a c a g g a a t c c t c t g a a a g c g g a g c c g t t t c 3'
                                                                 a-g-c---g-t-----c-a- ... osmE.-55.-54.-53.-51.-50.-47.-46.-45.-43
                                                                                      ... crp
```

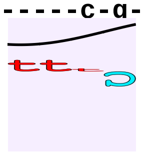

```
      *1820350  .         *1820340  .         *1820330  .         *1820320  .
5' g t t c a c g g g c t t g a a a a a g c g c c c a a t g t a t t c c a g g c t t 3'
... t---c-a-c-a----c--------------------------------------------------------------- ... osmE.-55.-54.-53.-51.-50.-47.-46.-45.-43
...                      crp  1.7 bits
                                   crp  0.0 bits
```

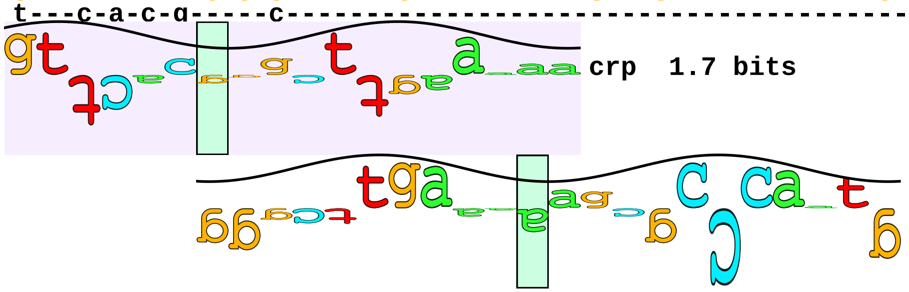

```
      *1820310  .         *1820300  .         *1820290
5' a t c t a a c a c g c t g a t a a a c a a g a g g a 3'
... ----------] osmE.-55.-54.-53.-51.-50.-47.-46.-45.-43.-42.-41.-40.-37
```

piece 45, NC_000913.2, otsBA, config: linear, direction: -, begin: 1980550, end: 1980446

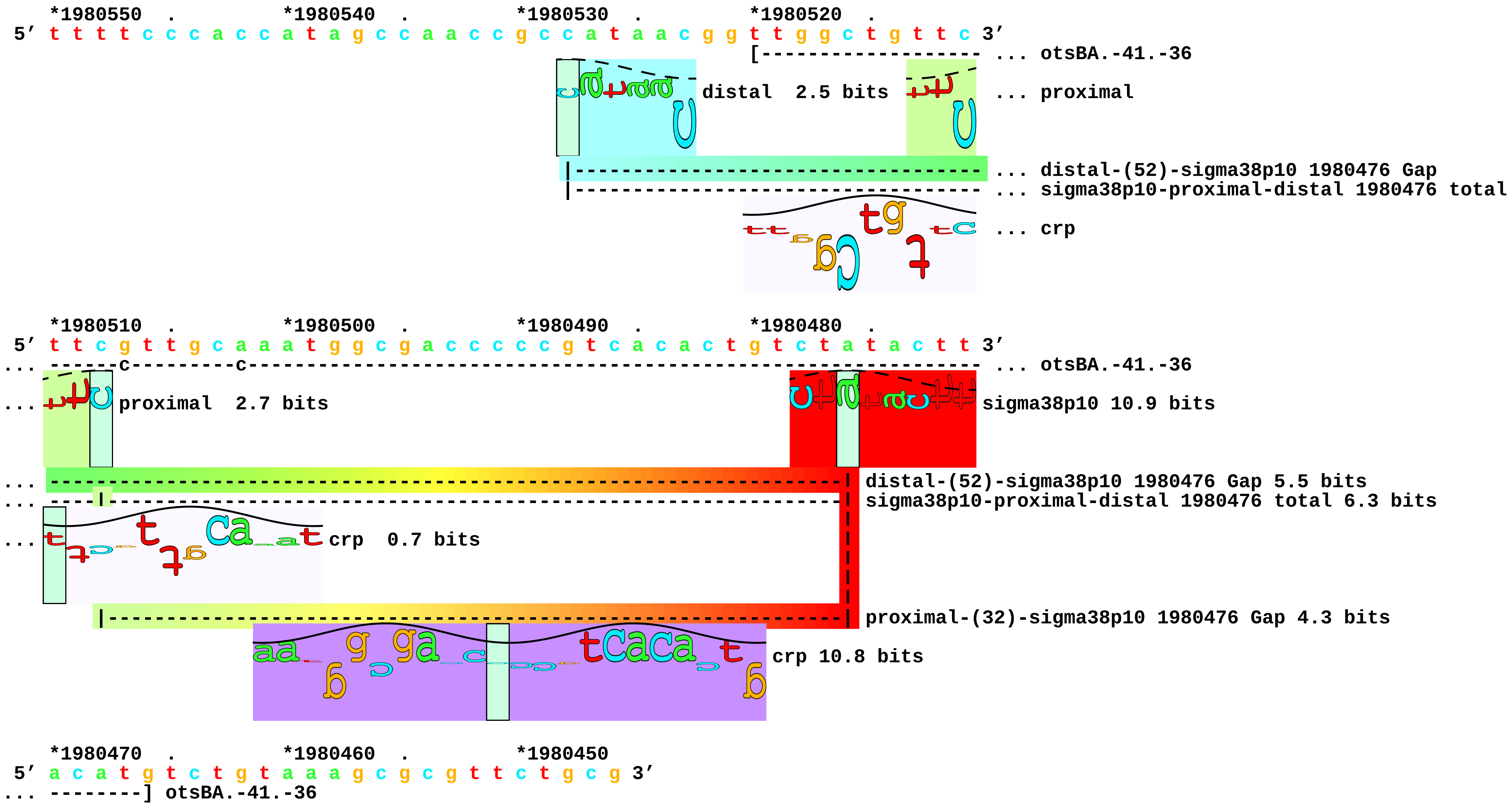

piece 46, NC_000913.2, poxB, config: linear, direction: -, begin: 910384, end: 910279

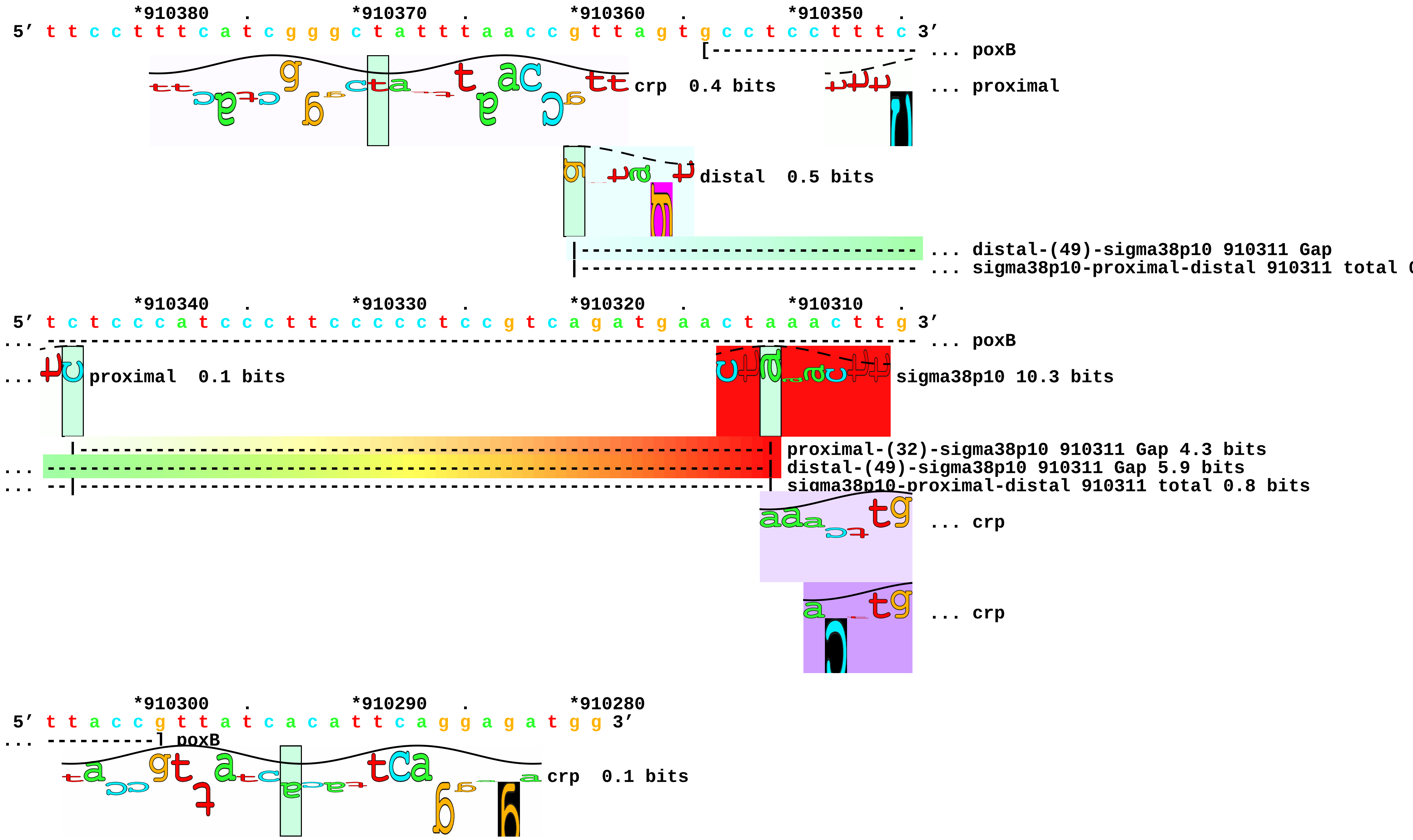

... 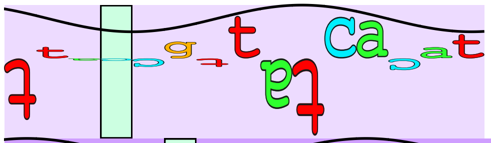 crp 3.4 bits

... 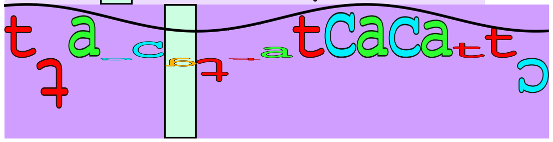 crp 9.3 bits piece 47, NC_000913.2, pqi5-P2, config: linear, direction: +, begin: 1010803, end: 1010909

```
      .         *1010810  .         *1010820  .         *1010830  .         *1010840
5' t g c g g t g a a a a a a a a c c g a a g a a g c c g c c g c g g c a a a a g c a 3'
```

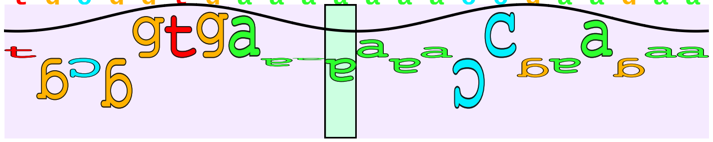

crp 2.0 bits [------------------- ... pqi5-P2

```
      .         *1010850  .         *1010860  .         *1010870  .         *1010880
5' g a a a c t g t a a a a c g c a g c a g t a g c a a a c t a a g c t a t a a a t 3'
... -------------------------------------------------------------------------------- ... pqi5-P2
```

```
      .         *1010890  .         *1010900  .
5' t g c a g c g c g a a c t g g a g c a g c t a c c g c 3'
... ------------] pqi5-P2
```

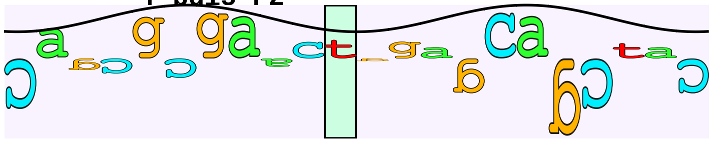

crp 1.3 bits

piece 48, NC_000913.2, proU-P1, config: linear, direction: +, begin: 2802498, end: 2802606

```
       *2802500  .         *2802510  .         *2802520  .         *2802530  .
5' a a a a t a t c a c t a c c c g c a g c a g g g a a a t a a t t c c c g c c a a 3'
              proximal  6.1 bits                         [-a----------------- ... proU-P1.-57.-3.-2.-1.
                                                                              ... proximal-(35)-sigma38p10 2802540 Gap
            distal  0.4 bits                    distal  5.9 bits              ... proximal
                                                                              ... distal-(41)-sigma38p10 2802540 Gap
                                                                              ... distal-proximal-sigma38p10 2802540 total
                                                                              ... distal-(53)-sigma38p10 2802574 Gap
                                                                              ... distal-proximal-sigma38p10 2802574 total
                                                                              ... distal
                                                                              ... distal-(45)-sigma38p10 2802580 Gap
                                                                              ... distal-proximal-sigma38p10 2802580 total
                                                                              ... crp
                                                                              ... crp

       *2802540  .         *2802550  .         *2802560  .         *2802570  .
5' a t a g c t t t t t a t c a c g c a a a t a a t t t g t g g t g a t c t a c a c 3'
... ---------------------------------------------------------------------------- ... proU-P1.-57.-3.-2.-1.
... ----| proximal-(35)-sigma38p10 2802540 Gap 3.1 bits                          ... sigma38p10
...        proximal  3.3 bits                                                    ... crp
... ----| distal-(41)-sigma38p10 2802540 Gap 3.1 bits
... ----| distal-proximal-sigma38p10 2802540 total 1.2 bits
```

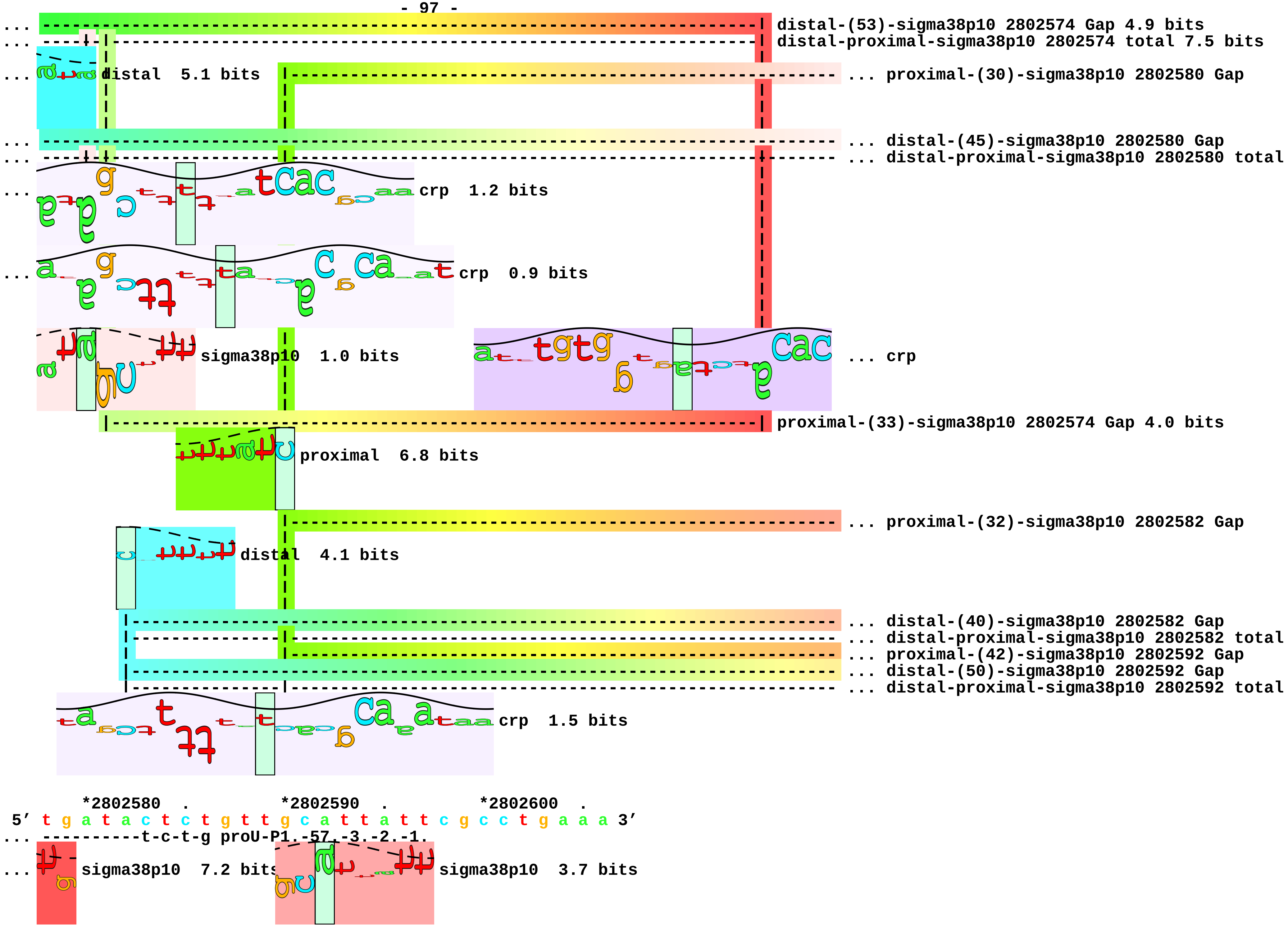
distal-(53)-sigma38p10 2802574 Gap 4.9 bits
distal-proximal-sigma38p10 2802574 total 7.5 bits
distal 5.1 bits
... proximal-(30)-sigma38p10 2802580 Gap
... distal-(45)-sigma38p10 2802580 Gap
... distal-proximal-sigma38p10 2802580 total
crp 1.2 bits
crp 0.9 bits
sigma38p10 1.0 bits
... crp
proximal-(33)-sigma38p10 2802574 Gap 4.0 bits
proximal 6.8 bits
... proximal-(32)-sigma38p10 2802582 Gap
distal 4.1 bits
... distal-(40)-sigma38p10 2802582 Gap
... distal-proximal-sigma38p10 2802582 total
... proximal-(42)-sigma38p10 2802592 Gap
... distal-(50)-sigma38p10 2802592 Gap
... distal-proximal-sigma38p10 2802592 total
crp 1.5 bits
*2802580 . *2802590 . *2802600 .
5′ t g a t a c t c t g t t g c a t t a t t c g c c t g a a a 3′
... ----------t-c-t-g proU-P1 -57 -3 -2 -1
sigma38p10 7.2 bits
sigma38p10 3.7 bits

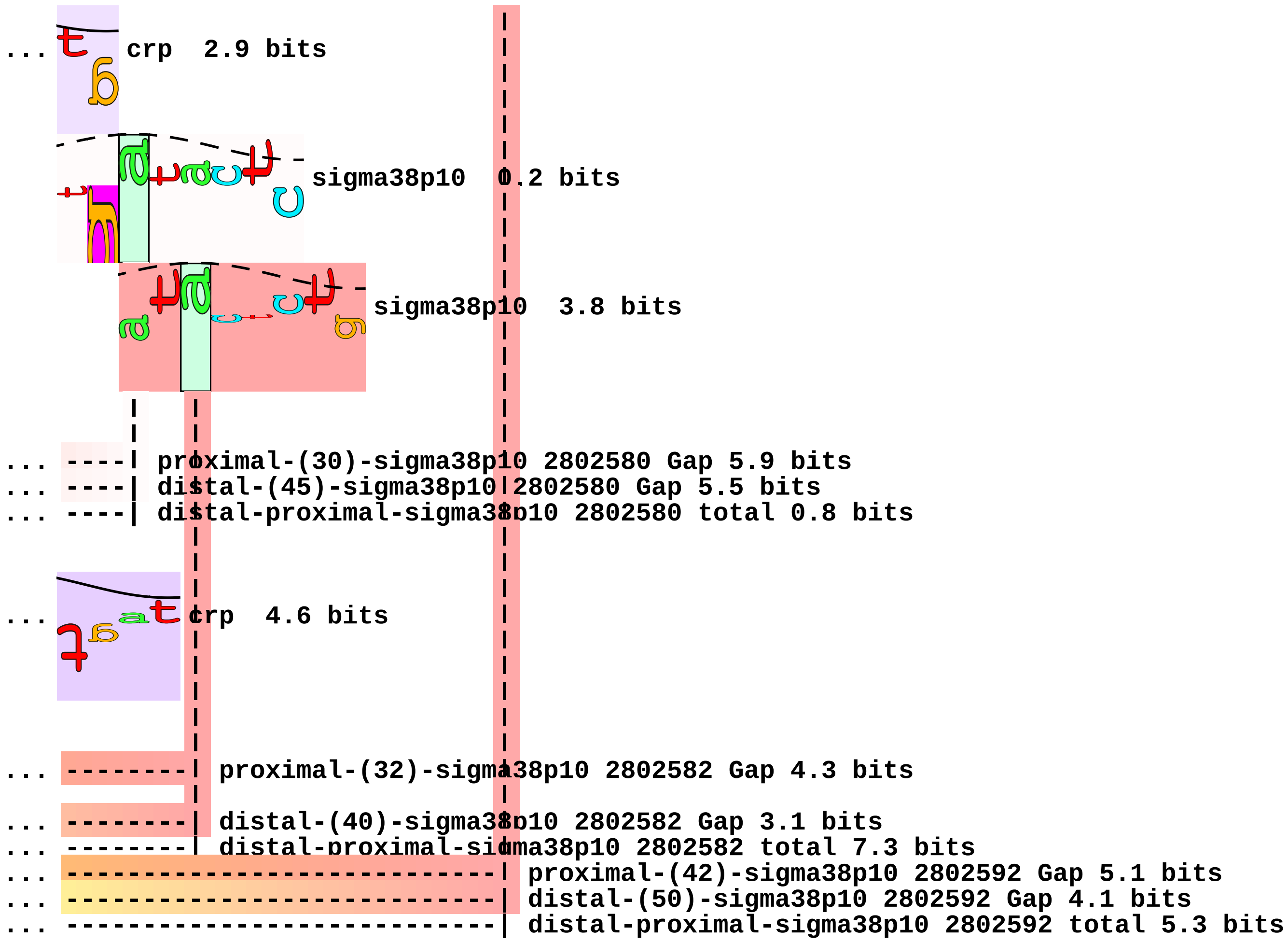
... crp 2.9 bits
sigma38p10 0.2 bits
sigma38p10 3.8 bits
... ----| proximal-(30)-sigma38p10 2802580 Gap 5.9 bits
... ----| distal-(45)-sigma38p10 2802580 Gap 5.5 bits
... ----| distal-proximal-sigma38p10 2802580 total 0.8 bits
... crp 4.6 bits
... --------| proximal-(32)-sigma38p10 2802582 Gap 4.3 bits
... --------| distal-(40)-sigma38p10 2802582 Gap 3.1 bits
... --------| distal-proximal-sigma38p10 2802582 total 7.3 bits
... --------------------------| proximal-(42)-sigma38p10 2802592 Gap 5.1 bits
... --------------------------| distal-(50)-sigma38p10 2802592 Gap 4.1 bits
... --------------------------| distal-proximal-sigma38p10 2802592 total 5.3 bits

```
piece 49, NC_000913.2, rssAB, config: linear, direction: +, begin: 1288241, end: 1288349

              .         *1288250  .         *1288260  .         *1288270  .         *1288280
5' a g t a a c g c g a a c g c a t g a g a t g t t c t g g a t c a g g t g c a a c 3'
                                                                ┌------------------- ... rssAB.-27.-24
                                                                                     ... crp
                                                                                     ... crp

              .         *1288290  .         *1288300  .         *1288310  .         *1288320
5' c t t t t c a c c a g a c a c a t a a g g g t g g c a a c a t a g g c t a t a c 3'
... -------------------------------------------c-----c-------------------------------- ... rssAB.-27.-24
...    crp  0.0 bits
...                                      crp  1.8 bits

              .         *1288330  .         *1288340  .
5' t c g a c a g c a c t a c c a c a g g g a c a a a g c t g 3'
... ----------------] rssAB.-27.-24
```

```
piece 50, NC_000913.2, topA-P1, config: linear, direction: +, begin: 1328727, end: 1328833

        *1328730   .       *1328740   .       *1328750   .       *1328760   .
5' c a g g c a t g g c t t g c a g a c a a a t a t a c c a c g c t g g t g g c a a 3'
                                                      [------------------- ... topA-P1
                                         proximal  4.9 bits
                                                        |----------------------------- ... proximal-(36)-sigma38p10 1328788 Gap
                                   distal  5.1 bits
                                    |------------------------------------------------ ... distal-(44)-sigma38p10 1328788 Gap
                                    |---------------|-------------------------------- ... distal-proximal-sigma38p10 1328788 total

        *1328770   .       *1328780   .       *1328790   .       *1328800   .
5' g a g c g c c t t a c t g g c a a c t t t g g a t t t t g c a t g c t a a t a a 3'
... ------------------------------------------------------------------------------ ... topA-P1
                                              sigma38p10  1.1 bits
... ------------------------------------------| proximal-(36)-sigma38p10 1328788 Gap 3.6 bits
                                                                                    ... crp
... ------------------------------------------| distal-(44)-sigma38p10 1328788 Gap 2.8 bits
... ------------------------------------------| distal-proximal-sigma38p10 1328788 total 4.7 bits

        *1328810   .       *1328820   .       *1328830
5' a g t t g c g t a t c g g a t t t t a t c a g g t a c 3'
... ------------] topA-P1

...       crp  1.5 bits
```

piece 51, NC_000913.2, treA, config: linear, direction: -, begin: 1246730, end: 1246625

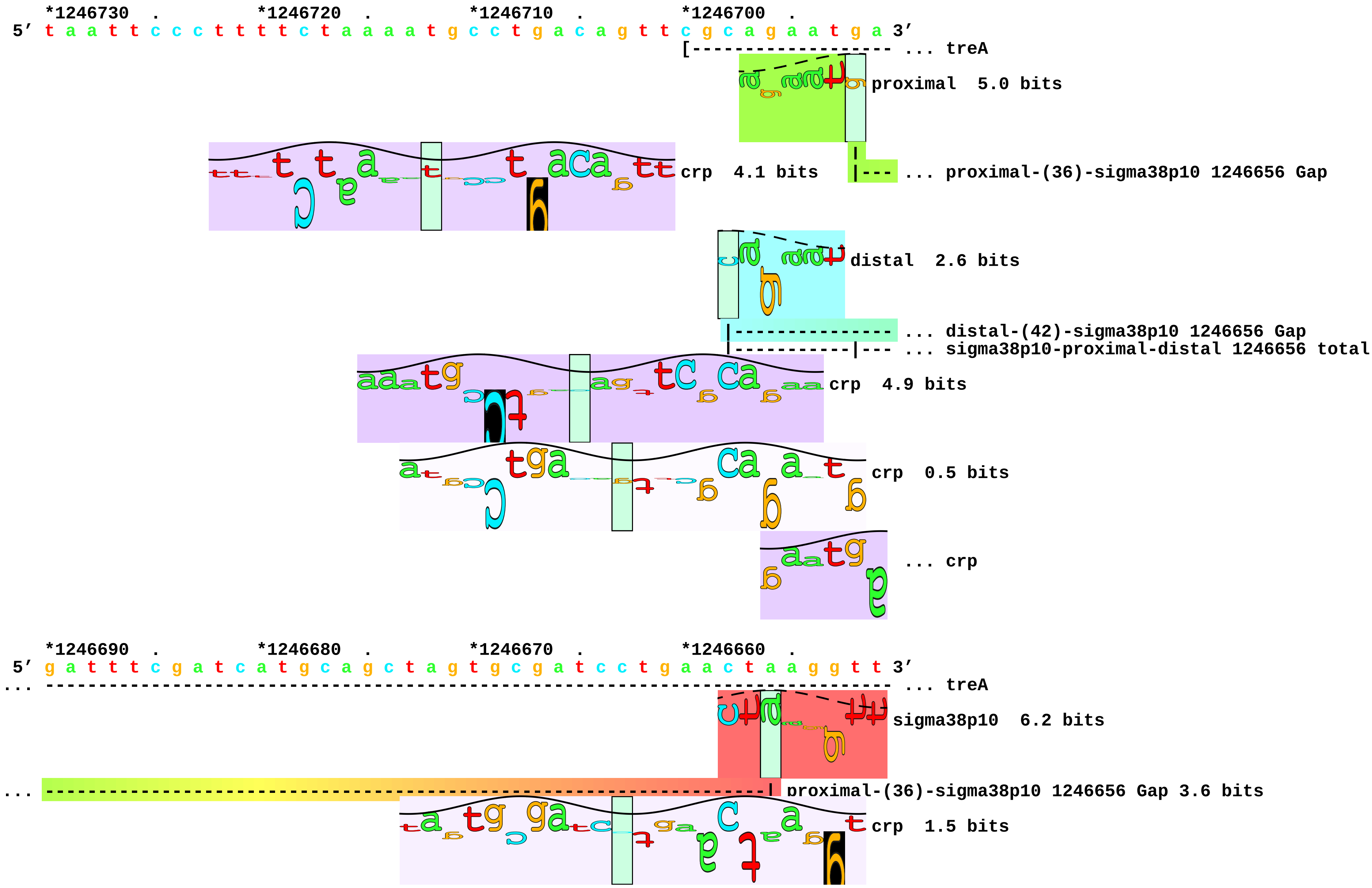

```
... ---------------------------------------------------------------------| distal-(42)-sigma38p10 1246656 Gap 3.3 bits
... ---------------------------------------------------------------------| sigma38p10-proximal-distal 1246656 total 7.0 bits
```

... 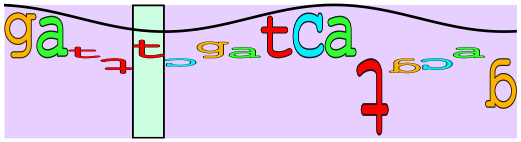 crp 4.6 bits

```
    *1246650  .         *1246640  .         *1246630  .
5' t t c t g a t a c t t g a a t a c c g t t t t a t t 3'
... ----------] treA
```

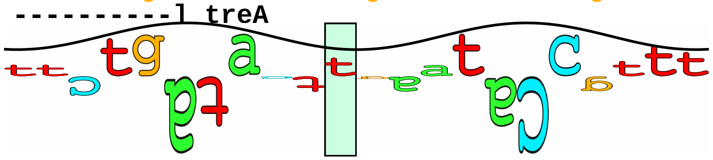 crp 0.1 bits
