## Supplement_for_sigma38_paper-directory for "*Escherichia coli σ*^38^ promoters use two UP elements instead of a −35 element: resolution of a paradox and discovery that *σ*^38^ transcribes ribosomal promoters": Typas.Hengge2005-table1-map.pdf

piece 1, NC_000913, rrnBP1

```
   *3939460  .       *3939470  .       *3939480  .       *3939490  .       *3939500  .       *3939510  .       *3939520  .       *3939530  .       *3939540  .       *3939550
5' t g c t g a a a a a a t g c g c g g t c a g a a a a t t a t t t t t a a a t t t c c t c t t g t c a g g c c g g a a t a a c t c c c t a t a a t g c g c c a c c a c t g a c a c g g a a 3'
```

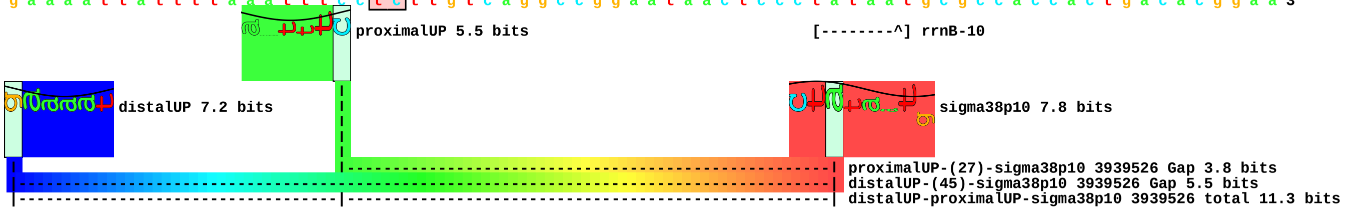


piece 2, NC_000913.i3939500,3939503ca, rrnBP1

```
   *3939460  .       *3939470  .       *3939480  .       *3939490  .       *3939500  .       *3939510  .       *3939520  .       *3939530  .       *3939540  .       *3939550
5' t g c t g a a a a a a t g c g c g g t c a g a a a a t t a t t t t t a a a t t t c c c a t t g t c a g g c c g g a a t a a c t c c c t a t a a t g c g c c a c c a c t g a c a c g g a a 3'
```

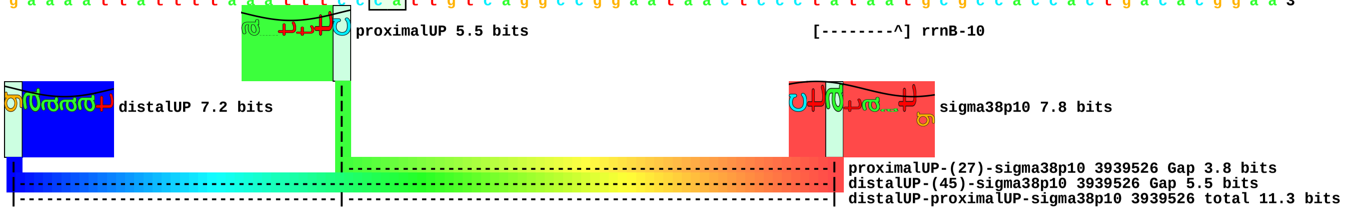

piece 3, NC_000913, bolAP1

```
   *453580    .         *453590    .         *453600    .         *453610    .         *453620    .         *453630    .         *453640    .         *453650    .         *453660    .         *453670
5' a c g g a a a a a c c a g g a c g a a a c c t a a a t a t t t g t t g t t a a g c t g c a a t g g a a a c g g t a a a a g c g g c t a g t a t t t a a a g g g a t g g a t g a c a t c 3'
```

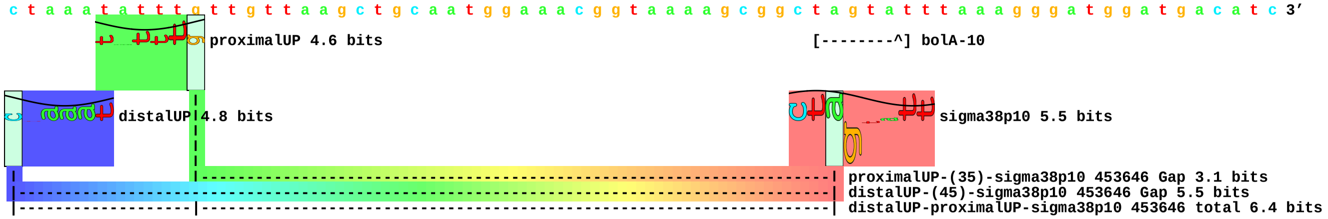


piece 4, NC_000913.a453604g.t453606c.t453608g.t453609c, bolAP1

```
   *453580    .         *453590    .         *453600    .         *453610    .         *453620    .         *453630    .         *453640    .         *453650    .         *453660    .         *453670
5' a c g g a a a a a c c a g g a c g a a a c c t a g a c a g c t g t t g t t a a g c t g c a a t g g a a a c g g t a a a a g c g g c t a g t a t t t a a a g g g a t g g a t g a c a t c 3'
                                                                                                                                                          [--------^] bolA-10
```

piece 5, NC_000913.t453613a.t453615a.t453616a, bolAP1

```
   *453580    .         *453590    .         *453600    .         *453610    .         *453620    .         *453630    .         *453640    .         *453650    .         *453660    .         *453670
5' a c g g a a a a a c c a g g a c g a a a c c t a a a t a t t t g t a g a a a a g c t g c a a t g g a a a c g g t a a a a g c g g c t a g t a t t t a a a g g g a t g g a t g a c a t c 3'
```

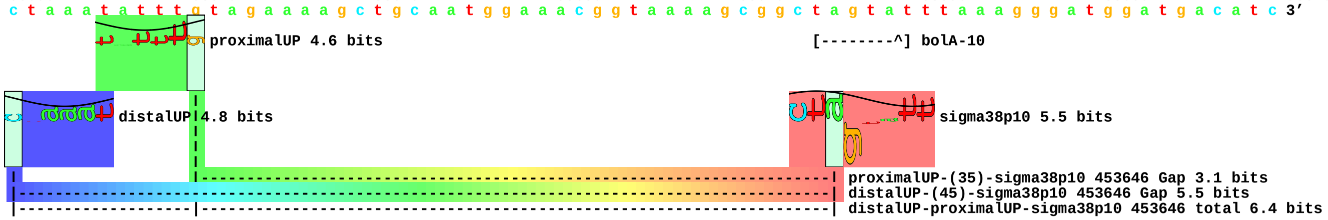

piece 6, NC_000913, csiE

```
           *2663350  .       *2663360  .       *2663370  .       *2663380  .       *2663390  .       *2663400  .       *2663410  .       *2663420  .       *2663430  .
5' a g a a a a a t t c t g t g a c g c t t g c c a a c a t t t c t g a t g a t t a g c a t t c c c t t c g c c a t t t c c t t g a g c a a a c t t t a g c t a t t c t t a t c a a t t a 3'
```

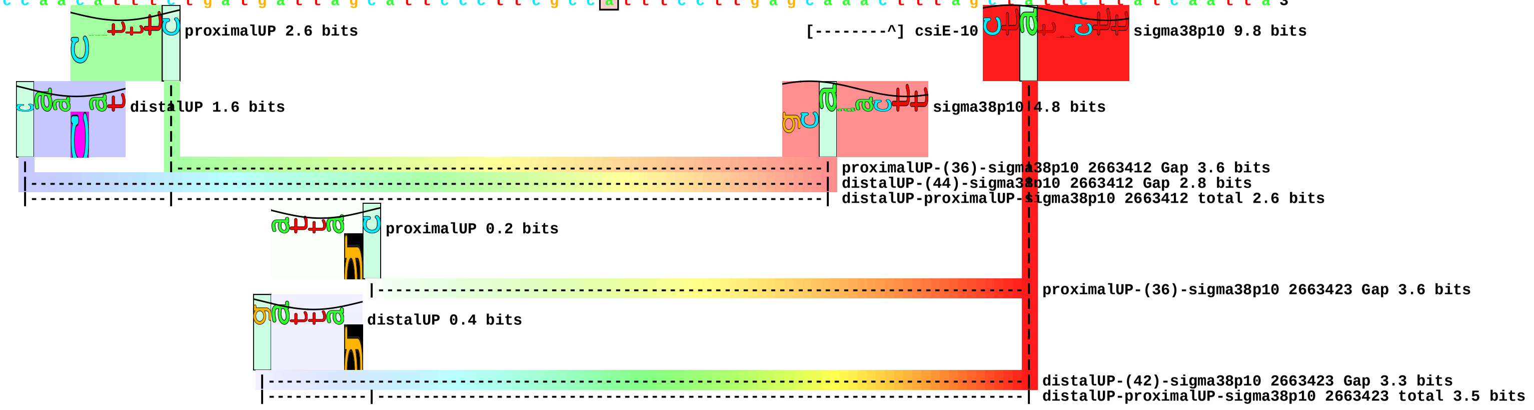


piece 7, NC_000913.t2663373g.t2663377g.i2663399,2663401a, csiE

```
           *2663350  .       *2663360  .       *2663370  .       *2663380  .       *2663390  .       *2663400  .       *2663410  .       *2663420  .       *2663430  .
5' a g a a a a a t t c t g t g a c g c t t g c c a a c a g t t c g g a t g a t t a g c a t t c c c t t c g c c a t t t c c t t g a g c a a a c t t t a g c t a t t c t t a t c a a t t a 3'
```

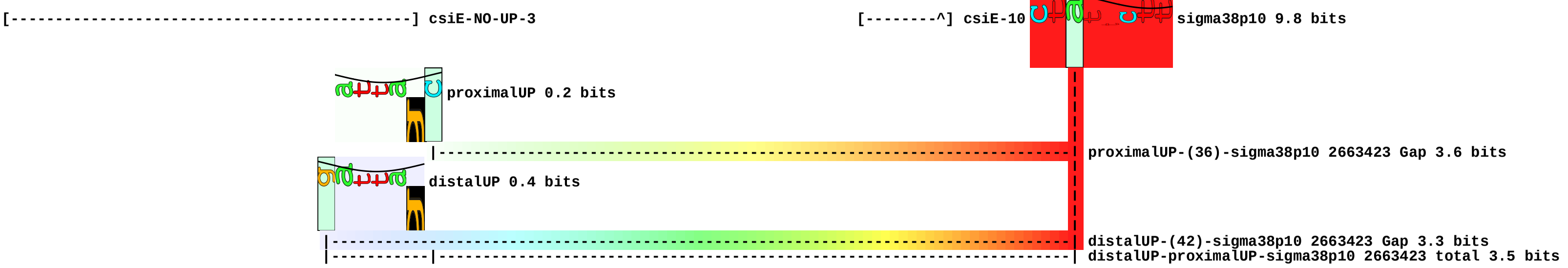


piece 8, NC_000913.g2663381a.t2663383a.t2663384a.i2663399,2663401a, csiE

```
           *2663350  .       *2663360  .       *2663370  .       *2663380  .       *2663390  .       *2663400  .       *2663410  .       *2663420  .       *2663430  .
5' a g a a a a a t t c t g t g a c g c t t g c c a a c a t t t c t g a t a a a a a g c a t t c c c t t c g c c a t t t c c t t g a g c a a a c t t t a g c t a t t c t t a t c a a t t a 3'
```

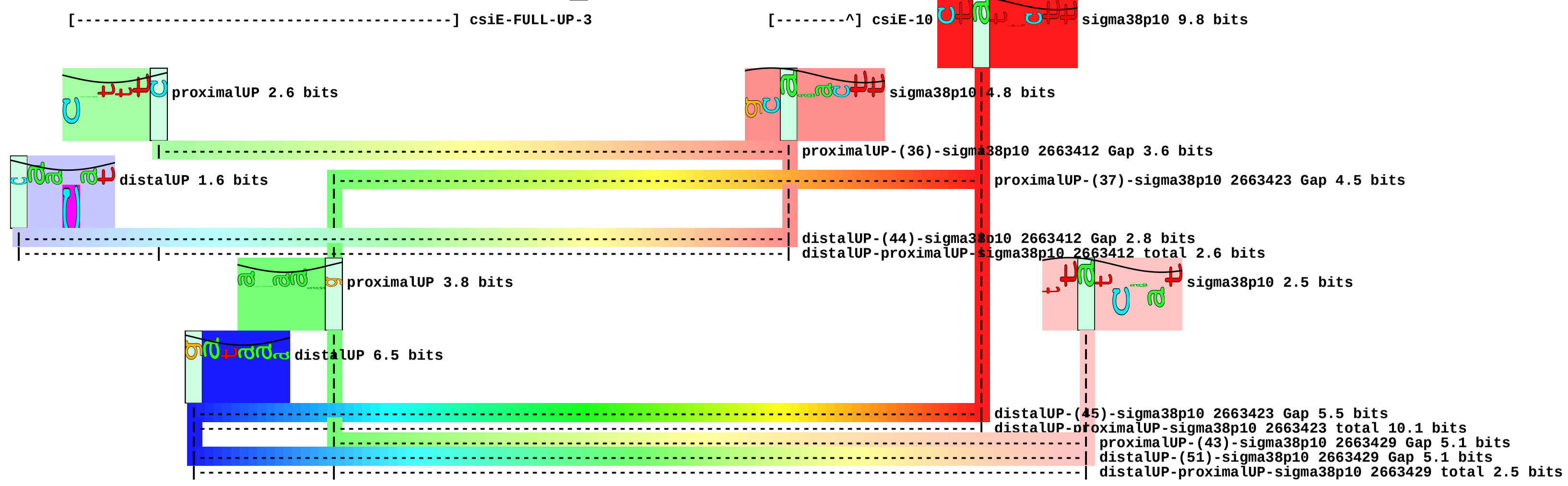

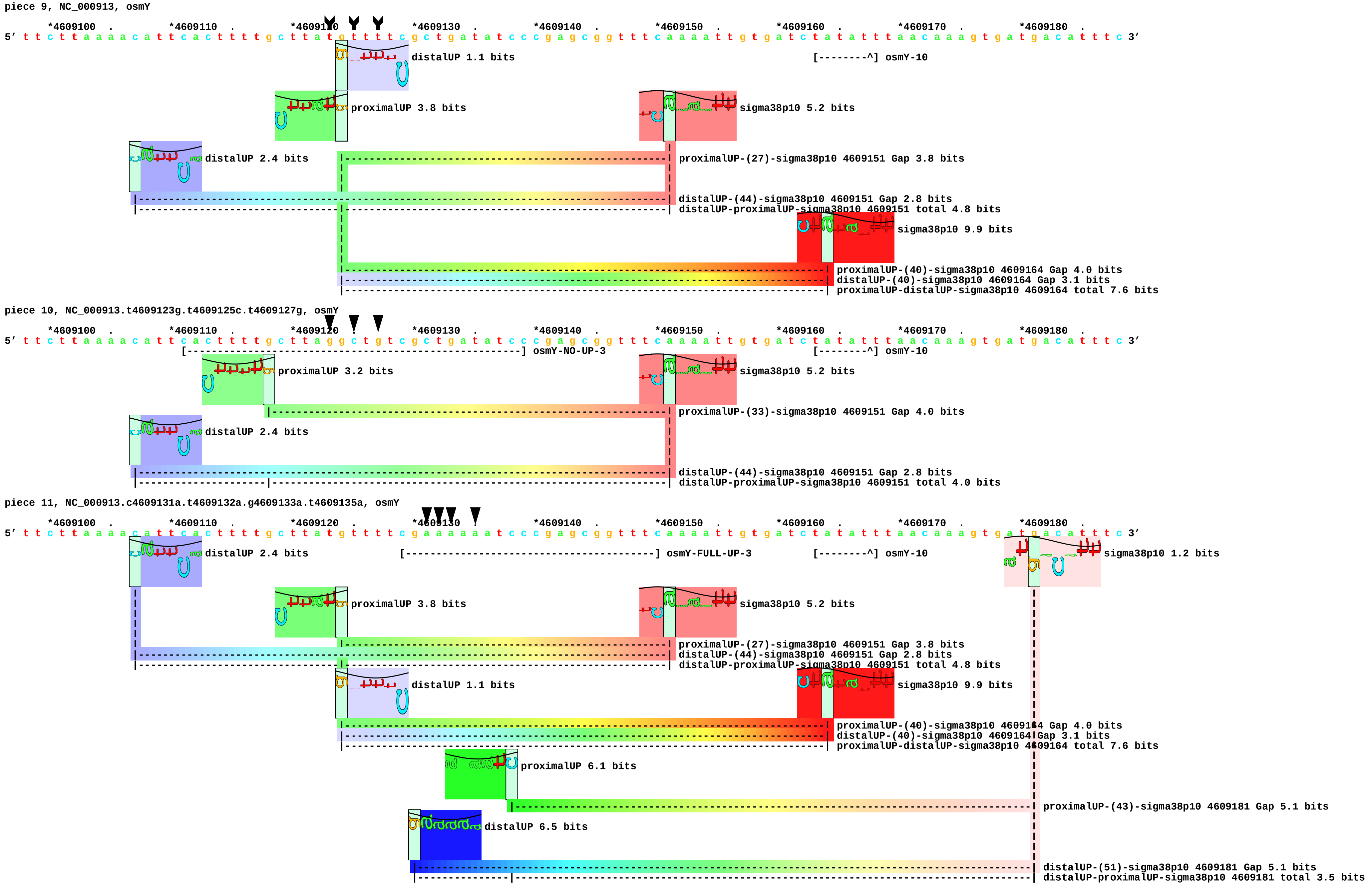
piece 9, NC_000913, osmY
*4609100 . *4609110 . *4609120 . *4609130 . *4609140 . *4609150 . *4609160 . *4609170 . *4609180 .
5' t t c t t a a a a c a t t c a c t t t t g c t t a t g t t t t c g c t g a t a t c c c g a g c g g t t t c a a a a t t g t g a t c t a t a t t t a a c a a a g t g a t g a c a t t t c 3'
distalUP 1.1 bits
[--------^] osmY-10
proximalUP 3.8 bits
sigma38p10 5.2 bits
distalUP 2.4 bits
proximalUP-(27)-sigma38p10 4609151 Gap 3.8 bits
distalUP-(44)-sigma38p10 4609151 Gap 2.8 bits
distalUP-proximalUP-sigma38p10 4609151 total 4.8 bits
sigma38p10 9.9 bits
proximalUP-(40)-sigma38p10 4609164 Gap 4.0 bits
distalUP-(40)-sigma38p10 4609164 Gap 3.1 bits
proximalUP-distalUP-sigma38p10 4609164 total 7.6 bits
piece 10, NC_000913.t4609123g.t4609125c.t4609127g, osmY
*4609100 . *4609110 . *4609120 . *4609130 . *4609140 . *4609150 . *4609160 . *4609170 . *4609180 .
5' t t c t t a a a a c a t t c a c t t t t g c t t a g g c t g t c g c t g a t a t c c c g a g c g g t t t c a a a a t t g t g a t c t a t a t t t a a c a a a g t g a t g a c a t t t c 3'
[--------------------------------------------------------] osmY-NO-UP-3
[--------^] osmY-10
proximalUP 3.2 bits
sigma38p10 5.2 bits
proximalUP-(33)-sigma38p10 4609151 Gap 4.0 bits
distalUP 2.4 bits
distalUP-(44)-sigma38p10 4609151 Gap 2.8 bits
distalUP-proximalUP-sigma38p10 4609151 total 4.0 bits
piece 11, NC_000913.c4609131a.t4609132a.g4609133a.t4609135a, osmY
*4609100 . *4609110 . *4609120 . *4609130 . *4609140 . *4609150 . *4609160 . *4609170 . *4609180 .
5' t t c t t a a a a c a t t c a c t t t t g c t t a t g t t t t c g a a a a a a t c c c g a g c g g t t t c a a a a t t g t g a t c t a t a t t t a a c a a a g t g a t g a c a t t t c 3'
distalUP 2.4 bits
[-------------------------------------------] osmY-FULL-UP-3
[--------^] osmY-10
sigma38p10 1.2 bits
proximalUP 3.8 bits
sigma38p10 5.2 bits
proximalUP-(27)-sigma38p10 4609151 Gap 3.8 bits
distalUP-(44)-sigma38p10 4609151 Gap 2.8 bits
distalUP-proximalUP-sigma38p10 4609151 total 4.8 bits
distalUP 1.1 bits
sigma38p10 9.9 bits
proximalUP-(40)-sigma38p10 4609164 Gap 4.0 bits
distalUP-(40)-sigma38p10 4609164 Gap 3.1 bits
proximalUP-distalUP-sigma38p10 4609164 total 7.6 bits
proximalUP 6.1 bits
proximalUP-(43)-sigma38p10 4609181 Gap 5.1 bits
distalUP 6.5 bits
distalUP-(51)-sigma38p10 4609181 Gap 5.1 bits
distalUP-proximalUP-sigma38p10 4609181 total 3.5 bits
