## Supplement_for_sigma38_paper-directory for "*Escherichia coli σ*^38^ promoters use two UP elements instead of a −35 element: resolution of a paradox and discovery that *σ*^38^ transcribes ribosomal promoters": Yasuno.Kyogoku2001-figure1b.pdf

piece 1, NC_000913, rrnB-P1, config: linear, direction: +, begin: 4164329, end: 4164353

```
     *4164330  .         *4164340  .         *4164350
5' t c a g a a a a t t a t t t t a a a t t t c c t c 3'
```

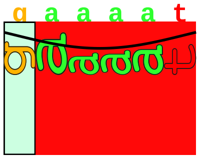

CombinedUP 7.0 bits

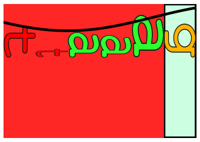

CombinedUP 6.1 bits

piece 1, NC_000913.i0,1gccg.i4164350,4164350gcgc, -55, config: linear, direction: +, begin: 4164335

```
    .         *4164340  .
5’ g c c g a a t t a t t g c g c 3’
```

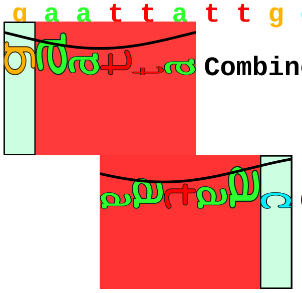

CombinedUP 5.6 bits

CombinedUP 5.8 bits

piece 1, NC_000913.i0,1gccg.i4164350,4164350gcgc, -53, config: linear, direction: +, begin: 4164337

*4164340 . *4164350

5′ g c c g t t a t t t t g c g c 3′

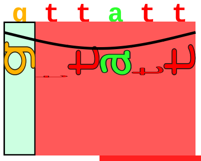
CombinedUP 4.7 bits

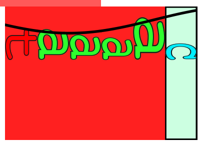
CombinedUP 6.3 bits

piece 1, NC_000913.i0,1cgcg.i4164350,4164350cgcg, -49, config: linear, direction: +, begin: 4164341

. *4164350

5’ c g c g t t t a a a c g c g 3’

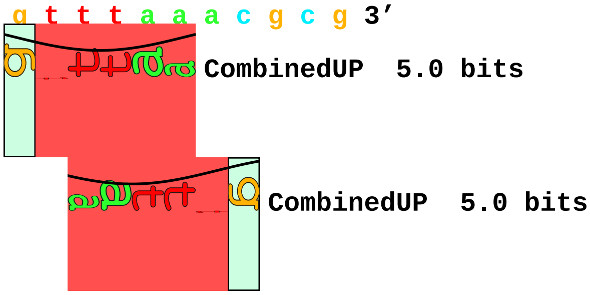

piece 1, NC_000913.i0,1cgcg.i4164350,4164350cgcg, -46, config: linear, direction: +, begin: 4164344

```
      .          *4164350  .
5' c g c g a a a t t t c g c g 3'
```

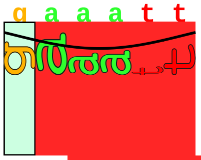

CombinedUP 6.1 bits

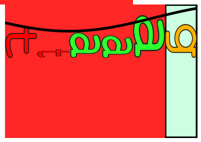

CombinedUP 6.1 bits

piece 1, NC_000913.i0,1cccggcgcgccggg.d4164329,4164329, NON, config: linear, direction: +, begin: 4

```
    *4164330  .         *4164340
5’ c c c g g c g c g c c g g g 3’
```
