## Supplementary figures and images for "*Escherichia coli σ*^38^ promoters use two UP elements instead of a −35 element: resolution of a paradox and discovery that *σ*^38^ transcribes ribosomal promoters"

### Estrem.Gourse1998-logo.pdf

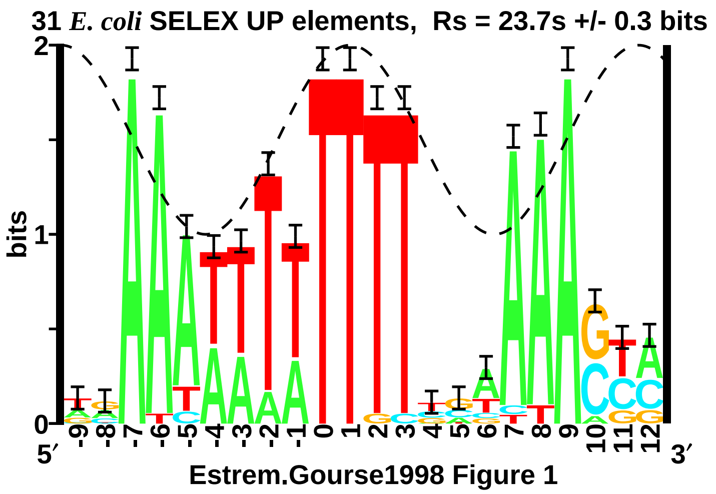

### Estrem.Gourse1999-fig1-map.pdf

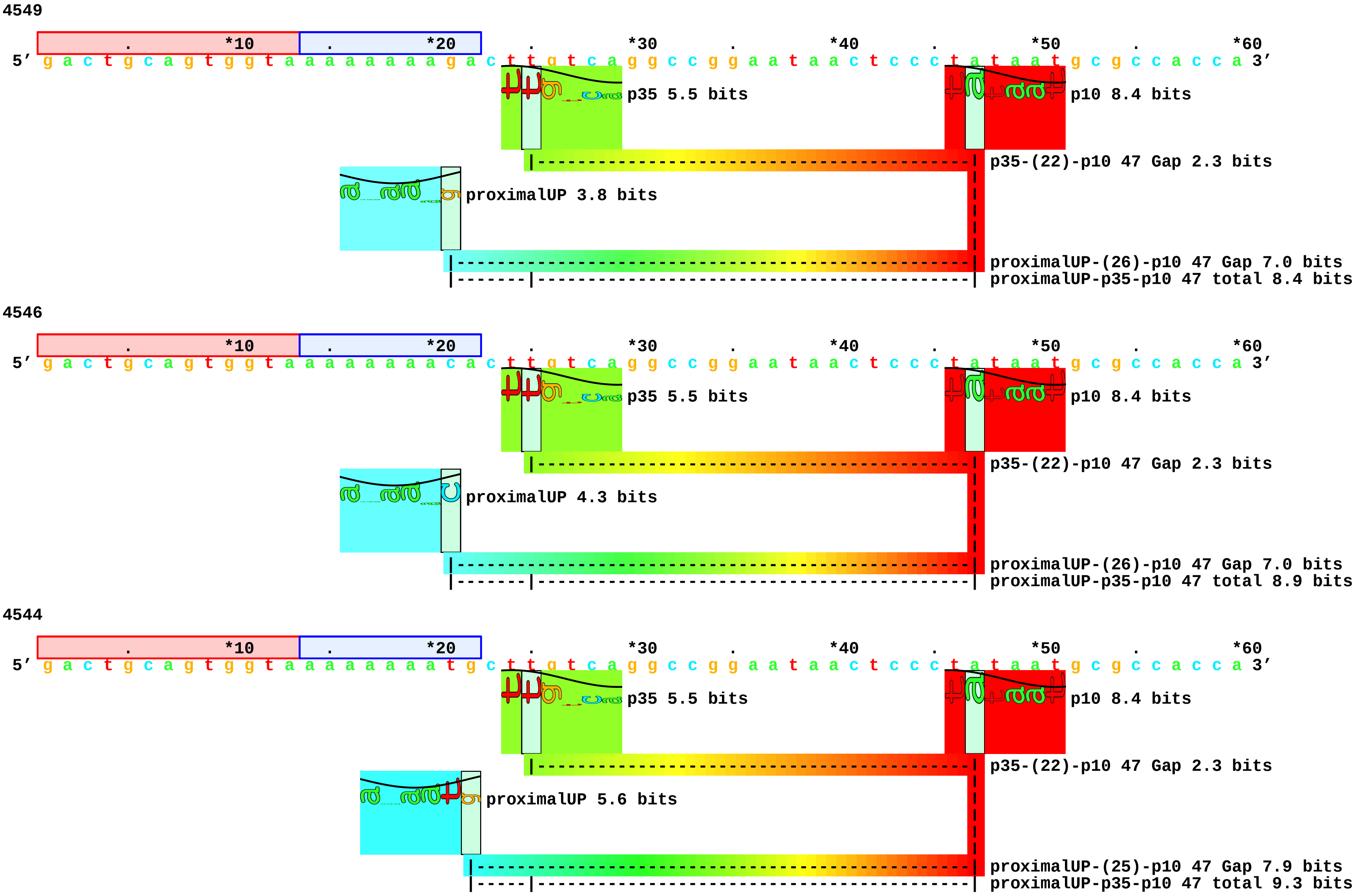

4547

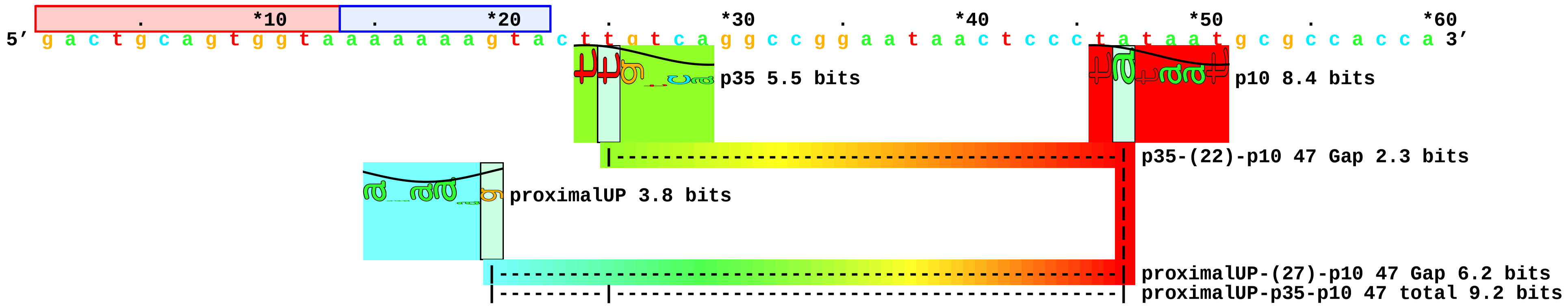


4548

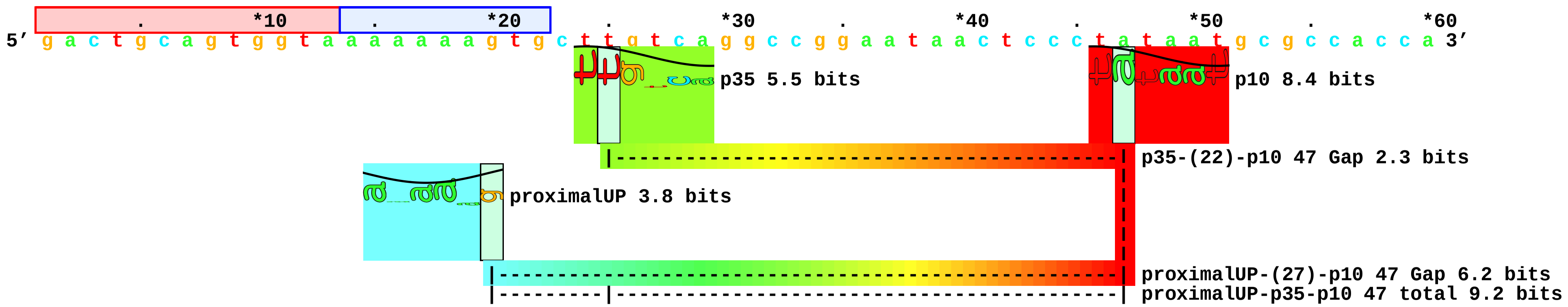


4542

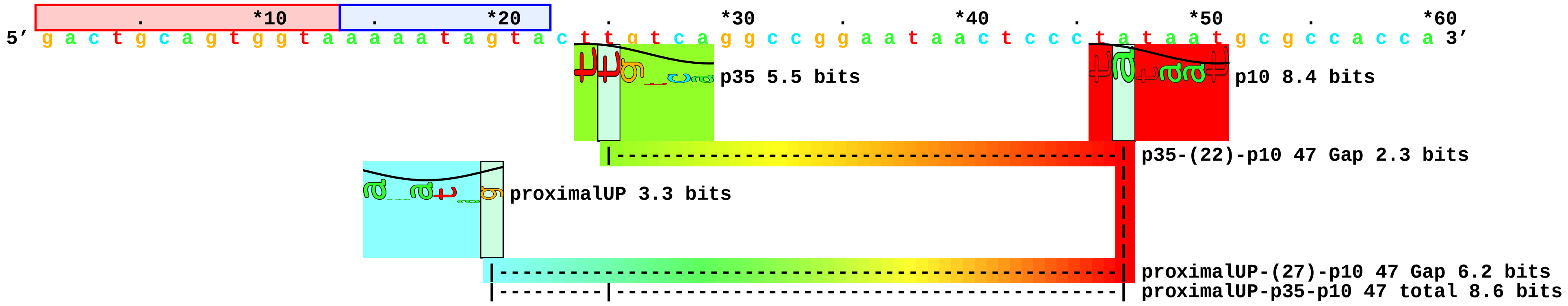

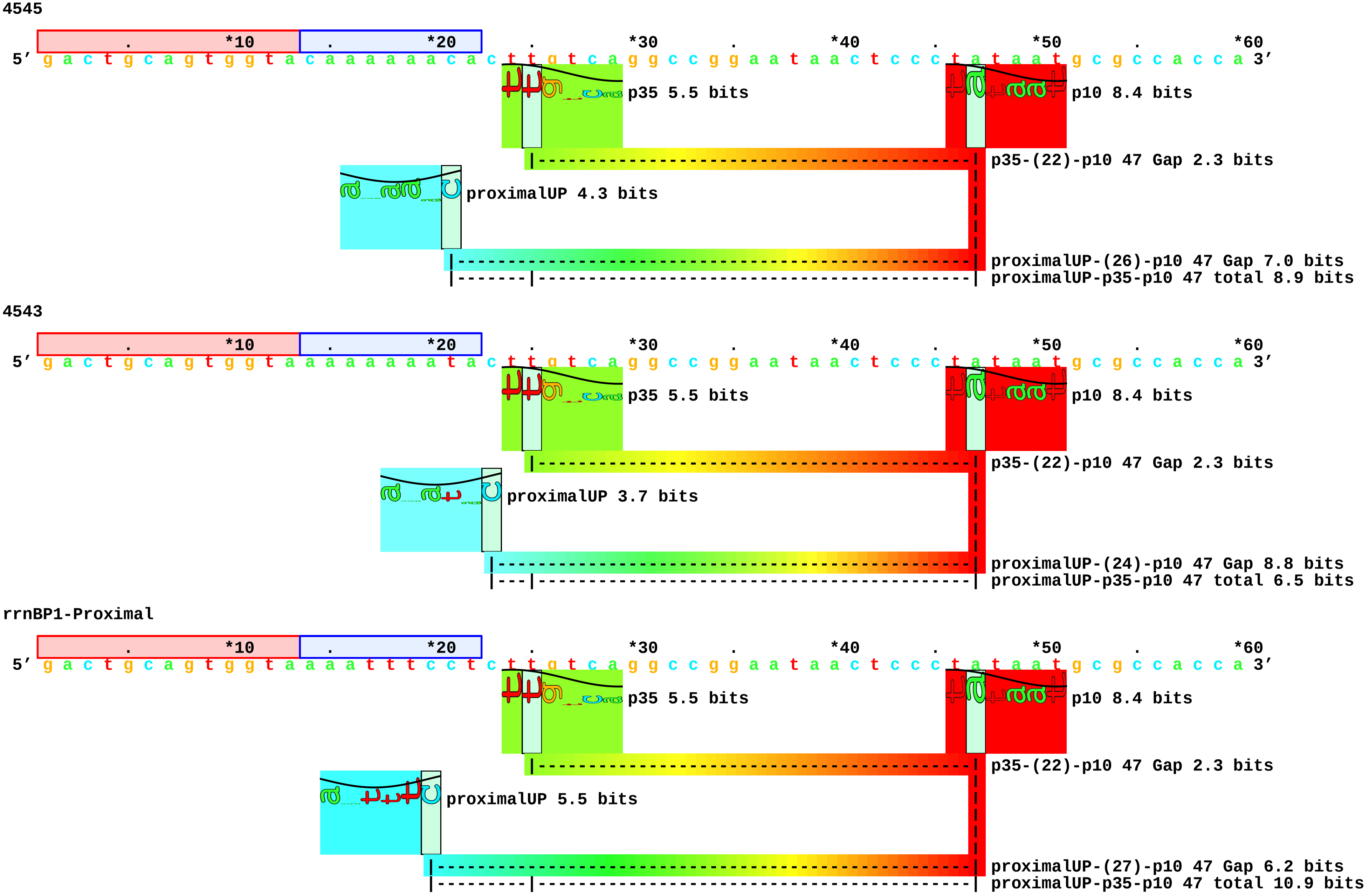

NoUP

```
                .         *10         .         *20         .         *30         .         *40         .         *50         .         *60
5' g a c t g c a g t g g t a c c t a g g a a t c t t g t c a g g c c g g a a t a a c t c c c t a t a a t g c g c c a c c a 3'
```

### Estrem.Gourse1999-fig3-map.pdf

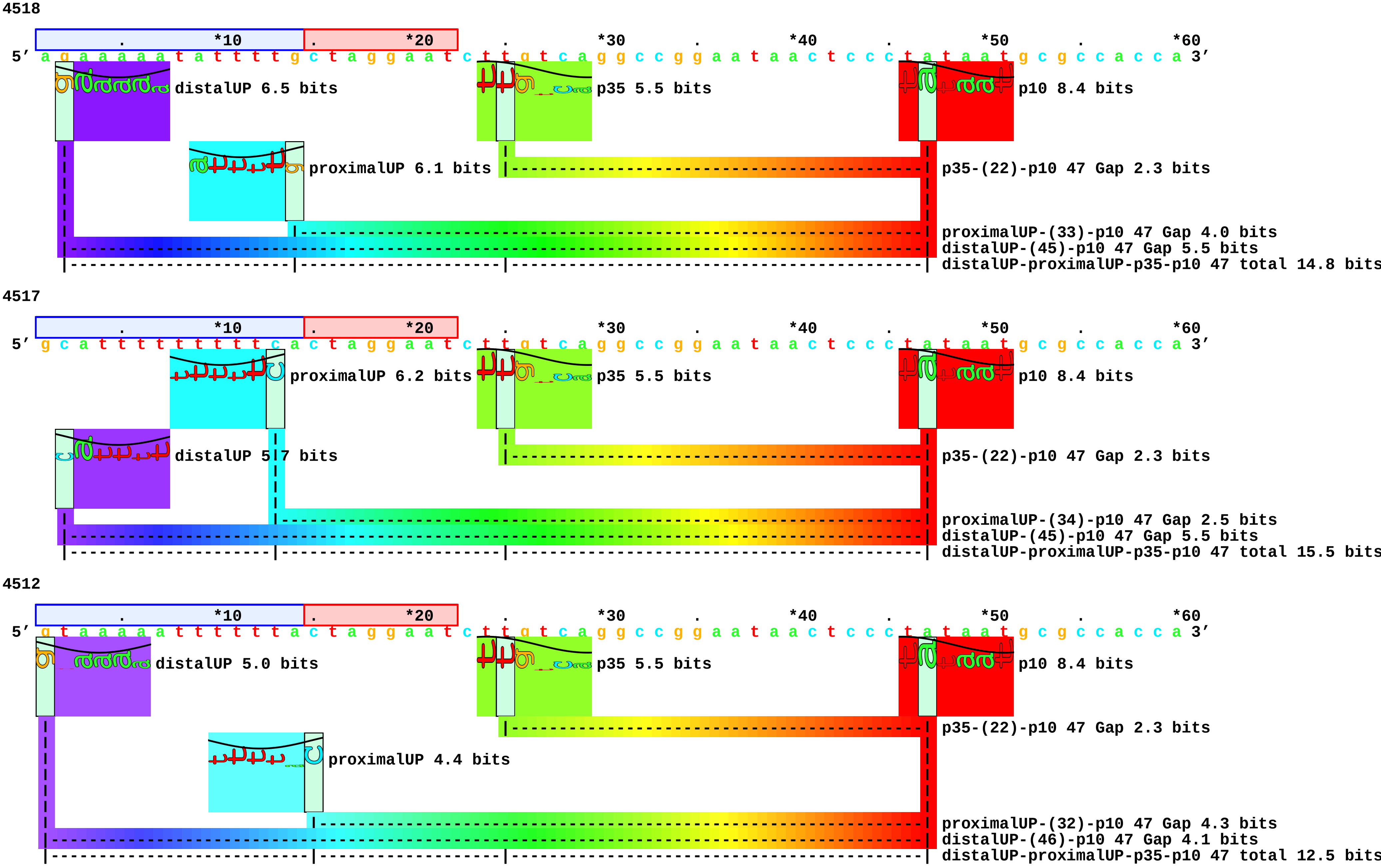

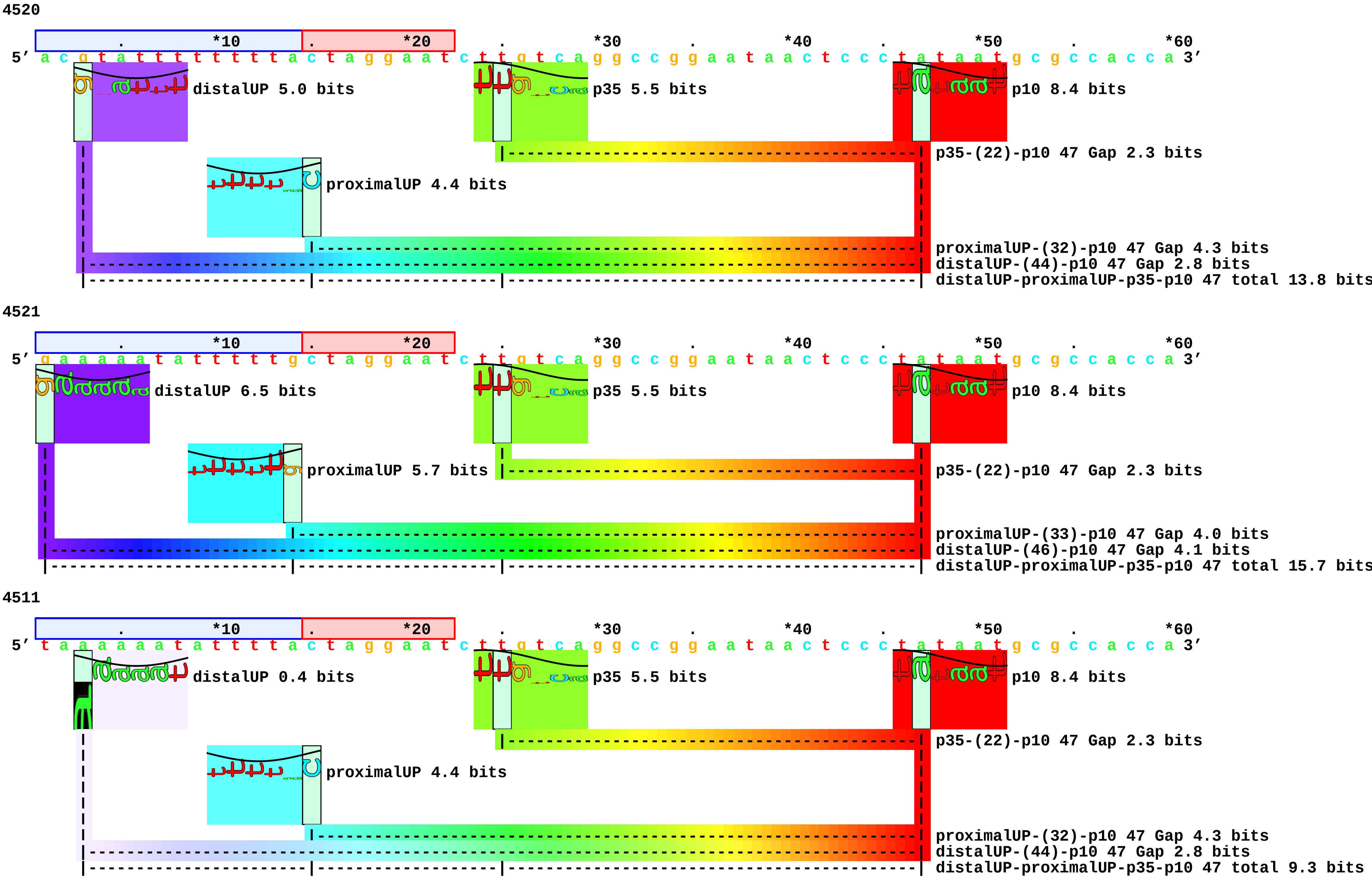

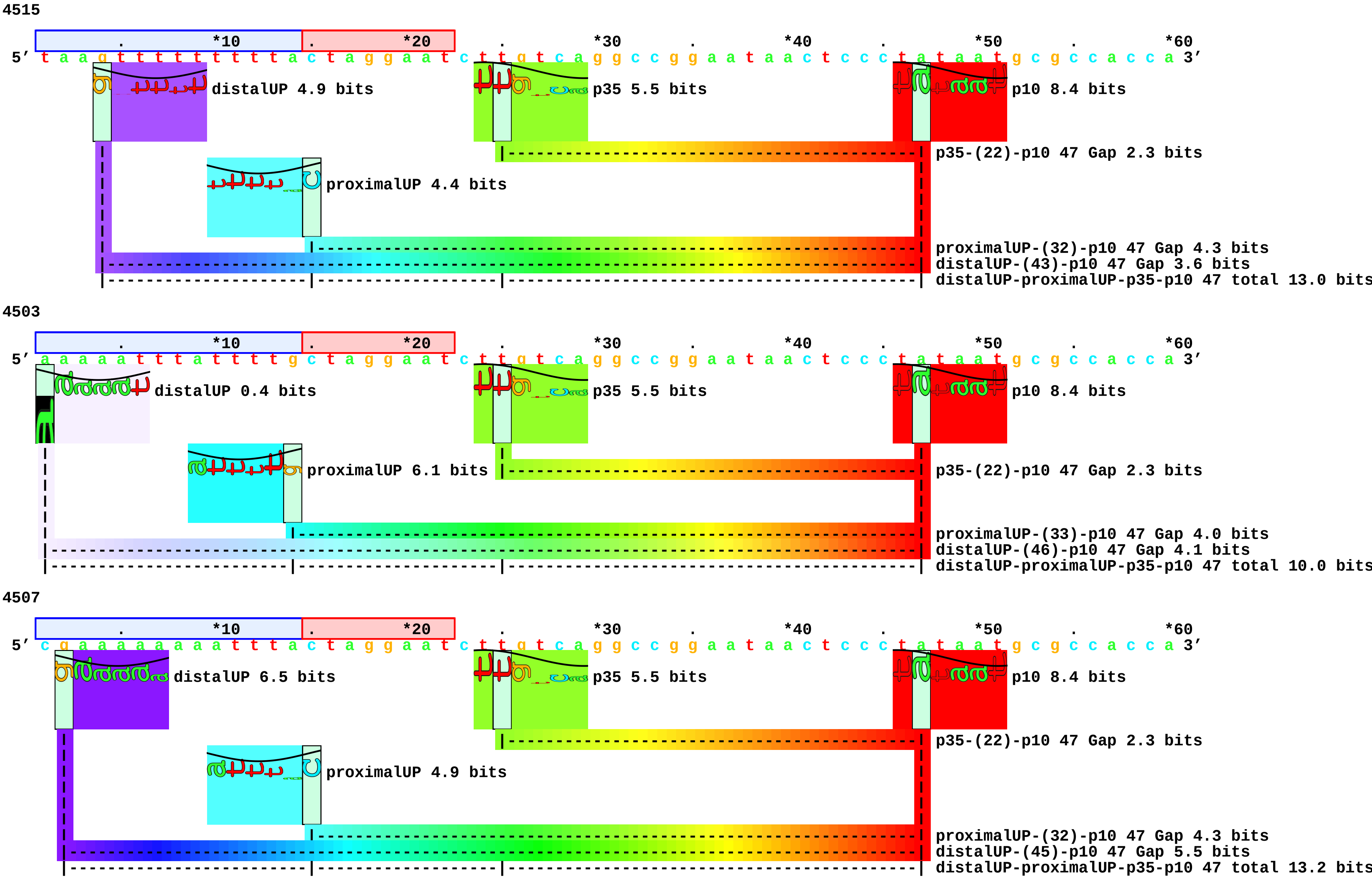

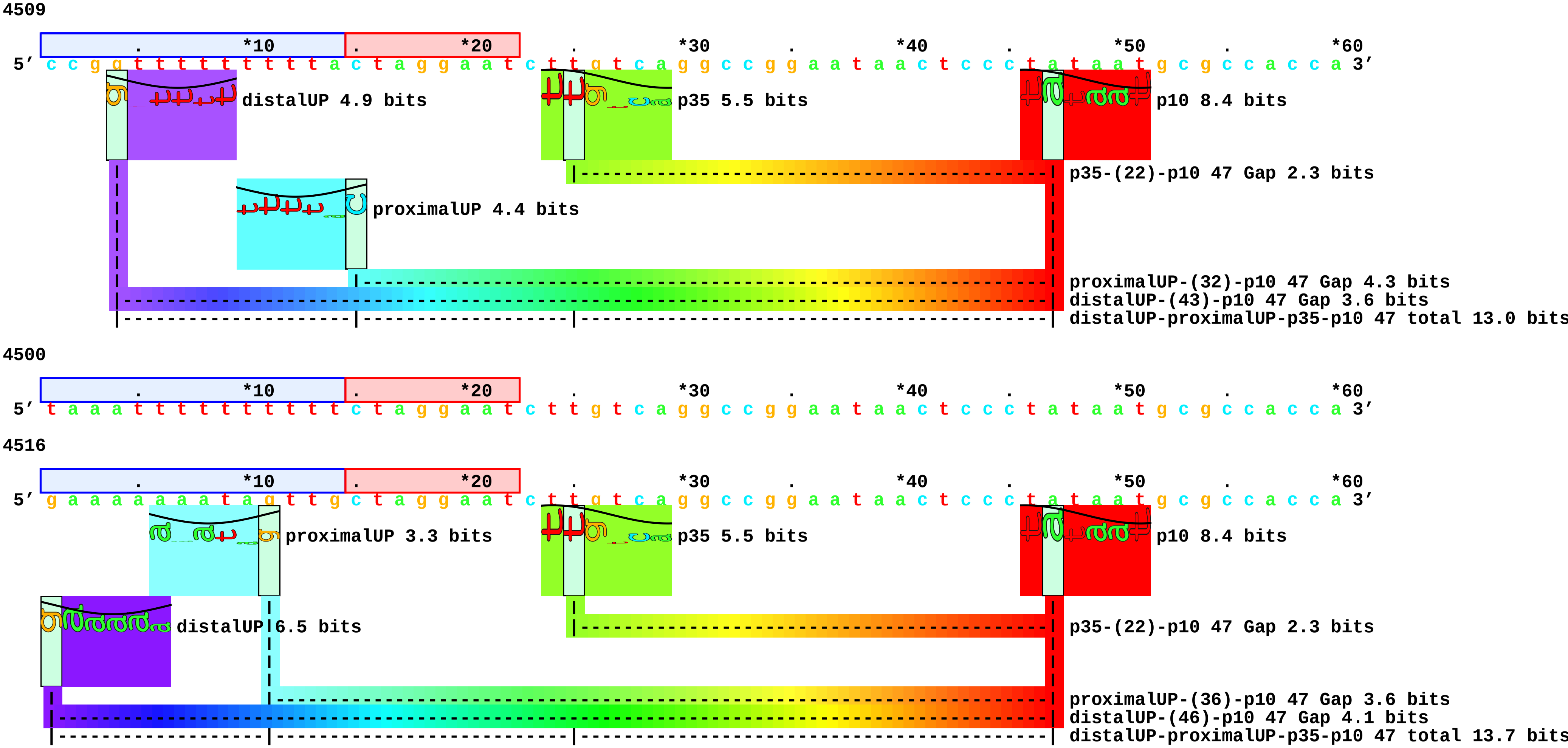

4514

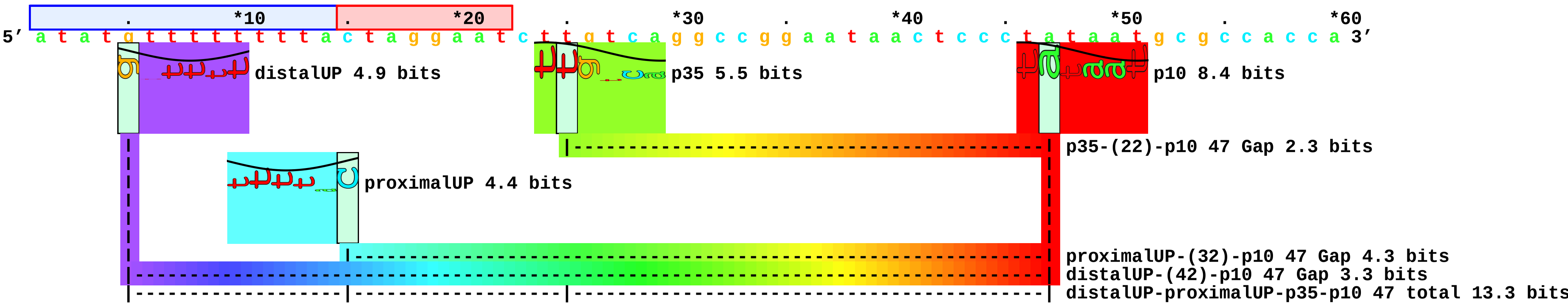


4505

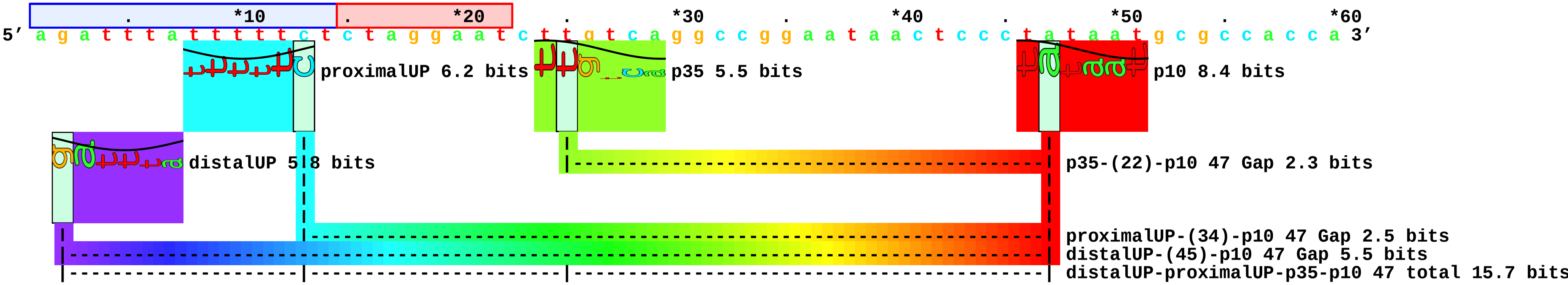


4504

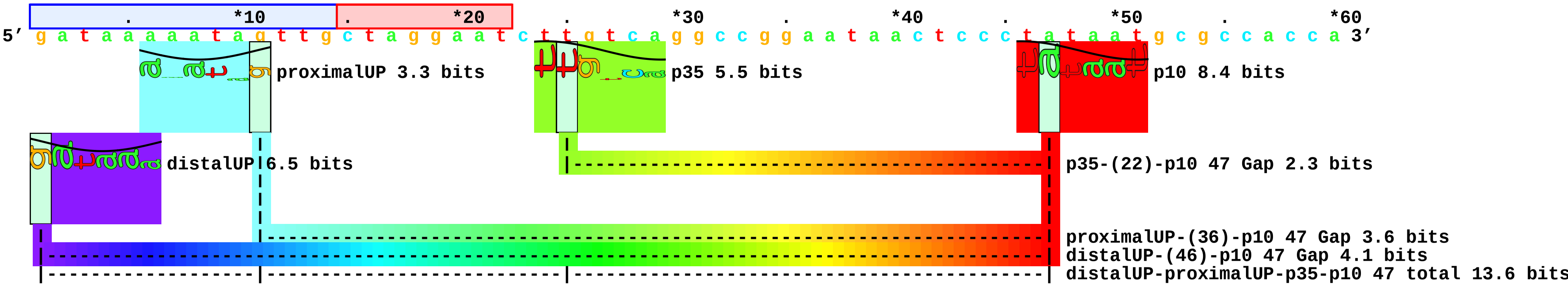

4510

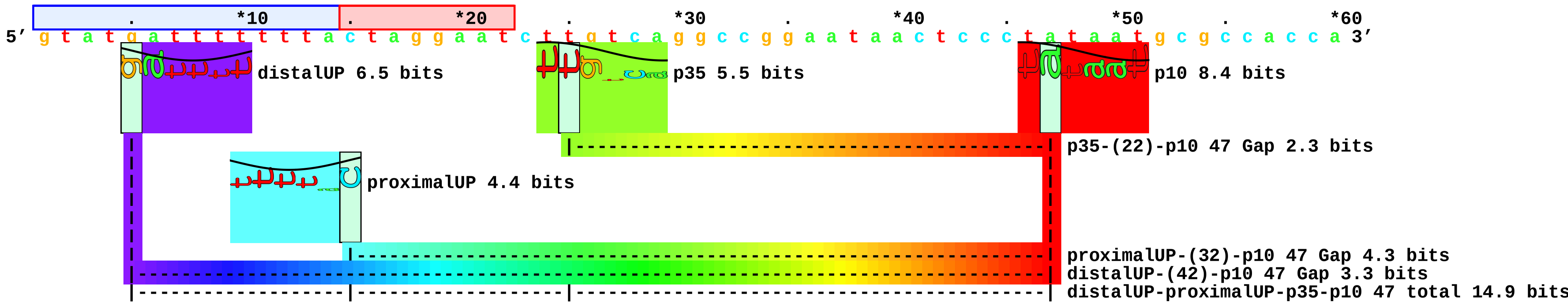


4508

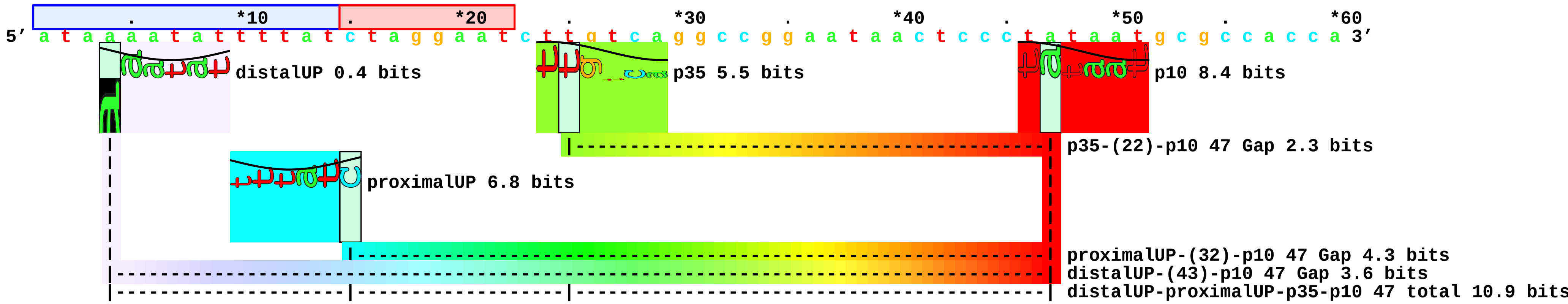


4519

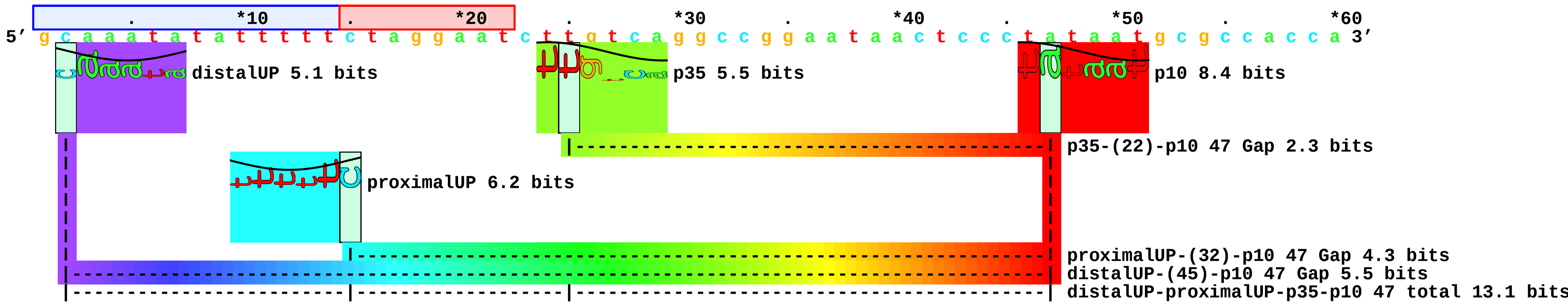

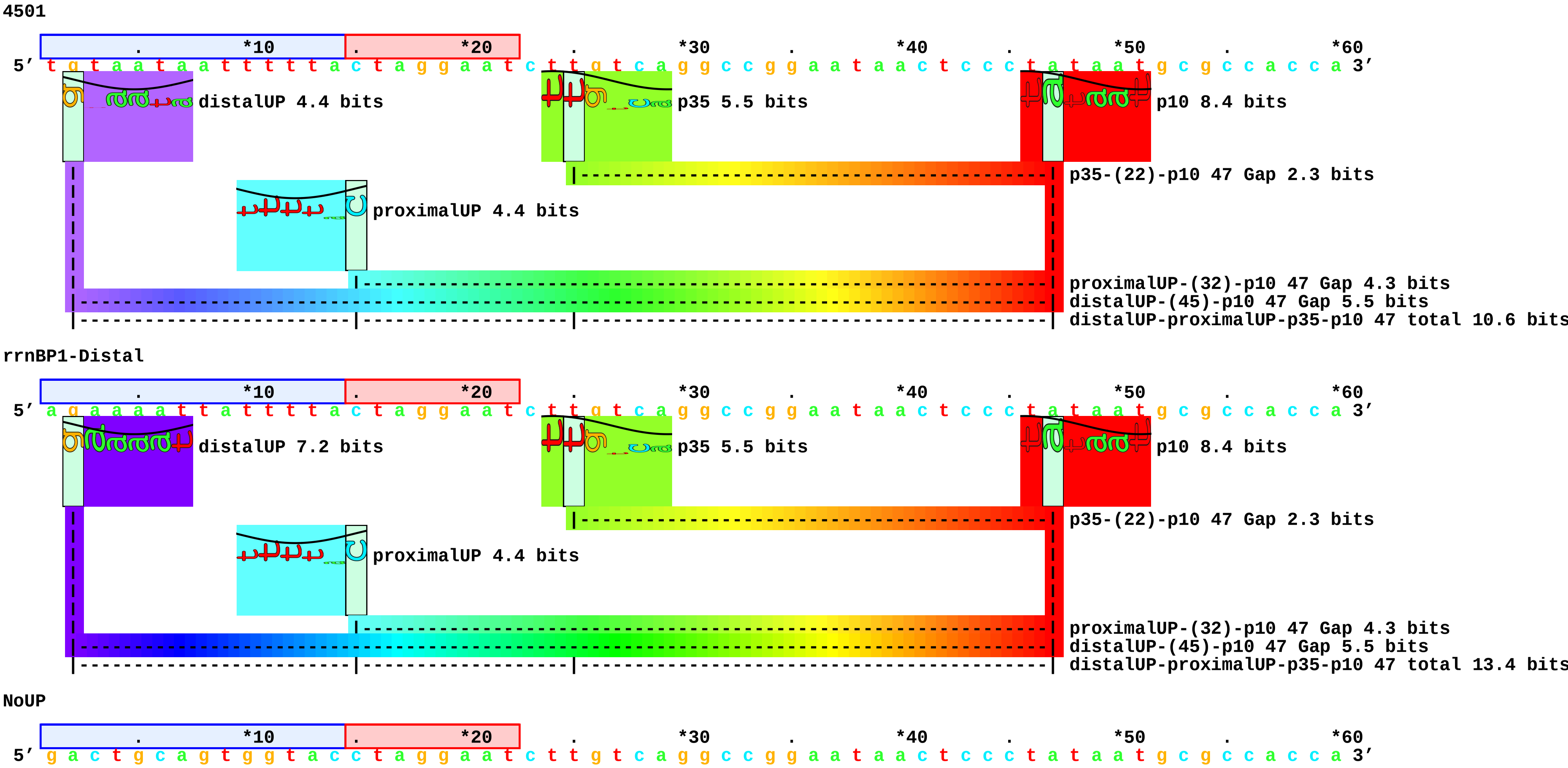

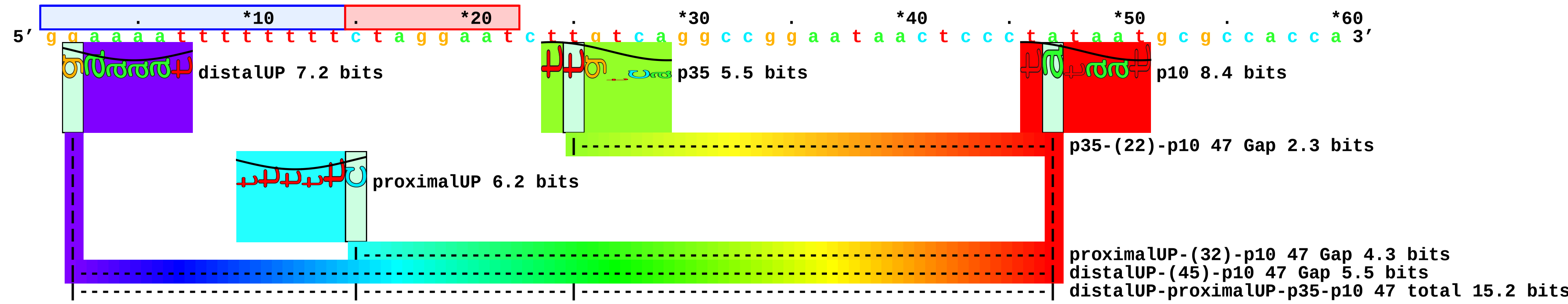

### EXPT-sigma38-bolAP1Mutants.tif

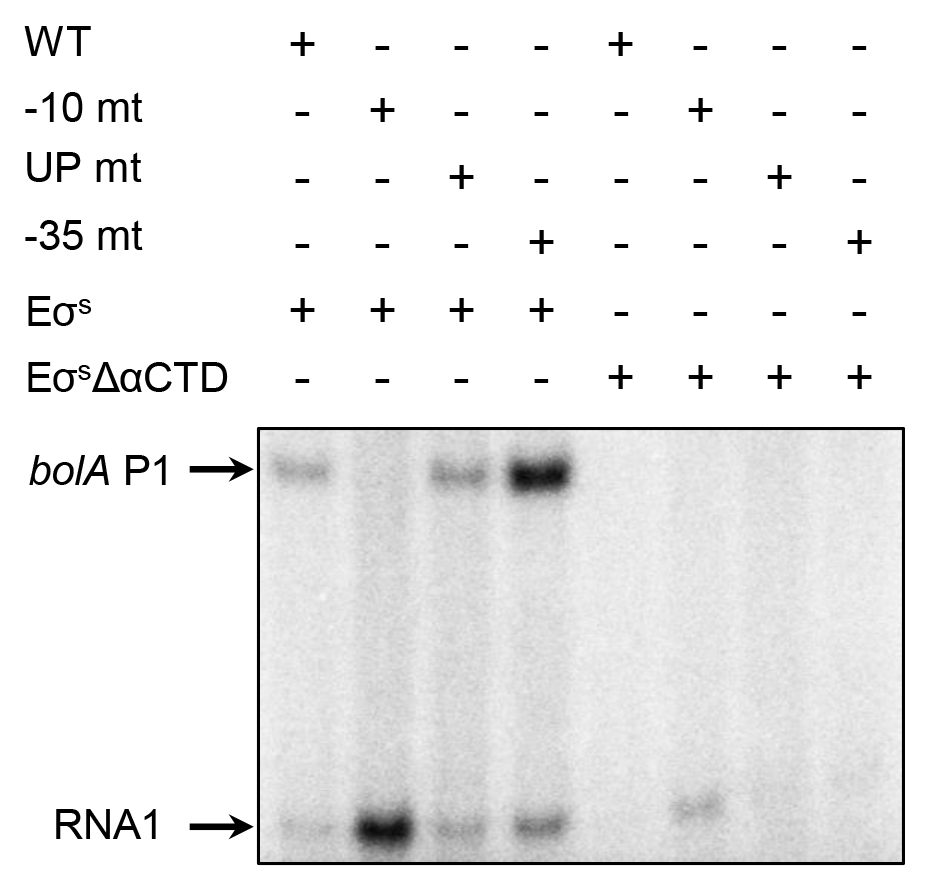

### EXPT-sigma38-rrnBP1.tif

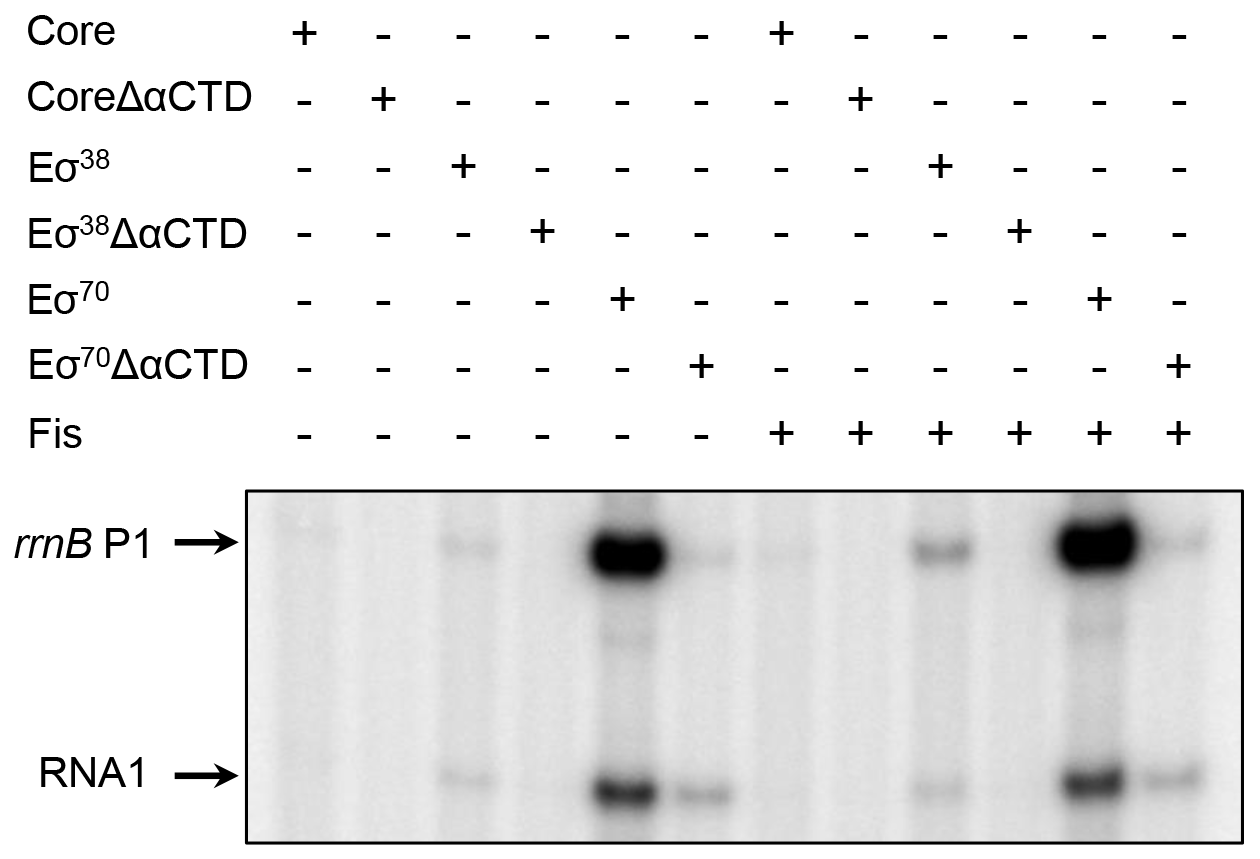

### EXPT-sigma38-rrnBP1Mutants.tif

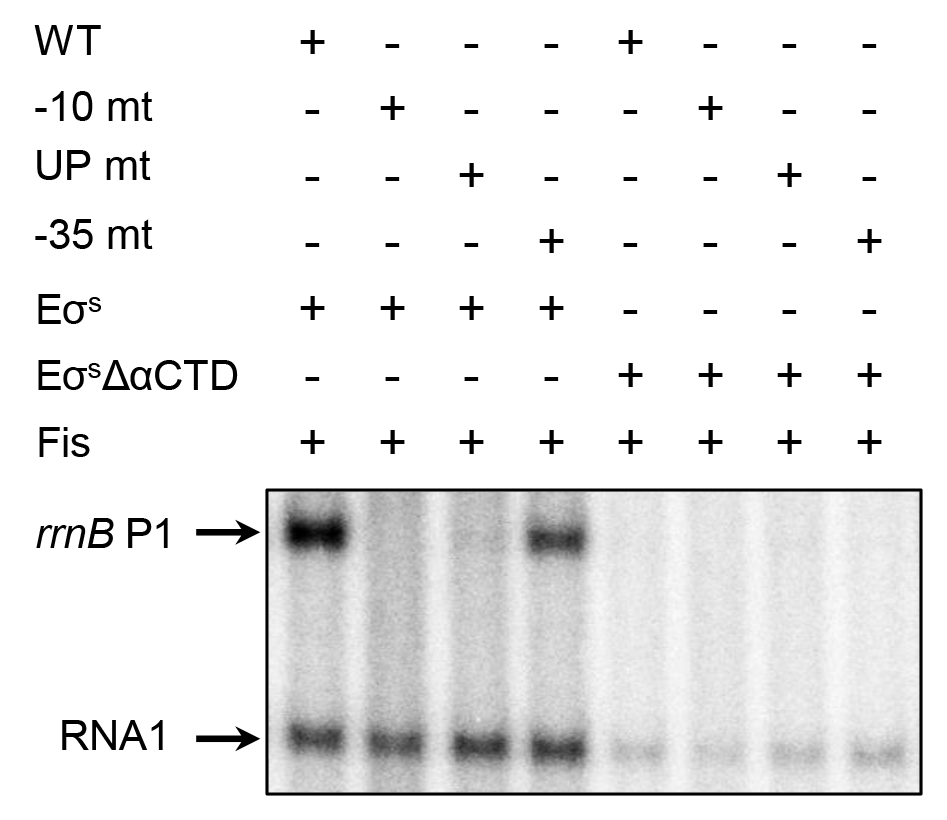

### figure-01.pdf

#### 78 Sigma38
##### transcriptional start sites
Rs = 0.13 +/- 0.06 bits [-2 to +2]

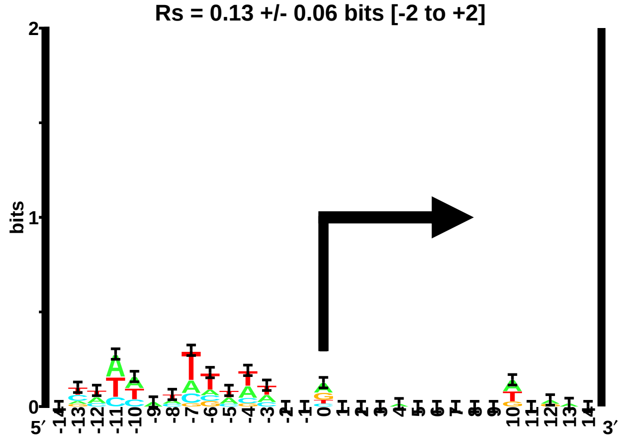

### figure-02.pdf

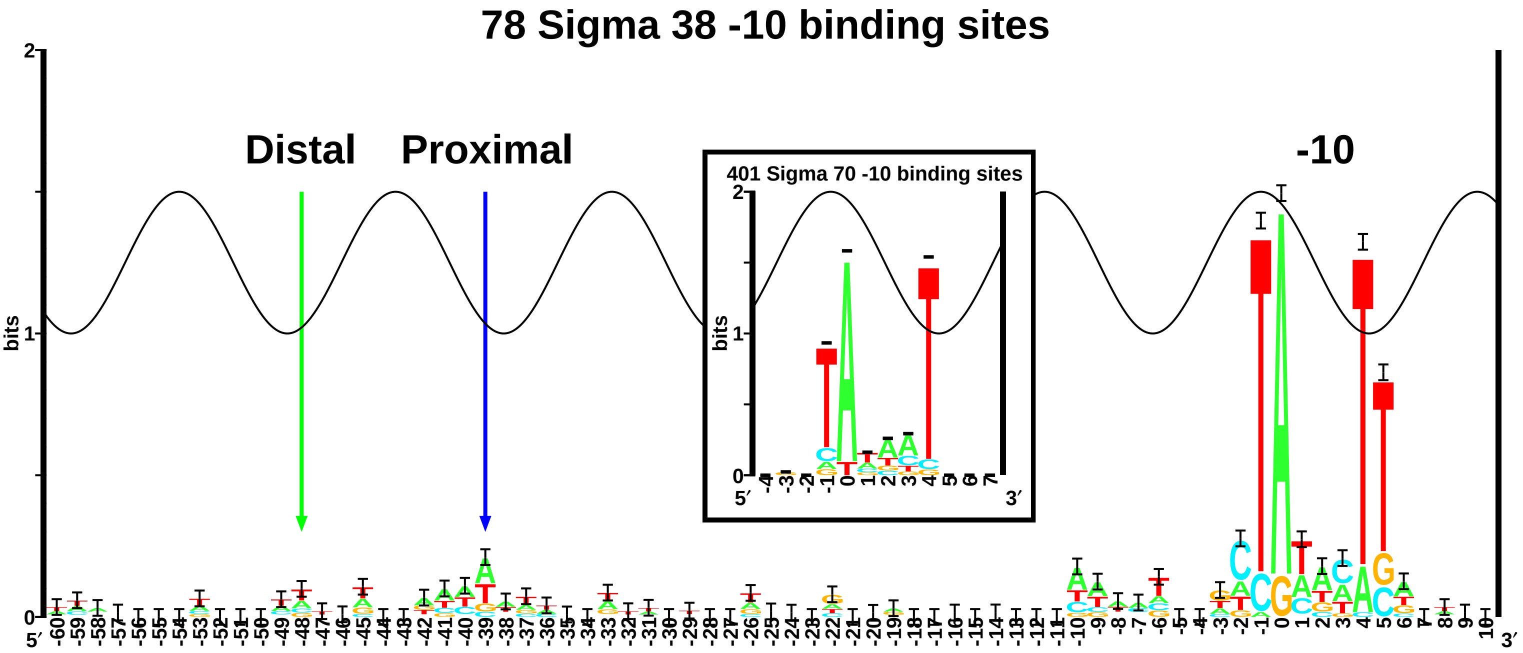

### figure-03.pdf

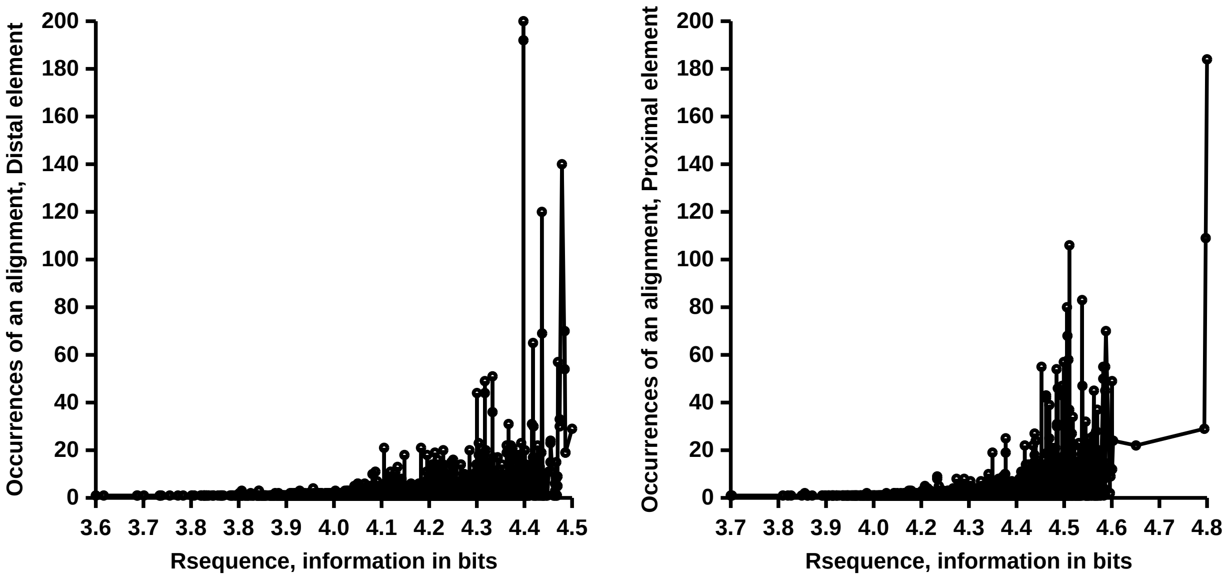

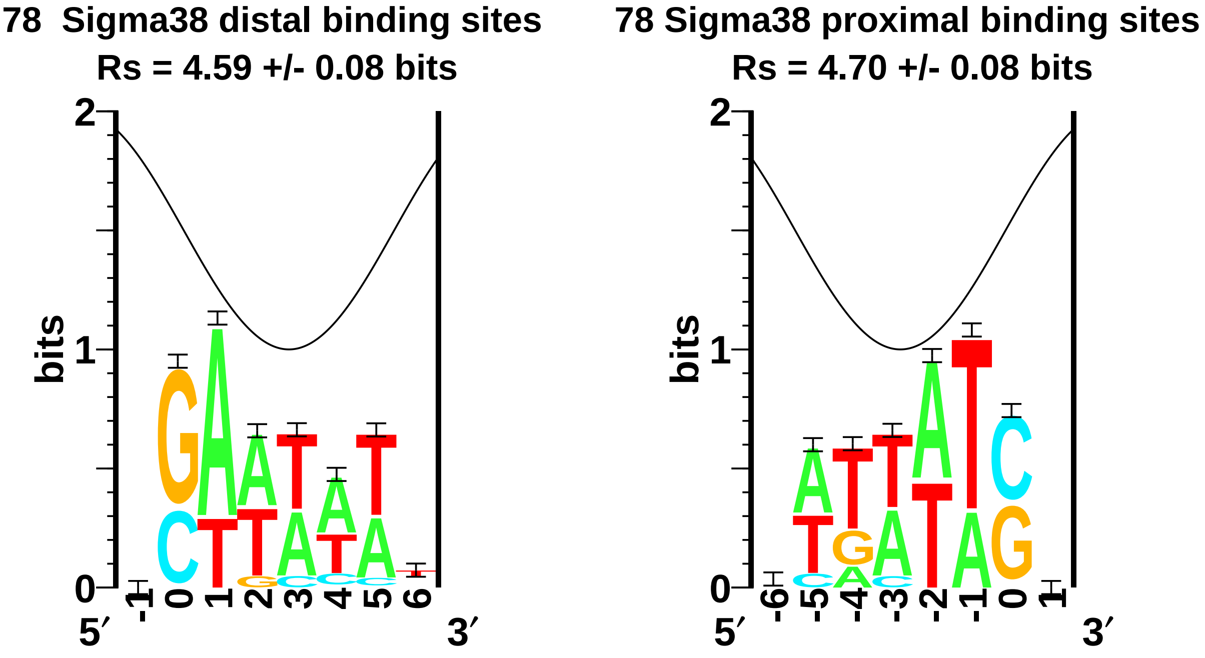

### figure-04.pdf

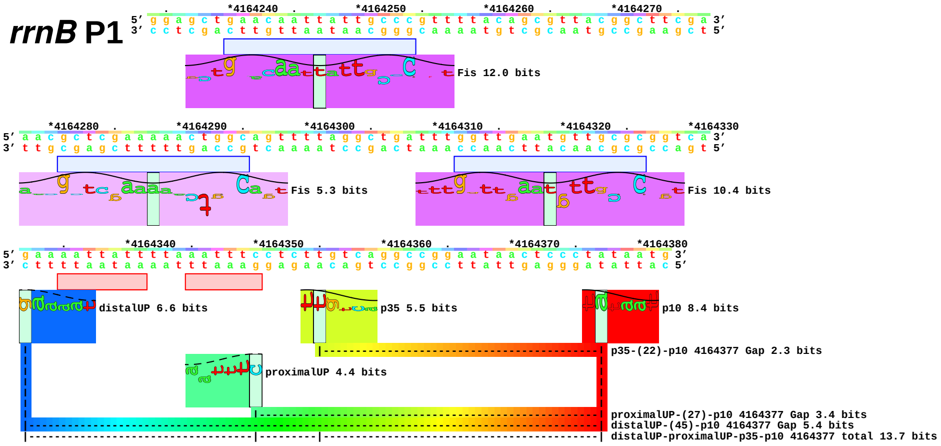

### figure-05.pdf

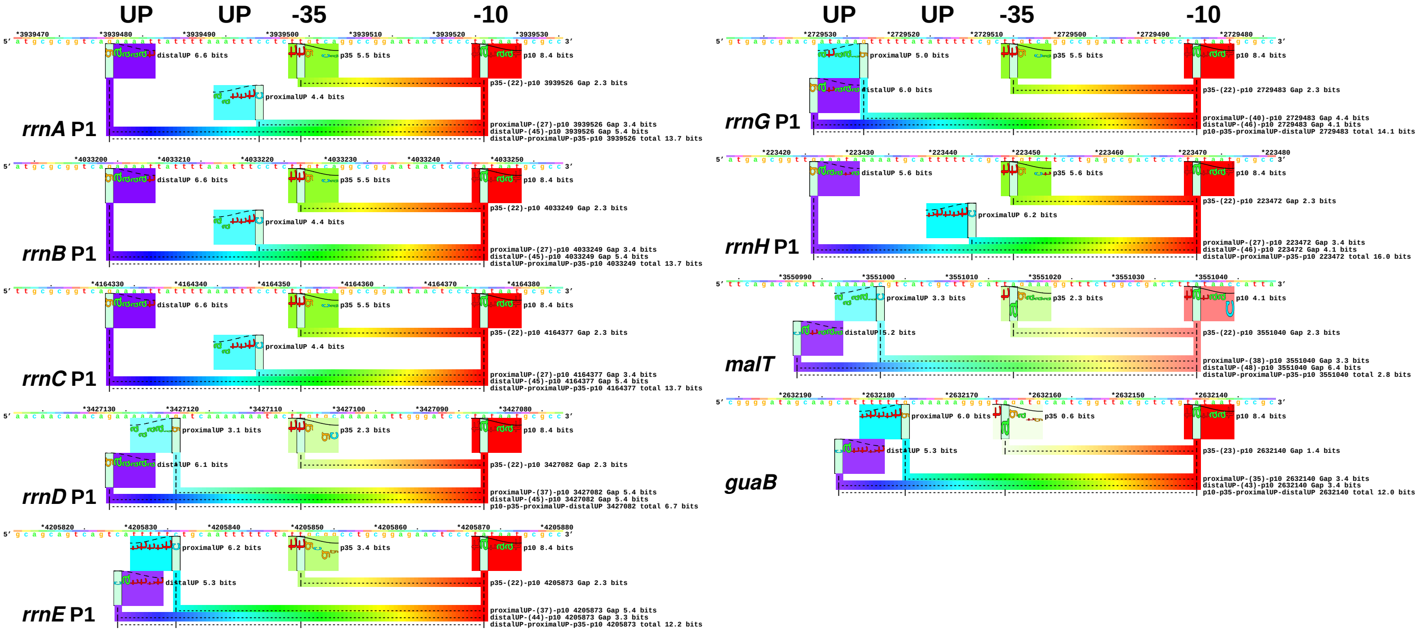

### figure-06.pdf

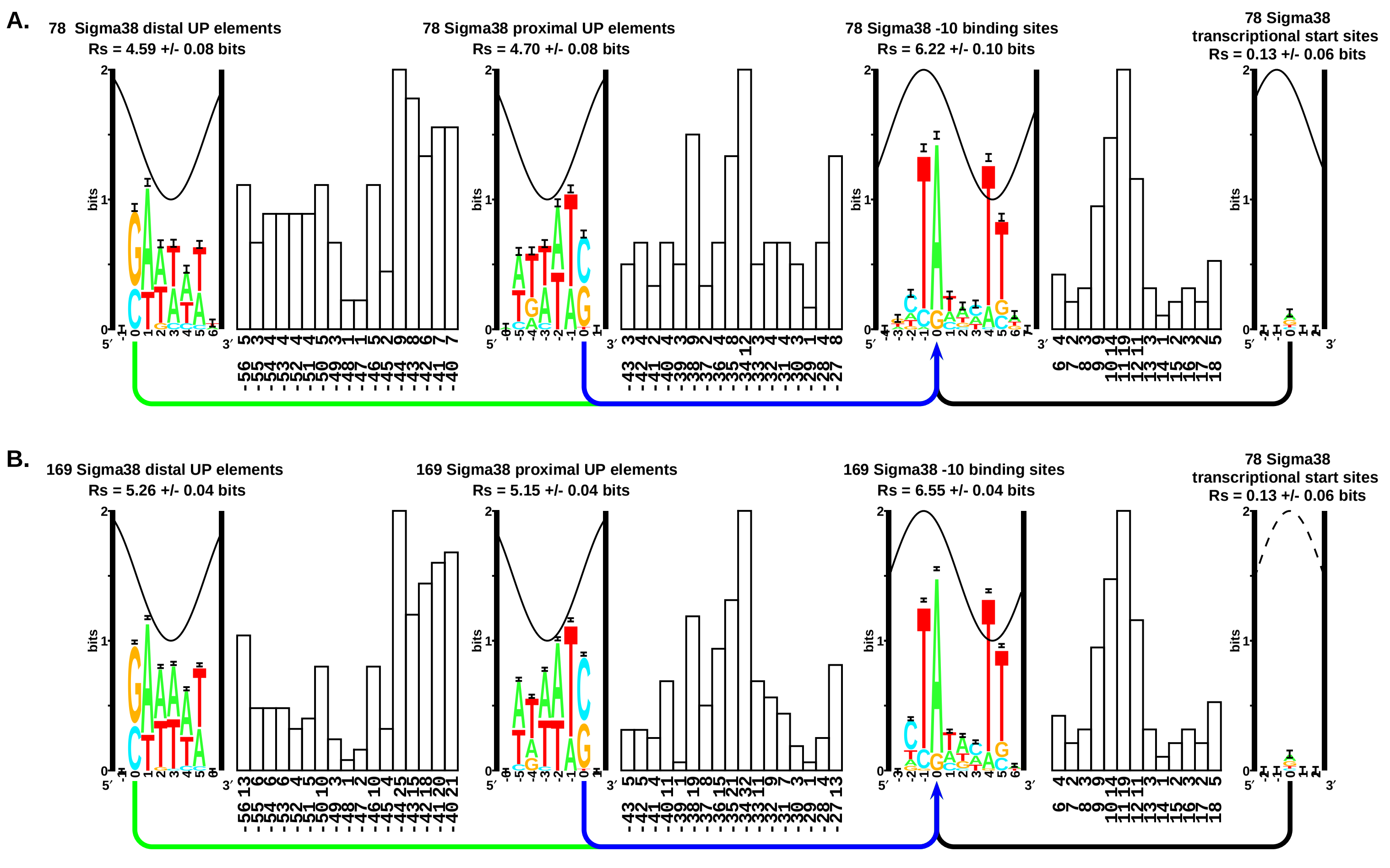

### figure-07.pdf

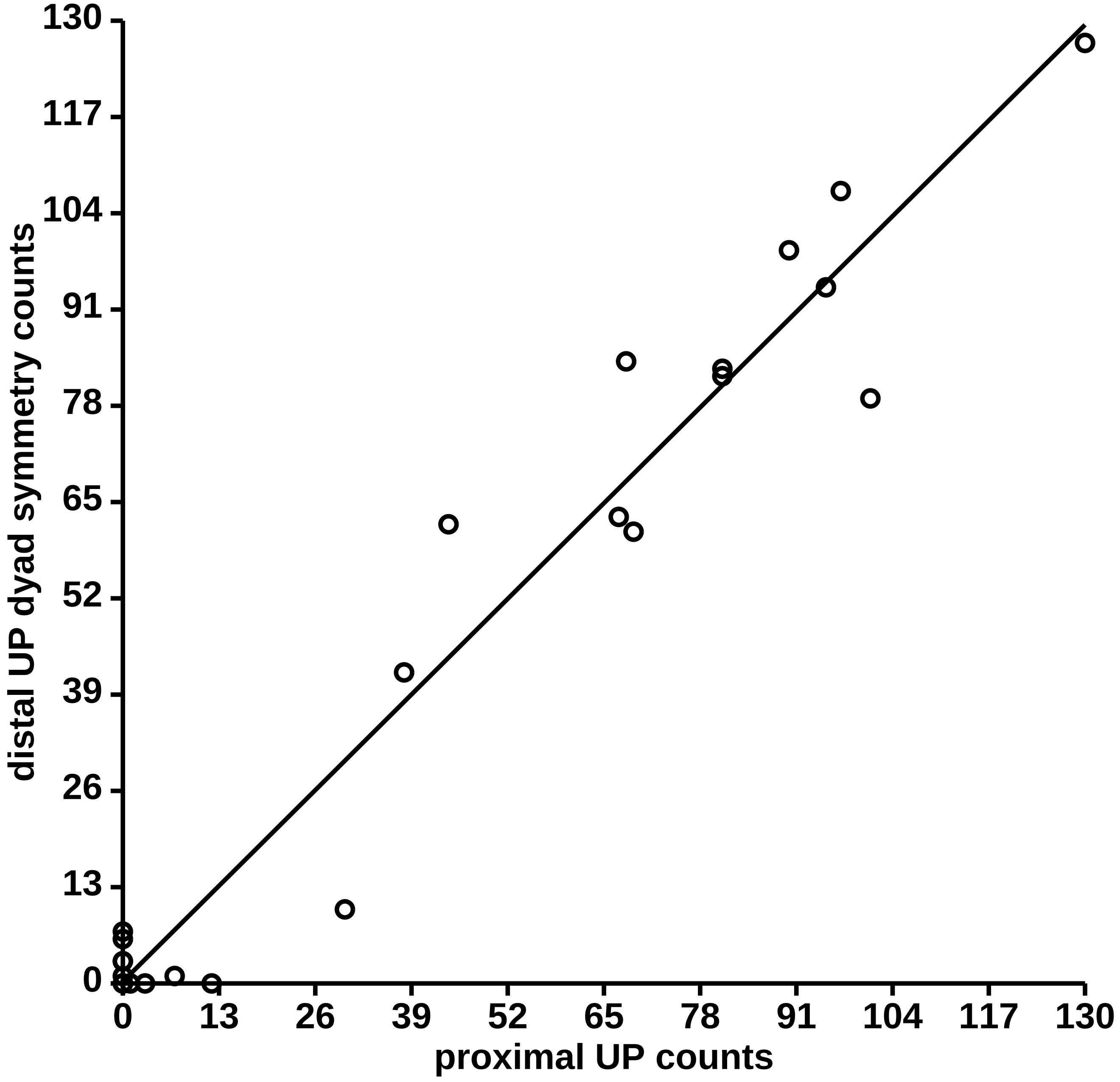

### figure-08.pdf

**338 Sigma38 UP element binding sites**
**(Combined distal and inverted proximal UP elements)**
**Rs = 5.09 +/- 0.02 bits**

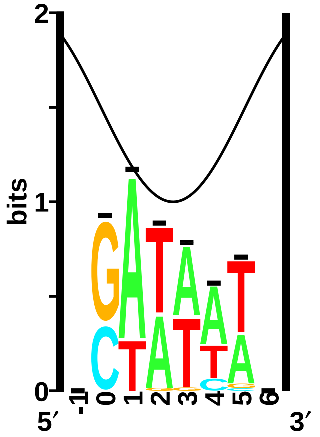

### figure-09.pdf

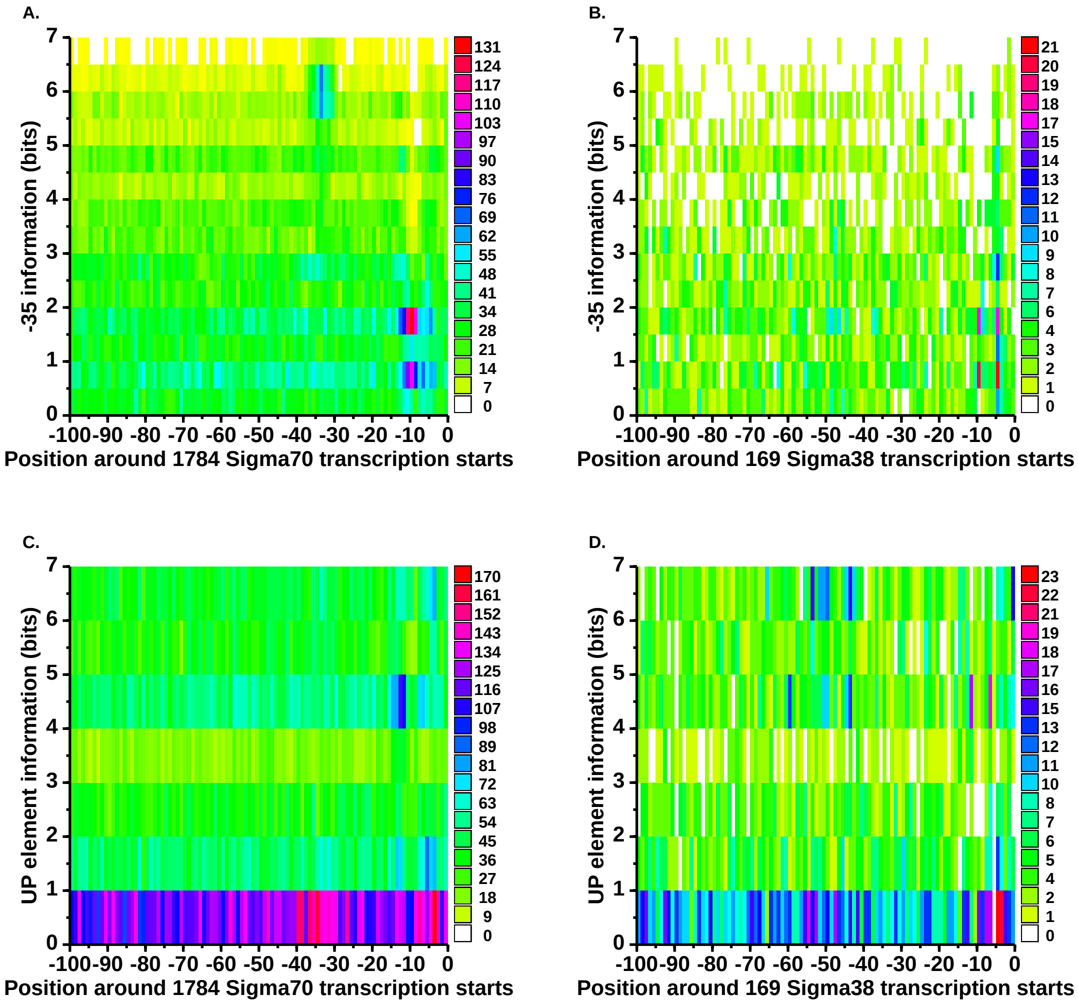

### figure-10.pdf

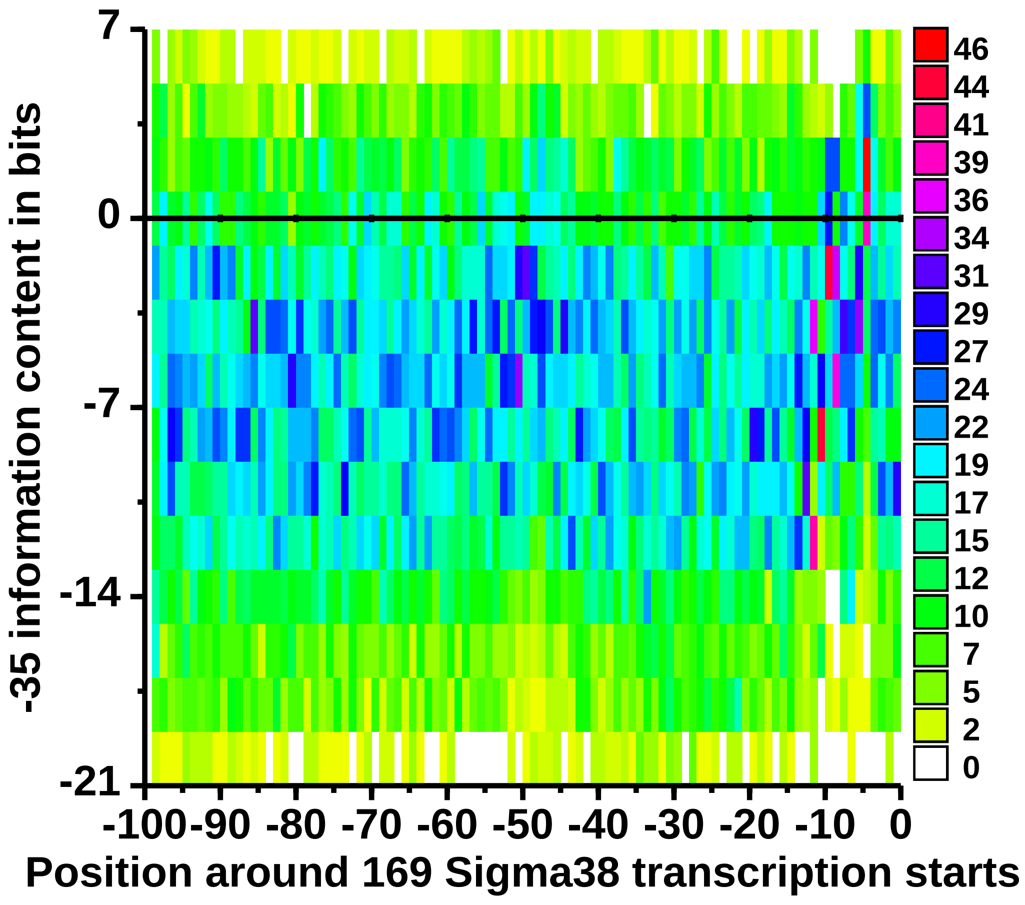

### figure-11.pdf

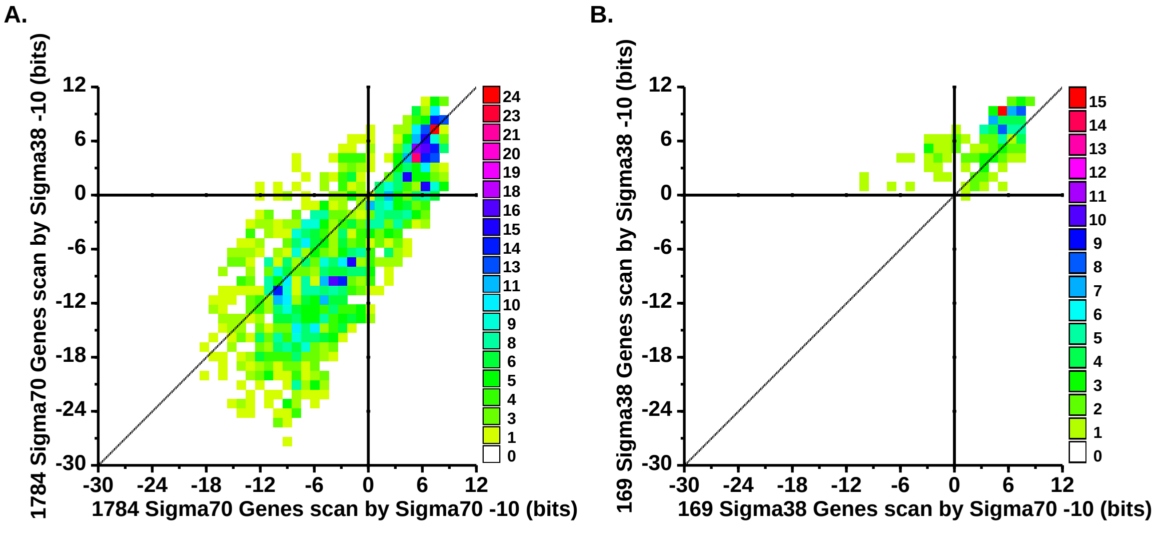

### figure-12.pdf

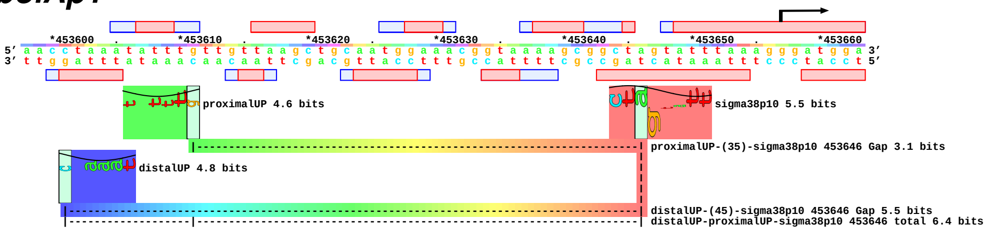

### figure-13.pdf

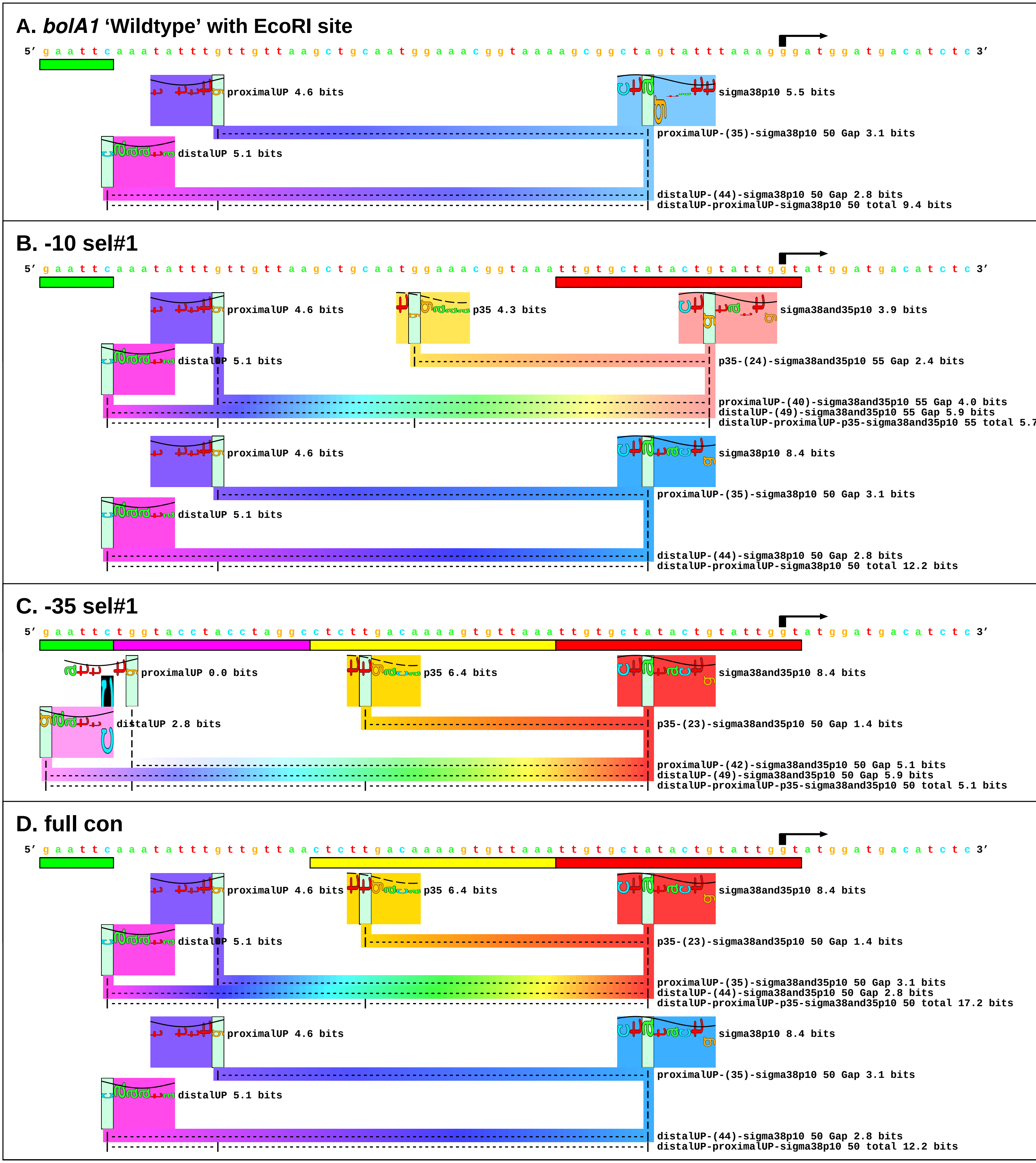

### figure-14.pdf

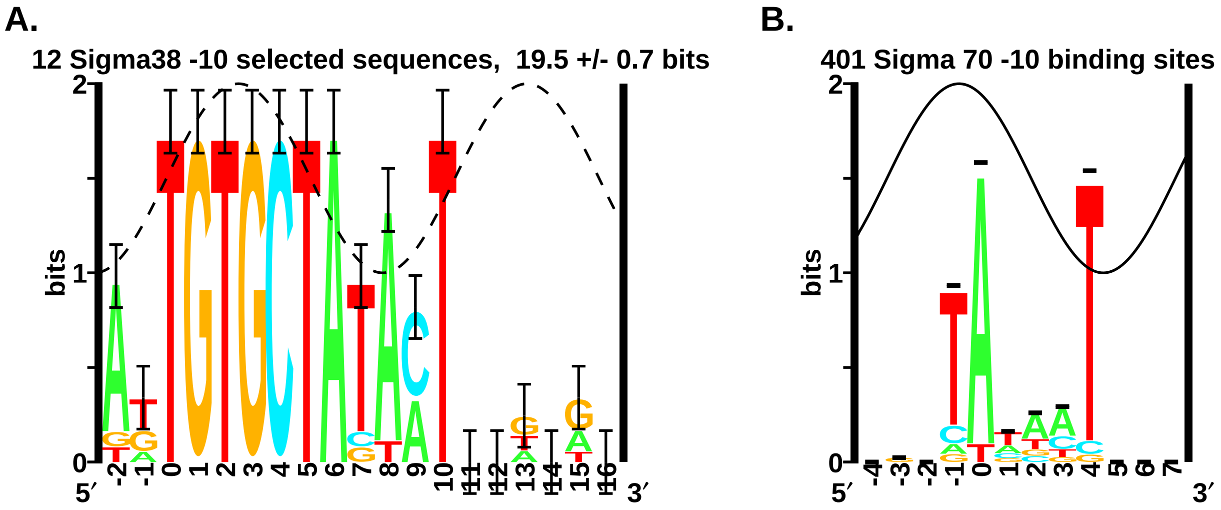

### figure-15.pdf

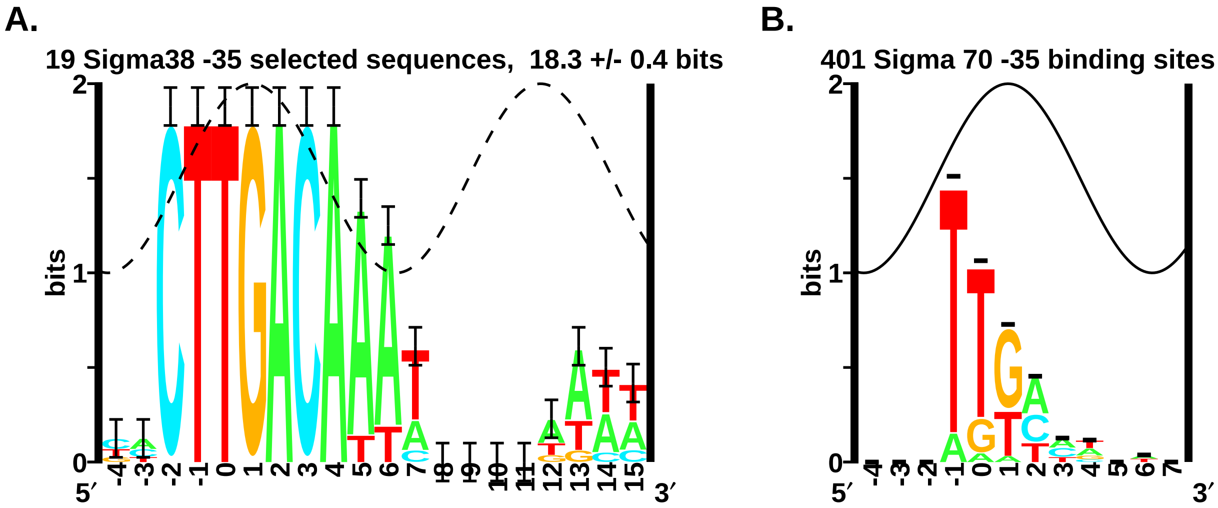

### figure-16.pdf

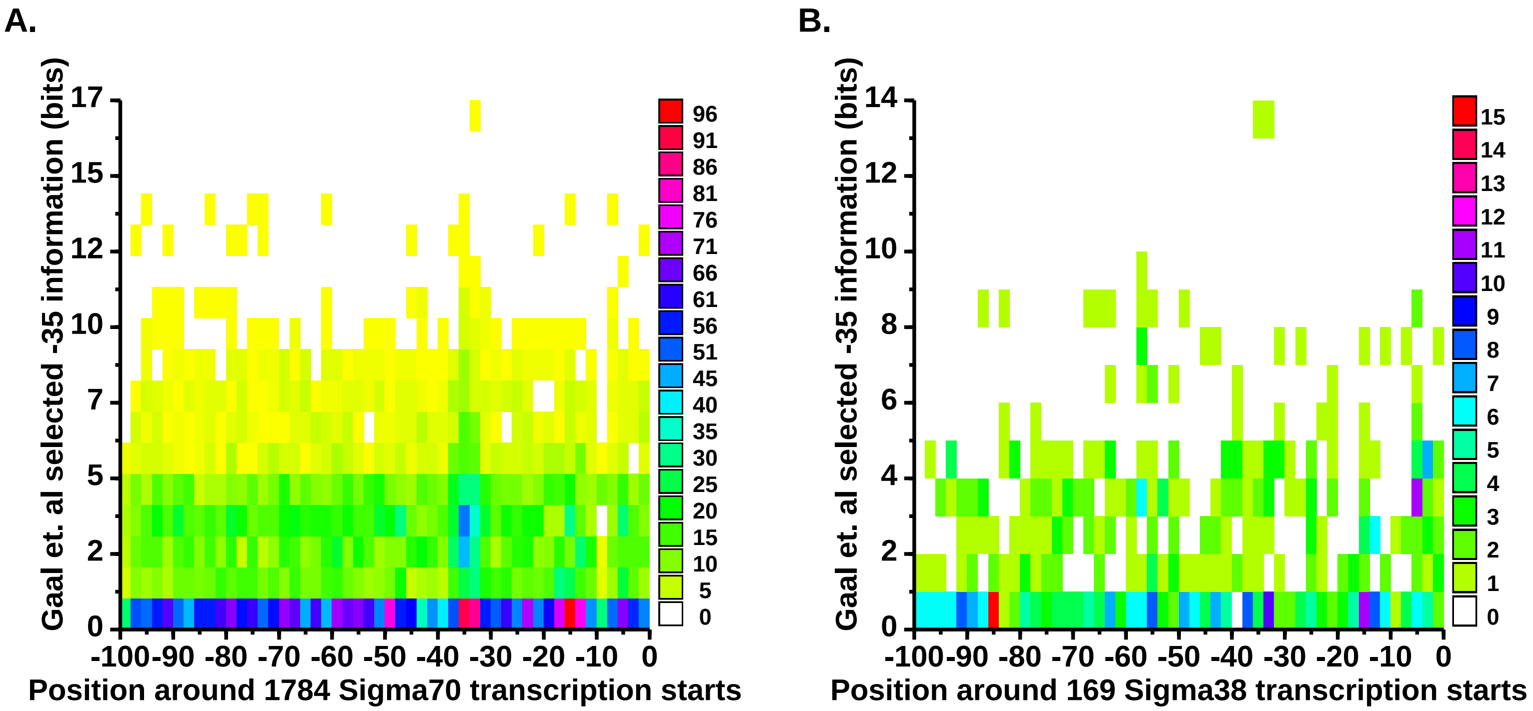

### figure-17.pdf

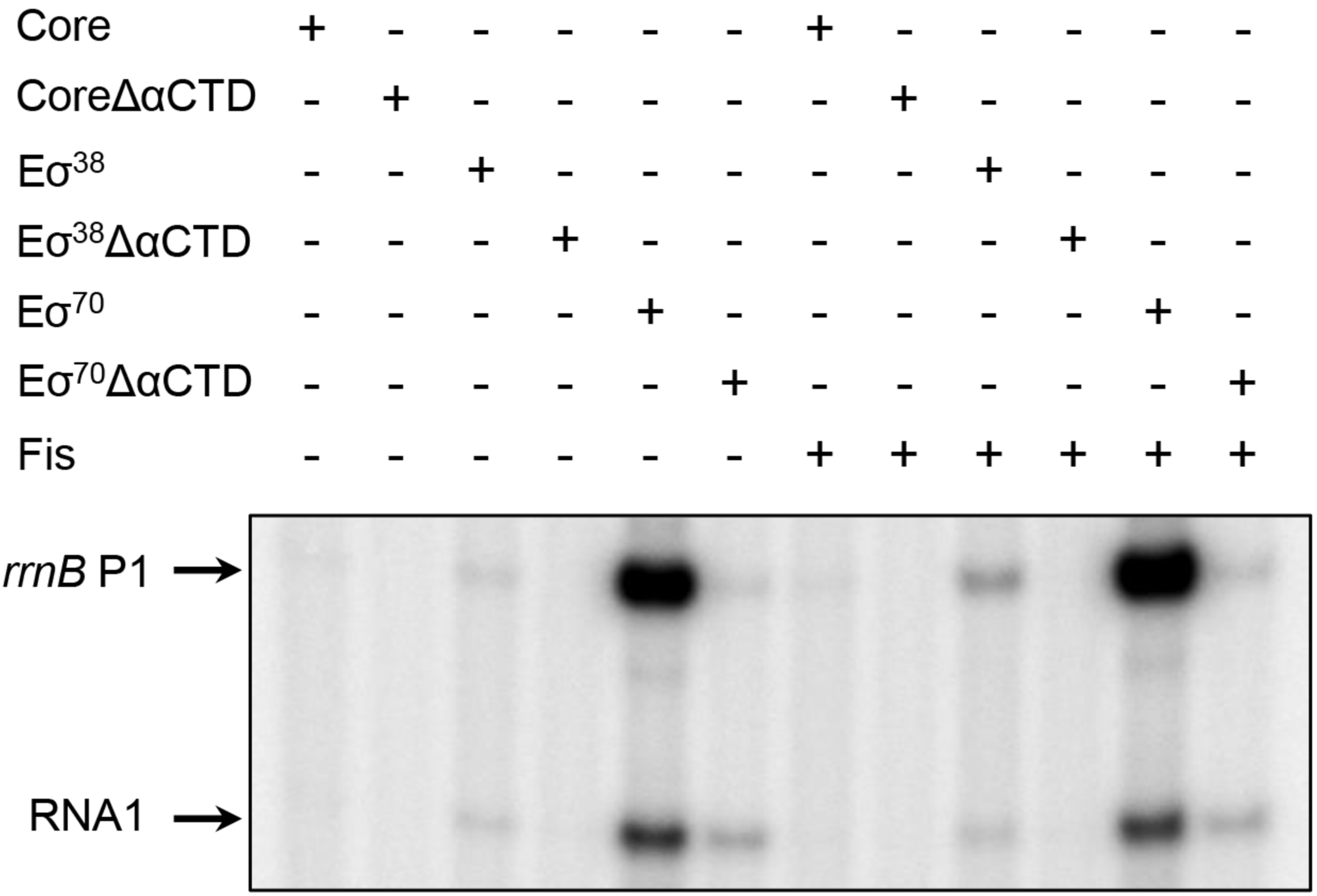

### figure-18.pdf

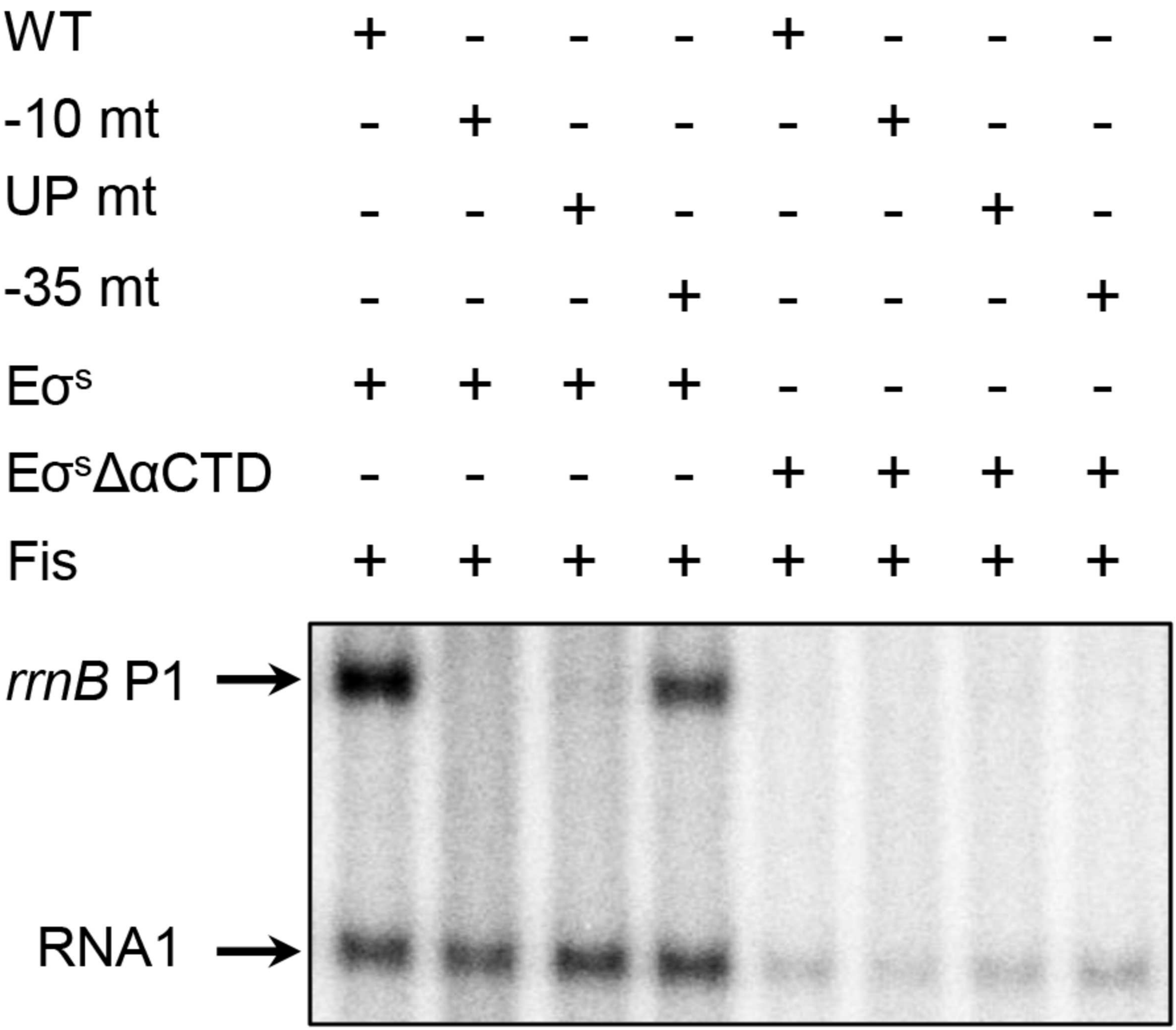

### figure-19.pdf

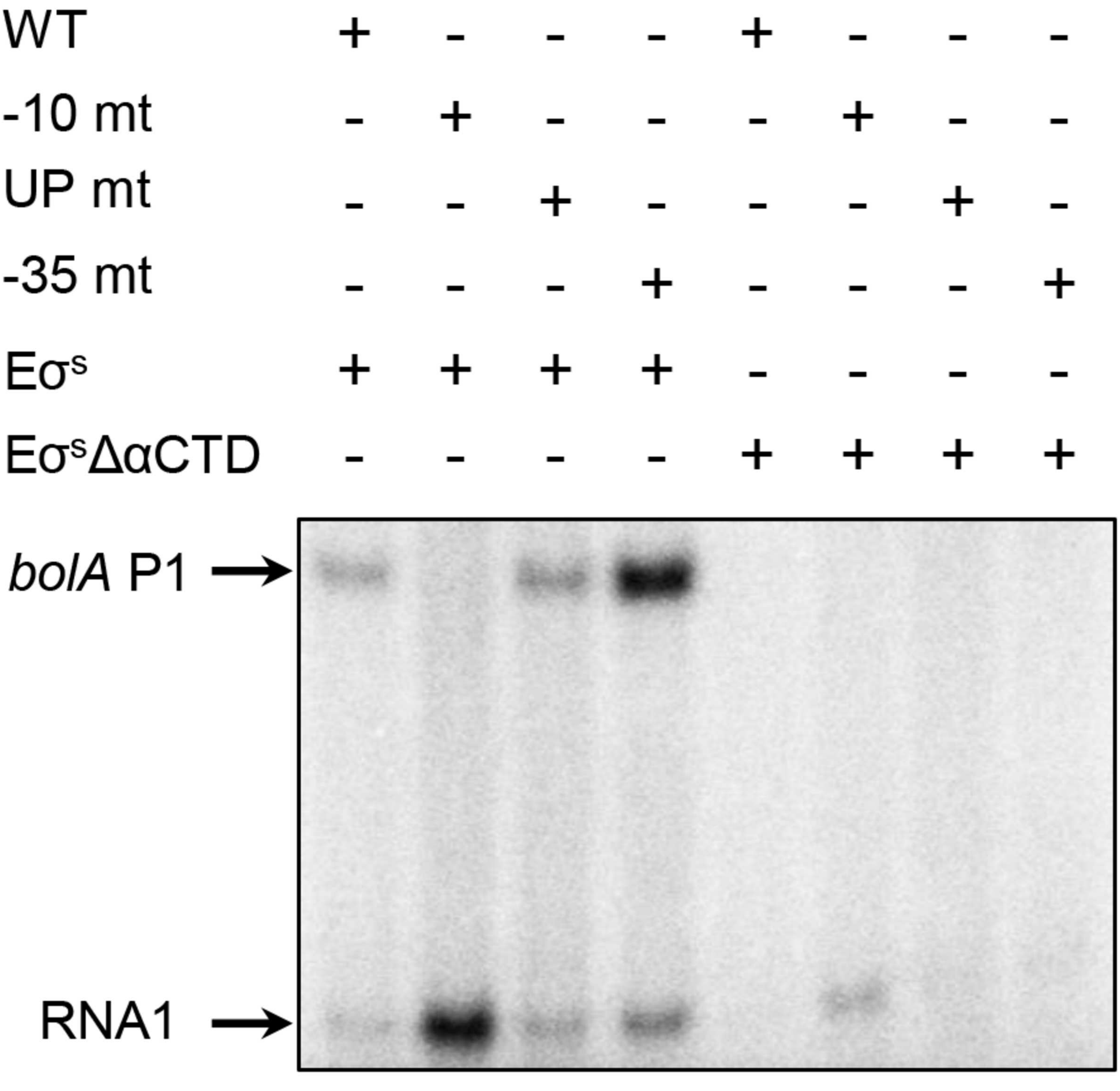

### figure-20.pdf

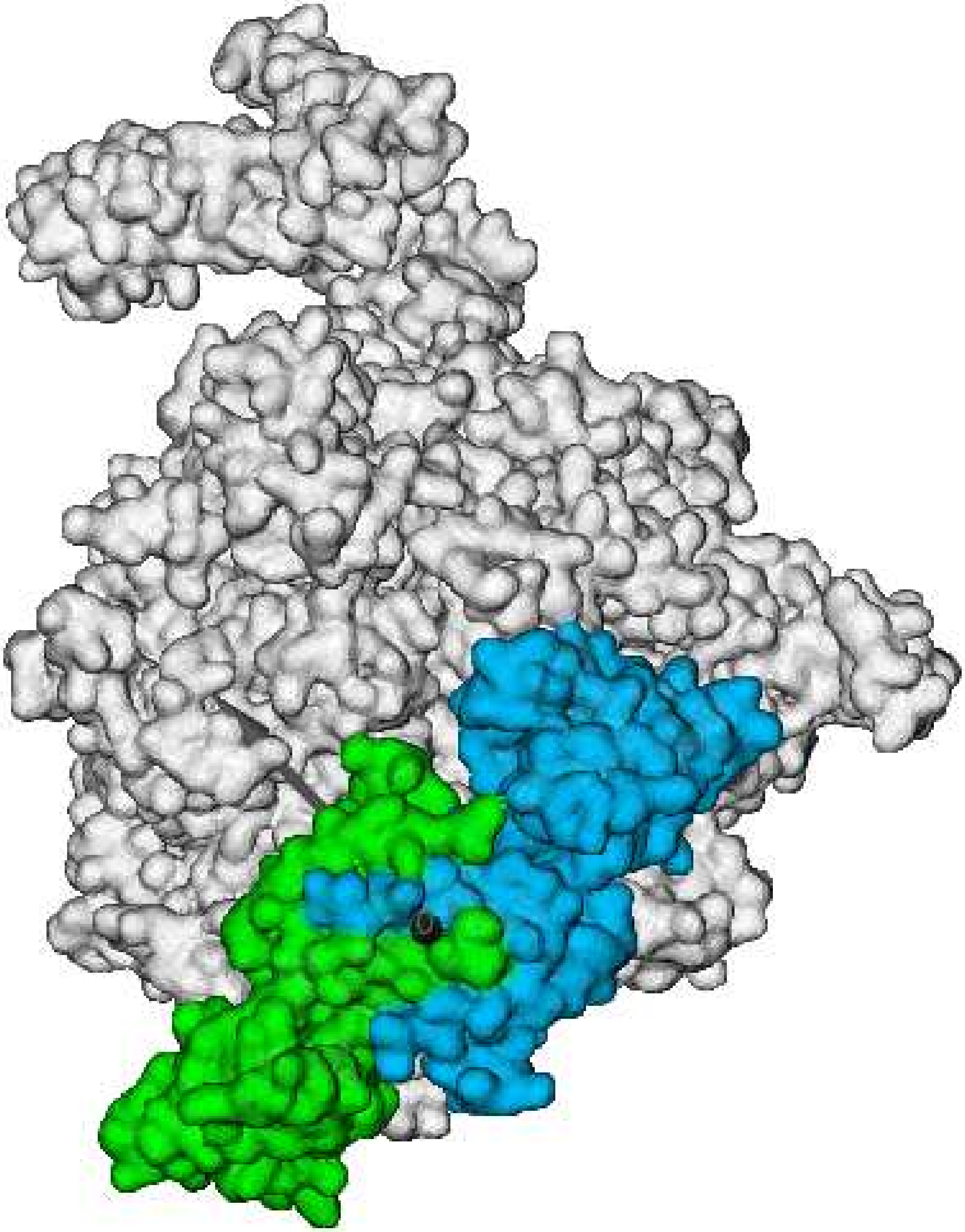

### sigma38cover.pdf

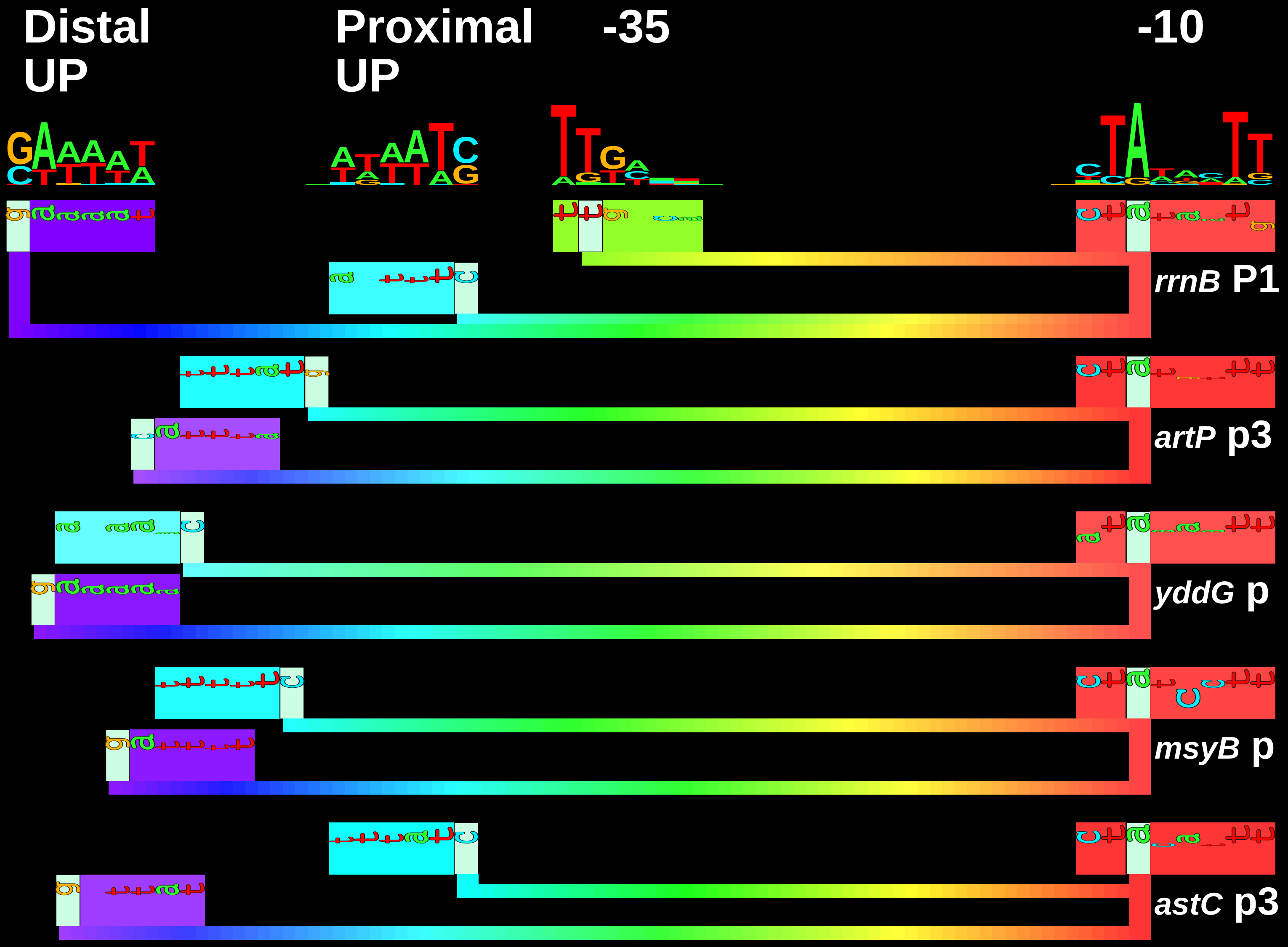
